# Functional constraints on insect immune system components govern their evolutionary trajectories

**DOI:** 10.1101/2021.07.02.450559

**Authors:** Livio Ruzzante, Romain Feron, Maarten J.M.F. Reijnders, Antonin Thiébaut, Robert M. Waterhouse

**Author notes:** Corresponding author (RMW).

## Abstract

The role of constraints in shaping evolutionary outcomes is often investigated in the contexts of developmental biology and population genetics, in terms of the capacity to generate new variants as well as how selection either limits or promotes consequent phenotypic change. Comparative genomics also recognises the role of constraints, in terms of shaping the evolution of gene and genome architectures, sequence evolutionary rates, and gene gains and losses, as well as on molecular phenotypes. Characterising patterns of genomic change where putative functions and interactions of system components are relatively well-described offers opportunities to explore whether genes with similar or analogous roles exhibit similar evolutionary trajectories, possibly governed by common constraints. Using insect innate immunity as our study system, we hypothesise that quantitative characterisation of gene evolutionary histories can define distinct dynamics associated with different functional roles. We develop metrics that quantify gene evolutionary histories, employ these to characterise evolutionary features of immune gene repertoires, and explore relationships between gene family evolutionary profiles and their roles in immunity to understand how different constraints may relate to distinct dynamics. We identified three main axes of evolutionary trajectories characterised by gene duplication and synteny, maintenance/stability and sequence conservation, and loss and sequence divergence, highlighting similar and contrasting patterns across these axes amongst subsets of immune genes. Our results indicate that where and how genes participate in immune responses limit the range of possible evolutionary scenarios they exhibit. Comparative genomics approaches therefore offer opportunities to characterise how functional constraints on different components of biological systems govern their evolutionary trajectories.

## Introduction

The concept of constraints in evolutionary biology encompasses a diverse array of interpretations and terminologies shaped by the approaches of different research fields (Antonovics & van Tienderen 1991). In general terms, constraints can be described as factors that limit or direct the process of natural selection leading to outcomes representing only a fraction of all theoretically possible scenarios. Constraints may impact the capacity to generate new variants as well as how selection either limits or promotes consequent phenotypic change, often considered in developmental biology (Richardson & Chipman 2003) and population genetics (Hoffmann 2013) contexts. Comparative genomics also recognises the role of constraints, in shaping the evolution of gene and genome architectures, sequence evolutionary rates, and gene gains and losses, as well as on the molecular phenotypes governed by their functional products (Koonin & Wolf 2010). For example, protein sequence evolution is constrained by requirements for maintaining proper protein structure and function, including during folding and interactions with other macromolecules (Worth et al. 2009). Functional constraints also impact the evolution of gene families, e.g. families of paralogs with or without essential genes exhibit dramatically different evolutionary regimes in terms of sequence divergence and duplication rates (Shakhnovich & Koonin 2006). These likely influence observed trends across the gene duplication spectrum that show a dichotomy of constrained single-copy control versus a multicopy licence for greatly relaxed copy-number restrictions (Waterhouse et al. 2011). Integrative analyses of evolutionary and functional constraints point to emergent properties such as a gene family’s “importance” or “status” characterised by low sequence divergence and propensity for gene loss with high expression levels, protein interactions, and essentiality; or a family’s “adaptability” manifested by high duplication levels, many genetic interaction partners, and a tendency of genes to be non-essential; or a family’s “reactivity” with high gain/loss and expression levels but low sequence divergence, a paucity of essential genes, and few physical or genetic interactions (Wolf et al. 2006). If such constraints limit the realm of possibilities in terms of allowed gene evolutionary trajectories then recurring patterns should be observable for genes evolving under similar constraints. Characterising these patterns in the context of a relatively well-studied system, where putative functional roles and interactions of member genes are well-described, offers an opportunity to explore whether genes with similar or analogous functions exhibit similar evolutionary trajectories, possibly governed by common constraints.

The insect innate immune system is relatively well-characterised with respect to the functional roles and evolutionary histories of key implicated pathways and component gene families. It confers remarkable resilience to encountered pathogens through the activation of powerful responses to neutralise and clear infections (Rolff & Reynolds 2009; Ligoxygakis 2017). The immune system comprises both humoral and cellular responses with components dedicated to recognising signs of infection, signalling cascades to activate primary defences and induce transcriptional responses, modulators that control the intensity and direction of responses, and effector proteins and biomolecules for pathogen killing. Many of the genes and their protein products implicated in these complex processes were first identified in the fruit fly, *Drosophila melanogaster* (Lemaitre & Hoffmann 2007; Imler 2014). Classical receptor proteins that recognise pathogen-associated molecular patterns include peptidoglycan recognition proteins (PGRPs) (Wang et al. 2019) and β-1,3-glucan recognition or gram-negative bacteria-binding proteins (GNBPs) (Rao et al. 2018). Pathogen recognition may then trigger immune signalling through the Toll (Valanne et al. 2011), Imd (Myllymäki et al. 2014), or the JAK/STAT (Myllymäki & Rämet 2014) pathways. Their activation leads to the translocation of transcription factors to the nucleus where the expression of effector genes such as those encoding antimicrobial peptides (AMPs) (Lazzaro et al. 2020) is upregulated. Defence responses are mediated by various cells and tissues including haemocytes, the fat body, and the midgut, and pathogen killing can occur via processes such as melanisation, phagocytosis, lysis, autophagy, and apoptosis (Hillyer 2016; King 2020), with RNA interference (RNAi) facilitating major antiviral defences (Mussabekova et al. 2017). These complex interactions collectively offer insects protection from a vast array of viruses, bacteria, fungi, protozoa, and nematodes.

Sequencing the *D. melanogaster* and *Anopheles gambiae* genomes provided the first opportunity for comparative genomic analysis of immune-related genes in insects (Christophides et al. 2002). Advances in genome sequencing technologies have facilitated an increasingly dense sampling of species to explore insect gene repertoires and perform cross-species comparisons to trace gene evolutionary histories (Waterhouse 2015). This has allowed comparisons beyond Diptera to include Hymenoptera (Evans et al. 2006; Brucker et al. 2012; Barribeau et al. 2015), Coleoptera (Zou et al. 2007), Lepidoptera (Tanaka et al. 2008), and Hemiptera (Gerardo et al. 2010), as well as expanded sampling of flies and mosquitoes (Sackton et al. 2007; Waterhouse et al. 2007; Bartholomay et al. 2010; Sackton et al. 2017). These comparative studies generally focused on the canonical immune-related gene repertoire, comprising genes that have been directly implicated in immune responses through experimental research, or indirectly linked to immunity through homology to known immune proteins (Bartholomay & Michel 2018; Waterhouse et al. 2020). Emerging patterns pointed to distinct evolutionary dynamics that characterise different immune phases: (*i*) gene and domain gains or losses (turnover) can create diversity in recognition modules; (*ii*) core signalling pathway members are almost always maintained as single-copy orthologues often with elevated levels of sequence divergence; (*iii*) modulators appear to form lineage-restricted units with members picked from large families often with high gene turnover rates; and (*iv*) effectors like AMPs show dynamic gains and losses or are lineage-restricted while oxidative defence effectors are widespread with low levels of sequence divergence. These observations provide specific examples and strong expectations of types of genes with similar functions that exhibit similar evolutionary trajectories, within the established framework of insect innate immunity that classifies genes and families into broad functional categories of recognition, signal transduction, modulation, and effector components.

These trends are based on observations from cross-species comparisons of insect immune gene repertoires. Here we hypothesise that comprehensive quantitative multi-feature characterisation of gene family evolutionary histories can define distinct dynamics associated with different functional roles in immune responses. Such detailed evolutionary profiling can then be used to address the question of whether gene families involved in common immune functional categories, modules, or processes exhibit similar evolutionary trajectories possibly driven by shared evolutionary constraints. We take advantage of genomic resources available for 22 mosquito species (Holt et al. 2002; Nene et al. 2007; Arensburger et al. 2010; Lawniczak et al. 2010; Marinotti et al. 2013; Jiang et al. 2014; Chen et al. 2015; Neafsey et al. 2015; Matthews et al. 2018; Ruzzante et al. 2019) to (*i*) develop a suite of metrics that quantify gene and gene family evolutionary histories, (*ii*) employ these metrics to characterise the evolutionary features of mosquito immune gene repertoires, and (*iii*) explore the relationships between gene family evolutionary profiles and their functional roles in immunity to understand how different constraints may relate to distinct dynamics. The resolution afforded by multi-species comparative analyses and our suite of gene sequence and copy-number evolutionary metrics reveals the evolutionary features that most clearly distinguish each family, and highlights similar and contrasting patterns across all immune gene families. Complementing knowledge-based functional categorisations with gene co-expression analyses identifies immune families that function in concert, revealing evolutionary-functional correspondences where most prominently, families involved in mosquito complement system responses show both high evolutionary similarities and high expression similarities.

## Materials and Methods

### Orthology, variation, alignment, and expression data

Orthologous Groups (OGs) of genes were defined using the OrthoDB (Kriventseva et al. 2015) orthology delineation procedure across 21 mosquitoes and 22 other insects. OrthoDB employs all-against-all protein sequence alignments to first identify all best reciprocal hits (BRHs) between all genes from each pair of species (Zdobnov et al. 2017). It then uses a graph-based clustering procedure that starts with BRH triangulation to progressively build OGs that include all genes descended from a single gene in the last common ancestor. Single nucleotide polymorphisms (SNPs) for *An. gambiae* PEST, including all synonymous and nonsynonymous SNPs in annotated coding regions, were retrieved using the BioMart data mining tool from VectorBase (Giraldo-Calderón et al. 2015). The SNPs derive from eight variation datasets hosted at VectorBase (Neafsey et al. 2010; White et al. 2011; Weetman et al. 2012; Markianos et al. 2016; Hammond et al. 2017; Miles et al. 2017; Wiltshire et al. 2018). Multi-species whole genome alignments were generated from the assemblies of 22 mosquitoes (Table S1) available from VectorBase. The alignment process starts with pairwise sequence comparisons that are then progressively combined following the species phylogeny using the MultiZ approach of the Threaded Blockset Aligner (Blanchette et al. 2004). Expression data for *An. gambiae* genes were retrieved from VectorBase (Expression Stats VB-2019-06) as log2 transformed expression values for 13’201 genes across 291 conditions (mean, variance, and number of replicates). Immune gene family co-expression analysis employed these expression statistics using a subset of the conditions to build co-expression modules. Co-expression analysis also employed clusters of genes defined by the VectorBase Expression Map (MacCallum et al. 2011), with gene membership of all clusters/cells retrieved from the AgamP4.11 VB-2019-02 map (comprising 12’672 genes and based on 202 conditions).

### *Anopheles gambiae* immunity gene catalogue

The catalogue of *An. gambiae* immune-related genes was built by combining and updating the results of previous comparative immunogenomics studies (Christophides et al. 2002; Waterhouse et al. 2007; Bartholomay et al. 2010; Neafsey et al. 2015). *An. gambiae* gene and orthologous group membership for 36 immune-related gene families and subfamilies are summarised in Table 1.

**Table 1.**
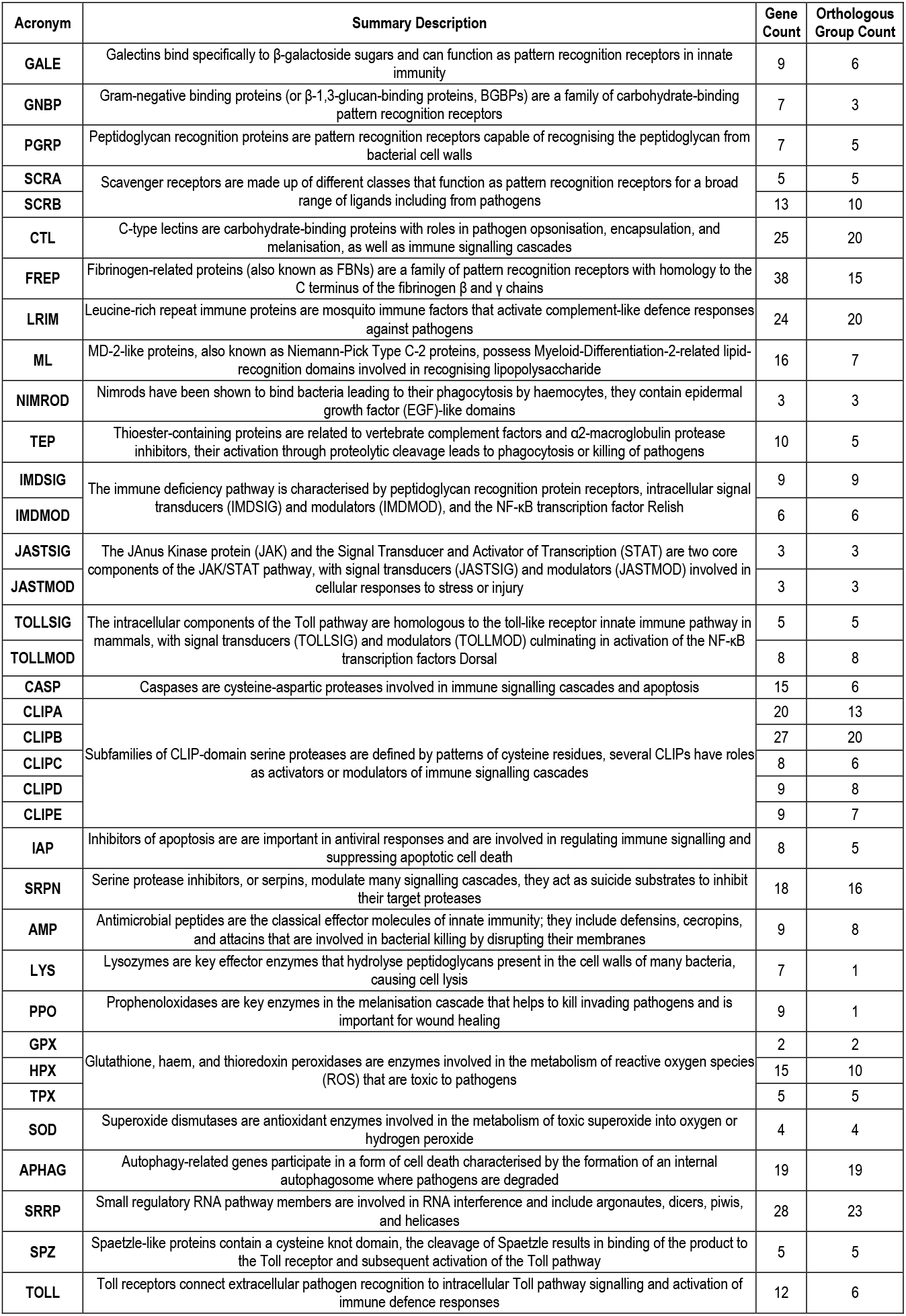
The *Anopheles gambiae* immunity gene catalogue.

### Orthology-based evolutionary features

Features were quantified as a suite of 13 orthology-based evolutionary metrics per OG that included: the evolutionary age (**AGE**) of the last common ancestor in terms of millions of years since divergence from the ultrametric species phylogeny; the universality (**UNI**) computed as the proportion of the total species present; the duplicability (**DUP**) computed as the proportion of species present with multi-copy orthologues; the average orthologue copy number (**ACN**); the copy number variation (**CNV**) computed as the standard deviation of orthologue counts per species present divided by the ACN. Phylogenetic analysis by maximum likelihood (PAML) (Yang 2007) was employed using the M0 model on the alignments of OG member sequences to compute the number of synonymous substitutions per synonymous site (**PDS**); the number of non-synonymous substitutions per non-synonymous site (**PDN**); and the non-synonymous to synonymous ratio (**SEL**). Gene turnover was estimated using the computational analysis of gene family evolution (CAFE) (Han et al. 2013) tool in order to quantify proportions gene gains (expansions, **EXP**), gene losses (contractions, **CON**), or no copy-number changes (stable, **STA**). Orthology data combined with genomic location data were used to quantify synteny conservation (**SYN**) as the proportion of orthologues that maintain their orthologous neighbours in the genomes of the other species. Finally, the evolutionary rate (**EVR**) of each OG corresponds to the average rate of protein sequence divergence normalized by the distance between each pair of species as computed by OrthoDB (Waterhouse et al. 2013).

### Variation-based and alignment-based evolutionary features

Five additional evolutionary feature metrics were computed from polymorphism data and whole genome alignments. The population genomics data for *An. gambiae* retrieved from VectorBase were used to compute per-gene metrics of the proportion of all coding-sequence SNPs that were non-synonymous (**NSP**), as well as the non-synonymous (**NSD**) and synonymous (**SSD**) SNP densities as the number of SNPs divided by the total coding-sequence length. Multi-species whole genome alignments were used to compute per-nucleotide metrics of conservation and constraint. Whole genome alignability (**WGA**) measures the proportion of the full set of 22 mosquitoes that were aligned to the *An. gambiae* reference genome for each nucleotide. PhastCons (Siepel et al. 2005) was used to estimate per-nucleotide levels of constraint (**PHC**) from the whole genome alignments. Per-nucleotide values were averaged over the full coding-sequence lengths of all genes to obtain per-gene metrics. The variation-based and alignment-based per-gene metrics were averaged over all genes in each OG to obtain the per-OG values for each of the metrics.

### Gene family metrics and comparisons

The *An. gambiae* canonical immunity gene catalogue defines immunity gene membership of subfamilies (e.g. cecropins, defensins, attacins), families (e.g. antimicrobial peptides), and broader categories (e.g. antimicrobial effectors), and the orthology datasets define gene membership of OGs. Thus, the gene family evolutionary metrics were computed by averaging values over all OGs containing genes belonging to each catalogued immune gene family. These family-level means for each metric were compared to the means of all other OGs that contain at least one *An. gambiae* immune gene to quantify the extent to which the metrics of the OGs of a given immune gene family differ from all other immune gene containing OGs, i.e. delta-mean 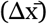. For graphical visualisation, 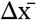 values were scaled by dividing by the absolute maximum 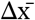 per evolutionary feature and plotted with the colour-blind safe RdYlBu palette from the *RColorBrewer* package from R (R Core Team 2021). The Wilcoxon rank-sum (Mann– Whitney U) test implemented in the *wilcox*.*test* function in R (default two-sided test) was used to test the significance of the difference of the distribution of each family’s OGs metric values (no scaling) compared to all other immune-related OGs for all metrics and each family. As several families contain few OGs, a permutation test implemented in R was also used to test the significance of the difference of the metric distributions. Observed 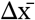 was compared to 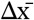 from permutations of all OG metric values randomly assigned to size-matched sets. The number of permutation differences that were greater than the observed difference, divided by the total number of permutations provides an empirical estimate of the probability of obtaining a 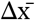 greater than the observed 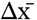 by chance.

### Clustering of gene family metrics

To assess and quantify the similarities of the evolutionary feature profiles, hierarchical clustering of the evolutionary features (n=18) and families (n=36) was performed with the *hclust* function in R. For all evolutionary feature metrics, both the means and the medians of all OGs per family were assessed. Prior to clustering, the *scale* function in R was used to normalise all metric values by subtracting the means and then dividing the (centered) values by their standard deviations. Dissimilarity matrices were computed with the normalised metric values using three correlation-based distance methods and the Euclidean distance method in R. Clustering with *hclust* was performed with all dissimilarity matrices using single, complete, average, and median linkage agglomeration methods. To estimate statistical support for the clustering of families and features, 10’000 bootstrap replicates were performed with the *pvclust* R package. In *pvclust*, the approximately unbiased (AU) p-values are computed using multiscale bootstrap resampling (Suzuki & Shimodaira 2006), and provide a confidence measure for each node of the cluster dendrograms of families and evolutionary features. The robustness of gene family clustering across all 16 tested distance-method combinations was further assessed by quantifying the co-occurrence of all pairs of families within subtrees of all 160’000 *pvclust* bootstrap replicates. This evolutionary profile similarity score (family subtree co-occurrence score) was computed and normalised as follows: (2 x co-occurrence of Family 1 & Family 2) / (co-occurrence of Family 1 with any Family + co-occurrence of Family 2 with any Family). Normalised scores of zero indicate that these pairs of families never appear in the same subtree and scores of one would indicate that they occur as sister lineages in all bootstrap samples from all distance-method combinations. Based on these assessments of clustering stability, the dissimilarity matrix from Pearson’s correlation method with the average linkage agglomeration method was selected. The hierarchical clustering results were visualised as heatmaps with corresponding family and evolutionary feature dendrograms showing AU support, plotted with the *gplots* and *dendextend* (Galili 2015) R packages. Principal Component Analysis (PCA) of the family by feature matrices of both median and mean metrics were performed with the *prcomp* function from the *stats* package in R.

### Gene and family co-expression analyses

Gene expression similarities amongst all pairs of immune-related families were quantified using the gene expression data and Expression Map (MacCallum et al. 2011) retrieved from VectorBase (Giraldo-Calderón et al. 2015). The map was analysed to quantify co-occurrences of gene family members in the same cell on the map (fine-scale resolution of gene co-expression), and in the same supercell, the cell and its immediate eight neighbouring cells on the map including toroidal neighbours (broader-scale resolution of gene co-expression). Pairwise family cell/supercell co-occurrence scores (expression similarity scores) were computed as the intersection, Family 1 ⋂Family 2, divided by the union, Family 1 ⋃Family 2 (i.e. number of cells with at least one gene from both Family 1 and Family 2 / number of cells with at least one gene from either Family 1 or Family 2). A score of zero: the pair of families have no member genes that cluster in the same cell/supercell. A score of one: all member genes from both families always cluster in cells/supercells with at least one member of the other family. Statistical significance of the family cell/supercell co-occurrence scores was assessed with a permutation test: scores were recomputed after gene to cell assignments were randomly shuffled (10’000 permutations) preserving the total number of cells and families, and the number of genes in each cell and each family. These were used to calculate an empirical estimate of the probability (p-value) of obtaining a co-occurrence score greater than the observed co-occurrence score by chance: the number of permutation scores that were greater than the observed score, divided by the total number of permutations. Complementary assessments of gene expression clustering were performed using the weighted correlation network analysis (WGCNA) approach (Langfelder & Horvath 2008) on a subset of 24 conditions selected from the VectorBase gene expression dataset including blood feeding experiments and tissues from (Marinotti et al. 2006; Neira Oviedo et al. 2008; Baker et al. 2011). Expression similarities of pairs of immune gene families and the significance of their co-occurrences were computed as for the Expression Map but using module membership rather than cell/supercell membership.

## Supplementary Materials and Methods

Additional details on the analysed species, genome assemblies, annotated gene sets, as well as software versions and analysis parameters are presented in the **Supplementary Materials**.

## Results and Discussion

### The evolutionary feature landscape of mosquito immunity

Profiles built from 18 quantified evolutionary features successfully delineate key similarities and differences amongst the catalogue of 36 canonical immune-related gene families and subfamilies (Figure 1). These features capture a broad spectrum of gene evolutionary dynamics including taxonomic spread, copy-number changes, protein- and DNA-level sequence divergence, conservation, and constraint, as well as genomic organisation, and population-level sequence variation. Family profiling employs a set of evolutionary features for which metrics are computed using gene orthology delineated across 43 insect species, whole genome alignments with 22 mosquitoes, and polymorphism data from *An. gambiae* (see Methods). The resulting evolutionary feature profiles for all families are visualised by averaging the metrics over all genes in orthologous groups (OGs) within each family. Compositions of families range from just a single OG for lysozymes (LYS, 7 genes) and prophenoloxidases (PPO, 9 genes), to 23 OGs with 28 genes for small regulatory RNA pathway members (SRRP) (Table 1). Contrasting the profile of a given family against the profiles of all other immune-related families reveals the evolutionary features that most clearly distinguish each family (Figure 1, Supplementary Figure S1).

**Figure 1.**
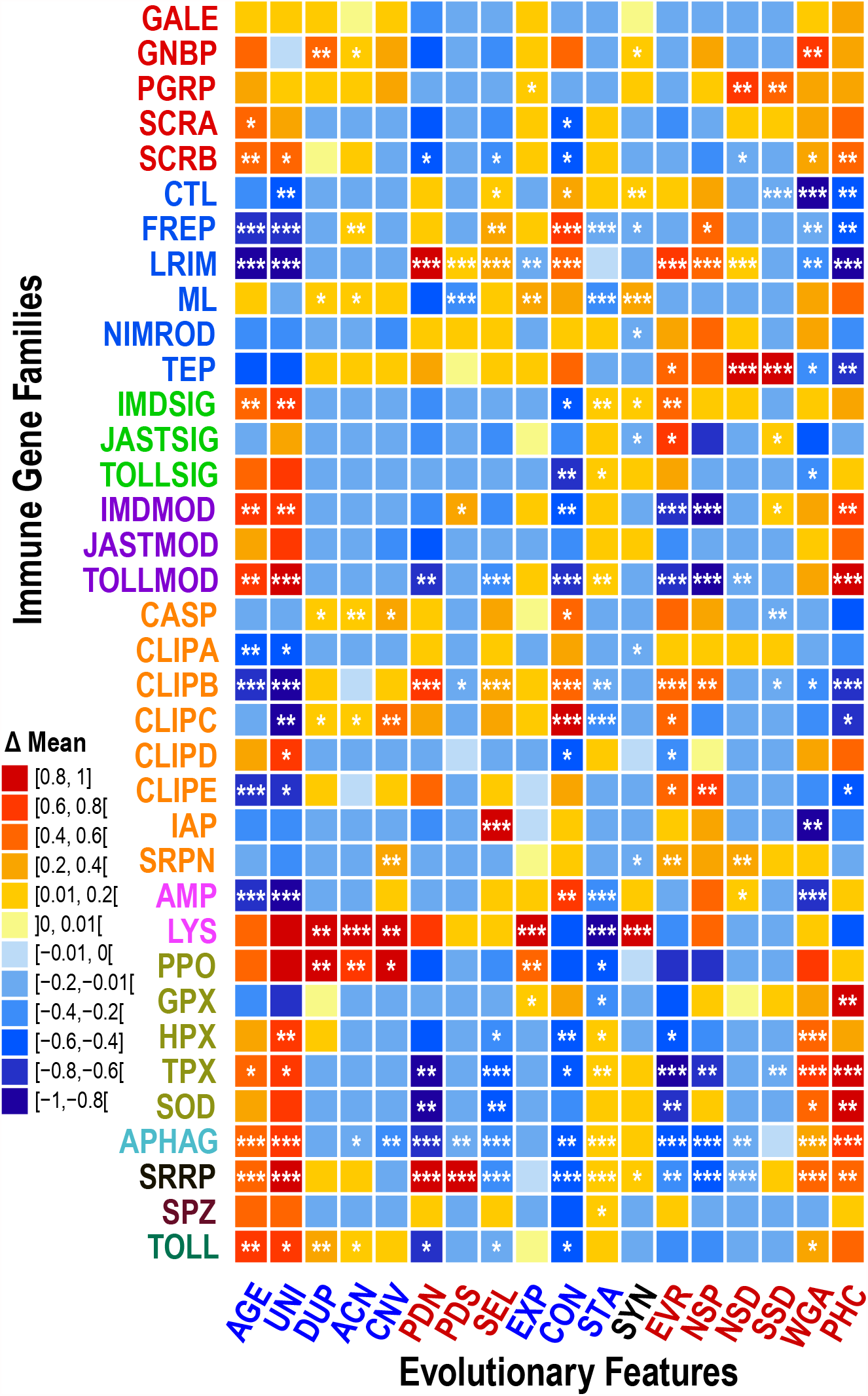
Evolutionary feature profiles of mosquito immune gene families. Evolutionary profiling highlights similar and contrasting patterns across all 36 immune gene families or subfamilies (rows). Deviations from the typical metric values for the suite of 18 evolutionary feature metrics (columns) are computed as the difference between the family mean and the average over all orthologous groups from other immune-related gene families 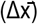. For visualisation, values of 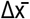 are scaled by the absolute maximum 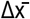 per metric, i.e. for each metric the distribution is transformed by dividing all values by the absolute maximum 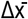. Values therefore range from a minimum of -1 for metrics where the largest deviation is below the mean, i.e. lower than other families, and the maximum of 1 for metrics where the largest deviation is above the mean, i.e. higher than other families. The significance of the difference of the distribution of metric values (no scaling) for each family compared to all other families was assessed using the Wilcoxon rank-sum (Mann– Whitney U) test and a permutation test (asterisks correspond to the lower p-value from these two tests; *** p≤0.01, ** p≤0.05, * p≤0.1). Family acronyms are defined in Table 1 and are coloured according to categories defined based on their putative roles in the principal immune phases: classical recognition (red), other recognition (blue), pathway signalling (bright green), pathway modulation (purple), cascade modulation (orange), antimicrobial effectors (pink), effector enzymes (olive green), autophagy (dark cyan), RNAi (black), cytokines (brown), and toll receptors (dark green). See text for definitions of evolutionary feature acronyms: taxonomic spread and copy-number features in blue; sequence-based features in red. Evolutionary feature profiles of mosquito immune gene families with median differences 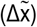 are presented in Figure S1.

This is clearly illustrated by the leucine-rich repeat immune genes (LRIMs) comprising 24 *An. gambiae* genes from 20 OGs, members of which interact with thioester-containing proteins (TEPs) to activate complement-like responses against pathogens (Povelones et al. 2009; Levashina & Baxter 2018). Their taxonomic age (AGE) and universality (UNI) are significantly lower, consistent with there being no detectable LRIM orthologues beyond mosquitoes (Waterhouse et al. 2010). They also exhibit fairly typical low duplicability (DUP), average copy-number (ACN), and copy-number variation (CNV), reflecting their mostly single-copy orthologue status across mosquitoes. These metrics describe the family as a whole while allowing for differences amongst members, e.g. the gene duplications that gave rise to three APL1/LRIM2 paralogs in one lineage of African *Anopheles* (Mitri et al. 2020). Estimates of non-synonymous substitutions per non-synonymous site (PDN) are high, and significantly so. They are not as extreme, but still significantly higher, for synonymous substitutions per synonymous site (PDS). Which together produce PDN:PDS ratios (SEL, i.e. dN/dS ratios) that are significantly higher, consistent with positive selection or relaxed constraint as observed in previous genus-wide analyses (Neafsey et al. 2015).

Gene gain/loss estimates for the LRIMs show significantly fewer expansions (EXP) and significantly more contractions (CON), but overall stability (STA) close to the mean, in agreement with the copy-number metrics. Conservation of genomic neighbourhood, or synteny (SYN) is slightly lower than average for LRIMs, while they notably show the most extreme significantly elevated protein sequence evolutionary divergence (EVR). Single nucleotide polymorphism data also show a significantly elevated proportion of non-synonymous SNPs (NSP) and significantly above average non-synonymous SNP density (NSD), with synonymous SNP density (SSD) slightly below the mean. The family as a whole thus appears to reflect the natural diversity and polymorphism observed for some family members (Rottschaefer et al. 2011; Holm et al. 2012). Finally, whole genome alignment data show that LRIMs are significantly less alignable (WGA) and significantly less constrained (PHC) than other immune gene families, reflecting the patterns observed with protein- and DNA-based measures of sequence divergence.

Family profiles highlight the extent to which each family deviates from or matches the typical metric values for each evolutionary feature. Gram-negative binding proteins (GNBPs) are characterised by high values for metrics capturing gene duplications (DUP and ACN) with high alignability across mosquito genomes (WGA), consistent with the birth of the B-type GNBP subfamily in the mosquito ancestor (Bartholomay et al. 2010). In contrast, Imd pathway signalling genes (IMDSIGs) are characterised as being relatively ancient (high AGE and UNI) and copy-number stable (low CON and high STA) with nevertheless a high protein sequence evolutionary rate (EVR), in agreement with previously observed evolutionary dynamics of immune signalling pathway members (Waterhouse et al. 2007). The subfamilies of CLIP-domain serine proteases are characteristically young (low AGE and UNI), except for CLIPDs which are older and significantly more taxonomically widespread (UNI), a contrast also reflected by several other evolutionary features. Differences amongst CLIP subfamilies could relate to the roles of catalytic and non-catalytic members in modulatory cascades and their hierarchies (El Moussawi et al. 2019).

The autophagy (APHAG) and small regulatory RNA (SRRP) pathway members share many features that are significantly different from the mean: they are ancient (high AGE and UNI), stable (low CON and high STA), and constrained (low SEL, EVR, NSP, NSD with high WGA and PHC). However they differ markedly with respect to estimates of dN and dS with both PDN and PDS being significantly lower for APHAGs and significantly higher for SRRPs. Their overall conservation and stability is consistent with both autophagy and RNAi being ancient cellular processes with roles beyond immunity, while their contrasting levels of substitutions could reflect different structural constraints on protein-protein versus protein-RNA interactions. The SRRPs do show above average DUP and ACN values, but not significantly so, consistent with reported single-copy orthologues of Argonautes 1 and 2 and duplications of Piwi/Aubergine in mosquitoes (Lewis et al. 2016). Indeed previous analyses of SRRPs suggested faster evolution in *Aedes* and *Culex* rather than *Anopheles* mosquitoes (Campbell et al. 2008), so conservative patterns observed here could be driven by the dataset consisting mainly of anophelines.

The distributions of computed OG metrics for all of the immune gene families for each evolutionary feature are presented in Additional File 1 together with statistical assessments of the significance of deviations from the typical metric values. The trends and significant differences observed across the suite of quantified features facilitate evolutionary profiling that recovers previous mostly qualitative observations and highlights similar and contrasting patterns across all immune gene families (Table 2).

**Table 2:**
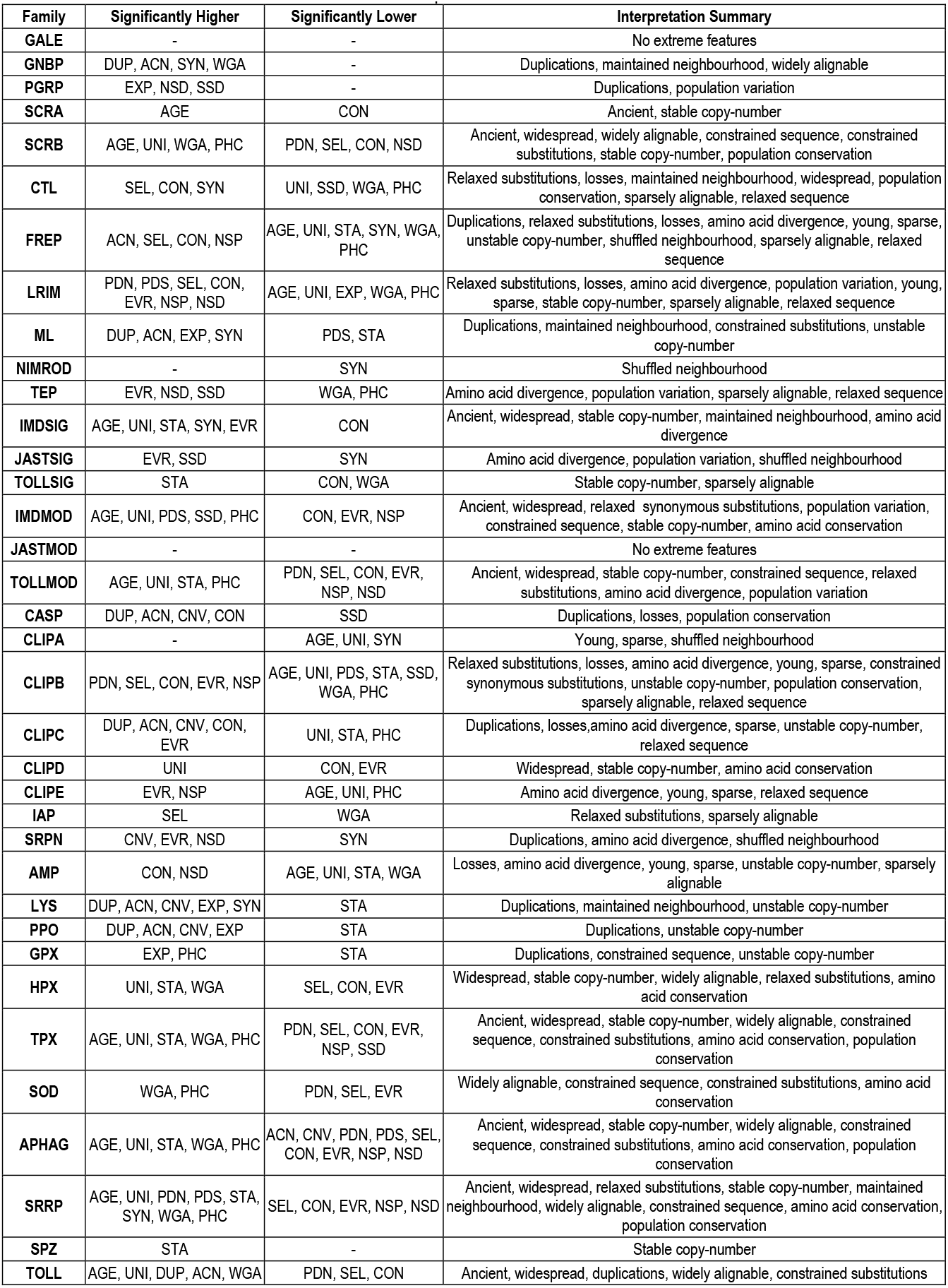
Characteristic evolutionary features of immune gene families and subfamilies. For each immune-related immune family, evolutionary features with significantly higher or significantly lower metrics compared to other immune families are listed with summarised interpretations.

### Families with similar functional roles exhibit similar evolutionary profiles

Several robust groupings of families and subsets of features are revealed when hierarchical clustering is applied to the matrix of evolutionary feature profiles of all immune gene families (Figure 2). Clustering aims to objectively delineate the hierarchical similarities amongst families and features to identify subsets of features that vary in concert, and groups of evolutionarily similar families (see Methods). Employing family median (Figure 2) and mean (Figure S2) metric values to build a dissimilarity matrix with Pearson’s correlation distances and performing bootstrapped clustering with the average linkage method results in several well-supported subsets and groupings. Using Pearson’s correlation distances for clustering aims to give weight to the metric directionalities rather than their magnitudes or ranks (Kassambara 2017), to identify families with similar evolutionary feature profiles. Nevertheless, clustering with alternative distance functions (Spearman’s and Kendall’s correlation, and Euclidean distances) and additional agglomerative clustering methods (Single, Complete, and Median linkage) confirms support for many of the observed hierarchical similarities (Supplementary Figures S3-6, Additional File 2). Furthermore, clustering using principal components instead of the metric values themselves also identifies several of the observed family groupings (Supplementary Figure S7). Overall, there are four main subsets of evolutionary features that consistently cluster together and somewhat more variable groupings of gene families depending on the combinations of metrics and methods applied.

**Figure 2.**
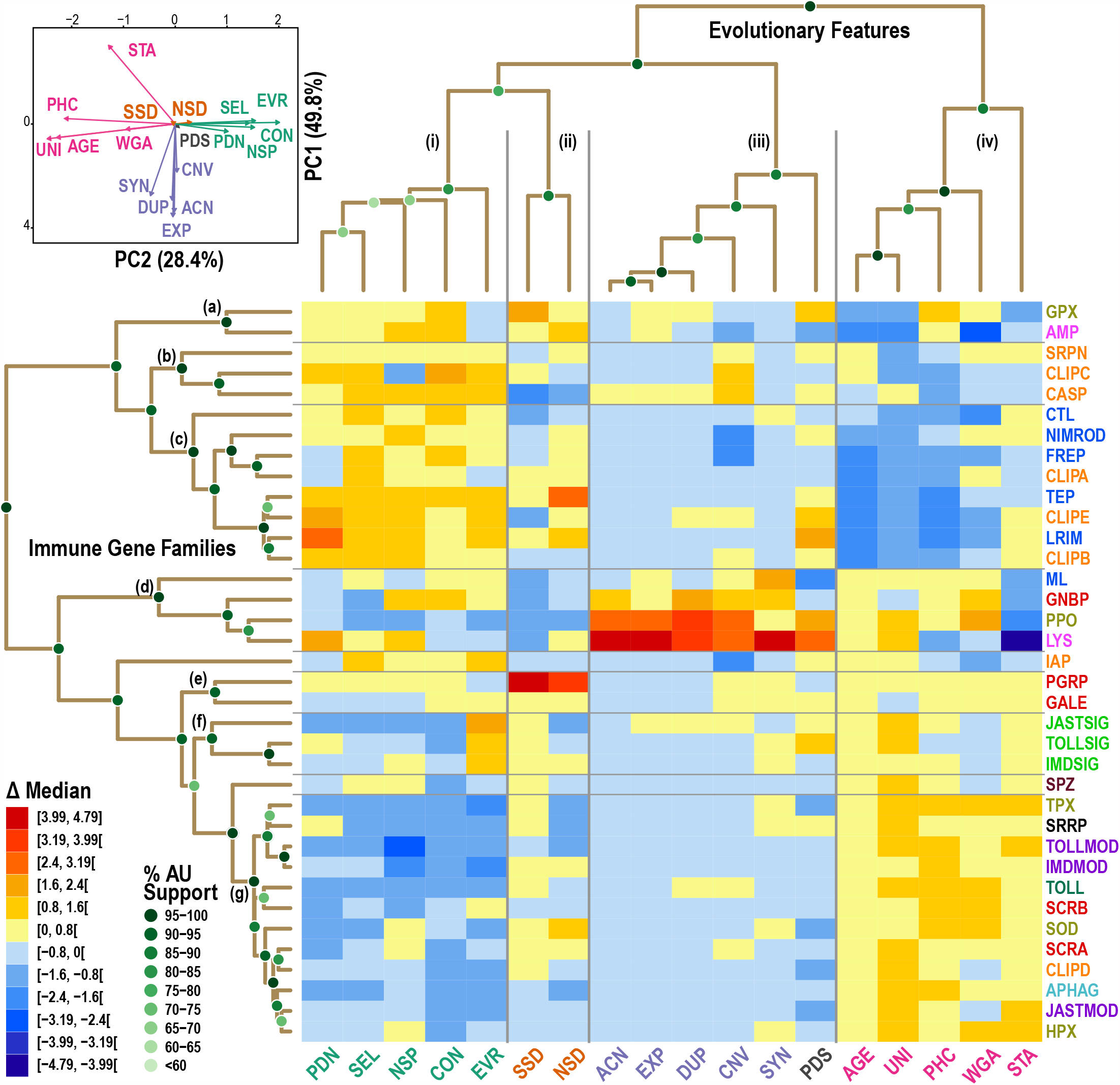
Clustering heatmap and dendrograms of immune families and their evolutionary features. Groupings of families and subsets of features delineated by hierarchical clustering using the matrix of evolutionary feature profiles of all immune gene families. Hierarchical clustering results are visualised for the immune families (n=36) and evolutionary features (n=18) using scaled median metrics with a Pearson’s correlation-based distance matrix and average linkage agglomerative clustering. The heatmap displays the relative values of the scaled metrics from low in blue to high in red. The dendrograms show the quantified distances (similarities) between each of the families, and between each of the features, and their groupings, determined by the clustering algorithm and distance method. Support for each node of the two dendrograms is shown with green-filled circles, using multiscale bootstrap resampling to estimate approximately unbiased (AU) support values. Principal Component Analysis (PCA) supports three major groupings of the four subsets of evolutionary features with PC1 and PC2 capturing 78.2% of the variance. Family acronyms are defined in Table 1 and are coloured according to categories defined based on their putative roles in the principal immune phases: classical recognition (red), other recognition (blue), pathway signalling (bright green), pathway modulation (purple), cascade modulation (orange), antimicrobial effectors (pink), effector enzymes (olive green), autophagy (dark cyan), RNAi (black), cytokines (brown), and toll receptors (dark green). See text for definitions of evolutionary feature acronyms, coloured according to groupings in the dendrogram and PCA. Clustering heatmap and dendrograms of immune families and their evolutionary features using mean metrics are presented in Figure S2.

First, with respect to evolutionary features, four subsets of features are repeatedly and robustly recovered: (i) PAML’s dN, PAML’s dN/dS (SEL), the proportion of non-synonymous SNPs, contractions (gene losses), and evolutionary rate (protein sequence divergence); (ii) densities of synonymous and non-synonymous SNPs; (iii) average copy-number, expansions, duplications, and copy-number variation, often also including synteny as in Figure 2; and (iv) age, universality, constraint, and alignability, often also including stability as in Figure 2. These subsets are also recovered when clustering using metric means rather than medians, with the exception of PAML’s dS (Figure S2). Principal Component Analysis (PCA) of both the median and mean metrics supports three major groupings of the four subsets, with PC1 dominated by set (iii) features contrasted by stability, and with PC2 clearly separating set (i) from set (iv) features (Figure 2, Figure S2, Figure S8).

Set (i) captures both gene losses and several features related to protein sequence divergence. PAML’s dN and dN/dS are computed from codon analysis of multiple protein sequence alignments, and the evolutionary rates are computed from amino acid similarities from all-against-all protein alignments, thus they are expected to vary in concert. The observed grouping of the proportion of non-synonymous SNPs with these protein-alignment-based metrics suggests that long-term divergence over millions of years of mosquito evolution is reflected in population-level polymorphism today. The grouping of gene losses with these sequence divergence and diversity features may appear less intuitive, however, correlations between the propensity for gene loss and sequence evolutionary rates have been observed previously from analyses of orthologues from seven distantly related eukaryotes (Krylov 2003). Here with a larger set of more closely related species (43 insects but mostly mosquitoes and other dipterans) this pattern is recovered while focusing exclusively on immune-related genes. Set (ii) groups together the expected correlated densities of genome-wide synonymous and non-synonymous SNPs.

Set (iii) captures all the copy-number features related to gene birth, linked to local genomic organisation (synteny). Gene duplications lead to higher and often more variable copy-numbers that are identified by CAFE as expansions, so these metrics should define different aspects of these features arising from the same underlying process and hence are expected to vary in concert. The link with maintained synteny suggests that duplicated genes often also maintain their local genomic neighbourhoods. However, this phenomenon is likely driven by only a small subset of families with both elevated duplication and synteny metrics: GNBPs, MD-2-like proteins (MLs), and particularly LYSs (Figure 1). For these immune genes it appears that retaining their relative genomic locations played an important role in maintaining their functionalities after duplicating in the mosquito or anopheline ancestor. Set (iv) captures the taxonomic spread features together with DNA-level sequence conservation and constraint, linked to gene family copy-number stability. This grouping clearly connects conservation at whole-gene and nucleotide levels, with older widespread immunity genes generally showing signs of greater constraints. In general, older genes do appear to evolve more slowly than younger ones (Albà & Castresana 2005), they are also longer, more highly expressed, and subject to stronger purifying selection (Wolf et al. 2009). In addition to constrained sequence evolution, genes functionally characterised as essential are more likely to be ancient and widespread (Waterhouse et al. 2011). This highlights the ancient origins and essential roles of several core components of the insect immune system that have been maintained over millennia.

The evolutionary profiles describe contrasting features of gene families and pathway members implicated in immune responses. The suite of features covers a wide spectrum of gene family evolutionary dynamics that can be broadly summarised by three main axes delineated by the major PCA groupings: axis 1, duplication and synteny; axis 2, maintenance/stability and sequence conservation; and axis 3, loss and sequence divergence. Axis 1 might be driven by only a subset of families, but the pattern is intuitive when considering the likely advantage of maintaining expression regulatory coordination across sets of duplicated genes. Axes 2 and 3 appear to reflect global trends in gene evolutionary dynamics observed in different taxa and over different timescales, suggesting that a broadly common set of rules also applies to the evolution of components of the immune system.

With respect to gene families (see Table 1 for family summary descriptions), several groupings of different sizes are recovered: from top to bottom in Figure 2 (a) antimicrobial peptides and glutathione peroxidases; (b) cysteine-aspartic and CLIPC proteases with serine protease inhibitors; (c) LRIMs, thioester-containing proteins, CLIPA protease homologues, CLIPB&E proteases, C-type lectins, and fibrinogen-related and Nimrod proteins; (d) GNBPs, MD-2-like proteins, lysozymes, and prophenoloxidases; (e) PGRPs and galectins; (f) Toll, Imd, and JAK/STAT signalling proteins; and (g) a large set comprising autophagy and RNA interference (RNAi) related proteins, Toll, Imd, and JAK/STAT pathway modulators, toll receptors, scavenger receptors A and B, CLIPD proteases, superoxide dismutases, as well as haem and thioredoxin peroxidases. Clustering with metric means rather than medians results in different hierarchies but with several broadly similar groupings including: LRIMs, CLIPA protease homologues, CLIPB&E proteases, and fibrinogen-related proteins; cysteine-aspartic and CLIPC proteases; GNBPs, MD-2-like proteins, lysozymes, and prophenoloxidases; and a large set comprising autophagy and RNAi related proteins, Toll, Imd, and JAK/STAT pathway modulators, toll receptors and galectins, scavenger receptors A and B, CLIPD proteases, superoxide dismutases, as well as haem and thioredoxin peroxidases (Figure S2). Similar variations of these groupings are obtained when clustering means or medians using alternative distance-clustering method combinations (Additional File 2). Combining this variation with results from bootstrapping provides a measure of evolutionary profile similarity between all pairs of families (see Methods). The families that most frequently cluster together using metric means (Figure S5) or medians (Figure S6) include: PGRPs, galectins, GNBPs, MD-2-like proteins, lysozymes, and prophenoloxidases; cysteine-aspartic and CLIPC proteases; LRIMs, thioester-containing proteins, CLIPA protease homologues, CLIPB&E proteases, and fibrinogen-related proteins; and a large set comprising autophagy and RNAi related proteins, Toll, Imd, and JAK/STAT pathway modulators, toll receptors, scavenger receptors A and B, CLIPD proteases, superoxide dismutases, as well as haem and thioredoxin peroxidases. Thus, while the gene family groupings are more variable across different distance-clustering method combinations than those of the evolutionary features, the results identify families with consistently similar evolutionary profiles.

Evolutionary profile clustering identifies features that are shared by genes and families within each of the major immune phases. Pairs of recognition protein families with similar profiles include PGRPs and galectins, A- and B-type scavenger receptors, and GNBPs and MD-2-like proteins, also indicating that MD-2-like proteins more closely resemble classical than other recognition families, thereby warranting their reclassification (Figure 2). PGRPs can bind bacterial cell wall Dap- or Lys-type peptidoglycans (Wang et al. 2019), while galectins can bind surface β-galactosides (Vasta 2020). Similarly, GNBPs can recognise β-1,3-glucans that make up structural polysaccharides of yeast cell walls (Rao et al. 2018), while MD-2-like proteins can bind lipopolysaccharides from the outer membrane of Gram-negative bacteria (Shi et al. 2012). A- and B-type scavenger receptors may have broader ligand specificities including lipoproteins and surface molecules of Gram-negative and Gram-positive bacteria (Alquraini & El Khoury 2020). As important pattern recognition receptors in animal immunity, these are all expectedly old families, however, despite interacting with pathogens they remain relatively constrained (DNA-level) and do not show extreme protein sequence divergence (Figure 2). This apparent lack of evidence for an arms race scenario may in fact reflect the relatively limited structural diversity of the main microbial ligands -peptidoglycan, β-1,3-glucan, lipopolysaccharide- they must bind to or cleave.

Signalling genes of the Toll, Imd, and JAK/STAT pathways group together, being generally ancient and stable but with remarkably elevated rates of protein sequence divergence. Their copy-number stability is possibly a reflection of constraints imposed by the large disruptive potential of duplicates on core signal transduction functionality. Their protein products work together as interacting partners, including the death-domain-mediated MyD88-Tube-Pelle complex of the Toll pathway (Valanne et al. 2011), the Imd pathway’s Imd-Fadd-Dredd, Tab2-Tak1, and IκB kinase complexes (Myllymäki et al. 2014), and the Domeless-Hopscotch complex of the JAK/STAT pathway (Myllymäki & Rämet 2014). Their greater sequence divergence could therefore be explained by the accumulation of compensatory amino acid changes that maintain key interactions amongst these partners, and overall pathway functionality. The signalling pathway modulators are also old and stable, but instead show constrained sequence evolution. These include several enzymes, such as ubiquitinases like Effete and Bendless, or E3 ligases like Pellino and Pias, which are likely under strong constraints to maintain their enzymatic activities. They are involved in proteasomal degradation and are therefore also critical for many other processes beyond immune signalling (Glickman & Ciechanover 2002). Other enzymes including the superoxide dismutases as well as the haem and thioredoxin peroxidases involved in reactive oxygen species metabolism (Hillyer 2016), show similarly conservative evolutionary profiles (Figure 2). Proteolytically activated PPOs oxidise phenols in the melanin production process (Nakhleh, El Moussawi, et al. 2017) and also show similar sequence constraints, however, multiple gene duplications result in an evolutionary profile that is radically different. Thus while there is some variation, in general the functional constraints on these types of enzymes appear to restrict their patterns of molecular divergence.

Members of ancient pathways controlling RNAi (SRRP) and autophagy (APHAG) responses group with other conservative evolutionary profiles characterised by low gene turnover and low sequence evolutionary rates (Figure 2). In contrast, much more dynamic evolutionary profiles characterise the grouping of families of immune cascade modulators like C-type lectins, CLIPA protease homologues and CLIPB&E proteases, regulators of melanisation responses like serine protease inhibitors, and key players in mosquito complement-like responses, like thioester-containing proteins and LRIMs. While melanisation is conserved across arthropods (Hillyer 2016), the proteolytic cascades that trigger or dampen melanin production often involve lineage-specific members of large gene families including these dynamically evolving modulators (Gulley et al. 2013; Meekins et al. 2017; El Moussawi et al. 2019; Bishnoi et al. 2019). The complement-like responses centred on TEPs and LRIMs are specific to mosquitoes, and are also triggered and regulated by members of these large and dynamic families (Blandin et al. 2008; Fraiture et al. 2009; Povelones et al. 2009, 2013, 2016). Based on understanding molecular functions of only a limited number of genes from these families, it appears that immune responses requiring such finely tuned activation, amplification, and deactivation processes source components from dynamically evolving families from which to build functional modules. The families involved are characterised with evolutionary profiles showing a pattern of younger and less widespread orthologues, with lower sequence constraints, and often elevated signatures of selection and population-level variation. This dynamism is more consistent with an arms race scenario, where the effectiveness of such functional modules is continuously being tested by evolving pathogen attacks and evasion strategies.

### Co-expression analyses identify immune families that function in concert

Analysis of multi-sample gene expression data shows that families with the strongest fine-scale or broad-scale expression similarities include many pairs whose members are known to function together *in vivo* (Figure 3, Figure S9). Thus, without presupposing any functional categorisations, the similar expression profiles highlight families whose members are likely working together across different conditions. Gene expression-based quantification of functional similarities amongst immune gene families provides an alternative objective classification that complements the classical categorisations based on their putative roles in key immune responses. The VectorBase Expression Map (MacCallum et al. 2011) defines clusters of genes with similar expression profiles for 12’672 genes using normalised data across 202 conditions, enabling the quantification of fine-scale or broad-scale gene expression similarities amongst all pairs of immune-related families (see Methods). Visualising pairwise family similarities as a spring model layout network optimised with the *neato* tool from the Graphviz package (Gansner & North 2000; Gansner et al. 2005) identifies subsets of families with putative roles in common immune processes (Figure 3). These prominently include a quintet of families with highly and significantly overlapping expression patterns: LRIMs, TEPs, FREPs, CLIPAs, and CLIPBs (Figure 3), with members implicated in coordinating and executing mosquito complement system responses (Povelones et al. 2016; Reyes Ruiz et al. 2019).

**Figure 3.**
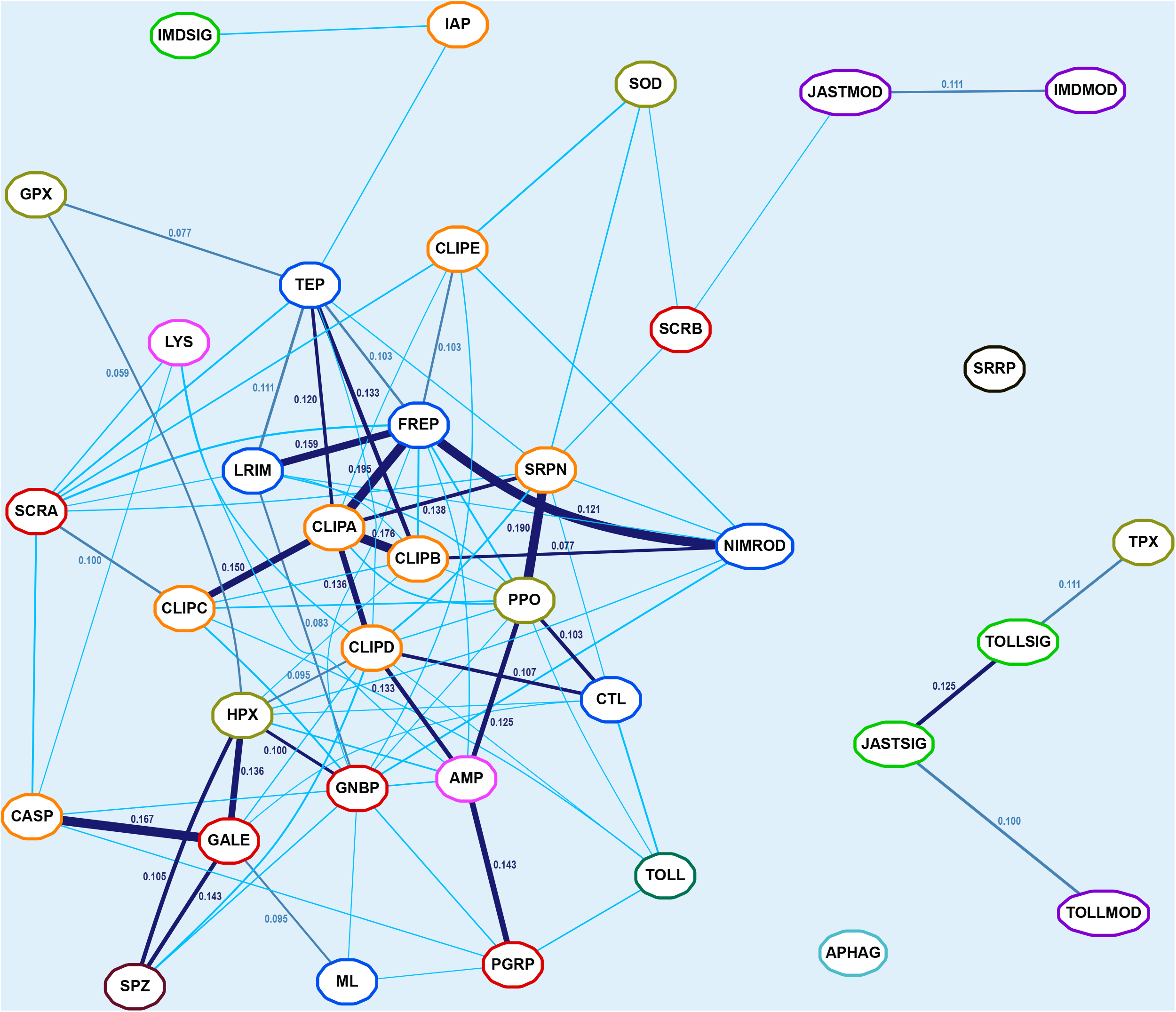
Network of immune family expression similarities based on the VectorBase Expression Map. The network layout optimised with a spring model provides a two-dimensional visualisation of expression similarities for pairwise comparisons of all 36 immune-related gene families computed as gene co-occurrence scores across the VectorBase Expression Map (AgamP4.11 VB-2019-02). Families with more similar gene expression profiles are placed closer together in the graph. Significant co-occurrences are indicated with connecting lines: light blue <0.05, royal blue <0.01, and dark blue <0.005 with line thickness scaled to the p-value and co-occurrence scores indicated for all pairs with p<0.01. Families are coloured according to categories defined based on their putative roles in the principal immune phases: classical recognition (red), other recognition (blue), pathway signalling (bright green), pathway modulation (purple), cascade modulation (orange), antimicrobial effectors (pink), effector enzymes (olive green), autophagy (dark cyan), RNAi (black), cytokines (brown), and toll receptors (dark green).

*An. gambiae* TEP1 forms a stable protein complex with a heterodimer of LRIM1 and APL1A/B/C (LRIM2 paralogues) in the haemolymph until the complement response is activated (Fraiture et al. 2009; Povelones et al. 2009; Williams et al. 2015), so coordinated co-expression of these genes is likely important for their functions. Like the LRIMs and TEPs, the FREPs are also found in the haemolymph, and several members are infection-responsive and important for defence, e.g. FREP57/FBN8, FREP13/FBN9, and FREP40/FBN39 (Dong et al. 2006; Dong & Dimopoulos 2009; Simões et al. 2017). FREPs themselves might dimerise or oligomerise, but whether they interact directly with TEPs and/or LRIMs in mosquitoes remains unknown, although evidence from snails indicates that FREPs and TEPs do interact (Li et al. 2020), and the observed expression similarities support at least some functional, if not physical, interaction. CLIPA serine protease homologues are positive and negative regulators of immune responses mediated by TEP1, e.g. CLIPA8 (Schnitger et al. 2007), SPCLIP1/CLIPA30 (Povelones et al. 2013), CLIPA2 (Yassine et al. 2014), CLIPA14 (Nakhleh, Christophides, et al. 2017), and CLIPA28 (El Moussawi et al. 2019). These regulatory modules also involve the catalytically active CLIPBs e.g. CLIPB14 and CLIPB15 (Volz et al. 2005), CLIPB8 (Zhang et al. 2016), and CLIPB10 (Zhang et al. 2021), and together CLIPAs and CLIPBs are also key modulators of melanisation responses (Volz et al. 2006). The available evidence therefore supports the family-level expression analyses that demonstrate highly and significantly overlapping expression patterns (Figure 3) of members of this quintet of families that function in concert.

Of this quintet, expression of CLIPA protease homologues is additionally strongly and significantly similar to that of CLIPCs, CLIPDs, and SRPNs (serpins, or serine protease inhibitors). The CLIPC9 protease has recently been shown to regulate melanisation downstream of SPCLIP1/CLIPA30, CLIPA8, and CLIPA28, and to be inhibited by SRPN2 (Sousa et al. 2020). CLIPC2 may function together with SRPN7 controlling the activation of effector mechanisms (Blumberg et al. 2013). Specific roles for CLIPDs, which show an evolutionary profile distinct from the other CLIPs (Figure 2), remain largely unknown. Serpins themselves are most similar in expression to prophenoloxidases, both of which would need to be replenished after being depleted during melanisation responses (Gulley et al. 2013; Nakhleh, El Moussawi, et al. 2017). The PPOs in turn appear significantly similar to the C-type lectins, which are generally considered glycan-binding recognition proteins, but at least two members -CTL4 and CTLMA2- are key regulators of melanisation downstream of immune recognition (Schnitger et al. 2009; Bishnoi et al. 2019). The family-level expression similarities (Figure 3) therefore likely reflect the functional links amongst the CLIP, CTL, and SRPN family members that modulate the activation of melanisation, and the PPO enzyme effectors of melanisation activity.

Amongst classical recognition proteins, PGRPs and GNBPs are most similar, and their expression patterns both closely match those of antimicrobial peptides and MD-2-like lipid recognition proteins. These similarities are likely driven by the upregulation of members of these gene families upon infection or following a blood meal, which promotes growth of the gut microbiota, e.g. in response to blood-feeding (Dana et al. 2005), microbes (Aguilar et al. 2005), *Plasmodium* (Dong et al. 2006), or fungi (Ramirez et al. 2020). They are nevertheless not as tightly interconnected as components of the complement and melanisation responses, possibly reflecting the contrast between broad-scope protection of these systems versus the generally much more pathogen-specific activities of different families of recognition proteins and antimicrobial effectors. Indeed feeding into and/or being transcriptionally activated by different immune signalling pathways means that these families may be thought of as performing analogous roles rather than functioning in concert *per se*. However, learning more about signalling crosstalk and response overlap has shifted thinking from traditional functional distinctions amongst immune pathways (Kounatidis & Ligoxygakis 2012). Thus, these similarities might reflect somewhat overlapping responses, but also a common readiness or priming to face newly perceived threats.

Notably, expression patterns of pathway signalling and modulation components remain distinct from the recognition and response families: Imd and JAK/STAT pathway modulators are significantly similar, while Toll pathway modulators group together with Toll and JAK/STAT pathway signalling members. Genes involved in RNAi (SRRP) and autophagy (APHAG) responses do not show significant similarities in expression patterns to other families, however, SRRP and APHAG genes have highly and significantly overlapping expression patterns at broad-scale resolution, and are most similar to modulators of all three pathways (Figure S10). At broad-scale resolution the distinction between pathway signalling/modulation and recognition/response families is accentuated, while the melanisation and complement responses become more closely interlinked. Many of the most similar families also show substantially overlapping expression patterns when quantifying similarities across co-expression modules built from a subset of immune-related experimental conditions (see Methods, Figures S11-S12, Table S2, Additional File 3). For example, families implicated in complement system responses again show similar expression patterns (Figure S13), and at a broader-scale resolution become more closely associated with melanisation responses (Figure S14). At broad-scale resolution pairs of similar recognition families include GNBPs and PGRPs, GNBPs and MD-2-like proteins, as well as galectins and B-type scavenger receptors, while at both resolutions Imd and JAK/STAT pathway signalling members are highly and significantly similar. Multi-condition co-expression analysis therefore identifies gene expression similarities amongst sets of immune-related families with members that are known or inferred to function in concert.

### Complement-related families exhibit elevated evolutionary-functional similarities

Immune gene family evolutionary-functional correspondences are revealed by employing quantifications of evolutionary similarities based on gene family feature profiling and of functional similarities based on gene family expression patterns (Figure 4, Additional File 4). Most prominently, families involved in mosquito complement system responses show both high evolutionary similarities and high fine-scale and broad-scale expression similarities: recognition family pairs of LRIMs-TEPs, FREPs-TEPs, and FREPs-LRIMs, as well as modulator-recognition family pairs of CLIPAs with FREPs and TEPs, and CLIPBs with FREPs, TEPs, and LRIMs. Members of these principal complement-response gene families exhibit common expression and evolutionary profiles suggestive of common constraints. Both TEPs and LRIMs are also highly evolutionarily similar to CLIPEs, for which specific roles in complement responses remain largely unknown, but with which their expression similarity increases at broad-scale resolution, albeit remaining non-significant. The CLIPA protease homologues and CLIPB proteases form a highly similar pair, but their strong and significant expression similarity is not maintained at broad-scale resolution, suggesting tight functional coupling of these key modulators. Conversely, CLIPB and CLIPE modulators also form a highly similar pair, but with strong and significant expression similarity only at broad-scale resolution. In contrast, FREP-NIMROD expression similarity is maintained at both resolutions and it is amongst the most significant of all family pairs that also show high evolutionary similarities. Although a much smaller gene family than the FREPs, NIMRODs including *draper, nimrod*, and *eater*, are also infection-responsive and important for defence (Midega et al. 2013; Estévez-Lao & Hillyer 2014). Combining results from evolutionary profiling and knowledge-blind functional clustering therefore identifies families that appear both evolutionarily and functionally similar. These similarities are notably pronounced for families with members known to function in concert to coordinate and execute mosquito complement system responses (Povelones et al. 2016; Reyes Ruiz et al. 2019).

**Figure 4.**
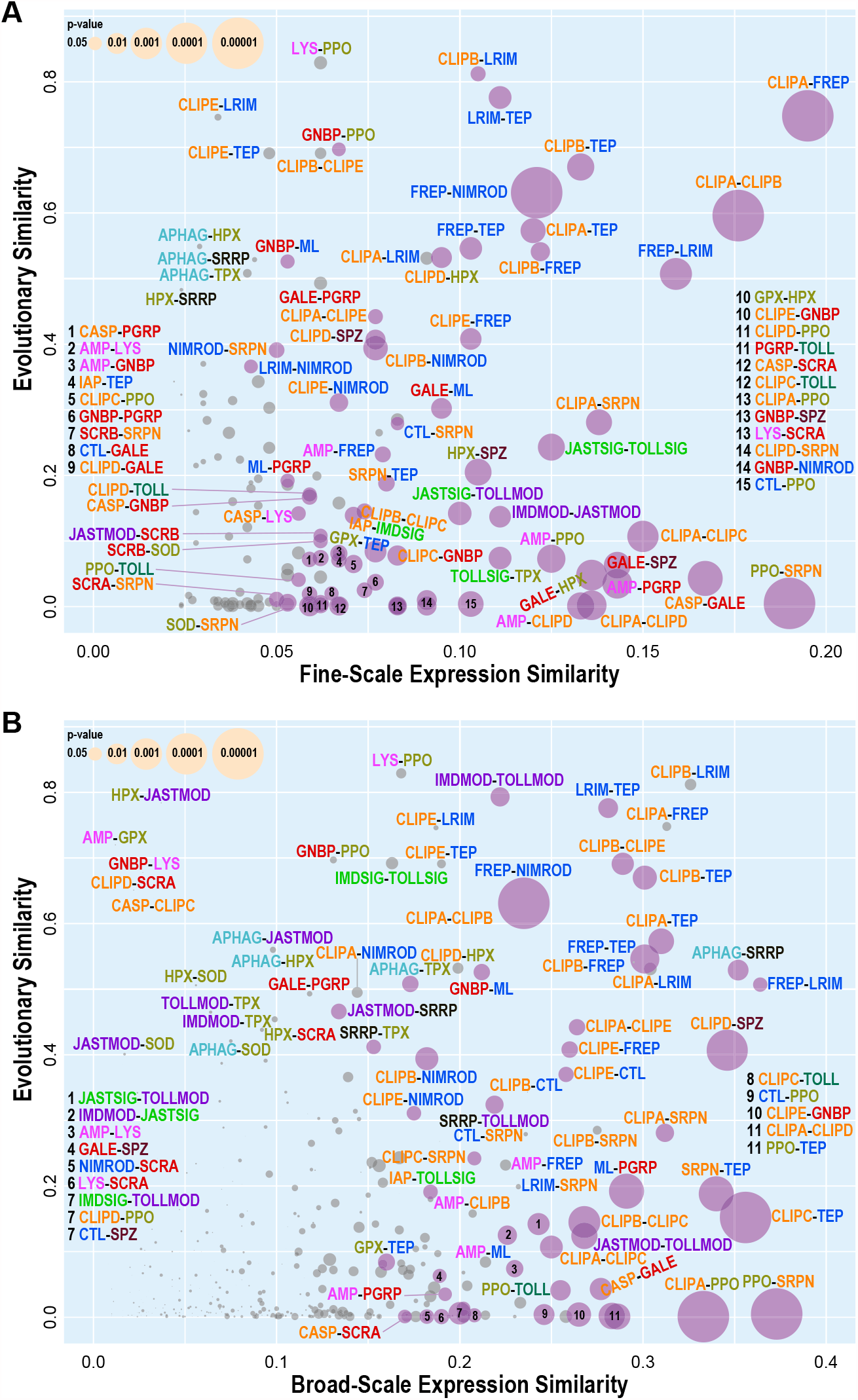
Pairwise comparisons of immune family expression similarity and evolutionary similarity. Evolutionary similarities (based on feature metric medians) of pairs of gene families are compared with their expression similarities at (A) fine-scale resolution and (B) broad-scale resolution. Pairs of families with significant (p<0.05) gene expression co-occurrence scores are shown with purple circles, with non-significant pairs shown in grey, and with circle sizes scaled by the p-value. Family acronyms are defined in Table 1 and are coloured according to categories defined based on their putative roles in the principal immune phases: classical recognition (red), other recognition (blue), pathway signalling (bright green), pathway modulation (purple), cascade modulation (orange), antimicrobial effectors (pink), effector enzymes (olive green), autophagy (dark cyan), RNAi (black), cytokines (brown), and toll receptors (dark green).

Additional families with above average evolutionary and expression similarities at both resolutions include another pair of modulators (CLIPA-SRPN), and another modulator-recognition pair (CLIPE-FREP). While CLIPAs and SRPNs are known to function together in cascades regulating melanisation (El Moussawi et al. 2019), potential functional interactions between CLIPEs and FREPs remain to be explored. The melanisation modulator-effector pair of SRPNs and PPOs shows the highest expression similarity at both resolutions, but with negligible evolutionary similarity, suggesting that regulating these responses and executing them are subject to different constraints. Amongst other recognition proteins, MD-2-like proteins (MLs) show above average evolutionary and expression similarities to the classical recognition families of galectins (GALEs) at fine-scale resolution, and PGRPs at broad-scale resolution with lower but still significant expression similarity at fine-scale resolution. Compared to galectins or PGRPs, the MLs are evolutionarily more similar to GNBPs, with which they show lower, but still significant, expression similarity. These patterns suggest analogous functionalities —recognition of foreign— with different specificities for lipopolysaccharides, β-galactosides, peptidoglycans, or β-1,3-glucans, that arise depending on the pathogen/microbe community composition. Common constraints faced by classical recognition phase families appear to produce similarities amongst their evolutionary trajectories, with functional similarities quantified through gene expression patterns possibly arising through immune pathway signalling crosstalk and priming (Kounatidis & Ligoxygakis 2012).

Evolutionarily similar families that only show high expression similarities at broad-scale resolution include modulators of the Imd and Toll pathways (IMDMOD-TOLLMOD) and genes involved in autophagy and RNAi responses (APHAG-SRRP). At fine-scale resolution, pathway components from JAK/STAT and Toll signalling (JASTSIG-TOLLSIG), Imd and JAK/STAT modulation (IMDMOD-JASTMOD), and JAK/STAT signalling and Toll modulation (JASTSIG-TOLLMOD) also show above average evolutionary and expression similarities. These pathways and responses play key roles in processes other than immunity, including in development and morphogenesis, so their gene expression-based functional similarities will vary depending on the conditions examined. Their functional similarities are more stably evident when the modules are abstracted to analogous phases of signal input, signal processing, and signal output. Whether functionally similar or analogous, these immune-related pathways and responses exhibit common conservative evolutionary profiles that distinguish them from other more dynamically evolving components of the immune system (Figure 2). These constrained evolutionary features could result from the effects of pleiotropy, and possibly the modular architectures, on the trade-offs during adaptive evolution producing a limited range of available trajectories (Mauro & Ghalambor 2020).

## Conclusions

Through quantitative evolutionary feature profiling of genes and gene families, integrated with knowledge- and expression-based functional categorisations, our multi-species comparative immunogenomic analyses identified evolutionary-functional correspondences suggesting that constraints on genes with similar or analogous functions govern their evolutionary trajectories. The profiles delineate whether and how each family deviates from the feature value distributions of other families, and provide the substrate for clustering to define similarities amongst families and features. We employed insect innate immunity as our study system because the key implicated pathways and component gene families have been well-characterised. While acknowledging that responses to infections involve diverse processes beyond the canonical immune system (Sackton 2019), this prior knowledge provided specific examples and strong expectations of types of genes with similar functions and distinguishing patterns of evolution, enabling the interpretation of observed correspondences within an established framework. Feature analysis identified three main axes of evolutionary trajectories characterised by gene duplication and synteny, gene maintenance/stability and sequence conservation, and gene loss and sequence divergence. Clustering highlighted similar and contrasting patterns across these axes amongst subsets of immune gene families. For example, classical recognition families, including the herein reclassified MD-2-like proteins, showed patterns likely reflecting the limited structural diversity of the principal microbial ligands with which they interact. Pathway signalling genes on the other hand exhibited trajectories that could relate to physical interactions of protein complexes and constraints from the effects of pleiotropy and disruptive effects of gene duplicates on signal transduction. Functional similarities defined by co-expression analyses recovered sets of immune-related families with members that are known or inferred to function in concert. Most prominently these included families involved in the complement system and melanisation responses, both of which occur mainly in the haemolymph. Comparing these with feature-based clustering results identified evolutionary-functional correspondences that were particularly striking amongst families with members known to function together in the coordination and execution of complement system responses. Our results provide quantitative evidence that where and how different genes participate in immune defence responses limit the range of possible evolutionary scenarios that are tolerated by natural selection. Comparative genomics approaches to explore constraints in evolutionary biology therefore offer opportunities to advance the understanding of how functional constraints on different components of biological systems govern their evolutionary trajectories.

## Acknowledgments

This research was supported by Novartis Foundation for medical-biological research grant #18B116 and Swiss National Science Foundation grants PP00P3_170664 and CRSII5_186397.

## Author Contributions

RMW conceived the study. LR, RF, MJMFR, AT, and RMW designed the analyses. LR analysed orthology data, quantified evolutionary features, and performed clustering analyses and statistical testing. RF built whole genome alignments, and analysed alignment and variation data. MR analysed variation data. AT curated data and assisted with gene expression data analysis. LR and RMW wrote the manuscript with input from all authors. All authors read and approved the manuscript.

## Conflict of Interest Statement

All authors declare no competing interests.

## Data Availability

No new genomic data were produced as part of this study and all data analysed herein are available from public databases.

Additional File 1: Distributions of computed OG metrics for all of the immune gene families for each evolutionary feature together with statistical assessments of the significance of deviations from the typical metric values.

Additional File 2: Heatmaps and dendrograms resulting from clustering with family mean metric values with different distance functions and agglomerative clustering methods.

Additional File 3: Visualisation of expression patterns per expression cluster defined by weighted correlation network analysis, with results from Gene Ontology term enrichment tests.

Additional File 4: Plots of evolutionary similarities versus expression similarities based on evolutionary feature metric means and medians for different sets of expression clusters.

## Notes

### Competing Interest Statement

The authors have declared no competing interest.

## References

Aguilar R et al. 2005. Global gene expression analysis of Anopheles gambiae responses to microbial challenge. Insect Biochem. Mol. Biol. 35:709–719. doi: 10.1016/j.ibmb.2005.02.019.

Albà MM, Castresana J. 2005. Inverse Relationship Between Evolutionary Rate and Age of Mammalian Genes. Mol. Biol. Evol. 22:598–606. doi: 10.1093/molbev/msi045.

Alquraini A, El Khoury J. 2020. Scavenger receptors. Curr. Biol. 30:R790–R795. doi: 10.1016/j.cub.2020.05.051.

Antonovics J, van Tienderen PH. 1991. Ontoecogenophyloconstraints? The chaos of constraint terminology. Trends Ecol. Evol. 6:166–168. doi: 10.1016/0169-5347(91)90059-7.

Arensburger P et al. 2010. Sequencing of Culex quinquefasciatus establishes a platform for mosquito comparative genomics. Science. 330:86–88. doi: 10.1126/science.1191864.

Baker DA et al. 2011. A comprehensive gene expression atlas of sex- and tissue-specificity in the malaria vector, Anopheles gambiae. BMC Genomics. 12:296. doi: 10.1186/1471-2164-12-296.

Barribeau SM et al. 2015. A depauperate immune repertoire precedes evolution of sociality in bees. Genome Biol. 16:83. doi: 10.1186/s13059-015-0628-y.

Bartholomay LC et al. 2010. Pathogenomics of Culex quinquefasciatus and meta-analysis of infection responses to diverse pathogens. Science. 330:88–90. doi: 10.1126/science.1193162.

Bartholomay LC, Michel K. 2018. Mosquito immunobiology: the intersection of vector health and vector competence. Annu. Rev. Entomol. 63:145–167. doi: 10.1146/annurev-ento-010715-023530.

Bishnoi R et al. 2019. Solution structure, glycan specificity and of phenol oxidase inhibitory activity of Anopheles C-type lectins CTL4 and CTLMA2. Sci. Rep. 9:15191. doi: 10.1038/s41598-019-51353-z.

Blanchette M et al. 2004. Aligning multiple genomic sequences with the threaded blockset aligner. Genome Res. 14:708– 715. doi: 10.1101/gr.1933104.

Blandin SA, Marois E, Levashina EA. 2008. Antimalarial Responses in Anopheles gambiae: From a Complement-like Protein to a Complement-like Pathway. Cell Host Microbe. 3:364–374. doi: 10.1016/j.chom.2008.05.007.

Blumberg BJ, Trop S, Das S, Dimopoulos G. 2013. Bacteria- and IMD Pathway-Independent Immune Defenses against Plasmodium falciparum in Anopheles gambiae Moreira, LA, editor. PLoS ONE. 8:e72130. doi: 10.1371/journal.pone.0072130.

Brucker RM, Funkhouser LJ, Setia S, Pauly R, Bordenstein SR. 2012. Insect innate immunity database (IIID): an annotation tool for identifying immune genes in insect genomes Promponas, VJ, editor. PLoS ONE. 7:e45125. doi: 10.1371/journal.pone.0045125.

Campbell CL, Black WC, Hess AM, Foy BD. 2008. Comparative genomics of small RNA regulatory pathway components in vector mosquitoes. BMC Genomics. 9:425. doi: 10.1186/1471-2164-9-425.

Chen X-G et al. 2015. Genome sequence of the Asian Tiger mosquito, Aedes albopictus, reveals insights into its biology, genetics, and evolution. Proc. Natl. Acad. Sci. 112:E5907–E5915. doi: 10.1073/pnas.1516410112.

Christophides GK et al. 2002. Immunity-related genes and gene families in Anopheles gambiae. Science. 298:159–165. doi: 10.1126/science.1077136.

Dana AN et al. 2005. Gene expression patterns associated with blood-feeding in the malaria mosquito Anopheles gambiae. BMC Genomics. 6:5. doi: 10.1186/1471-2164-6-5.

Dong Y et al. 2006. Anopheles gambiae Immune Responses to Human and Rodent Plasmodium Parasite Species Sher, A, editor. PLoS Pathog. 2:e52. doi: 10.1371/journal.ppat.0020052.

Dong Y, Dimopoulos G. 2009. Anopheles Fibrinogen-related Proteins Provide Expanded Pattern Recognition Capacity against Bacteria and Malaria Parasites. J. Biol. Chem. 284:9835–9844. doi: 10.1074/jbc.M807084200.

El Moussawi L, Nakhleh J, Kamareddine L, Osta MA. 2019. The mosquito melanization response requires hierarchical activation of non-catalytic clip domain serine protease homologs Dimopoulos, G, editor. PLOS Pathog. 15:e1008194. doi: 10.1371/journal.ppat.1008194.

Estévez-Lao TY, Hillyer JF. 2014. Involvement of the Anopheles gambiae Nimrod gene family in mosquito immune responses. Insect Biochem. Mol. Biol. 44:12–22. doi: 10.1016/j.ibmb.2013.10.008.

Evans JD et al. 2006. Immune pathways and defence mechanisms in honey bees Apis mellifera. Insect Mol. Biol. 15:645– 656. doi: 10.1111/j.1365-2583.2006.00682.x.

Fraiture M et al. 2009. Two Mosquito LRR Proteins Function as Complement Control Factors in the TEP1-Mediated Killing of Plasmodium. Cell Host Microbe. 5:273–284. doi: 10.1016/j.chom.2009.01.005.

Galili T. 2015. dendextend: an R package for visualizing, adjusting and comparing trees of hierarchical clustering. Bioinformatics. 31:3718–3720. doi: 10.1093/bioinformatics/btv428.

Gansner ER, Koren Y, North S. 2005. Graph Drawing by Stress Majorization. In: Graph Drawing. Pach, J, editor. Lecture Notes in Computer Science Vol. 3383 Springer Berlin Heidelberg: Berlin, Heidelberg pp. 239–250. doi: 10.1007/978-3-540-31843-9_25.

Gansner ER, North SC. 2000. An open graph visualization system and its applications to software engineering. Softw. -Pract. Exp. 30:1203–1233.

Gerardo NM et al. 2010. Immunity and other defenses in pea aphids, Acyrthosiphon pisum. Genome Biol. 11:R21. doi: 10.1186/gb-2010-11-2-r21.

Giraldo-Calderón GI et al. 2015. VectorBase: an updated bioinformatics resource for invertebrate vectors and other organisms related with human diseases. Nucleic Acids Res. 43:D707–D713. doi: 10.1093/nar/gku1117.

Glickman MH, Ciechanover A. 2002. The Ubiquitin-Proteasome Proteolytic Pathway: Destruction for the Sake of Construction. Physiol. Rev. 82:373–428. doi: 10.1152/physrev.00027.2001.

Gulley MM, Zhang X, Michel K. 2013. The roles of serpins in mosquito immunology and physiology. J. Insect Physiol. 59:138–147. doi: 10.1016/j.jinsphys.2012.08.015.

Hammond AM et al. 2017. The creation and selection of mutations resistant to a gene drive over multiple generations in the malaria mosquito Barton, NH, editor. PLOS Genet. 13:e1007039. doi: 10.1371/journal.pgen.1007039.

Han MV, Thomas GWC, Lugo-Martinez J, Hahn MW. 2013. Estimating gene gain and loss rates in the presence of error in genome assembly and annotation using CAFE 3. Mol. Biol. Evol. 30:1987–1997. doi: 10.1093/molbev/mst100.

Hillyer JF. 2016. Insect immunology and hematopoiesis. Dev. Comp. Immunol. 58:102–118. doi: 10.1016/j.dci.2015.12.006.

Hoffmann AA. 2013. III.8. Evolutionary Limits and Constraints. In: The Princeton Guide to Evolution. Losos, JBet al., editors. Princeton University Press pp. 247–252. doi: 10.1515/9781400848065-034.

Holm I et al. 2012. Diverged Alleles of the Anopheles gambiae Leucine-Rich Repeat Gene APL1A Display Distinct Protective Profiles against Plasmodium falciparum Michel, K, editor. PLoS ONE. 7:e52684. doi: 10.1371/journal.pone.0052684.

Holt RA et al. 2002. The genome sequence of the malaria mosquito Anopheles gambiae. Science. 298:129–149. doi: 10.1126/science.1076181.

Imler J-L. 2014. Overview of Drosophila immunity: A historical perspective. Dev. Comp. Immunol. 42:3–15. doi: 10.1016/j.dci.2013.08.018.

Jiang X et al. 2014. Genome analysis of a major urban malaria vector mosquito, Anopheles stephensi. Genome Biol. 15:459. doi: 10.1186/s13059-014-0459-2.

Kassambara A. 2017. Practical guide to cluster analysis in R: unsupervised machine learning. Edition 1. STHDA: France.

King JG. 2020. Developmental and comparative perspectives on mosquito immunity. Dev. Comp. Immunol. 103:103458. doi: 10.1016/j.dci.2019.103458.

Koonin EV, Wolf YI. 2010. Constraints and plasticity in genome and molecular-phenome evolution. Nat. Rev. Genet. 11:487–498. doi: 10.1038/nrg2810.

Kounatidis I, Ligoxygakis P. 2012. Drosophila as a model system to unravel the layers of innate immunity to infection. Open Biol. 2:120075. doi: 10.1098/rsob.120075.

Kriventseva EV et al. 2015. OrthoDB v8: update of the hierarchical catalog of orthologs and the underlying free software. Nucleic Acids Res. 43:D250–D256. doi: 10.1093/nar/gku1220.

Krylov DM. 2003. Gene Loss, Protein Sequence Divergence, Gene Dispensability, Expression Level, and Interactivity Are Correlated in Eukaryotic Evolution. Genome Res. 13:2229–2235. doi: 10.1101/gr.1589103.

Langfelder P, Horvath S. 2008. WGCNA: an R package for weighted correlation network analysis. BMC Bioinformatics. 9:559. doi: 10.1186/1471-2105-9-559.

Lawniczak MKN et al. 2010. Widespread divergence between incipient anopheles gambiae species revealed by whole genome sequences. Science. 330:512–514. doi: 10.1126/science.1195755.

Lazzaro BP, Zasloff M, Rolff J. 2020. Antimicrobial peptides: Application informed by evolution. Science. 368:eaau5480. doi: 10.1126/science.aau5480.

Lemaitre B, Hoffmann J. 2007. The host defense of Drosophila melanogaster. Annu. Rev. Immunol. 25:697–743. doi: 10.1146/annurev.immunol.25.022106.141615.

Levashina EA, Baxter RHG. 2018. Complement-Like System in the Mosquito Responses Against Malaria Parasites. In: Complement Activation in Malaria Immunity and Pathogenesis. Stoute, JA, editor. Springer International Publishing: Cham pp. 139–146. doi: 10.1007/978-3-319-77258-5_8.

Lewis SH, Salmela H, Obbard DJ. 2016. Duplication and Diversification of Dipteran Argonaute Genes, and the Evolutionary Divergence of Piwi and Aubergine. Genome Biol. Evol. 8:507–518. doi: 10.1093/gbe/evw018.

Li H et al. 2020. Coordination of humoral immune factors dictates compatibility between Schistosoma mansoni and Biomphalaria glabrata. eLife. 9:e51708. doi: 10.7554/eLife.51708.

Ligoxygakis P. 2017. Insect Immunity. Advances in insect physiology, Volume fifty two. https://search.ebscohost.com/login.aspx?direct=true&scope=site&db=nlebk&db=nlabk&AN=1433143 (Accessed May 13, 2020).

MacCallum RM, Redmond SN, Christophides GK. 2011. An expression map for Anopheles gambiae. BMC Genomics. 12:620. doi: 10.1186/1471-2164-12-620.

Marinotti O et al. 2006. Genome-wide analysis of gene expression in adult Anopheles gambiae. Insect Mol. Biol. 15:1–12. doi: 10.1111/j.1365-2583.2006.00610.x.

Marinotti O et al. 2013. The genome of Anopheles darlingi, the main neotropical malaria vector. Nucleic Acids Res. 41:7387–7400. doi: 10.1093/nar/gkt484.

Markianos K et al. 2016. Genetic structure of a local population of the Anopheles gambiae complex in Burkina Faso Brooke, B, editor. PLOS ONE. 11:e0145308. doi: 10.1371/journal.pone.0145308.

Matthews BJ et al. 2018. Improved reference genome of Aedes aegypti informs arbovirus vector control. Nature. 563:501– 507. doi: 10.1038/s41586-018-0692-z.

Mauro AA, Ghalambor CK. 2020. Trade-offs, Pleiotropy, and Shared Molecular Pathways: A Unified View of Constraints on Adaptation. Integr. Comp. Biol. 60:332–347. doi: 10.1093/icb/icaa056.

Meekins DA, Kanost MR, Michel K. 2017. Serpins in arthropod biology. Semin. Cell Dev. Biol. 62:105–119. doi: 10.1016/j.semcdb.2016.09.001.

Midega J et al. 2013. Discovery and characterization of two Nimrod superfamily members in Anopheles gambiae. Pathog. Glob. Health. 107:463–474. doi: 10.1179/204777213X13867543472674.

Miles A, The Anopheles gambiae 1000 Genomes Consortium, Kwiatkowski D. 2017. Genetic diversity of the African malaria vector Anopheles gambiae. Nature. 552:96–100. doi: 10.1038/nature24995.

Mitri C et al. 2020. Gene copy number and function of the APL1 immune factor changed during Anopheles evolution. Parasit. Vectors. 13:18. doi: 10.1186/s13071-019-3868-y.

Mussabekova A, Daeffler L, Imler J-L. 2017. Innate and intrinsic antiviral immunity in Drosophila. Cell. Mol. Life Sci. 74:2039–2054. doi: 10.1007/s00018-017-2453-9.

Myllymäki H, Rämet M. 2014. JAK/STAT pathway in Drosophila immunity. Scand. J. Immunol. 79:377–385. doi: 10.1111/sji.12170.

Myllymäki H, Valanne S, Rämet M. 2014. The Drosophila Imd signaling pathway. J. Immunol. 192:3455–3462. doi: 10.4049/jimmunol.1303309.

Nakhleh J, Christophides GK, Osta MA. 2017. The serine protease homolog CLIPA14 modulates the intensity of the immune response in the mosquito Anopheles gambiae. J. Biol. Chem. 292:18217–18226. doi: 10.1074/jbc.M117.797787.

Nakhleh J, El Moussawi L, Osta MA. 2017. The Melanization Response in Insect Immunity. In: Advances in Insect Physiology.Vol. 52 Elsevier pp. 83–109. doi: 10.1016/bs.aiip.2016.11.002.

Neafsey DE et al. 2015. Highly evolvable malaria vectors: The genomes of 16 Anopheles mosquitoes. Science. 347:1258522. doi: 10.1126/science.1258522.

Neafsey DE et al. 2010. SNP genotyping defines complex gene-flow boundaries among African malaria vector mosquitoes. Science. 330:514–517. doi: 10.1126/science.1193036.

Neira Oviedo M, VanEkeris L, Corena-Mcleod MDP, Linser PJ. 2008. A microarray-based analysis of transcriptional compartmentalization in the alimentary canal of Anopheles gambiae (Diptera: Culicidae) larvae: Gut transcriptome of larval An.gambiae. Insect Mol. Biol. 17:61–72. doi: 10.1111/j.1365-2583.2008.00779.x.

Nene V et al. 2007. Genome sequence of Aedes aegypti, a major arbovirus vector. Science. 316:1718–1723. doi: 10.1126/science.1138878.

Povelones M et al. 2013. The CLIP-Domain Serine Protease Homolog SPCLIP1 Regulates Complement Recruitment to Microbial Surfaces in the Malaria Mosquito Anopheles gambiae Vernick, KD, editor. PLoS Pathog. 9:e1003623. doi: 10.1371/journal.ppat.1003623.

Povelones M, Osta MA, Christophides GK. 2016. The complement system of malaria vector mosquitoes. In: Advances in Insect Physiology.Vol. 51 Elsevier pp. 223–242. doi: 10.1016/bs.aiip.2016.06.001.

Povelones M, Waterhouse RM, Kafatos FC, Christophides GK. 2009. Leucine-rich repeat protein complex activates mosquito complement in defense against Plasmodium parasites. Science. 324:258–261. doi: 10.1126/science.1171400.

R Core Team. 2021. R: A language and environment for statistical computing. R Foundation for Statistical Computing: Vienna, Austria https://www.R-project.org/.

Ramirez JL, Muturi EJ, Flor-Weiler LB, Vermillion K, Rooney AP. 2020. Peptidoglycan Recognition Proteins (PGRPs) Modulates Mosquito Resistance to Fungal Entomopathogens in a Fungal-Strain Specific Manner. Front. Cell. Infect. Microbiol. 9:465. doi: 10.3389/fcimb.2019.00465.

Rao X-J et al. 2018. Immune functions of insect βGRPs and their potential application. Dev. Comp. Immunol. 83:80–88. doi: 10.1016/j.dci.2017.12.007.

Reyes Ruiz VM et al. 2019. Stimulation of a protease targeting the LRIM1/APL1C complex reveals specificity in complement-like pathway activation in Anopheles gambiae Barillas-Mury, C, editor. PLOS ONE. 14:e0214753. doi: 10.1371/journal.pone.0214753.

Richardson MK, Chipman AD. 2003. Developmental constraints in a comparative framework: A test case using variations in phalanx number during amniote evolution. J. Exp. Zool. 296B:8–22. doi: 10.1002/jez.b.13.

Rolff J, Reynolds S. 2009. Insect Infection and Immunity. Oxford University Press doi: 10.1093/acprof:oso/9780199551354.001.0001.

Rottschaefer SM et al. 2011. Exceptional Diversity, Maintenance of Polymorphism, and Recent Directional Selection on the APL1 Malaria Resistance Genes of Anopheles gambiae Schneider, DS, editor. PLoS Biol. 9:e1000600. doi: 10.1371/journal.pbio.1000600.

Ruzzante L, Reijnders MJMF, Waterhouse RM. 2019. Of genes and genomes: mosquito evolution and diversity. Trends Parasitol. 35:32–51. doi: 10.1016/j.pt.2018.10.003.

Sackton TB. 2019. Comparative genomics and transcriptomics of host–pathogen interactions in insects: evolutionary insights and future directions. Curr. Opin. Insect Sci. 31:106–113. doi: 10.1016/j.cois.2018.12.007.

Sackton TB et al. 2007. Dynamic evolution of the innate immune system in Drosophila. Nat. Genet. 39:1461–1468. doi: 10.1038/ng.2007.60.

Sackton TB, Lazzaro BP, Clark AG. 2017. Rapid expansion of immune-related gene families in the house fly, Musca domestica. Mol. Biol. Evol. msw285. doi: 10.1093/molbev/msw285.

Schnitger AKD, Kafatos FC, Osta MA. 2007. The Melanization Reaction Is Not Required for Survival of Anopheles gambiae Mosquitoes after Bacterial Infections. J. Biol. Chem. 282:21884–21888. doi: 10.1074/jbc.M701635200.

Schnitger AKD, Yassine H, Kafatos FC, Osta MA. 2009. Two C-type Lectins Cooperate to Defend Anopheles gambiae against Gram-negative Bacteria. J. Biol. Chem. 284:17616–17624. doi: 10.1074/jbc.M808298200.

Shakhnovich BE, Koonin EV. 2006. Origins and impact of constraints in evolution of gene families. Genome Res. 16:1529– 1536. doi: 10.1101/gr.5346206.

Shi X-Z, Zhong X, Yu X-Q. 2012. Drosophila melanogaster NPC2 proteins bind bacterial cell wall components and may function in immune signal pathways. Insect Biochem. Mol. Biol. 42:545–556. doi: 10.1016/j.ibmb.2012.04.002.

Siepel A et al. 2005. Evolutionarily conserved elements in vertebrate, insect, worm, and yeast genomes. Genome Res. 15:1034–1050. doi: 10.1101/gr.3715005.

Simões ML et al. 2017. The Anopheles FBN9 immune factor mediates Plasmodium species-specific defense through transgenic fat body expression. Dev. Comp. Immunol. 67:257–265. doi: 10.1016/j.dci.2016.09.012.

Sousa GL, Bishnoi R, Baxter RHG, Povelones M. 2020. The CLIP-domain serine protease CLIPC9 regulates melanization downstream of SPCLIP1, CLIPA8, and CLIPA28 in the malaria vector Anopheles gambiae Hillyer, JF, editor. PLOS Pathog. 16:e1008985. doi: 10.1371/journal.ppat.1008985.

Suzuki R, Shimodaira H. 2006. Pvclust: an R package for assessing the uncertainty in hierarchical clustering. Bioinformatics. 22:1540–1542. doi: 10.1093/bioinformatics/btl117.

Tanaka H et al. 2008. A genome-wide analysis of genes and gene families involved in innate immunity of Bombyx mori. Insect Biochem. Mol. Biol. 38:1087–1110. doi: 10.1016/j.ibmb.2008.09.001.

Valanne S, Wang J-H, Rämet M. 2011. The Drosophila Toll signaling pathway. J. Immunol. 186:649–656. doi: 10.4049/jimmunol.1002302.

Vasta GR. 2020. Galectins in Host–Pathogen Interactions: Structural, Functional and Evolutionary Aspects. In: Lectin in Host Defense Against Microbial Infections. Hsieh, S-L, editor. Advances in Experimental Medicine and Biology Vol. 1204 Springer Singapore: Singapore pp. 169–196. doi: 10.1007/978-981-15-1580-4_7.

Volz J, Muller H-M, Zdanowicz A, Kafatos FC, Osta MA. 2006. A genetic module regulates the melanization response of Anopheles to Plasmodium. Cell. Microbiol. 8:1392–1405. doi: 10.1111/j.1462-5822.2006.00718.x.

Volz J, Osta MA, Kafatos FC, Müller H-M. 2005. The Roles of Two Clip Domain Serine Proteases in Innate Immune Responses of the Malaria Vector Anopheles gambiae. J. Biol. Chem. 280:40161–40168. doi: 10.1074/jbc.M506191200.

Wang Q, Ren M, Liu X, Xia H, Chen K. 2019. Peptidoglycan recognition proteins in insect immunity. Mol. Immunol. 106:69– 76. doi: 10.1016/j.molimm.2018.12.021.

Waterhouse RM. 2015. A maturing understanding of the composition of the insect gene repertoire. Curr. Opin. Insect Sci. 7:15–23. doi: 10.1016/j.cois.2015.01.004.

Waterhouse RM et al. 2007. Evolutionary dynamics of immune-related genes and pathways in disease-vector mosquitoes. Science. 316:1738–1743. doi: 10.1126/science.1139862.

Waterhouse RM, Lazzaro BP, Sackton TB. 2020. Characterization of insect immune systems from genomic data. In: Immunity in Insects. Sandrelli, F & Tettamanti, G, editors. Springer Protocols Handbooks Springer US: New York, NY pp. 3–34. doi: 10.1007/978-1-0716-0259-1_1.

Waterhouse RM, Povelones M, Christophides GK. 2010. Sequence-structure-function relations of the mosquito leucine-rich repeat immune proteins. BMC Genomics. 11:531. doi: 10.1186/1471-2164-11-531.

Waterhouse RM, Tegenfeldt F, Li J, Zdobnov EM, Kriventseva EV. 2013. OrthoDB: A hierarchical catalog of animal, fungal and bacterial orthologs. Nucleic Acids Res. 41:D358–65. doi: 10.1093/nar/gks1116.

Waterhouse RM, Zdobnov EM, Kriventseva EV. 2011. Correlating traits of gene retention, sequence divergence, duplicability and essentiality in vertebrates, arthropods, and fungi. Genome Biol. Evol. 3:75–86. doi: 10.1093/gbe/evq083.

Weetman D, Wilding CS, Steen K, Pinto J, Donnelly MJ. 2012. Gene flow-dependent genomic divergence between Anopheles gambiae M and S forms. Mol. Biol. Evol. 29:279–291. doi: 10.1093/molbev/msr199.

White BJ et al. 2011. Adaptive divergence between incipient species of Anopheles gambiae increases resistance to Plasmodium. Proc. Natl. Acad. Sci. 108:244–249. doi: 10.1073/pnas.1013648108.

Williams M, Summers BJ, Baxter RHG. 2015. Biophysical Analysis of Anopheles gambiae Leucine-Rich Repeat Proteins APL1A1, APL1B and APL1C and Their Interaction with LRIM1 Kobe, B, editor. PLOS ONE. 10:e0118911. doi: 10.1371/journal.pone.0118911.

Wiltshire RM et al. 2018. Reduced-representation sequencing identifies small effective population sizes of Anopheles gambiae in the north-western Lake Victoria basin, Uganda. Malar. J. 17:285. doi: 10.1186/s12936-018-2432-0.

Wolf YI, Carmel L, Koonin EV. 2006. Unifying measures of gene function and evolution. Proc. R. Soc. B Biol. Sci. 273:1507– 1515. doi: 10.1098/rspb.2006.3472.

Wolf YI, Novichkov PS, Karev GP, Koonin EV, Lipman DJ. 2009. The universal distribution of evolutionary rates of genes and distinct characteristics of eukaryotic genes of different apparent ages. Proc. Natl. Acad. Sci. 106:7273–7280. doi: 10.1073/pnas.0901808106.

Worth CL, Gong S, Blundell TL. 2009. Structural and functional constraints in the evolution of protein families. Nat. Rev. Mol. Cell Biol. 10:709–720. doi: 10.1038/nrm2762.

Yang Z. 2007. PAML 4: phylogenetic analysis by maximum likelihood. Mol. Biol. Evol. 24:1586–1591. doi: 10.1093/molbev/msm088.

Yassine H, Kamareddine L, Chamat S, Christophides GK, Osta MA. 2014. A Serine Protease Homolog Negatively Regulates TEP1 Consumption in Systemic Infections of the Malaria Vector Anopheles gambiae. J. Innate Immun. 6:806–818. doi: 10.1159/000363296.

Zdobnov EM et al. 2017. OrthoDB v9.1: cataloging evolutionary and functional annotations for animal, fungal, plant, archaeal, bacterial and viral orthologs. Nucleic Acids Res. 45:D744–D749. doi: 10.1093/nar/gkw1119.

Zhang X et al. 2021. CLIPB10 is a Terminal Protease in the Regulatory Network That Controls Melanization in the African Malaria Mosquito Anopheles gambiae. Front. Cell. Infect. Microbiol. 10:585986. doi: 10.3389/fcimb.2020.585986.

Zhang X, An C, Sprigg K, Michel K. 2016. CLIPB8 is part of the prophenoloxidase activation system in Anopheles gambiae mosquitoes. Insect Biochem. Mol. Biol. 71:106–115. doi: 10.1016/j.ibmb.2016.02.008.

Zou Z et al. 2007. Comparative genomic analysis of the Tribolium immune system. Genome Biol. 8:R177. doi: 10.1186/gb-2007-8-8-r177.

