## Supplementary Materials for "Functional constraints on insect immune system components govern their evolutionary trajectories"

### Contents


**Supplementary Data** 2
- Orthology data 2
- Alignment data 2
  - Table S1. Mosquito genome assemblies used for whole genome alignments. 3
- Variation data 4
- Expression data 4
- Anopheles gambiae immunity gene catalogue 4

**Supplementary Methods** 6
- Orthology-based evolutionary features 6
- Ultrametric tree estimation 6
- Variation-based evolutionary features 6
- Alignment-based evolutionary features 7
- Gene family metrics and comparisons 7
  - Figure S1. Evolutionary feature profiles of mosquito immune gene families (delta-medians). 9
- Additional File 1 description 10
- Clustering of gene family metrics 10
  - Figure S2. Clustering heatmap and dendrograms of immune families and their evolutionary features (means). 12
  - Figure S3. AU bootstrap support distributions with tested pairs of distance-agglomeration methods: Families. 13
  - Figure S4. AU bootstrap support distributions with tested pairs of distance-agglomeration methods: Metrics. 14
- Additional File 2 description 14
  - Figure S5. Gene family co-clustering using metric means with 16 distance-clustering method combinations. 15
  - Figure S6. Gene family co-clustering using metric medians with 16 distance-clustering method combinations. 16
  - Figure S7. Alternative family clustering by top 10 principal components of evolutionary features (medians). 17
- Principal Component Analysis (PCA) of gene family metrics 18
  - Figure S8. Principal Component Analysis of median (left) and mean (right) evolutionary feature metrics 18
- Gene and family co-expression analyses 19
  - Figure S9. Immune family expression similarities based on the VectorBase Expression Map. 20
  - Figure S10. Network of immune family expression similarities based on VectorBase Expression Map supercells. 21
  - Table S2. 24 experimental conditions selected for gene expression clustering with WGCNA. 22
- Additional File 3 description 23
  - Figure S11. WGCNA modules and expression tree. 24
  - Figure S12. Immune family co-expression across WGCNA modules. 25
  - Figure S13. Network of immune family expression similarities based on level 4 WGCNA modules. 26
  - Figure S14. Network of immune family expression similarities based on level 2 WGCNA modules. 27
- Additional File 4 description 27
- List of databases and software used 28
- R packages and functions 28

**Supplementary References** 30

### Supplementary Data

#### Orthology data

Orthologous Groups (OGs) of genes from 43 insects including 21 mosquitoes were delineated as part of the *Anopheles* 16 Genome Project (Neafsey et al. 2015). OrthoDB orthology delineation (Kriventseva et al. 2015) was employed to define orthologous groups of genes descended from each last common ancestor of the species phylogeny. The set of species comprised two culicine mosquitoes: *Aedes aegypti* and *Culex quinquefasciatus*, and 19 anophelines: *An. darlingi*, *An. albimanus*, *An. sinensis*, *An. atroparvus*, *An. farauti*, *An. dirus*, *An. funestus*, *An. minimus*, *An. culicifacies*, *An. maculatus*, *An. stephensi* (SDA-500), *An. stephensi* (INDIAN), *An. epiroticus*, *An. christyi*, *An. melas*, *An. quadriannulatus*, *An. arabiensis*, *An. merus*, and *An. gambiae* (PEST), as well as additional Diptera: *Lutzomyia longipalpis*, *Phlebotomus papatasi*, *Glossina morsitans*, and 12 *Drosophila* - *D. grimshawi*, *D. mojavensis*, *D. virilis*, *D. willistoni*, *D. persimilis*, *D. pseudoobscura*, *D. ananassae*, *D. erecta*, *D. yakuba*, *D. melanogaster*, *D. sechellia*, and *D. simulans*; and representatives of Lepidoptera: *Bombyx mori* and *Danaus plexippus*; Coleoptera: *Tribolium castaneum*; Hymenoptera: *Apis mellifera* and *Linepithema humile*, and Hemipterodea (or Exopterygota): *Pediculus humanus* and *Rhodnius prolixus*. From a total of 605'670 genes (mean 14'085; standard deviation 1'956), 530'519 genes were assigned to OGs with about 4'000 orthologs per species found in near-universal insect OGs and a further 4'000 to 6'000 orthologs in widespread insect OGs. Across the 21 mosquitoes, about 2'000 single-copy orthologs and 2'000-3'000 multi-copy orthologs per species were identified, for further details please see (Neafsey et al. 2015).

#### Alignment data

Genome assemblies of 19 *Anopheles*, two *Aedes*, and one *Culex* mosquitoes (Table S1) were retrieved from VectorBase (Giraldo-Calderón et al. 2015). The multi-species whole genome alignments were generated from these assemblies employing the LastZ–MultiZ approach of the Threaded Blockset Aligner (Blanchette et al. 2004). First, pairwise alignments were computed between all softmasked assemblies and that of *An. gambiae* (AgamP4) with LastZ version 1.04 (Harris 2007), using the lastz_32 binary, the HoxD55 substitution matrix, and the following parameters: "C 0 E 30 H 2000 K 2200 L 4000 M 0 O 400 Q T 1 Y 3400". These pairwise alignments were extended into chains with axtChain followed by chainAntiRepeat, and then into nets using chainNet. The axtChain, chainAntiRepeat, and chainNet binaries were obtained from http://hgdownload.cse.ucsc.edu/admin/exe/linux.x86_64/ on Jan. 23rd, 2019. The resulting files were filtered with single_cov2 from the MultiZ software package (single_cov2 version 11, https://github.com/multiz/multiz). Subsequently, the 21 pairwise alignments were combined in a single multi-species alignment with the ROAST software from MultiZ (ROAST version 3) (Blanchette et al. 2004) using AgamP4 as the reference assembly. ROAST requires a phylogenetic tree to guide the progressive alignment process. This species tree was generated by time-calibration and ultrametrisation of the molecular species phylogeny obtained from trimmed alignments of orthologues (see ***Ultrametric tree estimation***, below). Finally, regions from the AgamP4 assembly that were absent from the Multiple Alignment Format (MAF) file generated by ROAST (i.e. non-alignable regions) were added to the MAF file to build the final AgamP4 multispecies whole genome alignment.

**Table S1. Mosquito genome assemblies used for whole genome alignments.**

| Species strain | Assembly Version | Assembly size (bps) | Number of scaffolds |
|---|---|---|---|
| Anopheles gambiae PEST | AgamP4 | 273,109,044 | 8 |
| Anopheles quadriannulatus SANGWE | AquaS1 | 283,828,998 | 2,823 |
| Anopheles christyi ACHKN1017 | AchrA1 | 172,658,580 | 30,369 |
| Culex quinquefasciatus Johannesburg | CpipJ2 | 579,057,705 | 3,172 |
| Anopheles atroparvus EBRO | AatrE3 | 225,295,225 | 1,320 |
| Anopheles sinensis SINENSIS | AsinS2 | 375,763,635 | 10,448 |
| Anopheles epiroticus Epiroticus2 | AepiE1 | 223,486,714 | 2,673 |
| Anopheles stephensi SDA-500 | AsteS1 | 225,369,006 | 1,110 |
| Anopheles darlingi Coari | AdarC3 | 136,950,925 | 2,221 |
| Anopheles minimus MINIMUS1 | AminM1 | 201,793,324 | 678 |
| Aedes aegypti LVP | AaegL5 | 1,278,732,104 | 2,310 |
| Anopheles melas CM1001059 | AmelC2 | 224,162,116 | 20,229 |
| Anopheles arabiensis Dongola | AaraD1 | 246,567,867 | 1,214 |
| Anopheles coluzzii Mali-NIH | AcolM1 | 224,455,335 | 10,521 |
| Anopheles culicifacies A-37 | AculA1 | 202,998,806 | 16,162 |
| Anopheles merus MAF | AmerM2 | 288,048,996 | 2,027 |
| Anopheles dirus WRAIR2 | AdirW1 | 216,307,690 | 1,266 |
| Anopheles funestus FUMOZ | AfunF1 | 225,223,604 | 1,392 |
| Anopheles farauti FAR1 | AfarF2 | 183,103,254 | 310 |
| Anopheles maculatus maculatus3 | AmacM1 | 141,894,015 | 47,797 |
| Anopheles albimanus STECLA | AalbS2 | 173,339,240 | 201 |
| Aedes albopictus Foshan | AaloF1 | 1,923,476,627 | 154,782 |

**Variation data**

Variation data defining single nucleotide polymorphisms (SNPs) for *An. gambiae* PEST were retrieved from VectorBase using the BioMart data mining tool (Release VB-2019-04). The SNPs derive from eight variation datasets hosted at VectorBase: VBP0000002, sequencing studies of MR4 derived insect colonies; VBP0000003, *An. gambiae* M, S and Bamako populations (Neafsey et al. 2010); VBP0000004, genotyping and permethrin-resistance phenotyping of *An. gambiae* M and S form mosquitoes from Cameroon, Ghana, Uganda, and Guinea-Bissau (Weetman et al. 2012); VBP0000007, adaptive divergence between incipient species of *An. gambiae* (White et al. 2011); VBP0000124, genetic structure of a local population of the *An. gambiae* complex in Burkina Faso (Markianos et al. 2016); VBP0000163, the *Anopheles* 1000 genomes consortium phase 1 AR3.1 (Miles et al. 2017); VBP0000212, *An. gambiae* in the north-western Lake Victoria basin, Uganda (Wiltshire et al. 2018); and VBP0000227, creation and selection of mutations resistant to a gene drive G3 colony (Hammond et al. 2017). The VectorBase BioMart data mining tool (Giraldo-Calderón et al. 2015) was used to retrieve all synonymous and non-synonymous SNPs in coding regions for all *An. gambiae* genes (AgamP4.11).

**Expression data**

Expression data for *An. gambiae* genes were retrieved from VectorBase (Expression Stats VB-2019-06) as log2 transformed expression values for 13'201 genes across 291 conditions (mean, variance and number of replicates). These data are compiled by VectorBase as a repository for published or publication-quality RNA-seq and microarray experiments where they are processed through a standard normalisation and analysis pipeline that enables side-by-side comparison of results from a variety of experimental designs. These data form the basis of the VectorBase Expression Browser (Giraldo-Calderón et al. 2015) and Expression Maps (MacCallum et al. 2011). The downloaded expression statistics for a subset of the conditions were used to build co-expression modules in order to assess immune gene family expression similarities. The maps provide an overview of gene expression patterns by using the normalised gene expression data from different experimental conditions to construct clusters of genes with similar expression profiles. With Pearson correlation coefficient-based distance measurements to define gene expression profile similarities, clustering is achieved using a self-organising map algorithm. The map is set up to form a 25 by 20 grid resulting in a total of 500 possible clusters/cells, which facilitates data visualisation. Clustering adds genes to cells in the grid such that those with the most similar expression profiles will become members of the same cell, and neighbouring cells will usually show similar expression patterns. Gene membership for all cells was retrieved from the AgamP4.11 VB-2019-02 map (comprising 12'672 genes and based on 202 conditions) and used to perform immune gene family co-expression analysis.

***Anopheles gambiae* immunity gene catalogue**

The catalogue of *An. gambiae* immune-related genes was built by combining and updating the results of several previous comparative immunogenomics studies (Christophides et al. 2002; Waterhouse et al. 2007; Waterhouse 2009; Kafatos et al. 2009; Bartholomay et al. 2010; Neafsey et al. 2015). The protocols for building such a catalogue are detailed in (Waterhouse et al. 2020). The catalogue includes members of the canonical immune signalling pathways, Imd, JAK/STAT, and Toll, and their positive or

negative regulators. The examined gene families include antimicrobial peptides (AMPs, attacins, cecropins, defensins, gambicins etc.), autophagy-related proteins (APHAGs), caspases (CASPs) and their inhibitors (CASPAs), catalases (CATs), CLIP-domain serine proteases (CLIPs, families A-E), C-type lectins (CTLs), fibrinogen-related proteins (FREPs), galectins (GALEs), gram-negative binding proteins (GNBPs), inhibitors of apoptosis (IAPs), leucine-rich repeat immune proteins (LRIMs), lysozymes (LYSs), MD-2-like proteins (MLs), nimrod-related proteins (NIMRODs), peptidoglycan recognition proteins (PGRPs), prophenoloxidases (PPOs), peroxidases (haem HPXs, glutathione GPXs, and thioredoxin TPXs), scavenger receptors (SCRs, A, B, and C-types), superoxide dismutases (SODs), spaetzle-like proteins (SPZs), serine protease inhibitors (SRPNs), small regulatory RNA pathway members (SRRPs), thioester-containing proteins (TEPs), and toll-like receptors (TOLLs). For informative illustrations see the following articles:
https://doi.org/10.3390/v6114479 (Sim et al. 2014)
https://doi.org/10.1016/bs.aiip.2016.06.001 (Povelones et al. 2016)
https://doi.org/10.1038/srep43847 (Terradas et al. 2017)
https://doi.org/10.1016/B978-0-12-805350-8.00006-4 (Rodgers et al. 2017)
https://doi.org/10.1016/j.pt.2018.04.011 (Simões et al. 2018)

### Supplementary Methods

#### Orthology-based evolutionary features

Orthology data were used to compute a suite of 13 metrics for each orthologous group (OG) that capture different features of the evolutionary histories of all *An. gambiae* genes with identifiable orthologues in other species. These OG-based metrics were defined as follows, (*i*) species composition metrics: *evolutionary age* (**AGE**), the branch length (in million years) to the last common ancestor of all species in the OG, based on the full ultrametric species phylogeny (see below for phylogeny methods). *Universality* (**UNI**), OG species coverage across the full set of species, i.e. the proportion of species with at least one orthologue. (*ii*) Orthologue copy-number metrics: *duplicability* (**DUP**), the proportion of species exhibiting gene duplications within the OG, i.e. the number of species with more than one orthologue divided by the number of species with at least one orthologue. *Average-copy-number* (**ACN**), the mean number of orthologues per species within the OG, i.e. the total number of orthologues in the OG divided by the number of species with at least one orthologue. *Copy-number-variation* (**CNV**), the variability in the number of orthologues within the OG, i.e. the standard deviation of the number of orthologues per species in the OG divided by the mean (the ACN). (*iii*) PAML-based (Yang 2007) selection metrics (as described in (Neafsey et al. 2015)): using the M0 model on the alignments of OG member sequences to compute the number of *synonymous substitutions per synonymous site* (**PDS**), the number of *non-synonymous substitutions per non-synonymous site* (**PDN**), and the ratio of these two values, i.e. the *dN/dS* ratio (**SEL**). (iv) CAFE-based (Han et al. 2013) gene turnover metrics: the number of nodes on the species tree with *expansions* (**EXP**), *contractions* (**CON**), or that remain *stable* (**STA**), as computed by CAFE (v4.2.1; λ=0.00223, p-value=0.05, random seed=1000) using the ultrametric species tree and the full dataset of counts of orthologues per species. (*v*) Genomic organisation metric: orthology data combined with genomic location data were used to quantify synteny conservation (**SYN**) as the proportion of orthologues that maintain their orthologous neighbours in the genomes of the other species. (*vi*) Orthologue sequence conservation/divergence metric: *evolutionary-rate* (**EVR**), the relative level of protein sequence divergence amongst orthologues in the OG, i.e. the pairwise orthologue protein sequence identity normalized by the mean inter-species identity (computed by the OrthoDB orthology delineation pipeline, (Waterhouse et al. 2013)).

#### Ultrametric tree estimation

The Insecta 43-species molecular phylogeny was retrieved from the OrthoDBmoz2 project and produced from trimmed alignments of orthologues as detailed in (Neafsey et al. 2015). The tree was rooted using newick utilities (Junier & Zdobnov 2010). Time-calibration (358 million years for Insecta divergence time estimate obtained from the Time Tree of Life database, (Kumar et al. 2017)) and ultrametrisation was performed using the *ape* (Paradis & Schliep 2019) and *phytools* (Revell 2012) R packages.

#### Variation-based evolutionary features

Population-level genomic variation data were used to compute three metrics for each *An. gambiae* gene, using only annotated coding exons (i.e. excluding untranslated regions and introns) and the single longest protein-coding transcript per gene. SNPs were defined from the 'short' variation dataset filtering on non-synonymous variants and synonymous variants to create two subsets from which to count total non-synonymous and synonymous SNPs per gene. The metrics were defined as follows using: the *proportion of non-synonymous SNPs* (**NSP**), i.e. the number of non-synonymous SNPs divided by the total number of SNPs, the *non-synonymous SNP density* (**NSD**), i.e. the number of non-synonymous SNPs divided by the coding length of the gene, and the *synonymous SNP density* (**SSD**), i.e. the number of synonymous SNPs divided by the coding length of the gene.

**Alignment-based evolutionary features**

Whole-genome alignment data were used to compute two metrics for each *An. gambiae* gene, using only annotated coding exons (*i.e.* excluding untranslated regions and introns) from the union of all protein-coding transcripts per gene. The conservation and constraint metrics were computed for each genomic position and the average was then computed for each gene. They were defined as follows: *whole genome alignability* (**WGA**), *i.e.* the number of species in the alignment at the given genomic position, computed directly from the whole-genome alignment file excluding positions missing from the reference assembly (*i.e.* positions marked as "-" for the reference species). Non-reference assemblies were considered aligned at a genomic position if they were aligned without a gap at this position and the nucleotide was not N (*i.e.* the character in the alignment for this species at this position was either A, T, G, or C). A constraint score estimated with phastCons (Siepel et al. 2005) at the given genomic position, *phastCons constraint* (**PHC**). An initial tree model was computed from the entire whole-genome alignment file using phyloFit with the parameter --*tree*. This model was used to initialize the computation of tree models for conserved and nonconserved sites by running phastCons on a subset of the alignment (chromosome 2R) using the parameters --*target-coverage 0.3*, --*expected-length 35*, --*estimate-trees*, and --*no-post-probs*. These conserved and nonconserved models were used to compute a conservation score per nucleotide by running phastCons on the entire whole-genome alignment file using the parameters --*target-coverage 0.3* and --*expected-length 35*. The average score for each gene in AgamP4.11 annotation was computed from the per-nucleotide values for the **WGA** and **PHC** metrics.

**Gene family metrics and comparisons**

As outlined above, most of the evolutionary feature metrics are defined and computed per orthologous group (OG), while others are defined at nucleotide level and therefore are computed as averages over the protein-coding lengths of each *An. gambiae* gene. For OGs with more than one *An. gambiae* gene, these values are then averaged to obtain a mean metric value for the OG. Thus each of the subfamilies (e.g. cecropins, defensins, attacins), families (e.g. antimicrobial peptides), and broader categories (e.g. antimicrobial effectors), defined in the *An. gambiae* immunity gene catalogue consists of one or more OG, each with metric values for all of the defined evolutionary features. Family-level values for all metrics were then computed by averaging values over all OGs containing genes belonging to each catalogued immune gene family. For graphical visualisation, these family-level means for each metric were compared to the means of all other OGs that contain at least one *An. gambiae* immune gene to quantify the extent to which the metrics of the OGs of a given immune gene family differ from all other immune gene containing OGs, i.e. delta-mean ($\Delta\bar{x}$). To highlight how relatively far $\Delta\bar{x}$ values deviate from the mean for each family, the $\Delta\bar{x}$ values were scaled by the absolute maximum $\Delta\bar{x}$ per metric, i.e. for each metric the distribution was transformed by dividing all values by the absolute maximum $\Delta\bar{x}$. Values for visualisation therefore range from -1 to 1 with the minimum of -1 for metrics where the largest deviation is below the mean and the maximum of 1 for metrics where the largest deviation is above the mean. These were plotted with the colour-blind safe RdYlBu palette from the *RColorBrewer* package in R. Family-level values for all metrics were also compared and visualised using median family metrics, i.e. delta-median $\Delta\tilde{x}$ values (Figure S1). Finally, the significance of the difference of the distribution of each family's OGs metric values (no scaling) compared to all other immune-related OGs was assessed for all metrics and each family using the Wilcoxon rank-sum (Mann–Whitney U) test implemented in the *wilcox.test* function (default two-sided test) from the *stats* package in R. Several families comprise only a few OGs, rendering the Wilcoxon test inappropriate, thus a permutation test implemented in R was also used to test the significance of the difference of the distribution of each family's OGs metric values. The observed difference (absolute $\Delta\bar{x}$ or $\Delta\tilde{x}$) for each family was compared to absolute differences computed for 10'000 permutations of all OG metric values randomly assigned to size-matched sets. The p-value was computed as the number of permutation differences that were greater than the observed difference, divided by the total number of permutations, i.e. an empirical estimate of the probability of obtaining a $\Delta\bar{x}$ or $\Delta\tilde{x}$ greater than the observed $\Delta\bar{x}$ or $\Delta\tilde{x}$ by chance. The distributions of computed OG metrics for all of the immune gene families for each evolutionary feature together with results of both significance tests are presented in Additional File 1.

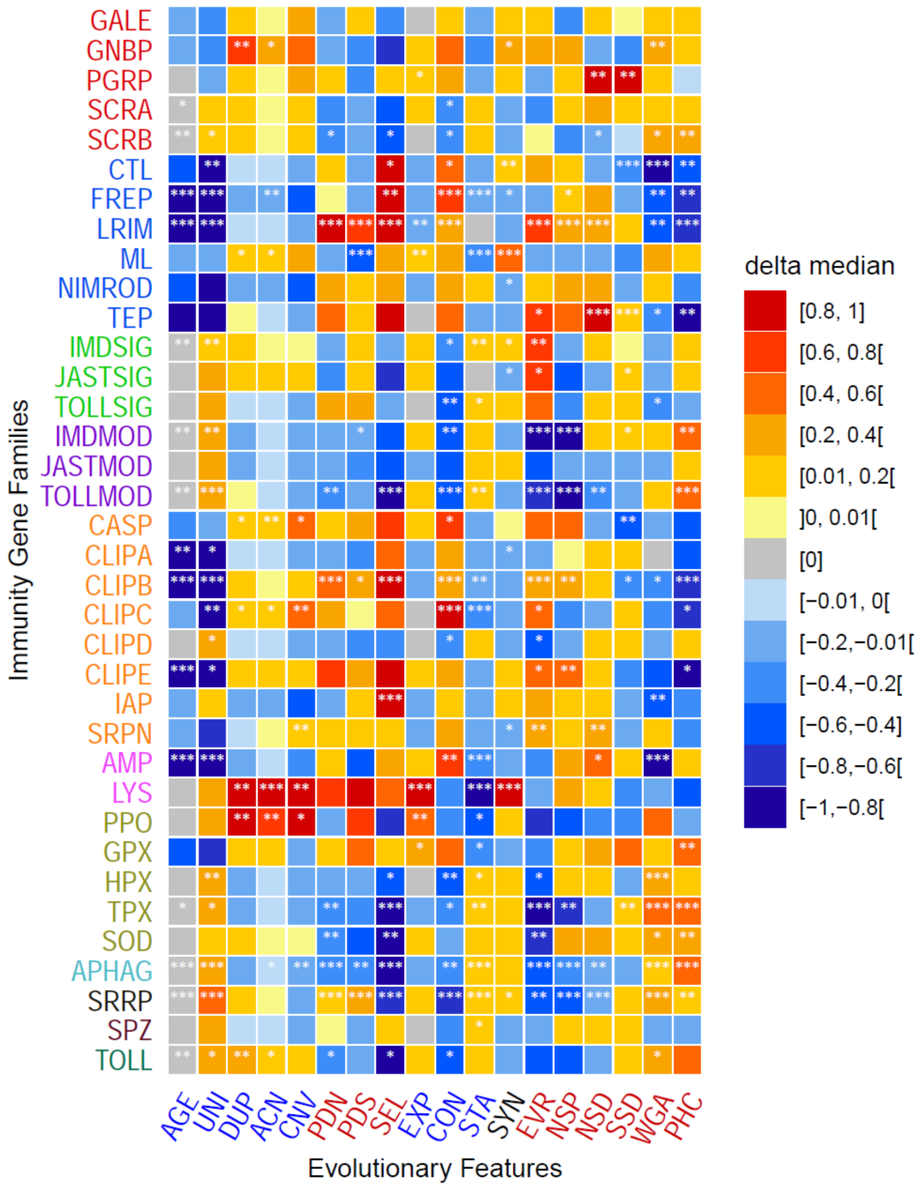


**Figure S1. Evolutionary feature profiles of mosquito immune gene families (delta-medians).**

Evolutionary profiling highlights similar and contrasting patterns across all 36 immune gene families or subfamilies (rows). Deviations from the typical metric values for the suite of 18 evolutionary feature metrics (columns) are computed as the difference between the family median and the average over all orthologous groups from other immune-related gene families ($\Delta\tilde{x}$). For visualisation, values of $\Delta\tilde{x}$ are scaled by the absolute maximum $\Delta\tilde{x}$ per metric, i.e. for each metric the distribution is transformed by dividing all values by the absolute maximum $\Delta\tilde{x}$. Values therefore range from a minimum of -1 for metrics where the largest deviation is below the mean, i.e. lower than other families, and the maximum of 1 for metrics where the largest deviation is above the median, i.e. higher than other families. The significance of the difference of the distribution of metric values (no scaling) for each family compared to all other families was assessed using the Wilcoxon rank-sum (Mann–Whitney U) test and a permutation test (asterisks correspond to the lower p-value from these two tests; *** $p\leq0.01$, ** $p\leq0.05$, * $p\leq0.1$). Family acronyms are defined in Table 1: classic recognition in red; other recognition in blue; core signalling in lime green; signal modulation in purple; cascade modulation in orange; antimicrobial effectors in pink; effector enzymes in olive green. See text for definitions of evolutionary feature acronyms: taxonomic spread and copy-number features in blue; sequence-based features in red.

**Additional File 1 description**

Distributions of computed OG metrics for all of the immune gene families for each evolutionary feature together with statistical assessments of the significance of deviations from the typical metric values. Per family data, coloured by superfamily/class, for each of the 18 evolutionary feature metrics: OGs, number of orthologous groups; Genes, number of genes; MW p-val, Mann Whitney U test p-value; PRM p-val, permutation test p-value.

**Clustering of gene family metrics**

Hierarchical clustering of the evolutionary feature (n=18) metric mean and median values per family (n=36) was performed to assess and quantify the similarities of the evolutionary feature profiles amongst all immune families. First, metric values were scaled with the *scale* function from the *base* package in R, which centers by subtracting the metric means and then scales by dividing the (centered) values by their standard deviations. Second, distance matrices were computed from the family means and medians using the *dist* and *cor* functions from the *stats* package in R, with distances defined as *1-correlation* for matrices computed with *cor*. Several correlation-based distances were examined and compared with Euclidean distance because the aim of clustering was to quantify similarities between evolutionary feature directions rather than magnitudes. Third, hierarchical clustering was performed using R's *hclust* function from the *stats* package and bootstrap support of the resulting hierarchies (trees) was obtained using R's *pvclust* package with 10'000 bootstrap replicates and a fixed seed of 1234, which calculates p-values for hierarchical clustering via multiscale bootstrap resampling (Suzuki & Shimodaira 2006). The bootstrap analyses enabled a comprehensive survey of the performance of combinations of different distance methods and clustering agglomeration methods (Figure S3, Figure S4). Resulting approximately unbiased (AU) p-values from *pvclust* for the 35 nodes of the gene family tree were compared for all pairs of distance metrics (Pearson's correlation, Spearman's correlation, Kendall's correlation, and Euclidean distance) and clustering agglomeration methods (Single, Complete, Average, and Median linkage). While some combinations resulted in overall lower bootstrap scores, no particular combination produced clearly better clustering stability. Using the Pearson's correlation distance method generally produced consistently well-supported clustering, and combined with the Average linkage agglomeration method resulted in only two of the 35 nodes (each for metric means and medians) with less than 75% AU support. The same assessments were applied to hierarchical clustering of the evolutionary features, where the Pearson+Average combination also performed well, with only two (metric means) and three (metric medians) of the 17 nodes receiving less than 75% AU support. The Pearson+Average combination was therefore selected for the main results shown in Figure 2 and Figure S2, with the heatmaps and dendrograms produced and combined with *dendextend* (Galili 2015) and *gplots* packages in R. The contributions of each evolutionary feature were assessed using Principal Component Analysis (PCA) of the family by feature matrices of both median and mean metrics with the *prcomp* function from the *stats* package in R. Results employing other distance-method combinations are presented in Additional File 2. To quantify the robustness of gene family clustering across all 16 tested distance-method combinations, all subtrees from each set of 10'000 *pvclust* bootstrap replicates were enumerated using the R package *ape* (Paradis & Schliep 2019). Because *pvclust*'s multiscale bootstrap resampling algorithm produces 10 sets with different sample sizes

(Suzuki & Shimodaira 2006), only the 10'000 bootstrap replicates resulting from set #6 and corresponding to the exact sample size of the original data were selected. The co-occurrence of all pairs of families was enumerated using all subtrees with a maximum of a quarter of all 36 families present, i.e. the number of times two families are clustered together within the same subtree with a maximum size of nine families. The normalised co-occurrence score (measure of evolutionary profile similarity) was then computed as follows: ( 2 x co-occurrence of Family 1 & Family 2 ) / ( co-occurrence of Family 1 with any Family + co-occurrence of Family 2 with any Family). Thus a normalised score of zero means that the two families never appear in the same quarter of the tree and a score of one means they occur as sister lineages in all bootstrap samples from all distance-method combinations. Pairwise scores, as well as the 30 pairs with the highest overall scores are shown for metric means in Figure S5 and metric medians in Figure S6. Complementary clustering of gene families was also performed using the top 10 Principal Components (PCs) from the metric median PCA results (capturing 97.15% of the variance) rather than the metric values themselves. This employed default squared dissimilarities of the Euclidean distances, with Ward's (ward.D2) agglomerative clustering algorithm (Murtagh & Legendre 2014) because it is based on multidimensional variance like PCA. As above, hierarchical clustering was performed using R's *hclust* function from the *stats* package and bootstrap support of the resulting hierarchies (trees) was obtained using R's *pvclust* package with 10'000 bootstrap replicates and a fixed seed of 1234 (Figure S7).

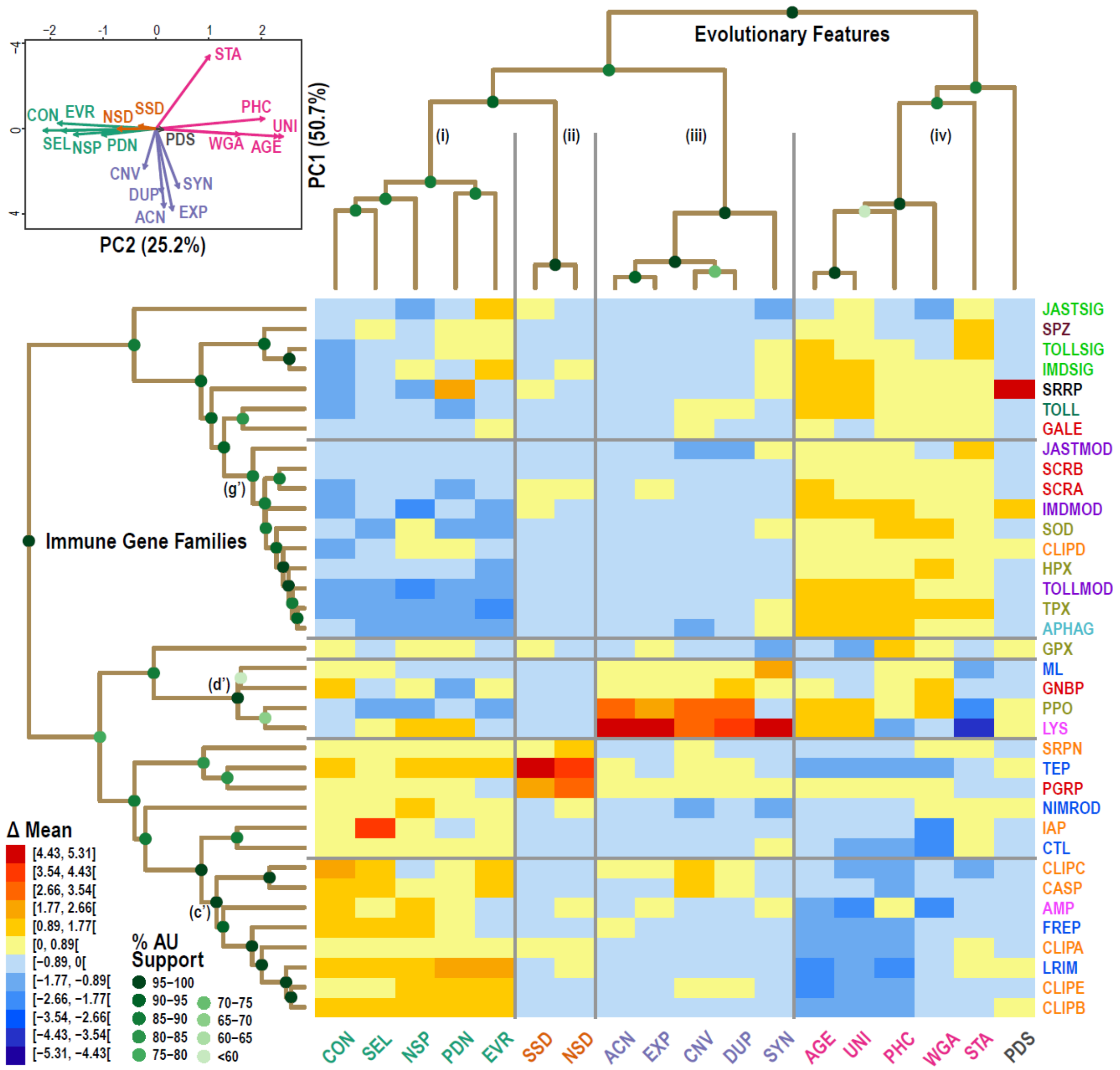


**Figure S2. Clustering heatmap and dendrograms of immune families and their evolutionary features (means).**

Groupings of families and subsets of features delineated by hierarchical clustering using the matrix of evolutionary feature profiles of all immune gene families. Hierarchical clustering results are visualised for the immune families (n=36) and evolutionary features (n=18) using scaled mean metrics with a Pearson's correlation-based distance matrix and average linkage agglomerative clustering. The heatmap displays the relative values of the scaled metrics from low in blue to high in red. The dendrograms show the quantified distances (similarities) between each of the families, and between each of the features, and their groupings, determined by the clustering algorithm and distance method. Support for each node of the two dendrograms is shown with green-filled circles, using multiscale bootstrap resampling to estimate approximately unbiased (AU) support values. Principal Component Analysis (PCA) supports three major groupings of the four subsets of evolutionary features with PC1 and PC2 capturing 75.9% of the variance. Family acronyms are defined in Table 1 and are coloured according to categories defined based on their putative roles in the principal immune phases: classical recognition (red), other recognition (blue), pathway signalling (bright green), pathway modulation (purple), cascade modulation (orange), antimicrobial effectors (pink), effector enzymes (olive green), autophagy (dark cyan), RNAi (black), cytokines (brown), and toll receptors (dark green). See text for definitions of evolutionary feature acronyms, coloured according to groupings in the dendrogram and PCA. Clustering heatmap and dendrograms of immune families and their evolutionary features using median metrics are presented in Figure 2.

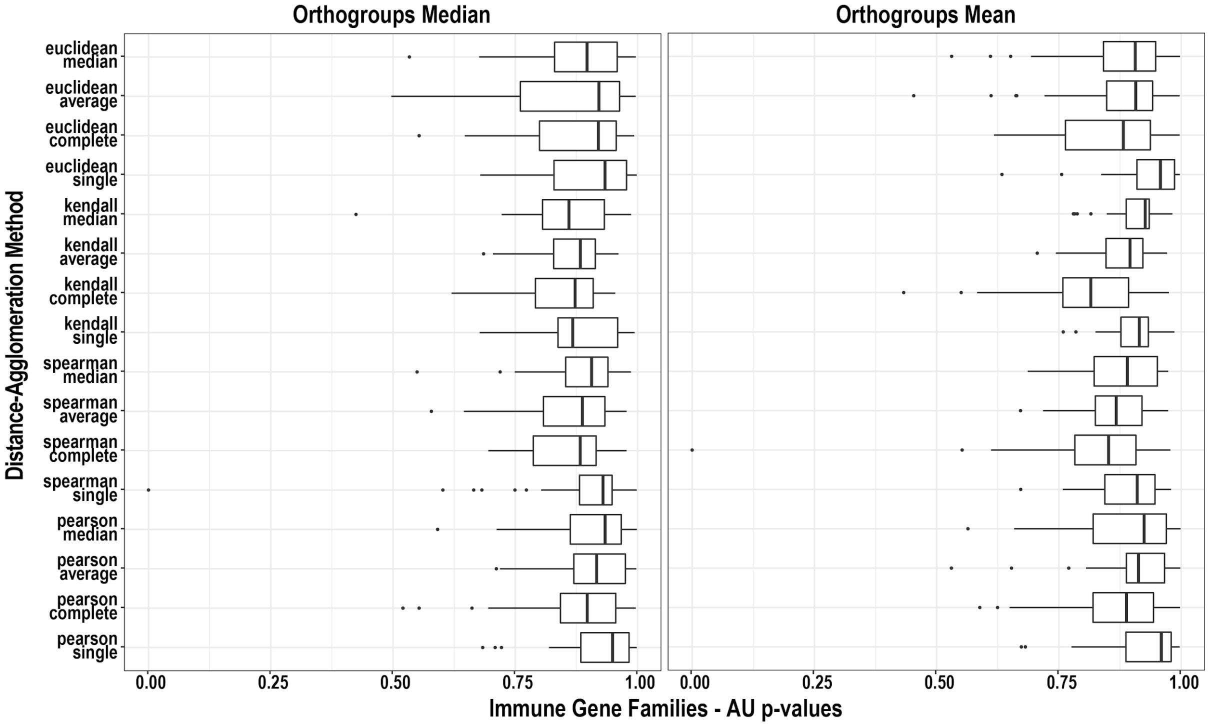


**Figure S3. AU bootstrap support distributions with tested pairs of distance-agglomeration methods: Families.**

Boxplots show the distribution of bootstrap support values (n=35, i.e. for each node in the computed families tree) for each combination of distance method and clustering agglomeration method (median left, mean right). Bootstrap support values, in the form of approximately unbiased (AU) p-values, were computed using the *pvclust* function in R. Distance methods: spearman, Spearman's correlation; kendall, Kendall's correlation; pearson, Pearson's correlation; uncentered, uncentered Pearson's correlation; abscor, absolute Pearson's correlation; and euclidean, Euclidean distance. Clustering agglomeration methods: single, complete, median, and centroid linkage.

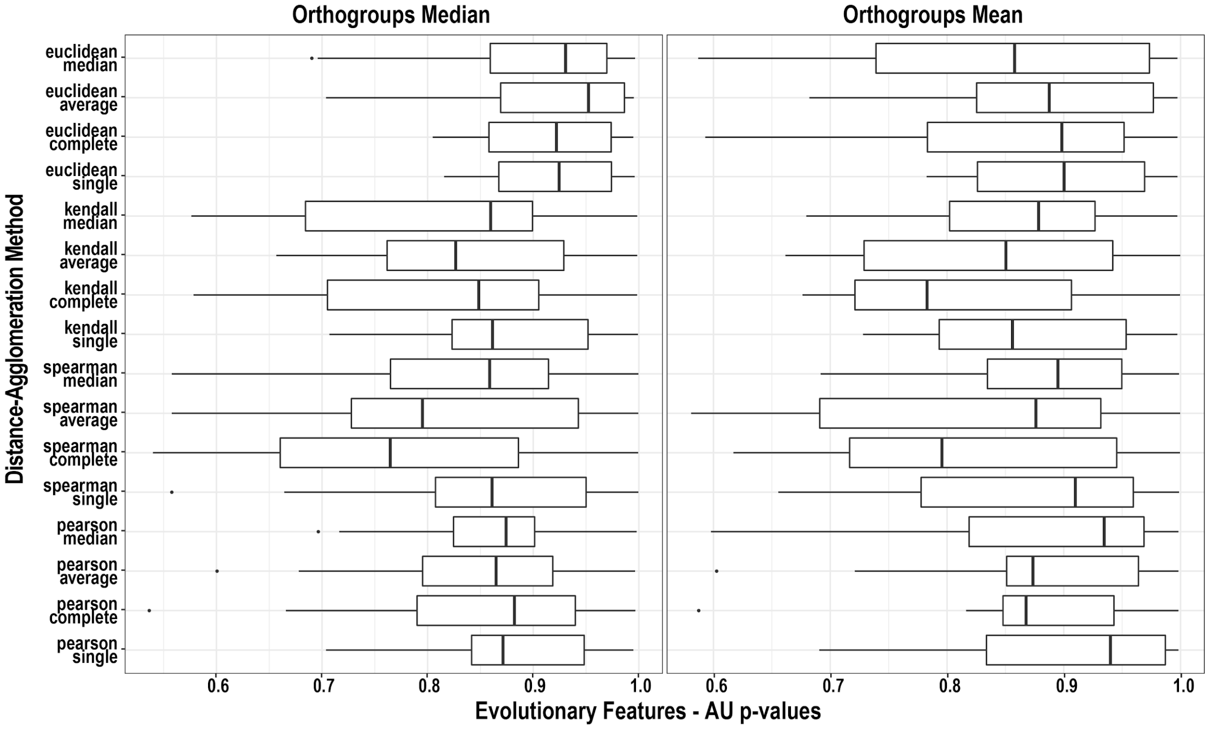

**Figure S4. AU bootstrap support distributions with tested pairs of distance-agglomeration methods: Metrics.**

Boxplots show the distribution of bootstrap support values (n=17, i.e. for each node in the computed metrics tree) for each combination of distance method and clustering agglomeration method (median left, mean right). See Figure S3 legend for details.

### Additional File 2 description

Heatmaps and dendrograms resulting from clustering with family mean metric values with different distance functions (Pearson's, Spearman's, and Kendall's correlation, and Euclidean distances) and agglomerative clustering methods (Average, Single, and Complete linkage) confirms support for many of the observed hierarchical similarities presented in Figures 2 and S2. Hierarchical clustering results are visualised for the immune families (n=36) and evolutionary features (n=18) using scaled median and mean metrics. The heatmaps display the relative values of the scaled metrics from low in blue to high in red. The dendrograms show the quantified distances (similarities) between each of the families, and between each of the features, and their groupings, determined by the clustering algorithm and distance method. Support for each node of the dendrograms is shown with green-filled circles, using multiscale bootstrap resampling to estimate approximately unbiased (AU) support values.

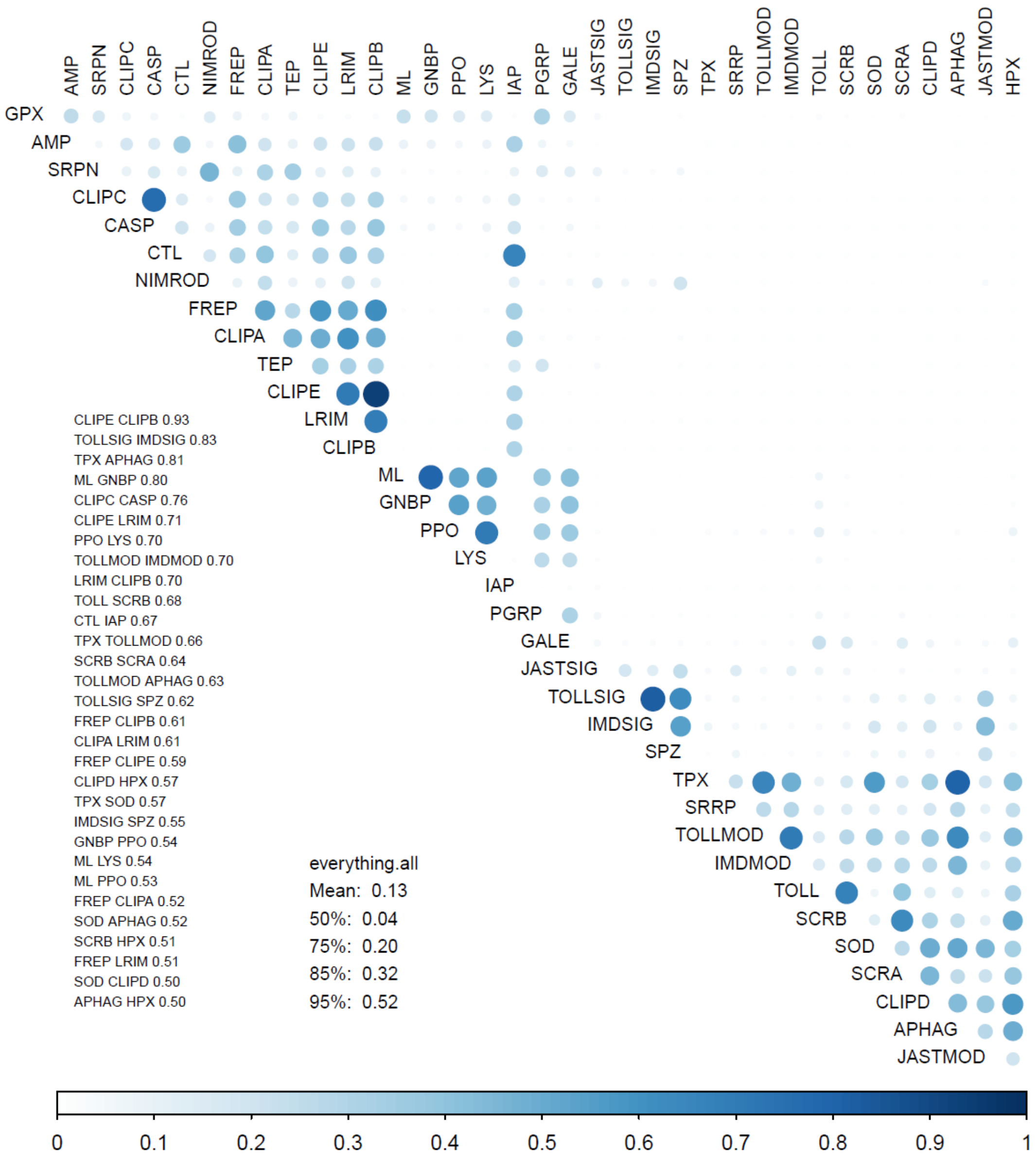


**Figure S5. Gene family co-clustering using metric means with 16 distance-clustering method combinations.**

The normalised co-occurrence score for each pair of families is shown above the diagonal with larger darker blue circles achieving the highest scores. A sorted list of the 30 highest-scoring family pairs with their scores is shown below the diagonal with the mean score, and scores for the 50th, 75th, 85th, and 95th percentiles for all pairs across all distance-clustering methods (everything.all). The order of families follows the order shown in Figure 2.

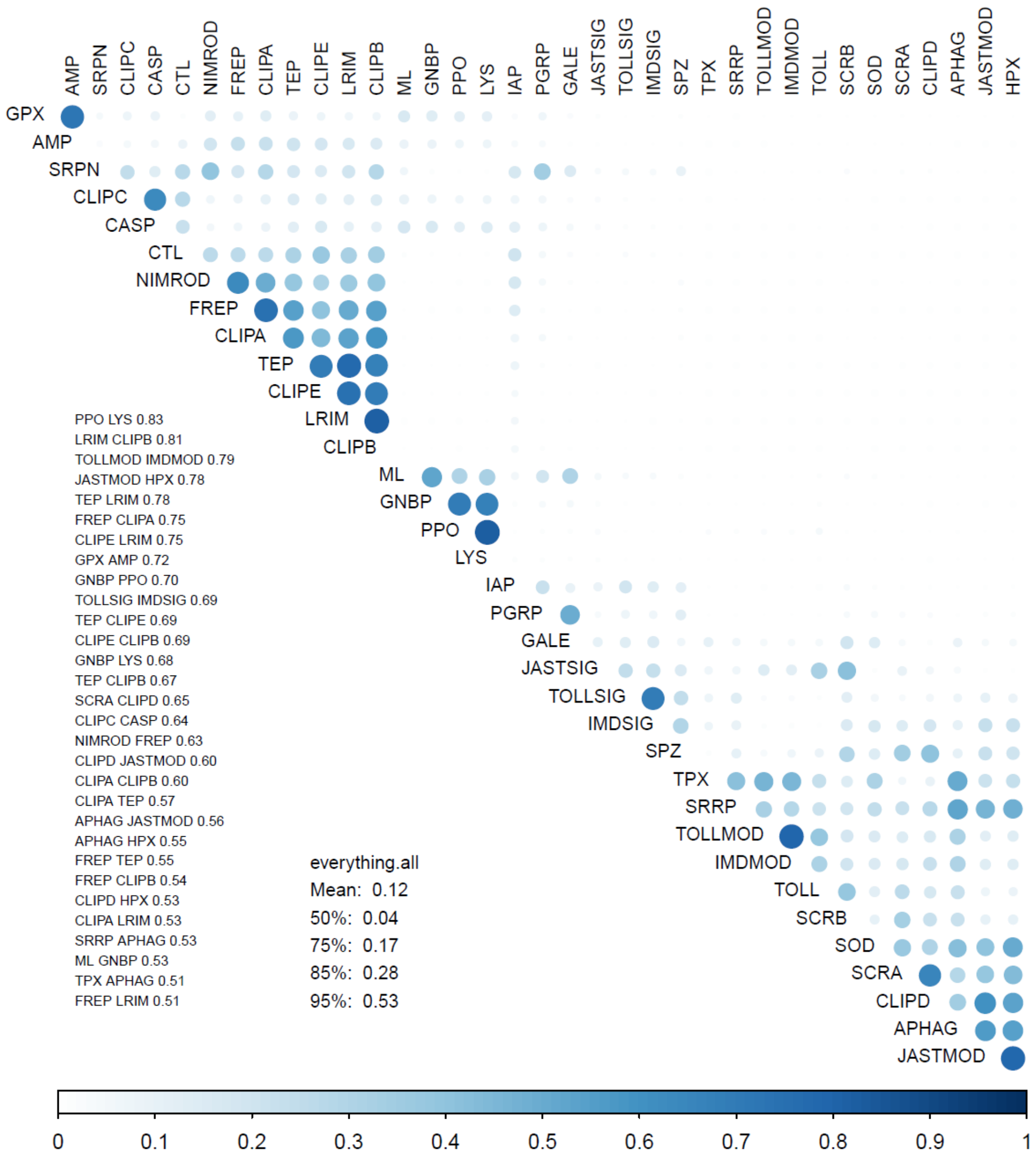


**Figure S6. Gene family co-clustering using metric medians with 16 distance-clustering method combinations.**

The normalised co-occurrence score for each pair of families is shown above the diagonal with larger darker blue circles achieving the highest scores. A sorted list of the 30 highest-scoring family pairs with their scores is shown below the diagonal with the mean score, and scores for the 50$^{th}$, 75$^{th}$, 85$^{th}$, and 95$^{th}$ percentiles for all pairs across all distance-clustering methods (everything.all). The order of families follows the order shown in Figure 2.

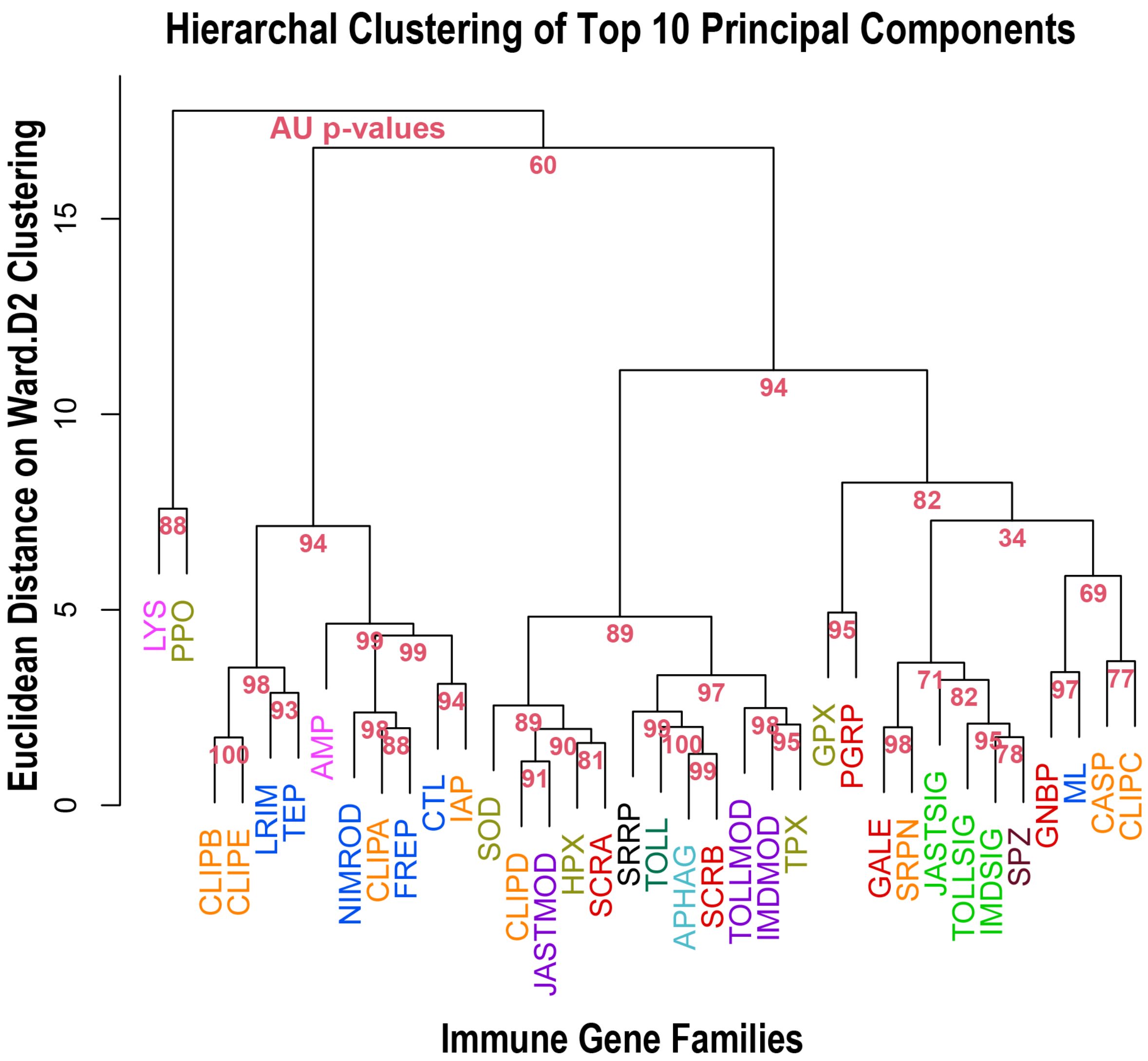


**Figure S7. Alternative family clustering by top 10 principal components of evolutionary features (medians).**

Groupings of immune gene families delineated by the alternative approach of hierarchical clustering using the top 10 principal components (PCs) from principal component analysis (PCA) of the matrix of evolutionary feature medians. Hierarchical clustering results are visualised for the immune families (n=36) using the squared dissimilarities of the Euclidean distance matrix of the scaled median metrics, with Ward's (ward.D2) agglomerative clustering algorithm. The proportions of explained variance were computed from the PCA standard deviations and resulted in: PC1: 0.35951; PC2: 0.30193; PC2: 0.08748, PC4: 0.05729, PC5: 0.04782, PC6: 0.04220, PC7: 0.02895, PC8: 0.02013, PC9: 0.01384, PC10: 0.01231; thus capturing 97.15% of the variance. The dendrogram shows the quantified distances (similarities) between each of the families and their groupings, determined by the clustering algorithm and distance method. Support for each node of the dendrogram is shown red, using multiscale bootstrap resampling to estimate approximately unbiased (AU) support values. Family acronyms are defined in Table 1 and are coloured according to categories defined based on their putative roles in the principal immune phases: classical recognition (red), other recognition (blue), pathway signalling (bright green), pathway modulation (purple), cascade modulation (orange), antimicrobial effectors (pink), effector enzymes (olive green), autophagy (dark cyan), RNAi (black), cytokines (brown), and toll receptors (dark green).

**Principal Component Analysis (PCA) of gene family metrics**

To assess complementary support for the sets of evolutionary features that group together in the hierarchical clustering results, the family by feature matrices of both median and mean metrics were evaluated using the *prcomp* function in R. Metrics were centered and scaled in the same manner as for the hierarchical clustering analyses. The first two principal components together capture 78.2% (medians) and 75.9% (means). For both metric summary statistics, PCA identifies distinct groupings of evolutionary features (Figure S8) that agree with those found with hierarchical clustering: (i) CON, EVR, NSP, PDN, SEL; (ii) SSD and NSD (weak); (iii) ACN, CNV, EXP, DUP, SYN; and (iv) AGE, UNI, WGA, PHC, and somewhat STA. Additionally, the evolutionary feature that did not robustly group with others, PDS, contributes least to PC1 or PC2. Set (iii) features are generally near-orthogonal to features in sets (i), (ii) and (iv), with the exception of STA. Set (iii) features correlate with PC1 and contribute to about half of the variance in the data, while features of sets (i), (ii) and (iv) correlate with PC2 and contribute to about a quarter of the variance in the data. The PCA results therefore demonstrate the extent of co-variance among the different evolutionary features and support their grouping into four major sets of related features.

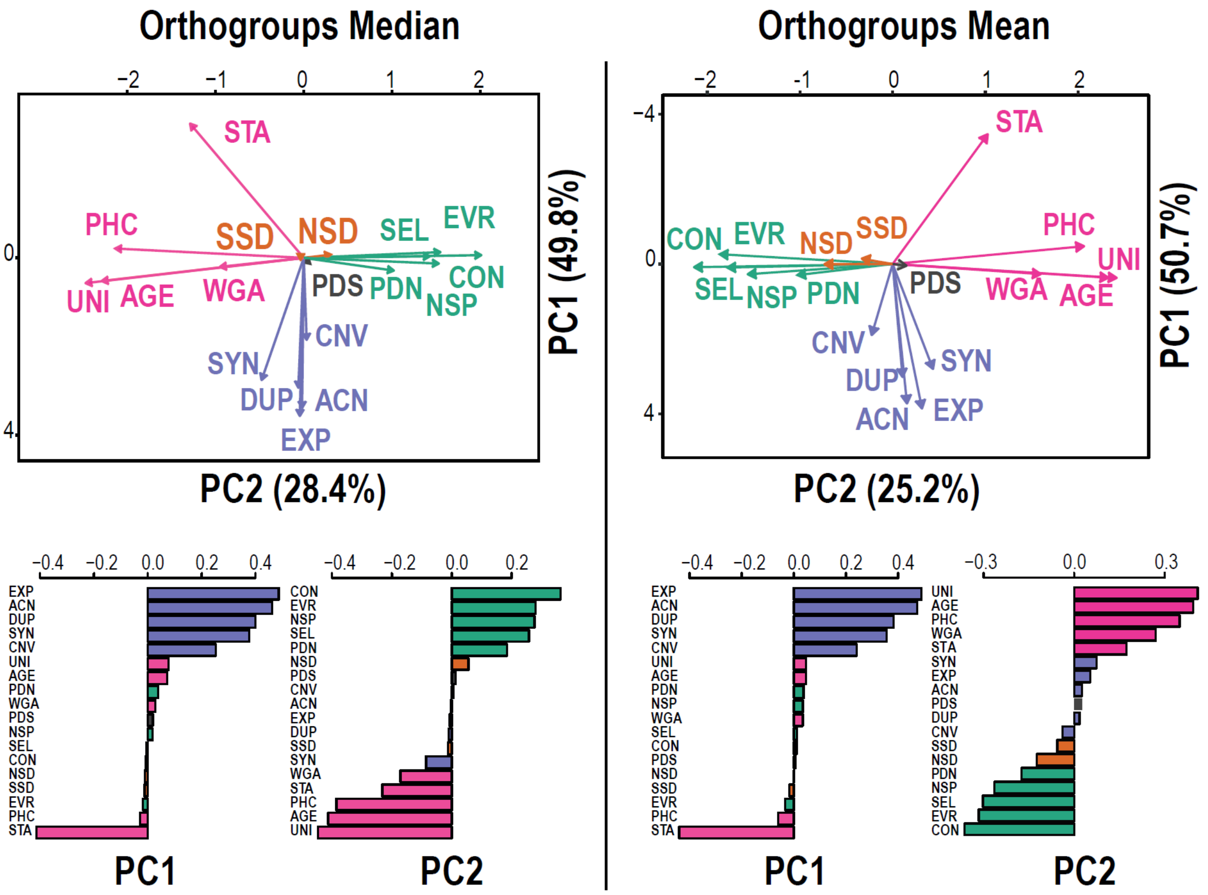


**Figure S8. Principal Component Analysis of median (left) and mean (right) evolutionary feature metrics**

The PCA loading plots show the strengths and directions of how each of the 18 evolutionary feature metrics contributes to the two principal components in two dimensions (above) and separately (below). These loadings support the subsets of features recovered by hierarchical clustering: (i, in green) contractions (gene losses), evolutionary rate (protein divergence), non-synonymous SNP proportion, PAML's dN, and PAML's dN/dS; (ii, in orange) densities of synonymous and non-synonymous

SNPs; (iii, in purple) average copy-number, copy-number variation, expansions, and duplications, often also including synteny as in Figure 2; and (iv, in pink) age, universality, alignability, and constraint, often also including stability as in Figure 2.

**Gene and family co-expression analyses**

The expression data and Expression Map (MacCallum et al. 2011) retrieved from VectorBase (Giraldo-Calderón et al. 2015) as described above, were used to quantify gene expression similarities among all pairs of immune-related families. First, the VectorBase Expression Map was analysed to quantify the frequencies with which each pair of families had member genes that clustered together in the same cell on the map. Next, this was extended to quantify the frequencies with which each pair of families had member genes that clustered together in the same supercell, i.e. the cell and its immediate eight neighbouring cells on the map (because neighbouring cells exhibit more similar expression profiles than more distantly placed cells). This provided a fine-scale (cell-level) and broad-scale (supercell) resolution of gene co-expression with which to assess gene family expression similarities. Specifically, the expression similarity scores (family cell/supercell co-occurrence scores) were computed as follows: the number of cells containing at least one gene from both Family 1 AND Family 2 / the unique number of cells containing at least one gene from either Family 1 OR Family 2 (i.e. intersection Family 1 ∩ Family 2 / union Family 1 ∪ Family 2). Extending this to supercells simply quantified gene membership of cells along with membership of the eight neighbouring cells including toroidal neighbours. Scores of zero indicate that these pairs of families have no member genes that cluster together in the same cell/supercell on the map, and scores of one would indicate that all member genes from both families always cluster together in cells/supercells on the map with at least one member of the other family. A permutation test was used to assess the statistical significance of the observed family cell/supercell co-occurrence scores: gene to cell assignments were randomly shuffled (Perl shuffle from List::Util) and co-occurrence scores were recomputed using the shuffled data 10'000 times. Shuffling preserves the total number of cells on the map, and the sizes (number of genes) of all cells and of all families, thereby taking into account the fact that larger families are more likely to have members clustered in the same cell by chance. For each pair of families, the number of times the permutation co-occurrence score was greater than the observed co-occurrence score was enumerated. The p-value was computed as the number of permutation scores that were greater than the observed score, divided by the total number of permutations, i.e. an empirical estimate of the probability of obtaining a co-occurrence score greater than the observed co-occurrence score by chance. Co-occurrence scores and their significance for all pairs of families are presented in Figure S9, plotted with the *corrplot* R package (Wei & Simko 2020) with results for cells above the diagonal and for supercells below the diagonal. A visualisation of all pairwise similarities reduced to two-dimensions was generated using a spring model layout network optimised with the *neato* tool from the Graphviz package (Gansner & North 2000; Gansner et al. 2005). Distances between nodes (families) in the network were provided as 1/co-occurrence score, where co-occurrence scores of zero were first re-set to the minimum observed co-occurrence score (separately for cells and supercells). Thus, a fully-connected set of nodes and edges was supplied from which to optimise the network using the spring model layout algorithm in *neato* with parameter *model=mds* such that the input length of an edge is used as the ideal distance between the nodes it connects.

Additional *neato* plotting parameters affecting layout optimisation: overlap=false (removes node overlaps); splines=true (draws edges as splines routed around nodes); epsilon=0.00001 (optimisation terminating condition); sep=0.5 (specifies margin to leave around nodes when removing overlaps). The resulting network (Figure S9) optimises the placement of families with the highest co-occurrence scores nearest to each other and the lowest co-occurrence scores farthest from each other, with node connections only plotted for the most significant ($p<0.005$) pairs of families in order to highlight the most important expression pattern similarities. For visualisation the thickness of the connecting lines was scaled to the p-value, with thicker lines indicating smaller p-values, and with p-values of <0.0005 first re-set to $p=0.0005$.

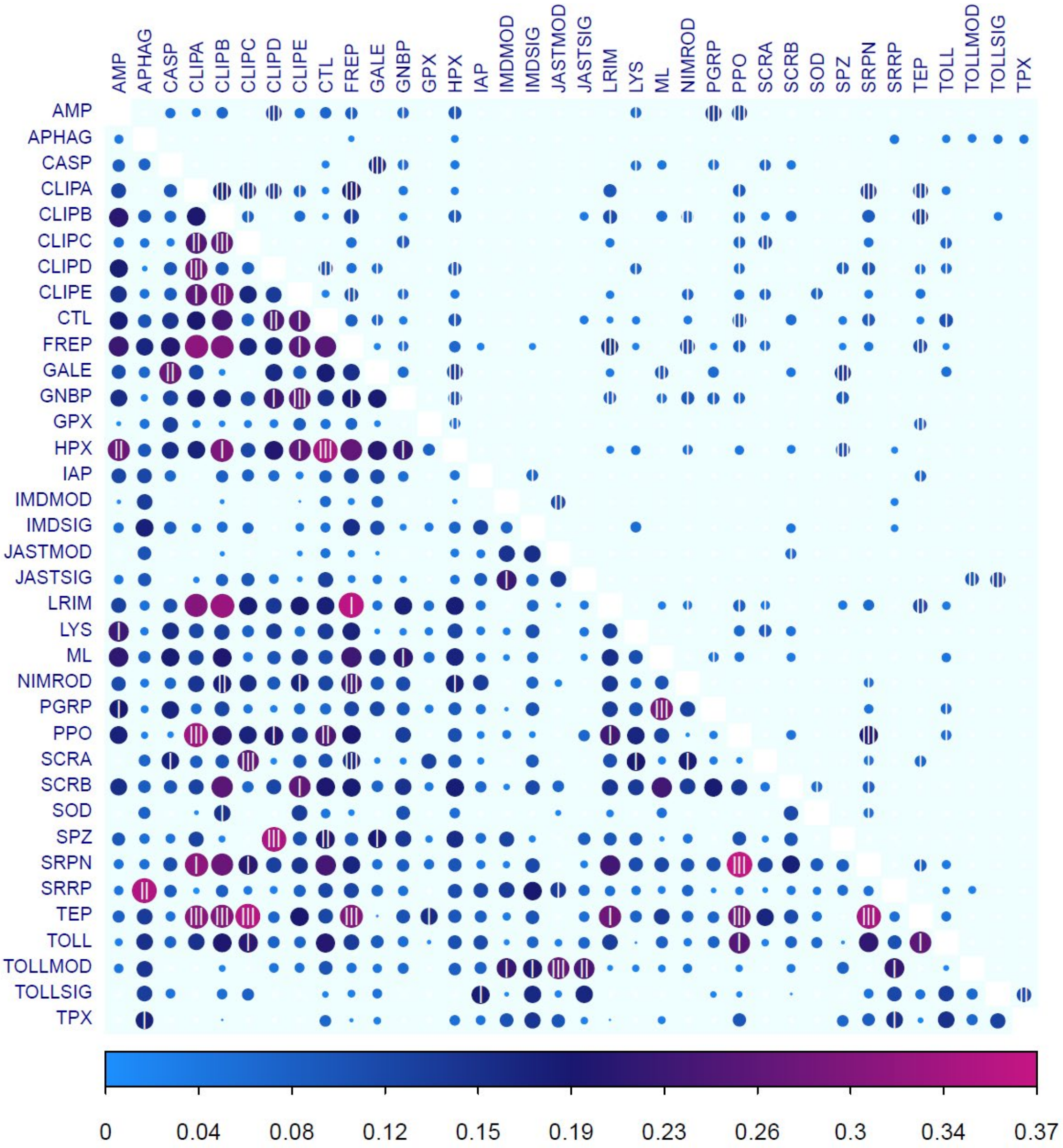


**Figure S9. Immune family expression similarities based on the VectorBase Expression Map.**
Pairwise comparisons of all 36 immune-related gene families (arranged in alphabetical order) showing their expression similarities computed as cell (above diagonal) and supercell (below diagonal) gene co-occurrence scores based on the VectorBase Expression Map (AgamP4.11 VB-2019-02). Circles are coloured and sized according to their co-occurrence score (see text for definition of score, cell, and supercell), and the significance of each co-occurrence is indicated with 3<0.005, 2<0.01, and 1<0.05 vertical lines.

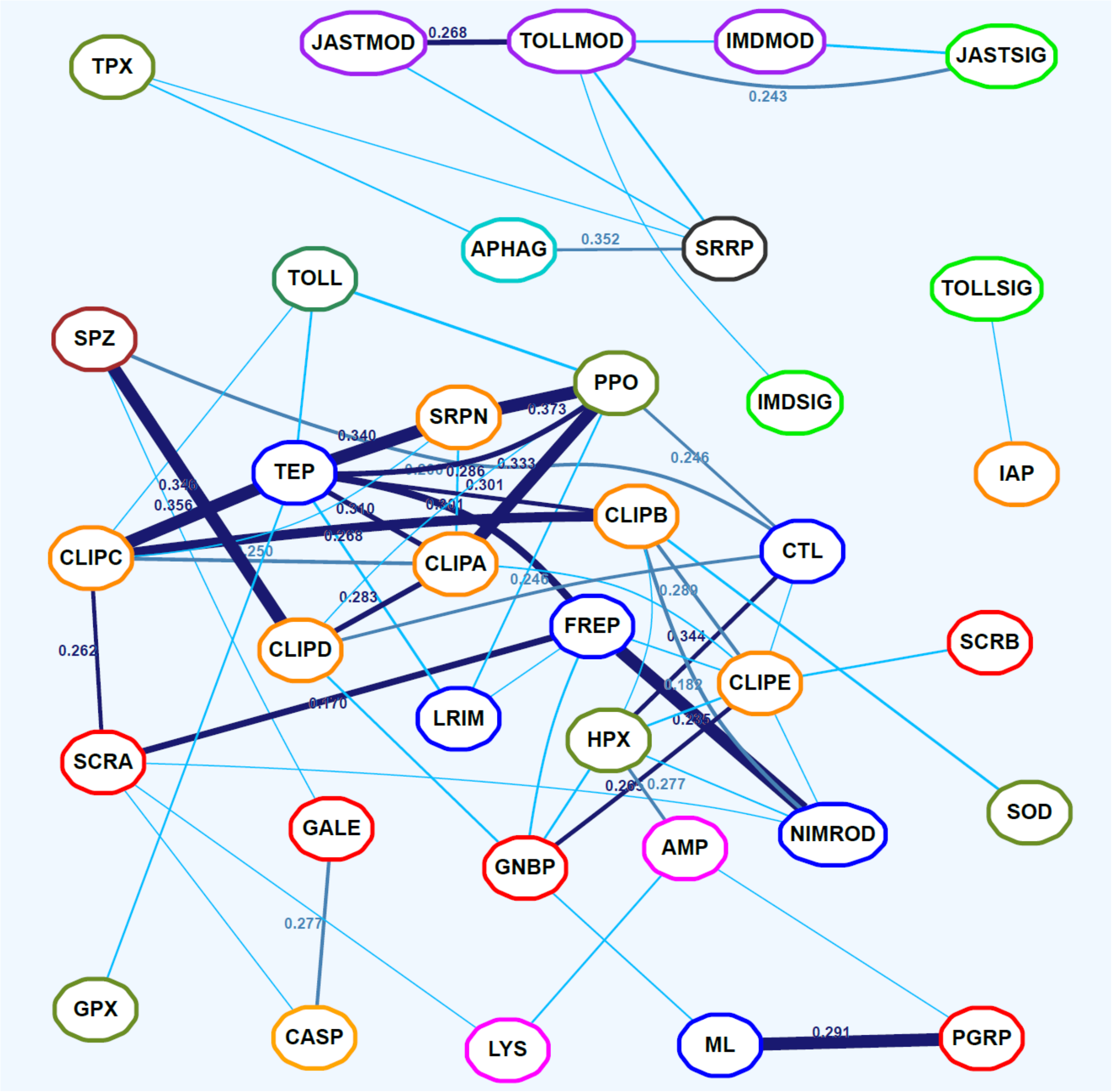


**Figure S10. Network of immune family expression similarities based on VectorBase Expression Map supercells.** The network layout optimised with a spring model provides a two-dimensional visualisation of expression similarities for pairwise comparisons of all 36 immune-related gene families computed as gene co-occurrence scores across supercells (cell + 8 neighbours) on the VectorBase Expression Map (AgamP4.11 VB-2019-02). Families with more similar gene expression profiles are placed closer together in the graph. Significant co-occurrences are indicated with connecting lines: light blue <0.005, royal blue <0.01, and dark blue <0.05 with line thickness scaled to the p-value and co-occurrence scores indicated for all pairs with p<0.01. Families are coloured according to categories defined based on their putative roles in the principal immune phases: classical recognition (red), other recognition (blue), pathway signalling (bright green), pathway modulation (purple), cascade modulation (orange), antimicrobial effectors (pink), effector enzymes (olive green), autophagy (dark cyan), RNAi (black), cytokines (brown), and toll receptors (dark green).

A subset of the conditions from the expression data retrieved from VectorBase was used to build co-expression modules using the weighted correlation network analysis (WGCNA) approach (Langfelder & Horvath 2008). This allowed for the quantification of gene expression similarities among all pairs of immune-related families based on selected conditions and using an alternative clustering approach to the self-organising map algorithm used to build the VectorBase Expression Map. Conditions were selected to focus on experiments assaying responses to blood feeding, because blood feeding is known to induce immune responses e.g. via haemocyte proliferation (Bryant & Michel 2014), or specific transcriptional changes (Dana et al. 2005; Bonizzoni et al. 2011; Upton et al. 2015). In addition, experiments assaying transcription in particularly immune-relevant tissues including the gut and fat body were selected, along with salivary glands, carcasses, and whole bodies in males and females. The 24 selected conditions (Table S2) limit the number of different data sources to just three publications (Marinotti et al. 2006; Neira Oviedo et al. 2008; Baker et al. 2011) in order to minimise technical variation and maximise the total number of genes across all conditions.

**Table S2. 24 experimental conditions selected for gene expression clustering with WGCNA.**

| Condition | Short Name |
|---|---|
| Blood meal time series (Marinotti et al., 2006) blood-fed 15d | Blood15d |
| Blood meal time series (Marinotti et al., 2006) blood-fed 24h | Blood24h |
| Blood meal time series (Marinotti et al., 2006) blood-fed 3h | Blood3h |
| Blood meal time series (Marinotti et al., 2006) blood_fed 48h | Blood48h |
| Blood meal time series (Marinotti et al., 2006) blood_fed 72h | Blood72h |
| Blood meal time series (Marinotti et al., 2006) blood-fed 96h | Blood96h |
| Blood meal time series (Marinotti et al., 2006) Non-blood-fed | BloodNBF |
| Blood meal after 15 days (Marinotti et al., 2006) Non-blood-fed 18d | BloodNBF18d |
| Two consecutive blood meals (Marinotti et al., 2006) blood-fed 24h | BloodTwo24h |
| Two consecutive blood meals (Marinotti et al., 2006) Non-blood-fed | BloodTwoNBF |
| Blood-fed adult female tissues (Marinotti et al., 2006) fat body | FatBodyBF |
| Alimentary canal compartments (Neira Oviedo et al., 2008) anterior midgut | GutAnt |
| Alimentary canal compartments (Neira Oviedo et al., 2008) gastric caeca | GutCaeca |
| Alimentary canal compartments (Neira Oviedo et al., 2008) hindgut | GutHind |
| Blood-fed adult female tissues (Marinotti et al., 2006) midgut | GutMidBF |
| Adult tissues (Baker et al., 2011) midgut:female | GutMidF |
| Adult tissues (Baker et al., 2011) midgut:male | GutMidM |
| Alimentary canal compartments (Neira Oviedo et al., 2008) posterior midgut | GutPost |
| Adult tissues (Baker et al., 2011) salivary gland:female | SalivF |
| Adult tissues (Baker et al., 2011) salivary gland:male | SalivM |
| Adult tissues (Baker et al., 2011) whole body:female | WholeF |
| Adult tissues (Baker et al., 2011) whole body:male | WholeM |
| Adult tissues (Baker et al., 2011) carcass:female | CarcassF |
| Adult tissues (Baker et al., 2011) carcass:male | CarcassM |

WGCNA proceeded first by assessing gene expression values across the 24 conditions using the *goodSamplesGenes* function to remove genes from the calculation with too many missing samples or zero variance, leaving a total of 10'434 genes for clustering. Next, a range of soft thresholding powers, the power to which co-expression similarity is raised to calculate adjacencies, was examined to select the most appropriate power. The *pickSoftThreshold* function was applied to compare powers 1-15 (increments of 1) and 16-32 (increments of 2) which showed power=10 as the lowest power with the greatest scale-free topology model fit. Adjacencies were computed using the *adjacency* function with power=10, which were then used to build a Topological Overlap Matrix (TOM) using the *TOMsimilarity* function, which was subsequently converted into a dissimilarity TOM for downstream analysis. Modules were then built using the *cutreeDynamic* function with the gene tree dendrogram generated with R's *hclust* function on the dissimilarity TOM with parameters pamRespectsDendro=FALSE, minClusterSize=12, and cutHeight=0.9, for five different levels of resolution defined by the parameter deepSplit set at 0, 1, 2, 3, and 4. For each of the five sets of modules, similar modules were then merged using the *mergeCloseModules* function with cutHeight=0.1 (chosen after inspecting hierarchical clustering of module eigengenes). The gene tree dendrogram is shown above the resulting 10 sets of modules in Figure S11 plotted with the *plotDendroAndColors* function. The lowest resolution (deepSplit=0) produced 42 modules (31 after merging), increasing resolution resulted in: deepSplit=1, 56 modules (35 after merging); deepSplit=2, 93 modules (47 after merging); deepSplit=3, 126 modules (53 after merging); deepSplit=4, 129 modules (55 after merging). Expression patterns per cluster were visualised as boxplots and coreplots (Additional File 3). Gene Ontology term enrichment was performed for each module using the classic Fisher and the weight01 Fisher tests implemented in the topGO package (Alexa et al. 2006) in R. Biological Process and Molecular Function ontologies for terms with at least 10 occurrences (nodeSize=10) were tested, results are presented for terms with at least two occurrences in the module and with p-values according to either test of $p<0.01$. Immune gene family expression similarities and significance of their co-occurrences were computed as above, here using module membership rather than cell/supercell membership, for the higher-resolution deepSplit=4 level modules and the lower-resolution deepSplit=2 level modules without merging (Figure S11).

**Additional File 3 description**

Visualisations of the 129 expression modules (deepSplit=4) built using the weighted correlation network analysis (WGCNA) approach with 24 selected conditions. Expression patterns per cluster are visualised as boxplots (showing meadians, 25th and 75th percentiles, lower and upper whiskers [1.5 x interquartile range], and outliers) and coreplots (showing the medians as a dotted line with coloured areas bounded by the ranges of the 25th and 75th percentiles), with the order of the conditions determined by hierarchical clustering with *hclust*. Tables show Biological Process and Molecular Function Gene Ontology terms for terms with at least two occurrences in the module and with p-values according to the classic Fisher or the weight01 Fisher tests of $p<0.01$.

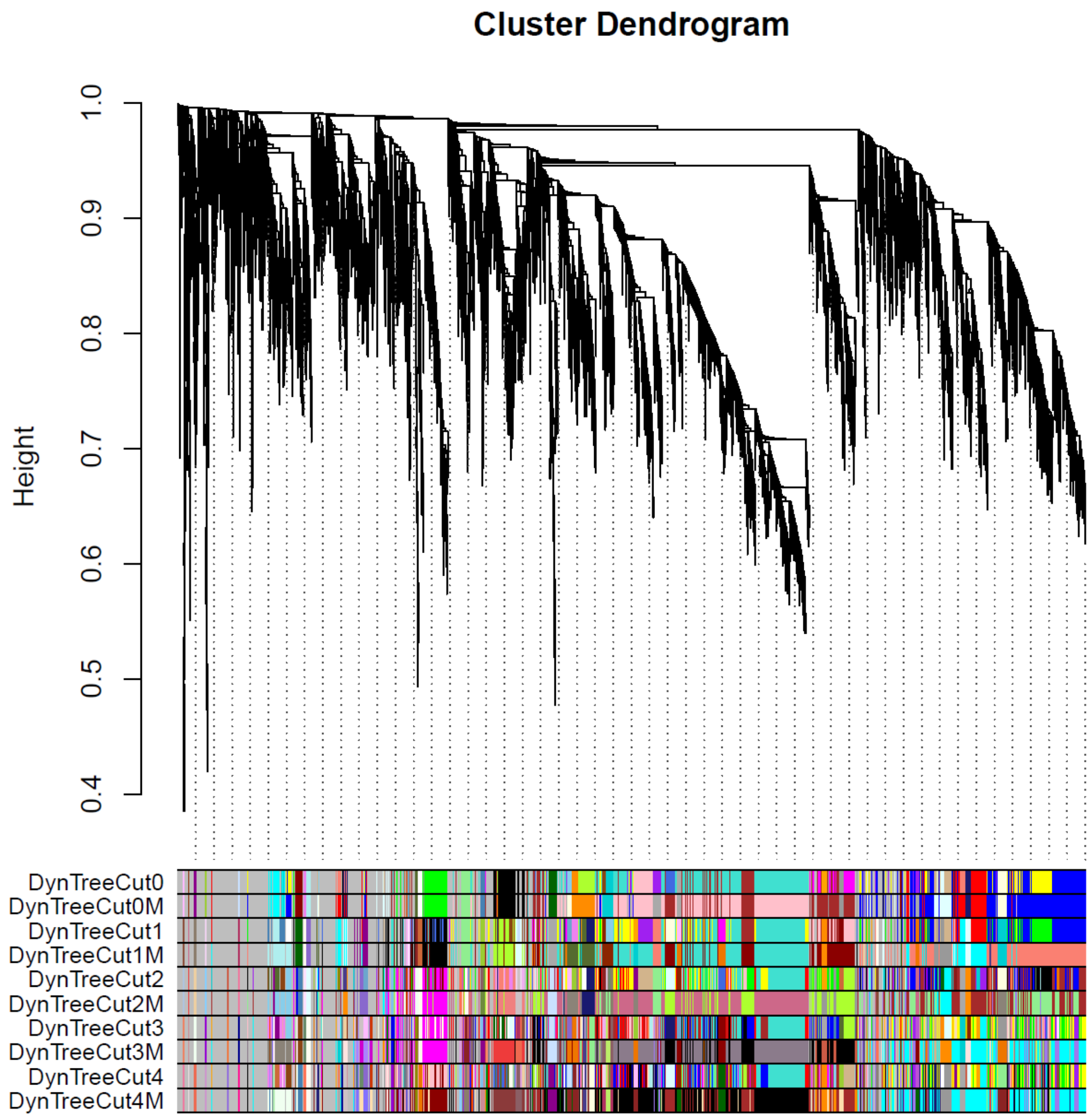


**Figure S11. WGCNA modules and expression tree.**

The hierarchical cluster dendrogram (top) from WGCNA analysis of 24 selected experimental conditions with gene module membership shown using coloured blocks (bottom) for ten different clustering resolutions. Each leaf of the tree corresponds to one of the 10'434 *An. gambiae* genes with enough variable expression data across the 24 conditions to be used as input for clustering with WGCNA. The Dynamic Tree Cut (DynTreeCut) algorithm was applied with five different spit levels (0 being the lowest and 4 being the highest) with subsequent merging (M) of similar modules at each level. The lowest resolution (deepSplit=0) produced 42 modules (31 after merging), increasing resolution resulted in: deepSplit=1, 56 modules (35 after merging); deepSplit=2, 93 modules (47 after merging); deepSplit=3, 126 modules (53 after merging); deepSplit=4, 129 modules (55 after merging).

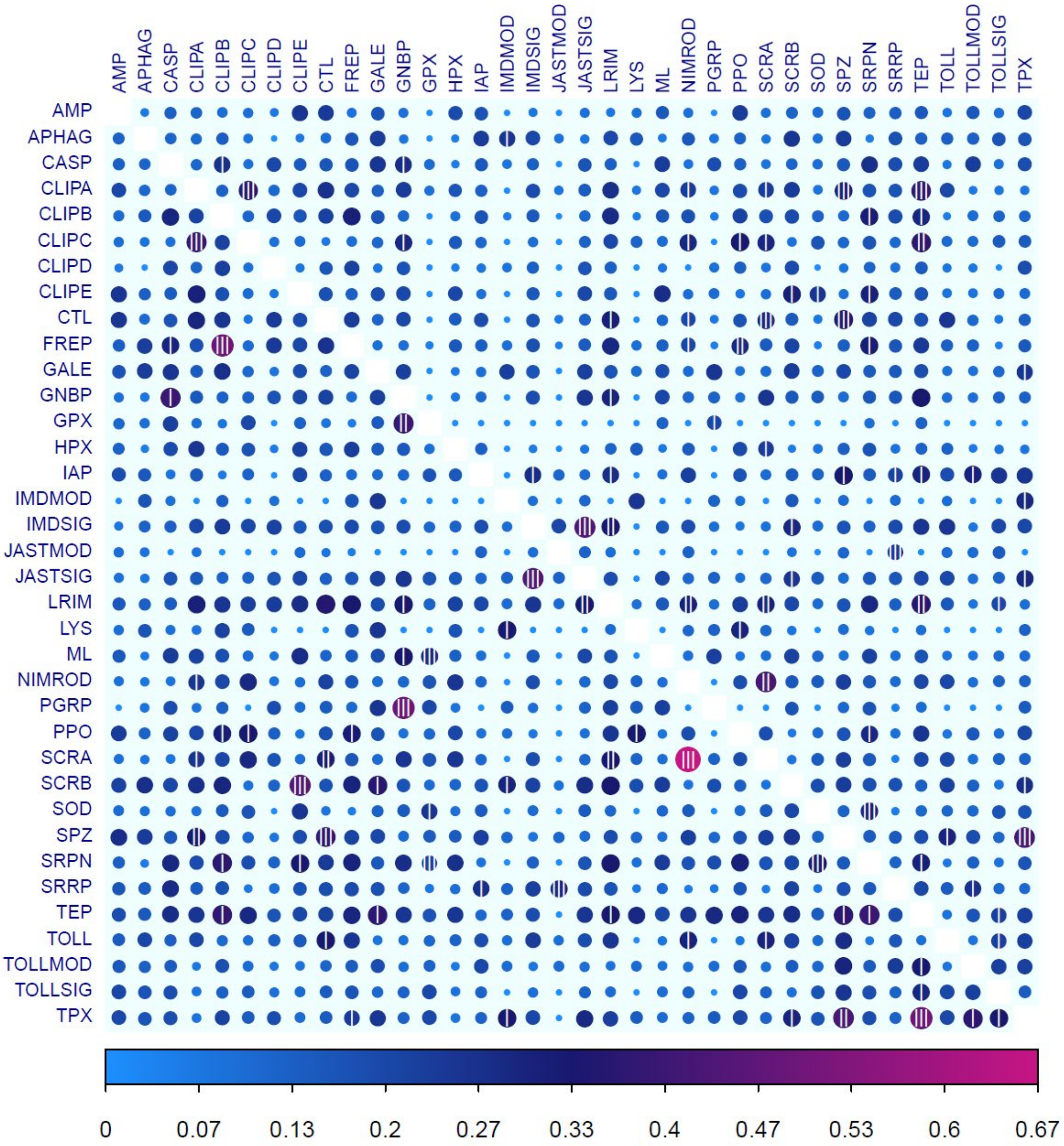


**Figure S12. Immune family co-expression across WGCNA modules.**

Pairwise comparisons of all 36 immune-related gene families (arranged in alphabetical order) showing their expression similarities computed as level 4 modules (above diagonal) and level 2 modules (below diagonal) gene co-occurrence scores based on WGCNA clustering of 24 selected conditions from the VectorBase expression dataset. Circles are coloured and sized according to their co-occurrence score (see text for definition of score, and module), and the significance of each co-occurrence is indicated with 3<0.005, 2<0.01, and 1<0.05 vertical lines.

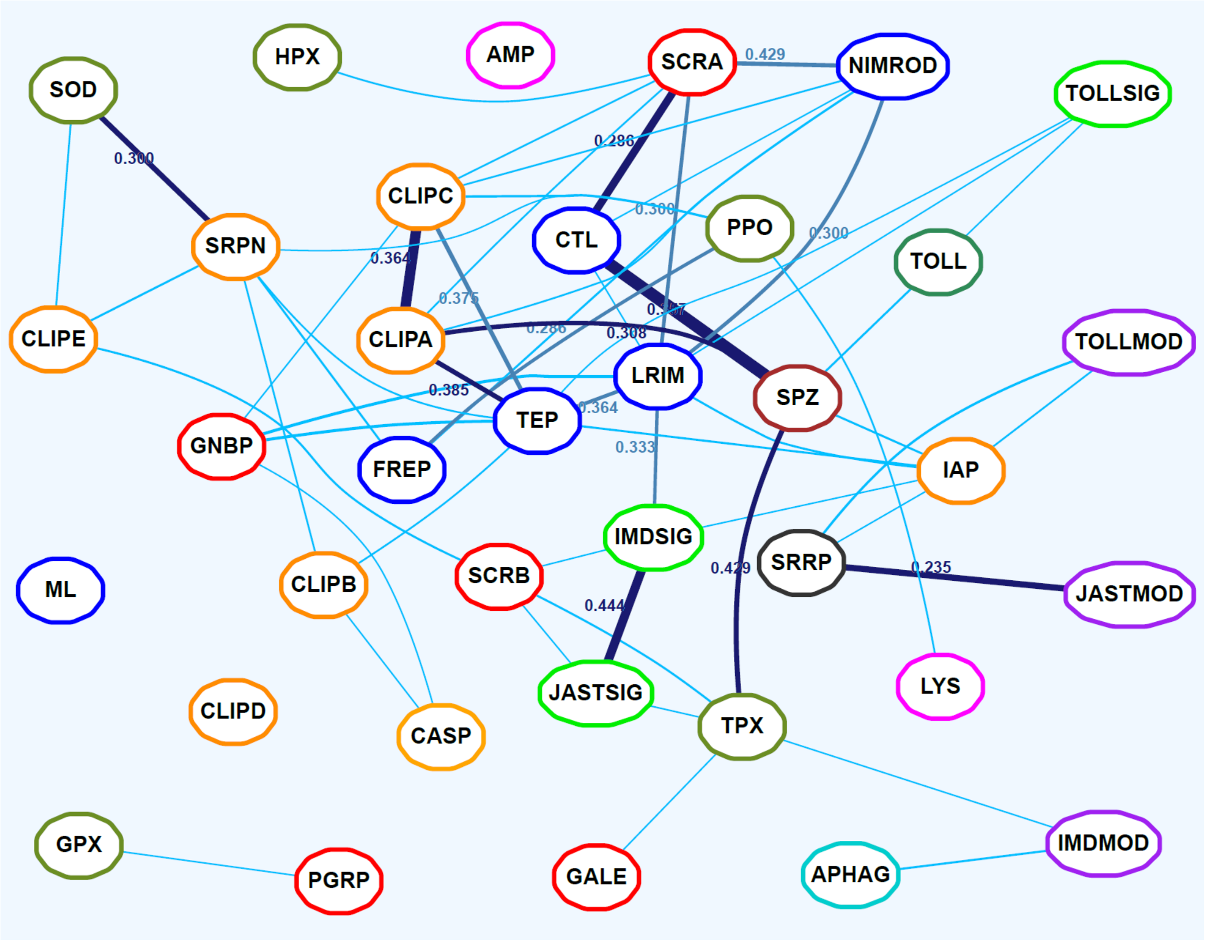


**Figure S13. Network of immune family expression similarities based on level 4 WGCNA modules.**
The network layout optimised with a spring model provides a two-dimensional visualisation of expression similarities for pairwise comparisons of all 36 immune-related gene families computed as gene co-occurrence scores level 4 modules delineated by WGCNA. Families with more similar gene expression profiles are placed closer together in the graph. Significant co-occurrences are indicated with connecting lines: light blue <0.005, royal blue <0.01, and dark blue <0.05 with line thickness scaled to the p-value and co-occurrence scores indicated for all pairs with p<0.01. Families are coloured according to categories defined based on their putative roles in the principal immune phases: classical recognition (red), other recognition (blue), pathway signalling (bright green), pathway modulation (purple), cascade modulation (orange), antimicrobial effectors (pink), effector enzymes (olive green), autophagy (dark cyan), RNAi (black), cytokines (brown), and toll receptors (dark green).

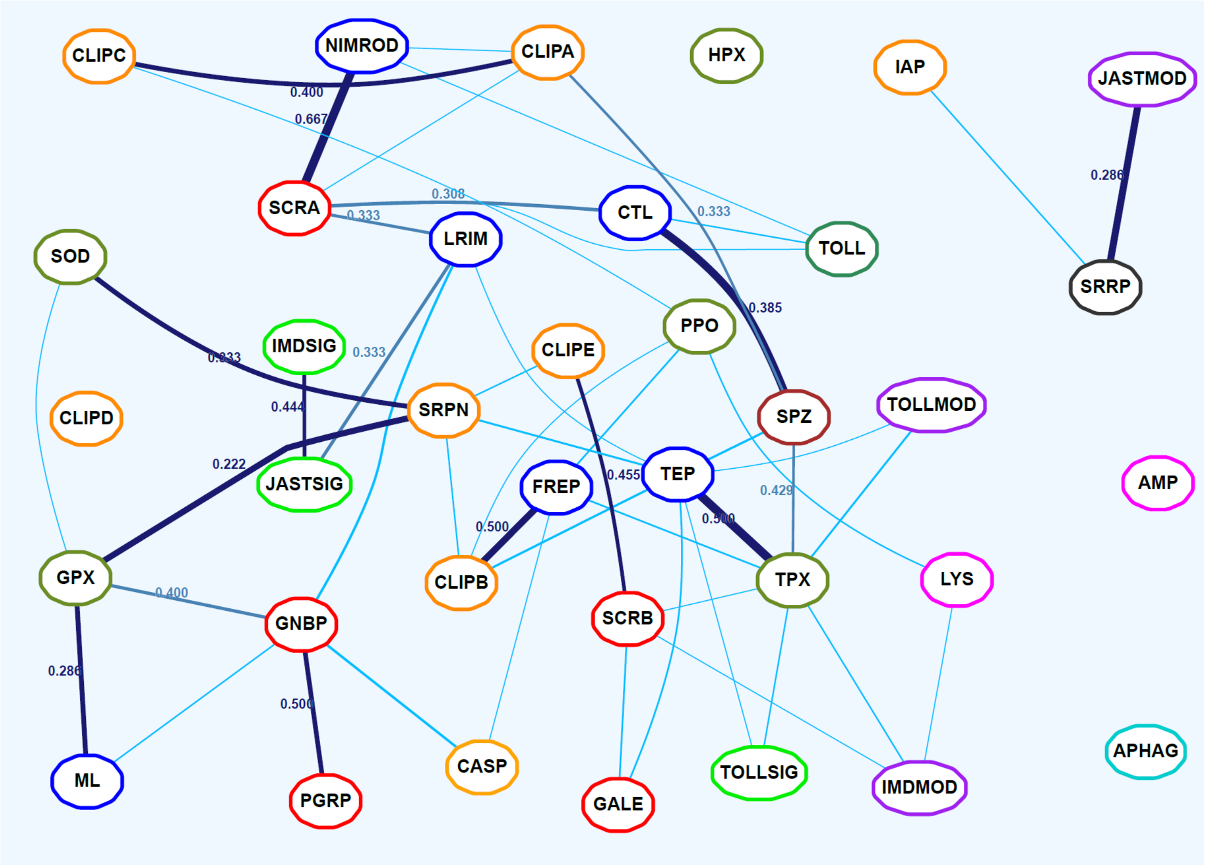


**Figure S14. Network of immune family expression similarities based on level 2 WGCNA modules.**
The network layout optimised with a spring model provides a two-dimensional visualisation of expression similarities for pairwise comparisons of all 36 immune-related gene families computed as gene co-occurrence scores level 2 modules delineated by WGCNA. Families with more similar gene expression profiles are placed closer together in the graph. Significant co-occurrences are indicated with connecting lines: light blue <0.005, royal blue <0.01, and dark blue <0.05 with line thickness scaled to the p-value and co-occurrence scores indicated for all pairs with p<0.01. Families are coloured according to categories defined based on their putative roles in the principal immune phases: classical recognition (red), other recognition (blue), pathway signalling (bright green), pathway modulation (purple), cascade modulation (orange), antimicrobial effectors (pink), effector enzymes (olive green), autophagy (dark cyan), RNAi (black), cytokines (brown), and toll receptors (dark green).

### Additional File 4 description

Additional comparisons of evolutionary similarities versus expression similarities are presented showing evolutionary similarities computed with mean (EvolMean) and median (EvolMedian) metrics for the VectorBase cells (ExprCell) and supercells (ExprSupercell) as well as for the WGCNA expression modules for levels 0-4 (ExprModuleN). Plots are built as for main text Figure 4, but with automatic label placement and without colouring by immune categories. Median values are shown as horizontal and vertical lines in yellow for all points and in orange for significant (purple) points.

### List of databases and software used

Computational Analysis of gene Family Evolution (CAFE) version: 4.2.1: (Han et al. 2013), https://www.ncbi.nlm.nih.gov/pubmed/23709260
Website: https://github.com/hahnlab/CAFE
Jim Kent utilities collection: UCSC, 2020, https://hgdownload.soe.ucsc.edu/downloads.html
axtChain, chainAntiRepeat and chainNet binaries, packaged in version 376 (Jan. 23rd, 2019), http://hgdownload.cse.ucsc.edu/admin/jksrc.v376.zip
LastZ, version 1.04.00 : Harris, 2007, http://www.bx.psu.edu/~rsharris/rsharris_phd_thesis_2007.pdf
Website: http://www.bx.psu.edu/~rsharris/lastz/
MultiZ - Threaded Blockset Aligner: (Blanchette et al. 2004), https://www.ncbi.nlm.nih.gov/pubmed/15060014
Website: https://github.com/multiz/multiz
ROAST, version 3 and single_cov2, version 11
Newick Utilities, version 1.6: (Junier & Zdobnov 2010), https://www.ncbi.nlm.nih.gov/pubmed/20472542
Website: https://github.com/tjunier/newick_utils/wiki
OrthoDB, version 8: (Kriventseva et al. 2015), https://www.ncbi.nlm.nih.gov/pubmed/25428351
Website: https://www.orthodb.org/v8/
OrthoDBmoz2: (Neafsey et al. 2015), https://www.ncbi.nlm.nih.gov/pubmed/25554792
PHAST (PHylogenetic Analysis with Space/Time models): https://www.ncbi.nlm.nih.gov/pubmed/21278375
Website: http://compgen.cshl.edu/phast/
PhastCons: Siepel *et al.*, 2005, https://www.ncbi.nlm.nih.gov/pubmed/16024819
Website: http://compgen.cshl.edu/phast/help-pages/phastCons.txt
phyloFit: Siepel *et al.*, 2004 https://www.ncbi.nlm.nih.gov/pubmed/14660683
Website address : http://compgen.cshl.edu/phast/help-pages/phyloFit.txt
Phylogenetic Analysis by Maximum Likelihood (PAML), version 4.8a: (Yang 2007), https://www.ncbi.nlm.nih.gov/pubmed/17483113
Website: http://abacus.gene.ucl.ac.uk/software/paml.html
R: Ihaka and Gentleman, 1996, https://www.tandfonline.com/doi/abs/10.1080/10618600.1996.10474713
Website: https://cran.r-project.org/ (4.0.2)
RAxML, version 8.2.11: (Stamatakis 2014), https://www.ncbi.nlm.nih.gov/pubmed/24451623
Website: https://cme.h-its.org/exelixis/web/software/raxml/
RStudio: https://rstudio.com/ (1.3.1093)
TimeTree: (Kumar et al. 2017), https://www.ncbi.nlm.nih.gov/pubmed/28387841
Website: http://www.timetree.org/
VectorBase: (Giraldo-Calderón et al. 2015), https://www.ncbi.nlm.nih.gov/pubmed/25510499
Website: https://www.vectorbase.org/

### R packages and functions

ape: (Paradis & Schliep 2019), https://www.ncbi.nlm.nih.gov/pubmed/30016406
ape function 5.4-1, https://www.rdocumentation.org/packages/ape/versions/5.4-1
dendextend: (Galili 2015), https://www.ncbi.nlm.nih.gov/pubmed/26209431
dendextend function 1.14.0, https://www.rdocumentation.org/packages/dendextend/versions/1.14.0
gplots: Warnes *et al.*, https://www.rdocumentation.org/packages/gplots/versions/3.1.1
3.1.1
phytools: (Revell 2012), https://besjournals.onlinelibrary.wiley.com/doi/10.1111/j.2041-210X.2011.00169.x

phytools function 0.7-20: https://www.rdocumentation.org/packages/phytools/versions/0.7-20

pvclust: (Suzuki & Shimodaira 2006), https://www.ncbi.nlm.nih.gov/pubmed/16595560

pvclust function 2.2-0, https://www.rdocumentation.org/packages/pvclust/versions/2.2-0/topics/pvclust

RColorBrewer: 1.1-2 Erich Neuwirth, https://www.rdocumentation.org/packages/RColorBrewer/versions/1.1-2/topics/RColorBrewer

Based on ColorBrewer2 (https://colorbrewer2.org/) by Brewer and Harrower, 2003, https://www.tandfonline.com/doi/abs/10.1179/000870403235002042

stats: R-core 3.6.3, https://www.rdocumentation.org/packages/stats/versions/3.6.2

cor function, https://www.rdocumentation.org/packages/stats/versions/3.6.2/topics/cor

dist function, https://www.rdocumentation.org/packages/stats/versions/3.6.2/topics/dist

hclust function, https://www.rdocumentation.org/packages/stats/versions/3.6.2/topics/hclust

wilcox.test function, https://www.rdocumentation.org/packages/stats/versions/3.6.2/topics/wilcox.test

### Supplementary References


Alexa A, Rahnenfuhrer J, Lengauer T. 2006. Improved scoring of functional groups from gene expression data by decorrelating GO graph structure. Bioinformatics. 22:1600–1607. doi: 10.1093/bioinformatics/btl140.

Baker DA et al. 2011. A comprehensive gene expression atlas of sex- and tissue-specificity in the malaria vector, Anopheles gambiae. BMC Genomics. 12:296. doi: 10.1186/1471-2164-12-296.

Bartholomay LC et al. 2010. Pathogenomics of Culex quinquefasciatus and meta-analysis of infection responses to diverse pathogens. Science. 330:88–90. doi: 10.1126/science.1193162.

Blanchette M et al. 2004. Aligning multiple genomic sequences with the threaded blockset aligner. Genome Res. 14:708–715. doi: 10.1101/gr.1933104.

Bonizzoni M et al. 2011. RNA-seq analyses of blood-induced changes in gene expression in the mosquito vector species, Aedes aegypti. BMC Genomics. 12:82. doi: 10.1186/1471-2164-12-82.

Bryant WB, Michel K. 2014. Blood feeding induces hemocyte proliferation and activation in the African malaria mosquito, Anopheles gambiae Giles. J. Exp. Biol. 217:1238–1245. doi: 10.1242/jeb.094573.

Christophides GK et al. 2002. Immunity-related genes and gene families in Anopheles gambiae. Science. 298:159–165. doi: 10.1126/science.1077136.

Dana AN et al. 2005. Gene expression patterns associated with blood-feeding in the malaria mosquito Anopheles gambiae. BMC Genomics. 6:5. doi: 10.1186/1471-2164-6-5.

Galili T. 2015. dendextend: an R package for visualizing, adjusting and comparing trees of hierarchical clustering. Bioinformatics. 31:3718–3720. doi: 10.1093/bioinformatics/btv428.

Gansner ER, Koren Y, North S. 2005. Graph Drawing by Stress Majorization. In: Graph Drawing. Pach, J, editor. Lecture Notes in Computer Science Vol. 3383 Springer Berlin Heidelberg: Berlin, Heidelberg pp. 239–250. doi: 10.1007/978-3-540-31843-9_25.

Gansner ER, North SC. 2000. An open graph visualization system and its applications to software engineering. Softw. - Pract. Exp. 30:1203–1233.

Giraldo-Calderón GI et al. 2015. VectorBase: an updated bioinformatics resource for invertebrate vectors and other organisms related with human diseases. Nucleic Acids Res. 43:D707–D713. doi: 10.1093/nar/gku1117.

Hammond AM et al. 2017. The creation and selection of mutations resistant to a gene drive over multiple generations in the malaria mosquito Barton, NH, editor. PLOS Genet. 13:e1007039. doi: 10.1371/journal.pgen.1007039.

Han MV, Thomas GWC, Lugo-Martinez J, Hahn MW. 2013. Estimating gene gain and loss rates in the presence of error in genome assembly and annotation using CAFE 3. Mol. Biol. Evol. 30:1987–1997. doi: 10.1093/molbev/mst100.

Harris RS. 2007. Improved pairwise alignment of genomic DNA. https://etda.libraries.psu.edu/catalog/7971 (Accessed May 13, 2020).

Junier T, Zdobnov EM. 2010. The Newick utilities: high-throughput phylogenetic tree processing in the Unix shell. Bioinformatics. 26:1669–1670. doi: 10.1093/bioinformatics/btq243.

Kafatos F, Waterhouse R, Zdobnov E, Christophides G. 2009. Comparative genomics of insect immunity. In: Insect Infection and Immunity: Evolution, Ecology, and Mechanisms. doi: 10.1093/acprof:oso/9780199551354.003.0006.

Kriventseva EV et al. 2015. OrthoDB v8: update of the hierarchical catalog of orthologs and the underlying free software. Nucleic Acids Res. 43:D250–D256. doi: 10.1093/nar/gku1220.

Kumar S, Stecher G, Suleski M, Hedges SB. 2017. TimeTree: a resource for timelines, timetrees, and divergence times. Mol. Biol. Evol. 34:1812–1819. doi: 10.1093/molbev/msx116.

Langfelder P, Horvath S. 2008. WGCNA: an R package for weighted correlation network analysis. BMC Bioinformatics. 9:559. doi: 10.1186/1471-2105-9-559.

MacCallum RM, Redmond SN, Christophides GK. 2011. An expression map for Anopheles gambiae. BMC Genomics. 12:620. doi: 10.1186/1471-2164-12-620.

Marinotti O et al. 2006. Genome-wide analysis of gene expression in adult Anopheles gambiae. Insect Mol. Biol. 15:1–12. doi: 10.1111/j.1365-2583.2006.00610.x.

Markianos K et al. 2016. Genetic structure of a local population of the Anopheles gambiae complex in Burkina Faso Brooke, B, editor. PLOS ONE. 11:e0145308. doi: 10.1371/journal.pone.0145308.

Miles A, The Anopheles gambiae 1000 Genomes Consortium, Kwiatkowski D. 2017. Genetic diversity of the African malaria vector Anopheles gambiae. Nature. 552:96–100. doi: 10.1038/nature24995.

Murtagh F, Legendre P. 2014. Ward's Hierarchical Agglomerative Clustering Method: Which Algorithms Implement Ward's Criterion? J. Classif. 31:274–295. doi: 10.1007/s00357-014-9161-z.

Neafsey DE et al. 2015. Highly evolvable malaria vectors: The genomes of 16 Anopheles mosquitoes. Science. 347:1258522. doi: 10.1126/science.1258522.

Neafsey DE et al. 2010. SNP genotyping defines complex gene-flow boundaries among African malaria vector mosquitoes. Science. 330:514–517. doi: 10.1126/science.1193036.
Neira Oviedo M, VanEkeris L, Corena-Mcleod MDP, Linser PJ. 2008. A microarray-based analysis of transcriptional compartmentalization in the alimentary canal of Anopheles gambiae (Diptera: Culicidae) larvae: Gut transcriptome of larval An.gambiae. Insect Mol. Biol. 17:61–72. doi: 10.1111/j.1365-2583.2008.00779.x.
Paradis E, Schliep K. 2019. ape 5.0: an environment for modern phylogenetics and evolutionary analyses in R Schwartz, R, editor. Bioinformatics. 35:526–528. doi: 10.1093/bioinformatics/bty633.
Povelones M, Osta MA, Christophides GK. 2016. The complement system of malaria vector mosquitoes. In: Advances in Insect Physiology.Vol. 51 Elsevier pp. 223–242. doi: 10.1016/bs.aiip.2016.06.001.
Revell LJ. 2012. phytools: an R package for phylogenetic comparative biology (and other things). Methods Ecol. Evol. 3:217–223. doi: 10.1111/j.2041-210X.2011.00169.x.
Rodgers FH, Gendrin M, Christophides GK. 2017. The Mosquito Immune System and Its Interactions With the Microbiota. In: Arthropod Vector: Controller of Disease Transmission, Volume 1. Elsevier pp. 101–122. doi: 10.1016/B978-0-12-805350-8.00006-4.
Siepel A et al. 2005. Evolutionarily conserved elements in vertebrate, insect, worm, and yeast genomes. Genome Res. 15:1034–1050. doi: 10.1101/gr.3715005.
Sim S, Jupatanakul N, Dimopoulos G. 2014. Mosquito Immunity against Arboviruses. Viruses. 6:4479–4504. doi: 10.3390/v6114479.
Simões ML, Caragata EP, Dimopoulos G. 2018. Diverse Host and Restriction Factors Regulate Mosquito–Pathogen Interactions. Trends Parasitol. 34:603–616. doi: 10.1016/j.pt.2018.04.011.
Stamatakis A. 2014. RAxML version 8: a tool for phylogenetic analysis and post-analysis of large phylogenies. Bioinformatics. 30:1312–1313. doi: 10.1093/bioinformatics/btu033.
Suzuki R, Shimodaira H. 2006. Pvclust: an R package for assessing the uncertainty in hierarchical clustering. Bioinformatics. 22:1540–1542. doi: 10.1093/bioinformatics/btl117.
Terradas G, Joubert DA, McGraw EA. 2017. The RNAi pathway plays a small part in Wolbachia-mediated blocking of dengue virus in mosquito cells. Sci. Rep. 7:43847. doi: 10.1038/srep43847.
Upton LM, Povelones M, Christophides GK. 2015. Anopheles gambiae blood feeding initiates an anticipatory defense response to Plasmodium berghei. J. Innate Immun. 7:74–86. doi: 10.1159/000365331.
Waterhouse RM. 2009. Computational comparative analysis of insect genomes. Ph.D., Imperial College London https://ethos.bl.uk/OrderDetails.do?uin=uk.bl.ethos.512005 (Accessed May 15, 2020).
Waterhouse RM et al. 2007. Evolutionary dynamics of immune-related genes and pathways in disease-vector mosquitoes. Science. 316:1738–1743. doi: 10.1126/science.1139862.
Waterhouse RM, Lazzaro BP, Sackton TB. 2020. Characterization of insect immune systems from genomic data. In: Immunity in Insects. Sandrelli, F & Tettamanti, G, editors. Springer Protocols Handbooks Springer US: New York, NY pp. 3–34. doi: 10.1007/978-1-0716-0259-1_1.
Waterhouse RM, Tegenfeldt F, Li J, Zdobnov EM, Kriventseva EV. 2013. OrthoDB: A hierarchical catalog of animal, fungal and bacterial orthologs. Nucleic Acids Res. 41:D358-65. doi: 10.1093/nar/gks1116.
Weetman D, Wilding CS, Steen K, Pinto J, Donnelly MJ. 2012. Gene flow-dependent genomic divergence between Anopheles gambiae M and S forms. Mol. Biol. Evol. 29:279–291. doi: 10.1093/molbev/msr199.
Wei T, Simko V. 2020. R package ‘corrplot’: Visualization of a Correlation Matrix. https://github.com/taiyun/corrplot.
White BJ et al. 2011. Adaptive divergence between incipient species of Anopheles gambiae increases resistance to Plasmodium. Proc. Natl. Acad. Sci. 108:244–249. doi: 10.1073/pnas.1013648108.
Wiltshire RM et al. 2018. Reduced-representation sequencing identifies small effective population sizes of Anopheles gambiae in the north-western Lake Victoria basin, Uganda. Malar. J. 17:285. doi: 10.1186/s12936-018-2432-0.
Yang Z. 2007. PAML 4: phylogenetic analysis by maximum likelihood. Mol. Biol. Evol. 24:1586–1591. doi: 10.1093/molbev/msm088.
