## Additional File 1 for "Functional constraints on insect immune system components govern their evolutionary trajectories"

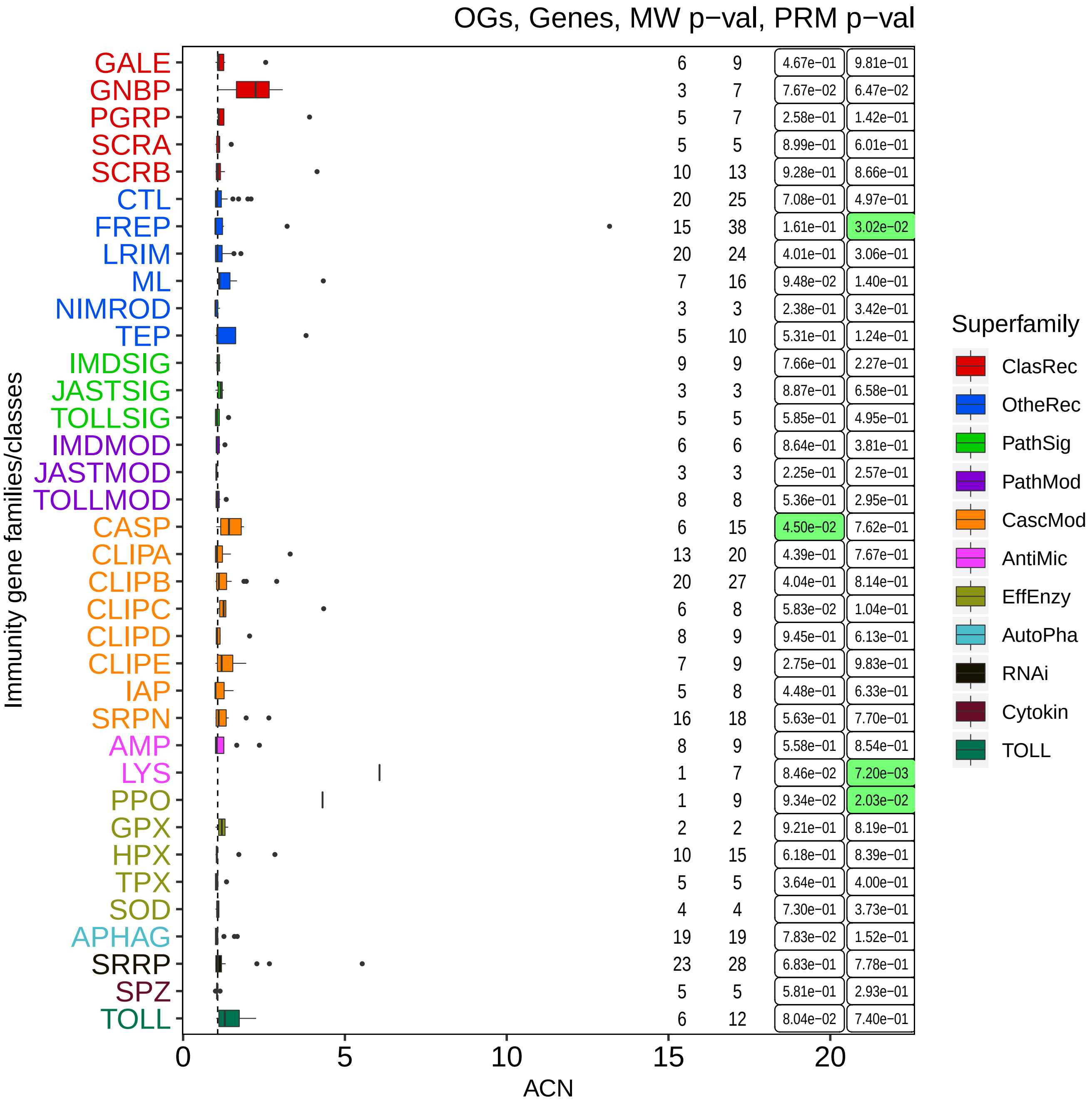
OGs, Genes, MW p−val, PRM p−val
GALE 6 9 4.67e−01 9.81e−01
GNBP 3 7 7.67e−02 6.47e−02
PGRP 5 7 2.58e−01 1.42e−01
SCRA 5 5 8.99e−01 6.01e−01
SCRB 10 13 9.28e−01 8.66e−01
CTL 20 25 7.08e−01 4.97e−01
FREP 15 38 1.61e−01 3.02e−02
LRIM 20 24 4.01e−01 3.06e−01
ML 7 16 9.48e−02 1.40e−01
NIMROD 3 3 2.38e−01 3.42e−01
TEP 5 10 5.31e−01 1.24e−01
IMDSIG 9 9 7.66e−01 2.27e−01
JASTSIG 3 3 8.87e−01 6.58e−01
TOLLSIG 5 5 5.85e−01 4.95e−01
IMDMOD 6 6 8.64e−01 3.81e−01
JASTMOD 3 3 2.25e−01 2.57e−01
TOLLMOD 8 8 5.36e−01 2.95e−01
CASP 6 15 4.50e−02 7.62e−01
CLIPA 13 20 4.39e−01 7.67e−01
CLIPB 20 27 4.04e−01 8.14e−01
CLIPC 6 8 5.83e−02 1.04e−01
CLIPD 8 9 9.45e−01 6.13e−01
CLIPE 7 9 2.75e−01 9.83e−01
IAP 5 8 4.48e−01 6.33e−01
SRPN 16 18 5.63e−01 7.70e−01
AMP 8 9 5.58e−01 8.54e−01
LYS 1 7 8.46e−02 7.20e−03
PPO 1 9 9.34e−02 2.03e−02
GPX 2 2 9.21e−01 8.19e−01
HPX 10 15 6.18e−01 8.39e−01
TPX 5 5 3.64e−01 4.00e−01
SOD 4 4 7.30e−01 3.73e−01
APHAG 19 19 7.83e−02 1.52e−01
SRRP 23 28 6.83e−01 7.78e−01
SPZ 5 5 5.81e−01 2.93e−01
TOLL 6 12 8.04e−02 7.40e−01
Immunity gene families/classes
0
5
10
15
20
ACN
Superfamily
ClasRec
OtheRec
PathSig
PathMod
CascMod
AntiMic
EffEnzy
AutoPha
RNAi
Cytokin
TOLL

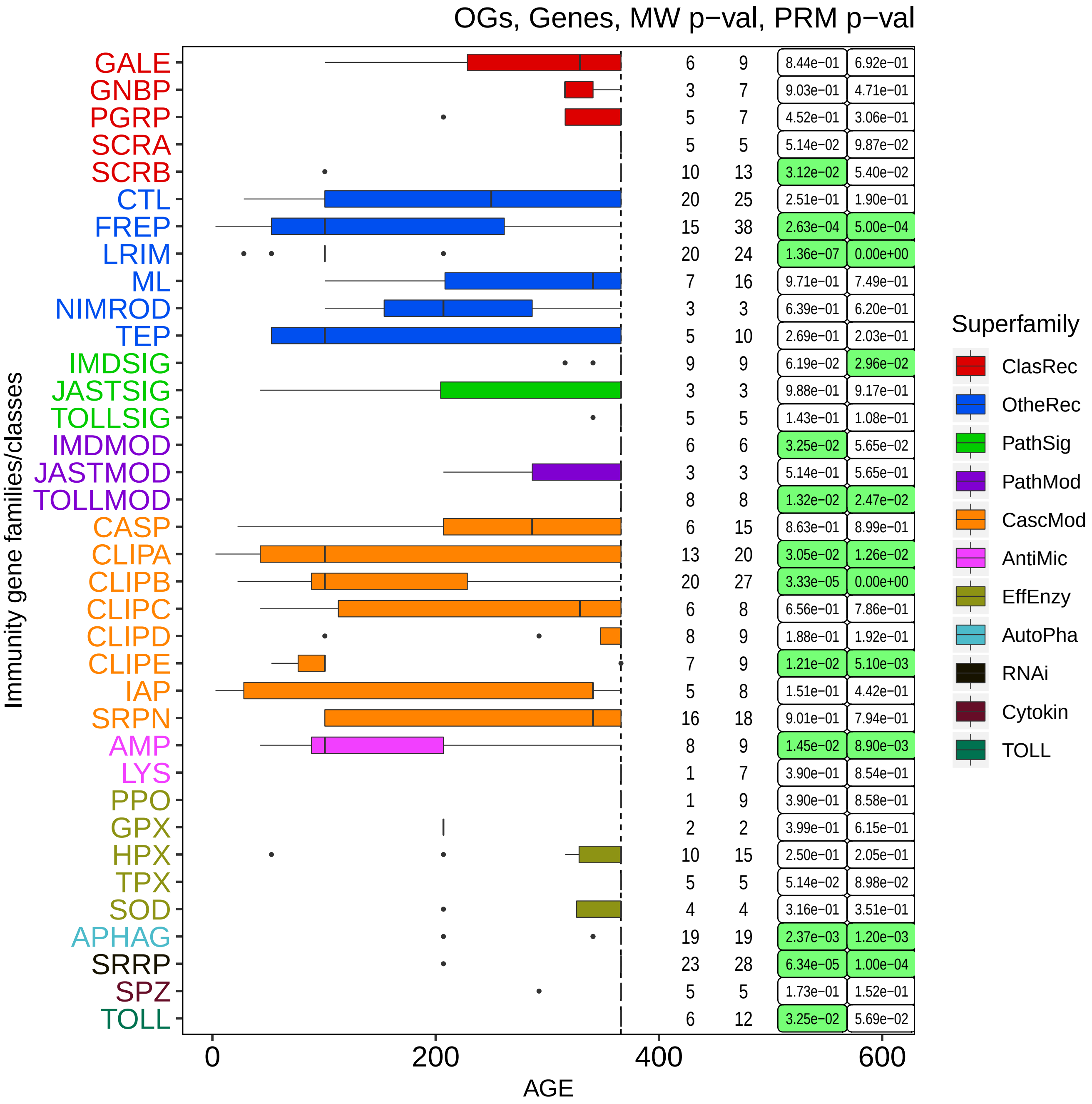
OGs, Genes, MW p−val, PRM p−val
Immunity gene families/classes
GALE 6 9 8.44e-01 6.92e-01
GNBP 3 7 9.03e-01 4.71e-01
PGRP 5 7 4.52e-01 3.06e-01
SCRA 5 5 5.14e-02 9.87e-02
SCRB 10 13 3.12e-02 5.40e-02
CTL 20 25 2.51e-01 1.90e-01
FREP 15 38 2.63e-04 5.00e-04
LRIM 20 24 1.36e-07 0.00e+00
ML 7 16 9.71e-01 7.49e-01
NIMROD 3 3 6.39e-01 6.20e-01
TEP 5 10 2.69e-01 2.03e-01
IMDSIG 9 9 6.19e-02 2.96e-02
JASTSIG 3 3 9.88e-01 9.17e-01
TOLLSIG 5 5 1.43e-01 1.08e-01
IMDMOD 6 6 3.25e-02 5.65e-02
JASTMOD 3 3 5.14e-01 5.65e-01
TOLLMOD 8 8 1.32e-02 2.47e-02
CASP 6 15 8.63e-01 8.99e-01
CLIPA 13 20 3.05e-02 1.26e-02
CLIPB 20 27 3.33e-05 0.00e+00
CLIPC 6 8 6.56e-01 7.86e-01
CLIPD 8 9 1.88e-01 1.92e-01
CLIPE 7 9 1.21e-02 5.10e-03
IAP 5 8 1.51e-01 4.42e-01
SRPN 16 18 9.01e-01 7.94e-01
AMP 8 9 1.45e-02 8.90e-03
LYS 1 7 3.90e-01 8.54e-01
PPO 1 9 3.90e-01 8.58e-01
GPX 2 2 3.99e-01 6.15e-01
HPX 10 15 2.50e-01 2.05e-01
TPX 5 5 5.14e-02 8.98e-02
SOD 4 4 3.16e-01 3.51e-01
APHAG 19 19 2.37e-03 1.20e-03
SRRP 23 28 6.34e-05 1.00e-04
SPZ 5 5 1.73e-01 1.52e-01
TOLL 6 12 3.25e-02 5.69e-02
0
200
400
600
AGE
Superfamily
ClasRec
OtheRec
PathSig
PathMod
CascMod
AntiMic
EffEnzy
AutoPha
RNAi
Cytokin
TOLL

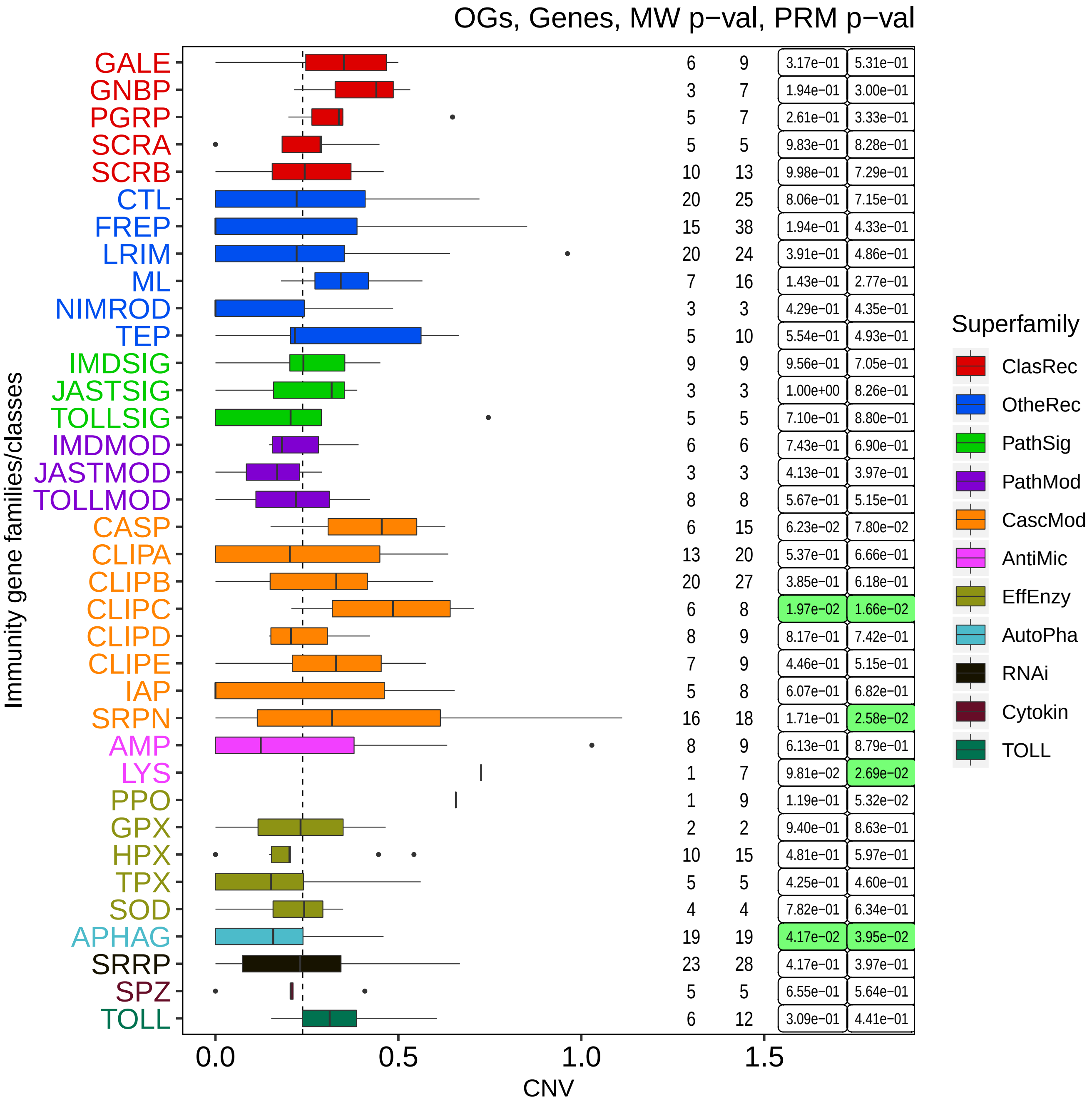
OGs, Genes, MW p−val, PRM p−val
GALE 6 9 3.17e-01 5.31e-01
GNBP 3 7 1.94e-01 3.00e-01
PGRP 5 7 2.61e-01 3.33e-01
SCRA 5 5 9.83e-01 8.28e-01
SCRB 10 13 9.98e-01 7.29e-01
CTL 20 25 8.06e-01 7.15e-01
FREP 15 38 1.94e-01 4.33e-01
LRIM 20 24 3.91e-01 4.86e-01
ML 7 16 1.43e-01 2.77e-01
NIMROD 3 3 4.29e-01 4.35e-01
TEP 5 10 5.54e-01 4.93e-01
IMDSIG 9 9 9.56e-01 7.05e-01
JASTSIG 3 3 1.00e+00 8.26e-01
TOLLSIG 5 5 7.10e-01 8.80e-01
IMDMOD 6 6 7.43e-01 6.90e-01
JASTMOD 3 3 4.13e-01 3.97e-01
TOLLMOD 8 8 5.67e-01 5.15e-01
CASP 6 15 6.23e-02 7.80e-02
CLIPA 13 20 5.37e-01 6.66e-01
CLIPB 20 27 3.85e-01 6.18e-01
CLIPC 6 8 1.97e-02 1.66e-02
CLIPD 8 9 8.17e-01 7.42e-01
CLIPE 7 9 4.46e-01 5.15e-01
IAP 5 8 6.07e-01 6.82e-01
SRPN 16 18 1.71e-01 2.58e-02
AMP 8 9 6.13e-01 8.79e-01
LYS 1 7 9.81e-02 2.69e-02
PPO 1 9 1.19e-01 5.32e-02
GPX 2 2 9.40e-01 8.63e-01
HPX 10 15 4.81e-01 5.97e-01
TPX 5 5 4.25e-01 4.60e-01
SOD 4 4 7.82e-01 6.34e-01
APHAG 19 19 4.17e-02 3.95e-02
SRRP 23 28 4.17e-01 3.97e-01
SPZ 5 5 6.55e-01 5.64e-01
TOLL 6 12 3.09e-01 4.41e-01
Immunity gene families/classes
0.0
0.5
1.0
1.5
CNV
Superfamily
ClasRec
OtheRec
PathSig
PathMod
CascMod
AntiMic
EffEnzy
AutoPha
RNAi
Cytokin
TOLL

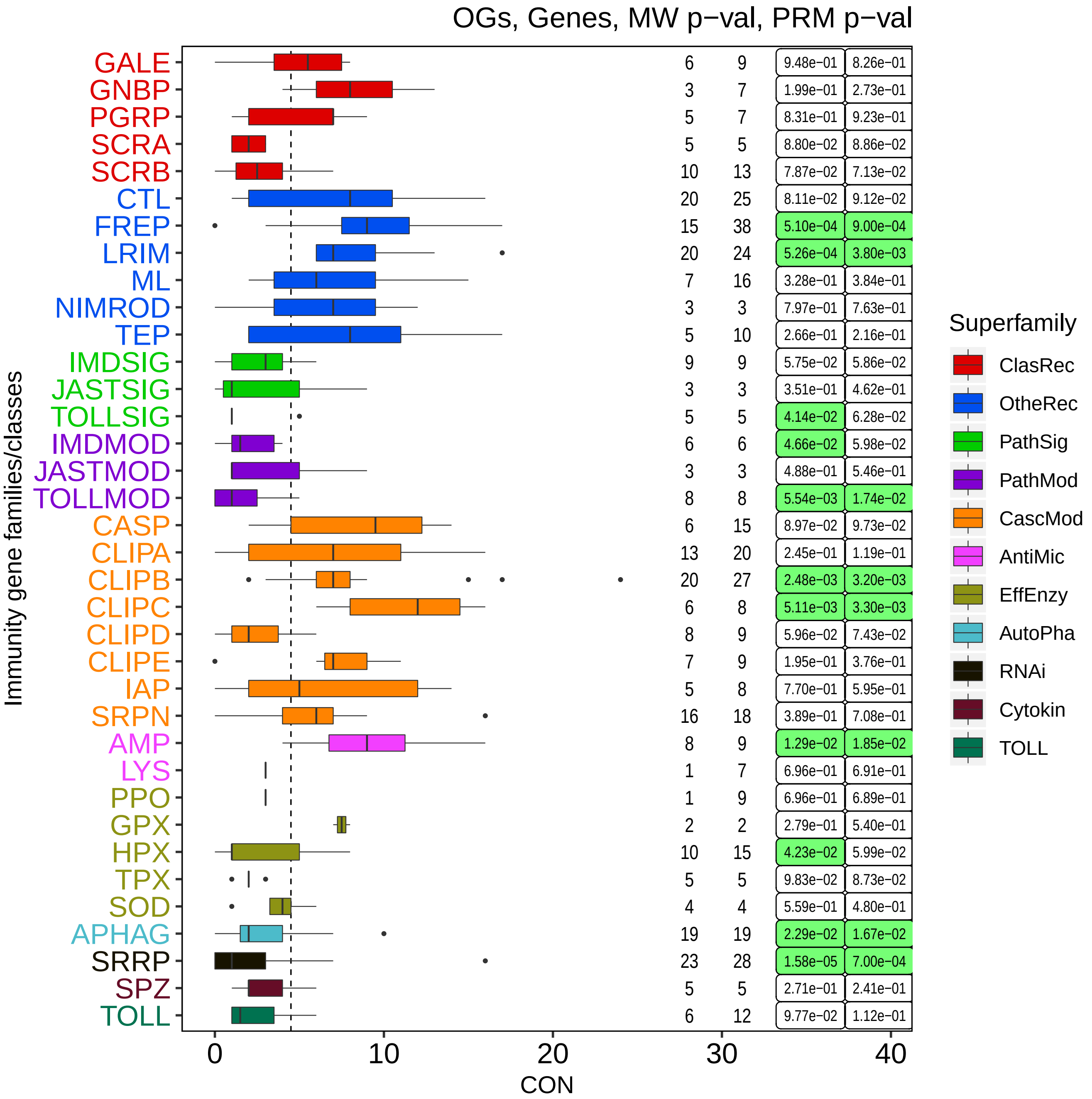
OGs, Genes, MW p−val, PRM p−val
Immunity gene families/classes
GALE 6 9 9.48e-01 8.26e-01
GNBP 3 7 1.99e-01 2.73e-01
PGRP 5 7 8.31e-01 9.23e-01
SCRA 5 5 8.80e-02 8.86e-02
SCRB 10 13 7.87e-02 7.13e-02
CTL 20 25 8.11e-02 9.12e-02
FREP 15 38 5.10e-04 9.00e-04
LRIM 20 24 5.26e-04 3.80e-03
ML 7 16 3.28e-01 3.84e-01
NIMROD 3 3 7.97e-01 7.63e-01
TEP 5 10 2.66e-01 2.16e-01
IMDSIG 9 9 5.75e-02 5.86e-02
JASTSIG 3 3 3.51e-01 4.62e-01
TOLLSIG 5 5 4.14e-02 6.28e-02
IMDMOD 6 6 4.66e-02 5.98e-02
JASTMOD 3 3 4.88e-01 5.46e-01
TOLLMOD 8 8 5.54e-03 1.74e-02
CASP 6 15 8.97e-02 9.73e-02
CLIPA 13 20 2.45e-01 1.19e-01
CLIPB 20 27 2.48e-03 3.20e-03
CLIPC 6 8 5.11e-03 3.30e-03
CLIPD 8 9 5.96e-02 7.43e-02
CLIPE 7 9 1.95e-01 3.76e-01
IAP 5 8 7.70e-01 5.95e-01
SRPN 16 18 3.89e-01 7.08e-01
AMP 8 9 1.29e-02 1.85e-02
LYS 1 7 6.96e-01 6.91e-01
PPO 1 9 6.96e-01 6.89e-01
GPX 2 2 2.79e-01 5.40e-01
HPX 10 15 4.23e-02 5.99e-02
TPX 5 5 9.83e-02 8.73e-02
SOD 4 4 5.59e-01 4.80e-01
APHAG 19 19 2.29e-02 1.67e-02
SRRP 23 28 1.58e-05 7.00e-04
SPZ 5 5 2.71e-01 2.41e-01
TOLL 6 12 9.77e-02 1.12e-01
0
10
20
30
40
CON
Superfamily
ClasRec
OtheRec
PathSig
PathMod
CascMod
AntiMic
EffEnzy
AutoPha
RNAi
Cytokin
TOLL

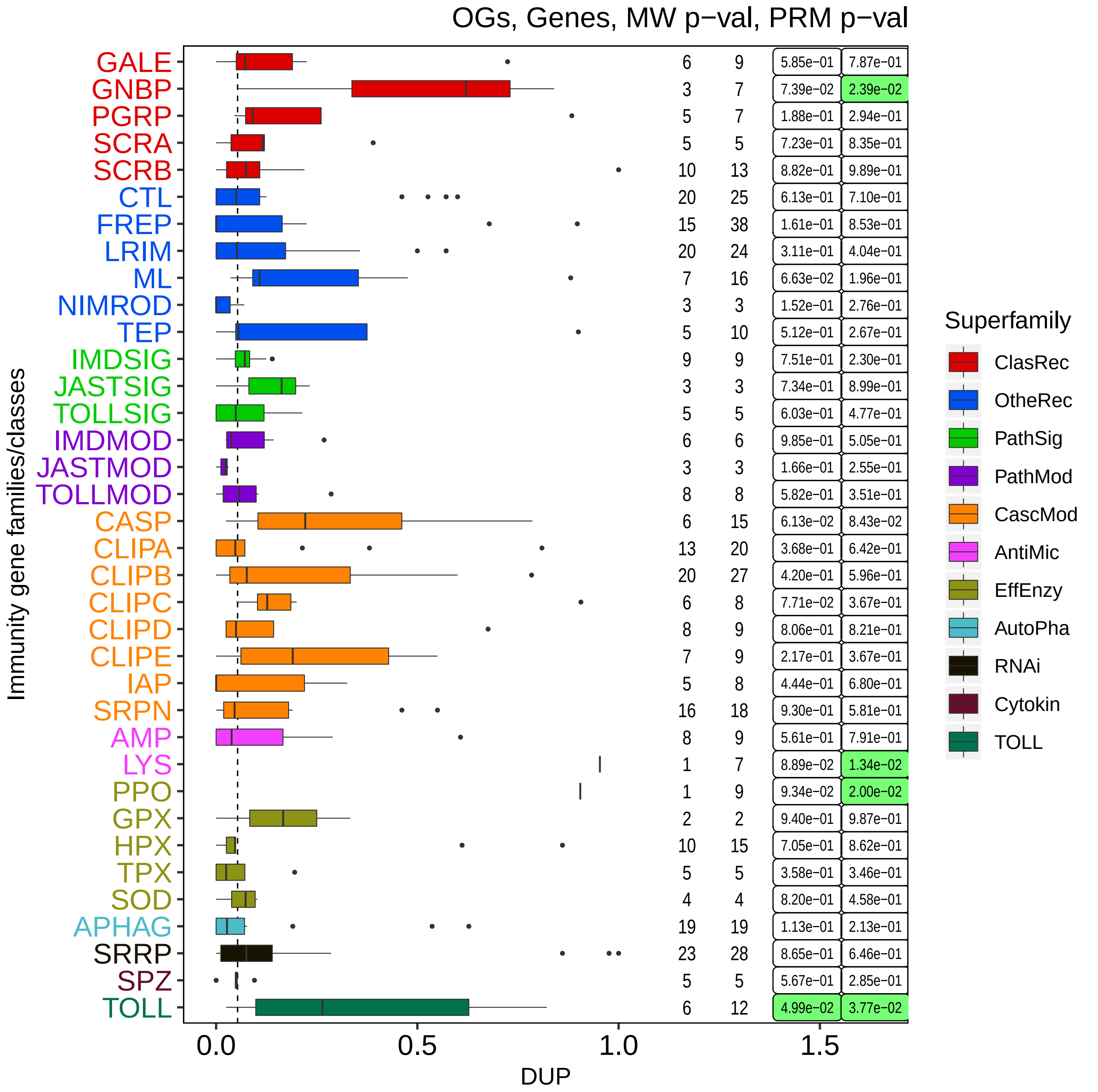
OGs, Genes, MW p−val, PRM p−val
Immunity gene families/classes
GALE 6 9 5.85e−01 7.87e−01
GNBP 3 7 7.39e−02 2.39e−02
PGRP 5 7 1.88e−01 2.94e−01
SCRA 5 5 7.23e−01 8.35e−01
SCRB 10 13 8.82e−01 9.89e−01
CTL 20 25 6.13e−01 7.10e−01
FREP 15 38 1.61e−01 8.53e−01
LRIM 20 24 3.11e−01 4.04e−01
ML 7 16 6.63e−02 1.96e−01
NIMROD 3 3 1.52e−01 2.76e−01
TEP 5 10 5.12e−01 2.67e−01
IMDSIG 9 9 7.51e−01 2.30e−01
JASTSIG 3 3 7.34e−01 8.99e−01
TOLLSIG 5 5 6.03e−01 4.77e−01
IMDMOD 6 6 9.85e−01 5.05e−01
JASTMOD 3 3 1.66e−01 2.55e−01
TOLLMOD 8 8 5.82e−01 3.51e−01
CASP 6 15 6.13e−02 8.43e−02
CLIPA 13 20 3.68e−01 6.42e−01
CLIPB 20 27 4.20e−01 5.96e−01
CLIPC 6 8 7.71e−02 3.67e−01
CLIPD 8 9 8.06e−01 8.21e−01
CLIPE 7 9 2.17e−01 3.67e−01
IAP 5 8 4.44e−01 6.80e−01
SRPN 16 18 9.30e−01 5.81e−01
AMP 8 9 5.61e−01 7.91e−01
LYS 1 7 8.89e−02 1.34e−02
PPO 1 9 9.34e−02 2.00e−02
GPX 2 2 9.40e−01 9.87e−01
HPX 10 15 7.05e−01 8.62e−01
TPX 5 5 3.58e−01 3.46e−01
SOD 4 4 8.20e−01 4.58e−01
APHAG 19 19 1.13e−01 2.13e−01
SRRP 23 28 8.65e−01 6.46e−01
SPZ 5 5 5.67e−01 2.85e−01
TOLL 6 12 4.99e−02 3.77e−02
0.0
0.5
1.0
1.5
DUP
Superfamily
ClasRec
OtheRec
PathSig
PathMod
CascMod
AntiMic
EffEnzy
AutoPha
RNAi
Cytokin
TOLL

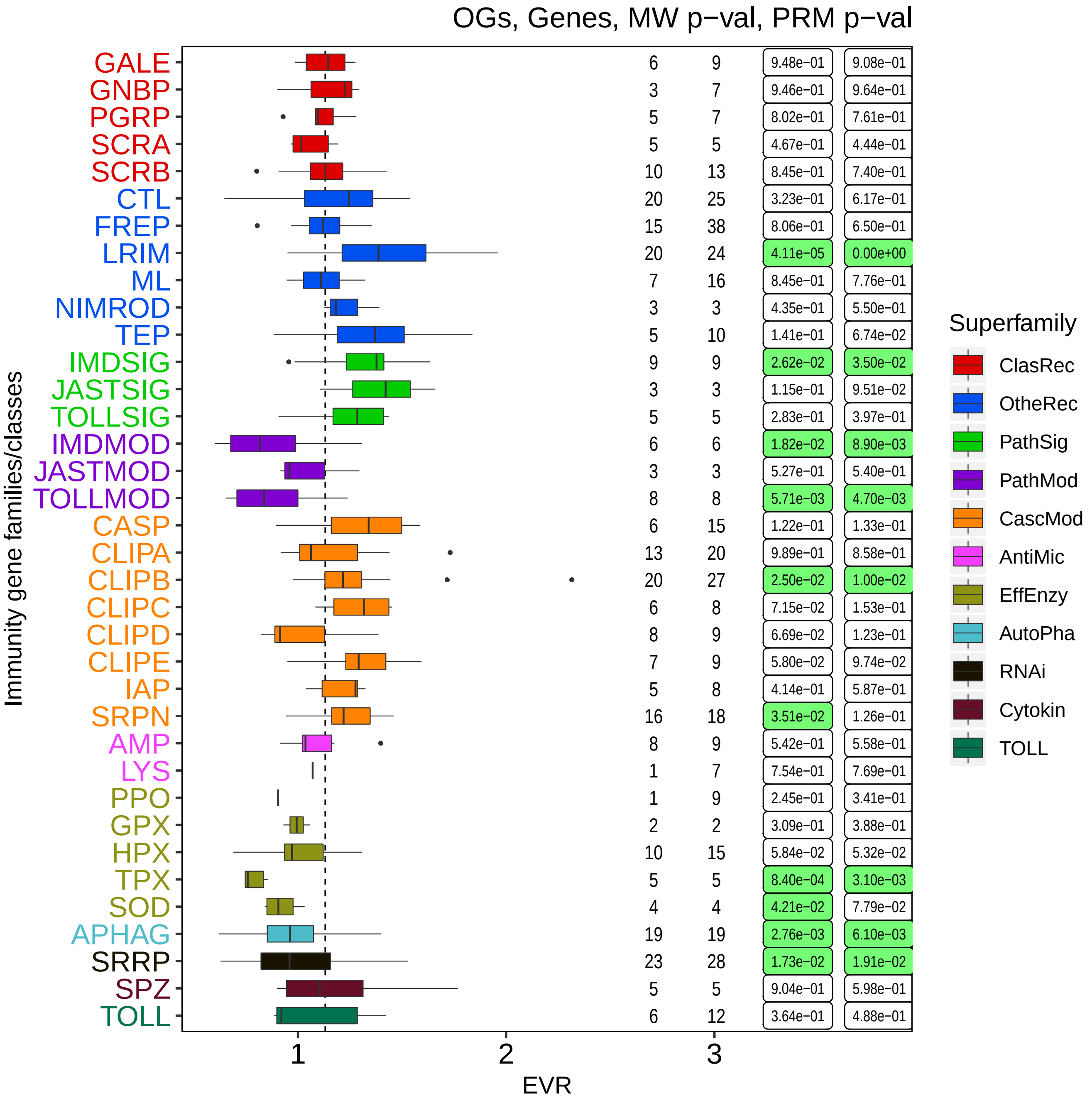

OGs, Genes, MW p−val, PRM p−val
Immunity gene families/classes
GALE 6 9 9.48e-01 9.08e-01
GNBP 3 7 9.46e-01 9.64e-01
PGRP 5 7 8.02e-01 7.61e-01
SCRA 5 5 4.67e-01 4.44e-01
SCRB 10 13 8.45e-01 7.40e-01
CTL 20 25 3.23e-01 6.17e-01
FREP 15 38 8.06e-01 6.50e-01
LRIM 20 24 4.11e-05 0.00e+00
ML 7 16 8.45e-01 7.76e-01
NIMROD 3 3 4.35e-01 5.50e-01
TEP 5 10 1.41e-01 6.74e-02
IMDSIG 9 9 2.62e-02 3.50e-02
JASTSIG 3 3 1.15e-01 9.51e-02
TOLLSIG 5 5 2.83e-01 3.97e-01
IMDMOD 6 6 1.82e-02 8.90e-03
JASTMOD 3 3 5.27e-01 5.40e-01
TOLLMOD 8 8 5.71e-03 4.70e-03
CASP 6 15 1.22e-01 1.33e-01
CLIPA 13 20 9.89e-01 8.58e-01
CLIPB 20 27 2.50e-02 1.00e-02
CLIPC 6 8 7.15e-02 1.53e-01
CLIPD 8 9 6.69e-02 1.23e-01
CLIPE 7 9 5.80e-02 9.74e-02
IAP 5 8 4.14e-01 5.87e-01
SRPN 16 18 3.51e-02 1.26e-01
AMP 8 9 5.42e-01 5.58e-01
LYS 1 7 7.54e-01 7.69e-01
PPO 1 9 2.45e-01 3.41e-01
GPX 2 2 3.09e-01 3.88e-01
HPX 10 15 5.84e-02 5.32e-02
TPX 5 5 8.40e-04 3.10e-03
SOD 4 4 4.21e-02 7.79e-02
APHAG 19 19 2.76e-03 6.10e-03
SRRP 23 28 1.73e-02 1.91e-02
SPZ 5 5 9.04e-01 5.98e-01
TOLL 6 12 3.64e-01 4.88e-01
1
2
3
EVR
Superfamily
ClasRec
OtheRec
PathSig
PathMod
CascMod
AntiMic
EffEnzy
AutoPha
RNAi
Cytokin
TOLL

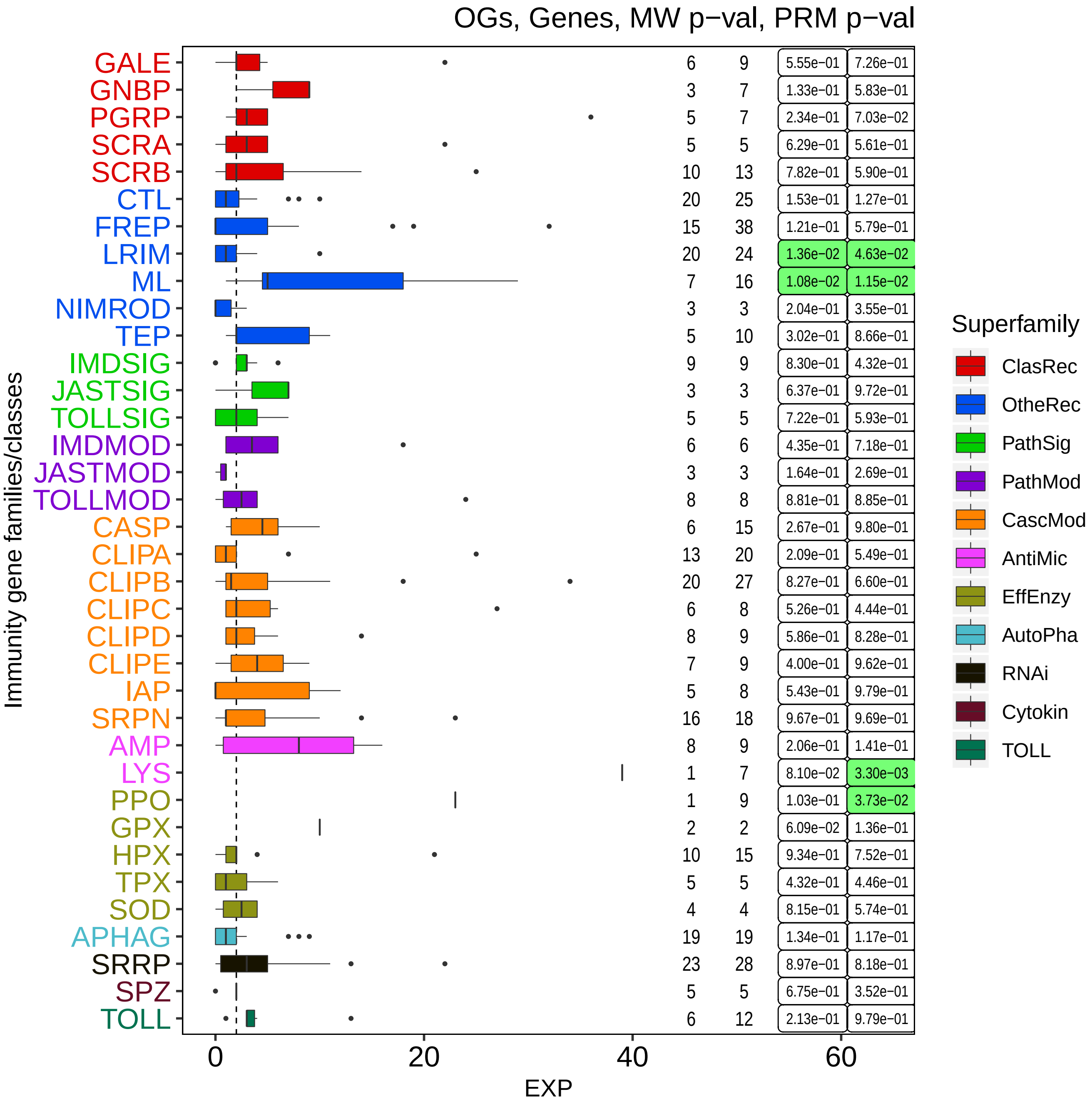
OGs, Genes, MW p−val, PRM p−val
Immunity gene families/classes
GALE 6 9 5.55e-01 7.26e-01
GNBP 3 7 1.33e-01 5.83e-01
PGRP 5 7 2.34e-01 7.03e-02
SCRA 5 5 6.29e-01 5.61e-01
SCRB 10 13 7.82e-01 5.90e-01
CTL 20 25 1.53e-01 1.27e-01
FREP 15 38 1.21e-01 5.79e-01
LRIM 20 24 1.36e-02 4.63e-02
ML 7 16 1.08e-02 1.15e-02
NIMROD 3 3 2.04e-01 3.55e-01
TEP 5 10 3.02e-01 8.66e-01
IMDSIG 9 9 8.30e-01 4.32e-01
JASTSIG 3 3 6.37e-01 9.72e-01
TOLLSIG 5 5 7.22e-01 5.93e-01
IMDMOD 6 6 4.35e-01 7.18e-01
JASTMOD 3 3 1.64e-01 2.69e-01
TOLLMOD 8 8 8.81e-01 8.85e-01
CASP 6 15 2.67e-01 9.80e-01
CLIPA 13 20 2.09e-01 5.49e-01
CLIPB 20 27 8.27e-01 6.60e-01
CLIPC 6 8 5.26e-01 4.44e-01
CLIPD 8 9 5.86e-01 8.28e-01
CLIPE 7 9 4.00e-01 9.62e-01
IAP 5 8 5.43e-01 9.79e-01
SRPN 16 18 9.67e-01 9.69e-01
AMP 8 9 2.06e-01 1.41e-01
LYS 1 7 8.10e-02 3.30e-03
PPO 1 9 1.03e-01 3.73e-02
GPX 2 2 6.09e-02 1.36e-01
HPX 10 15 9.34e-01 7.52e-01
TPX 5 5 4.32e-01 4.46e-01
SOD 4 4 8.15e-01 5.74e-01
APHAG 19 19 1.34e-01 1.17e-01
SRRP 23 28 8.97e-01 8.18e-01
SPZ 5 5 6.75e-01 3.52e-01
TOLL 6 12 2.13e-01 9.79e-01
0
20
40
60
EXP
Superfamily
ClasRec
OtheRec
PathSig
PathMod
CascMod
AntiMic
EffEnzy
AutoPha
RNAi
Cytokin
TOLL

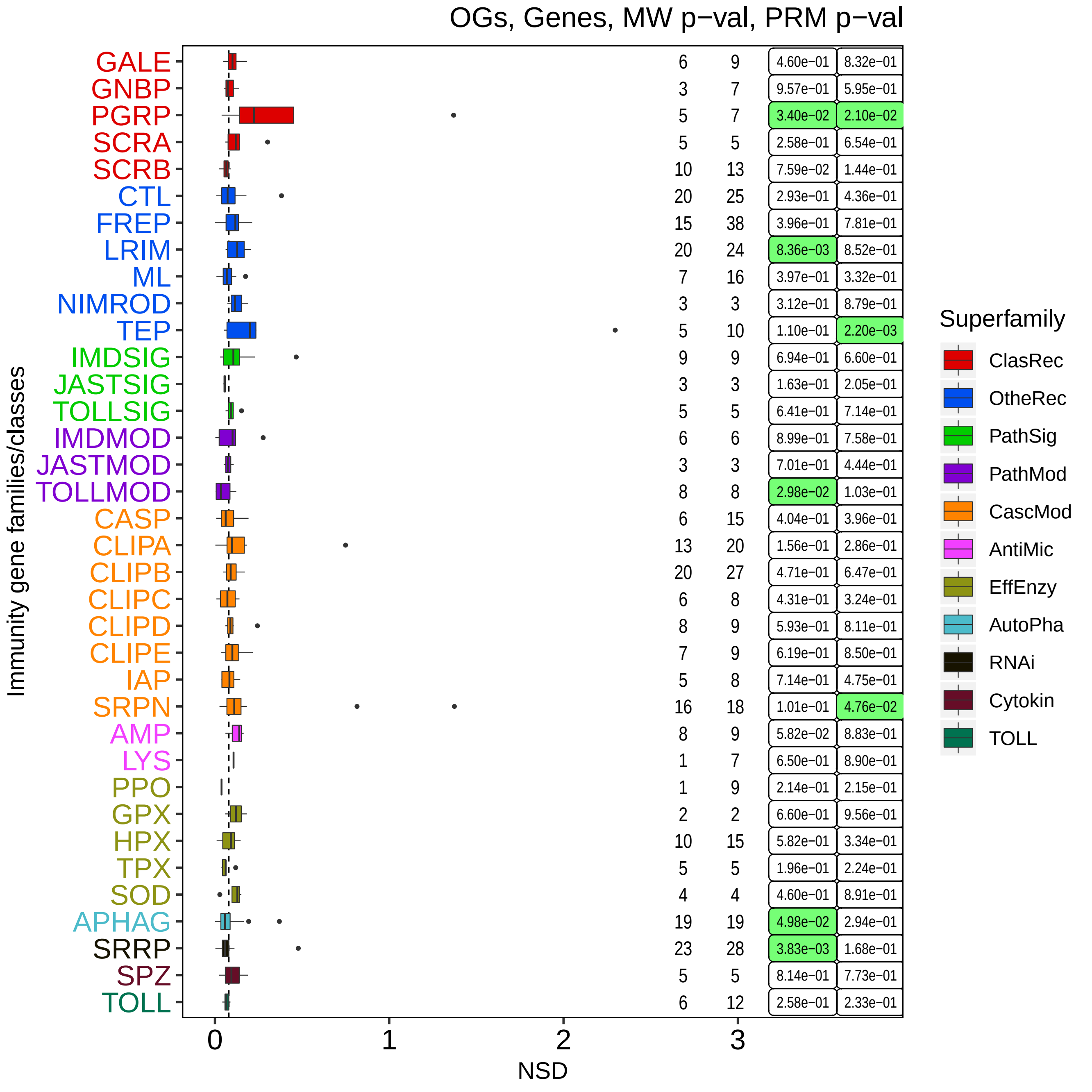

OGs, Genes, MW p−val, PRM p−val
Immunity gene families/classes
GALE 6 9 4.60e-01 8.32e-01
GNBP 3 7 9.57e-01 5.95e-01
PGRP 5 7 3.40e-02 2.10e-02
SCRA 5 5 2.58e-01 6.54e-01
SCRB 10 13 7.59e-02 1.44e-01
CTL 20 25 2.93e-01 4.36e-01
FREP 15 38 3.96e-01 7.81e-01
LRIM 20 24 8.36e-03 8.52e-01
ML 7 16 3.97e-01 3.32e-01
NIMROD 3 3 3.12e-01 8.79e-01
TEP 5 10 1.10e-01 2.20e-03
IMDSIG 9 9 6.94e-01 6.60e-01
JASTSIG 3 3 1.63e-01 2.05e-01
TOLLSIG 5 5 6.41e-01 7.14e-01
IMDMOD 6 6 8.99e-01 7.58e-01
JASTMOD 3 3 7.01e-01 4.44e-01
TOLLMOD 8 8 2.98e-02 1.03e-01
CASP 6 15 4.04e-01 3.96e-01
CLIPA 13 20 1.56e-01 2.86e-01
CLIPB 20 27 4.71e-01 6.47e-01
CLIPC 6 8 4.31e-01 3.24e-01
CLIPD 8 9 5.93e-01 8.11e-01
CLIPE 7 9 6.19e-01 8.50e-01
IAP 5 8 7.14e-01 4.75e-01
SRPN 16 18 1.01e-01 4.76e-02
AMP 8 9 5.82e-02 8.83e-01
LYS 1 7 6.50e-01 8.90e-01
PPO 1 9 2.14e-01 2.15e-01
GPX 2 2 6.60e-01 9.56e-01
HPX 10 15 5.82e-01 3.34e-01
TPX 5 5 1.96e-01 2.24e-01
SOD 4 4 4.60e-01 8.91e-01
APHAG 19 19 4.98e-02 2.94e-01
SRRP 23 28 3.83e-03 1.68e-01
SPZ 5 5 8.14e-01 7.73e-01
TOLL 6 12 2.58e-01 2.33e-01
0
1
2
3
NSD
Superfamily
ClasRec
OtheRec
PathSig
PathMod
CascMod
AntiMic
EffEnzy
AutoPha
RNAi
Cytokin
TOLL

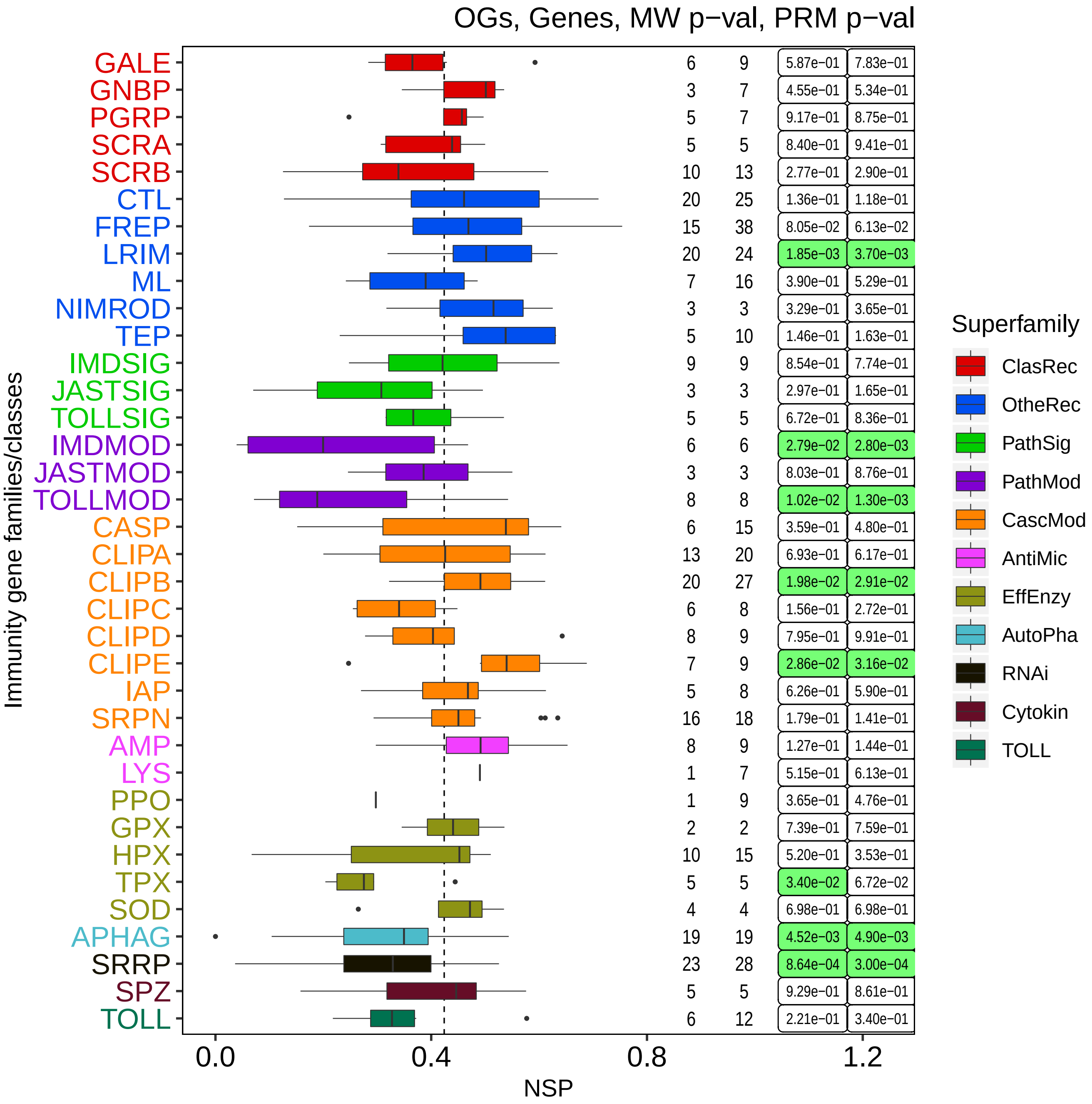

OGs, Genes, MW p−val, PRM p−val
GALE 6 9 5.87e−01 7.83e−01
GNBP 3 7 4.55e−01 5.34e−01
PGRP 5 7 9.17e−01 8.75e−01
SCRA 5 5 8.40e−01 9.41e−01
SCRB 10 13 2.77e−01 2.90e−01
CTL 20 25 1.36e−01 1.18e−01
FREP 15 38 8.05e−02 6.13e−02
LRIM 20 24 1.85e−03 3.70e−03
ML 7 16 3.90e−01 5.29e−01
NIMROD 3 3 3.29e−01 3.65e−01
TEP 5 10 1.46e−01 1.63e−01
IMDSIG 9 9 8.54e−01 7.74e−01
JASTSIG 3 3 2.97e−01 1.65e−01
TOLLSIG 5 5 6.72e−01 8.36e−01
IMDMOD 6 6 2.79e−02 2.80e−03
JASTMOD 3 3 8.03e−01 8.76e−01
TOLLMOD 8 8 1.02e−02 1.30e−03
CASP 6 15 3.59e−01 4.80e−01
CLIPA 13 20 6.93e−01 6.17e−01
CLIPB 20 27 1.98e−02 2.91e−02
CLIPC 6 8 1.56e−01 2.72e−01
CLIPD 8 9 7.95e−01 9.91e−01
CLIPE 7 9 2.86e−02 3.16e−02
IAP 5 8 6.26e−01 5.90e−01
SRPN 16 18 1.79e−01 1.41e−01
AMP 8 9 1.27e−01 1.44e−01
LYS 1 7 5.15e−01 6.13e−01
PPO 1 9 3.65e−01 4.76e−01
GPX 2 2 7.39e−01 7.59e−01
HPX 10 15 5.20e−01 3.53e−01
TPX 5 5 3.40e−02 6.72e−02
SOD 4 4 6.98e−01 6.98e−01
APHAG 19 19 4.52e−03 4.90e−03
SRRP 23 28 8.64e−04 3.00e−04
SPZ 5 5 9.29e−01 8.61e−01
TOLL 6 12 2.21e−01 3.40e−01
Immunity gene families/classes
0.0
0.4
0.8
1.2
NSP
Superfamily
ClasRec
OtheRec
PathSig
PathMod
CascMod
AntiMic
EffEnzy
AutoPha
RNAi
Cytokin
TOLL

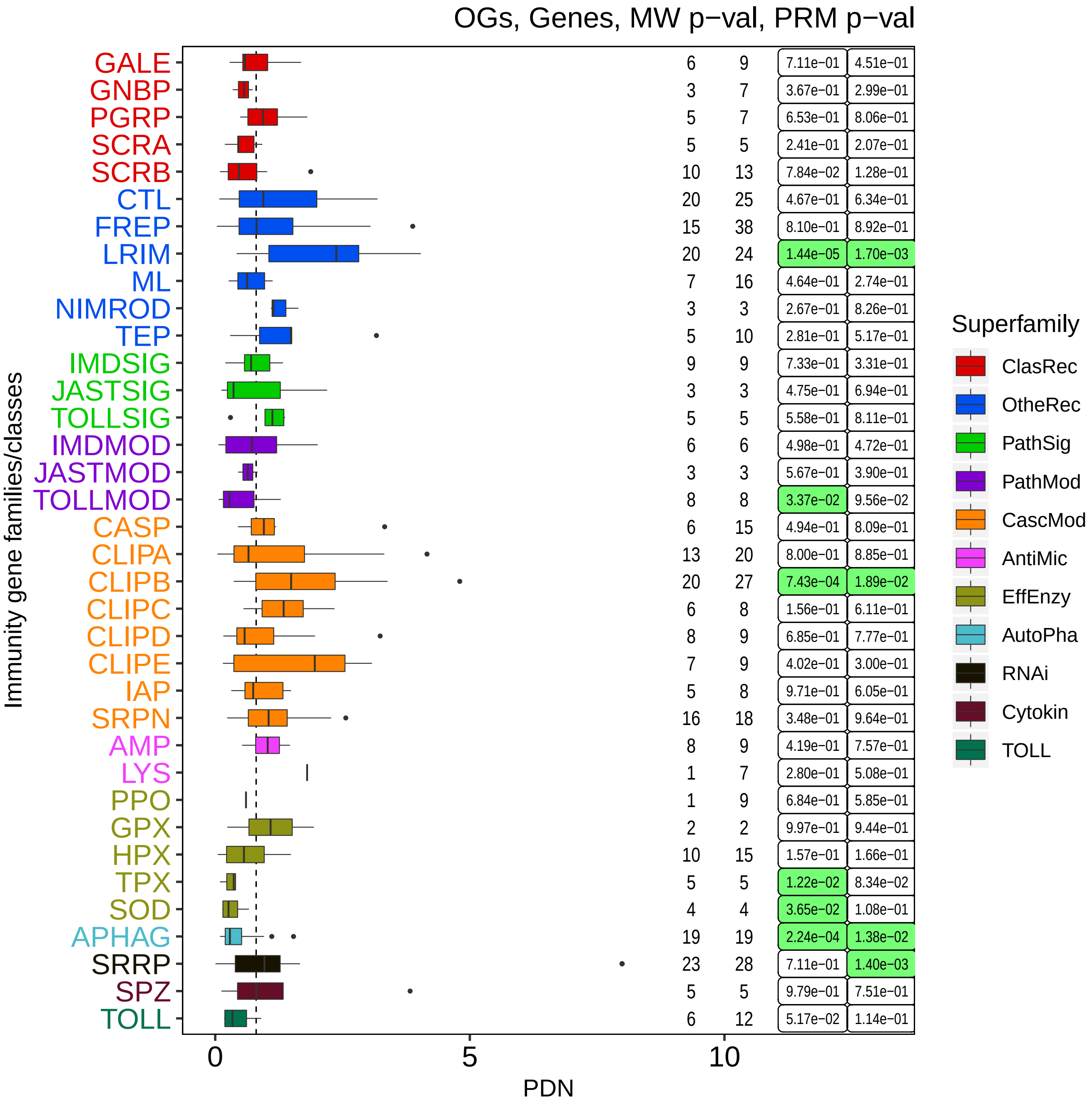

OGs, Genes, MW p−val, PRM p−val
Immunity gene families/classes
PDN
0
5
10
GALE 6 9 7.11e-01 4.51e-01
GNBP 3 7 3.67e-01 2.99e-01
PGRP 5 7 6.53e-01 8.06e-01
SCRA 5 5 2.41e-01 2.07e-01
SCRB 10 13 7.84e-02 1.28e-01
CTL 20 25 4.67e-01 6.34e-01
FREP 15 38 8.10e-01 8.92e-01
LRIM 20 24 1.44e-05 1.70e-03
ML 7 16 4.64e-01 2.74e-01
NIMROD 3 3 2.67e-01 8.26e-01
TEP 5 10 2.81e-01 5.17e-01
IMDSIG 9 9 7.33e-01 3.31e-01
JASTSIG 3 3 4.75e-01 6.94e-01
TOLLSIG 5 5 5.58e-01 8.11e-01
IMDMOD 6 6 4.98e-01 4.72e-01
JASTMOD 3 3 5.67e-01 3.90e-01
TOLLMOD 8 8 3.37e-02 9.56e-02
CASP 6 15 4.94e-01 8.09e-01
CLIPA 13 20 8.00e-01 8.85e-01
CLIPB 20 27 7.43e-04 1.89e-02
CLIPC 6 8 1.56e-01 6.11e-01
CLIPD 8 9 6.85e-01 7.77e-01
CLIPE 7 9 4.02e-01 3.00e-01
IAP 5 8 9.71e-01 6.05e-01
SRPN 16 18 3.48e-01 9.64e-01
AMP 8 9 4.19e-01 7.57e-01
LYS 1 7 2.80e-01 5.08e-01
PPO 1 9 6.84e-01 5.85e-01
GPX 2 2 9.97e-01 9.44e-01
HPX 10 15 1.57e-01 1.66e-01
TPX 5 5 1.22e-02 8.34e-02
SOD 4 4 3.65e-02 1.08e-01
APHAG 19 19 2.24e-04 1.38e-02
SRRP 23 28 7.11e-01 1.40e-03
SPZ 5 5 9.79e-01 7.51e-01
TOLL 6 12 5.17e-02 1.14e-01
Superfamily
ClasRec
OtheRec
PathSig
PathMod
CascMod
AntiMic
EffEnzy
AutoPha
RNAi
Cytokin
TOLL

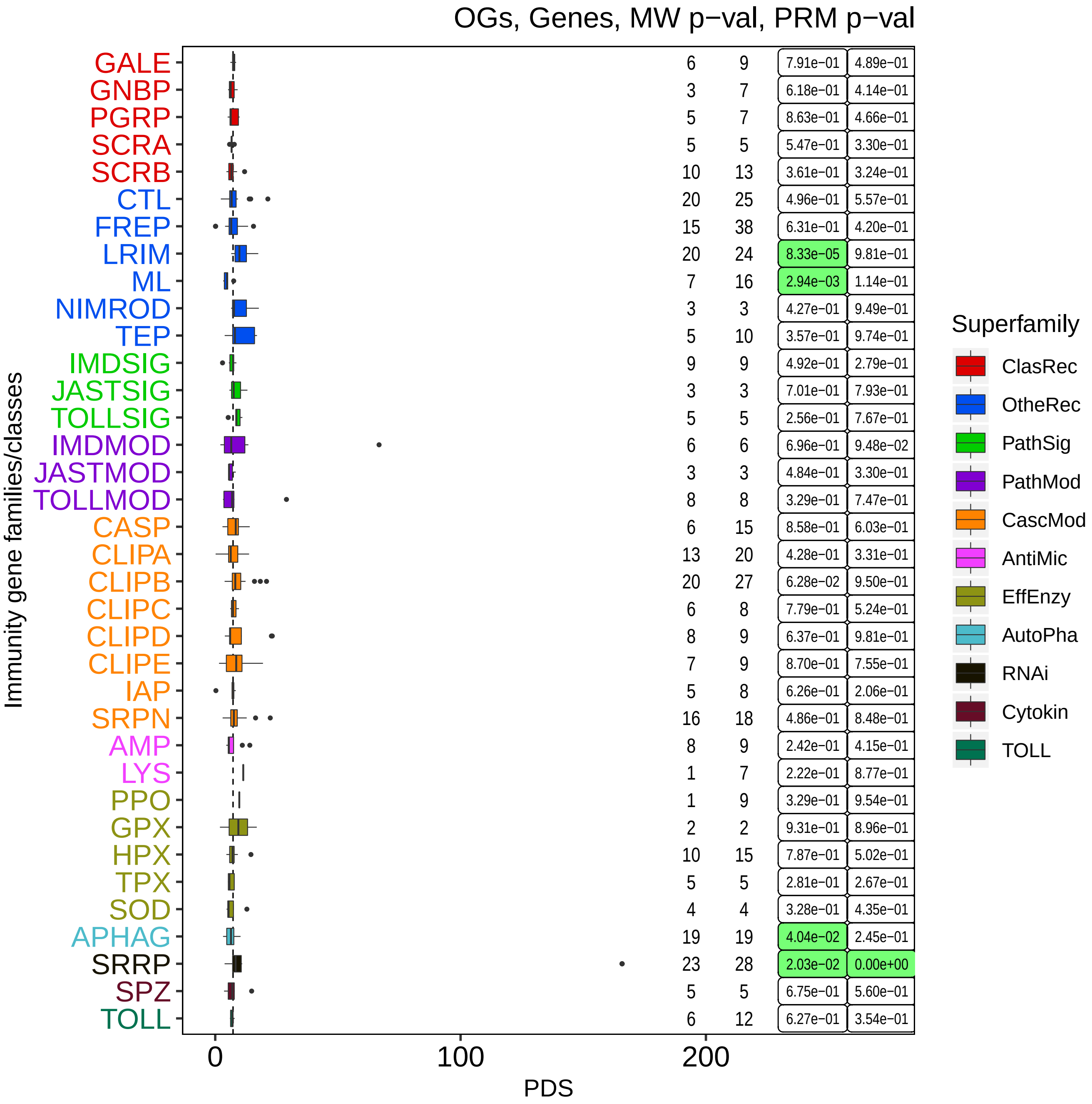

OGs, Genes, MW p−val, PRM p−val
Immunity gene families/classes
PDS
0
100
200
GALE 6 9 7.91e-01 4.89e-01
GNBP 3 7 6.18e-01 4.14e-01
PGRP 5 7 8.63e-01 4.66e-01
SCRA 5 5 5.47e-01 3.30e-01
SCRB 10 13 3.61e-01 3.24e-01
CTL 20 25 4.96e-01 5.57e-01
FREP 15 38 6.31e-01 4.20e-01
LRIM 20 24 8.33e-05 9.81e-01
ML 7 16 2.94e-03 1.14e-01
NIMROD 3 3 4.27e-01 9.49e-01
TEP 5 10 3.57e-01 9.74e-01
IMDSIG 9 9 4.92e-01 2.79e-01
JASTSIG 3 3 7.01e-01 7.93e-01
TOLLSIG 5 5 2.56e-01 7.67e-01
IMDMOD 6 6 6.96e-01 9.48e-02
JASTMOD 3 3 4.84e-01 3.30e-01
TOLLMOD 8 8 3.29e-01 7.47e-01
CASP 6 15 8.58e-01 6.03e-01
CLIPA 13 20 4.28e-01 3.31e-01
CLIPB 20 27 6.28e-02 9.50e-01
CLIPC 6 8 7.79e-01 5.24e-01
CLIPD 8 9 6.37e-01 9.81e-01
CLIPE 7 9 8.70e-01 7.55e-01
IAP 5 8 6.26e-01 2.06e-01
SRPN 16 18 4.86e-01 8.48e-01
AMP 8 9 2.42e-01 4.15e-01
LYS 1 7 2.22e-01 8.77e-01
PPO 1 9 3.29e-01 9.54e-01
GPX 2 2 9.31e-01 8.96e-01
HPX 10 15 7.87e-01 5.02e-01
TPX 5 5 2.81e-01 2.67e-01
SOD 4 4 3.28e-01 4.35e-01
APHAG 19 19 4.04e-02 2.45e-01
SRRP 23 28 2.03e-02 0.00e+00
SPZ 5 5 6.75e-01 5.60e-01
TOLL 6 12 6.27e-01 3.54e-01
Superfamily
ClasRec
OtheRec
PathSig
PathMod
CascMod
AntiMic
EffEnzy
AutoPha
RNAi
Cytokin
TOLL

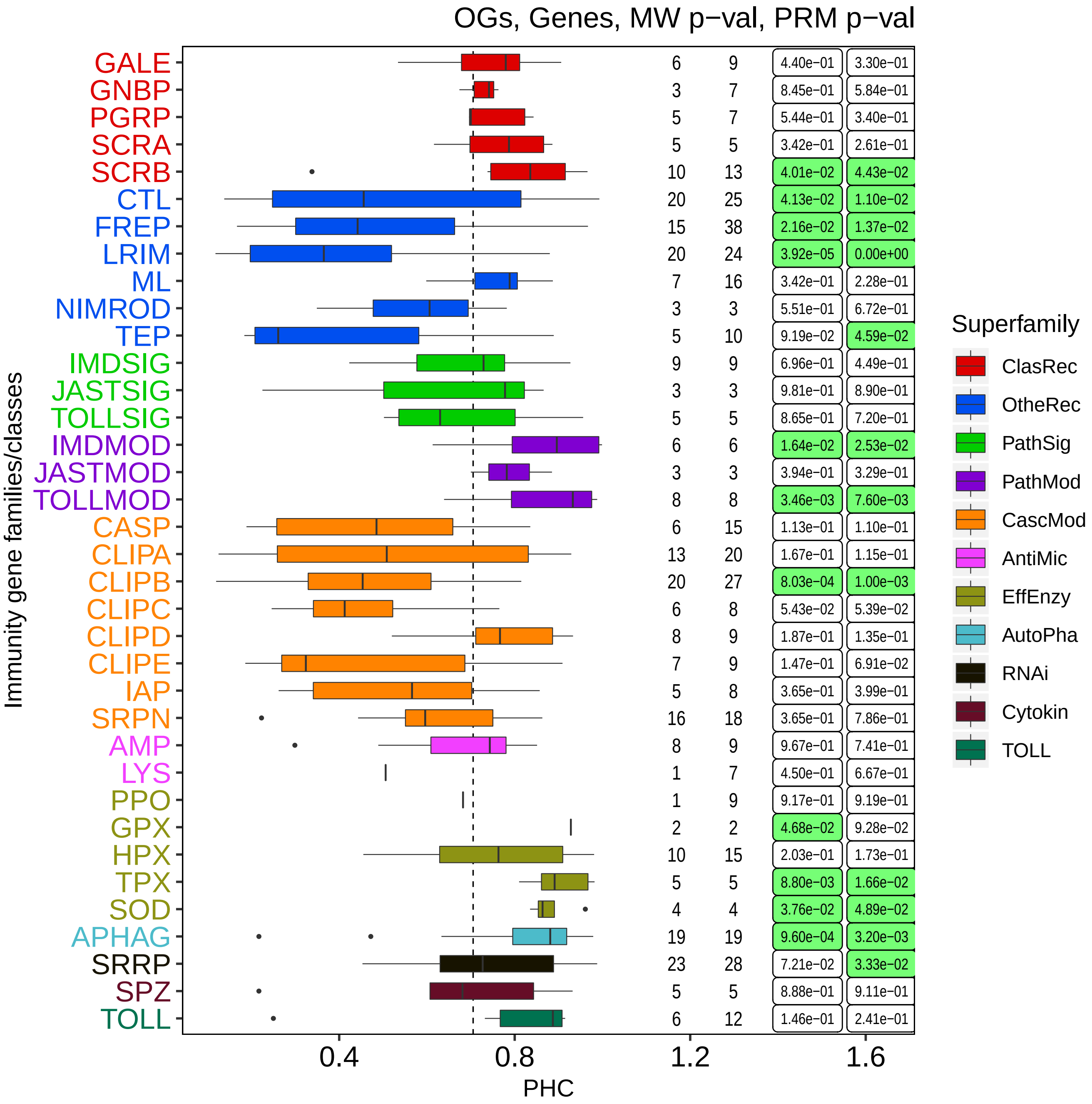

OGs, Genes, MW p−val, PRM p−val
Immunity gene families/classes
GALE 6 9 4.40e−01 3.30e−01
GNBP 3 7 8.45e−01 5.84e−01
PGRP 5 7 5.44e−01 3.40e−01
SCRA 5 5 3.42e−01 2.61e−01
SCRB 10 13 4.01e−02 4.43e−02
CTL 20 25 4.13e−02 1.10e−02
FREP 15 38 2.16e−02 1.37e−02
LRIM 20 24 3.92e−05 0.00e+00
ML 7 16 3.42e−01 2.28e−01
NIMROD 3 3 5.51e−01 6.72e−01
TEP 5 10 9.19e−02 4.59e−02
IMDSIG 9 9 6.96e−01 4.49e−01
JASTSIG 3 3 9.81e−01 8.90e−01
TOLLSIG 5 5 8.65e−01 7.20e−01
IMDMOD 6 6 1.64e−02 2.53e−02
JASTMOD 3 3 3.94e−01 3.29e−01
TOLLMOD 8 8 3.46e−03 7.60e−03
CASP 6 15 1.13e−01 1.10e−01
CLIPA 13 20 1.67e−01 1.15e−01
CLIPB 20 27 8.03e−04 1.00e−03
CLIPC 6 8 5.43e−02 5.39e−02
CLIPD 8 9 1.87e−01 1.35e−01
CLIPE 7 9 1.47e−01 6.91e−02
IAP 5 8 3.65e−01 3.99e−01
SRPN 16 18 3.65e−01 7.86e−01
AMP 8 9 9.67e−01 7.41e−01
LYS 1 7 4.50e−01 6.67e−01
PPO 1 9 9.17e−01 9.19e−01
GPX 2 2 4.68e−02 9.28e−02
HPX 10 15 2.03e−01 1.73e−01
TPX 5 5 8.80e−03 1.66e−02
SOD 4 4 3.76e−02 4.89e−02
APHAG 19 19 9.60e−04 3.20e−03
SRRP 23 28 7.21e−02 3.33e−02
SPZ 5 5 8.88e−01 9.11e−01
TOLL 6 12 1.46e−01 2.41e−01
0.4
0.8
1.2
1.6
PHC
Superfamily
ClasRec
OtheRec
PathSig
PathMod
CascMod
AntiMic
EffEnzy
AutoPha
RNAi
Cytokin
TOLL

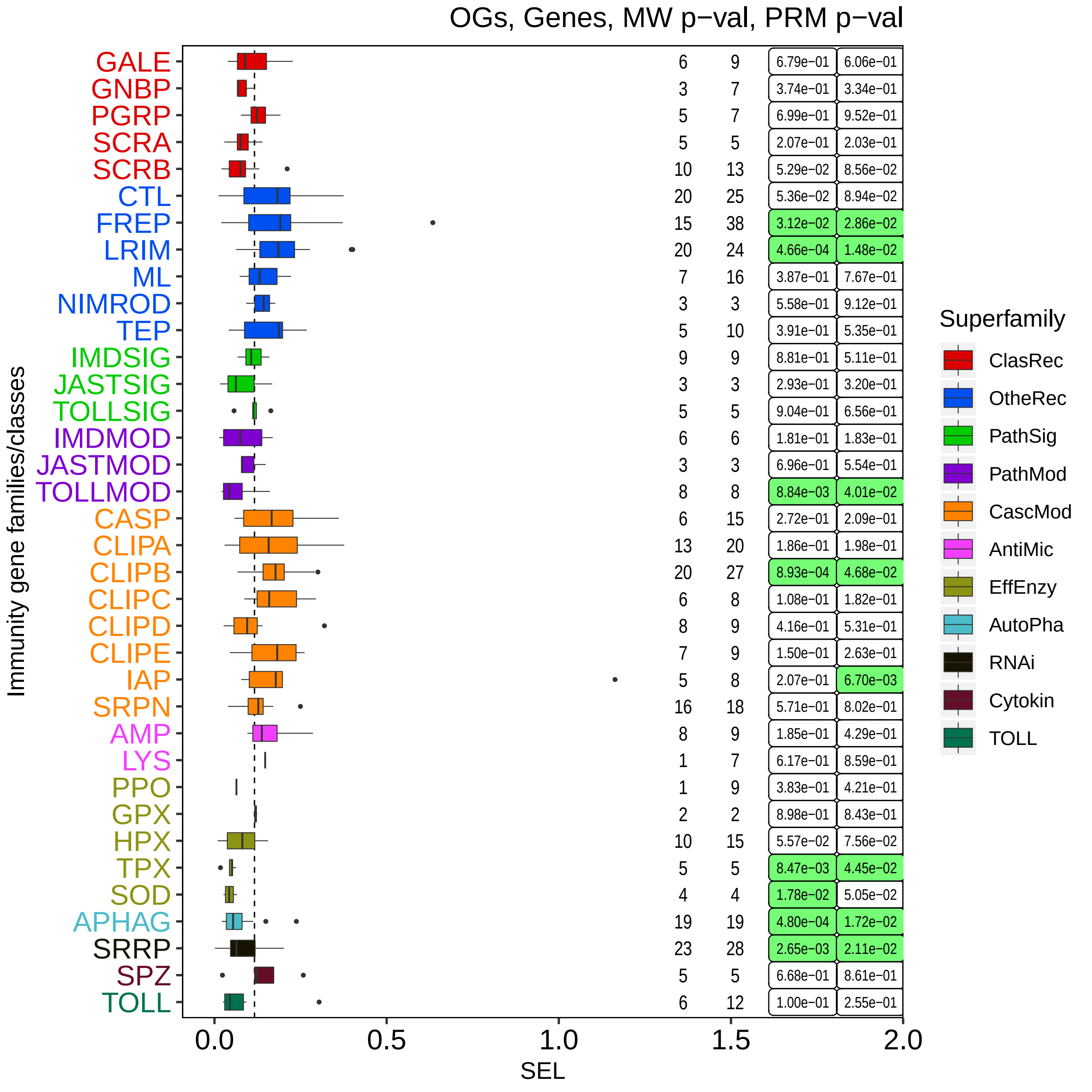

OGs, Genes, MW p−val, PRM p−val
Immunity gene families/classes
GALE 6 9 6.79e-01 6.06e-01
GNBP 3 7 3.74e-01 3.34e-01
PGRP 5 7 6.99e-01 9.52e-01
SCRA 5 5 2.07e-01 2.03e-01
SCRB 10 13 5.29e-02 8.56e-02
CTL 20 25 5.36e-02 8.94e-02
FREP 15 38 3.12e-02 2.86e-02
LRIM 20 24 4.66e-04 1.48e-02
ML 7 16 3.87e-01 7.67e-01
NIMROD 3 3 5.58e-01 9.12e-01
TEP 5 10 3.91e-01 5.35e-01
IMDSIG 9 9 8.81e-01 5.11e-01
JASTSIG 3 3 2.93e-01 3.20e-01
TOLLSIG 5 5 9.04e-01 6.56e-01
IMDMOD 6 6 1.81e-01 1.83e-01
JASTMOD 3 3 6.96e-01 5.54e-01
TOLLMOD 8 8 8.84e-03 4.01e-02
CASP 6 15 2.72e-01 2.09e-01
CLIPA 13 20 1.86e-01 1.98e-01
CLIPB 20 27 8.93e-04 4.68e-02
CLIPC 6 8 1.08e-01 1.82e-01
CLIPD 8 9 4.16e-01 5.31e-01
CLIPE 7 9 1.50e-01 2.63e-01
IAP 5 8 2.07e-01 6.70e-03
SRPN 16 18 5.71e-01 8.02e-01
AMP 8 9 1.85e-01 4.29e-01
LYS 1 7 6.17e-01 8.59e-01
PPO 1 9 3.83e-01 4.21e-01
GPX 2 2 8.98e-01 8.43e-01
HPX 10 15 5.57e-02 7.56e-02
TPX 5 5 8.47e-03 4.45e-02
SOD 4 4 1.78e-02 5.05e-02
APHAG 19 19 4.80e-04 1.72e-02
SRRP 23 28 2.65e-03 2.11e-02
SPZ 5 5 6.68e-01 8.61e-01
TOLL 6 12 1.00e-01 2.55e-01
0.0
0.5
1.0
1.5
2.0
SEL
Superfamily
ClasRec
OtheRec
PathSig
PathMod
CascMod
AntiMic
EffEnzy
AutoPha
RNAi
Cytokin
TOLL

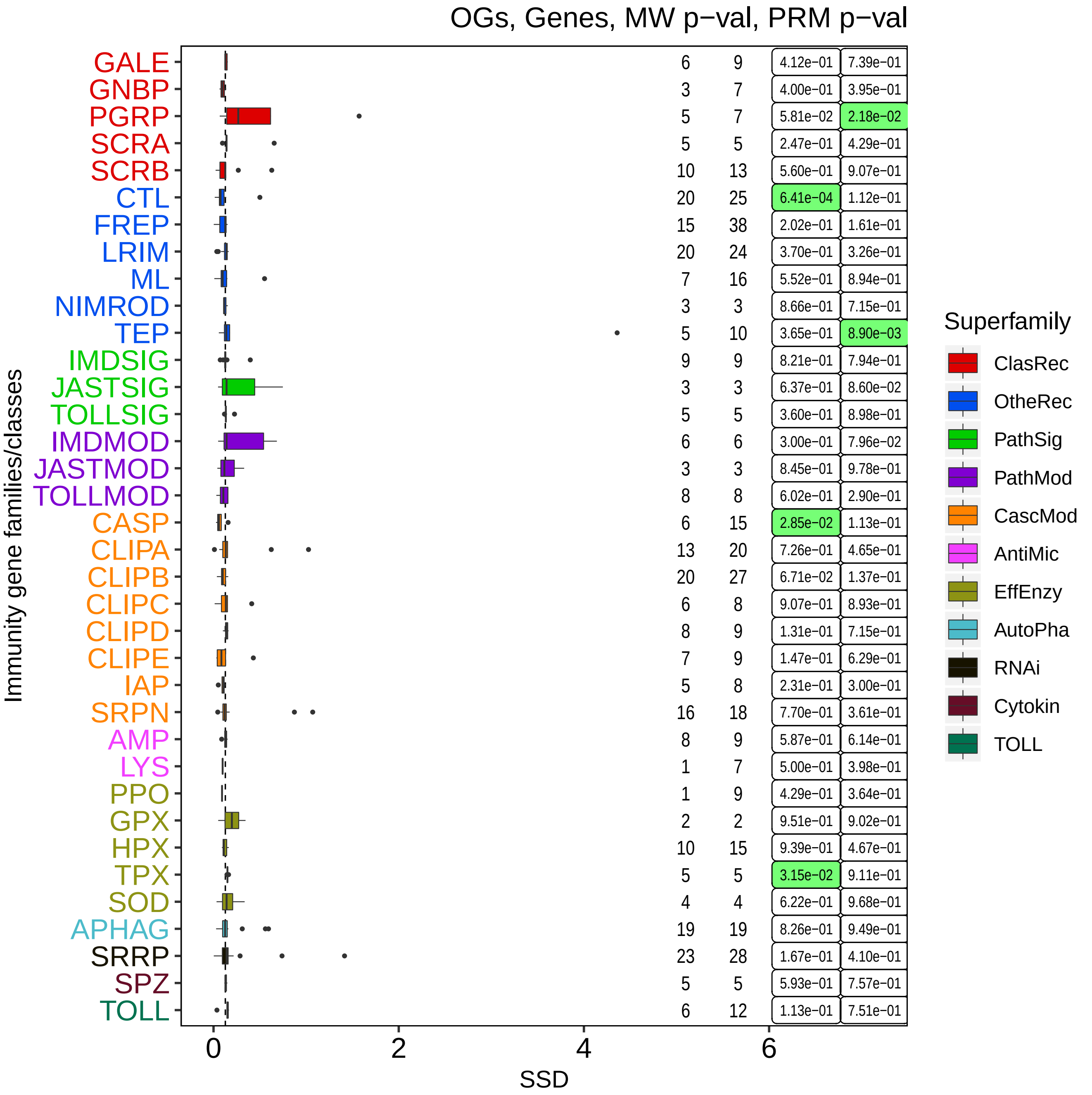
OGs, Genes, MW p−val, PRM p−val
Immunity gene families/classes
SSD
0
2
4
6
GALE 6 9 4.12e-01 7.39e-01
GNBP 3 7 4.00e-01 3.95e-01
PGRP 5 7 5.81e-02 2.18e-02
SCRA 5 5 2.47e-01 4.29e-01
SCRB 10 13 5.60e-01 9.07e-01
CTL 20 25 6.41e-04 1.12e-01
FREP 15 38 2.02e-01 1.61e-01
LRIM 20 24 3.70e-01 3.26e-01
ML 7 16 5.52e-01 8.94e-01
NIMROD 3 3 8.66e-01 7.15e-01
TEP 5 10 3.65e-01 8.90e-03
IMDSIG 9 9 8.21e-01 7.94e-01
JASTSIG 3 3 6.37e-01 8.60e-02
TOLLSIG 5 5 3.60e-01 8.98e-01
IMDMOD 6 6 3.00e-01 7.96e-02
JASTMOD 3 3 8.45e-01 9.78e-01
TOLLMOD 8 8 6.02e-01 2.90e-01
CASP 6 15 2.85e-02 1.13e-01
CLIPA 13 20 7.26e-01 4.65e-01
CLIPB 20 27 6.71e-02 1.37e-01
CLIPC 6 8 9.07e-01 8.93e-01
CLIPD 8 9 1.31e-01 7.15e-01
CLIPE 7 9 1.47e-01 6.29e-01
IAP 5 8 2.31e-01 3.00e-01
SRPN 16 18 7.70e-01 3.61e-01
AMP 8 9 5.87e-01 6.14e-01
LYS 1 7 5.00e-01 3.98e-01
PPO 1 9 4.29e-01 3.64e-01
GPX 2 2 9.51e-01 9.02e-01
HPX 10 15 9.39e-01 4.67e-01
TPX 5 5 3.15e-02 9.11e-01
SOD 4 4 6.22e-01 9.68e-01
APHAG 19 19 8.26e-01 9.49e-01
SRRP 23 28 1.67e-01 4.10e-01
SPZ 5 5 5.93e-01 7.57e-01
TOLL 6 12 1.13e-01 7.51e-01
Superfamily
ClasRec
OtheRec
PathSig
PathMod
CascMod
AntiMic
EffEnzy
AutoPha
RNAi
Cytokin
TOLL

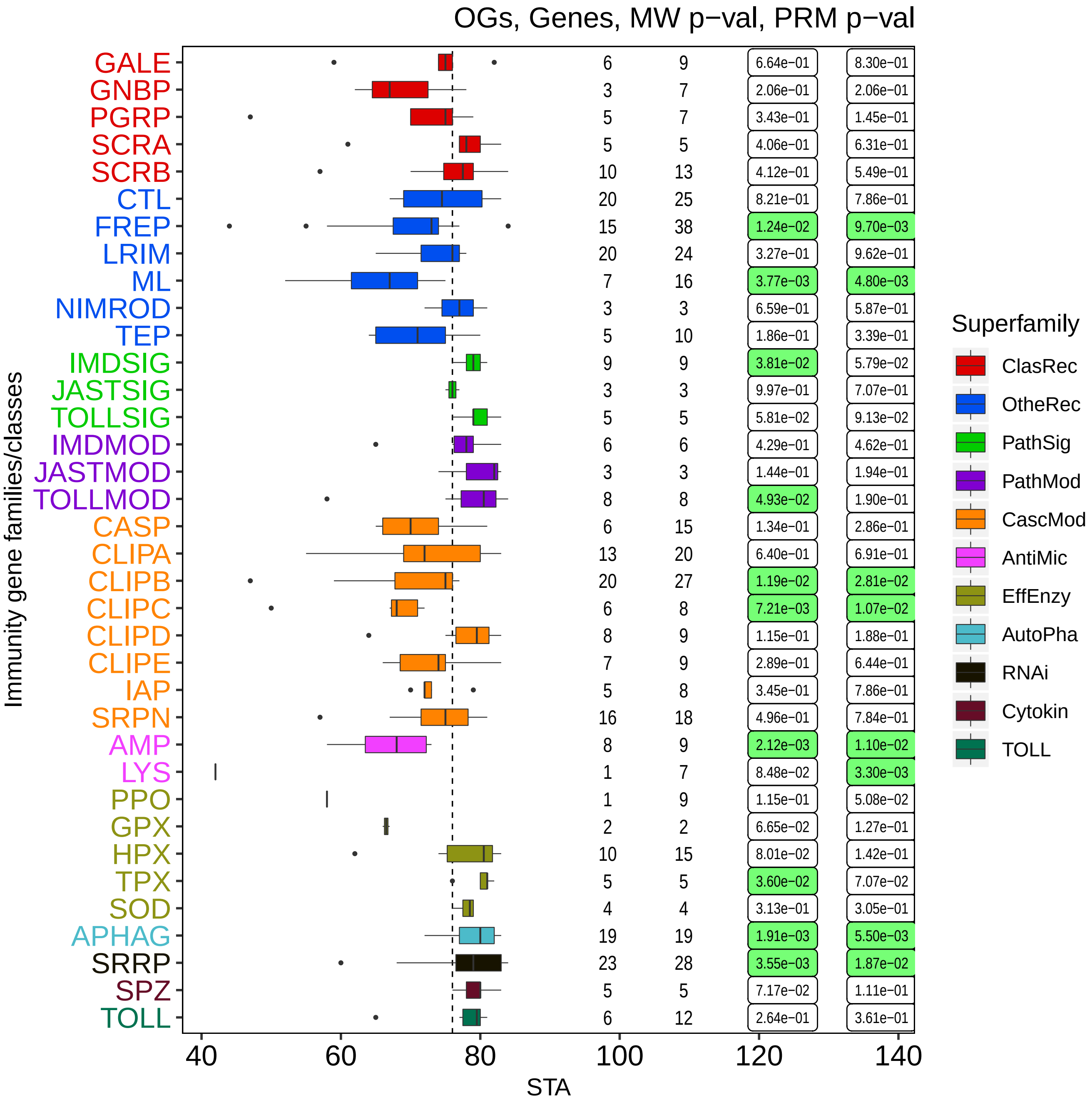
OGs, Genes, MW p−val, PRM p−val
Immunity gene families/classes
GALE 6 9 6.64e-01 8.30e-01
GNBP 3 7 2.06e-01 2.06e-01
PGRP 5 7 3.43e-01 1.45e-01
SCRA 5 5 4.06e-01 6.31e-01
SCRB 10 13 4.12e-01 5.49e-01
CTL 20 25 8.21e-01 7.86e-01
FREP 15 38 1.24e-02 9.70e-03
LRIM 20 24 3.27e-01 9.62e-01
ML 7 16 3.77e-03 4.80e-03
NIMROD 3 3 6.59e-01 5.87e-01
TEP 5 10 1.86e-01 3.39e-01
IMDSIG 9 9 3.81e-02 5.79e-02
JASTSIG 3 3 9.97e-01 7.07e-01
TOLLSIG 5 5 5.81e-02 9.13e-02
IMDMOD 6 6 4.29e-01 4.62e-01
JASTMOD 3 3 1.44e-01 1.94e-01
TOLLMOD 8 8 4.93e-02 1.90e-01
CASP 6 15 1.34e-01 2.86e-01
CLIPA 13 20 6.40e-01 6.91e-01
CLIPB 20 27 1.19e-02 2.81e-02
CLIPC 6 8 7.21e-03 1.07e-02
CLIPD 8 9 1.15e-01 1.88e-01
CLIPE 7 9 2.89e-01 6.44e-01
IAP 5 8 3.45e-01 7.86e-01
SRPN 16 18 4.96e-01 7.84e-01
AMP 8 9 2.12e-03 1.10e-02
LYS 1 7 8.48e-02 3.30e-03
PPO 1 9 1.15e-01 5.08e-02
GPX 2 2 6.65e-02 1.27e-01
HPX 10 15 8.01e-02 1.42e-01
TPX 5 5 3.60e-02 7.07e-02
SOD 4 4 3.13e-01 3.05e-01
APHAG 19 19 1.91e-03 5.50e-03
SRRP 23 28 3.55e-03 1.87e-02
SPZ 5 5 7.17e-02 1.11e-01
TOLL 6 12 2.64e-01 3.61e-01
40 60 80 100 120 140
STA
Superfamily
ClasRec
OtheRec
PathSig
PathMod
CascMod
AntiMic
EffEnzy
AutoPha
RNAi
Cytokin
TOLL

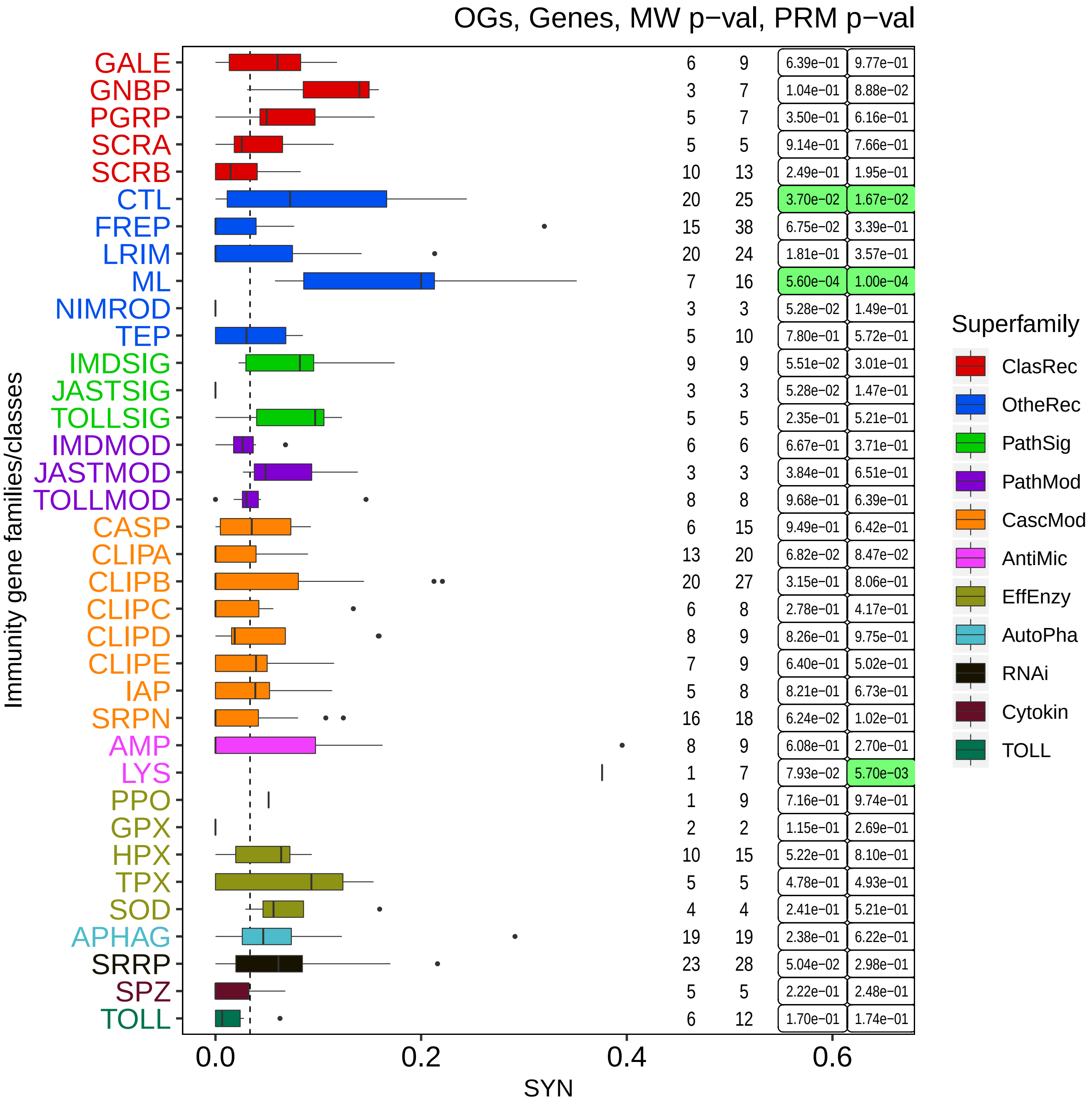

OGs, Genes, MW p−val, PRM p−val
Immunity gene families/classes
GALE 6 9 6.39e-01 9.77e-01
GNBP 3 7 1.04e-01 8.88e-02
PGRP 5 7 3.50e-01 6.16e-01
SCRA 5 5 9.14e-01 7.66e-01
SCRB 10 13 2.49e-01 1.95e-01
CTL 20 25 3.70e-02 1.67e-02
FREP 15 38 6.75e-02 3.39e-01
LRIM 20 24 1.81e-01 3.57e-01
ML 7 16 5.60e-04 1.00e-04
NIMROD 3 3 5.28e-02 1.49e-01
TEP 5 10 7.80e-01 5.72e-01
IMDSIG 9 9 5.51e-02 3.01e-01
JASTSIG 3 3 5.28e-02 1.47e-01
TOLLSIG 5 5 2.35e-01 5.21e-01
IMDMOD 6 6 6.67e-01 3.71e-01
JASTMOD 3 3 3.84e-01 6.51e-01
TOLLMOD 8 8 9.68e-01 6.39e-01
CASP 6 15 9.49e-01 6.42e-01
CLIPA 13 20 6.82e-02 8.47e-02
CLIPB 20 27 3.15e-01 8.06e-01
CLIPC 6 8 2.78e-01 4.17e-01
CLIPD 8 9 8.26e-01 9.75e-01
CLIPE 7 9 6.40e-01 5.02e-01
IAP 5 8 8.21e-01 6.73e-01
SRPN 16 18 6.24e-02 1.02e-01
AMP 8 9 6.08e-01 2.70e-01
LYS 1 7 7.93e-02 5.70e-03
PPO 1 9 7.16e-01 9.74e-01
GPX 2 2 1.15e-01 2.69e-01
HPX 10 15 5.22e-01 8.10e-01
TPX 5 5 4.78e-01 4.93e-01
SOD 4 4 2.41e-01 5.21e-01
APHAG 19 19 2.38e-01 6.22e-01
SRRP 23 28 5.04e-02 2.98e-01
SPZ 5 5 2.22e-01 2.48e-01
TOLL 6 12 1.70e-01 1.74e-01
0.0
0.2
0.4
0.6
SYN
Superfamily
ClasRec
OtheRec
PathSig
PathMod
CascMod
AntiMic
EffEnzy
AutoPha
RNAi
Cytokin
TOLL

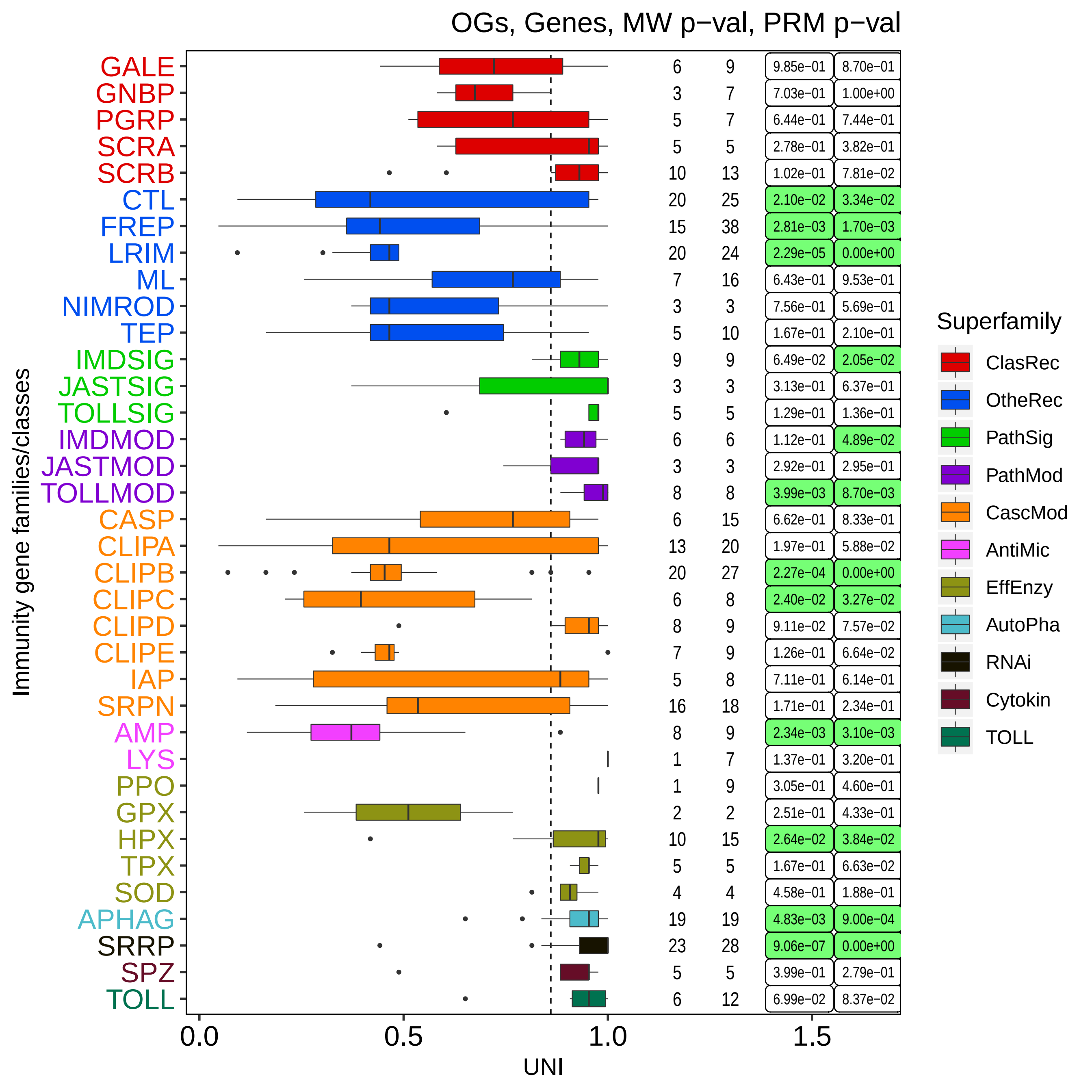

OGs, Genes, MW p−val, PRM p−val
Immunity gene families/classes
GALE
GNBP
PGRP
SCRA
SCRB
CTL
FREP
LRIM
ML
NIMROD
TEP
IMDSIG
JASTSIG
TOLLSIG
IMDMOD
JASTMOD
TOLLMOD
CASP
CLIPA
CLIPB
CLIPC
CLIPD
CLIPE
IAP
SRPN
AMP
LYS
PPO
GPX
HPX
TPX
SOD
APHAG
SRRP
SPZ
TOLL
6 9 9.85e−01 8.70e−01
3 7 7.03e−01 1.00e+00
5 7 6.44e−01 7.44e−01
5 5 2.78e−01 3.82e−01
10 13 1.02e−01 7.81e−02
20 25 2.10e−02 3.34e−02
15 38 2.81e−03 1.70e−03
20 24 2.29e−05 0.00e+00
7 16 6.43e−01 9.53e−01
3 3 7.56e−01 5.69e−01
5 10 1.67e−01 2.10e−01
9 9 6.49e−02 2.05e−02
3 3 3.13e−01 6.37e−01
5 5 1.29e−01 1.36e−01
6 6 1.12e−01 4.89e−02
3 3 2.92e−01 2.95e−01
8 8 3.99e−03 8.70e−03
6 15 6.62e−01 8.33e−01
13 20 1.97e−01 5.88e−02
20 27 2.27e−04 0.00e+00
6 8 2.40e−02 3.27e−02
8 9 9.11e−02 7.57e−02
7 9 1.26e−01 6.64e−02
5 8 7.11e−01 6.14e−01
16 18 1.71e−01 2.34e−01
8 9 2.34e−03 3.10e−03
1 7 1.37e−01 3.20e−01
1 9 3.05e−01 4.60e−01
2 2 2.51e−01 4.33e−01
10 15 2.64e−02 3.84e−02
5 5 1.67e−01 6.63e−02
4 4 4.58e−01 1.88e−01
19 19 4.83e−03 9.00e−04
23 28 9.06e−07 0.00e+00
5 5 3.99e−01 2.79e−01
6 12 6.99e−02 8.37e−02
0.0
0.5
1.0
1.5
UNI
Superfamily
ClasRec
OtheRec
PathSig
PathMod
CascMod
AntiMic
EffEnzy
AutoPha
RNAi
Cytokin
TOLL

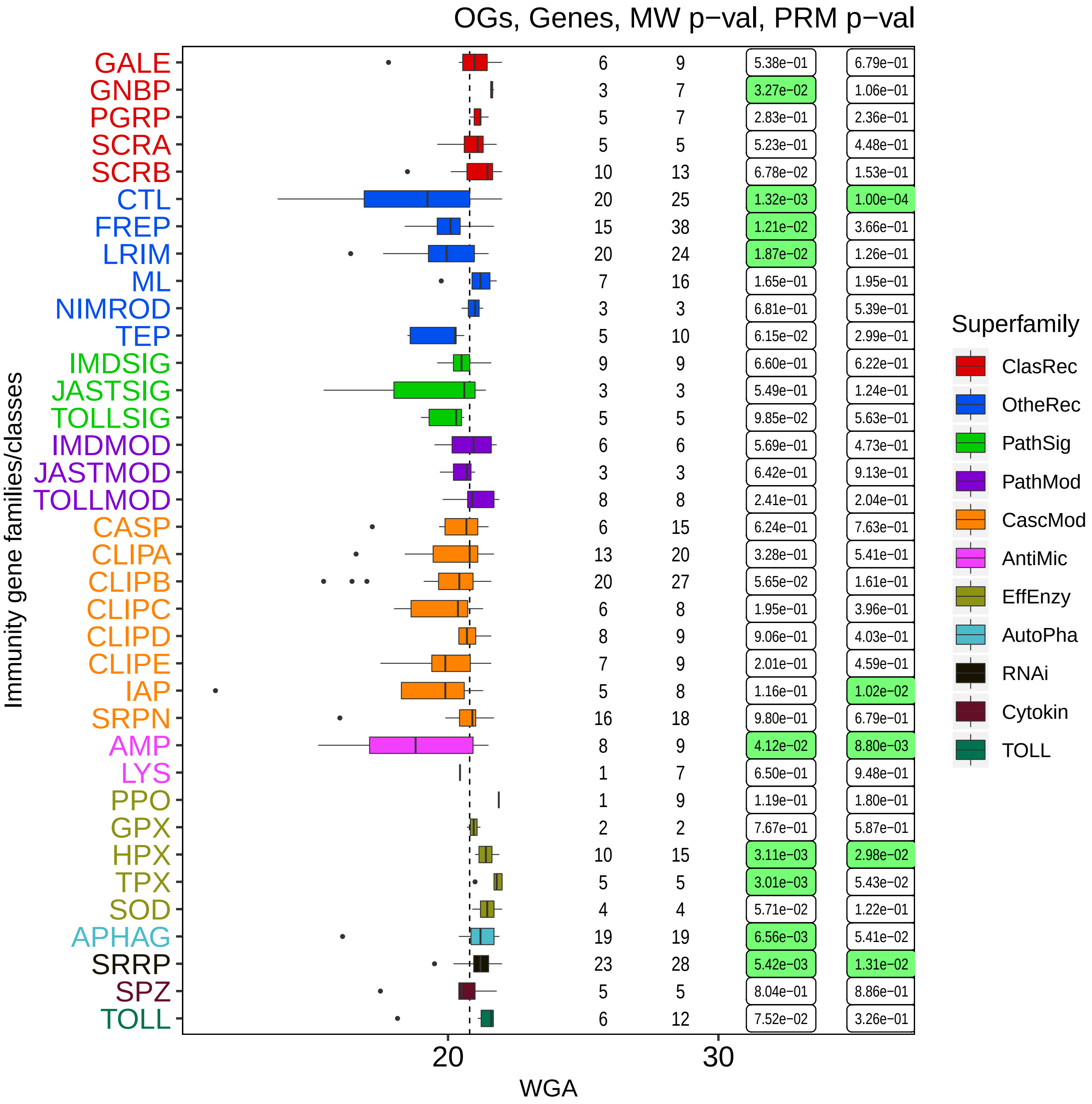

OGs, Genes, MW p−val, PRM p−val
Immunity gene families/classes
GALE 6 9 5.38e-01 6.79e-01
GNBP 3 7 3.27e-02 1.06e-01
PGRP 5 7 2.83e-01 2.36e-01
SCRA 5 5 5.23e-01 4.48e-01
SCRB 10 13 6.78e-02 1.53e-01
CTL 20 25 1.32e-03 1.00e-04
FREP 15 38 1.21e-02 3.66e-01
LRIM 20 24 1.87e-02 1.26e-01
ML 7 16 1.65e-01 1.95e-01
NIMROD 3 3 6.81e-01 5.39e-01
TEP 5 10 6.15e-02 2.99e-01
IMDSIG 9 9 6.60e-01 6.22e-01
JASTSIG 3 3 5.49e-01 1.24e-01
TOLLSIG 5 5 9.85e-02 5.63e-01
IMDMOD 6 6 5.69e-01 4.73e-01
JASTMOD 3 3 6.42e-01 9.13e-01
TOLLMOD 8 8 2.41e-01 2.04e-01
CASP 6 15 6.24e-01 7.63e-01
CLIPA 13 20 3.28e-01 5.41e-01
CLIPB 20 27 5.65e-02 1.61e-01
CLIPC 6 8 1.95e-01 3.96e-01
CLIPD 8 9 9.06e-01 4.03e-01
CLIPE 7 9 2.01e-01 4.59e-01
IAP 5 8 1.16e-01 1.02e-02
SRPN 16 18 9.80e-01 6.79e-01
AMP 8 9 4.12e-02 8.80e-03
LYS 1 7 6.50e-01 9.48e-01
PPO 1 9 1.19e-01 1.80e-01
GPX 2 2 7.67e-01 5.87e-01
HPX 10 15 3.11e-03 2.98e-02
TPX 5 5 3.01e-03 5.43e-02
SOD 4 4 5.71e-02 1.22e-01
APHAG 19 19 6.56e-03 5.41e-02
SRRP 23 28 5.42e-03 1.31e-02
SPZ 5 5 8.04e-01 8.86e-01
TOLL 6 12 7.52e-02 3.26e-01
20
30
WGA
Superfamily
ClasRec
OtheRec
PathSig
PathMod
CascMod
AntiMic
EffEnzy
AutoPha
RNAi
Cytokin
TOLL
