## Additional File 2 for "Functional constraints on insect immune system components govern their evolutionary trajectories"

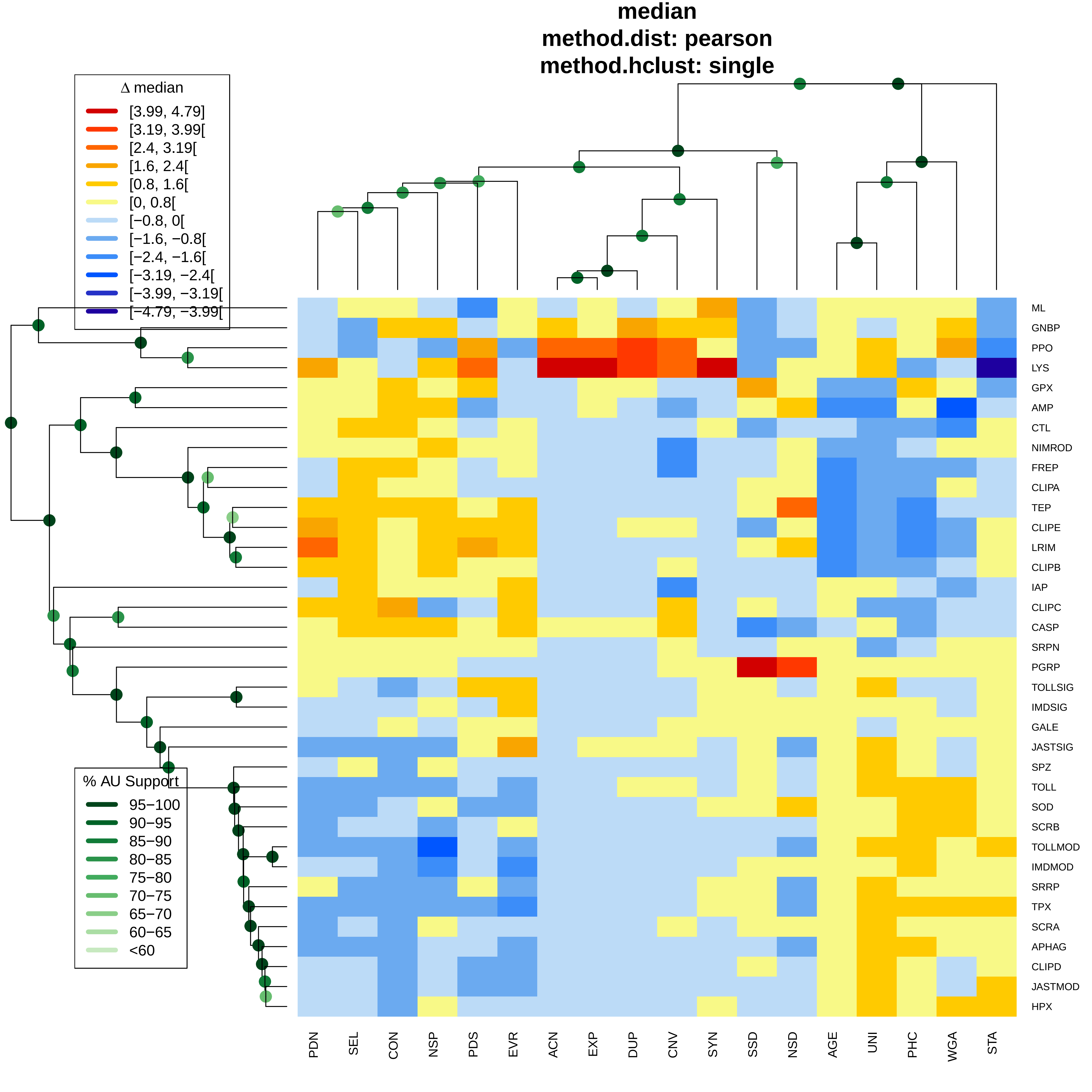
median
method.dist: pearson
method.hclust: single
Δ median
[3.99, 4.79]
[3.19, 3.99[
[2.4, 3.19[
[1.6, 2.4[
[0.8, 1.6[
[0, 0.8[
[−0.8, 0[
[−1.6, −0.8[
[−2.4, −1.6[
[−3.19, −2.4[
[−3.99, −3.19[
[−4.79, −3.99[
% AU Support
95−100
90−95
85−90
80−85
75−80
70−75
65−70
60−65
<60
ML
GNBP
PPO
LYS
GPX
AMP
CTL
NIMROD
FREP
CLIPA
TEP
CLIPE
LRIM
CLIPB
IAP
CLIPC
CASP
SRPN
PGRP
TOLLSIG
IMDSIG
GALE
JASTSIG
SPZ
TOLL
SOD
SCRB
TOLLMOD
IMDMOD
SRRP
TPX
SCRA
APHAG
CLIPD
JASTMOD
HPX
PDN
SEL
CON
NSP
PDS
EVR
ACN
EXP
DUP
CNV
SYN
SSD
NSD
AGE
UNI
PHC
WGA
STA

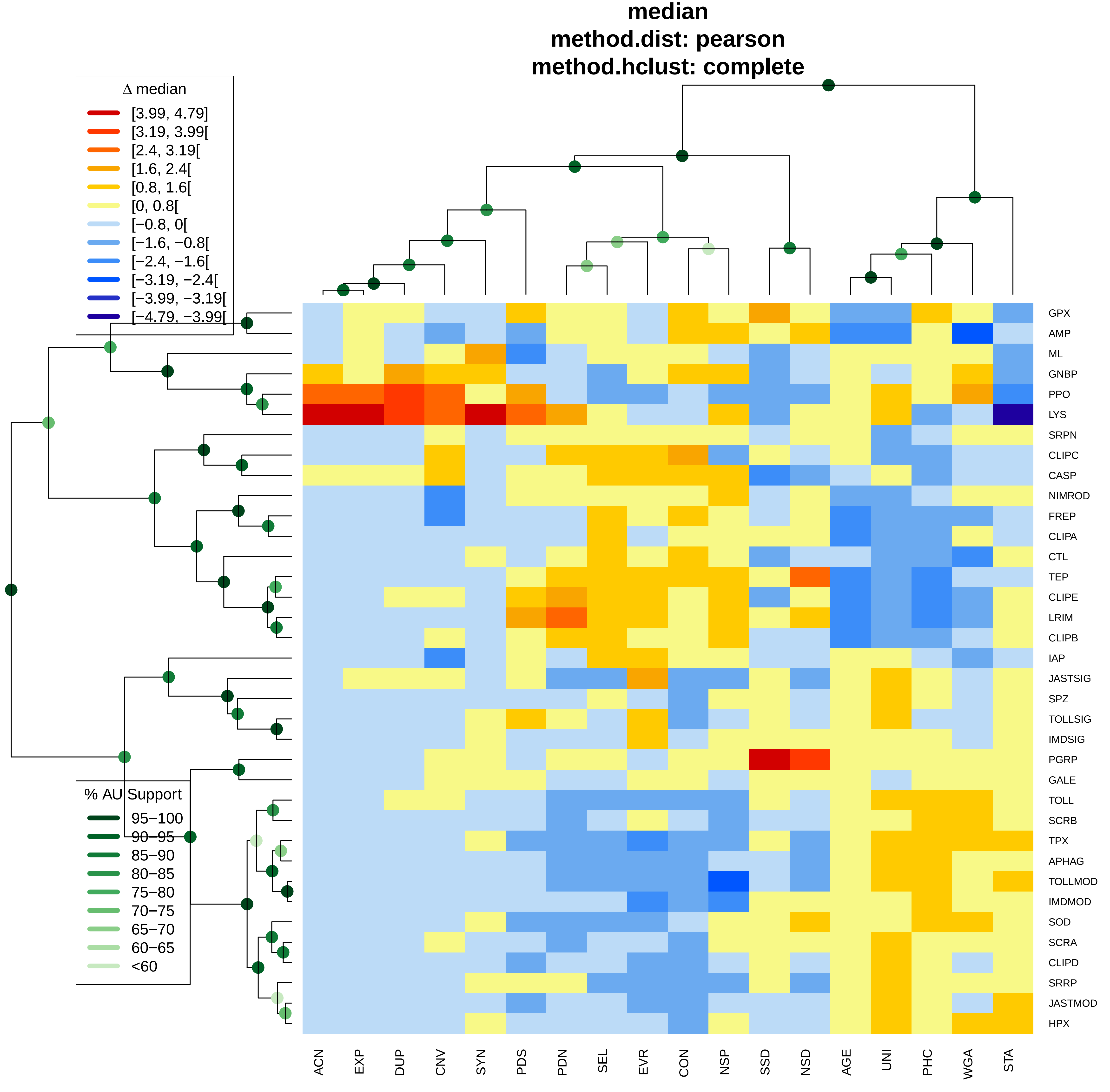
median
method.dist: pearson
method.hclust: complete
Δ median
[3.99, 4.79]
[3.19, 3.99[
[2.4, 3.19[
[1.6, 2.4[
[0.8, 1.6[
[0, 0.8[
[−0.8, 0[
[−1.6, −0.8[
[−2.4, −1.6[
[−3.19, −2.4[
[−3.99, −3.19[
[−4.79, −3.99[
% AU Support
95−100
90−95
85−90
80−85
75−80
70−75
65−70
60−65
<60
GPX
AMP
ML
GNBP
PPO
LYS
SRPN
CLIPC
CASP
NIMROD
FREP
CLIPA
CTL
TEP
CLIPE
LRIM
CLIPB
IAP
JASTSIG
SPZ
TOLLSIG
IMDSIG
PGRP
GALE
TOLL
SCRB
TPX
APHAG
TOLLMOD
IMDMOD
SOD
SCRA
CLIPD
SRRP
JASTMOD
HPX
ACN
EXP
DUP
CNV
SYN
PDS
PDN
SEL
EVR
CON
NSP
SSD
NSD
AGE
UNI
PHC
WGA
STA

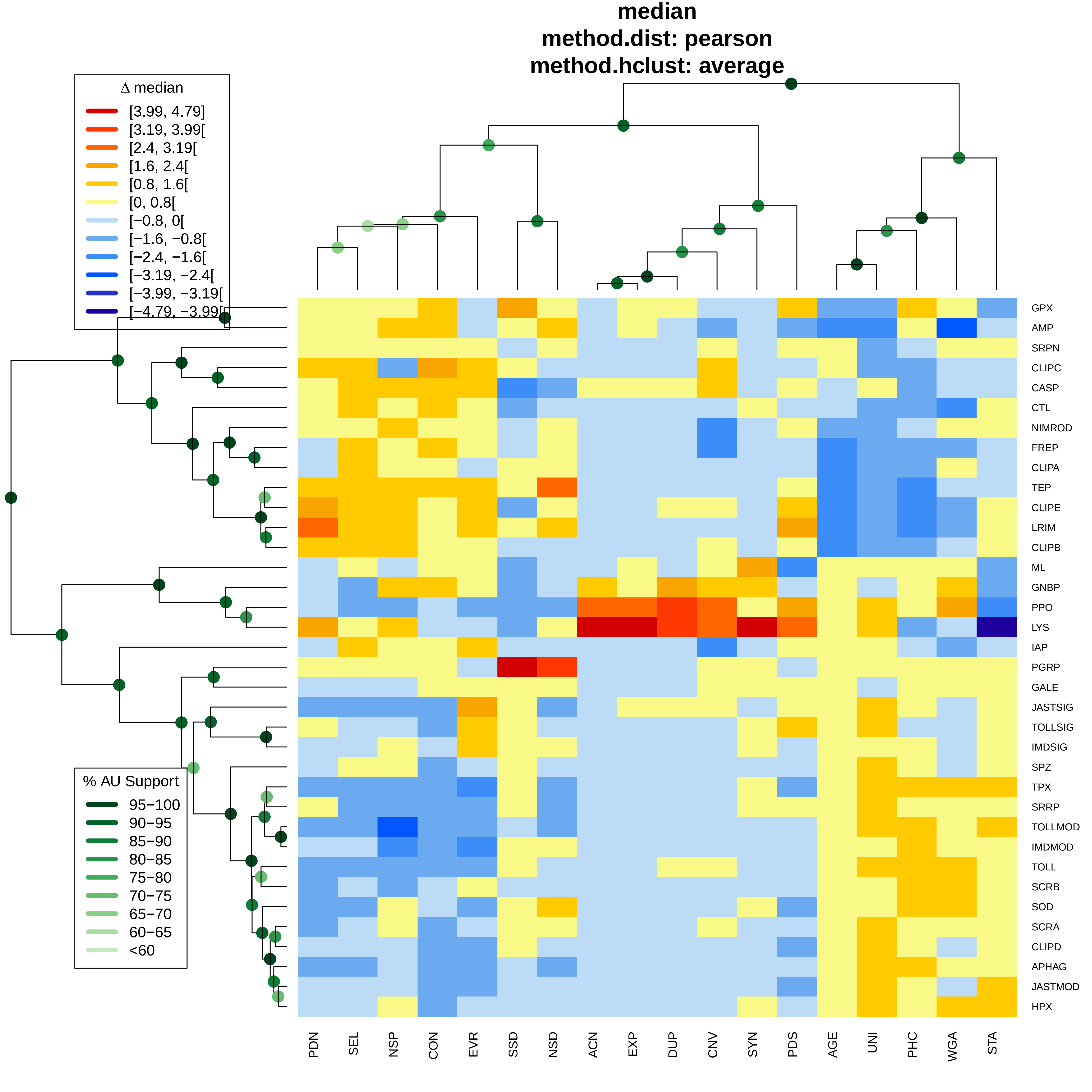
median
method.dist: pearson
method.hclust: average
Δ median
[3.99, 4.79]
[3.19, 3.99[
[2.4, 3.19[
[1.6, 2.4[
[0.8, 1.6[
[0, 0.8[
[−0.8, 0[
[−1.6, −0.8[
[−2.4, −1.6[
[−3.19, −2.4[
[−3.99, −3.19[
[−4.79, −3.99[
% AU Support
95−100
90−95
85−90
80−85
75−80
70−75
65−70
60−65
<60
GPX
AMP
SRPN
CLIPC
CASP
CTL
NIMROD
FREP
CLIPA
TEP
CLIPE
LRIM
CLIPB
ML
GNBP
PPO
LYS
IAP
PGRP
GALE
JASTSIG
TOLLSIG
IMDSIG
SPZ
TPX
SRRP
TOLLMOD
IMDMOD
TOLL
SCRB
SOD
SCRA
CLIPD
APHAG
JASTMOD
HPX
PDN
SEL
NSP
CON
EVR
SSD
NSD
ACN
EXP
DUP
CNV
SYN
PDS
AGE
UNI
PHC
WGA
STA

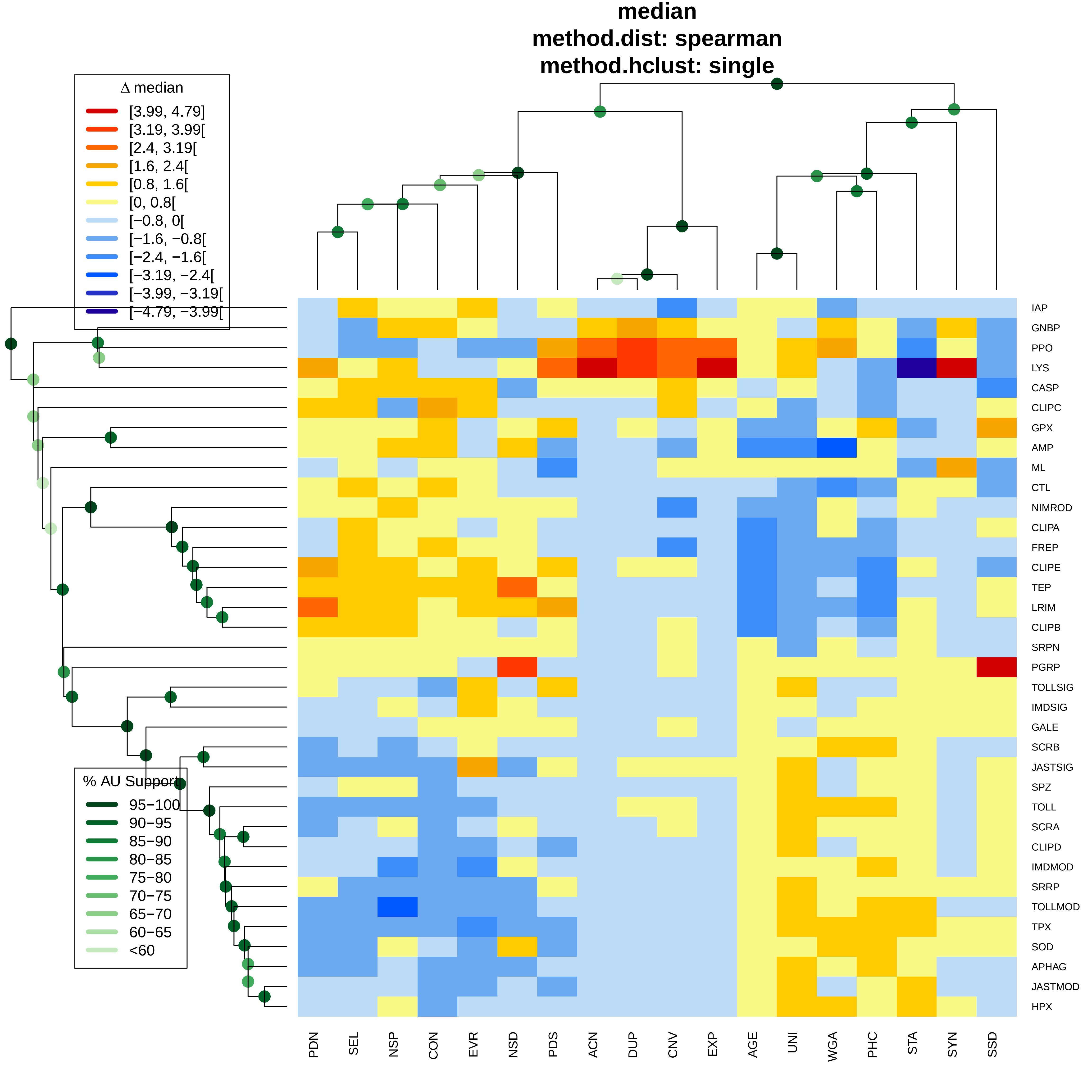
median
method.dist: spearman
method.hclust: single
Δ median
[3.99, 4.79]
[3.19, 3.99[
[2.4, 3.19[
[1.6, 2.4[
[0.8, 1.6[
[0, 0.8[
[−0.8, 0[
[−1.6, −0.8[
[−2.4, −1.6[
[−3.19, −2.4[
[−3.99, −3.19[
[−4.79, −3.99[
% AU Support
95−100
90−95
85−90
80−85
75−80
70−75
65−70
60−65
<60
IAP
GNBP
PPO
LYS
CASP
CLIPC
GPX
AMP
ML
CTL
NIMROD
CLIPA
FREP
CLIPE
TEP
LRIM
CLIPB
SRPN
PGRP
TOLLSIG
IMDSIG
GALE
SCRB
JASTSIG
SPZ
TOLL
SCRA
CLIPD
IMDMOD
SRRP
TOLLMOD
TPX
SOD
APHAG
JASTMOD
HPX
PDN
SEL
NSP
CON
EVR
NSD
PDS
ACN
DUP
CNV
EXP
AGE
UNI
WGA
PHC
STA
SYN
SSD

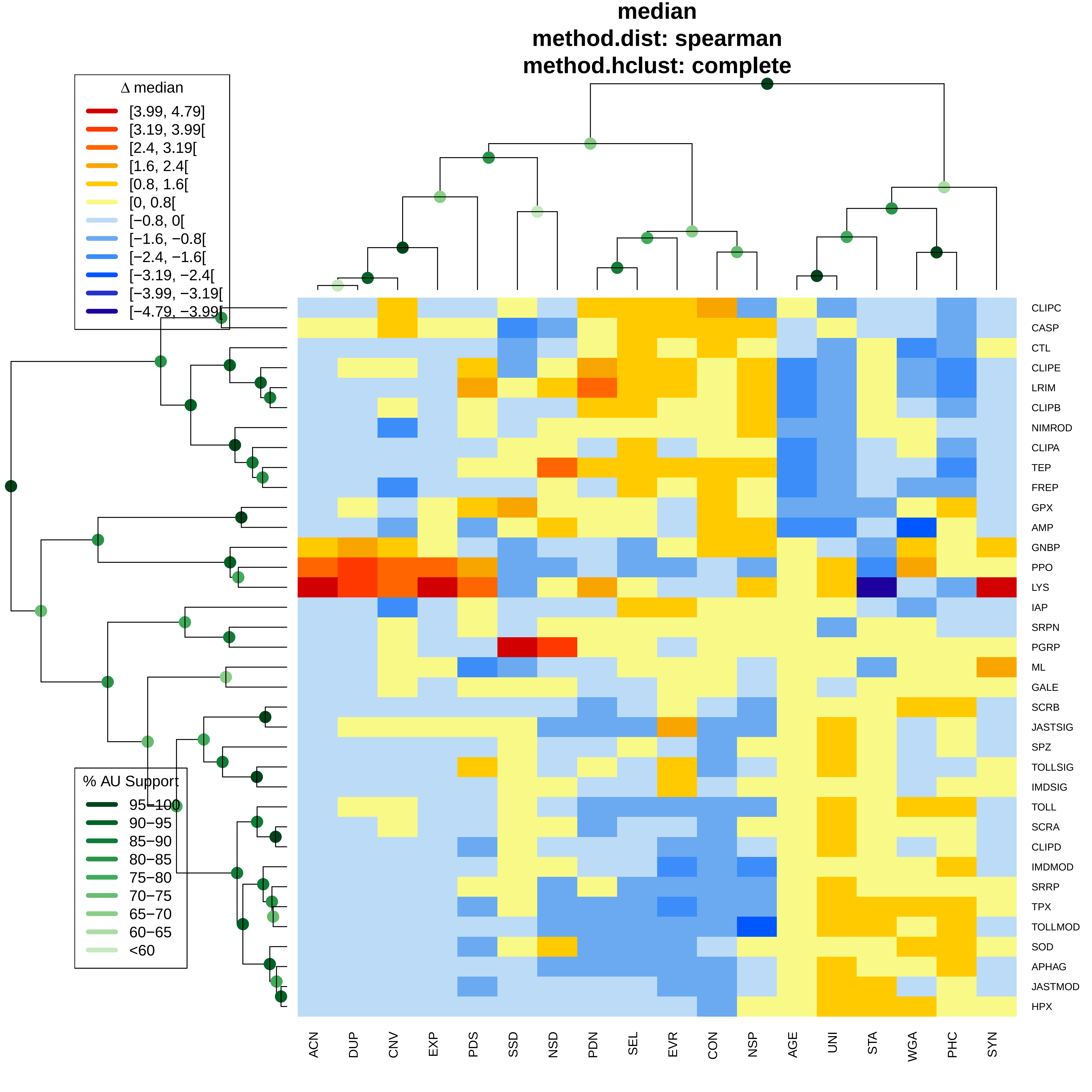
median
method.dist: spearman
method.hclust: complete
Δ median
[3.99, 4.79]
[3.19, 3.99[
[2.4, 3.19[
[1.6, 2.4[
[0.8, 1.6[
[0, 0.8[
[−0.8, 0[
[−1.6, −0.8[
[−2.4, −1.6[
[−3.19, −2.4[
[−3.99, −3.19[
[−4.79, −3.99[
% AU Support
95−100
90−95
85−90
80−85
75−80
70−75
65−70
60−65
<60
CLIPC
CASP
CTL
CLIPE
LRIM
CLIPB
NIMROD
CLIPA
TEP
FREP
GPX
AMP
GNBP
PPO
LYS
IAP
SRPN
PGRP
ML
GALE
SCRB
JASTSIG
SPZ
TOLLSIG
IMDSIG
TOLL
SCRA
CLIPD
IMDMOD
SRRP
TPX
TOLLMOD
SOD
APHAG
JASTMOD
HPX
ACN
DUP
CNV
EXP
PDS
SSD
NSD
PDN
SEL
EVR
CON
NSP
AGE
UNI
STA
WGA
PHC
SYN

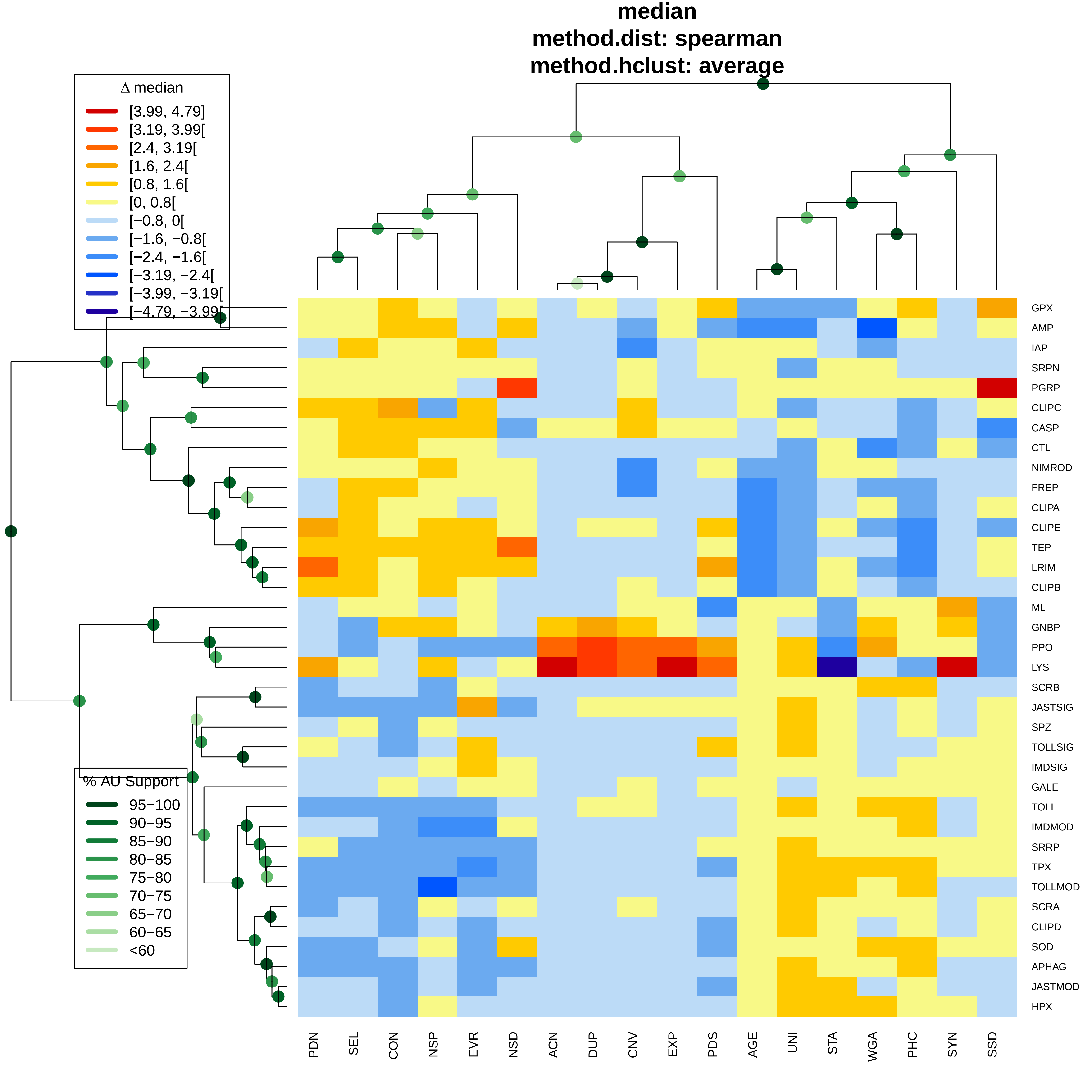

median
method.dist: spearman
method.hclust: average
Δ median
[3.99, 4.79]
[3.19, 3.99[
[2.4, 3.19[
[1.6, 2.4[
[0.8, 1.6[
[0, 0.8[
[−0.8, 0[
[−1.6, −0.8[
[−2.4, −1.6[
[−3.19, −2.4[
[−3.99, −3.19[
[−4.79, −3.99[
% AU Support
95−100
90−95
85−90
80−85
75−80
70−75
65−70
60−65
<60
GPX
AMP
IAP
SRPN
PGRP
CLIPC
CASP
CTL
NIMROD
FREP
CLIPA
CLIPE
TEP
LRIM
CLIPB
ML
GNBP
PPO
LYS
SCRB
JASTSIG
SPZ
TOLLSIG
IMDSIG
GALE
TOLL
IMDMOD
SRRP
TPX
TOLLMOD
SCRA
CLIPD
SOD
APHAG
JASTMOD
HPX
PDN
SEL
CON
NSP
EVR
NSD
ACN
DUP
CNV
EXP
PDS
AGE
UNI
STA
WGA
PHC
SYN
SSD

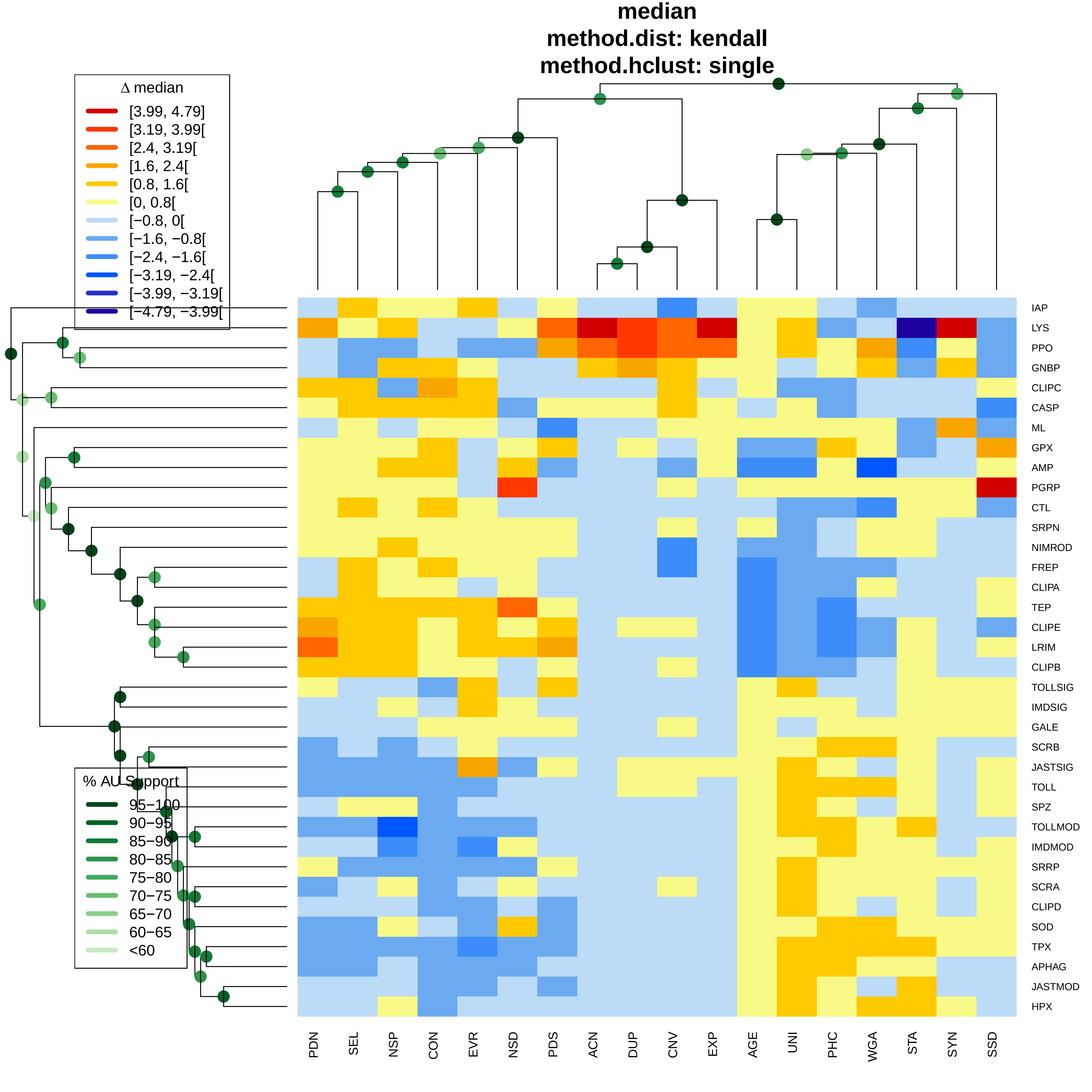
median
method.dist: kendall
method.hclust: single
Δ median
[3.99, 4.79]
[3.19, 3.99[
[2.4, 3.19[
[1.6, 2.4[
[0.8, 1.6[
[0, 0.8[
[−0.8, 0[
[−1.6, −0.8[
[−2.4, −1.6[
[−3.19, −2.4[
[−3.99, −3.19[
[−4.79, −3.99[
% AU Support
95−100
90−95
85−90
80−85
75−80
70−75
65−70
60−65
<60
IAP
LYS
PPO
GNBP
CLIPC
CASP
ML
GPX
AMP
PGRP
CTL
SRPN
NIMROD
FREP
CLIPA
TEP
CLIPE
LRIM
CLIPB
TOLLSIG
IMDSIG
GALE
SCRB
JASTSIG
TOLL
SPZ
TOLLMOD
IMDMOD
SRRP
SCRA
CLIPD
SOD
TPX
APHAG
JASTMOD
HPX
PDN
SEL
NSP
CON
EVR
NSD
PDS
ACN
DUP
CNV
EXP
AGE
UNI
PHC
WGA
STA
SYN
SSD

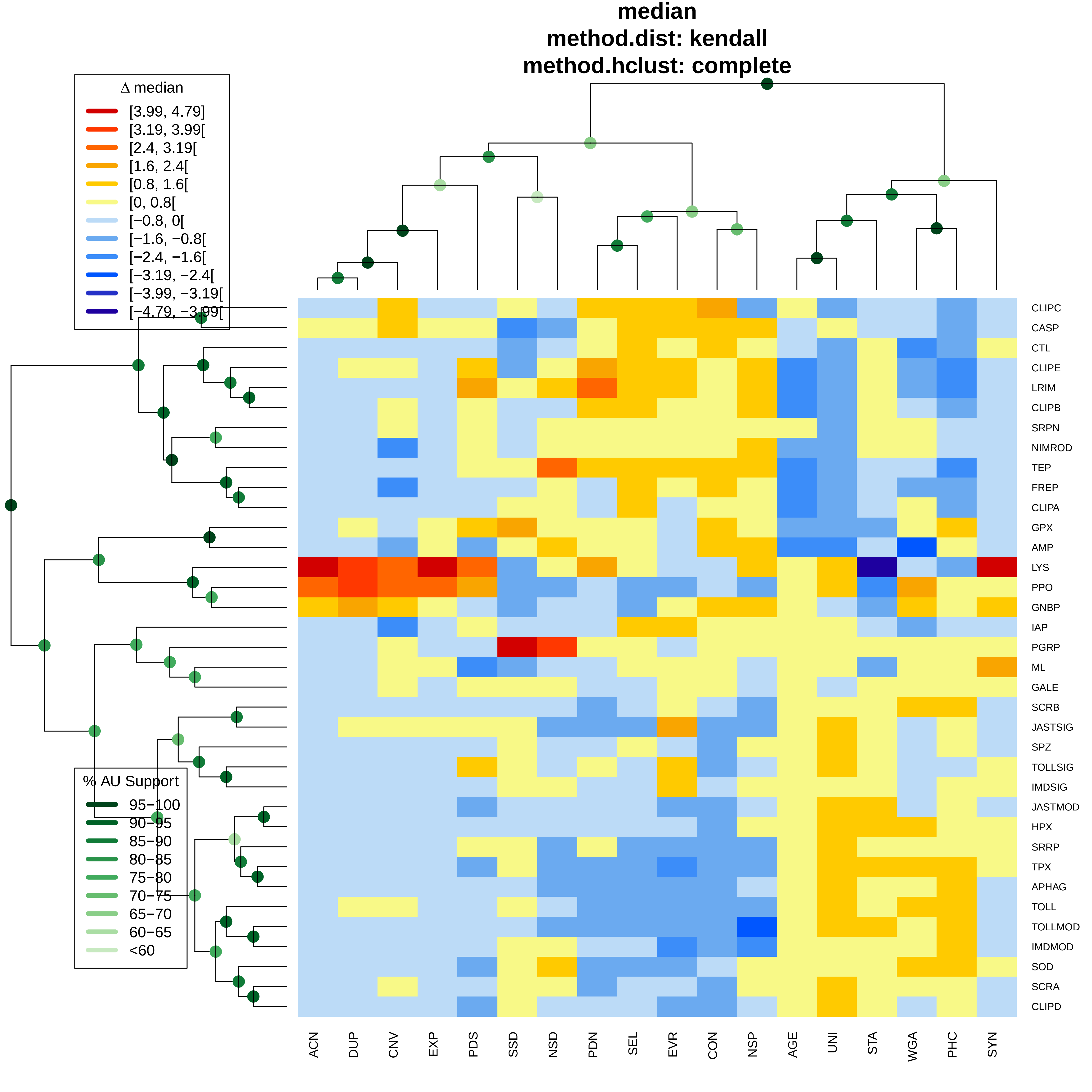
median
method.dist: kendall
method.hclust: complete
Δ median
[3.99, 4.79]
[3.19, 3.99[
[2.4, 3.19[
[1.6, 2.4[
[0.8, 1.6[
[0, 0.8[
[−0.8, 0[
[−1.6, −0.8[
[−2.4, −1.6[
[−3.19, −2.4[
[−3.99, −3.19[
[−4.79, −3.99[
% AU Support
95−100
90−95
85−90
80−85
75−80
70−75
65−70
60−65
<60
CLIPC
CASP
CTL
CLIPE
LRIM
CLIPB
SRPN
NIMROD
TEP
FREP
CLIPA
GPX
AMP
LYS
PPO
GNBP
IAP
PGRP
ML
GALE
SCRB
JASTSIG
SPZ
TOLLSIG
IMDSIG
JASTMOD
HPX
SRRP
TPX
APHAG
TOLL
TOLLMOD
IMDMOD
SOD
SCRA
CLIPD
ACN
DUP
CNV
EXP
PDS
SSD
NSD
PDN
SEL
EVR
CON
NSP
AGE
UNI
STA
WGA
PHC
SYN

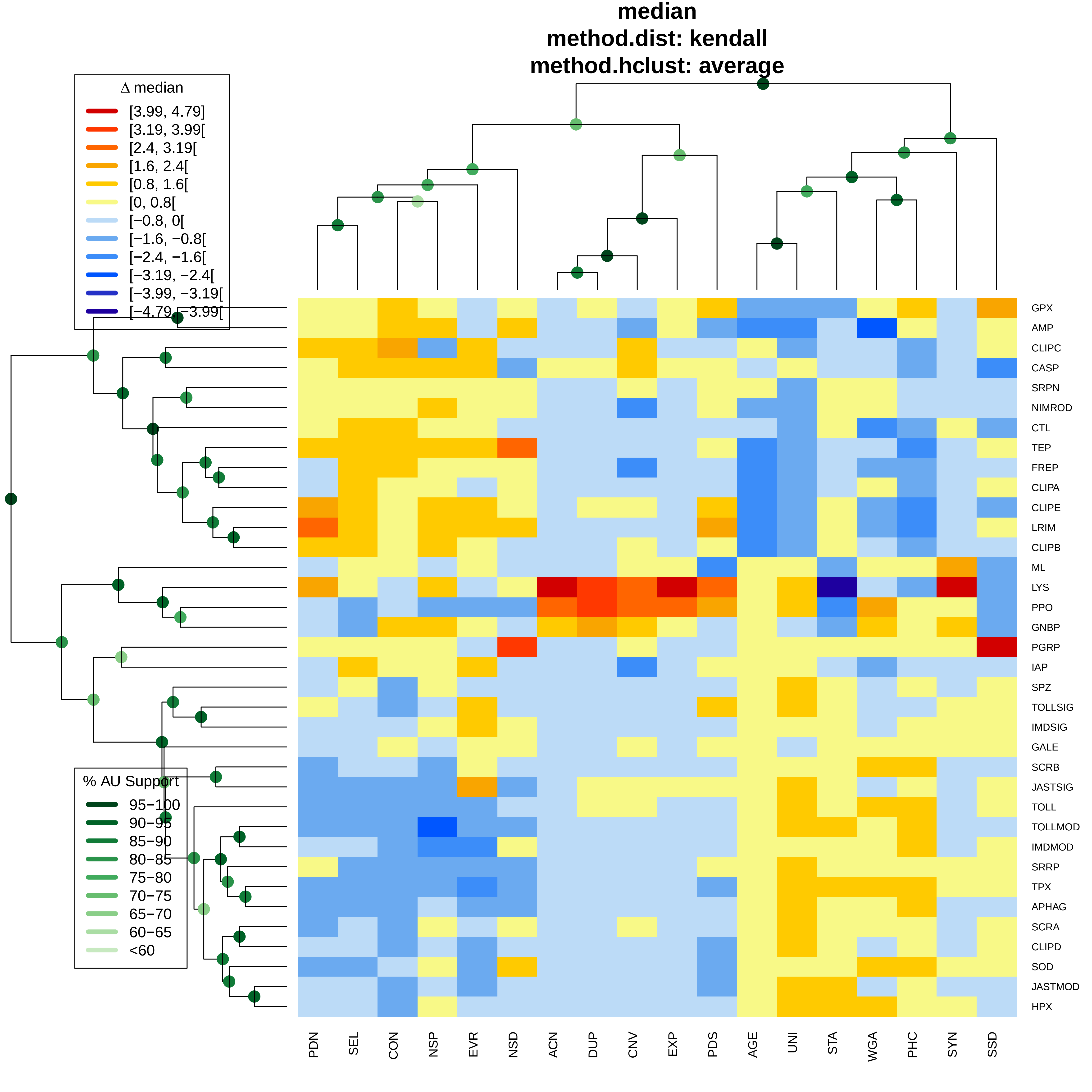

median
method.dist: kendall
method.hclust: average
Δ median
[3.99, 4.79]
[3.19, 3.99[
[2.4, 3.19[
[1.6, 2.4[
[0.8, 1.6[
[0, 0.8[
[−0.8, 0[
[−1.6, −0.8[
[−2.4, −1.6[
[−3.19, −2.4[
[−3.99, −3.19[
[−4.79, −3.99[
% AU Support
95−100
90−95
85−90
80−85
75−80
70−75
65−70
60−65
<60
PDN
SEL
CON
NSP
EVR
NSD
ACN
DUP
CNV
EXP
PDS
AGE
UNI
STA
WGA
PHC
SYN
SSD
GPX
AMP
CLIPC
CASP
SRPN
NIMROD
CTL
TEP
FREP
CLIPA
CLIPE
LRIM
CLIPB
ML
LYS
PPO
GNBP
PGRP
IAP
SPZ
TOLLSIG
IMDSIG
GALE
SCRB
JASTSIG
TOLL
TOLLMOD
IMDMOD
SRRP
TPX
APHAG
SCRA
CLIPD
SOD
JASTMOD
HPX

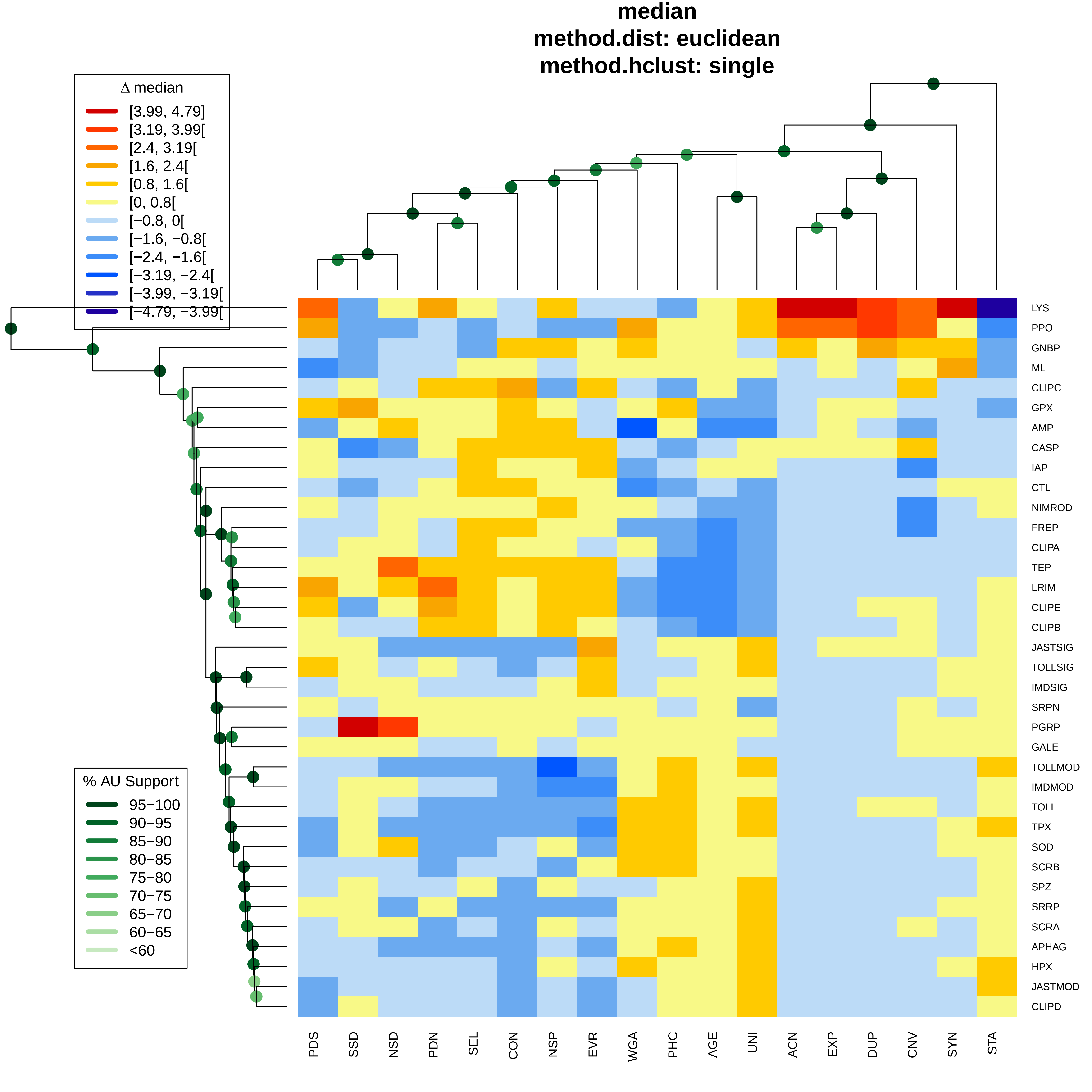

median
method.dist: euclidean
method.hclust: single
Δ median
[3.99, 4.79]
[3.19, 3.99[
[2.4, 3.19[
[1.6, 2.4[
[0.8, 1.6[
[0, 0.8[
[−0.8, 0[
[−1.6, −0.8[
[−2.4, −1.6[
[−3.19, −2.4[
[−3.99, −3.19[
[−4.79, −3.99[
% AU Support
95−100
90−95
85−90
80−85
75−80
70−75
65−70
60−65
<60
LYS
PPO
GNBP
ML
CLIPC
GPX
AMP
CASP
IAP
CTL
NIMROD
FREP
CLIPA
TEP
LRIM
CLIPE
CLIPB
JASTSIG
TOLLSIG
IMDSIG
SRPN
PGRP
GALE
TOLLMOD
IMDMOD
TOLL
TPX
SOD
SCRB
SPZ
SRRP
SCRA
APHAG
HPX
JASTMOD
CLIPD
PDS
SSD
NSD
PDN
SEL
CON
NSP
EVR
WGA
PHC
AGE
UNI
ACN
EXP
DUP
CNV
SYN
STA

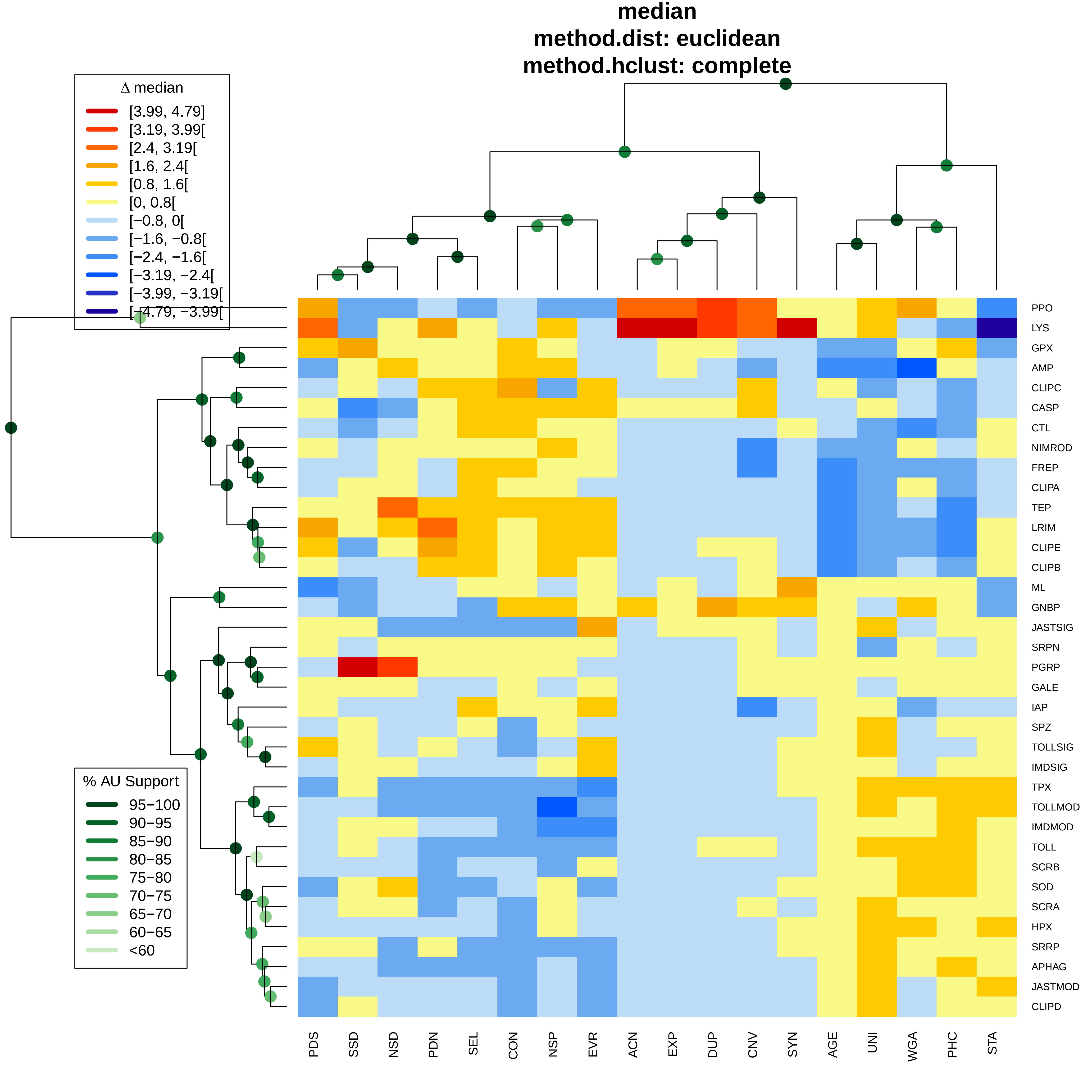
median
method.dist: euclidean
method.hclust: complete
Δ median
[3.99, 4.79]
[3.19, 3.99[
[2.4, 3.19[
[1.6, 2.4[
[0.8, 1.6[
[0, 0.8[
[−0.8, 0[
[−1.6, −0.8[
[−2.4, −1.6[
[−3.19, −2.4[
[−3.99, −3.19[
[−4.79, −3.99[
% AU Support
95−100
90−95
85−90
80−85
75−80
70−75
65−70
60−65
<60
PPO
LYS
GPX
AMP
CLIPC
CASP
CTL
NIMROD
FREP
CLIPA
TEP
LRIM
CLIPE
CLIPB
ML
GNBP
JASTSIG
SRPN
PGRP
GALE
IAP
SPZ
TOLLSIG
IMDSIG
TPX
TOLLMOD
IMDMOD
TOLL
SCRB
SOD
SCRA
HPX
SRRP
APHAG
JASTMOD
CLIPD
PDS
SSD
NSD
PDN
SEL
CON
NSP
EVR
ACN
EXP
DUP
CNV
SYN
AGE
UNI
WGA
PHC
STA

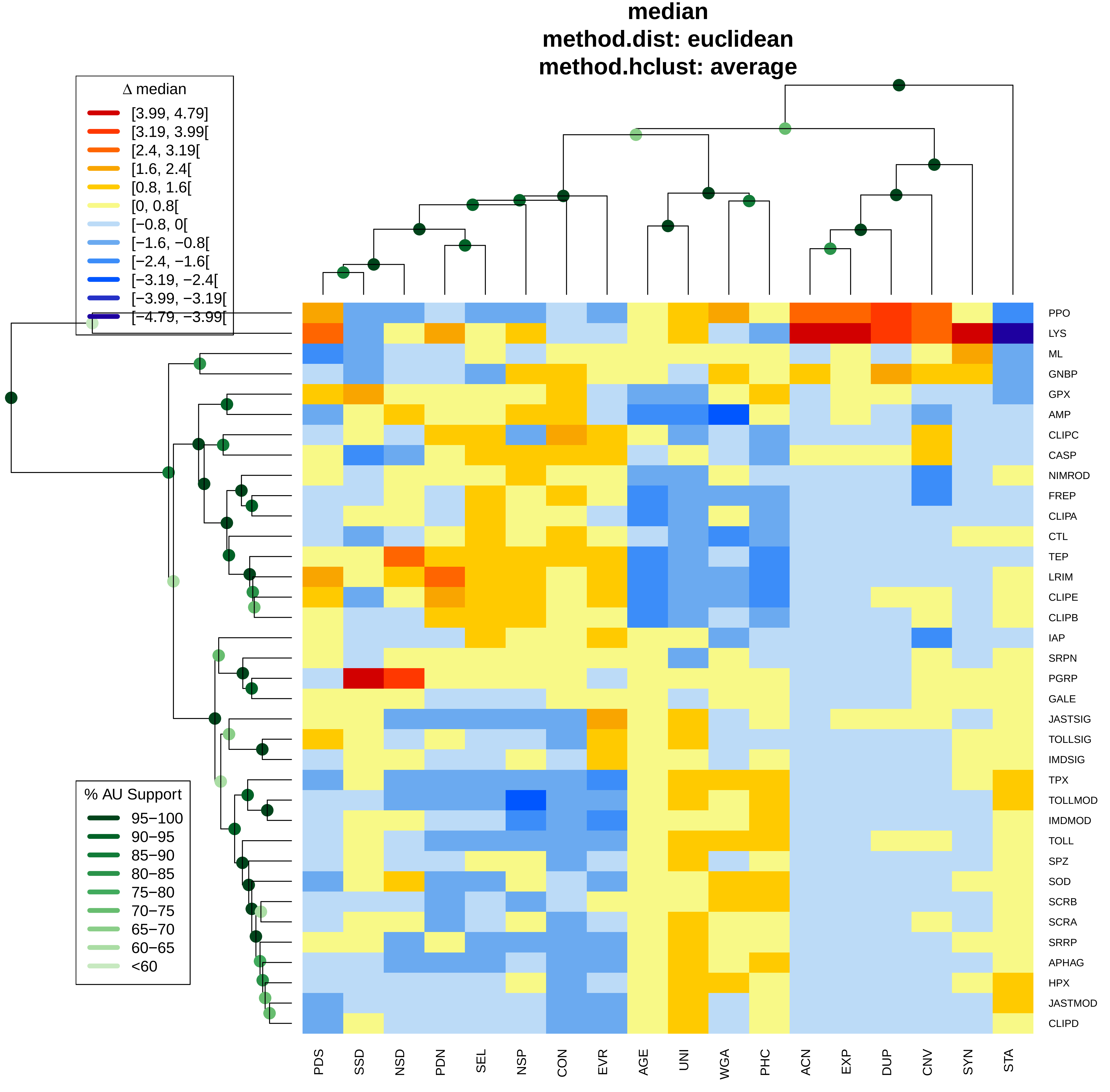
median
method.dist: euclidean
method.hclust: average
Δ median
[3.99, 4.79]
[3.19, 3.99[
[2.4, 3.19[
[1.6, 2.4[
[0.8, 1.6[
[0, 0.8[
[−0.8, 0[
[−1.6, −0.8[
[−2.4, −1.6[
[−3.19, −2.4[
[−3.99, −3.19[
[−4.79, −3.99[
% AU Support
95−100
90−95
85−90
80−85
75−80
70−75
65−70
60−65
<60
PPO
LYS
ML
GNBP
GPX
AMP
CLIPC
CASP
NIMROD
FREP
CLIPA
CTL
TEP
LRIM
CLIPE
CLIPB
IAP
SRPN
PGRP
GALE
JASTSIG
TOLLSIG
IMDSIG
TPX
TOLLMOD
IMDMOD
TOLL
SPZ
SOD
SCRB
SCRA
SRRP
APHAG
HPX
JASTMOD
CLIPD
PDS
SSD
NSD
PDN
SEL
NSP
CON
EVR
AGE
UNI
WGA
PHC
ACN
EXP
DUP
CNV
SYN
STA

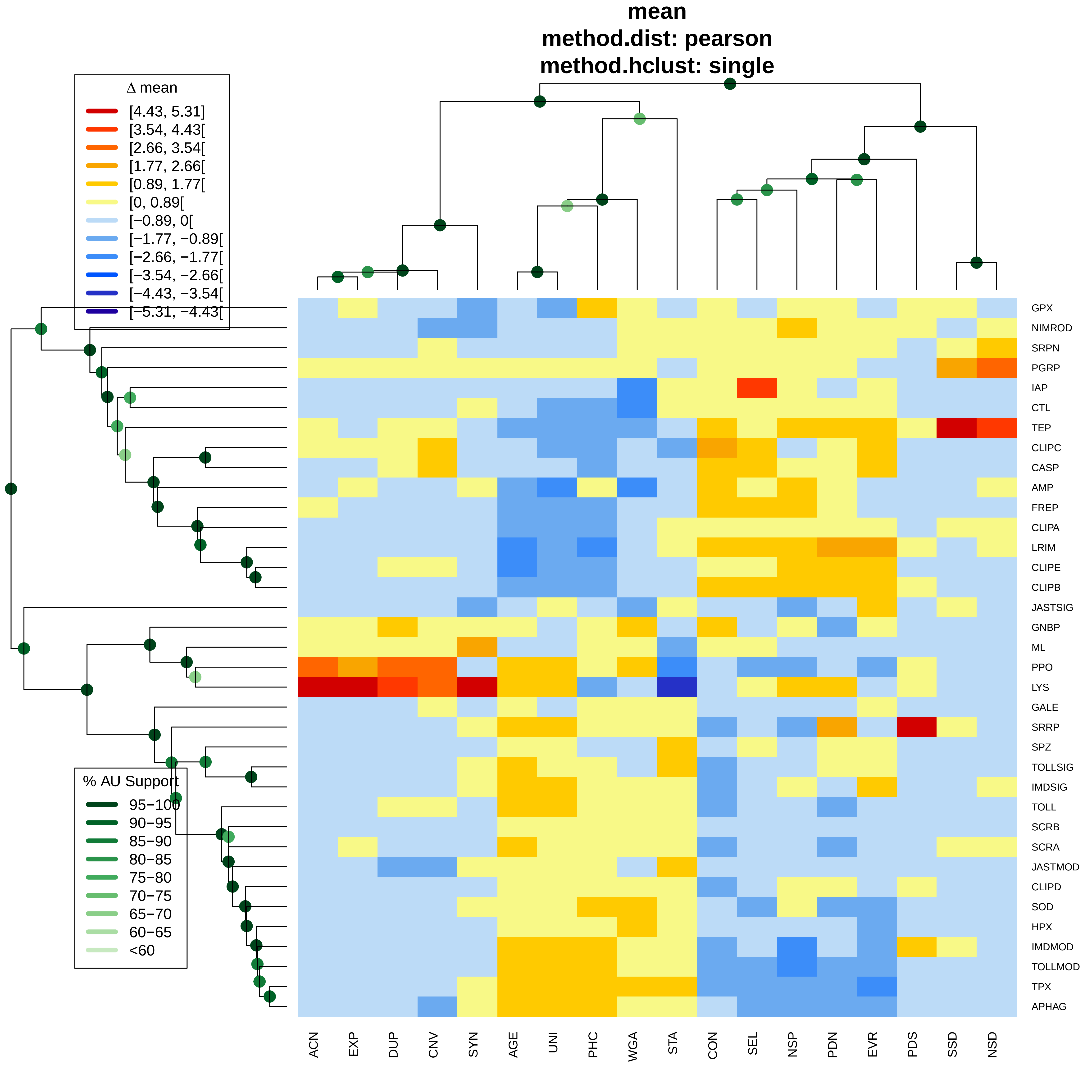

mean
method.dist: pearson
method.hclust: single
Δ mean
[4.43, 5.31]
[3.54, 4.43[
[2.66, 3.54[
[1.77, 2.66[
[0.89, 1.77[
[0, 0.89[
[−0.89, 0[
[−1.77, −0.89[
[−2.66, −1.77[
[−3.54, −2.66[
[−4.43, −3.54[
[−5.31, −4.43[
% AU Support
95−100
90−95
85−90
80−85
75−80
70−75
65−70
60−65
<60
GPX
NIMROD
SRPN
PGRP
IAP
CTL
TEP
CLIPC
CASP
AMP
FREP
CLIPA
LRIM
CLIPE
CLIPB
JASTSIG
GNBP
ML
PPO
LYS
GALE
SRRP
SPZ
TOLLSIG
IMDSIG
TOLL
SCRB
SCRA
JASTMOD
CLIPD
SOD
HPX
IMDMOD
TOLLMOD
TPX
APHAG
ACN
EXP
DUP
CNV
SYN
AGE
UNI
PHC
WGA
STA
CON
SEL
NSP
PDN
EVR
PDS
SSD
NSD

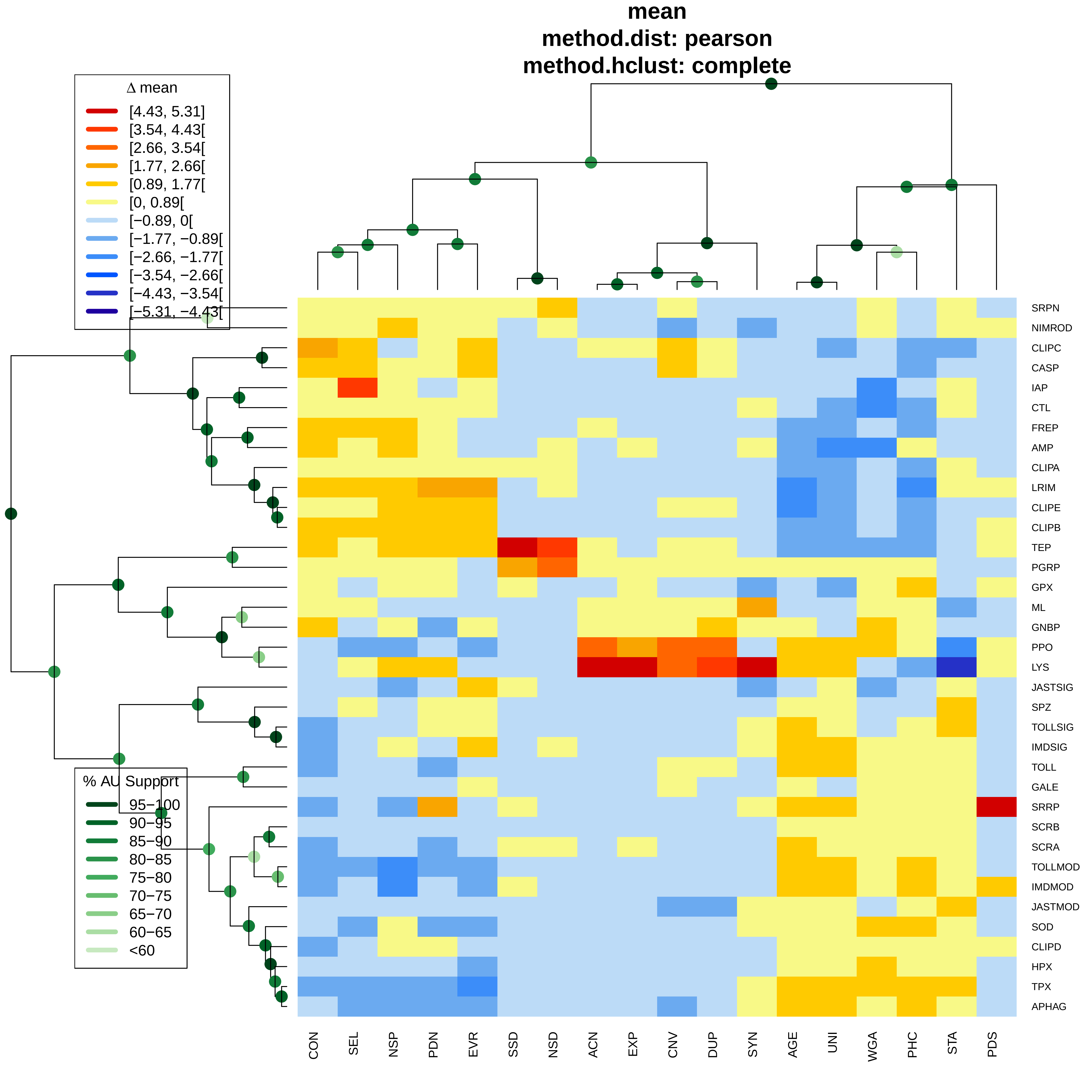
mean
method.dist: pearson
method.hclust: complete
Δ mean
[4.43, 5.31]
[3.54, 4.43[
[2.66, 3.54[
[1.77, 2.66[
[0.89, 1.77[
[0, 0.89[
[−0.89, 0[
[−1.77, −0.89[
[−2.66, −1.77[
[−3.54, −2.66[
[−4.43, −3.54[
[−5.31, −4.43[
% AU Support
95−100
90−95
85−90
80−85
75−80
70−75
65−70
60−65
<60
SRPN
NIMROD
CLIPC
CASP
IAP
CTL
FREP
AMP
CLIPA
LRIM
CLIPE
CLIPB
TEP
PGRP
GPX
ML
GNBP
PPO
LYS
JASTSIG
SPZ
TOLLSIG
IMDSIG
TOLL
GALE
SRRP
SCRB
SCRA
TOLLMOD
IMDMOD
JASTMOD
SOD
CLIPD
HPX
TPX
APHAG
CON
SEL
NSP
PDN
EVR
SSD
NSD
ACN
EXP
CNV
DUP
SYN
AGE
UNI
WGA
PHC
STA
PDS

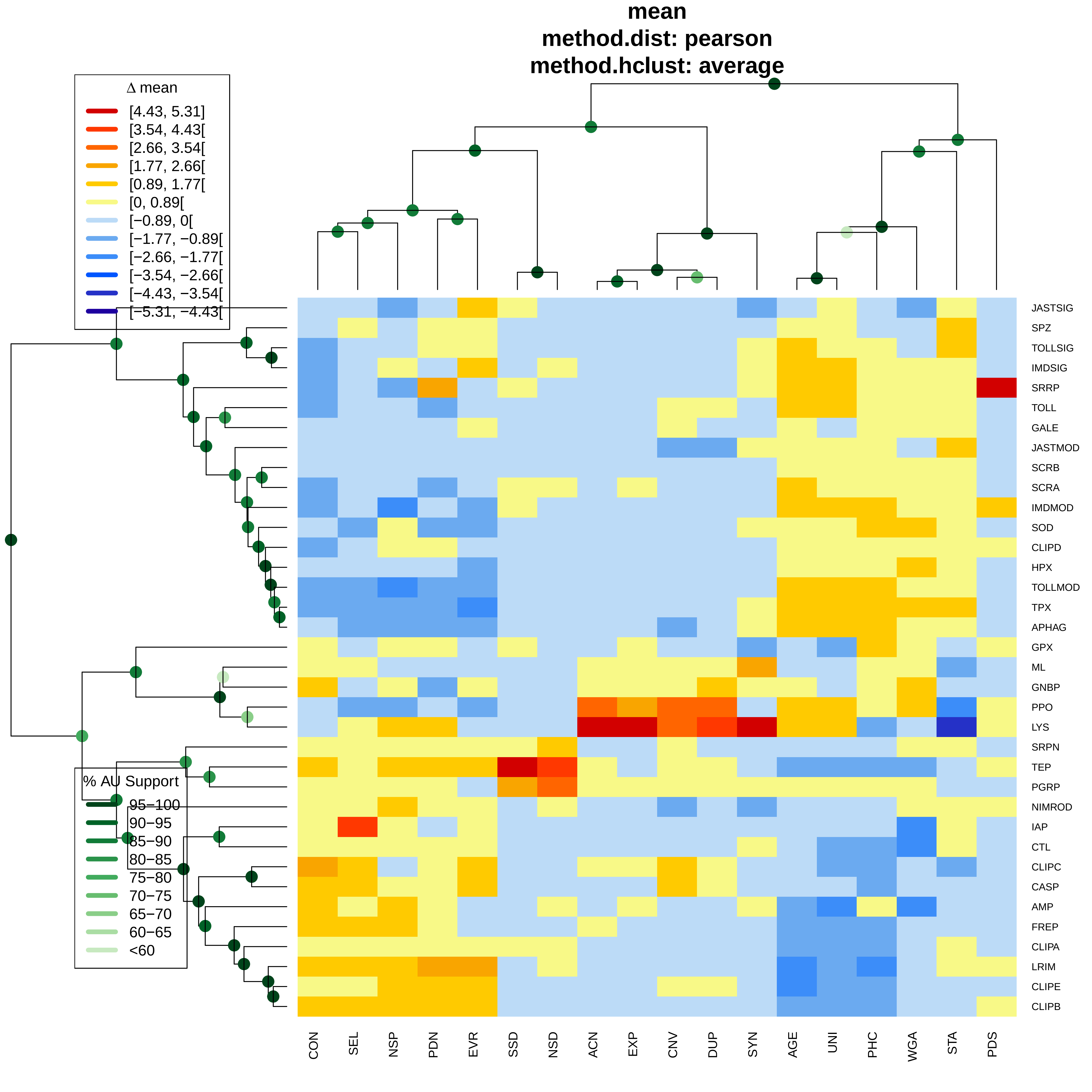

mean
method.dist: pearson
method.hclust: average
Δ mean
[4.43, 5.31]
[3.54, 4.43[
[2.66, 3.54[
[1.77, 2.66[
[0.89, 1.77[
[0, 0.89[
[−0.89, 0[
[−1.77, −0.89[
[−2.66, −1.77[
[−3.54, −2.66[
[−4.43, −3.54[
[−5.31, −4.43[
% AU Support
95−100
90−95
85−90
80−85
75−80
70−75
65−70
60−65
<60
JASTSIG
SPZ
TOLLSIG
IMDSIG
SRRP
TOLL
GALE
JASTMOD
SCRB
SCRA
IMDMOD
SOD
CLIPD
HPX
TOLLMOD
TPX
APHAG
GPX
ML
GNBP
PPO
LYS
SRPN
TEP
PGRP
NIMROD
IAP
CTL
CLIPC
CASP
AMP
FREP
CLIPA
LRIM
CLIPE
CLIPB
CON
SEL
NSP
PDN
EVR
SSD
NSD
ACN
EXP
CNV
DUP
SYN
AGE
UNI
PHC
WGA
STA
PDS

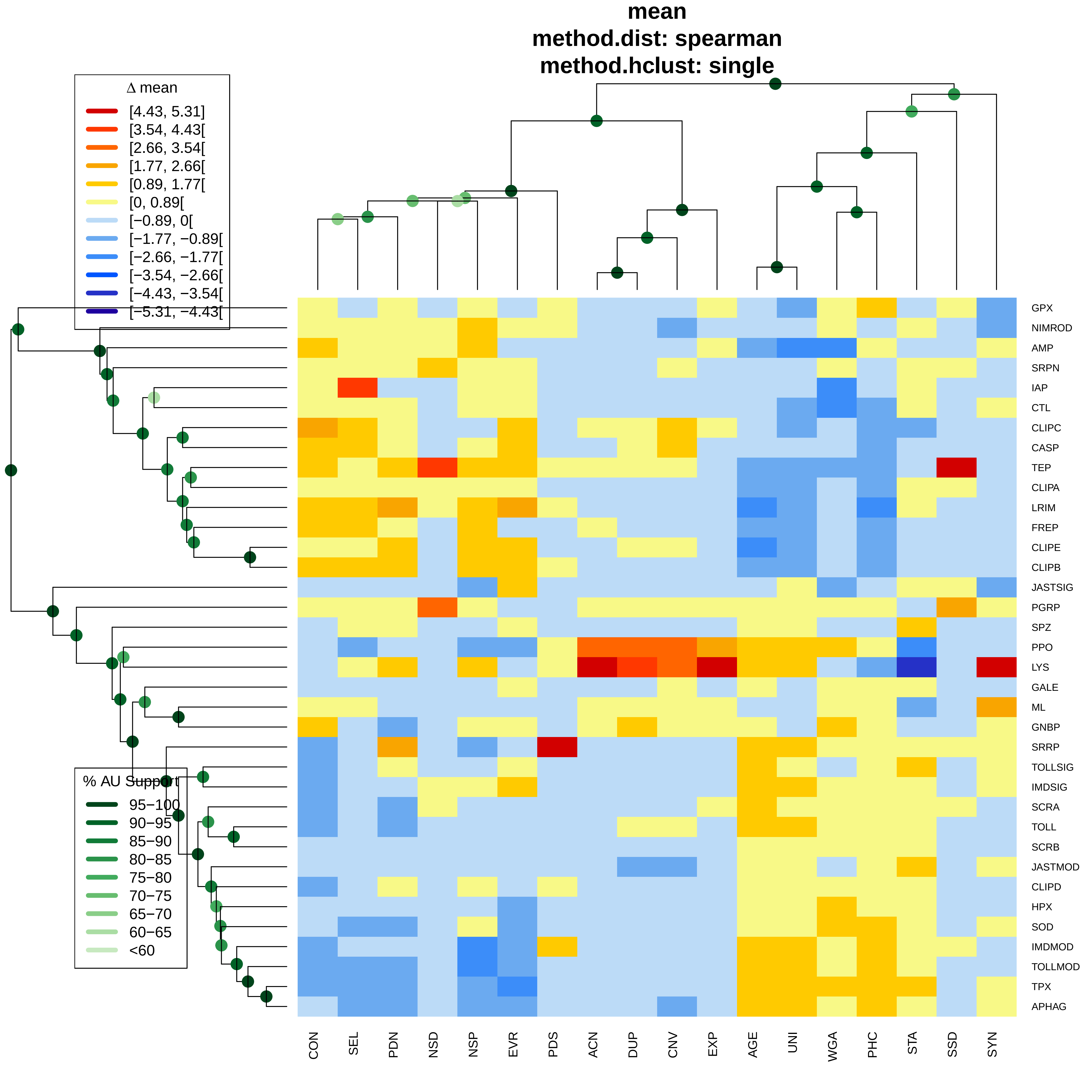

mean
method.dist: spearman
method.hclust: single
Δ mean
[4.43, 5.31]
[3.54, 4.43[
[2.66, 3.54[
[1.77, 2.66[
[0.89, 1.77[
[0, 0.89[
[−0.89, 0[
[−1.77, −0.89[
[−2.66, −1.77[
[−3.54, −2.66[
[−4.43, −3.54[
[−5.31, −4.43[
% AU Support
95−100
90−95
85−90
80−85
75−80
70−75
65−70
60−65
<60
GPX
NIMROD
AMP
SRPN
IAP
CTL
CLIPC
CASP
TEP
CLIPA
LRIM
FREP
CLIPE
CLIPB
JASTSIG
PGRP
SPZ
PPO
LYS
GALE
ML
GNBP
SRRP
TOLLSIG
IMDSIG
SCRA
TOLL
SCRB
JASTMOD
CLIPD
HPX
SOD
IMDMOD
TOLLMOD
TPX
APHAG
CON
SEL
PDN
NSD
NSP
EVR
PDS
ACN
DUP
CNV
EXP
AGE
UNI
WGA
PHC
STA
SSD
SYN

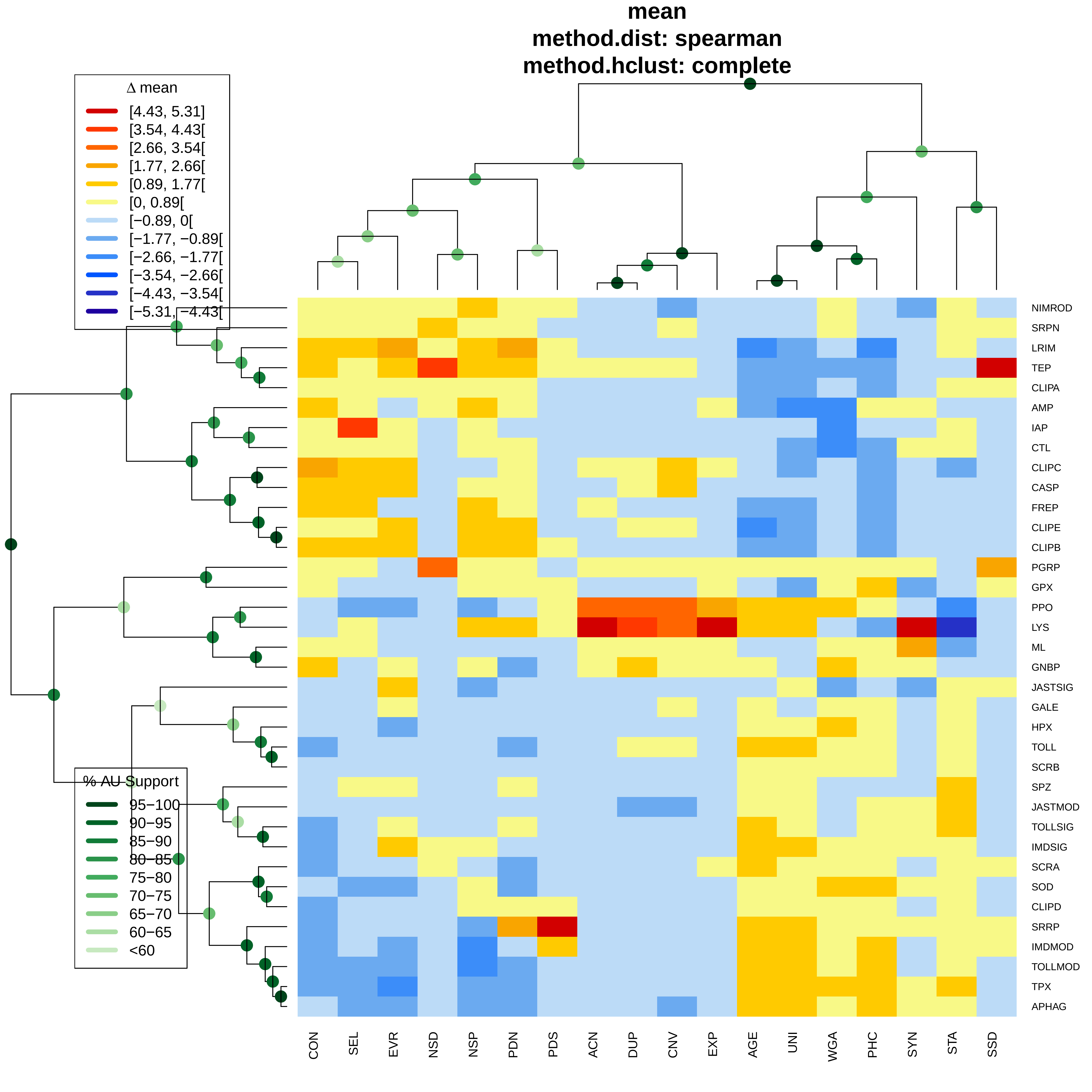

mean
method.dist: spearman
method.hclust: complete
Δ mean
[4.43, 5.31]
[3.54, 4.43[
[2.66, 3.54[
[1.77, 2.66[
[0.89, 1.77[
[0, 0.89[
[−0.89, 0[
[−1.77, −0.89[
[−2.66, −1.77[
[−3.54, −2.66[
[−4.43, −3.54[
[−5.31, −4.43[
% AU Support
95−100
90−95
85−90
80−85
75−80
70−75
65−70
60−65
<60
NIMROD
SRPN
LRIM
TEP
CLIPA
AMP
IAP
CTL
CLIPC
CASP
FREP
CLIPE
CLIPB
PGRP
GPX
PPO
LYS
ML
GNBP
JASTSIG
GALE
HPX
TOLL
SCRB
SPZ
JASTMOD
TOLLSIG
IMDSIG
SCRA
SOD
CLIPD
SRRP
IMDMOD
TOLLMOD
TPX
APHAG
CON
SEL
EVR
NSD
NSP
PDN
PDS
ACN
DUP
CNV
EXP
AGE
UNI
WGA
PHC
SYN
STA
SSD

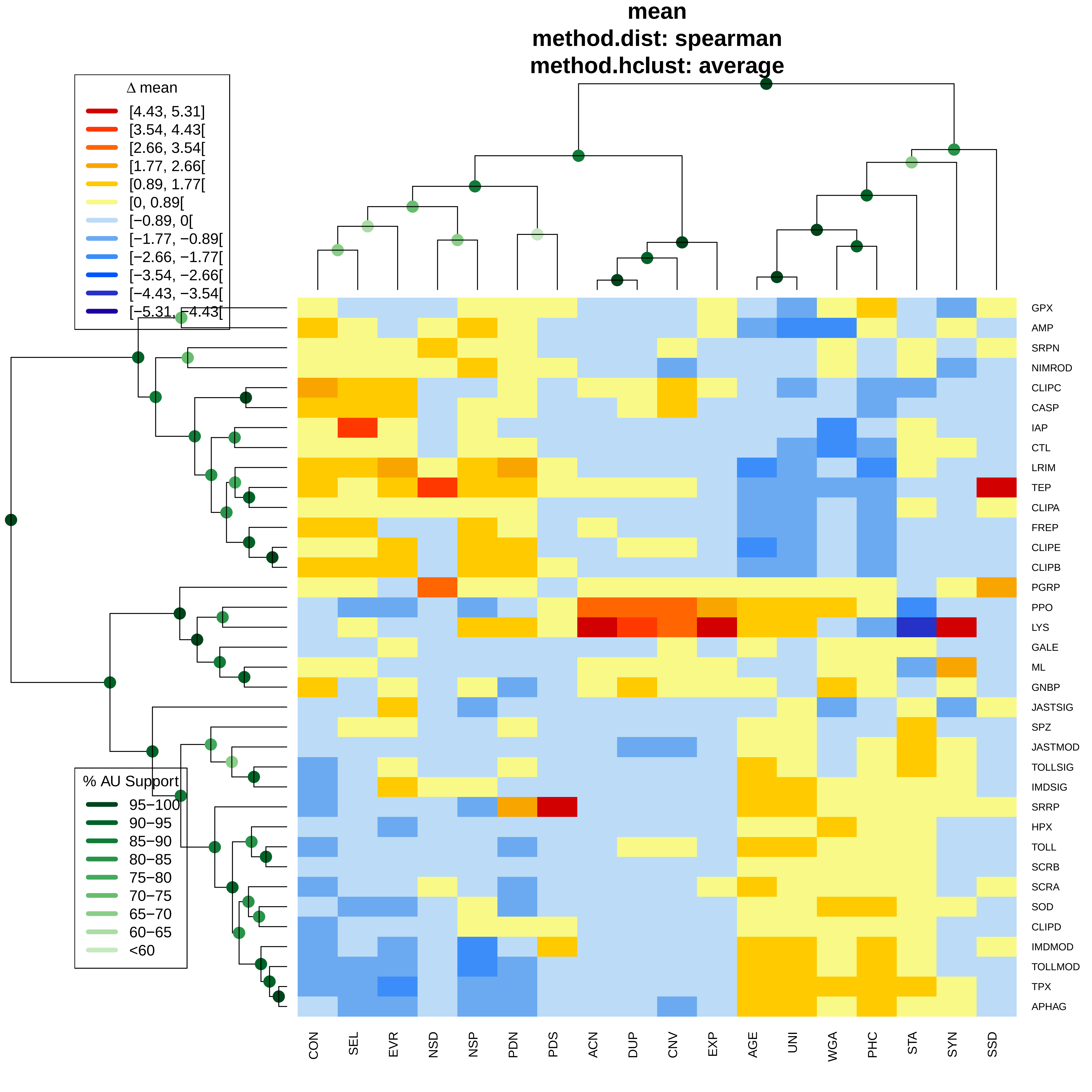

mean
method.dist: spearman
method.hclust: average
Δ mean
[4.43, 5.31]
[3.54, 4.43[
[2.66, 3.54[
[1.77, 2.66[
[0.89, 1.77[
[0, 0.89[
[−0.89, 0[
[−1.77, −0.89[
[−2.66, −1.77[
[−3.54, −2.66[
[−4.43, −3.54[
[−5.31, −4.43[
% AU Support
95−100
90−95
85−90
80−85
75−80
70−75
65−70
60−65
<60
GPX
AMP
SRPN
NIMROD
CLIPC
CASP
IAP
CTL
LRIM
TEP
CLIPA
FREP
CLIPE
CLIPB
PGRP
PPO
LYS
GALE
ML
GNBP
JASTSIG
SPZ
JASTMOD
TOLLSIG
IMDSIG
SRRP
HPX
TOLL
SCRB
SCRA
SOD
CLIPD
IMDMOD
TOLLMOD
TPX
APHAG
CON
SEL
EVR
NSD
NSP
PDN
PDS
ACN
DUP
CNV
EXP
AGE
UNI
WGA
PHC
STA
SYN
SSD

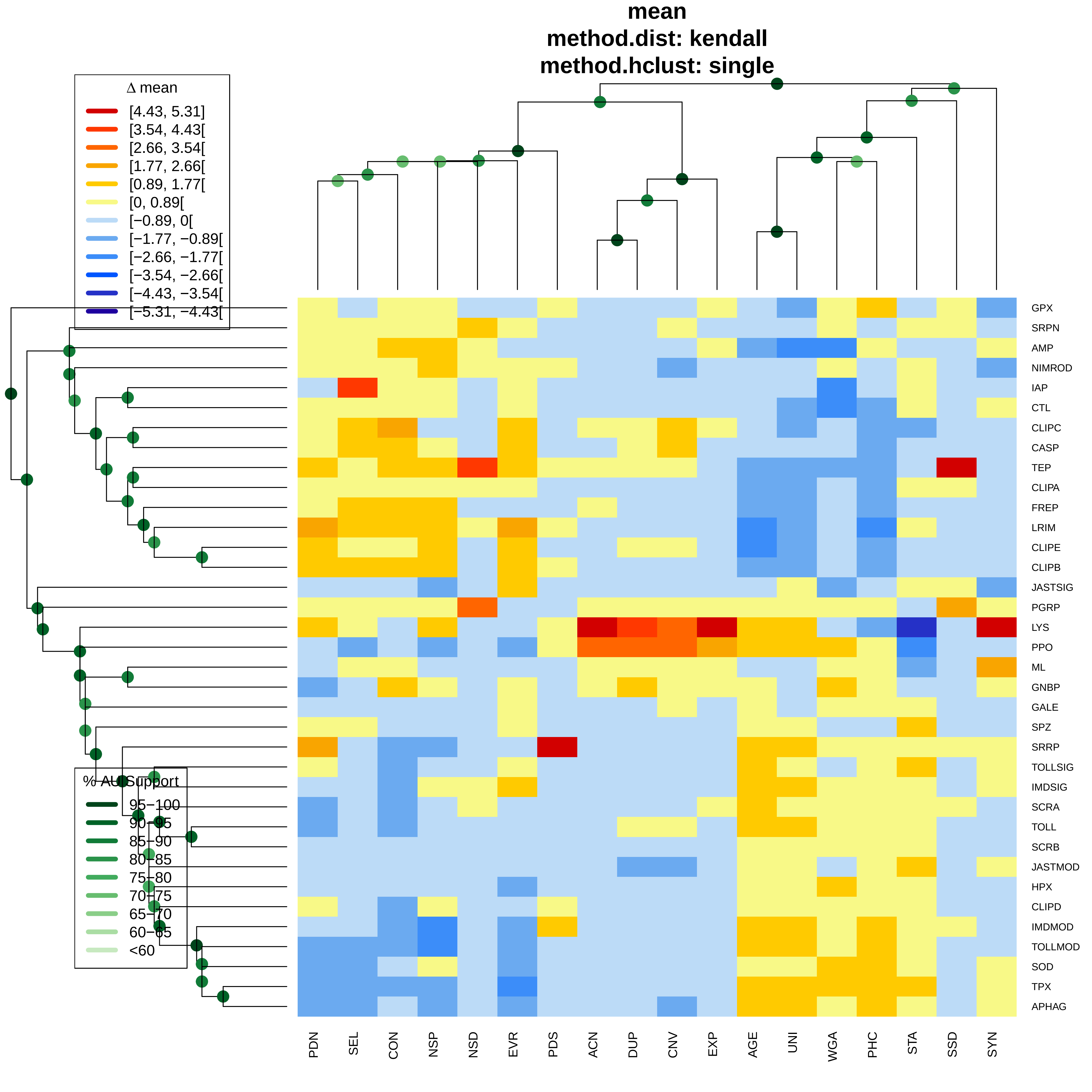
mean
method.dist: kendall
method.hclust: single
Δ mean
[4.43, 5.31]
[3.54, 4.43[
[2.66, 3.54[
[1.77, 2.66[
[0.89, 1.77[
[0, 0.89[
[−0.89, 0[
[−1.77, −0.89[
[−2.66, −1.77[
[−3.54, −2.66[
[−4.43, −3.54[
[−5.31, −4.43[
% AU Support
95−100
90−95
85−90
80−85
75−80
70−75
65−70
60−65
<60
GPX
SRPN
AMP
NIMROD
IAP
CTL
CLIPC
CASP
TEP
CLIPA
FREP
LRIM
CLIPE
CLIPB
JASTSIG
PGRP
LYS
PPO
ML
GNBP
GALE
SPZ
SRRP
TOLLSIG
IMDSIG
SCRA
TOLL
SCRB
JASTMOD
HPX
CLIPD
IMDMOD
TOLLMOD
SOD
TPX
APHAG
PDN
SEL
CON
NSP
NSD
EVR
PDS
ACN
DUP
CNV
EXP
AGE
UNI
WGA
PHC
STA
SSD
SYN

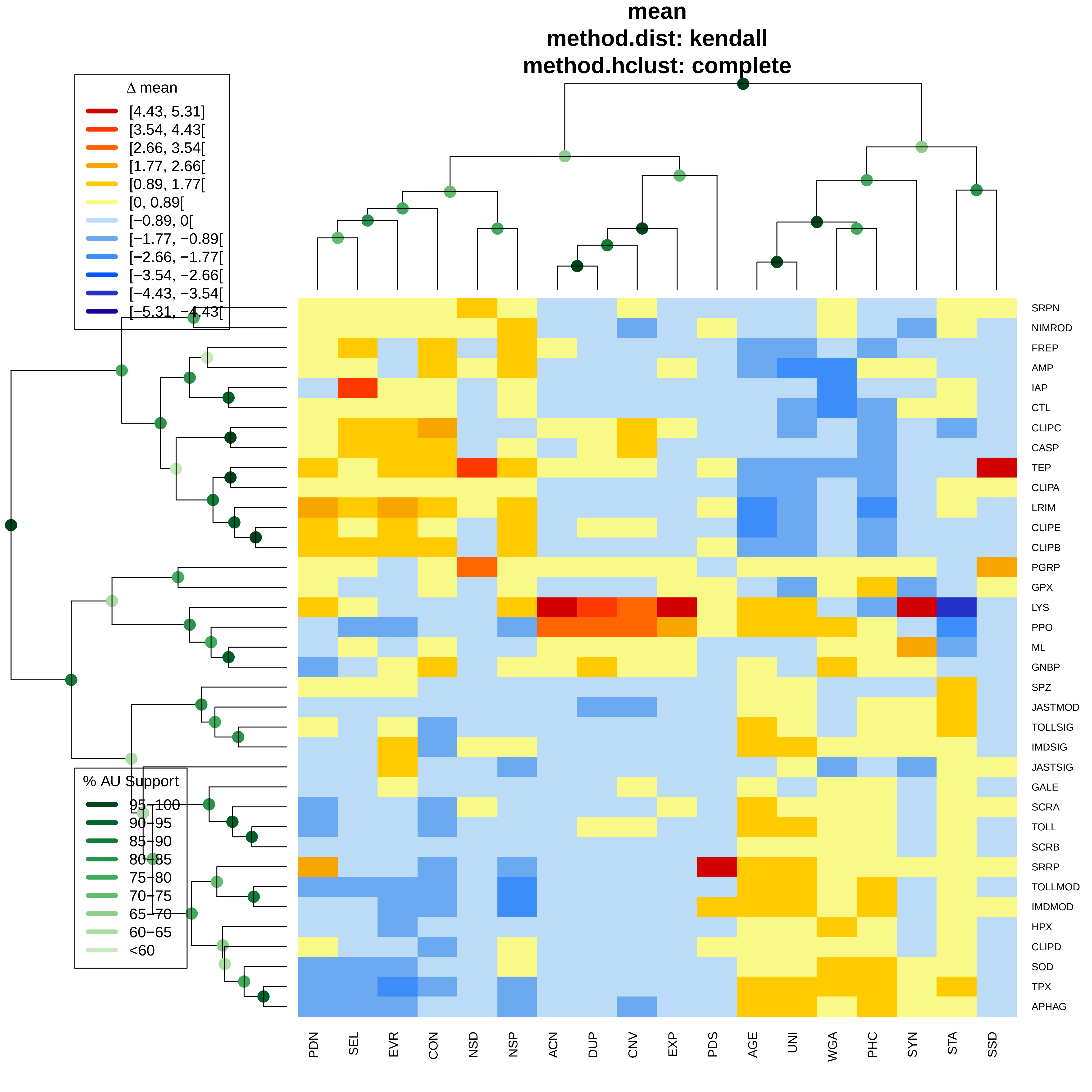
mean
method.dist: kendall
method.hclust: complete
Δ mean
[4.43, 5.31]
[3.54, 4.43[
[2.66, 3.54[
[1.77, 2.66[
[0.89, 1.77[
[0, 0.89[
[−0.89, 0[
[−1.77, −0.89[
[−2.66, −1.77[
[−3.54, −2.66[
[−4.43, −3.54[
[−5.31, −4.43[
% AU Support
95−100
90−95
85−90
80−85
75−80
70−75
65−70
60−65
<60
SRPN
NIMROD
FREP
AMP
IAP
CTL
CLIPC
CASP
TEP
CLIPA
LRIM
CLIPE
CLIPB
PGRP
GPX
LYS
PPO
ML
GNBP
SPZ
JASTMOD
TOLLSIG
IMDSIG
JASTSIG
GALE
SCRA
TOLL
SCRB
SRRP
TOLLMOD
IMDMOD
HPX
CLIPD
SOD
TPX
APHAG
PDN
SEL
EVR
CON
NSD
NSP
ACN
DUP
CNV
EXP
PDS
AGE
UNI
WGA
PHC
SYN
STA
SSD

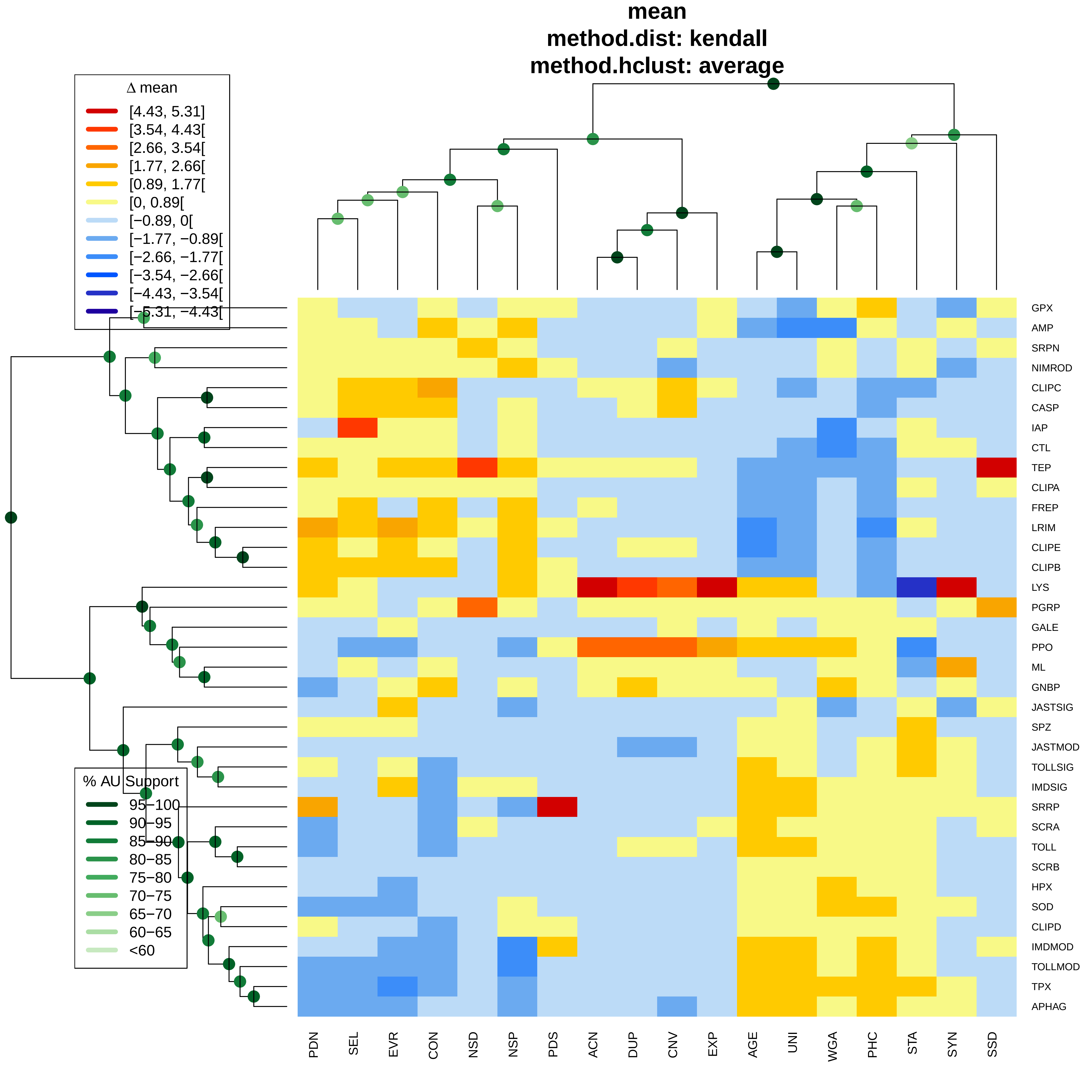

mean
method.dist: kendall
method.hclust: average
Δ mean
[4.43, 5.31]
[3.54, 4.43[
[2.66, 3.54[
[1.77, 2.66[
[0.89, 1.77[
[0, 0.89[
[−0.89, 0[
[−1.77, −0.89[
[−2.66, −1.77[
[−3.54, −2.66[
[−4.43, −3.54[
[−5.31, −4.43[
% AU Support
95−100
90−95
85−90
80−85
75−80
70−75
65−70
60−65
<60
GPX
AMP
SRPN
NIMROD
CLIPC
CASP
IAP
CTL
TEP
CLIPA
FREP
LRIM
CLIPE
CLIPB
LYS
PGRP
GALE
PPO
ML
GNBP
JASTSIG
SPZ
JASTMOD
TOLLSIG
IMDSIG
SRRP
SCRA
TOLL
SCRB
HPX
SOD
CLIPD
IMDMOD
TOLLMOD
TPX
APHAG
PDN
SEL
EVR
CON
NSD
NSP
PDS
ACN
DUP
CNV
EXP
AGE
UNI
WGA
PHC
STA
SYN
SSD

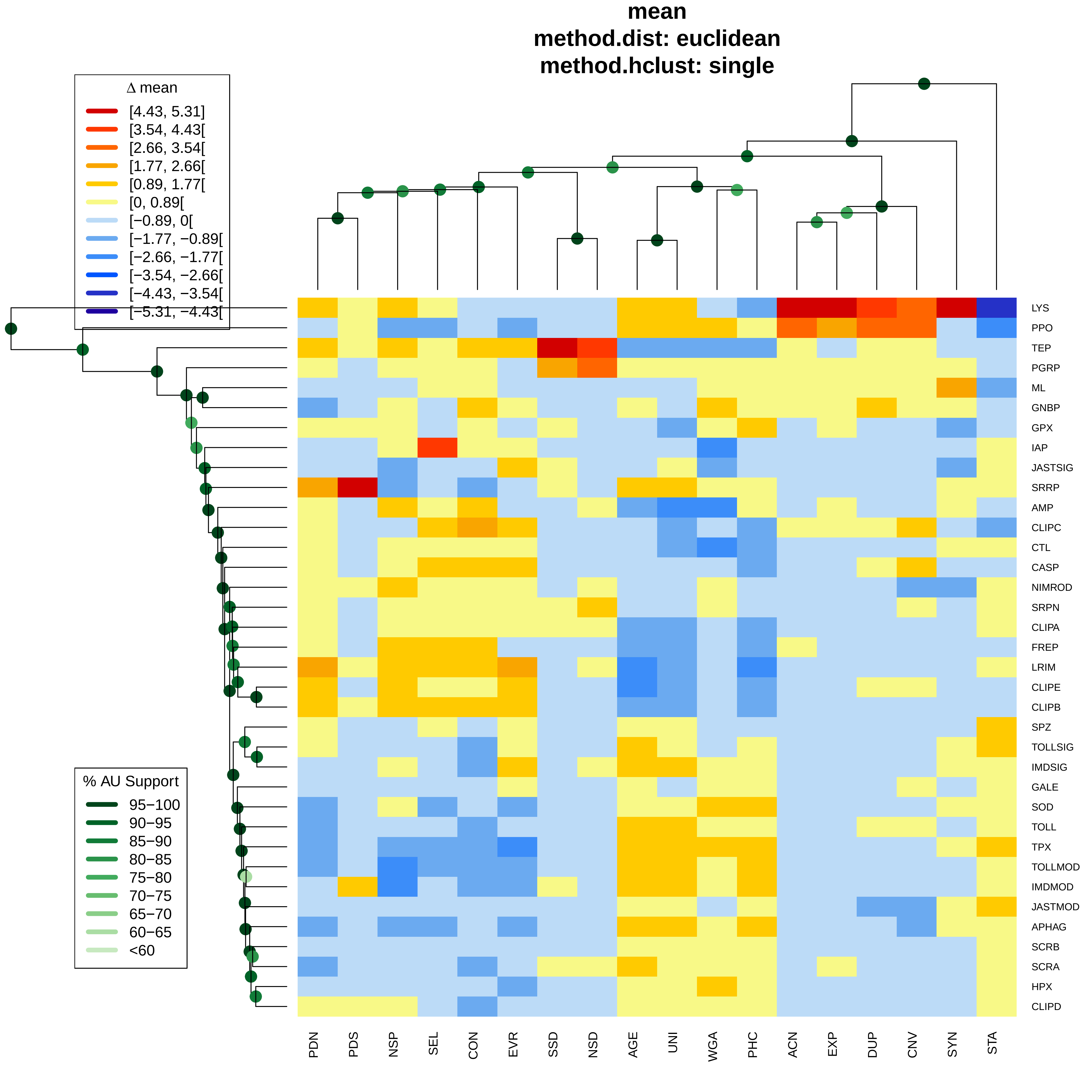
mean
method.dist: euclidean
method.hclust: single
Δ mean
[4.43, 5.31]
[3.54, 4.43[
[2.66, 3.54[
[1.77, 2.66[
[0.89, 1.77[
[0, 0.89[
[−0.89, 0[
[−1.77, −0.89[
[−2.66, −1.77[
[−3.54, −2.66[
[−4.43, −3.54[
[−5.31, −4.43[
% AU Support
95−100
90−95
85−90
80−85
75−80
70−75
65−70
60−65
<60
LYS
PPO
TEP
PGRP
ML
GNBP
GPX
IAP
JASTSIG
SRRP
AMP
CLIPC
CTL
CASP
NIMROD
SRPN
CLIPA
FREP
LRIM
CLIPE
CLIPB
SPZ
TOLLSIG
IMDSIG
GALE
SOD
TOLL
TPX
TOLLMOD
IMDMOD
JASTMOD
APHAG
SCRB
SCRA
HPX
CLIPD
PDN
PDS
NSP
SEL
CON
EVR
SSD
NSD
AGE
UNI
WGA
PHC
ACN
EXP
DUP
CNV
SYN
STA

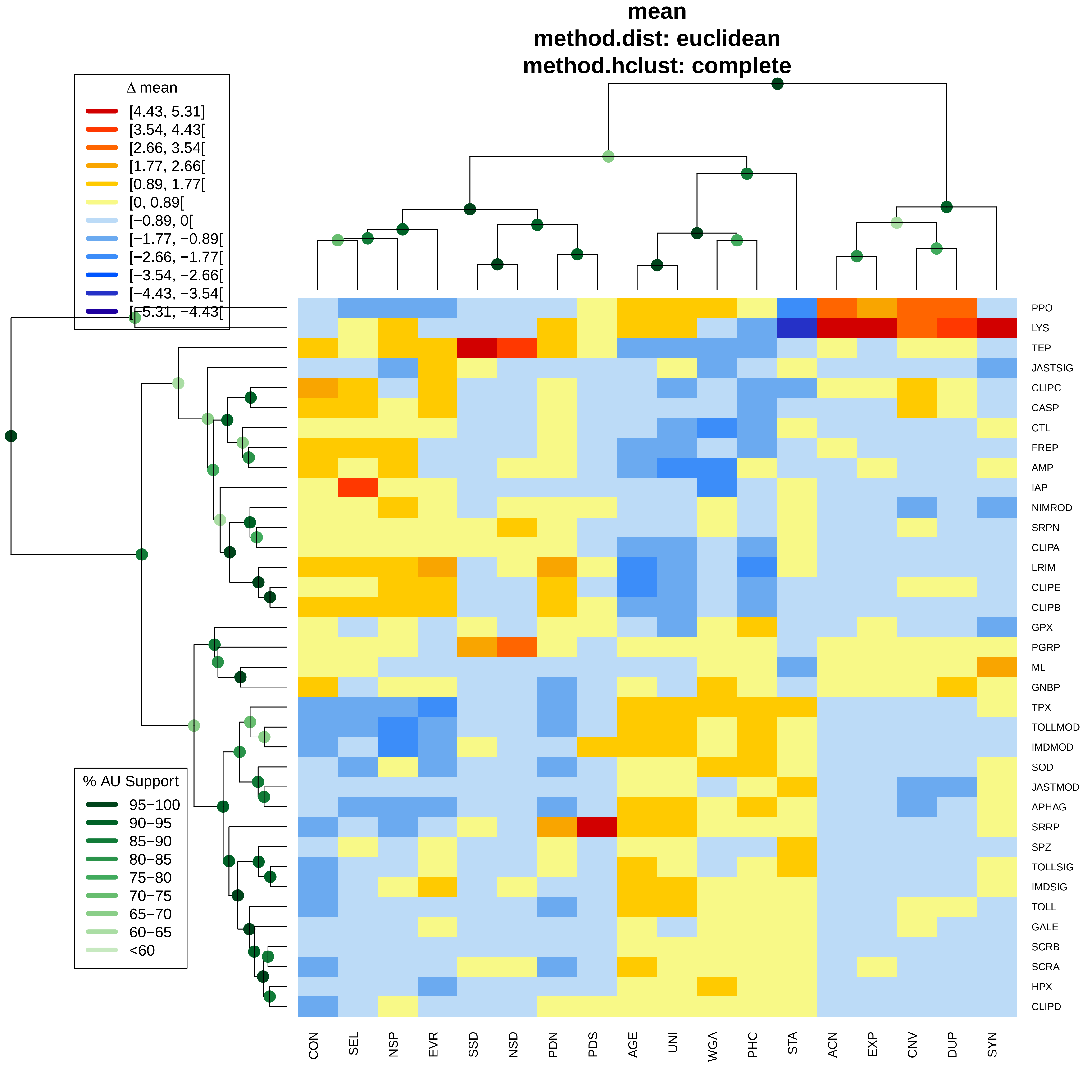
mean
method.dist: euclidean
method.hclust: complete
Δ mean
[4.43, 5.31]
[3.54, 4.43[
[2.66, 3.54[
[1.77, 2.66[
[0.89, 1.77[
[0, 0.89[
[−0.89, 0[
[−1.77, −0.89[
[−2.66, −1.77[
[−3.54, −2.66[
[−4.43, −3.54[
[−5.31, −4.43[
% AU Support
95−100
90−95
85−90
80−85
75−80
70−75
65−70
60−65
<60
PPO
LYS
TEP
JASTSIG
CLIPC
CASP
CTL
FREP
AMP
IAP
NIMROD
SRPN
CLIPA
LRIM
CLIPE
CLIPB
GPX
PGRP
ML
GNBP
TPX
TOLLMOD
IMDMOD
SOD
JASTMOD
APHAG
SRRP
SPZ
TOLLSIG
IMDSIG
TOLL
GALE
SCRB
SCRA
HPX
CLIPD
CON
SEL
NSP
EVR
SSD
NSD
PDN
PDS
AGE
UNI
WGA
PHC
STA
ACN
EXP
CNV
DUP
SYN

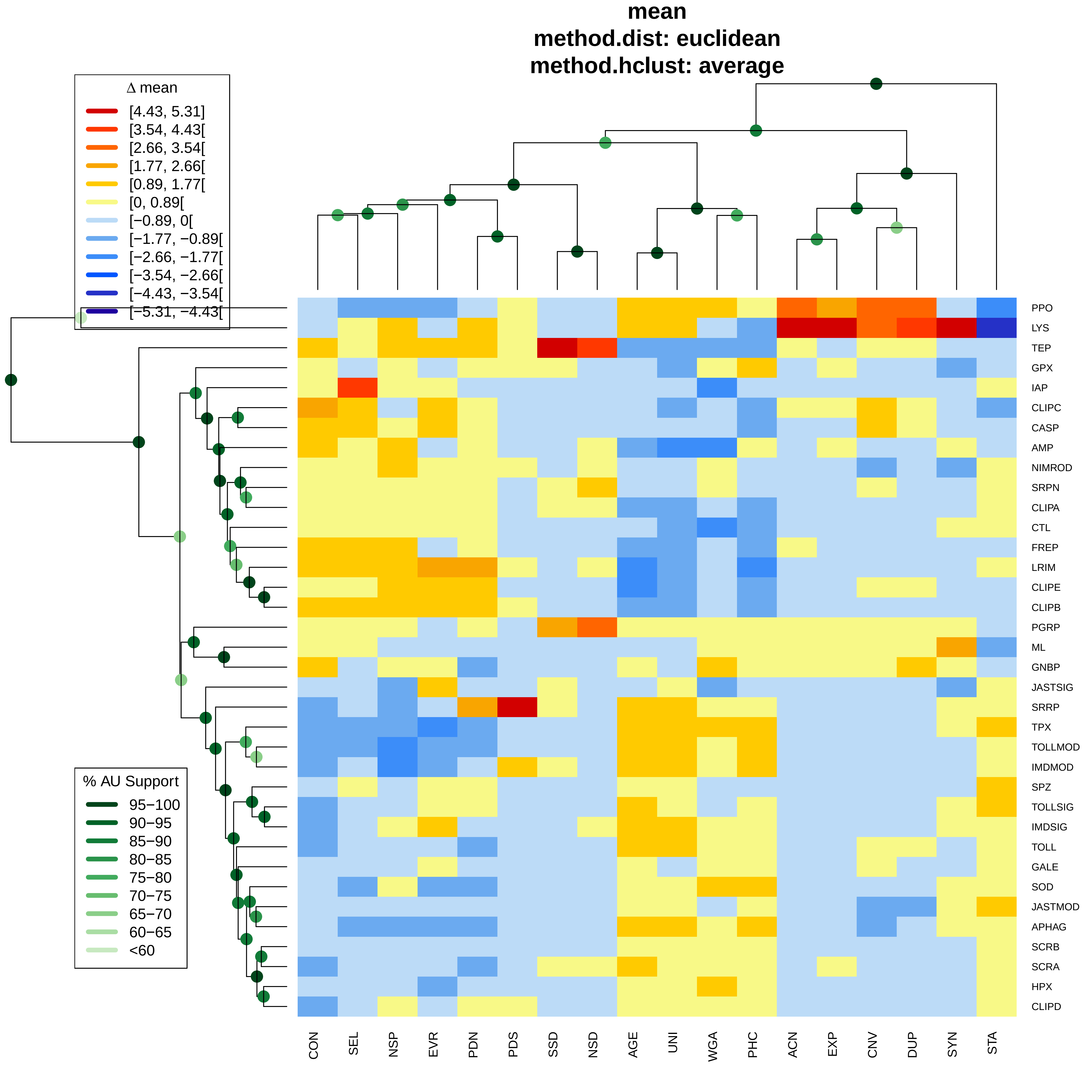
mean
method.dist: euclidean
method.hclust: average
Δ mean
[4.43, 5.31]
[3.54, 4.43[
[2.66, 3.54[
[1.77, 2.66[
[0.89, 1.77[
[0, 0.89[
[−0.89, 0[
[−1.77, −0.89[
[−2.66, −1.77[
[−3.54, −2.66[
[−4.43, −3.54[
[−5.31, −4.43[
% AU Support
95−100
90−95
85−90
80−85
75−80
70−75
65−70
60−65
<60
PPO
LYS
TEP
GPX
IAP
CLIPC
CASP
AMP
NIMROD
SRPN
CLIPA
CTL
FREP
LRIM
CLIPE
CLIPB
PGRP
ML
GNBP
JASTSIG
SRRP
TPX
TOLLMOD
IMDMOD
SPZ
TOLLSIG
IMDSIG
TOLL
GALE
SOD
JASTMOD
APHAG
SCRB
SCRA
HPX
CLIPD
CON
SEL
NSP
EVR
PDN
PDS
SSD
NSD
AGE
UNI
WGA
PHC
ACN
EXP
CNV
DUP
SYN
STA
