## Additional File 3 for "Functional constraints on insect immune system components govern their evolutionary trajectories"

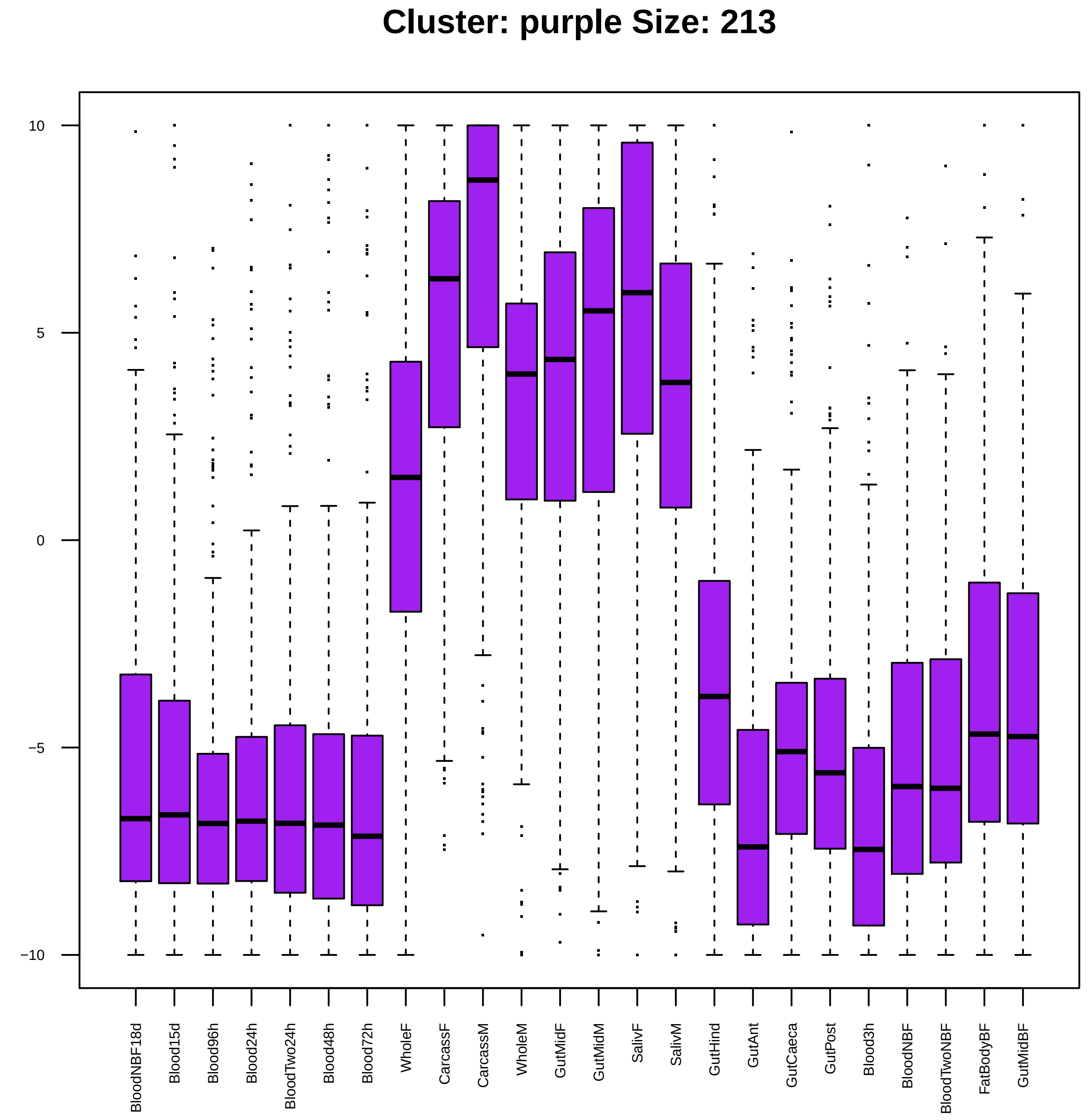

Cluster: purple Size: 213
10
5
0
−5
−10
BloodNBF18d
Blood15d
Blood96h
Blood24h
BloodTwo24h
Blood48h
Blood72h
WholeF
CarcassF
CarcassM
WholeM
GutMidF
GutMidM
SalivF
SalivM
GutHind
GutAnt
GutCaeca
GutPost
Blood3h
BloodNBF
BloodTwoNBF
FatBodyBF
GutMidBF

| | GO.ID | BPCluster: purple Size: 213 | Annotated | Significant | Expected | Rank in ClassicF | Weight01F | ClassicF |
|---|---|---|---|---|---|---|---|---|
| *1* | GO:0048871 | multicellular organismal homeostasis | 19 | 4 | 0.40 | 8 | 0.00043 | 0.00058 |
| *2* | GO:0030203 | glycosaminoglycan metabolic process | 18 | 4 | 0.38 | 7 | 0.00047 | 0.00047 |
| *3* | GO:0006906 | vesicle fusion | 22 | 5 | 0.47 | 4 | 0.00130 | 7.8e−05 |
| *4* | GO:0008333 | endosome to lysosome transport | 11 | 3 | 0.23 | 13 | 0.00135 | 0.00135 |
| *5* | GO:0006026 | aminoglycan catabolic process | 33 | 4 | 0.70 | 23 | 0.00488 | 0.00488 |
| *6* | GO:0051606 | detection of stimulus | 121 | 3 | 2.56 | 758 | 0.00644 | 0.47472 |
| *7* | GO:0006886 | intracellular protein transport | 167 | 9 | 3.54 | 33 | 0.00752 | 0.00870 |
| *8* | GO:0016236 | macroautophagy | 21 | 3 | 0.44 | 35 | 0.00934 | 0.00934 |
| *20* | GO:0006955 | immune response | 95 | 7 | 2.01 | 22 | 0.02440 | 0.00383 |
| *29* | GO:0031347 | regulation of defense response | 40 | 6 | 0.85 | 5 | 0.04060 | 0.00017 |

| | GO.ID | MFCluster: purple Size: 213 | Annotated | Significant | Expected | Rank in ClassicF | Weight01F | ClassicF |
|---|---|---|---|---|---|---|---|---|
| *2* | GO:0016811 | hydrolase activity, acting on carbon−nit... | 32 | 4 | 0.63 | 2 | 0.0034 | 0.0034 |
| *3* | GO:0003924 | GTPase activity | 116 | 7 | 2.28 | 3 | 0.0078 | 0.0078 |

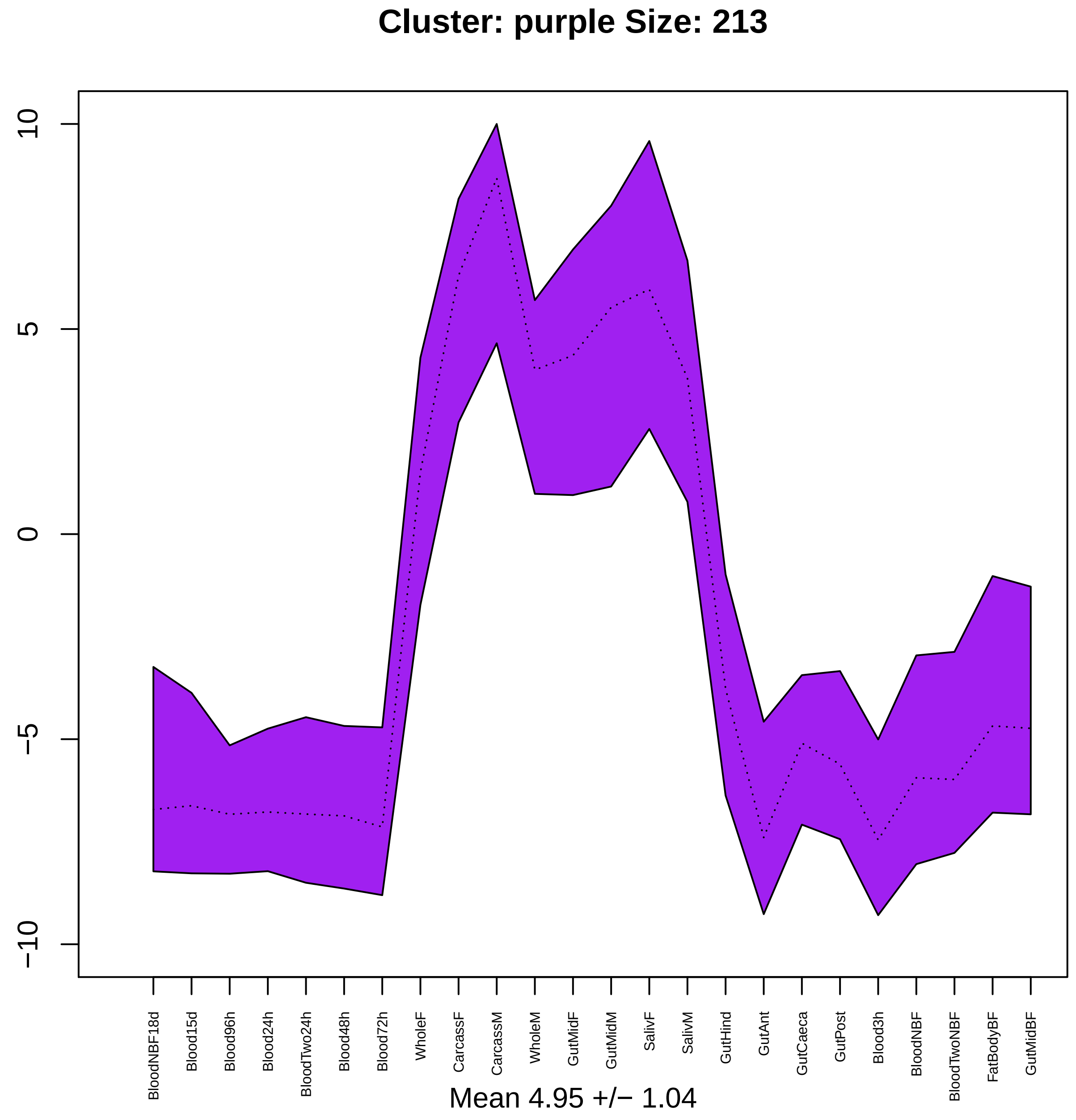

Cluster: purple Size: 213
10
5
0
−5
−10
BloodNBF18d
Blood15d
Blood96h
Blood24h
BloodTwo24h
Blood48h
Blood72h
WholeF
CarcassF
CarcassM
WholeM
GutMidF
GutMidM
SalivF
SalivM
GutHind
GutAnt
GutCaeca
GutPost
Blood3h
BloodNBF
BloodTwoNBF
FatBodyBF
GutMidBF
Mean 4.95 +/− 1.04

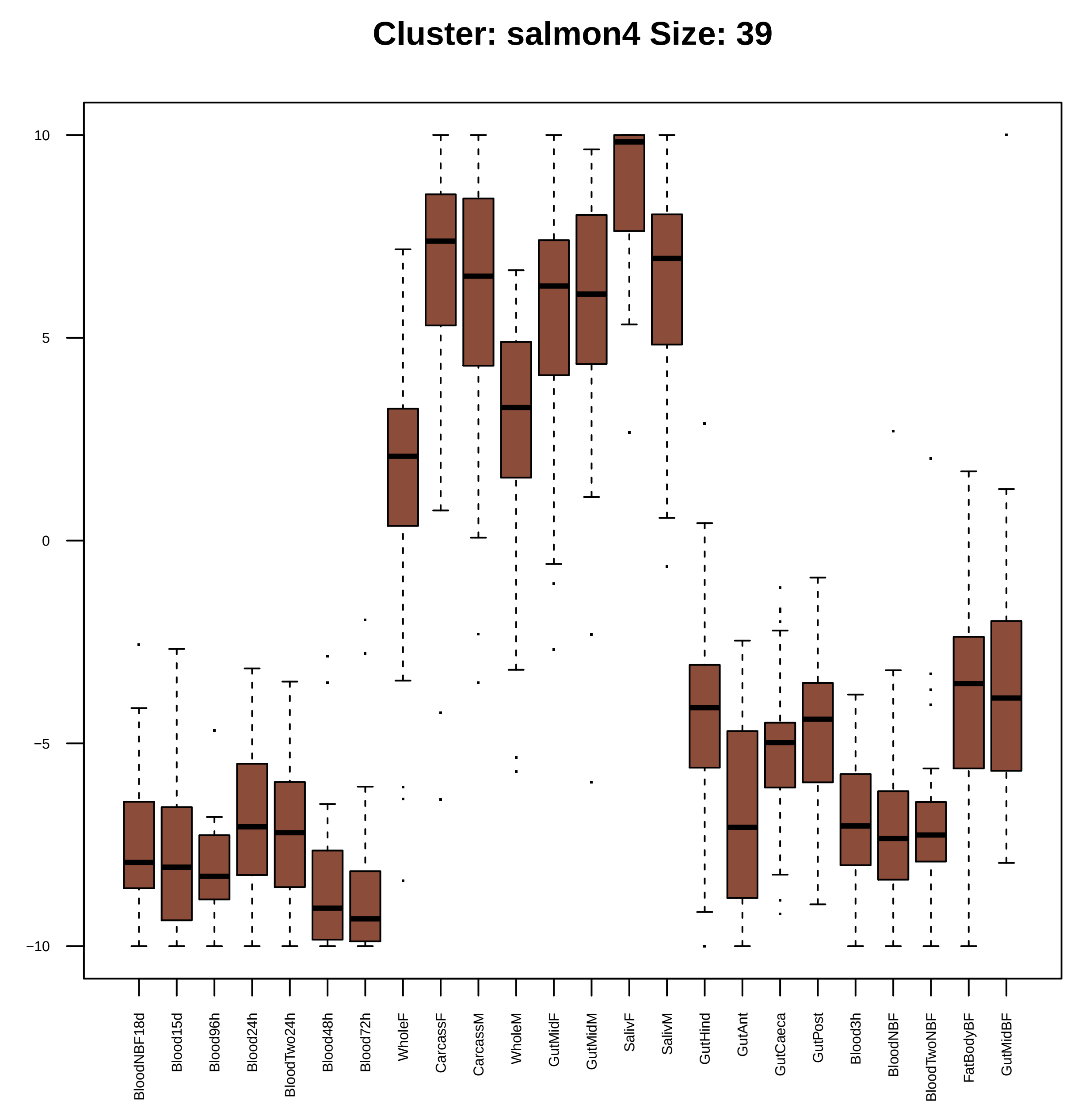
Cluster: salmon4 Size: 39
10
5
0
−5
−10
BloodNBF18d
Blood15d
Blood96h
Blood24h
BloodTwo24h
Blood48h
Blood72h
WholeF
CarcassF
CarcassM
WholeM
GutMidF
GutMidM
SalivF
SalivM
GutHind
GutAnt
GutCaeca
GutPost
Blood3h
BloodNBF
BloodTwoNBF
FatBodyBF
GutMidBF

| | GO.ID | BPCluster: salmon4 Size: 39 | Annotated | Significant | Expected | Rank in ClassicF | Weight01F | ClassicF |
|---|---|---|---|---|---|---|---|---|
| *1* | GO:0006904 | vesicle docking involved in exocytosis | 15 | 2 | 0.05 | 1 | 0.0011 | 0.0011 |
| *3* | GO:0007033 | vacuole organization | 20 | 2 | 0.07 | 5 | 0.0249 | 0.0019 |
| *7* | GO:0006906 | vesicle fusion | 22 | 2 | 0.07 | 6 | 0.0340 | 0.0023 |

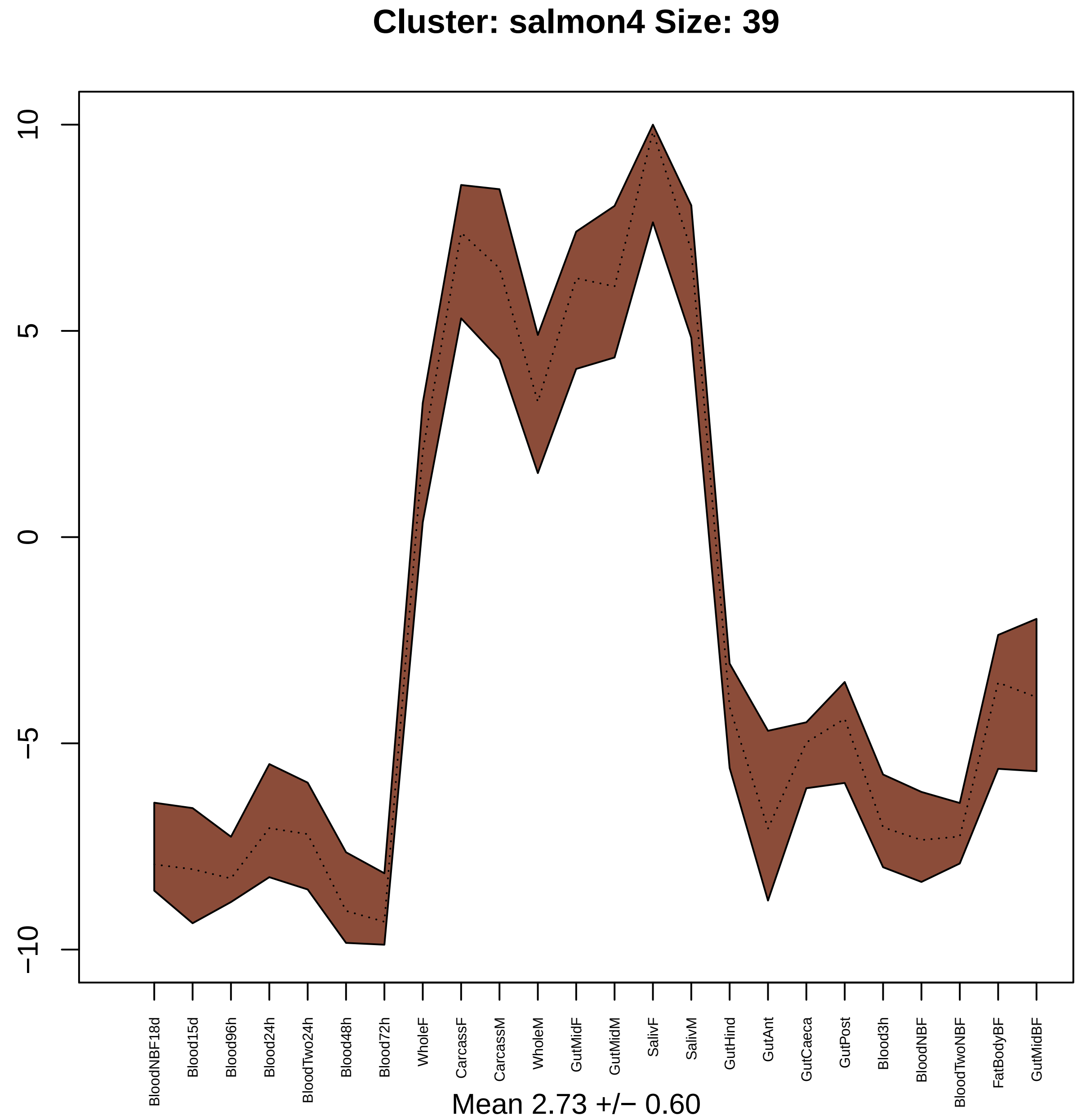

Cluster: salmon4 Size: 39
10
5
0
−5
−10
BloodNBF18d
Blood15d
Blood96h
Blood24h
BloodTwo24h
Blood48h
Blood72h
WholeF
CarcassF
CarcassM
WholeM
GutMidF
GutMidM
SalivF
SalivM
GutHind
GutAnt
GutCaeca
GutPost
Blood3h
BloodNBF
BloodTwoNBF
FatBodyBF
GutMidBF
Mean 2.73 +/− 0.60

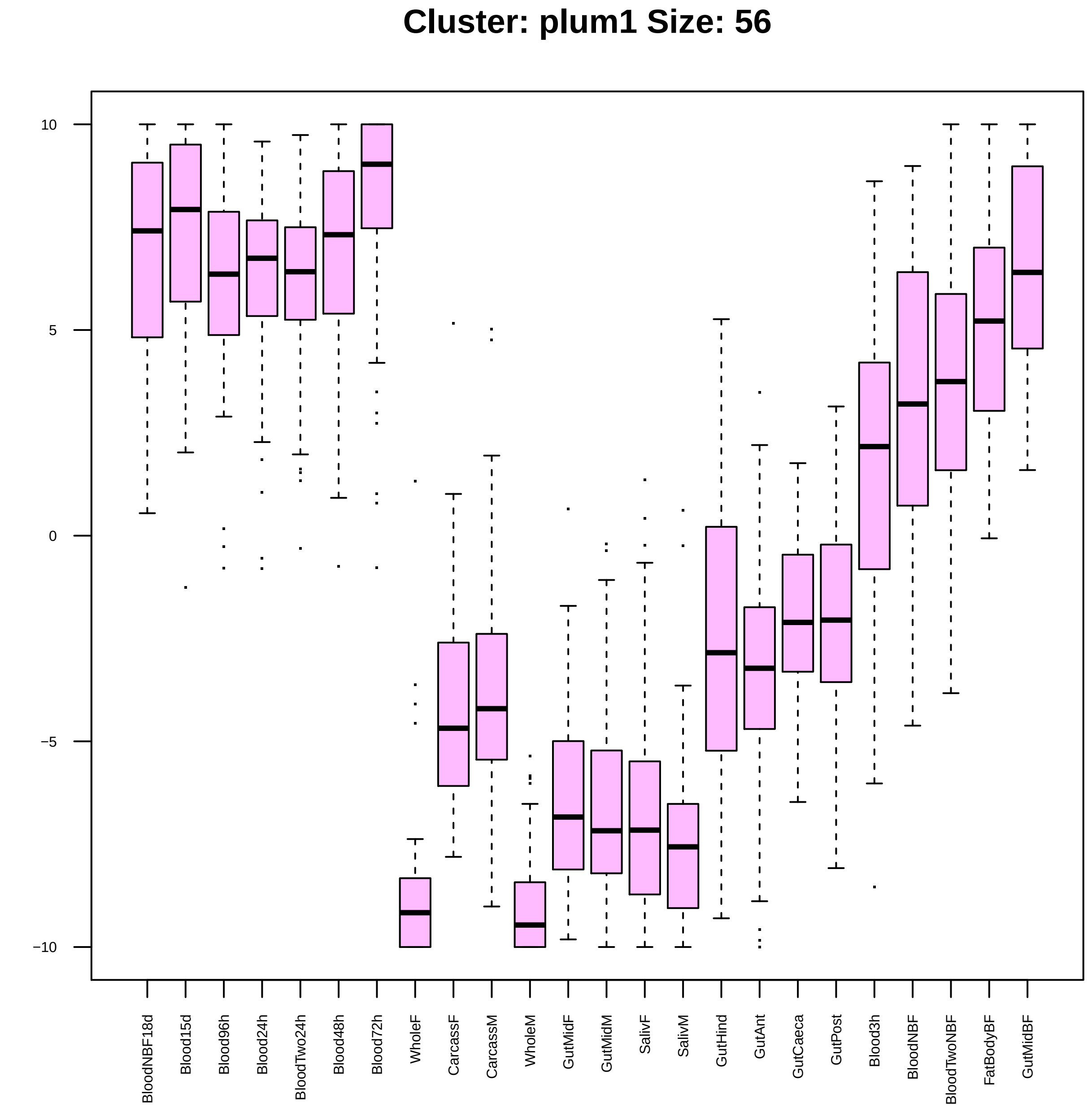

Cluster: plum1 Size: 56
10
5
0
−5
−10
BloodNBF18d
Blood15d
Blood96h
Blood24h
BloodTwo24h
Blood48h
Blood72h
WholeF
CarcassF
CarcassM
WholeM
GutMidF
GutMidM
SalivF
SalivM
GutHind
GutAnt
GutCaeca
GutPost
Blood3h
BloodNBF
BloodTwoNBF
FatBodyBF
GutMidBF

| | GO.ID | BPCluster: plum1 Size: 56 | Annotated | Significant | Expected | Rank in ClassicF | Weight01F | ClassicF |
|---|---|---|---|---|---|---|---|---|
| *1* | GO:0007277 | pole cell development | 10 | 2 | 0.05 | 2 | 0.0011 | 0.00107 |
| *2* | GO:0070507 | regulation of microtubule cytoskeleton o... | 19 | 2 | 0.10 | 8 | 0.0040 | 0.00396 |
| *3* | GO:0016325 | oocyte microtubule cytoskeleton organiza... | 22 | 2 | 0.11 | 10 | 0.0053 | 0.00529 |
| *4* | GO:0060811 | intracellular mRNA localization involved... | 30 | 2 | 0.15 | 19 | 0.0146 | 0.00972 |
| *9* | GO:0008298 | intracellular mRNA localization | 39 | 3 | 0.20 | 1 | 0.0372 | 0.00093 |
| *16* | GO:0007143 | female meiotic nuclear division | 26 | 2 | 0.13 | 17 | 0.0616 | 0.00736 |

| | GO.ID | MFCluster: plum1 Size: 56 | Annotated | Significant | Expected | Rank in ClassicF | Weight01F | ClassicF |
|---|---|---|---|---|---|---|---|---|
| *1* | GO:0008270 | zinc ion binding | 522 | 8 | 3.04 | 2 | 0.0094 | 0.0094 |
| *2* | GO:0005515 | protein binding | 2143 | 21 | 12.49 | 1 | 0.0112 | 0.0045 |

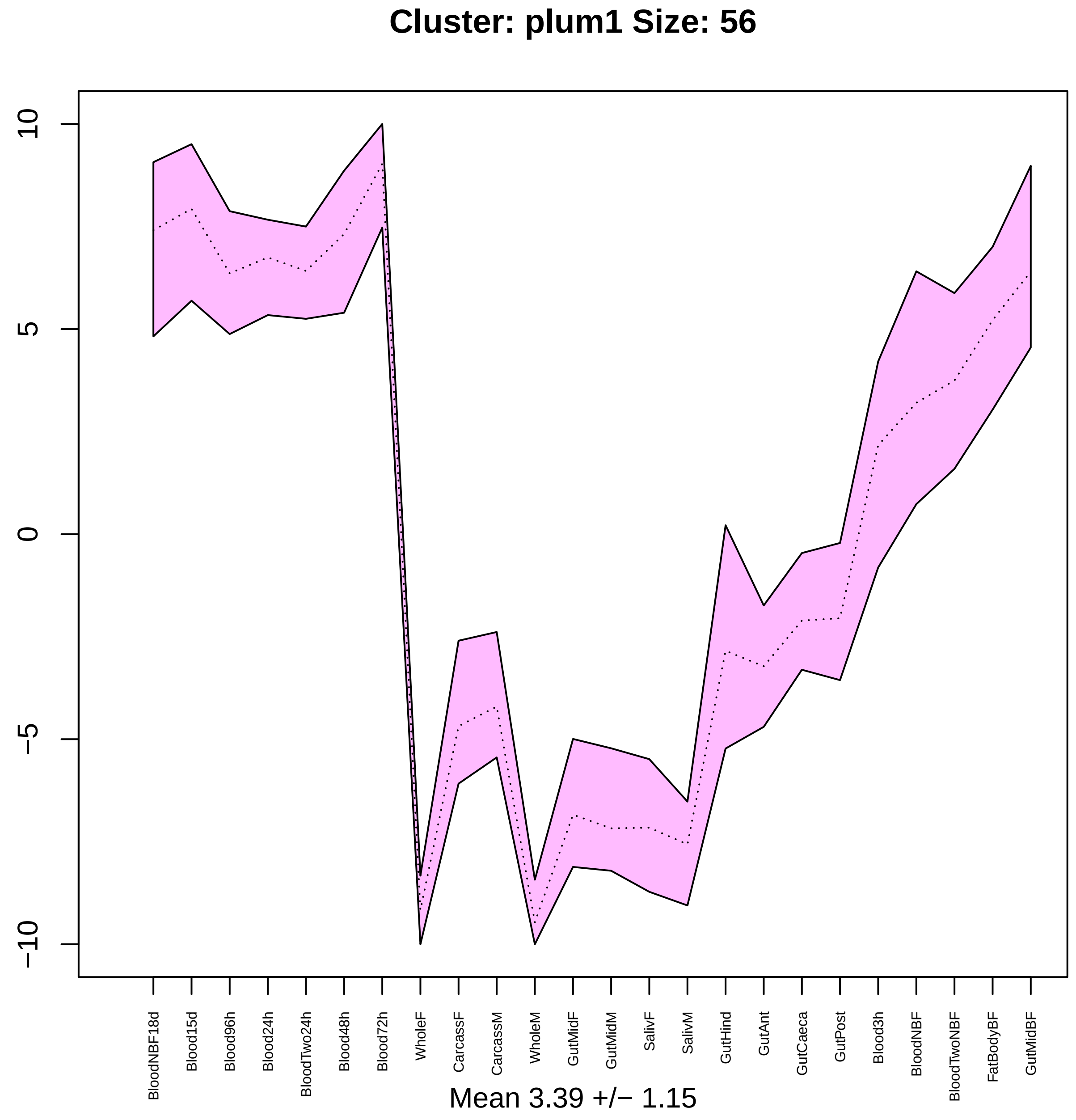

Cluster: plum1 Size: 56
10
5
0
−5
−10
BloodNBF18d
Blood15d
Blood96h
Blood24h
BloodTwo24h
Blood48h
Blood72h
WholeF
CarcassF
CarcassM
WholeM
GutMidF
GutMidM
SalivF
SalivM
GutHind
GutAnt
GutCaeca
GutPost
Blood3h
BloodNBF
BloodTwoNBF
FatBodyBF
GutMidBF
Mean 3.39 +/− 1.15

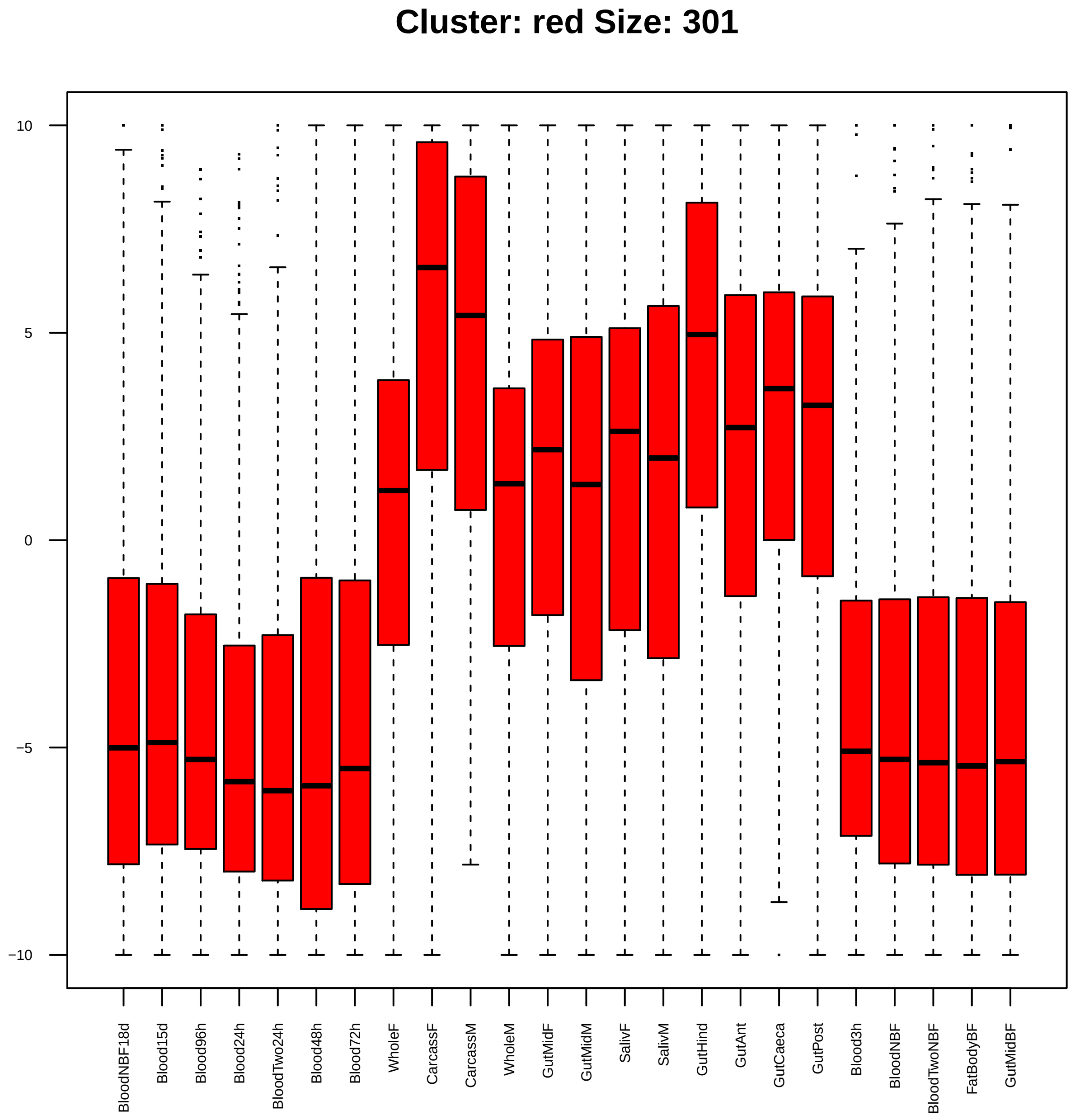

Cluster: red Size: 301
10
5
0
−5
−10
BloodNBF18d
Blood15d
Blood96h
Blood24h
BloodTwo24h
Blood48h
Blood72h
WholeF
CarcassF
CarcassM
WholeM
GutMidF
GutMidM
SalivF
SalivM
GutHind
GutAnt
GutCaeca
GutPost
Blood3h
BloodNBF
BloodTwoNBF
FatBodyBF
GutMidBF

| | GO.ID | BPCluster: red Size: 301 | Annotated | Significant | Expected | Rank in ClassicF | Weight01F | ClassicF |
|---|---|---|---|---|---|---|---|---|
| *1* | GO:0006412 | translation | 289 | 21 | 8.70 | 2 | 0.00017 | 0.00014 |
| *2* | GO:0015893 | drug transport | 11 | 3 | 0.33 | 7 | 0.00371 | 0.00371 |
| *10* | GO:0007098 | centrosome cycle | 60 | 6 | 1.81 | 8 | 0.02960 | 0.00894 |

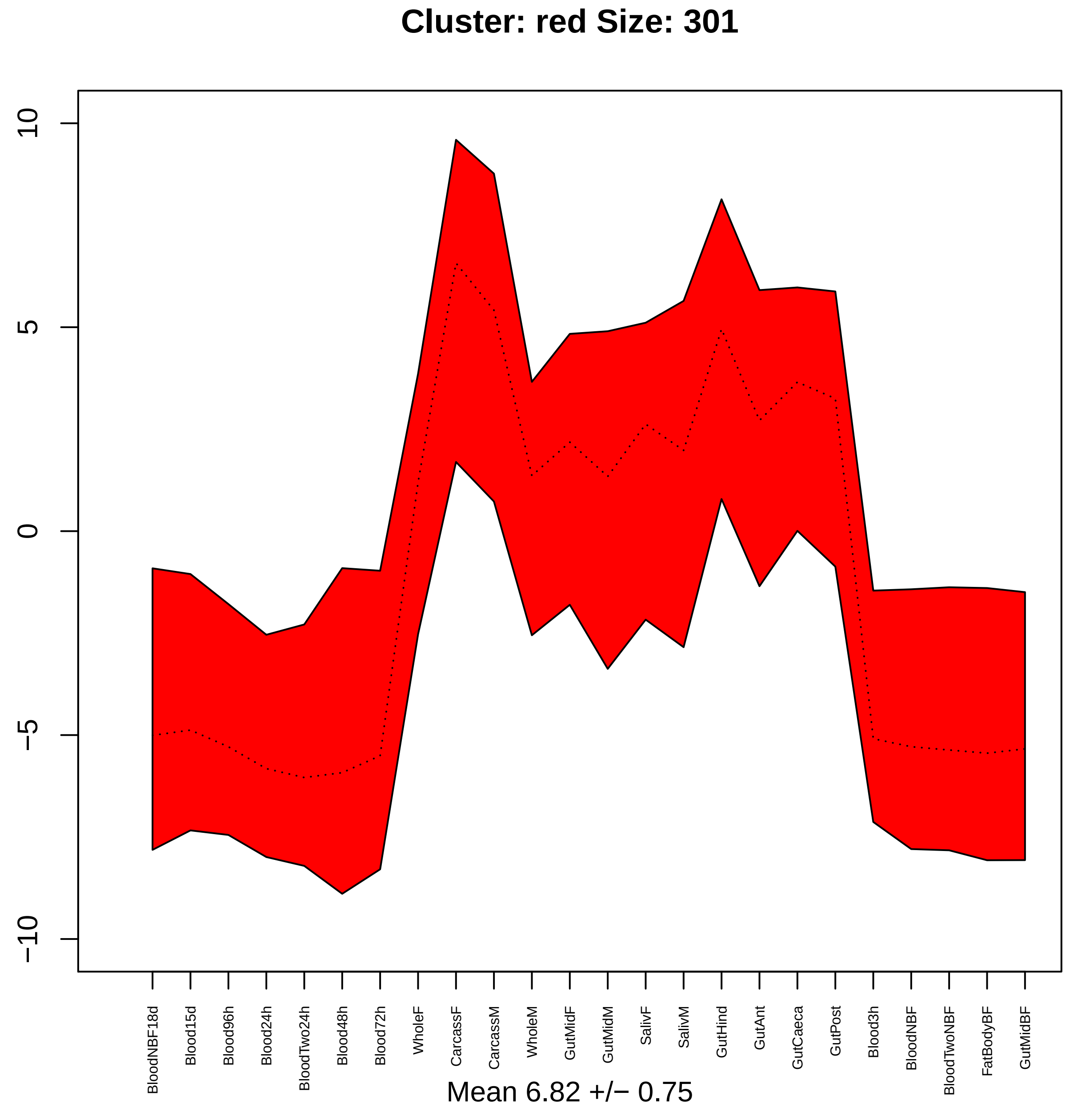

Cluster: red Size: 301
10
5
0
−5
−10
BloodNBF18d
Blood15d
Blood96h
Blood24h
BloodTwo24h
Blood48h
Blood72h
WholeF
CarcassF
CarcassM
WholeM
GutMidF
GutMidM
SalivF
SalivM
GutHind
GutAnt
GutCaeca
GutPost
Blood3h
BloodNBF
BloodTwoNBF
FatBodyBF
GutMidBF
Mean 6.82 +/− 0.75

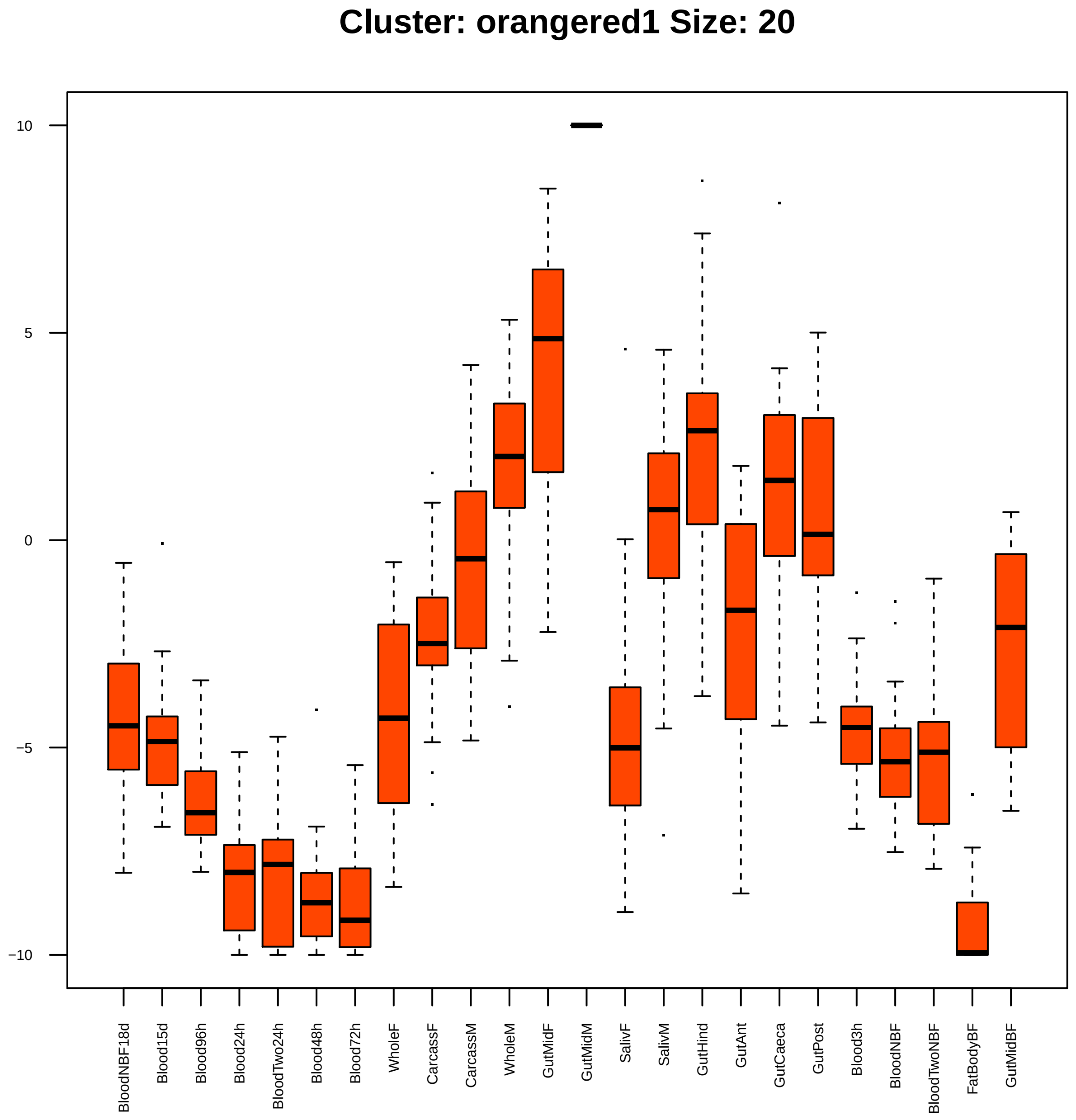

Cluster: orangered1 Size: 20
10
5
0
−5
−10
BloodNBF18d
Blood15d
Blood96h
Blood24h
BloodTwo24h
Blood48h
Blood72h
WholeF
CarcassF
CarcassM
WholeM
GutMidF
GutMidM
SalivF
SalivM
GutHind
GutAnt
GutCaeca
GutPost
Blood3h
BloodNBF
BloodTwoNBF
FatBodyBF
GutMidBF

| | GO.ID | BPCluster: orangered1 Size: 20 | Annotated | Significant | Expected | Rank in ClassicF | Weight01F | ClassicF |
|---|---|---|---|---|---|---|---|---|
| *1* | GO:0008152 | metabolic process | 4285 | 14 | 12.10 | 58 | 0.0053 | 0.2454 |
| *28* | GO:0030534 | adult behavior | 49 | 2 | 0.14 | 1 | 1.0000 | 0.0082 |

| | GO.ID | MFCluster: orangered1 Size: 20 | Annotated | Significant | Expected | Rank in ClassicF | Weight01F | ClassicF |
|---|---|---|---|---|---|---|---|---|
| 3 | GO:0016705 | oxidoreductase activity, acting on paire... | 133 | 4 | 0.30 | 4 | 0.00017 | 0.00017 |
| 4 | GO:0020037 | heme binding | 133 | 4 | 0.30 | 5 | 0.00017 | 0.00017 |
| 16 | GO:0003824 | catalytic activity | 3160 | 13 | 7.01 | 7 | 0.22045 | 0.00257 |
| 23 | GO:0046906 | tetrapyrrole binding | 134 | 4 | 0.30 | 6 | 1.00000 | 0.00017 |

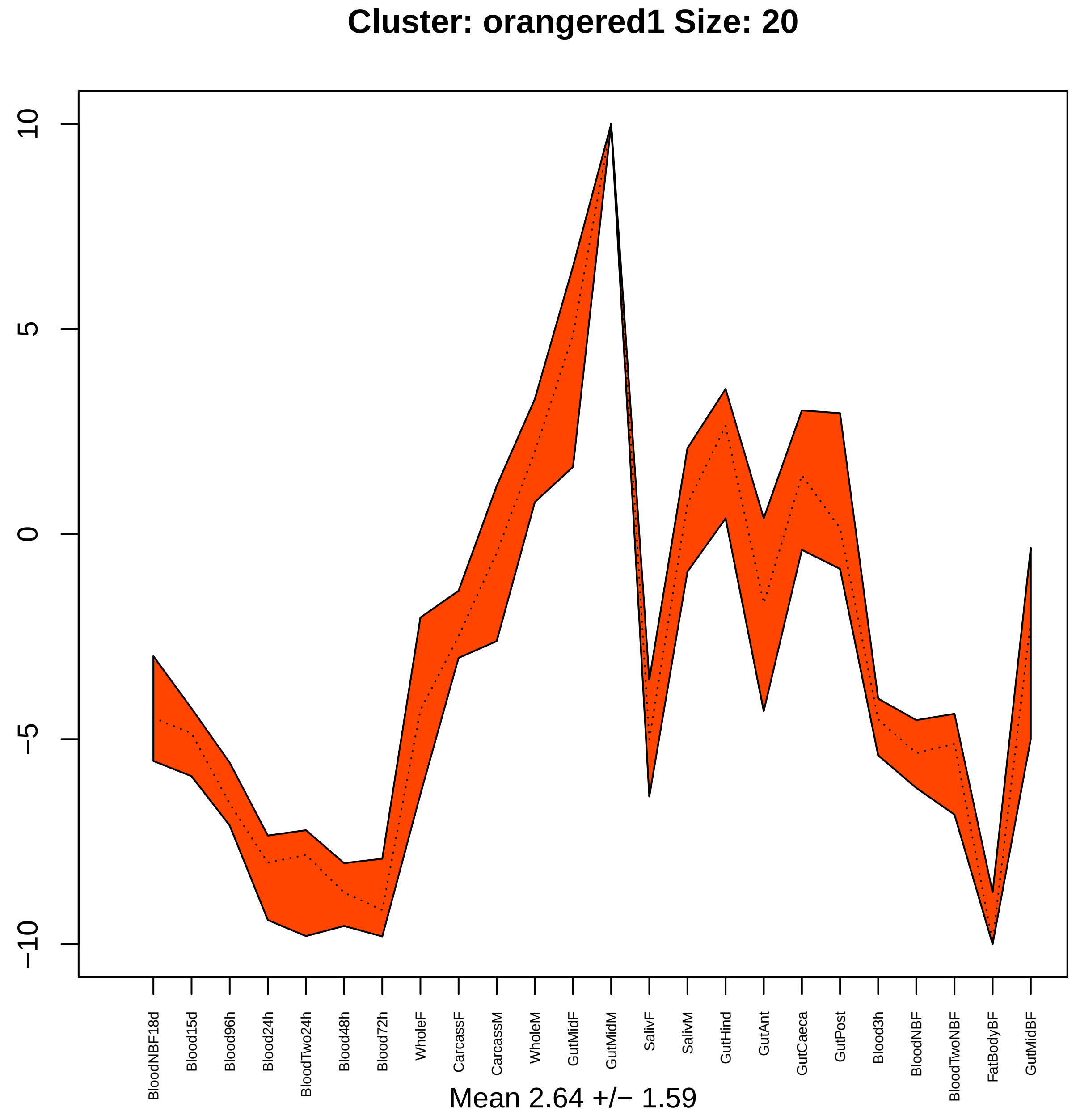
Cluster: orangered1 Size: 20
10
5
0
−5
−10
BloodNBF18d
Blood15d
Blood96h
Blood24h
BloodTwo24h
Blood48h
Blood72h
WholeF
CarcassF
CarcassM
WholeM
GutMidF
GutMidM
SalivF
SalivM
GutHind
GutAnt
GutCaeca
GutPost
Blood3h
BloodNBF
BloodTwoNBF
FatBodyBF
GutMidBF
Mean 2.64 +/− 1.59

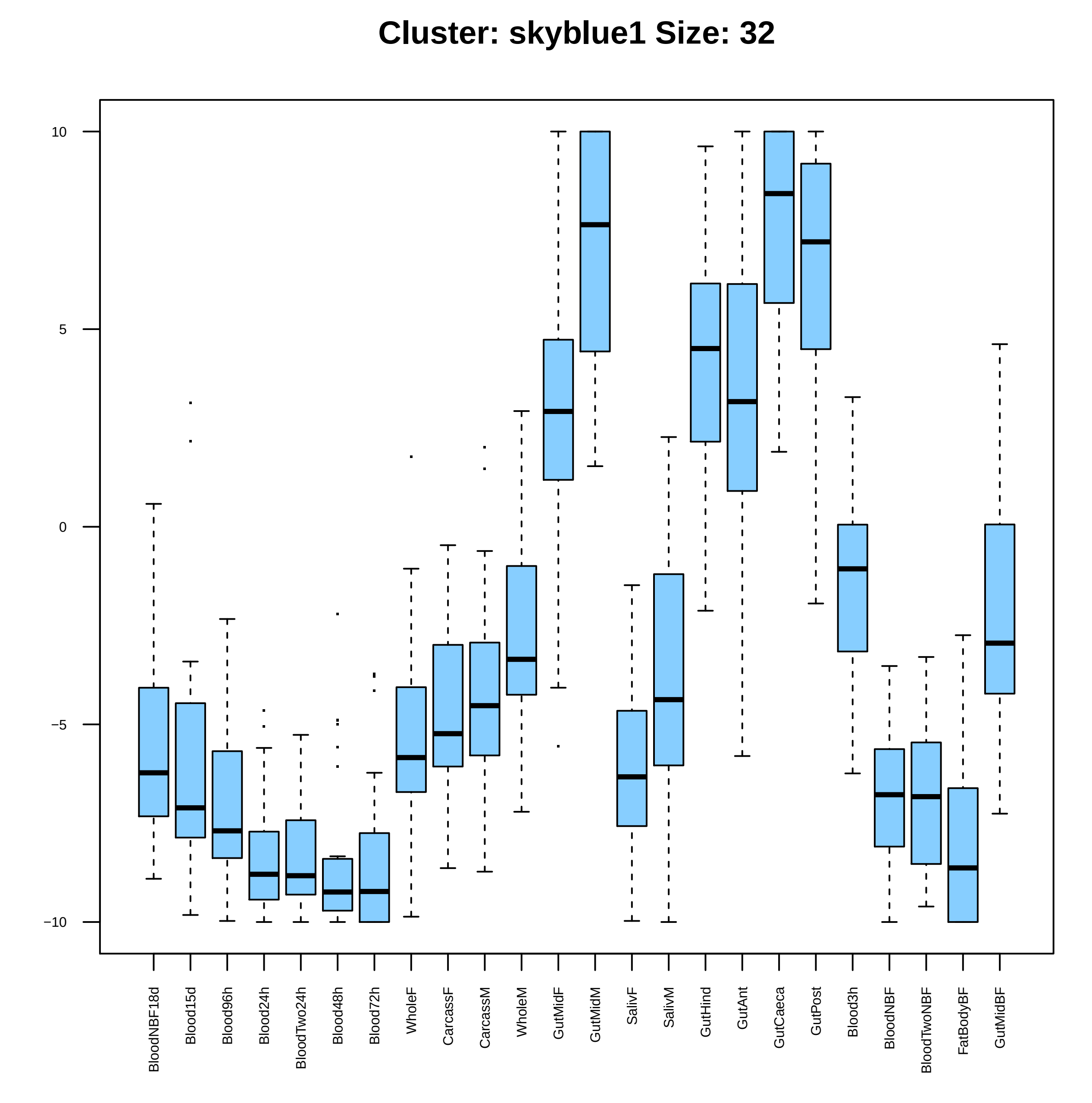

Cluster: skyblue1 Size: 32
10
5
0
−5
−10
BloodNBF18d
Blood15d
Blood96h
Blood24h
BloodTwo24h
Blood48h
Blood72h
WholeF
CarcassF
CarcassM
WholeM
GutMidF
GutMidM
SalivF
SalivM
GutHind
GutAnt
GutCaeca
GutPost
Blood3h
BloodNBF
BloodTwoNBF
FatBodyBF
GutMidBF

| | GO.ID | BPCluster: skyblue1 Size: 32 | Annotated | Significant | Expected | Rank in ClassicF | Weight01F | ClassicF |
|---|---|---|---|---|---|---|---|---|
| *2* | GO:0008152 | metabolic process | 4285 | 22 | 16.8 | 2 | 0.0022 | 0.017 |

| | GO.ID | MFCluster: skyblue1 Size: 32 | Annotated | Significant | Expected | Rank in ClassicF | Weight01F | ClassicF |
|---|---|---|---|---|---|---|---|---|
| *2* | GO:0004497 | monooxygenase activity | 110 | 3 | 0.4 | 5 | 0.007 | 0.007 |

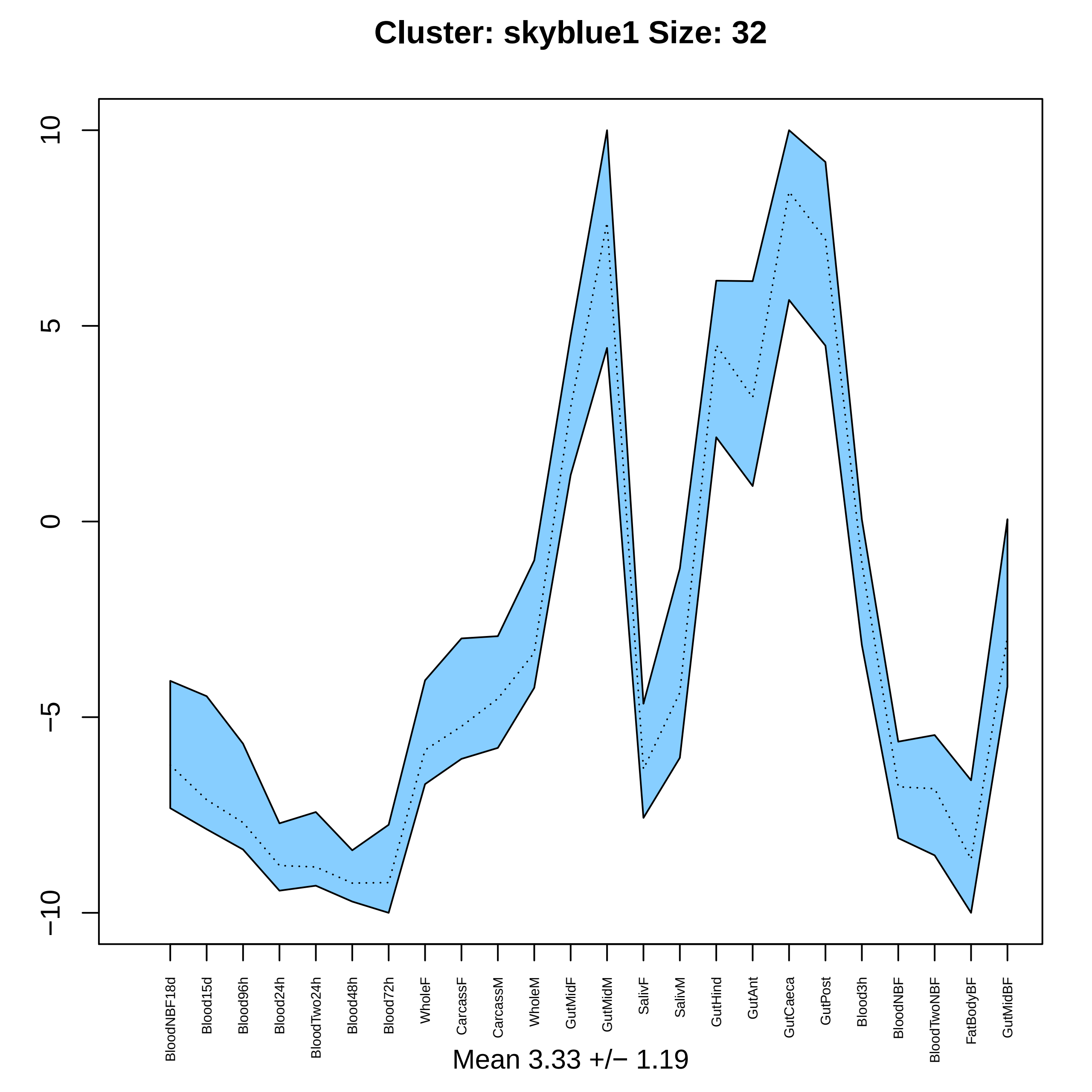
Cluster: skyblue1 Size: 32
10
5
0
−5
−10
BloodNBF18d
Blood15d
Blood96h
Blood24h
BloodTwo24h
Blood48h
Blood72h
WholeF
CarcassF
CarcassM
WholeM
GutMidF
GutMidM
SalivF
SalivM
GutHind
GutAnt
GutCaeca
GutPost
Blood3h
BloodNBF
BloodTwoNBF
FatBodyBF
GutMidBF
Mean 3.33 +/− 1.19

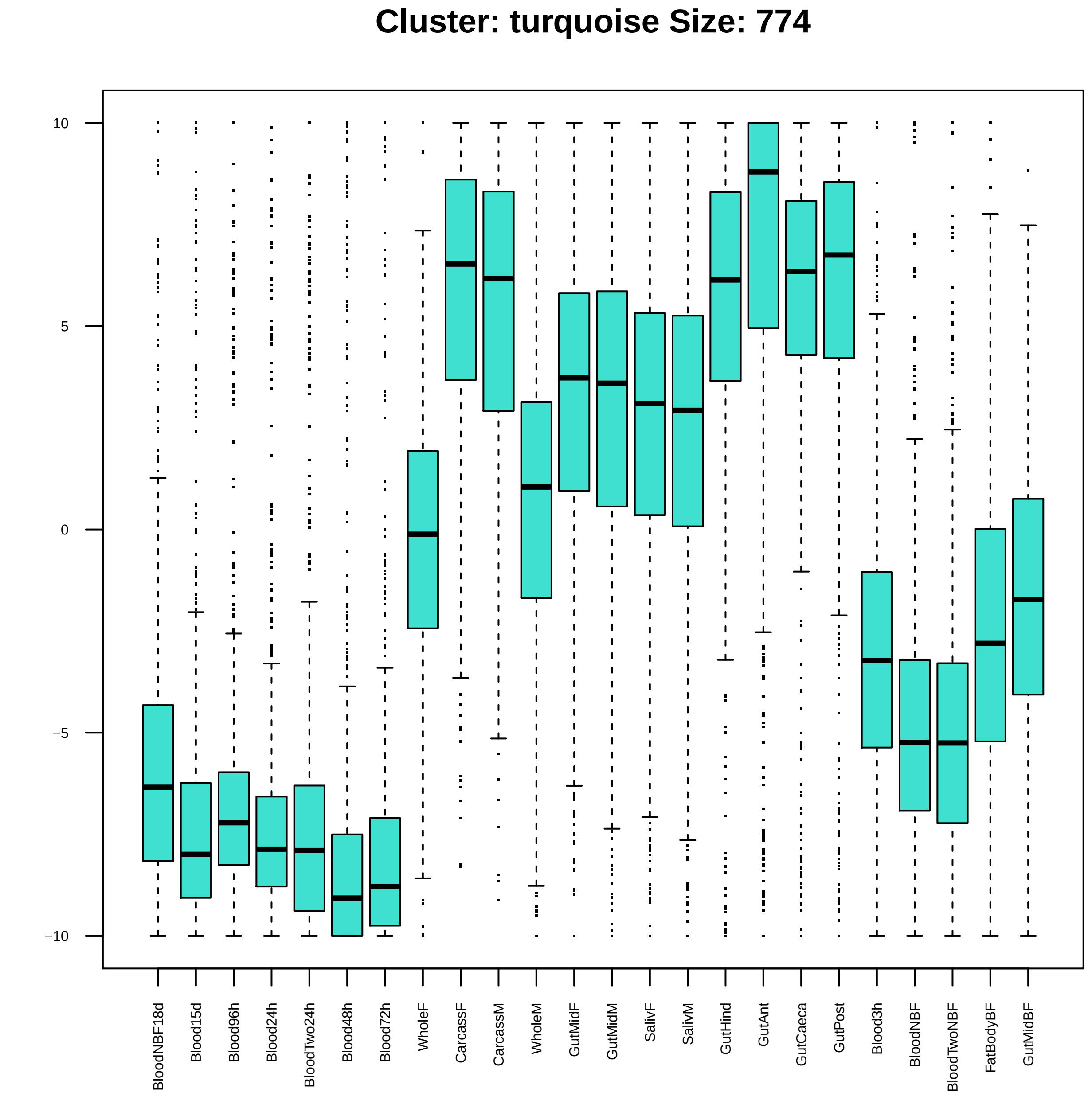
Cluster: turquoise Size: 774
10
5
0
−5
−10
BloodNBF18d
Blood15d
Blood96h
Blood24h
BloodTwo24h
Blood48h
Blood72h
WholeF
CarcassF
CarcassM
WholeM
GutMidF
GutMidM
SalivF
SalivM
GutHind
GutAnt
GutCaeca
GutPost
Blood3h
BloodNBF
BloodTwoNBF
FatBodyBF
GutMidBF

|  | GO.ID | BPCluster: turquoise Size: 774 | Annotated | Significant | Expected | Rank in ClassicF | Weight01F | ClassicF |
|---|---|---|---|---|---|---|---|---|
| *4* | GO:0034220 | ion transmembrane transport | 275 | 32 | 19.92 | 18 | 0.0030 | 0.00478 |
| *5* | GO:0050909 | sensory perception of taste | 56 | 13 | 4.06 | 14 | 0.0035 | 0.00013 |
| *6* | GO:0006066 | alcohol metabolic process | 37 | 9 | 2.68 | 15 | 0.0038 | 0.00101 |
| *7* | GO:0071805 | potassium ion transmembrane transport | 23 | 6 | 1.67 | 19 | 0.0049 | 0.00488 |
| *8* | GO:0050912 | detection of chemical stimulus involved ... | 12 | 4 | 0.87 | 22 | 0.0084 | 0.00843 |

| | GO.ID | MFCluster: turquoise Size: 774 | Annotated | Significant | Expected | Rank in ClassicF | Weight01F | ClassicF |
|---|---|---|---|---|---|---|---|---|
| 6 | GO:0008527 | taste receptor activity | 12 | 5 | 0.90 | 25 | 0.0012 | 0.0012 |
| 7 | GO:0003995 | acyl−CoA dehydrogenase activity | 14 | 5 | 1.05 | 27 | 0.0026 | 0.0026 |
| 8 | GO:0050660 | flavin adenine dinucleotide binding | 58 | 11 | 4.35 | 28 | 0.0034 | 0.0034 |
| 9 | GO:0030594 | neurotransmitter receptor activity | 45 | 16 | 3.38 | 6 | 0.0035 | 6.8e−08 |
| 10 | GO:0005249 | voltage−gated potassium channel activity | 15 | 5 | 1.13 | 29 | 0.0037 | 0.0037 |
| 11 | GO:0003705 | transcription factor activity, RNA polym... | 32 | 7 | 2.40 | 33 | 0.0083 | 0.0083 |
| 12 | GO:0008812 | choline dehydrogenase activity | 18 | 5 | 1.35 | 35 | 0.0088 | 0.0088 |
| 14 | GO:0043565 | sequence−specific DNA binding | 212 | 27 | 15.91 | 30 | 0.0194 | 0.0045 |
| 16 | GO:0042302 | structural constituent of cuticle | 99 | 15 | 7.43 | 32 | 0.0270 | 0.0066 |

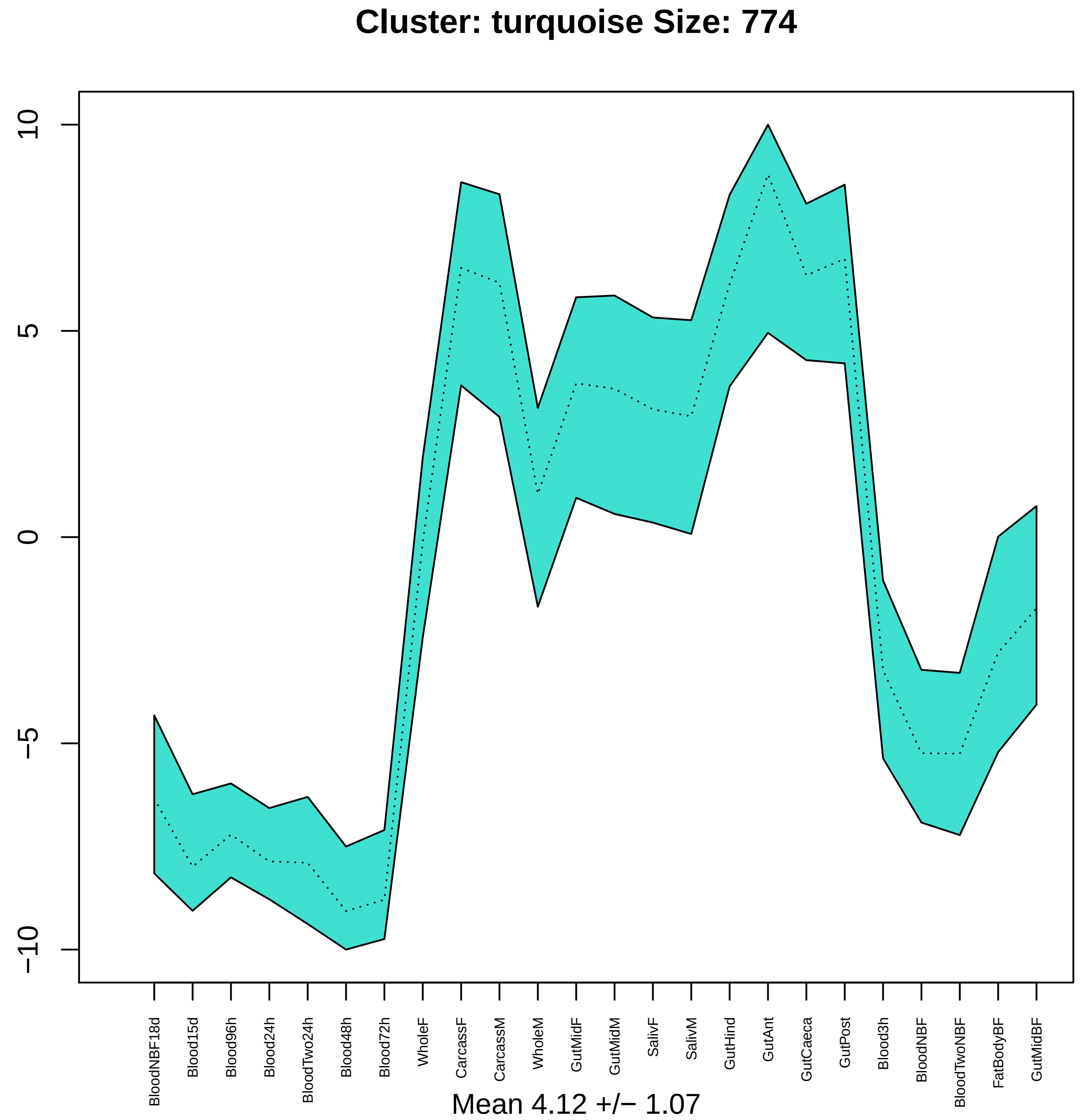
Cluster: turquoise Size: 774
10
5
0
−5
−10
BloodNBF18d
Blood15d
Blood96h
Blood24h
BloodTwo24h
Blood48h
Blood72h
WholeF
CarcassF
CarcassM
WholeM
GutMidF
GutMidM
SalivF
SalivM
GutHind
GutAnt
GutCaeca
GutPost
Blood3h
BloodNBF
BloodTwoNBF
FatBodyBF
GutMidBF
Mean 4.12 +/− 1.07

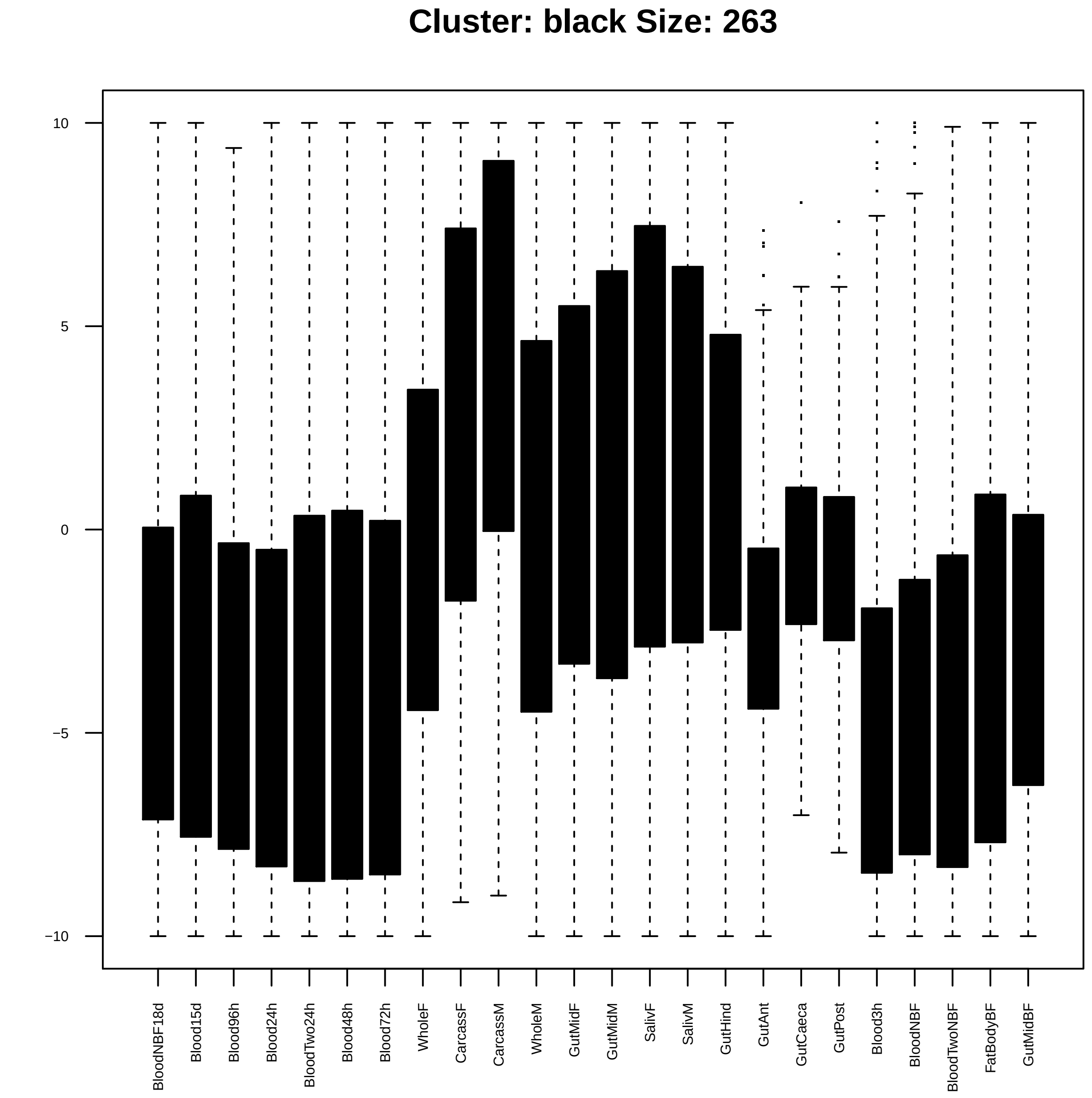

Cluster: black Size: 263
10
5
0
−5
−10
BloodNBF18d
Blood15d
Blood96h
Blood24h
BloodTwo24h
Blood48h
Blood72h
WholeF
CarcassF
CarcassM
WholeM
GutMidF
GutMidM
SalivF
SalivM
GutHind
GutAnt
GutCaeca
GutPost
Blood3h
BloodNBF
BloodTwoNBF
FatBodyBF
GutMidBF

| | GO.ID | BPCluster: black Size: 263 | Annotated | Significant | Expected | Rank in ClassicF | Weight01F | ClassicF |
|---|---|---|---|---|---|---|---|---|
| *1* | GO:0046580 | negative regulation of Ras protein signa... | 12 | 3 | 0.32 | 1 | 0.0035 | 0.0035 |
| *2* | GO:0045475 | locomotor rhythm | 24 | 4 | 0.65 | 3 | 0.0036 | 0.0036 |
| *3* | GO:0006904 | vesicle docking involved in exocytosis | 15 | 3 | 0.40 | 4 | 0.0069 | 0.0069 |
| *4* | GO:0030163 | protein catabolic process | 111 | 6 | 2.99 | 88 | 0.0098 | 0.0788 |

| | GO.ID | MFCluster: black Size: 263 | Annotated | Significant | Expected | Rank in ClassicF | Weight01F | ClassicF |
|---|---|---|---|---|---|---|---|---|
| *1* | GO:0004520 | endodeoxyribonuclease activity | 12 | 3 | 0.30 | 1 | 0.0029 | 0.0029 |
| *2* | GO:0016893 | endonuclease activity, active with eithe... | 16 | 3 | 0.40 | 3 | 0.0059 | 0.0069 |
| *18* | GO:0019887 | protein kinase regulator activity | 29 | 4 | 0.73 | 2 | 0.0725 | 0.0056 |

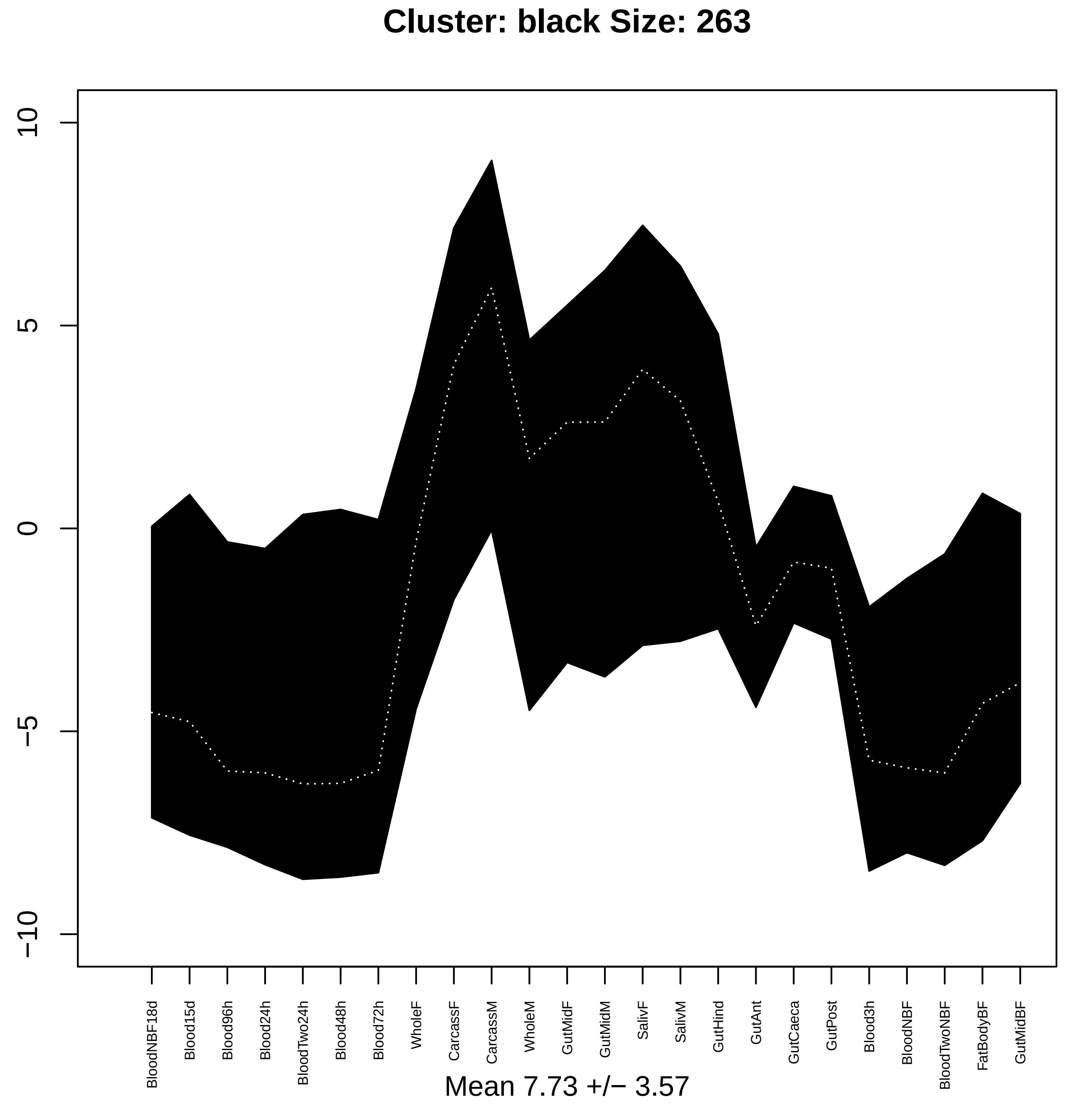

Cluster: black Size: 263
10
5
0
−5
−10
BloodNBF18d
Blood15d
Blood96h
Blood24h
BloodTwo24h
Blood48h
Blood72h
WholeF
CarcassF
CarcassM
WholeM
GutMidF
GutMidM
SalivF
SalivM
GutHind
GutAnt
GutCaeca
GutPost
Blood3h
BloodNBF
BloodTwoNBF
FatBodyBF
GutMidBF
Mean 7.73 +/− 3.57

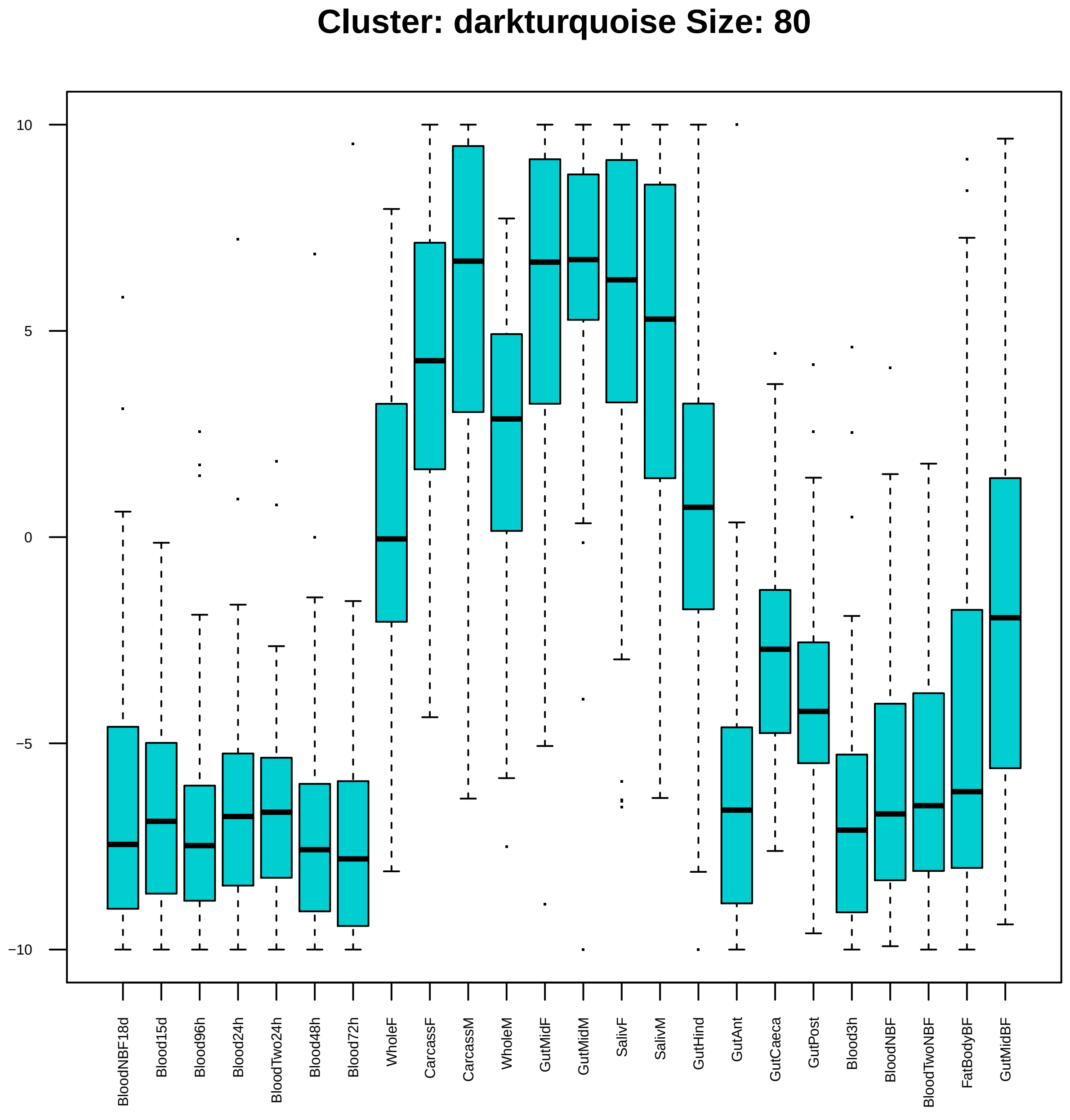

Cluster: darkturquoise Size: 80
10
5
0
−5
−10
BloodNBF18d
Blood15d
Blood96h
Blood24h
BloodTwo24h
Blood48h
Blood72h
WholeF
CarcassF
CarcassM
WholeM
GutMidF
GutMidM
SalivF
SalivM
GutHind
GutAnt
GutCaeca
GutPost
Blood3h
BloodNBF
BloodTwoNBF
FatBodyBF
GutMidBF

| | GO.ID | BPCluster: darkturquoise Size: 80 | Annotated | Significant | Expected | Rank in ClassicF | Weight01F | ClassicF |
|---|---|---|---|---|---|---|---|---|
| *1* | GO:0009792 | embryo development ending in birth or eg... | 96 | 3 | 0.89 | 62 | 0.00082 | 0.05871 |
| *2* | GO:0043901 | negative regulation of multi−organism pr... | 10 | 2 | 0.09 | 5 | 0.00361 | 0.00361 |
| *3* | GO:0034976 | response to endoplasmic reticulum stress | 11 | 2 | 0.10 | 6 | 0.00439 | 0.00439 |
| *4* | GO:0008593 | regulation of Notch signaling pathway | 52 | 4 | 0.48 | 2 | 0.00495 | 0.00129 |
| *28* | GO:0007219 | Notch signaling pathway | 63 | 5 | 0.58 | 1 | 0.09167 | 0.00027 |

| | GO.ID | MFCluster: darkturquoise Size: 80 | Annotated | Significant | Expected | Rank in ClassicF | Weight01F | ClassicF |
|---|---|---|---|---|---|---|---|---|
| *1* | GO:0003730 | mRNA 3'−UTR binding | 16 | 2 | 0.12 | 1 | 0.0064 | 0.0064 |
| *2* | GO:0046982 | protein heterodimerization activity | 55 | 3 | 0.42 | 2 | 0.0083 | 0.0083 |

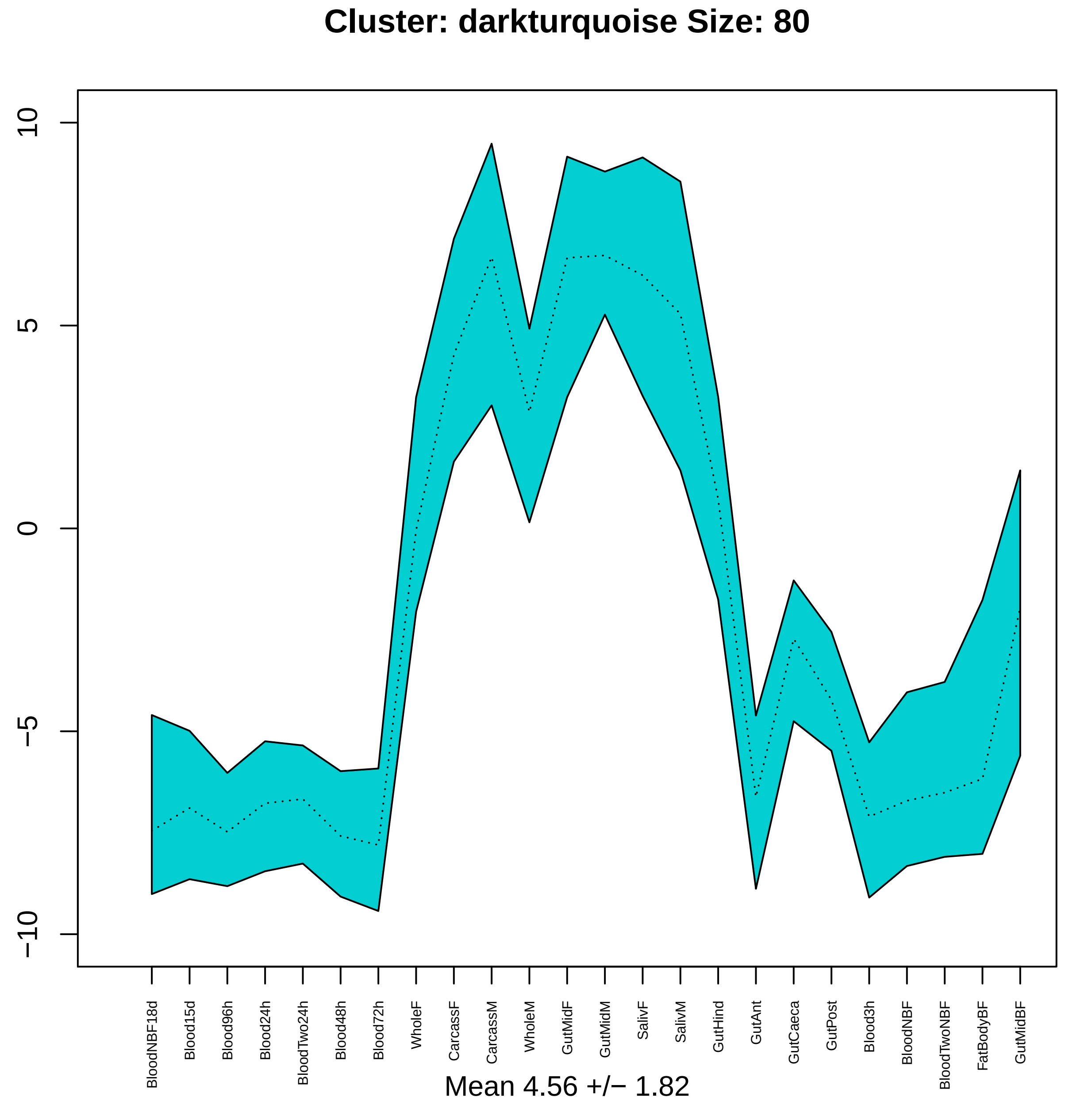

Cluster: darkturquoise Size: 80
10
5
0
−5
−10
BloodNBF18d
Blood15d
Blood96h
Blood24h
BloodTwo24h
Blood48h
Blood72h
WholeF
CarcassF
CarcassM
WholeM
GutMidF
GutMidM
SalivF
SalivM
GutHind
GutAnt
GutCaeca
GutPost
Blood3h
BloodNBF
BloodTwoNBF
FatBodyBF
GutMidBF
Mean 4.56 +/− 1.82

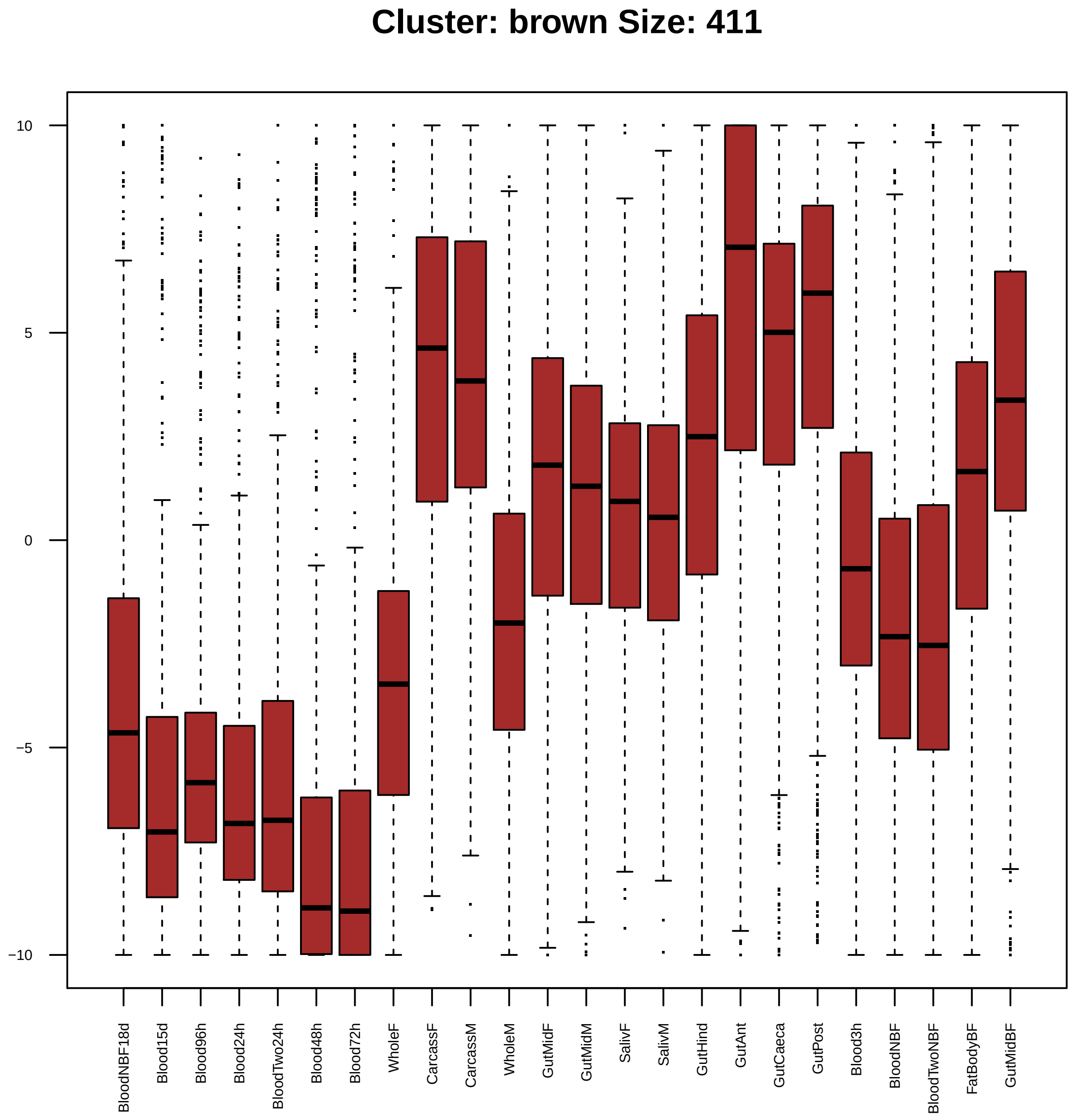

Cluster: brown Size: 411
10
5
0
−5
−10
BloodNBF18d
Blood15d
Blood96h
Blood24h
BloodTwo24h
Blood48h
Blood72h
WholeF
CarcassF
CarcassM
WholeM
GutMidF
GutMidM
SalivF
SalivM
GutHind
GutAnt
GutCaeca
GutPost
Blood3h
BloodNBF
BloodTwoNBF
FatBodyBF
GutMidBF

| | GO.ID | BPCluster: brown Size: 411 | Annotated | Significant | Expected | Rank in ClassicF | Weight01F | ClassicF |
|---|---|---|---|---|---|---|---|---|
| *1* | GO:0006355 | regulation of transcription, DNA−templat... | 501 | 32 | 17.60 | 7 | 0.00017 | 0.00061 |
| *2* | GO:0007186 | G protein−coupled receptor signaling pat... | 228 | 20 | 8.01 | 3 | 0.00211 | 0.00013 |
| *3* | GO:0050911 | detection of chemical stimulus involved ... | 71 | 8 | 2.49 | 23 | 0.00323 | 0.00323 |
| *4* | GO:0003333 | amino acid transmembrane transport | 23 | 4 | 0.81 | 35 | 0.00776 | 0.00776 |
| *5* | GO:0034765 | regulation of ion transmembrane transpor... | 23 | 4 | 0.81 | 36 | 0.00776 | 0.00776 |

| | GO.ID | MFCluster: brown Size: 411 | Annotated | Significant | Expected | Rank in ClassicF | Weight01F | ClassicF |
|---|---|---|---|---|---|---|---|---|
| 1 | GO:0022843 | voltage−gated cation channel activity | 23 | 5 | 0.87 | 8 | 0.0026 | 0.00144 |
| 2 | GO:0005262 | calcium channel activity | 17 | 4 | 0.64 | 11 | 0.0032 | 0.00324 |
| 3 | GO:0000976 | transcription regulatory region sequence... | 45 | 8 | 1.70 | 1 | 0.0039 | 0.00024 |
| 4 | GO:0004984 | olfactory receptor activity | 69 | 8 | 2.61 | 14 | 0.0043 | 0.00430 |
| 5 | GO:0005549 | odorant binding | 118 | 11 | 4.47 | 15 | 0.0049 | 0.00494 |
| 6 | GO:0003700 | DNA−binding transcription factor activit... | 282 | 20 | 10.68 | 16 | 0.0049 | 0.00497 |
| 9 | GO:0004930 | G protein−coupled receptor activity | 171 | 14 | 6.48 | 17 | 0.0222 | 0.00526 |
| 18 | GO:0043565 | sequence−specific DNA binding | 212 | 18 | 8.03 | 7 | 0.0824 | 0.00108 |

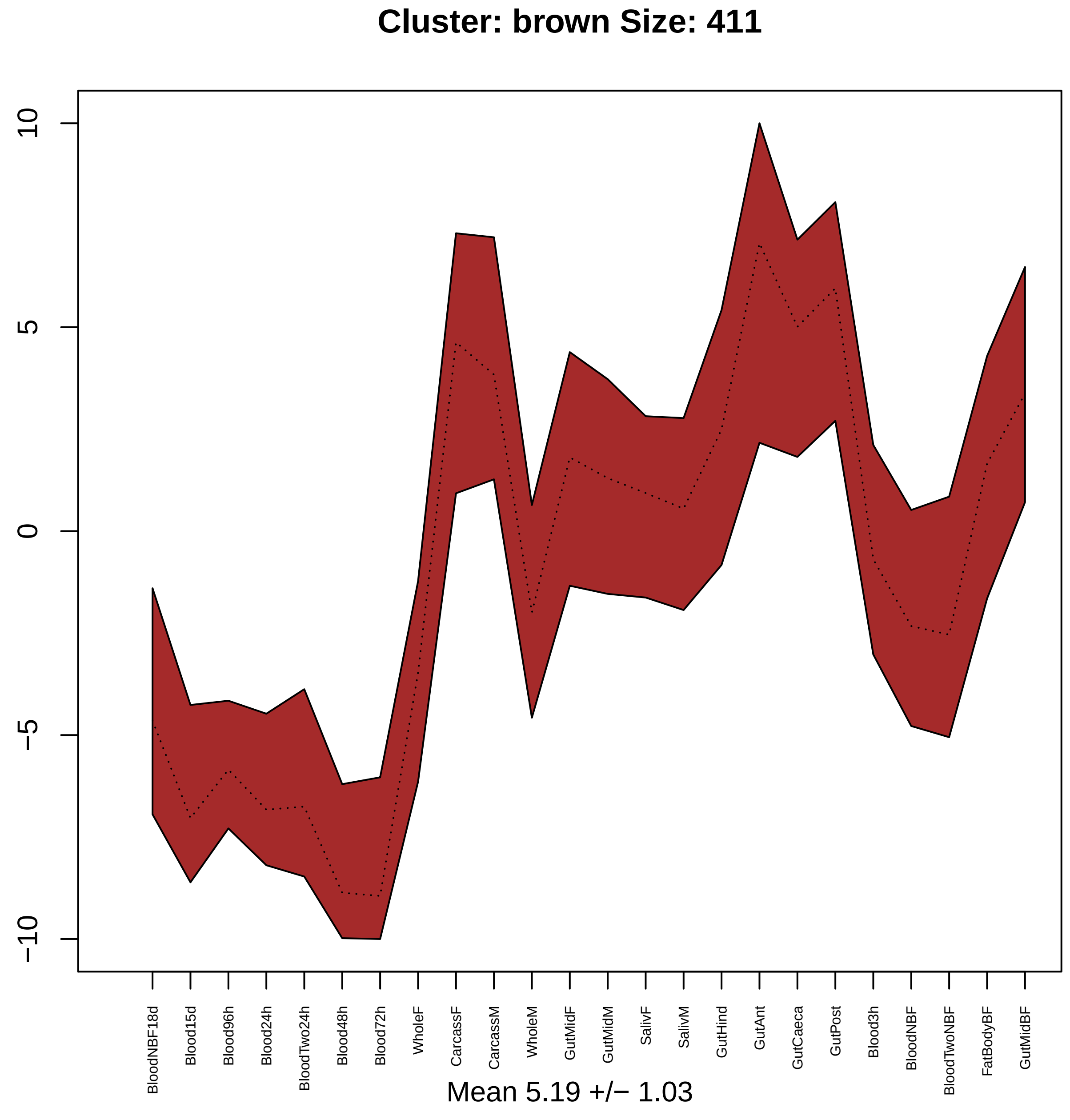

Cluster: brown Size: 411
10
5
0
−5
−10
BloodNBF18d
Blood15d
Blood96h
Blood24h
BloodTwo24h
Blood48h
Blood72h
WholeF
CarcassF
CarcassM
WholeM
GutMidF
GutMidM
SalivF
SalivM
GutHind
GutAnt
GutCaeca
GutPost
Blood3h
BloodNBF
BloodTwoNBF
FatBodyBF
GutMidBF
Mean 5.19 +/− 1.03

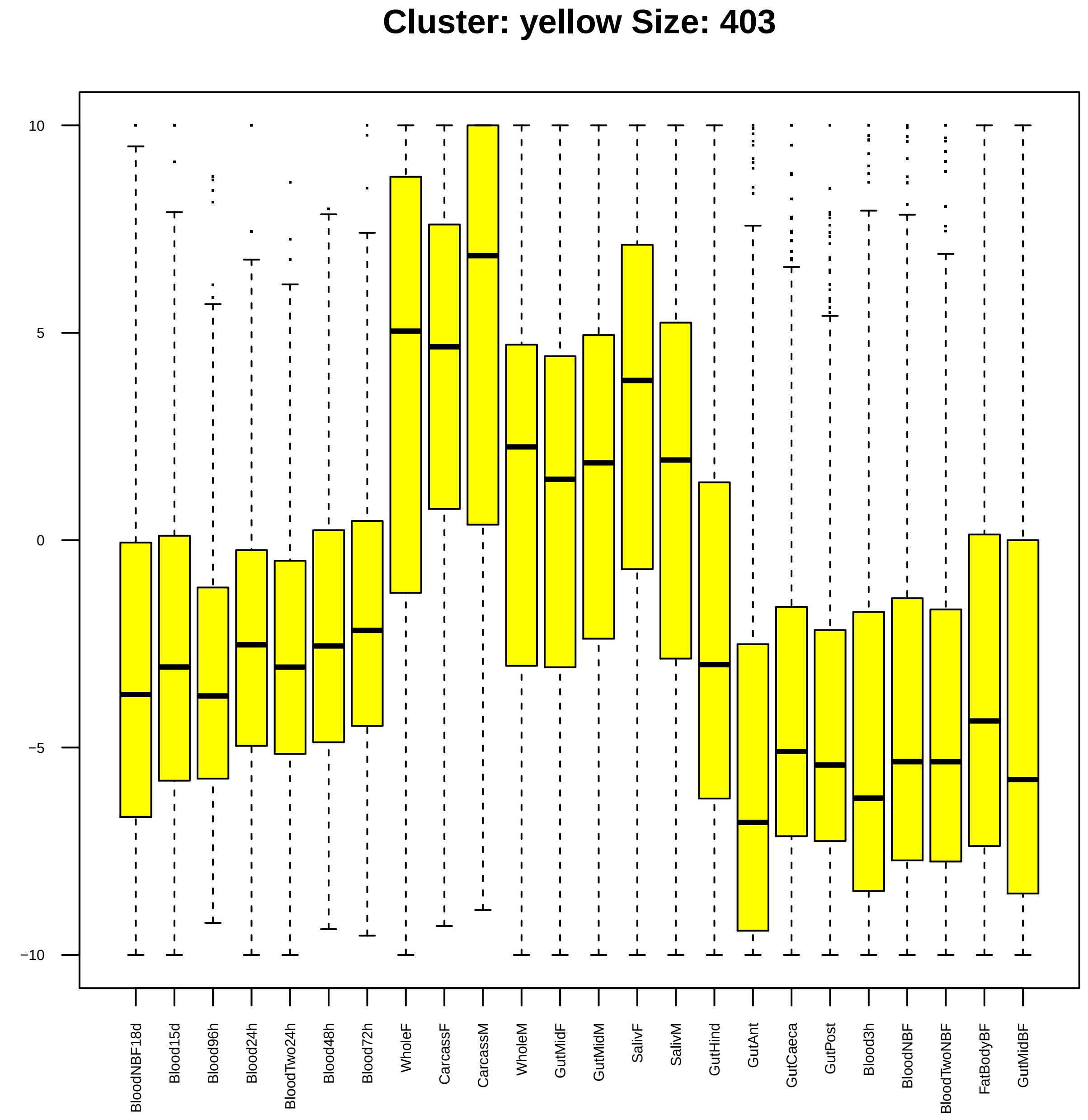

Cluster: yellow Size: 403
10
5
0
−5
−10
BloodNBF18d
Blood15d
Blood96h
Blood24h
BloodTwo24h
Blood48h
Blood72h
WholeF
CarcassF
CarcassM
WholeM
GutMidF
GutMidM
SalivF
SalivM
GutHind
GutAnt
GutCaeca
GutPost
Blood3h
BloodNBF
BloodTwoNBF
FatBodyBF
GutMidBF

| | GO.ID | BPCluster: yellow Size: 403 | Annotated | Significant | Expected | Rank in ClassicF | Weight01F | ClassicF |
|---|---|---|---|---|---|---|---|---|
| *1* | GO:0090502 | RNA phosphodiester bond hydrolysis, endo... | 14 | 5 | 0.59 | 16 | 0.00019 | 0.00019 |
| *2* | GO:0007095 | mitotic G2 DNA damage checkpoint | 43 | 8 | 1.81 | 20 | 0.00036 | 0.00036 |
| *3* | GO:0006511 | ubiquitin−dependent protein catabolic pr... | 88 | 9 | 3.71 | 80 | 0.00120 | 0.01144 |
| *4* | GO:0016236 | macroautophagy | 21 | 5 | 0.89 | 38 | 0.00150 | 0.00150 |
| *5* | GO:0016567 | protein ubiquitination | 55 | 8 | 2.32 | 46 | 0.00196 | 0.00196 |
| *6* | GO:0007033 | vacuole organization | 20 | 6 | 0.84 | 13 | 0.00346 | 0.00013 |
| *7* | GO:0006396 | RNA processing | 267 | 31 | 11.26 | 1 | 0.00697 | 2.0e−07 |
| *8* | GO:0042044 | fluid transport | 10 | 3 | 0.42 | 66 | 0.00714 | 0.00714 |
| *9* | GO:0033227 | dsRNA transport | 19 | 4 | 0.80 | 69 | 0.00727 | 0.00727 |
| *10* | GO:0048640 | negative regulation of developmental gro... | 28 | 5 | 1.18 | 59 | 0.00939 | 0.00569 |
| *11* | GO:0010883 | regulation of lipid storage | 11 | 3 | 0.46 | 77 | 0.00952 | 0.00952 |
| *15* | GO:0000398 | mRNA splicing, via spliceosome | 134 | 14 | 5.65 | 36 | 0.01398 | 0.00146 |

| | GO.ID | MFCluster: yellow Size: 403 | Annotated | Significant | Expected | Rank in ClassicF | Weight01F | ClassicF |
|---|---|---|---|---|---|---|---|---|
| *1* | GO:0016891 | endoribonuclease activity, producing 5'−... | 11 | 4 | 0.48 | 4 | 0.0009 | 0.00090 |
| *2* | GO:0005548 | phospholipid transporter activity | 10 | 3 | 0.43 | 16 | 0.0077 | 0.00775 |
| *6* | GO:0003723 | RNA binding | 307 | 26 | 13.33 | 2 | 0.0164 | 0.00078 |
| *7* | GO:0005488 | binding | 4627 | 226 | 200.89 | 5 | 0.0191 | 0.00129 |
| *10* | GO:0016874 | ligase activity | 161 | 15 | 6.99 | 11 | 0.0267 | 0.00416 |

### Cluster: yellow Size: 403

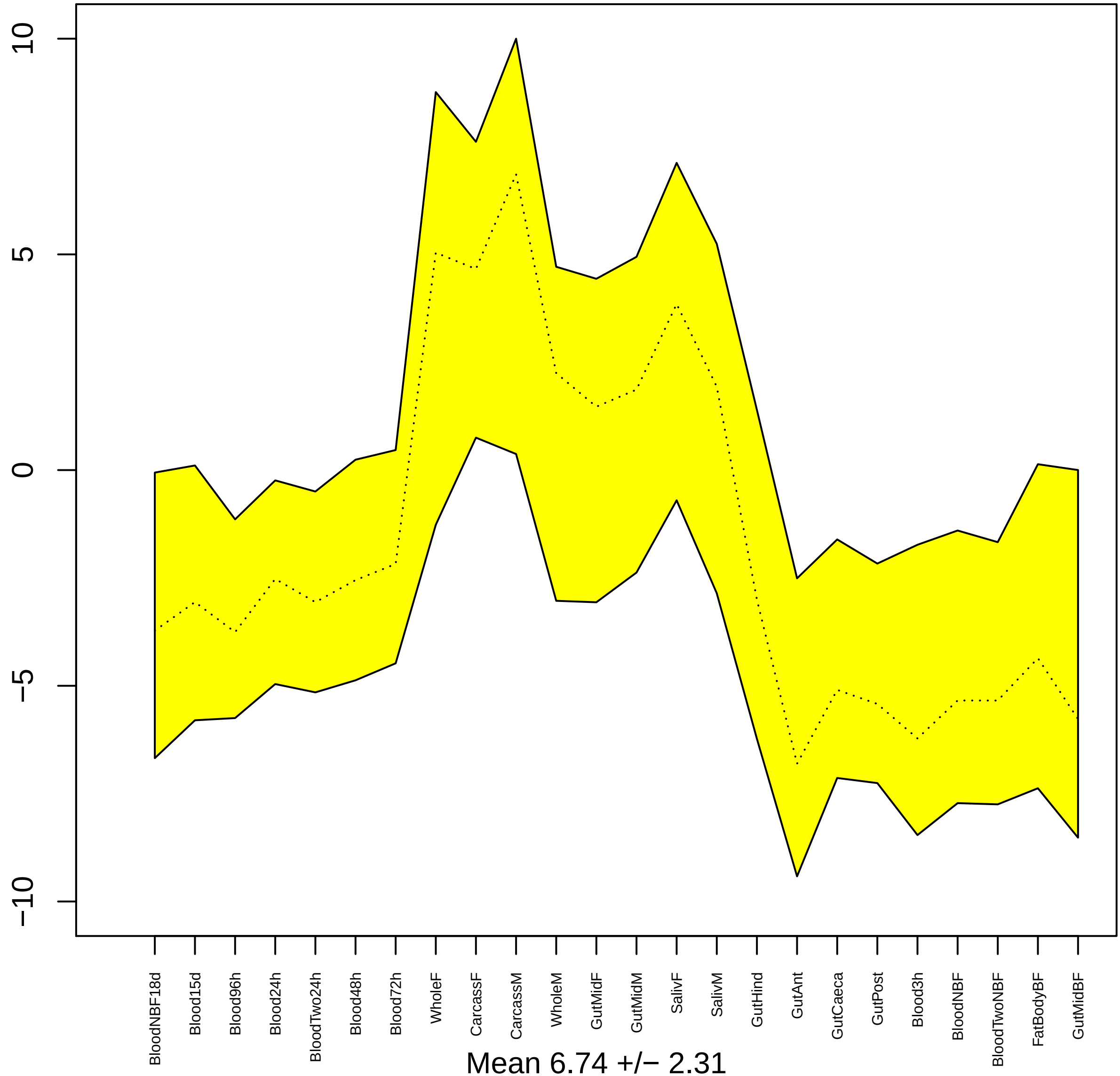


Mean 6.74 +/− 2.31

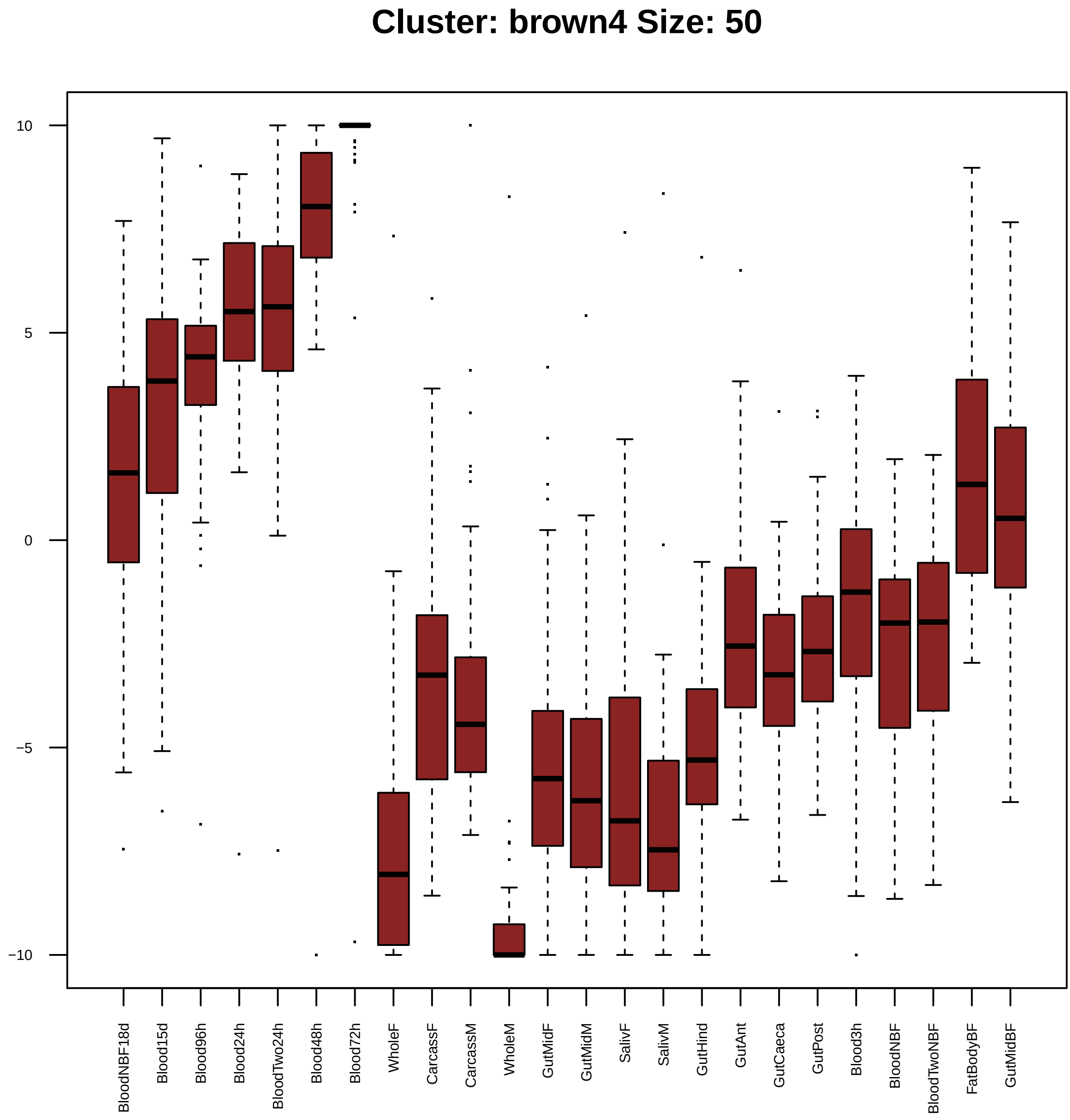

Cluster: brown4 Size: 50
10
5
0
−5
−10
BloodNBF18d
Blood15d
Blood96h
Blood24h
BloodTwo24h
Blood48h
Blood72h
WholeF
CarcassF
CarcassM
WholeM
GutMidF
GutMidM
SalivF
SalivM
GutHind
GutAnt
GutCaeca
GutPost
Blood3h
BloodNBF
BloodTwoNBF
FatBodyBF
GutMidBF

| | GO.ID | BPCluster: brown4 Size: 50 | Annotated | Significant | Expected | Rank in ClassicF | Weight01F | ClassicF |
|---|---|---|---|---|---|---|---|---|
| *1* | GO:0007062 | sister chromatid cohesion | 13 | 2 | 0.06 | 38 | 0.0013 | 0.0013 |
| *2* | GO:0006355 | regulation of transcription, DNA−templat... | 501 | 10 | 2.12 | 8 | 0.0014 | 2e−05 |
| *3* | GO:0000122 | negative regulation of transcription by ... | 57 | 3 | 0.24 | 46 | 0.0017 | 0.0017 |
| *4* | GO:0043065 | positive regulation of apoptotic process | 21 | 2 | 0.09 | 56 | 0.0035 | 0.0035 |
| *5* | GO:0045944 | positive regulation of transcription by ... | 88 | 3 | 0.37 | 69 | 0.0058 | 0.0058 |
| *7* | GO:0000082 | G1/S transition of mitotic cell cycle | 23 | 2 | 0.10 | 59 | 0.0283 | 0.0041 |

| | GO.ID | MFCluster: brown4 Size: 50 | Annotated | Significant | Expected | Rank in ClassicF | Weight01F | ClassicF |
|---|---|---|---|---|---|---|---|---|
| 2 | GO:0016896 | exoribonuclease activity, producing 5'−p... | 11 | 2 | 0.06 | 6 | 0.0015 | 0.00152 |
| 3 | GO:0001077 | proximal promoter DNA−binding transcript... | 17 | 2 | 0.09 | 9 | 0.0037 | 0.00368 |
| 4 | GO:0003676 | nucleic acid binding | 1152 | 15 | 6.23 | 4 | 0.0040 | 0.00058 |
| 5 | GO:0016817 | hydrolase activity, acting on acid anhyd... | 442 | 3 | 2.39 | 67 | 0.0055 | 0.43144 |
| 6 | GO:0003682 | chromatin binding | 69 | 3 | 0.37 | 15 | 0.0060 | 0.00600 |
| 7 | GO:0001012 | RNA polymerase II regulatory region DNA ... | 26 | 2 | 0.14 | 18 | 0.0106 | 0.00854 |
| 8 | GO:0043565 | sequence−specific DNA binding | 212 | 5 | 1.15 | 13 | 0.0107 | 0.00535 |
| 9 | GO:0003700 | DNA−binding transcription factor activit... | 282 | 6 | 1.53 | 10 | 0.0215 | 0.00373 |
| 10 | GO:0005488 | binding | 4627 | 35 | 25.03 | 3 | 0.0271 | 0.00029 |
| 21 | GO:0003677 | DNA binding | 523 | 8 | 2.83 | 16 | 0.1767 | 0.00602 |

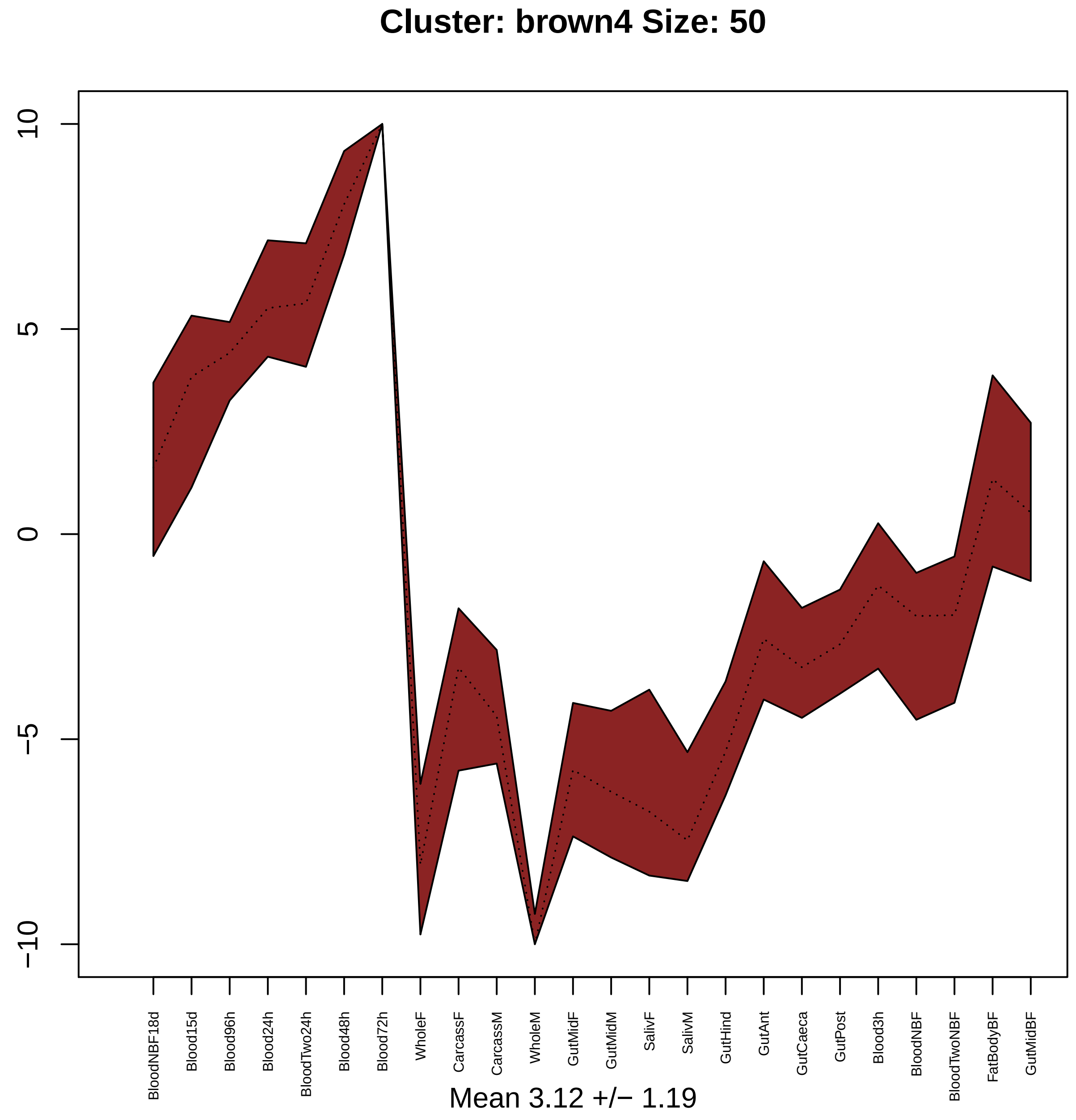

Cluster: brown4 Size: 50
10
5
0
−5
−10
BloodNBF18d
Blood15d
Blood96h
Blood24h
BloodTwo24h
Blood48h
Blood72h
WholeF
CarcassF
CarcassM
WholeM
GutMidF
GutMidM
SalivF
SalivM
GutHind
GutAnt
GutCaeca
GutPost
Blood3h
BloodNBF
BloodTwoNBF
FatBodyBF
GutMidBF
Mean 3.12 +/− 1.19

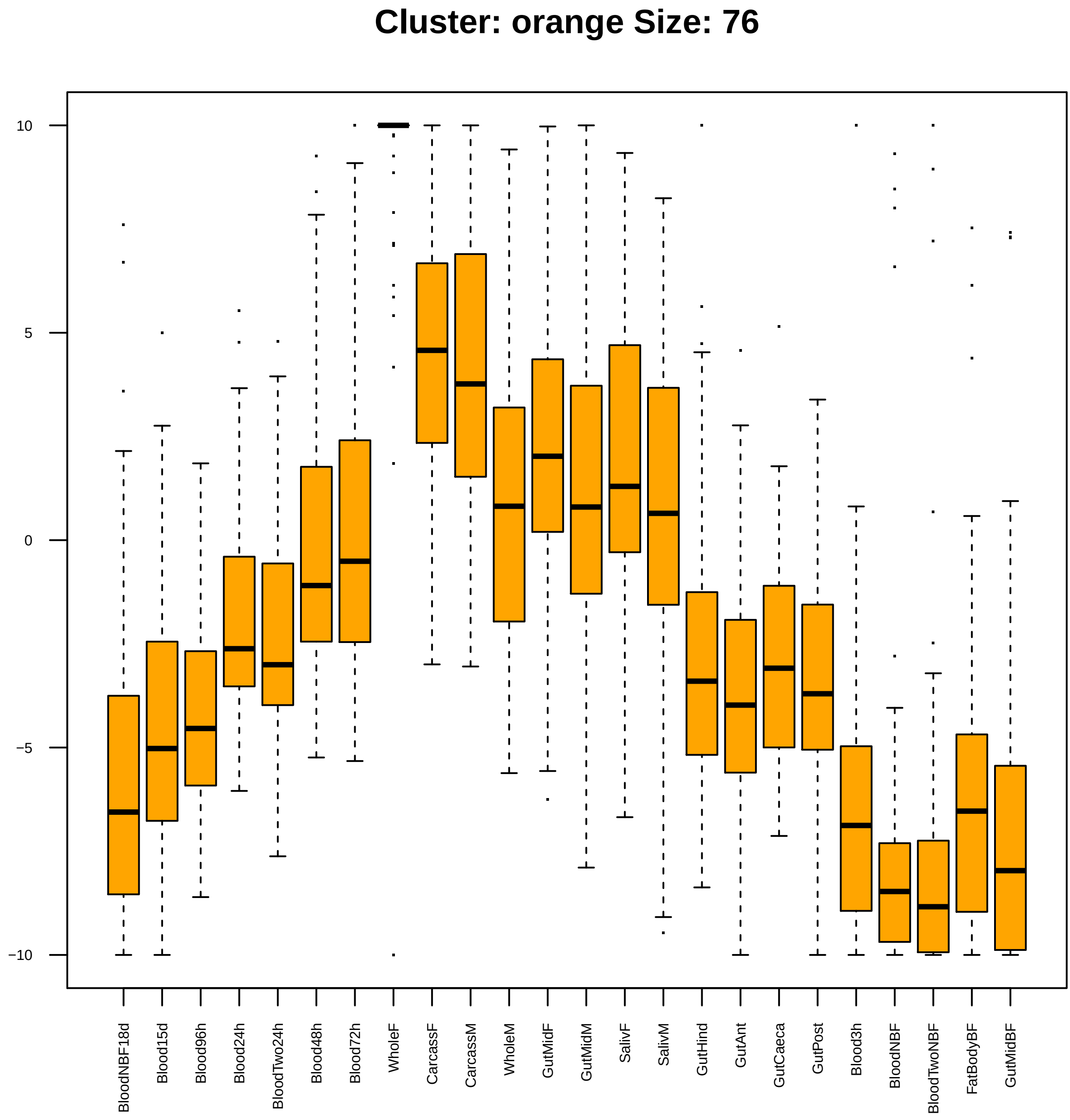

Cluster: orange Size: 76
10
5
0
−5
−10
BloodNBF18d
Blood15d
Blood96h
Blood24h
BloodTwo24h
Blood48h
Blood72h
WholeF
CarcassF
CarcassM
WholeM
GutMidF
GutMidM
SalivF
SalivM
GutHind
GutAnt
GutCaeca
GutPost
Blood3h
BloodNBF
BloodTwoNBF
FatBodyBF
GutMidBF

| | GO.ID | BPCluster: orange Size: 76 | Annotated | Significant | Expected | Rank in ClassicF | Weight01F | ClassicF |
|---|---|---|---|---|---|---|---|---|
| *1* | GO:0006406 | mRNA export from nucleus | 13 | 3 | 0.10 | 2 | 0.0001 | 0.00010 |
| *2* | GO:0065004 | protein−DNA complex assembly | 29 | 3 | 0.21 | 19 | 0.0032 | 0.00120 |
| *3* | GO:0051726 | regulation of cell cycle | 180 | 5 | 1.33 | 47 | 0.0045 | 0.00996 |
| *4* | GO:0045814 | negative regulation of gene expression, ... | 27 | 3 | 0.20 | 16 | 0.0071 | 0.00097 |
| *5* | GO:0006260 | DNA replication | 77 | 4 | 0.57 | 24 | 0.0134 | 0.00237 |
| *8* | GO:0071824 | protein−DNA complex subunit organization | 40 | 4 | 0.29 | 4 | 0.0275 | 0.00019 |
| *9* | GO:0060968 | regulation of gene silencing | 19 | 2 | 0.14 | 43 | 0.0356 | 0.00839 |
| *10* | GO:0044786 | cell cycle DNA replication | 18 | 2 | 0.13 | 41 | 0.0426 | 0.00755 |

| | GO.ID | MFCluster: orange Size: 76 | Annotated | Significant | Expected | Rank in ClassicF | Weight01F | ClassicF |
|---|---|---|---|---|---|---|---|---|
| *1* | GO:0003676 | nucleic acid binding | 1152 | 19 | 9.27 | 1 | 0.00073 | 0.0012 |
| *2* | GO:0008270 | zinc ion binding | 522 | 11 | 4.20 | 2 | 0.00259 | 0.0026 |

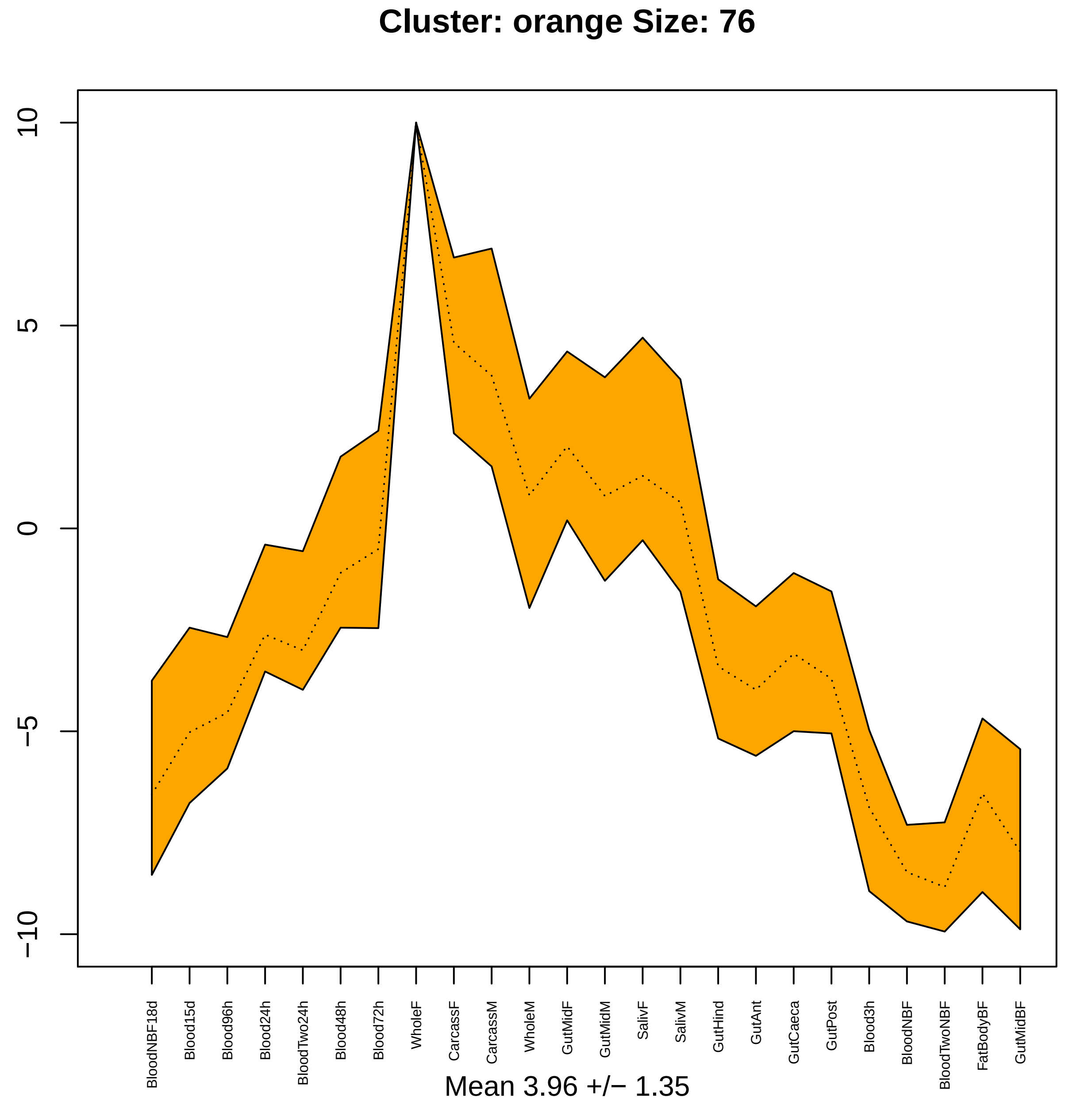
Cluster: orange Size: 76
10
5
0
−5
−10
BloodNBF18d
Blood15d
Blood96h
Blood24h
BloodTwo24h
Blood48h
Blood72h
WholeF
CarcassF
CarcassM
WholeM
GutMidF
GutMidM
SalivF
SalivM
GutHind
GutAnt
GutCaeca
GutPost
Blood3h
BloodNBF
BloodTwoNBF
FatBodyBF
GutMidBF
Mean 3.96 +/− 1.35

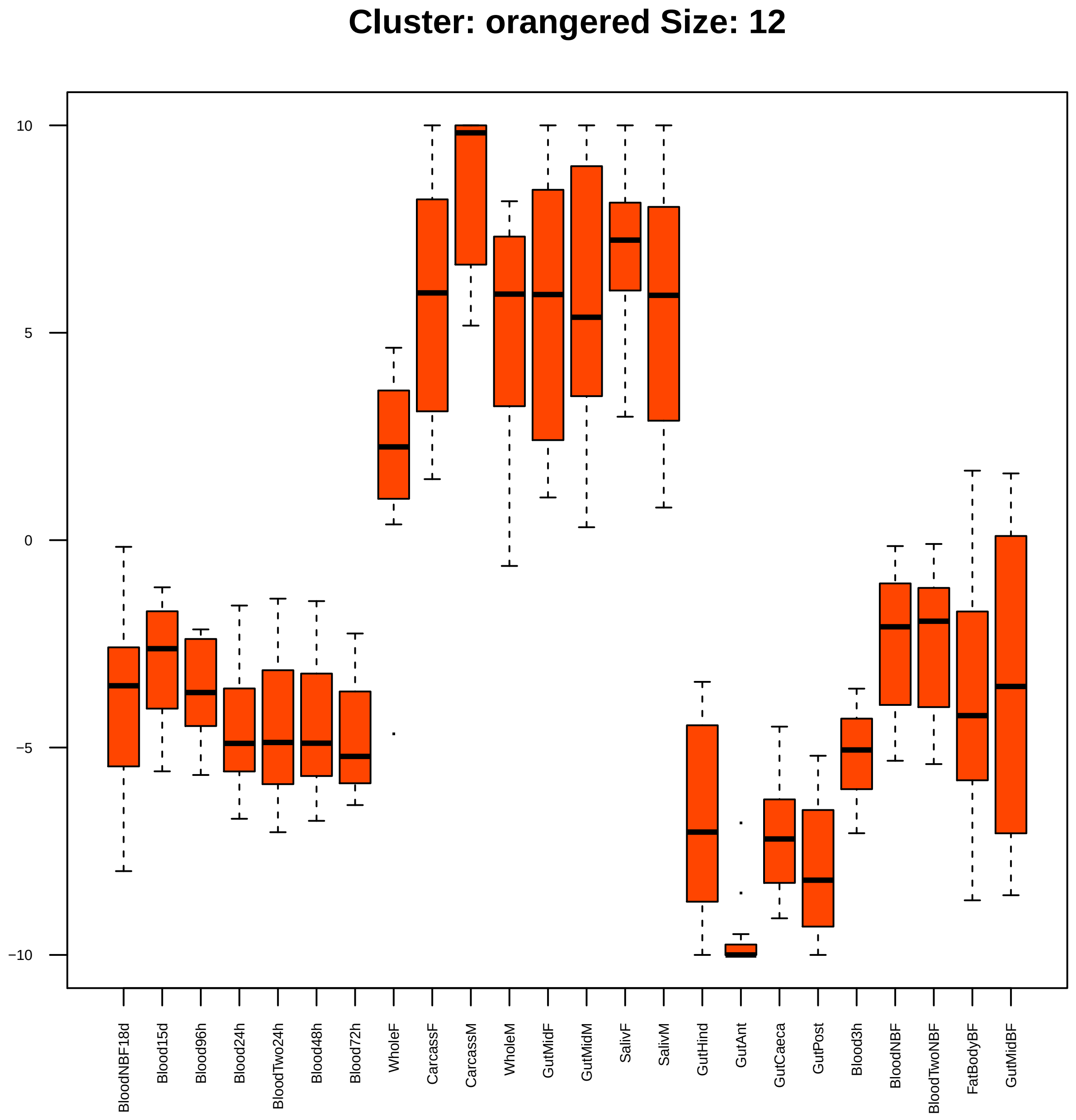
Cluster: orangered Size: 12
10
5
0
−5
−10
BloodNBF18d
Blood15d
Blood96h
Blood24h
BloodTwo24h
Blood48h
Blood72h
WholeF
CarcassF
CarcassM
WholeM
GutMidF
GutMidM
SalivF
SalivM
GutHind
GutAnt
GutCaeca
GutPost
Blood3h
BloodNBF
BloodTwoNBF
FatBodyBF
GutMidBF

| | GO.ID | BPCluster: orangered Size: 12 | Annotated | Significant | Expected | Rank in ClassicF | Weight01F | ClassicF |
|---|---|---|---|---|---|---|---|---|
| *13* | GO:0044262 | cellular carbohydrate metabolic process | 39 | 2 | 0.05 | 1 | 0.023 | 0.001 |

| | GO.ID | MFCluster: orangered Size: 12 | Annotated | Significant | Expected | Rank in ClassicF | Weight01F | ClassicF |
|---|---|---|---|---|---|---|---|---|
| *1* | GO:0019887 | protein kinase regulator activity | 29 | 2 | 0.04 | 1 | 0.00055 | 0.00055 |

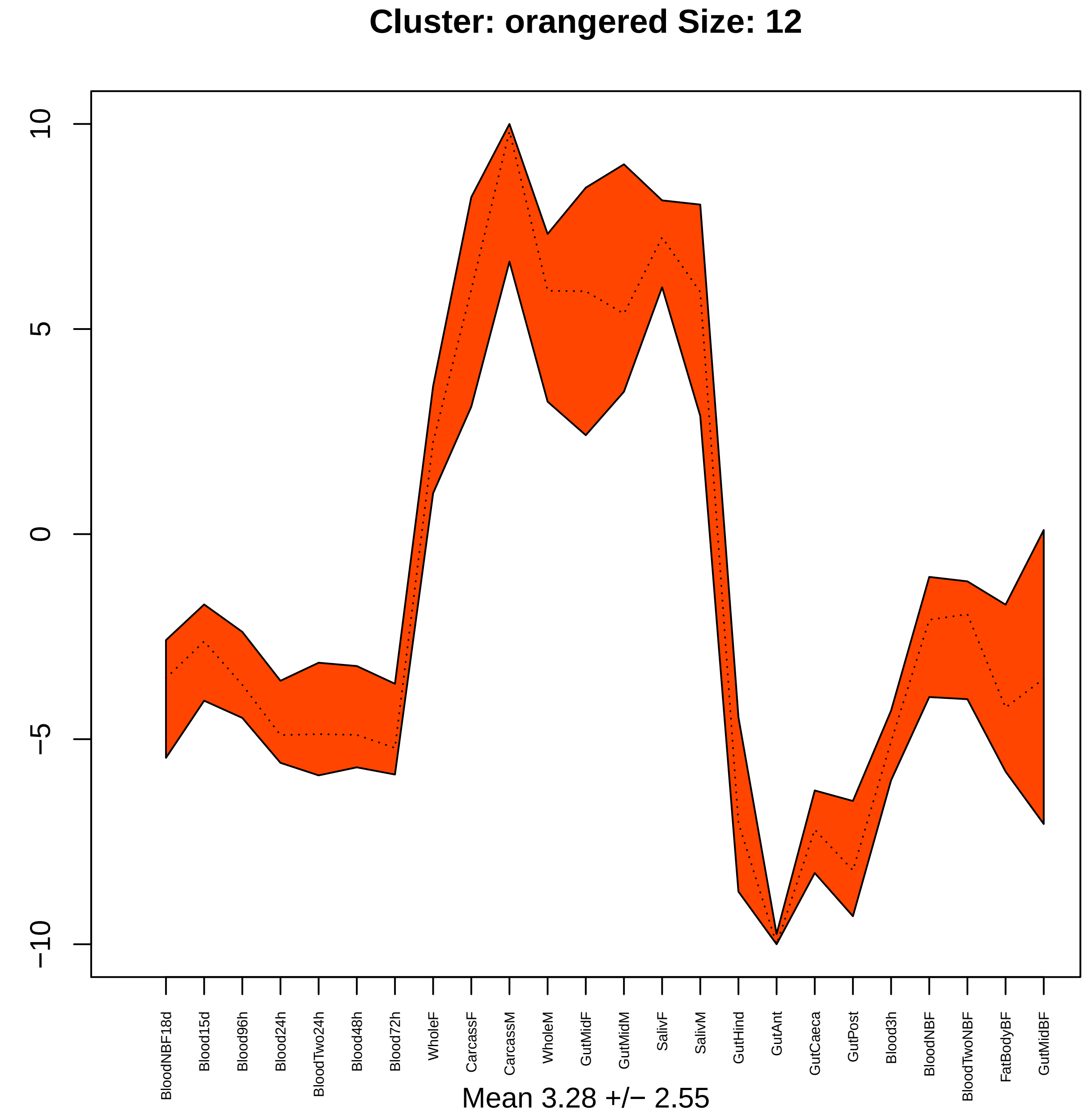

Cluster: orangered Size: 12
10
5
0
−5
−10
BloodNBF18d
Blood15d
Blood96h
Blood24h
BloodTwo24h
Blood48h
Blood72h
WholeF
CarcassF
CarcassM
WholeM
GutMidF
GutMidM
SalivF
SalivM
GutHind
GutAnt
GutCaeca
GutPost
Blood3h
BloodNBF
BloodTwoNBF
FatBodyBF
GutMidBF
Mean 3.28 +/− 2.55

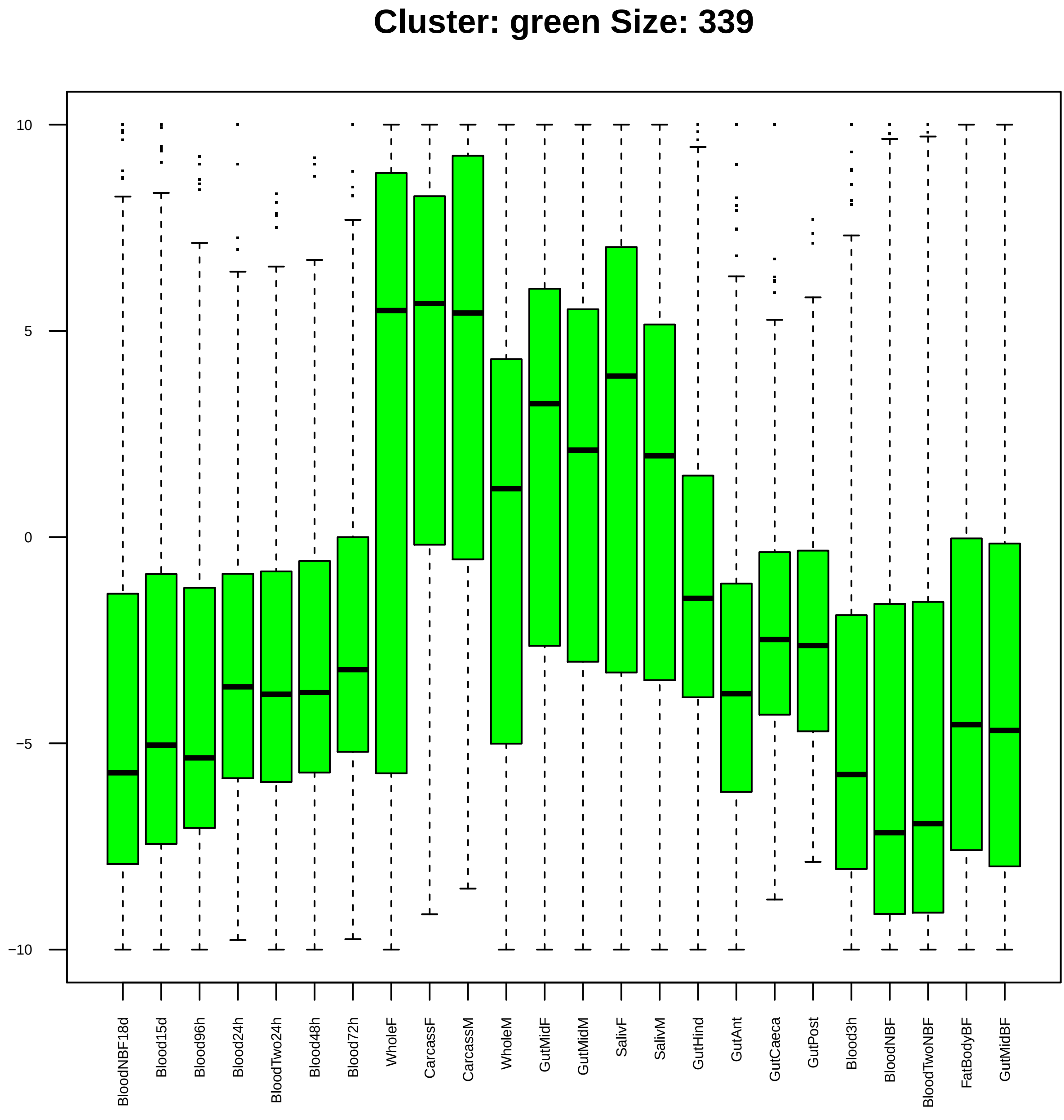

Cluster: green Size: 339
10
5
0
−5
−10
BloodNBF18d
Blood15d
Blood96h
Blood24h
BloodTwo24h
Blood48h
Blood72h
WholeF
CarcassF
CarcassM
WholeM
GutMidF
GutMidM
SalivF
SalivM
GutHind
GutAnt
GutCaeca
GutPost
Blood3h
BloodNBF
BloodTwoNBF
FatBodyBF
GutMidBF

| | GO.ID | BPCluster: green Size: 339 | Annotated | Significant | Expected | Rank in ClassicF | Weight01F | ClassicF |
|---|---|---|---|---|---|---|---|---|
| 3 | GO:0015031 | protein transport | 242 | 18 | 8.88 | 36 | 0.0022 | 0.00325 |
| 4 | GO:0001510 | RNA methylation | 17 | 4 | 0.62 | 34 | 0.0029 | 0.00288 |
| 5 | GO:0016319 | mushroom body development | 28 | 5 | 1.03 | 35 | 0.0031 | 0.00314 |
| 6 | GO:0042254 | ribosome biogenesis | 64 | 7 | 2.35 | 55 | 0.0042 | 0.00857 |
| 7 | GO:0006891 | intra−Golgi vesicle−mediated transport | 10 | 3 | 0.37 | 44 | 0.0048 | 0.00483 |
| 8 | GO:0000723 | telomere maintenance | 10 | 3 | 0.37 | 45 | 0.0048 | 0.00483 |
| 9 | GO:0000381 | regulation of alternative mRNA splicing,... | 32 | 5 | 1.17 | 50 | 0.0057 | 0.00570 |
| 10 | GO:0000398 | mRNA splicing, via spliceosome | 134 | 14 | 4.92 | 27 | 0.0058 | 0.00037 |
| 11 | GO:0006281 | DNA repair | 89 | 9 | 3.27 | 49 | 0.0098 | 0.00510 |
| 19 | GO:0043484 | regulation of RNA splicing | 38 | 6 | 1.39 | 33 | 0.0361 | 0.00236 |

| | GO.ID | MFCluster: green Size: 339 | Annotated | Significant | Expected | Rank in ClassicF | Weight01F | ClassicF |
|---|---|---|---|---|---|---|---|---|
| *3* | GO:0008173 | RNA methyltransferase activity | 19 | 5 | 0.67 | 18 | 0.0004 | 0.00040 |
| *4* | GO:0005524 | ATP binding | 592 | 35 | 20.86 | 20 | 0.0015 | 0.00153 |
| *5* | GO:0003676 | nucleic acid binding | 1152 | 62 | 40.59 | 17 | 0.0018 | 0.00027 |
| *6* | GO:0008408 | 3'−5' exonuclease activity | 18 | 4 | 0.63 | 26 | 0.0031 | 0.00312 |
| *7* | GO:0015297 | antiporter activity | 10 | 3 | 0.35 | 29 | 0.0043 | 0.00432 |
| *8* | GO:0000049 | tRNA binding | 10 | 3 | 0.35 | 30 | 0.0043 | 0.00432 |
| *9* | GO:0016706 | oxidoreductase activity, acting on paire... | 12 | 3 | 0.42 | 32 | 0.0075 | 0.00751 |
| *13* | GO:0004518 | nuclease activity | 66 | 8 | 2.33 | 23 | 0.0224 | 0.00208 |
| *14* | GO:0016796 | exonuclease activity, active with either... | 18 | 4 | 0.63 | 27 | 0.0228 | 0.00312 |
| *18* | GO:0019001 | guanyl nucleotide binding | 137 | 11 | 4.83 | 33 | 0.0345 | 0.00880 |

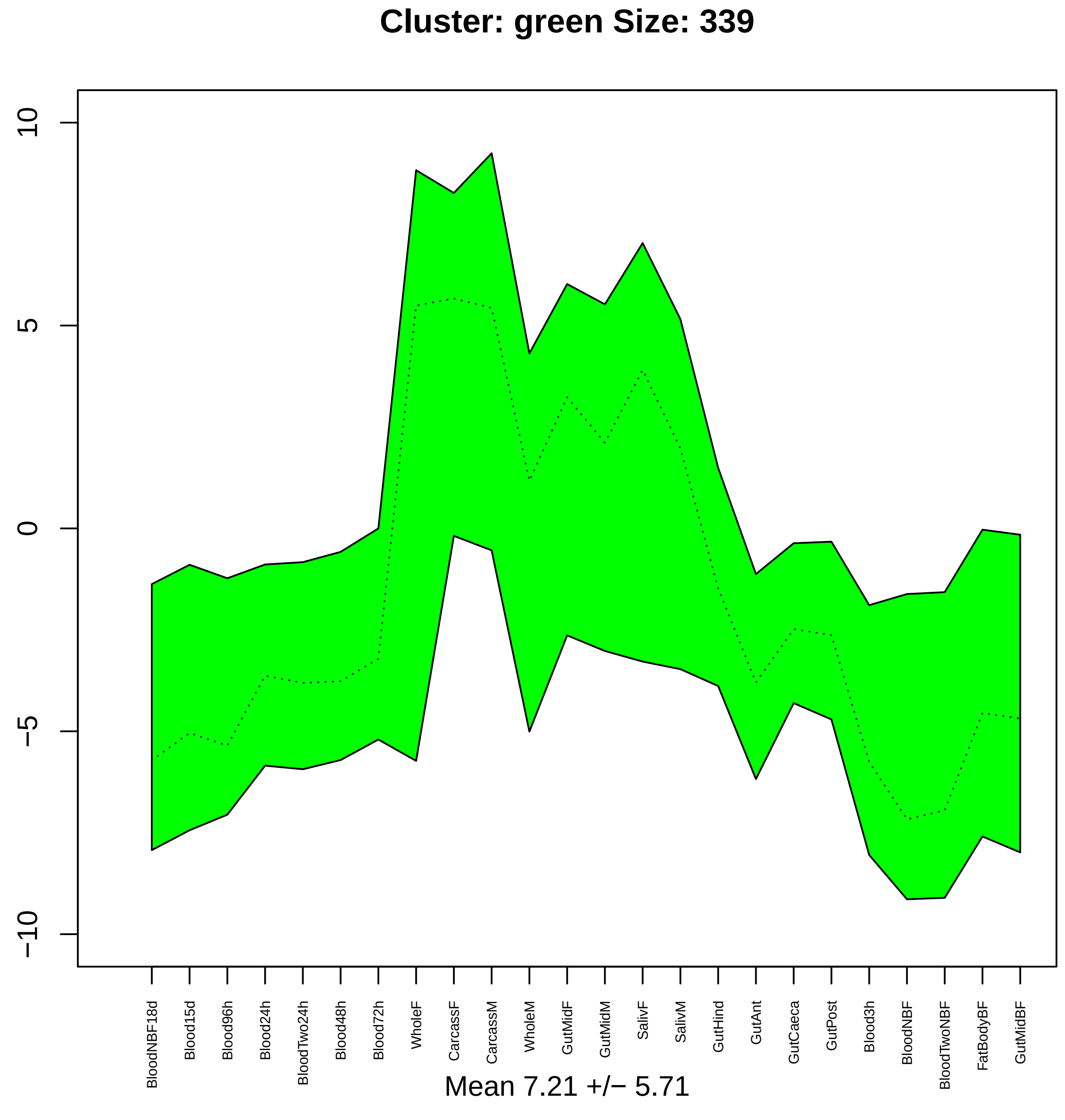
Cluster: green Size: 339
10
5
0
−5
−10
BloodNBF18d
Blood15d
Blood96h
Blood24h
BloodTwo24h
Blood48h
Blood72h
WholeF
CarcassF
CarcassM
WholeM
GutMidF
GutMidM
SalivF
SalivM
GutHind
GutAnt
GutCaeca
GutPost
Blood3h
BloodNBF
BloodTwoNBF
FatBodyBF
GutMidBF
Mean 7.21 +/− 5.71

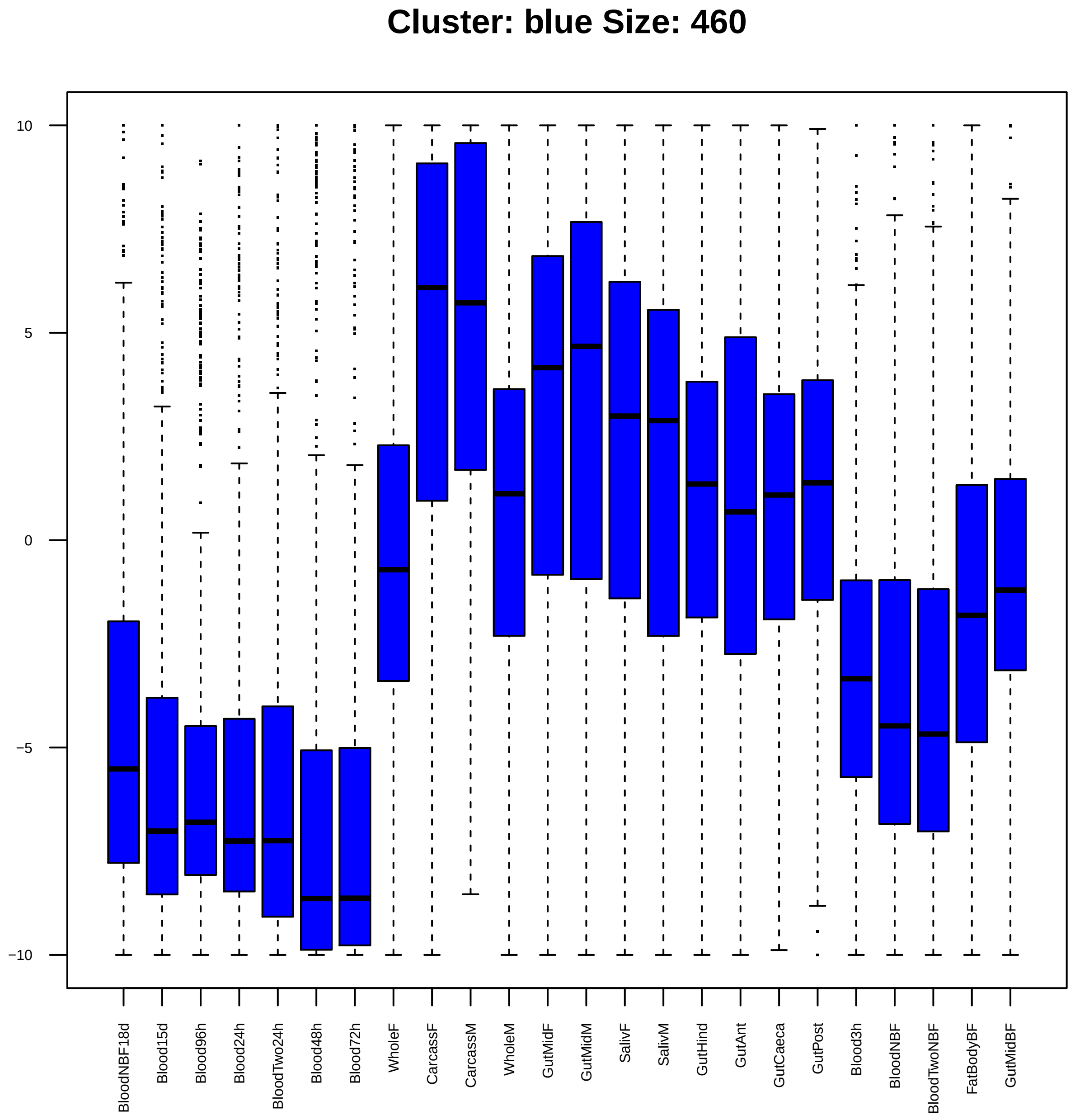

Cluster: blue Size: 460
10
5
0
−5
−10
BloodNBF18d
Blood15d
Blood96h
Blood24h
BloodTwo24h
Blood48h
Blood72h
WholeF
CarcassF
CarcassM
WholeM
GutMidF
GutMidM
SalivF
SalivM
GutHind
GutAnt
GutCaeca
GutPost
Blood3h
BloodNBF
BloodTwoNBF
FatBodyBF
GutMidBF

| | GO.ID | BPCluster: blue Size: 460 | Annotated | Significant | Expected | Rank in ClassicF | Weight01F | ClassicF |
|---|---|---|---|---|---|---|---|---|
| *1* | GO:0008045 | motor neuron axon guidance | 27 | 6 | 1.16 | 1 | 0.00082 | 0.00082 |
| *2* | GO:1901565 | organonitrogen compound catabolic proces... | 194 | 12 | 8.34 | 146 | 0.00205 | 0.12949 |
| *3* | GO:0060070 | canonical Wnt signaling pathway | 18 | 4 | 0.77 | 5 | 0.00633 | 0.00633 |
| *4* | GO:0042749 | regulation of circadian sleep/wake cycle | 10 | 3 | 0.43 | 6 | 0.00752 | 0.00752 |
| *29* | GO:0000902 | cell morphogenesis | 335 | 24 | 14.39 | 9 | 0.06867 | 0.00906 |

| | GO.ID | MFCluster: blue Size: 460 | Annotated | Significant | Expected | Rank in ClassicF | Weight01F | ClassicF |
|---|---|---|---|---|---|---|---|---|
| *1* | GO:0016746 | transferase activity, transferring acyl ... | 141 | 10 | 6.02 | 24 | 0.0012 | 0.0790 |
| *2* | GO:0001664 | G protein−coupled receptor binding | 16 | 4 | 0.68 | 1 | 0.0040 | 0.0040 |
| *3* | GO:0005506 | iron ion binding | 137 | 13 | 5.85 | 2 | 0.0056 | 0.0056 |
| *4* | GO:0004497 | monooxygenase activity | 110 | 11 | 4.70 | 3 | 0.0071 | 0.0071 |
| *5* | GO:0046983 | protein dimerization activity | 136 | 9 | 5.81 | 37 | 0.0075 | 0.1271 |

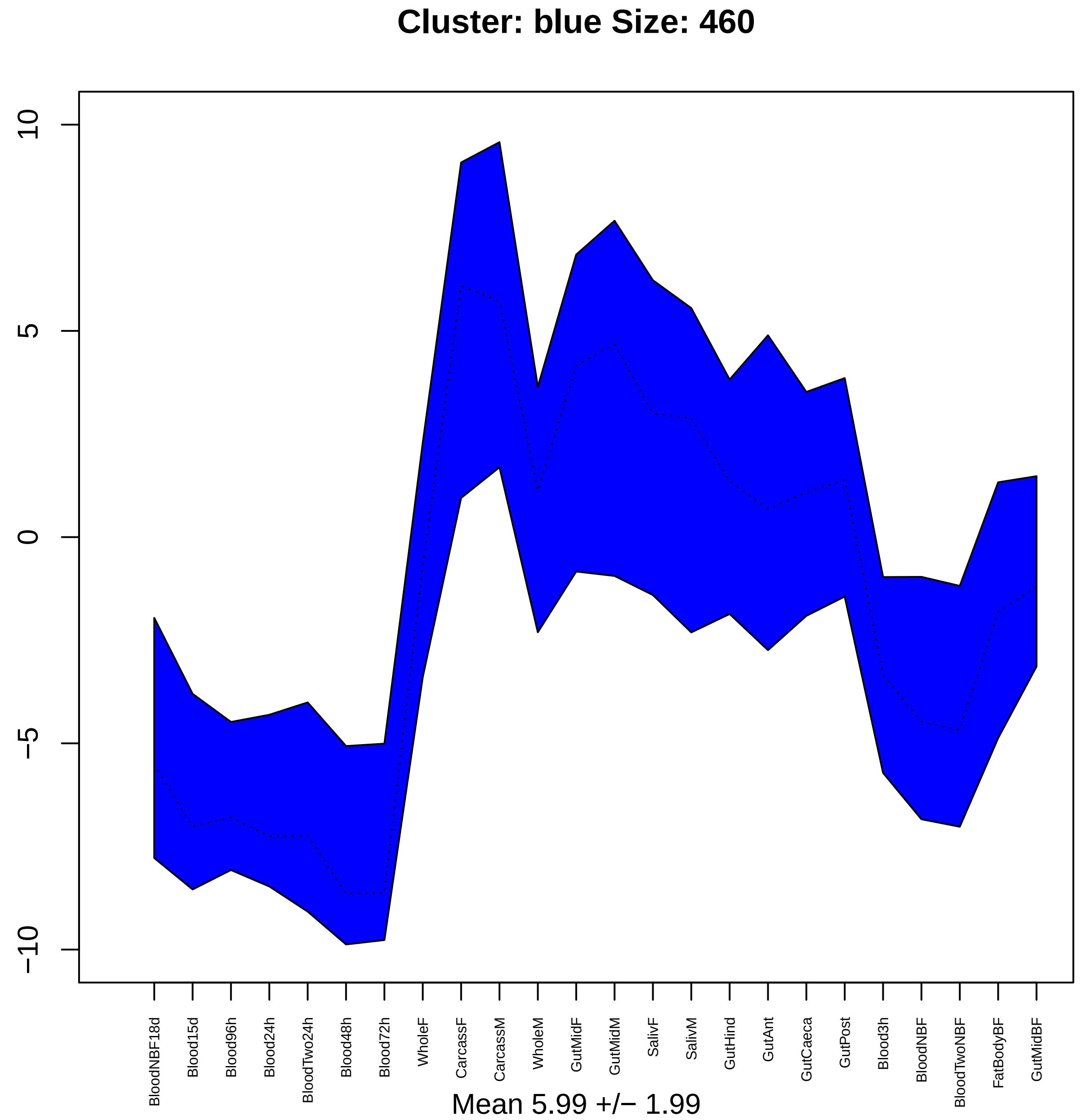

Cluster: blue Size: 460
10
5
0
−5
−10
BloodNBF18d
Blood15d
Blood96h
Blood24h
BloodTwo24h
Blood48h
Blood72h
WholeF
CarcassF
CarcassM
WholeM
GutMidF
GutMidM
SalivF
SalivM
GutHind
GutAnt
GutCaeca
GutPost
Blood3h
BloodNBF
BloodTwoNBF
FatBodyBF
GutMidBF
Mean 5.99 +/− 1.99

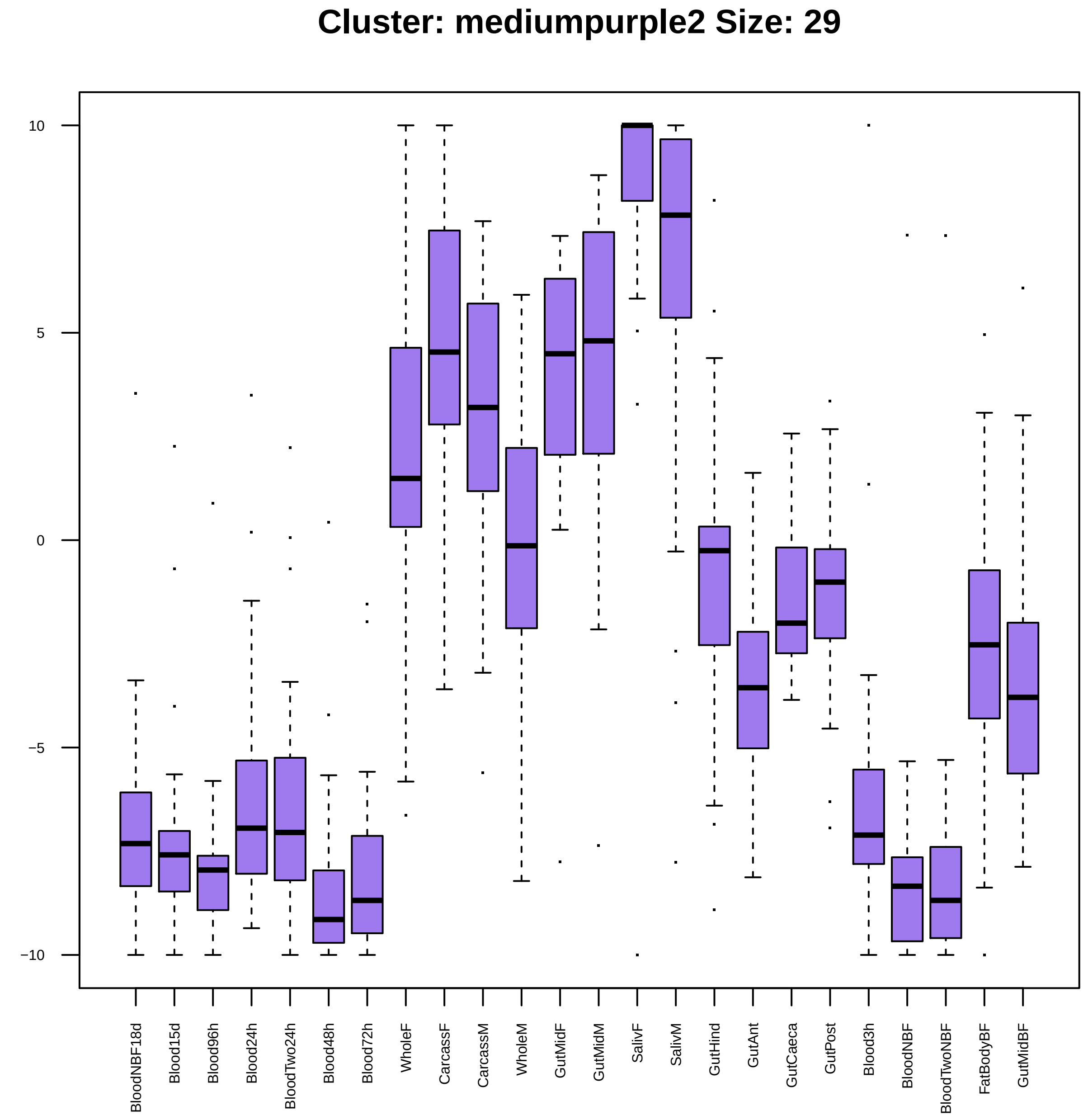

Cluster: mediumpurple2 Size: 29
10
5
0
−5
−10
BloodNBF18d
Blood15d
Blood96h
Blood24h
BloodTwo24h
Blood48h
Blood72h
WholeF
CarcassF
CarcassM
WholeM
GutMidF
GutMidM
SalivF
SalivM
GutHind
GutAnt
GutCaeca
GutPost
Blood3h
BloodNBF
BloodTwoNBF
FatBodyBF
GutMidBF

| | GO.ID | BPCluster: mediumpurple2 Size: 29 | Annotated | Significant | Expected | Rank in ClassicF | Weight01F | ClassicF |
|---|---|---|---|---|---|---|---|---|
| 3 | GO:0000022 | mitotic spindle elongation | 46 | 3 | 0.17 | 9 | 0.00056 | 0.00056 |
| 4 | GO:0006412 | translation | 289 | 7 | 1.04 | 4 | 0.00071 | 4.8e−05 |
| 5 | GO:0007030 | Golgi organization | 37 | 2 | 0.13 | 29 | 0.00768 | 0.00768 |
| 20 | GO:0010243 | response to organonitrogen compound | 37 | 2 | 0.13 | 30 | 0.05730 | 0.00768 |

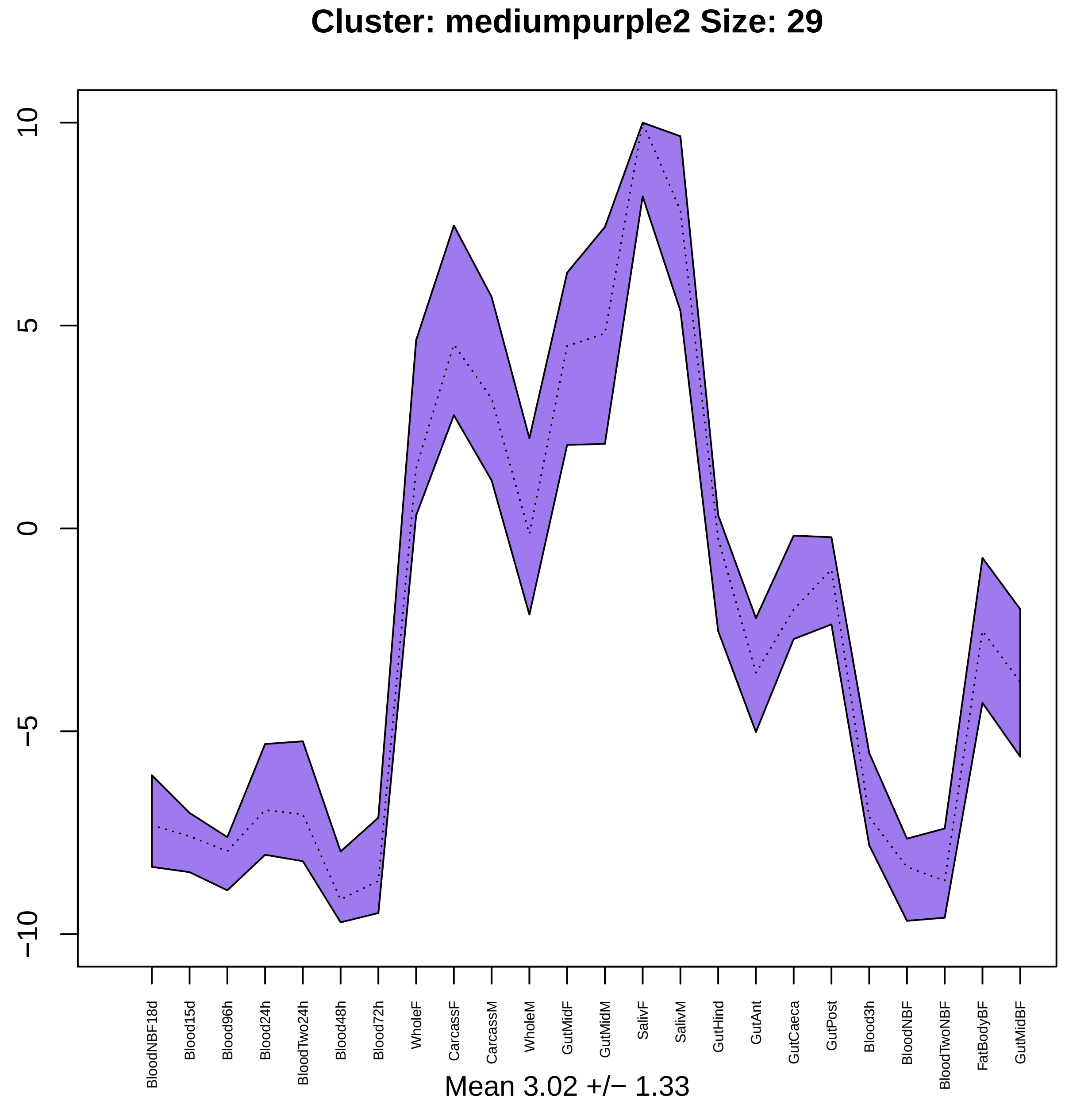
Cluster: mediumpurple2 Size: 29
10
5
0
−5
−10
BloodNBF18d
Blood15d
Blood96h
Blood24h
BloodTwo24h
Blood48h
Blood72h
WholeF
CarcassF
CarcassM
WholeM
GutMidF
GutMidM
SalivF
SalivM
GutHind
GutAnt
GutCaeca
GutPost
Blood3h
BloodNBF
BloodTwoNBF
FatBodyBF
GutMidBF
Mean 3.02 +/− 1.33

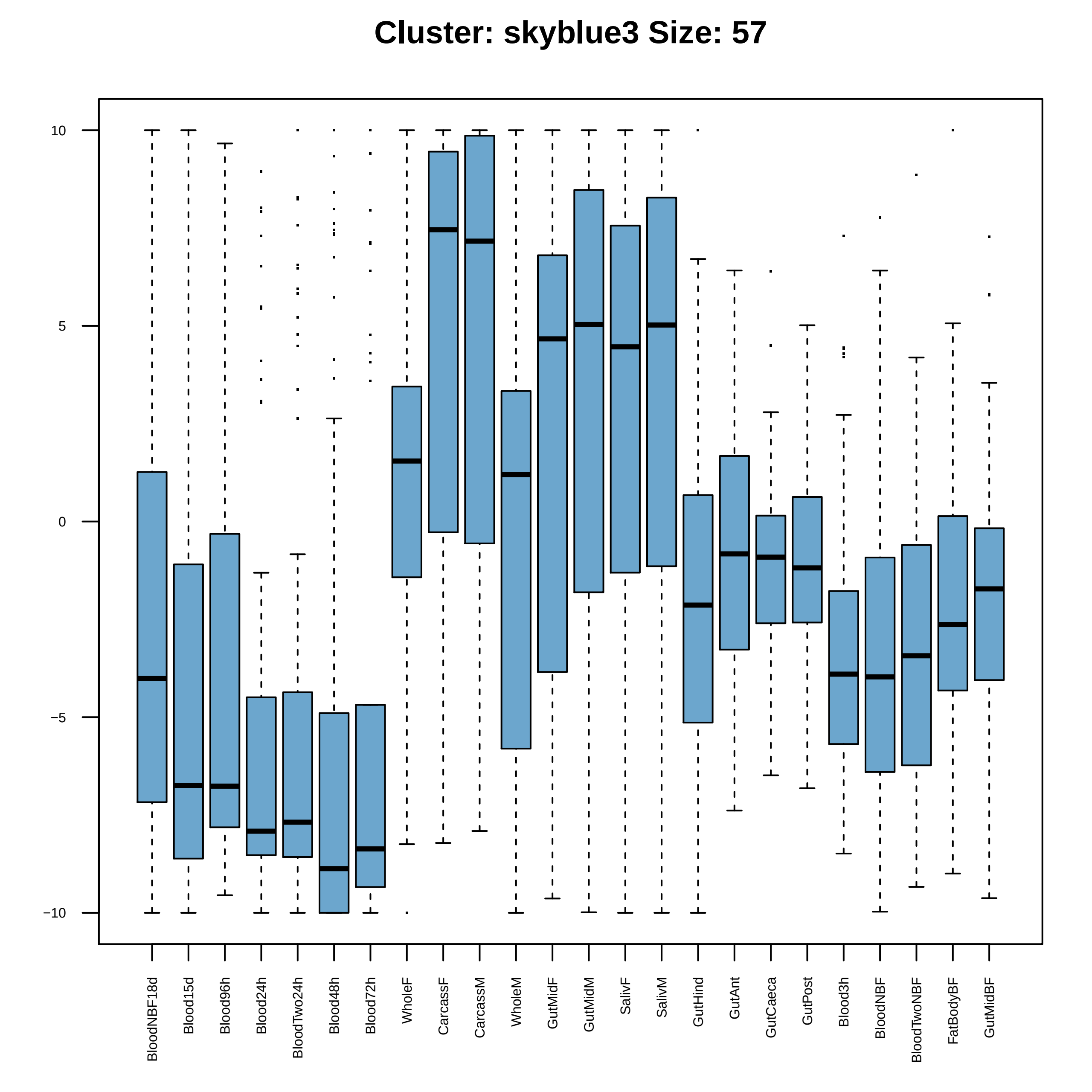

Cluster: skyblue3 Size: 57
10
5
0
−5
−10
BloodNBF18d
Blood15d
Blood96h
Blood24h
BloodTwo24h
Blood48h
Blood72h
WholeF
CarcassF
CarcassM
WholeM
GutMidF
GutMidM
SalivF
SalivM
GutHind
GutAnt
GutCaeca
GutPost
Blood3h
BloodNBF
BloodTwoNBF
FatBodyBF
GutMidBF

| | GO.ID | BPCluster: skyblue3 Size: 57 | Annotated | Significant | Expected | Rank in ClassicF | Weight01F | ClassicF |
|---|---|---|---|---|---|---|---|---|
| *1* | GO:0055088 | lipid homeostasis | 12 | 2 | 0.08 | 2 | 0.0024 | 0.0024 |

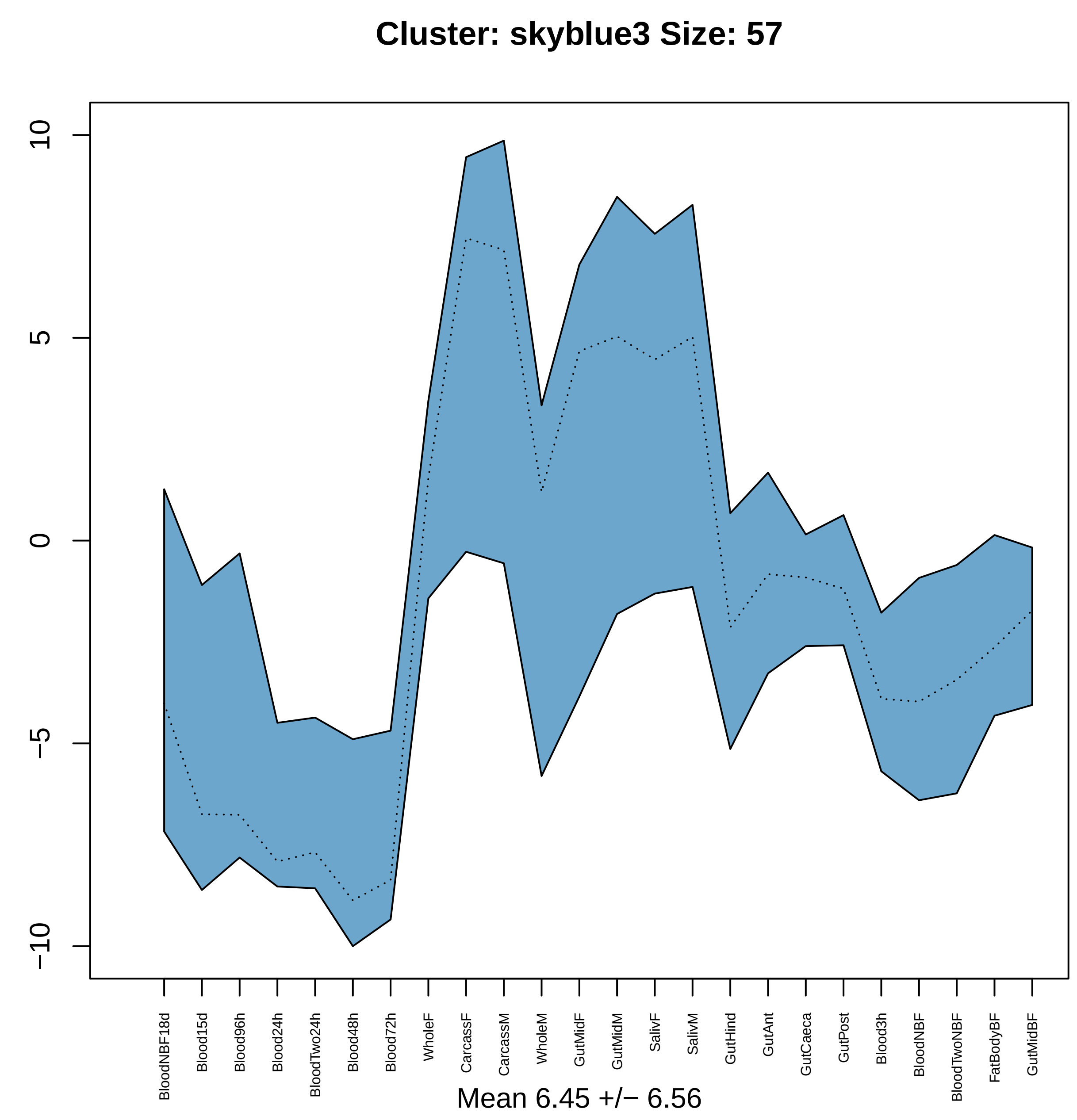
Cluster: skyblue3 Size: 57
10
5
0
−5
−10
BloodNBF18d
Blood15d
Blood96h
Blood24h
BloodTwo24h
Blood48h
Blood72h
WholeF
CarcassF
CarcassM
WholeM
GutMidF
GutMidM
SalivF
SalivM
GutHind
GutAnt
GutCaeca
GutPost
Blood3h
BloodNBF
BloodTwoNBF
FatBodyBF
GutMidBF
Mean 6.45 +/− 6.56

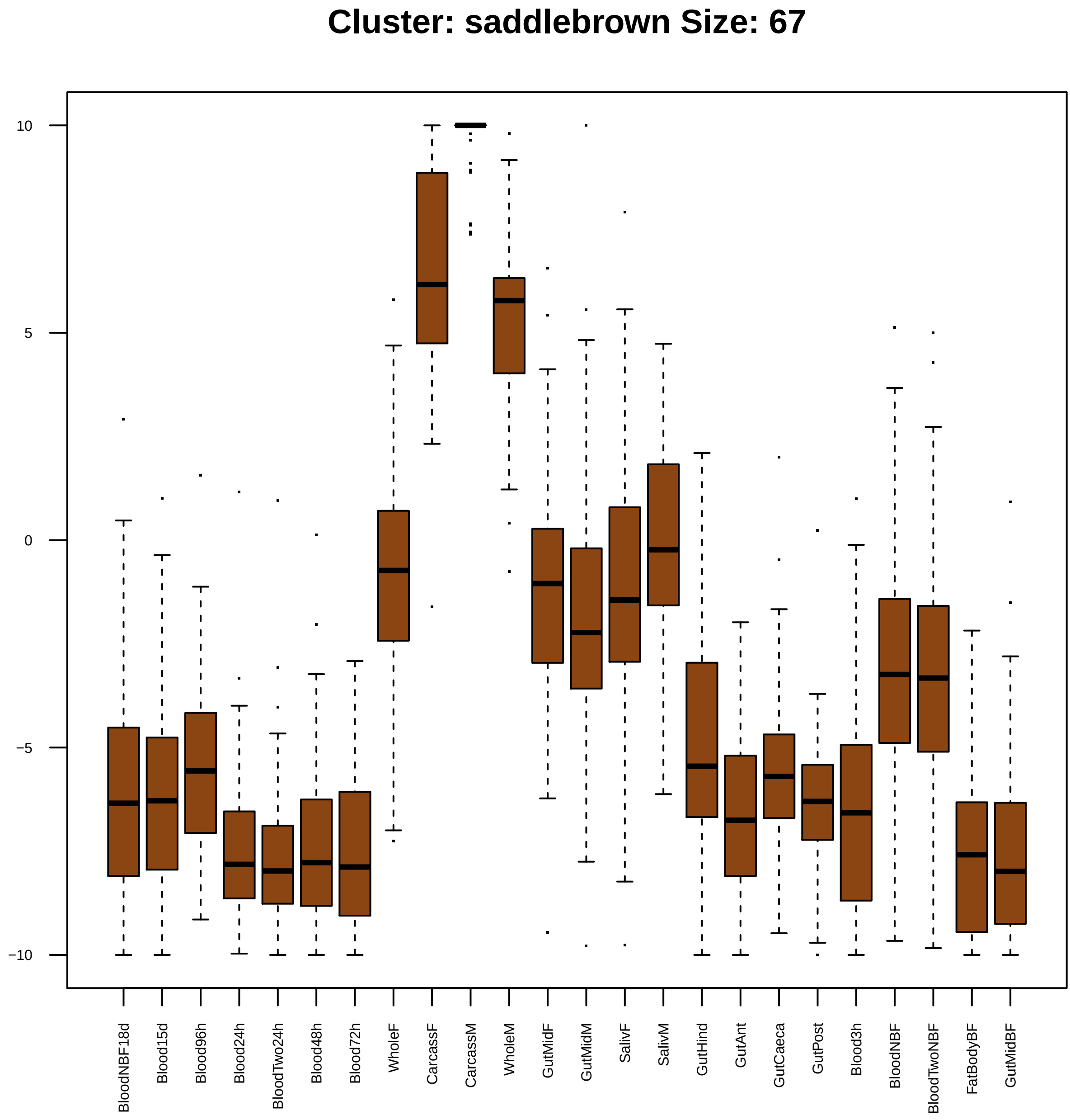
Cluster: saddlebrown Size: 67
10
5
0
−5
−10
BloodNBF18d
Blood15d
Blood96h
Blood24h
BloodTwo24h
Blood48h
Blood72h
WholeF
CarcassF
CarcassM
WholeM
GutMidF
GutMidM
SalivF
SalivM
GutHind
GutAnt
GutCaeca
GutPost
Blood3h
BloodNBF
BloodTwoNBF
FatBodyBF
GutMidBF

| | GO.ID | BPCluster: saddlebrown Size: 67 | Annotated | Significant | Expected | Rank in ClassicF | Weight01F | ClassicF |
|---|---|---|---|---|---|---|---|---|
| 2 | GO:0055114 | oxidation−reduction process | 545 | 16 | 3.67 | 2 | 0.0010 | 2.0e−07 |
| 3 | GO:0006544 | glycine metabolic process | 10 | 2 | 0.07 | 19 | 0.0019 | 0.00193 |
| 4 | GO:0016226 | iron−sulfur cluster assembly | 11 | 2 | 0.07 | 20 | 0.0024 | 0.00235 |
| 5 | GO:0005977 | glycogen metabolic process | 11 | 2 | 0.07 | 21 | 0.0024 | 0.00235 |
| 6 | GO:0042135 | neurotransmitter catabolic process | 12 | 2 | 0.08 | 29 | 0.0028 | 0.00281 |
| 7 | GO:0006096 | glycolytic process | 12 | 2 | 0.08 | 30 | 0.0028 | 0.00281 |
| 11 | GO:0009117 | nucleotide metabolic process | 165 | 5 | 1.11 | 39 | 0.0310 | 0.00475 |
| 12 | GO:0016052 | carbohydrate catabolic process | 17 | 3 | 0.11 | 14 | 0.0318 | 0.00018 |
| 15 | GO:0006090 | pyruvate metabolic process | 20 | 3 | 0.13 | 16 | 0.0504 | 0.00030 |

| | GO.ID | MFCluster: saddlebrown Size: 67 | Annotated | Significant | Expected | Rank in ClassicF | Weight01F | ClassicF |
|---|---|---|---|---|---|---|---|---|
| *1* | GO:0016616 | oxidoreductase activity, acting on the C... | 41 | 4 | 0.28 | 2 | 0.00017 | 0.00017 |
| *2* | GO:0051287 | NAD binding | 26 | 3 | 0.18 | 4 | 0.00073 | 0.00073 |
| *3* | GO:0000287 | magnesium ion binding | 34 | 3 | 0.24 | 7 | 0.00161 | 0.00161 |
| *5* | GO:0016903 | oxidoreductase activity, acting on the a... | 33 | 3 | 0.23 | 5 | 0.03945 | 0.00148 |
| *17* | GO:0043169 | cation binding | 1275 | 17 | 8.84 | 9 | 0.10838 | 0.00403 |

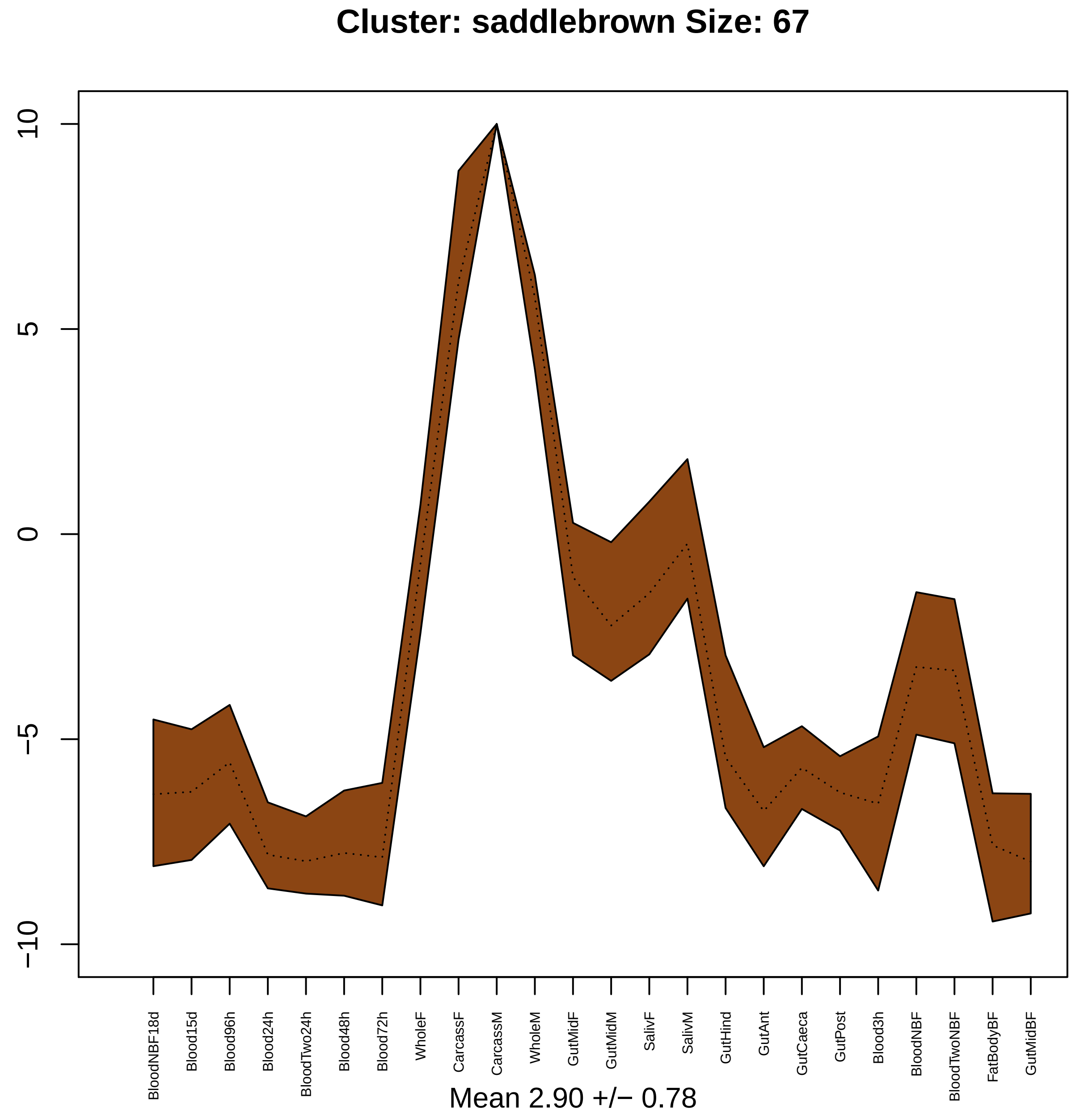

Cluster: saddlebrown Size: 67
10
5
0
−5
−10
BloodNBF18d
Blood15d
Blood96h
Blood24h
BloodTwo24h
Blood48h
Blood72h
WholeF
CarcassF
CarcassM
WholeM
GutMidF
GutMidM
SalivF
SalivM
GutHind
GutAnt
GutCaeca
GutPost
Blood3h
BloodNBF
BloodTwoNBF
FatBodyBF
GutMidBF
Mean 2.90 +/− 0.78

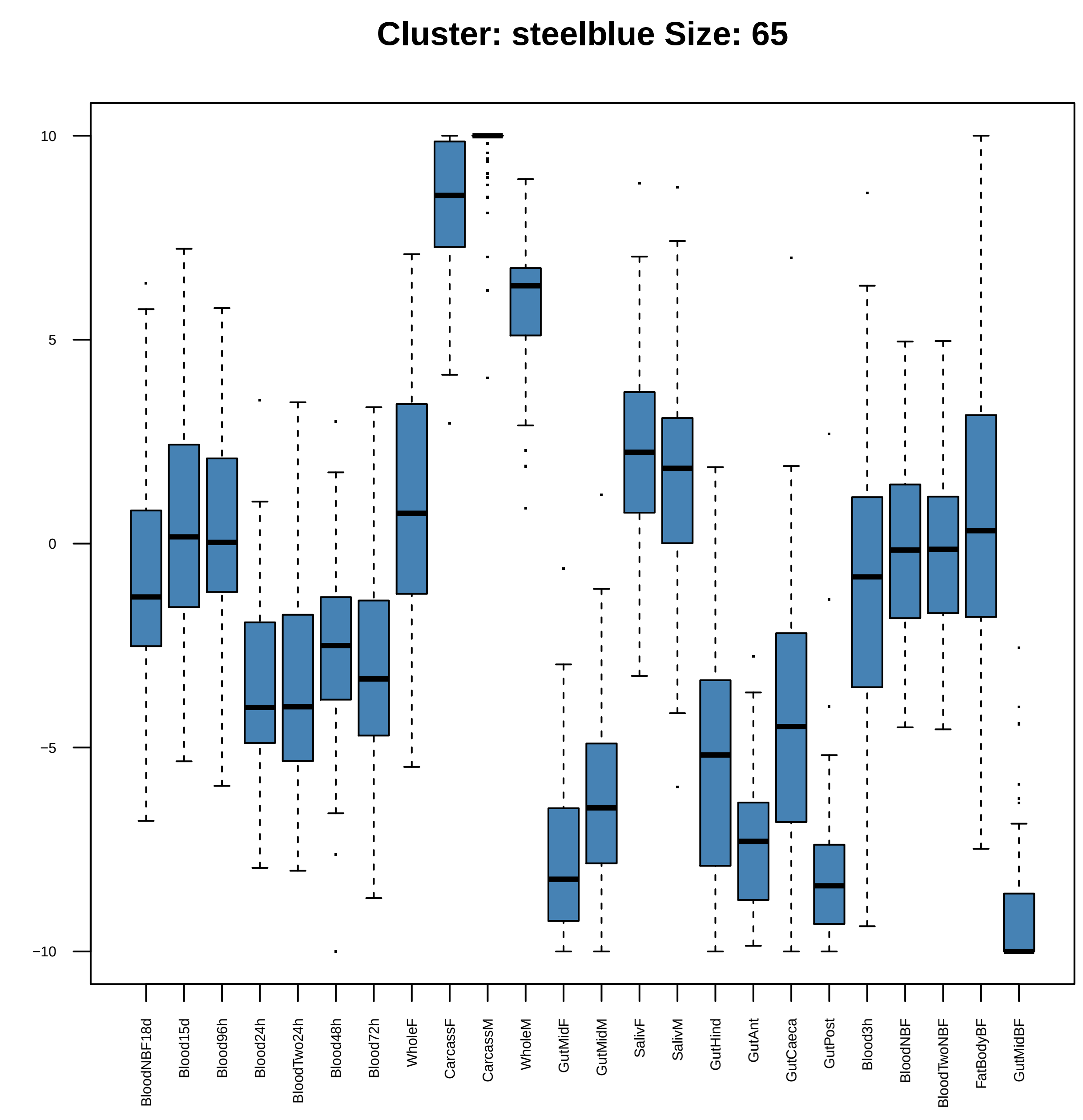

Cluster: steelblue Size: 65
10
5
0
−5
−10
BloodNBF18d
Blood15d
Blood96h
Blood24h
BloodTwo24h
Blood48h
Blood72h
WholeF
CarcassF
CarcassM
WholeM
GutMidF
GutMidM
SalivF
SalivM
GutHind
GutAnt
GutCaeca
GutPost
Blood3h
BloodNBF
BloodTwoNBF
FatBodyBF
GutMidBF

| | GO.ID | BPCluster: steelblue Size: 65 | Annotated | Significant | Expected | Rank in ClassicF | Weight01F | ClassicF |
|---|---|---|---|---|---|---|---|---|
| *3* | GO:0009084 | glutamine family amino acid biosynthetic... | 12 | 2 | 0.07 | 18 | 0.0021 | 0.00208 |
| *10* | GO:0008152 | metabolic process | 4285 | 34 | 24.86 | 11 | 0.0858 | 0.00043 |

| | GO.ID | MFCluster: steelblue Size: 65 | Annotated | Significant | Expected | Rank in ClassicF | Weight01F | ClassicF |
|---|---|---|---|---|---|---|---|---|
| 2 | GO:0004866 | endopeptidase inhibitor activity | 40 | 4 | 0.23 | 9 | 0.0039 | 7.8e−05 |
| 3 | GO:0008233 | peptidase activity | 500 | 22 | 2.91 | 4 | 0.0042 | 2.8e−15 |
| 4 | GO:0004867 | serine−type endopeptidase inhibitor acti... | 23 | 2 | 0.13 | 15 | 0.0078 | 0.0078 |

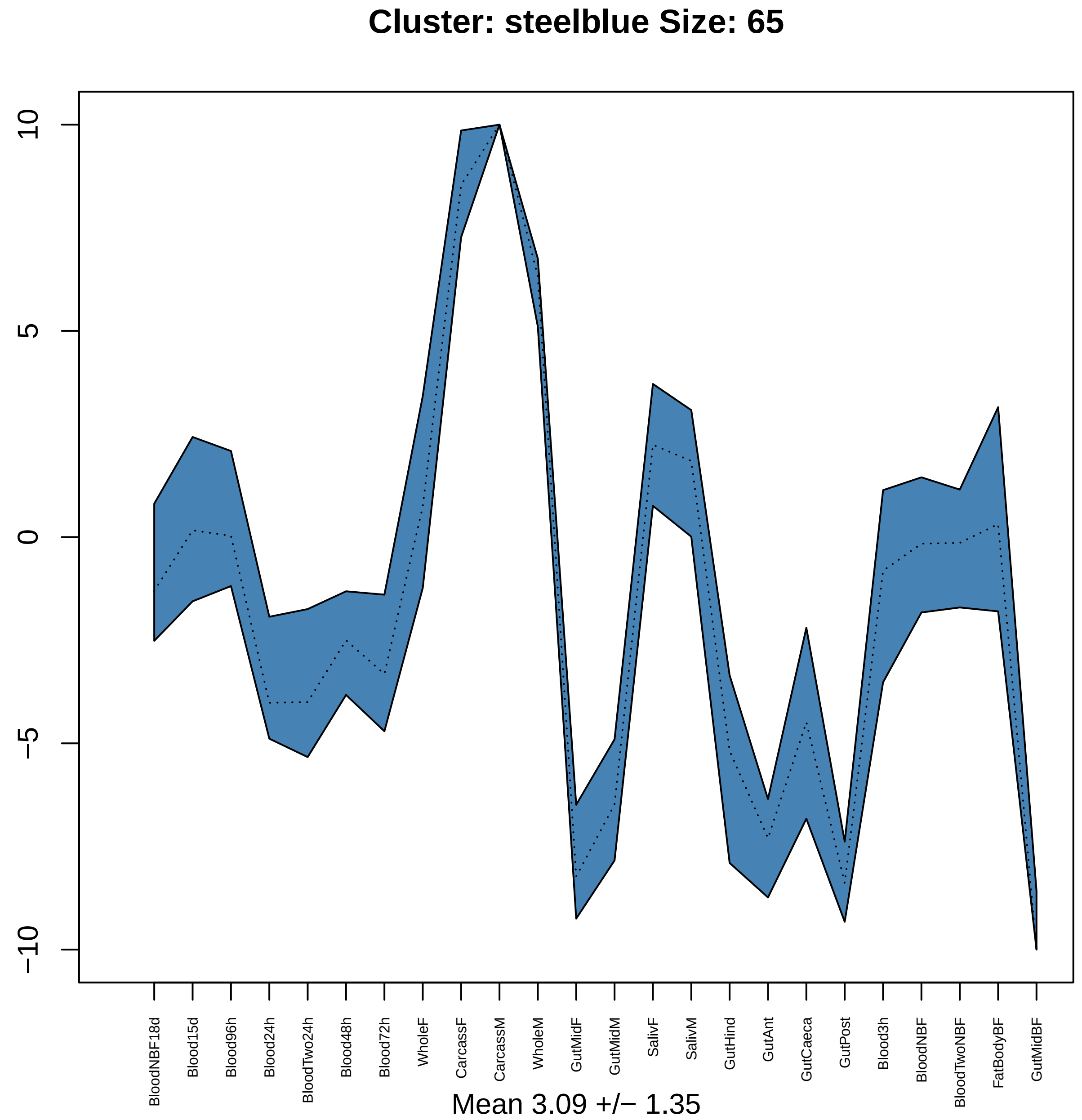
Cluster: steelblue Size: 65
10
5
0
−5
−10
BloodNBF18d
Blood15d
Blood96h
Blood24h
BloodTwo24h
Blood48h
Blood72h
WholeF
CarcassF
CarcassM
WholeM
GutMidF
GutMidM
SalivF
SalivM
GutHind
GutAnt
GutCaeca
GutPost
Blood3h
BloodNBF
BloodTwoNBF
FatBodyBF
GutMidBF
Mean 3.09 +/− 1.35

### Cluster: lightgreen Size: 100

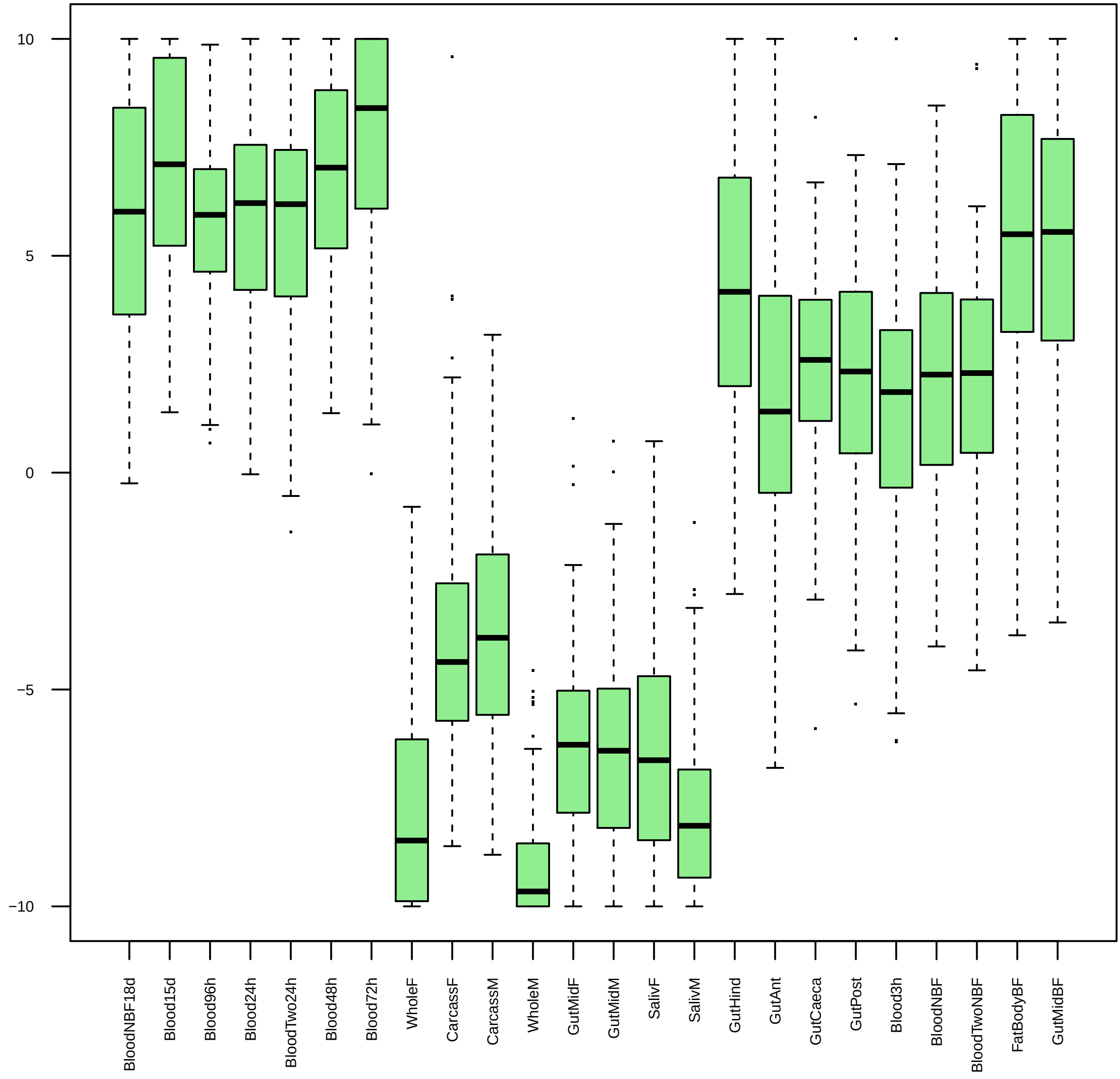

| | GO.ID | BPCluster: lightgreen Size: 100 | Annotated | Significant | Expected | Rank in ClassicF | Weight01F | ClassicF |
|---|---|---|---|---|---|---|---|---|
| 1 | GO:0016482 | cytosolic transport | 15 | 3 | 0.16 | 30 | 0.00052 | 0.00052 |
| 2 | GO:0042078 | germ−line stem cell division | 18 | 3 | 0.20 | 37 | 0.00092 | 0.00092 |
| 3 | GO:0050829 | defense response to Gram−negative bacter... | 43 | 4 | 0.47 | 43 | 0.00119 | 0.00119 |
| 4 | GO:0022416 | chaeta development | 44 | 4 | 0.48 | 46 | 0.00130 | 0.00130 |
| 5 | GO:0030718 | germ−line stem cell population maintenan... | 26 | 3 | 0.29 | 62 | 0.00275 | 0.00275 |
| 6 | GO:0007391 | dorsal closure | 58 | 4 | 0.64 | 75 | 0.00362 | 0.00362 |
| 7 | GO:0022411 | cellular component disassembly | 36 | 3 | 0.40 | 113 | 0.00399 | 0.00698 |
| 8 | GO:0042044 | fluid transport | 10 | 2 | 0.11 | 89 | 0.00505 | 0.00505 |
| 9 | GO:0045887 | positive regulation of synaptic growth a... | 10 | 2 | 0.11 | 90 | 0.00505 | 0.00505 |
| 10 | GO:0098930 | axonal transport | 10 | 2 | 0.11 | 91 | 0.00505 | 0.00505 |
| 11 | GO:0046330 | positive regulation of JNK cascade | 11 | 2 | 0.12 | 102 | 0.00613 | 0.00613 |
| 12 | GO:0046529 | imaginal disc fusion, thorax closure | 12 | 2 | 0.13 | 117 | 0.00730 | 0.00730 |
| 13 | GO:0007432 | salivary gland boundary specification | 12 | 2 | 0.13 | 118 | 0.00730 | 0.00730 |
| 14 | GO:0007517 | muscle organ development | 66 | 4 | 0.72 | 99 | 0.00790 | 0.00575 |
| 15 | GO:0110111 | negative regulation of animal organ morp... | 14 | 2 | 0.15 | 142 | 0.00993 | 0.00993 |
| 16 | GO:0048542 | lymph gland development | 14 | 2 | 0.15 | 143 | 0.00993 | 0.00993 |
| 17 | GO:0007400 | neuroblast fate determination | 14 | 2 | 0.15 | 144 | 0.00993 | 0.00993 |
| 26 | GO:0001763 | morphogenesis of a branching structure | 38 | 3 | 0.42 | 126 | 0.02133 | 0.00812 |
| 27 | GO:0072001 | renal system development | 40 | 3 | 0.44 | 138 | 0.02134 | 0.00936 |
| 28 | GO:0035051 | cardiocyte differentiation | 14 | 2 | 0.15 | 145 | 0.02157 | 0.00993 |
| 30 | GO:0006468 | protein phosphorylation | 271 | 9 | 2.97 | 61 | 0.02210 | 0.00264 |

| | GO.ID | MFCluster: lightgreen Size: 100 | Annotated | Significant | Expected | Rank in ClassicF | Weight01F | ClassicF |
|---|---|---|---|---|---|---|---|---|
| 1 | GO:0004674 | protein serine/threonine kinase activity | 150 | 6 | 1.56 | 4 | 0.0045 | 0.00454 |
| 2 | GO:0004879 | nuclear receptor activity | 10 | 2 | 0.10 | 5 | 0.0046 | 0.00455 |
| 3 | GO:0008092 | cytoskeletal protein binding | 170 | 8 | 1.77 | 1 | 0.0057 | 0.00036 |
| 7 | GO:0005515 | protein binding | 2143 | 36 | 22.30 | 2 | 0.0341 | 0.00062 |

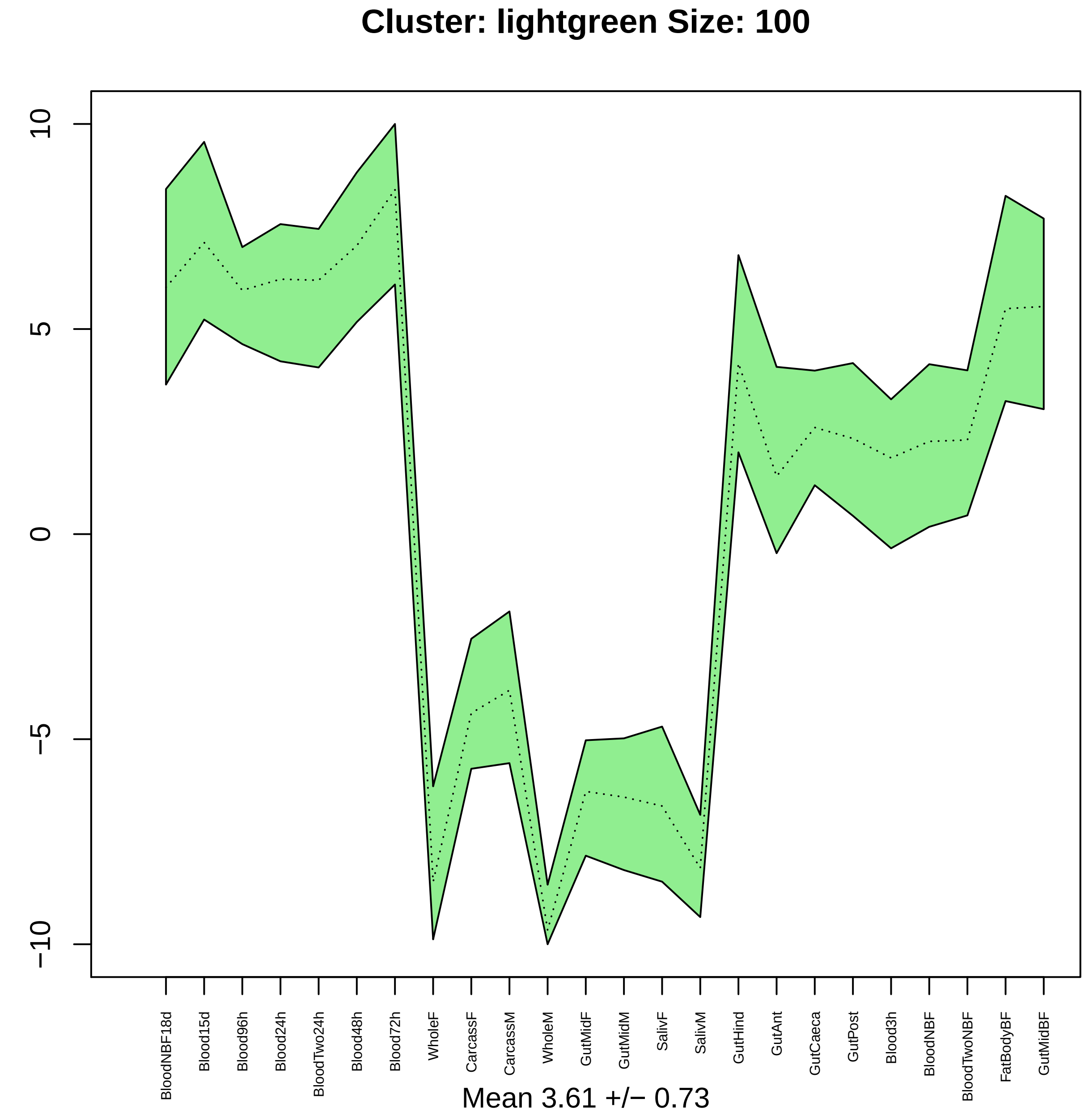
Cluster: lightgreen Size: 100
10
5
0
−5
−10
BloodNBF18d
Blood15d
Blood96h
Blood24h
BloodTwo24h
Blood48h
Blood72h
WholeF
CarcassF
CarcassM
WholeM
GutMidF
GutMidM
SalivF
SalivM
GutHind
GutAnt
GutCaeca
GutPost
Blood3h
BloodNBF
BloodTwoNBF
FatBodyBF
GutMidBF
Mean 3.61 +/− 0.73

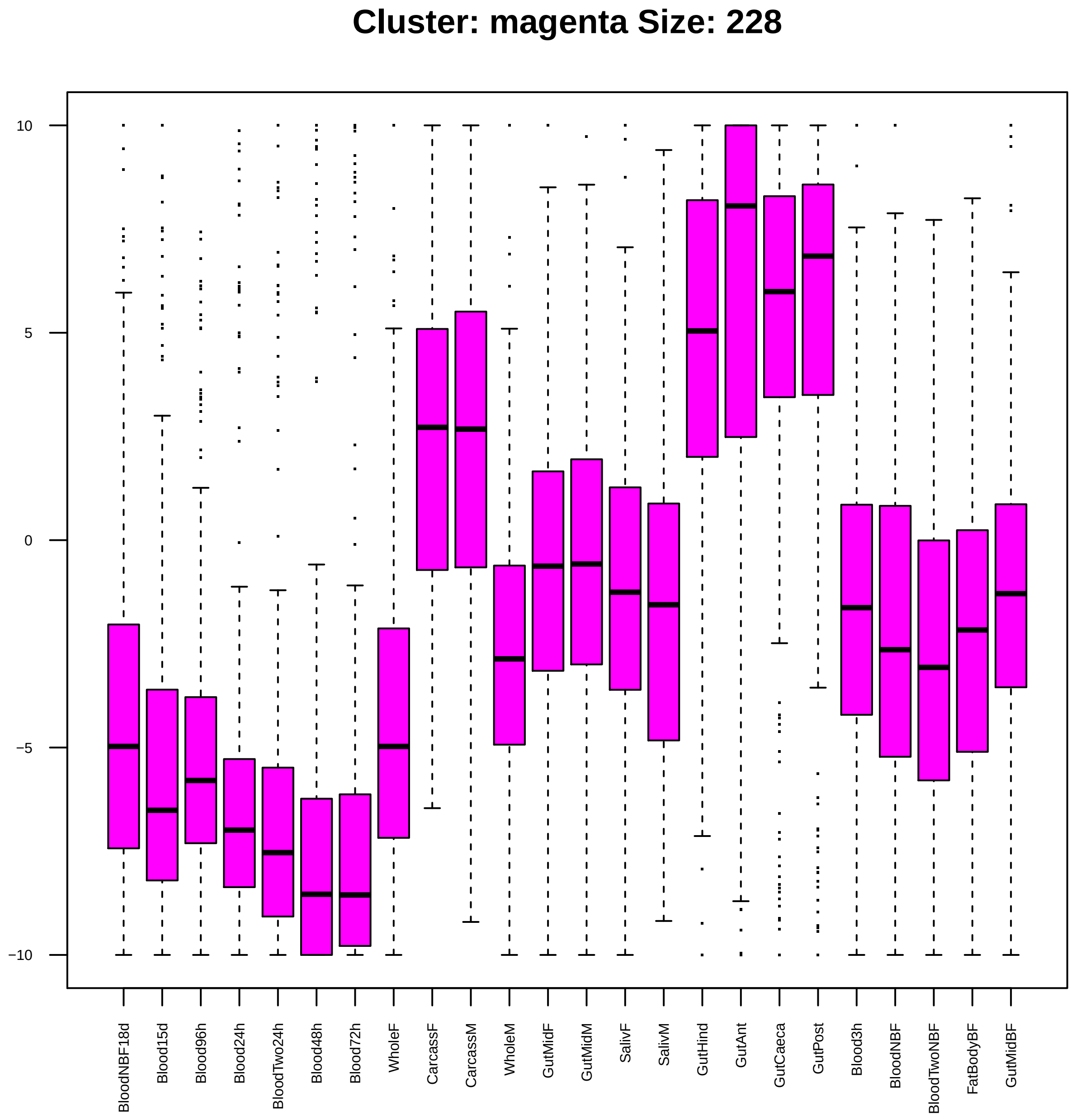
Cluster: magenta Size: 228
10
5
0
−5
−10
BloodNBF18d
Blood15d
Blood96h
Blood24h
BloodTwo24h
Blood48h
Blood72h
WholeF
CarcassF
CarcassM
WholeM
GutMidF
GutMidM
SalivF
SalivM
GutHind
GutAnt
GutCaeca
GutPost
Blood3h
BloodNBF
BloodTwoNBF
FatBodyBF
GutMidBF

| | GO.ID | BPCluster: magenta Size: 228 | Annotated | Significant | Expected | Rank in ClassicF | Weight01F | ClassicF |
|---|---|---|---|---|---|---|---|---|
| 1 | GO:0050907 | detection of chemical stimulus involved ... | 83 | 7 | 1.65 | 3 | 0.0011 | 0.0012 |
| 2 | GO:0008152 | metabolic process | 4285 | 87 | 85.34 | 403 | 0.0031 | 0.4161 |
| 3 | GO:0006836 | neurotransmitter transport | 81 | 4 | 1.61 | 61 | 0.0069 | 0.0774 |

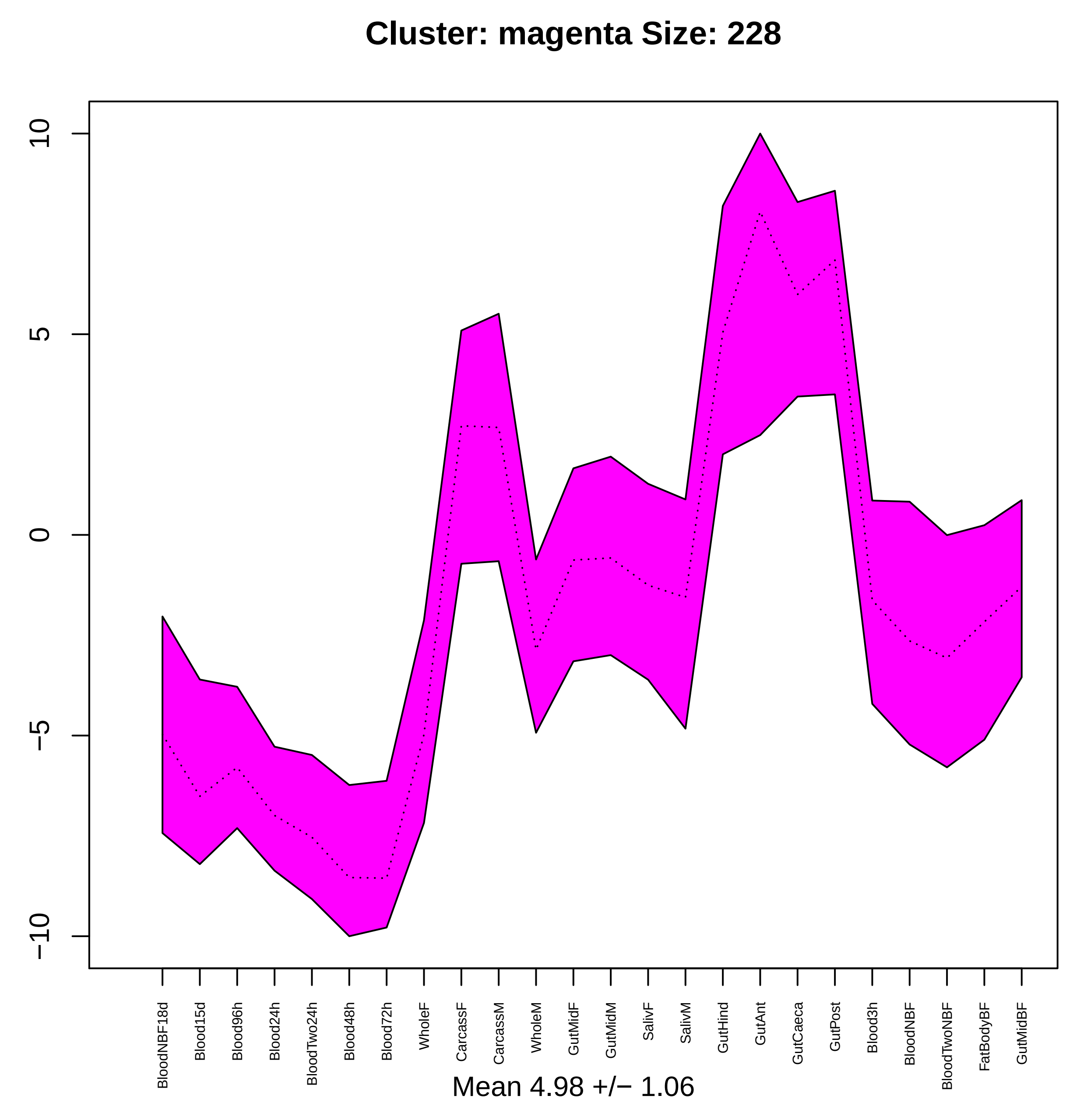

Cluster: magenta Size: 228
10
5
0
−5
−10
BloodNBF18d
Blood15d
Blood96h
Blood24h
BloodTwo24h
Blood48h
Blood72h
WholeF
CarcassF
CarcassM
WholeM
GutMidF
GutMidM
SalivF
SalivM
GutHind
GutAnt
GutCaeca
GutPost
Blood3h
BloodNBF
BloodTwoNBF
FatBodyBF
GutMidBF
Mean 4.98 +/− 1.06

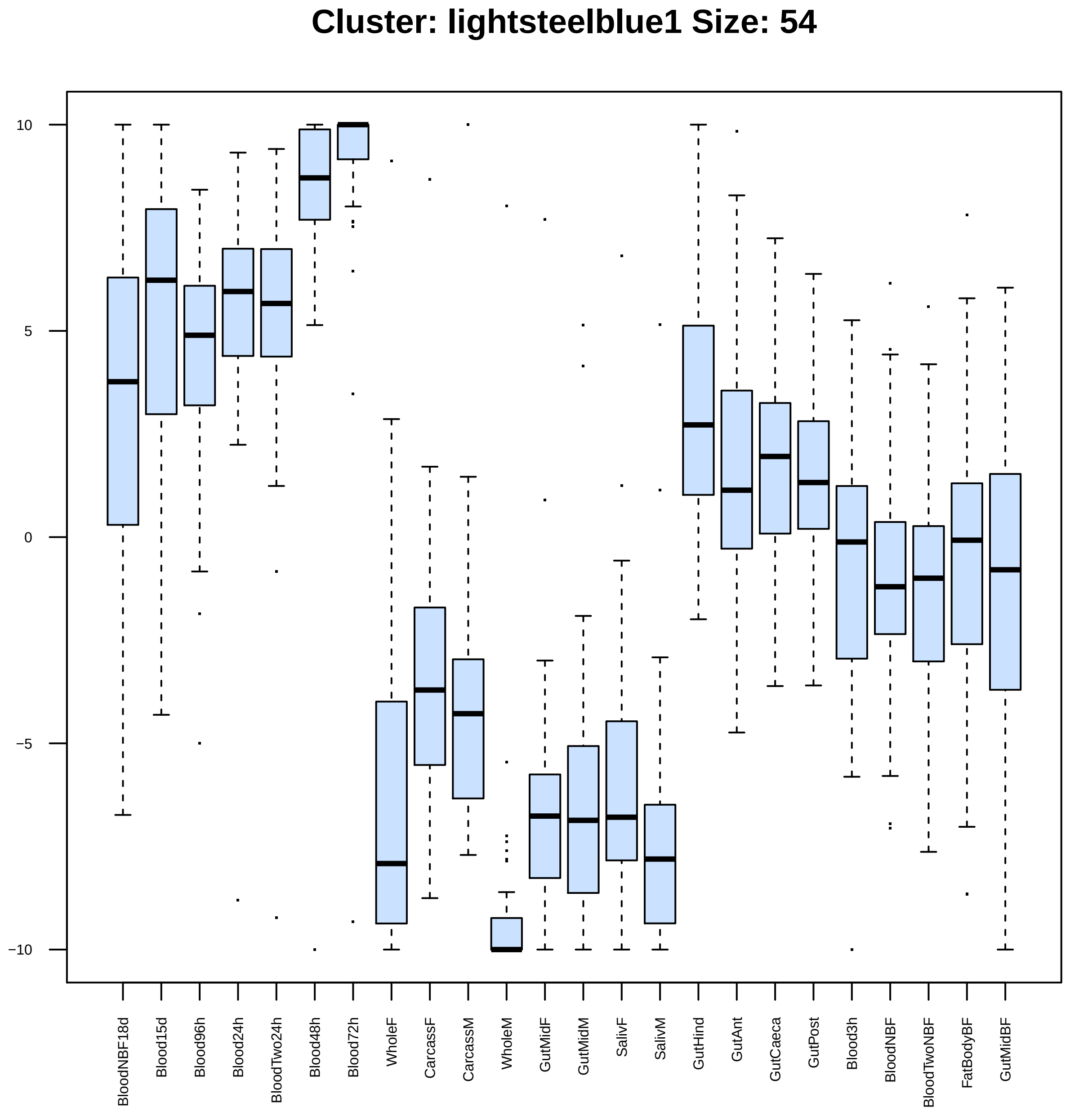
Cluster: lightsteelblue1 Size: 54
10
5
0
−5
−10
BloodNBF18d
Blood15d
Blood96h
Blood24h
BloodTwo24h
Blood48h
Blood72h
WholeF
CarcassF
CarcassM
WholeM
GutMidF
GutMidM
SalivF
SalivM
GutHind
GutAnt
GutCaeca
GutPost
Blood3h
BloodNBF
BloodTwoNBF
FatBodyBF
GutMidBF

| | GO.ID | BPCluster: lightsteelblue1 Size: 54 | Annotated | Significant | Expected | Rank in ClassicF | Weight01F | ClassicF |
|---|---|---|---|---|---|---|---|---|
| *1* | GO:2000045 | regulation of G1/S transition of mitotic... | 16 | 2 | 0.09 | 20 | 0.0033 | 0.0033 |
| *2* | GO:0120032 | regulation of plasma membrane bounded ce... | 13 | 2 | 0.07 | 16 | 0.0053 | 0.0022 |

| | GO.ID | MFCluster: lightsteelblue1 Size: 54 | Annotated | Significant | Expected | Rank in ClassicF | Weight01F | ClassicF |
|---|---|---|---|---|---|---|---|---|
| 1 | GO:0003729 | mRNA binding | 99 | 4 | 0.56 | 6 | 0.0023 | 0.00230 |
| 2 | GO:0000166 | nucleotide binding | 998 | 10 | 5.68 | 16 | 0.0025 | 0.04892 |
| 3 | GO:0019901 | protein kinase binding | 24 | 2 | 0.14 | 7 | 0.0080 | 0.00805 |
| 5 | GO:0003723 | RNA binding | 307 | 8 | 1.75 | 3 | 0.0198 | 0.00028 |

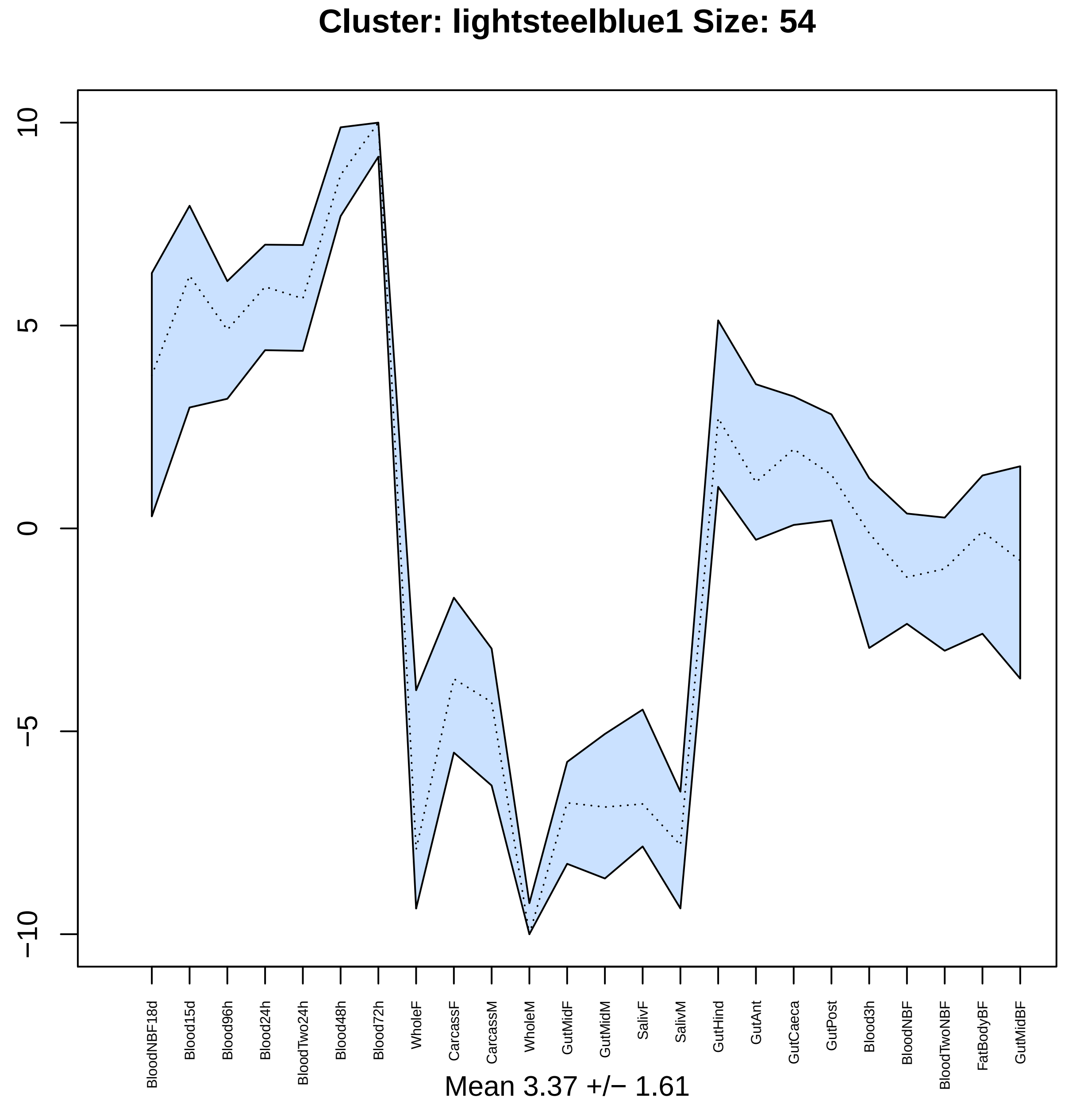
Cluster: lightsteelblue1 Size: 54
10
5
0
−5
−10
BloodNBF18d
Blood15d
Blood96h
Blood24h
BloodTwo24h
Blood48h
Blood72h
WholeF
CarcassF
CarcassM
WholeM
GutMidF
GutMidM
SalivF
SalivM
GutHind
GutAnt
GutCaeca
GutPost
Blood3h
BloodNBF
BloodTwoNBF
FatBodyBF
GutMidBF
Mean 3.37 +/− 1.61

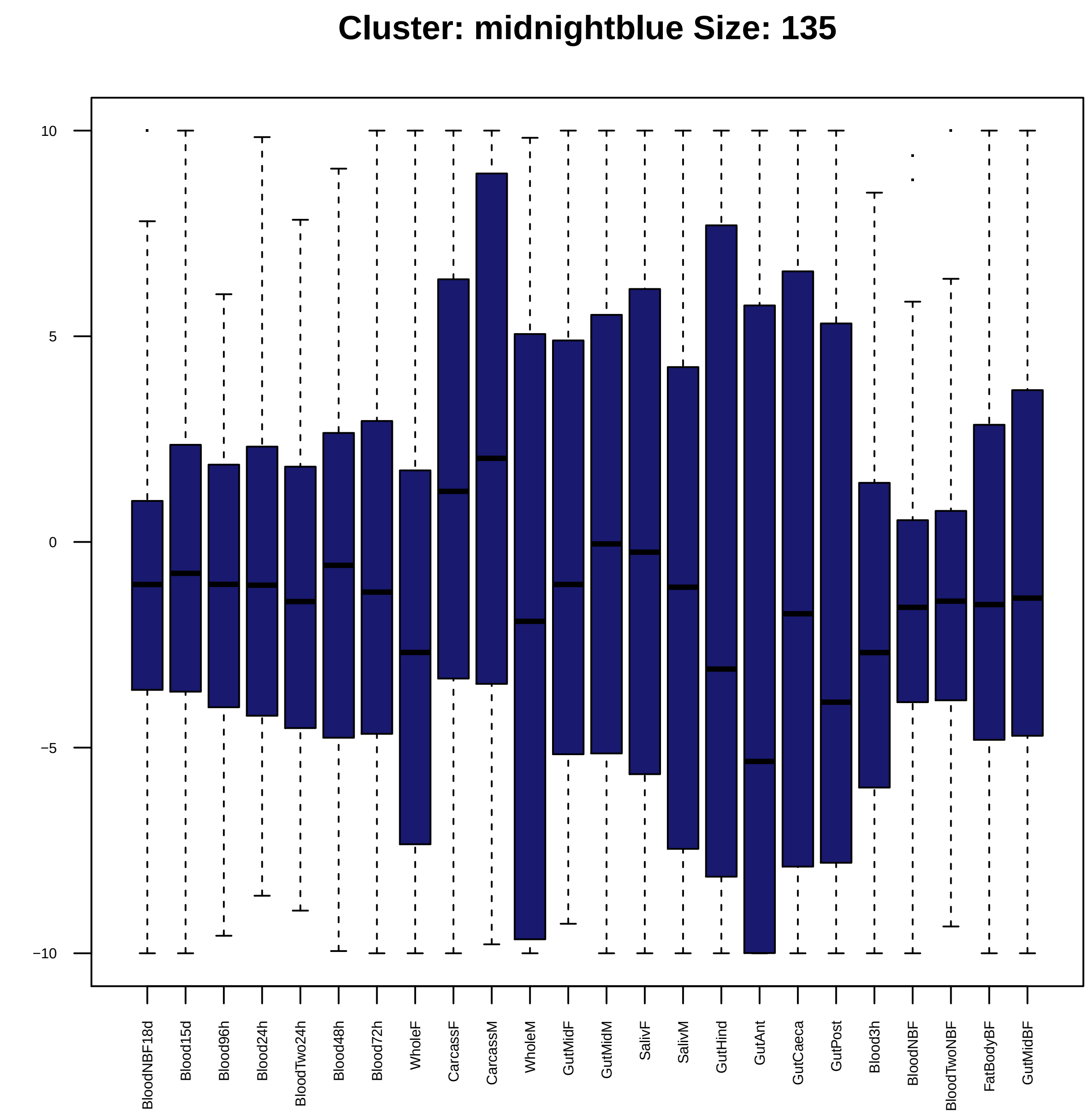

Cluster: midnightblue Size: 135
10
5
0
−5
−10
BloodNBF18d
Blood15d
Blood96h
Blood24h
BloodTwo24h
Blood48h
Blood72h
WholeF
CarcassF
CarcassM
WholeM
GutMidF
GutMidM
SalivF
SalivM
GutHind
GutAnt
GutCaeca
GutPost
Blood3h
BloodNBF
BloodTwoNBF
FatBodyBF
GutMidBF

| | GO.ID | BPCluster: midnightblue Size: 135 | Annotated | Significant | Expected | Rank in ClassicF | Weight01F | ClassicF |
|---|---|---|---|---|---|---|---|---|
| *1* | GO:0009100 | glycoprotein metabolic process | 60 | 5 | 0.88 | 9 | 0.0012 | 0.0018 |
| *2* | GO:0000281 | mitotic cytokinesis | 26 | 3 | 0.38 | 18 | 0.0063 | 0.0063 |
| *3* | GO:0007469 | antennal development | 10 | 2 | 0.15 | 21 | 0.0090 | 0.0090 |
| *4* | GO:0007314 | oocyte anterior/posterior axis specifica... | 44 | 4 | 0.65 | 13 | 0.0146 | 0.0038 |
| *12* | GO:0030707 | ovarian follicle cell development | 121 | 6 | 1.78 | 20 | 0.0268 | 0.0086 |
| *20* | GO:0009798 | axis specification | 108 | 7 | 1.59 | 5 | 0.0415 | 0.0010 |

| | GO.ID | MFCluster: midnightblue Size: 135 | Annotated | Significant | Expected | Rank in ClassicF | Weight01F | ClassicF |
|---|---|---|---|---|---|---|---|---|
| *1* | GO:0008324 | cation transmembrane transporter activit... | 210 | 4 | 2.80 | 61 | 0.0014 | 0.3062 |
| *2* | GO:0030554 | adenyl nucleotide binding | 606 | 11 | 8.07 | 33 | 0.0034 | 0.1810 |
| *6* | GO:0016791 | phosphatase activity | 97 | 5 | 1.29 | 2 | 0.0870 | 0.0093 |

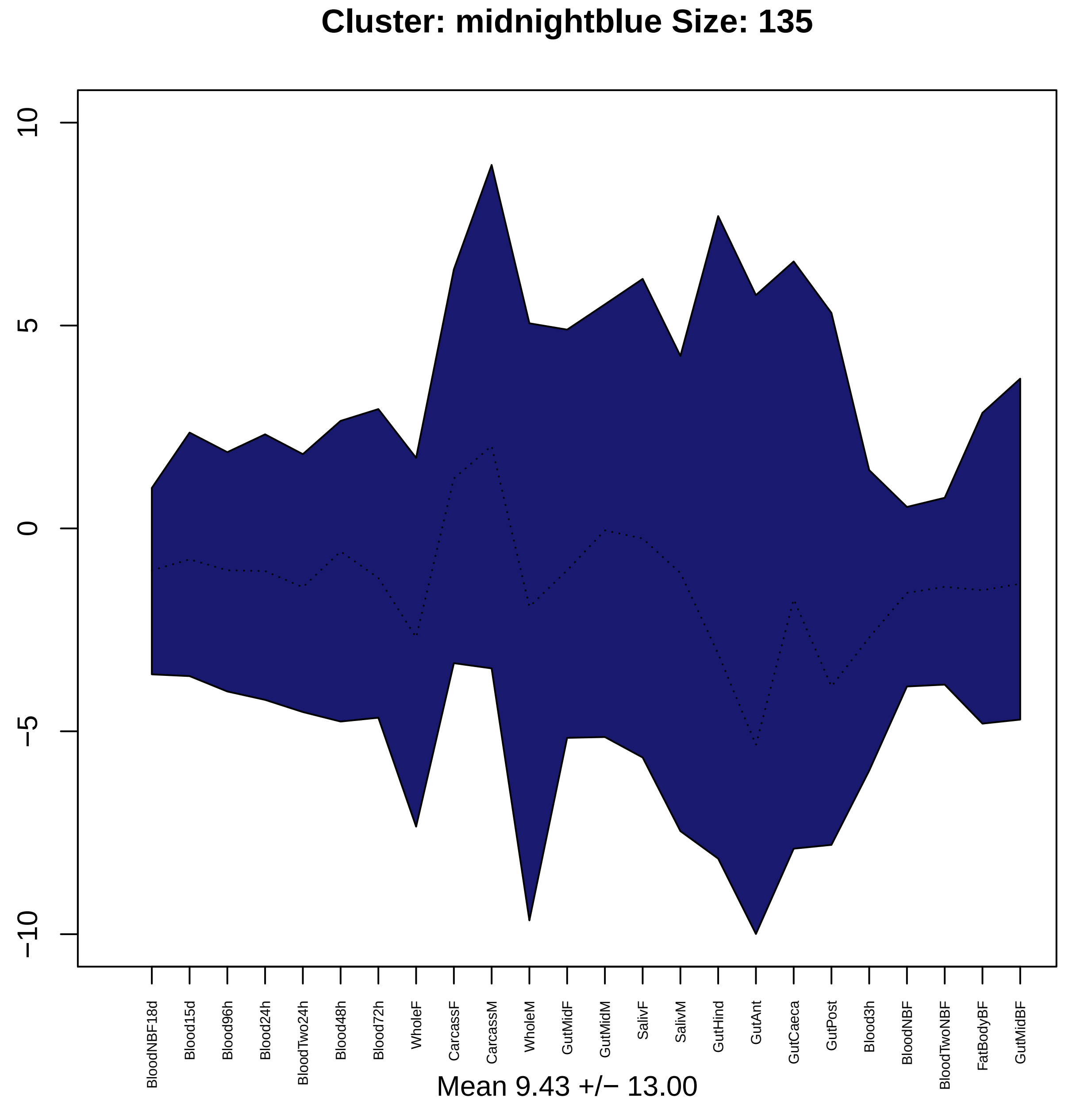
Cluster: midnightblue Size: 135
10
5
0
−5
−10
BloodNBF18d
Blood15d
Blood96h
Blood24h
BloodTwo24h
Blood48h
Blood72h
WholeF
CarcassF
CarcassM
WholeM
GutMidF
GutMidM
SalivF
SalivM
GutHind
GutAnt
GutCaeca
GutPost
Blood3h
BloodNBF
BloodTwoNBF
FatBodyBF
GutMidBF
Mean 9.43 +/− 13.00

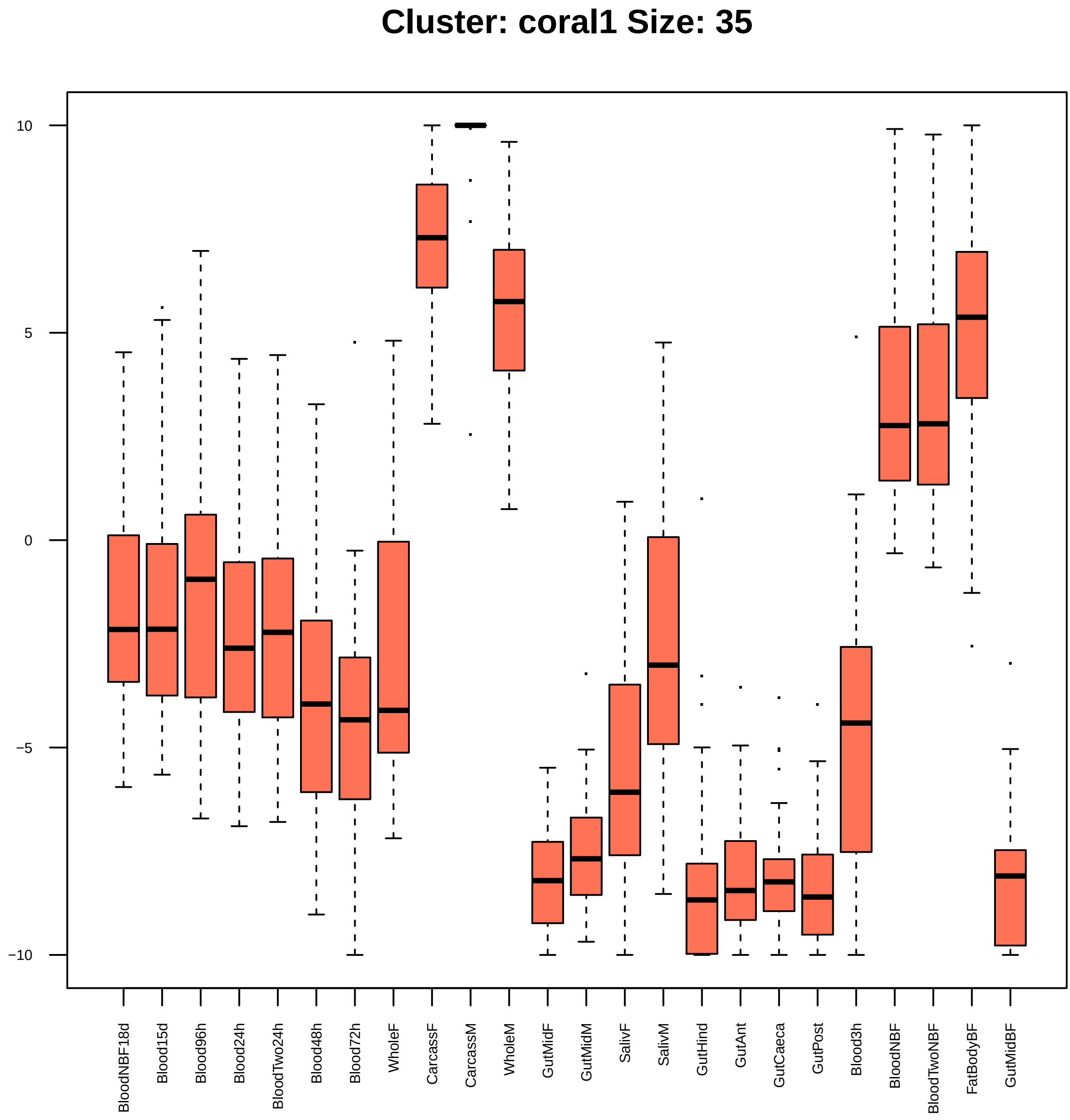
Cluster: coral1 Size: 35
10
5
0
−5
−10
BloodNBF18d
Blood15d
Blood96h
Blood24h
BloodTwo24h
Blood48h
Blood72h
WholeF
CarcassF
CarcassM
WholeM
GutMidF
GutMidM
SalivF
SalivM
GutHind
GutAnt
GutCaeca
GutPost
Blood3h
BloodNBF
BloodTwoNBF
FatBodyBF
GutMidBF

|  | GO.ID | BPCluster: coral1 Size: 35 | Annotated | Significant | Expected | Rank in ClassicF | Weight01F | ClassicF |
|---|---|---|---|---|---|---|---|---|
| *1* | GO:0055114 | oxidation−reduction process | 545 | 6 | 1.54 | 1 | 0.0029 | 0.0029 |
| *2* | GO:0009058 | biosynthetic process | 1357 | 4 | 3.83 | 41 | 0.0053 | 0.5533 |

| | GO.ID | MFCluster: coral1 Size: 35 | Annotated | Significant | Expected | Rank in ClassicF | Weight01F | ClassicF |
|---|---|---|---|---|---|---|---|---|
| *1* | GO:0042302 | structural constituent of cuticle | 99 | 3 | 0.29 | 1 | 0.019 | 0.0028 |
| *3* | GO:0016747 | transferase activity, transferring acyl ... | 112 | 3 | 0.33 | 3 | 0.032 | 0.0040 |
| *12* | GO:0016491 | oxidoreductase activity | 523 | 6 | 1.52 | 2 | 0.059 | 0.0030 |

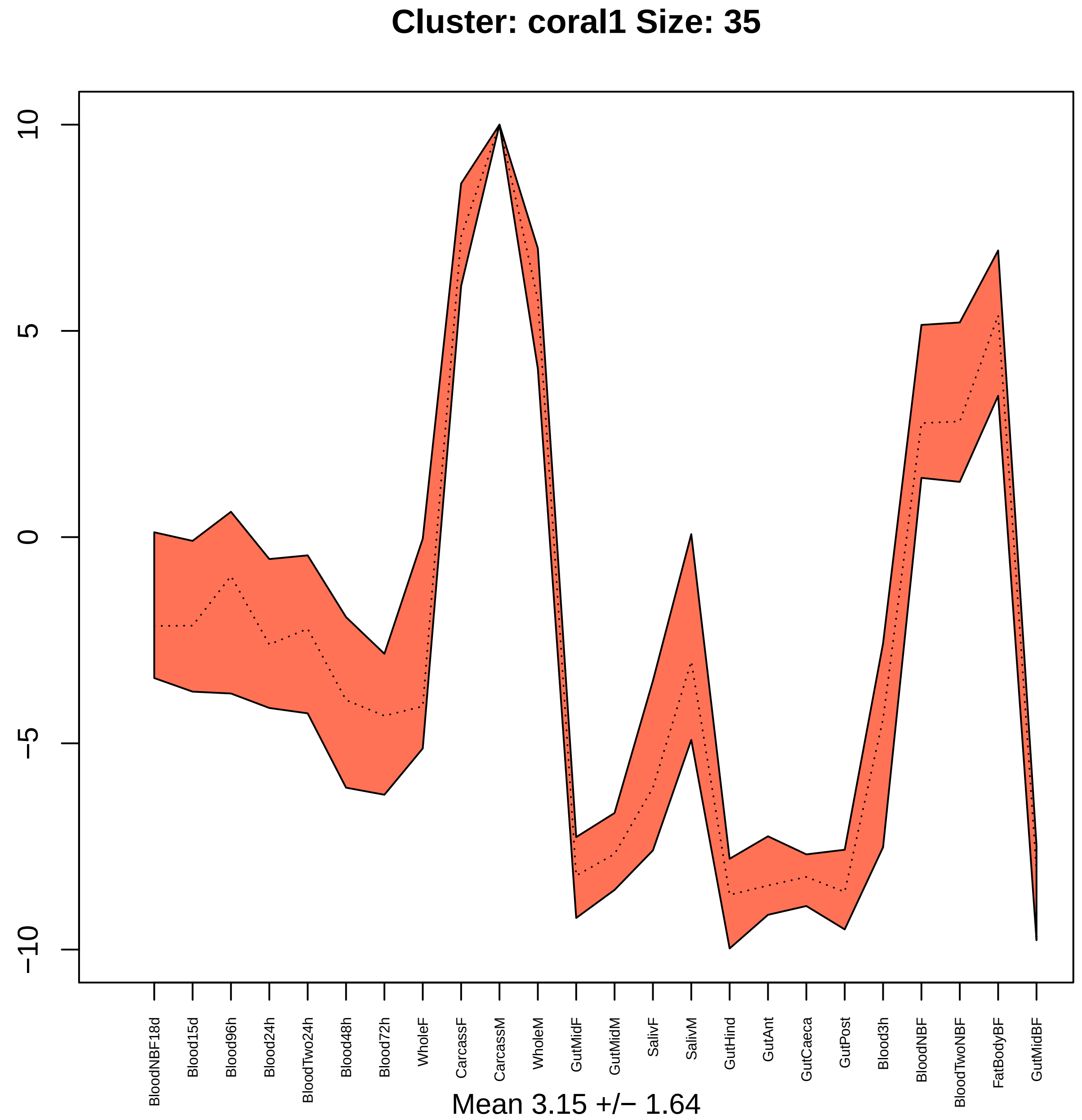
Cluster: coral1 Size: 35
10
5
0
−5
−10
BloodNBF18d
Blood15d
Blood96h
Blood24h
BloodTwo24h
Blood48h
Blood72h
WholeF
CarcassF
CarcassM
WholeM
GutMidF
GutMidM
SalivF
SalivM
GutHind
GutAnt
GutCaeca
GutPost
Blood3h
BloodNBF
BloodTwoNBF
FatBodyBF
GutMidBF
Mean 3.15 +/− 1.64

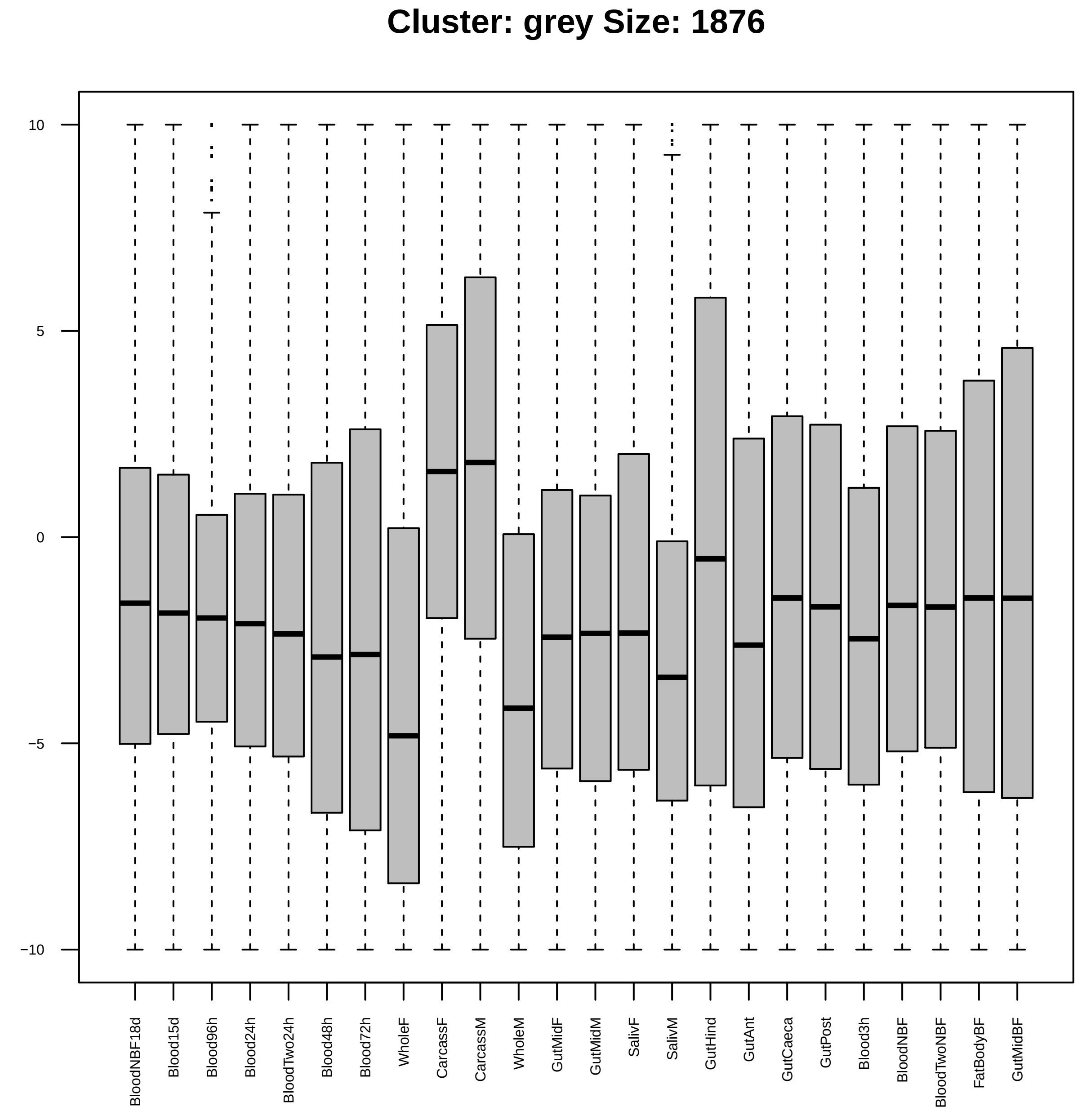
Cluster: grey Size: 1876
10
5
0
−5
−10
BloodNBF18d
Blood15d
Blood96h
Blood24h
BloodTwo24h
Blood48h
Blood72h
WholeF
CarcassF
CarcassM
WholeM
GutMidF
GutMidM
SalivF
SalivM
GutHind
GutAnt
GutCaeca
GutPost
Blood3h
BloodNBF
BloodTwoNBF
FatBodyBF
GutMidBF

| | GO.ID | BPCluster: grey Size: 1876 | Annotated | Significant | Expected | Rank in ClassicF | Weight01F | ClassicF |
|---|---|---|---|---|---|---|---|---|
| *1* | GO:0055085 | transmembrane transport | 469 | 105 | 84.95 | 36 | 6.7e−05 | 0.00852 |
| *2* | GO:1901606 | alpha−amino acid catabolic process | 27 | 12 | 4.89 | 7 | 0.0014 | 0.00140 |
| *3* | GO:0055059 | asymmetric neuroblast division | 21 | 10 | 3.80 | 10 | 0.0019 | 0.00185 |
| *4* | GO:0007298 | border follicle cell migration | 58 | 20 | 10.50 | 13 | 0.0021 | 0.00208 |
| *5* | GO:0050803 | regulation of synapse structure or activ... | 56 | 15 | 10.14 | 247 | 0.0029 | 0.06925 |
| *6* | GO:0006576 | cellular biogenic amine metabolic proces... | 16 | 8 | 2.90 | 17 | 0.0037 | 0.00367 |
| *7* | GO:0019318 | hexose metabolic process | 23 | 10 | 4.17 | 18 | 0.0042 | 0.00424 |
| *8* | GO:0017144 | drug metabolic process | 246 | 48 | 44.56 | 705 | 0.0057 | 0.30520 |
| *9* | GO:0007309 | oocyte axis specification | 60 | 16 | 10.87 | 235 | 0.0059 | 0.06418 |
| *10* | GO:0010721 | negative regulation of cell development | 23 | 9 | 4.17 | 59 | 0.0059 | 0.01460 |
| *11* | GO:0007391 | dorsal closure | 58 | 17 | 10.50 | 90 | 0.0065 | 0.02471 |
| *12* | GO:0035099 | hemocyte migration | 11 | 6 | 1.99 | 25 | 0.0070 | 0.00702 |
| *13* | GO:0006637 | acyl−CoA metabolic process | 11 | 6 | 1.99 | 26 | 0.0070 | 0.00702 |
| *14* | GO:0008340 | determination of adult lifespan | 68 | 21 | 12.32 | 28 | 0.0072 | 0.00716 |
| *15* | GO:0005975 | carbohydrate metabolic process | 210 | 63 | 38.04 | 1 | 0.0083 | 1.4e−05 |
| *21* | GO:0035088 | establishment or maintenance of apical/b... | 27 | 11 | 4.89 | 19 | 0.0126 | 0.00501 |
| *28* | GO:0006810 | transport | 1142 | 251 | 206.84 | 3 | 0.0193 | 0.00014 |

| | GO.ID | MFCluster: grey Size: 1876 | Annotated | Significant | Expected | Rank in ClassicF | Weight01F | ClassicF |
|---|---|---|---|---|---|---|---|---|
| *1* | GO:0030246 | carbohydrate binding | 60 | 22 | 10.69 | 1 | 0.00039 | 0.00039 |
| *2* | GO:0016903 | oxidoreductase activity, acting on the a... | 33 | 13 | 5.88 | 2 | 0.00090 | 0.00286 |
| *3* | GO:0005215 | transporter activity | 548 | 117 | 97.60 | 6 | 0.00721 | 0.01546 |
| *25* | GO:0038024 | cargo receptor activity | 17 | 8 | 3.03 | 3 | 0.07333 | 0.00525 |

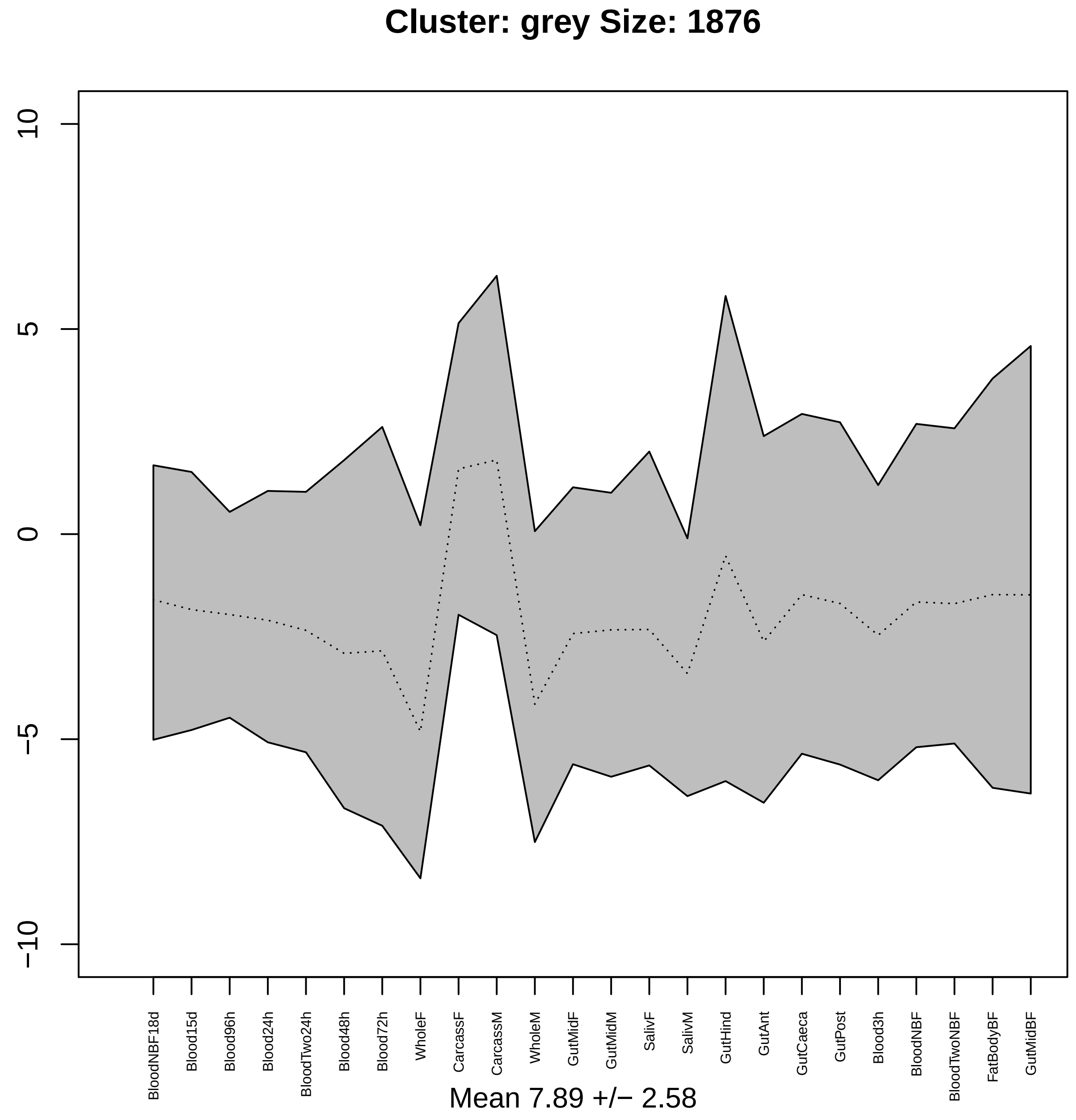

Cluster: grey Size: 1876
10
5
0
−5
−10
BloodNBF18d
Blood15d
Blood96h
Blood24h
BloodTwo24h
Blood48h
Blood72h
WholeF
CarcassF
CarcassM
WholeM
GutMidF
GutMidM
SalivF
SalivM
GutHind
GutAnt
GutCaeca
GutPost
Blood3h
BloodNBF
BloodTwoNBF
FatBodyBF
GutMidBF
Mean 7.89 +/− 2.58

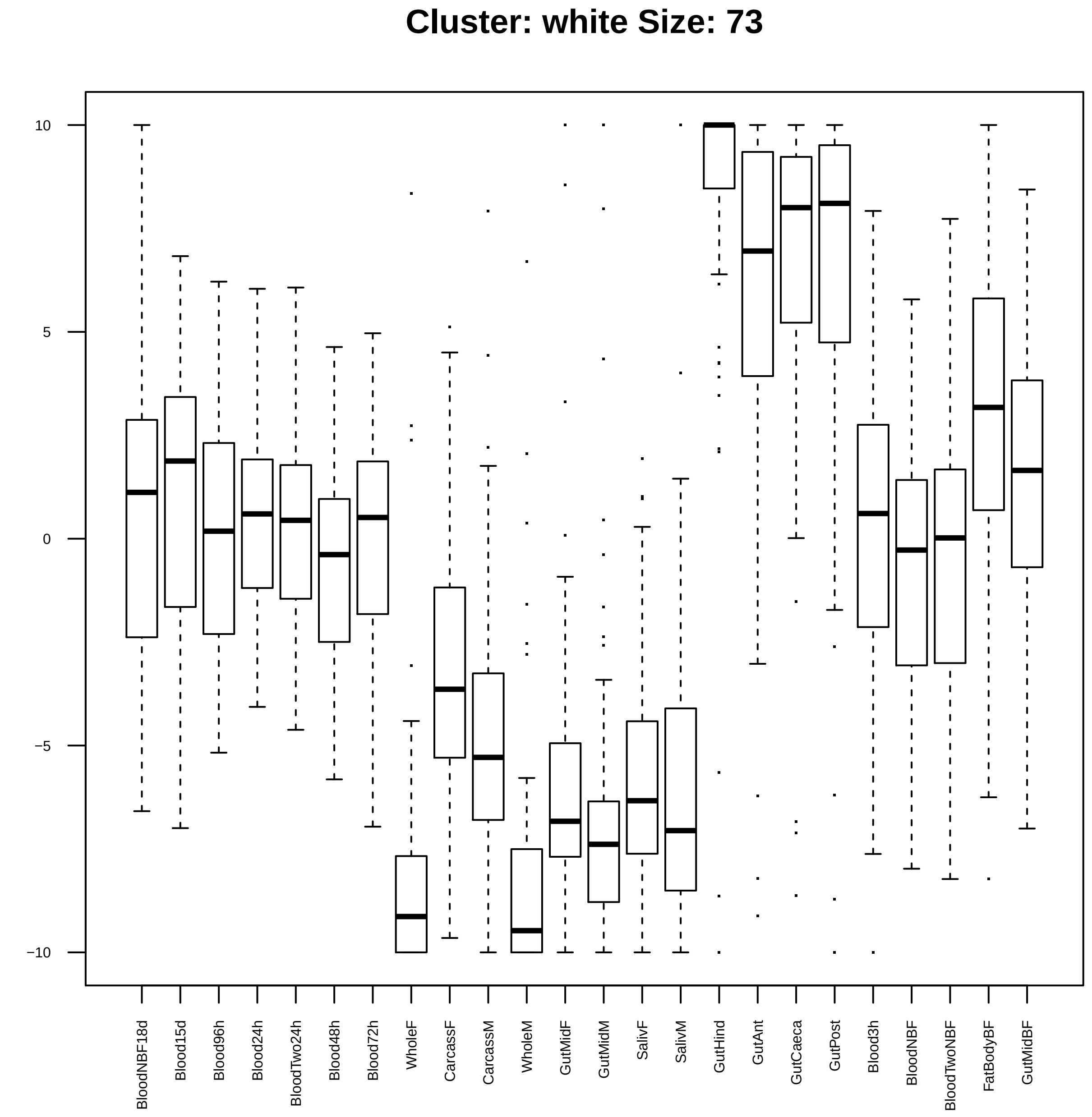
Cluster: white Size: 73
10
5
0
−5
−10
BloodNBF18d
Blood15d
Blood96h
Blood24h
BloodTwo24h
Blood48h
Blood72h
WholeF
CarcassF
CarcassM
WholeM
GutMidF
GutMidM
SalivF
SalivM
GutHind
GutAnt
GutCaeca
GutPost
Blood3h
BloodNBF
BloodTwoNBF
FatBodyBF
GutMidBF

| | GO.ID | BPCluster: white Size: 73 | Annotated | Significant | Expected | Rank in ClassicF | Weight01F | ClassicF |
|---|---|---|---|---|---|---|---|---|
| 3 | GO:0051298 | centrosome duplication | 51 | 4 | 0.42 | 33 | 0.0008 | 0.0008 |
| 4 | GO:0043161 | proteasome−mediated ubiquitin−dependent ... | 37 | 3 | 0.31 | 40 | 0.0035 | 0.0035 |
| 5 | GO:0009893 | positive regulation of metabolic process | 218 | 3 | 1.81 | 217 | 0.0083 | 0.2714 |

| | GO.ID | MFCluster: white Size: 73 | Annotated | Significant | Expected | Rank in ClassicF | Weight01F | ClassicF |
|---|---|---|---|---|---|---|---|---|
| *2* | GO:0004298 | threonine−type endopeptidase activity | 14 | 2 | 0.1 | 3 | 0.0041 | 0.0041 |

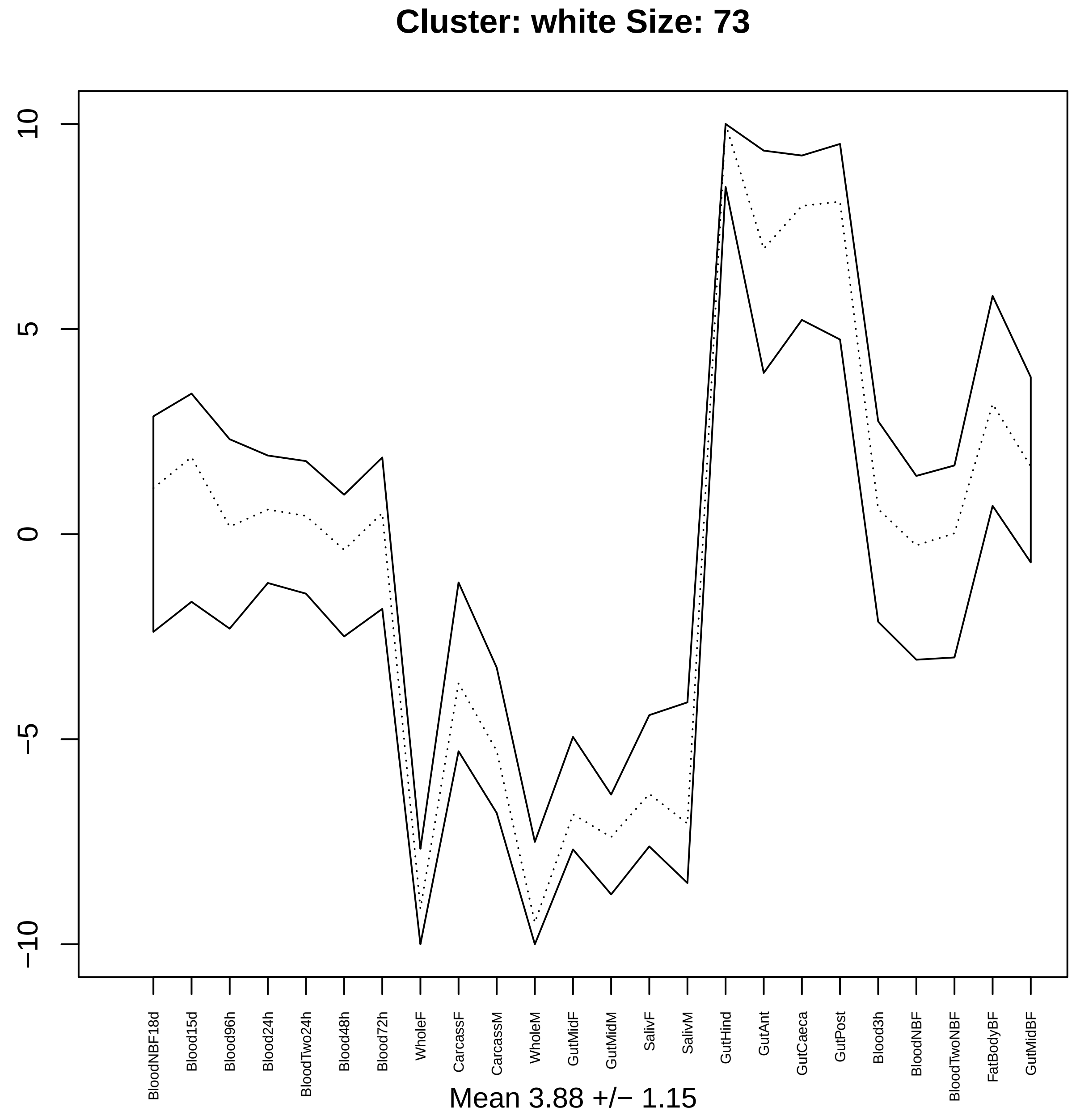

Cluster: white Size: 73
10
5
0
−5
−10
BloodNBF18d
Blood15d
Blood96h
Blood24h
BloodTwo24h
Blood48h
Blood72h
WholeF
CarcassF
CarcassM
WholeM
GutMidF
GutMidM
SalivF
SalivM
GutHind
GutAnt
GutCaeca
GutPost
Blood3h
BloodNBF
BloodTwoNBF
FatBodyBF
GutMidBF
Mean 3.88 +/− 1.15

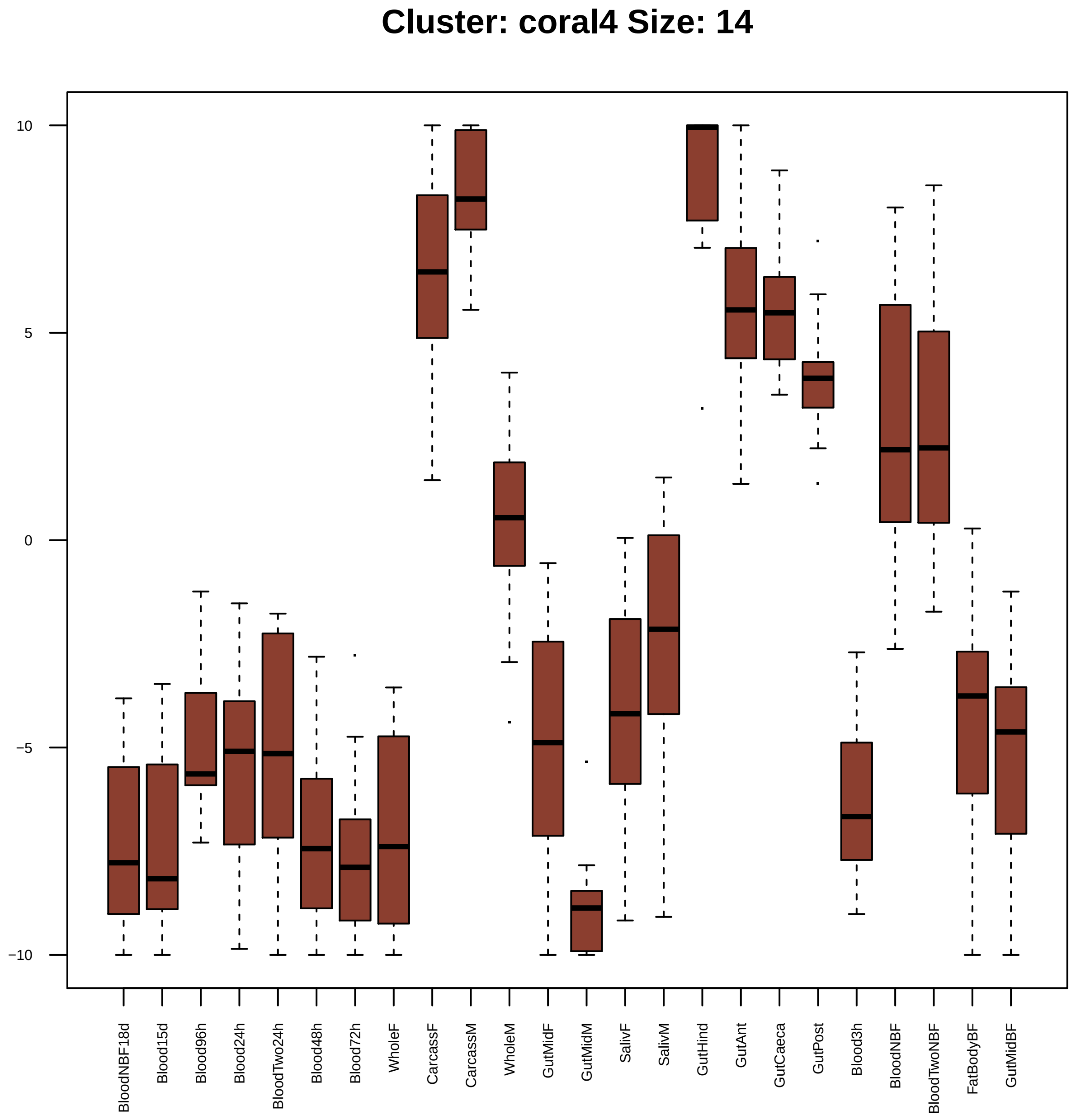

Cluster: coral4 Size: 14
10
5
0
−5
−10
BloodNBF18d
Blood15d
Blood96h
Blood24h
BloodTwo24h
Blood48h
Blood72h
WholeF
CarcassF
CarcassM
WholeM
GutMidF
GutMidM
SalivF
SalivM
GutHind
GutAnt
GutCaeca
GutPost
Blood3h
BloodNBF
BloodTwoNBF
FatBodyBF
GutMidBF

| | GO.ID | BPCluster: coral4 Size: 14 | Annotated | Significant | Expected | Rank in ClassicF | Weight01F | ClassicF |
|---|---|---|---|---|---|---|---|---|
| *3* | GO:0022900 | electron transport chain | 37 | 5 | 0.06 | 17 | 0.0033 | 2.2e−09 |
| *11* | GO:0006164 | purine nucleotide biosynthetic process | 68 | 3 | 0.12 | 47 | 1.0000 | 0.00018 |
| *12* | GO:0006139 | nucleobase−containing compound metabolic... | 1429 | 7 | 2.46 | 62 | 1.0000 | 0.00390 |
| *30* | GO:1901137 | carbohydrate derivative biosynthetic pro... | 161 | 3 | 0.28 | 59 | 1.0000 | 0.00225 |

| | GO.ID | MFCluster: coral4 Size: 14 | Annotated | Significant | Expected | Rank in ClassicF | Weight01F | ClassicF |
|---|---|---|---|---|---|---|---|---|
| *5* | GO:0003954 | NADH dehydrogenase activity | 25 | 6 | 0.04 | 1 | 0.0042 | 4.2e−13 |
| *19* | GO:0022804 | active transmembrane transporter activit... | 136 | 3 | 0.21 | 17 | 1.0000 | 0.00097 |
| *26* | GO:0015077 | monovalent inorganic cation transmembran... | 138 | 3 | 0.21 | 18 | 1.0000 | 0.00101 |
| *28* | GO:0015399 | primary active transmembrane transporter... | 76 | 3 | 0.12 | 15 | 1.0000 | 0.00017 |
| *30* | GO:0043492 | ATPase activity, coupled to movement of ... | 73 | 3 | 0.11 | 13 | 1.0000 | 0.00016 |

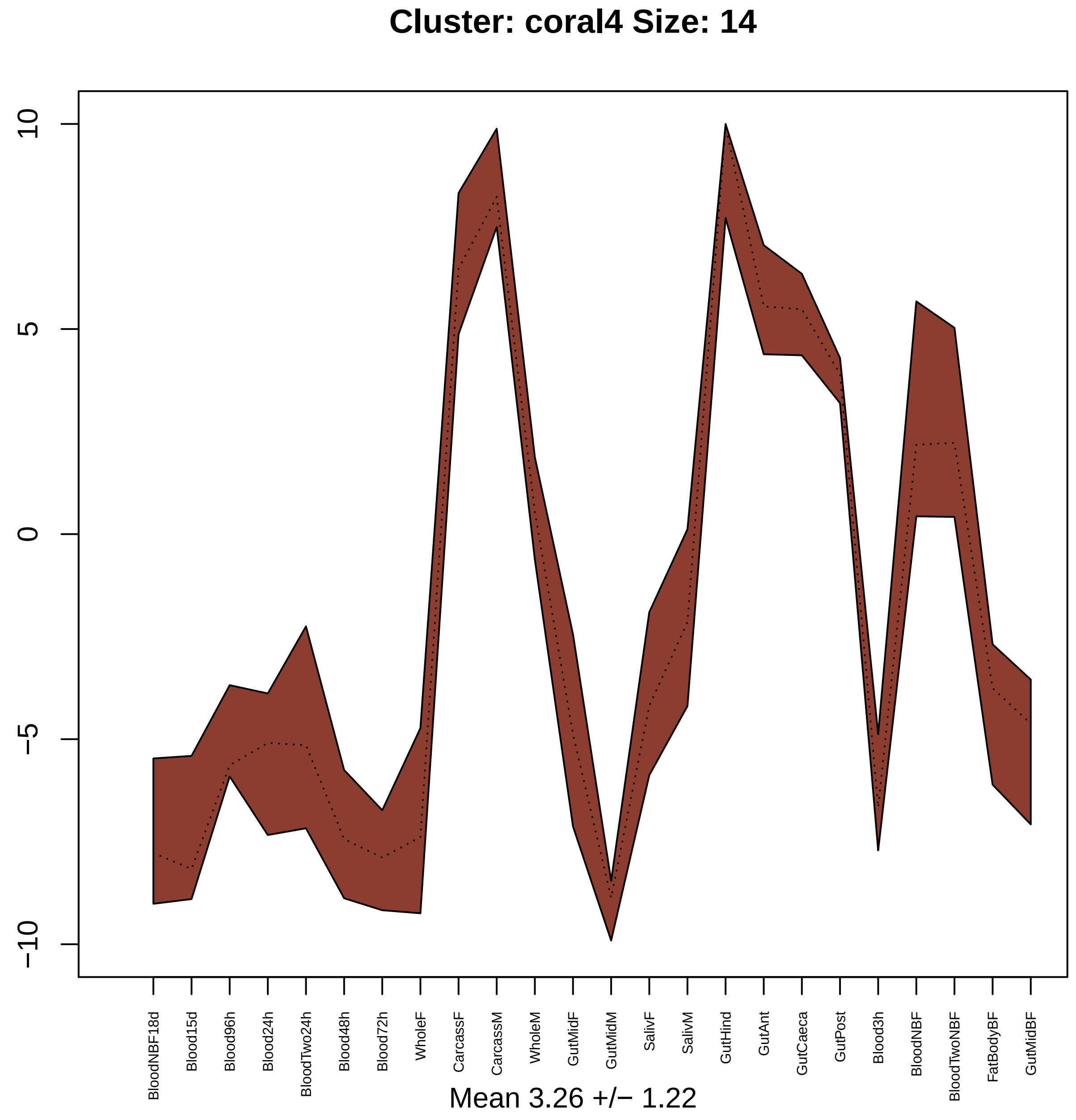

Cluster: coral4 Size: 14
10
5
0
−5
−10
BloodNBF18d
Blood15d
Blood96h
Blood24h
BloodTwo24h
Blood48h
Blood72h
WholeF
CarcassF
CarcassM
WholeM
GutMidF
GutMidM
SalivF
SalivM
GutHind
GutAnt
GutCaeca
GutPost
Blood3h
BloodNBF
BloodTwoNBF
FatBodyBF
GutMidBF
Mean 3.26 +/− 1.22

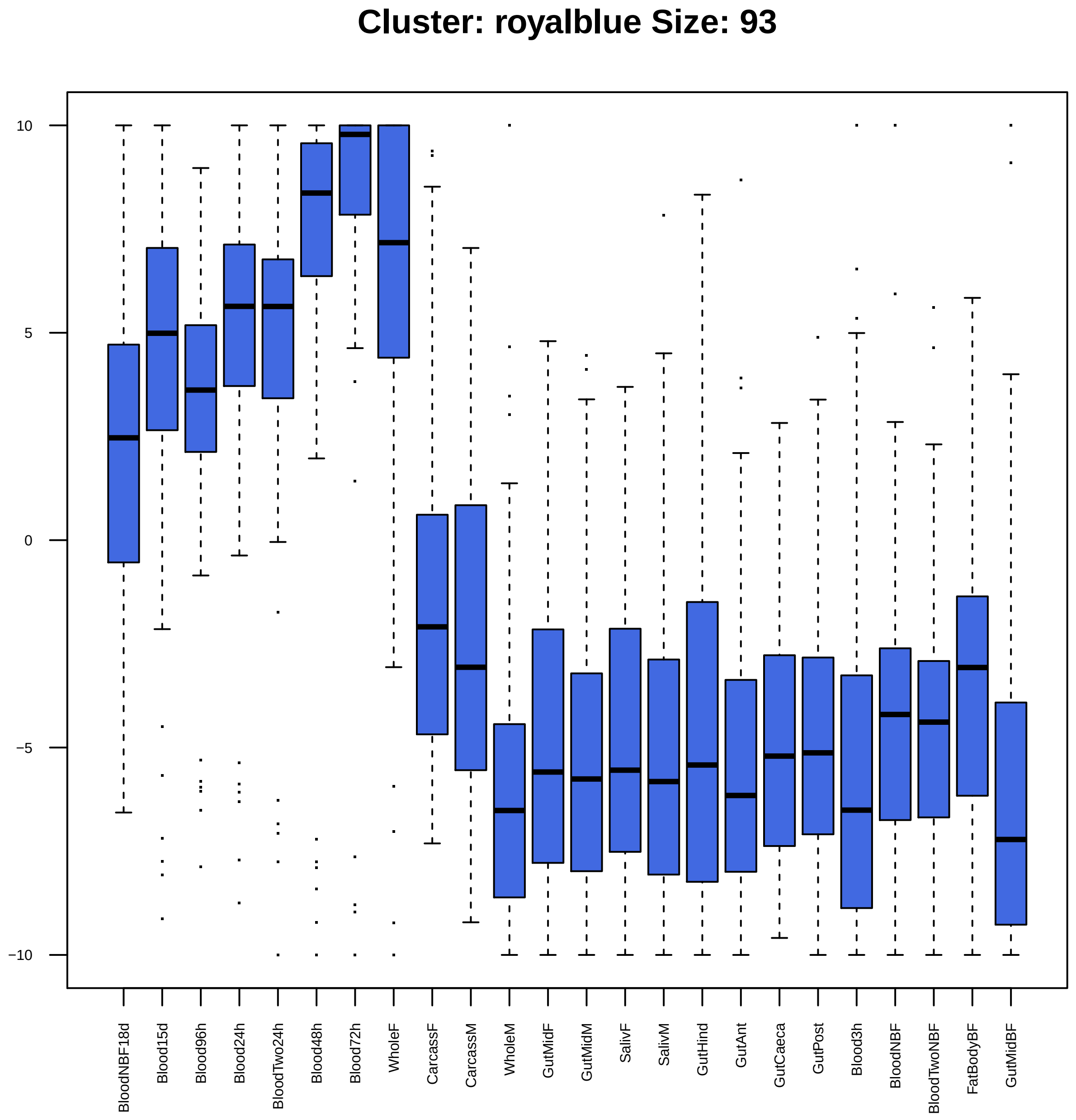
Cluster: royalblue Size: 93
10
5
0
−5
−10
BloodNBF18d
Blood15d
Blood96h
Blood24h
BloodTwo24h
Blood48h
Blood72h
WholeF
CarcassF
CarcassM
WholeM
GutMidF
GutMidM
SalivF
SalivM
GutHind
GutAnt
GutCaeca
GutPost
Blood3h
BloodNBF
BloodTwoNBF
FatBodyBF
GutMidBF

| | GO.ID | BPCluster: royalblue Size: 93 | Annotated | Significant | Expected | Rank in ClassicF | Weight01F | ClassicF |
|---|---|---|---|---|---|---|---|---|
| *1* | GO:0007447 | imaginal disc pattern formation | 47 | 3 | 0.36 | 17 | 0.00083 | 0.00545 |
| *2* | GO:0006606 | protein import into nucleus | 26 | 3 | 0.20 | 4 | 0.00098 | 0.00098 |
| *3* | GO:0033301 | cell cycle comprising mitosis without cy... | 11 | 2 | 0.08 | 12 | 0.00304 | 0.00304 |
| *4* | GO:0007099 | centriole replication | 15 | 2 | 0.12 | 18 | 0.00570 | 0.00570 |
| *5* | GO:0016318 | ommatidial rotation | 16 | 2 | 0.12 | 20 | 0.00648 | 0.00648 |
| *6* | GO:0006334 | nucleosome assembly | 17 | 2 | 0.13 | 22 | 0.00731 | 0.00731 |
| *7* | GO:0045448 | mitotic cell cycle, embryonic | 17 | 2 | 0.13 | 23 | 0.00731 | 0.00731 |
| *8* | GO:0007611 | learning or memory | 57 | 3 | 0.44 | 28 | 0.00744 | 0.00933 |
| *9* | GO:2000113 | negative regulation of cellular macromol... | 129 | 3 | 0.99 | 158 | 0.00752 | 0.07608 |
| *10* | GO:0008283 | cell proliferation | 153 | 4 | 1.18 | 70 | 0.00792 | 0.02928 |
| *11* | GO:0042059 | negative regulation of epidermal growth ... | 19 | 2 | 0.15 | 26 | 0.00910 | 0.00910 |

| | GO.ID | MFCluster: royalblue Size: 93 | Annotated | Significant | Expected | Rank in ClassicF | Weight01F | ClassicF |
|---|---|---|---|---|---|---|---|---|
| 2 | GO:0008536 | Ran GTPase binding | 16 | 2 | 0.15 | 11 | 0.0097 | 0.00966 |
| 3 | GO:0005515 | protein binding | 2143 | 33 | 20.21 | 7 | 0.0130 | 0.00080 |
| 5 | GO:0003676 | nucleic acid binding | 1152 | 21 | 10.87 | 8 | 0.0216 | 0.00154 |
| 19 | GO:0008565 | protein transporter activity | 28 | 4 | 0.26 | 4 | 0.1508 | 0.00013 |

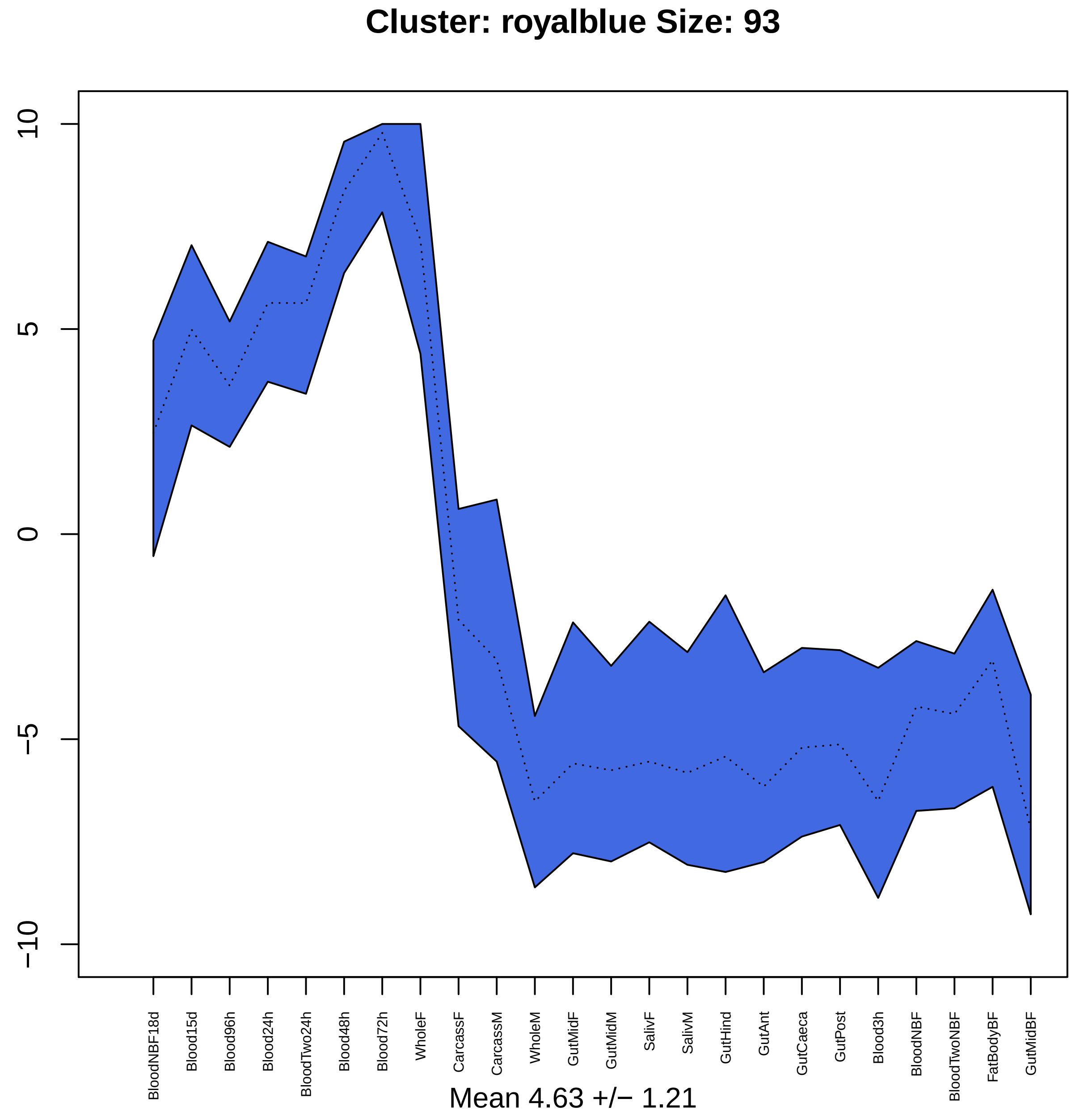

Cluster: royalblue Size: 93
10
5
0
−5
−10
BloodNBF18d
Blood15d
Blood96h
Blood24h
BloodTwo24h
Blood48h
Blood72h
WholeF
CarcassF
CarcassM
WholeM
GutMidF
GutMidM
SalivF
SalivM
GutHind
GutAnt
GutCaeca
GutPost
Blood3h
BloodNBF
BloodTwoNBF
FatBodyBF
GutMidBF
Mean 4.63 +/− 1.21

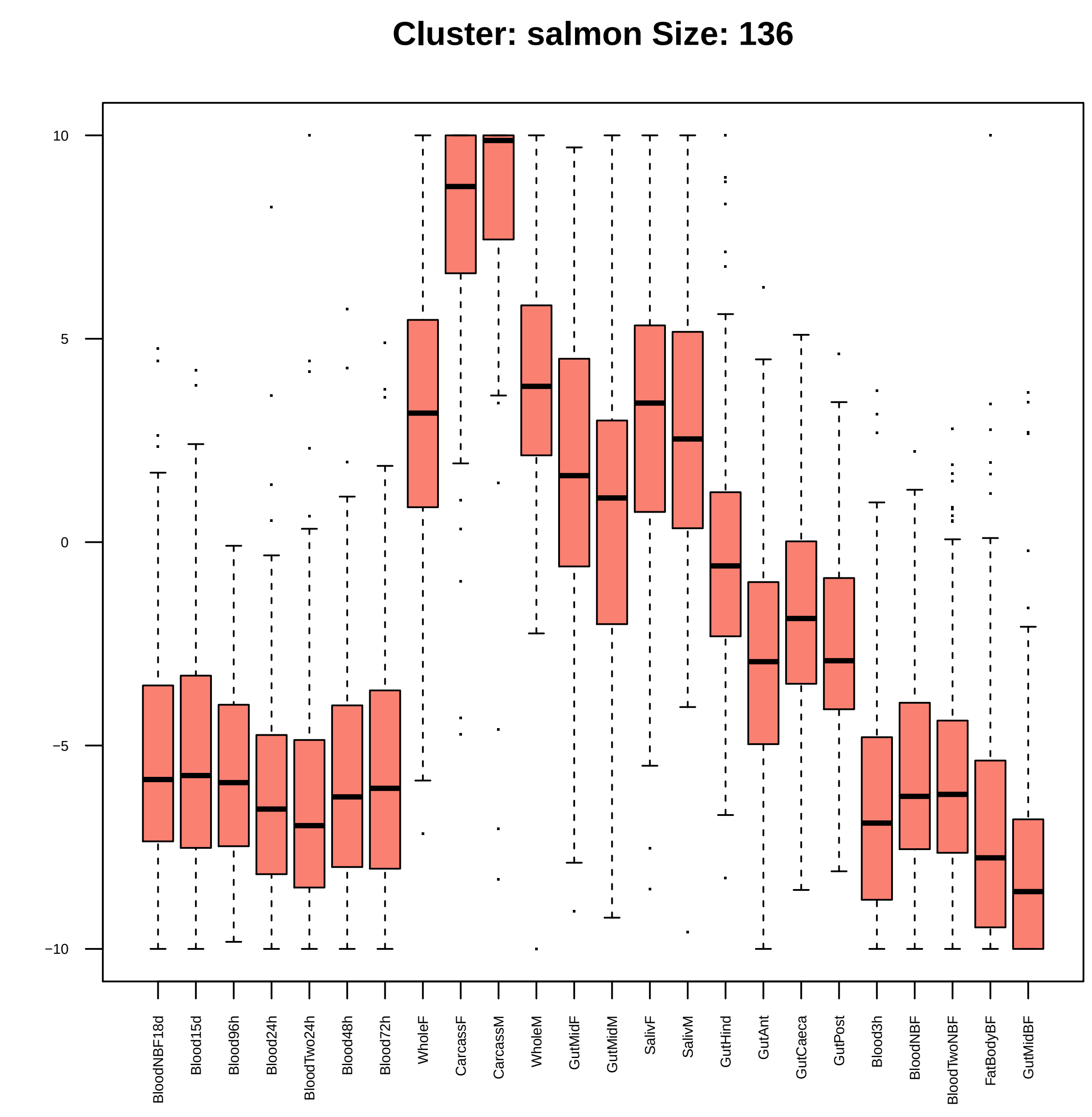

Cluster: salmon Size: 136
10
5
0
−5
−10
BloodNBF18d
Blood15d
Blood96h
Blood24h
BloodTwo24h
Blood48h
Blood72h
WholeF
CarcassF
CarcassM
WholeM
GutMidF
GutMidM
SalivF
SalivM
GutHind
GutAnt
GutCaeca
GutPost
Blood3h
BloodNBF
BloodTwoNBF
FatBodyBF
GutMidBF

| | GO.ID | BPCluster: salmon Size: 136 | Annotated | Significant | Expected | Rank in ClassicF | Weight01F | ClassicF |
|---|---|---|---|---|---|---|---|---|
| *3* | GO:0009108 | coenzyme biosynthetic process | 60 | 6 | 0.88 | 11 | 0.00033 | 0.00023 |
| *4* | GO:0006412 | translation | 289 | 19 | 4.26 | 1 | 0.00066 | 2.7e−08 |
| *5* | GO:0006414 | translational elongation | 22 | 3 | 0.32 | 23 | 0.00390 | 0.00390 |
| *6* | GO:0006733 | oxidoreduction coenzyme metabolic proces... | 34 | 3 | 0.50 | 32 | 0.00425 | 0.01334 |
| *14* | GO:0032787 | monocarboxylic acid metabolic process | 88 | 5 | 1.30 | 27 | 0.05600 | 0.00936 |
| *16* | GO:0006732 | coenzyme metabolic process | 77 | 7 | 1.14 | 10 | 0.06789 | 0.00013 |
| *17* | GO:0009058 | biosynthetic process | 1357 | 34 | 20.00 | 16 | 0.06801 | 0.00060 |

| | GO.ID | MFCluster: salmon Size: 136 | Annotated | Significant | Expected | Rank in ClassicF | Weight01F | ClassicF |
|---|---|---|---|---|---|---|---|---|
| *2* | GO:0003746 | translation elongation factor activity | 14 | 3 | 0.18 | 3 | 0.00066 | 0.00066 |
| *3* | GO:0000287 | magnesium ion binding | 34 | 4 | 0.43 | 5 | 0.00086 | 0.00086 |
| *4* | GO:0016860 | intramolecular oxidoreductase activity | 23 | 3 | 0.29 | 7 | 0.00296 | 0.00296 |
| *5* | GO:0000049 | tRNA binding | 10 | 2 | 0.13 | 9 | 0.00678 | 0.00678 |
| *6* | GO:0016878 | acid−thiol ligase activity | 11 | 2 | 0.14 | 11 | 0.00822 | 0.00822 |
| *7* | GO:0016779 | nucleotidyltransferase activity | 61 | 4 | 0.78 | 10 | 0.00887 | 0.00745 |
| *8* | GO:0016645 | oxidoreductase activity, acting on the C... | 12 | 2 | 0.15 | 12 | 0.00978 | 0.00978 |
| *17* | GO:0016877 | ligase activity, forming carbon−sulfur b... | 17 | 3 | 0.22 | 6 | 0.07274 | 0.00120 |

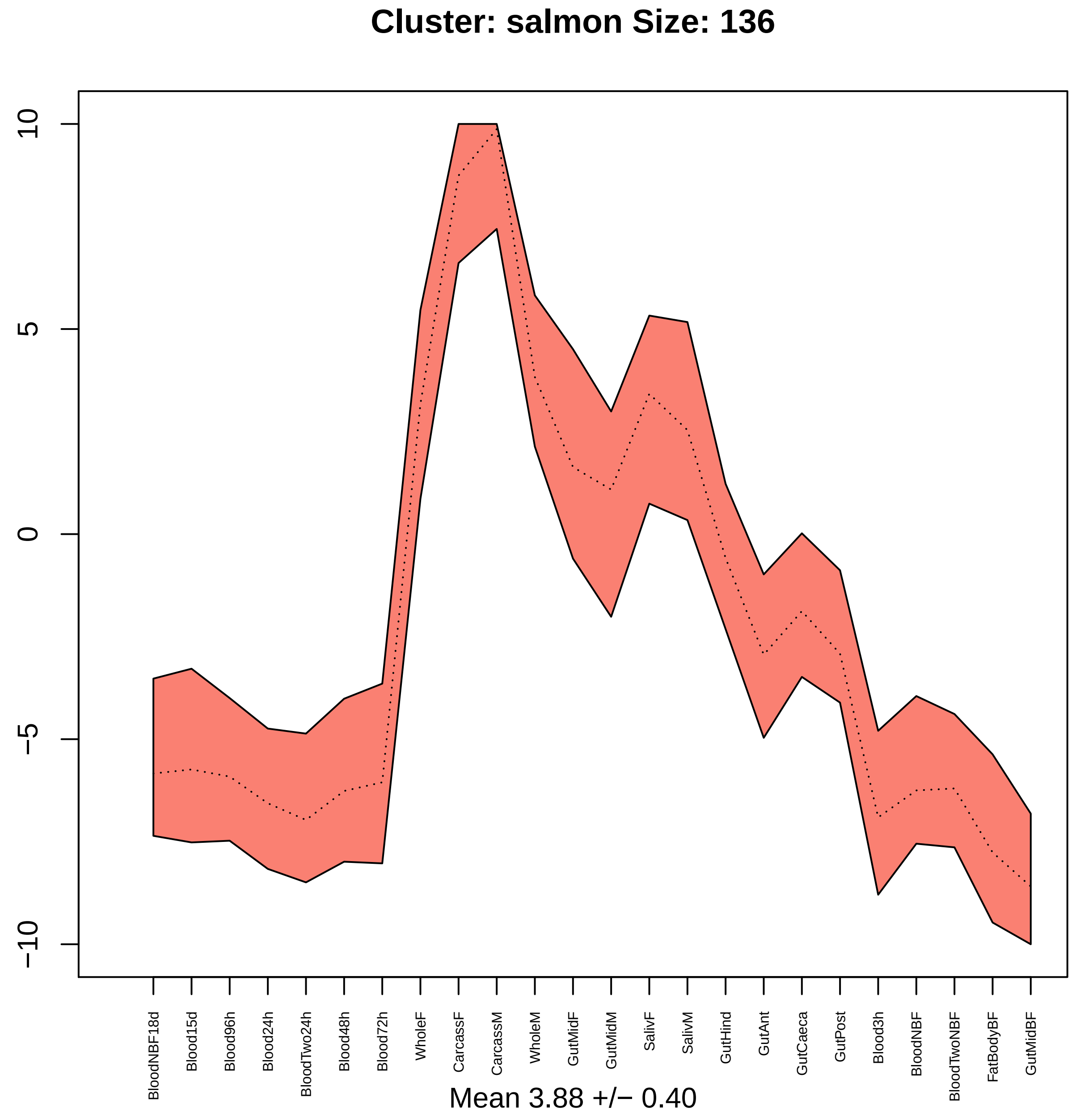

Cluster: salmon Size: 136
10
5
0
−5
−10
BloodNBF18d
Blood15d
Blood96h
Blood24h
BloodTwo24h
Blood48h
Blood72h
WholeF
CarcassF
CarcassM
WholeM
GutMidF
GutMidM
SalivF
SalivM
GutHind
GutAnt
GutCaeca
GutPost
Blood3h
BloodNBF
BloodTwoNBF
FatBodyBF
GutMidBF
Mean 3.88 +/− 0.40

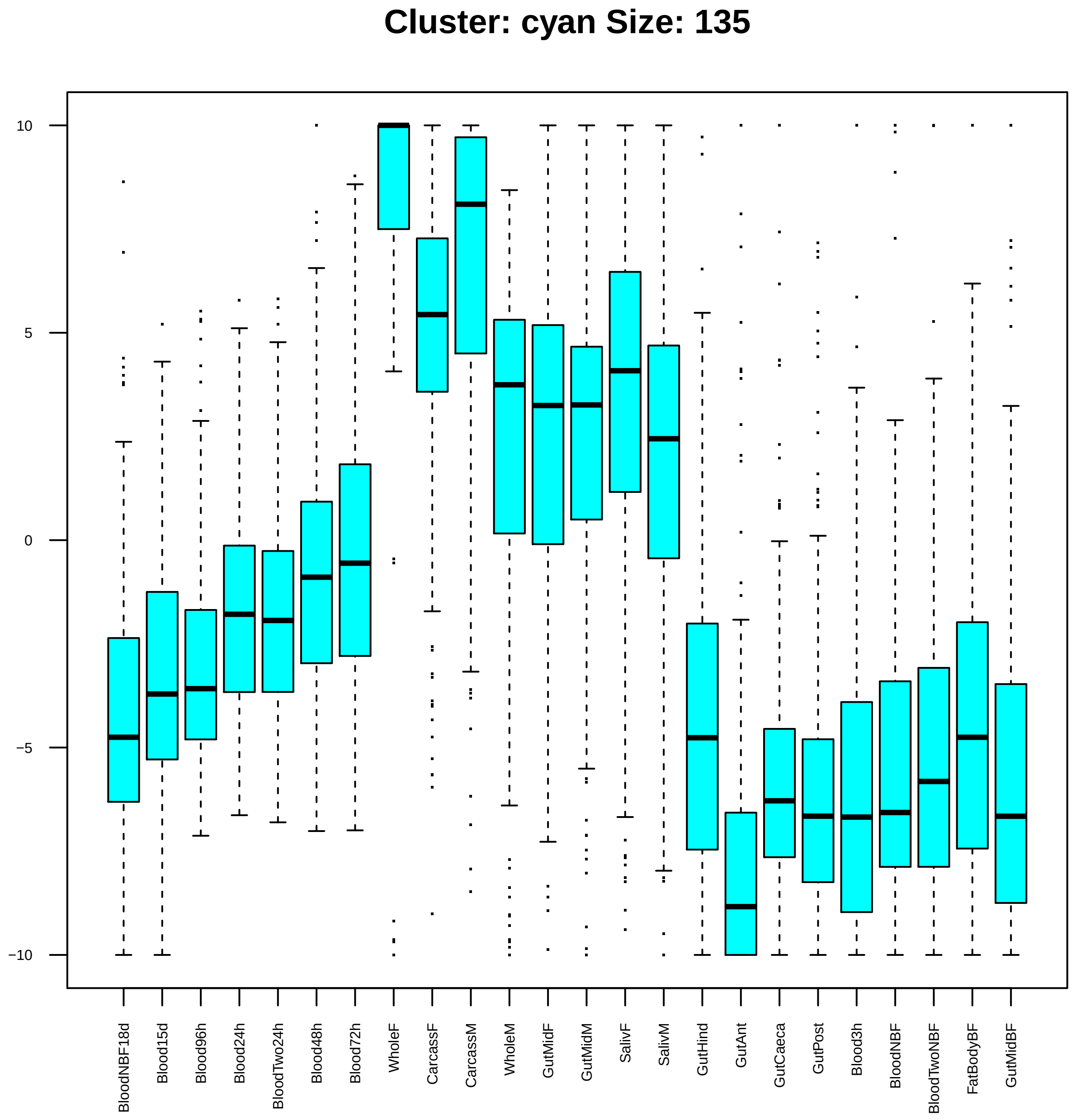
Cluster: cyan Size: 135
10
5
0
−5
−10
BloodNBF18d
Blood15d
Blood96h
Blood24h
BloodTwo24h
Blood48h
Blood72h
WholeF
CarcassF
CarcassM
WholeM
GutMidF
GutMidM
SalivF
SalivM
GutHind
GutAnt
GutCaeca
GutPost
Blood3h
BloodNBF
BloodTwoNBF
FatBodyBF
GutMidBF

| | GO.ID | BPCluster: cyan Size: 135 | Annotated | Significant | Expected | Rank in ClassicF | Weight01F | ClassicF |
|---|---|---|---|---|---|---|---|---|
| *1* | GO:0022008 | neurogenesis | 706 | 23 | 10.07 | 19 | 6.7e−06 | 0.00010 |
| *2* | GO:0000398 | mRNA splicing, via spliceosome | 134 | 11 | 1.91 | 9 | 0.00034 | 2.6e−06 |
| *3* | GO:0034472 | snRNA 3'−end processing | 11 | 3 | 0.16 | 32 | 0.00043 | 0.00043 |
| *4* | GO:0000381 | regulation of alternative mRNA splicing,... | 32 | 4 | 0.46 | 42 | 0.00103 | 0.00103 |
| *5* | GO:0016246 | RNA interference | 19 | 3 | 0.27 | 57 | 0.00231 | 0.00231 |
| *6* | GO:0007517 | muscle organ development | 66 | 5 | 0.94 | 58 | 0.00239 | 0.00239 |
| *7* | GO:0007095 | mitotic G2 DNA damage checkpoint | 43 | 4 | 0.61 | 62 | 0.00313 | 0.00313 |
| *8* | GO:0000956 | nuclear−transcribed mRNA catabolic proce... | 20 | 4 | 0.29 | 20 | 0.00808 | 0.00016 |
| *9* | GO:0019751 | polyol metabolic process | 10 | 2 | 0.14 | 91 | 0.00841 | 0.00841 |
| *10* | GO:0045815 | positive regulation of gene expression, ... | 10 | 2 | 0.14 | 92 | 0.00841 | 0.00841 |
| *11* | GO:0000288 | nuclear−transcribed mRNA catabolic proce... | 10 | 2 | 0.14 | 93 | 0.00841 | 0.00841 |
| *13* | GO:0000075 | cell cycle checkpoint | 64 | 6 | 0.91 | 27 | 0.01362 | 0.00028 |
| *22* | GO:1901991 | negative regulation of mitotic cell cycl... | 60 | 6 | 0.86 | 23 | 0.01976 | 0.00019 |

| | GO.ID | MFCluster: cyan Size: 135 | Annotated | Significant | Expected | Rank in ClassicF | Weight01F | ClassicF |
|---|---|---|---|---|---|---|---|---|
| *1* | GO:0004004 | ATP−dependent RNA helicase activity | 28 | 3 | 0.40 | 4 | 0.0074 | 0.0074 |
| *12* | GO:0003676 | nucleic acid binding | 1152 | 28 | 16.62 | 1 | 0.0720 | 0.0029 |
| *13* | GO:0000166 | nucleotide binding | 998 | 24 | 14.40 | 2 | 0.0758 | 0.0071 |

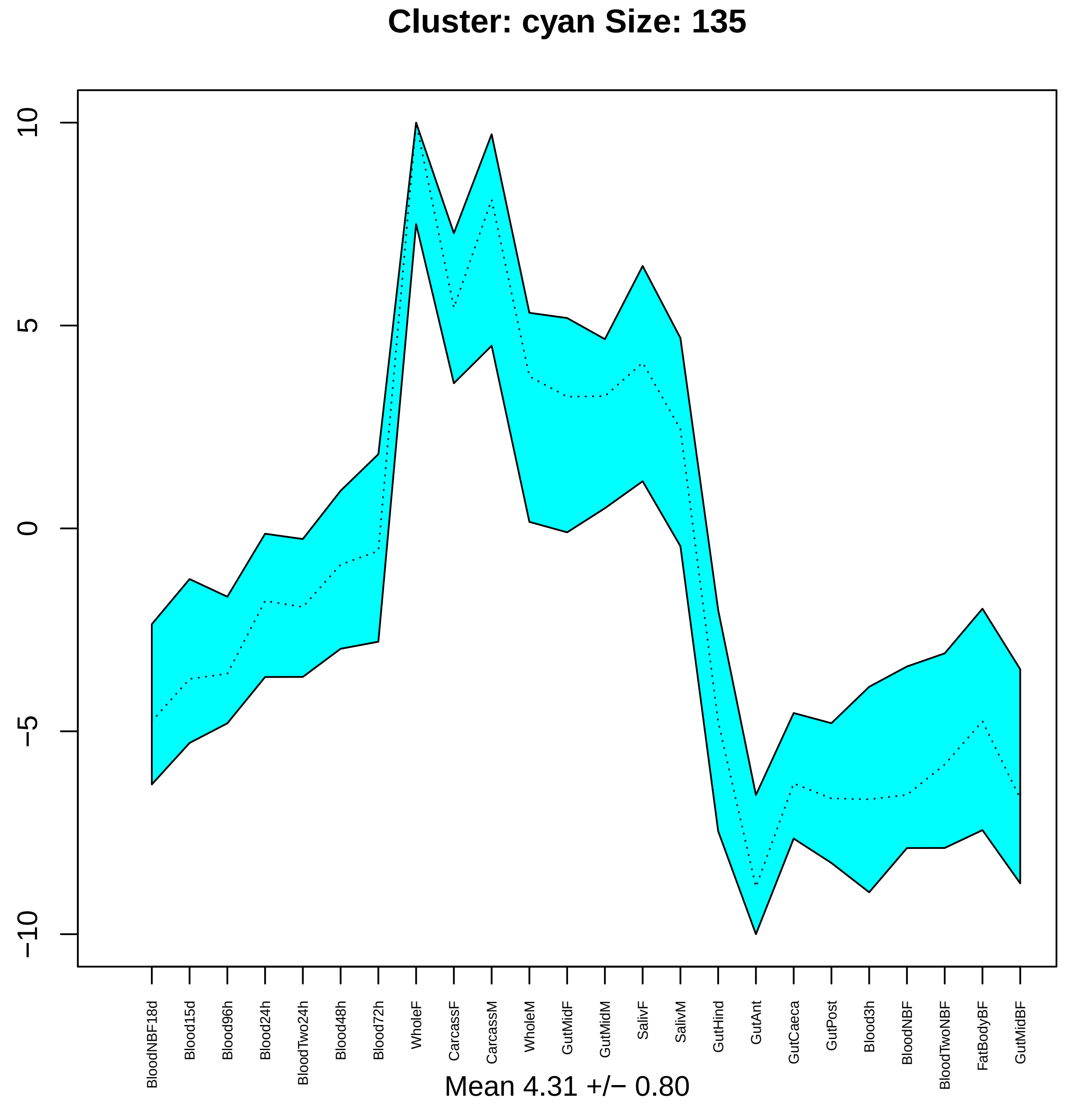

Cluster: cyan Size: 135
10
5
0
−5
−10
BloodNBF18d
Blood15d
Blood96h
Blood24h
BloodTwo24h
Blood48h
Blood72h
WholeF
CarcassF
CarcassM
WholeM
GutMidF
GutMidM
SalivF
SalivM
GutHind
GutAnt
GutCaeca
GutPost
Blood3h
BloodNBF
BloodTwoNBF
FatBodyBF
GutMidBF
Mean 4.31 +/− 0.80

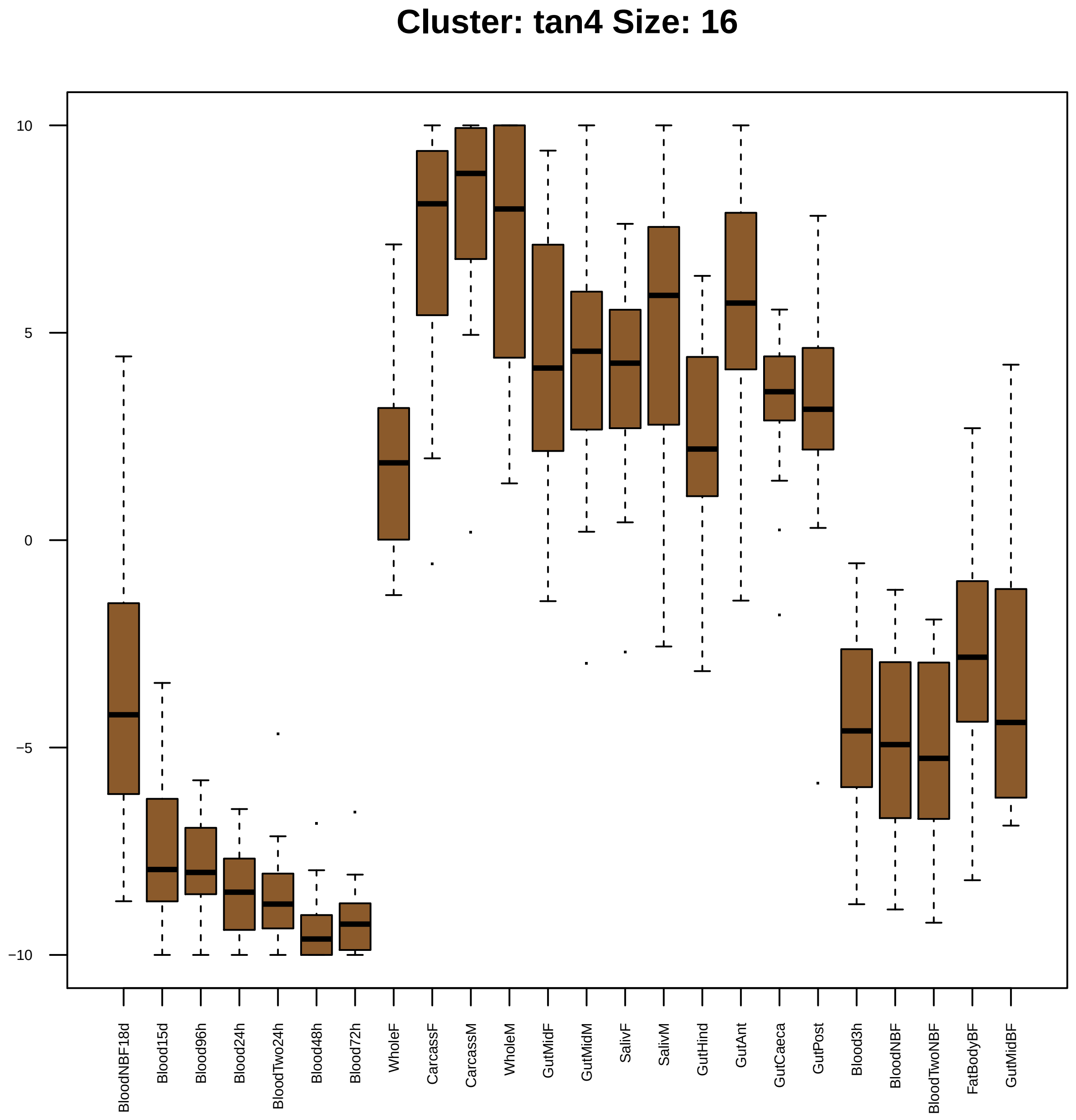
Cluster: tan4 Size: 16
10
5
0
−5
−10
BloodNBF18d
Blood15d
Blood96h
Blood24h
BloodTwo24h
Blood48h
Blood72h
WholeF
CarcassF
CarcassM
WholeM
GutMidF
GutMidM
SalivF
SalivM
GutHind
GutAnt
GutCaeca
GutPost
Blood3h
BloodNBF
BloodTwoNBF
FatBodyBF
GutMidBF

| | GO.ID | BPCluster: tan4 Size: 16 | Annotated | Significant | Expected | Rank in ClassicF | Weight01F | ClassicF |
|---|---|---|---|---|---|---|---|---|
| *10* | GO:0009416 | response to light stimulus | 65 | 2 | 0.08 | 2 | 0.057 | 0.0028 |

| | GO.ID | MFCluster: tan4 Size: 16 | Annotated | Significant | Expected | Rank in ClassicF | Weight01F | ClassicF |
|---|---|---|---|---|---|---|---|---|
| *1* | GO:0046961 | proton−transporting ATPase activity, rot... | 21 | 2 | 0.03 | 1 | 0.00029 | 0.00029 |
| *23* | GO:0043492 | ATPase activity, coupled to movement of ... | 73 | 2 | 0.09 | 8 | 1.00000 | 0.00348 |

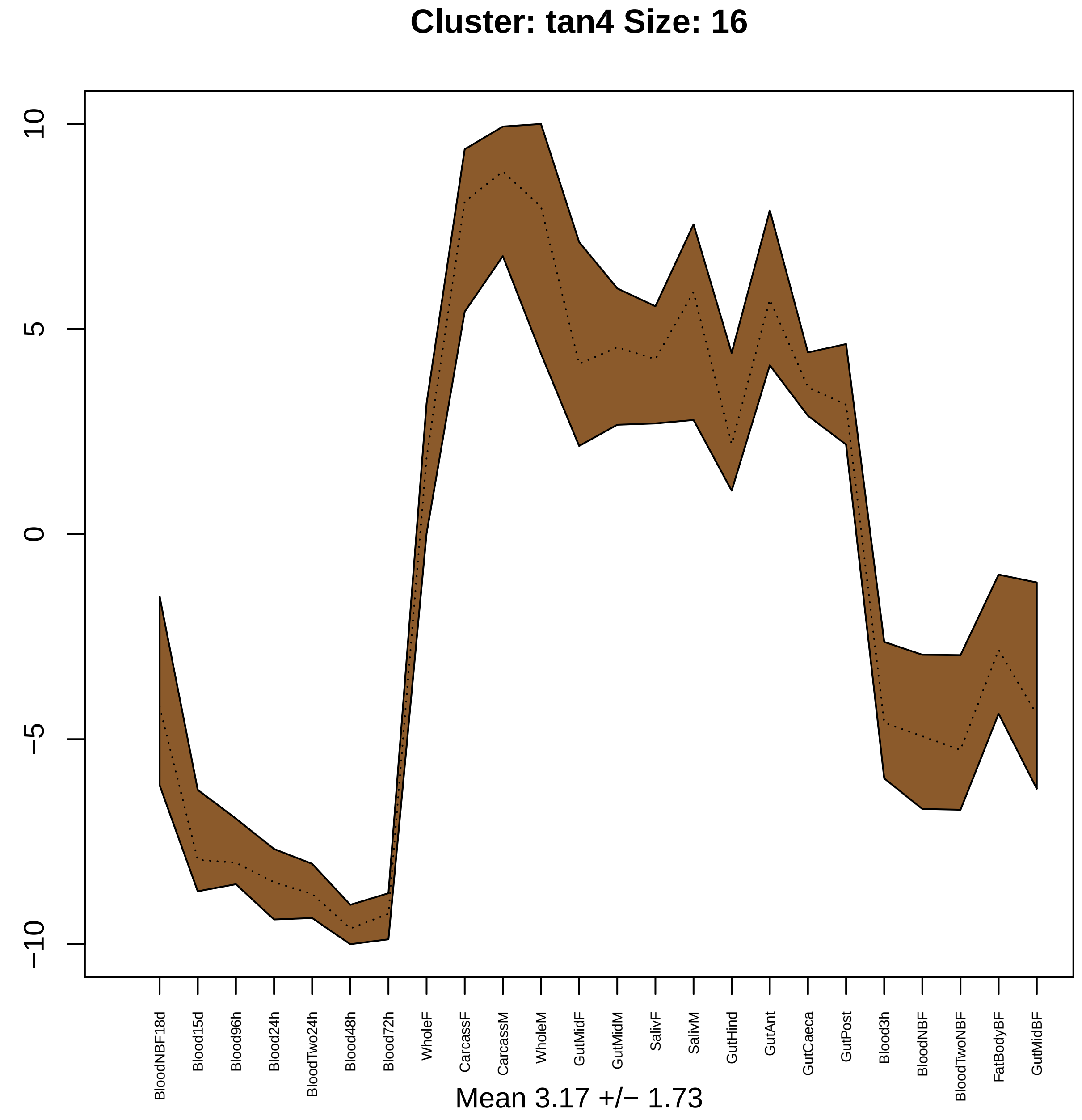
Cluster: tan4 Size: 16
10
5
0
−5
−10
BloodNBF18d
Blood15d
Blood96h
Blood24h
BloodTwo24h
Blood48h
Blood72h
WholeF
CarcassF
CarcassM
WholeM
GutMidF
GutMidM
SalivF
SalivM
GutHind
GutAnt
GutCaeca
GutPost
Blood3h
BloodNBF
BloodTwoNBF
FatBodyBF
GutMidBF
Mean 3.17 +/− 1.73

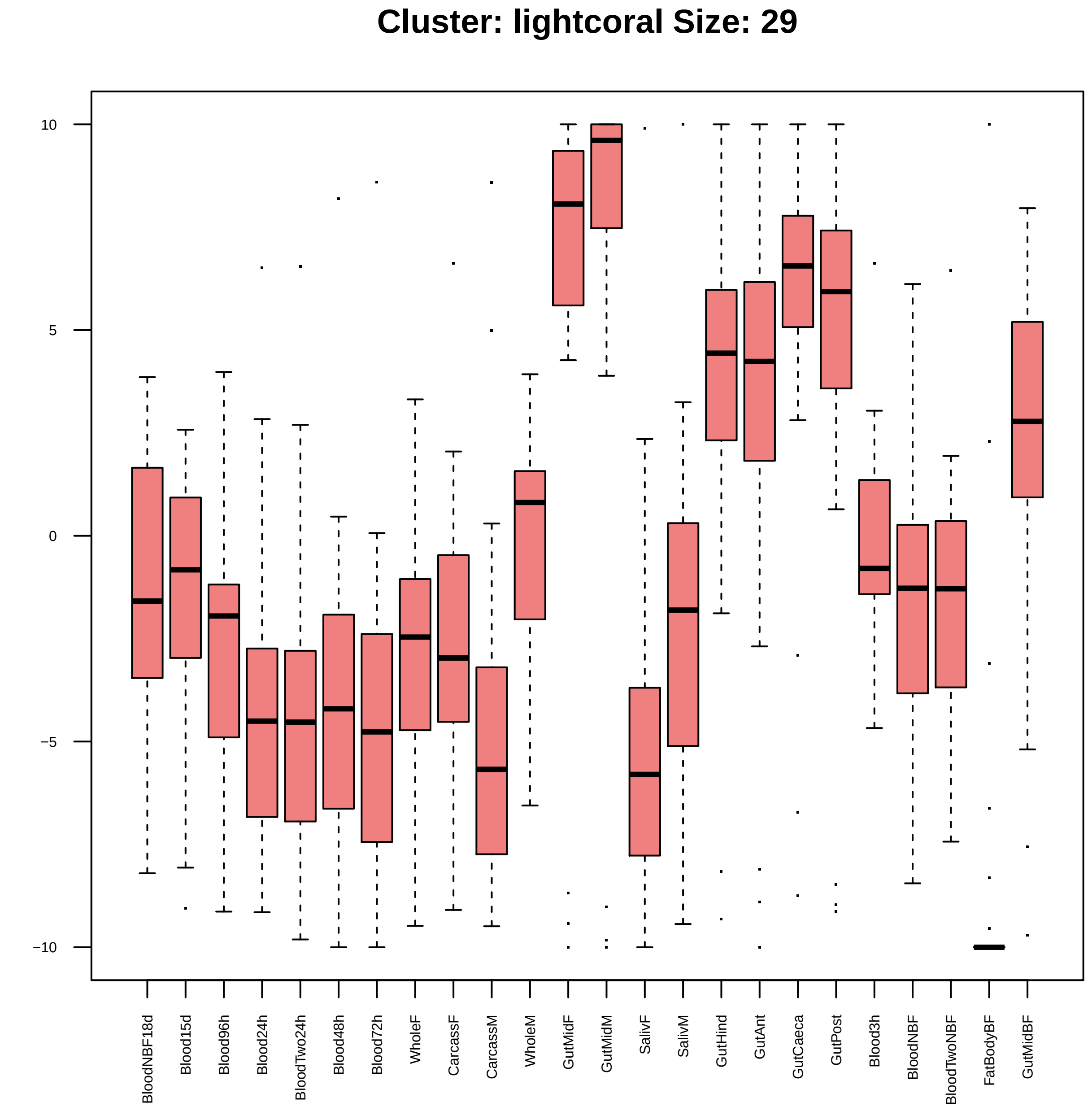

Cluster: lightcoral Size: 29
10
5
0
−5
−10
BloodNBF18d
Blood15d
Blood96h
Blood24h
BloodTwo24h
Blood48h
Blood72h
WholeF
CarcassF
CarcassM
WholeM
GutMidF
GutMidM
SalivF
SalivM
GutHind
GutAnt
GutCaeca
GutPost
Blood3h
BloodNBF
BloodTwoNBF
FatBodyBF
GutMidBF

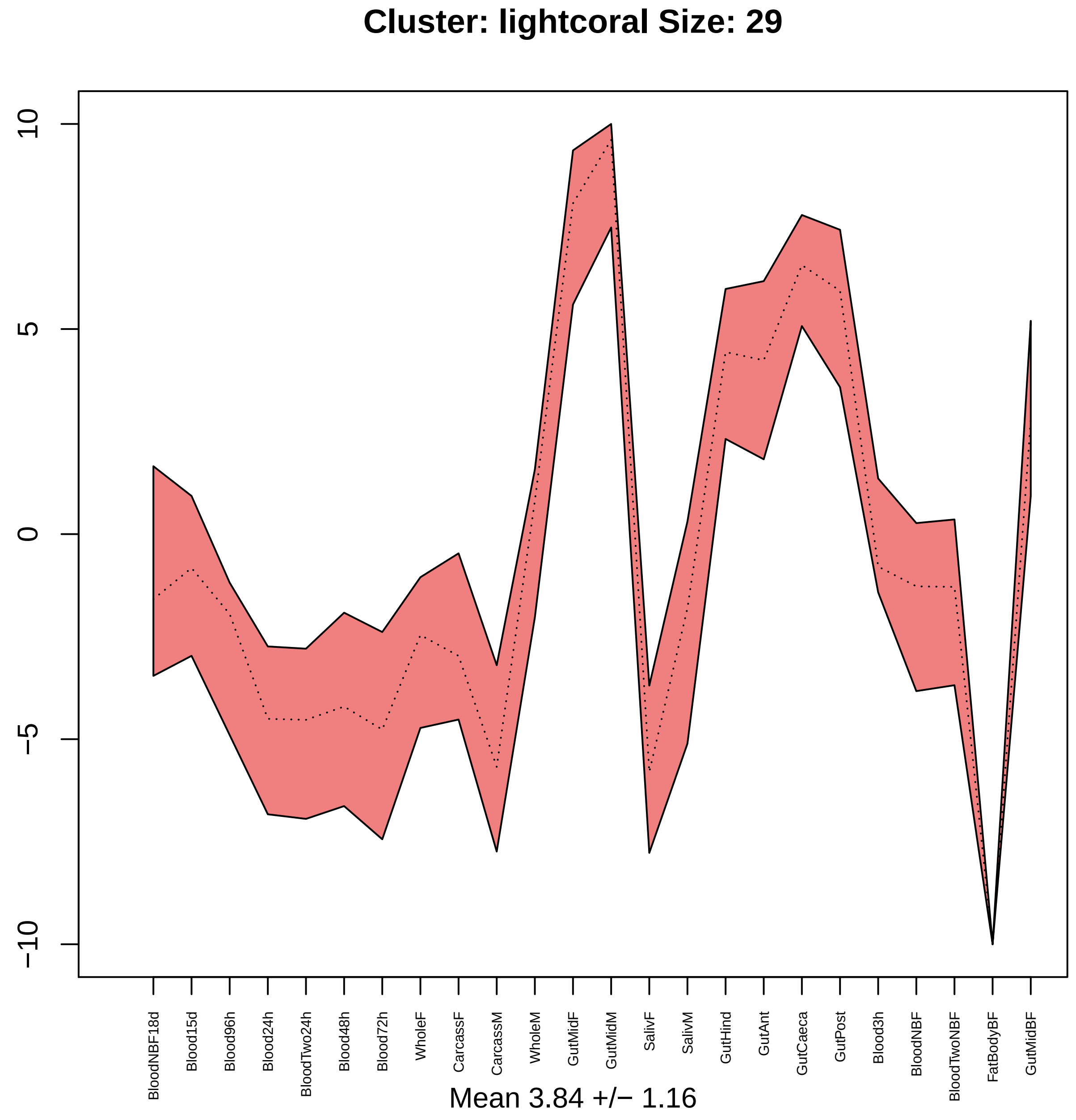

Cluster: lightcoral Size: 29
10
5
0
−5
−10
BloodNBF18d
Blood15d
Blood96h
Blood24h
BloodTwo24h
Blood48h
Blood72h
WholeF
CarcassF
CarcassM
WholeM
GutMidF
GutMidM
SalivF
SalivM
GutHind
GutAnt
GutCaeca
GutPost
Blood3h
BloodNBF
BloodTwoNBF
FatBodyBF
GutMidBF
Mean 3.84 +/− 1.16

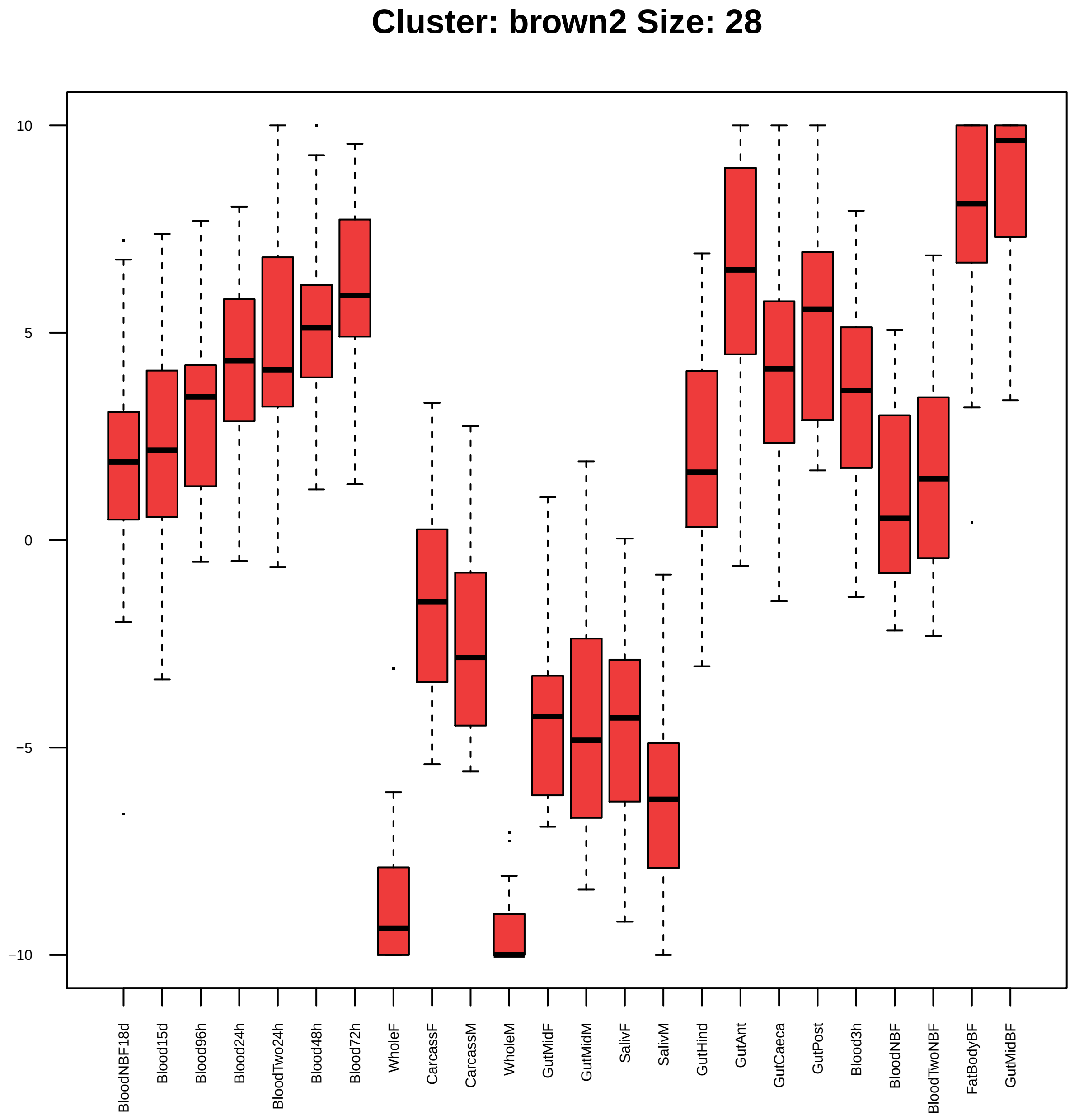
Cluster: brown2 Size: 28
10
5
0
−5
−10
BloodNBF18d
Blood15d
Blood96h
Blood24h
BloodTwo24h
Blood48h
Blood72h
WholeF
CarcassF
CarcassM
WholeM
GutMidF
GutMidM
SalivF
SalivM
GutHind
GutAnt
GutCaeca
GutPost
Blood3h
BloodNBF
BloodTwoNBF
FatBodyBF
GutMidBF

| | GO.ID | BPCluster: brown2 Size: 28 | Annotated | Significant | Expected | Rank in ClassicF | Weight01F | ClassicF |
|---|---|---|---|---|---|---|---|---|
| *1* | GO:0030177 | positive regulation of Wnt signaling pat... | 11 | 2 | 0.02 | 1 | 0.00018 | 0.00018 |
| *2* | GO:0006367 | transcription initiation from RNA polyme... | 39 | 2 | 0.07 | 3 | 0.00231 | 0.00231 |
| *3* | GO:0048569 | post−embryonic animal organ development | 222 | 3 | 0.42 | 9 | 0.01131 | 0.00725 |

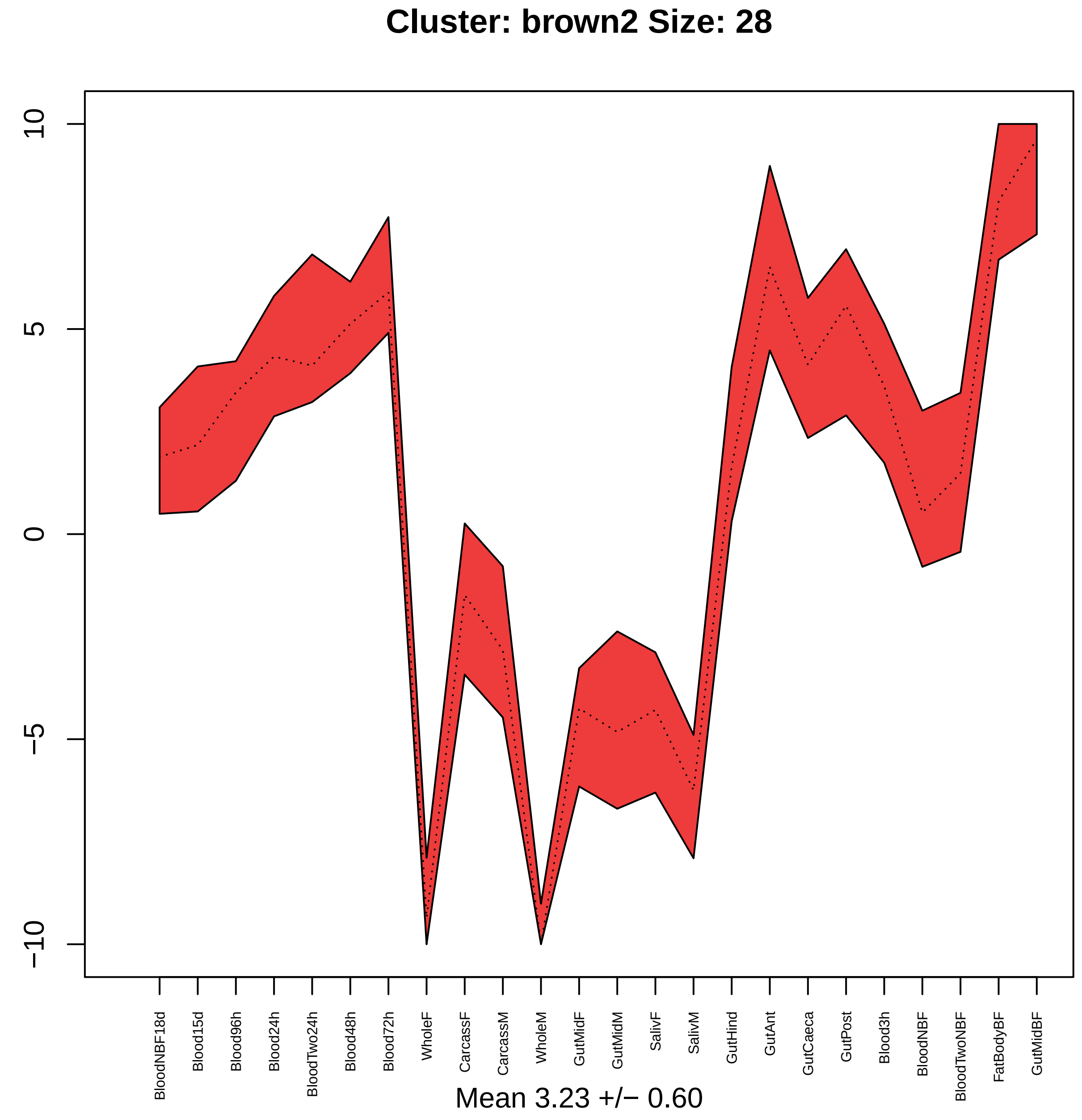
Cluster: brown2 Size: 28
10
5
0
−5
−10
BloodNBF18d
Blood15d
Blood96h
Blood24h
BloodTwo24h
Blood48h
Blood72h
WholeF
CarcassF
CarcassM
WholeM
GutMidF
GutMidM
SalivF
SalivM
GutHind
GutAnt
GutCaeca
GutPost
Blood3h
BloodNBF
BloodTwoNBF
FatBodyBF
GutMidBF
Mean 3.23 +/− 0.60

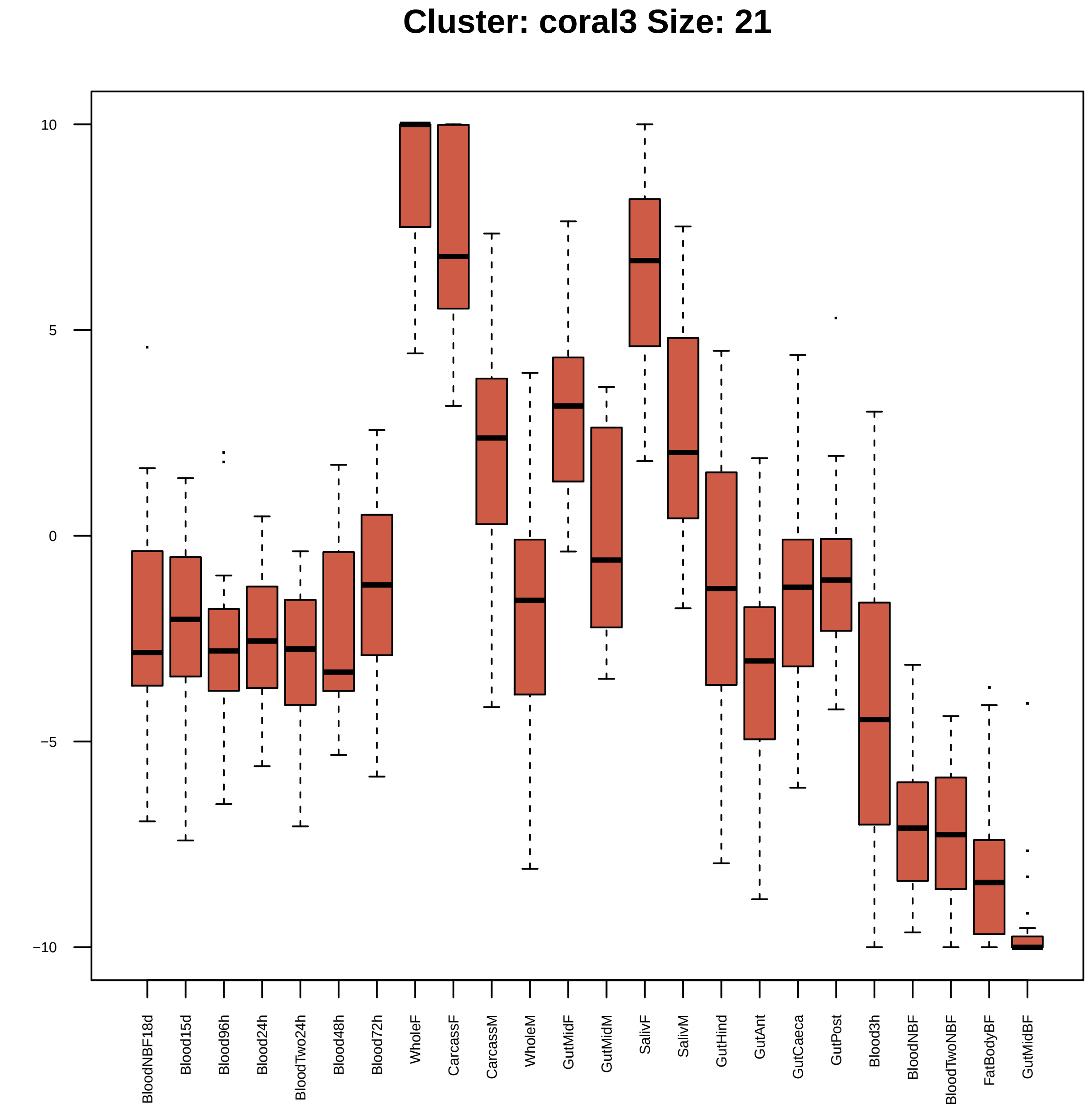
Cluster: coral3 Size: 21
10
5
0
−5
−10
BloodNBF18d
Blood15d
Blood96h
Blood24h
BloodTwo24h
Blood48h
Blood72h
WholeF
CarcassF
CarcassM
WholeM
GutMidF
GutMidM
SalivF
SalivM
GutHind
GutAnt
GutCaeca
GutPost
Blood3h
BloodNBF
BloodTwoNBF
FatBodyBF
GutMidBF

| | GO.ID | BPCluster: coral3 Size: 21 | Annotated | Significant | Expected | Rank in ClassicF | Weight01F | ClassicF |
|---|---|---|---|---|---|---|---|---|
| *1* | GO:0022008 | neurogenesis | 706 | 8 | 1.88 | 2 | 5.5e−06 | 0.00021 |
| *2* | GO:0001731 | formation of translation preinitiation c... | 13 | 2 | 0.03 | 4 | 0.00051 | 0.00051 |
| *3* | GO:0031122 | cytoplasmic microtubule organization | 16 | 2 | 0.04 | 8 | 0.00079 | 0.00079 |
| *4* | GO:0006457 | protein folding | 72 | 3 | 0.19 | 9 | 0.00084 | 0.00084 |
| *5* | GO:0006446 | regulation of translational initiation | 19 | 2 | 0.05 | 11 | 0.00111 | 0.00111 |
| *6* | GO:0044267 | cellular protein metabolic process | 1027 | 6 | 2.74 | 43 | 0.00266 | 0.04328 |
| *7* | GO:0007052 | mitotic spindle organization | 114 | 3 | 0.30 | 14 | 0.00315 | 0.00315 |
| *8* | GO:0006364 | rRNA processing | 34 | 2 | 0.09 | 16 | 0.00357 | 0.00357 |
| *12* | GO:0042254 | ribosome biogenesis | 64 | 3 | 0.17 | 6 | 0.06872 | 0.00059 |

| | GO.ID | MFCluster: coral3 Size: 21 | Annotated | Significant | Expected | Rank in ClassicF | Weight01F | ClassicF |
|---|---|---|---|---|---|---|---|---|
| *1* | GO:0051082 | unfolded protein binding | 41 | 3 | 0.10 | 1 | 0.00013 | 0.00013 |
| *2* | GO:0003743 | translation initiation factor activity | 35 | 2 | 0.09 | 2 | 0.00334 | 0.00334 |
| *3* | GO:0000166 | nucleotide binding | 998 | 7 | 2.49 | 4 | 0.01471 | 0.00755 |
| *6* | GO:0003723 | RNA binding | 307 | 4 | 0.77 | 3 | 0.04756 | 0.00614 |
| *17* | GO:0008135 | translation factor activity, RNA binding | 54 | 2 | 0.13 | 6 | 1.00000 | 0.00780 |
| *22* | GO:1901265 | nucleoside phosphate binding | 998 | 7 | 2.49 | 5 | 1.00000 | 0.00755 |

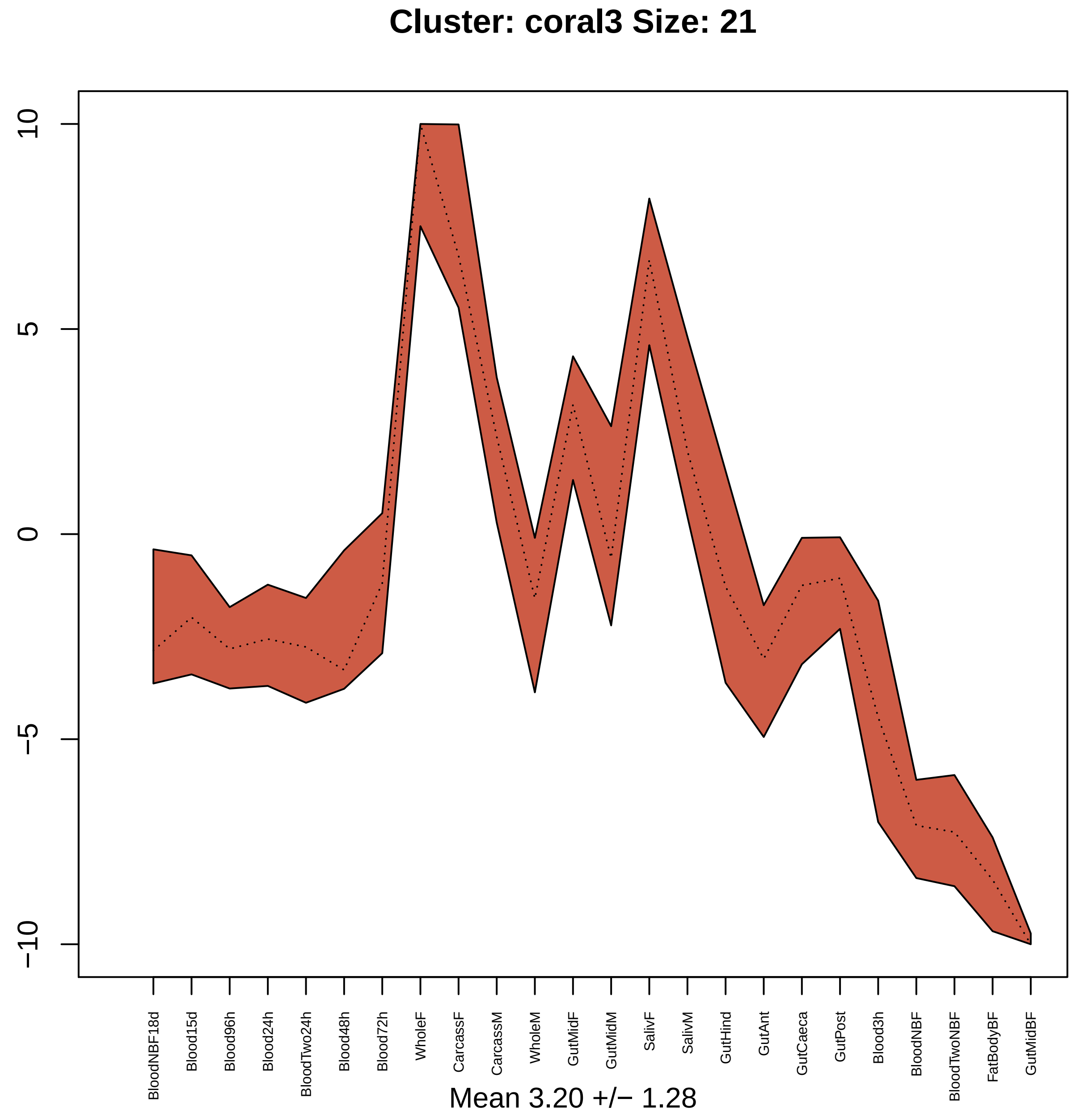

Cluster: coral3 Size: 21
10
5
0
−5
−10
BloodNBF18d
Blood15d
Blood96h
Blood24h
BloodTwo24h
Blood48h
Blood72h
WholeF
CarcassF
CarcassM
WholeM
GutMidF
GutMidM
SalivF
SalivM
GutHind
GutAnt
GutCaeca
GutPost
Blood3h
BloodNBF
BloodTwoNBF
FatBodyBF
GutMidBF
Mean 3.20 +/− 1.28

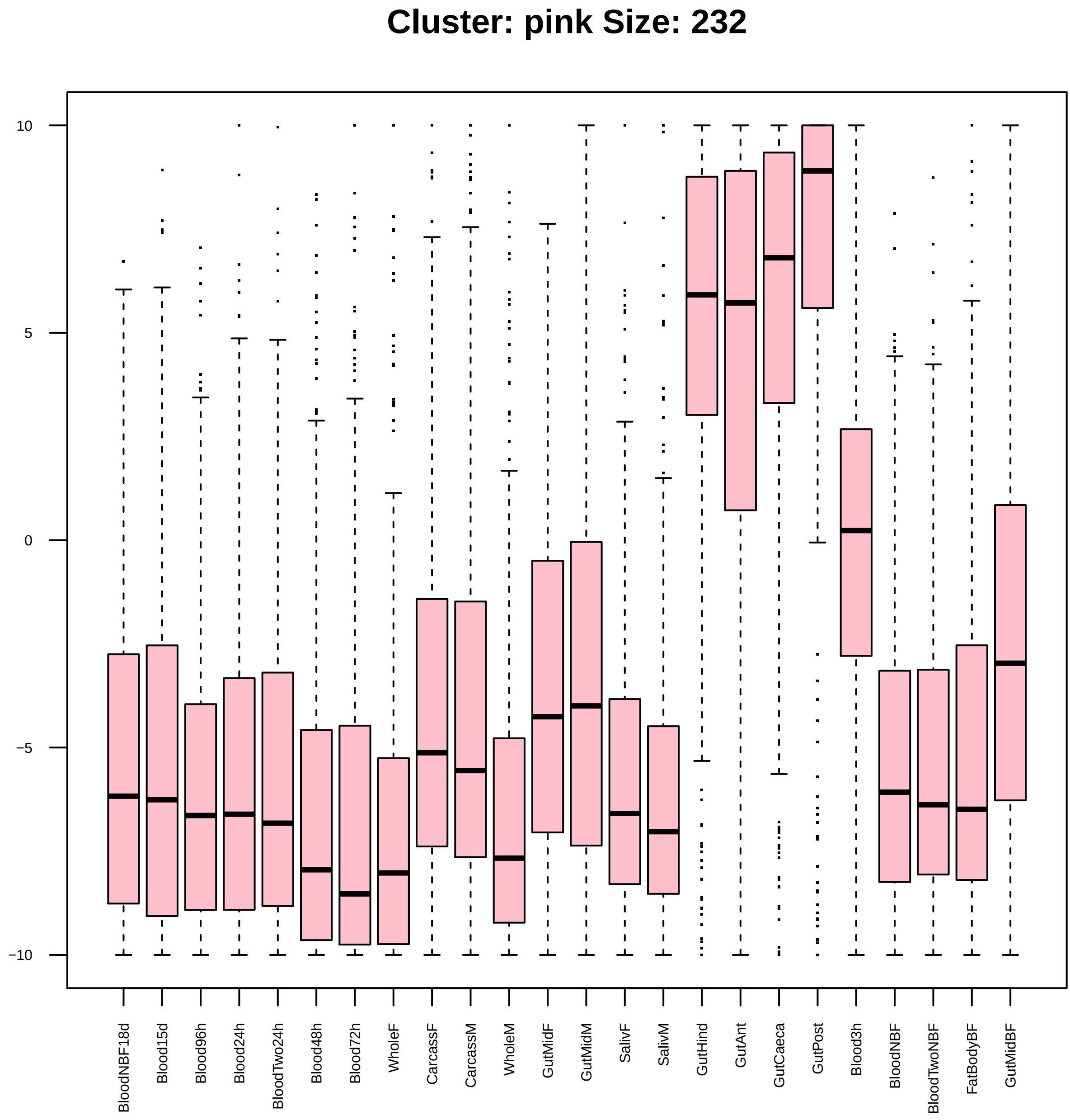

Cluster: pink Size: 232
10
5
0
−5
−10
BloodNBF18d
Blood15d
Blood96h
Blood24h
BloodTwo24h
Blood48h
Blood72h
WholeF
CarcassF
CarcassM
WholeM
GutMidF
GutMidM
SalivF
SalivM
GutHind
GutAnt
GutCaeca
GutPost
Blood3h
BloodNBF
BloodTwoNBF
FatBodyBF
GutMidBF

| | GO.ID | BPCluster: pink Size: 232 | Annotated | Significant | Expected | Rank in ClassicF | Weight01F | ClassicF |
|---|---|---|---|---|---|---|---|---|
| 3 | GO:0015991 | ATP hydrolysis coupled proton transport | 19 | 4 | 0.46 | 3 | 0.00096 | 0.00096 |
| 4 | GO:0044270 | cellular nitrogen compound catabolic pro... | 304 | 9 | 7.34 | 195 | 0.00202 | 0.31293 |
| 5 | GO:0046700 | heterocycle catabolic process | 305 | 9 | 7.37 | 197 | 0.00264 | 0.31632 |
| 6 | GO:1901361 | organic cyclic compound catabolic proces... | 308 | 9 | 7.44 | 207 | 0.00518 | 0.32652 |
| 22 | GO:0032787 | monocarboxylic acid metabolic process | 88 | 7 | 2.13 | 10 | 0.09083 | 0.00517 |

| | GO.ID | MFCluster: pink Size: 232 | Annotated | Significant | Expected | Rank in ClassicF | Weight01F | ClassicF |
|---|---|---|---|---|---|---|---|---|
| *3* | GO:0008236 | serine−type peptidase activity | 278 | 24 | 6.40 | 2 | 0.0019 | 1.7e−08 |
| *4* | GO:0032934 | sterol binding | 12 | 3 | 0.28 | 10 | 0.0023 | 0.0023 |
| *5* | GO:0003824 | catalytic activity | 3160 | 118 | 72.76 | 1 | 0.0025 | 6.4e−13 |
| *6* | GO:0016787 | hydrolase activity | 1445 | 56 | 33.27 | 8 | 0.0033 | 2.0e−05 |
| *13* | GO:0016627 | oxidoreductase activity, acting on the C... | 30 | 4 | 0.69 | 13 | 0.0503 | 0.0047 |

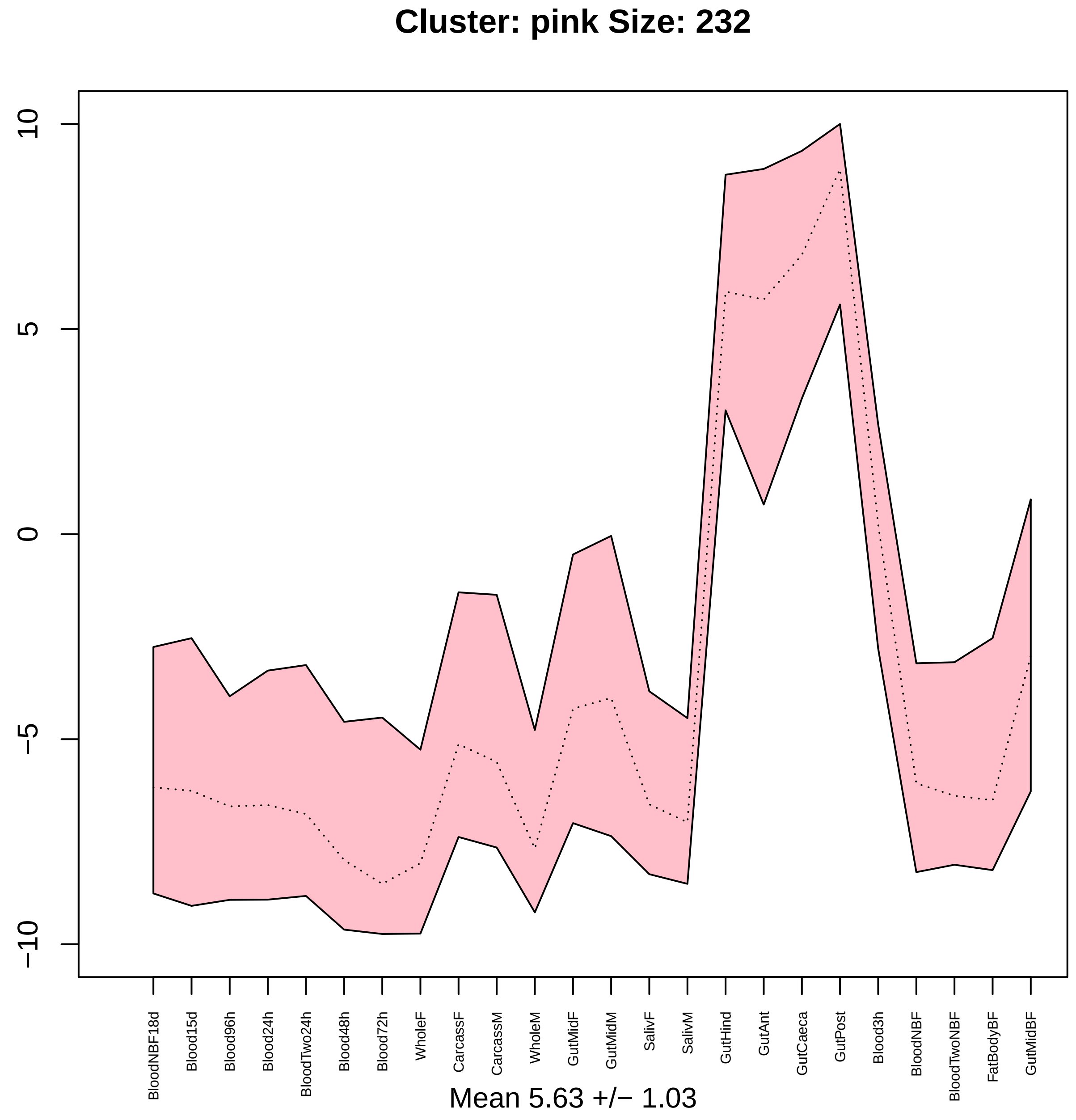

Cluster: pink Size: 232
10
5
0
−5
−10
BloodNBF18d
Blood15d
Blood96h
Blood24h
BloodTwo24h
Blood48h
Blood72h
WholeF
CarcassF
CarcassM
WholeM
GutMidF
GutMidM
SalivF
SalivM
GutHind
GutAnt
GutCaeca
GutPost
Blood3h
BloodNBF
BloodTwoNBF
FatBodyBF
GutMidBF
Mean 5.63 +/− 1.03

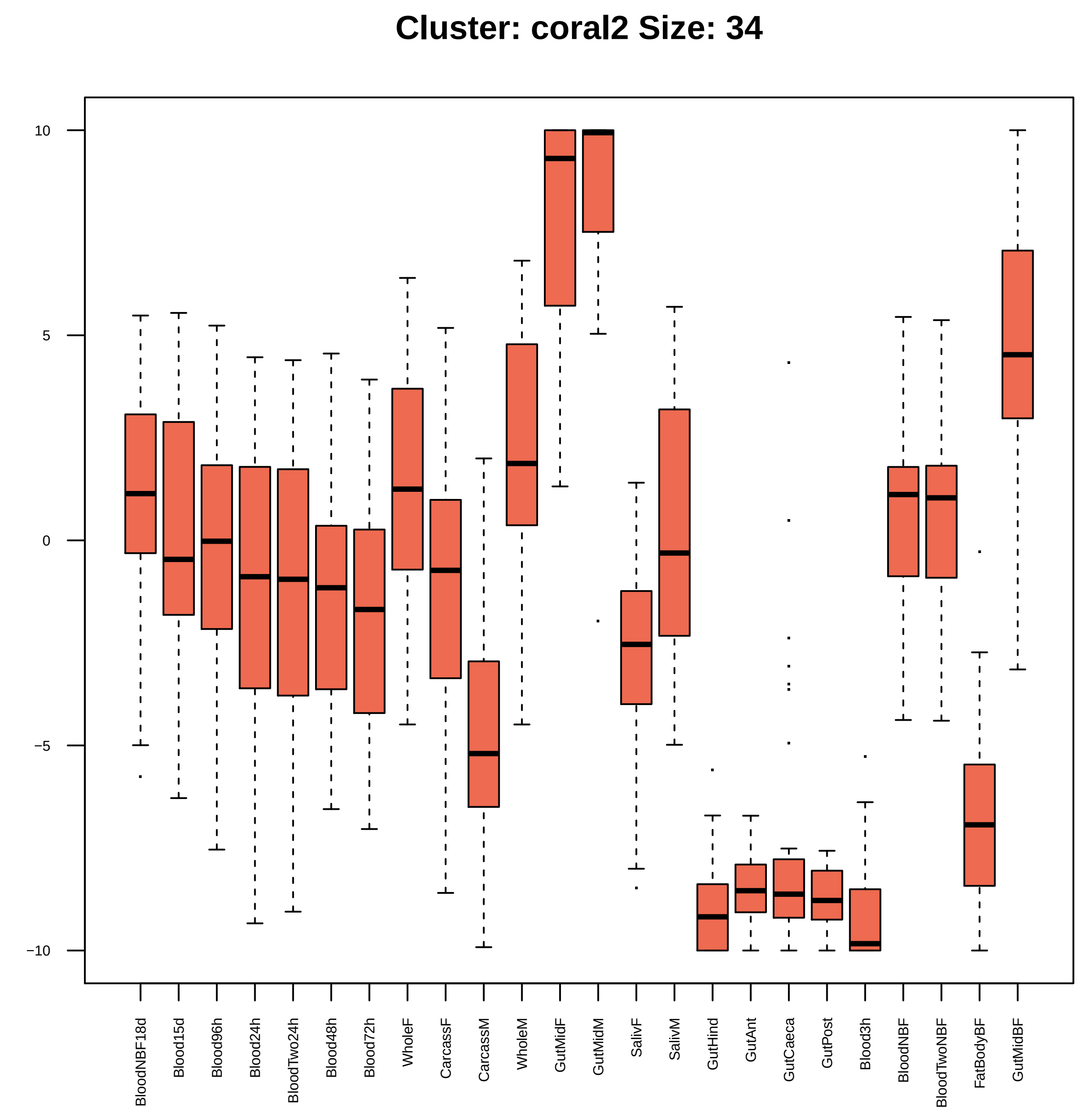
Cluster: coral2 Size: 34
10
5
0
−5
−10
BloodNBF18d
Blood15d
Blood96h
Blood24h
BloodTwo24h
Blood48h
Blood72h
WholeF
CarcassF
CarcassM
WholeM
GutMidF
GutMidM
SalivF
SalivM
GutHind
GutAnt
GutCaeca
GutPost
Blood3h
BloodNBF
BloodTwoNBF
FatBodyBF
GutMidBF

| | GO.ID | BPCluster: coral2 Size: 34 | Annotated | Significant | Expected | Rank in ClassicF | Weight01F | ClassicF |
|---|---|---|---|---|---|---|---|---|
| 3 | GO:0006508 | proteolysis | 594 | 8 | 2.05 | 6 | 0.00053 | 0.00053 |
| 4 | GO:0005975 | carbohydrate metabolic process | 210 | 5 | 0.72 | 7 | 0.00061 | 0.00061 |
| 5 | GO:0045087 | innate immune response | 60 | 3 | 0.21 | 8 | 0.00107 | 0.00107 |
| 6 | GO:0042742 | defense response to bacterium | 61 | 3 | 0.21 | 9 | 0.00113 | 0.00113 |
| 15 | GO:1901564 | organonitrogen compound metabolic proces... | 1889 | 13 | 6.52 | 15 | 1.00000 | 0.00377 |
| 16 | GO:1901135 | carbohydrate derivative metabolic proces... | 344 | 5 | 1.19 | 18 | 1.00000 | 0.00546 |
| 22 | GO:0051707 | response to other organism | 92 | 3 | 0.32 | 13 | 1.00000 | 0.00367 |
| 23 | GO:0009617 | response to bacterium | 64 | 3 | 0.22 | 11 | 1.00000 | 0.00130 |
| 29 | GO:0006952 | defense response | 108 | 3 | 0.37 | 19 | 1.00000 | 0.00576 |

| | GO.ID | MFCluster: coral2 Size: 34 | Annotated | Significant | Expected | Rank in ClassicF | Weight01F | ClassicF |
|---|---|---|---|---|---|---|---|---|
| *8* | GO:0003824 | catalytic activity | 3160 | 14 | 8.33 | 10 | 0.145 | 0.0082 |
| *22* | GO:0140096 | catalytic activity, acting on a protein | 947 | 8 | 2.50 | 9 | 1.000 | 0.0017 |
| *26* | GO:0016787 | hydrolase activity | 1445 | 10 | 3.81 | 8 | 1.000 | 0.0016 |

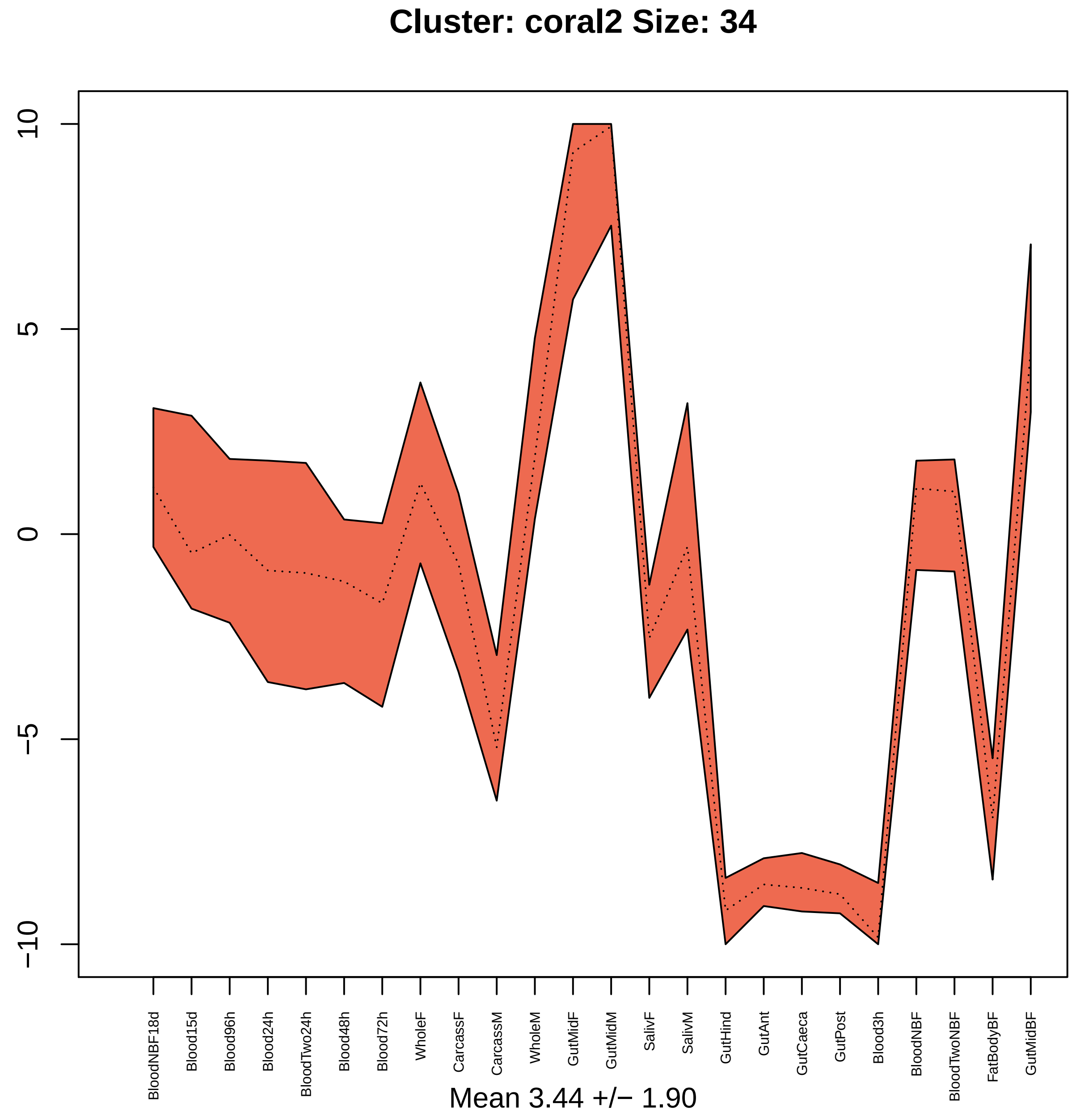

Cluster: coral2 Size: 34
10
5
0
−5
−10
BloodNBF18d
Blood15d
Blood96h
Blood24h
BloodTwo24h
Blood48h
Blood72h
WholeF
CarcassF
CarcassM
WholeM
GutMidF
GutMidM
SalivF
SalivM
GutHind
GutAnt
GutCaeca
GutPost
Blood3h
BloodNBF
BloodTwoNBF
FatBodyBF
GutMidBF
Mean 3.44 +/− 1.90

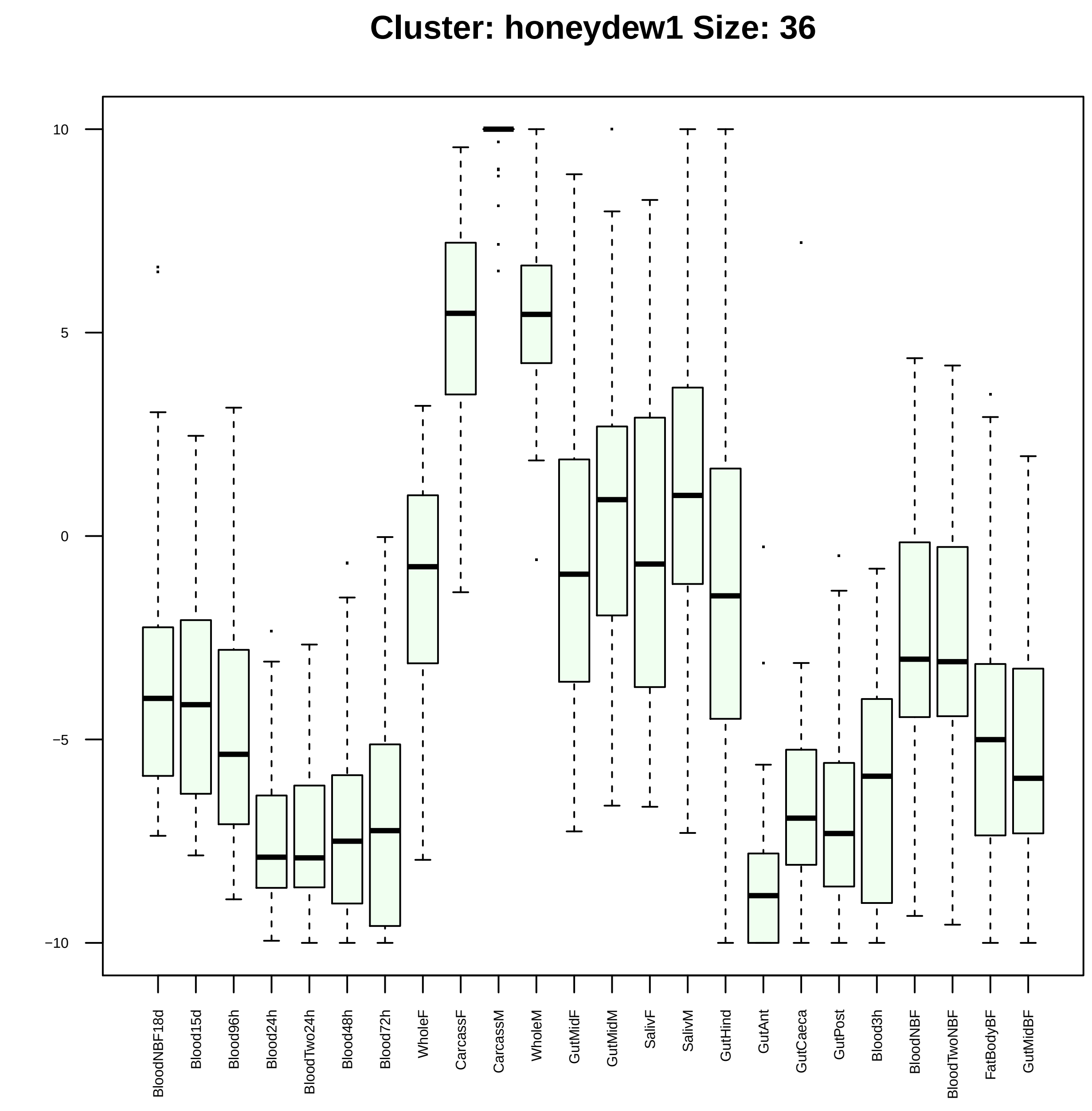

Cluster: honeydew1 Size: 36
10
5
0
−5
−10
BloodNBF18d
Blood15d
Blood96h
Blood24h
BloodTwo24h
Blood48h
Blood72h
WholeF
CarcassF
CarcassM
WholeM
GutMidF
GutMidM
SalivF
SalivM
GutHind
GutAnt
GutCaeca
GutPost
Blood3h
BloodNBF
BloodTwoNBF
FatBodyBF
GutMidBF

| | GO.ID | BPCluster: honeydew1 Size: 36 | Annotated | Significant | Expected | Rank in ClassicF | Weight01F | ClassicF |
|---|---|---|---|---|---|---|---|---|
| *1* | GO:0070997 | neuron death | 15 | 2 | 0.06 | 1 | 0.0036 | 0.0014 |
| *2* | GO:0048284 | organelle fusion | 33 | 2 | 0.12 | 3 | 0.0286 | 0.0067 |
| *3* | GO:0022411 | cellular component disassembly | 36 | 2 | 0.14 | 4 | 0.0322 | 0.0079 |
| *21* | GO:0010639 | negative regulation of organelle organiz... | 29 | 2 | 0.11 | 2 | 0.0632 | 0.0052 |

| | GO.ID | MFCluster: honeydew1 Size: 36 | Annotated | Significant | Expected | Rank in ClassicF | Weight01F | ClassicF |
|---|---|---|---|---|---|---|---|---|
| *1* | GO:0003779 | actin binding | 82 | 3 | 0.30 | 1 | 0.0031 | 0.0031 |
| *2* | GO:0015294 | solute:cation symporter activity | 37 | 2 | 0.13 | 2 | 0.0207 | 0.0077 |

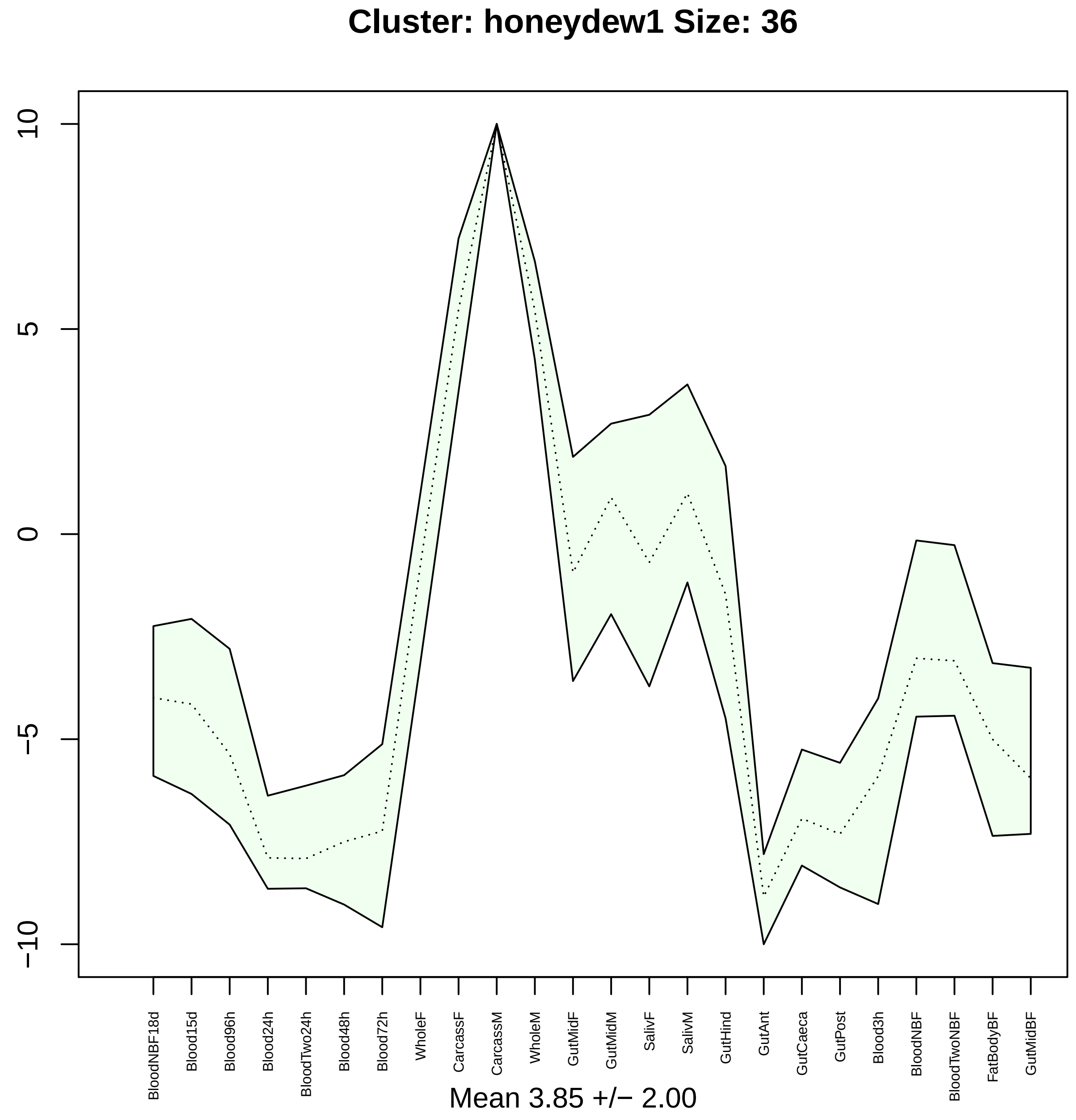
Cluster: honeydew1 Size: 36
10
5
0
−5
−10
BloodNBF18d
Blood15d
Blood96h
Blood24h
BloodTwo24h
Blood48h
Blood72h
WholeF
CarcassF
CarcassM
WholeM
GutMidF
GutMidM
SalivF
SalivM
GutHind
GutAnt
GutCaeca
GutPost
Blood3h
BloodNBF
BloodTwoNBF
FatBodyBF
GutMidBF
Mean 3.85 +/− 2.00

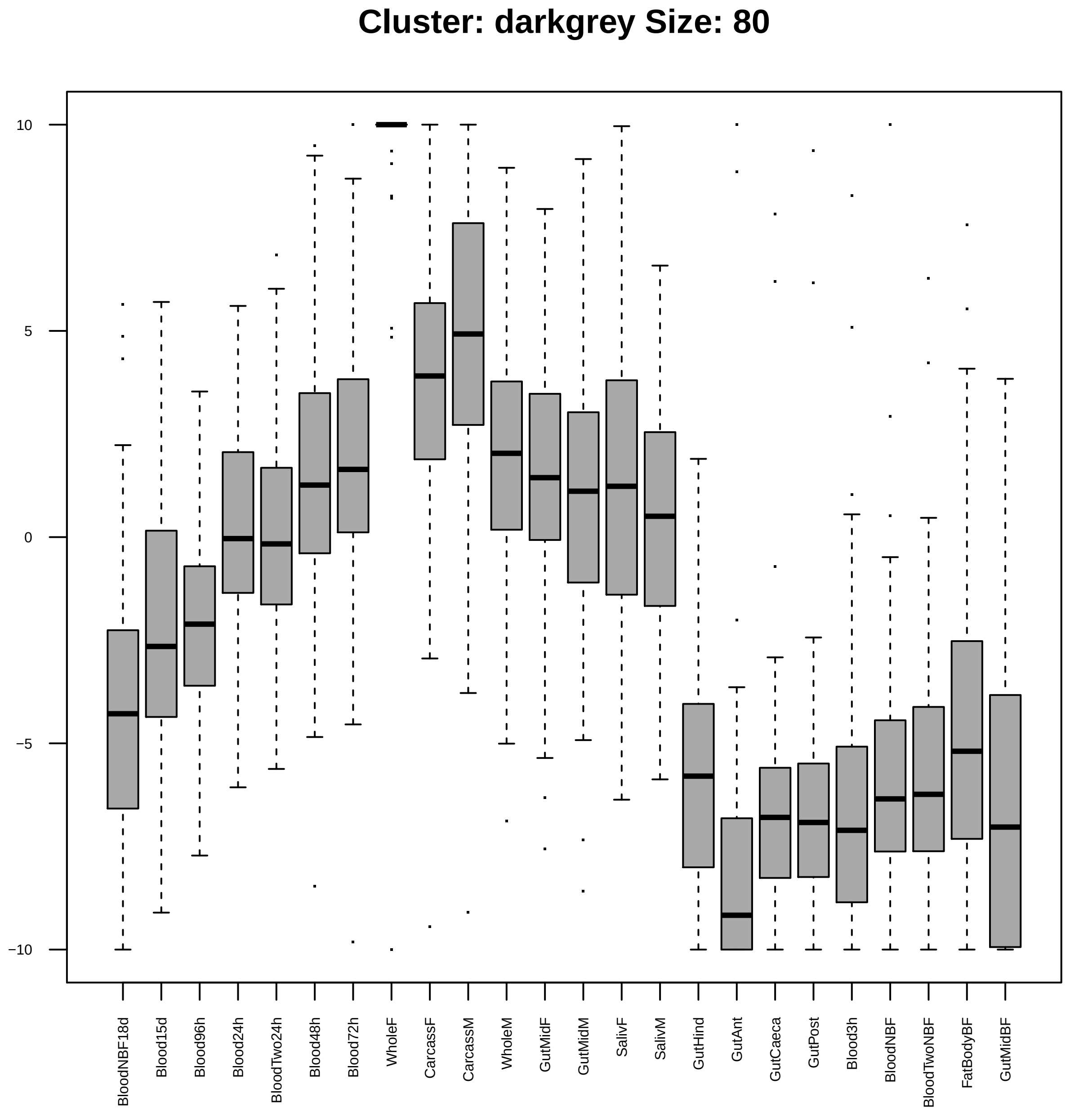
Cluster: darkgrey Size: 80
10
5
0
−5
−10
BloodNBF18d
Blood15d
Blood96h
Blood24h
BloodTwo24h
Blood48h
Blood72h
WholeF
CarcassF
CarcassM
WholeM
GutMidF
GutMidM
SalivF
SalivM
GutHind
GutAnt
GutCaeca
GutPost
Blood3h
BloodNBF
BloodTwoNBF
FatBodyBF
GutMidBF

|  | GO.ID | BPCluster: darkgrey Size: 80 | Annotated | Significant | Expected | Rank in ClassicF | Weight01F | ClassicF |
|---|---|---|---|---|---|---|---|---|
| 2 | GO:2001251 | negative regulation of chromosome organi... | 14 | 3 | 0.11 | 46 | 0.00017 | 0.00017 |
| 3 | GO:0007052 | mitotic spindle organization | 114 | 6 | 0.93 | 51 | 0.00030 | 0.00030 |
| 4 | GO:0022008 | neurogenesis | 706 | 13 | 5.76 | 98 | 0.00050 | 0.00354 |
| 5 | GO:0000077 | DNA damage checkpoint | 53 | 4 | 0.43 | 66 | 0.00060 | 0.00086 |
| 6 | GO:0006261 | DNA−dependent DNA replication | 52 | 4 | 0.42 | 64 | 0.00080 | 0.00080 |
| 7 | GO:0007474 | imaginal disc−derived wing vein specific... | 26 | 3 | 0.21 | 75 | 0.00116 | 0.00116 |
| 8 | GO:0007093 | mitotic cell cycle checkpoint | 56 | 4 | 0.46 | 71 | 0.00212 | 0.00106 |
| 9 | GO:0008544 | epidermis development | 34 | 3 | 0.28 | 90 | 0.00220 | 0.00256 |
| 10 | GO:0022403 | cell cycle phase | 10 | 2 | 0.08 | 91 | 0.00281 | 0.00281 |
| 11 | GO:0030261 | chromosome condensation | 25 | 4 | 0.20 | 32 | 0.00318 | 4.4e−05 |
| 12 | GO:2000241 | regulation of reproductive process | 30 | 3 | 0.24 | 79 | 0.00331 | 0.00178 |
| 13 | GO:0007076 | mitotic chromosome condensation | 14 | 2 | 0.11 | 110 | 0.00557 | 0.00557 |
| 14 | GO:0051649 | establishment of localization in cell | 317 | 6 | 2.58 | 208 | 0.00601 | 0.04281 |
| 15 | GO:0000413 | protein peptidyl−prolyl isomerization | 15 | 2 | 0.12 | 113 | 0.00640 | 0.00640 |
| 16 | GO:1901991 | negative regulation of mitotic cell cycl... | 60 | 4 | 0.49 | 77 | 0.00683 | 0.00137 |
| 20 | GO:0006338 | chromatin remodeling | 37 | 3 | 0.30 | 96 | 0.01943 | 0.00327 |
| 23 | GO:0140013 | meiotic nuclear division | 68 | 4 | 0.55 | 88 | 0.02311 | 0.00218 |
| 28 | GO:0010389 | regulation of G2/M transition of mitotic... | 48 | 3 | 0.39 | 116 | 0.03122 | 0.00683 |

| | GO.ID | MFCluster: darkgrey Size: 80 | Annotated | Significant | Expected | Rank in ClassicF | Weight01F | ClassicF |
|---|---|---|---|---|---|---|---|---|
| 1 | GO:0003677 | DNA binding | 523 | 13 | 4.43 | 4 | 0.00053 | 0.00034 |
| 2 | GO:0008270 | zinc ion binding | 522 | 12 | 4.42 | 7 | 0.00120 | 0.00120 |
| 3 | GO:0008094 | DNA−dependent ATPase activity | 33 | 4 | 0.28 | 3 | 0.00415 | 0.00016 |
| 4 | GO:0005524 | ATP binding | 592 | 11 | 5.01 | 15 | 0.00993 | 0.00993 |
| 8 | GO:0005515 | protein binding | 2143 | 28 | 18.13 | 11 | 0.02336 | 0.00534 |
| 10 | GO:0140097 | catalytic activity, acting on DNA | 81 | 5 | 0.69 | 5 | 0.02899 | 0.00057 |
| 13 | GO:0008026 | ATP−dependent helicase activity | 76 | 4 | 0.64 | 9 | 0.07458 | 0.00378 |
| 18 | GO:0016853 | isomerase activity | 84 | 4 | 0.71 | 12 | 0.09415 | 0.00541 |

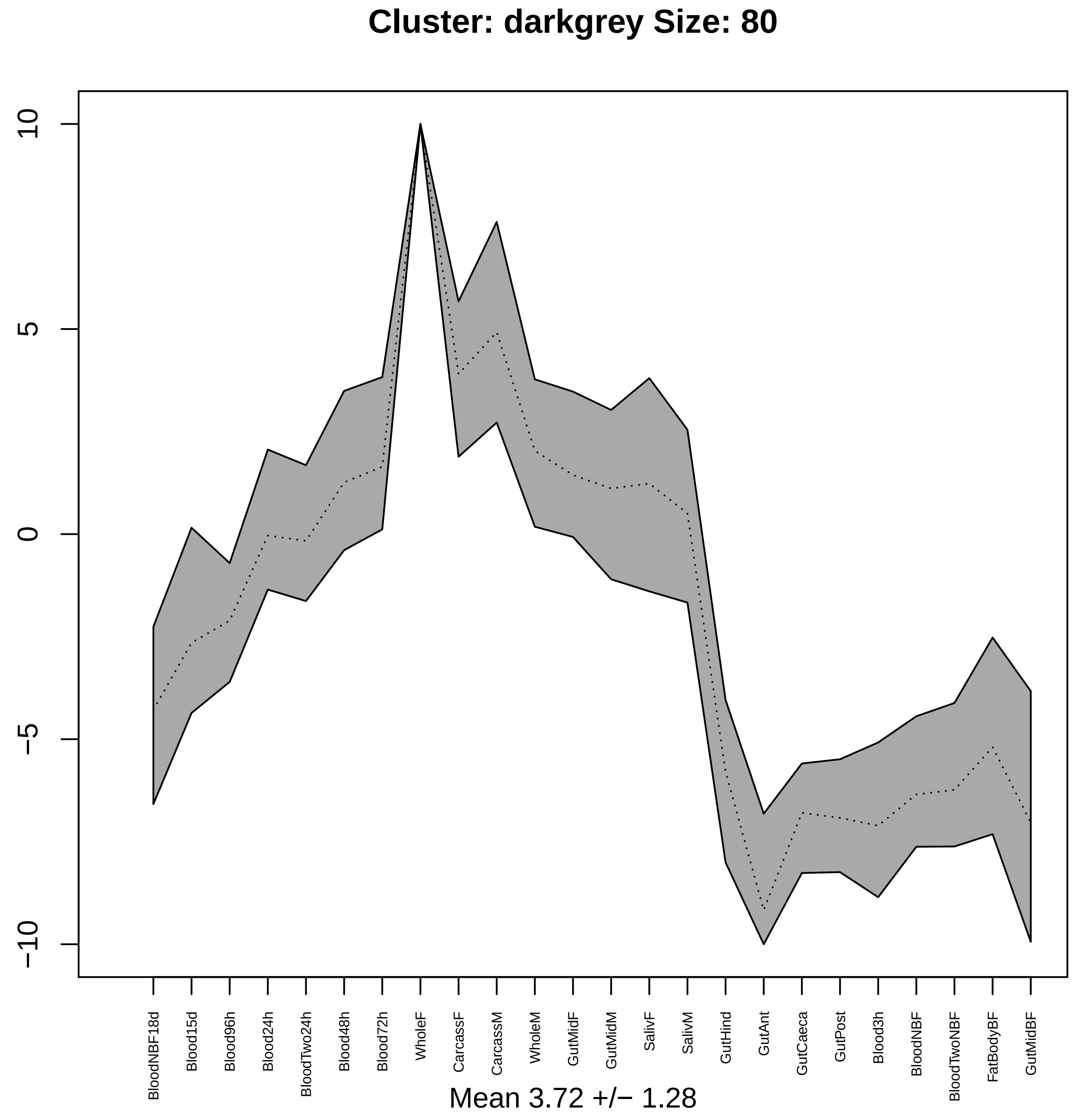
Cluster: darkgrey Size: 80
10
5
0
−5
−10
BloodNBF18d
Blood15d
Blood96h
Blood24h
BloodTwo24h
Blood48h
Blood72h
WholeF
CarcassF
CarcassM
WholeM
GutMidF
GutMidM
SalivF
SalivM
GutHind
GutAnt
GutCaeca
GutPost
Blood3h
BloodNBF
BloodTwoNBF
FatBodyBF
GutMidBF
Mean 3.72 +/− 1.28

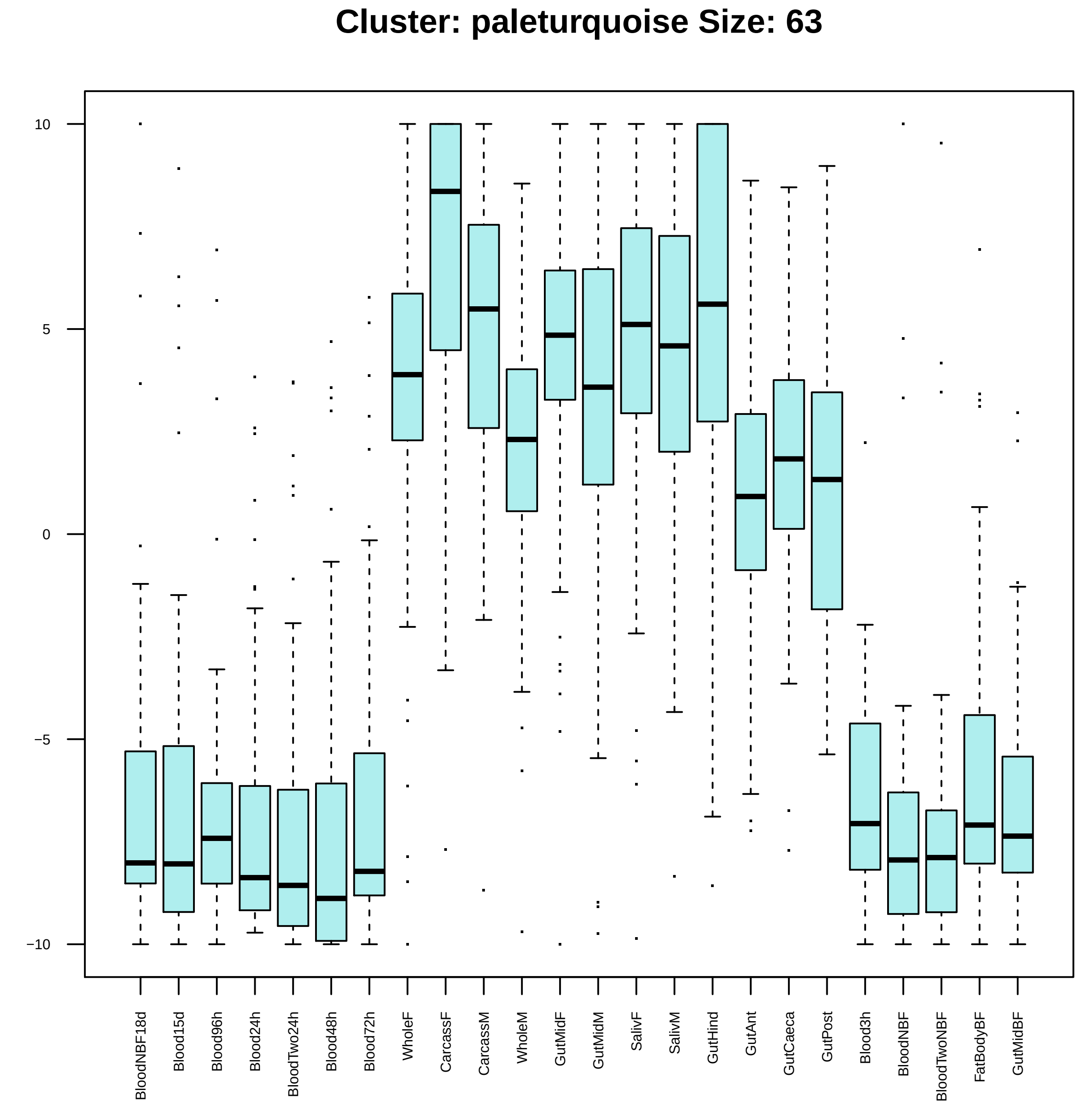

Cluster: paleturquoise Size: 63
10
5
0
−5
−10
BloodNBF18d
Blood15d
Blood96h
Blood24h
BloodTwo24h
Blood48h
Blood72h
WholeF
CarcassF
CarcassM
WholeM
GutMidF
GutMidM
SalivF
SalivM
GutHind
GutAnt
GutCaeca
GutPost
Blood3h
BloodNBF
BloodTwoNBF
FatBodyBF
GutMidBF

| | GO.ID | BPCluster: paleturquoise Size: 63 | Annotated | Significant | Expected | Rank in ClassicF | Weight01F | ClassicF |
|---|---|---|---|---|---|---|---|---|
| 3 | GO:0006413 | translational initiation | 39 | 3 | 0.18 | 16 | 0.0033 | 0.00077 |
| 4 | GO:0009058 | biosynthetic process | 1357 | 14 | 6.38 | 21 | 0.0066 | 0.00166 |
| 13 | GO:0140014 | mitotic nuclear division | 105 | 5 | 0.49 | 10 | 0.0758 | 0.00011 |

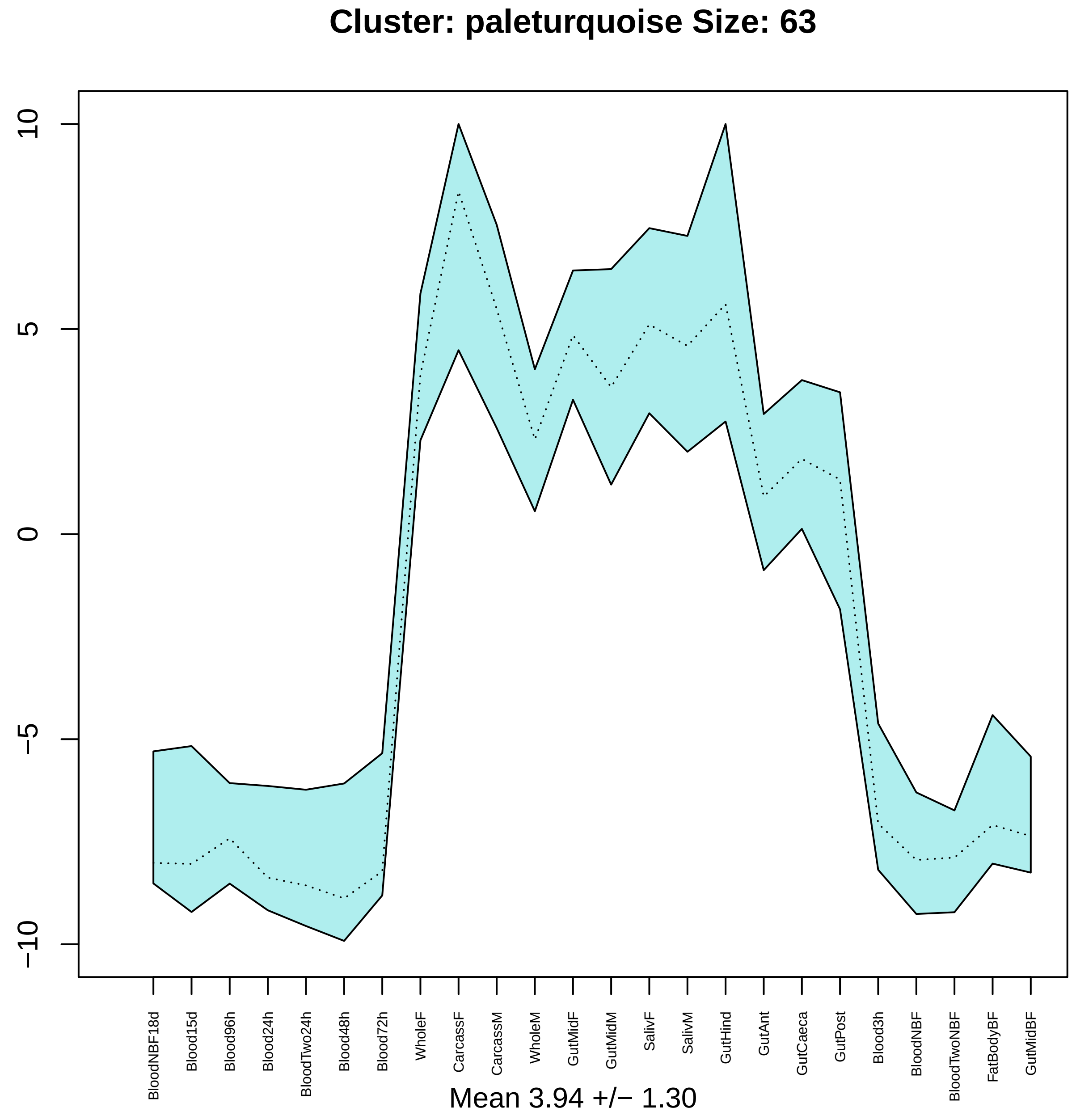

Cluster: paleturquoise Size: 63
10
5
0
−5
−10
BloodNBF18d
Blood15d
Blood96h
Blood24h
BloodTwo24h
Blood48h
Blood72h
WholeF
CarcassF
CarcassM
WholeM
GutMidF
GutMidM
SalivF
SalivM
GutHind
GutAnt
GutCaeca
GutPost
Blood3h
BloodNBF
BloodTwoNBF
FatBodyBF
GutMidBF
Mean 3.94 +/− 1.30

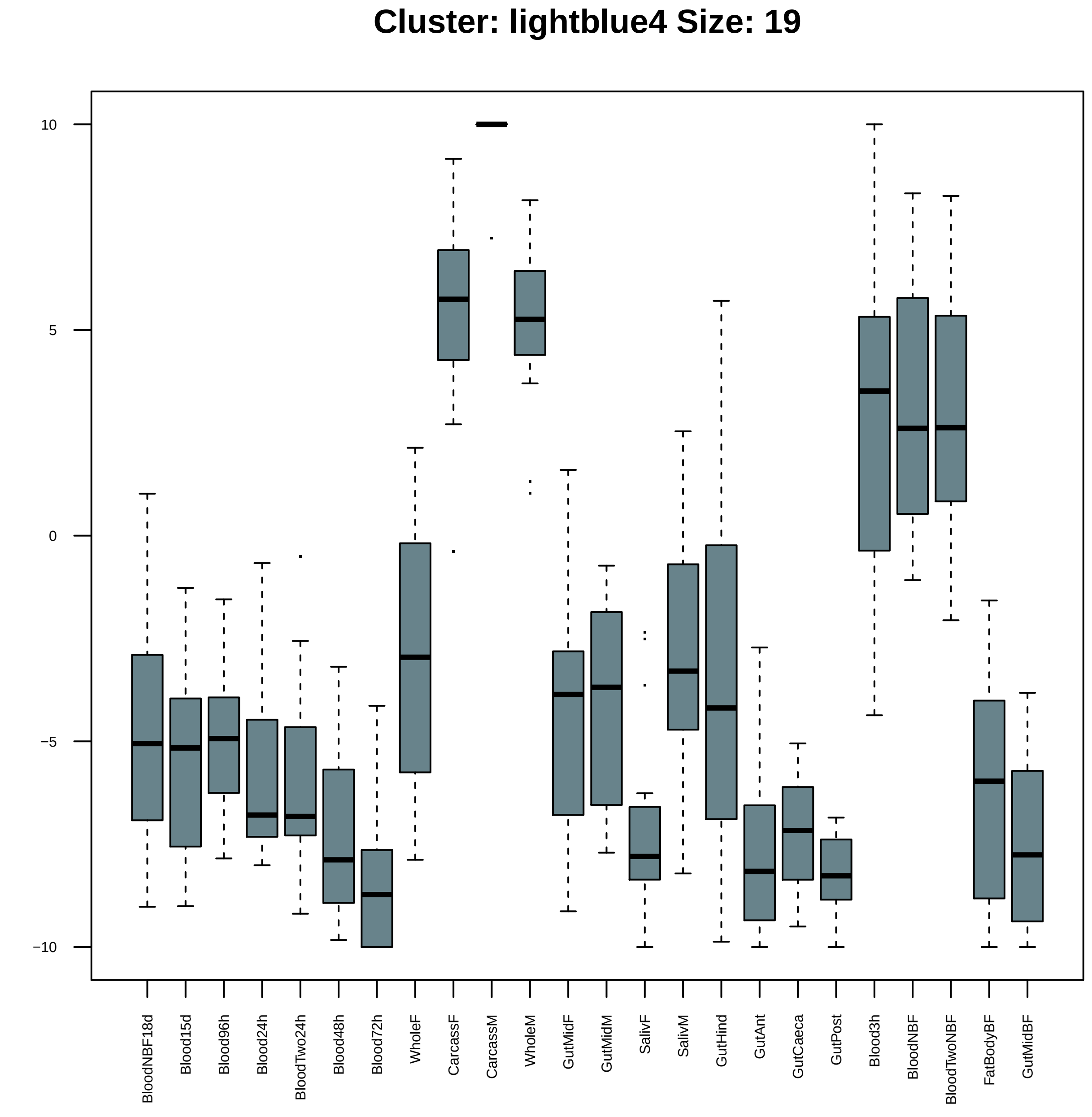

Cluster: lightblue4 Size: 19
10
5
0
−5
−10
BloodNBF18d
Blood15d
Blood96h
Blood24h
BloodTwo24h
Blood48h
Blood72h
WholeF
CarcassF
CarcassM
WholeM
GutMidF
GutMidM
SalivF
SalivM
GutHind
GutAnt
GutCaeca
GutPost
Blood3h
BloodNBF
BloodTwoNBF
FatBodyBF
GutMidBF

| | GO.ID | BPCluster: lightblue4 Size: 19 | Annotated | Significant | Expected | Rank in ClassicF | Weight01F | ClassicF |
|---|---|---|---|---|---|---|---|---|
| *1* | GO:0003012 | muscle system process | 14 | 2 | 0.02 | 1 | 0.0063 | 0.00024 |
| *29* | GO:0070925 | organelle assembly | 89 | 2 | 0.15 | 2 | 1.0000 | 0.00976 |

| | GO.ID | MFCluster: lightblue4 Size: 19 | Annotated | Significant | Expected | Rank in ClassicF | Weight01F | ClassicF |
|---|---|---|---|---|---|---|---|---|
| 1 | GO:0003774 | motor activity | 60 | 3 | 0.11 | 1 | 0.00015 | 0.00015 |
| 2 | GO:0005509 | calcium ion binding | 173 | 4 | 0.31 | 2 | 0.00019 | 0.00019 |

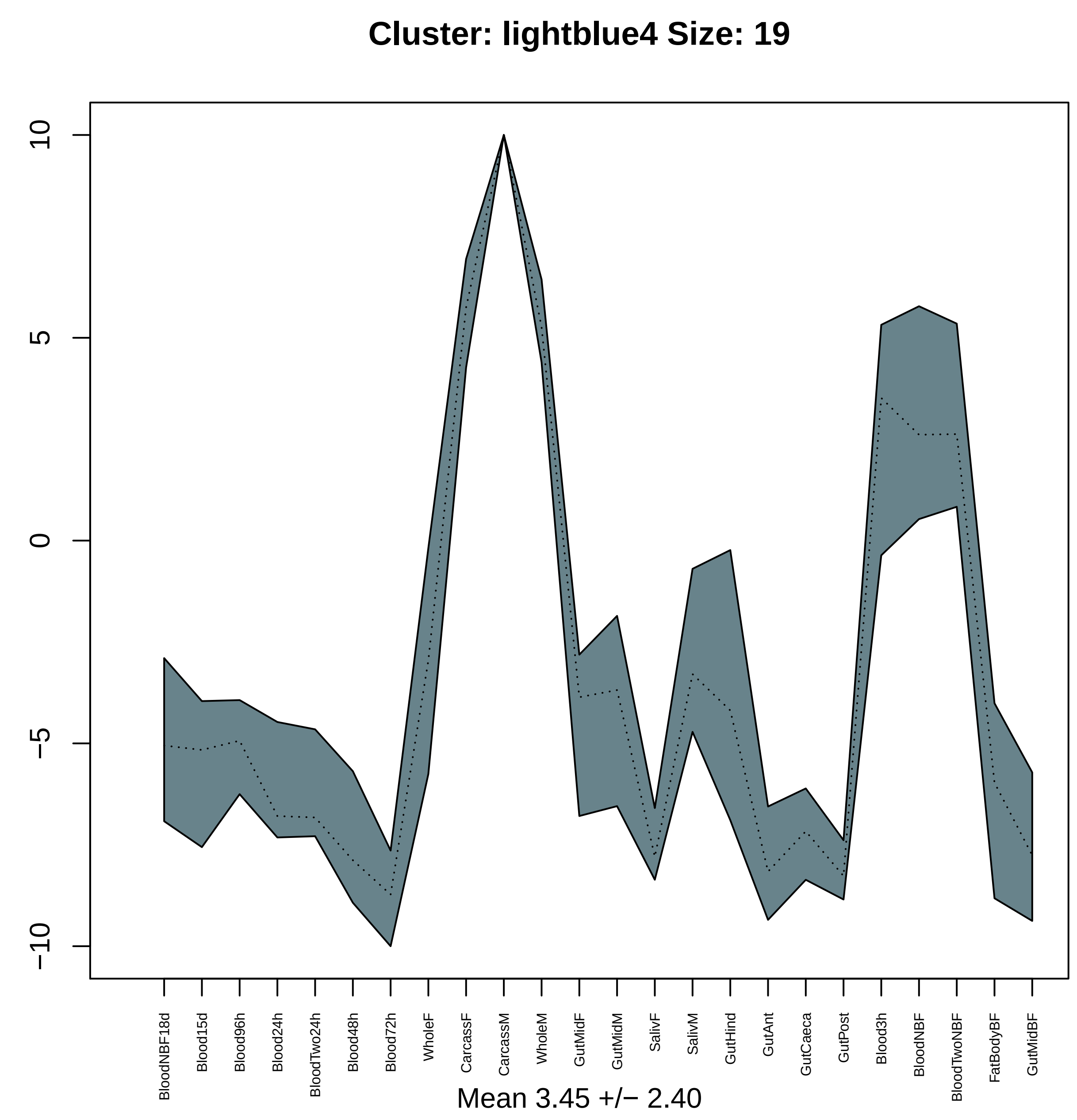
Cluster: lightblue4 Size: 19
10
5
0
−5
−10
BloodNBF18d
Blood15d
Blood96h
Blood24h
BloodTwo24h
Blood48h
Blood72h
WholeF
CarcassF
CarcassM
WholeM
GutMidF
GutMidM
SalivF
SalivM
GutHind
GutAnt
GutCaeca
GutPost
Blood3h
BloodNBF
BloodTwoNBF
FatBodyBF
GutMidBF
Mean 3.45 +/− 2.40

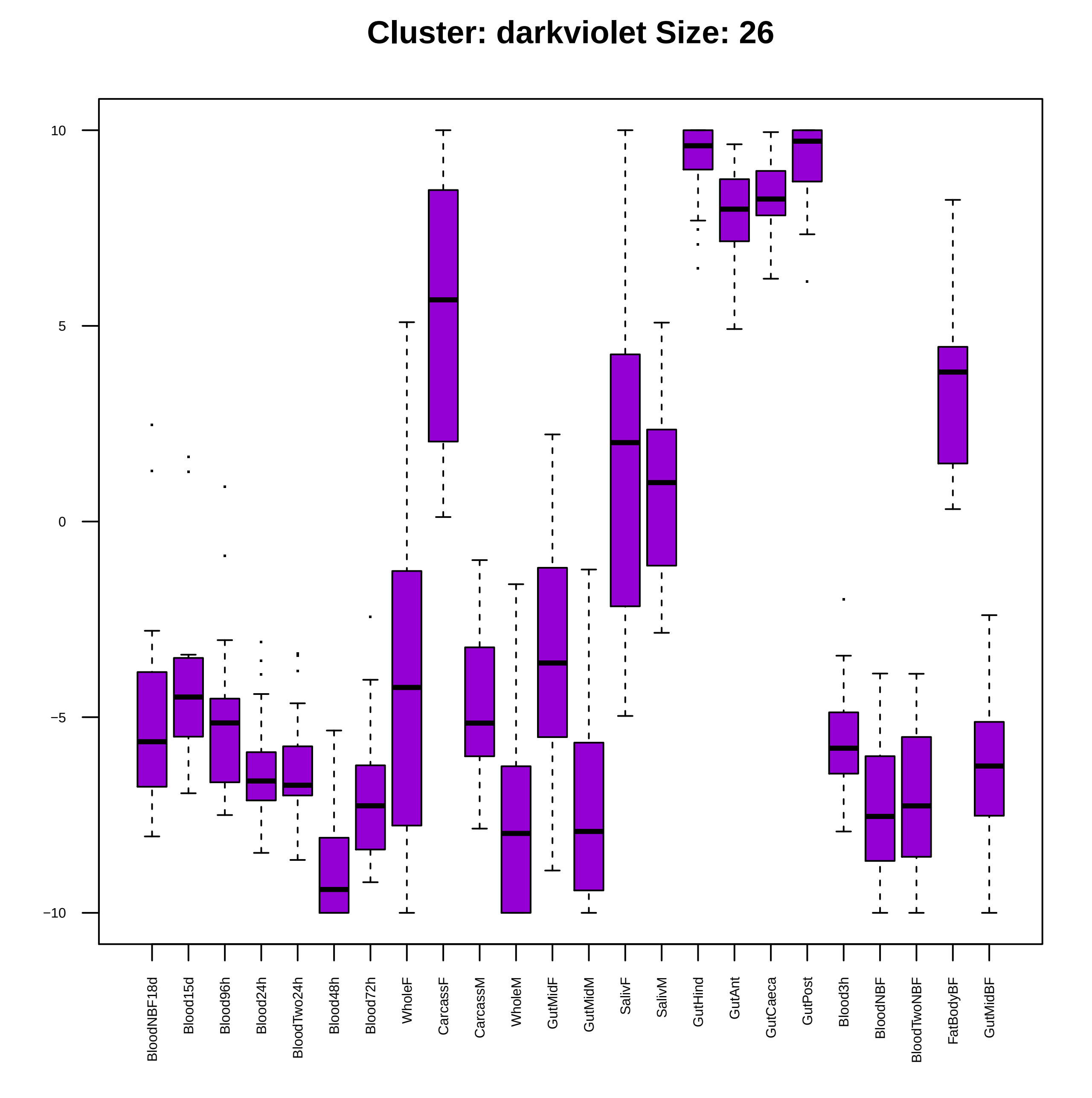
Cluster: darkviolet Size: 26
10
5
0
−5
−10
BloodNBF18d
Blood15d
Blood96h
Blood24h
BloodTwo24h
Blood48h
Blood72h
WholeF
CarcassF
CarcassM
WholeM
GutMidF
GutMidM
SalivF
SalivM
GutHind
GutAnt
GutCaeca
GutPost
Blood3h
BloodNBF
BloodTwoNBF
FatBodyBF
GutMidBF

| | GO.ID | BPCluster: darkviolet Size: 26 | Annotated | Significant | Expected | Rank in ClassicF | Weight01F | ClassicF |
|---|---|---|---|---|---|---|---|---|
| *1* | GO:0006412 | translation | 289 | 25 | 1.13 | 1 | < 1e−30 | < 1e−30 |
| *4* | GO:0006414 | translational elongation | 22 | 2 | 0.09 | 50 | 0.0032 | 0.0032 |
| *29* | GO:0071840 | cellular component organization or bioge... | 1209 | 11 | 4.74 | 51 | 1.0000 | 0.0035 |

| | GO.ID | MFCluster: darkviolet Size: 26 | Annotated | Significant | Expected | Rank in ClassicF | Weight01F | ClassicF |
|---|---|---|---|---|---|---|---|---|
| *1* | GO:0003735 | structural constituent of ribosome | 146 | 24 | 0.51 | 1 | < 1e−30 | < 1e−30 |
| *11* | GO:0003723 | RNA binding | 307 | 5 | 1.06 | 4 | 1.000 | 0.0036 |
| *14* | GO:0005198 | structural molecule activity | 318 | 24 | 1.10 | 2 | 1.000 | < 1e−30 |

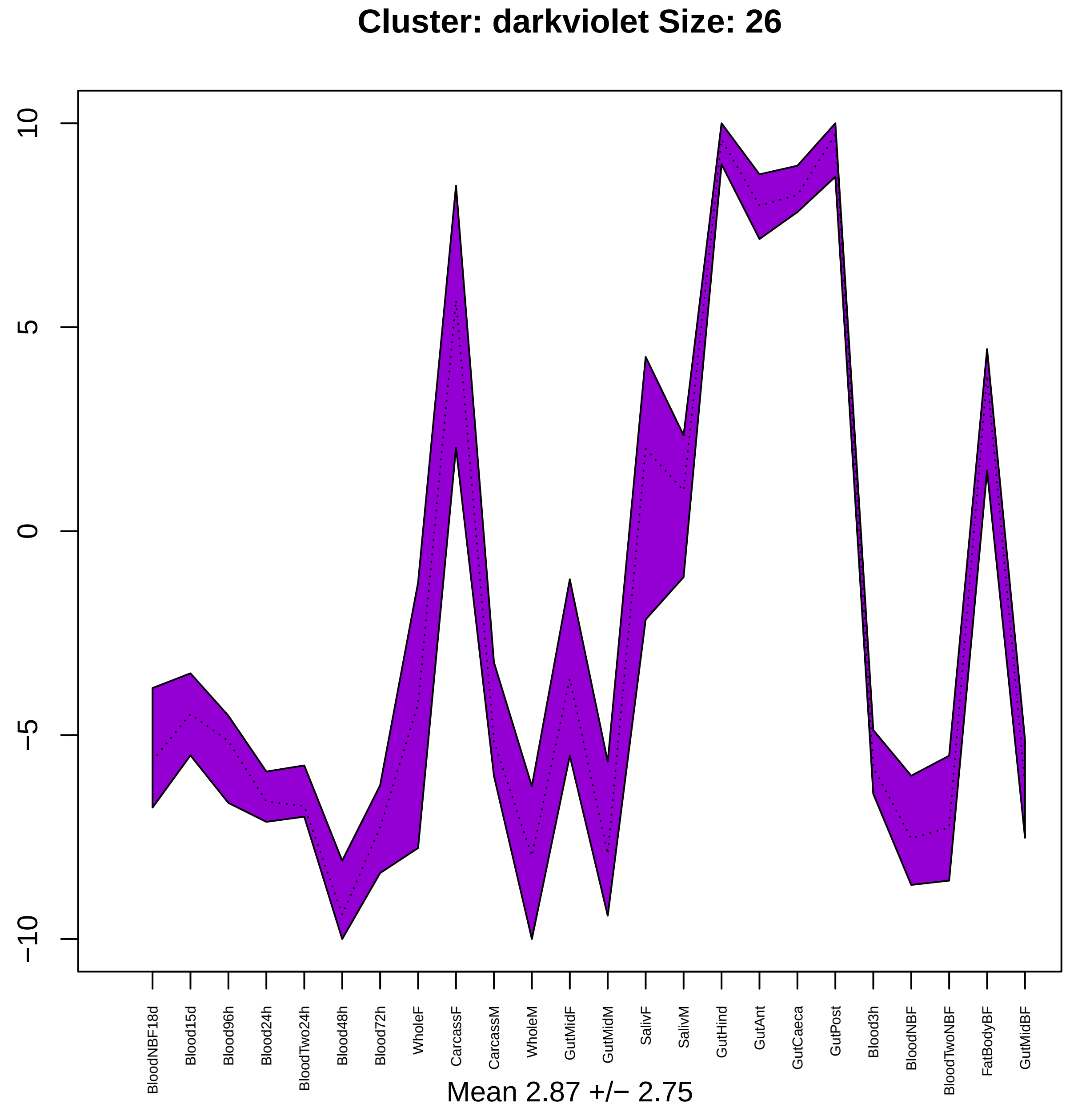

Cluster: darkviolet Size: 26
10
5
0
−5
−10
BloodNBF18d
Blood15d
Blood96h
Blood24h
BloodTwo24h
Blood48h
Blood72h
WholeF
CarcassF
CarcassM
WholeM
GutMidF
GutMidM
SalivF
SalivM
GutHind
GutAnt
GutCaeca
GutPost
Blood3h
BloodNBF
BloodTwoNBF
FatBodyBF
GutMidBF
Mean 2.87 +/− 2.75

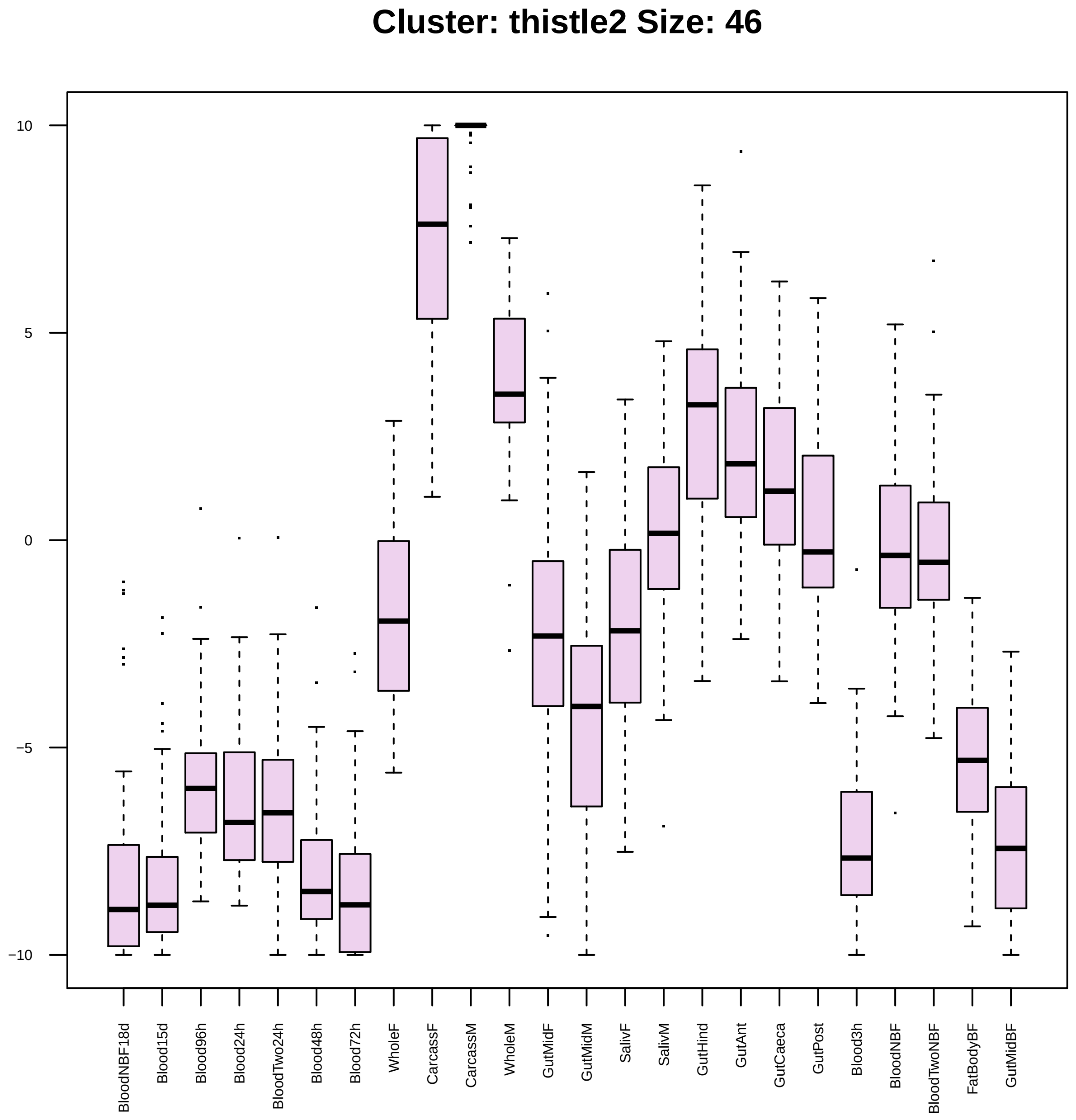

Cluster: thistle2 Size: 46
10
5
0
−5
−10
BloodNBF18d
Blood15d
Blood96h
Blood24h
BloodTwo24h
Blood48h
Blood72h
WholeF
CarcassF
CarcassM
WholeM
GutMidF
GutMidM
SalivF
SalivM
GutHind
GutAnt
GutCaeca
GutPost
Blood3h
BloodNBF
BloodTwoNBF
FatBodyBF
GutMidBF

| | GO.ID | BPCluster: thistle2 Size: 46 | Annotated | Significant | Expected | Rank in ClassicF | Weight01F | ClassicF |
|---|---|---|---|---|---|---|---|---|
| *29* | GO:0009165 | nucleotide biosynthetic process | 90 | 3 | 0.44 | 63 | 1.000 | 0.0092 |

| | GO.ID | MFCluster: thistle2 Size: 46 | Annotated | Significant | Expected | Rank in ClassicF | Weight01F | ClassicF |
|---|---|---|---|---|---|---|---|---|
| 5 | GO:0051539 | 4 iron, 4 sulfur cluster binding | 14 | 2 | 0.06 | 22 | 0.0015 | 0.00148 |
| 6 | GO:0022857 | transmembrane transporter activity | 442 | 10 | 1.84 | 9 | 0.0015 | 6.7e−06 |
| 7 | GO:0015078 | proton transmembrane transporter activit... | 52 | 6 | 0.22 | 8 | 0.0023 | 5.4e−08 |
| 8 | GO:0015662 | ATPase activity, coupled to transmembran... | 20 | 2 | 0.08 | 23 | 0.0030 | 0.00304 |
| 9 | GO:0044769 | ATPase activity, coupled to transmembran... | 22 | 2 | 0.09 | 25 | 0.0040 | 0.00367 |
| 10 | GO:0036442 | proton−exporting ATPase activity | 27 | 2 | 0.11 | 26 | 0.0240 | 0.00551 |
| 26 | GO:0008324 | cation transmembrane transporter activit... | 210 | 6 | 0.87 | 17 | 1.0000 | 0.00019 |
| 27 | GO:0003824 | catalytic activity | 3160 | 21 | 13.15 | 24 | 1.0000 | 0.00336 |
| 29 | GO:0051540 | metal cluster binding | 44 | 3 | 0.18 | 20 | 1.0000 | 0.00077 |

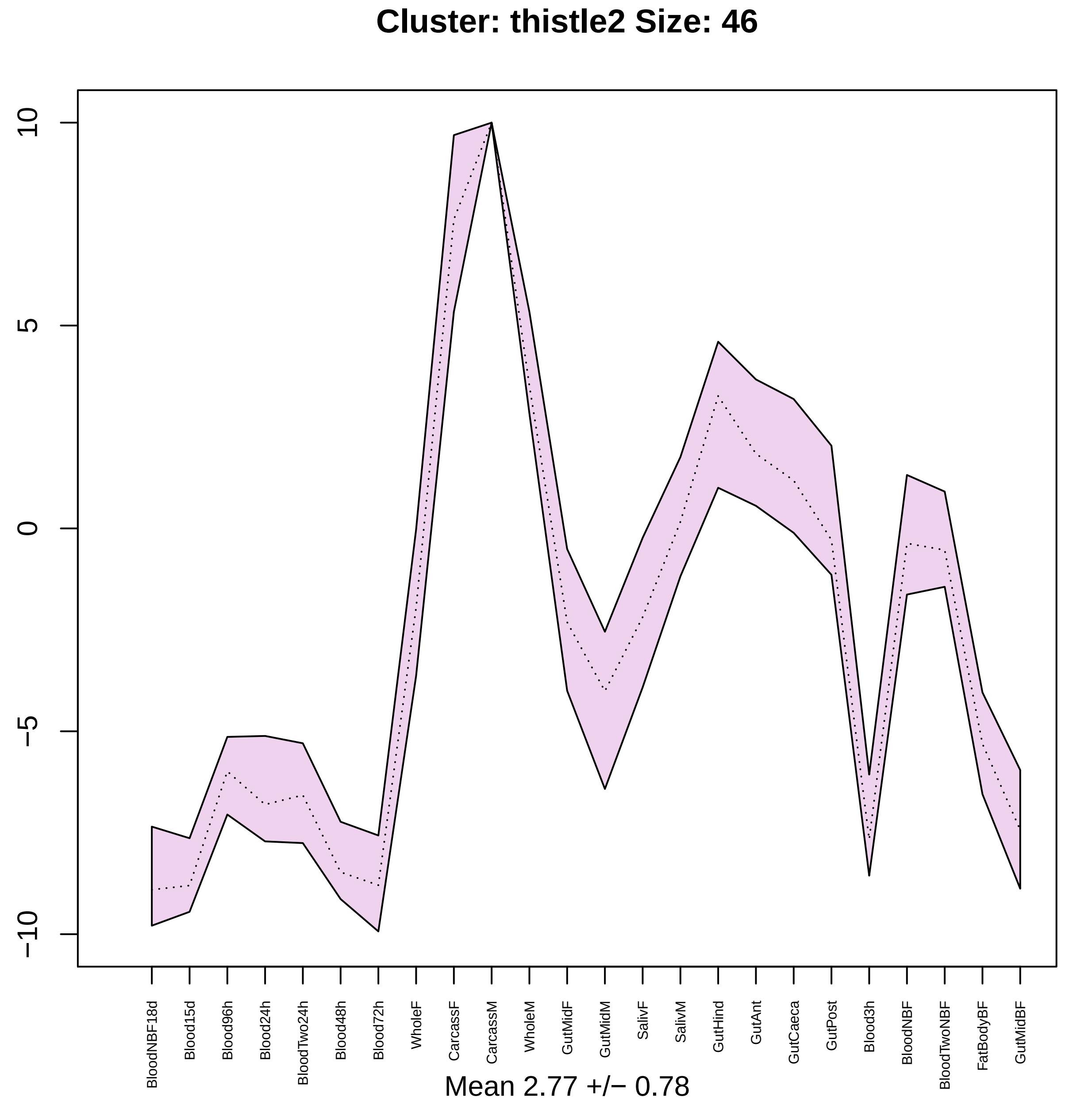
Cluster: thistle2 Size: 46
10
5
0
−5
−10
BloodNBF18d
Blood15d
Blood96h
Blood24h
BloodTwo24h
Blood48h
Blood72h
WholeF
CarcassF
CarcassM
WholeM
GutMidF
GutMidM
SalivF
SalivM
GutHind
GutAnt
GutCaeca
GutPost
Blood3h
BloodNBF
BloodTwoNBF
FatBodyBF
GutMidBF
Mean 2.77 +/− 0.78

### Cluster: palevioletred3 Size: 39

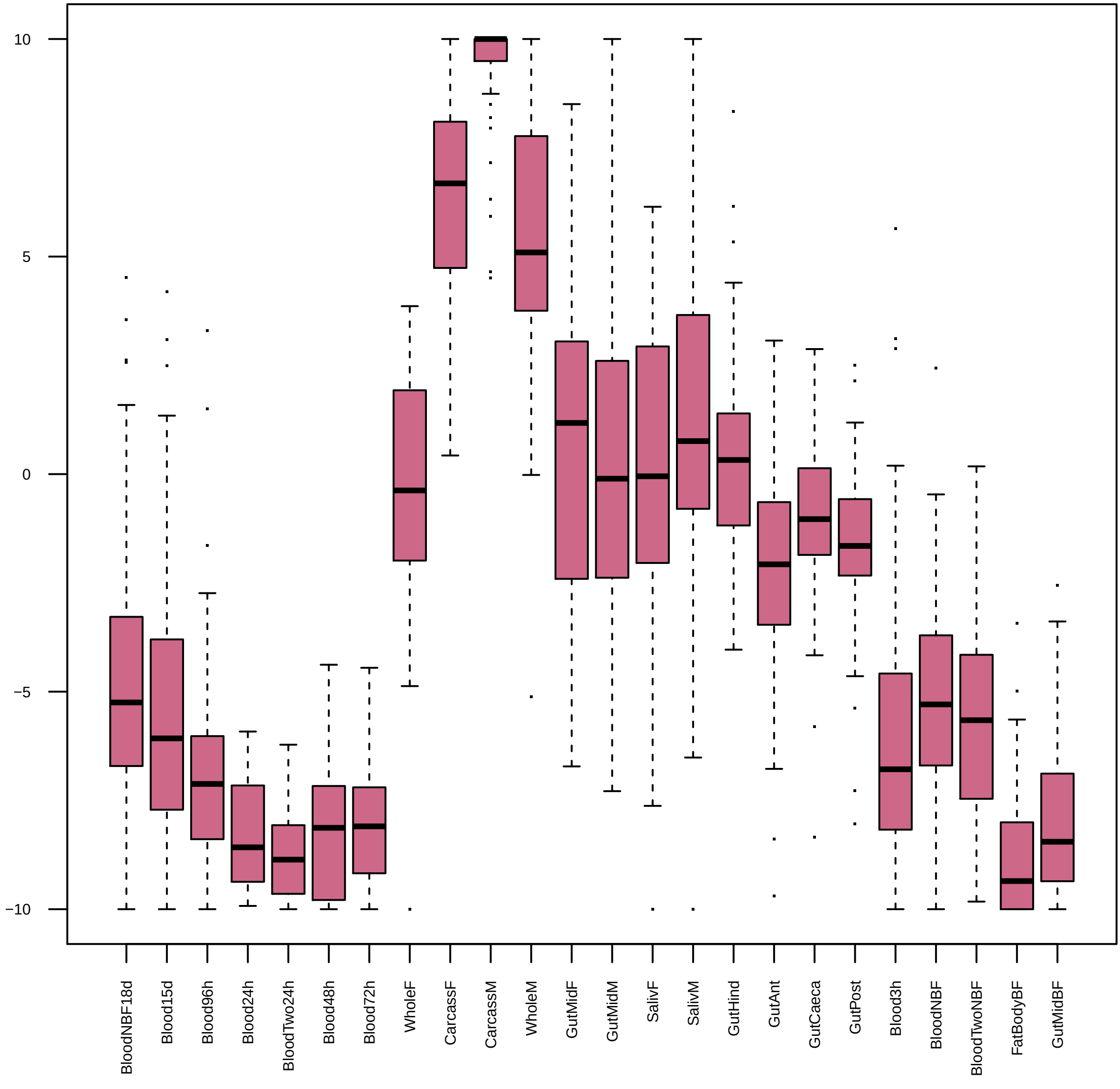

| | GO.ID | BPCluster: palevioletred3 Size: 39 | Annotated | Significant | Expected | Rank in ClassicF | Weight01F | ClassicF |
|---|---|---|---|---|---|---|---|---|
| 1 | GO:0016226 | iron−sulfur cluster assembly | 11 | 2 | 0.04 | 3 | 0.00061 | 0.00061 |
| 2 | GO:0042181 | ketone biosynthetic process | 14 | 2 | 0.05 | 5 | 0.00101 | 0.00101 |
| 3 | GO:0006733 | oxidoreduction coenzyme metabolic proces... | 34 | 2 | 0.12 | 8 | 0.00596 | 0.00596 |
| 4 | GO:0045454 | cell redox homeostasis | 36 | 2 | 0.12 | 9 | 0.00667 | 0.00667 |
| 5 | GO:0009058 | biosynthetic process | 1357 | 5 | 4.68 | 149 | 0.00740 | 0.51731 |
| 6 | GO:0055114 | oxidation−reduction process | 545 | 6 | 1.88 | 10 | 0.00863 | 0.00863 |
| 14 | GO:0019725 | cellular homeostasis | 84 | 4 | 0.29 | 1 | 0.06142 | 0.00017 |

|  | GO.ID | MFCluster: palevioletred3 Size: 39 | Annotated | Significant | Expected | Rank in ClassicF | Weight01F | ClassicF |
|---|---|---|---|---|---|---|---|---|
| *1* | GO:0015035 | protein disulfide oxidoreductase activit... | 14 | 2 | 0.04 | 1 | 0.00072 | 0.00072 |
| *2* | GO:0051536 | iron−sulfur cluster binding | 44 | 2 | 0.13 | 4 | 0.00710 | 0.00710 |
| *3* | GO:0009055 | electron transfer activity | 48 | 2 | 0.14 | 6 | 0.00841 | 0.00841 |

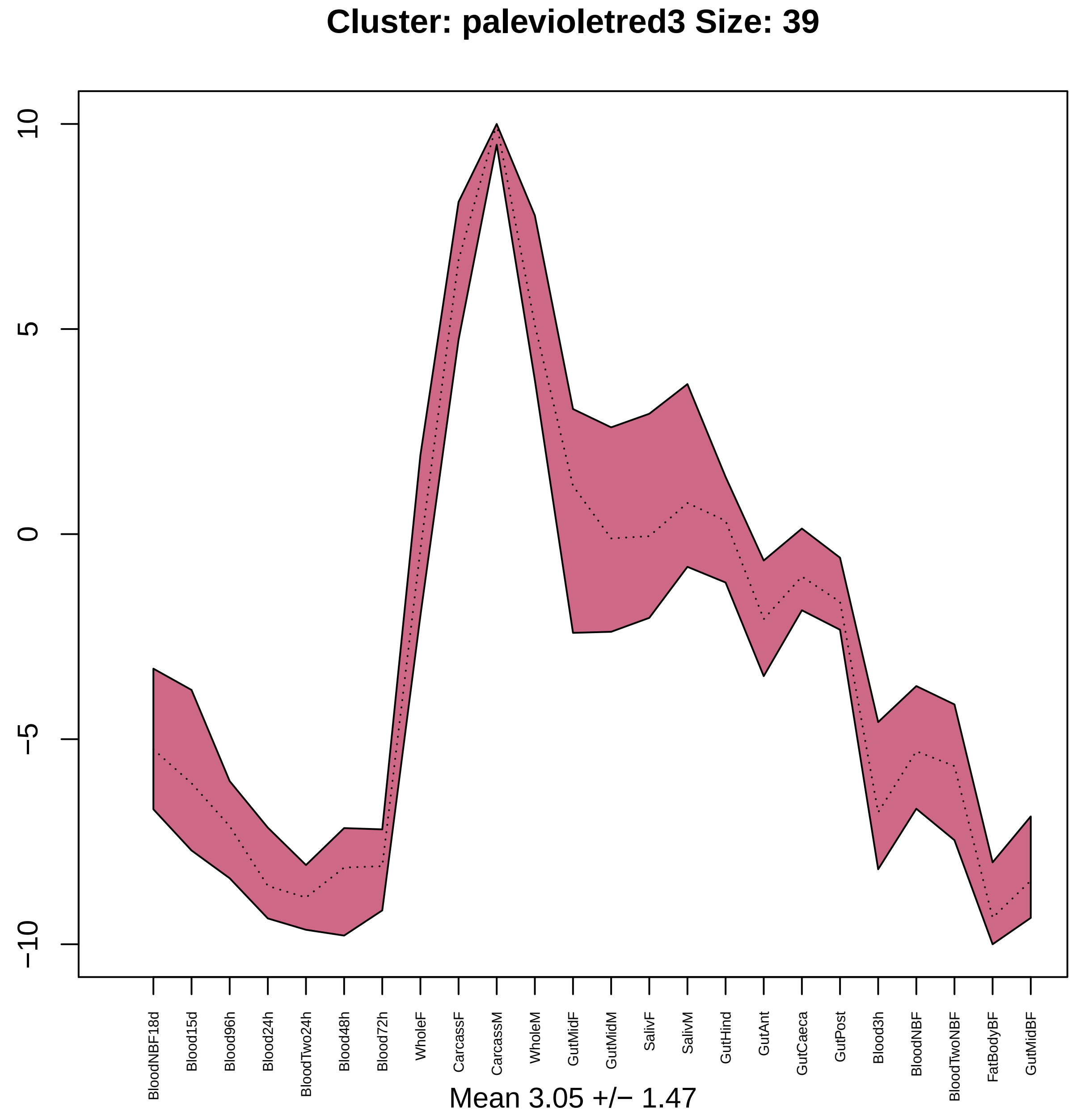
Cluster: palevioletred3 Size: 39
10
5
0
−5
−10
BloodNBF18d
Blood15d
Blood96h
Blood24h
BloodTwo24h
Blood48h
Blood72h
WholeF
CarcassF
CarcassM
WholeM
GutMidF
GutMidM
SalivF
SalivM
GutHind
GutAnt
GutCaeca
GutPost
Blood3h
BloodNBF
BloodTwoNBF
FatBodyBF
GutMidBF
Mean 3.05 +/− 1.47

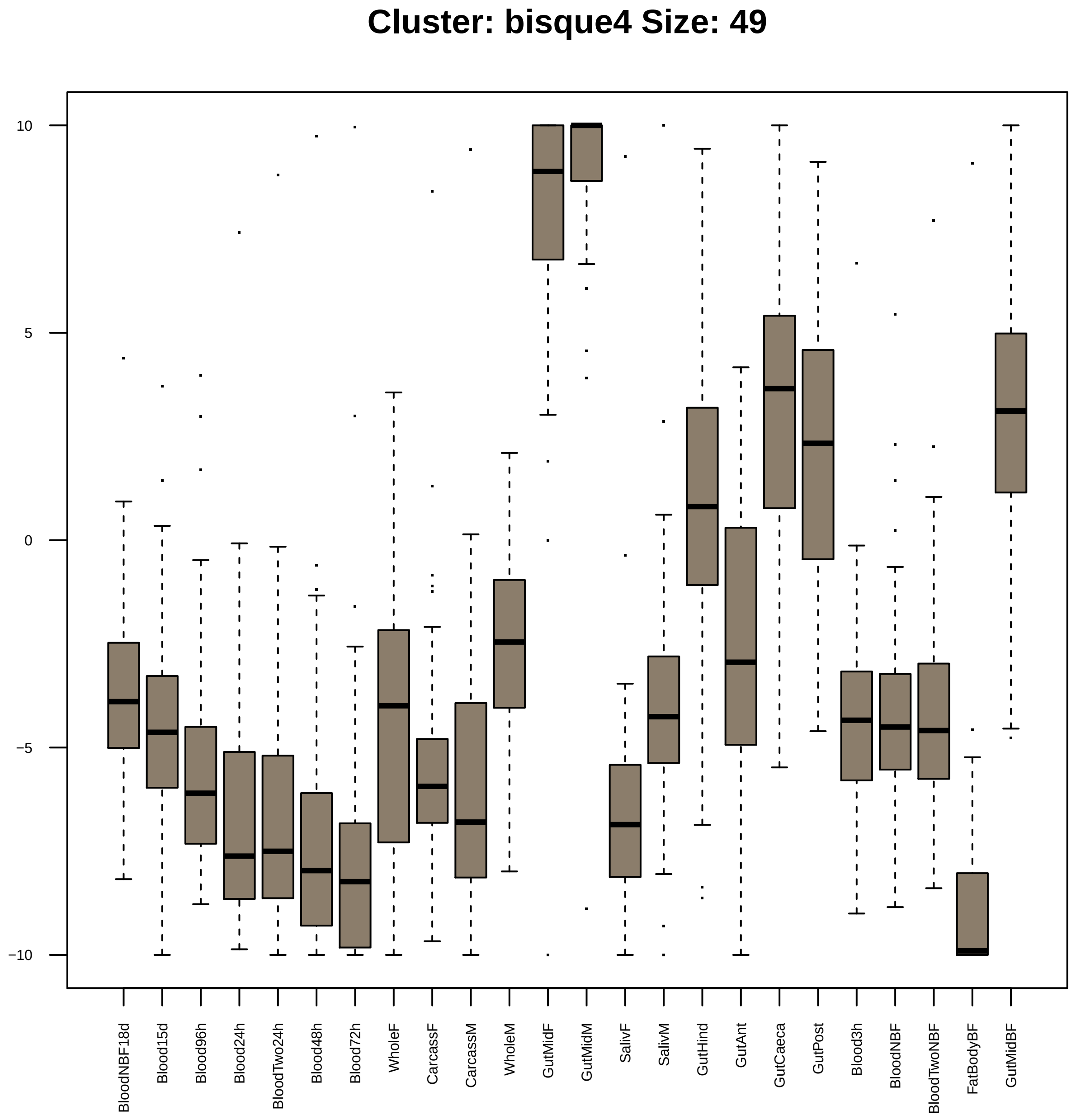
Cluster: bisque4 Size: 49
10
5
0
−5
−10
BloodNBF18d
Blood15d
Blood96h
Blood24h
BloodTwo24h
Blood48h
Blood72h
WholeF
CarcassF
CarcassM
WholeM
GutMidF
GutMidM
SalivF
SalivM
GutHind
GutAnt
GutCaeca
GutPost
Blood3h
BloodNBF
BloodTwoNBF
FatBodyBF
GutMidBF

| | GO.ID | BPCluster: bisque4 Size: 49 | Annotated | Significant | Expected | Rank in ClassicF | Weight01F | ClassicF |
|---|---|---|---|---|---|---|---|---|
| *1* | GO:0007156 | homophilic cell adhesion via plasma memb... | 23 | 2 | 0.1 | 1 | 0.0048 | 0.0048 |

| | GO.ID | MFCluster: bisque4 Size: 49 | Annotated | Significant | Expected | Rank in ClassicF | Weight01F | ClassicF |
|---|---|---|---|---|---|---|---|---|
| *1* | GO:0016705 | oxidoreductase activity, acting on paire... | 133 | 4 | 0.54 | 2 | 0.0018 | 0.00184 |
| *2* | GO:0005509 | calcium ion binding | 173 | 4 | 0.70 | 5 | 0.0048 | 0.00476 |
| *3* | GO:0008237 | metallopeptidase activity | 104 | 4 | 0.42 | 1 | 0.0056 | 0.00074 |
| *4* | GO:0001882 | nucleoside binding | 142 | 3 | 0.57 | 15 | 0.0076 | 0.01880 |
| *5* | GO:0004497 | monooxygenase activity | 110 | 3 | 0.44 | 10 | 0.0095 | 0.00947 |
| *29* | GO:0003824 | catalytic activity | 3160 | 20 | 12.71 | 6 | 0.4275 | 0.00549 |

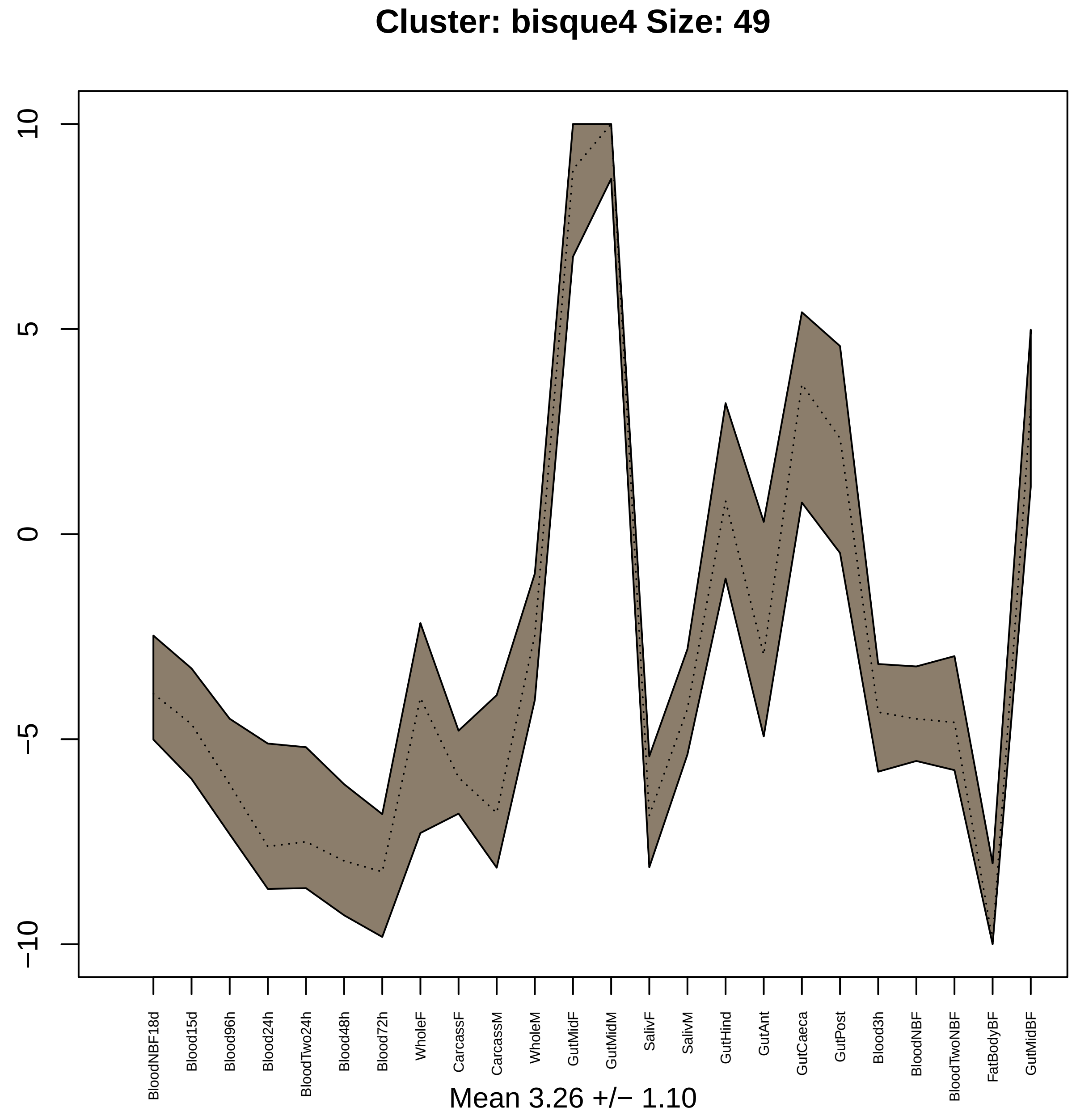

Cluster: bisque4 Size: 49
10
5
0
−5
−10
BloodNBF18d
Blood15d
Blood96h
Blood24h
BloodTwo24h
Blood48h
Blood72h
WholeF
CarcassF
CarcassM
WholeM
GutMidF
GutMidM
SalivF
SalivM
GutHind
GutAnt
GutCaeca
GutPost
Blood3h
BloodNBF
BloodTwoNBF
FatBodyBF
GutMidBF
Mean 3.26 +/− 1.10

### Cluster: lightcyan1 Size: 53

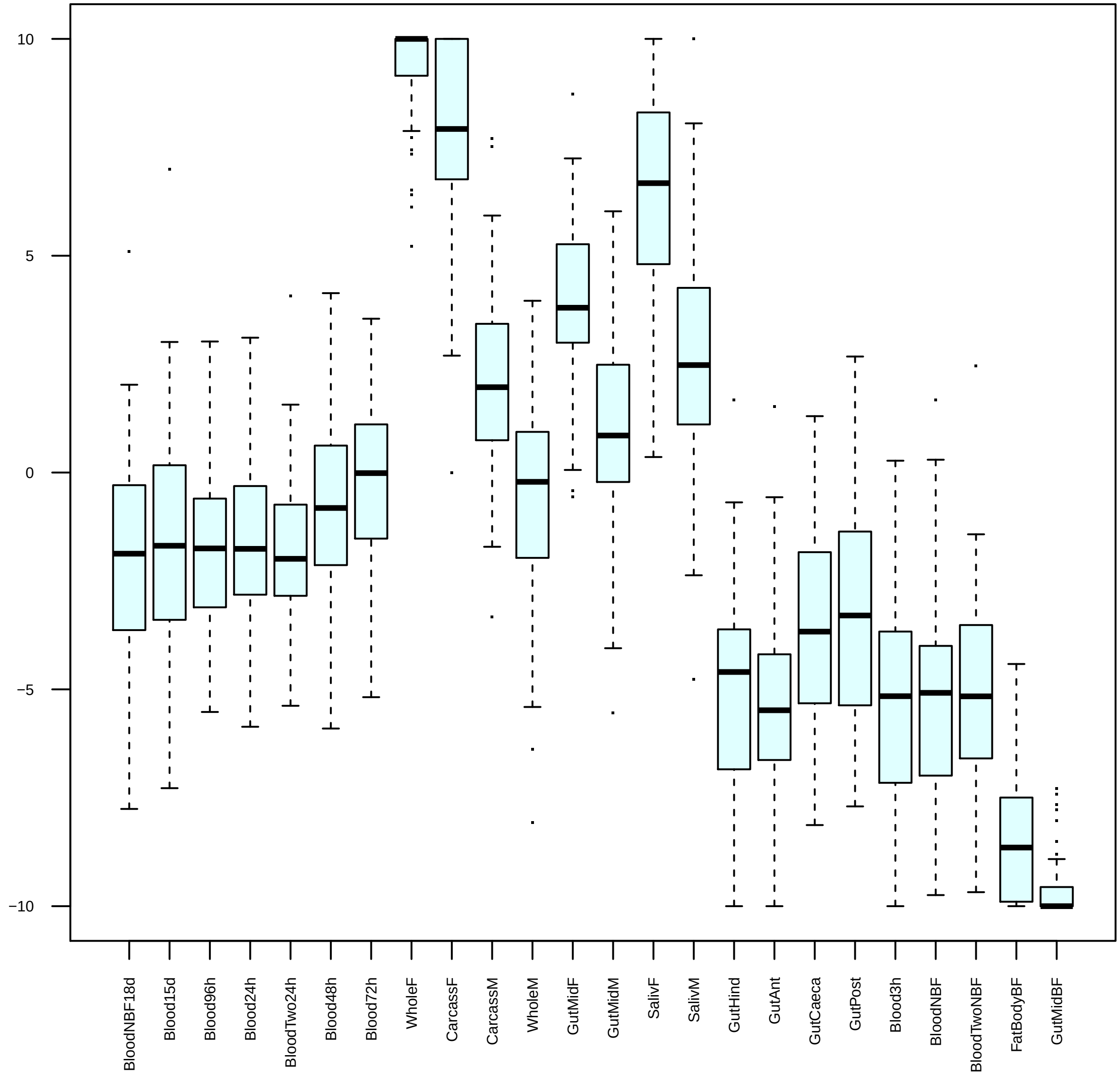

|  | GO.ID | BPCluster: lightcyan1 Size: 53 | Annotated | Significant | Expected | Rank in ClassicF | Weight01F | ClassicF |
|---|---|---|---|---|---|---|---|---|
| *4* | GO:0042254 | ribosome biogenesis | 64 | 10 | 0.37 | 1 | 0.00036 | 1.4e−12 |
| *5* | GO:0006378 | mRNA polyadenylation | 13 | 2 | 0.08 | 27 | 0.00245 | 0.0025 |
| *6* | GO:0006379 | mRNA cleavage | 13 | 2 | 0.08 | 28 | 0.00245 | 0.0025 |

| | GO.ID | MFCluster: lightcyan1 Size: 53 | Annotated | Significant | Expected | Rank in ClassicF | Weight01F | ClassicF |
|---|---|---|---|---|---|---|---|---|
| *2* | GO:0003729 | mRNA binding | 99 | 4 | 0.56 | 6 | 0.011 | 0.00230 |
| *4* | GO:0003723 | RNA binding | 307 | 8 | 1.75 | 2 | 0.022 | 0.00028 |
| *6* | GO:0008026 | ATP−dependent helicase activity | 76 | 4 | 0.43 | 3 | 0.028 | 0.00086 |
| *18* | GO:0016887 | ATPase activity | 233 | 5 | 1.33 | 12 | 0.191 | 0.00980 |
| *19* | GO:0005488 | binding | 4627 | 35 | 26.32 | 7 | 0.193 | 0.00234 |

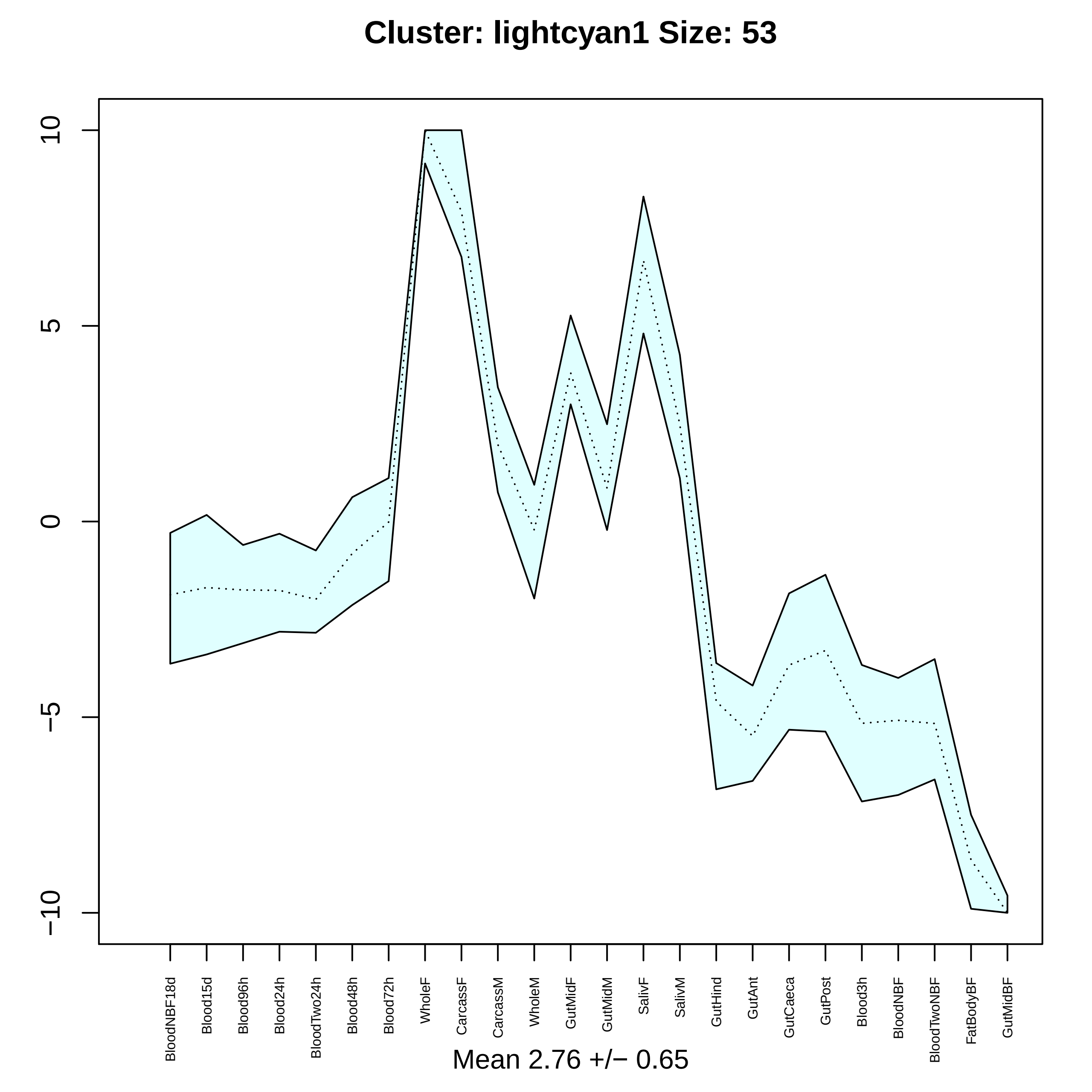
Cluster: lightcyan1 Size: 53
10
5
0
−5
−10
BloodNBF18d
Blood15d
Blood96h
Blood24h
BloodTwo24h
Blood48h
Blood72h
WholeF
CarcassF
CarcassM
WholeM
GutMidF
GutMidM
SalivF
SalivM
GutHind
GutAnt
GutCaeca
GutPost
Blood3h
BloodNBF
BloodTwoNBF
FatBodyBF
GutMidBF
Mean 2.76 +/− 0.65

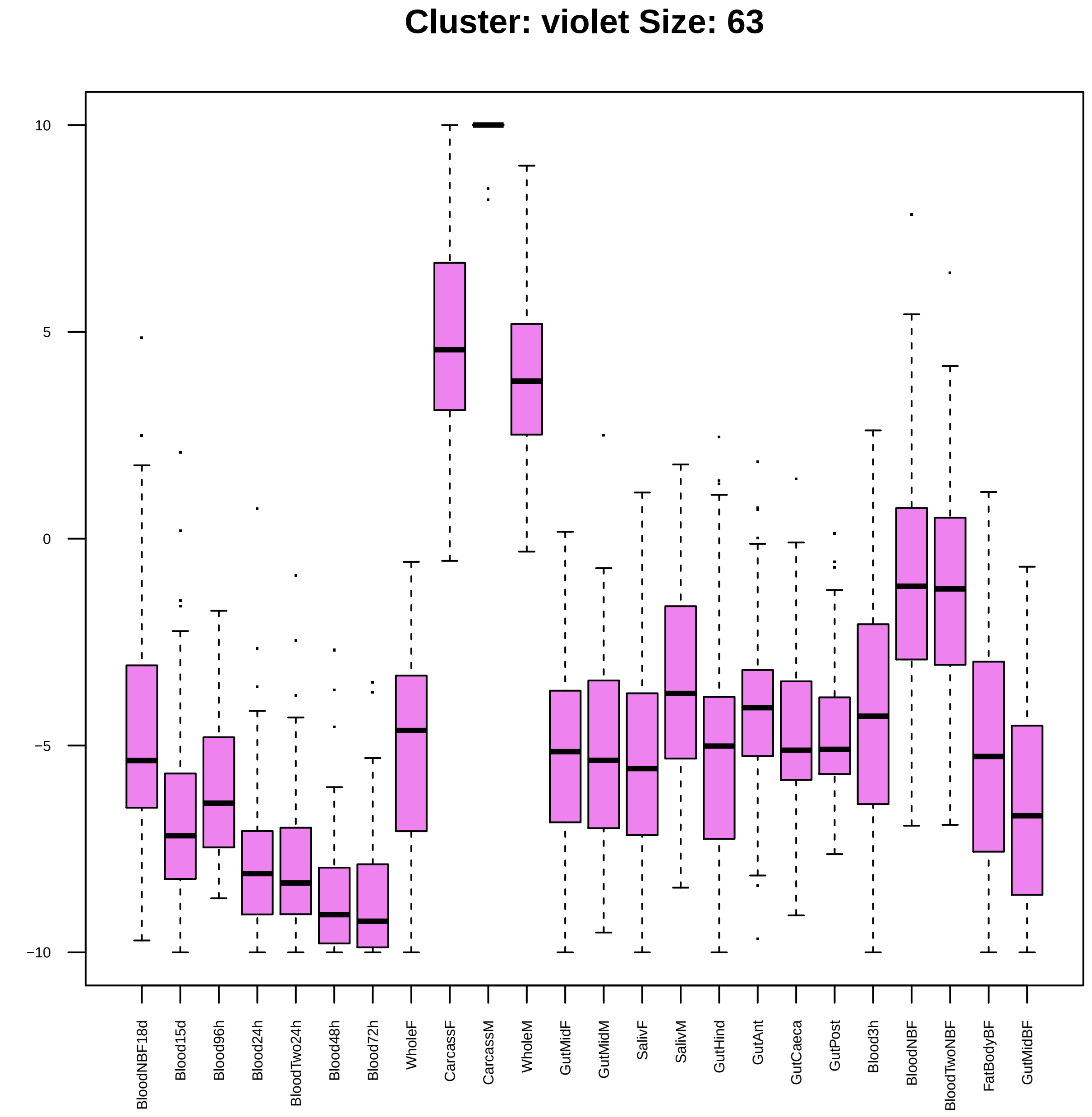
Cluster: violet Size: 63
10
5
0
−5
−10
BloodNBF18d
Blood15d
Blood96h
Blood24h
BloodTwo24h
Blood48h
Blood72h
WholeF
CarcassF
CarcassM
WholeM
GutMidF
GutMidM
SalivF
SalivM
GutHind
GutAnt
GutCaeca
GutPost
Blood3h
BloodNBF
BloodTwoNBF
FatBodyBF
GutMidBF

| | GO.ID | BPCluster: violet Size: 63 | Annotated | Significant | Expected | Rank in ClassicF | Weight01F | ClassicF |
|---|---|---|---|---|---|---|---|---|
| *1* | GO:0042246 | tissue regeneration | 10 | 2 | 0.06 | 1 | 0.0014 | 0.0014 |
| *2* | GO:0007155 | cell adhesion | 136 | 4 | 0.77 | 3 | 0.0069 | 0.0069 |
| *3* | GO:0071805 | potassium ion transmembrane transport | 23 | 2 | 0.13 | 5 | 0.0073 | 0.0073 |

| | GO.ID | MFCluster: violet Size: 63 | Annotated | Significant | Expected | Rank in ClassicF | Weight01F | ClassicF |
|---|---|---|---|---|---|---|---|---|
| *1* | GO:0005267 | potassium channel activity | 25 | 2 | 0.12 | 11 | 0.0068 | 0.00677 |
| *2* | GO:0005215 | transporter activity | 548 | 9 | 2.74 | 7 | 0.0112 | 0.00116 |
| *12* | GO:0022839 | ion gated channel activity | 87 | 4 | 0.43 | 5 | 0.1135 | 0.00087 |

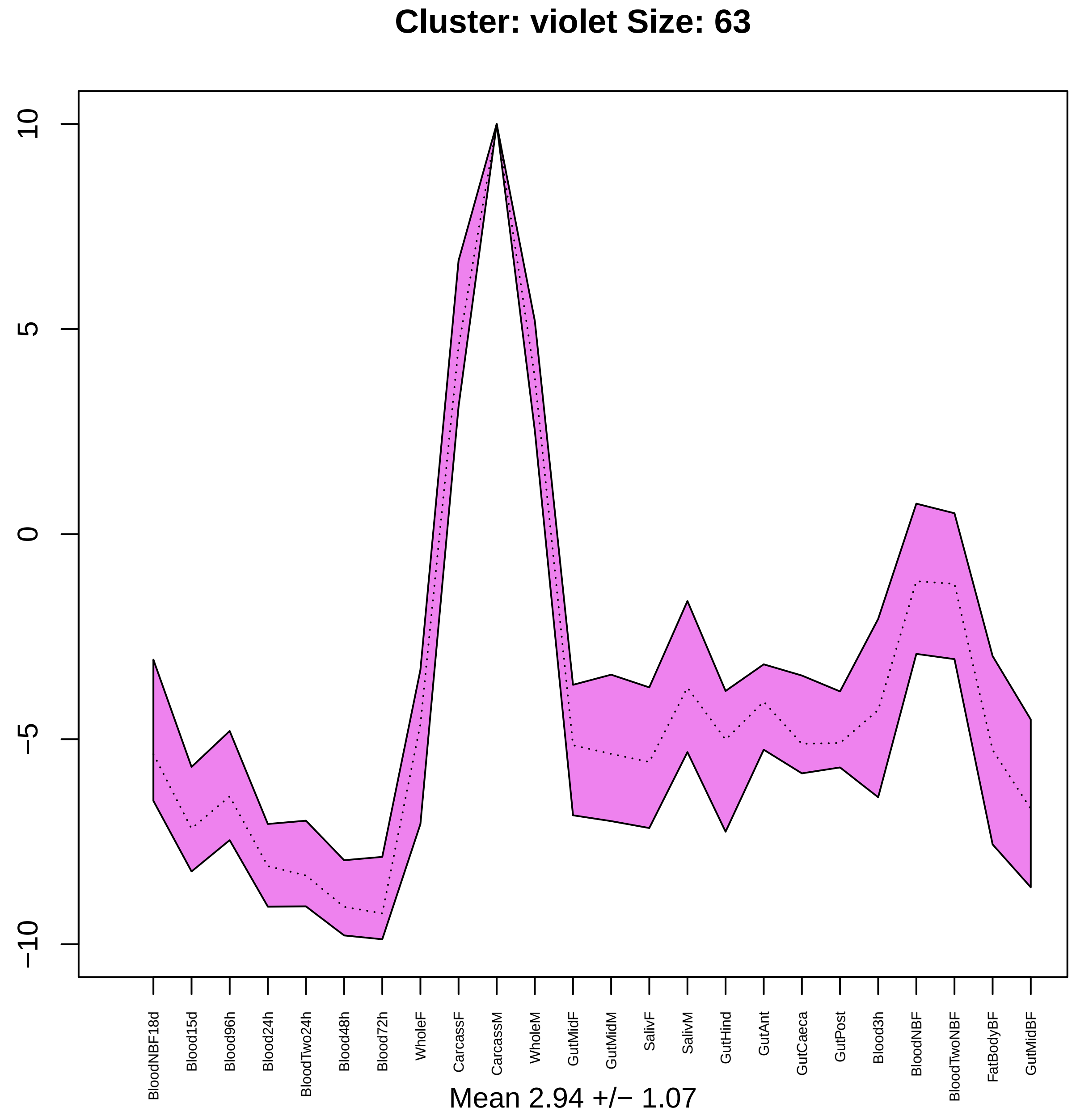

Cluster: violet Size: 63
10
5
0
−5
−10
BloodNBF18d
Blood15d
Blood96h
Blood24h
BloodTwo24h
Blood48h
Blood72h
WholeF
CarcassF
CarcassM
WholeM
GutMidF
GutMidM
SalivF
SalivM
GutHind
GutAnt
GutCaeca
GutPost
Blood3h
BloodNBF
BloodTwoNBF
FatBodyBF
GutMidBF
Mean 2.94 +/− 1.07

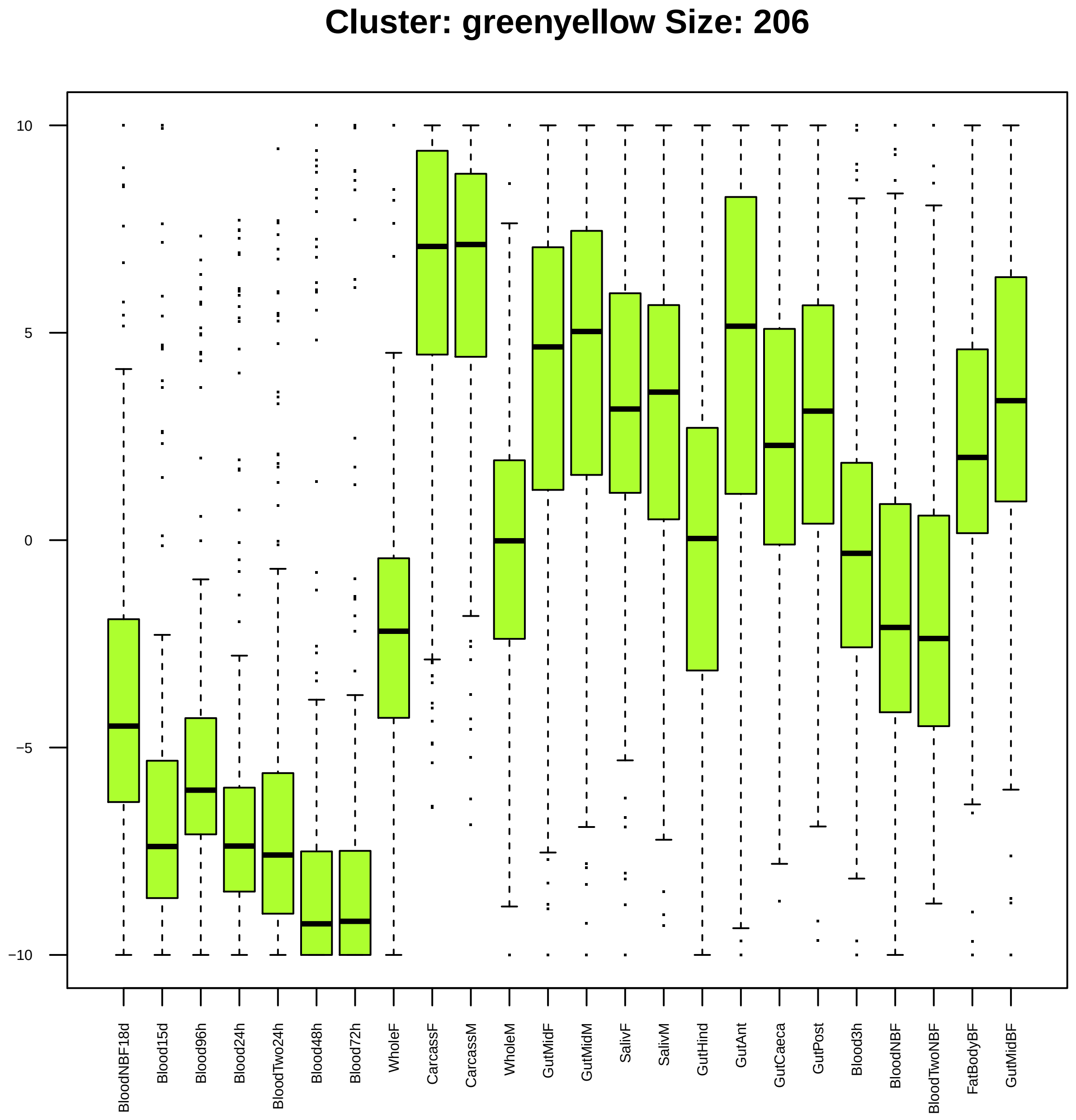

Cluster: greenyellow Size: 206
10
5
0
−5
−10
BloodNBF18d
Blood15d
Blood96h
Blood24h
BloodTwo24h
Blood48h
Blood72h
WholeF
CarcassF
CarcassM
WholeM
GutMidF
GutMidM
SalivF
SalivM
GutHind
GutAnt
GutCaeca
GutPost
Blood3h
BloodNBF
BloodTwoNBF
FatBodyBF
GutMidBF

| | GO.ID | BPCluster: greenyellow Size: 206 | Annotated | Significant | Expected | Rank in ClassicF | Weight01F | ClassicF |
|---|---|---|---|---|---|---|---|---|
| *27* | GO:0007606 | sensory perception of chemical stimulus | 149 | 8 | 2.62 | 1 | 0.097 | 0.0045 |

| | GO.ID | MFCluster: greenyellow Size: 206 | Annotated | Significant | Expected | Rank in ClassicF | Weight01F | ClassicF |
|---|---|---|---|---|---|---|---|---|
| 1 | GO:0042626 | ATPase activity, coupled to transmembran... | 73 | 5 | 1.42 | 11 | 0.0032 | 0.01343 |
| 2 | GO:0003705 | transcription factor activity, RNA polym... | 32 | 4 | 0.62 | 6 | 0.0032 | 0.00321 |
| 16 | GO:0043565 | sequence−specific DNA binding | 212 | 10 | 4.12 | 8 | 0.0916 | 0.00806 |
| 17 | GO:0019199 | transmembrane receptor protein kinase ac... | 15 | 3 | 0.29 | 4 | 0.0923 | 0.00275 |
| 18 | GO:0003700 | DNA−binding transcription factor activit... | 282 | 13 | 5.48 | 5 | 0.0943 | 0.00319 |
| 25 | GO:0004888 | transmembrane signaling receptor activit... | 255 | 14 | 4.95 | 1 | 0.1530 | 0.00041 |

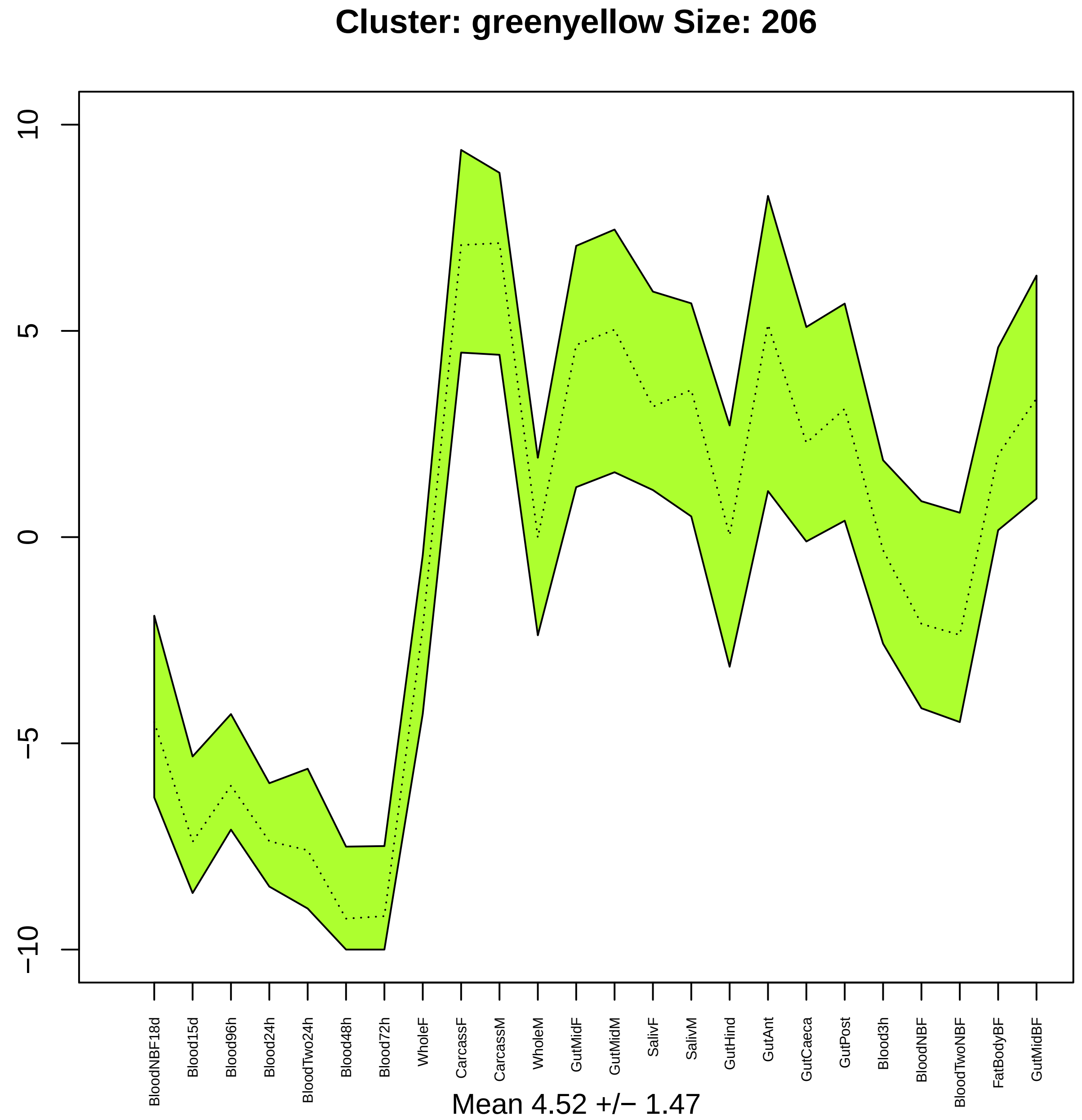

Cluster: greenyellow Size: 206
10
5
0
−5
−10
BloodNBF18d
Blood15d
Blood96h
Blood24h
BloodTwo24h
Blood48h
Blood72h
WholeF
CarcassF
CarcassM
WholeM
GutMidF
GutMidM
SalivF
SalivM
GutHind
GutAnt
GutCaeca
GutPost
Blood3h
BloodNBF
BloodTwoNBF
FatBodyBF
GutMidBF
Mean 4.52 +/− 1.47

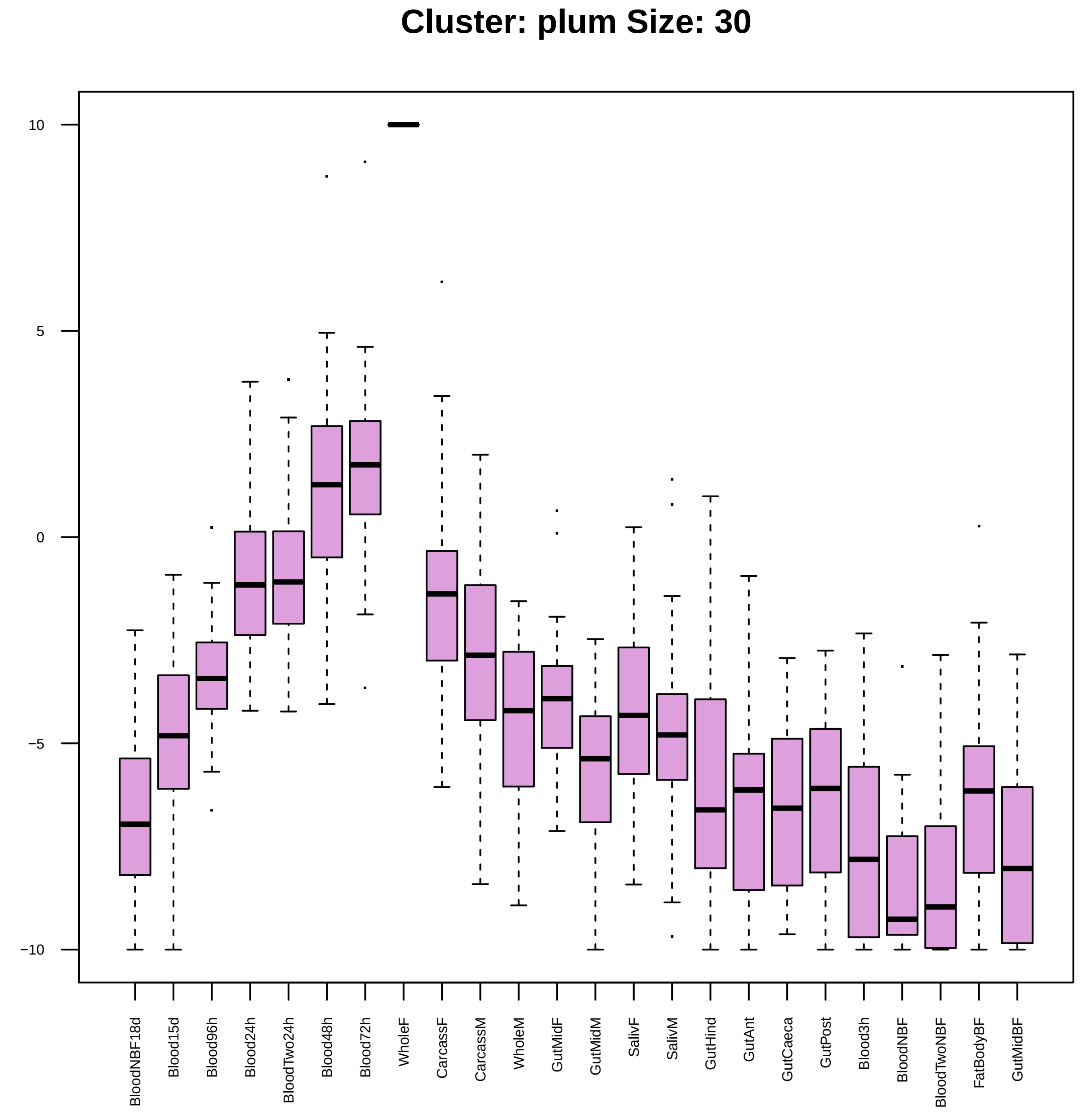

Cluster: plum Size: 30
10
5
0
−5
−10
BloodNBF18d
Blood15d
Blood96h
Blood24h
BloodTwo24h
Blood48h
Blood72h
WholeF
CarcassF
CarcassM
WholeM
GutMidF
GutMidM
SalivF
SalivM
GutHind
GutAnt
GutCaeca
GutPost
Blood3h
BloodNBF
BloodTwoNBF
FatBodyBF
GutMidBF

| | GO.ID | BPCluster: plum Size: 30 | Annotated | Significant | Expected | Rank in ClassicF | Weight01F | ClassicF |
|---|---|---|---|---|---|---|---|---|
| 1 | GO:0008152 | metabolic process | 4285 | 13 | 10.08 | 69 | 0.0030 | 0.0855 |
| 2 | GO:0007018 | microtubule−based movement | 65 | 2 | 0.15 | 4 | 0.0099 | 0.0099 |
| 3 | GO:0006913 | nucleocytoplasmic transport | 58 | 2 | 0.14 | 1 | 0.0176 | 0.0079 |

| | GO.ID | MFCluster: plum Size: 30 | Annotated | Significant | Expected | Rank in ClassicF | Weight01F | ClassicF |
|---|---|---|---|---|---|---|---|---|
| *1* | GO:0005524 | ATP binding | 592 | 7 | 1.48 | 5 | 0.00035 | 0.00035 |
| *2* | GO:0003777 | microtubule motor activity | 40 | 2 | 0.10 | 22 | 0.00434 | 0.00434 |
| *3* | GO:0008017 | microtubule binding | 55 | 2 | 0.14 | 24 | 0.00809 | 0.00809 |
| *4* | GO:0008094 | DNA−dependent ATPase activity | 33 | 2 | 0.08 | 20 | 0.02804 | 0.00297 |
| *6* | GO:0004386 | helicase activity | 103 | 3 | 0.26 | 16 | 0.03538 | 0.00198 |
| *15* | GO:0042623 | ATPase activity, coupled | 193 | 4 | 0.48 | 8 | 0.19992 | 0.00113 |
| *18* | GO:0000166 | nucleotide binding | 998 | 8 | 2.49 | 14 | 0.34904 | 0.00158 |
| *25* | GO:0016887 | ATPase activity | 233 | 4 | 0.58 | 17 | 1.00000 | 0.00228 |
| *26* | GO:1901265 | nucleoside phosphate binding | 998 | 8 | 2.49 | 15 | 1.00000 | 0.00158 |
| *27* | GO:0035639 | purine ribonucleoside triphosphate bindi... | 725 | 7 | 1.81 | 10 | 1.00000 | 0.00119 |
| *29* | GO:0032553 | ribonucleotide binding | 740 | 7 | 1.85 | 12 | 1.00000 | 0.00135 |
| *30* | GO:0032555 | purine ribonucleotide binding | 732 | 7 | 1.83 | 11 | 1.00000 | 0.00126 |

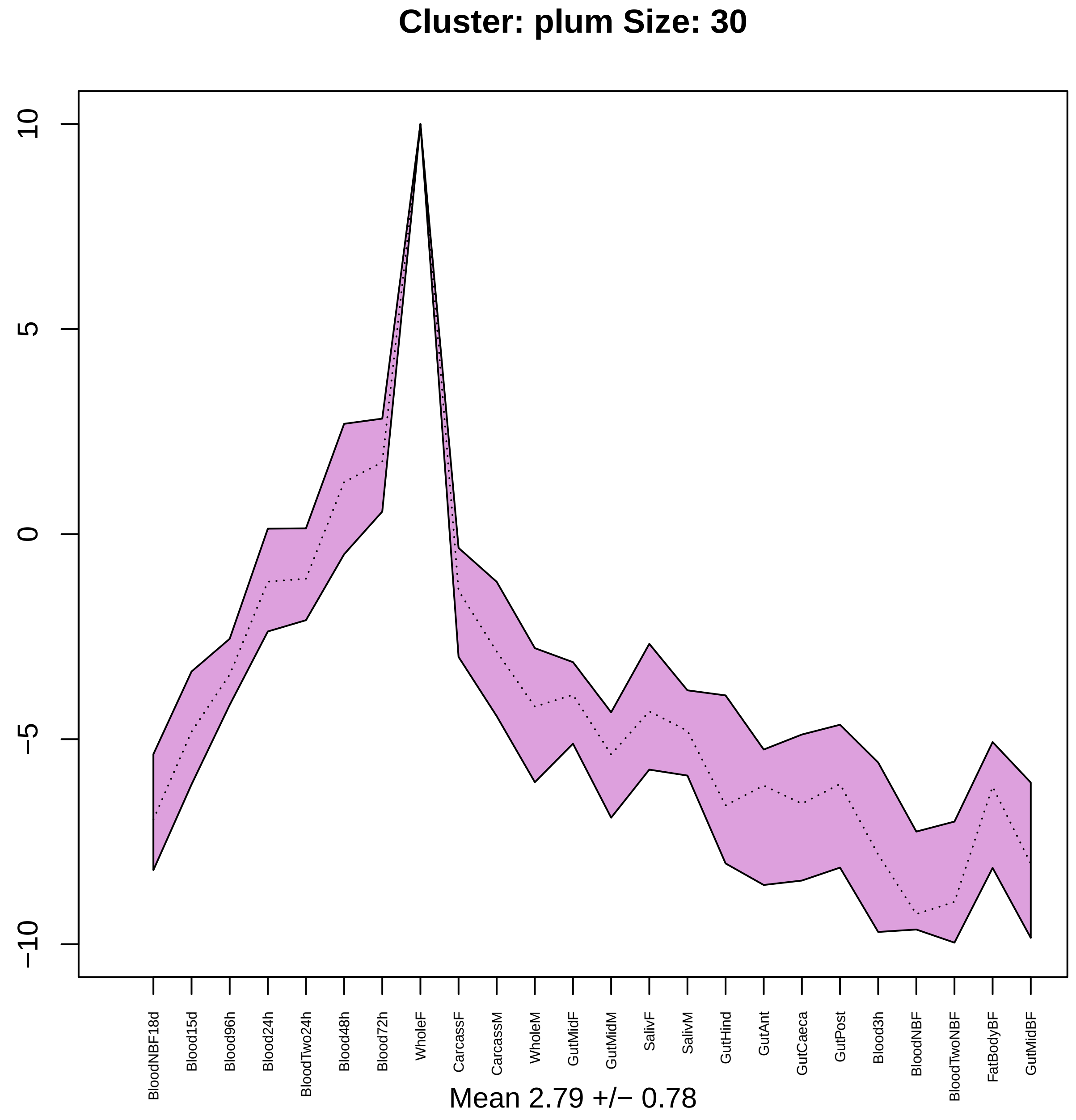
Cluster: plum Size: 30
10
5
0
−5
−10
BloodNBF18d
Blood15d
Blood96h
Blood24h
BloodTwo24h
Blood48h
Blood72h
WholeF
CarcassF
CarcassM
WholeM
GutMidF
GutMidM
SalivF
SalivM
GutHind
GutAnt
GutCaeca
GutPost
Blood3h
BloodNBF
BloodTwoNBF
FatBodyBF
GutMidBF
Mean 2.79 +/− 0.78

#### Cluster: grey60 Size: 104

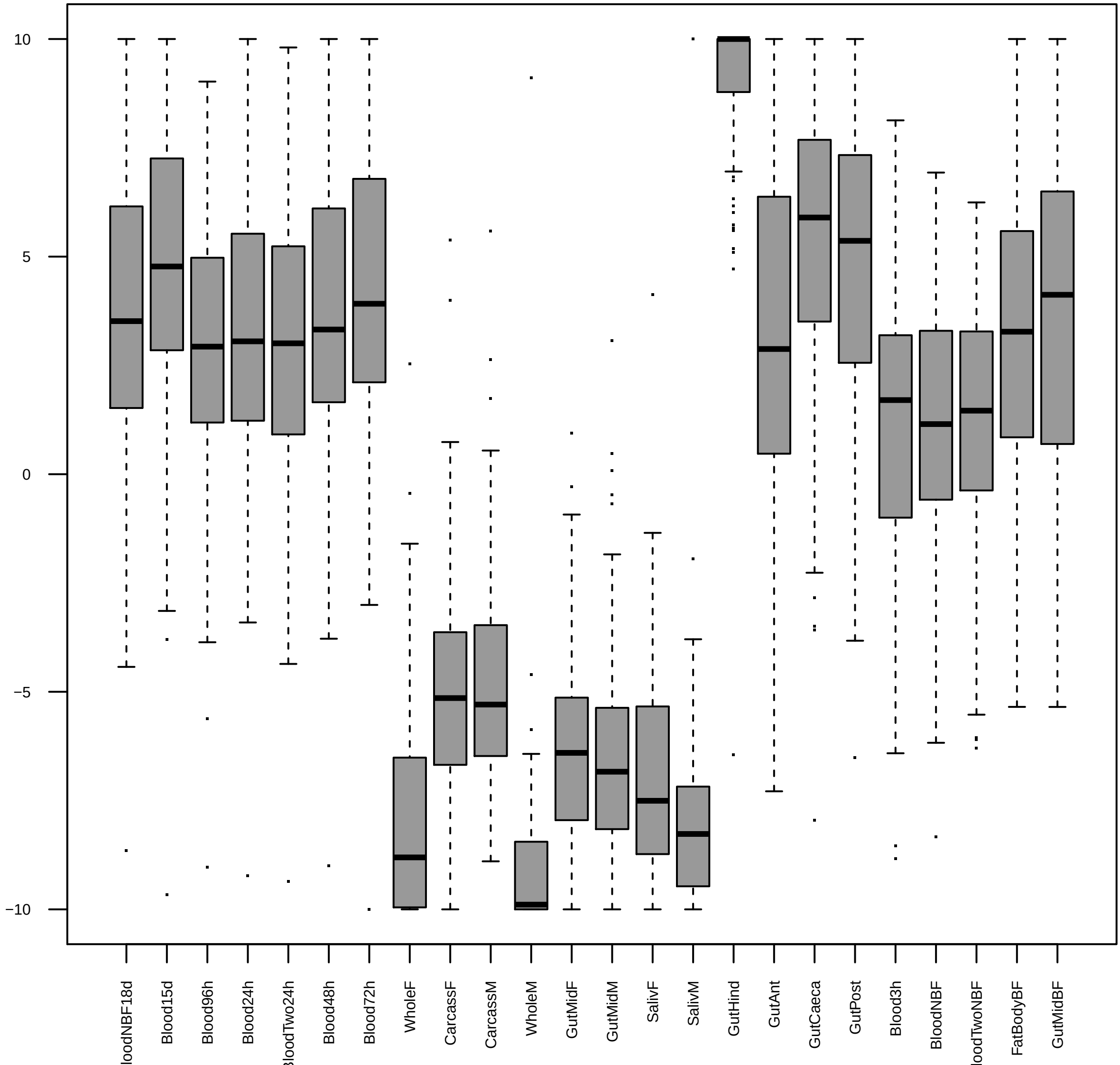

| | GO.ID | BPCluster: grey60 Size: 104 | Annotated | Significant | Expected | Rank in ClassicF | Weight01F | ClassicF |
|---|---|---|---|---|---|---|---|---|
| *2* | GO:0032482 | Rab protein signal transduction | 27 | 4 | 0.28 | 14 | 0.00014 | 0.00014 |
| *3* | GO:0007264 | small GTPase mediated signal transductio... | 153 | 9 | 1.56 | 4 | 0.00021 | 2.1e−05 |
| *4* | GO:0048488 | synaptic vesicle endocytosis | 38 | 5 | 0.39 | 9 | 0.00142 | 3.6e−05 |
| *5* | GO:0007269 | neurotransmitter secretion | 61 | 4 | 0.62 | 31 | 0.00332 | 0.00332 |
| *6* | GO:0043161 | proteasome−mediated ubiquitin−dependent ... | 37 | 3 | 0.38 | 39 | 0.00613 | 0.00613 |
| *7* | GO:0007040 | lysosome organization | 12 | 2 | 0.12 | 40 | 0.00632 | 0.00632 |
| *8* | GO:0008347 | glial cell migration | 13 | 2 | 0.13 | 44 | 0.00742 | 0.00742 |
| *9* | GO:0016183 | synaptic vesicle coating | 13 | 2 | 0.13 | 45 | 0.00742 | 0.00742 |
| *10* | GO:0006796 | phosphate−containing compound metabolic ... | 642 | 6 | 6.54 | 831 | 0.00802 | 0.65027 |
| *11* | GO:0051124 | synaptic growth at neuromuscular junctio... | 57 | 3 | 0.58 | 79 | 0.00969 | 0.02000 |
| *12* | GO:0000413 | protein peptidyl−prolyl isomerization | 15 | 2 | 0.15 | 49 | 0.00986 | 0.00986 |
| *16* | GO:0046907 | intracellular transport | 280 | 10 | 2.85 | 20 | 0.02062 | 0.00048 |
| *20* | GO:0072528 | pyrimidine−containing compound biosynthe... | 15 | 2 | 0.15 | 50 | 0.02987 | 0.00986 |

| | GO.ID | MFCluster: grey60 Size: 104 | Annotated | Significant | Expected | Rank in ClassicF | Weight01F | ClassicF |
|---|---|---|---|---|---|---|---|---|
| *3* | GO:0004298 | threonine−type endopeptidase activity | 14 | 2 | 0.14 | 13 | 0.0085 | 0.0085 |
| *8* | GO:0016776 | phosphotransferase activity, phosphate g... | 14 | 2 | 0.14 | 14 | 0.0394 | 0.0085 |

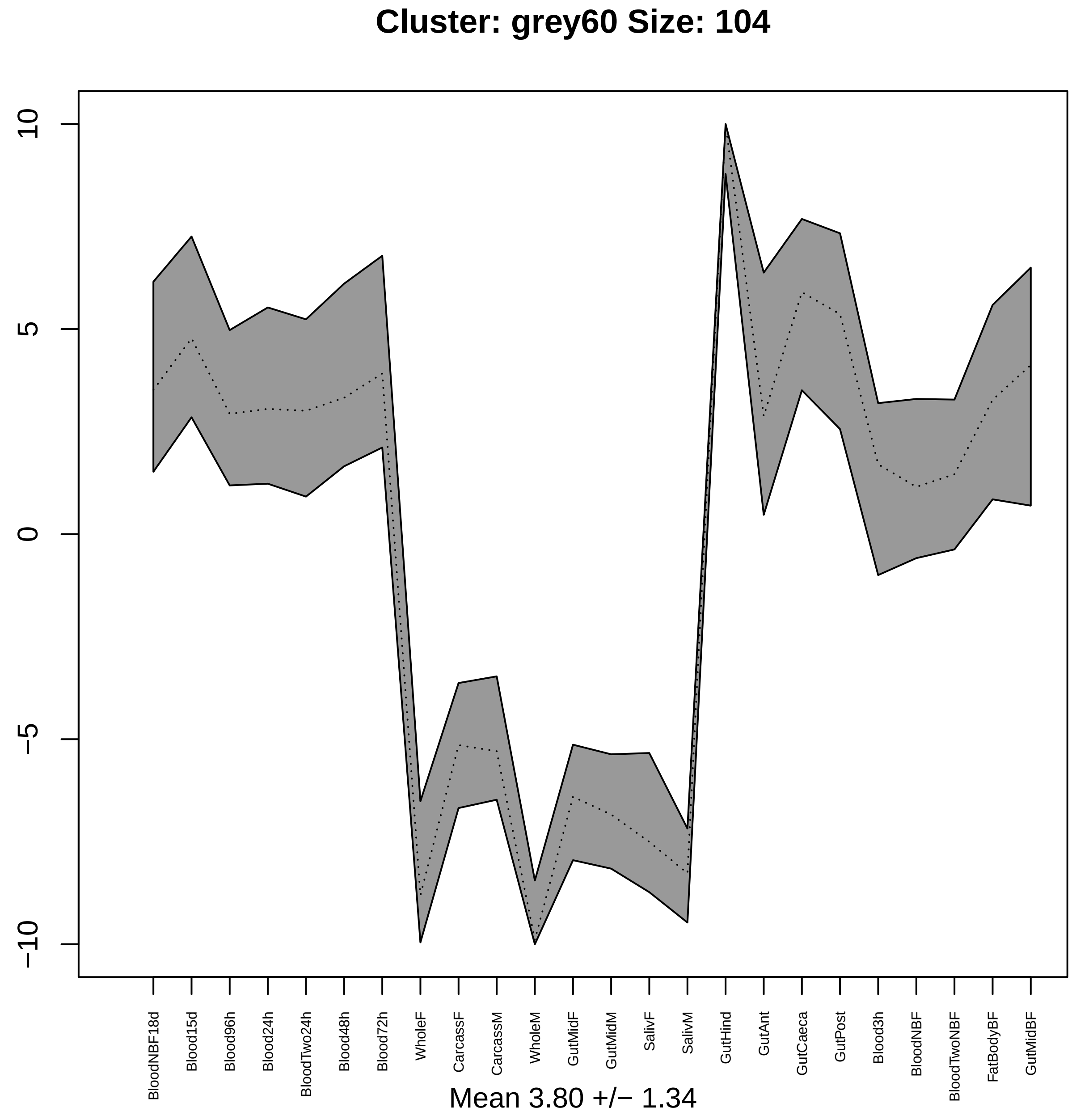
Cluster: grey60 Size: 104
10
5
0
−5
−10
BloodNBF18d
Blood15d
Blood96h
Blood24h
BloodTwo24h
Blood48h
Blood72h
WholeF
CarcassF
CarcassM
WholeM
GutMidF
GutMidM
SalivF
SalivM
GutHind
GutAnt
GutCaeca
GutPost
Blood3h
BloodNBF
BloodTwoNBF
FatBodyBF
GutMidBF
Mean 3.80 +/− 1.34

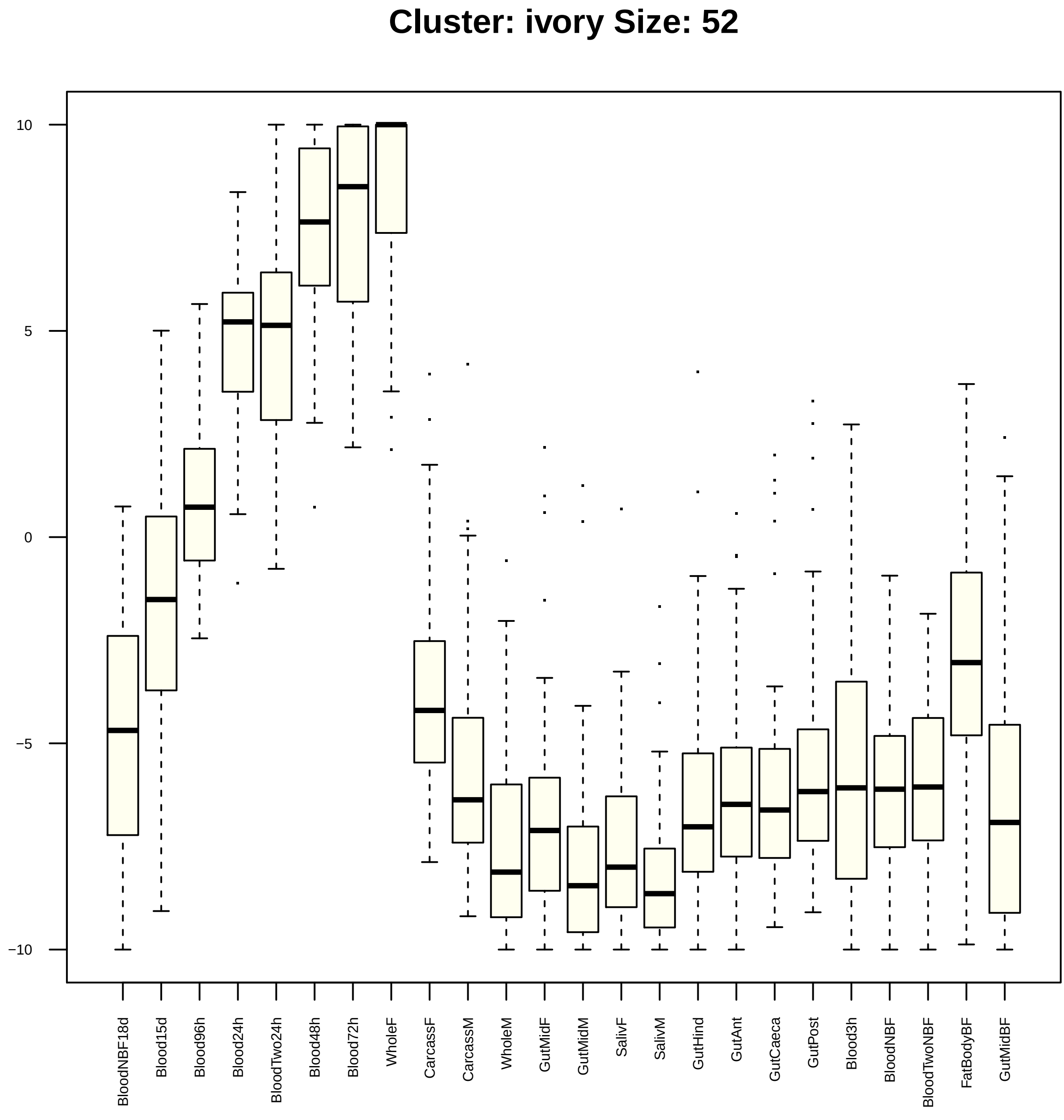

Cluster: ivory Size: 52
10
5
0
−5
−10
BloodNBF18d
Blood15d
Blood96h
Blood24h
BloodTwo24h
Blood48h
Blood72h
WholeF
CarcassF
CarcassM
WholeM
GutMidF
GutMidM
SalivF
SalivM
GutHind
GutAnt
GutCaeca
GutPost
Blood3h
BloodNBF
BloodTwoNBF
FatBodyBF
GutMidBF

| | GO.ID | BPCluster: ivory Size: 52 | Annotated | Significant | Expected | Rank in ClassicF | Weight01F | ClassicF |
|---|---|---|---|---|---|---|---|---|
| *8* | GO:2000112 | regulation of cellular macromolecule bio... | 560 | 7 | 2.81 | 156 | 0.00012 | 0.01878 |
| *9* | GO:0006261 | DNA−dependent DNA replication | 52 | 11 | 0.26 | 3 | 0.00013 | 3.9e−16 |
| *10* | GO:0007088 | regulation of mitotic nuclear division | 31 | 3 | 0.16 | 84 | 0.00047 | 0.00047 |
| *11* | GO:0042023 | DNA endoreduplication | 12 | 2 | 0.06 | 92 | 0.00156 | 0.00156 |
| *12* | GO:0007052 | mitotic spindle organization | 114 | 4 | 0.57 | 99 | 0.00237 | 0.00237 |
| *13* | GO:0007131 | reciprocal meiotic recombination | 15 | 2 | 0.08 | 101 | 0.00246 | 0.00246 |
| *14* | GO:0007099 | centriole replication | 15 | 2 | 0.08 | 102 | 0.00246 | 0.00246 |
| *15* | GO:0051276 | chromosome organization | 251 | 9 | 1.26 | 40 | 0.00260 | 2.5e−06 |
| *16* | GO:0006260 | DNA replication | 77 | 13 | 0.39 | 2 | 0.00301 | 1.2e−17 |
| *17* | GO:0045448 | mitotic cell cycle, embryonic | 17 | 2 | 0.09 | 109 | 0.00317 | 0.00317 |
| *18* | GO:0031570 | DNA integrity checkpoint | 54 | 4 | 0.27 | 73 | 0.00459 | 0.00014 |
| *19* | GO:0043065 | positive regulation of apoptotic process | 21 | 2 | 0.11 | 117 | 0.00483 | 0.00483 |
| *20* | GO:0008284 | positive regulation of cell proliferatio... | 23 | 2 | 0.12 | 122 | 0.00578 | 0.00578 |
| *21* | GO:0006342 | chromatin silencing | 26 | 2 | 0.13 | 127 | 0.00736 | 0.00736 |
| *22* | GO:0006277 | DNA amplification | 15 | 6 | 0.08 | 6 | 0.00847 | 4.7e−11 |
| *23* | GO:0051783 | regulation of nuclear division | 33 | 4 | 0.17 | 53 | 0.00912 | 1.9e−05 |
| *24* | GO:0006974 | cellular response to DNA damage stimulus | 168 | 10 | 0.84 | 10 | 0.00961 | 4.8e−09 |
| *25* | GO:0019730 | antimicrobial humoral response | 30 | 2 | 0.15 | 137 | 0.00972 | 0.00972 |
| *26* | GO:0051298 | centrosome duplication | 51 | 4 | 0.26 | 72 | 0.01226 | 0.00011 |
| *27* | GO:0045930 | negative regulation of mitotic cell cycl... | 67 | 4 | 0.34 | 79 | 0.01372 | 0.00032 |
| *28* | GO:0007259 | JAK−STAT cascade | 22 | 2 | 0.11 | 119 | 0.01456 | 0.00529 |
| *29* | GO:0044773 | mitotic DNA damage checkpoint | 47 | 3 | 0.24 | 93 | 0.01882 | 0.00160 |
| *30* | GO:0000086 | G2/M transition of mitotic cell cycle | 52 | 3 | 0.26 | 97 | 0.01883 | 0.00215 |

| | GO.ID | MFCluster: ivory Size: 52 | Annotated | Significant | Expected | Rank in ClassicF | Weight01F | ClassicF |
|---|---|---|---|---|---|---|---|---|
| *4* | GO:0005524 | ATP binding | 592 | 10 | 3.04 | 14 | 0.00059 | 0.00059 |
| *5* | GO:0003887 | DNA−directed DNA polymerase activity | 16 | 2 | 0.08 | 23 | 0.00294 | 0.00294 |
| *6* | GO:0016796 | exonuclease activity, active with either... | 18 | 2 | 0.09 | 28 | 0.00372 | 0.00372 |
| *7* | GO:0004536 | deoxyribonuclease activity | 19 | 2 | 0.10 | 29 | 0.00415 | 0.00415 |
| *9* | GO:0016462 | pyrophosphatase activity | 439 | 9 | 2.25 | 10 | 0.03787 | 0.00029 |
| *14* | GO:0000166 | nucleotide binding | 998 | 13 | 5.12 | 17 | 0.08237 | 0.00090 |
| *22* | GO:0017111 | nucleoside−triphosphatase activity | 430 | 8 | 2.21 | 19 | 0.17815 | 0.00125 |
| *26* | GO:0004386 | helicase activity | 103 | 5 | 0.53 | 9 | 0.28023 | 0.00016 |

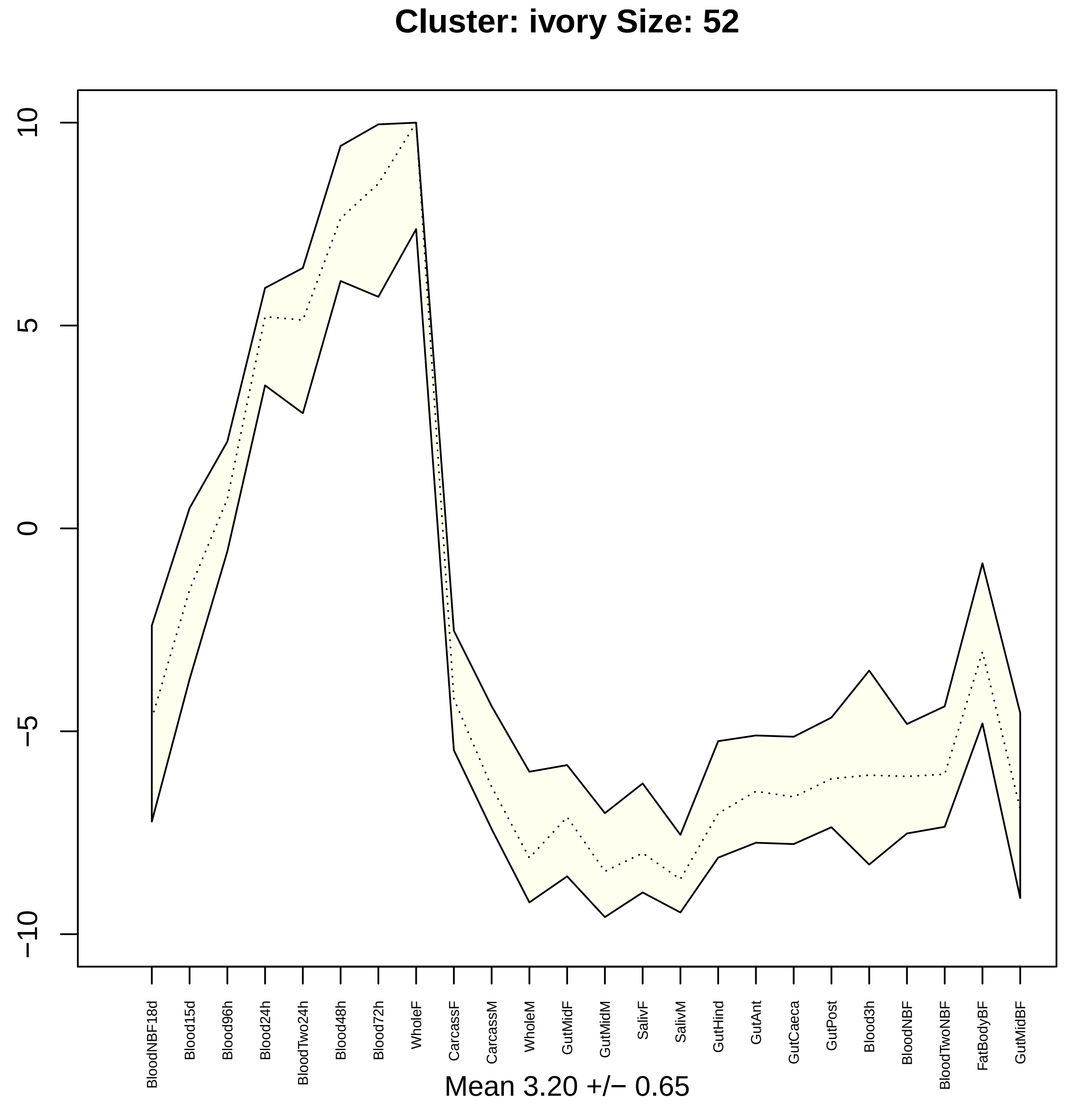

Cluster: ivory Size: 52
10
5
0
−5
−10
BloodNBF18d
Blood15d
Blood96h
Blood24h
BloodTwo24h
Blood48h
Blood72h
WholeF
CarcassF
CarcassM
WholeM
GutMidF
GutMidM
SalivF
SalivM
GutHind
GutAnt
GutCaeca
GutPost
Blood3h
BloodNBF
BloodTwoNBF
FatBodyBF
GutMidBF
Mean 3.20 +/− 0.65

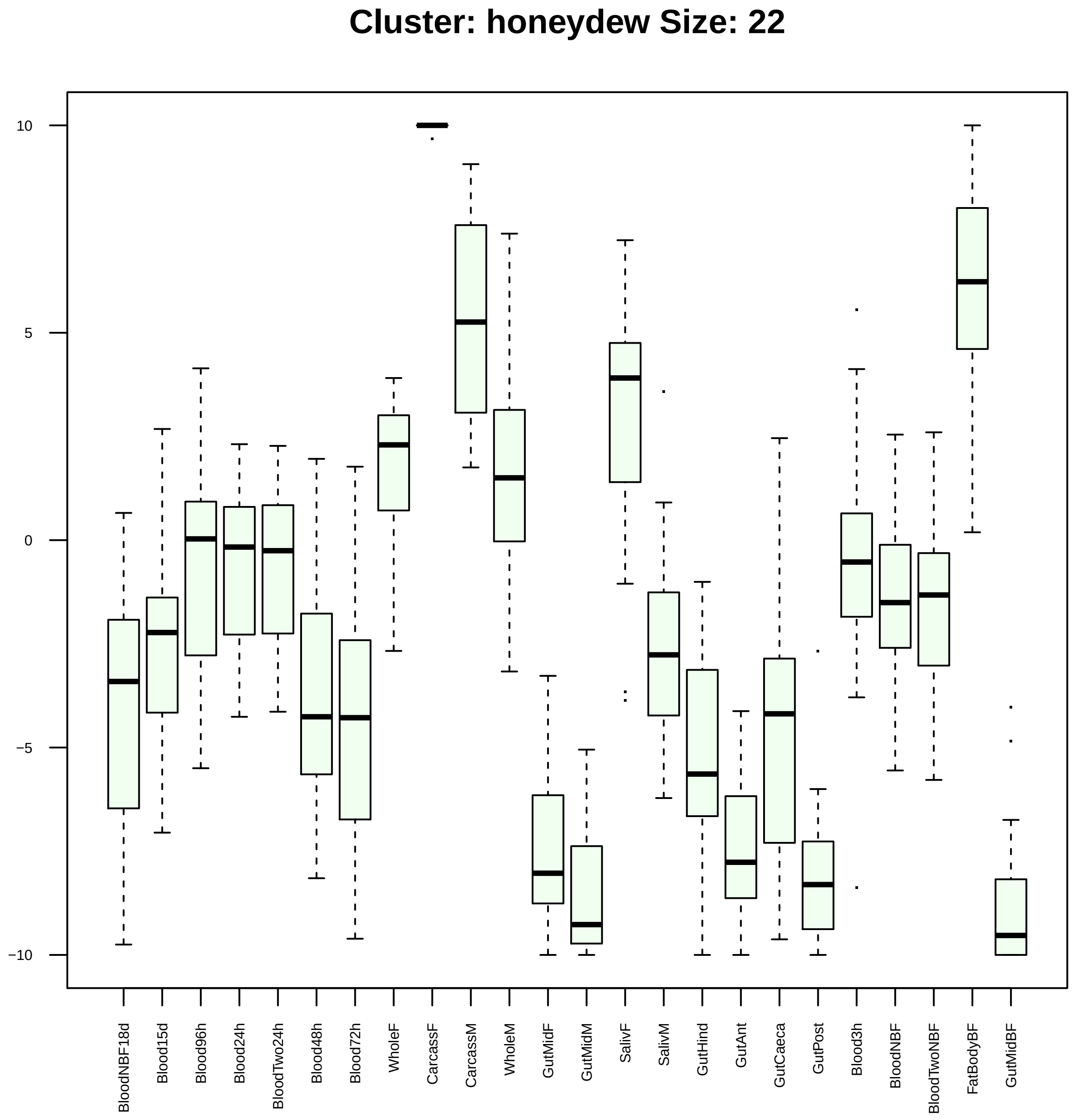

Cluster: honeydew Size: 22
10
5
0
−5
−10
BloodNBF18d
Blood15d
Blood96h
Blood24h
BloodTwo24h
Blood48h
Blood72h
WholeF
CarcassF
CarcassM
WholeM
GutMidF
GutMidM
SalivF
SalivM
GutHind
GutAnt
GutCaeca
GutPost
Blood3h
BloodNBF
BloodTwoNBF
FatBodyBF
GutMidBF

| | GO.ID | BPCluster: honeydew Size: 22 | Annotated | Significant | Expected | Rank in ClassicF | Weight01F | ClassicF |
|---|---|---|---|---|---|---|---|---|
| *2* | GO:0042135 | neurotransmitter catabolic process | 12 | 2 | 0.03 | 6 | 0.00043 | 0.00043 |
| *3* | GO:0009072 | aromatic amino acid family metabolic pro... | 14 | 2 | 0.04 | 11 | 0.00060 | 0.00060 |
| *4* | GO:0042737 | drug catabolic process | 48 | 3 | 0.13 | 5 | 0.00159 | 0.00025 |
| *5* | GO:1901606 | alpha−amino acid catabolic process | 27 | 2 | 0.07 | 15 | 0.00226 | 0.00226 |
| *6* | GO:0006807 | nitrogen compound metabolic process | 2967 | 13 | 7.91 | 23 | 0.00876 | 0.01199 |

| | GO.ID | MFCluster: honeydew Size: 22 | Annotated | Significant | Expected | Rank in ClassicF | Weight01F | ClassicF |
|---|---|---|---|---|---|---|---|---|
| 1 | GO:0016741 | transferase activity, transferring one−c... | 92 | 3 | 0.22 | 1 | 0.0084 | 0.0012 |

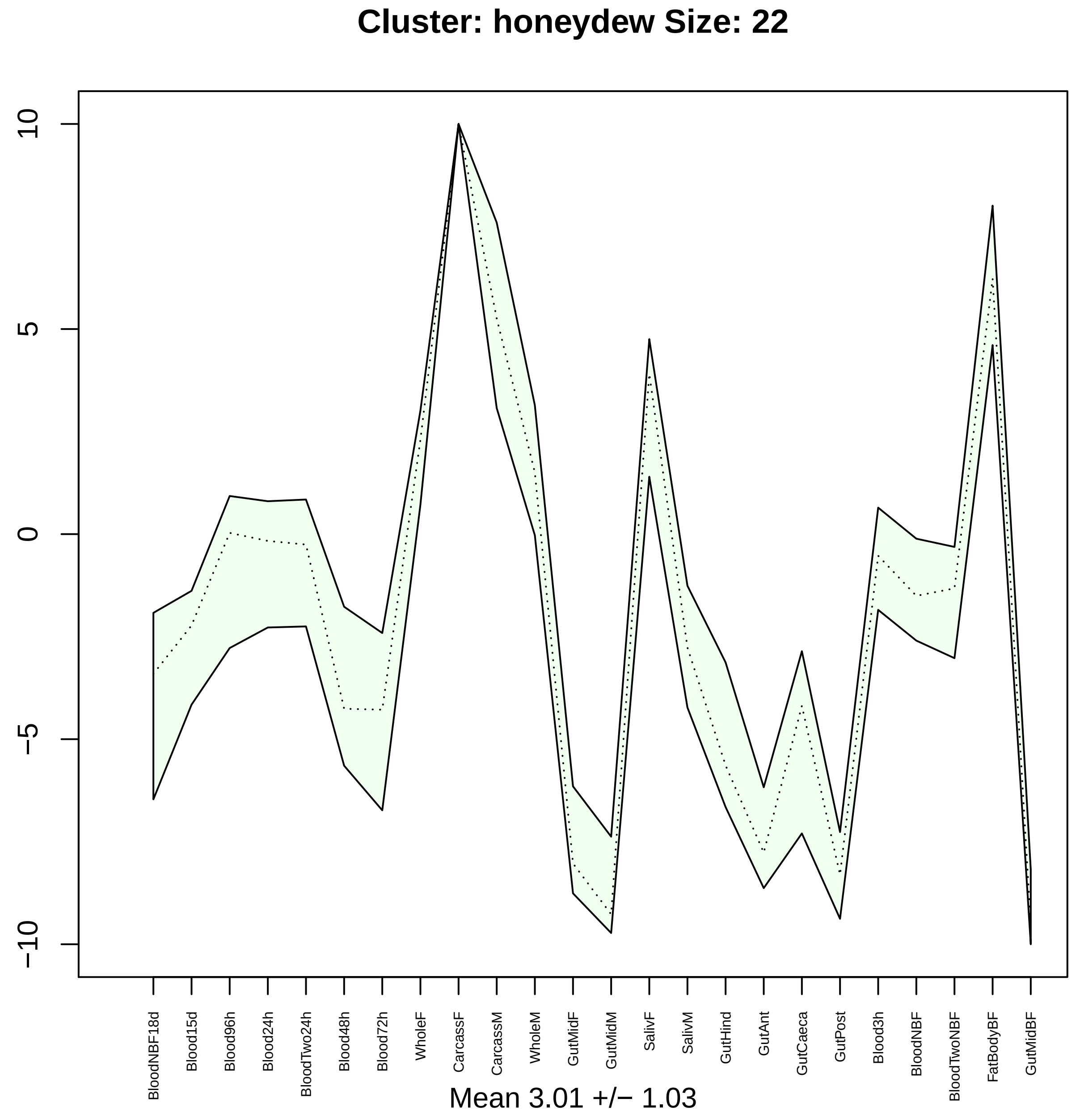
Cluster: honeydew Size: 22
10
5
0
−5
−10
BloodNBF18d
Blood15d
Blood96h
Blood24h
BloodTwo24h
Blood48h
Blood72h
WholeF
CarcassF
CarcassM
WholeM
GutMidF
GutMidM
SalivF
SalivM
GutHind
GutAnt
GutCaeca
GutPost
Blood3h
BloodNBF
BloodTwoNBF
FatBodyBF
GutMidBF
Mean 3.01 +/− 1.03

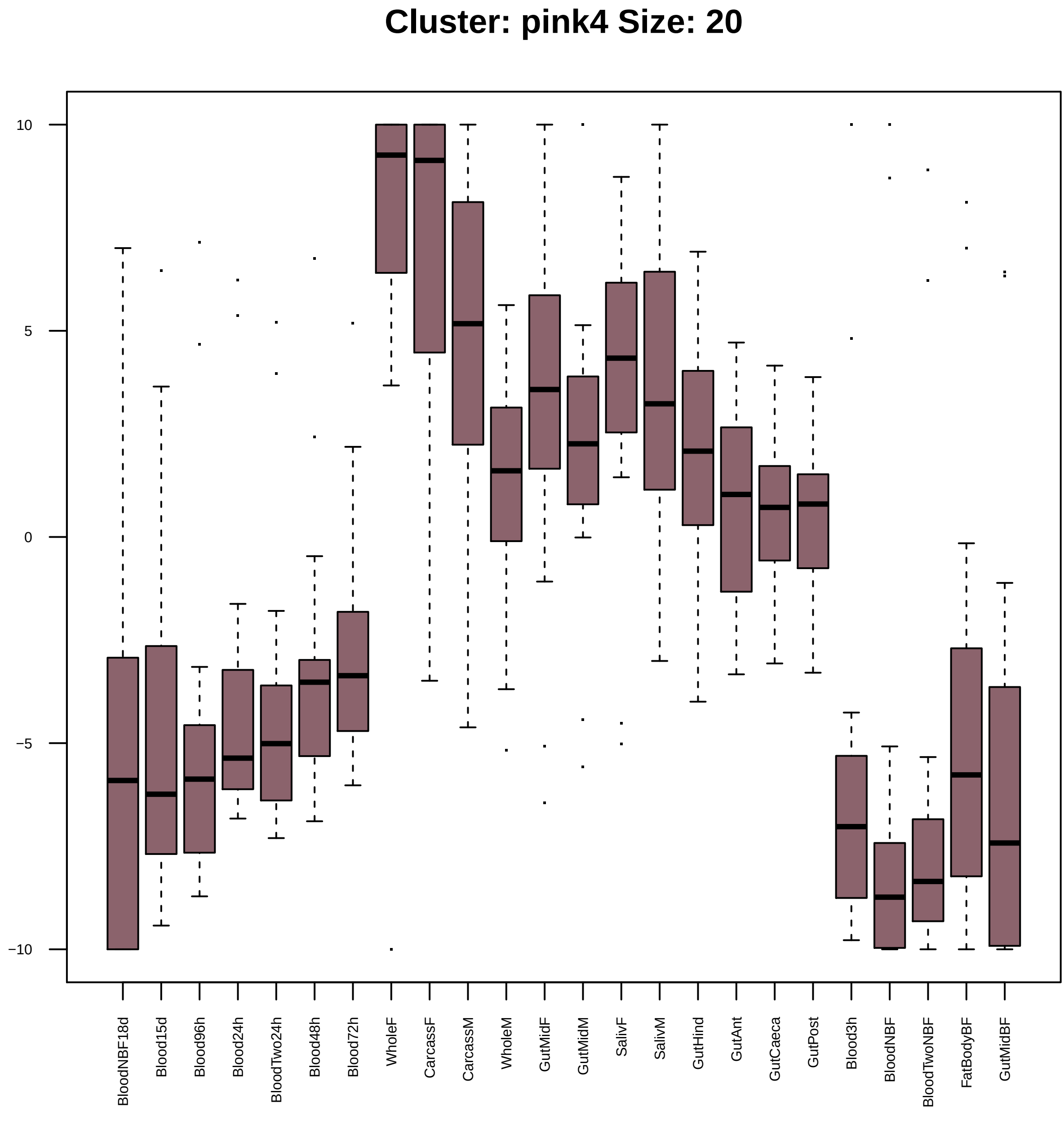
Cluster: pink4 Size: 20
10
5
0
−5
−10
BloodNBF18d
Blood15d
Blood96h
Blood24h
BloodTwo24h
Blood48h
Blood72h
WholeF
CarcassF
CarcassM
WholeM
GutMidF
GutMidM
SalivF
SalivM
GutHind
GutAnt
GutCaeca
GutPost
Blood3h
BloodNBF
BloodTwoNBF
FatBodyBF
GutMidBF

| | GO.ID | BPCluster: pink4 Size: 20 | Annotated | Significant | Expected | Rank in ClassicF | Weight01F | ClassicF |
|---|---|---|---|---|---|---|---|---|
| *13* | GO:0051298 | centrosome duplication | 51 | 2 | 0.14 | 2 | 0.087 | 0.0079 |

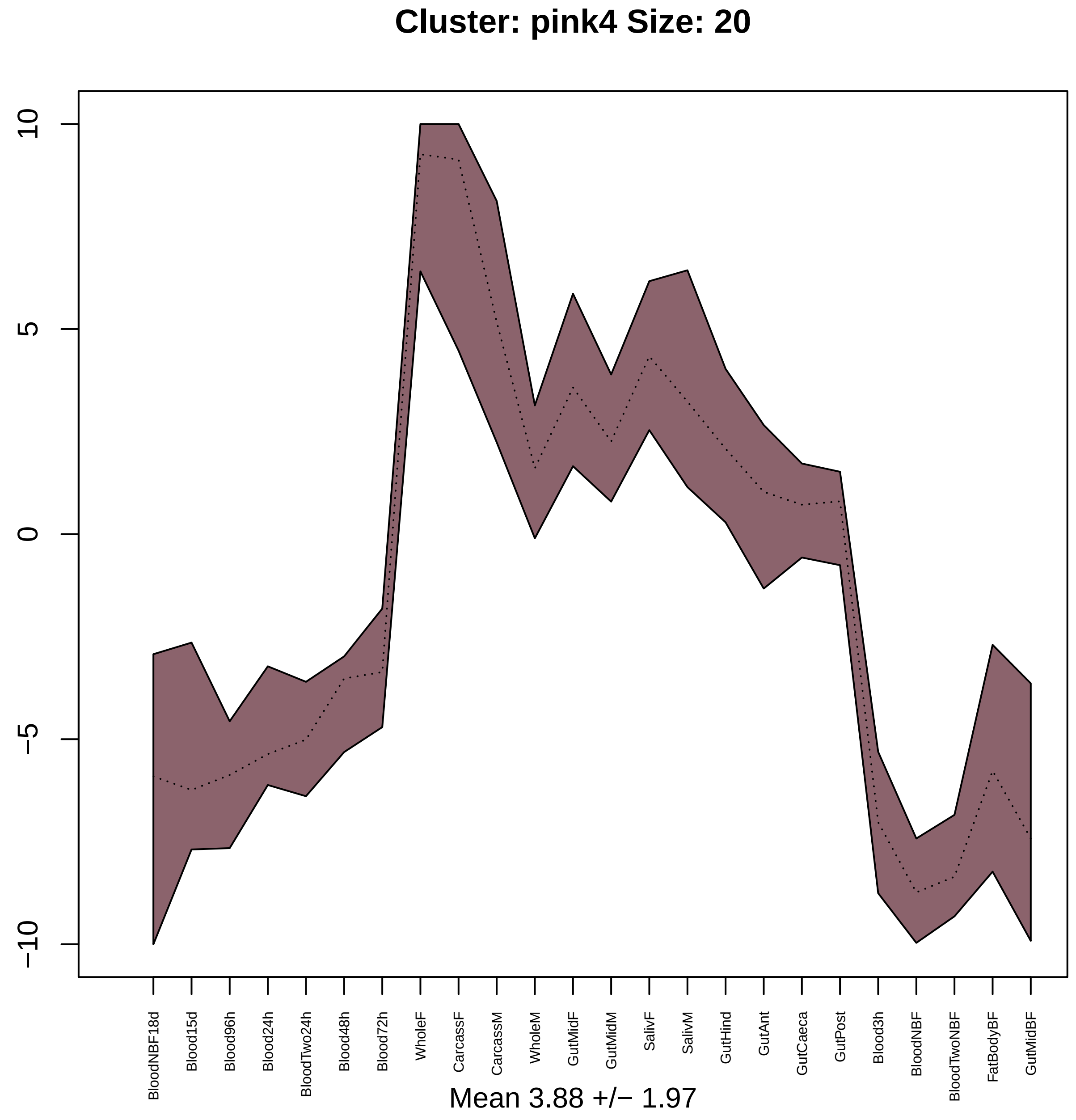
Cluster: pink4 Size: 20
10
5
0
−5
−10
BloodNBF18d
Blood15d
Blood96h
Blood24h
BloodTwo24h
Blood48h
Blood72h
WholeF
CarcassF
CarcassM
WholeM
GutMidF
GutMidM
SalivF
SalivM
GutHind
GutAnt
GutCaeca
GutPost
Blood3h
BloodNBF
BloodTwoNBF
FatBodyBF
GutMidBF
Mean 3.88 +/− 1.97

#### Cluster: plum2 Size: 47

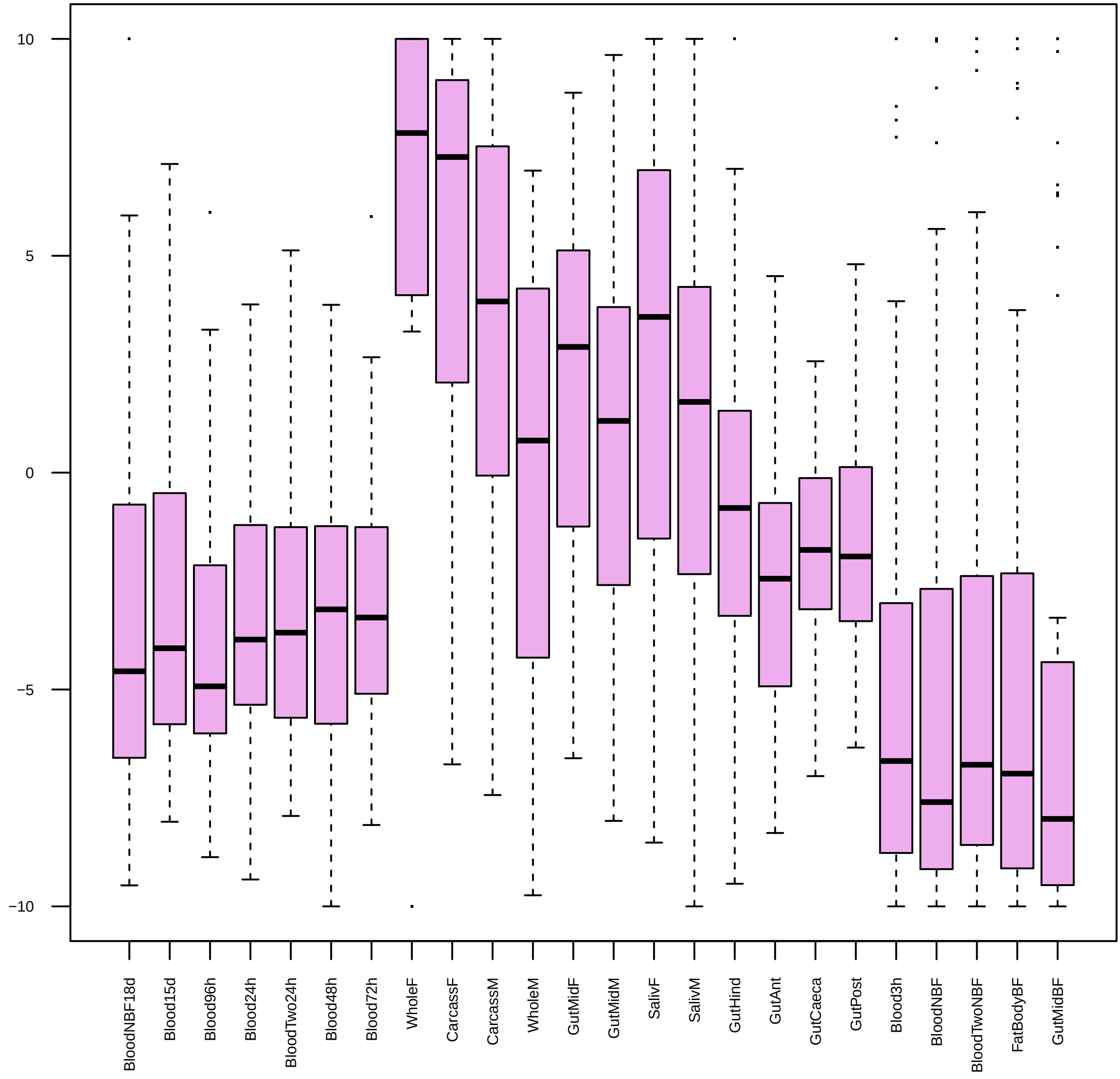

| | GO.ID | BPCluster: plum2 Size: 47 | Annotated | Significant | Expected | Rank in ClassicF | Weight01F | ClassicF |
|---|---|---|---|---|---|---|---|---|
| *2* | GO:0006479 | protein methylation | 34 | 2 | 0.13 | 3 | 0.029 | 0.0071 |

| | GO.ID | MFCluster: plum2 Size: 47 | Annotated | Significant | Expected | Rank in ClassicF | Weight01F | ClassicF |
|---|---|---|---|---|---|---|---|---|
| *1* | GO:0016417 | S−acyltransferase activity | 11 | 2 | 0.05 | 1 | 0.0012 | 0.0012 |
| *3* | GO:0008276 | protein methyltransferase activity | 25 | 2 | 0.12 | 3 | 0.0372 | 0.0064 |
| *9* | GO:0008170 | N−methyltransferase activity | 25 | 2 | 0.12 | 4 | 0.0508 | 0.0064 |

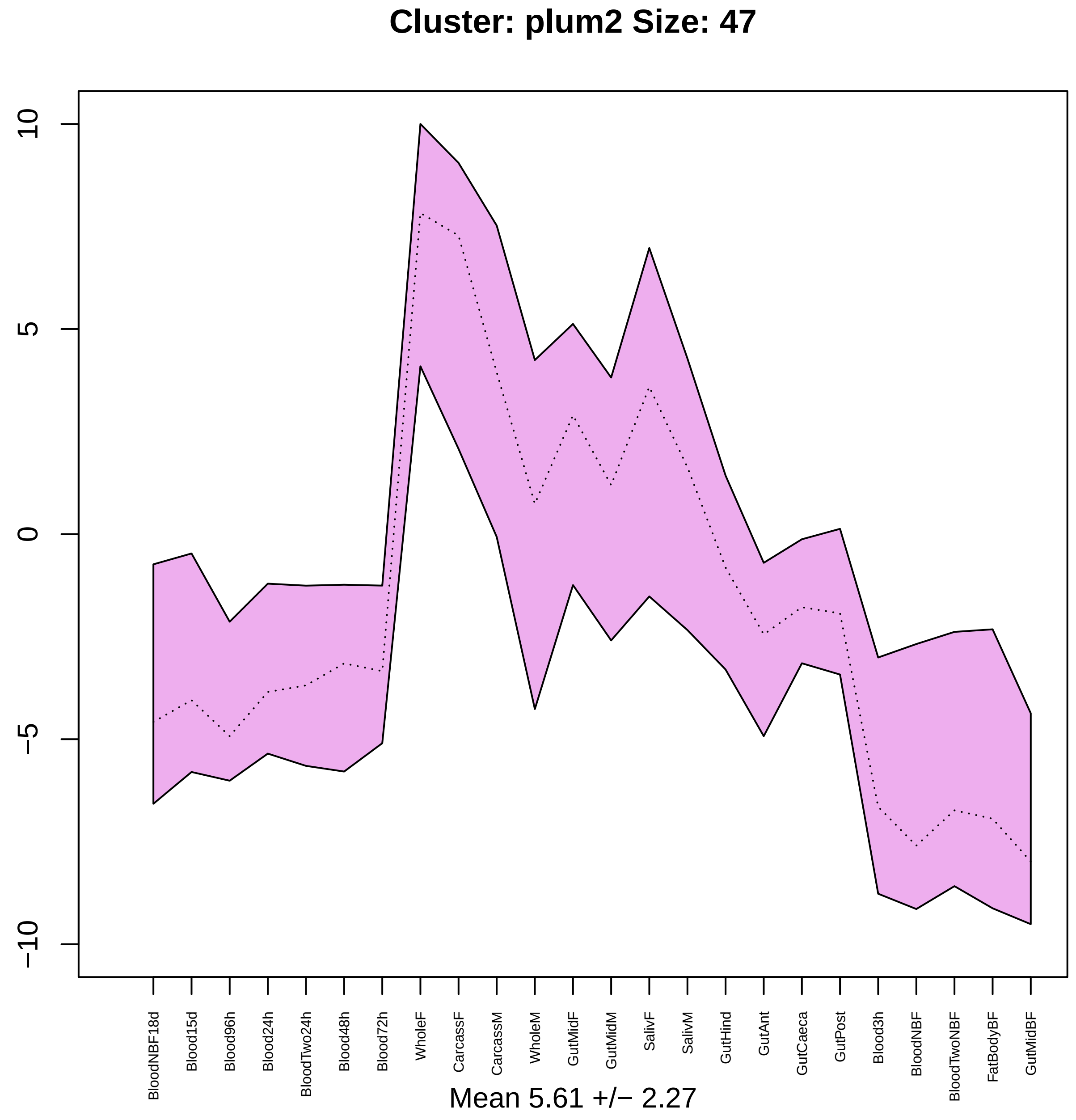

Cluster: plum2 Size: 47
10
5
0
−5
−10
BloodNBF18d
Blood15d
Blood96h
Blood24h
BloodTwo24h
Blood48h
Blood72h
WholeF
CarcassF
CarcassM
WholeM
GutMidF
GutMidM
SalivF
SalivM
GutHind
GutAnt
GutCaeca
GutPost
Blood3h
BloodNBF
BloodTwoNBF
FatBodyBF
GutMidBF
Mean 5.61 +/− 2.27

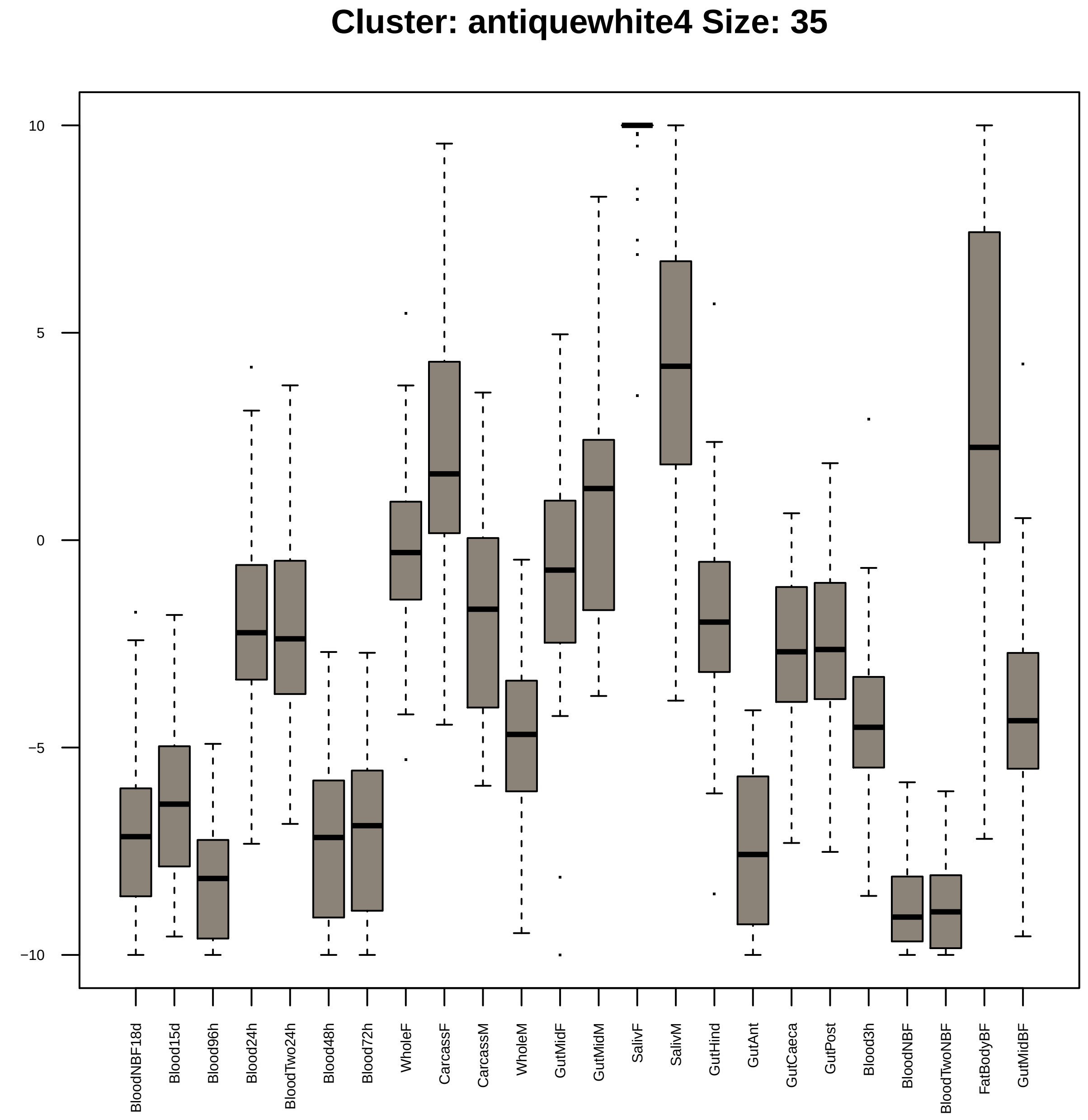
Cluster: antiquewhite4 Size: 35
10
5
0
−5
−10
BloodNBF18d
Blood15d
Blood96h
Blood24h
BloodTwo24h
Blood48h
Blood72h
WholeF
CarcassF
CarcassM
WholeM
GutMidF
GutMidM
SalivF
SalivM
GutHind
GutAnt
GutCaeca
GutPost
Blood3h
BloodNBF
BloodTwoNBF
FatBodyBF
GutMidBF

| | GO.ID | BPCluster: antiquewhite4 Size: 35 | Annotated | Significant | Expected | Rank in ClassicF | Weight01F | ClassicF |
|---|---|---|---|---|---|---|---|---|
| 3 | GO:0006662 | glycerol ether metabolic process | 11 | 2 | 0.05 | 10 | 0.0011 | 0.0011 |
| 4 | GO:0048193 | Golgi vesicle transport | 54 | 3 | 0.25 | 15 | 0.0018 | 0.0018 |
| 5 | GO:0045047 | protein targeting to ER | 11 | 2 | 0.05 | 11 | 0.0044 | 0.0011 |
| 6 | GO:0016485 | protein processing | 24 | 2 | 0.11 | 25 | 0.0052 | 0.0052 |
| 7 | GO:0070972 | protein localization to endoplasmic reti... | 13 | 3 | 0.06 | 8 | 0.0085 | 2.3e−05 |
| 10 | GO:0006612 | protein targeting to membrane | 14 | 2 | 0.06 | 14 | 0.0131 | 0.0018 |
| 16 | GO:0061024 | membrane organization | 94 | 3 | 0.43 | 29 | 0.0467 | 0.0086 |
| 28 | GO:0006886 | intracellular protein transport | 167 | 4 | 0.76 | 27 | 0.0920 | 0.0065 |

| | GO.ID | MFCluster: antiquewhite4 Size: 35 | Annotated | Significant | Expected | Rank in ClassicF | Weight01F | ClassicF |
|---|---|---|---|---|---|---|---|---|
| *2* | GO:0008194 | UDP−glycosyltransferase activity | 30 | 3 | 0.12 | 4 | 0.00020 | 0.00020 |
| *3* | GO:0005048 | signal sequence binding | 11 | 2 | 0.04 | 5 | 0.00078 | 0.00078 |
| *4* | GO:0043021 | ribonucleoprotein complex binding | 14 | 2 | 0.05 | 6 | 0.00129 | 0.00129 |
| *5* | GO:0015035 | protein disulfide oxidoreductase activit... | 14 | 2 | 0.05 | 7 | 0.00129 | 0.00129 |
| *6* | GO:0016860 | intramolecular oxidoreductase activity | 23 | 2 | 0.09 | 10 | 0.00350 | 0.00350 |

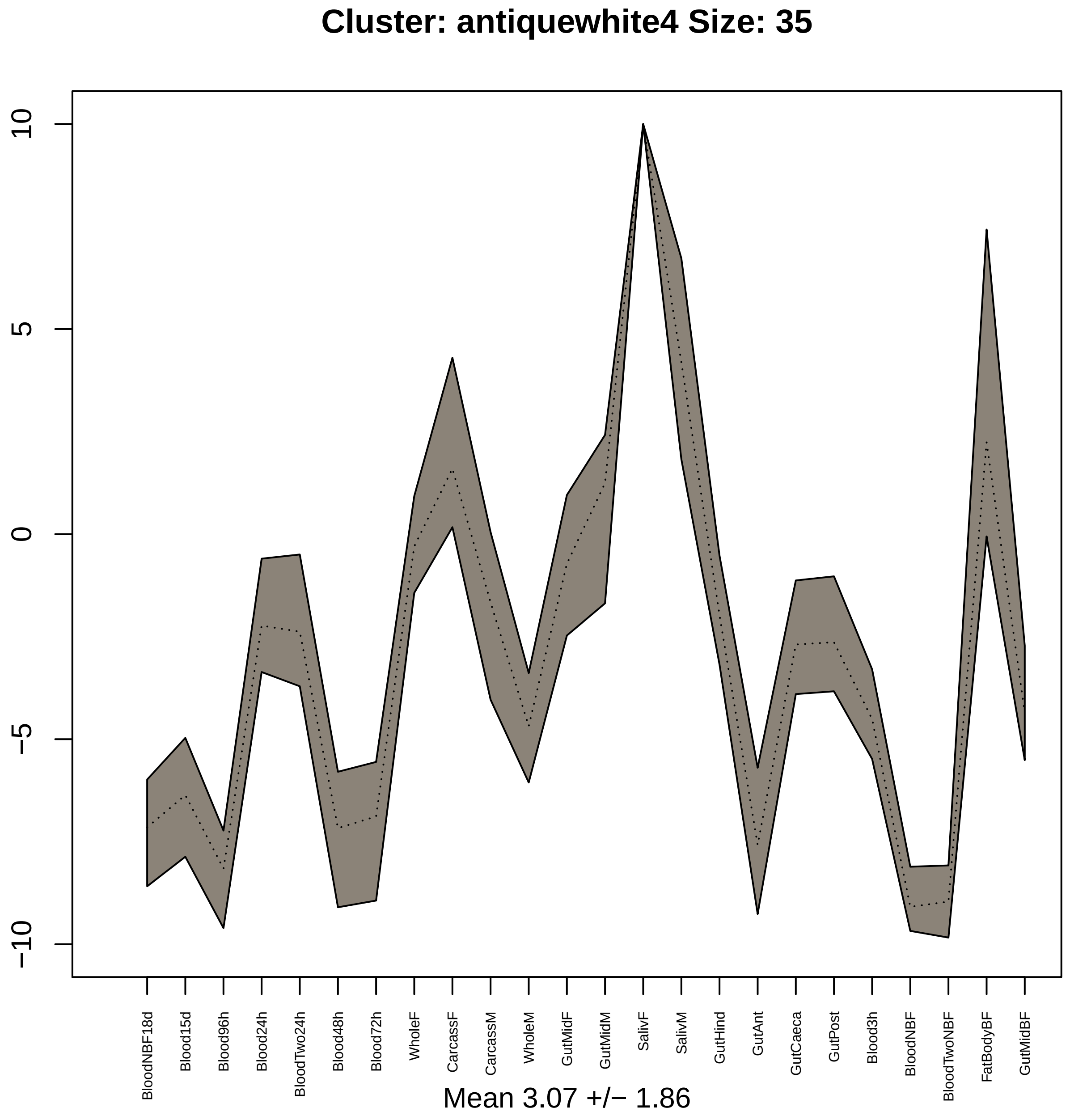
Cluster: antiquewhite4 Size: 35
10
5
0
−5
−10
BloodNBF18d
Blood15d
Blood96h
Blood24h
BloodTwo24h
Blood48h
Blood72h
WholeF
CarcassF
CarcassM
WholeM
GutMidF
GutMidM
SalivF
SalivM
GutHind
GutAnt
GutCaeca
GutPost
Blood3h
BloodNBF
BloodTwoNBF
FatBodyBF
GutMidBF
Mean 3.07 +/− 1.86

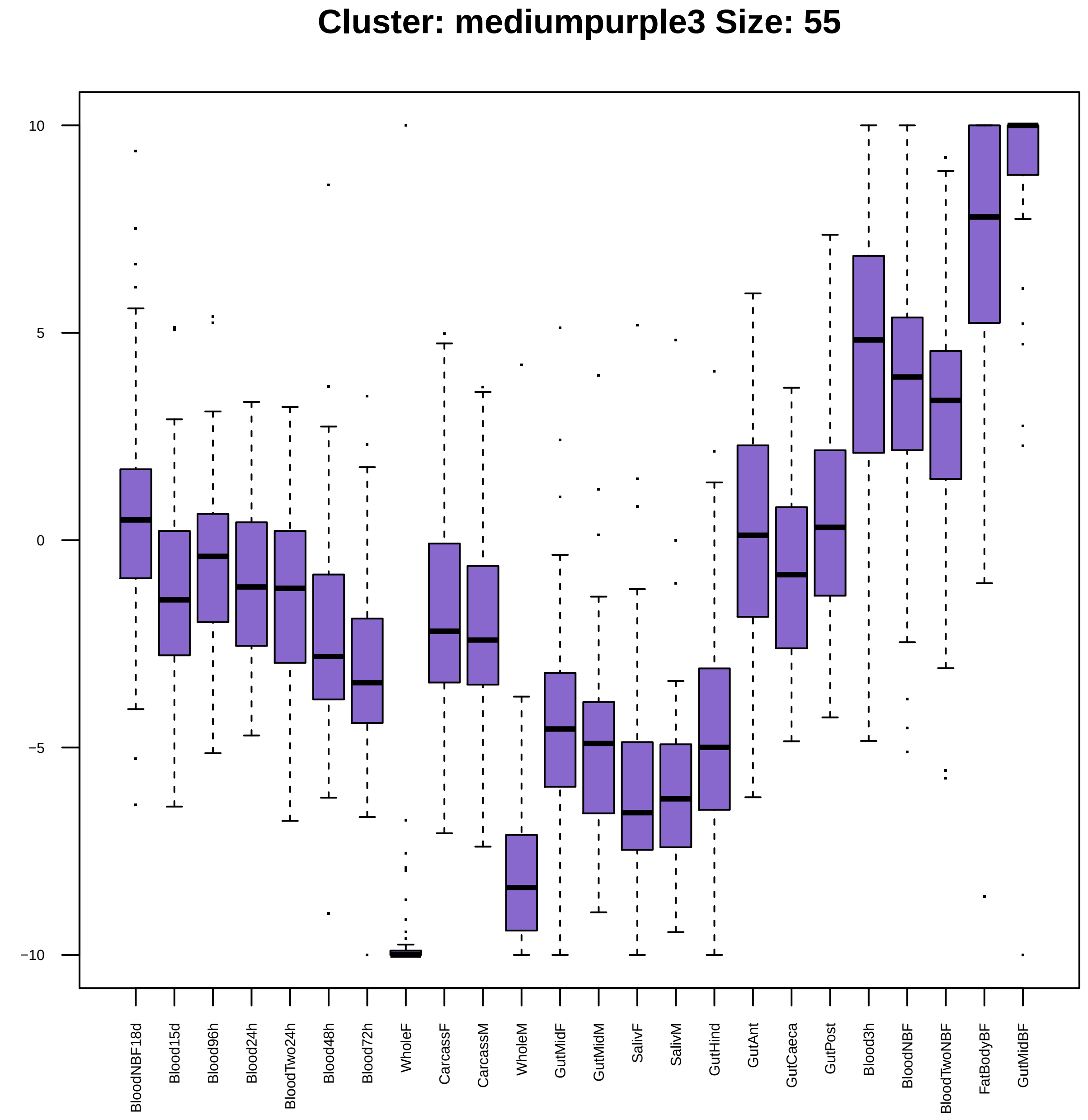
Cluster: mediumpurple3 Size: 55
10
5
0
−5
−10
BloodNBF18d
Blood15d
Blood96h
Blood24h
BloodTwo24h
Blood48h
Blood72h
WholeF
CarcassF
CarcassM
WholeM
GutMidF
GutMidM
SalivF
SalivM
GutHind
GutAnt
GutCaeca
GutPost
Blood3h
BloodNBF
BloodTwoNBF
FatBodyBF
GutMidBF

| | GO.ID | BPCluster: mediumpurple3 Size: 55 | Annotated | Significant | Expected | Rank in ClassicF | Weight01F | ClassicF |
|---|---|---|---|---|---|---|---|---|
| *4* | GO:0050909 | sensory perception of taste | 56 | 3 | 0.32 | 21 | 0.024 | 0.0037 |

| | GO.ID | MFCluster: mediumpurple3 Size: 55 | Annotated | Significant | Expected | Rank in ClassicF | Weight01F | ClassicF |
|---|---|---|---|---|---|---|---|---|
| *3* | GO:0005549 | odorant binding | 118 | 5 | 0.65 | 6 | 0.00045 | 0.00045 |
| *4* | GO:0005272 | sodium channel activity | 26 | 2 | 0.14 | 12 | 0.00897 | 0.00897 |

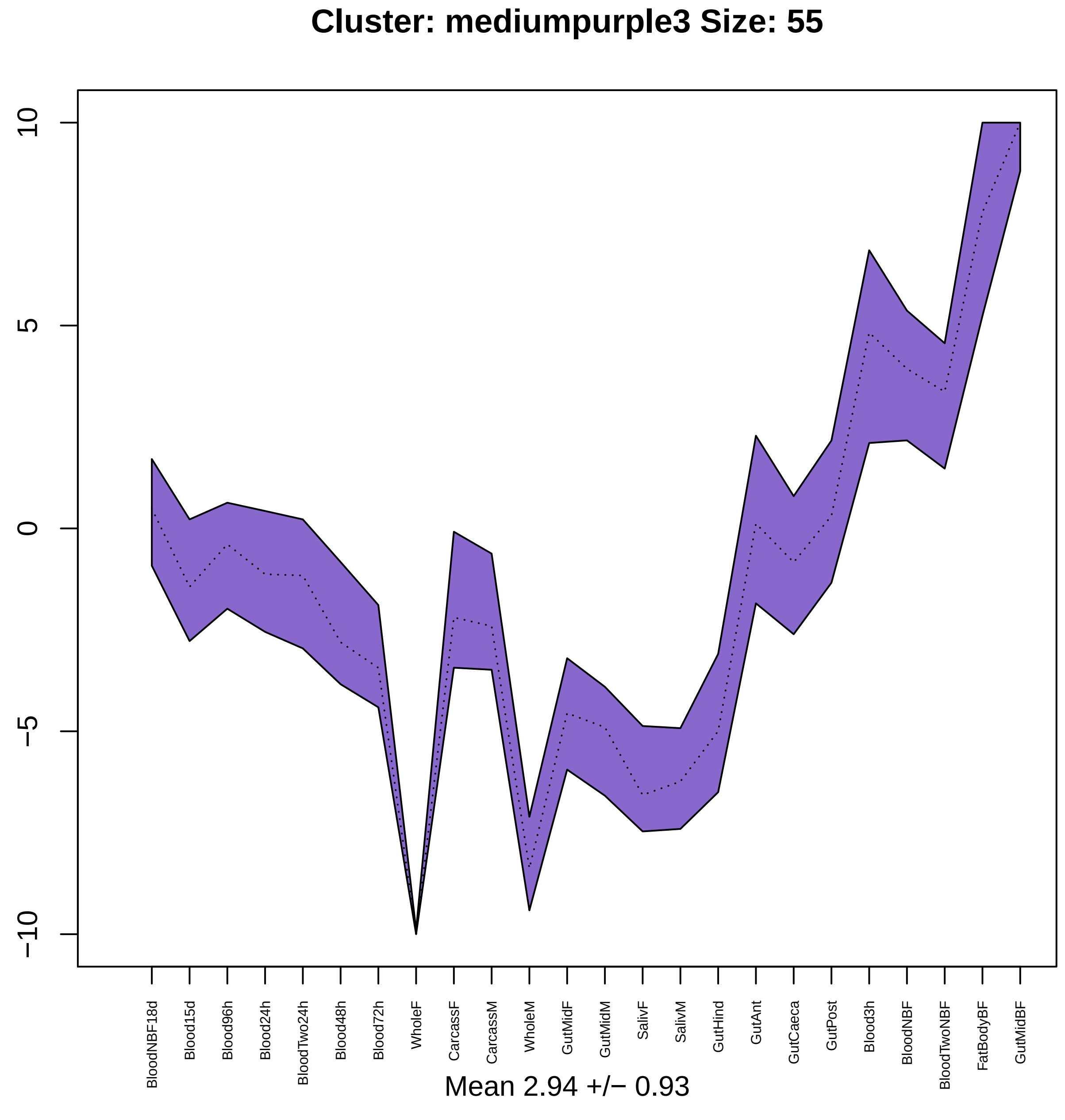

Cluster: mediumpurple3 Size: 55
10
5
0
−5
−10
BloodNBF18d
Blood15d
Blood96h
Blood24h
BloodTwo24h
Blood48h
Blood72h
WholeF
CarcassF
CarcassM
WholeM
GutMidF
GutMidM
SalivF
SalivM
GutHind
GutAnt
GutCaeca
GutPost
Blood3h
BloodNBF
BloodTwoNBF
FatBodyBF
GutMidBF
Mean 2.94 +/− 0.93

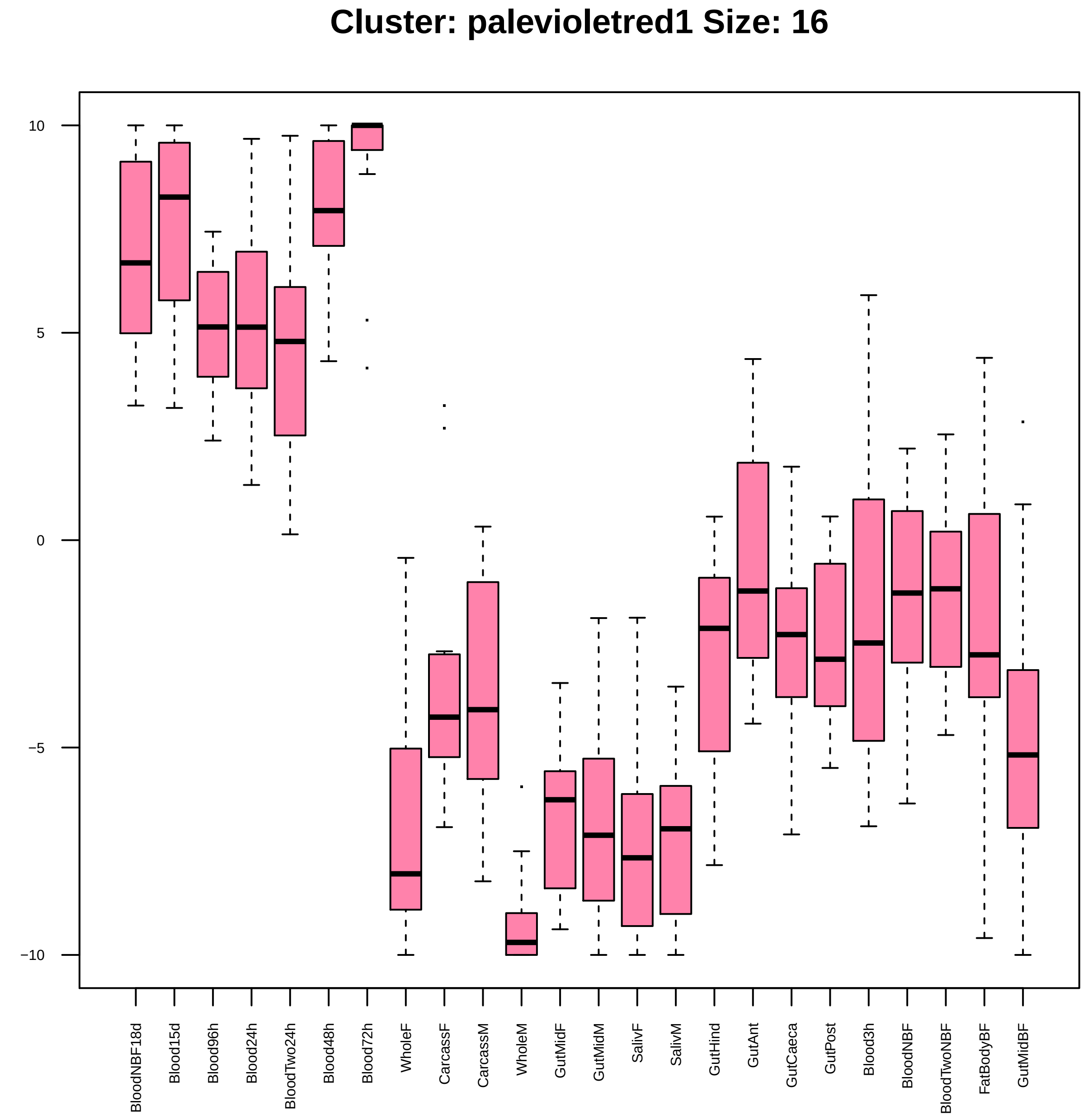
Cluster: palevioletred1 Size: 16
10
5
0
−5
−10
BloodNBF18d
Blood15d
Blood96h
Blood24h
BloodTwo24h
Blood48h
Blood72h
WholeF
CarcassF
CarcassM
WholeM
GutMidF
GutMidM
SalivF
SalivM
GutHind
GutAnt
GutCaeca
GutPost
Blood3h
BloodNBF
BloodTwoNBF
FatBodyBF
GutMidBF

| | GO.ID | BPCluster: palevioletred1 Size: 16 | Annotated | Significant | Expected | Rank in ClassicF | Weight01F | ClassicF |
|---|---|---|---|---|---|---|---|---|
| *1* | GO:0006342 | chromatin silencing | 26 | 2 | 0.04 | 6 | 0.00057 | 0.00057 |
| *2* | GO:0016570 | histone modification | 58 | 2 | 0.08 | 18 | 0.00281 | 0.00281 |

| | GO.ID | MFCluster: palevioletred1 Size: 16 | Annotated | Significant | Expected | Rank in ClassicF | Weight01F | ClassicF |
|---|---|---|---|---|---|---|---|---|
| *1* | GO:0019904 | protein domain specific binding | 37 | 2 | 0.05 | 3 | 0.0011 | 0.00112 |
| *2* | GO:0043565 | sequence−specific DNA binding | 212 | 3 | 0.29 | 5 | 0.0026 | 0.00258 |
| *3* | GO:0003677 | DNA binding | 523 | 6 | 0.73 | 1 | 0.0027 | 2.3e−05 |
| *4* | GO:0003700 | DNA−binding transcription factor activit... | 282 | 3 | 0.39 | 6 | 0.0058 | 0.00579 |
| *18* | GO:1901363 | heterocyclic compound binding | 2046 | 7 | 2.84 | 7 | 1.0000 | 0.00756 |
| *23* | GO:0140110 | transcription regulator activity | 322 | 4 | 0.45 | 2 | 1.0000 | 0.00066 |

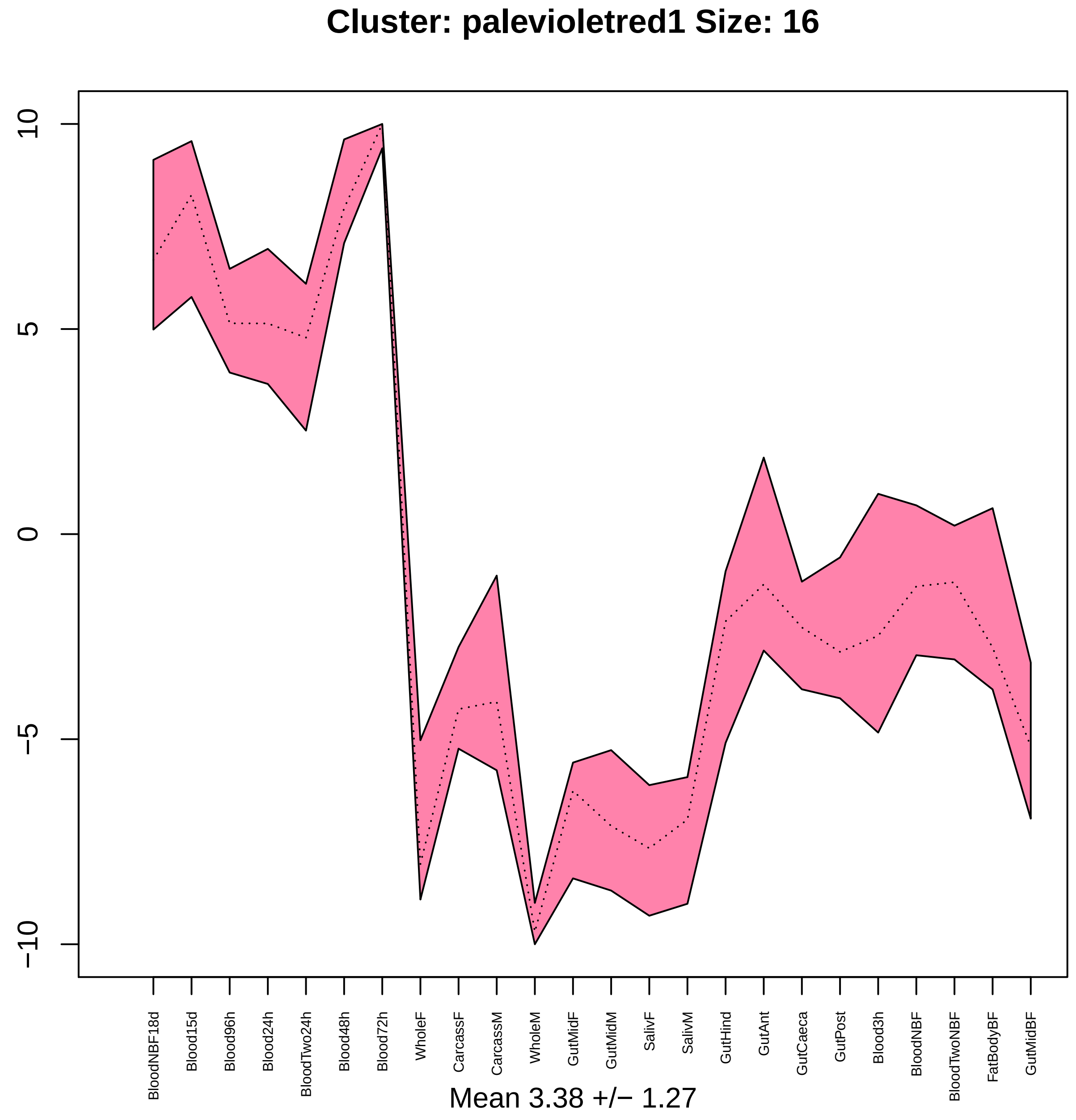
Cluster: palevioletred1 Size: 16
10
5
0
−5
−10
BloodNBF18d
Blood15d
Blood96h
Blood24h
BloodTwo24h
Blood48h
Blood72h
WholeF
CarcassF
CarcassM
WholeM
GutMidF
GutMidM
SalivF
SalivM
GutHind
GutAnt
GutCaeca
GutPost
Blood3h
BloodNBF
BloodTwoNBF
FatBodyBF
GutMidBF
Mean 3.38 +/− 1.27

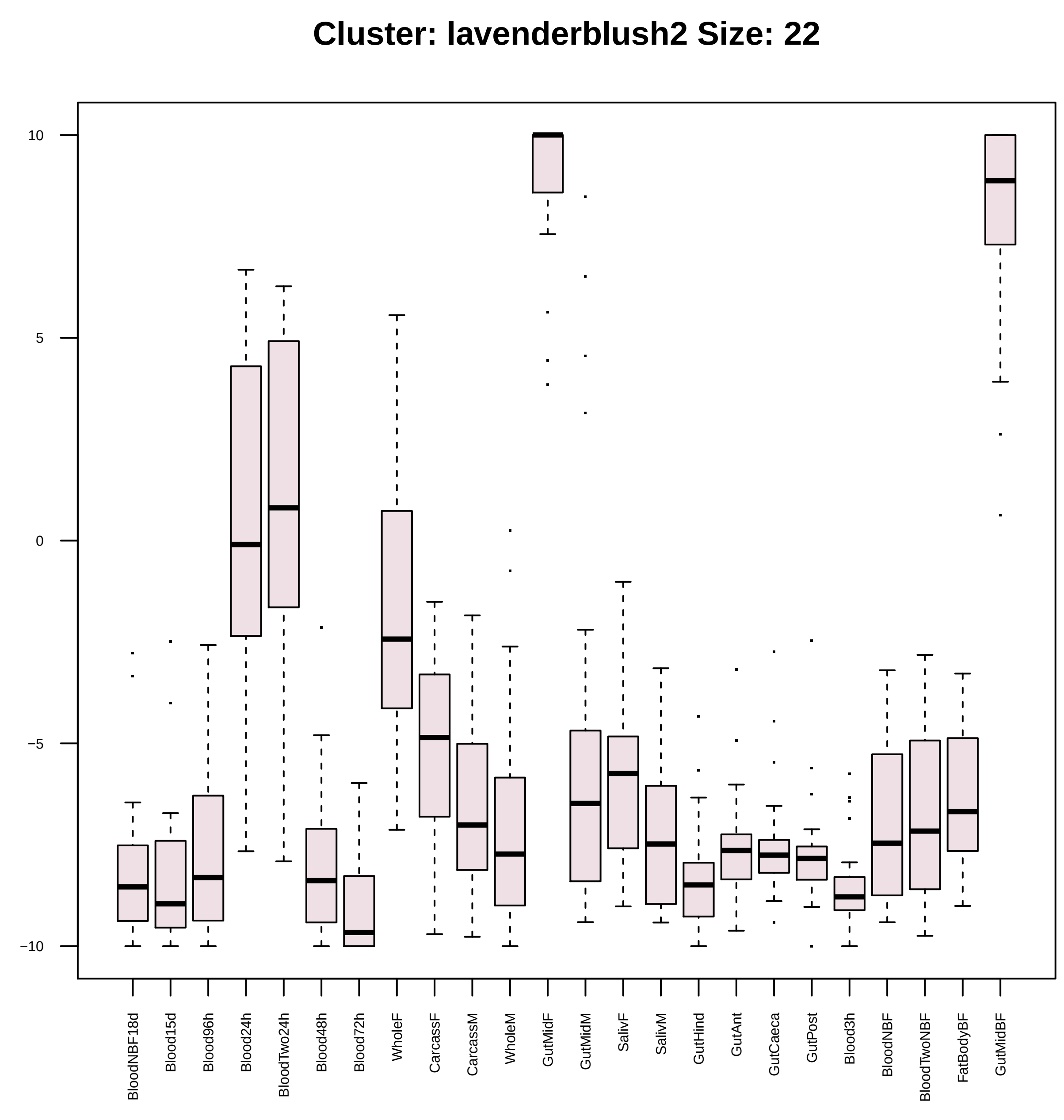
Cluster: lavenderblush2 Size: 22
10
5
0
−5
−10
BloodNBF18d
Blood15d
Blood96h
Blood24h
BloodTwo24h
Blood48h
Blood72h
WholeF
CarcassF
CarcassM
WholeM
GutMidF
GutMidM
SalivF
SalivM
GutHind
GutAnt
GutCaeca
GutPost
Blood3h
BloodNBF
BloodTwoNBF
FatBodyBF
GutMidBF

| | GO.ID | BPCluster: lavenderblush2 Size: 22 | Annotated | Significant | Expected | Rank in ClassicF | Weight01F | ClassicF |
|---|---|---|---|---|---|---|---|---|
| 7 | GO:0008152 | metabolic process | 4285 | 14 | 9.41 | 6 | 0.274 | 0.00380 |
| 19 | GO:0019538 | protein metabolic process | 1480 | 10 | 3.25 | 3 | 1.000 | 0.00017 |

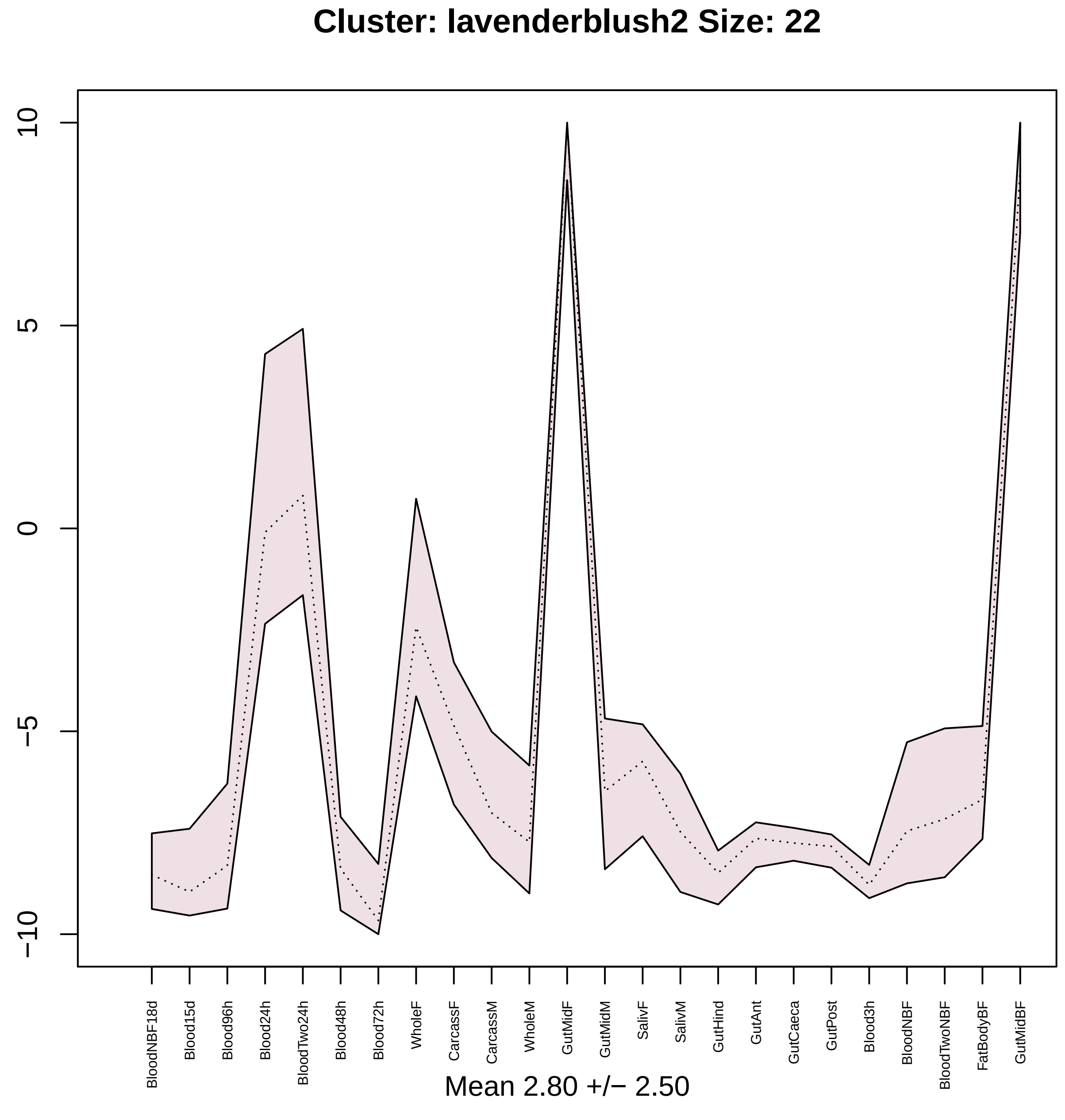

Cluster: lavenderblush2 Size: 22
10
5
0
−5
−10
BloodNBF18d
Blood15d
Blood96h
Blood24h
BloodTwo24h
Blood48h
Blood72h
WholeF
CarcassF
CarcassM
WholeM
GutMidF
GutMidM
SalivF
SalivM
GutHind
GutAnt
GutCaeca
GutPost
Blood3h
BloodNBF
BloodTwoNBF
FatBodyBF
GutMidBF
Mean 2.80 +/− 2.50

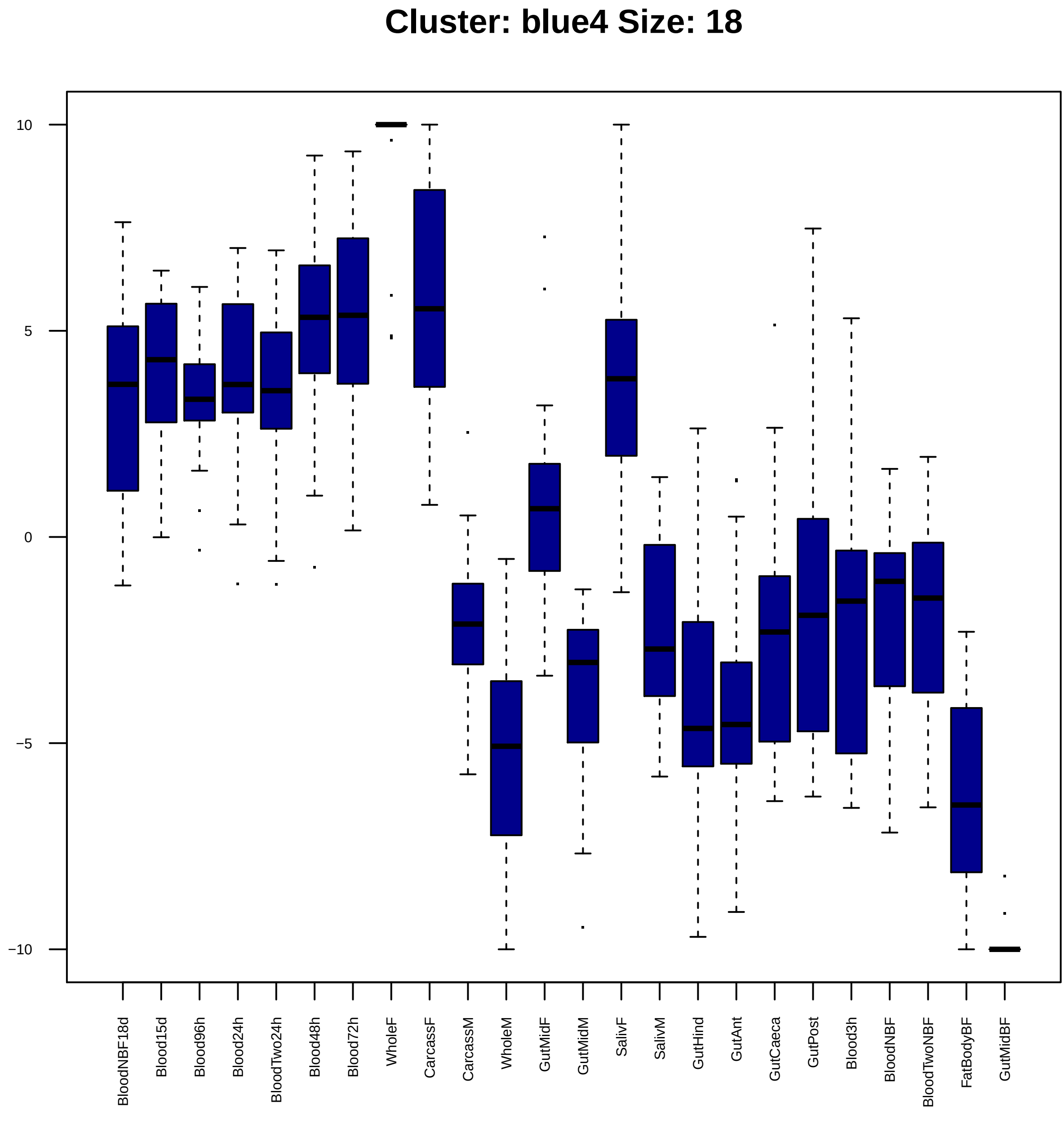
Cluster: blue4 Size: 18
10
5
0
−5
−10
BloodNBF18d
Blood15d
Blood96h
Blood24h
BloodTwo24h
Blood48h
Blood72h
WholeF
CarcassF
CarcassM
WholeM
GutMidF
GutMidM
SalivF
SalivM
GutHind
GutAnt
GutCaeca
GutPost
Blood3h
BloodNBF
BloodTwoNBF
FatBodyBF
GutMidBF

| | GO.ID | BPCluster: blue4 Size: 18 | Annotated | Significant | Expected | Rank in ClassicF | Weight01F | ClassicF |
|---|---|---|---|---|---|---|---|---|
| *1* | GO:0042254 | ribosome biogenesis | 64 | 3 | 0.09 | 1 | 0.00059 | 7.8e−05 |
| *8* | GO:0006396 | RNA processing | 267 | 3 | 0.38 | 4 | 0.03702 | 0.0051 |

|  | GO.ID | MFCluster: blue4 Size: 18 | Annotated | Significant | Expected | Rank in ClassicF | Weight01F | ClassicF |
|---|---|---|---|---|---|---|---|---|
| *1* | GO:0003723 | RNA binding | 307 | 4 | 0.60 | 1 | 0.0057 | 0.0023 |
| *27* | GO:0140098 | catalytic activity, acting on RNA | 165 | 3 | 0.32 | 2 | 1.0000 | 0.0036 |

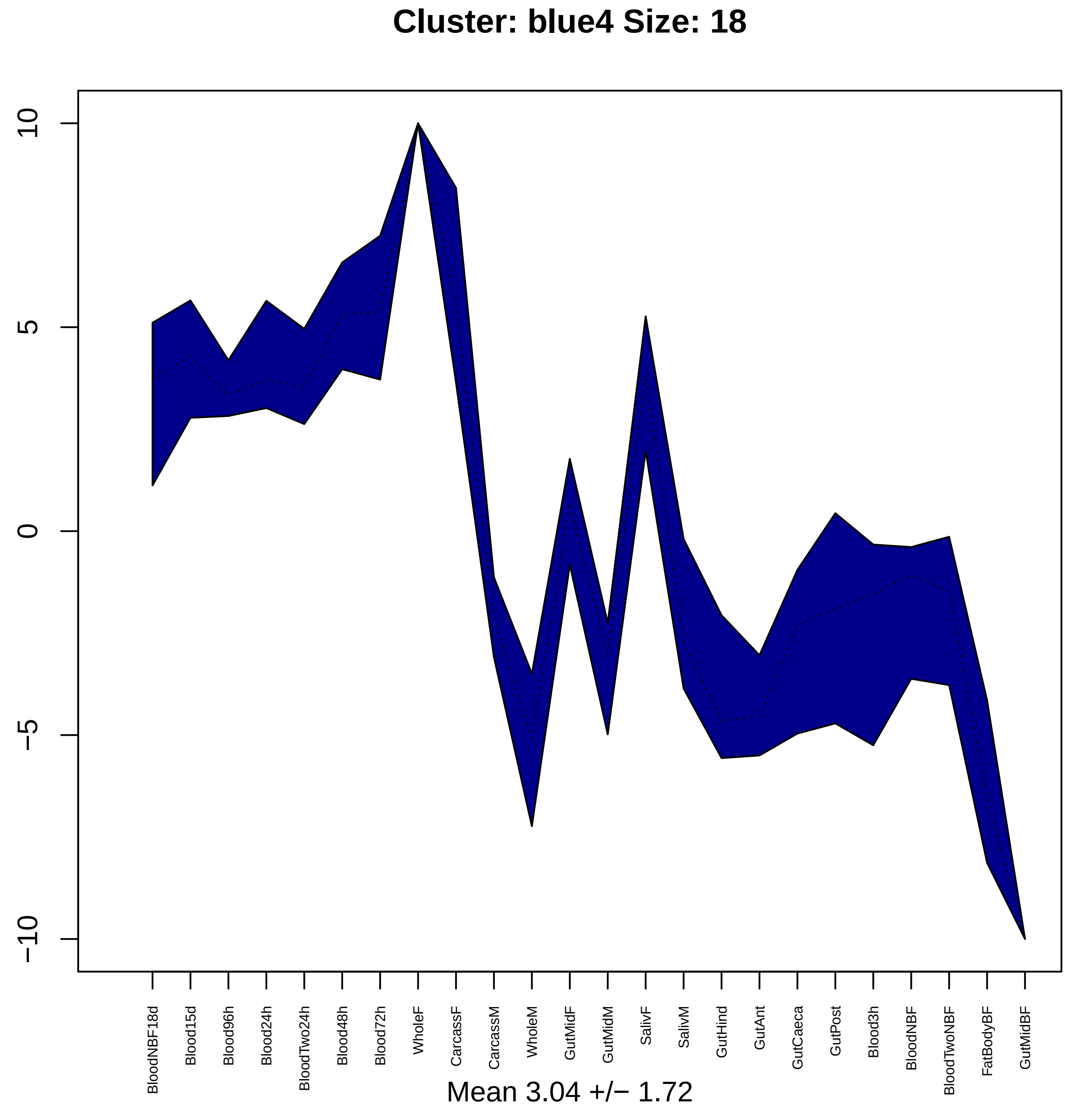

Cluster: blue4 Size: 18
10
5
0
−5
−10
BloodNBF18d
Blood15d
Blood96h
Blood24h
BloodTwo24h
Blood48h
Blood72h
WholeF
CarcassF
CarcassM
WholeM
GutMidF
GutMidM
SalivF
SalivM
GutHind
GutAnt
GutCaeca
GutPost
Blood3h
BloodNBF
BloodTwoNBF
FatBodyBF
GutMidBF
Mean 3.04 +/− 1.72

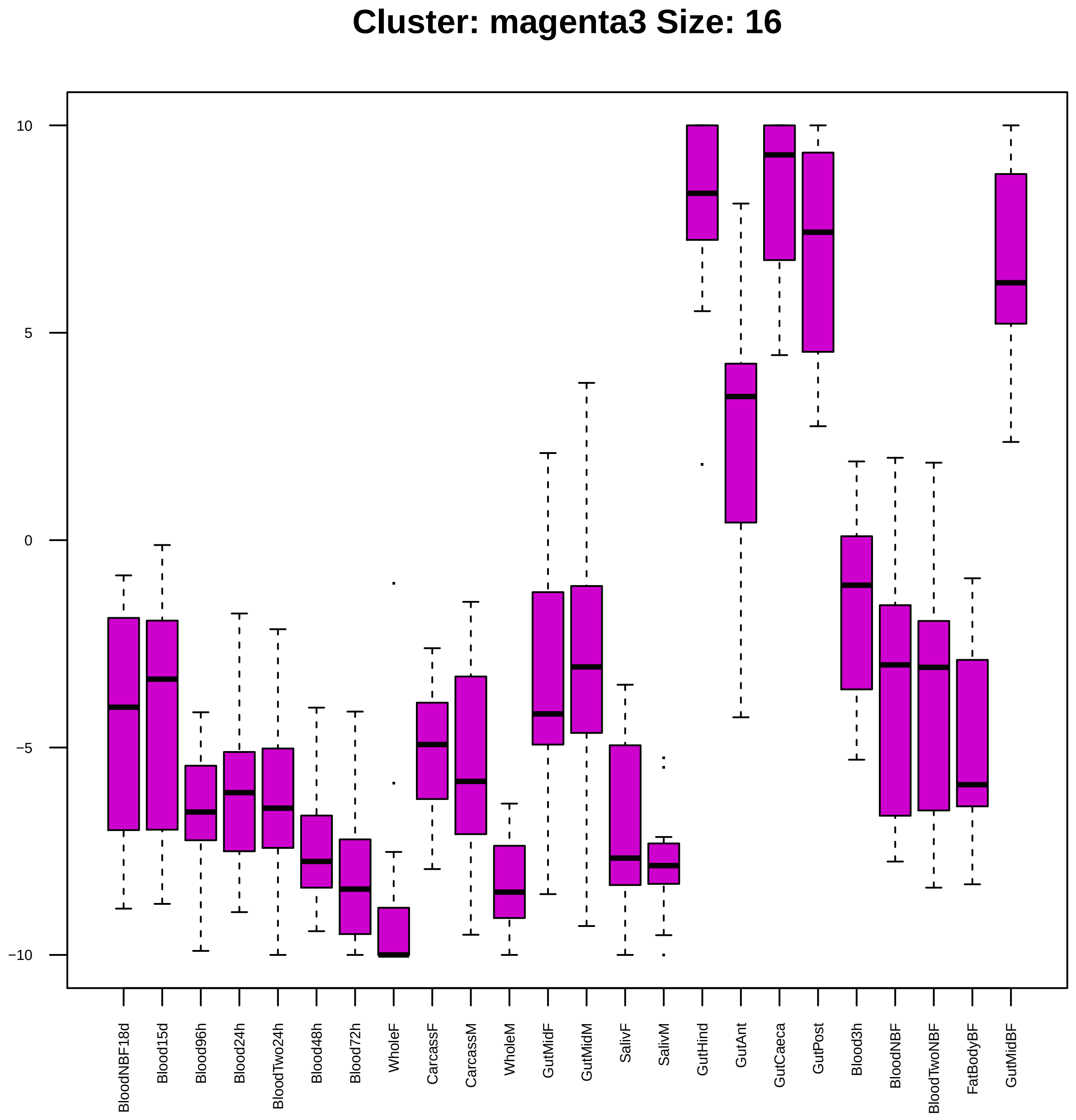

Cluster: magenta3 Size: 16
10
5
0
−5
−10
BloodNBF18d
Blood15d
Blood96h
Blood24h
BloodTwo24h
Blood48h
Blood72h
WholeF
CarcassF
CarcassM
WholeM
GutMidF
GutMidM
SalivF
SalivM
GutHind
GutAnt
GutCaeca
GutPost
Blood3h
BloodNBF
BloodTwoNBF
FatBodyBF
GutMidBF

| | GO.ID | MFCluster: magenta3 Size: 16 | Annotated | Significant | Expected | Rank in ClassicF | Weight01F | ClassicF |
|---|---|---|---|---|---|---|---|---|
| *1* | GO:0004553 | hydrolase activity, hydrolyzing O−glycos... | 84 | 2 | 0.10 | 1 | 0.0046 | 0.0046 |
| *17* | GO:0016798 | hydrolase activity, acting on glycosyl b... | 95 | 2 | 0.12 | 2 | 1.0000 | 0.0058 |

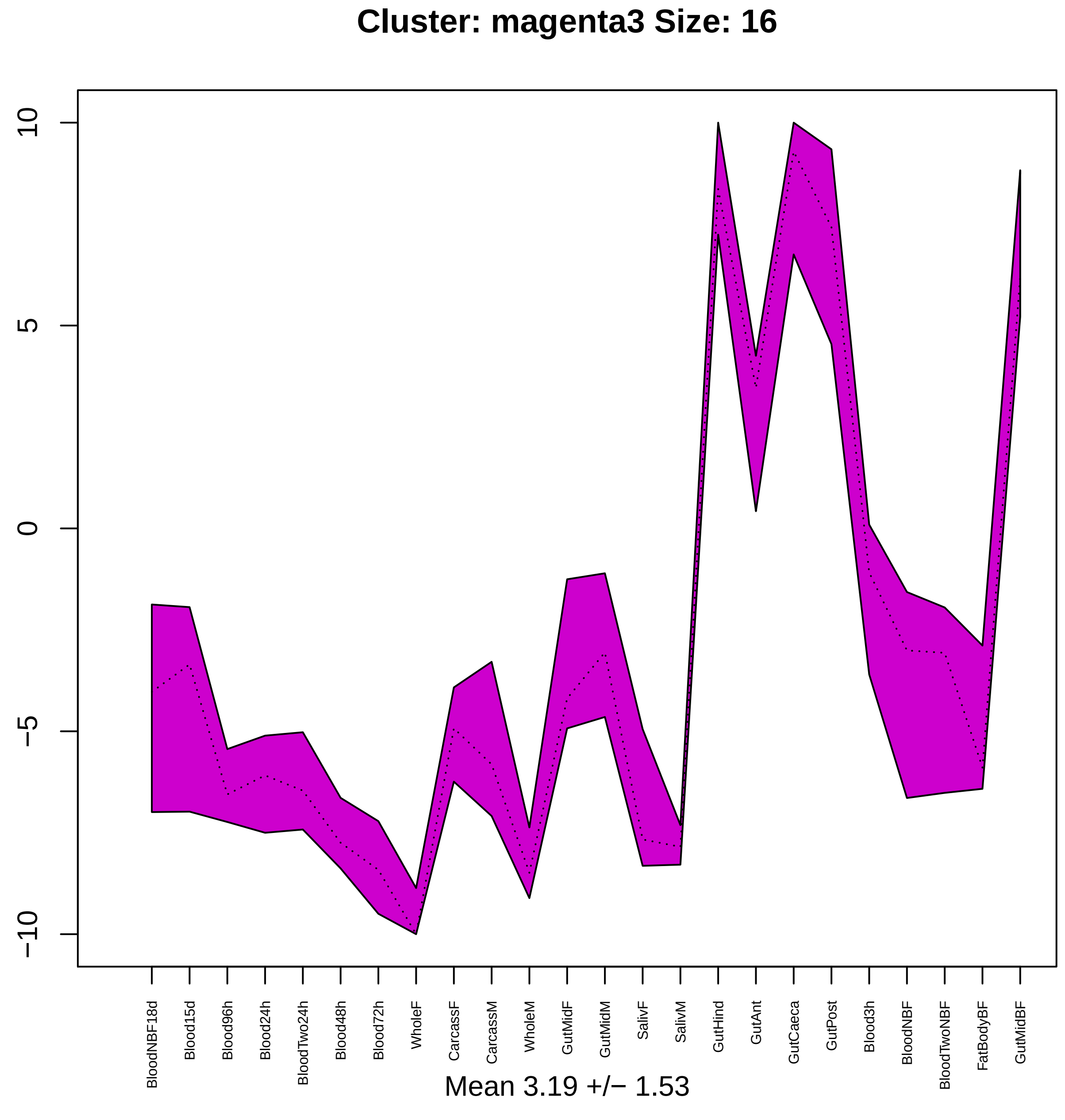
Cluster: magenta3 Size: 16
10
5
0
−5
−10
BloodNBF18d
Blood15d
Blood96h
Blood24h
BloodTwo24h
Blood48h
Blood72h
WholeF
CarcassF
CarcassM
WholeM
GutMidF
GutMidM
SalivF
SalivM
GutHind
GutAnt
GutCaeca
GutPost
Blood3h
BloodNBF
BloodTwoNBF
FatBodyBF
GutMidBF
Mean 3.19 +/− 1.53

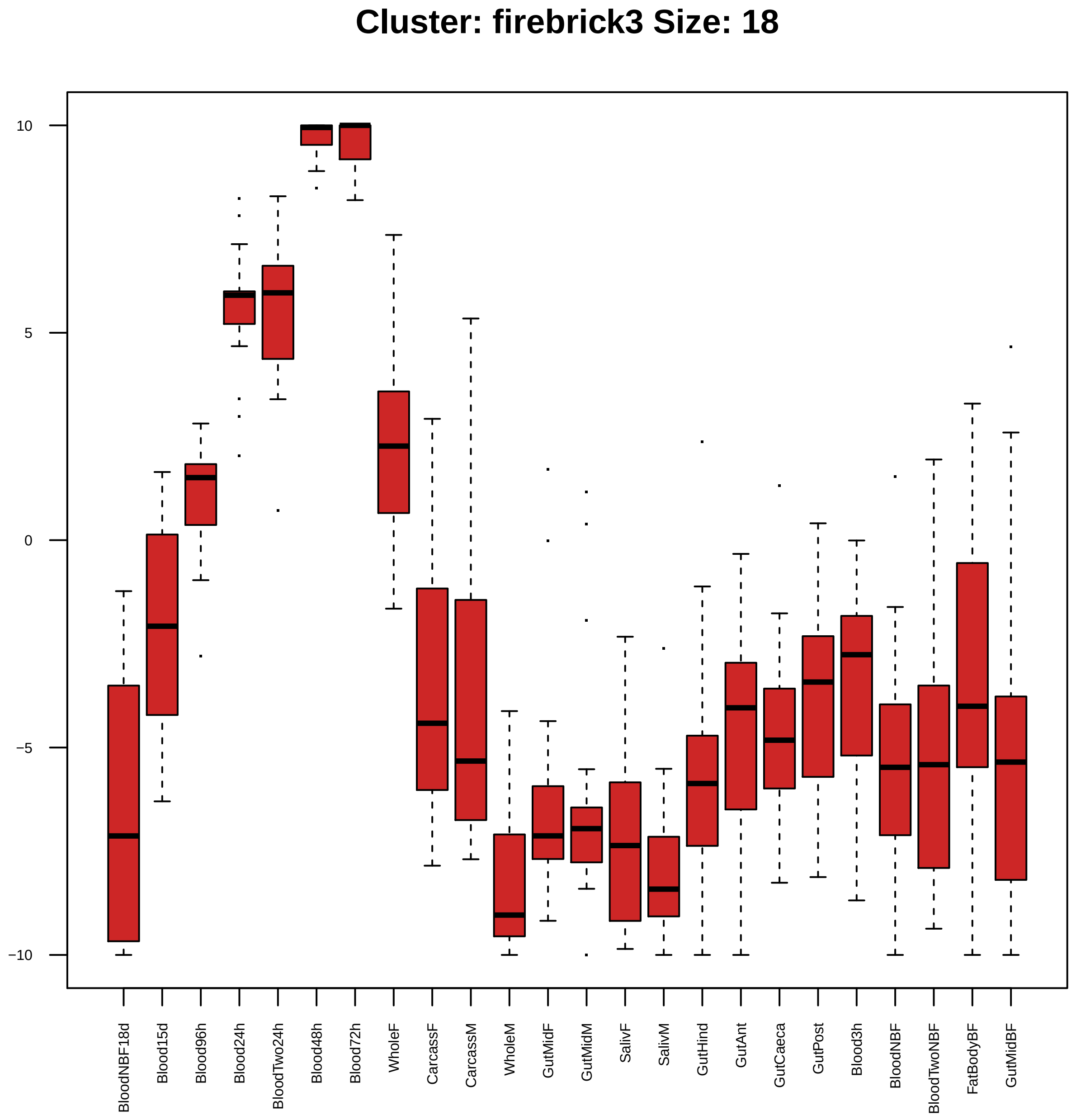
Cluster: firebrick3 Size: 18
10
5
0
−5
−10
BloodNBF18d
Blood15d
Blood96h
Blood24h
BloodTwo24h
Blood48h
Blood72h
WholeF
CarcassF
CarcassM
WholeM
GutMidF
GutMidM
SalivF
SalivM
GutHind
GutAnt
GutCaeca
GutPost
Blood3h
BloodNBF
BloodTwoNBF
FatBodyBF
GutMidBF

| | GO.ID | BPCluster: firebrick3 Size: 18 | Annotated | Significant | Expected | Rank in ClassicF | Weight01F | ClassicF |
|---|---|---|---|---|---|---|---|---|
| *1* | GO:0051298 | centrosome duplication | 51 | 2 | 0.11 | 17 | 0.0054 | 0.0054 |
| *2* | GO:0140014 | mitotic nuclear division | 105 | 3 | 0.23 | 2 | 0.0057 | 0.0014 |
| *3* | GO:0051301 | cell division | 126 | 3 | 0.28 | 7 | 0.0105 | 0.0023 |
| *4* | GO:0007049 | cell cycle | 432 | 5 | 0.95 | 3 | 0.0167 | 0.0017 |
| *22* | GO:0007088 | regulation of mitotic nuclear division | 31 | 2 | 0.07 | 5 | 0.0322 | 0.0020 |

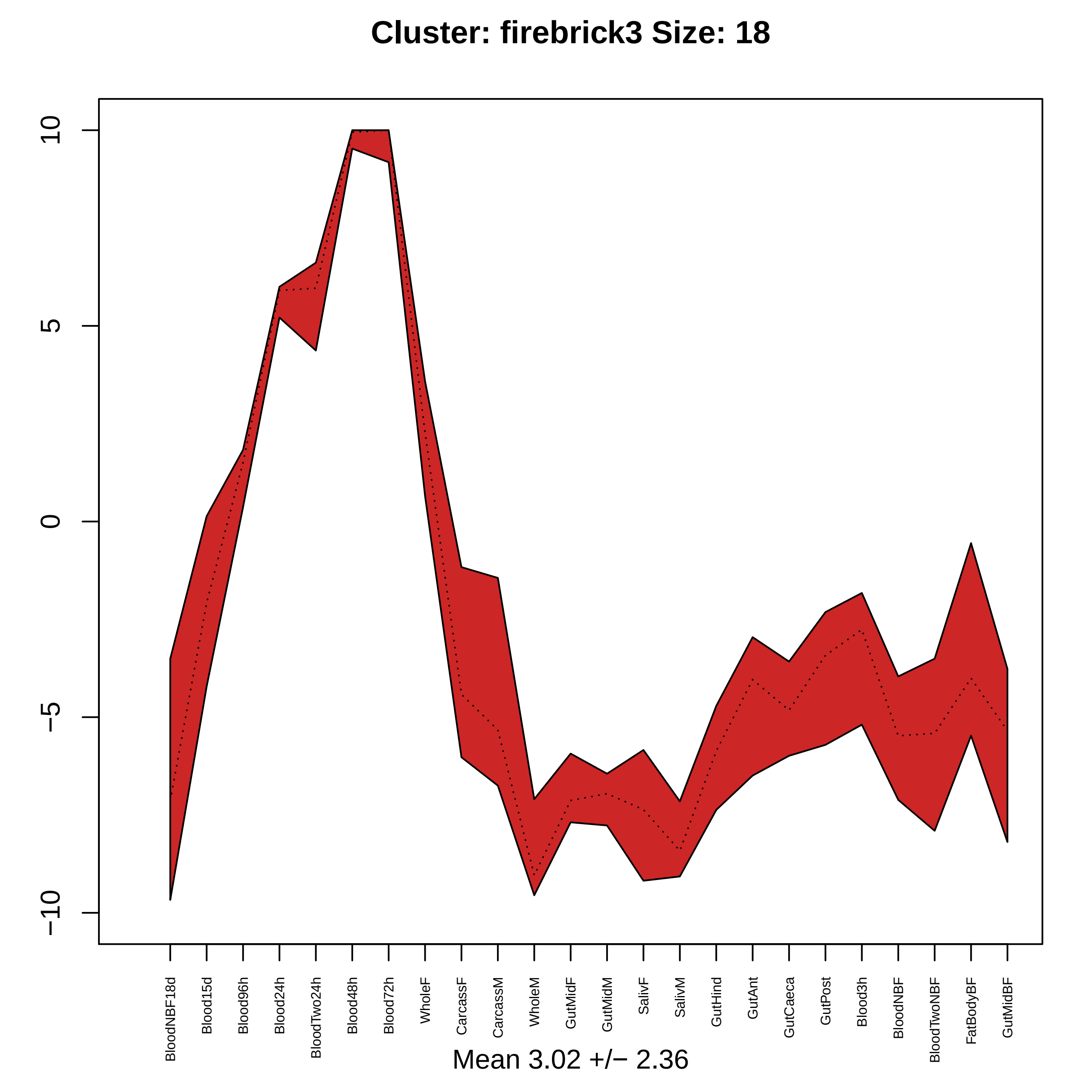

Cluster: firebrick3 Size: 18
10
5
0
−5
−10
BloodNBF18d
Blood15d
Blood96h
Blood24h
BloodTwo24h
Blood48h
Blood72h
WholeF
CarcassF
CarcassM
WholeM
GutMidF
GutMidM
SalivF
SalivM
GutHind
GutAnt
GutCaeca
GutPost
Blood3h
BloodNBF
BloodTwoNBF
FatBodyBF
GutMidBF
Mean 3.02 +/− 2.36

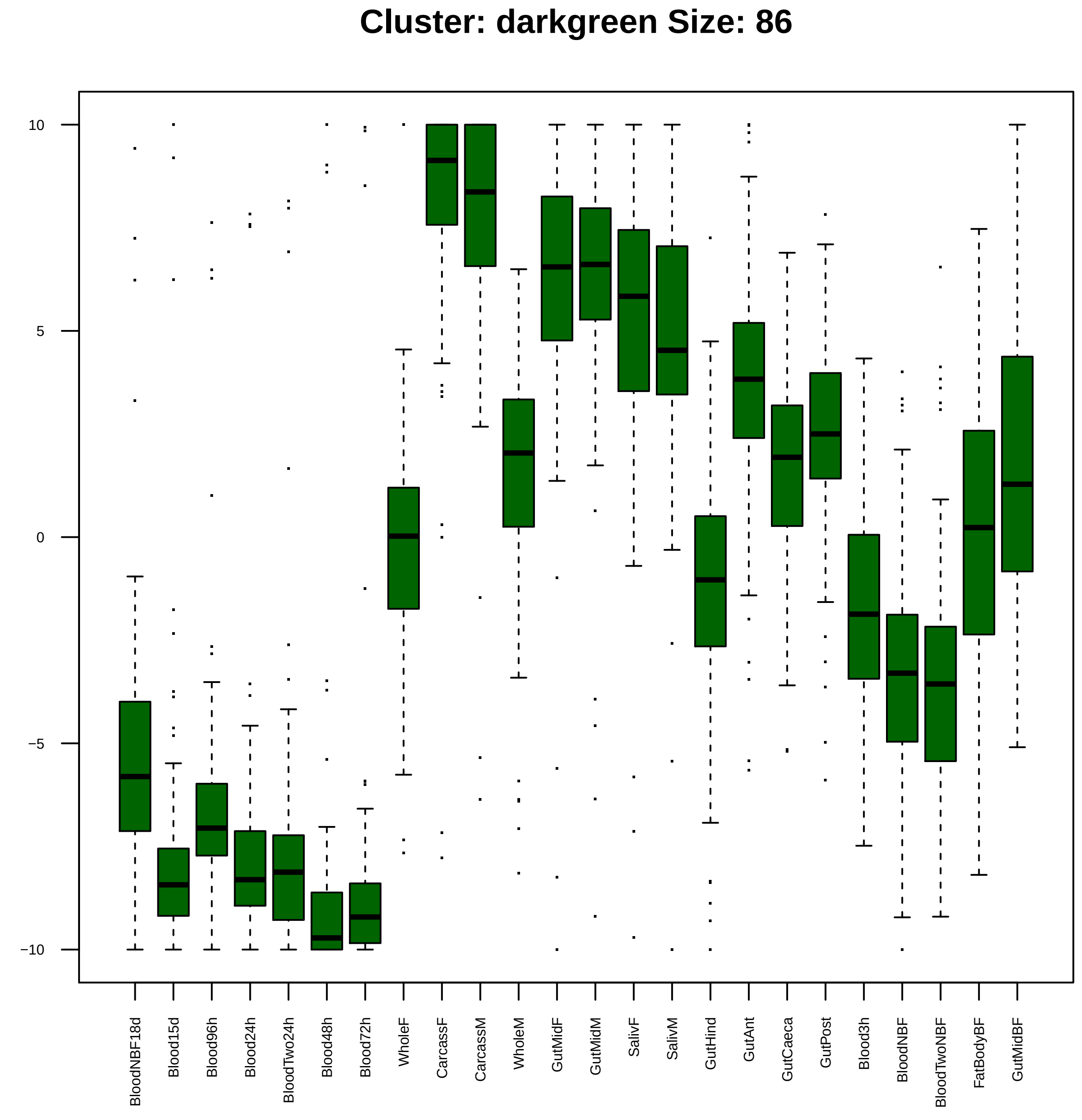
Cluster: darkgreen Size: 86
10
5
0
−5
−10
BloodNBF18d
Blood15d
Blood96h
Blood24h
BloodTwo24h
Blood48h
Blood72h
WholeF
CarcassF
CarcassM
WholeM
GutMidF
GutMidM
SalivF
SalivM
GutHind
GutAnt
GutCaeca
GutPost
Blood3h
BloodNBF
BloodTwoNBF
FatBodyBF
GutMidBF

| | GO.ID | BPCluster: darkgreen Size: 86 | Annotated | Significant | Expected | Rank in ClassicF | Weight01F | ClassicF |
|---|---|---|---|---|---|---|---|---|
| *1* | GO:0022404 | molting cycle process | 10 | 2 | 0.08 | 2 | 0.0025 | 0.0025 |
| *2* | GO:0007432 | salivary gland boundary specification | 12 | 2 | 0.09 | 3 | 0.0036 | 0.0036 |
| *3* | GO:0016042 | lipid catabolic process | 53 | 3 | 0.41 | 6 | 0.0076 | 0.0076 |

| | GO.ID | MFCluster: darkgreen Size: 86 | Annotated | Significant | Expected | Rank in ClassicF | Weight01F | ClassicF |
|---|---|---|---|---|---|---|---|---|
| *1* | GO:0016298 | lipase activity | 36 | 3 | 0.26 | 2 | 0.0038 | 0.0023 |
| *3* | GO:0052689 | carboxylic ester hydrolase activity | 35 | 3 | 0.26 | 1 | 0.0138 | 0.0021 |
| *12* | GO:0004930 | G protein−coupled receptor activity | 171 | 5 | 1.26 | 3 | 0.0859 | 0.0081 |

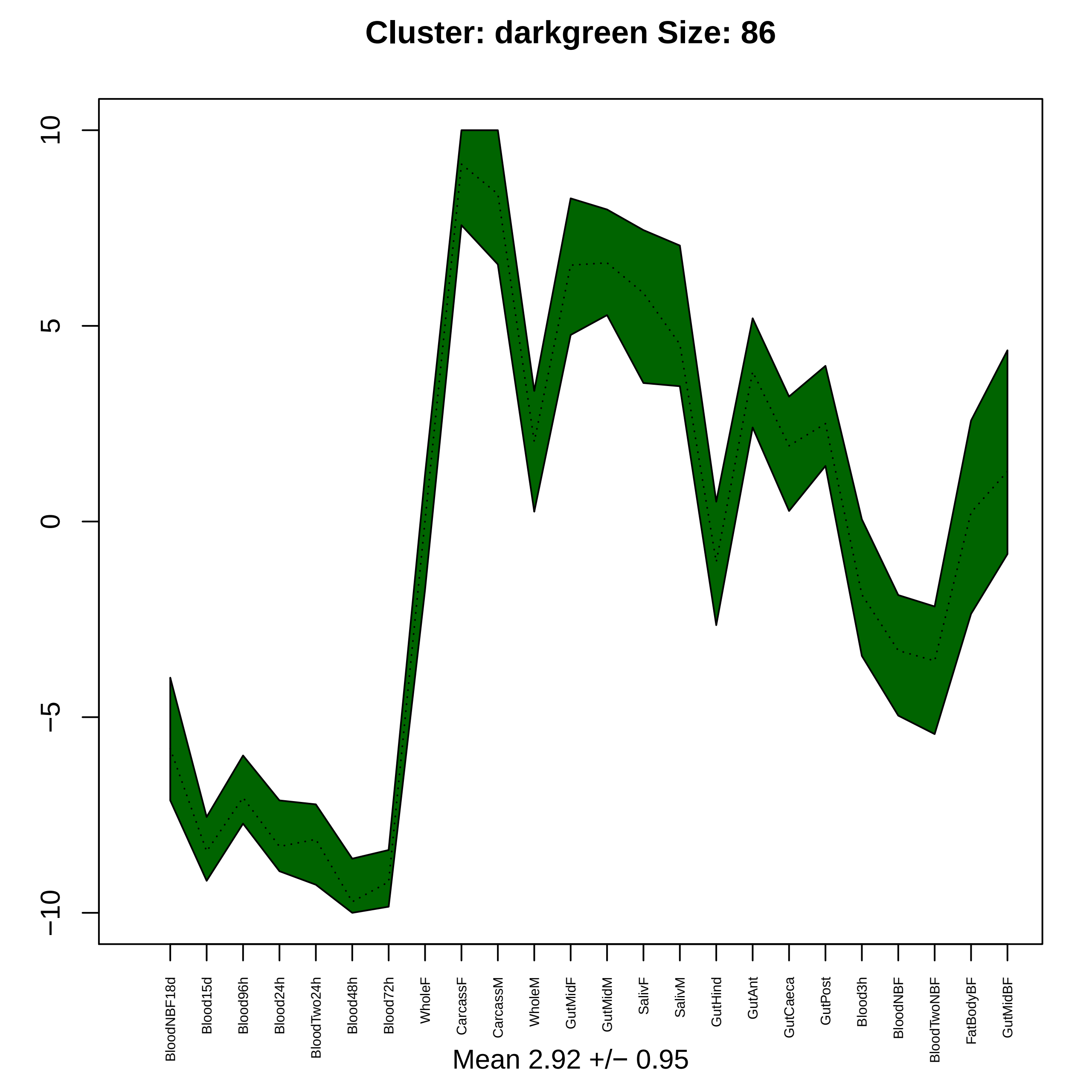

Cluster: darkgreen Size: 86
10
5
0
−5
−10
BloodNBF18d
Blood15d
Blood96h
Blood24h
BloodTwo24h
Blood48h
Blood72h
WholeF
CarcassF
CarcassM
WholeM
GutMidF
GutMidM
SalivF
SalivM
GutHind
GutAnt
GutCaeca
GutPost
Blood3h
BloodNBF
BloodTwoNBF
FatBodyBF
GutMidBF
Mean 2.92 +/− 0.95

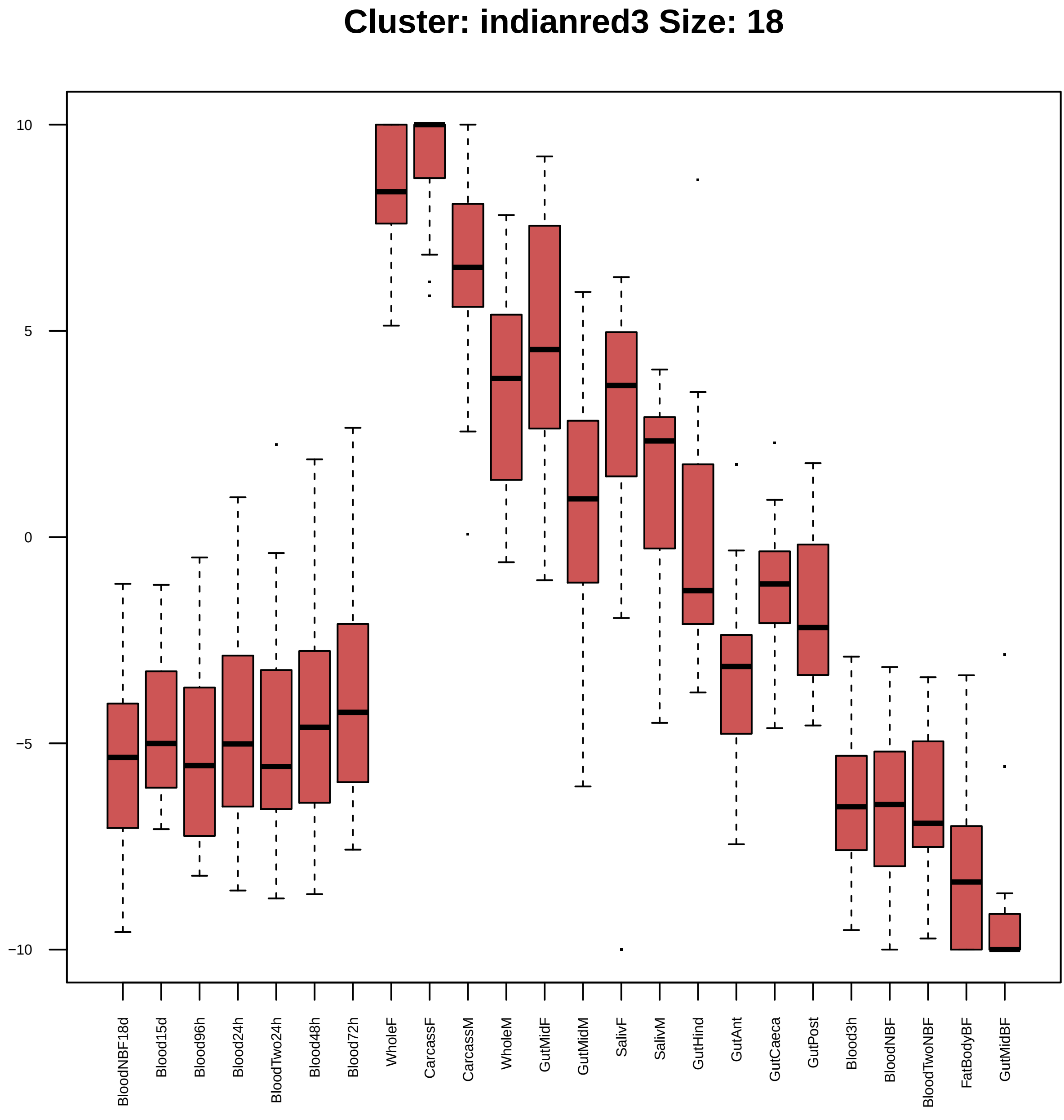

Cluster: indianred3 Size: 18
10
5
0
−5
−10
BloodNBF18d
Blood15d
Blood96h
Blood24h
BloodTwo24h
Blood48h
Blood72h
WholeF
CarcassF
CarcassM
WholeM
GutMidF
GutMidM
SalivF
SalivM
GutHind
GutAnt
GutCaeca
GutPost
Blood3h
BloodNBF
BloodTwoNBF
FatBodyBF
GutMidBF

| | GO.ID | BPCluster: indianred3 Size: 18 | Annotated | Significant | Expected | Rank in ClassicF | Weight01F | ClassicF |
|---|---|---|---|---|---|---|---|---|
| 1 | GO:0006457 | protein folding | 72 | 3 | 0.19 | 1 | 0.00084 | 0.00084 |
| 2 | GO:0006412 | translation | 289 | 4 | 0.77 | 5 | 0.00615 | 0.00615 |

| | GO.ID | MFCluster: indianred3 Size: 18 | Annotated | Significant | Expected | Rank in ClassicF | Weight01F | ClassicF |
|---|---|---|---|---|---|---|---|---|
| *2* | GO:0003735 | structural constituent of ribosome | 146 | 3 | 0.3 | 2 | 0.0031 | 0.0031 |

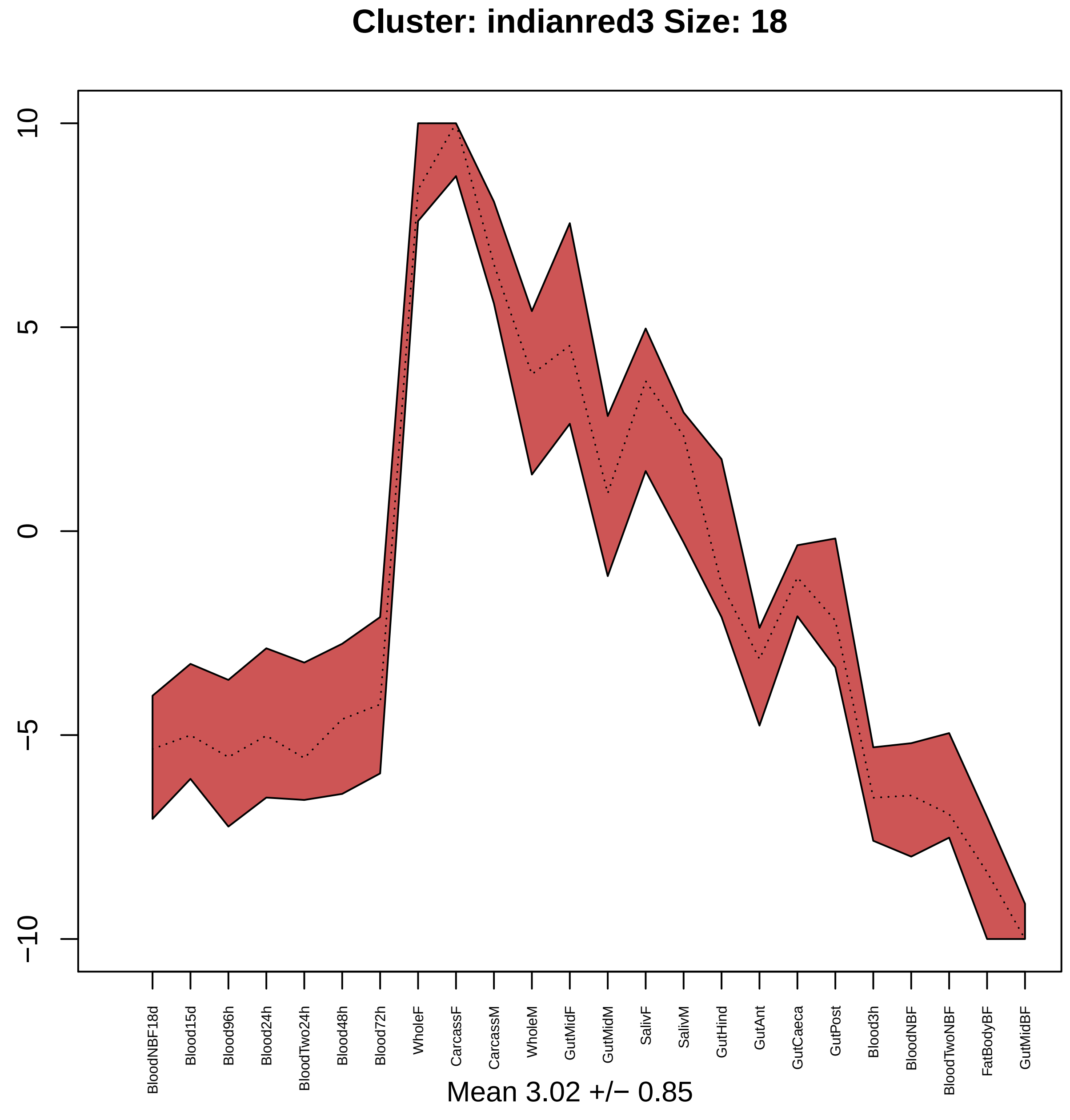

Cluster: indianred3 Size: 18
10
5
0
−5
−10
BloodNBF18d
Blood15d
Blood96h
Blood24h
BloodTwo24h
Blood48h
Blood72h
WholeF
CarcassF
CarcassM
WholeM
GutMidF
GutMidM
SalivF
SalivM
GutHind
GutAnt
GutCaeca
GutPost
Blood3h
BloodNBF
BloodTwoNBF
FatBodyBF
GutMidBF
Mean 3.02 +/− 0.85

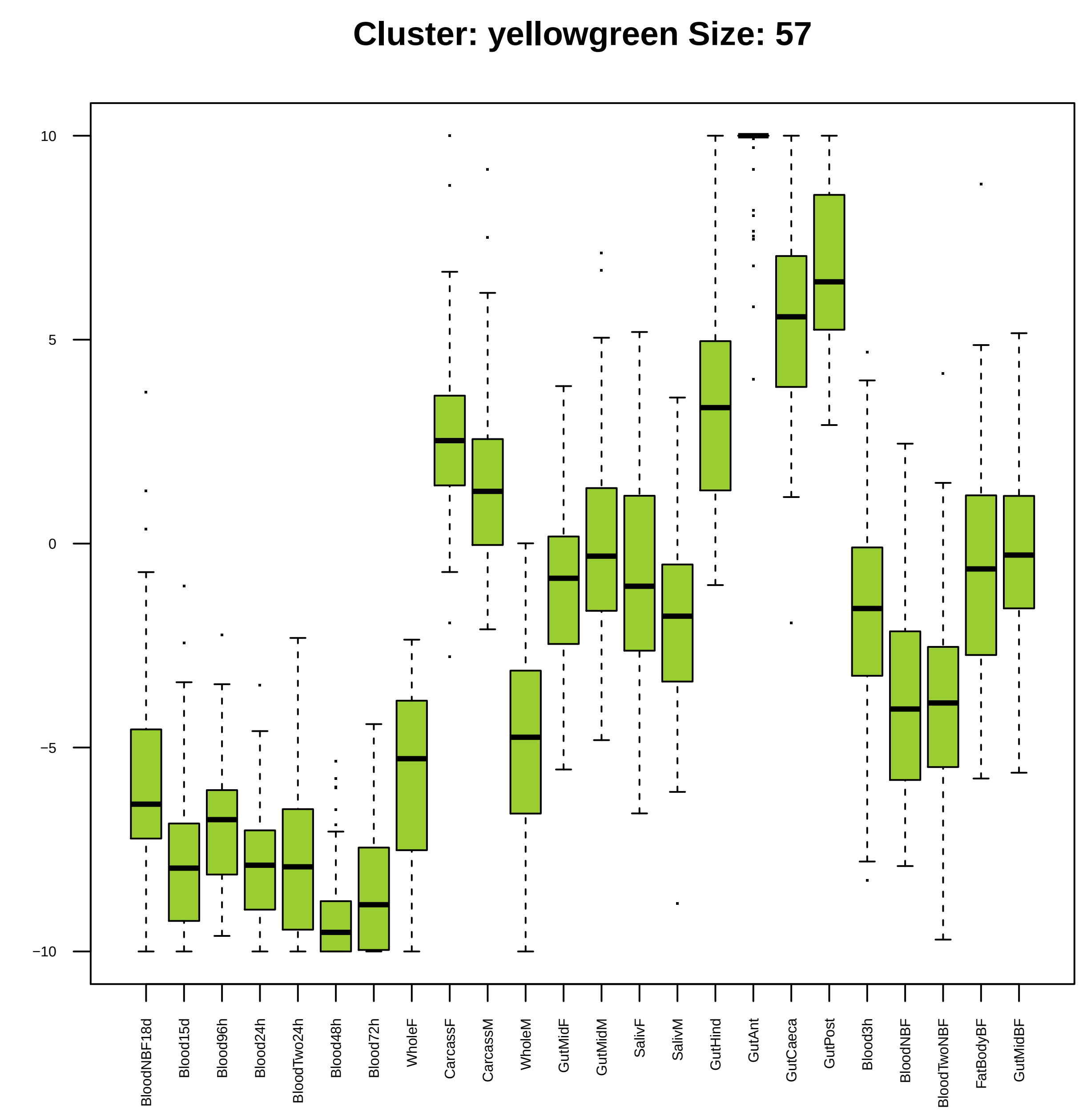

Cluster: yellowgreen Size: 57
10
5
0
−5
−10
BloodNBF18d
Blood15d
Blood96h
Blood24h
BloodTwo24h
Blood48h
Blood72h
WholeF
CarcassF
CarcassM
WholeM
GutMidF
GutMidM
SalivF
SalivM
GutHind
GutAnt
GutCaeca
GutPost
Blood3h
BloodNBF
BloodTwoNBF
FatBodyBF
GutMidBF

| | GO.ID | BPCluster: yellowgreen Size: 57 | Annotated | Significant | Expected | Rank in ClassicF | Weight01F | ClassicF |
|---|---|---|---|---|---|---|---|---|
| *1* | GO:0002805 | regulation of antimicrobial peptide bios... | 18 | 2 | 0.08 | 2 | 0.014 | 0.0031 |

| | GO.ID | MFCluster: yellowgreen Size: 57 | Annotated | Significant | Expected | Rank in ClassicF | Weight01F | ClassicF |
|---|---|---|---|---|---|---|---|---|
| 6 | GO:0005216 | ion channel activity | 139 | 4 | 0.69 | 1 | 0.076 | 0.0048 |

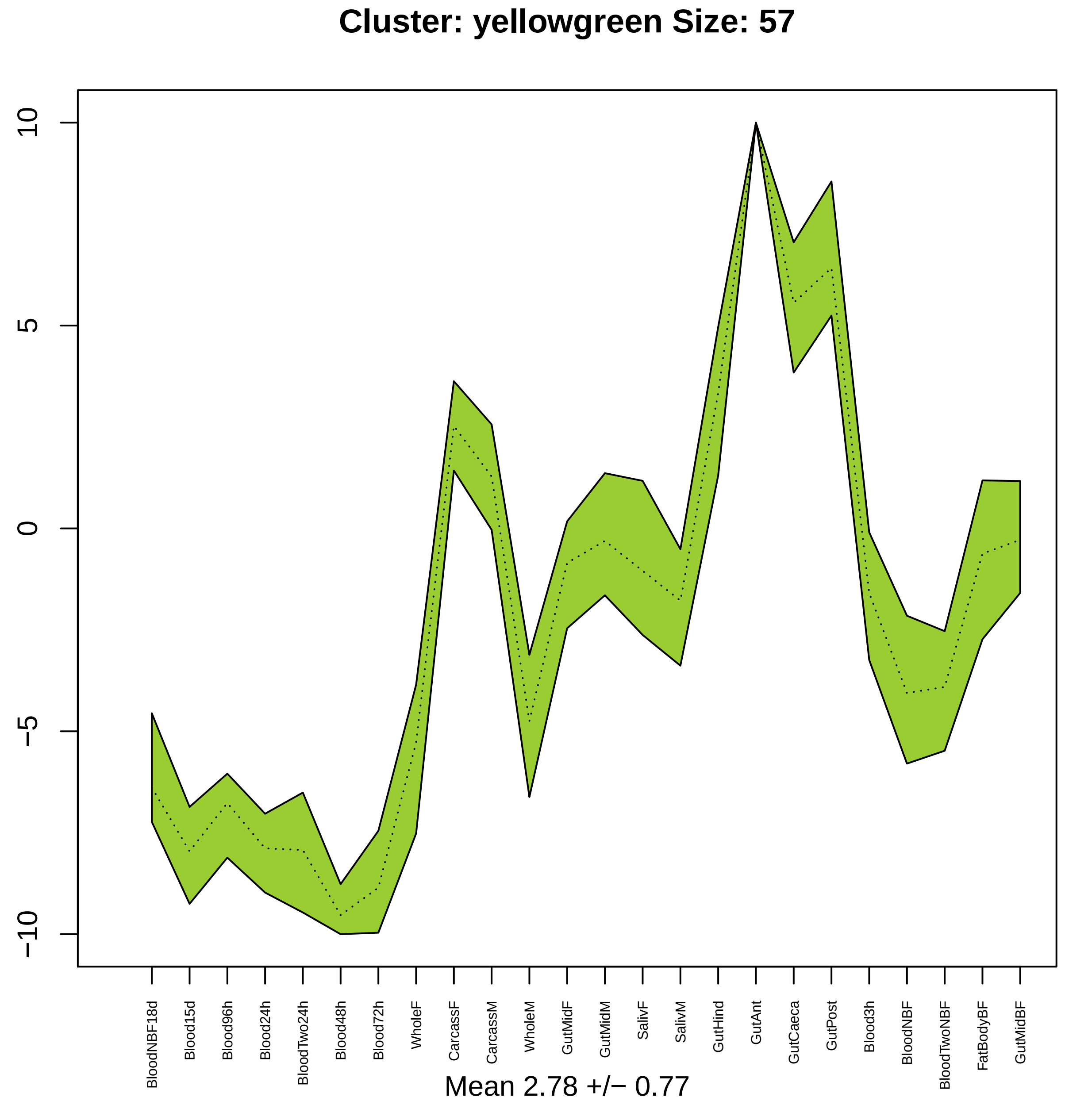
Cluster: yellowgreen Size: 57
10
5
0
−5
−10
BloodNBF18d
Blood15d
Blood96h
Blood24h
BloodTwo24h
Blood48h
Blood72h
WholeF
CarcassF
CarcassM
WholeM
GutMidF
GutMidM
SalivF
SalivM
GutHind
GutAnt
GutCaeca
GutPost
Blood3h
BloodNBF
BloodTwoNBF
FatBodyBF
GutMidBF
Mean 2.78 +/− 0.77

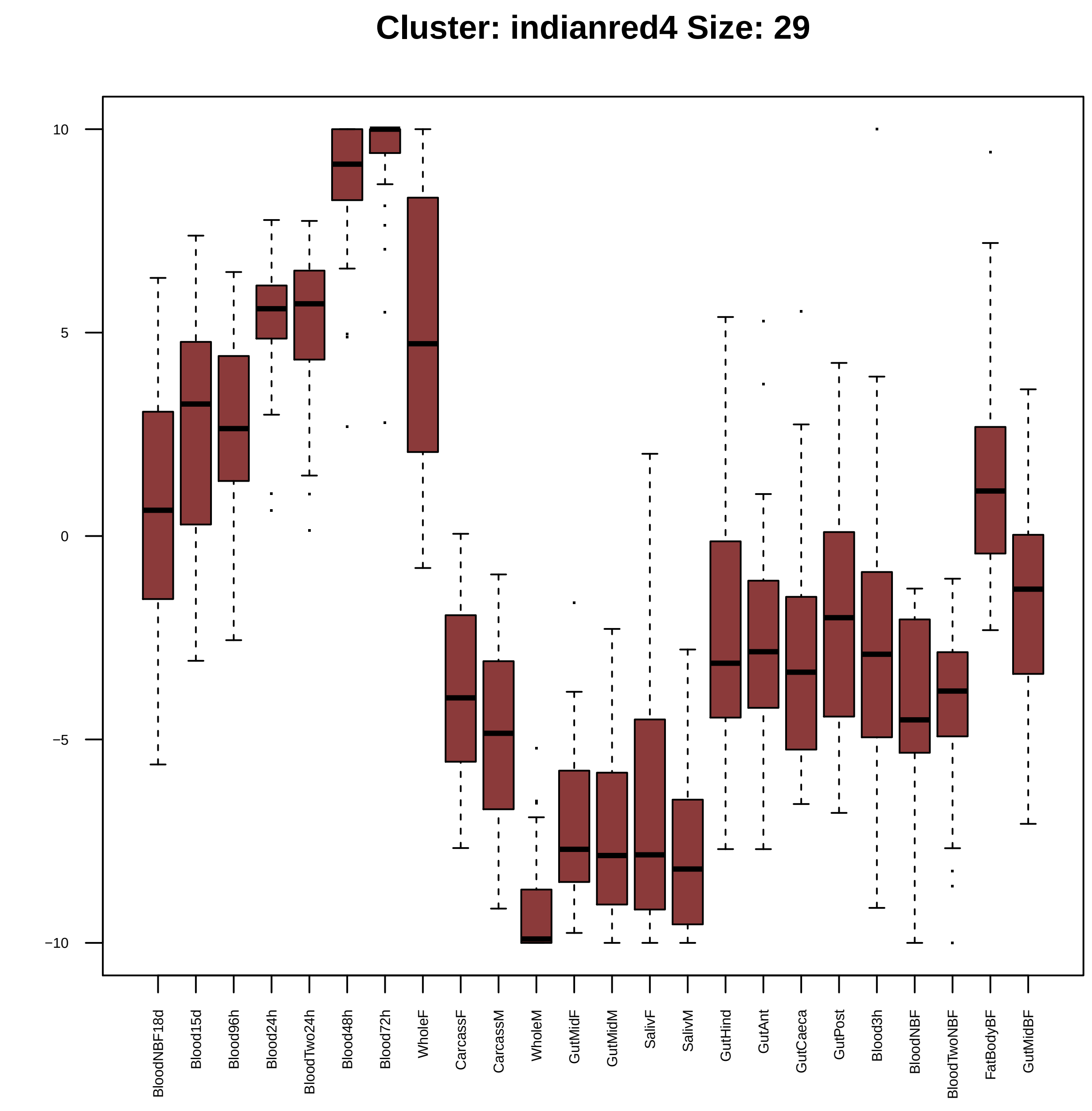
Cluster: indianred4 Size: 29
10
5
0
−5
−10
BloodNBF18d
Blood15d
Blood96h
Blood24h
BloodTwo24h
Blood48h
Blood72h
WholeF
CarcassF
CarcassM
WholeM
GutMidF
GutMidM
SalivF
SalivM
GutHind
GutAnt
GutCaeca
GutPost
Blood3h
BloodNBF
BloodTwoNBF
FatBodyBF
GutMidBF

| | GO.ID | BPCluster: indianred4 Size: 29 | Annotated | Significant | Expected | Rank in ClassicF | Weight01F | ClassicF |
|---|---|---|---|---|---|---|---|---|
| *1* | GO:0046331 | lateral inhibition | 92 | 3 | 0.29 | 11 | 0.0028 | 0.0028 |
| *2* | GO:0007030 | Golgi organization | 37 | 2 | 0.12 | 18 | 0.0058 | 0.0058 |
| *3* | GO:0018108 | peptidyl−tyrosine phosphorylation | 38 | 2 | 0.12 | 21 | 0.0061 | 0.0061 |
| *4* | GO:0010389 | regulation of G2/M transition of mitotic... | 48 | 2 | 0.15 | 26 | 0.0119 | 0.0097 |

| | GO.ID | MFCluster: indianred4 Size: 29 | Annotated | Significant | Expected | Rank in ClassicF | Weight01F | ClassicF |
|---|---|---|---|---|---|---|---|---|
| *1* | GO:0043138 | 3'−5' DNA helicase activity | 11 | 2 | 0.04 | 1 | 0.00053 | 0.00053 |
| *9* | GO:0004713 | protein tyrosine kinase activity | 36 | 2 | 0.11 | 5 | 0.07662 | 0.00574 |

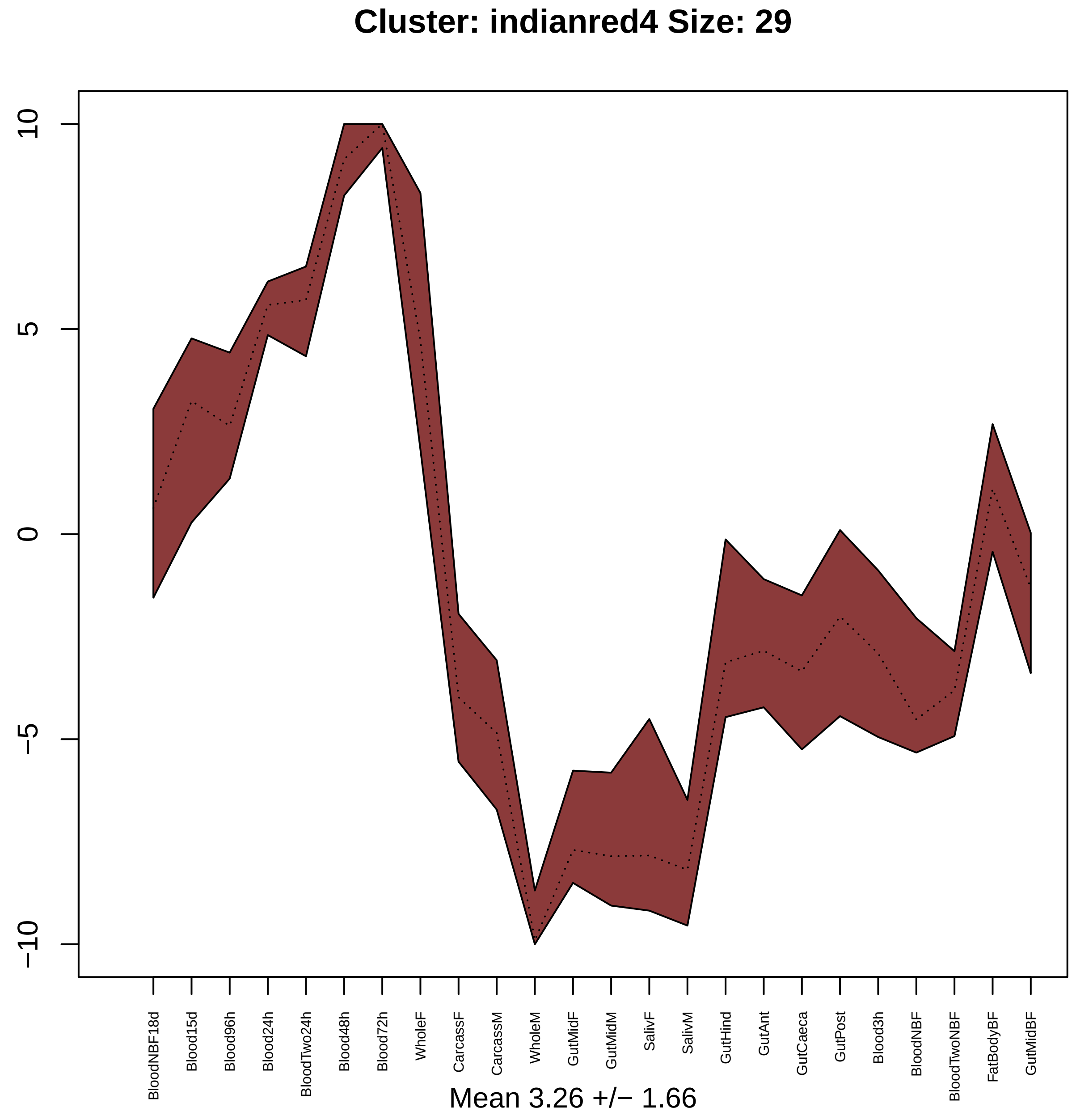

Cluster: indianred4 Size: 29
10
5
0
−5
−10
BloodNBF18d
Blood15d
Blood96h
Blood24h
BloodTwo24h
Blood48h
Blood72h
WholeF
CarcassF
CarcassM
WholeM
GutMidF
GutMidM
SalivF
SalivM
GutHind
GutAnt
GutCaeca
GutPost
Blood3h
BloodNBF
BloodTwoNBF
FatBodyBF
GutMidBF
Mean 3.26 +/− 1.66

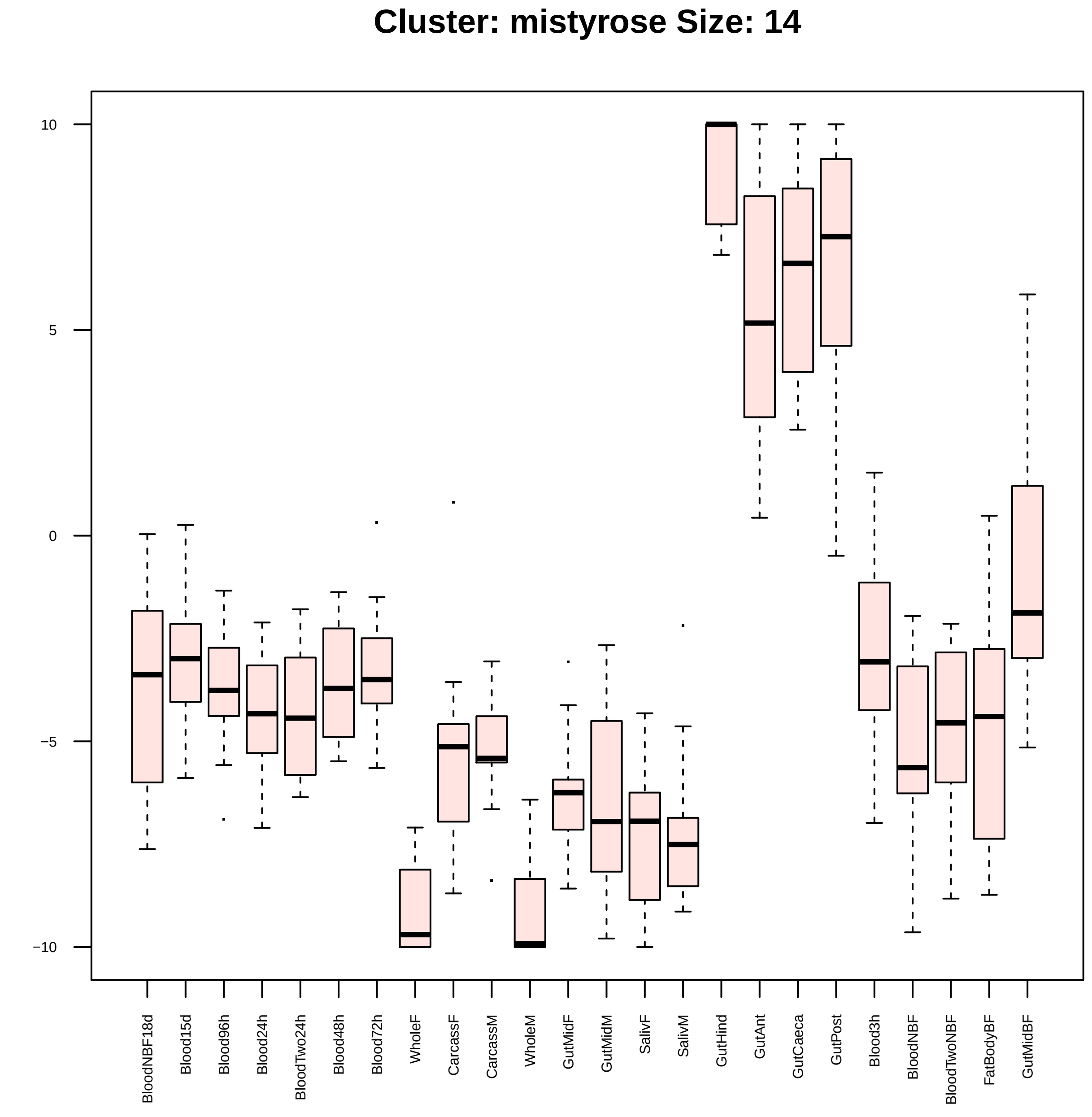
Cluster: mistyrose Size: 14
10
5
0
−5
−10
BloodNBF18d
Blood15d
Blood96h
Blood24h
BloodTwo24h
Blood48h
Blood72h
WholeF
CarcassF
CarcassM
WholeM
GutMidF
GutMidM
SalivF
SalivM
GutHind
GutAnt
GutCaeca
GutPost
Blood3h
BloodNBF
BloodTwoNBF
FatBodyBF
GutMidBF

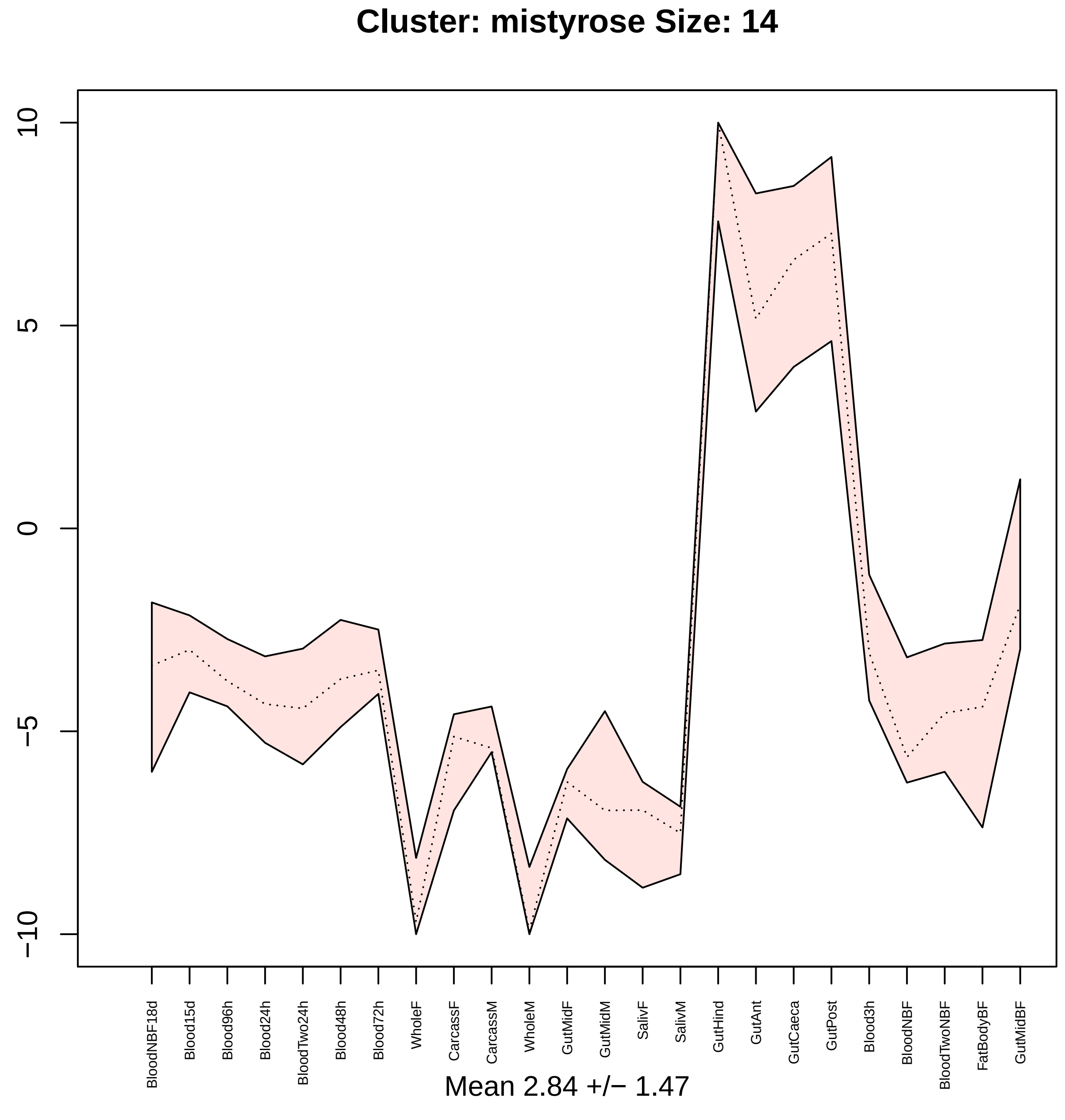

Cluster: mistyrose Size: 14
10
5
0
−5
−10
BloodNBF18d
Blood15d
Blood96h
Blood24h
BloodTwo24h
Blood48h
Blood72h
WholeF
CarcassF
CarcassM
WholeM
GutMidF
GutMidM
SalivF
SalivM
GutHind
GutAnt
GutCaeca
GutPost
Blood3h
BloodNBF
BloodTwoNBF
FatBodyBF
GutMidBF
Mean 2.84 +/− 1.47

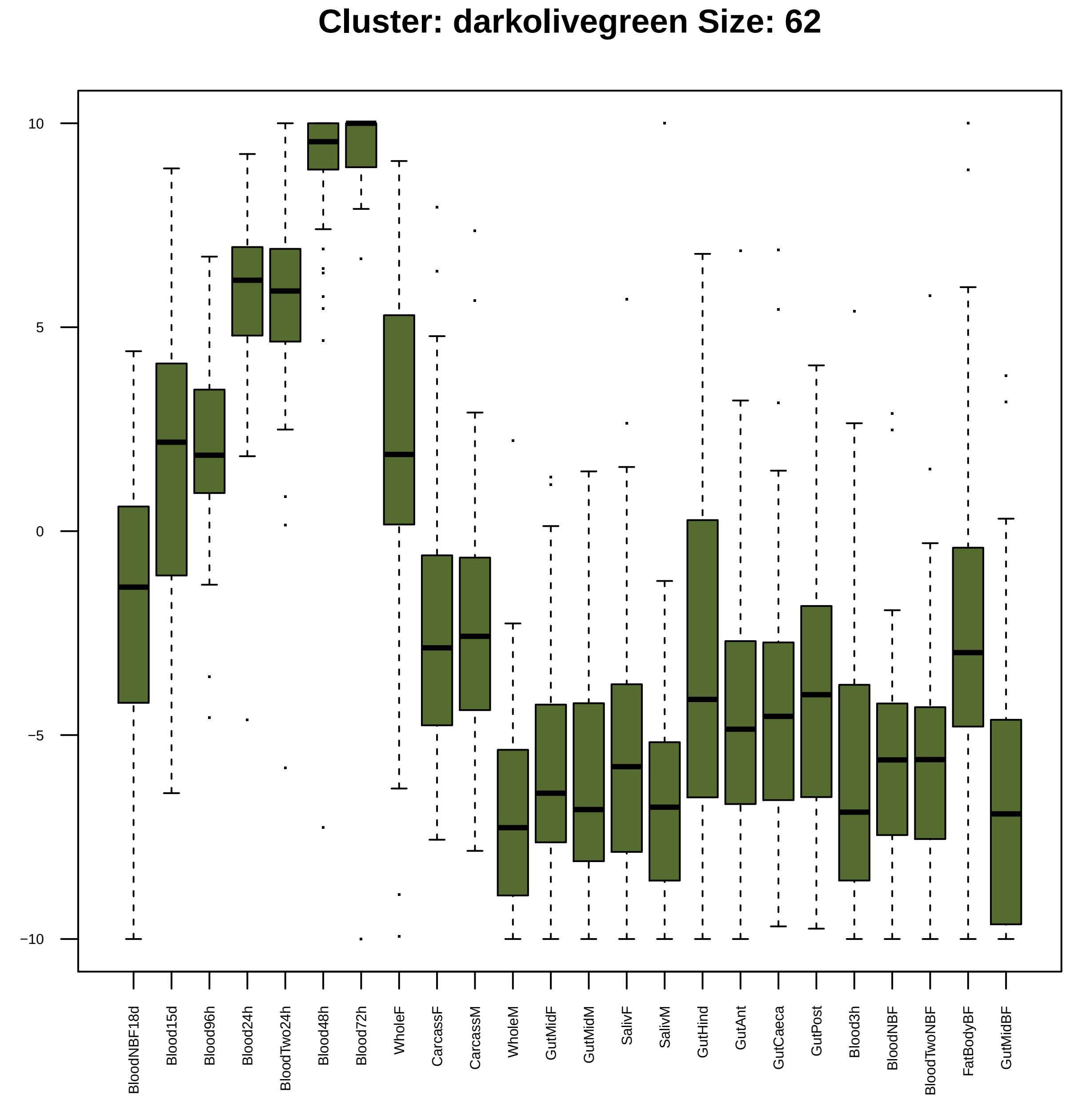
Cluster: darkolivegreen Size: 62
10
5
0
−5
−10
BloodNBF18d
Blood15d
Blood96h
Blood24h
BloodTwo24h
Blood48h
Blood72h
WholeF
CarcassF
CarcassM
WholeM
GutMidF
GutMidM
SalivF
SalivM
GutHind
GutAnt
GutCaeca
GutPost
Blood3h
BloodNBF
BloodTwoNBF
FatBodyBF
GutMidBF

| | GO.ID | BPCluster: darkolivegreen Size: 62 | Annotated | Significant | Expected | Rank in ClassicF | Weight01F | ClassicF |
|---|---|---|---|---|---|---|---|---|
| *1* | GO:0007093 | mitotic cell cycle checkpoint | 56 | 3 | 0.30 | 11 | 0.00093 | 0.00317 |
| *2* | GO:0007346 | regulation of mitotic cell cycle | 114 | 5 | 0.61 | 3 | 0.00238 | 0.00031 |
| *3* | GO:0030334 | regulation of cell migration | 14 | 2 | 0.07 | 8 | 0.00241 | 0.00241 |
| *4* | GO:0051179 | localization | 1336 | 10 | 7.12 | 247 | 0.00496 | 0.15710 |
| *5* | GO:0022008 | neurogenesis | 706 | 10 | 3.76 | 10 | 0.01309 | 0.00286 |
| *11* | GO:0040012 | regulation of locomotion | 21 | 3 | 0.11 | 2 | 0.03469 | 0.00017 |

| | GO.ID | MFCluster: darkolivegreen Size: 62 | Annotated | Significant | Expected | Rank in ClassicF | Weight01F | ClassicF |
|---|---|---|---|---|---|---|---|---|
| *1* | GO:0008017 | microtubule binding | 55 | 4 | 0.33 | 1 | 0.0003 | 0.0003 |
| *5* | GO:0004536 | deoxyribonuclease activity | 19 | 2 | 0.11 | 4 | 0.0402 | 0.0056 |

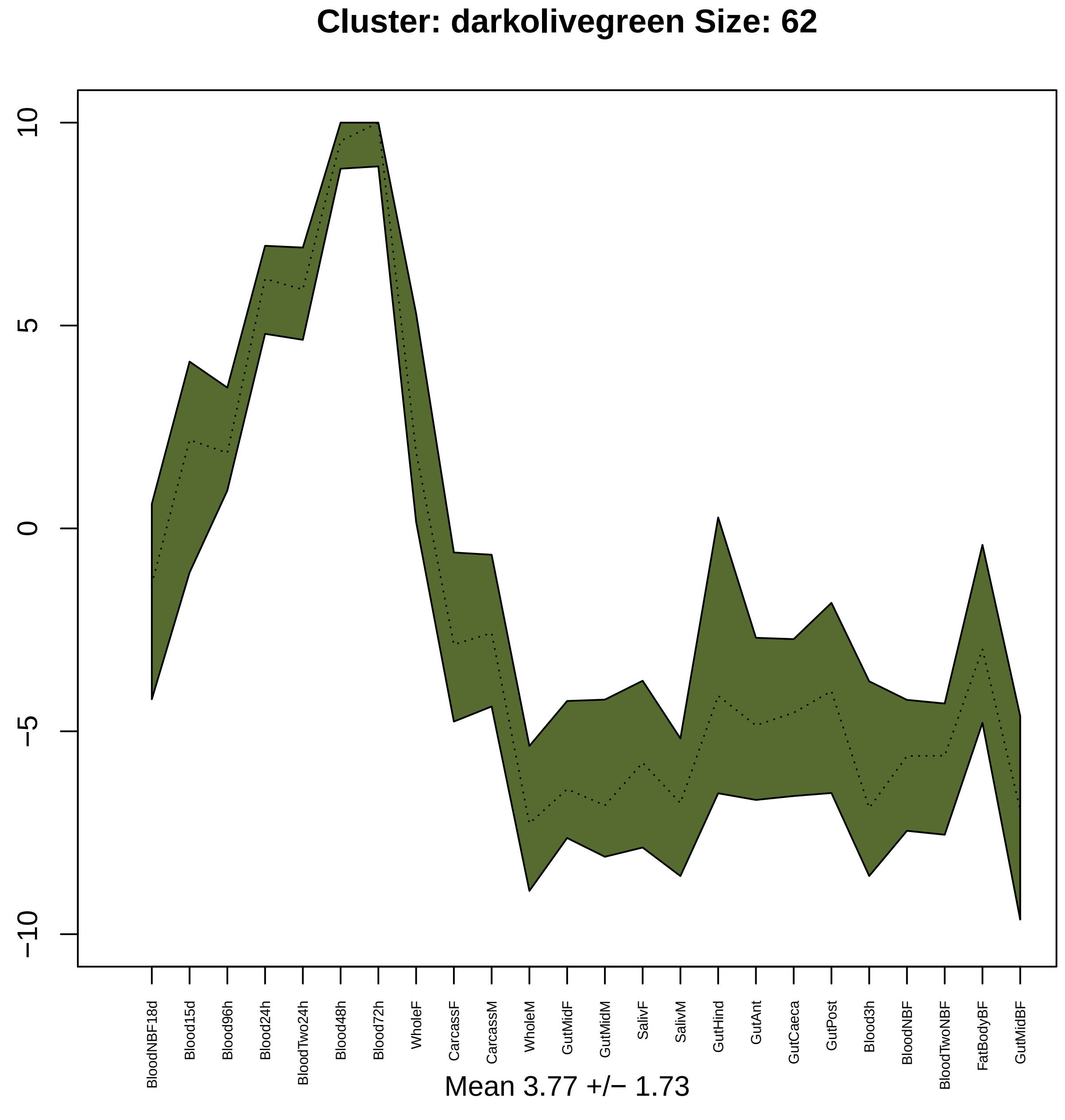
Cluster: darkolivegreen Size: 62
10
5
0
−5
−10
BloodNBF18d
Blood15d
Blood96h
Blood24h
BloodTwo24h
Blood48h
Blood72h
WholeF
CarcassF
CarcassM
WholeM
GutMidF
GutMidM
SalivF
SalivM
GutHind
GutAnt
GutCaeca
GutPost
Blood3h
BloodNBF
BloodTwoNBF
FatBodyBF
GutMidBF
Mean 3.77 +/− 1.73

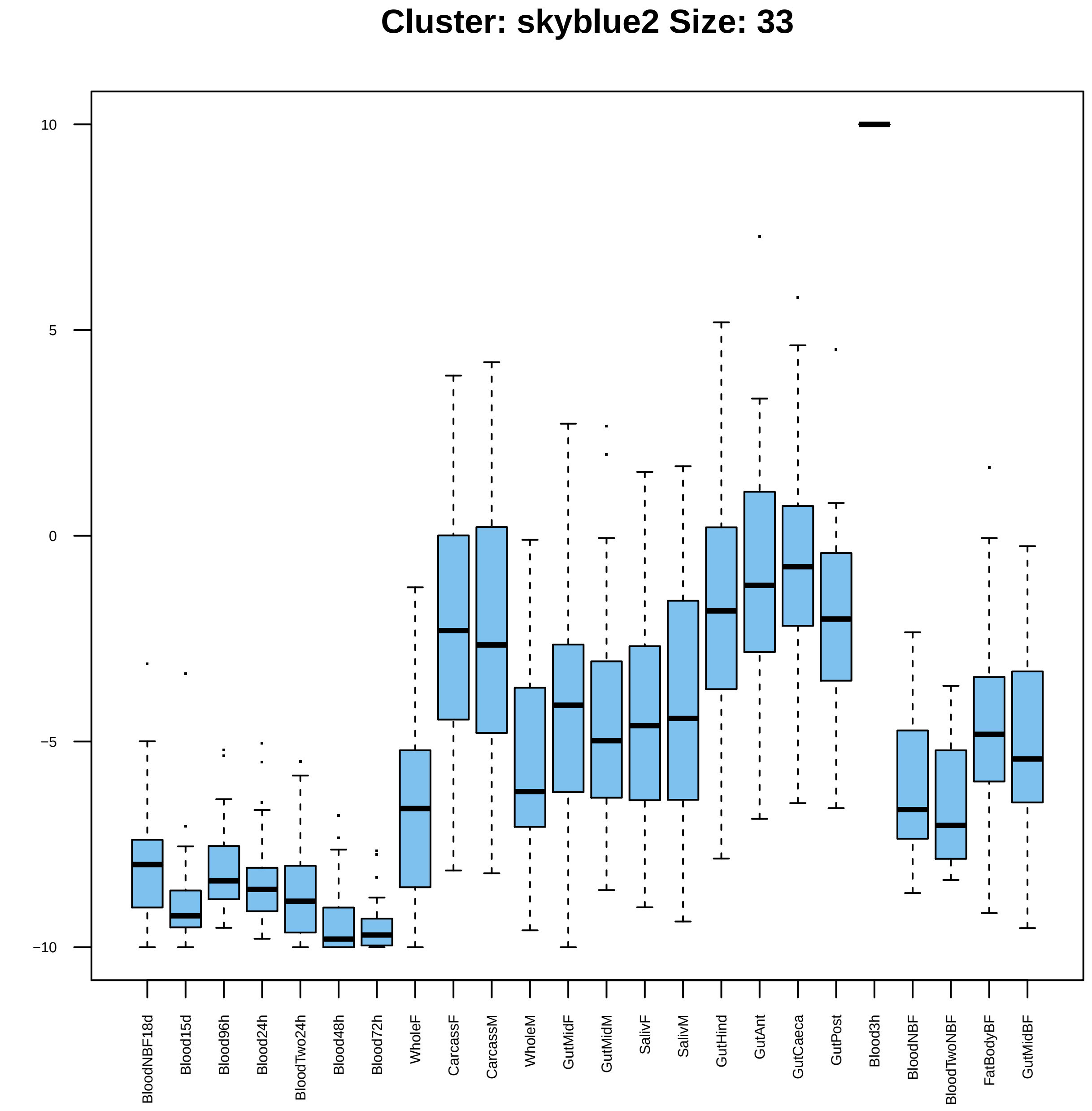

Cluster: skyblue2 Size: 33
10
5
0
−5
−10
BloodNBF18d
Blood15d
Blood96h
Blood24h
BloodTwo24h
Blood48h
Blood72h
WholeF
CarcassF
CarcassM
WholeM
GutMidF
GutMidM
SalivF
SalivM
GutHind
GutAnt
GutCaeca
GutPost
Blood3h
BloodNBF
BloodTwoNBF
FatBodyBF
GutMidBF

| | GO.ID | BPCluster: skyblue2 Size: 33 | Annotated | Significant | Expected | Rank in ClassicF | Weight01F | ClassicF |
|---|---|---|---|---|---|---|---|---|
| *3* | GO:0042737 | drug catabolic process | 48 | 2 | 0.08 | 1 | 0.034 | 0.0024 |
| *9* | GO:0006030 | chitin metabolic process | 95 | 2 | 0.15 | 3 | 0.096 | 0.0091 |
| *17* | GO:1901071 | glucosamine−containing compound metaboli... | 99 | 2 | 0.16 | 5 | 1.000 | 0.0099 |
| *22* | GO:1901564 | organonitrogen compound metabolic proces... | 1889 | 7 | 2.96 | 4 | 1.000 | 0.0098 |

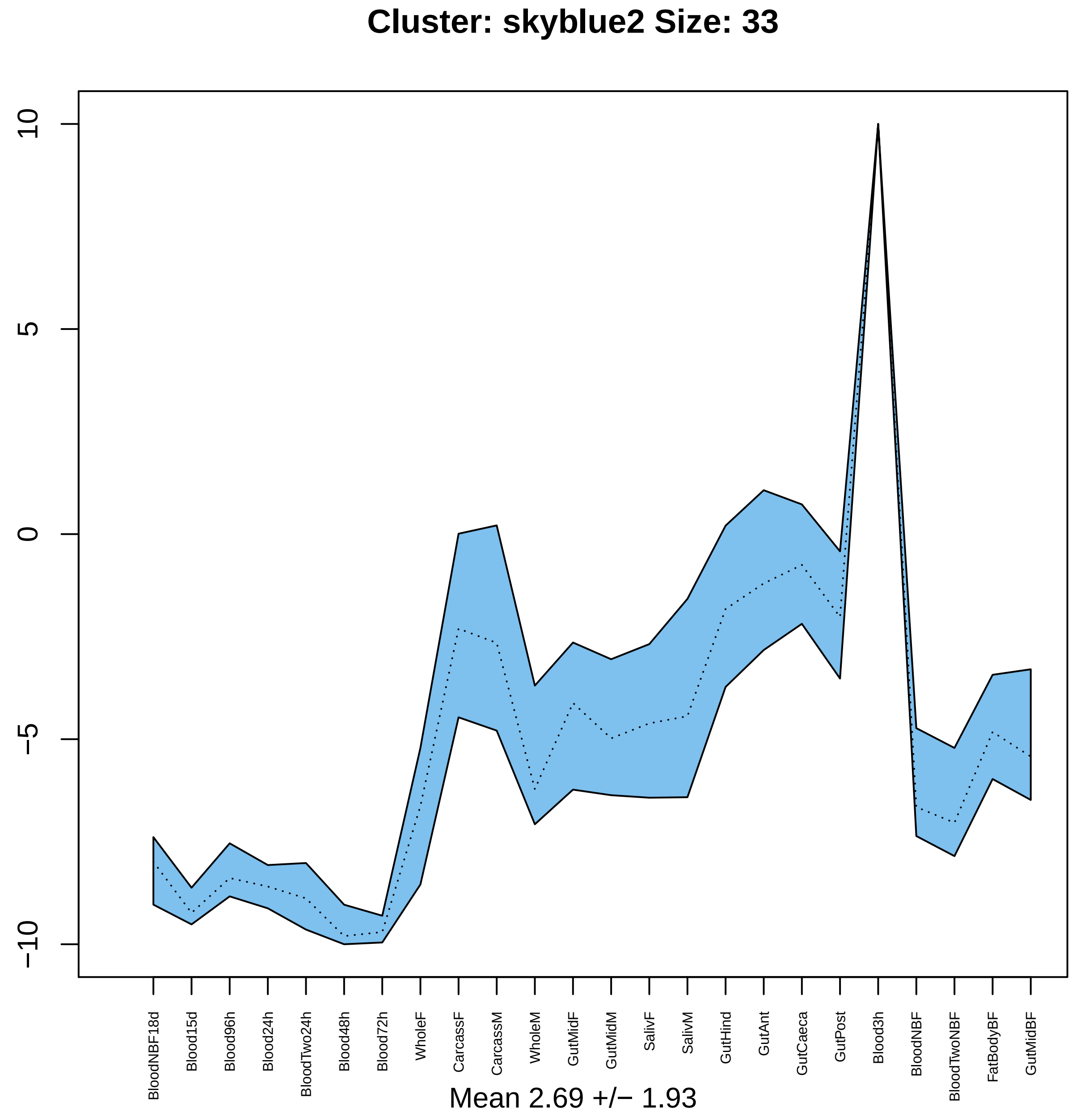

Cluster: skyblue2 Size: 33
10
5
0
−5
−10
BloodNBF18d
Blood15d
Blood96h
Blood24h
BloodTwo24h
Blood48h
Blood72h
WholeF
CarcassF
CarcassM
WholeM
GutMidF
GutMidM
SalivF
SalivM
GutHind
GutAnt
GutCaeca
GutPost
Blood3h
BloodNBF
BloodTwoNBF
FatBodyBF
GutMidBF
Mean 2.69 +/− 1.93

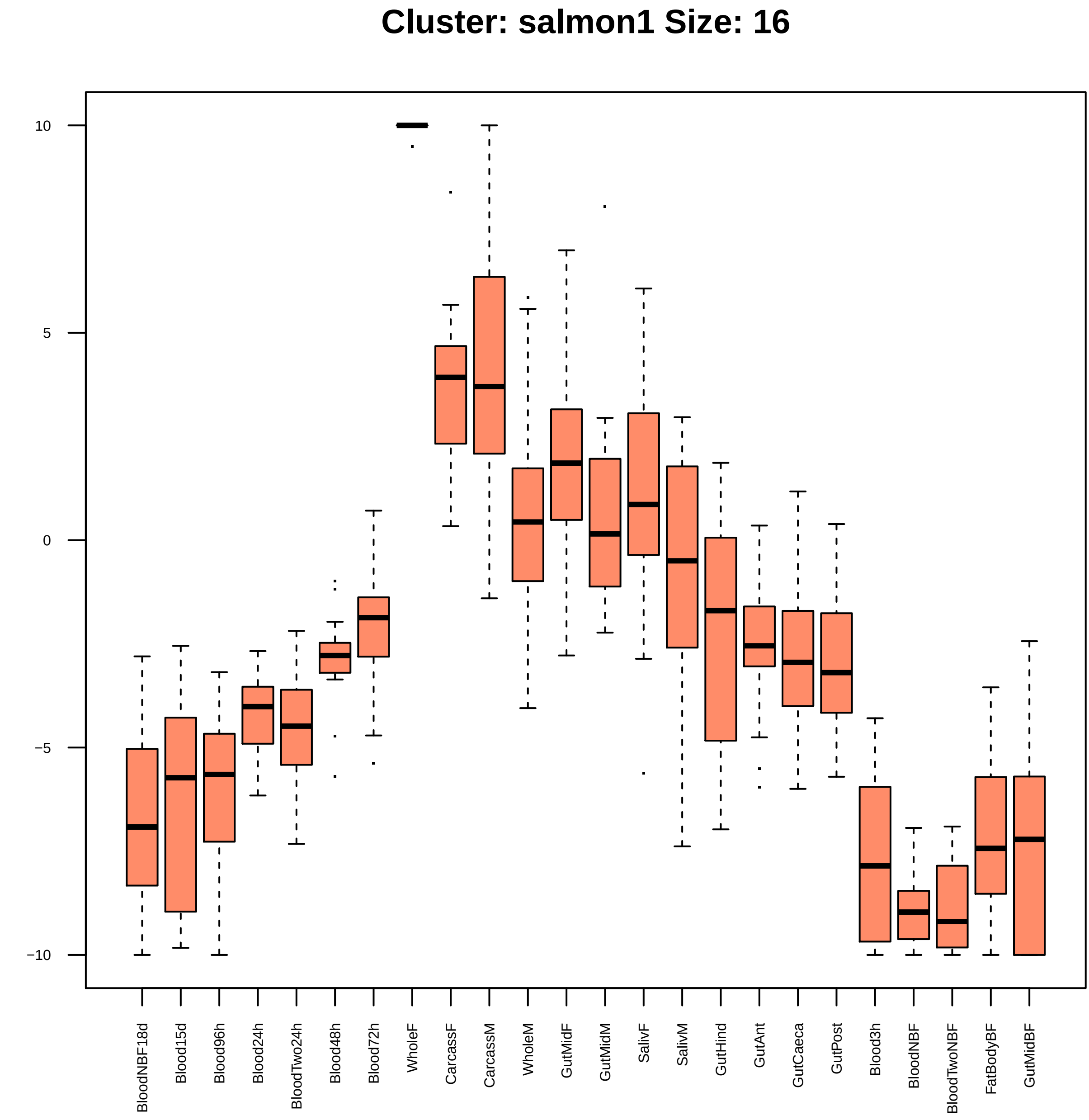

Cluster: salmon1 Size: 16
10
5
0
−5
−10
BloodNBF18d
Blood15d
Blood96h
Blood24h
BloodTwo24h
Blood48h
Blood72h
WholeF
CarcassF
CarcassM
WholeM
GutMidF
GutMidM
SalivF
SalivM
GutHind
GutAnt
GutCaeca
GutPost
Blood3h
BloodNBF
BloodTwoNBF
FatBodyBF
GutMidBF

| | GO.ID | BPCluster: salmon1 Size: 16 | Annotated | Significant | Expected | Rank in ClassicF | Weight01F | ClassicF |
|---|---|---|---|---|---|---|---|---|
| *1* | GO:0006338 | chromatin remodeling | 37 | 2 | 0.05 | 1 | 0.0011 | 0.0011 |
| *13* | GO:0006396 | RNA processing | 267 | 3 | 0.38 | 4 | 0.0932 | 0.0051 |
| *26* | GO:0090304 | nucleic acid metabolic process | 1059 | 5 | 1.49 | 5 | 1.0000 | 0.0088 |

| | GO.ID | MFCluster: salmon1 Size: 16 | Annotated | Significant | Expected | Rank in ClassicF | Weight01F | ClassicF |
|---|---|---|---|---|---|---|---|---|
| *1* | GO:0043138 | 3'−5' DNA helicase activity | 11 | 2 | 0.02 | 1 | 0.00012 | 0.00012 |
| *2* | GO:0004003 | ATP−dependent DNA helicase activity | 21 | 2 | 0.03 | 2 | 0.00044 | 0.00044 |
| *27* | GO:0140097 | catalytic activity, acting on DNA | 81 | 2 | 0.12 | 7 | 1.00000 | 0.00642 |

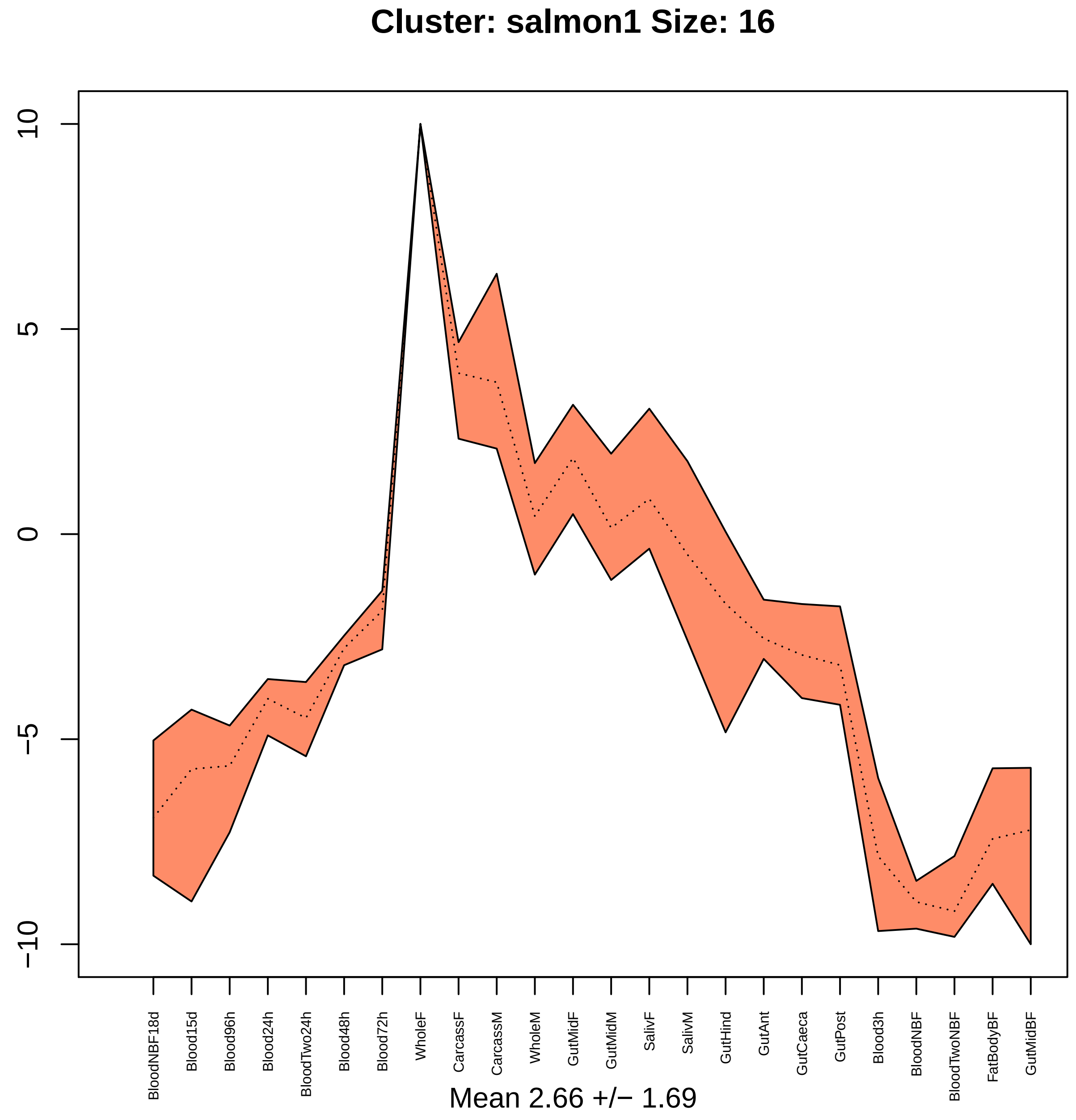
Cluster: salmon1 Size: 16
10
5
0
−5
−10
BloodNBF18d
Blood15d
Blood96h
Blood24h
BloodTwo24h
Blood48h
Blood72h
WholeF
CarcassF
CarcassM
WholeM
GutMidF
GutMidM
SalivF
SalivM
GutHind
GutAnt
GutCaeca
GutPost
Blood3h
BloodNBF
BloodTwoNBF
FatBodyBF
GutMidBF
Mean 2.66 +/− 1.69

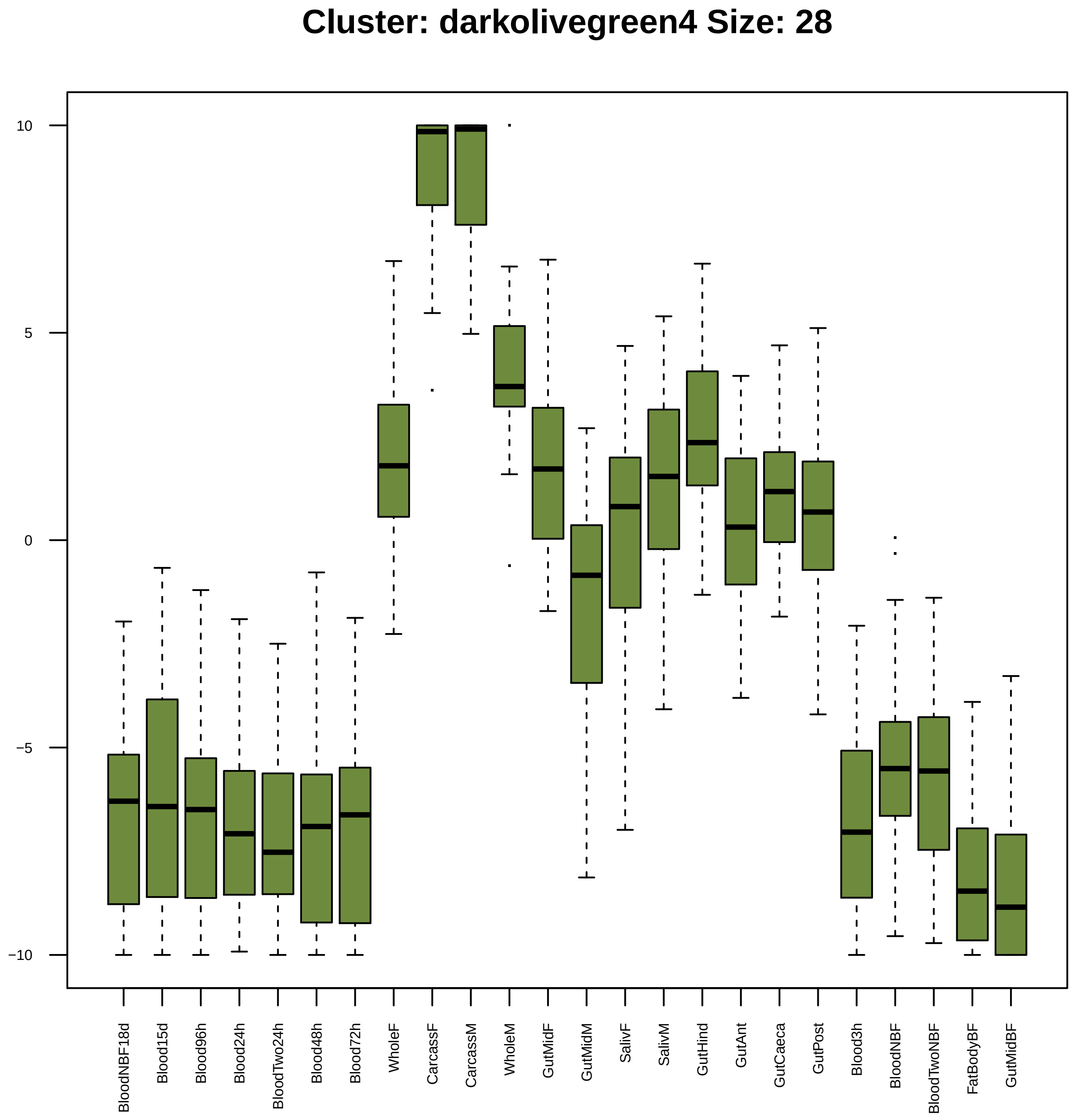
Cluster: darkolivegreen4 Size: 28
10
5
0
−5
−10
BloodNBF18d
Blood15d
Blood96h
Blood24h
BloodTwo24h
Blood48h
Blood72h
WholeF
CarcassF
CarcassM
WholeM
GutMidF
GutMidM
SalivF
SalivM
GutHind
GutAnt
GutCaeca
GutPost
Blood3h
BloodNBF
BloodTwoNBF
FatBodyBF
GutMidBF

| | GO.ID | BPCluster: darkolivegreen4 Size: 28 | Annotated | Significant | Expected | Rank in ClassicF | Weight01F | ClassicF |
|---|---|---|---|---|---|---|---|---|
| *24* | GO:0044249 | cellular biosynthetic process | 1306 | 11 | 3.89 | 12 | 1.000 | 0.00038 |

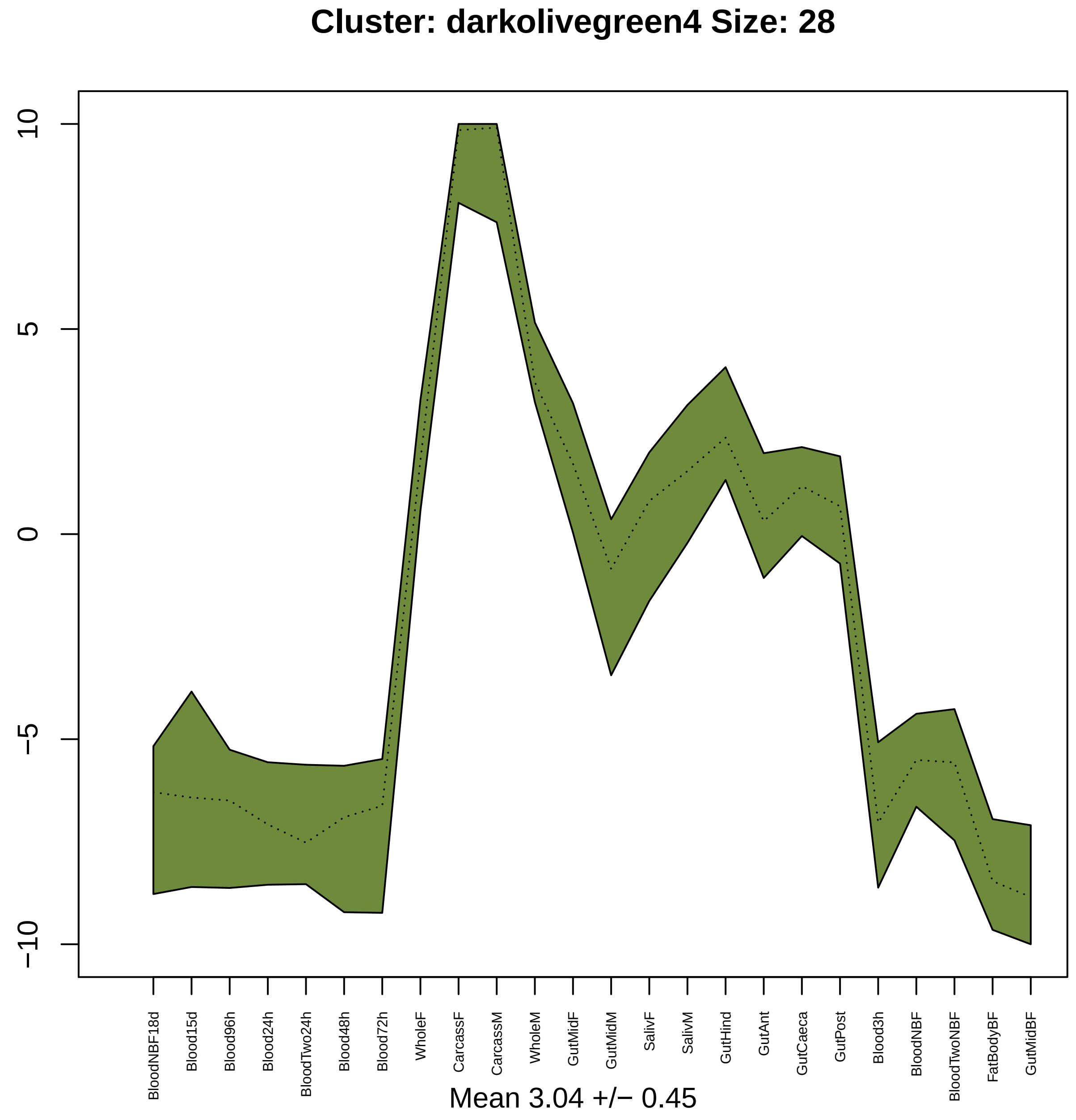

Cluster: darkolivegreen4 Size: 28
10
5
0
−5
−10
BloodNBF18d
Blood15d
Blood96h
Blood24h
BloodTwo24h
Blood48h
Blood72h
WholeF
CarcassF
CarcassM
WholeM
GutMidF
GutMidM
SalivF
SalivM
GutHind
GutAnt
GutCaeca
GutPost
Blood3h
BloodNBF
BloodTwoNBF
FatBodyBF
GutMidBF
Mean 3.04 +/− 0.45

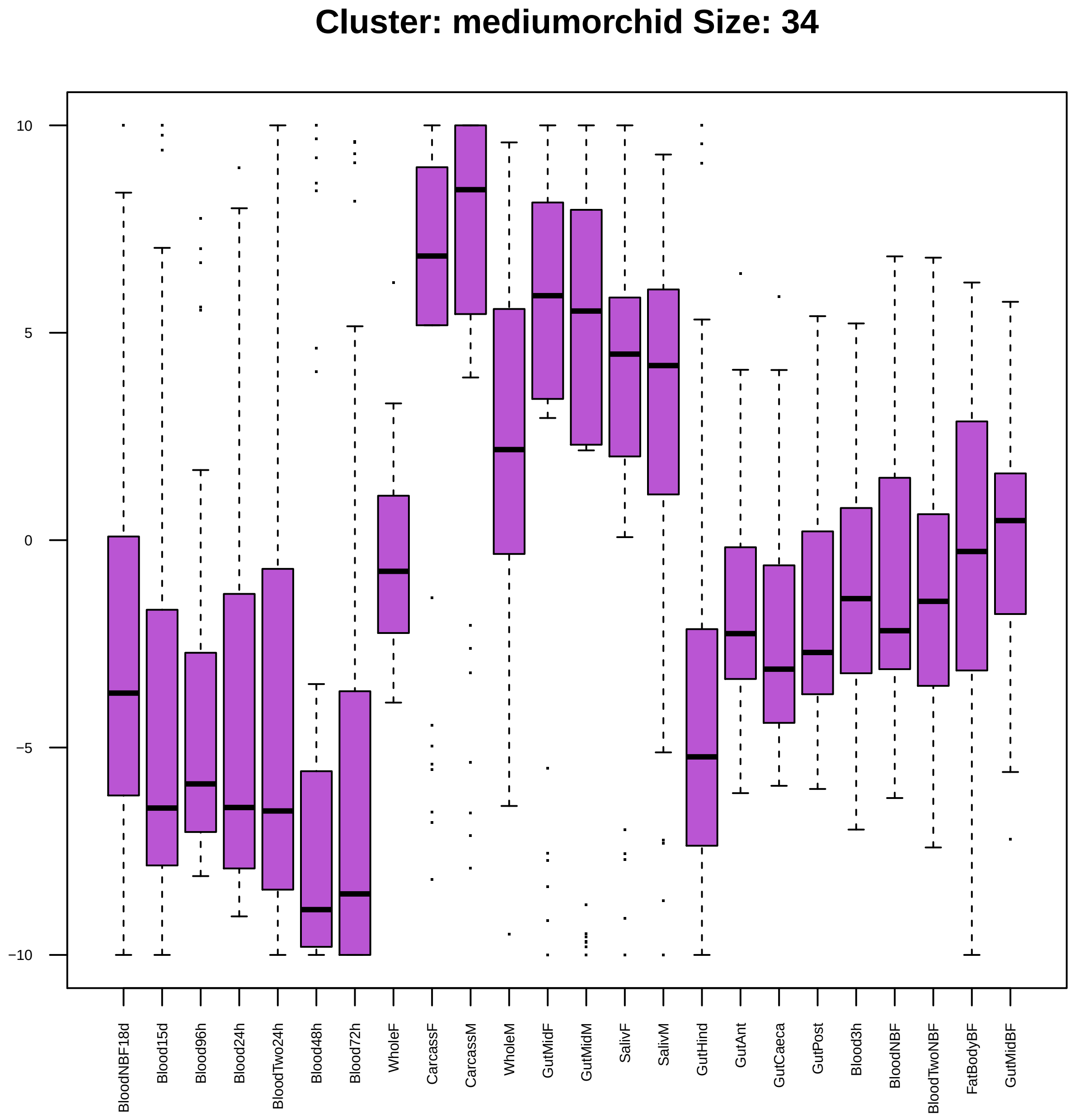
Cluster: mediumorchid Size: 34
10
5
0
−5
−10
BloodNBF18d
Blood15d
Blood96h
Blood24h
BloodTwo24h
Blood48h
Blood72h
WholeF
CarcassF
CarcassM
WholeM
GutMidF
GutMidM
SalivF
SalivM
GutHind
GutAnt
GutCaeca
GutPost
Blood3h
BloodNBF
BloodTwoNBF
FatBodyBF
GutMidBF

| | GO.ID | BPCluster: mediumorchid Size: 34 | Annotated | Significant | Expected | Rank in ClassicF | Weight01F | ClassicF |
|---|---|---|---|---|---|---|---|---|
| 1 | GO:0007430 | terminal branching, open tracheal system | 12 | 2 | 0.04 | 2 | 0.00067 | 0.00067 |
| 2 | GO:0042059 | negative regulation of epidermal growth ... | 19 | 2 | 0.06 | 9 | 0.00171 | 0.00171 |
| 3 | GO:0008355 | olfactory learning | 26 | 2 | 0.09 | 18 | 0.00320 | 0.00320 |
| 4 | GO:0006633 | fatty acid biosynthetic process | 32 | 2 | 0.11 | 26 | 0.00483 | 0.00483 |
| 8 | GO:0022603 | regulation of anatomical structure morph... | 133 | 3 | 0.44 | 48 | 0.02407 | 0.00896 |
| 9 | GO:0007613 | memory | 34 | 2 | 0.11 | 29 | 0.02493 | 0.00544 |
| 15 | GO:2000241 | regulation of reproductive process | 30 | 2 | 0.10 | 23 | 0.03409 | 0.00425 |
| 20 | GO:0045595 | regulation of cell differentiation | 112 | 3 | 0.37 | 32 | 0.03576 | 0.00557 |

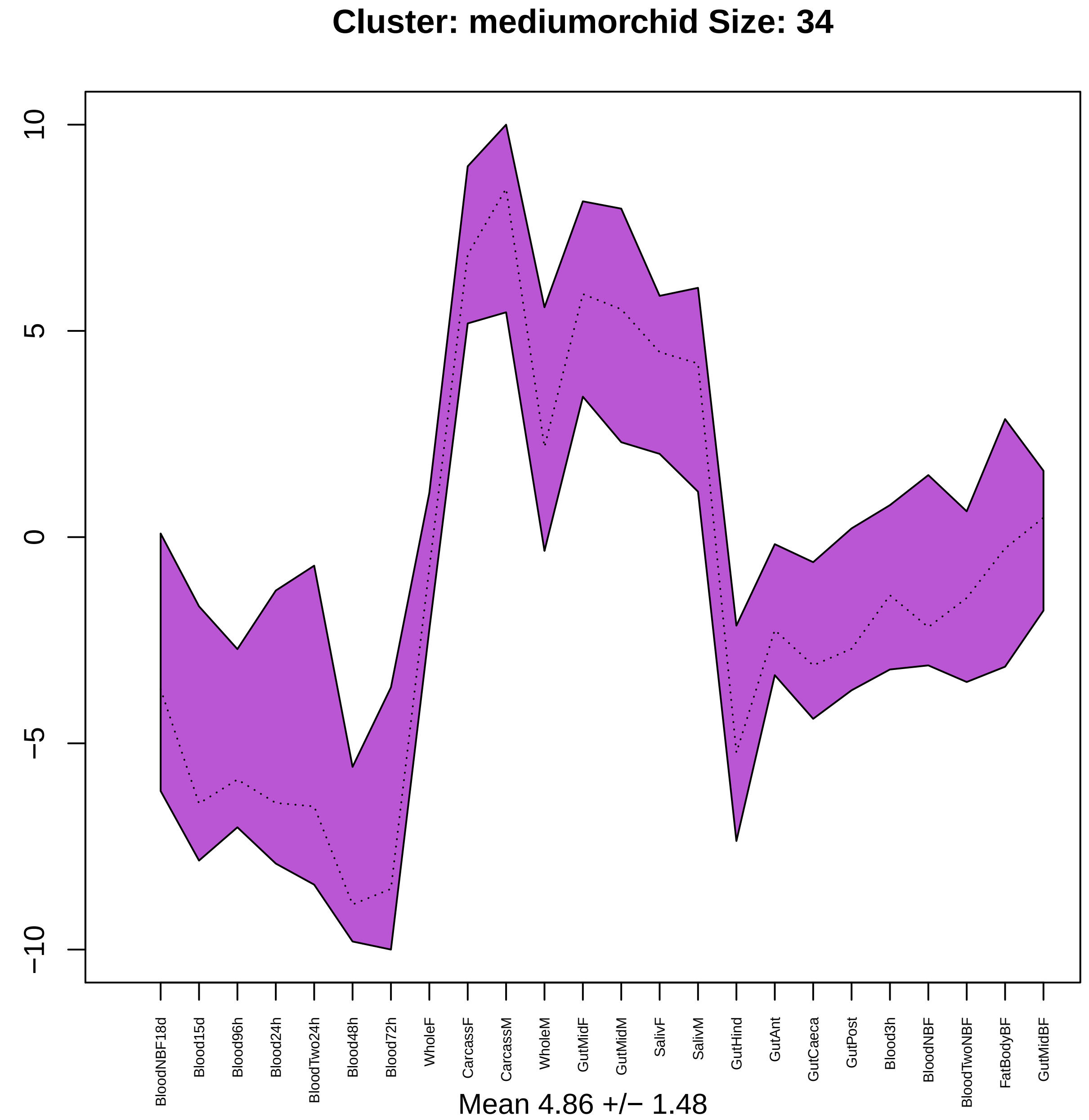

Cluster: mediumorchid Size: 34
10
5
0
−5
−10
BloodNBF18d
Blood15d
Blood96h
Blood24h
BloodTwo24h
Blood48h
Blood72h
WholeF
CarcassF
CarcassM
WholeM
GutMidF
GutMidM
SalivF
SalivM
GutHind
GutAnt
GutCaeca
GutPost
Blood3h
BloodNBF
BloodTwoNBF
FatBodyBF
GutMidBF
Mean 4.86 +/− 1.48

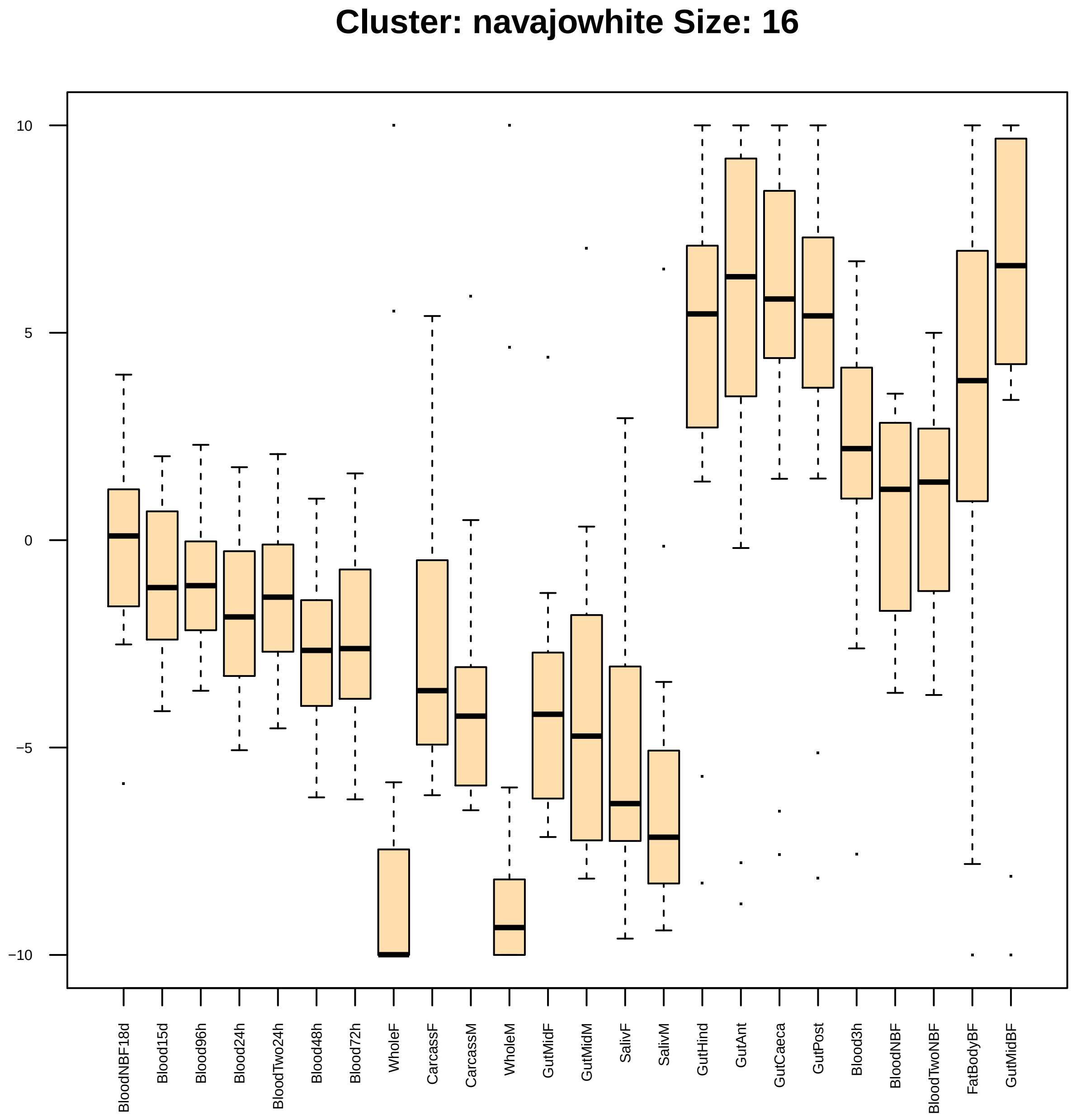

Cluster: navajowhite Size: 16
10
5
0
−5
−10
BloodNBF18d
Blood15d
Blood96h
Blood24h
BloodTwo24h
Blood48h
Blood72h
WholeF
CarcassF
CarcassM
WholeM
GutMidF
GutMidM
SalivF
SalivM
GutHind
GutAnt
GutCaeca
GutPost
Blood3h
BloodNBF
BloodTwoNBF
FatBodyBF
GutMidBF

| | GO.ID | BPCluster: navajowhite Size: 16 | Annotated | Significant | Expected | Rank in ClassicF | Weight01F | ClassicF |
|---|---|---|---|---|---|---|---|---|
| *1* | GO:0006605 | protein targeting | 47 | 2 | 0.07 | 2 | 0.0019 | 0.0019 |
| *2* | GO:0017038 | protein import | 35 | 2 | 0.05 | 1 | 0.0113 | 0.0010 |
| *11* | GO:0072594 | establishment of protein localization to... | 55 | 2 | 0.08 | 3 | 0.0360 | 0.0025 |

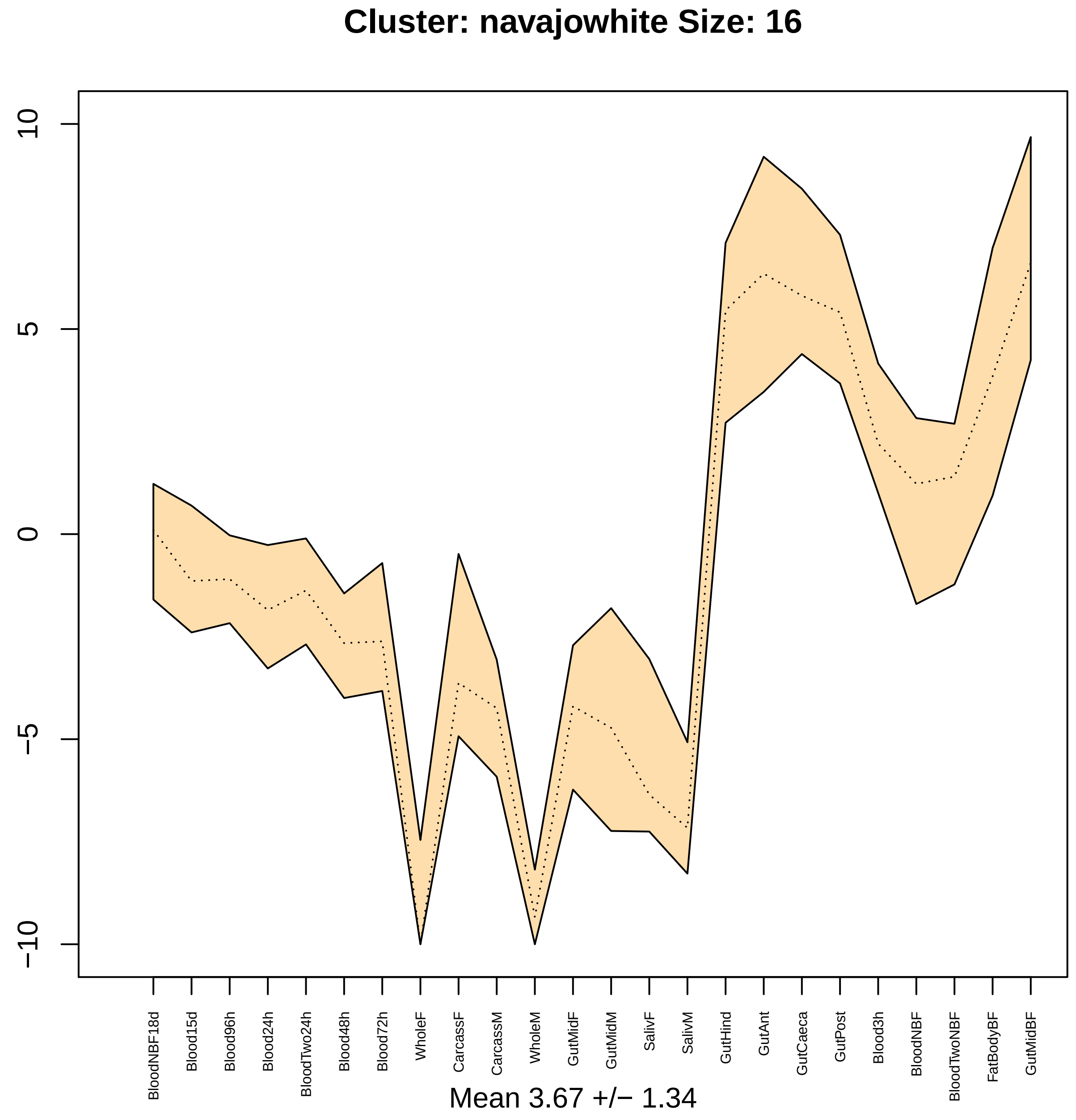

Cluster: navajowhite Size: 16
10
5
0
−5
−10
BloodNBF18d
Blood15d
Blood96h
Blood24h
BloodTwo24h
Blood48h
Blood72h
WholeF
CarcassF
CarcassM
WholeM
GutMidF
GutMidM
SalivF
SalivM
GutHind
GutAnt
GutCaeca
GutPost
Blood3h
BloodNBF
BloodTwoNBF
FatBodyBF
GutMidBF
Mean 3.67 +/− 1.34

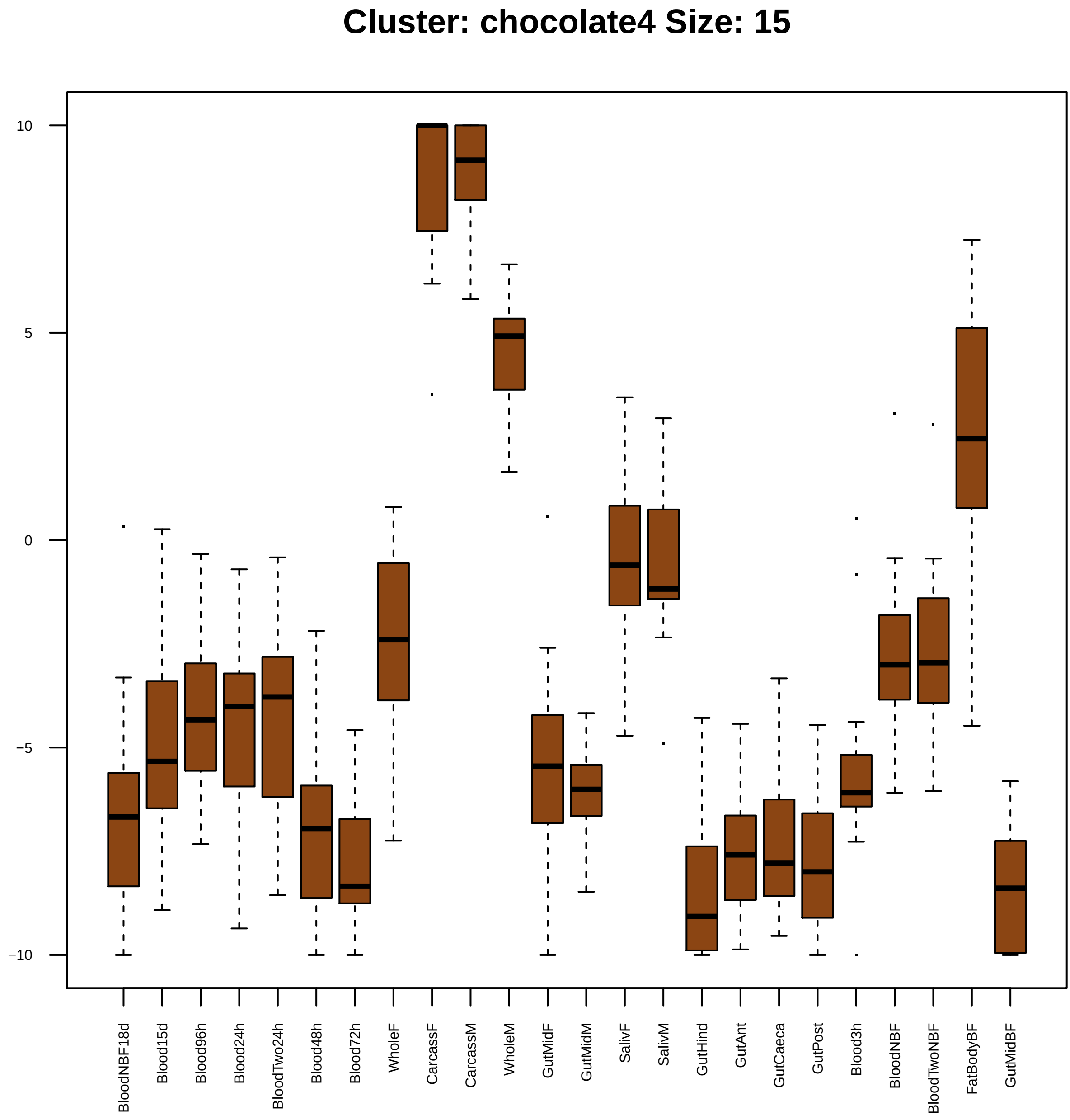

Cluster: chocolate4 Size: 15
10
5
0
−5
−10
BloodNBF18d
Blood15d
Blood96h
Blood24h
BloodTwo24h
Blood48h
Blood72h
WholeF
CarcassF
CarcassM
WholeM
GutMidF
GutMidM
SalivF
SalivM
GutHind
GutAnt
GutCaeca
GutPost
Blood3h
BloodNBF
BloodTwoNBF
FatBodyBF
GutMidBF

| | GO.ID | BPCluster: chocolate4 Size: 15 | Annotated | Significant | Expected | Rank in ClassicF | Weight01F | ClassicF |
|---|---|---|---|---|---|---|---|---|
| *1* | GO:0006508 | proteolysis | 594 | 4 | 0.56 | 1 | 0.0049 | 0.00096 |

| | GO.ID | MFCluster: chocolate4 Size: 15 | Annotated | Significant | Expected | Rank in ClassicF | Weight01F | ClassicF |
|---|---|---|---|---|---|---|---|---|
| 1 | GO:0004252 | serine−type endopeptidase activity | 252 | 3 | 0.35 | 2 | 0.0042 | 0.0042 |
| 3 | GO:0008233 | peptidase activity | 500 | 4 | 0.69 | 1 | 0.0308 | 0.0034 |
| 9 | GO:0008236 | serine−type peptidase activity | 278 | 3 | 0.39 | 3 | 1.0000 | 0.0056 |
| 20 | GO:0017171 | serine hydrolase activity | 279 | 3 | 0.39 | 4 | 1.0000 | 0.0056 |

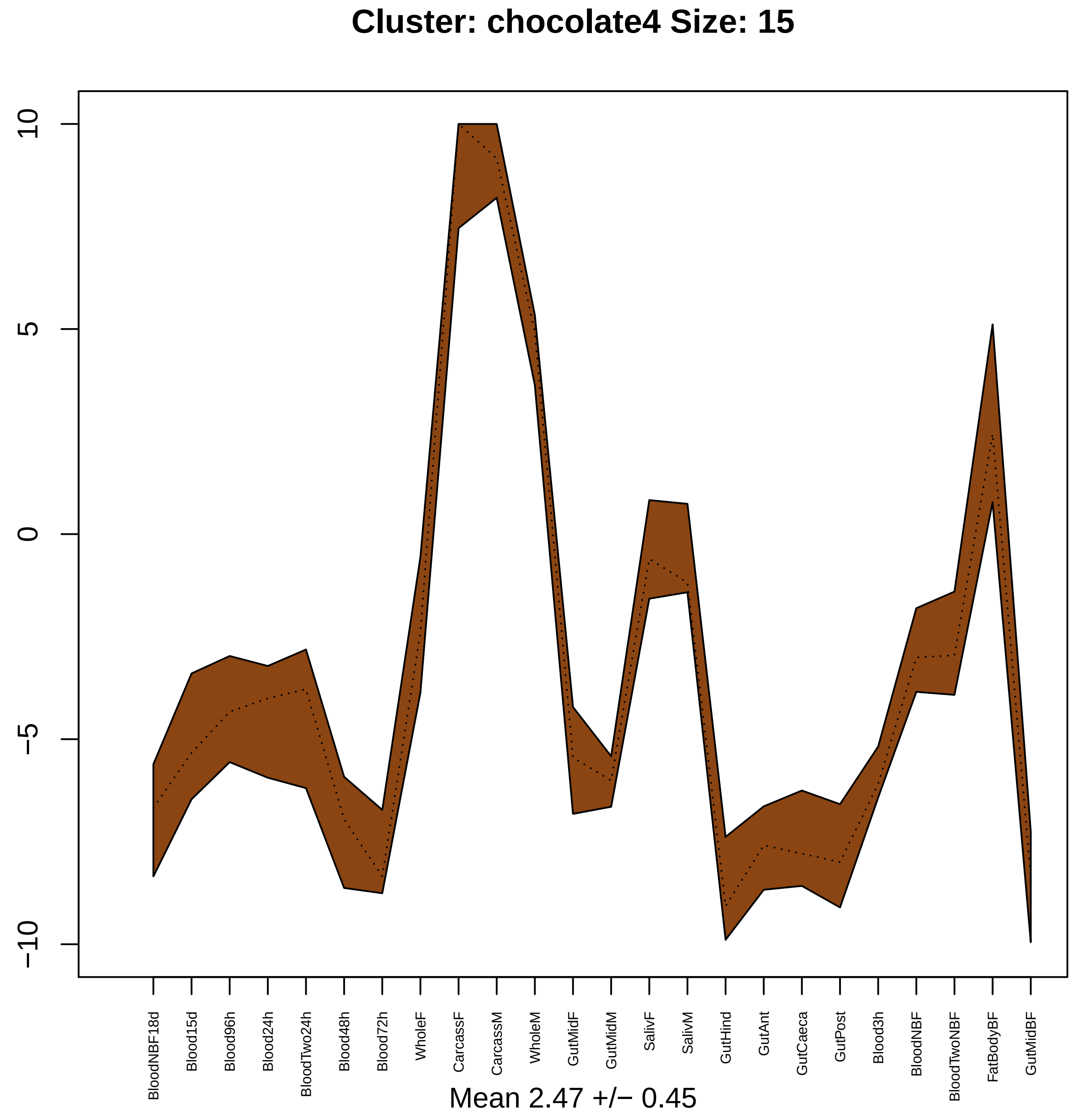

Cluster: chocolate4 Size: 15
10
5
0
−5
−10
BloodNBF18d
Blood15d
Blood96h
Blood24h
BloodTwo24h
Blood48h
Blood72h
WholeF
CarcassF
CarcassM
WholeM
GutMidF
GutMidM
SalivF
SalivM
GutHind
GutAnt
GutCaeca
GutPost
Blood3h
BloodNBF
BloodTwoNBF
FatBodyBF
GutMidBF
Mean 2.47 +/− 0.45

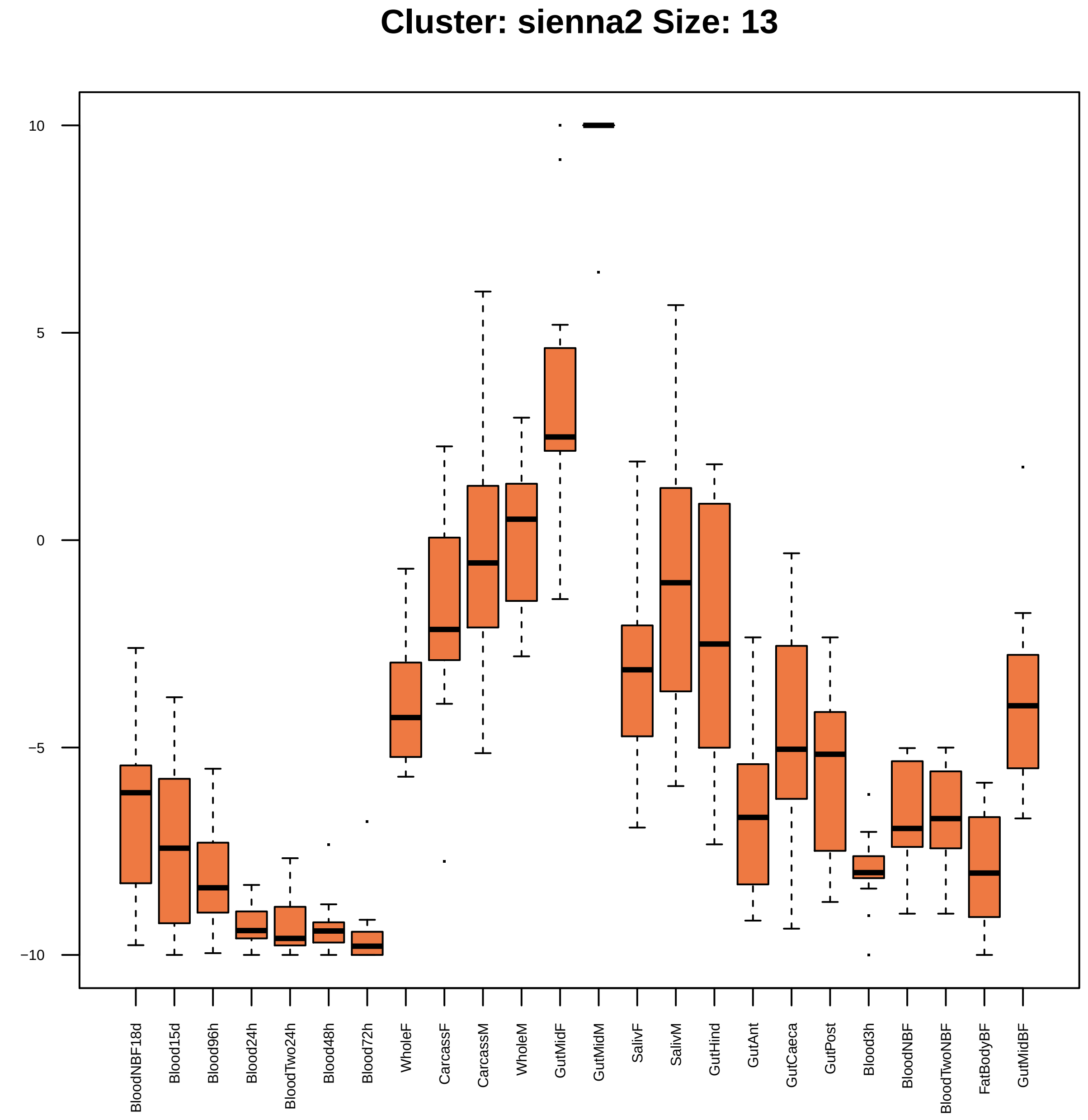

Cluster: sienna2 Size: 13
10
5
0
−5
−10
BloodNBF18d
Blood15d
Blood96h
Blood24h
BloodTwo24h
Blood48h
Blood72h
WholeF
CarcassF
CarcassM
WholeM
GutMidF
GutMidM
SalivF
SalivM
GutHind
GutAnt
GutCaeca
GutPost
Blood3h
BloodNBF
BloodTwoNBF
FatBodyBF
GutMidBF

| | GO.ID | BPCluster: sienna2 Size: 13 | Annotated | Significant | Expected | Rank in ClassicF | Weight01F | ClassicF |
|---|---|---|---|---|---|---|---|---|
| *1* | GO:0035725 | sodium ion transmembrane transport | 46 | 2 | 0.08 | 2 | 0.0027 | 0.00269 |
| *4* | GO:0055085 | transmembrane transport | 469 | 5 | 0.81 | 1 | 0.0248 | 0.00067 |

|  | GO.ID | MFCluster: sienna2 Size: 13 | Annotated | Significant | Expected | Rank in ClassicF | Weight01F | ClassicF |
|---|---|---|---|---|---|---|---|---|
| *8* | GO:0015370 | solute:sodium symporter activity | 31 | 2 | 0.05 | 3 | 0.023 | 0.00096 |
| *12* | GO:0022857 | transmembrane transporter activity | 442 | 5 | 0.67 | 1 | 0.066 | 0.00029 |
| *28* | GO:0015291 | secondary active transmembrane transport... | 60 | 2 | 0.09 | 7 | 1.000 | 0.00357 |
| *30* | GO:0015293 | symporter activity | 37 | 2 | 0.06 | 5 | 1.000 | 0.00137 |

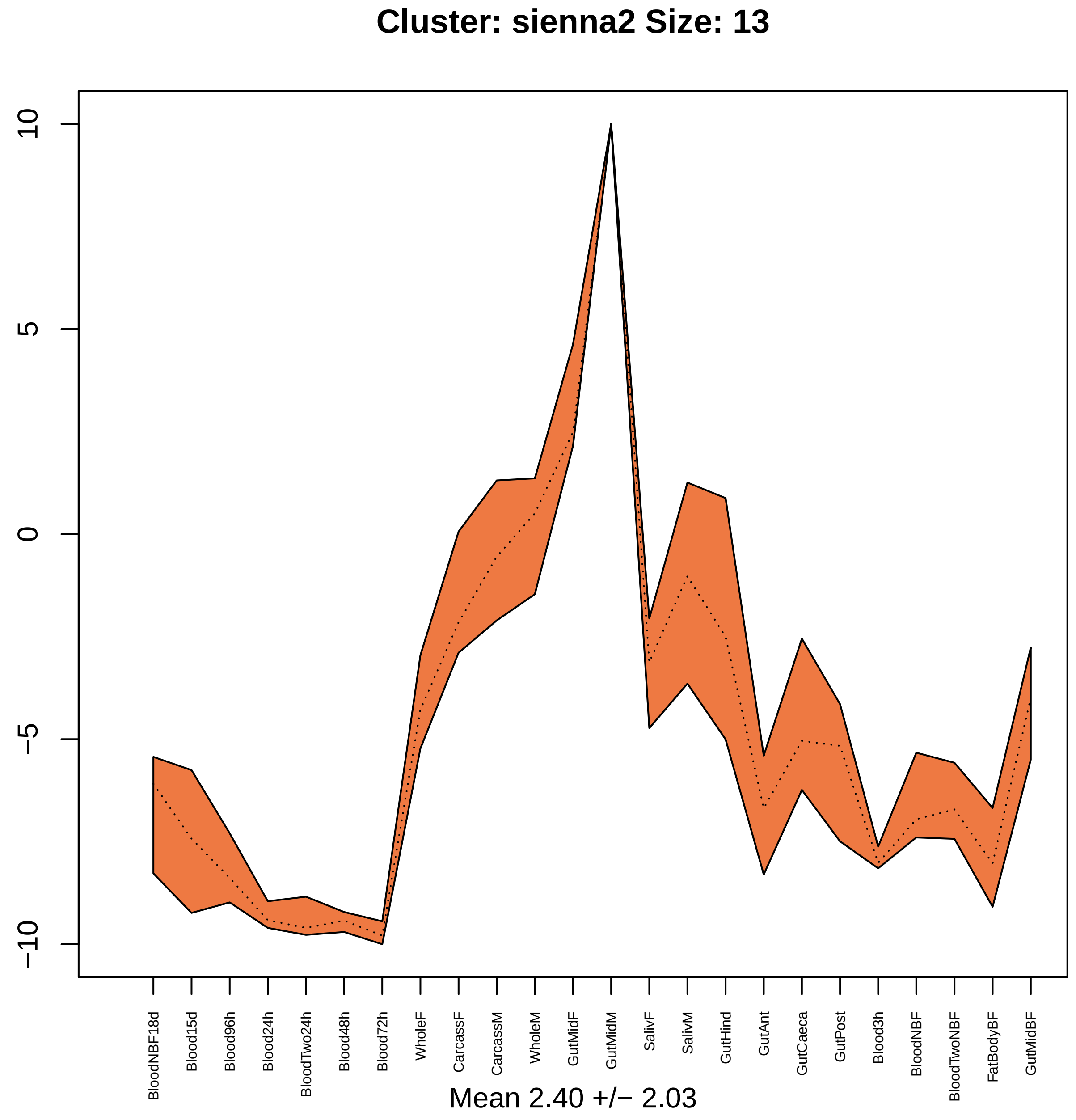

Cluster: sienna2 Size: 13
10
5
0
−5
−10
BloodNBF18d
Blood15d
Blood96h
Blood24h
BloodTwo24h
Blood48h
Blood72h
WholeF
CarcassF
CarcassM
WholeM
GutMidF
GutMidM
SalivF
SalivM
GutHind
GutAnt
GutCaeca
GutPost
Blood3h
BloodNBF
BloodTwoNBF
FatBodyBF
GutMidBF
Mean 2.40 +/− 2.03

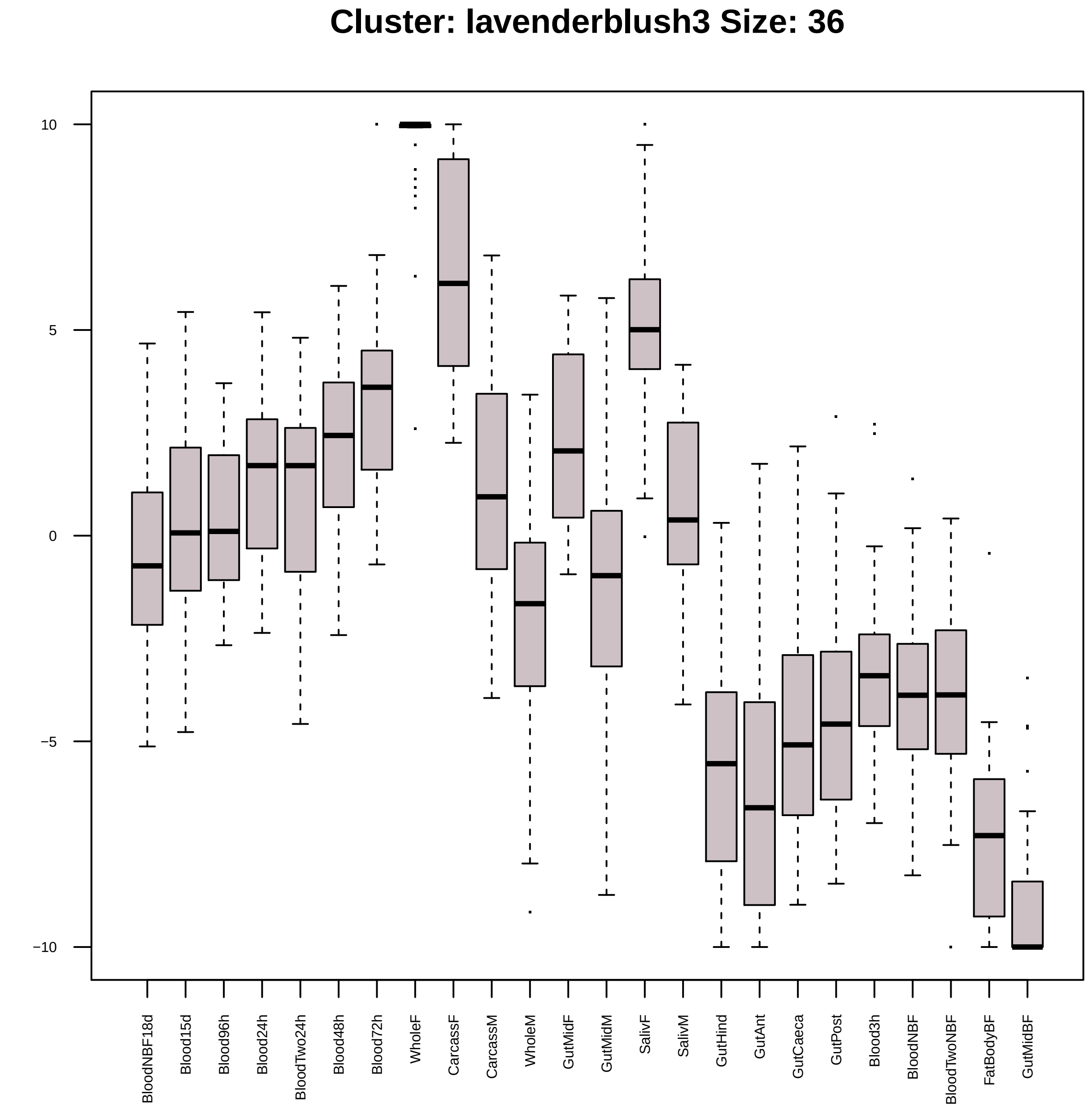
Cluster: lavenderblush3 Size: 36
10
5
0
−5
−10
BloodNBF18d
Blood15d
Blood96h
Blood24h
BloodTwo24h
Blood48h
Blood72h
WholeF
CarcassF
CarcassM
WholeM
GutMidF
GutMidM
SalivF
SalivM
GutHind
GutAnt
GutCaeca
GutPost
Blood3h
BloodNBF
BloodTwoNBF
FatBodyBF
GutMidBF

| | GO.ID | BPCluster: lavenderblush3 Size: 36 | Annotated | Significant | Expected | Rank in ClassicF | Weight01F | ClassicF |
|---|---|---|---|---|---|---|---|---|
| *3* | GO:0032508 | DNA duplex unwinding | 18 | 2 | 0.08 | 24 | 0.0031 | 0.00312 |
| *4* | GO:0010972 | negative regulation of G2/M transition o... | 44 | 2 | 0.21 | 37 | 0.0046 | 0.01791 |
| *5* | GO:0042254 | ribosome biogenesis | 64 | 4 | 0.30 | 13 | 0.0076 | 0.00021 |
| *6* | GO:0000398 | mRNA splicing, via spliceosome | 134 | 4 | 0.63 | 26 | 0.0095 | 0.00334 |
| *22* | GO:0006397 | mRNA processing | 164 | 5 | 0.77 | 20 | 0.1105 | 0.00090 |

| | GO.ID | MFCluster: lavenderblush3 Size: 36 | Annotated | Significant | Expected | Rank in ClassicF | Weight01F | ClassicF |
|---|---|---|---|---|---|---|---|---|
| 1 | GO:0003729 | mRNA binding | 99 | 4 | 0.45 | 6 | 0.0010 | 0.0010 |
| 2 | GO:0003676 | nucleic acid binding | 1152 | 17 | 5.27 | 1 | 0.0039 | 2.3e−06 |
| 3 | GO:0004003 | ATP−dependent DNA helicase activity | 21 | 2 | 0.10 | 7 | 0.0040 | 0.0040 |
| 4 | GO:0003899 | DNA−directed 5'−3' RNA polymerase activi... | 27 | 2 | 0.12 | 10 | 0.0066 | 0.0066 |
| 5 | GO:0003723 | RNA binding | 307 | 8 | 1.41 | 2 | 0.0079 | 5.4e−05 |

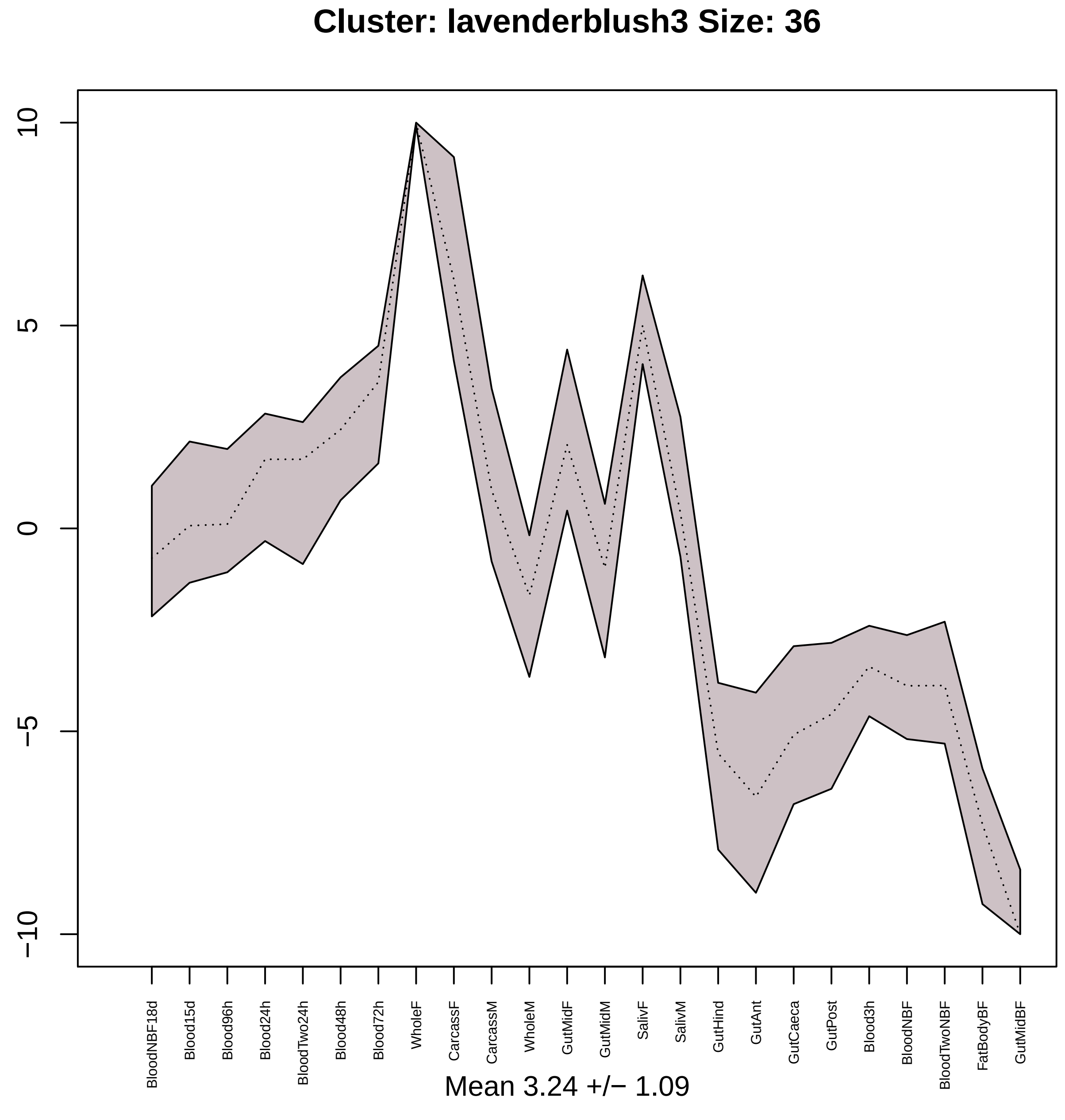

Cluster: lavenderblush3 Size: 36
10
5
0
−5
−10
BloodNBF18d
Blood15d
Blood96h
Blood24h
BloodTwo24h
Blood48h
Blood72h
WholeF
CarcassF
CarcassM
WholeM
GutMidF
GutMidM
SalivF
SalivM
GutHind
GutAnt
GutCaeca
GutPost
Blood3h
BloodNBF
BloodTwoNBF
FatBodyBF
GutMidBF
Mean 3.24 +/− 1.09

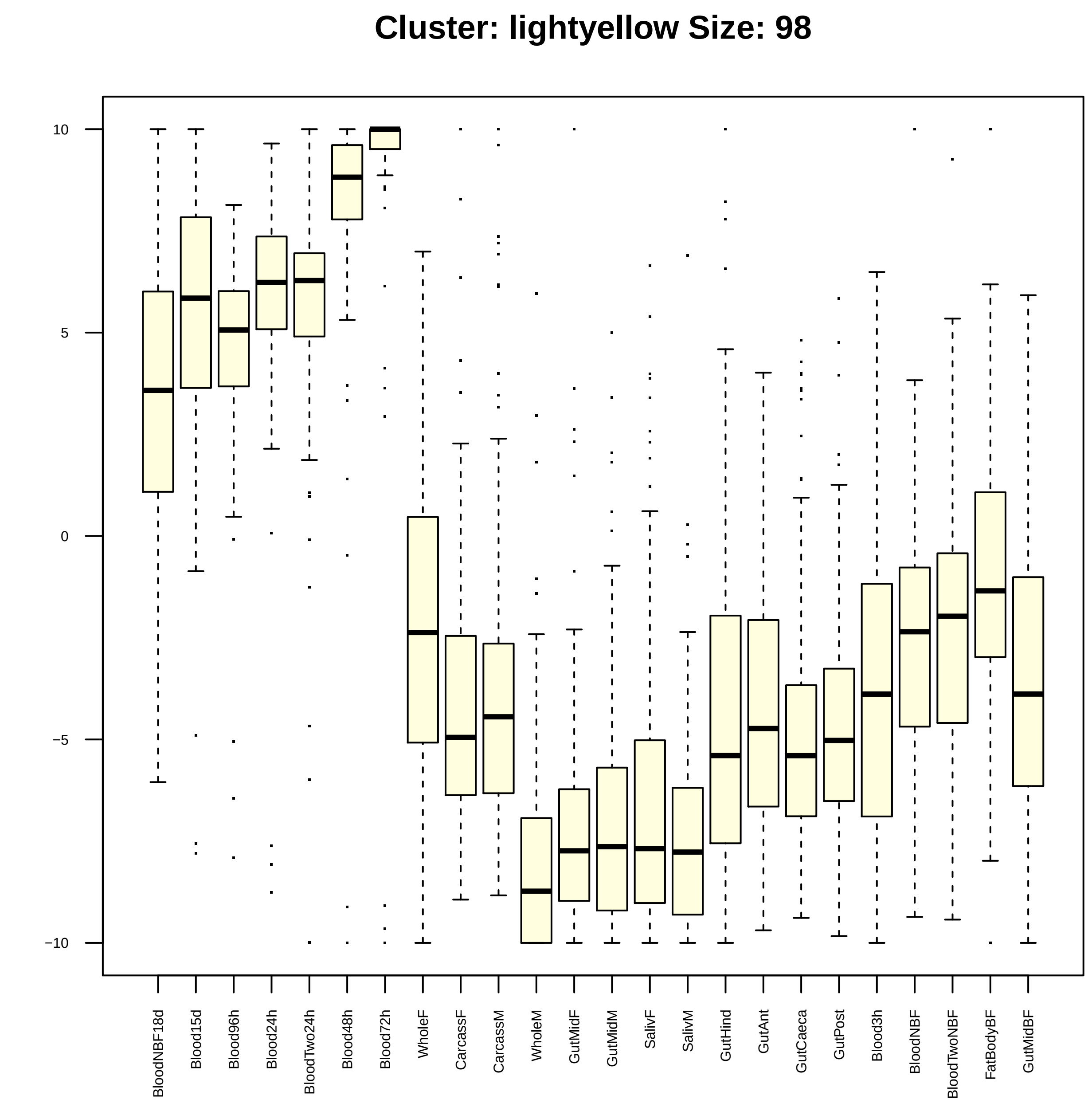

Cluster: lightyellow Size: 98
10
5
0
−5
−10
BloodNBF18d
Blood15d
Blood96h
Blood24h
BloodTwo24h
Blood48h
Blood72h
WholeF
CarcassF
CarcassM
WholeM
GutMidF
GutMidM
SalivF
SalivM
GutHind
GutAnt
GutCaeca
GutPost
Blood3h
BloodNBF
BloodTwoNBF
FatBodyBF
GutMidBF

| | GO.ID | BPCluster: lightyellow Size: 98 | Annotated | Significant | Expected | Rank in ClassicF | Weight01F | ClassicF |
|---|---|---|---|---|---|---|---|---|
| 1 | GO:0032006 | regulation of TOR signaling | 10 | 3 | 0.10 | 29 | 0.00010 | 0.00010 |
| 2 | GO:0046626 | regulation of insulin receptor signaling... | 12 | 3 | 0.12 | 36 | 0.00018 | 0.00018 |
| 3 | GO:0016476 | regulation of embryonic cell shape | 18 | 3 | 0.18 | 70 | 0.00064 | 0.00064 |
| 4 | GO:0007447 | imaginal disc pattern formation | 47 | 3 | 0.46 | 180 | 0.00133 | 0.01047 |
| 5 | GO:0017148 | negative regulation of translation | 24 | 3 | 0.23 | 90 | 0.00153 | 0.00153 |
| 6 | GO:0008285 | negative regulation of cell proliferatio... | 30 | 3 | 0.29 | 114 | 0.00295 | 0.00295 |
| 7 | GO:0007446 | imaginal disc growth | 28 | 3 | 0.27 | 104 | 0.00388 | 0.00241 |
| 8 | GO:0030713 | ovarian follicle cell stalk formation | 10 | 2 | 0.10 | 126 | 0.00398 | 0.00398 |
| 9 | GO:0022403 | cell cycle phase | 10 | 2 | 0.10 | 127 | 0.00398 | 0.00398 |
| 10 | GO:1902532 | negative regulation of intracellular sig... | 34 | 3 | 0.33 | 131 | 0.00423 | 0.00423 |
| 11 | GO:0009954 | proximal/distal pattern formation | 11 | 2 | 0.11 | 138 | 0.00483 | 0.00483 |
| 12 | GO:0035099 | hemocyte migration | 11 | 2 | 0.11 | 139 | 0.00483 | 0.00483 |
| 13 | GO:0048518 | positive regulation of biological proces... | 430 | 14 | 4.18 | 20 | 0.00507 | 4.7e−05 |
| 14 | GO:1902533 | positive regulation of intracellular sig... | 41 | 4 | 0.40 | 69 | 0.00546 | 0.00063 |
| 15 | GO:0016322 | neuron remodeling | 12 | 2 | 0.12 | 149 | 0.00577 | 0.00577 |
| 16 | GO:0030178 | negative regulation of Wnt signaling pat... | 13 | 2 | 0.13 | 160 | 0.00677 | 0.00677 |
| 20 | GO:0007423 | sensory organ development | 219 | 8 | 2.13 | 84 | 0.01517 | 0.00116 |
| 21 | GO:0000003 | reproduction | 452 | 11 | 4.39 | 125 | 0.01747 | 0.00376 |
| 24 | GO:0007420 | brain development | 54 | 4 | 0.53 | 94 | 0.02468 | 0.00178 |
| 30 | GO:0048477 | oogenesis | 263 | 8 | 2.56 | 123 | 0.02826 | 0.00368 |

| | GO.ID | MFCluster: lightyellow Size: 98 | Annotated | Significant | Expected | Rank in ClassicF | Weight01F | ClassicF |
|---|---|---|---|---|---|---|---|---|
| *1* | GO:0005515 | protein binding | 2143 | 43 | 23.48 | 2 | 0.00028 | 3.6e−06 |
| *2* | GO:0043021 | ribonucleoprotein complex binding | 14 | 2 | 0.15 | 7 | 0.00991 | 0.00991 |
| *11* | GO:0044877 | protein−containing complex binding | 50 | 4 | 0.55 | 4 | 0.05636 | 0.00210 |
| *16* | GO:0003676 | nucleic acid binding | 1152 | 25 | 12.62 | 3 | 0.09265 | 0.00038 |

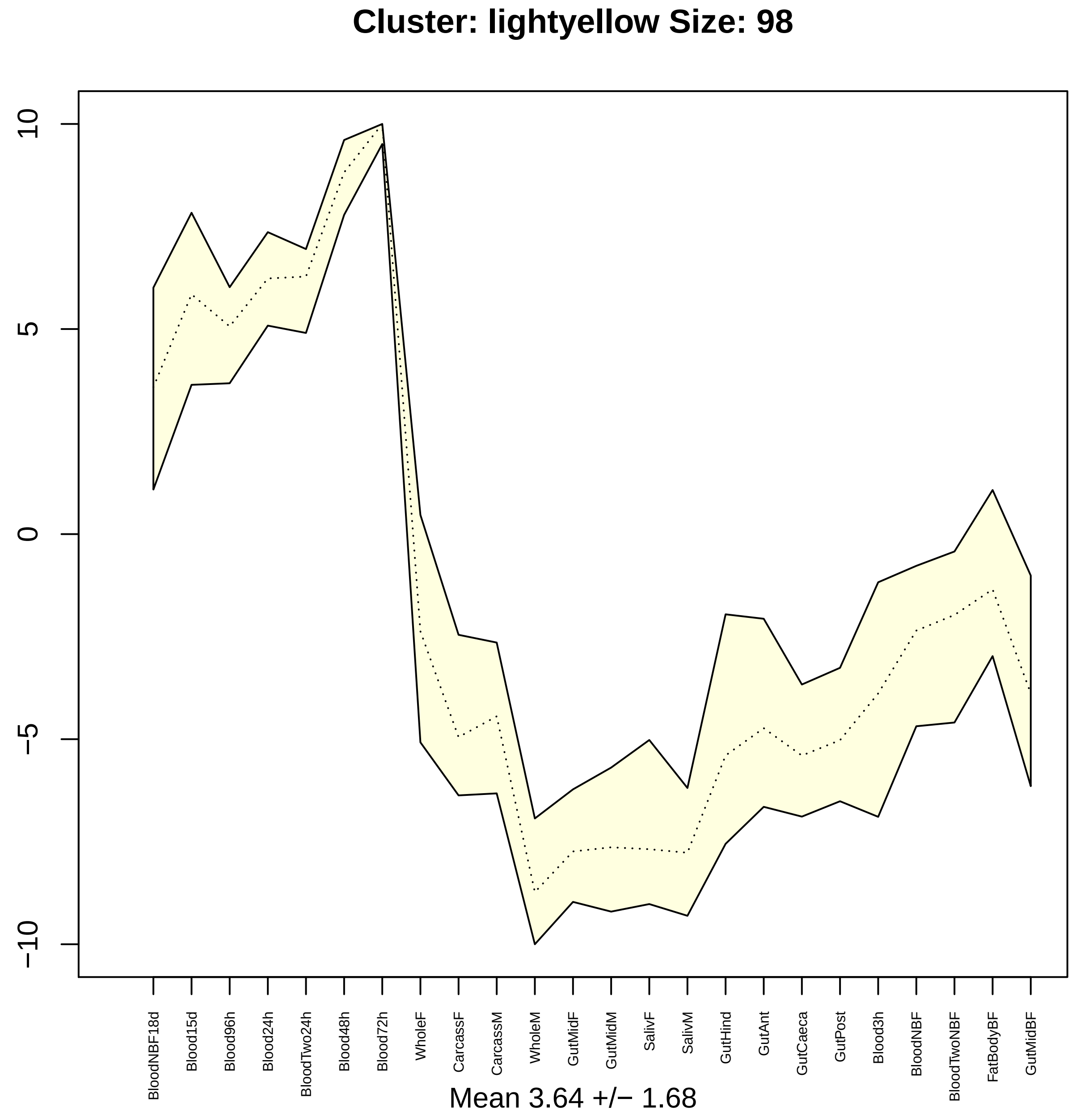
Cluster: lightyellow Size: 98
10
5
0
−5
−10
BloodNBF18d
Blood15d
Blood96h
Blood24h
BloodTwo24h
Blood48h
Blood72h
WholeF
CarcassF
CarcassM
WholeM
GutMidF
GutMidM
SalivF
SalivM
GutHind
GutAnt
GutCaeca
GutPost
Blood3h
BloodNBF
BloodTwoNBF
FatBodyBF
GutMidBF
Mean 3.64 +/− 1.68

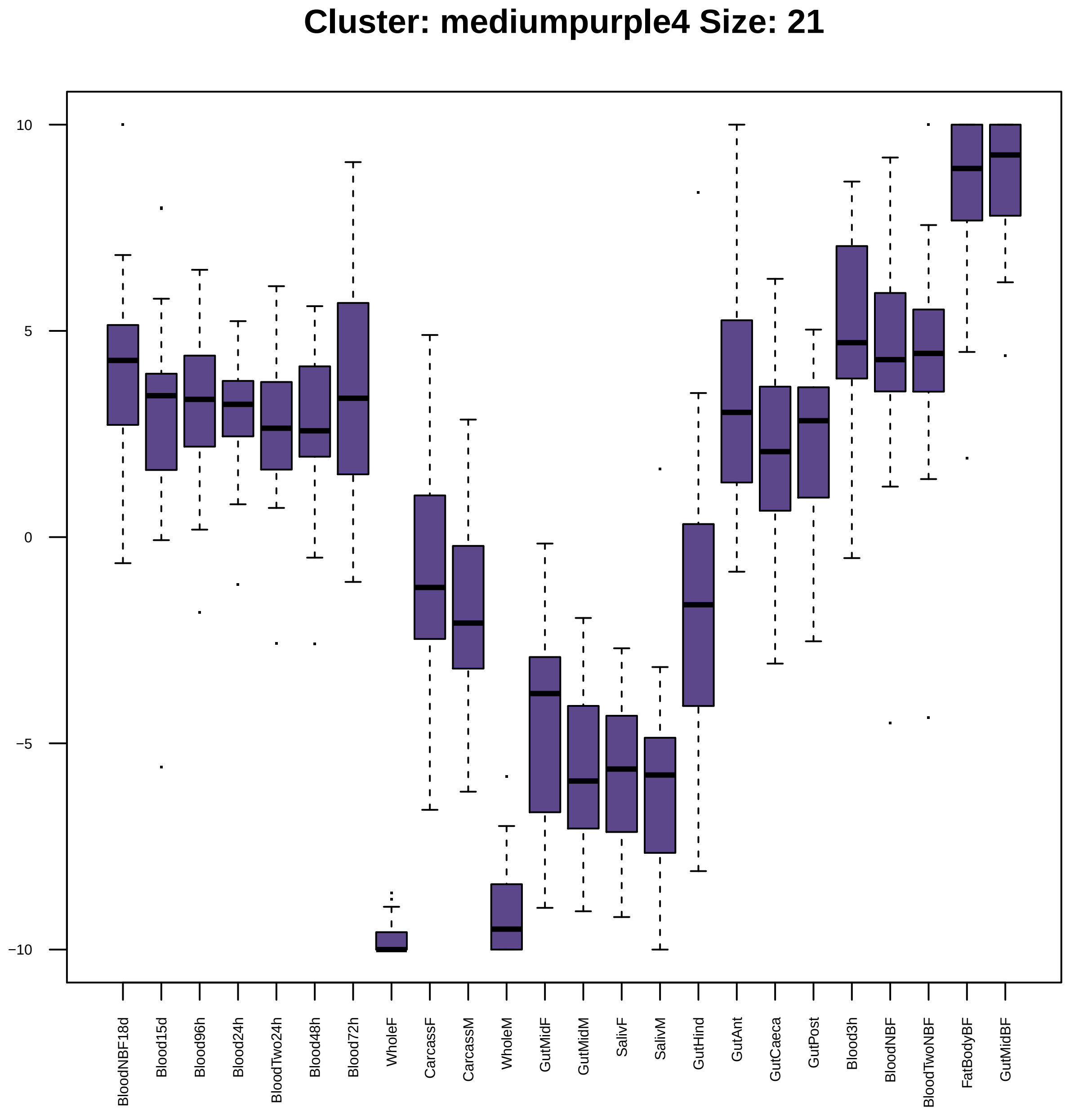

Cluster: mediumpurple4 Size: 21
10
5
0
−5
−10
BloodNBF18d
Blood15d
Blood96h
Blood24h
BloodTwo24h
Blood48h
Blood72h
WholeF
CarcassF
CarcassM
WholeM
GutMidF
GutMidM
SalivF
SalivM
GutHind
GutAnt
GutCaeca
GutPost
Blood3h
BloodNBF
BloodTwoNBF
FatBodyBF
GutMidBF

| | GO.ID | BPCluster: mediumpurple4 Size: 21 | Annotated | Significant | Expected | Rank in ClassicF | Weight01F | ClassicF |
|---|---|---|---|---|---|---|---|---|
| *23* | GO:0016192 | vesicle−mediated transport | 333 | 3 | 0.47 | 2 | 0.114 | 0.0094 |

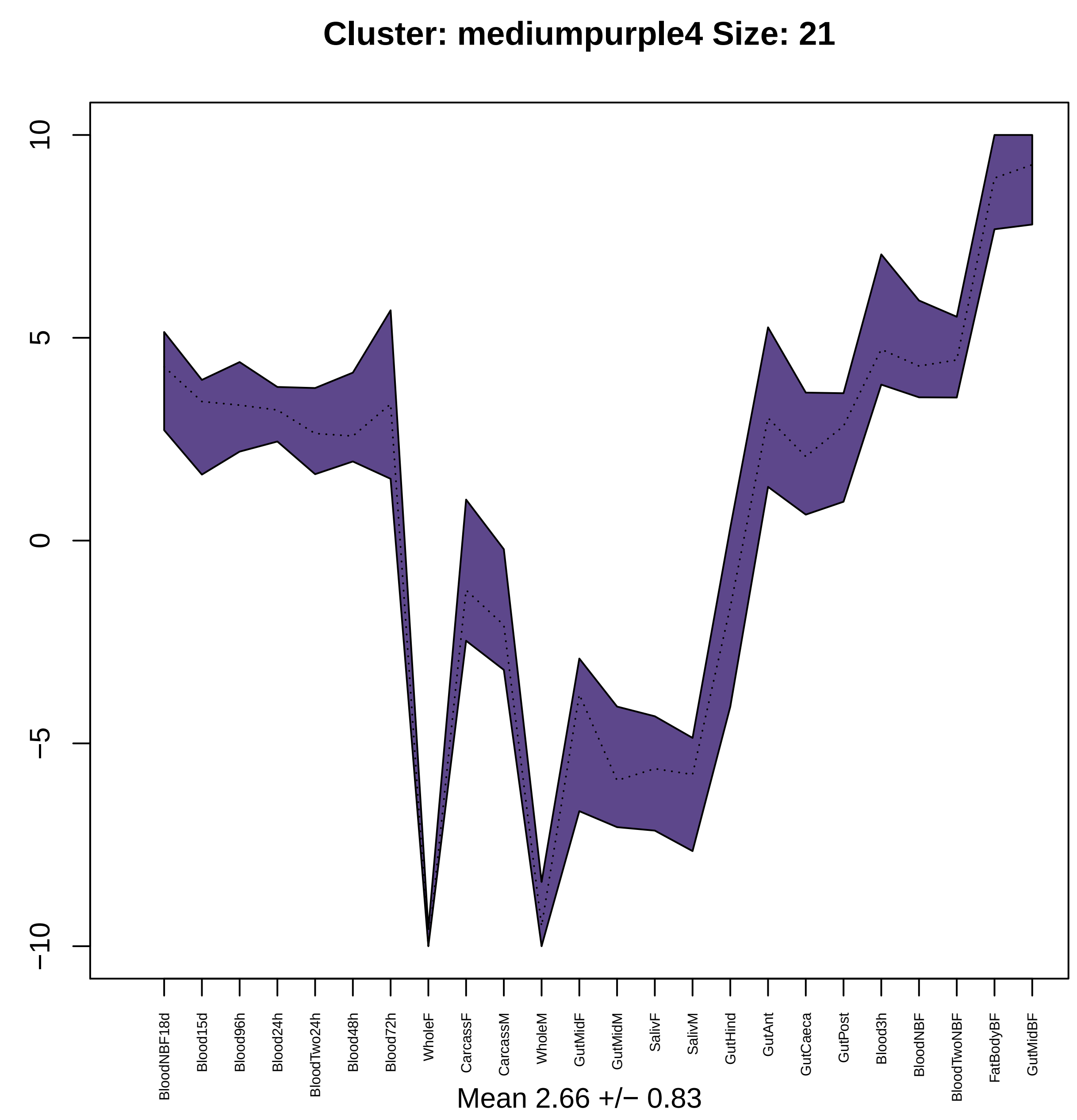
Cluster: mediumpurple4 Size: 21
10
5
0
−5
−10
BloodNBF18d
Blood15d
Blood96h
Blood24h
BloodTwo24h
Blood48h
Blood72h
WholeF
CarcassF
CarcassM
WholeM
GutMidF
GutMidM
SalivF
SalivM
GutHind
GutAnt
GutCaeca
GutPost
Blood3h
BloodNBF
BloodTwoNBF
FatBodyBF
GutMidBF
Mean 2.66 +/− 0.83

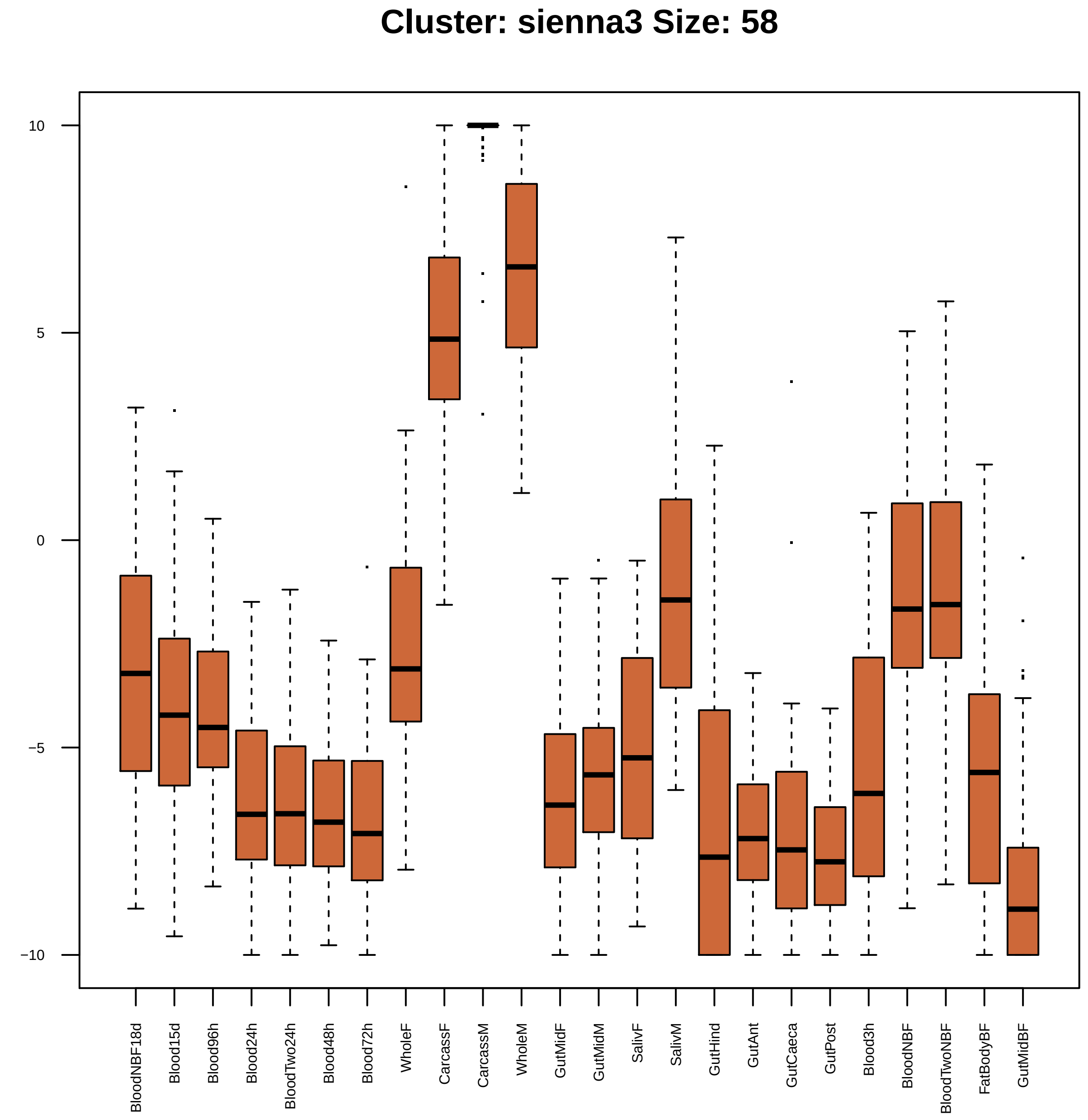
Cluster: sienna3 Size: 58
10
5
0
−5
−10
BloodNBF18d
Blood15d
Blood96h
Blood24h
BloodTwo24h
Blood48h
Blood72h
WholeF
CarcassF
CarcassM
WholeM
GutMidF
GutMidM
SalivF
SalivM
GutHind
GutAnt
GutCaeca
GutPost
Blood3h
BloodNBF
BloodTwoNBF
FatBodyBF
GutMidBF

| | GO.ID | BPCluster: sienna3 Size: 58 | Annotated | Significant | Expected | Rank in ClassicF | Weight01F | ClassicF |
|---|---|---|---|---|---|---|---|---|
| *1* | GO:0055085 | transmembrane transport | 469 | 8 | 2.94 | 1 | 0.0059 | 0.0076 |

| | GO.ID | MFCluster: sienna3 Size: 58 | Annotated | Significant | Expected | Rank in ClassicF | Weight01F | ClassicF |
|---|---|---|---|---|---|---|---|---|
| 1 | GO:0022857 | transmembrane transporter activity | 442 | 6 | 2.70 | 8 | 0.0077 | 0.0502 |
| 8 | GO:1901681 | sulfur compound binding | 23 | 2 | 0.14 | 2 | 0.0750 | 0.0085 |
| 12 | GO:0005215 | transporter activity | 548 | 9 | 3.34 | 1 | 0.0900 | 0.0051 |

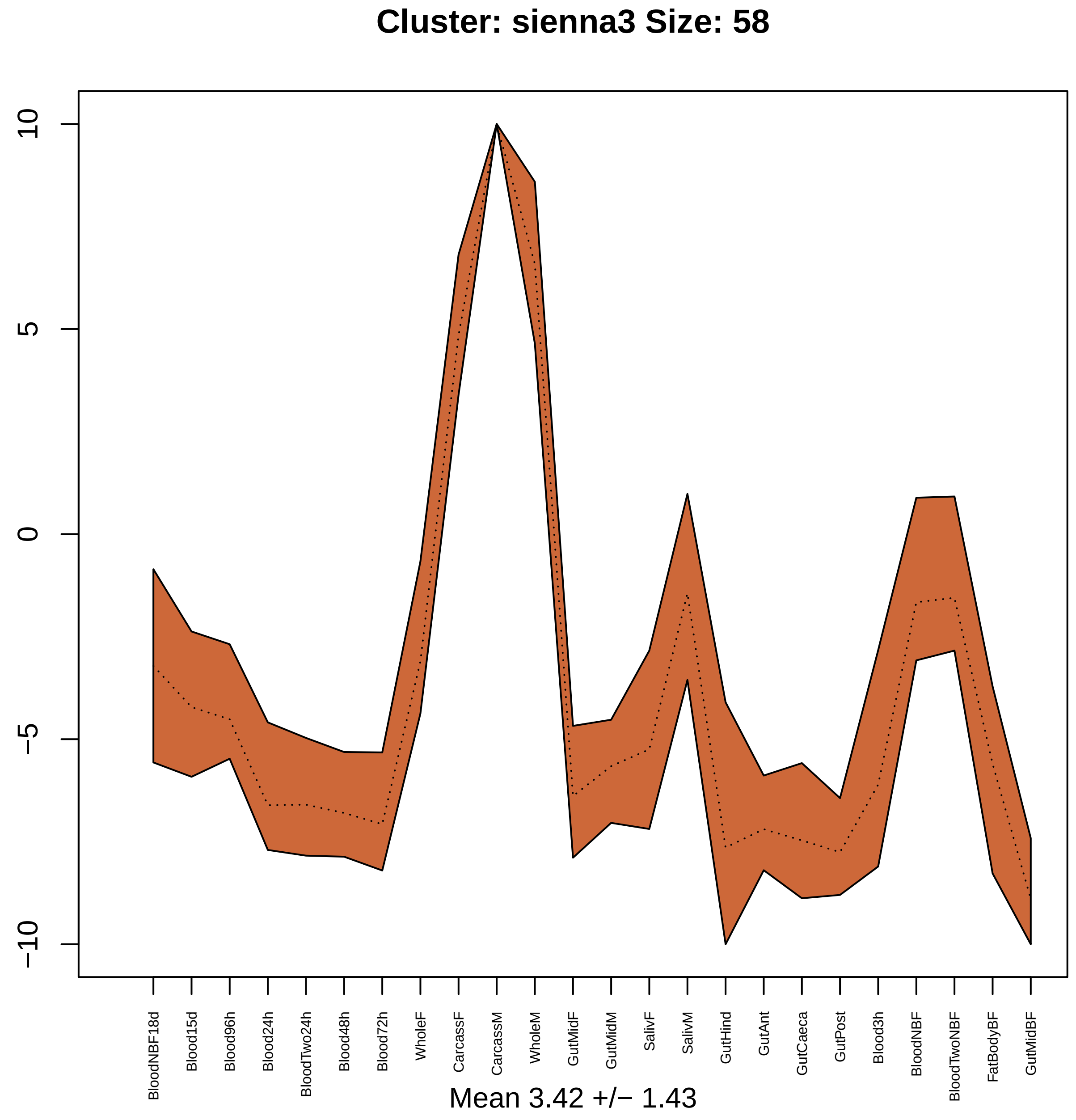

Cluster: sienna3 Size: 58
10
5
0
−5
−10
BloodNBF18d
Blood15d
Blood96h
Blood24h
BloodTwo24h
Blood48h
Blood72h
WholeF
CarcassF
CarcassM
WholeM
GutMidF
GutMidM
SalivF
SalivM
GutHind
GutAnt
GutCaeca
GutPost
Blood3h
BloodNBF
BloodTwoNBF
FatBodyBF
GutMidBF
Mean 3.42 +/− 1.43

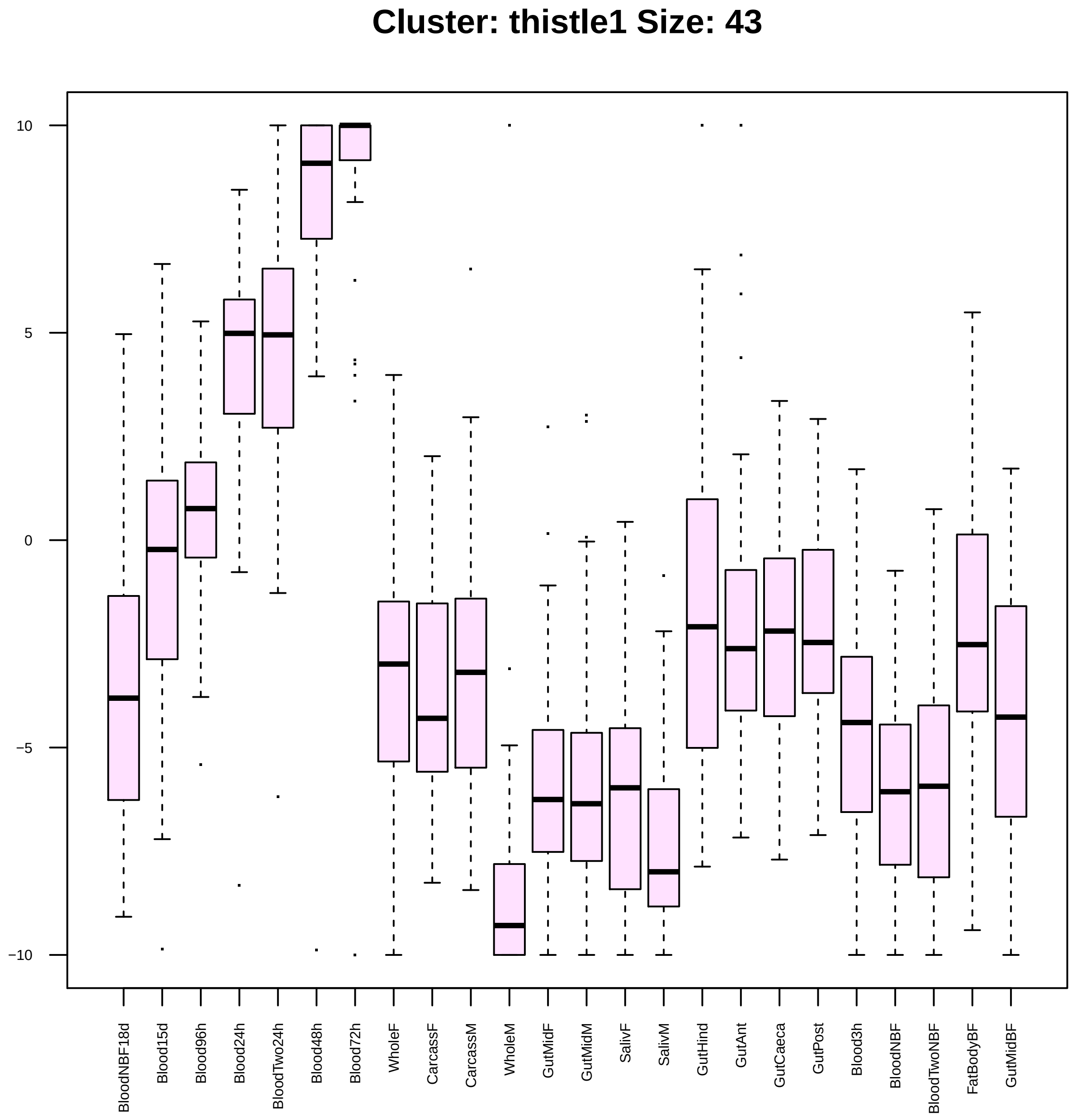

Cluster: thistle1 Size: 43
10
5
0
−5
−10
BloodNBF18d
Blood15d
Blood96h
Blood24h
BloodTwo24h
Blood48h
Blood72h
WholeF
CarcassF
CarcassM
WholeM
GutMidF
GutMidM
SalivF
SalivM
GutHind
GutAnt
GutCaeca
GutPost
Blood3h
BloodNBF
BloodTwoNBF
FatBodyBF
GutMidBF

|  | GO.ID | BPCluster: thistle1 Size: 43 | Annotated | Significant | Expected | Rank in ClassicF | Weight01F | ClassicF |
|---|---|---|---|---|---|---|---|---|
| 1 | GO:0000375 | RNA splicing, via transesterification re... | 135 | 2 | 0.47 | 108 | 0.0034 | 0.07802 |
| 2 | GO:0007143 | female meiotic nuclear division | 26 | 2 | 0.09 | 5 | 0.0035 | 0.00351 |
| 3 | GO:0016570 | histone modification | 58 | 3 | 0.20 | 2 | 0.0072 | 0.00097 |
| 4 | GO:0032774 | RNA biosynthetic process | 565 | 4 | 1.95 | 161 | 0.0098 | 0.12460 |

| | GO.ID | MFCluster: thistle1 Size: 43 | Annotated | Significant | Expected | Rank in ClassicF | Weight01F | ClassicF |
|---|---|---|---|---|---|---|---|---|
| 1 | GO:0003682 | chromatin binding | 69 | 3 | 0.27 | 3 | 0.0023 | 0.0023 |
| 2 | GO:0003676 | nucleic acid binding | 1152 | 11 | 4.47 | 4 | 0.0048 | 0.0026 |

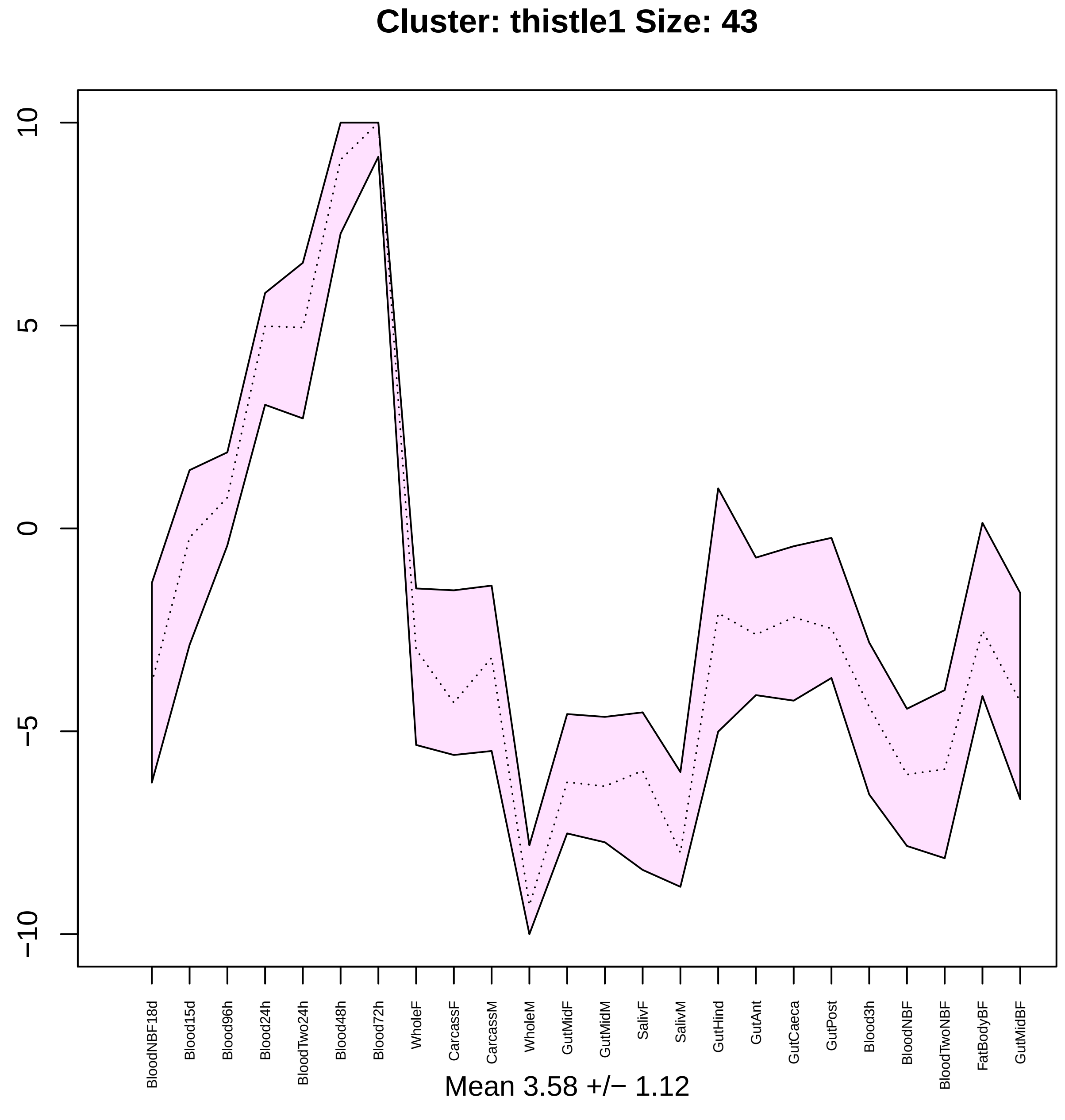
Cluster: thistle1 Size: 43
10
5
0
−5
−10
BloodNBF18d
Blood15d
Blood96h
Blood24h
BloodTwo24h
Blood48h
Blood72h
WholeF
CarcassF
CarcassM
WholeM
GutMidF
GutMidM
SalivF
SalivM
GutHind
GutAnt
GutCaeca
GutPost
Blood3h
BloodNBF
BloodTwoNBF
FatBodyBF
GutMidBF
Mean 3.58 +/− 1.12

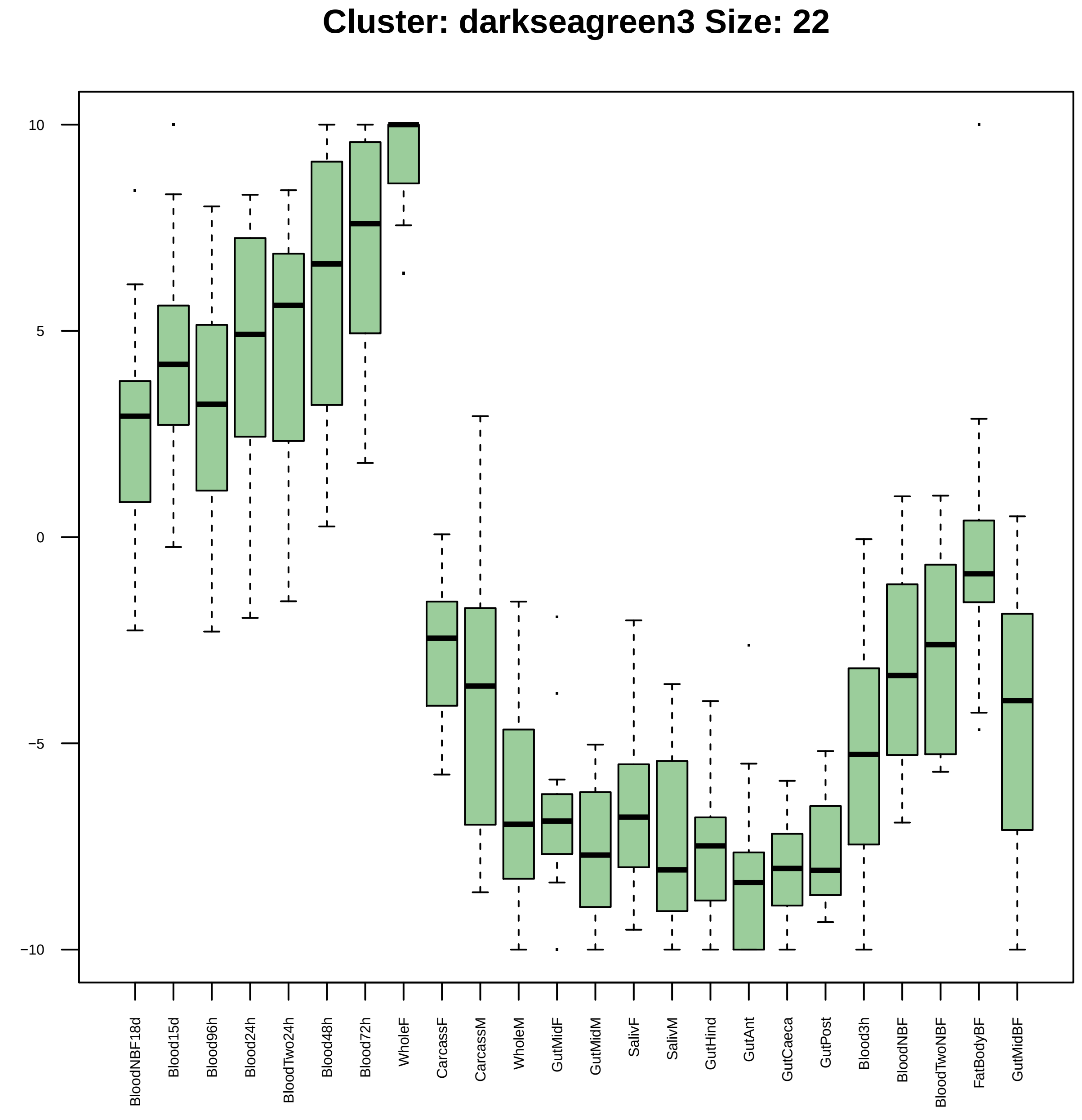
Cluster: darkseagreen3 Size: 22
10
5
0
−5
−10
BloodNBF18d
Blood15d
Blood96h
Blood24h
BloodTwo24h
Blood48h
Blood72h
WholeF
CarcassF
CarcassM
WholeM
GutMidF
GutMidM
SalivF
SalivM
GutHind
GutAnt
GutCaeca
GutPost
Blood3h
BloodNBF
BloodTwoNBF
FatBodyBF
GutMidBF

| | GO.ID | BPCluster: darkseagreen3 Size: 22 | Annotated | Significant | Expected | Rank in ClassicF | Weight01F | ClassicF |
|---|---|---|---|---|---|---|---|---|
| *1* | GO:0061983 | meiosis II cell cycle process | 10 | 2 | 0.02 | 1 | 0.00017 | 0.00017 |
| *2* | GO:0044837 | actomyosin contractile ring organization | 14 | 2 | 0.03 | 2 | 0.00034 | 0.00034 |
| *3* | GO:0032506 | cytokinetic process | 15 | 2 | 0.03 | 3 | 0.00040 | 0.00040 |
| *4* | GO:0072593 | reactive oxygen species metabolic proces... | 21 | 2 | 0.04 | 10 | 0.00079 | 0.00079 |
| *5* | GO:0051225 | spindle assembly | 27 | 2 | 0.06 | 13 | 0.00131 | 0.00131 |
| *6* | GO:0007283 | spermatogenesis | 76 | 3 | 0.15 | 5 | 0.00189 | 0.00043 |
| *7* | GO:0043066 | negative regulation of apoptotic process | 34 | 2 | 0.07 | 15 | 0.00207 | 0.00207 |
| *8* | GO:0006979 | response to oxidative stress | 51 | 2 | 0.10 | 23 | 0.00462 | 0.00462 |
| *9* | GO:0033206 | meiotic cytokinesis | 20 | 2 | 0.04 | 8 | 0.00753 | 0.00071 |

| | GO.ID | MFCluster: darkseagreen3 Size: 22 | Annotated | Significant | Expected | Rank in ClassicF | Weight01F | ClassicF |
|---|---|---|---|---|---|---|---|---|
| 1 | GO:0004601 | peroxidase activity | 26 | 2 | 0.05 | 1 | 0.0013 | 0.0013 |
| 2 | GO:0004674 | protein serine/threonine kinase activity | 150 | 3 | 0.31 | 5 | 0.0033 | 0.0033 |
| 21 | GO:0016684 | oxidoreductase activity, acting on perox... | 26 | 2 | 0.05 | 2 | 1.0000 | 0.0013 |

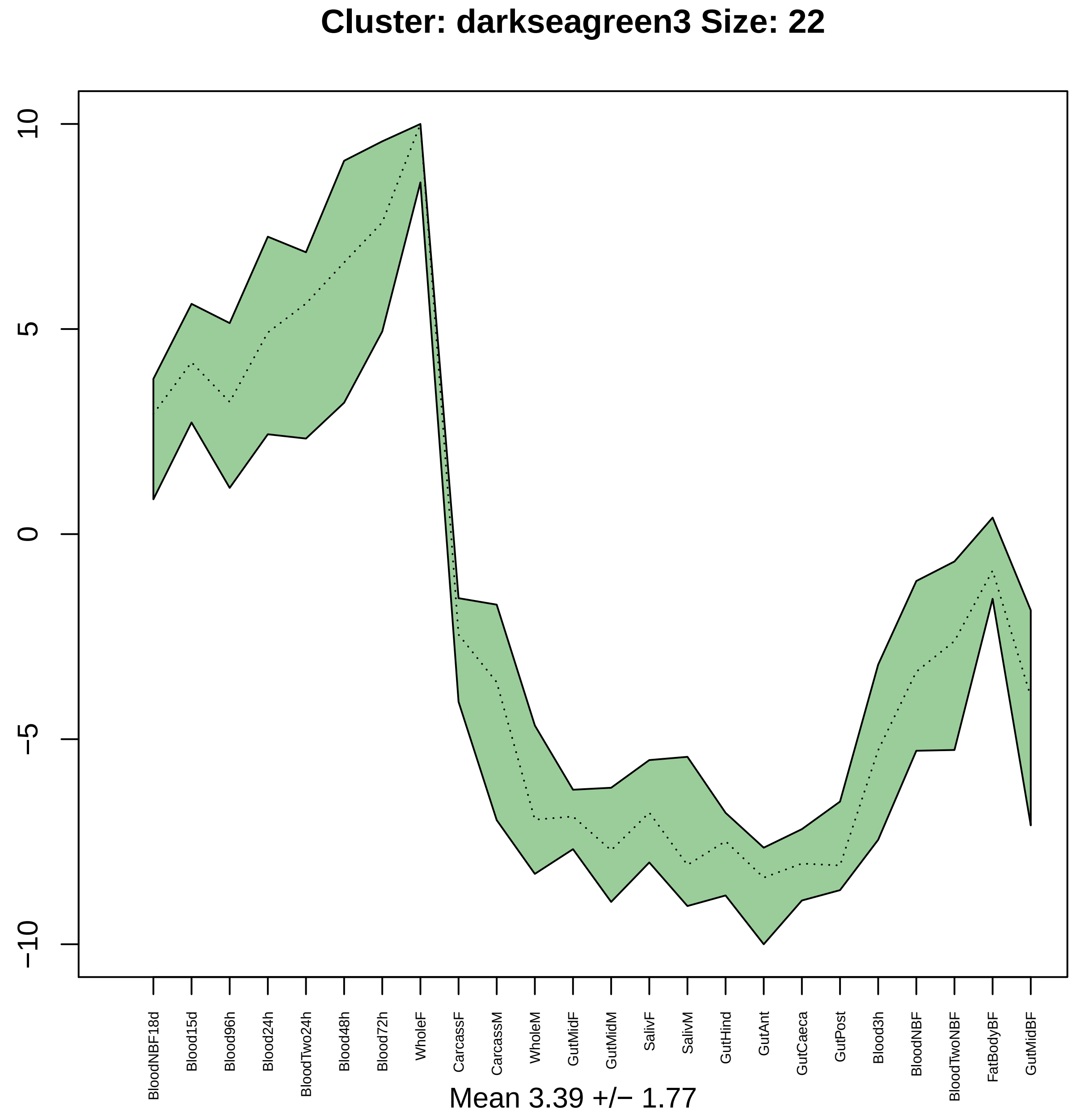
Cluster: darkseagreen3 Size: 22
10
5
0
−5
−10
BloodNBF18d
Blood15d
Blood96h
Blood24h
BloodTwo24h
Blood48h
Blood72h
WholeF
CarcassF
CarcassM
WholeM
GutMidF
GutMidM
SalivF
SalivM
GutHind
GutAnt
GutCaeca
GutPost
Blood3h
BloodNBF
BloodTwoNBF
FatBodyBF
GutMidBF
Mean 3.39 +/− 1.77

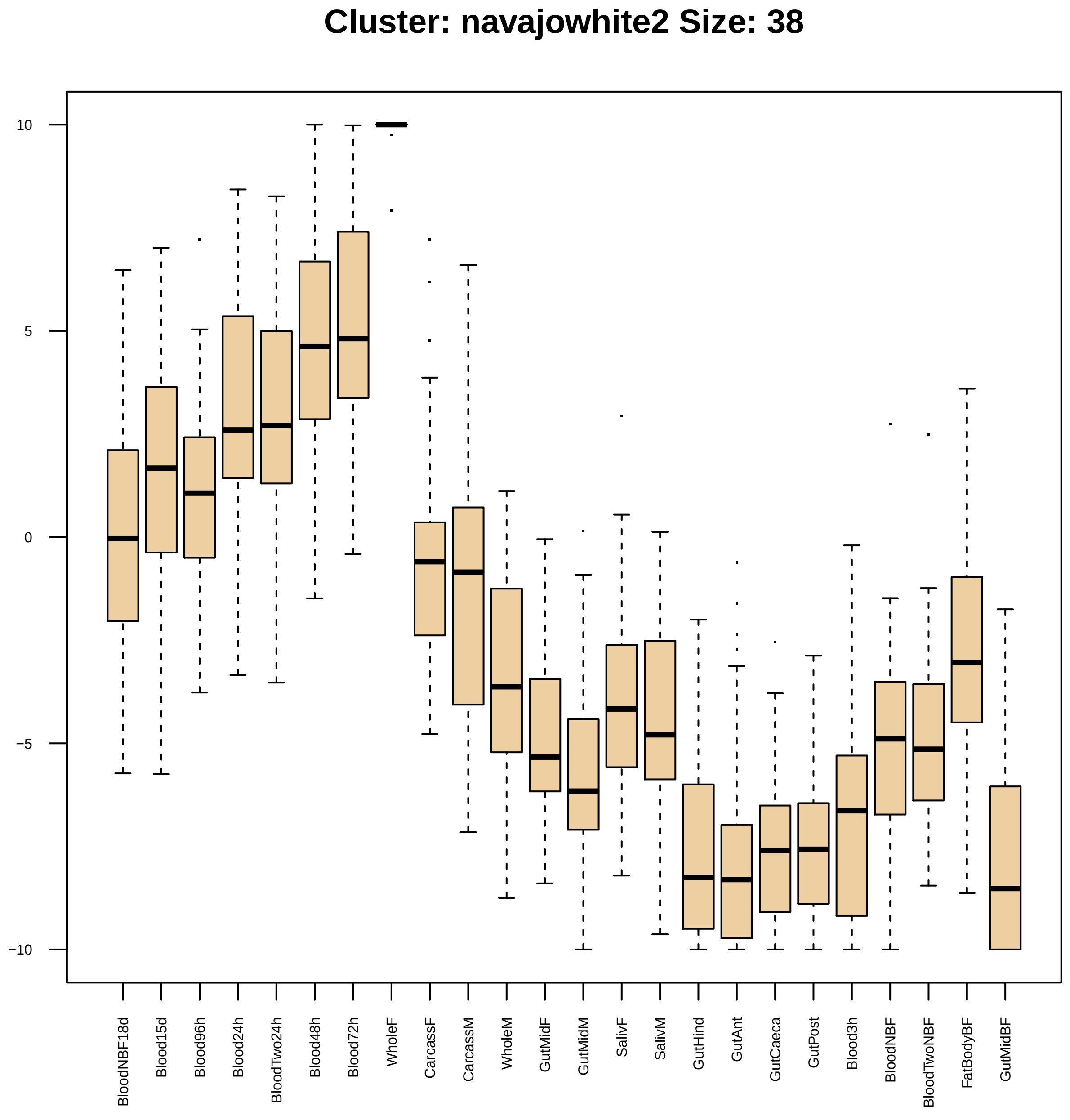

Cluster: navajowhite2 Size: 38
10
5
0
−5
−10
BloodNBF18d
Blood15d
Blood96h
Blood24h
BloodTwo24h
Blood48h
Blood72h
WholeF
CarcassF
CarcassM
WholeM
GutMidF
GutMidM
SalivF
SalivM
GutHind
GutAnt
GutCaeca
GutPost
Blood3h
BloodNBF
BloodTwoNBF
FatBodyBF
GutMidBF

| | GO.ID | BPCluster: navajowhite2 Size: 38 | Annotated | Significant | Expected | Rank in ClassicF | Weight01F | ClassicF |
|---|---|---|---|---|---|---|---|---|
| 1 | GO:0030727 | germarium−derived female germ−line cyst ... | 12 | 2 | 0.04 | 7 | 0.00061 | 0.00061 |
| 2 | GO:0000079 | regulation of cyclin−dependent protein s... | 13 | 2 | 0.04 | 10 | 0.00071 | 0.00071 |
| 3 | GO:0030100 | regulation of endocytosis | 15 | 2 | 0.05 | 13 | 0.00096 | 0.00096 |
| 4 | GO:0022008 | neurogenesis | 706 | 6 | 2.21 | 79 | 0.00193 | 0.01790 |
| 5 | GO:0051171 | regulation of nitrogen compound metaboli... | 775 | 9 | 2.43 | 3 | 0.00641 | 0.00027 |
| 6 | GO:0007314 | oocyte anterior/posterior axis specifica... | 44 | 2 | 0.14 | 43 | 0.00817 | 0.00817 |
| 7 | GO:0051247 | positive regulation of protein metabolic... | 63 | 2 | 0.20 | 75 | 0.00900 | 0.01628 |
| 9 | GO:0007127 | meiosis I | 19 | 2 | 0.06 | 17 | 0.01190 | 0.00155 |
| 11 | GO:0080090 | regulation of primary metabolic process | 774 | 9 | 2.43 | 2 | 0.02119 | 0.00026 |
| 12 | GO:0051028 | mRNA transport | 21 | 2 | 0.07 | 20 | 0.02365 | 0.00189 |
| 15 | GO:0031323 | regulation of cellular metabolic process | 792 | 8 | 2.48 | 19 | 0.03206 | 0.00174 |
| 27 | GO:0021700 | developmental maturation | 91 | 3 | 0.29 | 25 | 0.04205 | 0.00269 |

|  | GO.ID | MFCluster: navajowhite2 Size: 38 | Annotated | Significant | Expected | Rank in ClassicF | Weight01F | ClassicF |
|---|---|---|---|---|---|---|---|---|
| *1* | GO:0004693 | cyclin−dependent protein serine/threonin... | 10 | 2 | 0.03 | 1 | 0.00047 | 0.00047 |
| *2* | GO:0019901 | protein kinase binding | 24 | 2 | 0.08 | 7 | 0.00280 | 0.00280 |
| *3* | GO:0016881 | acid−amino acid ligase activity | 38 | 2 | 0.13 | 13 | 0.00694 | 0.00694 |
| *4* | GO:0004536 | deoxyribonuclease activity | 19 | 2 | 0.06 | 4 | 0.02217 | 0.00175 |
| *6* | GO:0008094 | DNA−dependent ATPase activity | 33 | 2 | 0.11 | 11 | 0.03776 | 0.00527 |
| *17* | GO:0008026 | ATP−dependent helicase activity | 76 | 3 | 0.25 | 5 | 0.07987 | 0.00194 |

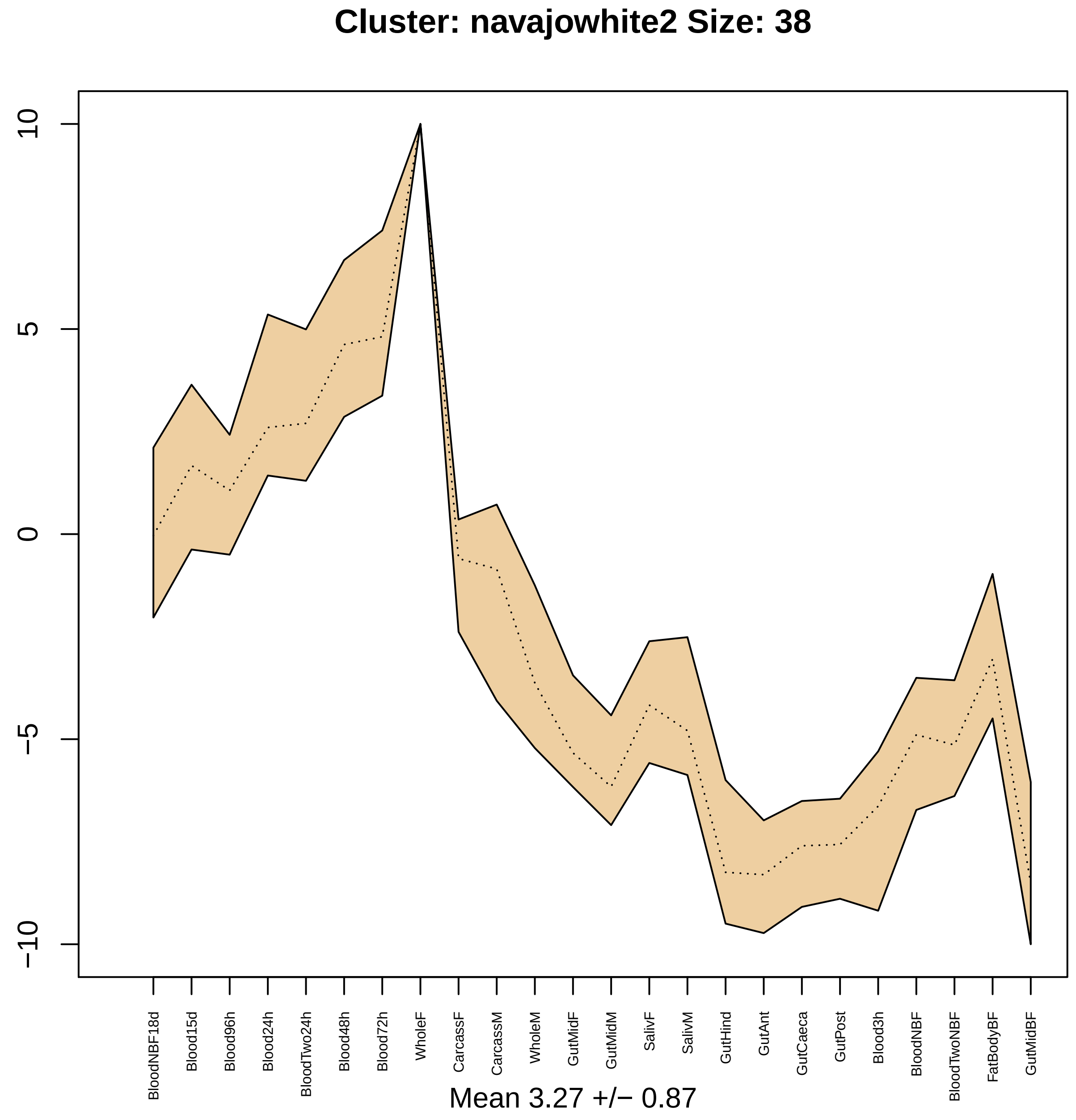
Cluster: navajowhite2 Size: 38
10
5
0
−5
−10
BloodNBF18d
Blood15d
Blood96h
Blood24h
BloodTwo24h
Blood48h
Blood72h
WholeF
CarcassF
CarcassM
WholeM
GutMidF
GutMidM
SalivF
SalivM
GutHind
GutAnt
GutCaeca
GutPost
Blood3h
BloodNBF
BloodTwoNBF
FatBodyBF
GutMidBF
Mean 3.27 +/− 0.87

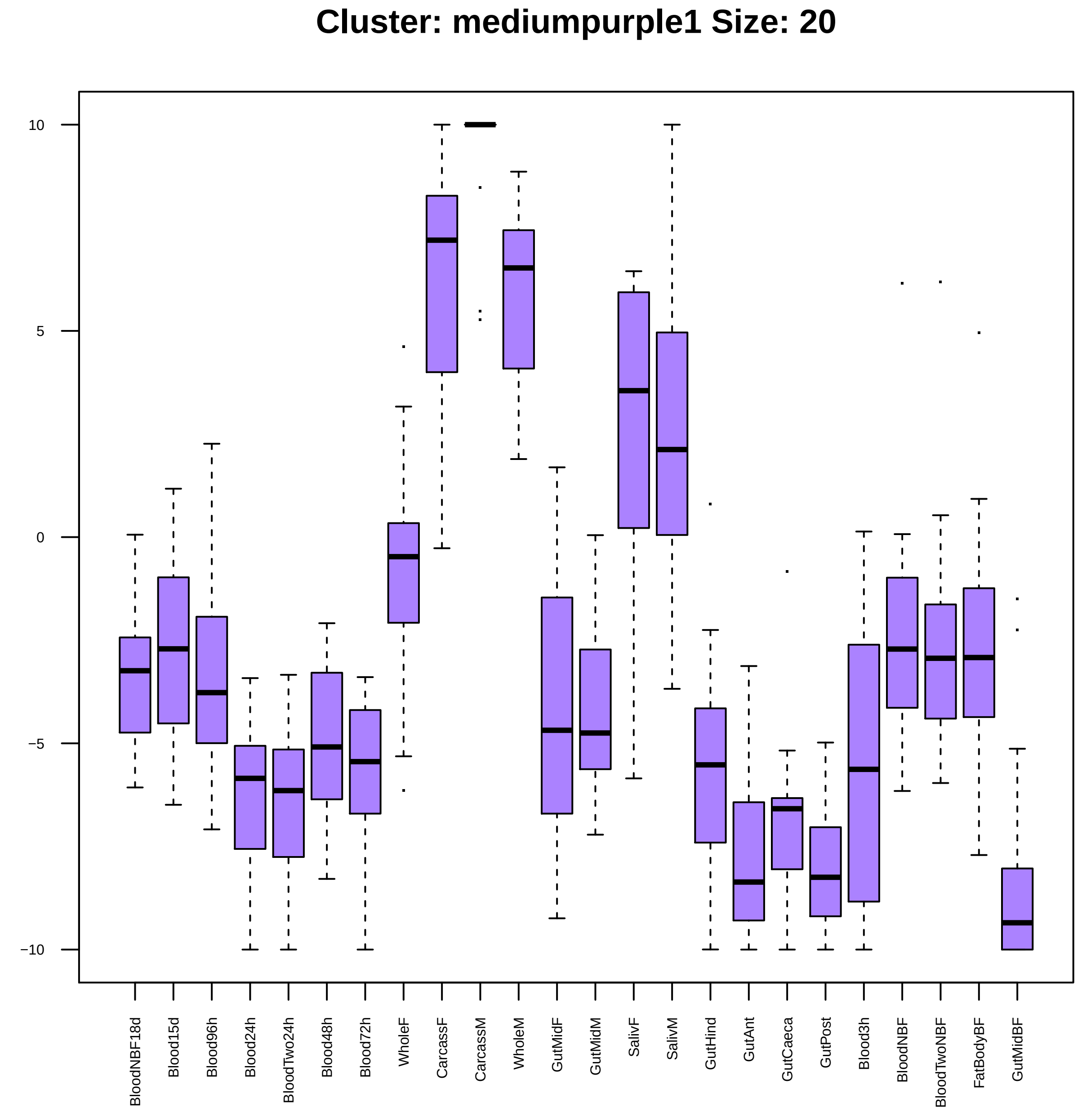

Cluster: mediumpurple1 Size: 20
10
5
0
−5
−10
BloodNBF18d
Blood15d
Blood96h
Blood24h
BloodTwo24h
Blood48h
Blood72h
WholeF
CarcassF
CarcassM
WholeM
GutMidF
GutMidM
SalivF
SalivM
GutHind
GutAnt
GutCaeca
GutPost
Blood3h
BloodNBF
BloodTwoNBF
FatBodyBF
GutMidBF

|  | GO.ID | BPCluster: mediumpurple1 Size: 20 | Annotated | Significant | Expected | Rank in ClassicF | Weight01F | ClassicF |
|---|---|---|---|---|---|---|---|---|
| 1 | GO:0044092 | negative regulation of molecular functio... | 65 | 2 | 0.11 | 1 | 0.0095 | 0.0053 |

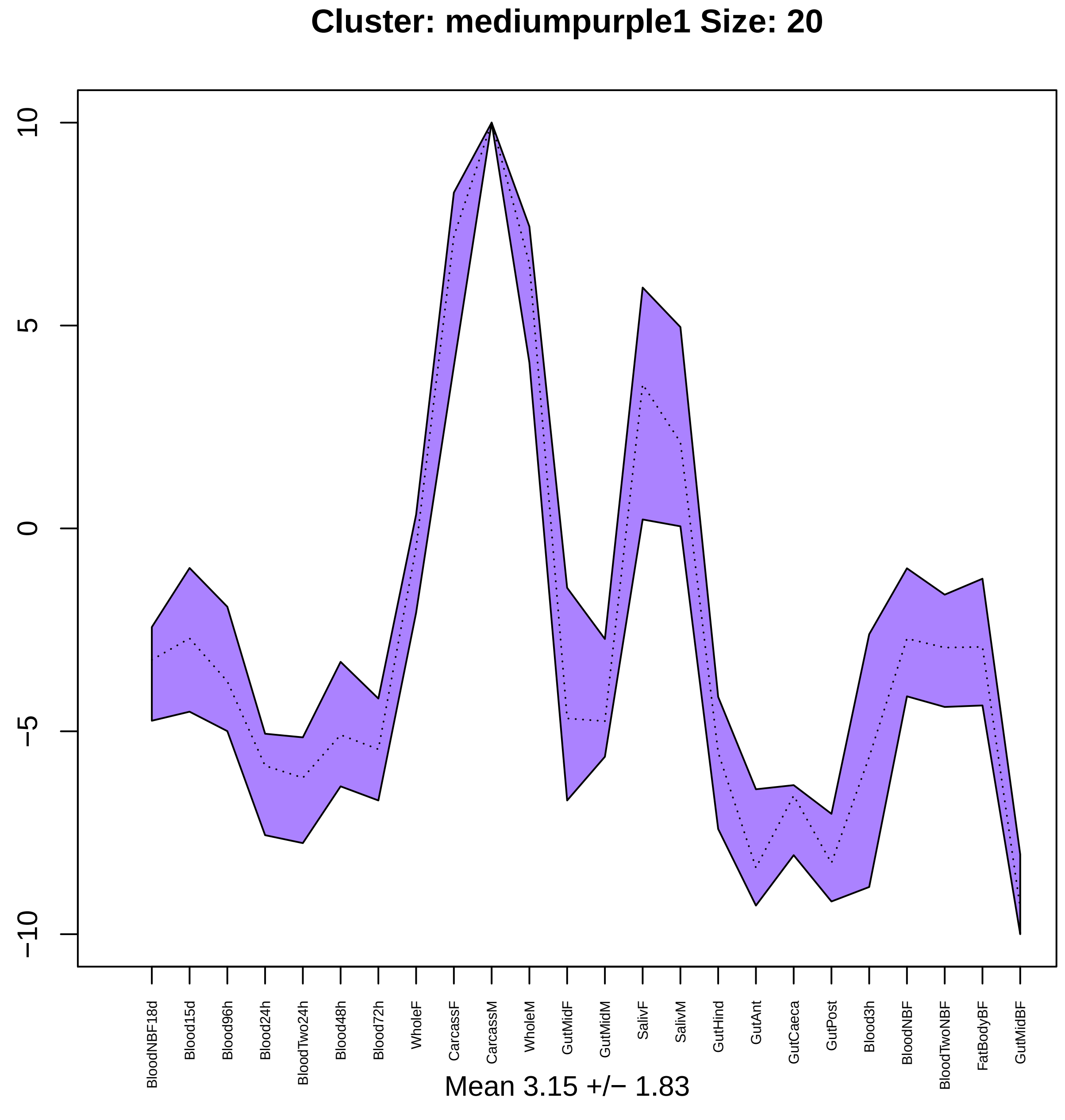

Cluster: mediumpurple1 Size: 20
10
5
0
−5
−10
BloodNBF18d
Blood15d
Blood96h
Blood24h
BloodTwo24h
Blood48h
Blood72h
WholeF
CarcassF
CarcassM
WholeM
GutMidF
GutMidM
SalivF
SalivM
GutHind
GutAnt
GutCaeca
GutPost
Blood3h
BloodNBF
BloodTwoNBF
FatBodyBF
GutMidBF
Mean 3.15 +/− 1.83

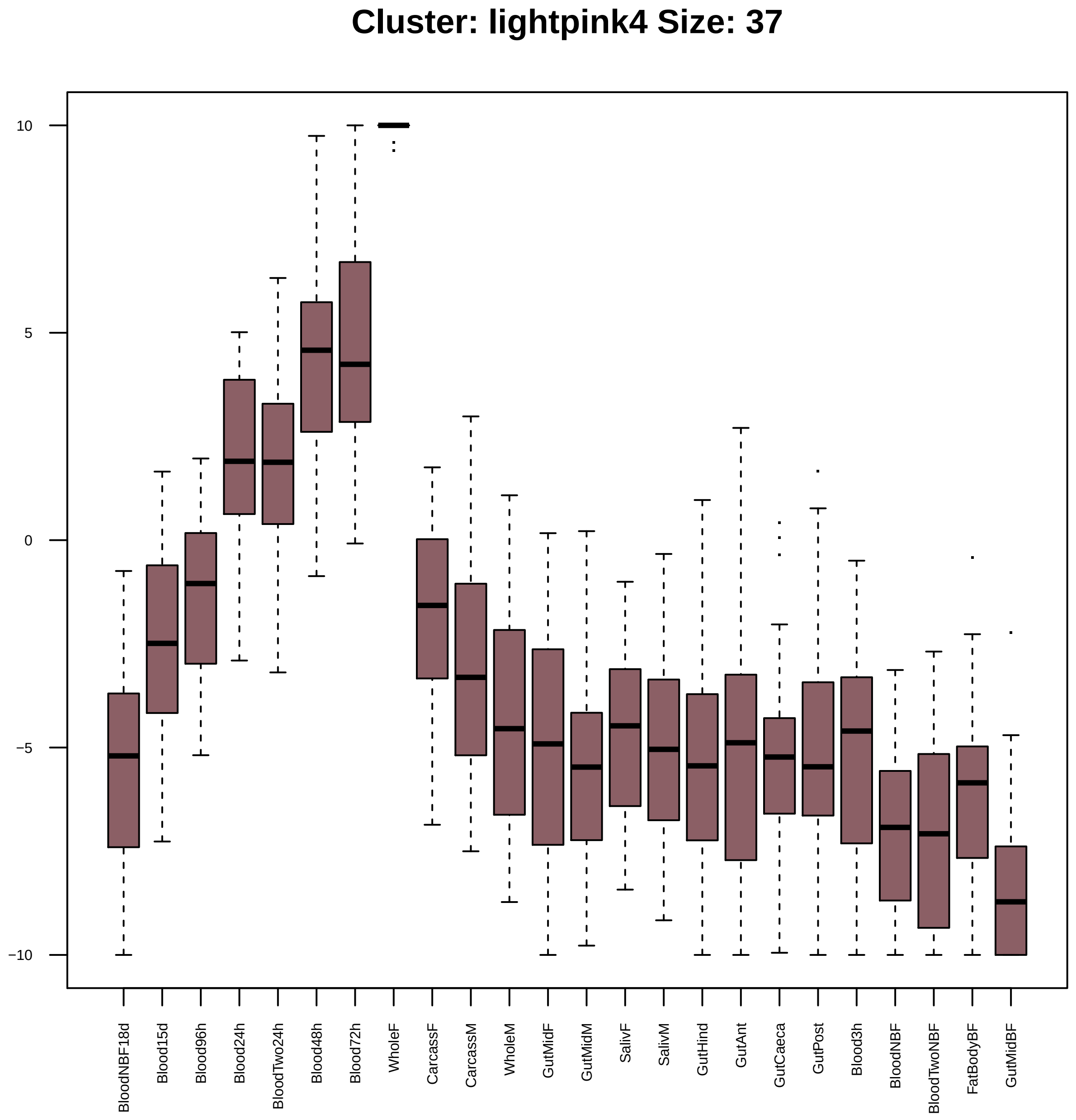

Cluster: lightpink4 Size: 37
10
5
0
−5
−10
BloodNBF18d
Blood15d
Blood96h
Blood24h
BloodTwo24h
Blood48h
Blood72h
WholeF
CarcassF
CarcassM
WholeM
GutMidF
GutMidM
SalivF
SalivM
GutHind
GutAnt
GutCaeca
GutPost
Blood3h
BloodNBF
BloodTwoNBF
FatBodyBF
GutMidBF

| | GO.ID | BPCluster: lightpink4 Size: 37 | Annotated | Significant | Expected | Rank in ClassicF | Weight01F | ClassicF |
|---|---|---|---|---|---|---|---|---|
| 2 | GO:0044786 | cell cycle DNA replication | 18 | 3 | 0.06 | 8 | 0.00013 | 2.1e−05 |
| 3 | GO:0065004 | protein−DNA complex assembly | 29 | 3 | 0.09 | 11 | 0.00055 | 9.2e−05 |
| 4 | GO:0006270 | DNA replication initiation | 12 | 2 | 0.04 | 15 | 0.00061 | 0.00061 |
| 5 | GO:0000724 | double−strand break repair via homologou... | 12 | 2 | 0.04 | 16 | 0.00061 | 0.00061 |
| 6 | GO:0007131 | reciprocal meiotic recombination | 15 | 2 | 0.05 | 18 | 0.00096 | 0.00096 |
| 7 | GO:0032508 | DNA duplex unwinding | 18 | 2 | 0.06 | 24 | 0.00139 | 0.00139 |
| 8 | GO:0046331 | lateral inhibition | 92 | 3 | 0.29 | 35 | 0.00277 | 0.00277 |

| | GO.ID | MFCluster: lightpink4 Size: 37 | Annotated | Significant | Expected | Rank in ClassicF | Weight01F | ClassicF |
|---|---|---|---|---|---|---|---|---|
| 1 | GO:0005524 | ATP binding | 592 | 8 | 1.64 | 2 | 0.00010 | 0.00010 |
| 2 | GO:0003677 | DNA binding | 523 | 8 | 1.45 | 1 | 0.00051 | 4.2e−05 |
| 3 | GO:0003697 | single−stranded DNA binding | 15 | 2 | 0.04 | 11 | 0.00075 | 0.00075 |
| 4 | GO:0016887 | ATPase activity | 233 | 3 | 0.65 | 30 | 0.00510 | 0.02531 |
| 5 | GO:0044877 | protein−containing complex binding | 50 | 2 | 0.14 | 22 | 0.00827 | 0.00827 |
| 11 | GO:0003678 | DNA helicase activity | 32 | 2 | 0.09 | 18 | 0.05407 | 0.00345 |
| 19 | GO:0017111 | nucleoside−triphosphatase activity | 430 | 6 | 1.19 | 12 | 0.31355 | 0.00082 |
| 25 | GO:1901265 | nucleoside phosphate binding | 998 | 8 | 2.77 | 19 | 1.00000 | 0.00353 |
| 26 | GO:0035639 | purine ribonucleoside triphosphate bindi... | 725 | 8 | 2.01 | 7 | 1.00000 | 0.00042 |
| 27 | GO:0032553 | ribonucleotide binding | 740 | 8 | 2.05 | 9 | 1.00000 | 0.00049 |
| 29 | GO:0032555 | purine ribonucleotide binding | 732 | 8 | 2.03 | 8 | 1.00000 | 0.00045 |
| 30 | GO:0036094 | small molecule binding | 1068 | 8 | 2.96 | 21 | 1.00000 | 0.00541 |

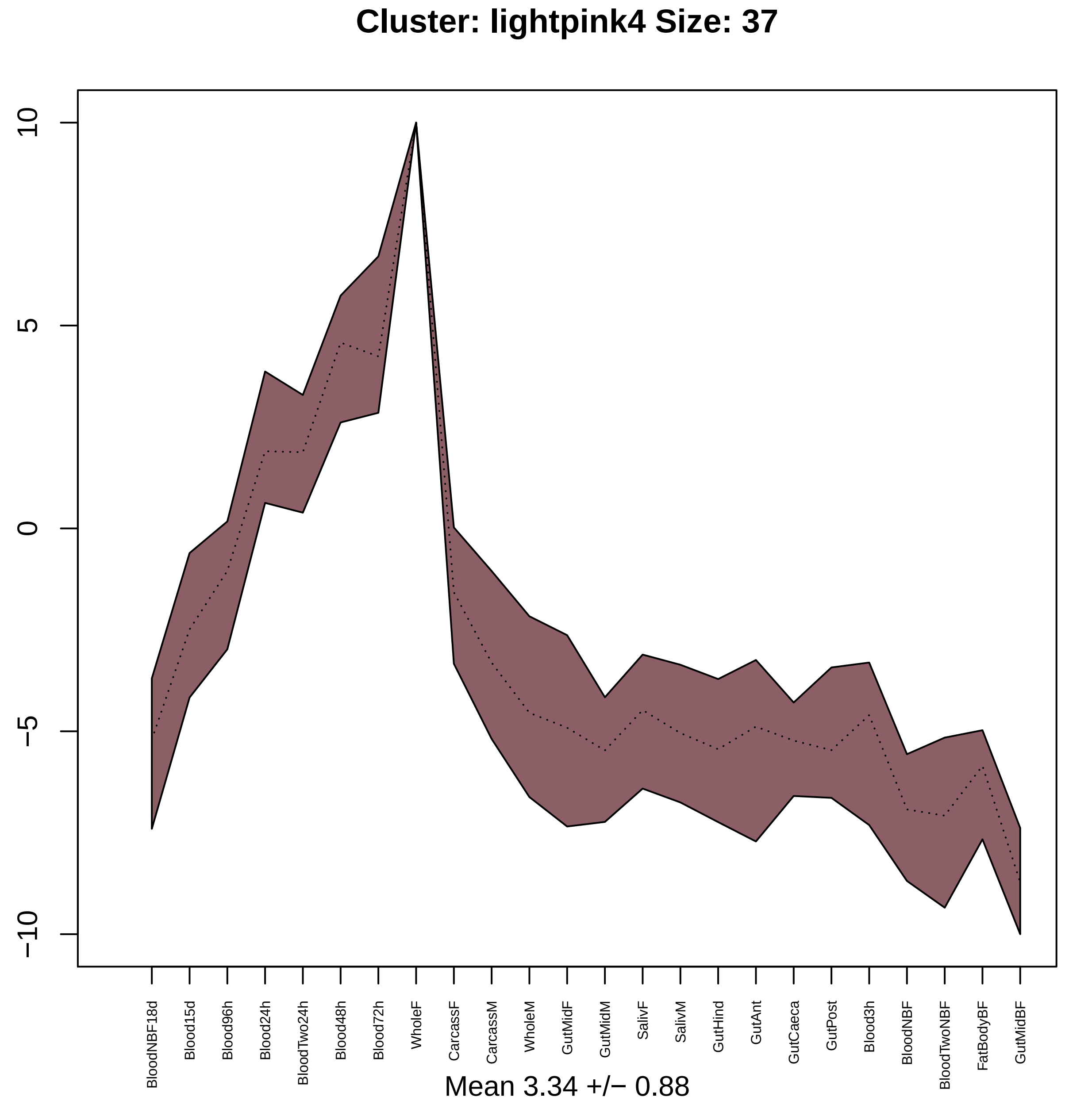
Cluster: lightpink4 Size: 37
10
5
0
−5
−10
BloodNBF18d
Blood15d
Blood96h
Blood24h
BloodTwo24h
Blood48h
Blood72h
WholeF
CarcassF
CarcassM
WholeM
GutMidF
GutMidM
SalivF
SalivM
GutHind
GutAnt
GutCaeca
GutPost
Blood3h
BloodNBF
BloodTwoNBF
FatBodyBF
GutMidBF
Mean 3.34 +/− 0.88

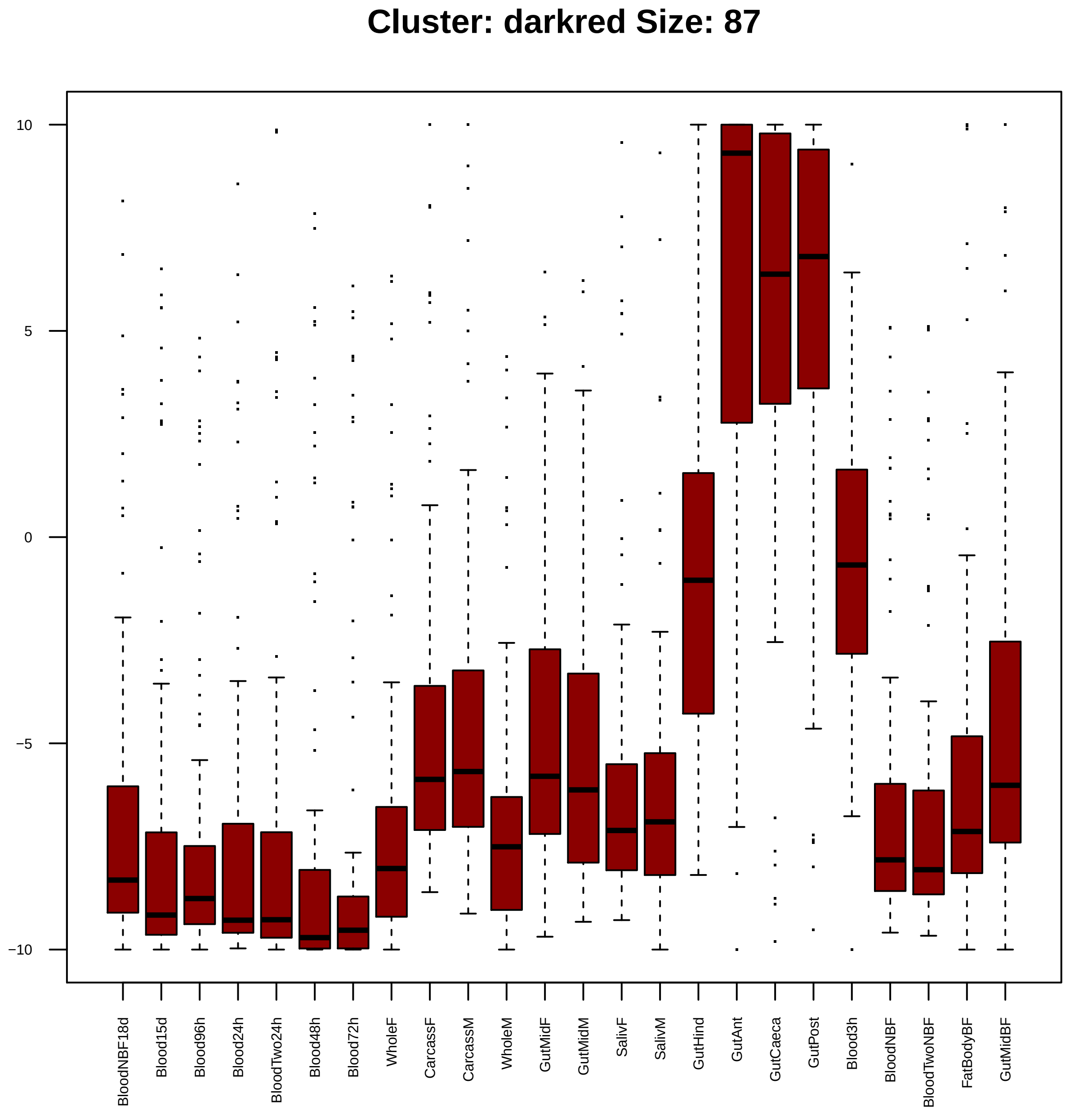

Cluster: darkred Size: 87
10
5
0
−5
−10
BloodNBF18d
Blood15d
Blood96h
Blood24h
BloodTwo24h
Blood48h
Blood72h
WholeF
CarcassF
CarcassM
WholeM
GutMidF
GutMidM
SalivF
SalivM
GutHind
GutAnt
GutCaeca
GutPost
Blood3h
BloodNBF
BloodTwoNBF
FatBodyBF
GutMidBF

| | GO.ID | BPCluster: darkred Size: 87 | Annotated | Significant | Expected | Rank in ClassicF | Weight01F | ClassicF |
|---|---|---|---|---|---|---|---|---|
| *3* | GO:0008152 | metabolic process | 4285 | 59 | 46.36 | 6 | 0.00056 | 0.00045 |

| | GO.ID | MFCluster: darkred Size: 87 | Annotated | Significant | Expected | Rank in ClassicF | Weight01F | ClassicF |
|---|---|---|---|---|---|---|---|---|
| 2 | GO:0008061 | chitin binding | 76 | 6 | 0.76 | 9 | 0.0001 | 0.0001 |
| 3 | GO:0004497 | monooxygenase activity | 110 | 5 | 1.10 | 11 | 0.0047 | 0.0047 |
| 4 | GO:0032934 | sterol binding | 12 | 2 | 0.12 | 12 | 0.0061 | 0.0061 |

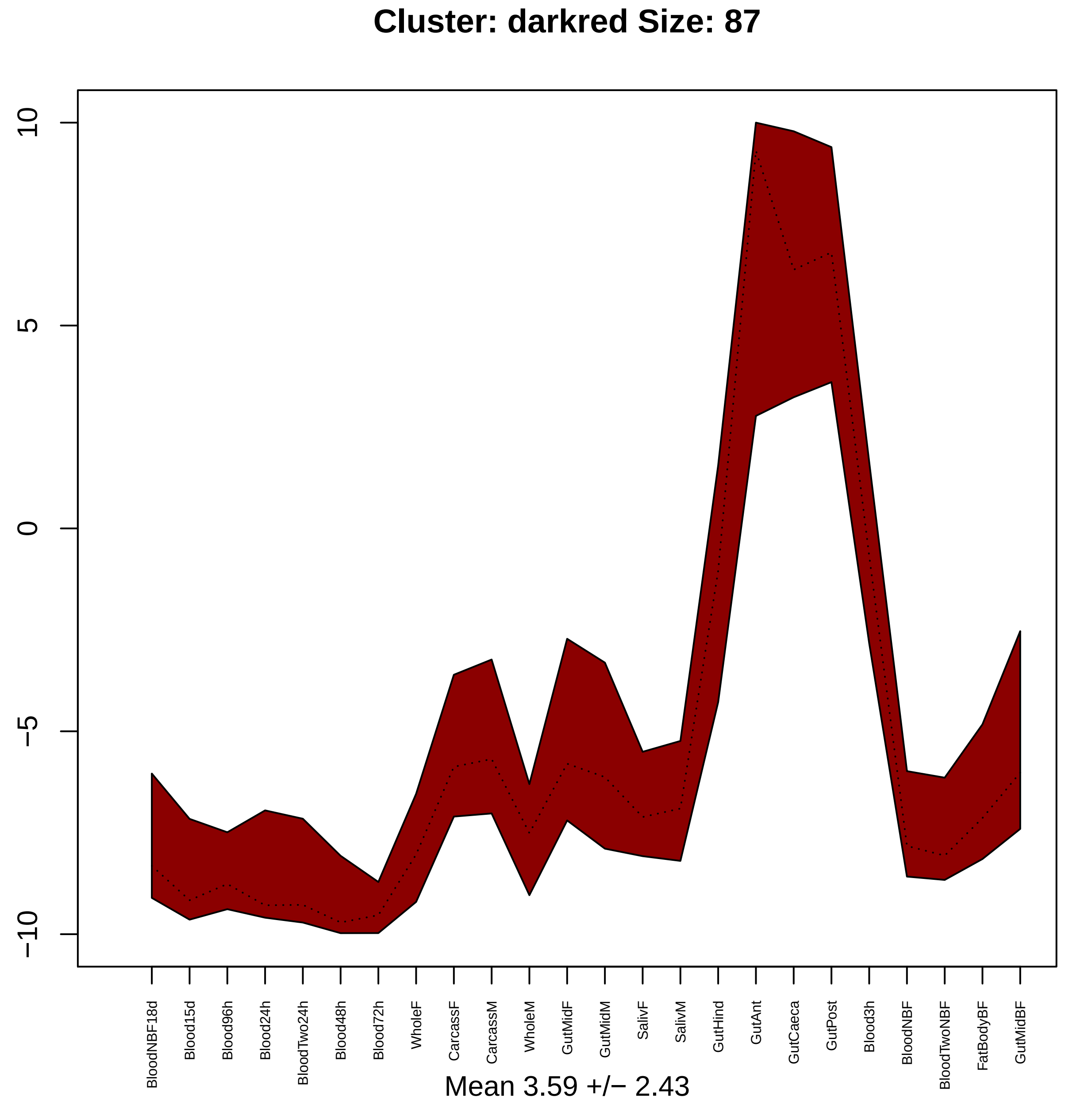

Cluster: darkred Size: 87
10
5
0
−5
−10
BloodNBF18d
Blood15d
Blood96h
Blood24h
BloodTwo24h
Blood48h
Blood72h
WholeF
CarcassF
CarcassM
WholeM
GutMidF
GutMidM
SalivF
SalivM
GutHind
GutAnt
GutCaeca
GutPost
Blood3h
BloodNBF
BloodTwoNBF
FatBodyBF
GutMidBF
Mean 3.59 +/− 2.43

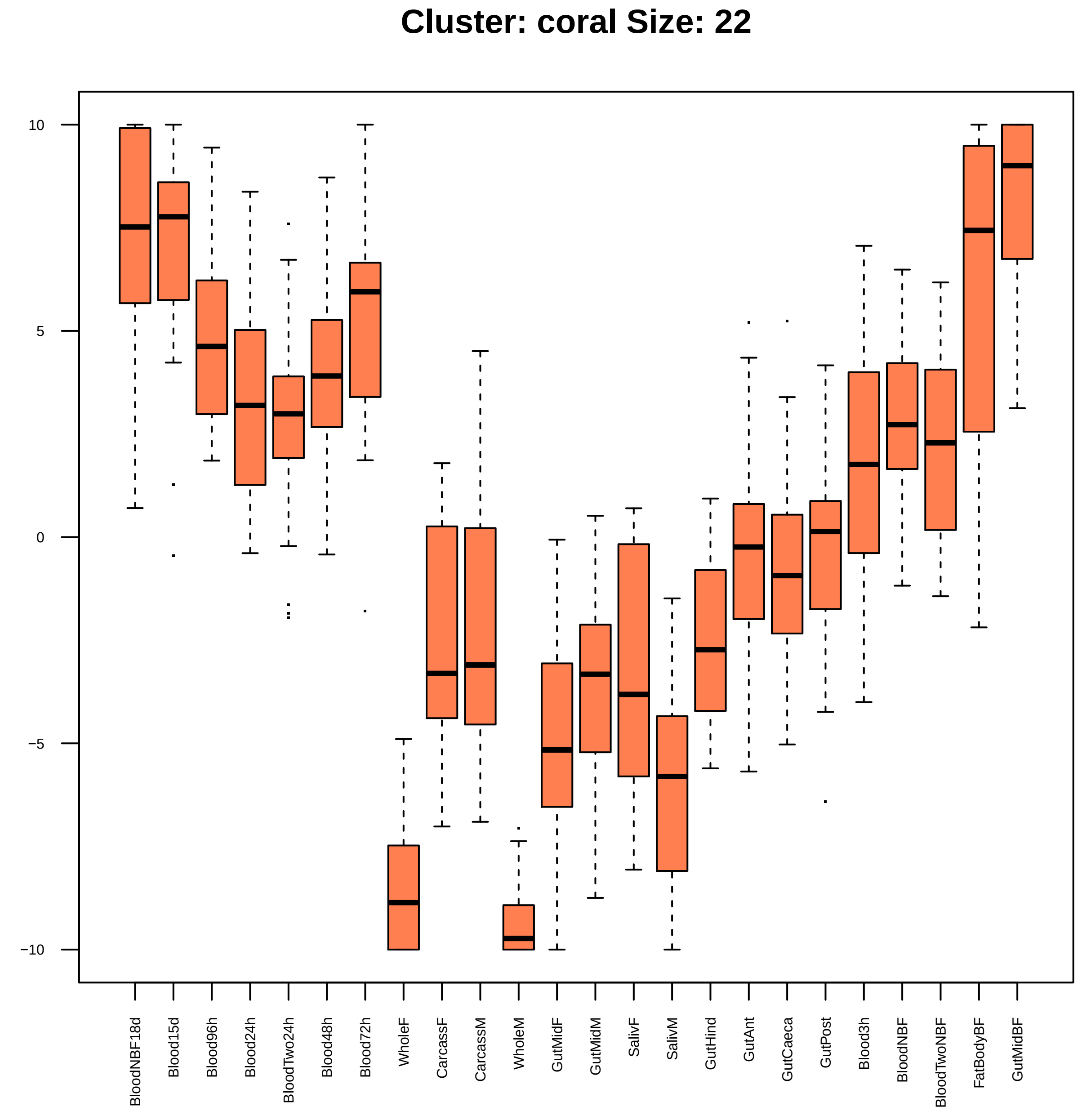
Cluster: coral Size: 22
10
5
0
−5
−10
BloodNBF18d
Blood15d
Blood96h
Blood24h
BloodTwo24h
Blood48h
Blood72h
WholeF
CarcassF
CarcassM
WholeM
GutMidF
GutMidM
SalivF
SalivM
GutHind
GutAnt
GutCaeca
GutPost
Blood3h
BloodNBF
BloodTwoNBF
FatBodyBF
GutMidBF

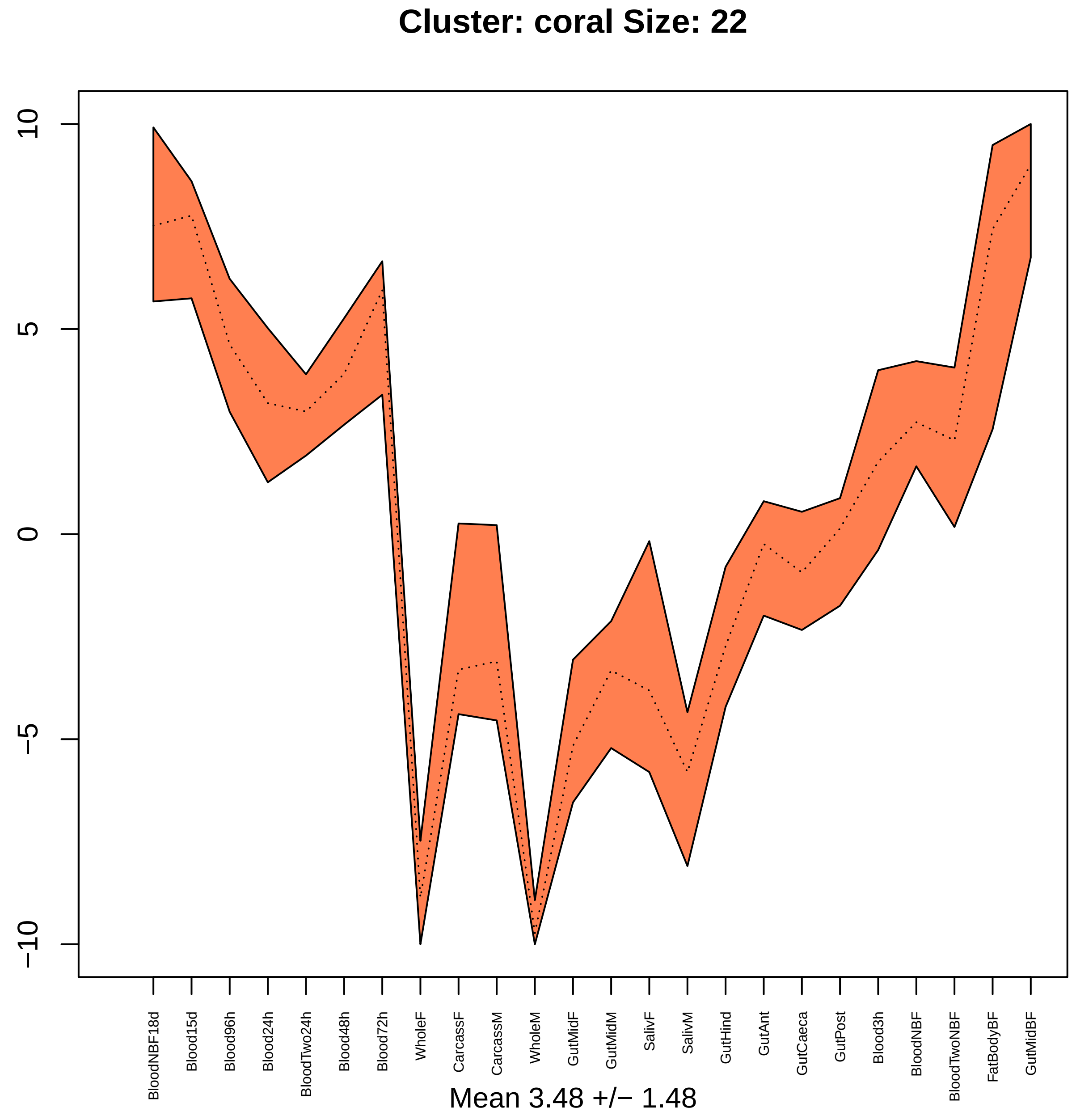

Cluster: coral Size: 22
10
5
0
−5
−10
BloodNBF18d
Blood15d
Blood96h
Blood24h
BloodTwo24h
Blood48h
Blood72h
WholeF
CarcassF
CarcassM
WholeM
GutMidF
GutMidM
SalivF
SalivM
GutHind
GutAnt
GutCaeca
GutPost
Blood3h
BloodNBF
BloodTwoNBF
FatBodyBF
GutMidBF
Mean 3.48 +/− 1.48

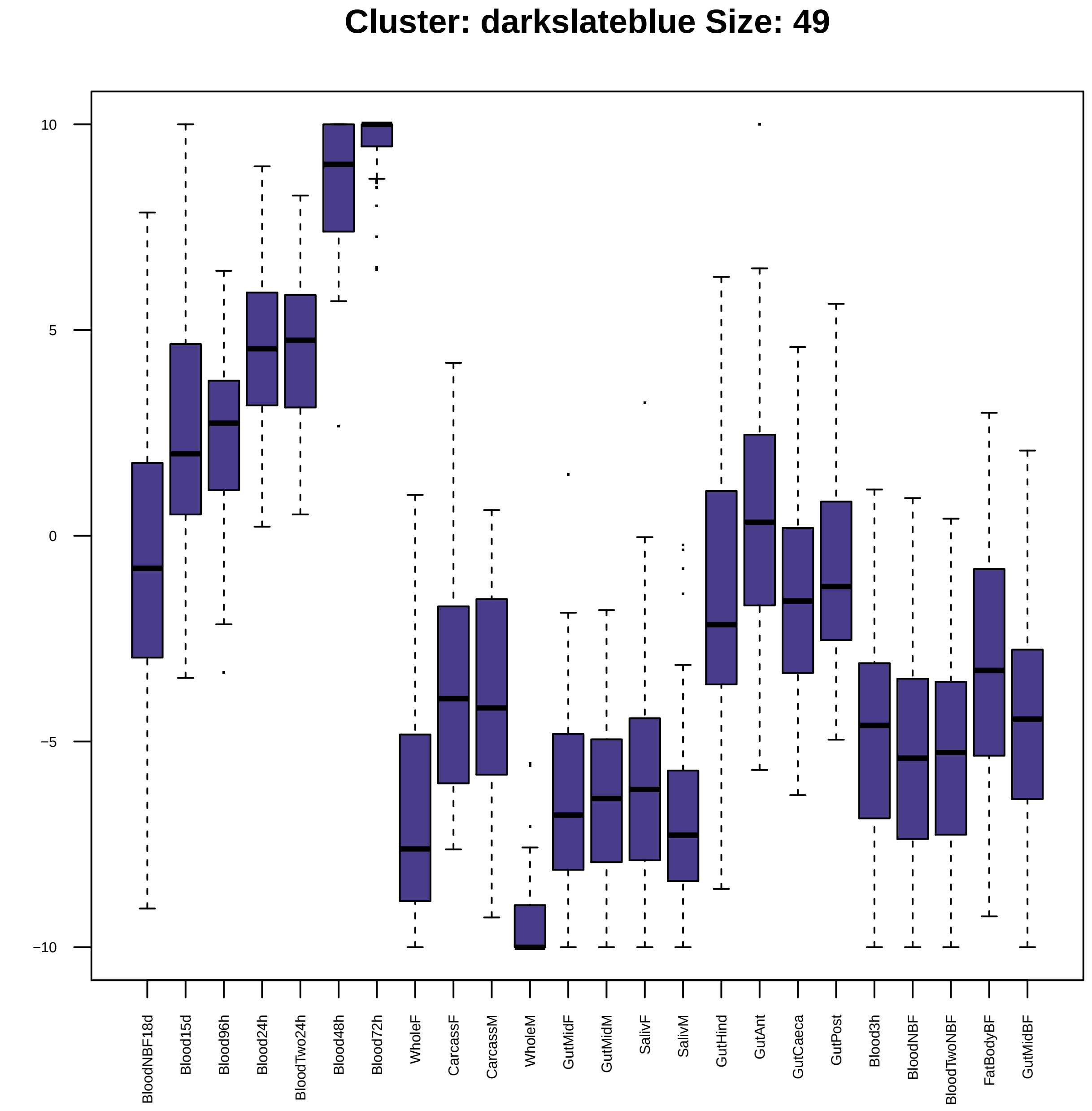

Cluster: darkslateblue Size: 49
10
5
0
−5
−10
BloodNBF18d
Blood15d
Blood96h
Blood24h
BloodTwo24h
Blood48h
Blood72h
WholeF
CarcassF
CarcassM
WholeM
GutMidF
GutMidM
SalivF
SalivM
GutHind
GutAnt
GutCaeca
GutPost
Blood3h
BloodNBF
BloodTwoNBF
FatBodyBF
GutMidBF

| | GO.ID | BPCluster: darkslateblue Size: 49 | Annotated | Significant | Expected | Rank in ClassicF | Weight01F | ClassicF |
|---|---|---|---|---|---|---|---|---|
| *1* | GO:0008589 | regulation of smoothened signaling pathw... | 17 | 2 | 0.07 | 2 | 0.0019 | 0.0019 |
| *2* | GO:0006338 | chromatin remodeling | 37 | 2 | 0.15 | 9 | 0.0090 | 0.0090 |
| *3* | GO:0016571 | histone methylation | 26 | 2 | 0.10 | 4 | 0.0150 | 0.0045 |

| | GO.ID | MFCluster: darkslateblue Size: 49 | Annotated | Significant | Expected | Rank in ClassicF | Weight01F | ClassicF |
|---|---|---|---|---|---|---|---|---|
| 1 | GO:0003729 | mRNA binding | 99 | 4 | 0.43 | 1 | 0.00079 | 0.00079 |
| 2 | GO:0008017 | microtubule binding | 55 | 3 | 0.24 | 2 | 0.00162 | 0.00162 |
| 4 | GO:0042054 | histone methyltransferase activity | 17 | 2 | 0.07 | 6 | 0.01246 | 0.00234 |
| 6 | GO:0003677 | DNA binding | 523 | 7 | 2.25 | 10 | 0.01612 | 0.00577 |
| 9 | GO:0008170 | N−methyltransferase activity | 25 | 2 | 0.11 | 7 | 0.04495 | 0.00505 |
| 20 | GO:0008168 | methyltransferase activity | 88 | 3 | 0.38 | 11 | 0.14328 | 0.00617 |
| 30 | GO:0003676 | nucleic acid binding | 1152 | 12 | 4.95 | 4 | 0.65983 | 0.00191 |

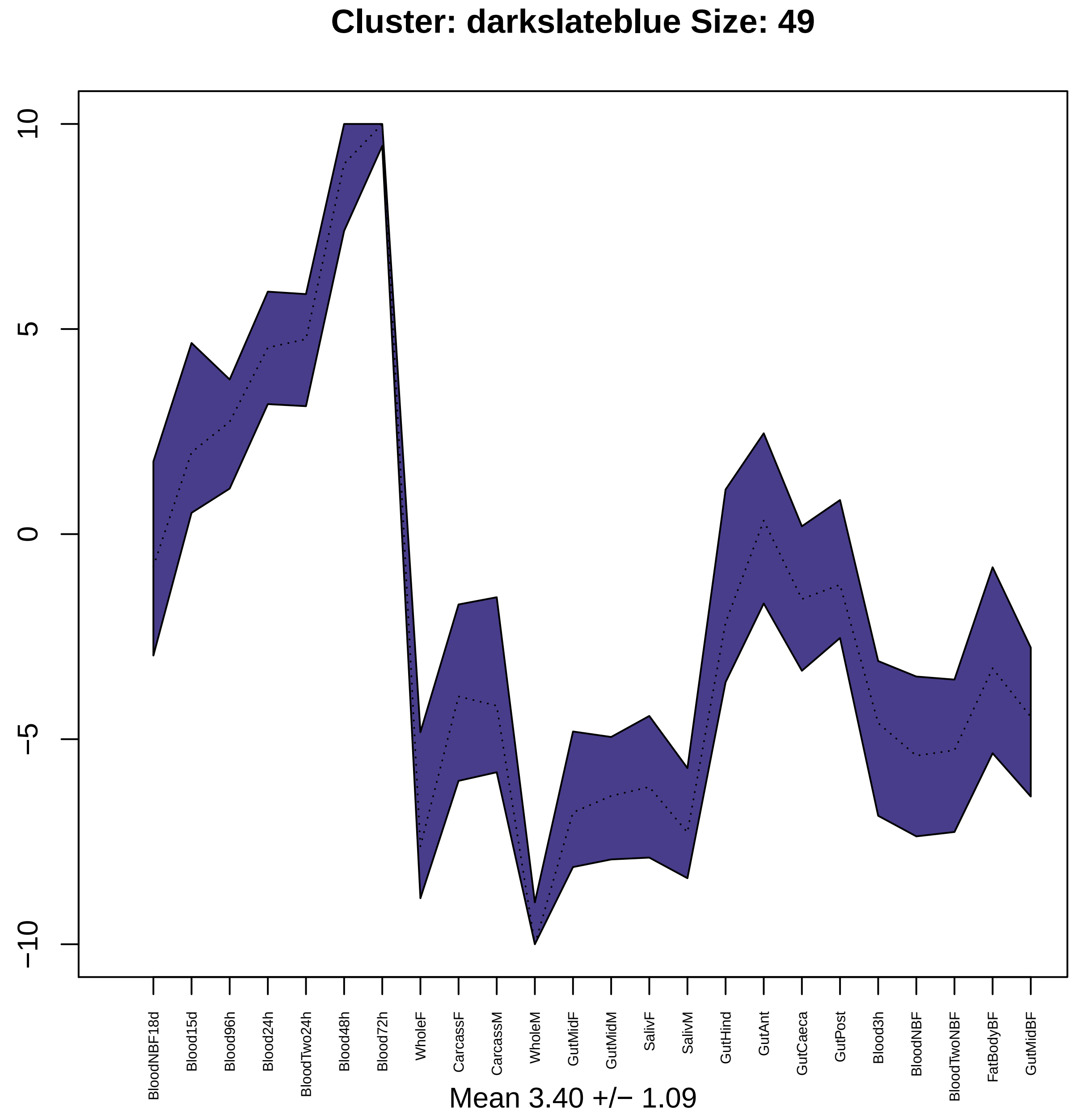

Cluster: darkslateblue Size: 49
10
5
0
−5
−10
BloodNBF18d
Blood15d
Blood96h
Blood24h
BloodTwo24h
Blood48h
Blood72h
WholeF
CarcassF
CarcassM
WholeM
GutMidF
GutMidM
SalivF
SalivM
GutHind
GutAnt
GutCaeca
GutPost
Blood3h
BloodNBF
BloodTwoNBF
FatBodyBF
GutMidBF
Mean 3.40 +/− 1.09

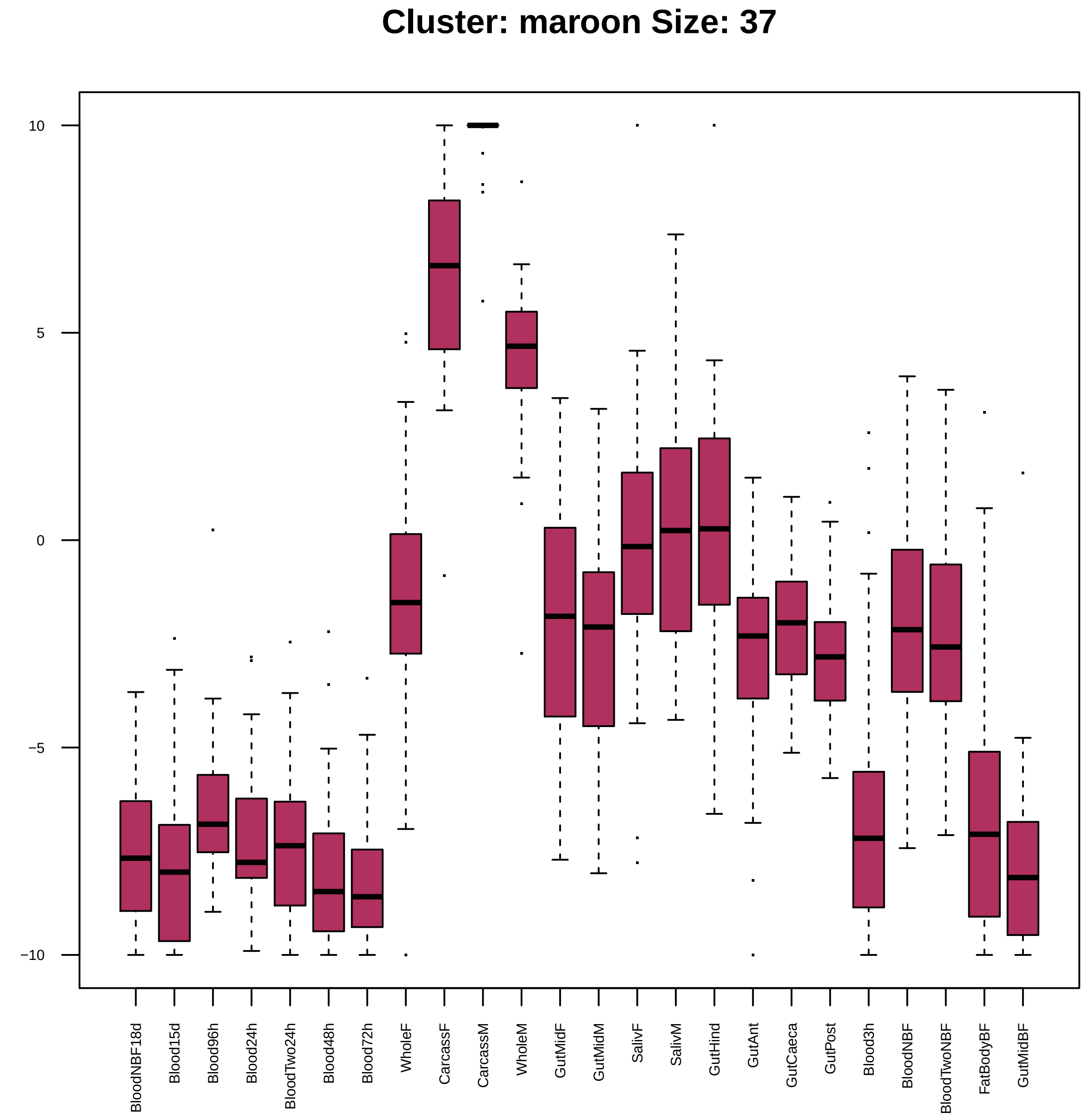
Cluster: maroon Size: 37
10
5
0
−5
−10
BloodNBF18d
Blood15d
Blood96h
Blood24h
BloodTwo24h
Blood48h
Blood72h
WholeF
CarcassF
CarcassM
WholeM
GutMidF
GutMidM
SalivF
SalivM
GutHind
GutAnt
GutCaeca
GutPost
Blood3h
BloodNBF
BloodTwoNBF
FatBodyBF
GutMidBF

| | GO.ID | BPCluster: maroon Size: 37 | Annotated | Significant | Expected | Rank in ClassicF | Weight01F | ClassicF |
|---|---|---|---|---|---|---|---|---|
| *1* | GO:0042775 | mitochondrial ATP synthesis coupled elec... | 29 | 4 | 0.11 | 17 | 0.00074 | 3.4e−06 |
| *2* | GO:0043648 | dicarboxylic acid metabolic process | 16 | 2 | 0.06 | 32 | 0.00158 | 0.0016 |
| *3* | GO:0006120 | mitochondrial electron transport, NADH t... | 17 | 2 | 0.06 | 33 | 0.00178 | 0.0018 |
| *4* | GO:0006099 | tricarboxylic acid cycle | 27 | 2 | 0.10 | 37 | 0.00450 | 0.0045 |
| *5* | GO:0022900 | electron transport chain | 37 | 5 | 0.14 | 9 | 0.00943 | 1.9e−07 |
| *6* | GO:0009060 | aerobic respiration | 31 | 3 | 0.12 | 30 | 0.01379 | 0.0002 |
| *7* | GO:0006733 | oxidoreduction coenzyme metabolic proces... | 34 | 2 | 0.13 | 45 | 0.02509 | 0.0071 |
| *16* | GO:0044237 | cellular metabolic process | 2862 | 17 | 10.77 | 47 | 0.05303 | 0.0091 |

| | GO.ID | MFCluster: maroon Size: 37 | Annotated | Significant | Expected | Rank in ClassicF | Weight01F | ClassicF |
|---|---|---|---|---|---|---|---|---|
| *1* | GO:0008137 | NADH dehydrogenase (ubiquinone) activity | 20 | 2 | 0.06 | 4 | 0.0018 | 0.00179 |
| *2* | GO:0016836 | hydro−lyase activity | 28 | 2 | 0.09 | 8 | 0.0035 | 0.00350 |
| *5* | GO:0016651 | oxidoreductase activity, acting on NAD(P... | 35 | 3 | 0.11 | 2 | 0.0289 | 0.00017 |
| *24* | GO:0003824 | catalytic activity | 3160 | 18 | 10.08 | 3 | 0.2428 | 0.00082 |

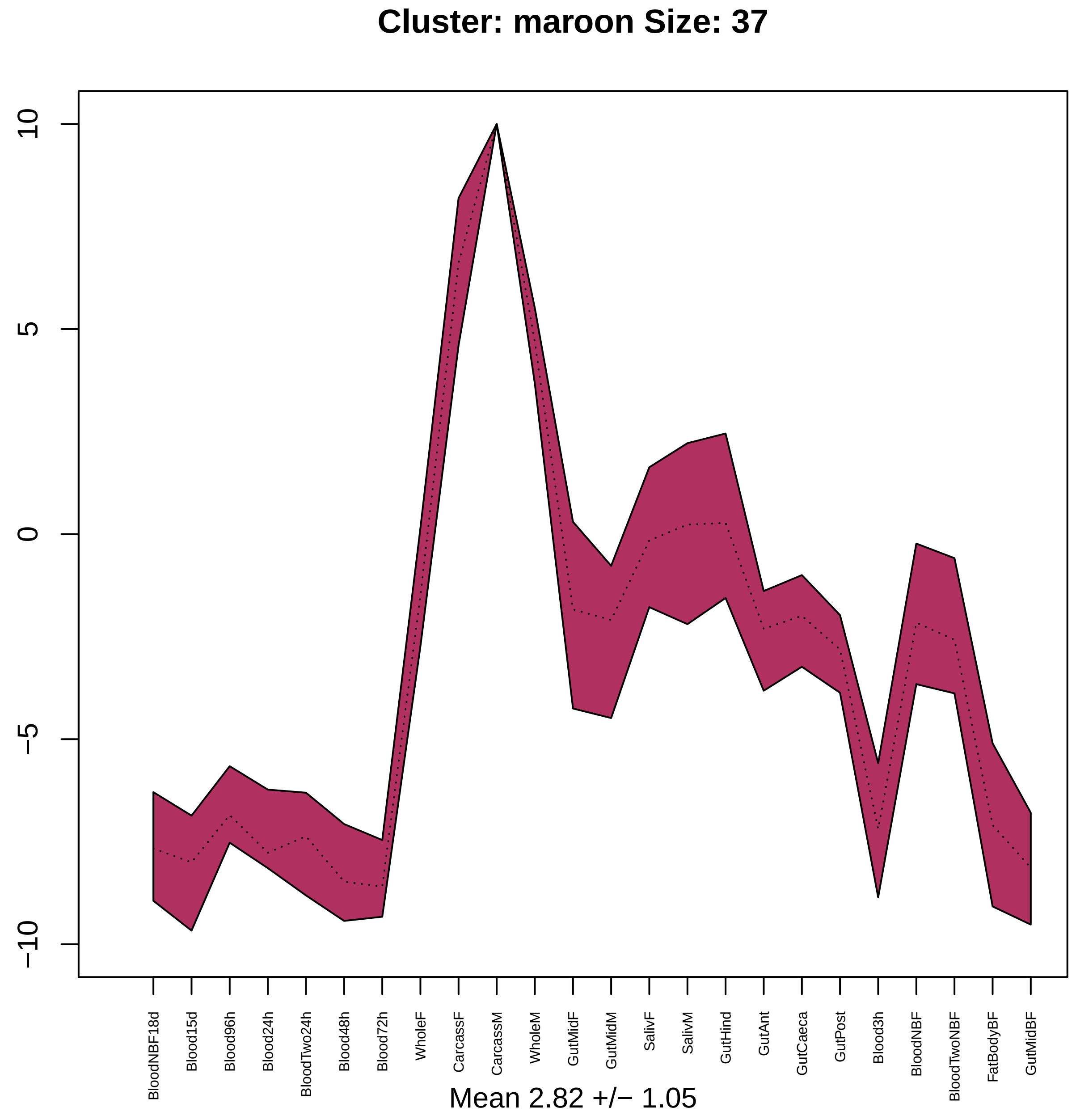

Cluster: maroon Size: 37
10
5
0
−5
−10
BloodNBF18d
Blood15d
Blood96h
Blood24h
BloodTwo24h
Blood48h
Blood72h
WholeF
CarcassF
CarcassM
WholeM
GutMidF
GutMidM
SalivF
SalivM
GutHind
GutAnt
GutCaeca
GutPost
Blood3h
BloodNBF
BloodTwoNBF
FatBodyBF
GutMidBF
Mean 2.82 +/− 1.05

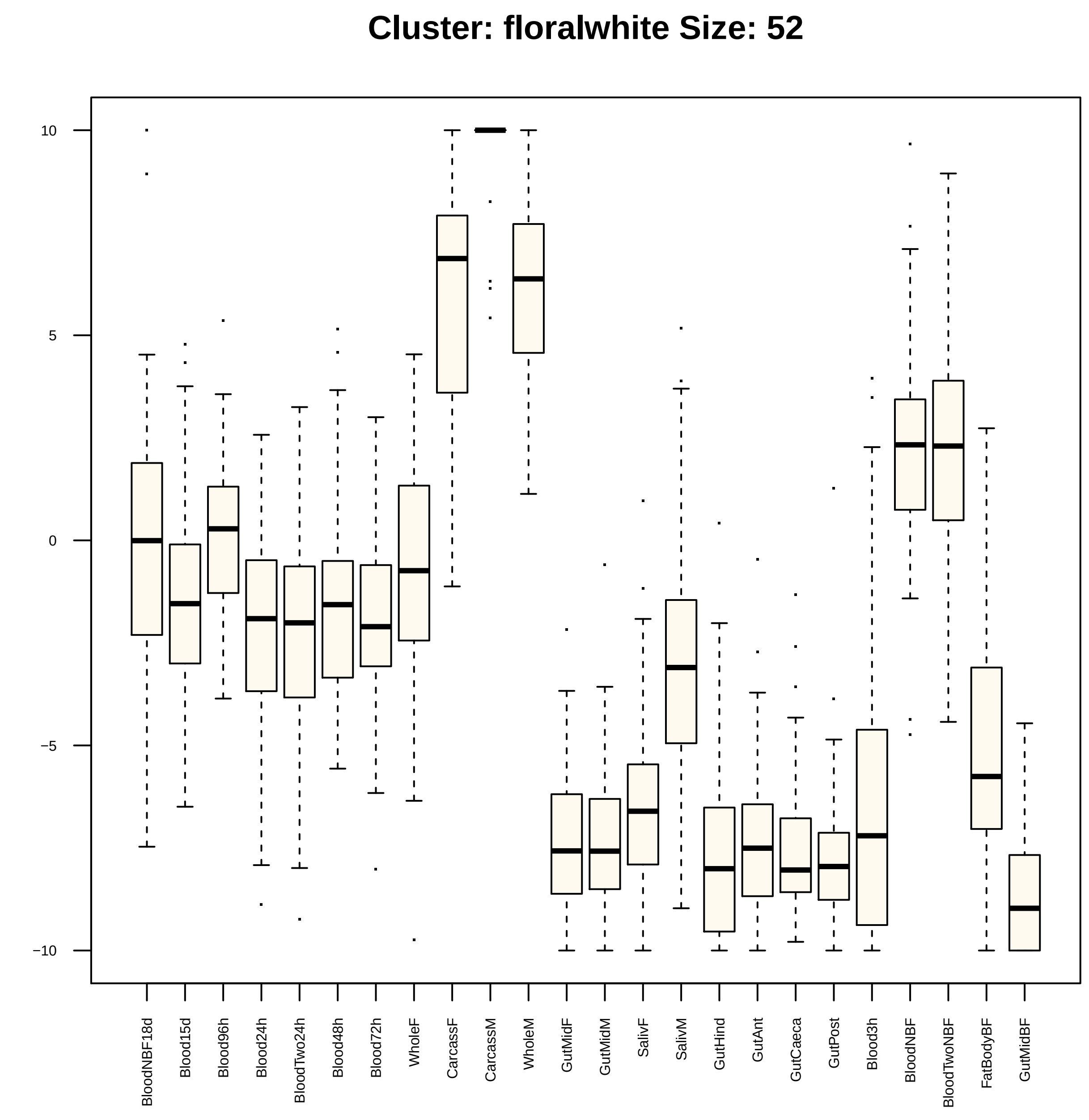

Cluster: floralwhite Size: 52
10
5
0
−5
−10
BloodNBF18d
Blood15d
Blood96h
Blood24h
BloodTwo24h
Blood48h
Blood72h
WholeF
CarcassF
CarcassM
WholeM
GutMidF
GutMidM
SalivF
SalivM
GutHind
GutAnt
GutCaeca
GutPost
Blood3h
BloodNBF
BloodTwoNBF
FatBodyBF
GutMidBF

| | GO.ID | BPCluster: floralwhite Size: 52 | Annotated | Significant | Expected | Rank in ClassicF | Weight01F | ClassicF |
|---|---|---|---|---|---|---|---|---|
| 1 | GO:0035335 | peptidyl−tyrosine dephosphorylation | 30 | 2 | 0.10 | 3 | 0.0047 | 0.0047 |
| 13 | GO:0009166 | nucleotide catabolic process | 25 | 2 | 0.09 | 1 | 0.0421 | 0.0032 |

| | GO.ID | MFCluster: floralwhite Size: 52 | Annotated | Significant | Expected | Rank in ClassicF | Weight01F | ClassicF |
|---|---|---|---|---|---|---|---|---|
| *1* | GO:0004725 | protein tyrosine phosphatase activity | 32 | 2 | 0.12 | 2 | 0.0063 | 0.0063 |
| *17* | GO:0016791 | phosphatase activity | 97 | 3 | 0.36 | 1 | 0.1249 | 0.0055 |

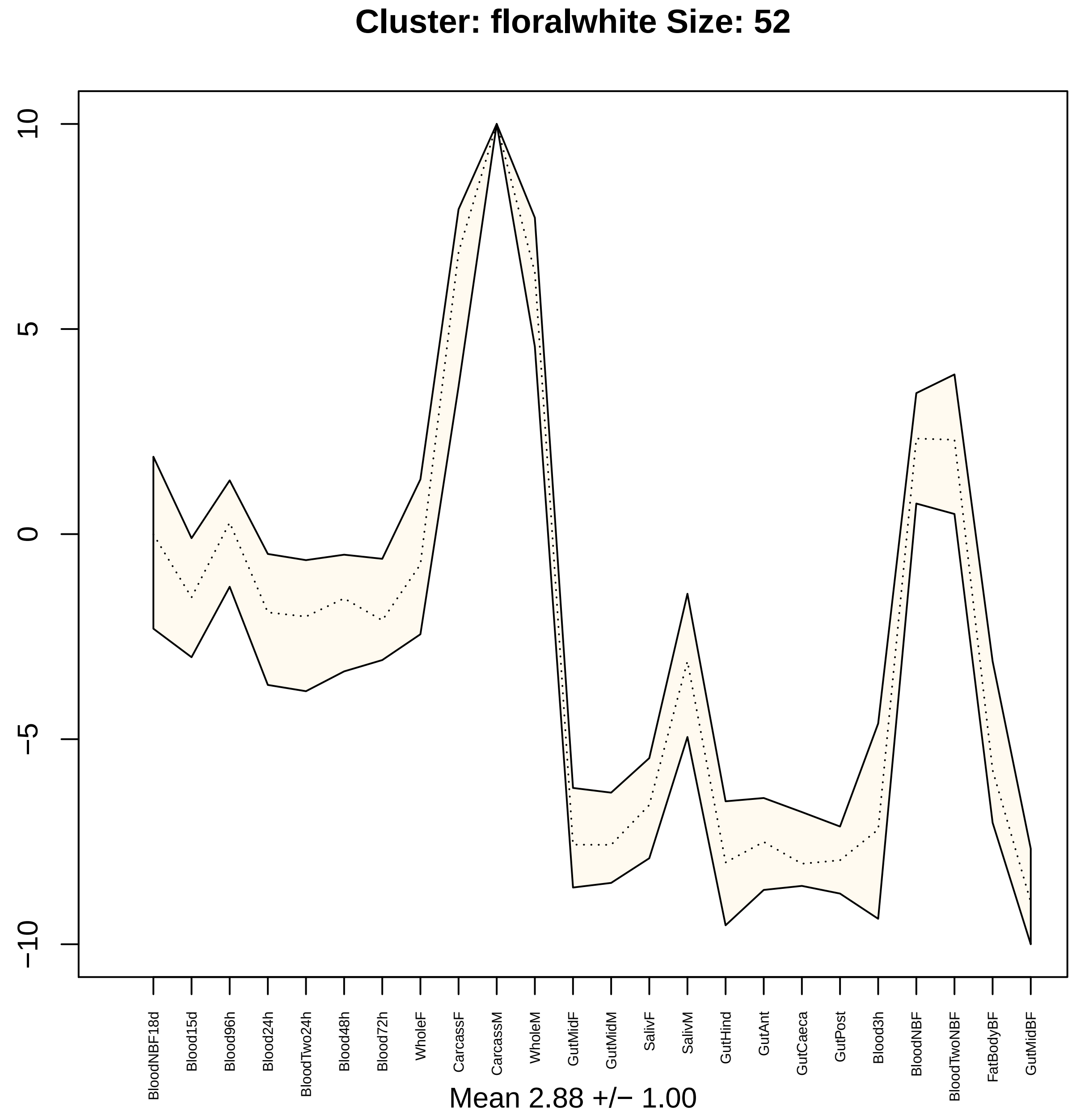

Cluster: floralwhite Size: 52
10
5
0
−5
−10
BloodNBF18d
Blood15d
Blood96h
Blood24h
BloodTwo24h
Blood48h
Blood72h
WholeF
CarcassF
CarcassM
WholeM
GutMidF
GutMidM
SalivF
SalivM
GutHind
GutAnt
GutCaeca
GutPost
Blood3h
BloodNBF
BloodTwoNBF
FatBodyBF
GutMidBF
Mean 2.88 +/− 1.00

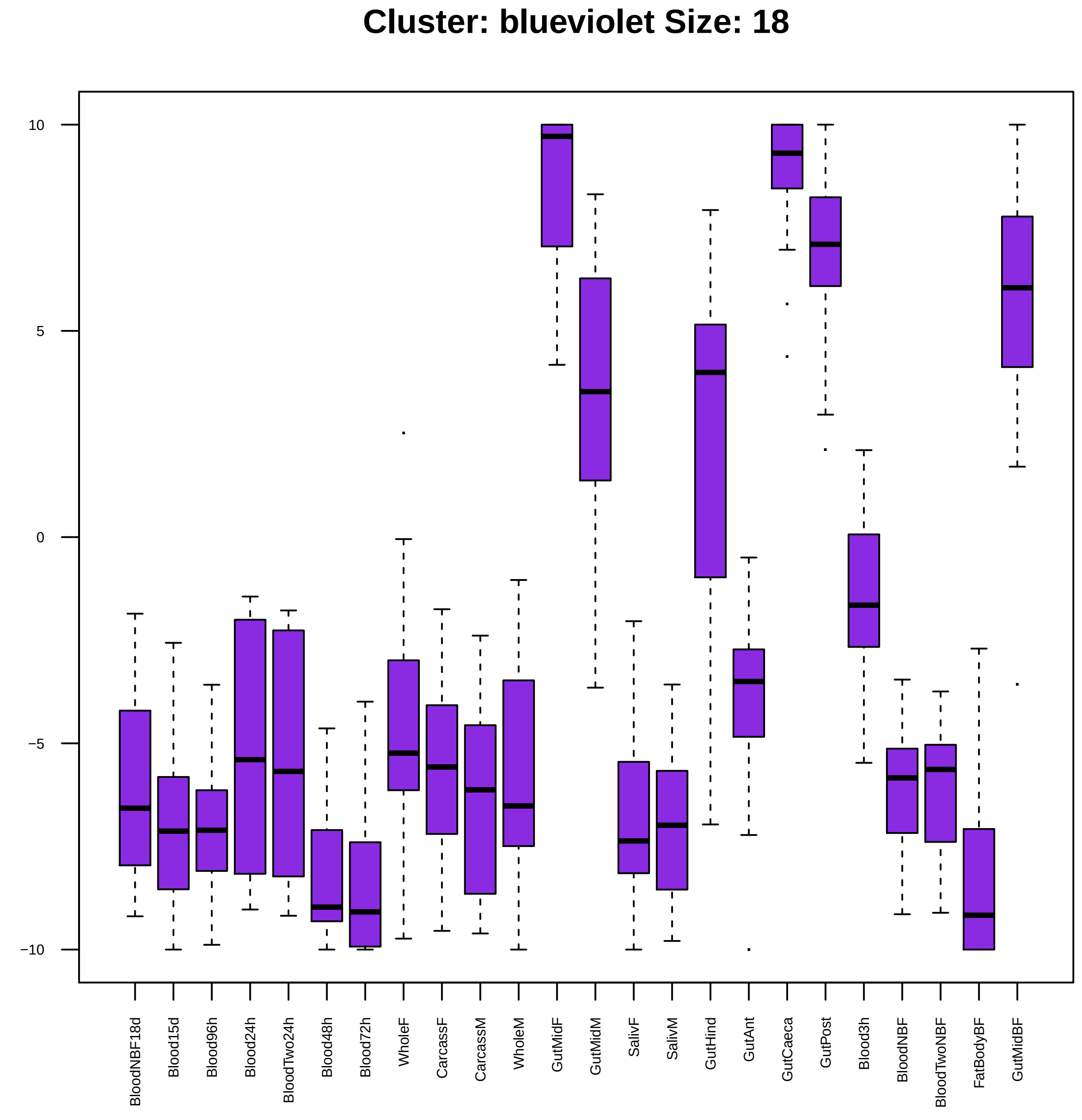

Cluster: blueviolet Size: 18
10
5
0
−5
−10
BloodNBF18d
Blood15d
Blood96h
Blood24h
BloodTwo24h
Blood48h
Blood72h
WholeF
CarcassF
CarcassM
WholeM
GutMidF
GutMidM
SalivF
SalivM
GutHind
GutAnt
GutCaeca
GutPost
Blood3h
BloodNBF
BloodTwoNBF
FatBodyBF
GutMidBF

| | GO.ID | BPCluster: blueviolet Size: 18 | Annotated | Significant | Expected | Rank in ClassicF | Weight01F | ClassicF |
|---|---|---|---|---|---|---|---|---|
| 1 | GO:0016311 | dephosphorylation | 98 | 3 | 0.25 | 1 | 0.0017 | 0.0017 |

| | GO.ID | MFCluster: blueviolet Size: 18 | Annotated | Significant | Expected | Rank in ClassicF | Weight01F | ClassicF |
|---|---|---|---|---|---|---|---|---|
| *1* | GO:0016791 | phosphatase activity | 97 | 3 | 0.19 | 1 | 0.00077 | 0.00077 |
| *9* | GO:0016788 | hydrolase activity, acting on ester bond... | 269 | 4 | 0.52 | 2 | 0.20357 | 0.00141 |
| *11* | GO:0016787 | hydrolase activity | 1445 | 8 | 2.81 | 4 | 0.54006 | 0.00241 |
| *19* | GO:0008238 | exopeptidase activity | 66 | 2 | 0.13 | 5 | 1.00000 | 0.00700 |

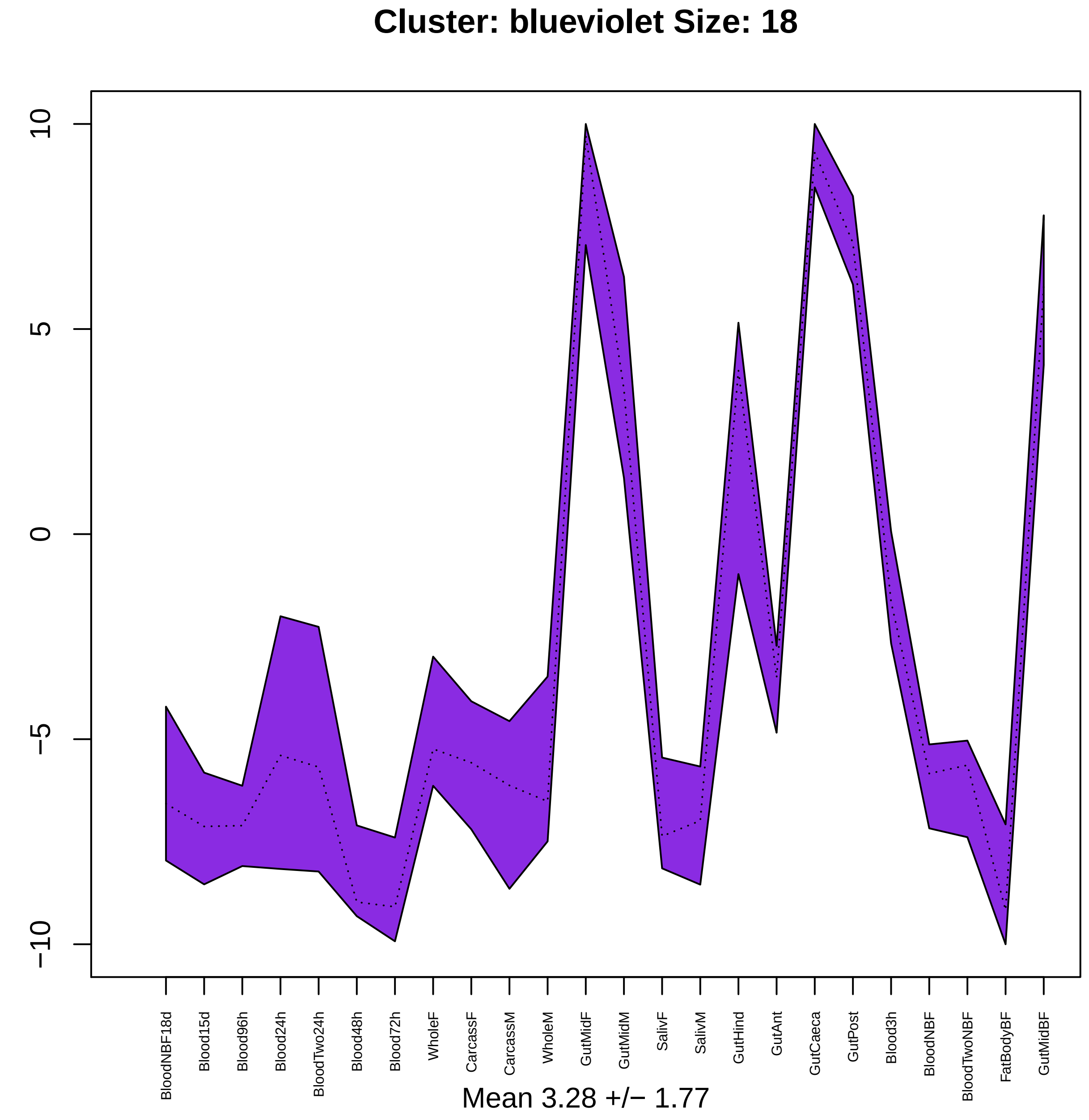
Cluster: blueviolet Size: 18
10
5
0
−5
−10
BloodNBF18d
Blood15d
Blood96h
Blood24h
BloodTwo24h
Blood48h
Blood72h
WholeF
CarcassF
CarcassM
WholeM
GutMidF
GutMidM
SalivF
SalivM
GutHind
GutAnt
GutCaeca
GutPost
Blood3h
BloodNBF
BloodTwoNBF
FatBodyBF
GutMidBF
Mean 3.28 +/− 1.77

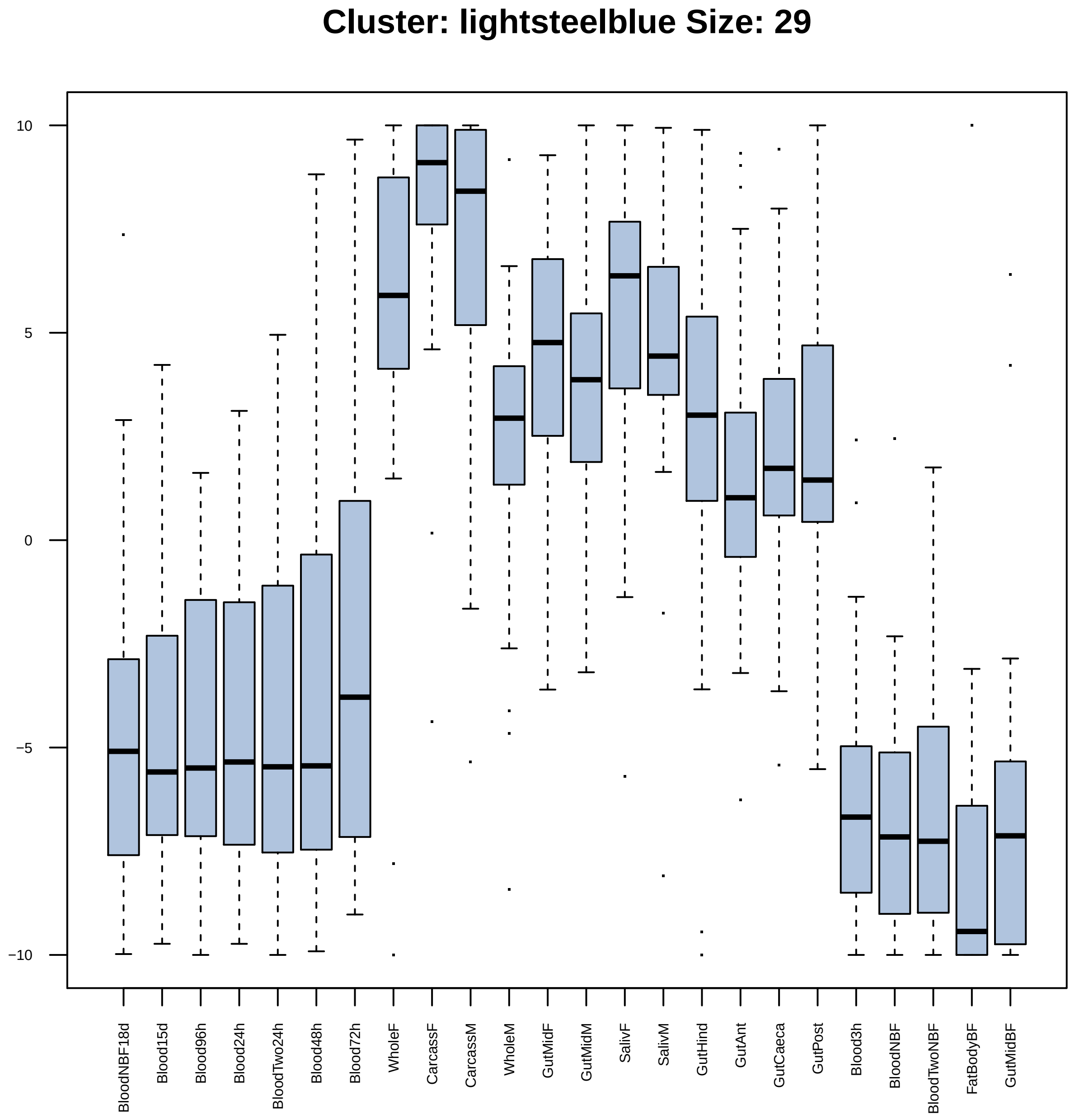
Cluster: lightsteelblue Size: 29
10
5
0
−5
−10
BloodNBF18d
Blood15d
Blood96h
Blood24h
BloodTwo24h
Blood48h
Blood72h
WholeF
CarcassF
CarcassM
WholeM
GutMidF
GutMidM
SalivF
SalivM
GutHind
GutAnt
GutCaeca
GutPost
Blood3h
BloodNBF
BloodTwoNBF
FatBodyBF
GutMidBF

| | GO.ID | BPCluster: lightsteelblue Size: 29 | Annotated | Significant | Expected | Rank in ClassicF | Weight01F | ClassicF |
|---|---|---|---|---|---|---|---|---|
| 1 | GO:0006979 | response to oxidative stress | 51 | 2 | 0.14 | 22 | 0.0088 | 0.00884 |
| 2 | GO:0006412 | translation | 289 | 5 | 0.82 | 6 | 0.0152 | 0.00097 |
| 3 | GO:0000398 | mRNA splicing, via spliceosome | 134 | 3 | 0.38 | 14 | 0.0246 | 0.00588 |

| | GO.ID | MFCluster: lightsteelblue Size: 29 | Annotated | Significant | Expected | Rank in ClassicF | Weight01F | ClassicF |
|---|---|---|---|---|---|---|---|---|
| *1* | GO:0003735 | structural constituent of ribosome | 146 | 3 | 0.43 | 1 | 0.0083 | 0.0083 |

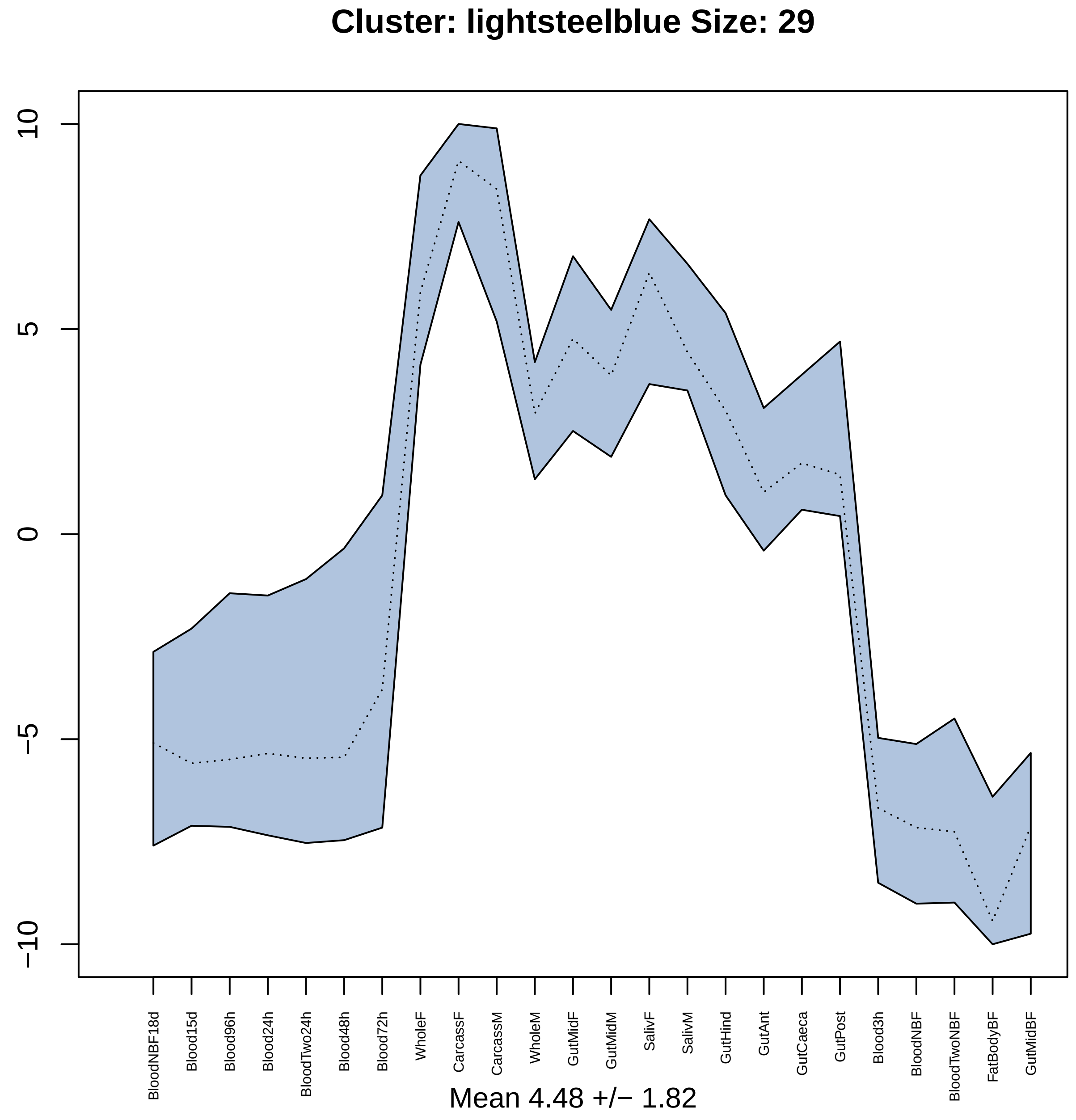
Cluster: lightsteelblue Size: 29
10
5
0
−5
−10
BloodNBF18d
Blood15d
Blood96h
Blood24h
BloodTwo24h
Blood48h
Blood72h
WholeF
CarcassF
CarcassM
WholeM
GutMidF
GutMidM
SalivF
SalivM
GutHind
GutAnt
GutCaeca
GutPost
Blood3h
BloodNBF
BloodTwoNBF
FatBodyBF
GutMidBF
Mean 4.48 +/− 1.82

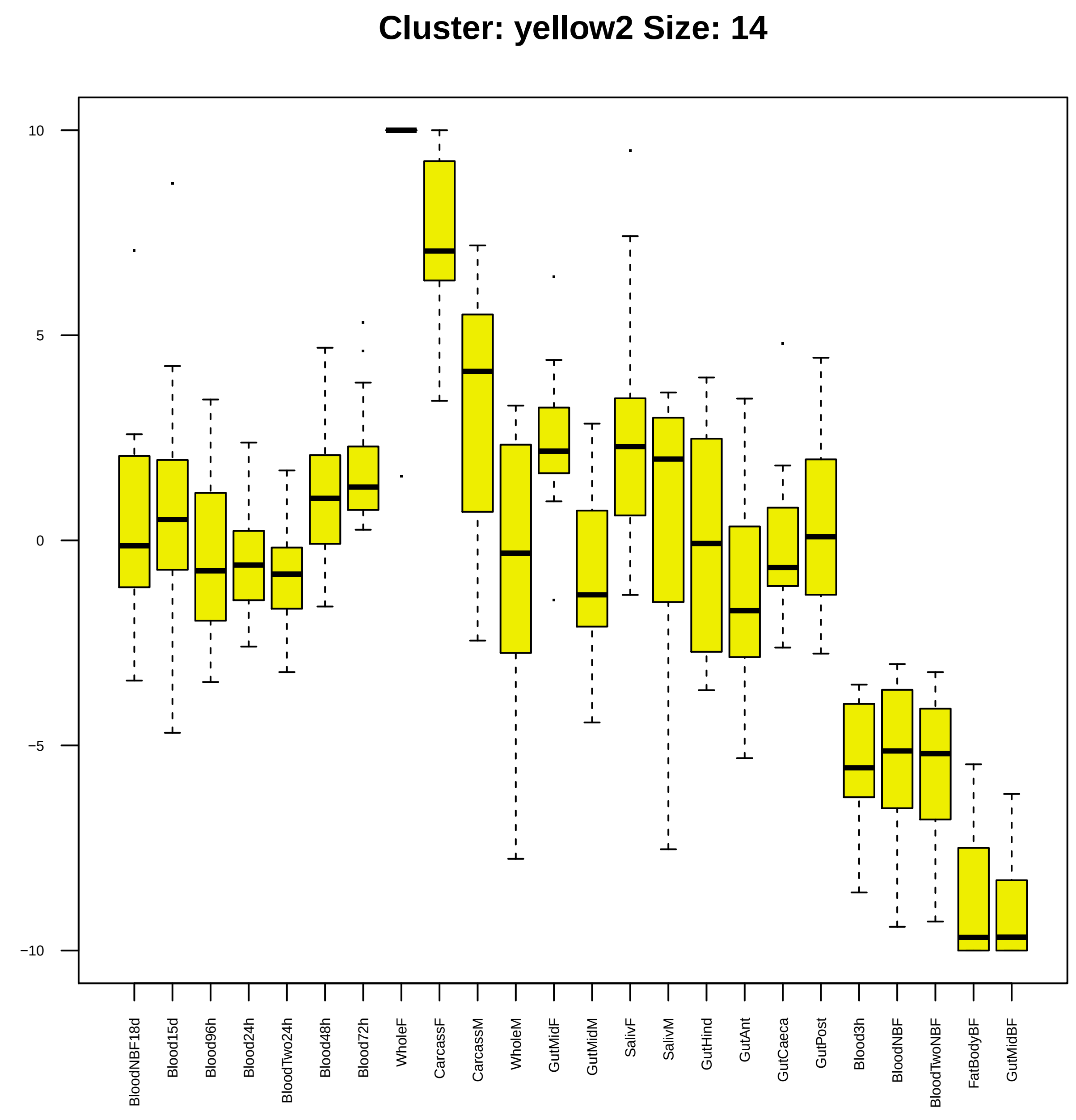

Cluster: yellow2 Size: 14
10
5
0
−5
−10
BloodNBF18d
Blood15d
Blood96h
Blood24h
BloodTwo24h
Blood48h
Blood72h
WholeF
CarcassF
CarcassM
WholeM
GutMidF
GutMidM
SalivF
SalivM
GutHind
GutAnt
GutCaeca
GutPost
Blood3h
BloodNBF
BloodTwoNBF
FatBodyBF
GutMidBF

| | GO.ID | BPCluster: yellow2 Size: 14 | Annotated | Significant | Expected | Rank in ClassicF | Weight01F | ClassicF |
|---|---|---|---|---|---|---|---|---|
| *13* | GO:0016070 | RNA metabolic process | 906 | 4 | 0.85 | 1 | 1.000 | 0.0048 |

|  | GO.ID | MFCluster: yellow2 Size: 14 | Annotated | Significant | Expected | Rank in ClassicF | Weight01F | ClassicF |
|---|---|---|---|---|---|---|---|---|
| 7 | GO:0140098 | catalytic activity, acting on RNA | 165 | 2 | 0.09 | 5 | 1.000 | 0.00303 |
| 11 | GO:0016779 | nucleotidyltransferase activity | 61 | 2 | 0.03 | 4 | 1.000 | 0.00042 |
| 13 | GO:0016740 | transferase activity | 911 | 3 | 0.51 | 6 | 1.000 | 0.00729 |

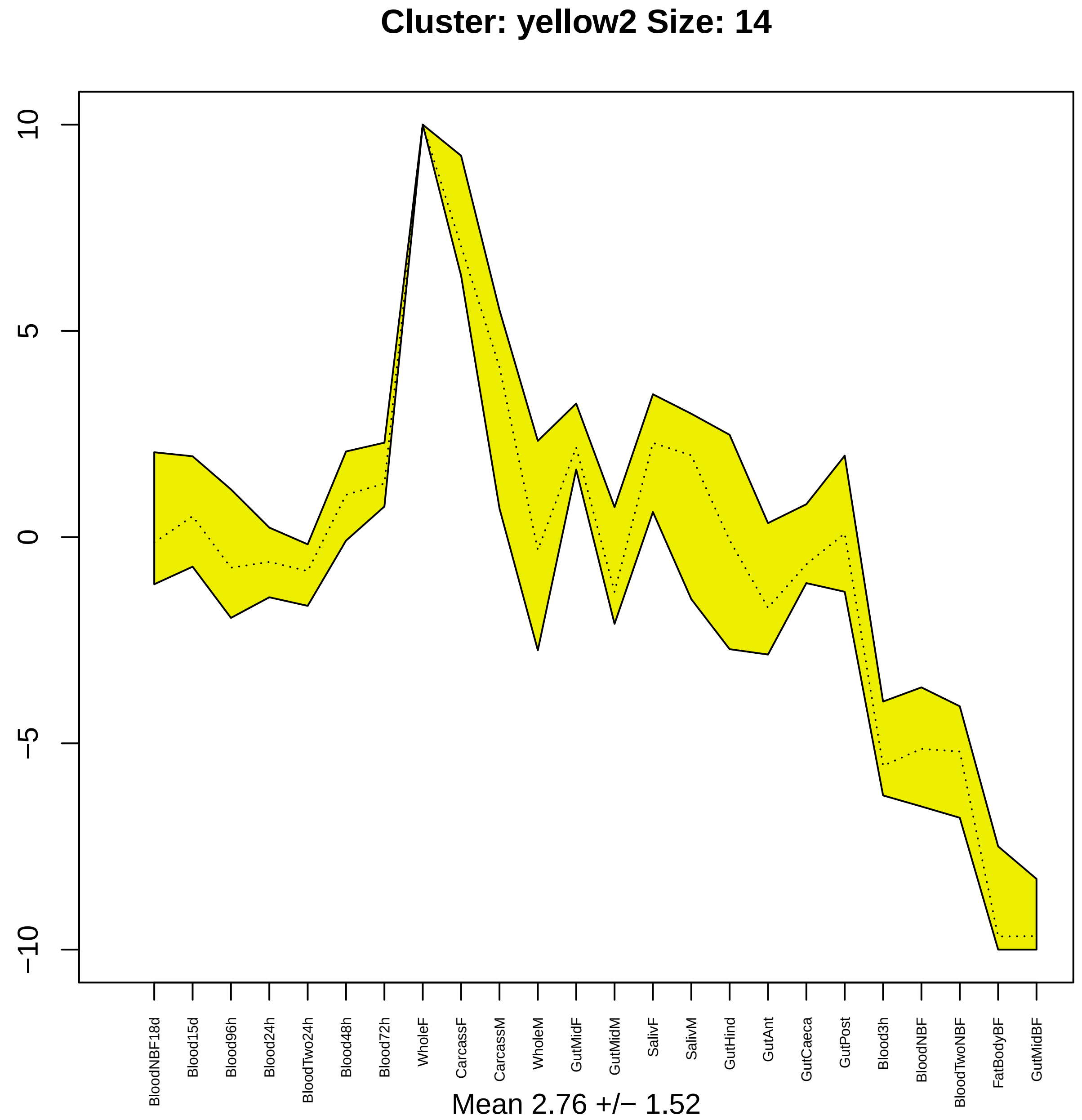

Cluster: yellow2 Size: 14
10
5
0
−5
−10
BloodNBF18d
Blood15d
Blood96h
Blood24h
BloodTwo24h
Blood48h
Blood72h
WholeF
CarcassF
CarcassM
WholeM
GutMidF
GutMidM
SalivF
SalivM
GutHind
GutAnt
GutCaeca
GutPost
Blood3h
BloodNBF
BloodTwoNBF
FatBodyBF
GutMidBF
Mean 2.76 +/− 1.52

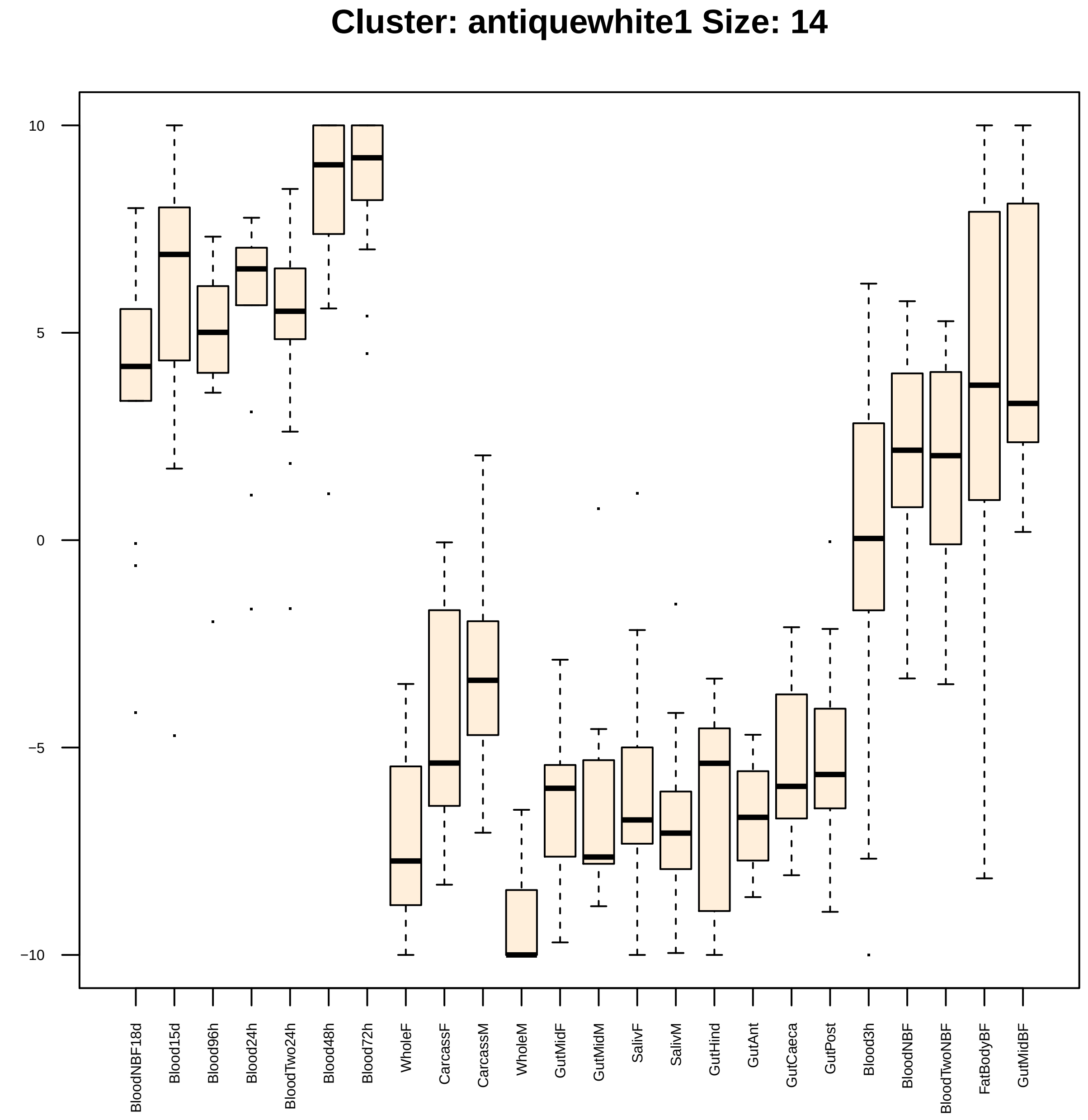
Cluster: antiquewhite1 Size: 14
10
5
0
−5
−10
BloodNBF18d
Blood15d
Blood96h
Blood24h
BloodTwo24h
Blood48h
Blood72h
WholeF
CarcassF
CarcassM
WholeM
GutMidF
GutMidM
SalivF
SalivM
GutHind
GutAnt
GutCaeca
GutPost
Blood3h
BloodNBF
BloodTwoNBF
FatBodyBF
GutMidBF

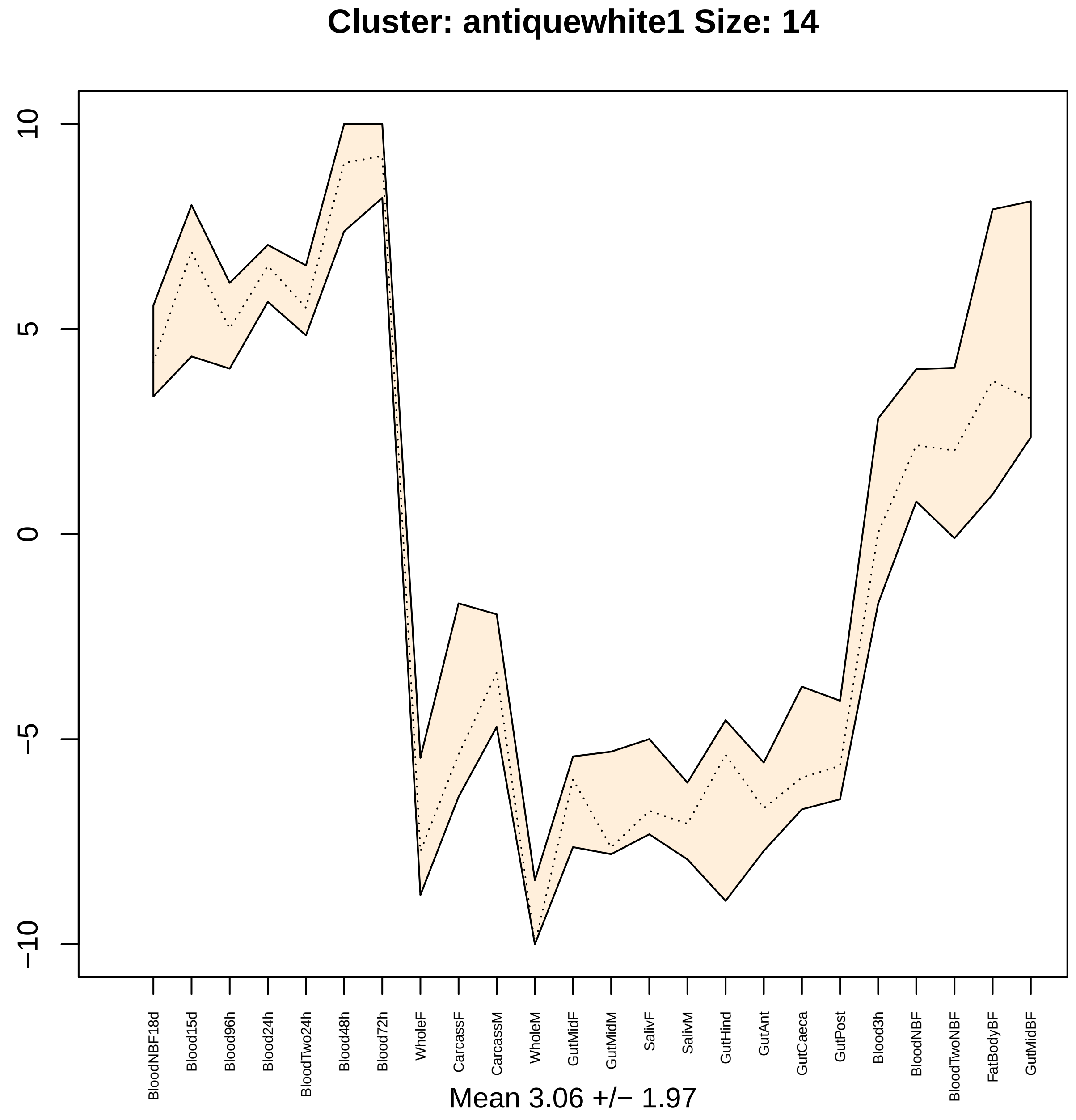

Cluster: antiquewhite1 Size: 14
10
5
0
−5
−10
BloodNBF18d
Blood15d
Blood96h
Blood24h
BloodTwo24h
Blood48h
Blood72h
WholeF
CarcassF
CarcassM
WholeM
GutMidF
GutMidM
SalivF
SalivM
GutHind
GutAnt
GutCaeca
GutPost
Blood3h
BloodNBF
BloodTwoNBF
FatBodyBF
GutMidBF
Mean 3.06 +/− 1.97

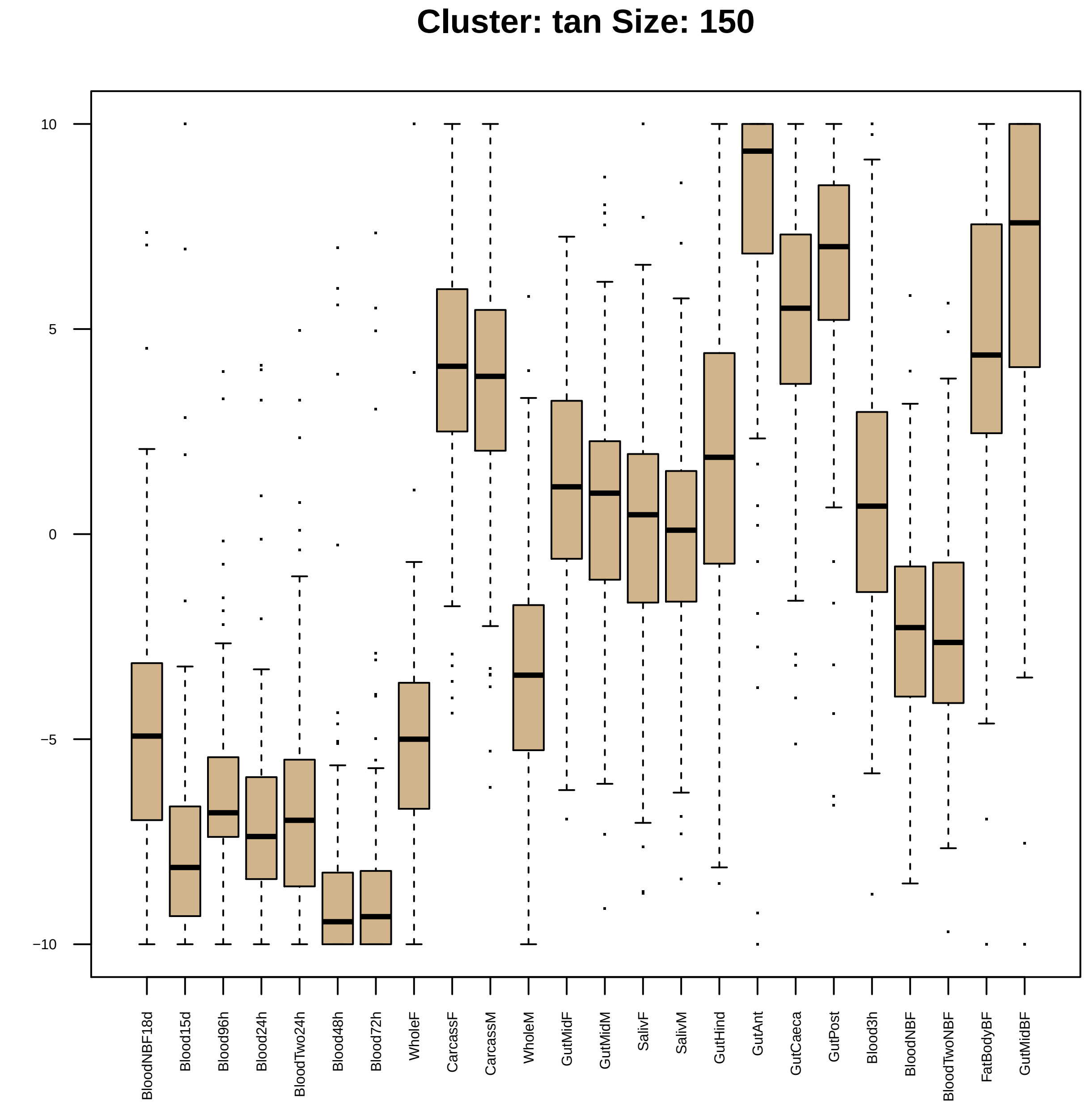
Cluster: tan Size: 150
10
5
0
−5
−10
BloodNBF18d
Blood15d
Blood96h
Blood24h
BloodTwo24h
Blood48h
Blood72h
WholeF
CarcassF
CarcassM
WholeM
GutMidF
GutMidM
SalivF
SalivM
GutHind
GutAnt
GutCaeca
GutPost
Blood3h
BloodNBF
BloodTwoNBF
FatBodyBF
GutMidBF

| | GO.ID | BPCluster: tan Size: 150 | Annotated | Significant | Expected | Rank in ClassicF | Weight01F | ClassicF |
|---|---|---|---|---|---|---|---|---|
| *4* | GO:0035235 | ionotropic glutamate receptor signaling ... | 28 | 3 | 0.34 | 22 | 0.0046 | 0.0046 |
| *11* | GO:0048749 | compound eye development | 157 | 7 | 1.92 | 17 | 0.0419 | 0.0029 |

| | GO.ID | MFCluster: tan Size: 150 | Annotated | Significant | Expected | Rank in ClassicF | Weight01F | ClassicF |
|---|---|---|---|---|---|---|---|---|
| *3* | GO:0005549 | odorant binding | 118 | 8 | 1.57 | 6 | 0.00016 | 0.00016 |
| *4* | GO:0004970 | ionotropic glutamate receptor activity | 29 | 3 | 0.39 | 19 | 0.00651 | 0.00651 |
| *5* | GO:0005234 | extracellularly glutamate−gated ion chan... | 29 | 3 | 0.39 | 20 | 0.00651 | 0.00651 |
| *6* | GO:0022835 | transmitter−gated channel activity | 30 | 4 | 0.40 | 9 | 0.01295 | 0.00062 |
| *8* | GO:0005230 | extracellular ligand−gated ion channel a... | 51 | 5 | 0.68 | 8 | 0.02972 | 0.00055 |
| *11* | GO:0003700 | DNA−binding transcription factor activit... | 282 | 10 | 3.76 | 16 | 0.05047 | 0.00413 |
| *23* | GO:0030594 | neurotransmitter receptor activity | 45 | 5 | 0.60 | 7 | 0.17606 | 0.00030 |

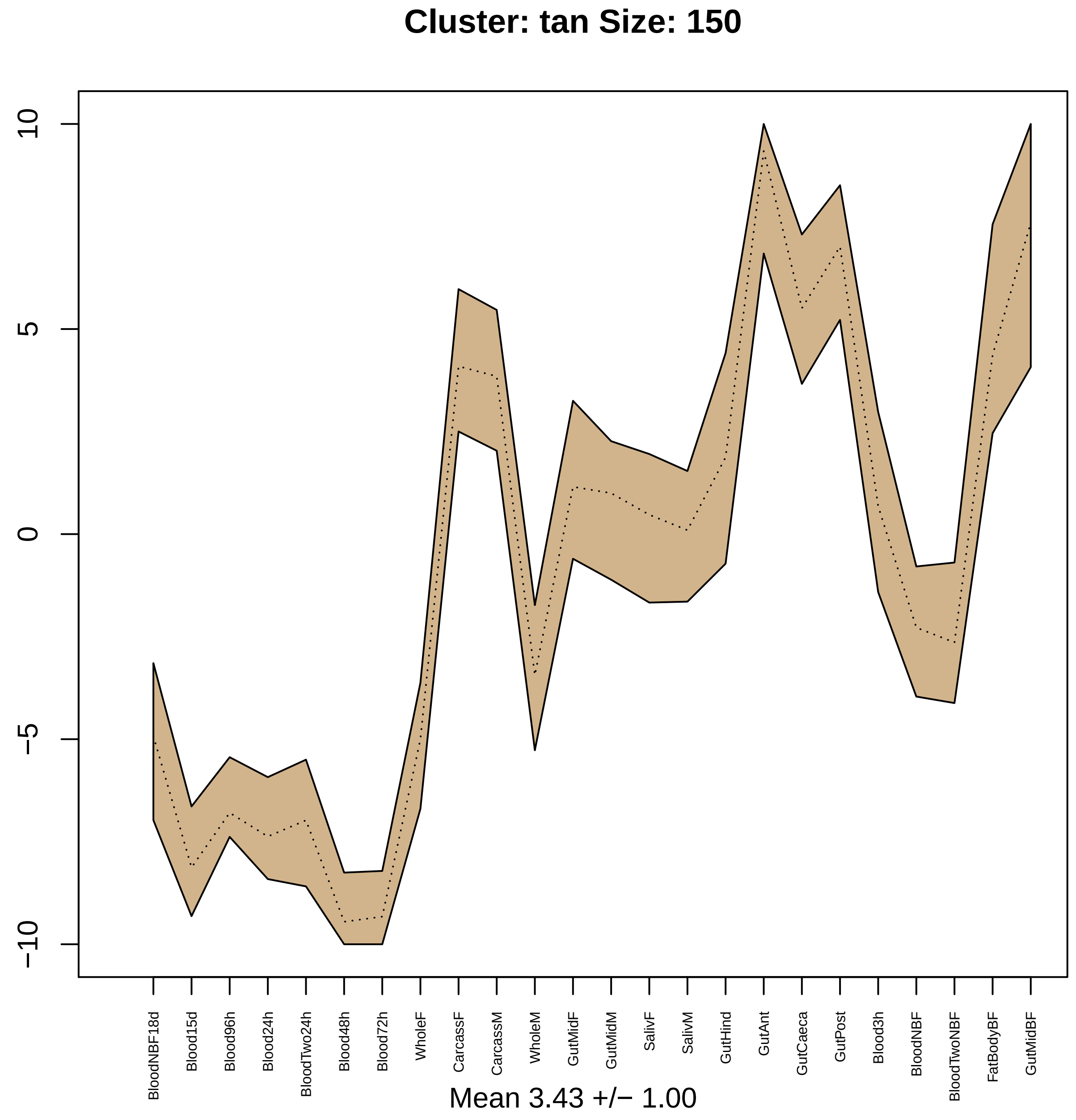

Cluster: tan Size: 150
10
5
0
−5
−10
BloodNBF18d
Blood15d
Blood96h
Blood24h
BloodTwo24h
Blood48h
Blood72h
WholeF
CarcassF
CarcassM
WholeM
GutMidF
GutMidM
SalivF
SalivM
GutHind
GutAnt
GutCaeca
GutPost
Blood3h
BloodNBF
BloodTwoNBF
FatBodyBF
GutMidBF
Mean 3.43 +/− 1.00

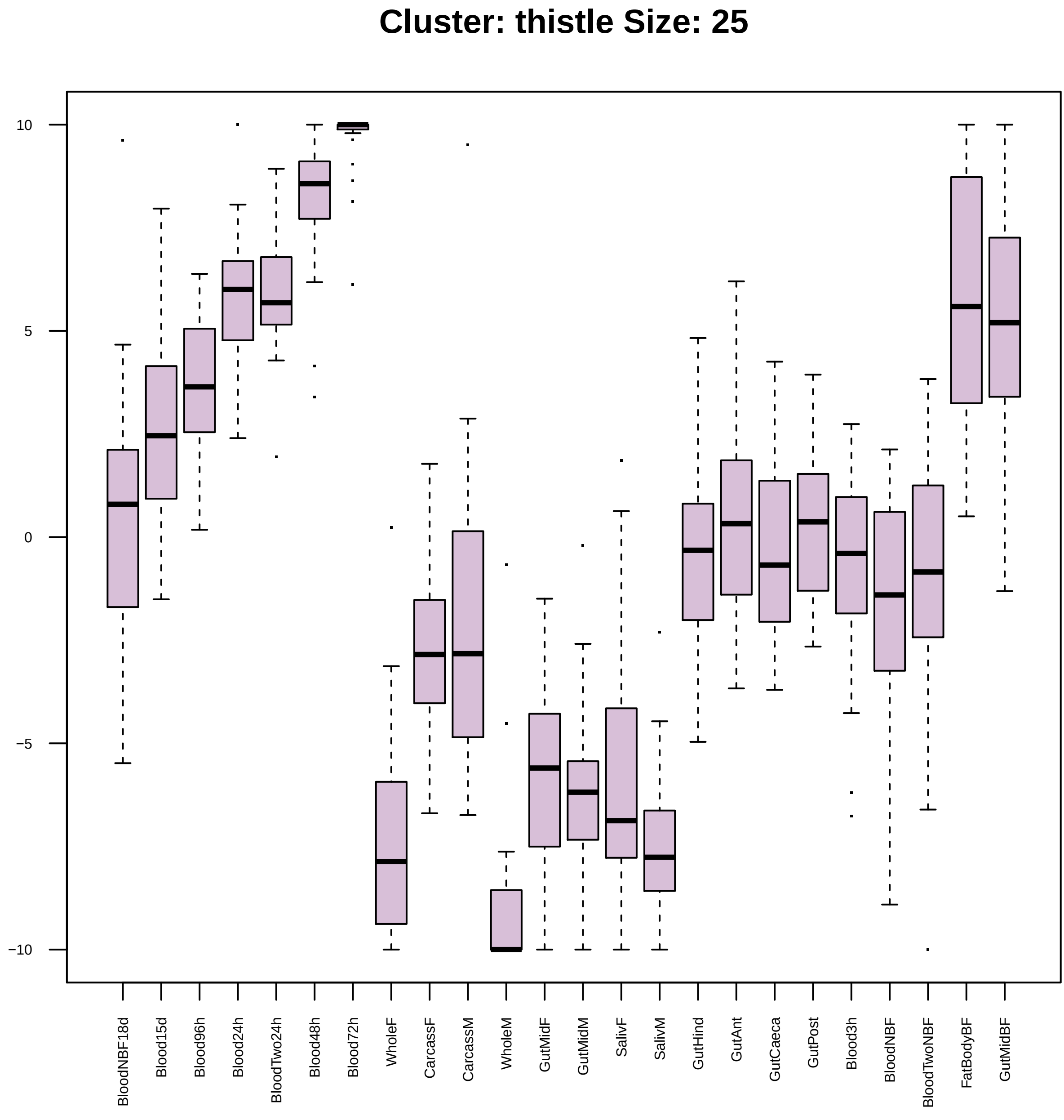

Cluster: thistle Size: 25
10
5
0
−5
−10
BloodNBF18d
Blood15d
Blood96h
Blood24h
BloodTwo24h
Blood48h
Blood72h
WholeF
CarcassF
CarcassM
WholeM
GutMidF
GutMidM
SalivF
SalivM
GutHind
GutAnt
GutCaeca
GutPost
Blood3h
BloodNBF
BloodTwoNBF
FatBodyBF
GutMidBF

| | GO.ID | BPCluster: thistle Size: 25 | Annotated | Significant | Expected | Rank in ClassicF | Weight01F | ClassicF |
|---|---|---|---|---|---|---|---|---|
| *1* | GO:0110110 | positive regulation of animal organ morp... | 12 | 2 | 0.03 | 6 | 0.00043 | 0.00043 |
| *2* | GO:0030833 | regulation of actin filament polymerizat... | 21 | 2 | 0.06 | 13 | 0.00136 | 0.00136 |
| *3* | GO:0008039 | synaptic target recognition | 21 | 2 | 0.06 | 14 | 0.00136 | 0.00136 |
| *4* | GO:0008104 | protein localization | 313 | 3 | 0.83 | 174 | 0.00429 | 0.04777 |
| *5* | GO:0043547 | positive regulation of GTPase activity | 42 | 2 | 0.11 | 48 | 0.00541 | 0.00541 |
| *6* | GO:0007264 | small GTPase mediated signal transductio... | 153 | 3 | 0.41 | 56 | 0.00696 | 0.00719 |
| *7* | GO:0110053 | regulation of actin filament organizatio... | 27 | 3 | 0.07 | 1 | 0.00941 | 4.4e−05 |
| *9* | GO:0044093 | positive regulation of molecular functio... | 79 | 3 | 0.21 | 10 | 0.02122 | 0.00110 |
| *30* | GO:0009266 | response to temperature stimulus | 38 | 2 | 0.10 | 37 | 0.03472 | 0.00444 |

| | GO.ID | MFCluster: thistle Size: 25 | Annotated | Significant | Expected | Rank in ClassicF | Weight01F | ClassicF |
|---|---|---|---|---|---|---|---|---|
| *1* | GO:0008270 | zinc ion binding | 522 | 6 | 1.45 | 1 | 0.0023 | 0.0023 |

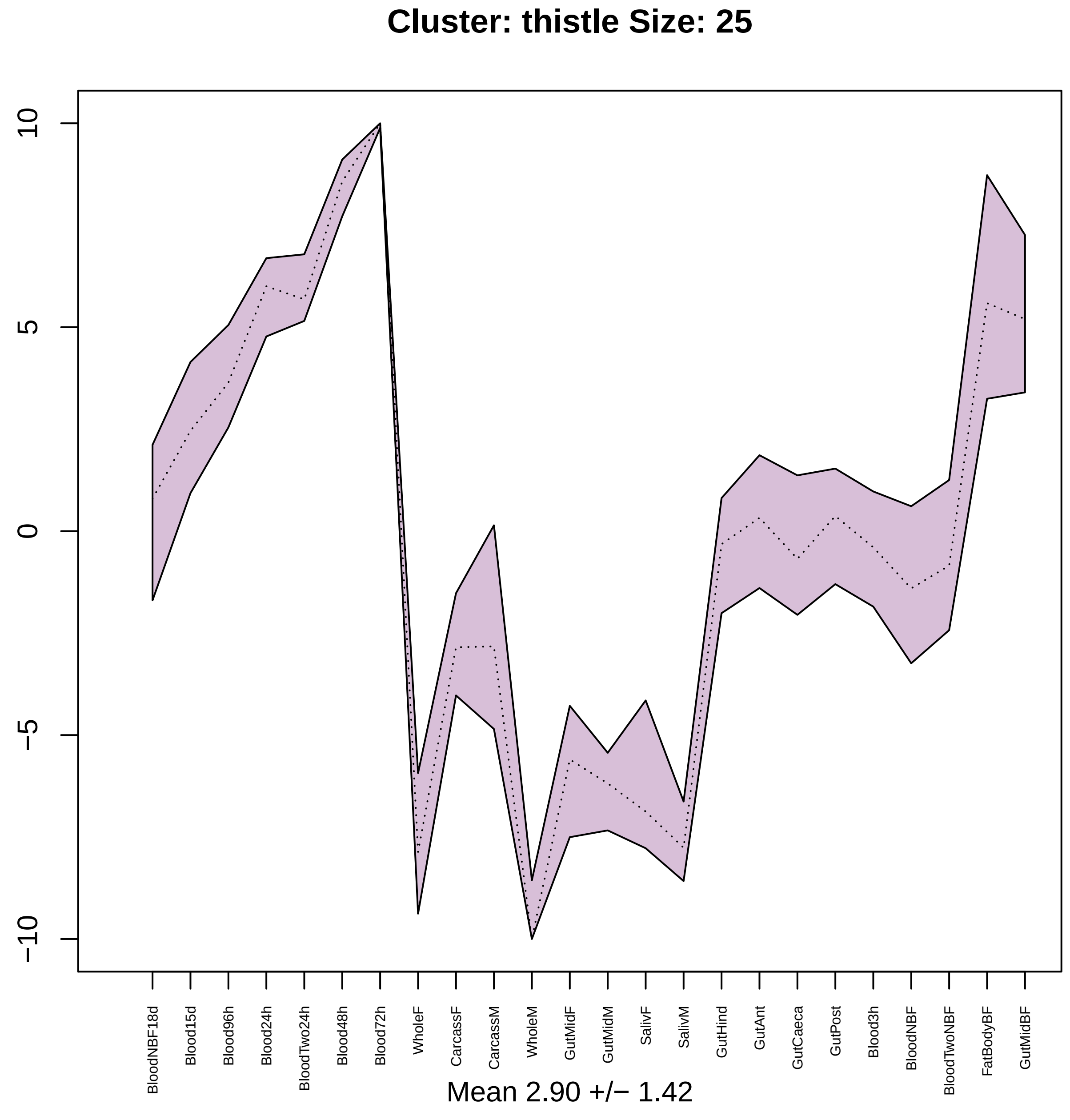

Cluster: thistle Size: 25
10
5
0
−5
−10
BloodNBF18d
Blood15d
Blood96h
Blood24h
BloodTwo24h
Blood48h
Blood72h
WholeF
CarcassF
CarcassM
WholeM
GutMidF
GutMidM
SalivF
SalivM
GutHind
GutAnt
GutCaeca
GutPost
Blood3h
BloodNBF
BloodTwoNBF
FatBodyBF
GutMidBF
Mean 2.90 +/− 1.42

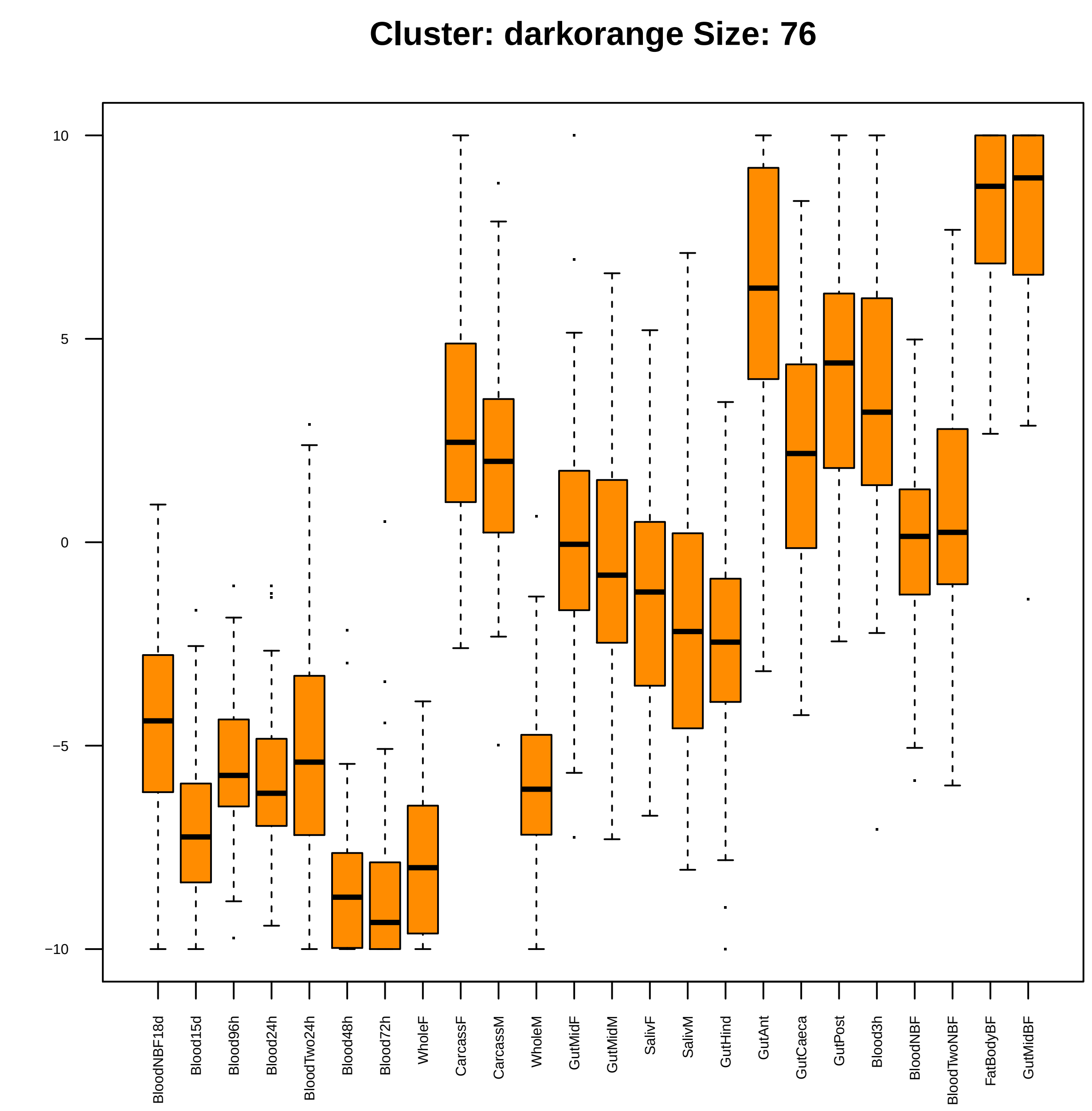

Cluster: darkorange Size: 76
10
5
0
−5
−10
BloodNBF18d
Blood15d
Blood96h
Blood24h
BloodTwo24h
Blood48h
Blood72h
WholeF
CarcassF
CarcassM
WholeM
GutMidF
GutMidM
SalivF
SalivM
GutHind
GutAnt
GutCaeca
GutPost
Blood3h
BloodNBF
BloodTwoNBF
FatBodyBF
GutMidBF

| | GO.ID | BPCluster: darkorange Size: 76 | Annotated | Significant | Expected | Rank in ClassicF | Weight01F | ClassicF |
|---|---|---|---|---|---|---|---|---|
| 1 | GO:0000212 | meiotic spindle organization | 13 | 2 | 0.08 | 4 | 0.0030 | 0.0030 |
| 2 | GO:0007419 | ventral cord development | 15 | 2 | 0.10 | 5 | 0.0040 | 0.0040 |
| 3 | GO:0007112 | male meiosis cytokinesis | 16 | 2 | 0.10 | 6 | 0.0046 | 0.0046 |
| 4 | GO:0003007 | heart morphogenesis | 17 | 2 | 0.11 | 7 | 0.0052 | 0.0052 |

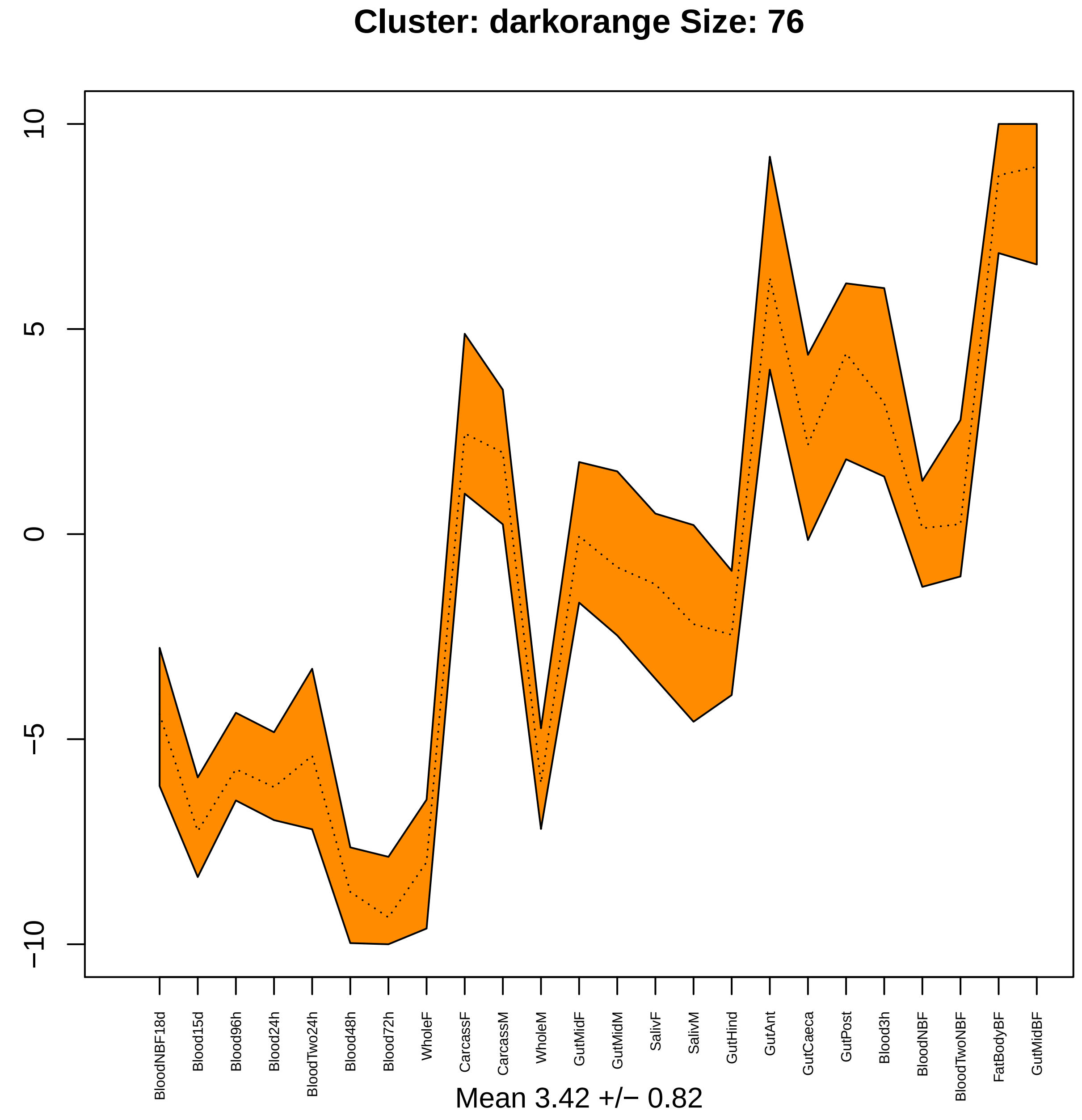

Cluster: darkorange Size: 76
10
5
0
−5
−10
BloodNBF18d
Blood15d
Blood96h
Blood24h
BloodTwo24h
Blood48h
Blood72h
WholeF
CarcassF
CarcassM
WholeM
GutMidF
GutMidM
SalivF
SalivM
GutHind
GutAnt
GutCaeca
GutPost
Blood3h
BloodNBF
BloodTwoNBF
FatBodyBF
GutMidBF
Mean 3.42 +/− 0.82

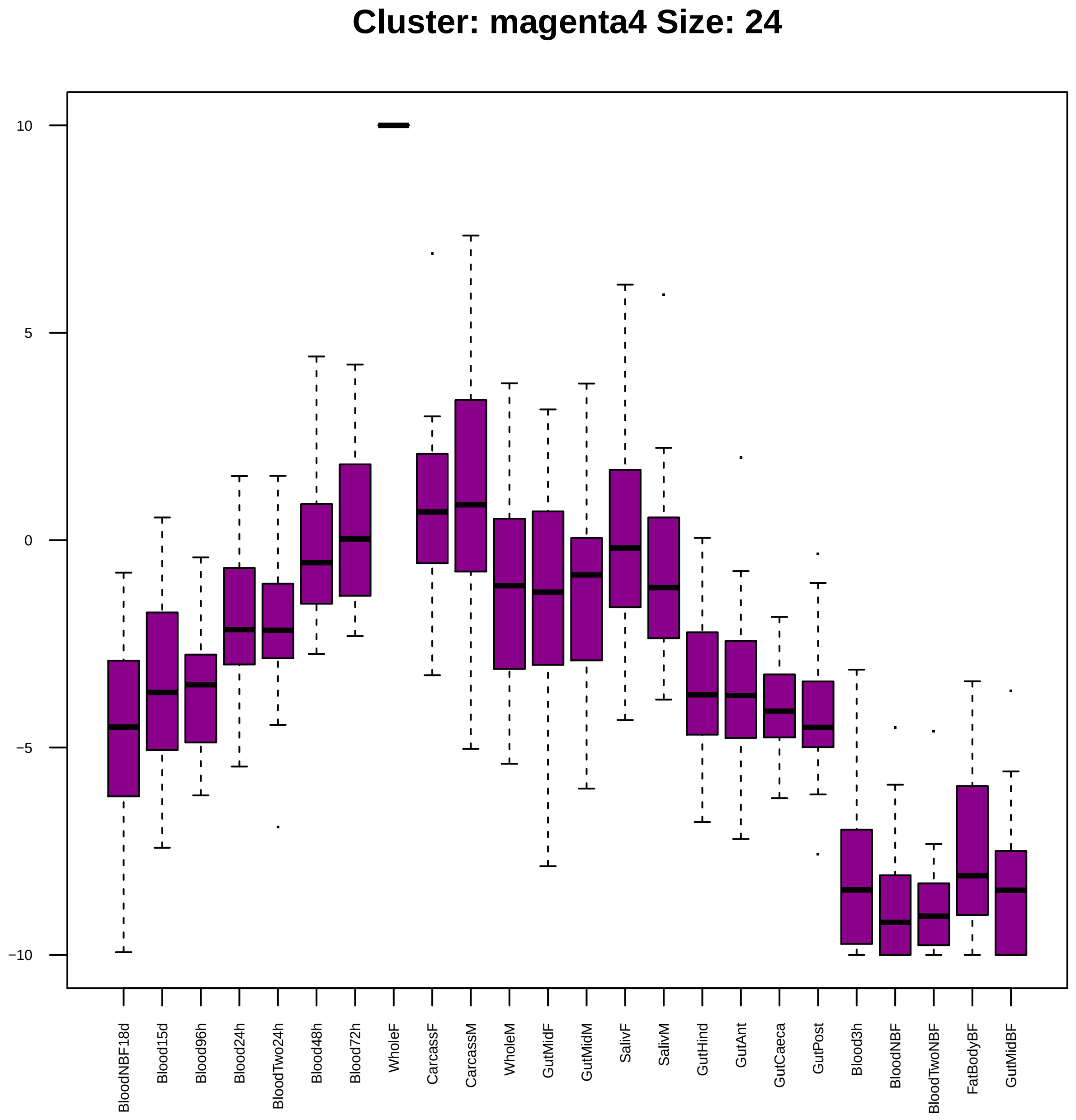

Cluster: magenta4 Size: 24
10
5
0
−5
−10
BloodNBF18d
Blood15d
Blood96h
Blood24h
BloodTwo24h
Blood48h
Blood72h
WholeF
CarcassF
CarcassM
WholeM
GutMidF
GutMidM
SalivF
SalivM
GutHind
GutAnt
GutCaeca
GutPost
Blood3h
BloodNBF
BloodTwoNBF
FatBodyBF
GutMidBF

| | GO.ID | BPCluster: magenta4 Size: 24 | Annotated | Significant | Expected | Rank in ClassicF | Weight01F | ClassicF |
|---|---|---|---|---|---|---|---|---|
| *1* | GO:0032774 | RNA biosynthetic process | 565 | 5 | 1.51 | 23 | 0.0067 | 0.0135 |
| *2* | GO:0019941 | modification−dependent protein catabolic... | 91 | 3 | 0.24 | 3 | 0.0071 | 0.0017 |
| *6* | GO:0031647 | regulation of protein stability | 34 | 2 | 0.09 | 9 | 0.0323 | 0.0036 |
| *14* | GO:0051276 | chromosome organization | 251 | 4 | 0.67 | 10 | 0.0425 | 0.0037 |

| | GO.ID | MFCluster: magenta4 Size: 24 | Annotated | Significant | Expected | Rank in ClassicF | Weight01F | ClassicF |
|---|---|---|---|---|---|---|---|---|
| *1* | GO:0003713 | transcription coactivator activity | 29 | 2 | 0.07 | 2 | 0.0023 | 0.00230 |
| *5* | GO:0003712 | transcription coregulator activity | 44 | 3 | 0.11 | 1 | 0.0329 | 0.00016 |
| *9* | GO:0016779 | nucleotidyltransferase activity | 61 | 2 | 0.15 | 4 | 0.0464 | 0.00988 |

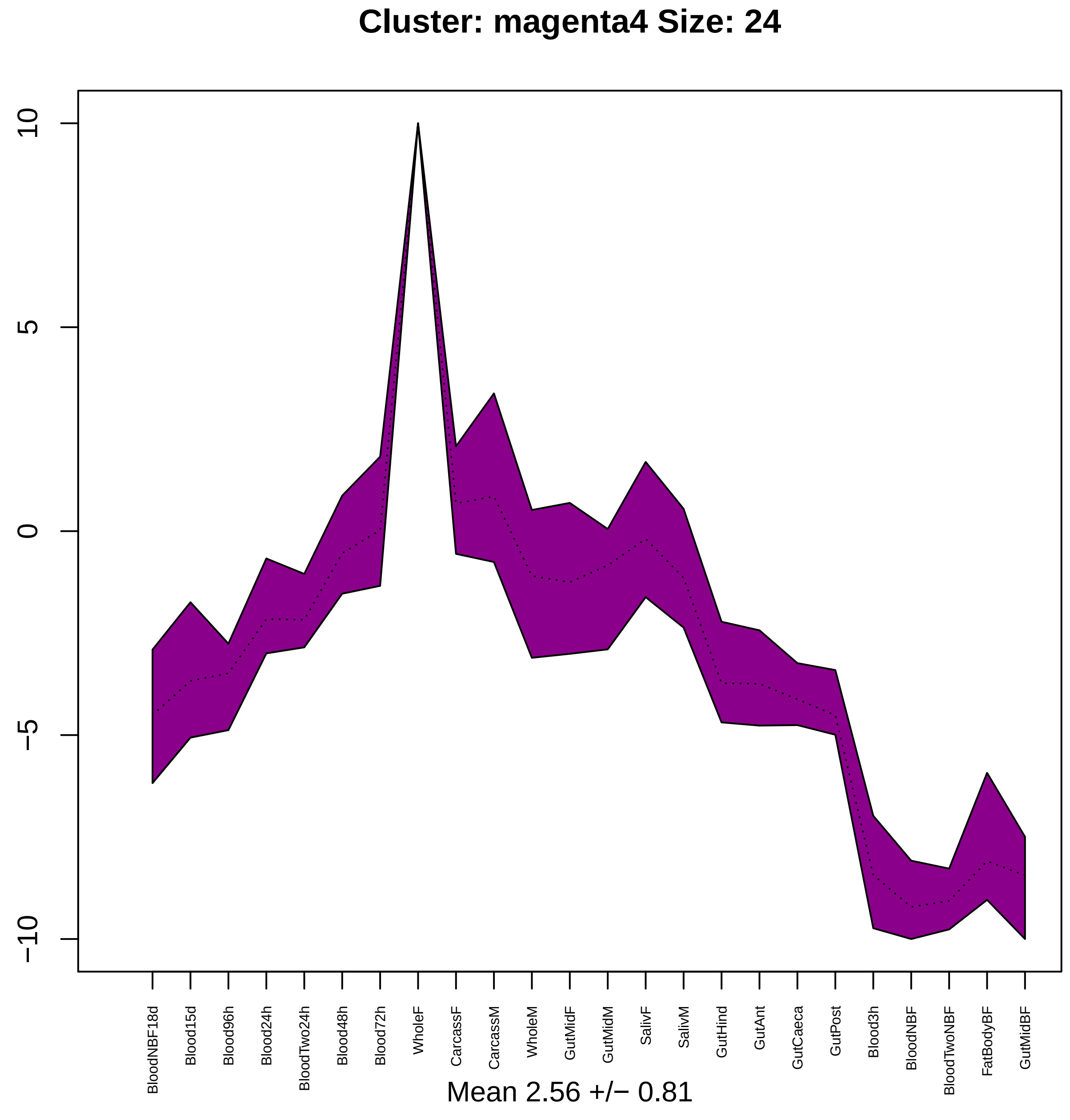

Cluster: magenta4 Size: 24
10
5
0
−5
−10
BloodNBF18d
Blood15d
Blood96h
Blood24h
BloodTwo24h
Blood48h
Blood72h
WholeF
CarcassF
CarcassM
WholeM
GutMidF
GutMidM
SalivF
SalivM
GutHind
GutAnt
GutCaeca
GutPost
Blood3h
BloodNBF
BloodTwoNBF
FatBodyBF
GutMidBF
Mean 2.56 +/− 0.81

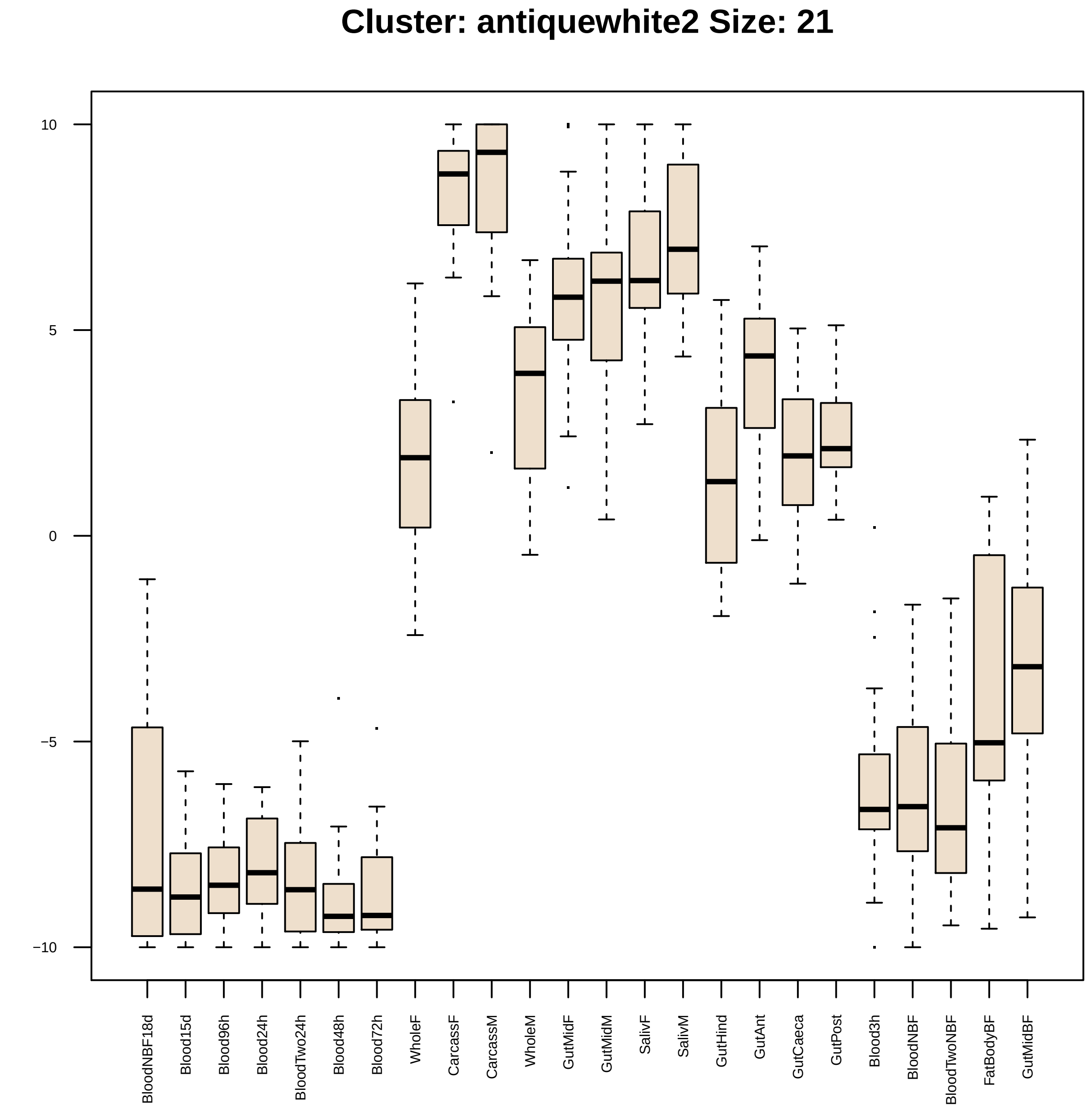
Cluster: antiquewhite2 Size: 21
10
5
0
−5
−10
BloodNBF18d
Blood15d
Blood96h
Blood24h
BloodTwo24h
Blood48h
Blood72h
WholeF
CarcassF
CarcassM
WholeM
GutMidF
GutMidM
SalivF
SalivM
GutHind
GutAnt
GutCaeca
GutPost
Blood3h
BloodNBF
BloodTwoNBF
FatBodyBF
GutMidBF

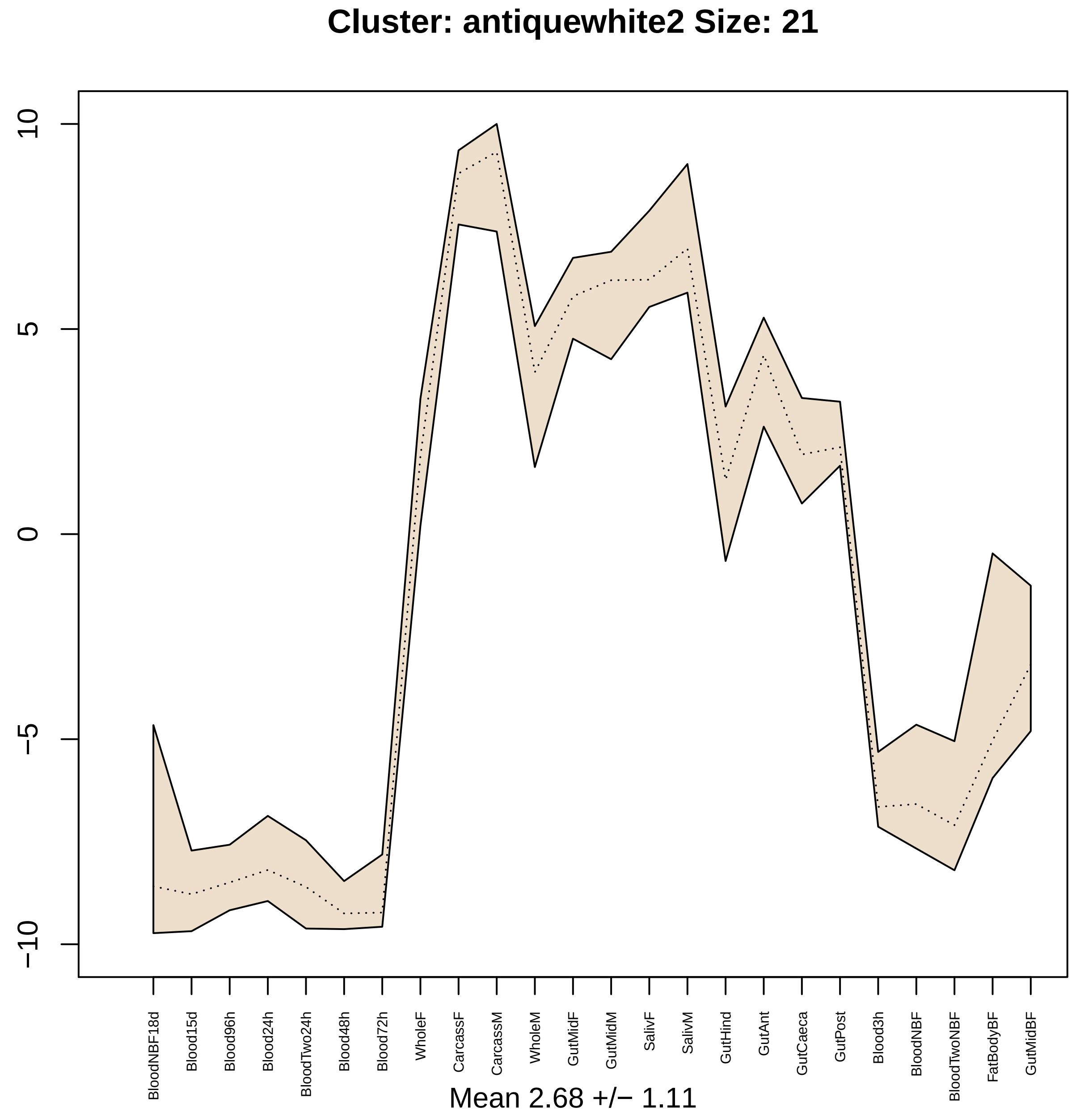

Cluster: antiquewhite2 Size: 21
10
5
0
−5
−10
BloodNBF18d
Blood15d
Blood96h
Blood24h
BloodTwo24h
Blood48h
Blood72h
WholeF
CarcassF
CarcassM
WholeM
GutMidF
GutMidM
SalivF
SalivM
GutHind
GutAnt
GutCaeca
GutPost
Blood3h
BloodNBF
BloodTwoNBF
FatBodyBF
GutMidBF
Mean 2.68 +/− 1.11

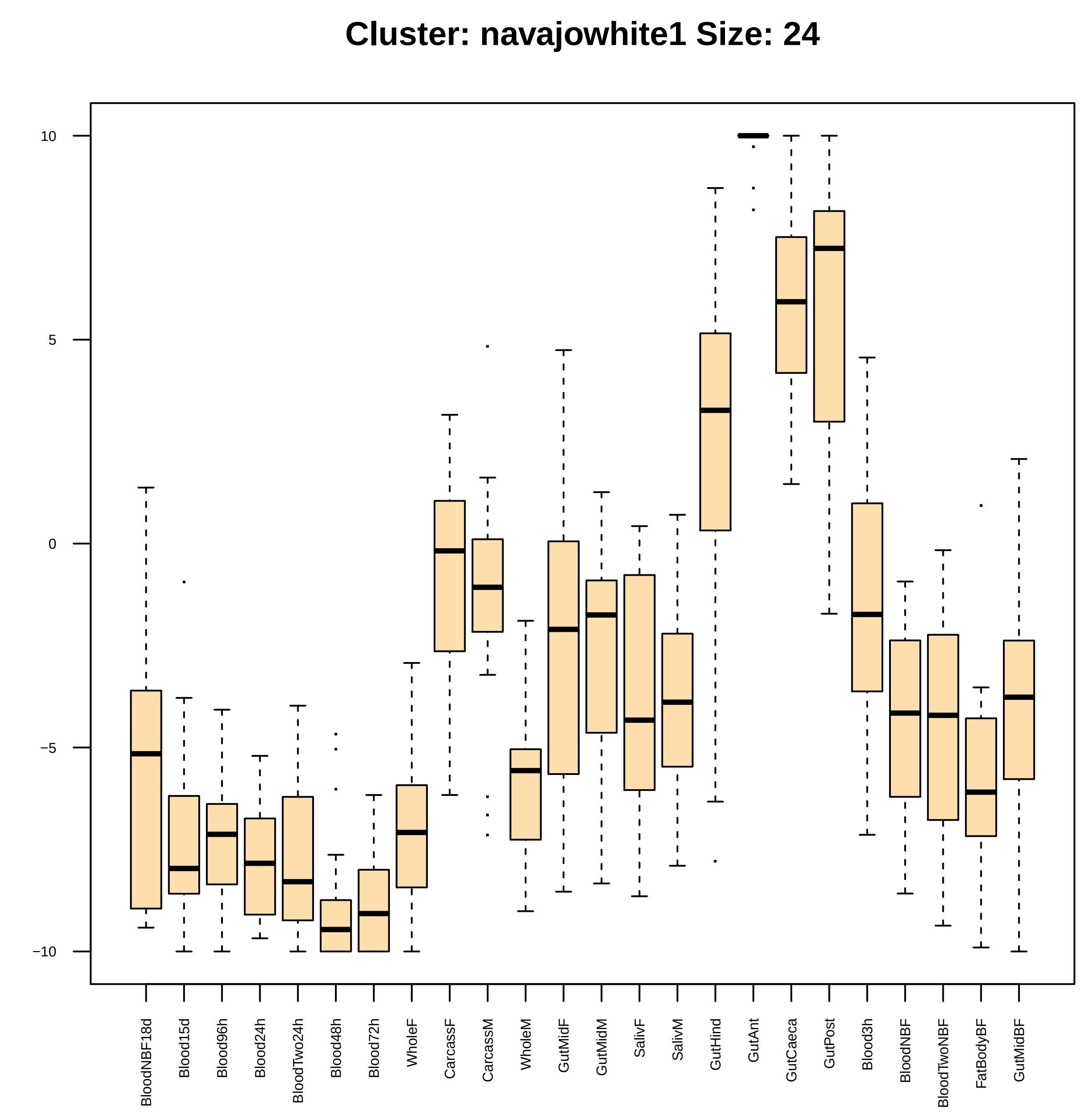

Cluster: navajowhite1 Size: 24
10
5
0
−5
−10
BloodNBF18d
Blood15d
Blood96h
Blood24h
BloodTwo24h
Blood48h
Blood72h
WholeF
CarcassF
CarcassM
WholeM
GutMidF
GutMidM
SalivF
SalivM
GutHind
GutAnt
GutCaeca
GutPost
Blood3h
BloodNBF
BloodTwoNBF
FatBodyBF
GutMidBF

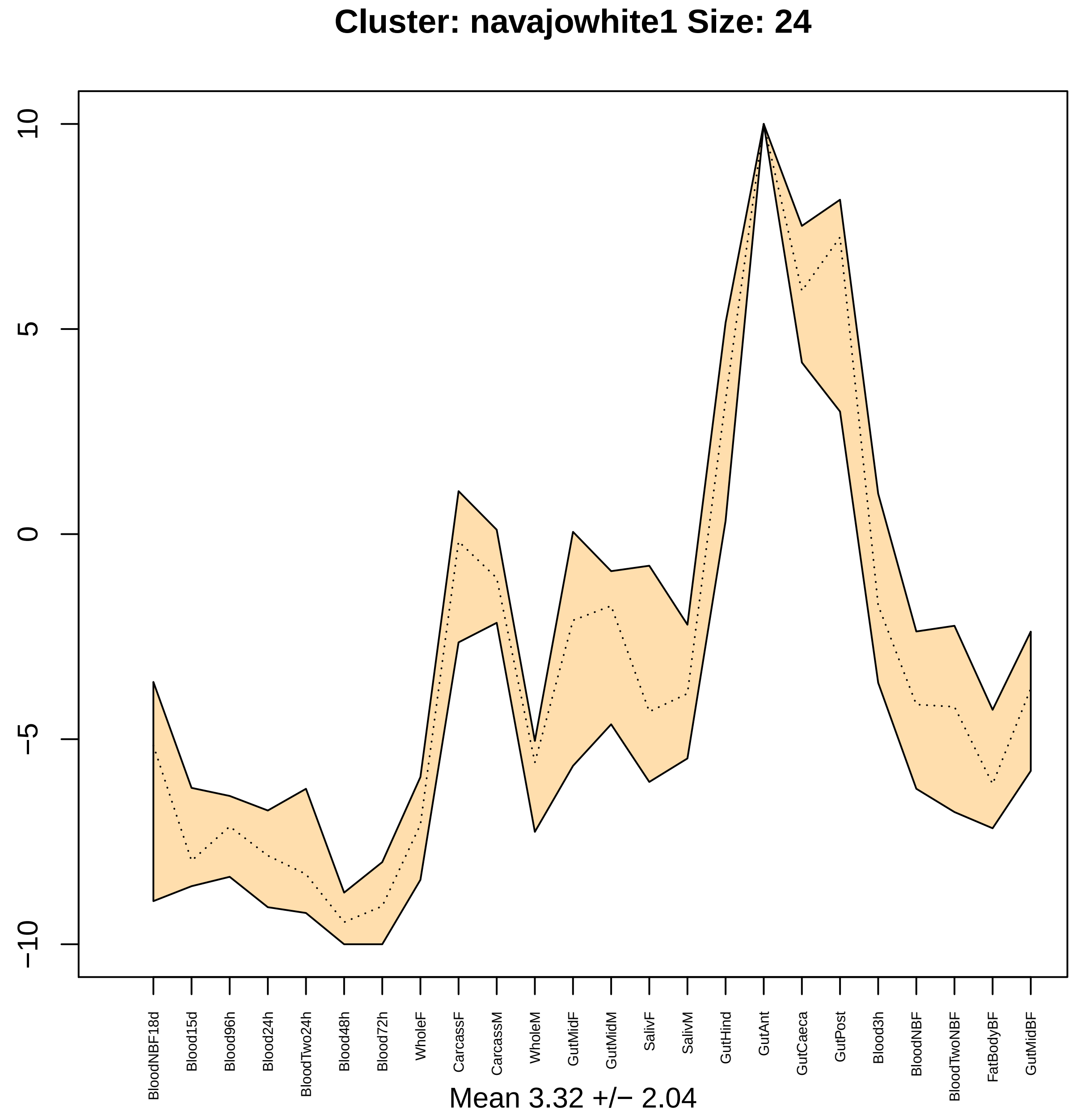

Cluster: navajowhite1 Size: 24
10
5
0
−5
−10
BloodNBF18d
Blood15d
Blood96h
Blood24h
BloodTwo24h
Blood48h
Blood72h
WholeF
CarcassF
CarcassM
WholeM
GutMidF
GutMidM
SalivF
SalivM
GutHind
GutAnt
GutCaeca
GutPost
Blood3h
BloodNBF
BloodTwoNBF
FatBodyBF
GutMidBF
Mean 3.32 +/− 2.04

### Cluster: darkolivegreen2 Size: 18

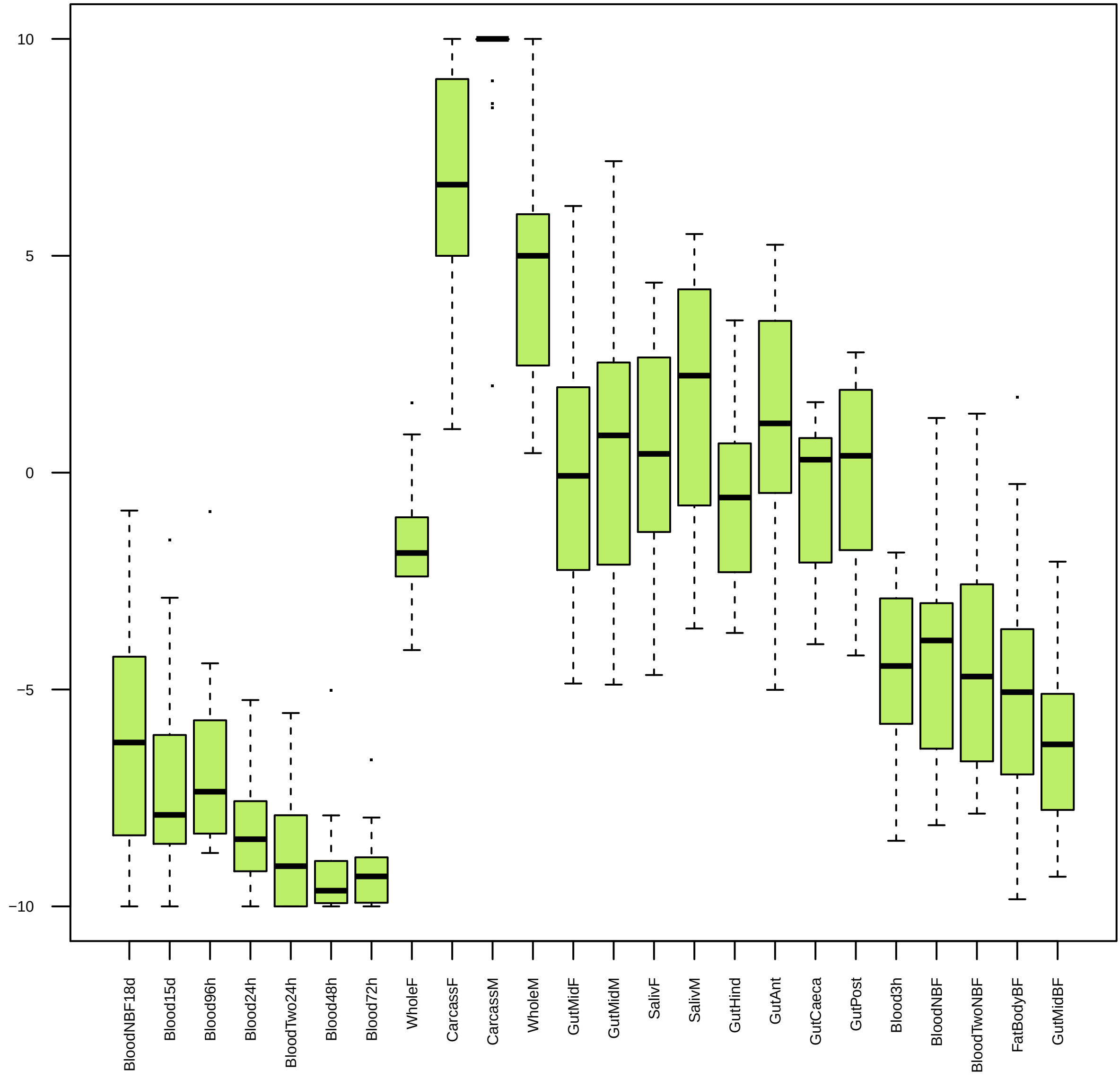

| | GO.ID | MFCluster: darkolivegreen2 Size: 18 | Annotated | Significant | Expected | Rank in ClassicF | Weight01F | ClassicF |
|---|---|---|---|---|---|---|---|---|
| 5 | GO:0008238 | exopeptidase activity | 66 | 2 | 0.09 | 1 | 0.051 | 0.0035 |

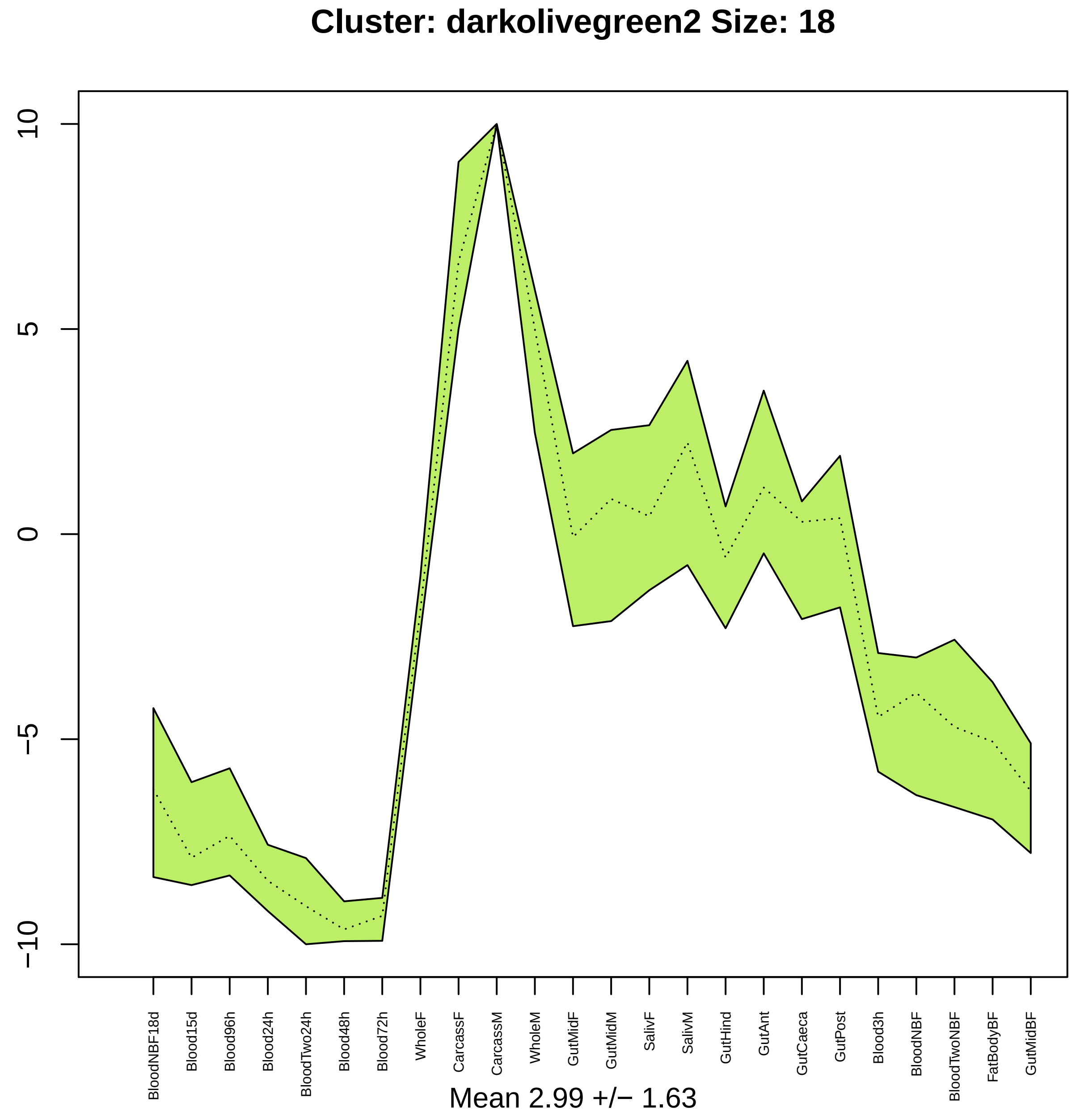

Cluster: darkolivegreen2 Size: 18
10
5
0
−5
−10
BloodNBF18d
Blood15d
Blood96h
Blood24h
BloodTwo24h
Blood48h
Blood72h
WholeF
CarcassF
CarcassM
WholeM
GutMidF
GutMidM
SalivF
SalivM
GutHind
GutAnt
GutCaeca
GutPost
Blood3h
BloodNBF
BloodTwoNBF
FatBodyBF
GutMidBF
Mean 2.99 +/− 1.63

### Cluster: sienna4 Size: 20

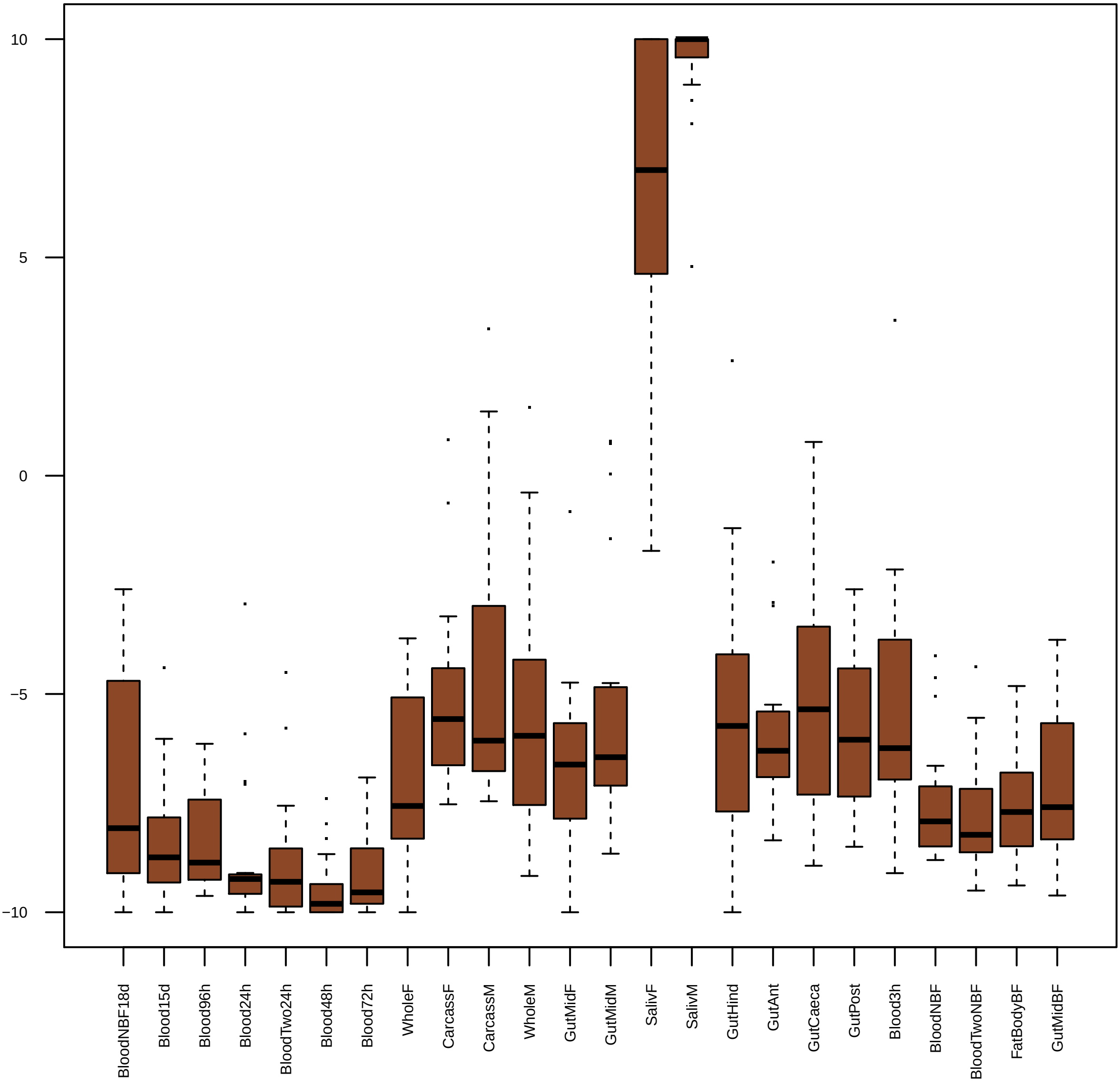

|  | GO.ID | BPCluster: sienna4 Size: 20 | Annotated | Significant | Expected | Rank in ClassicF | Weight01F | ClassicF |
|---|---|---|---|---|---|---|---|---|
| *1* | GO:0006508 | proteolysis | 594 | 6 | 1.3 | 1 | 0.001 | 0.001 |

| | GO.ID | MFCluster: sienna4 Size: 20 | Annotated | Significant | Expected | Rank in ClassicF | Weight01F | ClassicF |
|---|---|---|---|---|---|---|---|---|
| *17* | GO:0070011 | peptidase activity, acting on L−amino ac... | 470 | 6 | 0.91 | 5 | 1.000 | 0.00014 |
| *21* | GO:0016787 | hydrolase activity | 1445 | 8 | 2.81 | 7 | 1.000 | 0.00241 |
| *24* | GO:0008233 | peptidase activity | 500 | 6 | 0.97 | 6 | 1.000 | 0.00020 |

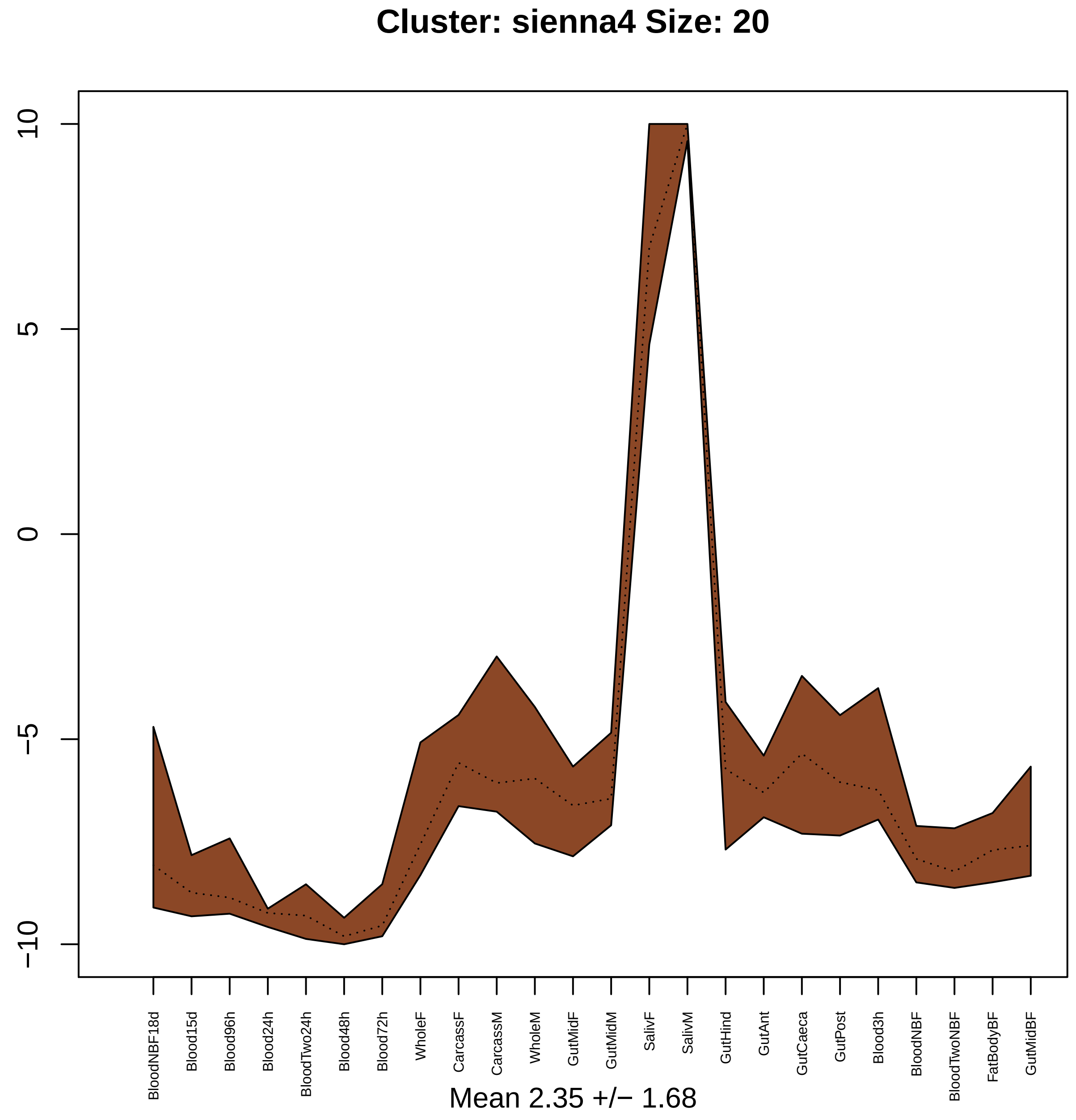

Cluster: sienna4 Size: 20
10
5
0
−5
−10
BloodNBF18d
Blood15d
Blood96h
Blood24h
BloodTwo24h
Blood48h
Blood72h
WholeF
CarcassF
CarcassM
WholeM
GutMidF
GutMidM
SalivF
SalivM
GutHind
GutAnt
GutCaeca
GutPost
Blood3h
BloodNBF
BloodTwoNBF
FatBodyBF
GutMidBF
Mean 2.35 +/− 1.68

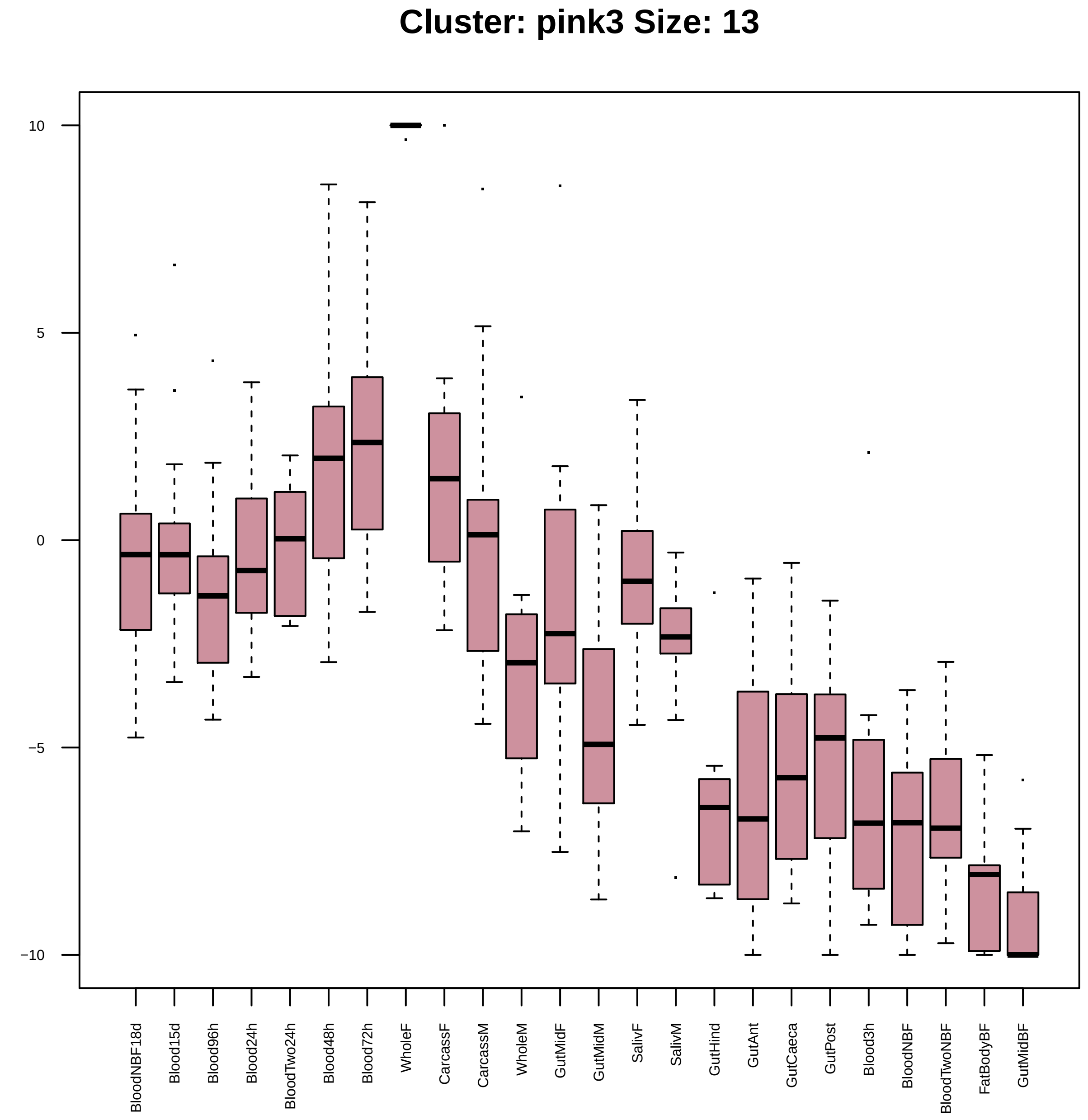
Cluster: pink3 Size: 13
10
5
0
−5
−10
BloodNBF18d
Blood15d
Blood96h
Blood24h
BloodTwo24h
Blood48h
Blood72h
WholeF
CarcassF
CarcassM
WholeM
GutMidF
GutMidM
SalivF
SalivM
GutHind
GutAnt
GutCaeca
GutPost
Blood3h
BloodNBF
BloodTwoNBF
FatBodyBF
GutMidBF

| | GO.ID | BPCluster: pink3 Size: 13 | Annotated | Significant | Expected | Rank in ClassicF | Weight01F | ClassicF |
|---|---|---|---|---|---|---|---|---|
| *1* | GO:0000398 | mRNA splicing, via spliceosome | 134 | 3 | 0.21 | 3 | 0.0078 | 0.00098 |
| *8* | GO:0008033 | tRNA processing | 39 | 2 | 0.06 | 6 | 0.0266 | 0.00159 |
| *26* | GO:0006399 | tRNA metabolic process | 82 | 2 | 0.13 | 11 | 1.0000 | 0.00687 |

| | GO.ID | MFCluster: pink3 Size: 13 | Annotated | Significant | Expected | Rank in ClassicF | Weight01F | ClassicF |
|---|---|---|---|---|---|---|---|---|
| 1 | GO:0140101 | catalytic activity, acting on a tRNA | 66 | 2 | 0.09 | 1 | 0.0035 | 0.0035 |

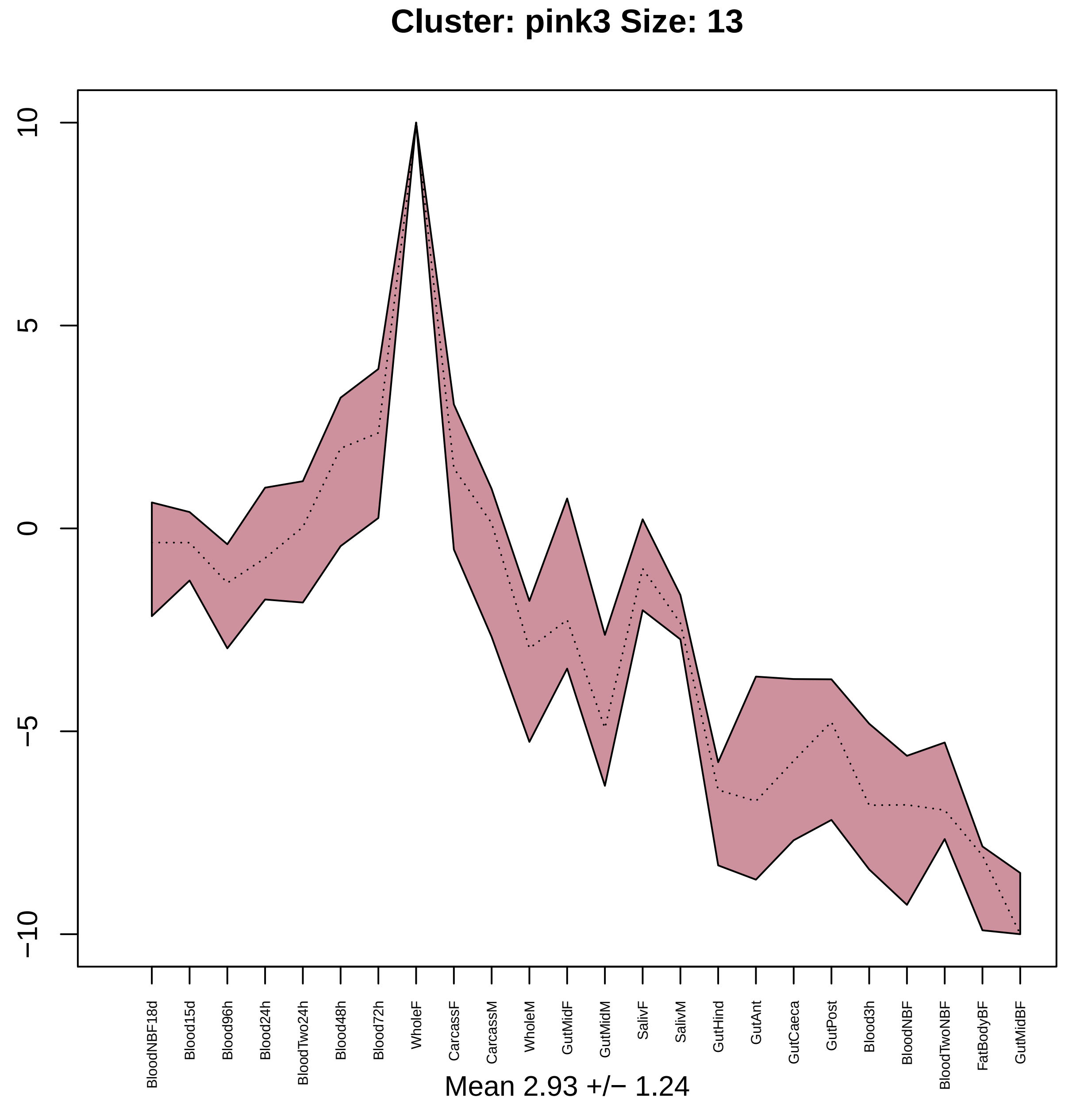

Cluster: pink3 Size: 13
10
5
0
−5
−10
BloodNBF18d
Blood15d
Blood96h
Blood24h
BloodTwo24h
Blood48h
Blood72h
WholeF
CarcassF
CarcassM
WholeM
GutMidF
GutMidM
SalivF
SalivM
GutHind
GutAnt
GutCaeca
GutPost
Blood3h
BloodNBF
BloodTwoNBF
FatBodyBF
GutMidBF
Mean 2.93 +/− 1.24

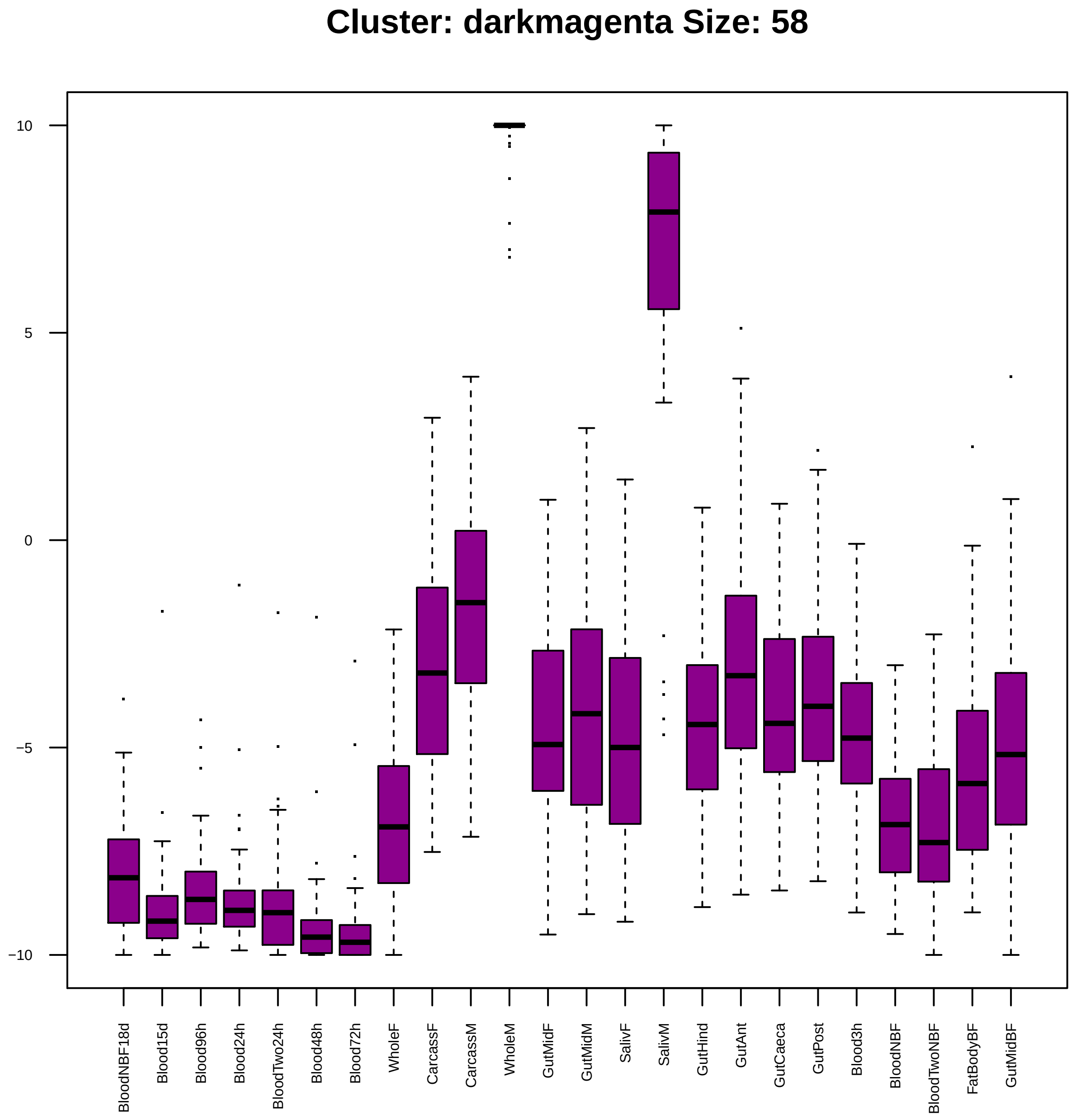
Cluster: darkmagenta Size: 58
10
5
0
−5
−10
BloodNBF18d
Blood15d
Blood96h
Blood24h
BloodTwo24h
Blood48h
Blood72h
WholeF
CarcassF
CarcassM
WholeM
GutMidF
GutMidM
SalivF
SalivM
GutHind
GutAnt
GutCaeca
GutPost
Blood3h
BloodNBF
BloodTwoNBF
FatBodyBF
GutMidBF

| | GO.ID | BPCluster: darkmagenta Size: 58 | Annotated | Significant | Expected | Rank in ClassicF | Weight01F | ClassicF |
|---|---|---|---|---|---|---|---|---|
| *1* | GO:0007017 | microtubule−based process | 282 | 7 | 0.93 | 1 | 0.0011 | 2.1e−05 |
| *2* | GO:0007018 | microtubule−based movement | 65 | 3 | 0.21 | 2 | 0.0012 | 0.0012 |
| *3* | GO:0051258 | protein polymerization | 39 | 2 | 0.13 | 3 | 0.0071 | 0.0071 |

| | GO.ID | MFCluster: darkmagenta Size: 58 | Annotated | Significant | Expected | Rank in ClassicF | Weight01F | ClassicF |
|---|---|---|---|---|---|---|---|---|
| 1 | GO:0030145 | manganese ion binding | 12 | 2 | 0.04 | 2 | 0.00069 | 0.00069 |
| 2 | GO:0005200 | structural constituent of cytoskeleton | 19 | 2 | 0.06 | 4 | 0.00175 | 0.00175 |
| 3 | GO:0004177 | aminopeptidase activity | 25 | 2 | 0.08 | 5 | 0.00304 | 0.00304 |
| 4 | GO:0008235 | metalloexopeptidase activity | 29 | 2 | 0.10 | 6 | 0.00408 | 0.00408 |
| 5 | GO:0016887 | ATPase activity | 233 | 3 | 0.78 | 31 | 0.00743 | 0.04091 |
| 6 | GO:0003777 | microtubule motor activity | 40 | 2 | 0.13 | 9 | 0.00767 | 0.00767 |
| 12 | GO:0003774 | motor activity | 60 | 3 | 0.20 | 3 | 0.05970 | 0.00098 |

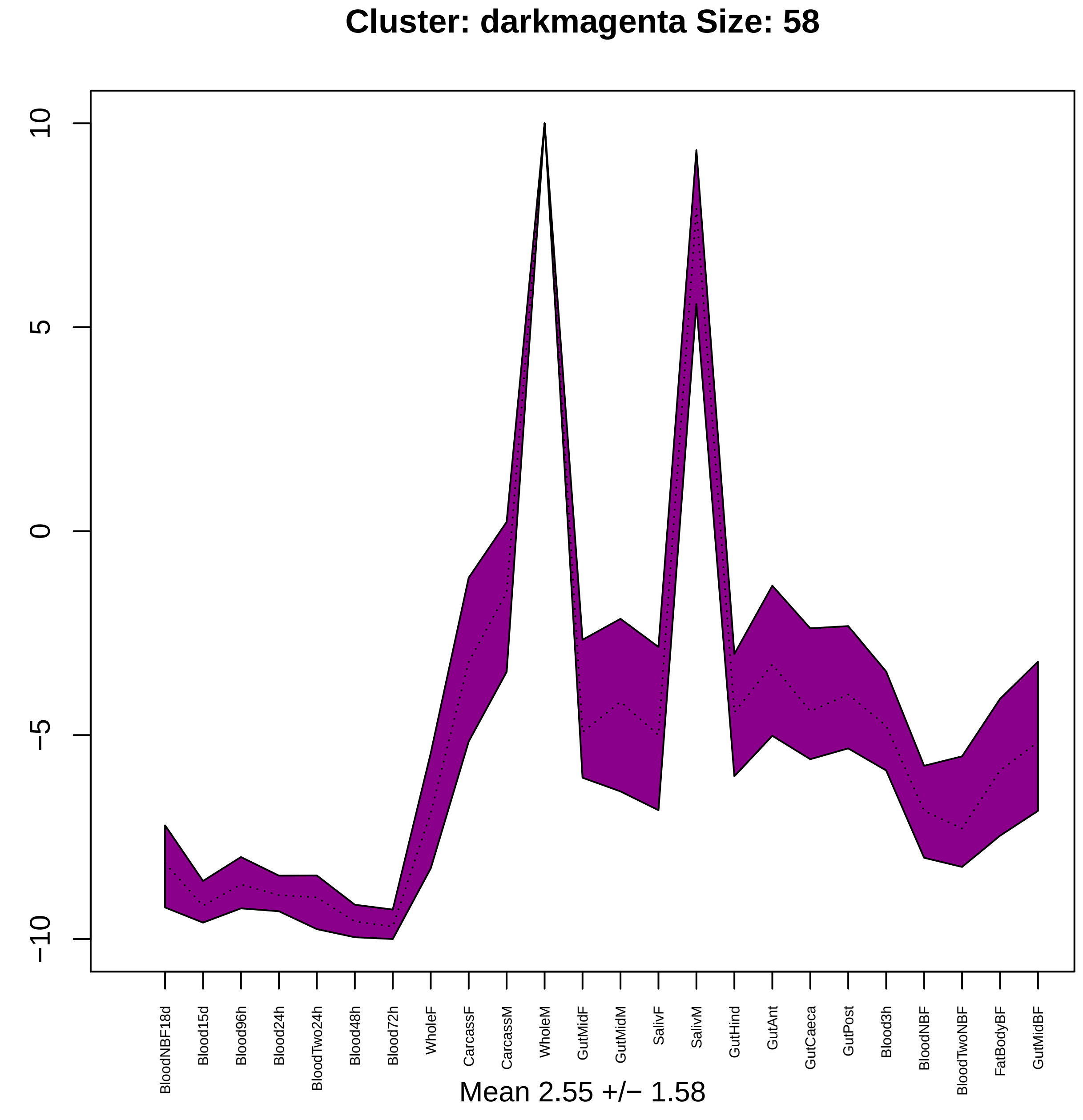
Cluster: darkmagenta Size: 58
10
5
0
−5
−10
BloodNBF18d
Blood15d
Blood96h
Blood24h
BloodTwo24h
Blood48h
Blood72h
WholeF
CarcassF
CarcassM
WholeM
GutMidF
GutMidM
SalivF
SalivM
GutHind
GutAnt
GutCaeca
GutPost
Blood3h
BloodNBF
BloodTwoNBF
FatBodyBF
GutMidBF
Mean 2.55 +/− 1.58

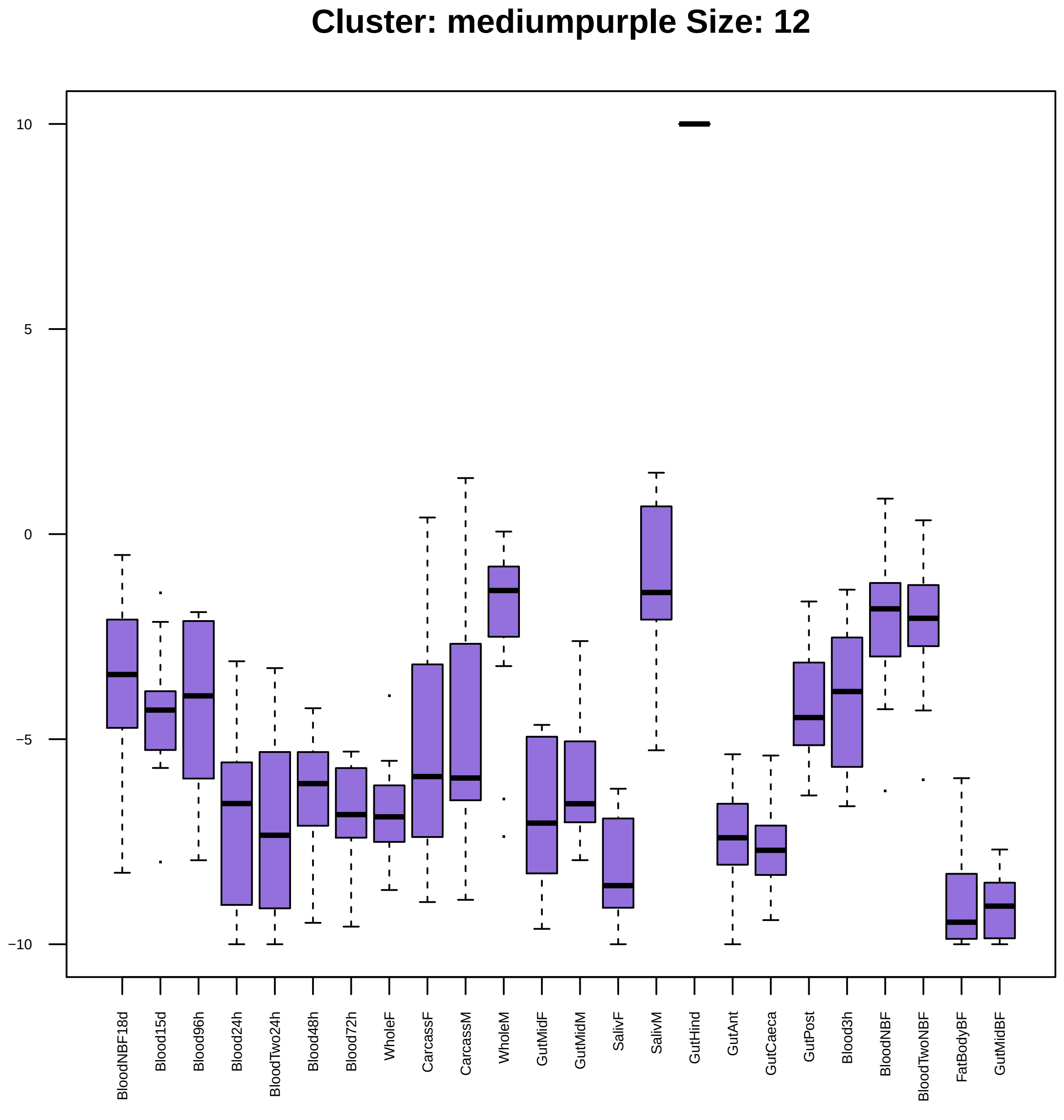
Cluster: mediumpurple Size: 12
10
5
0
−5
−10
BloodNBF18d
Blood15d
Blood96h
Blood24h
BloodTwo24h
Blood48h
Blood72h
WholeF
CarcassF
CarcassM
WholeM
GutMidF
GutMidM
SalivF
SalivM
GutHind
GutAnt
GutCaeca
GutPost
Blood3h
BloodNBF
BloodTwoNBF
FatBodyBF
GutMidBF

| | GO.ID | MFCluster: mediumpurple Size: 12 | Annotated | Significant | Expected | Rank in ClassicF | Weight01F | ClassicF |
|---|---|---|---|---|---|---|---|---|
| *2* | GO:0022857 | transmembrane transporter activity | 442 | 4 | 0.61 | 5 | 0.0031 | 0.00218 |
| *5* | GO:0015370 | solute:sodium symporter activity | 31 | 2 | 0.04 | 2 | 0.0211 | 0.00079 |
| *17* | GO:0015291 | secondary active transmembrane transport... | 60 | 2 | 0.08 | 6 | 1.0000 | 0.00294 |
| *20* | GO:0015293 | symporter activity | 37 | 2 | 0.05 | 3 | 1.0000 | 0.00112 |
| *21* | GO:0015294 | solute:cation symporter activity | 37 | 2 | 0.05 | 4 | 1.0000 | 0.00112 |

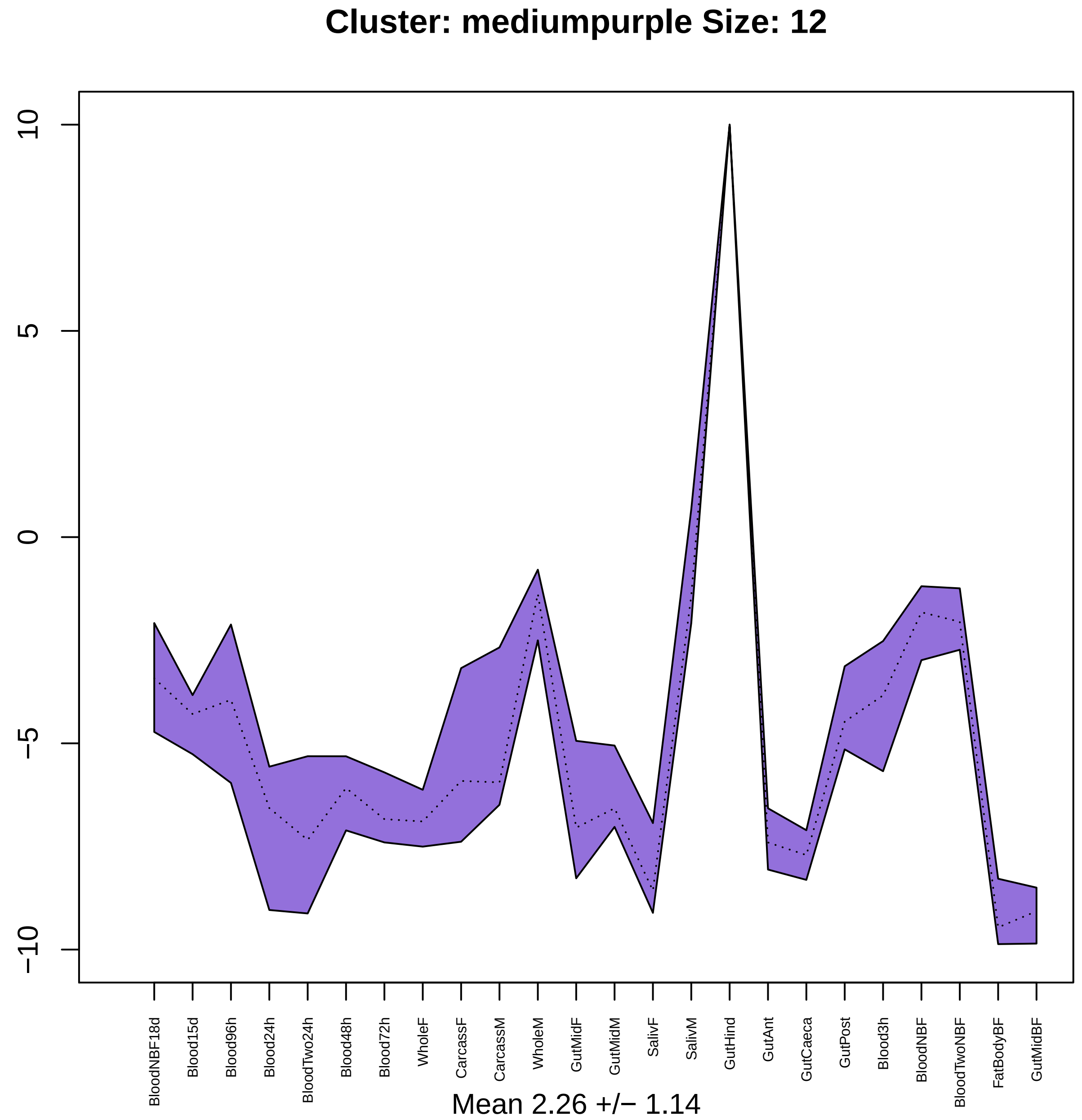

Cluster: mediumpurple Size: 12
10
5
0
−5
−10
BloodNBF18d
Blood15d
Blood96h
Blood24h
BloodTwo24h
Blood48h
Blood72h
WholeF
CarcassF
CarcassM
WholeM
GutMidF
GutMidM
SalivF
SalivM
GutHind
GutAnt
GutCaeca
GutPost
Blood3h
BloodNBF
BloodTwoNBF
FatBodyBF
GutMidBF
Mean 2.26 +/− 1.14

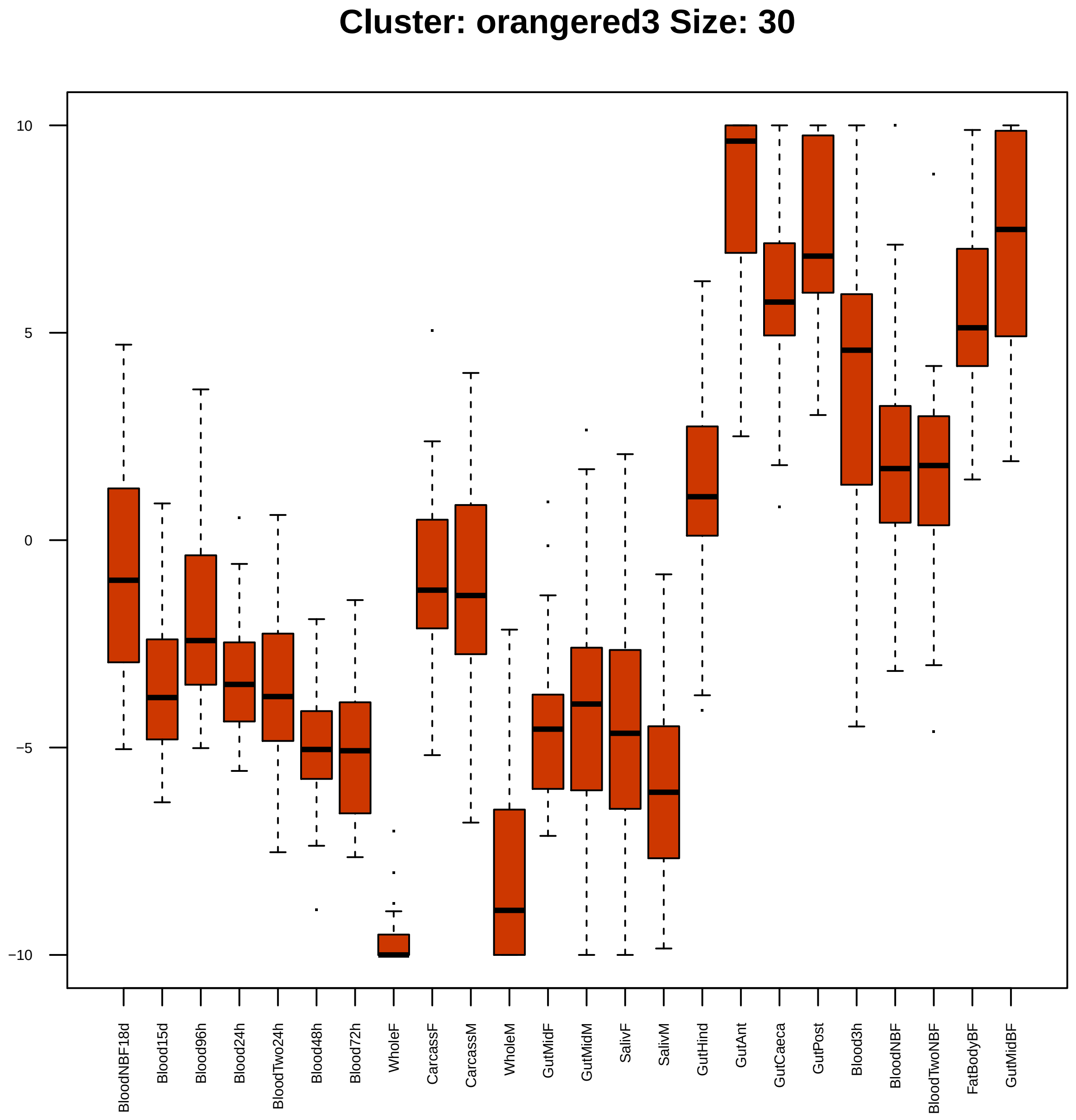

Cluster: orangered3 Size: 30
10
5
0
−5
−10
BloodNBF18d
Blood15d
Blood96h
Blood24h
BloodTwo24h
Blood48h
Blood72h
WholeF
CarcassF
CarcassM
WholeM
GutMidF
GutMidM
SalivF
SalivM
GutHind
GutAnt
GutCaeca
GutPost
Blood3h
BloodNBF
BloodTwoNBF
FatBodyBF
GutMidBF

| | GO.ID | BPCluster: orangered3 Size: 30 | Annotated | Significant | Expected | Rank in ClassicF | Weight01F | ClassicF |
|---|---|---|---|---|---|---|---|---|
| 1 | GO:0007469 | antennal development | 10 | 2 | 0.03 | 3 | 0.00037 | 0.00037 |
| 2 | GO:0009954 | proximal/distal pattern formation | 11 | 2 | 0.03 | 5 | 0.00046 | 0.00046 |
| 3 | GO:0007455 | eye−antennal disc morphogenesis | 17 | 2 | 0.05 | 8 | 0.00111 | 0.00111 |
| 4 | GO:0001708 | cell fate specification | 38 | 3 | 0.11 | 2 | 0.00169 | 0.00018 |
| 5 | GO:0007427 | epithelial cell migration, open tracheal... | 27 | 2 | 0.08 | 21 | 0.00282 | 0.00282 |
| 6 | GO:0008586 | imaginal disc−derived wing vein morphoge... | 28 | 2 | 0.08 | 22 | 0.00304 | 0.00304 |
| 7 | GO:0007548 | sex differentiation | 40 | 2 | 0.12 | 41 | 0.00849 | 0.00613 |
| 8 | GO:0007447 | imaginal disc pattern formation | 47 | 2 | 0.14 | 51 | 0.01413 | 0.00839 |
| 10 | GO:0035218 | leg disc development | 43 | 2 | 0.13 | 45 | 0.01971 | 0.00706 |
| 12 | GO:0045466 | R7 cell differentiation | 29 | 2 | 0.09 | 23 | 0.02521 | 0.00326 |
| 22 | GO:0045595 | regulation of cell differentiation | 112 | 3 | 0.33 | 31 | 0.03467 | 0.00416 |

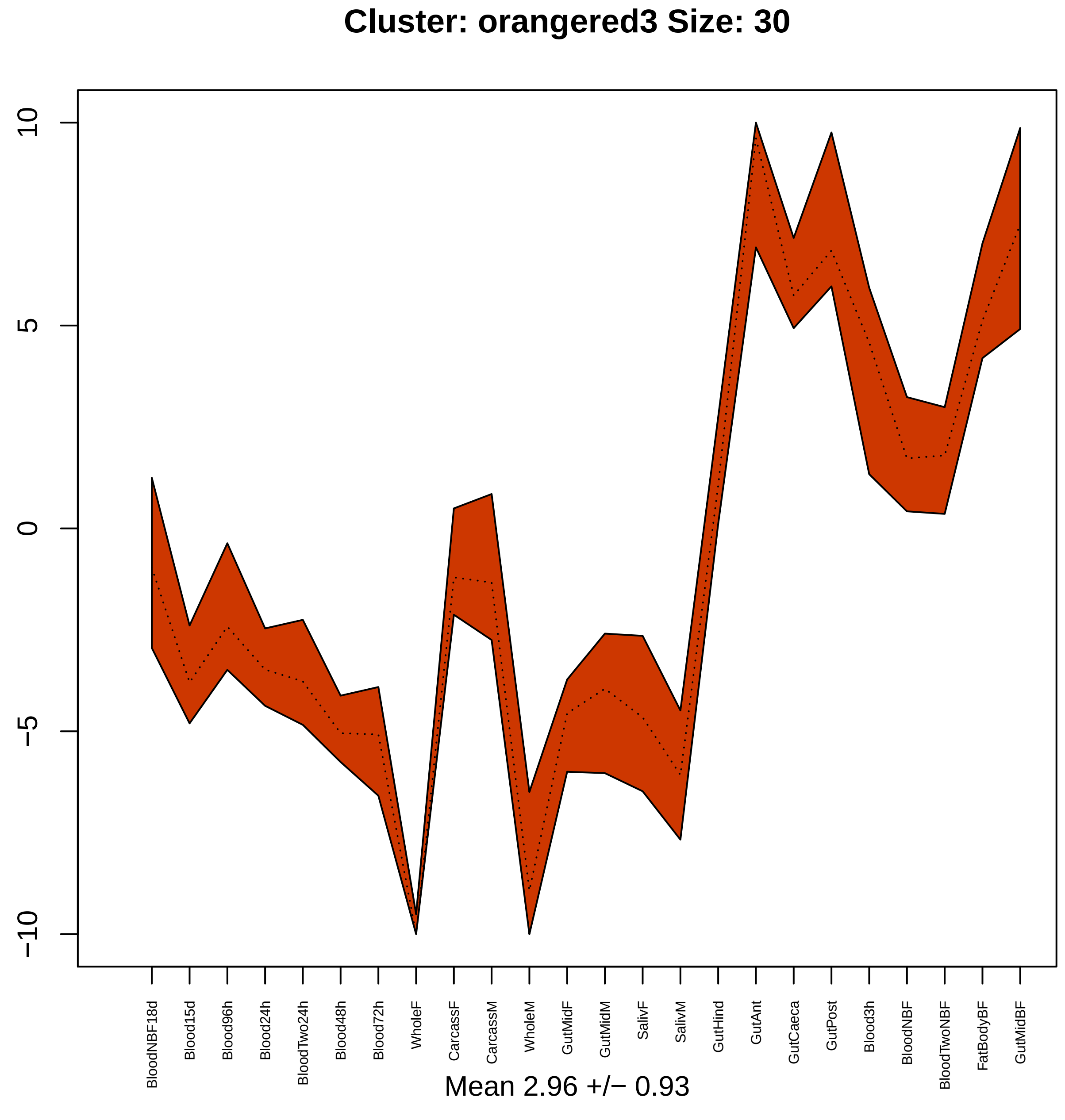

Cluster: orangered3 Size: 30
10
5
0
−5
−10
BloodNBF18d
Blood15d
Blood96h
Blood24h
BloodTwo24h
Blood48h
Blood72h
WholeF
CarcassF
CarcassM
WholeM
GutMidF
GutMidM
SalivF
SalivM
GutHind
GutAnt
GutCaeca
GutPost
Blood3h
BloodNBF
BloodTwoNBF
FatBodyBF
GutMidBF
Mean 2.96 +/− 0.93

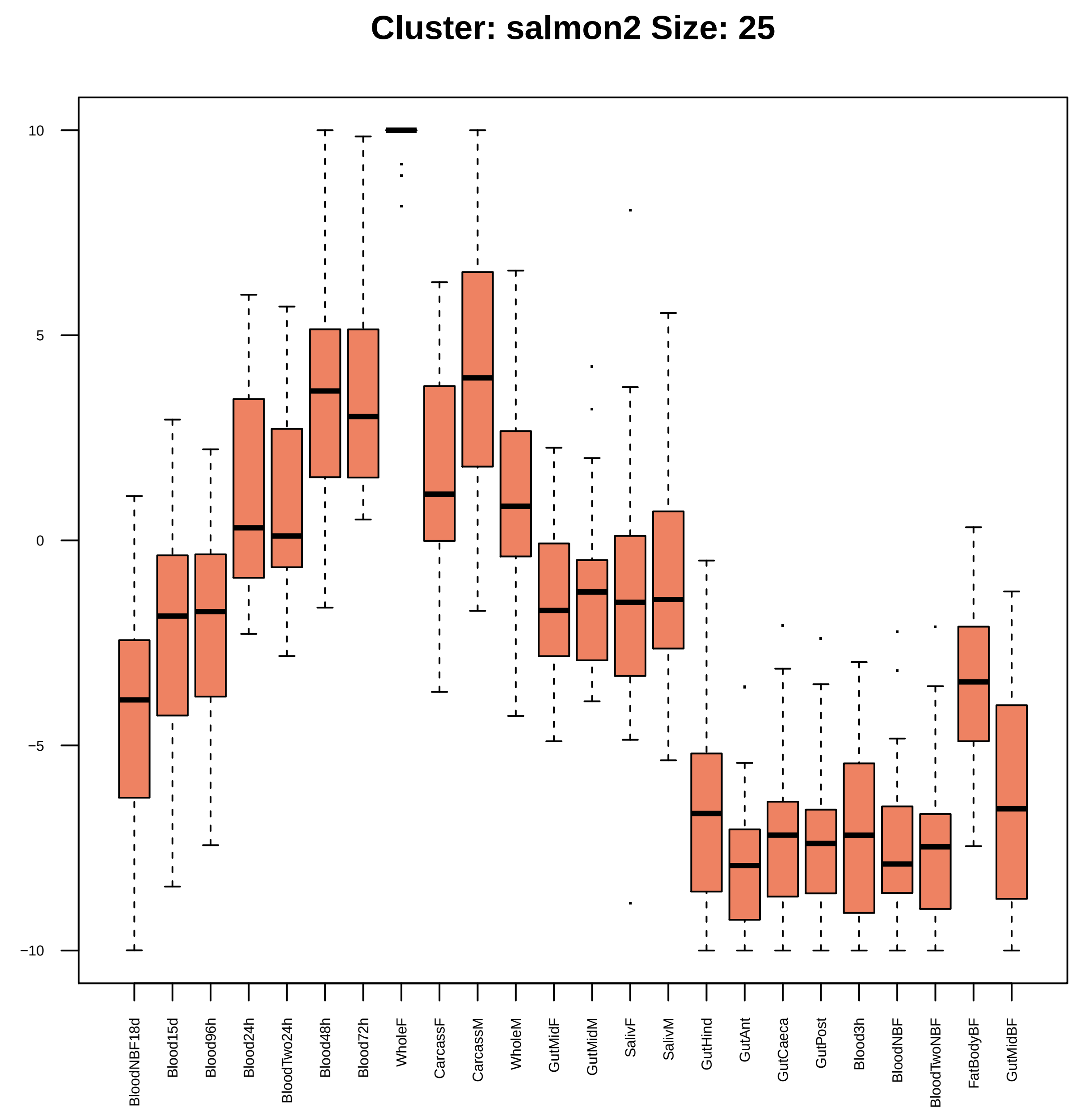

Cluster: salmon2 Size: 25
10
5
0
−5
−10
BloodNBF18d
Blood15d
Blood96h
Blood24h
BloodTwo24h
Blood48h
Blood72h
WholeF
CarcassF
CarcassM
WholeM
GutMidF
GutMidM
SalivF
SalivM
GutHind
GutAnt
GutCaeca
GutPost
Blood3h
BloodNBF
BloodTwoNBF
FatBodyBF
GutMidBF

| | GO.ID | BPCluster: salmon2 Size: 25 | Annotated | Significant | Expected | Rank in ClassicF | Weight01F | ClassicF |
|---|---|---|---|---|---|---|---|---|
| *1* | GO:0006261 | DNA−dependent DNA replication | 52 | 2 | 0.09 | 2 | 0.0034 | 0.0034 |
| *2* | GO:0006281 | DNA repair | 89 | 2 | 0.15 | 5 | 0.0098 | 0.0098 |
| *12* | GO:0007399 | nervous system development | 801 | 5 | 1.38 | 4 | 0.0260 | 0.0074 |

| | GO.ID | MFCluster: salmon2 Size: 25 | Annotated | Significant | Expected | Rank in ClassicF | Weight01F | ClassicF |
|---|---|---|---|---|---|---|---|---|
| *1* | GO:0008270 | zinc ion binding | 522 | 7 | 1.45 | 3 | 0.00034 | 0.00034 |
| *2* | GO:0003677 | DNA binding | 523 | 6 | 1.45 | 7 | 0.00228 | 0.00228 |
| *7* | GO:0046872 | metal ion binding | 1255 | 10 | 3.48 | 4 | 0.04363 | 0.00084 |
| *28* | GO:0046914 | transition metal ion binding | 692 | 7 | 1.92 | 6 | 1.00000 | 0.00185 |

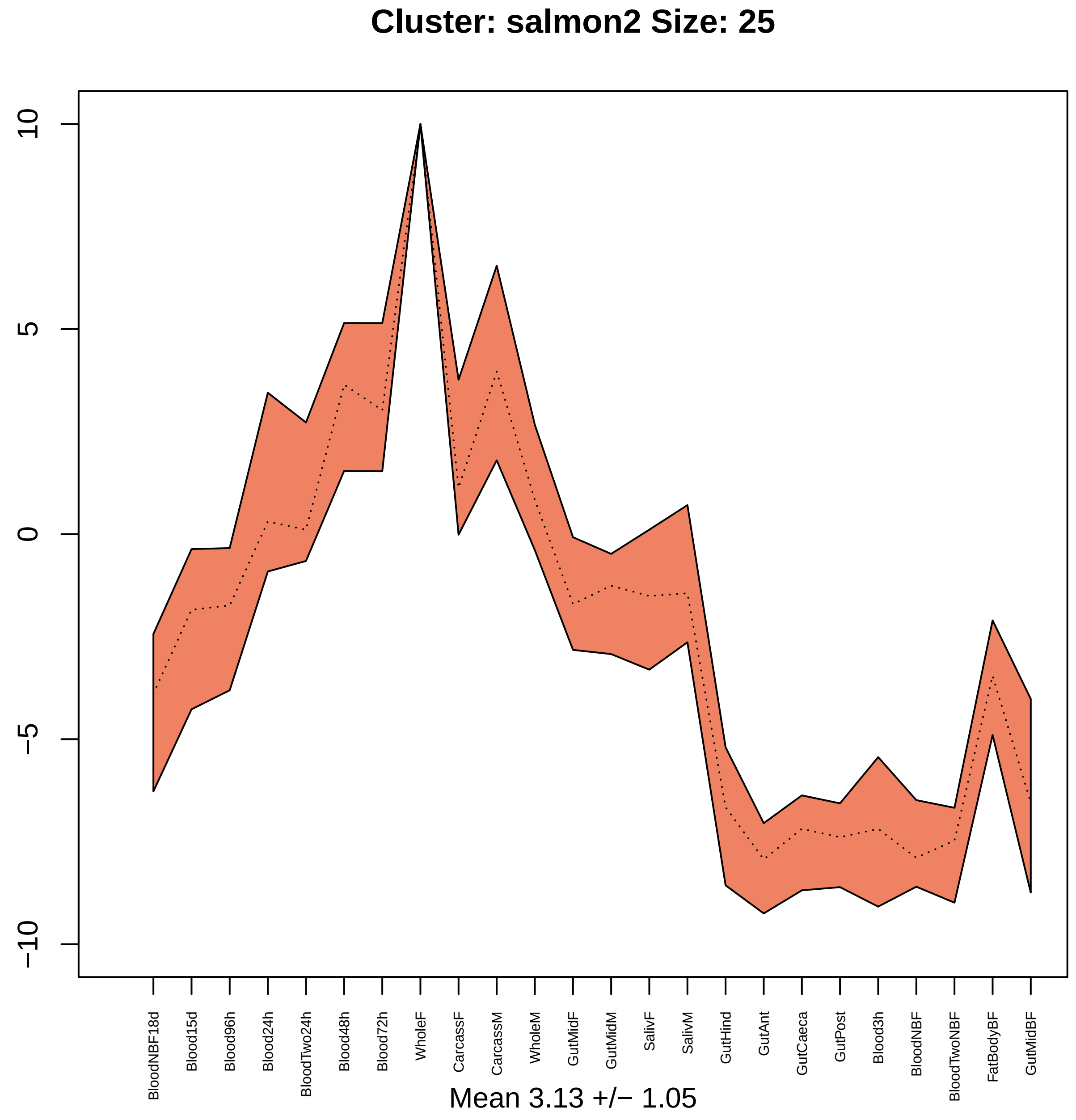

Cluster: salmon2 Size: 25
10
5
0
−5
−10
BloodNBF18d
Blood15d
Blood96h
Blood24h
BloodTwo24h
Blood48h
Blood72h
WholeF
CarcassF
CarcassM
WholeM
GutMidF
GutMidM
SalivF
SalivM
GutHind
GutAnt
GutCaeca
GutPost
Blood3h
BloodNBF
BloodTwoNBF
FatBodyBF
GutMidBF
Mean 3.13 +/− 1.05

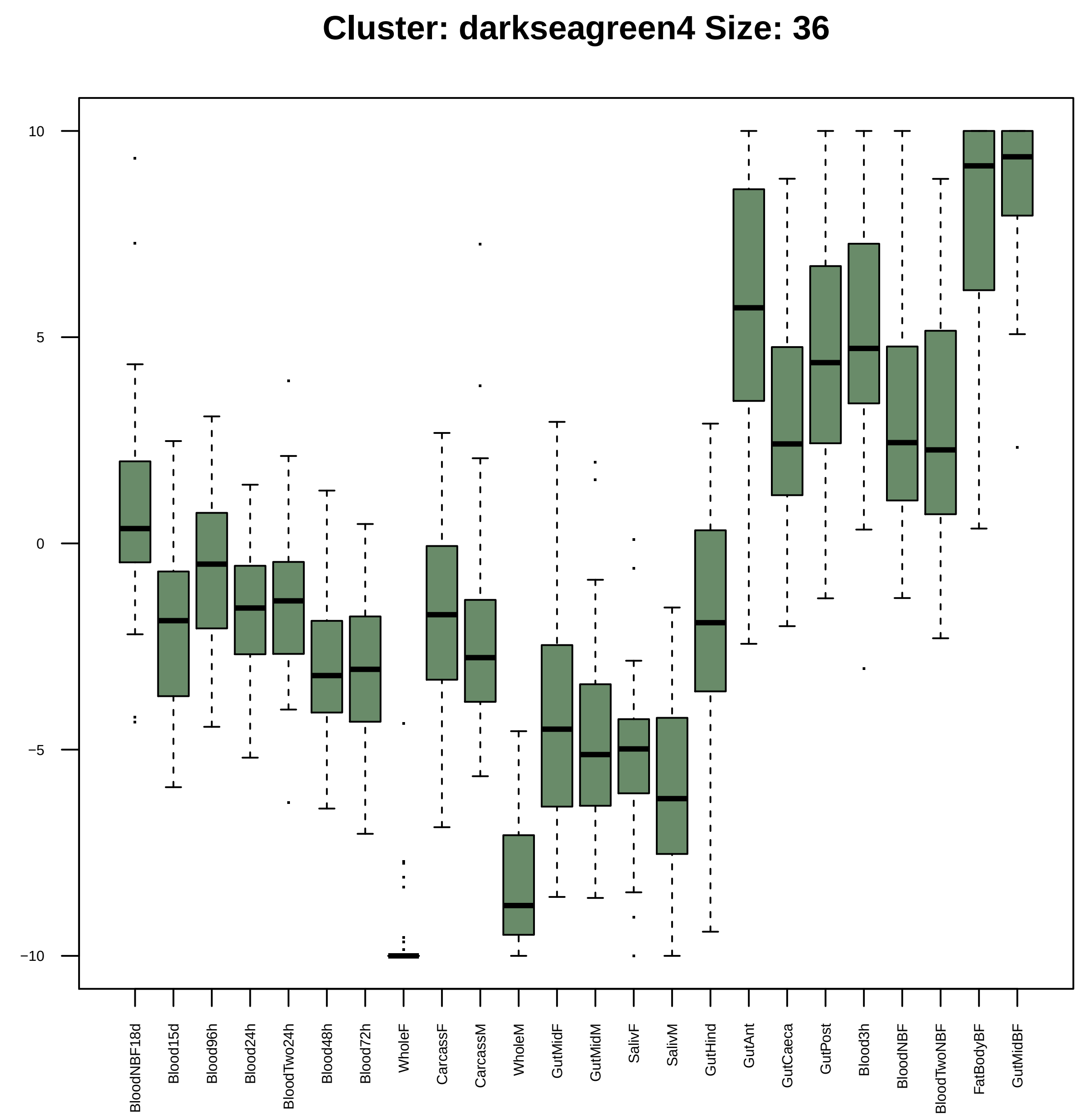

Cluster: darkseagreen4 Size: 36
10
5
0
−5
−10
BloodNBF18d
Blood15d
Blood96h
Blood24h
BloodTwo24h
Blood48h
Blood72h
WholeF
CarcassF
CarcassM
WholeM
GutMidF
GutMidM
SalivF
SalivM
GutHind
GutAnt
GutCaeca
GutPost
Blood3h
BloodNBF
BloodTwoNBF
FatBodyBF
GutMidBF

| | GO.ID | BPCluster: darkseagreen4 Size: 36 | Annotated | Significant | Expected | Rank in ClassicF | Weight01F | ClassicF |
|---|---|---|---|---|---|---|---|---|
| *1* | GO:0043269 | regulation of ion transport | 31 | 2 | 0.09 | 1 | 0.0037 | 0.0037 |

| | GO.ID | MFCluster: darkseagreen4 Size: 36 | Annotated | Significant | Expected | Rank in ClassicF | Weight01F | ClassicF |
|---|---|---|---|---|---|---|---|---|
| *29* | GO:0038023 | signaling receptor activity | 282 | 4 | 0.86 | 2 | 1.000 | 0.0096 |

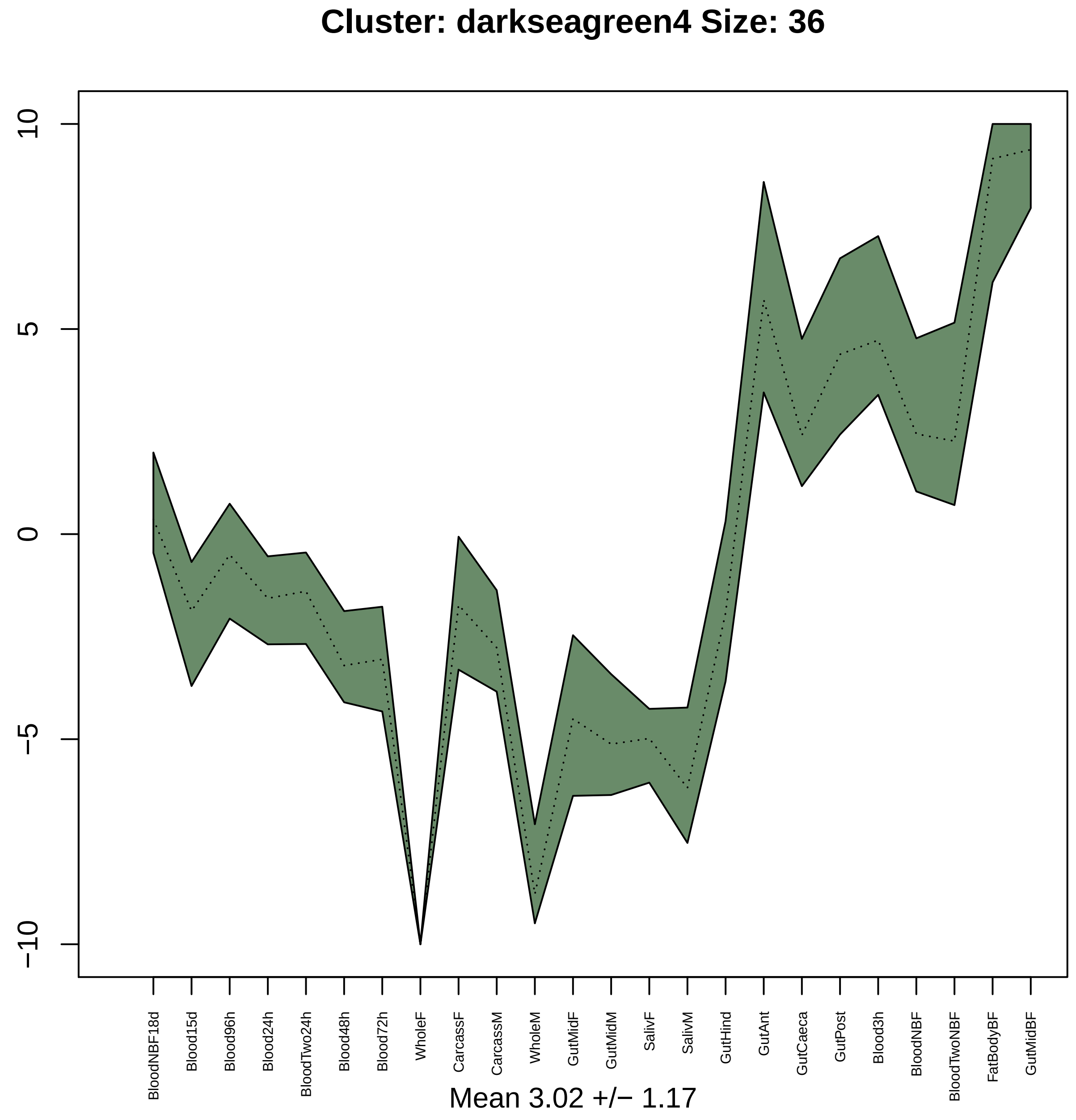
Cluster: darkseagreen4 Size: 36
10
5
0
−5
−10
BloodNBF18d
Blood15d
Blood96h
Blood24h
BloodTwo24h
Blood48h
Blood72h
WholeF
CarcassF
CarcassM
WholeM
GutMidF
GutMidM
SalivF
SalivM
GutHind
GutAnt
GutCaeca
GutPost
Blood3h
BloodNBF
BloodTwoNBF
FatBodyBF
GutMidBF
Mean 3.02 +/− 1.17

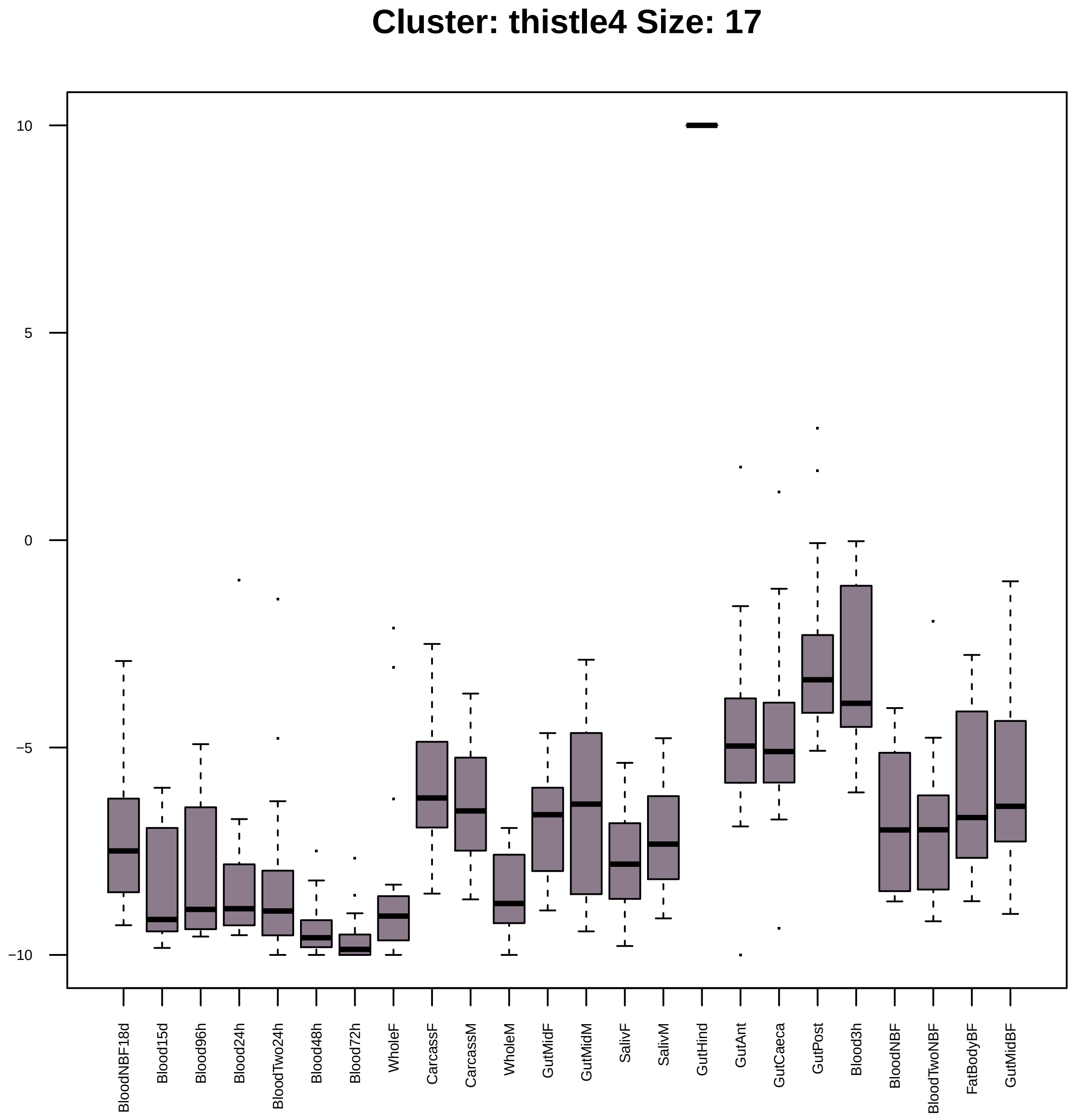
Cluster: thistle4 Size: 17
10
5
0
−5
−10
BloodNBF18d
Blood15d
Blood96h
Blood24h
BloodTwo24h
Blood48h
Blood72h
WholeF
CarcassF
CarcassM
WholeM
GutMidF
GutMidM
SalivF
SalivM
GutHind
GutAnt
GutCaeca
GutPost
Blood3h
BloodNBF
BloodTwoNBF
FatBodyBF
GutMidBF

| | GO.ID | MFCluster: thistle4 Size: 17 | Annotated | Significant | Expected | Rank in ClassicF | Weight01F | ClassicF |
|---|---|---|---|---|---|---|---|---|
| *1* | GO:0008061 | chitin binding | 76 | 2 | 0.15 | 1 | 0.0092 | 0.0092 |

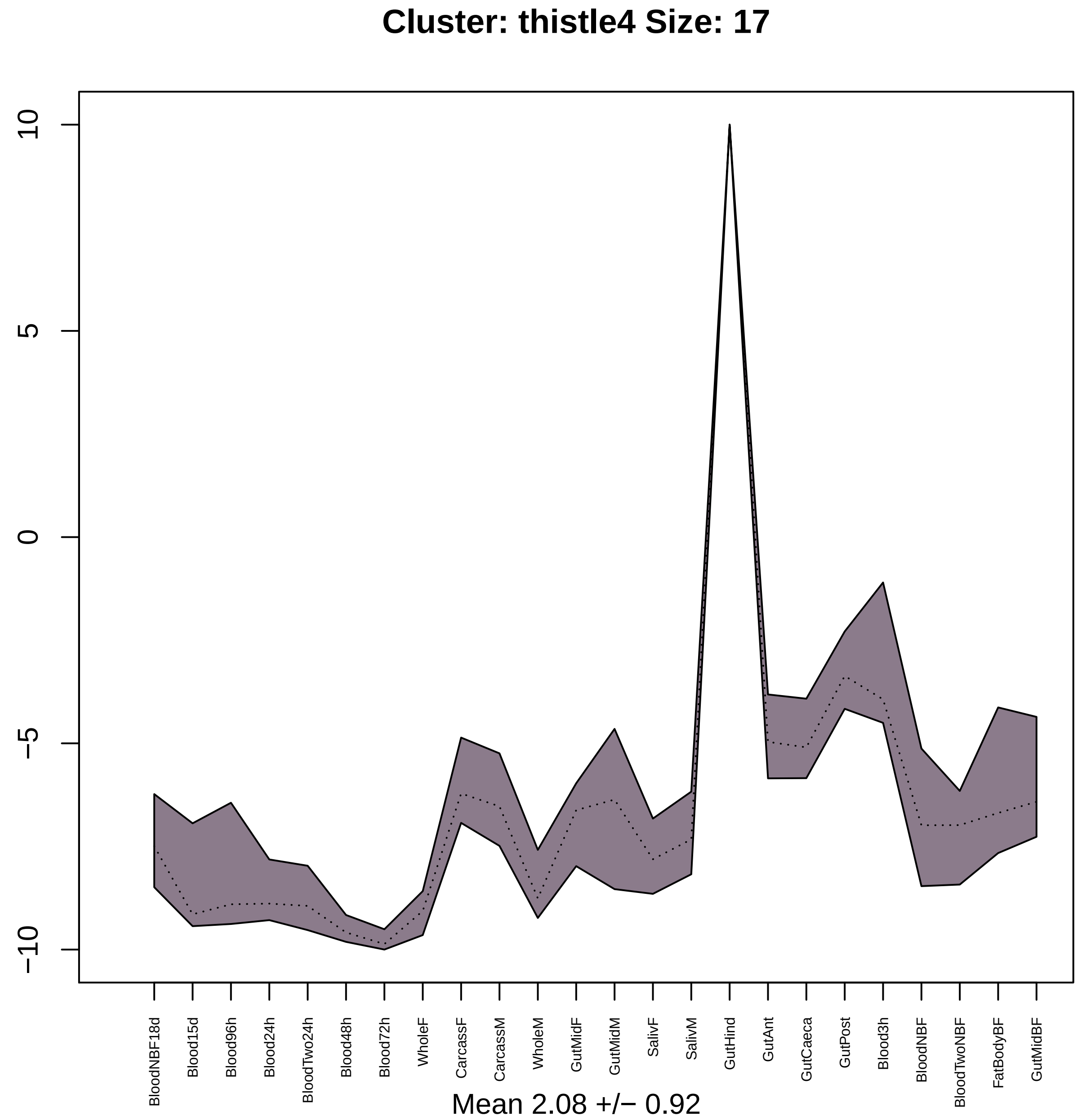

Cluster: thistle4 Size: 17
10
5
0
−5
−10
BloodNBF18d
Blood15d
Blood96h
Blood24h
BloodTwo24h
Blood48h
Blood72h
WholeF
CarcassF
CarcassM
WholeM
GutMidF
GutMidM
SalivF
SalivM
GutHind
GutAnt
GutCaeca
GutPost
Blood3h
BloodNBF
BloodTwoNBF
FatBodyBF
GutMidBF
Mean 2.08 +/− 0.92

### Cluster: plum4 Size: 17

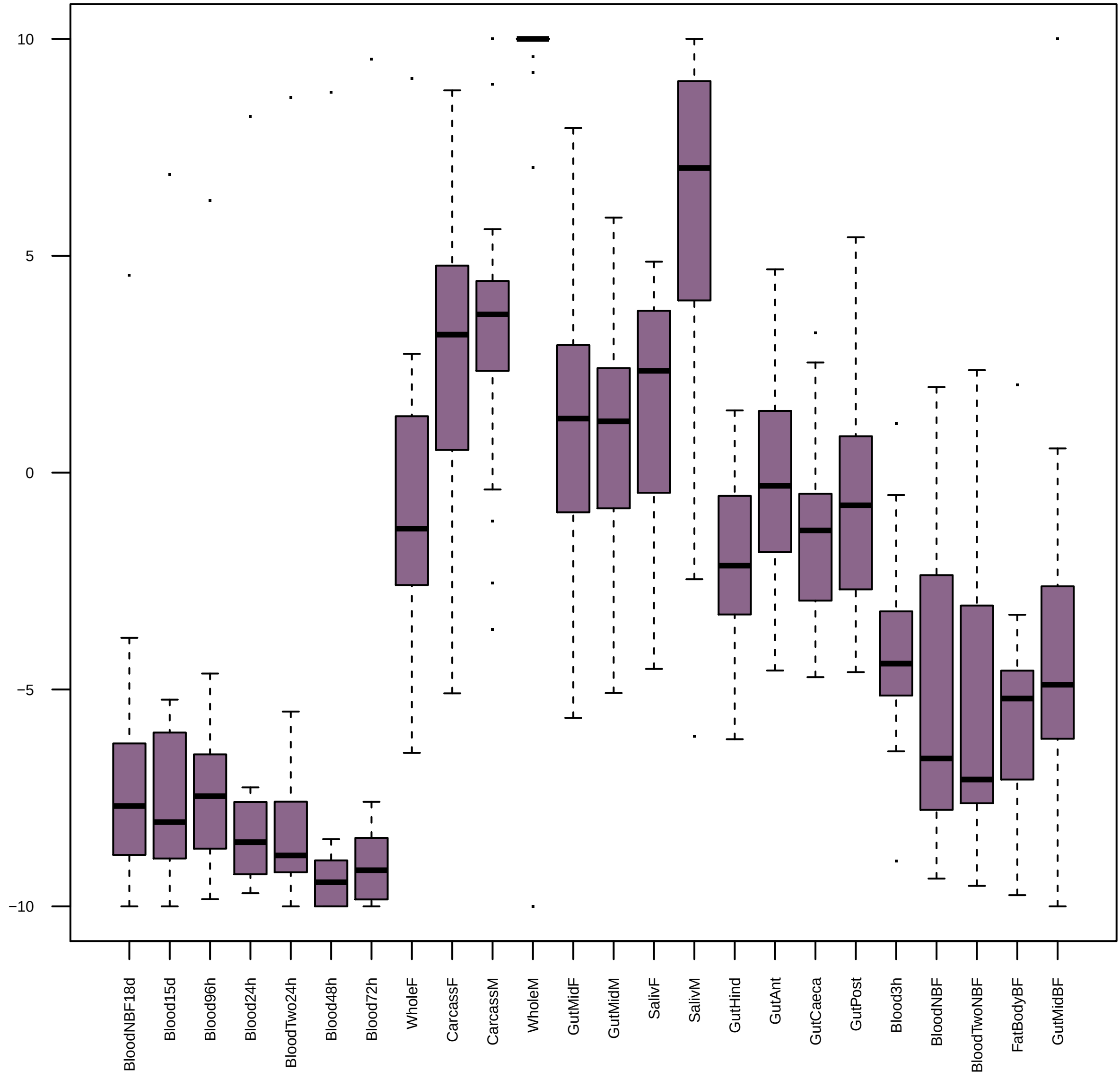

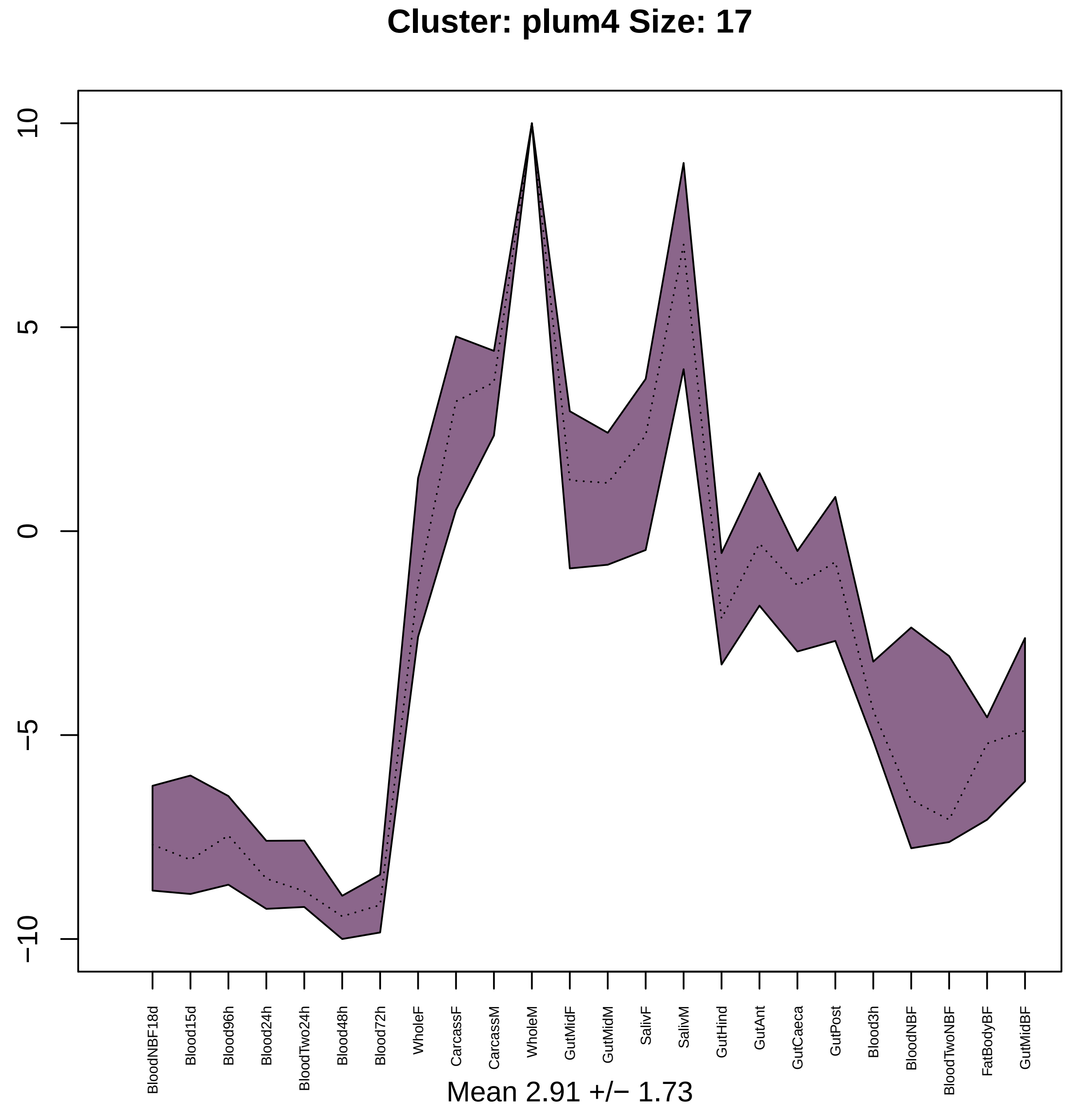

Cluster: plum4 Size: 17
10
5
0
−5
−10
BloodNBF18d
Blood15d
Blood96h
Blood24h
BloodTwo24h
Blood48h
Blood72h
WholeF
CarcassF
CarcassM
WholeM
GutMidF
GutMidM
SalivF
SalivM
GutHind
GutAnt
GutCaeca
GutPost
Blood3h
BloodNBF
BloodTwoNBF
FatBodyBF
GutMidBF
Mean 2.91 +/− 1.73

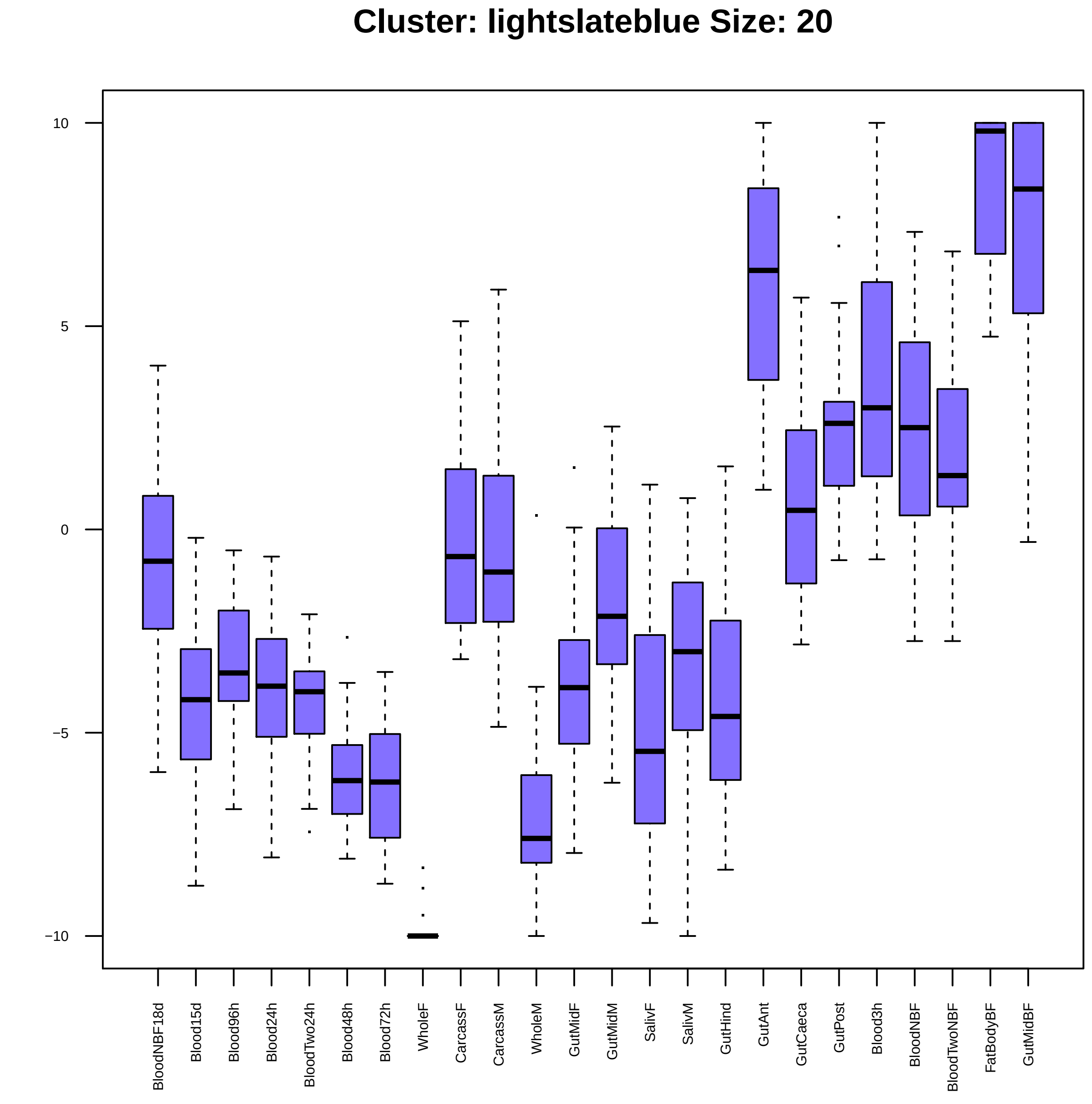
Cluster: lightslateblue Size: 20
10
5
0
−5
−10
BloodNBF18d
Blood15d
Blood96h
Blood24h
BloodTwo24h
Blood48h
Blood72h
WholeF
CarcassF
CarcassM
WholeM
GutMidF
GutMidM
SalivF
SalivM
GutHind
GutAnt
GutCaeca
GutPost
Blood3h
BloodNBF
BloodTwoNBF
FatBodyBF
GutMidBF

| | GO.ID | BPCluster: lightslateblue Size: 20 | Annotated | Significant | Expected | Rank in ClassicF | Weight01F | ClassicF |
|---|---|---|---|---|---|---|---|---|
| 1 | GO:0048639 | positive regulation of developmental gro... | 26 | 2 | 0.05 | 7 | 0.0010 | 0.0010 |
| 2 | GO:0006355 | regulation of transcription, DNA−templat... | 501 | 5 | 0.94 | 10 | 0.0011 | 0.0015 |
| 3 | GO:0048813 | dendrite morphogenesis | 93 | 3 | 0.18 | 5 | 0.0080 | 0.0006 |
| 5 | GO:0006935 | chemotaxis | 129 | 3 | 0.24 | 15 | 0.0143 | 0.0016 |
| 7 | GO:0048518 | positive regulation of biological proces... | 430 | 4 | 0.81 | 43 | 0.0195 | 0.0065 |
| 29 | GO:0007186 | G protein−coupled receptor signaling pat... | 228 | 3 | 0.43 | 47 | 0.0439 | 0.0078 |

| | GO.ID | MFCluster: lightslateblue Size: 20 | Annotated | Significant | Expected | Rank in ClassicF | Weight01F | ClassicF |
|---|---|---|---|---|---|---|---|---|
| *1* | GO:0003700 | DNA−binding transcription factor activit... | 282 | 4 | 0.51 | 3 | 0.0044 | 0.00124 |
| *2* | GO:0008528 | G protein−coupled peptide receptor activ... | 24 | 2 | 0.04 | 1 | 0.0050 | 0.00081 |
| *3* | GO:0003677 | DNA binding | 523 | 5 | 0.94 | 4 | 0.0099 | 0.00155 |
| *11* | GO:0004930 | G protein−coupled receptor activity | 171 | 3 | 0.31 | 6 | 0.2035 | 0.00315 |
| *17* | GO:0140110 | transcription regulator activity | 322 | 4 | 0.58 | 5 | 1.0000 | 0.00203 |

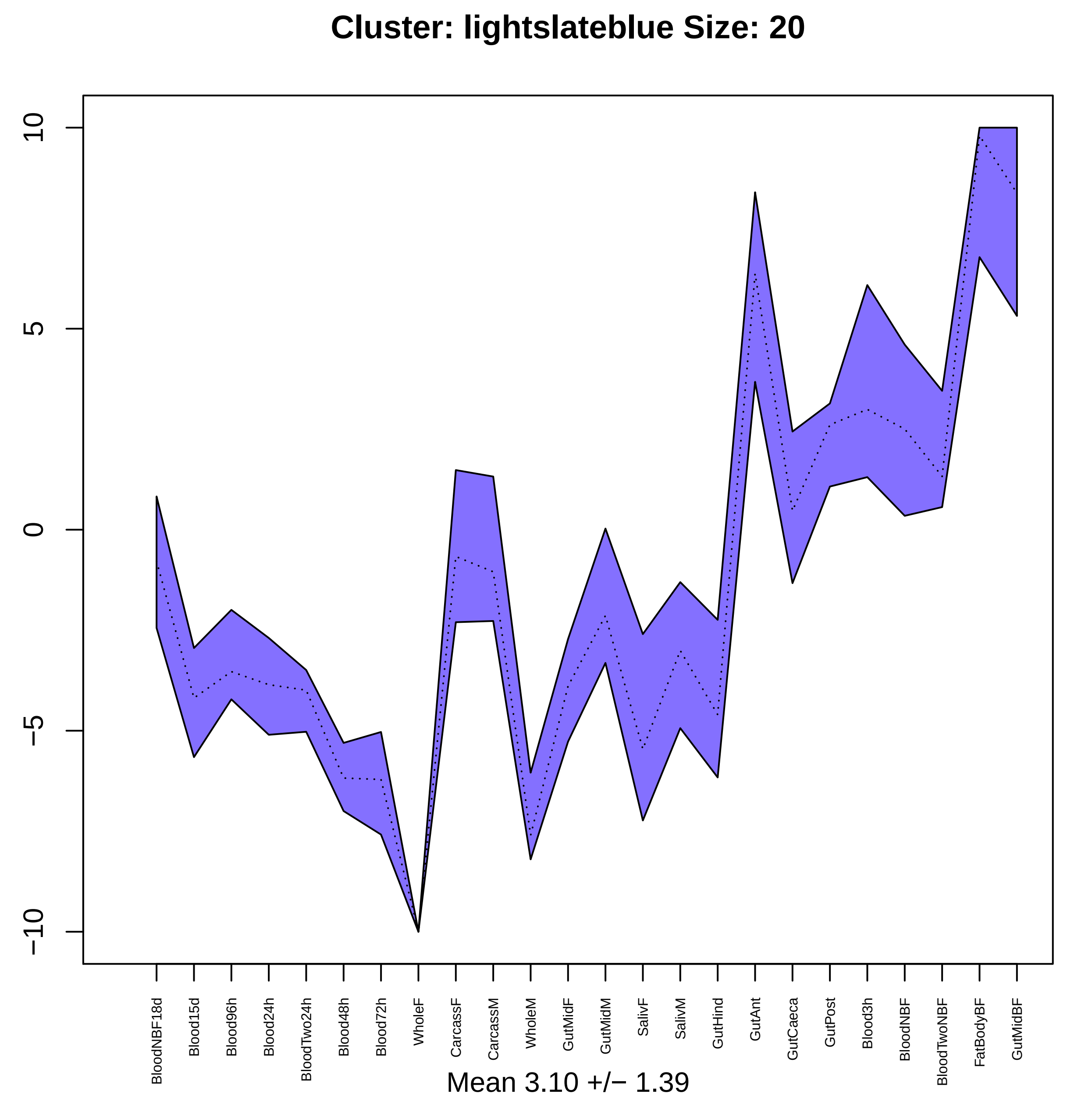

Cluster: lightslateblue Size: 20
10
5
0
−5
−10
BloodNBF18d
Blood15d
Blood96h
Blood24h
BloodTwo24h
Blood48h
Blood72h
WholeF
CarcassF
CarcassM
WholeM
GutMidF
GutMidM
SalivF
SalivM
GutHind
GutAnt
GutCaeca
GutPost
Blood3h
BloodNBF
BloodTwoNBF
FatBodyBF
GutMidBF
Mean 3.10 +/− 1.39

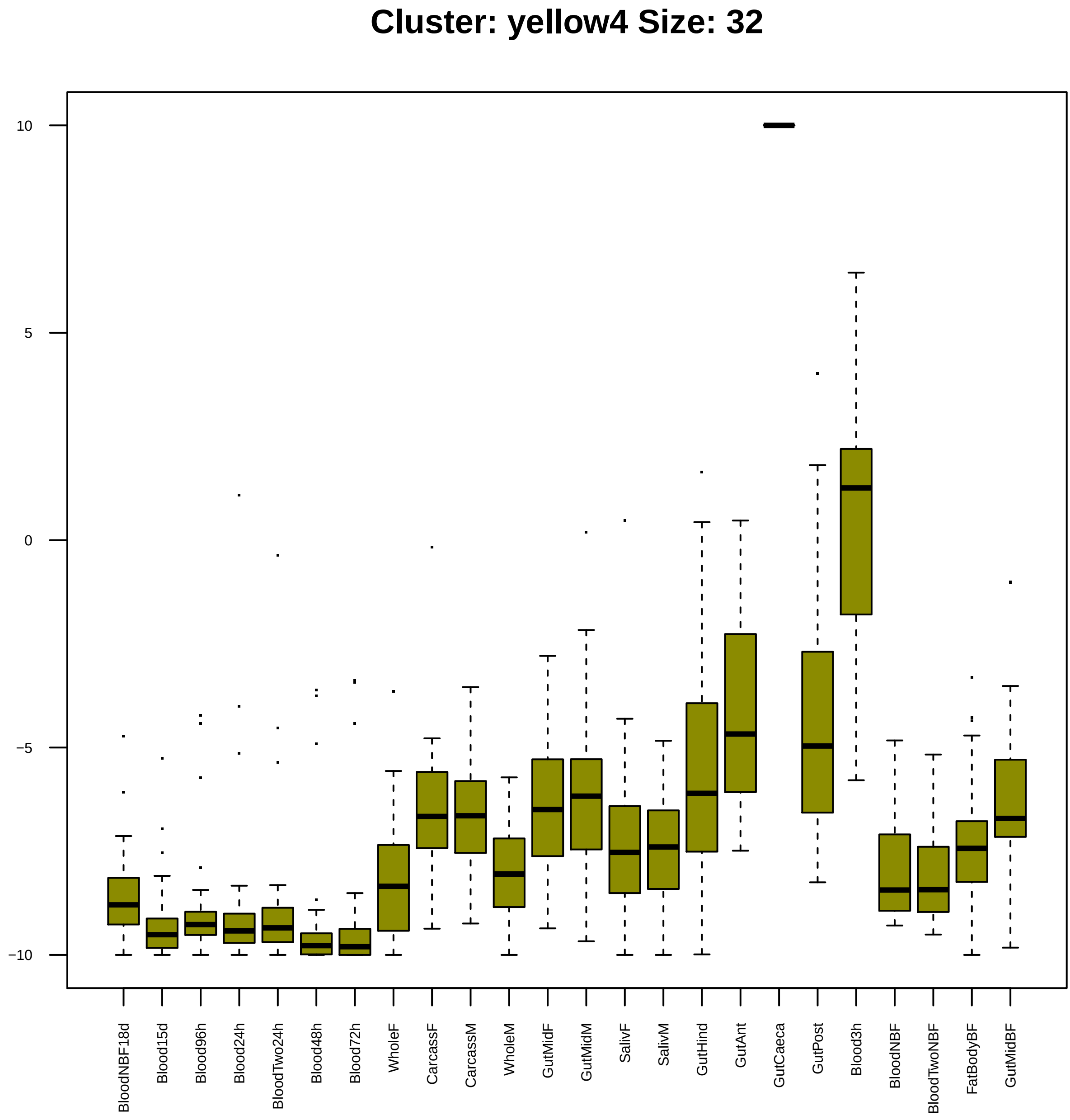

Cluster: yellow4 Size: 32
10
5
0
−5
−10
BloodNBF18d
Blood15d
Blood96h
Blood24h
BloodTwo24h
Blood48h
Blood72h
WholeF
CarcassF
CarcassM
WholeM
GutMidF
GutMidM
SalivF
SalivM
GutHind
GutAnt
GutCaeca
GutPost
Blood3h
BloodNBF
BloodTwoNBF
FatBodyBF
GutMidBF

| | GO.ID | BPCluster: yellow4 Size: 32 | Annotated | Significant | Expected | Rank in ClassicF | Weight01F | ClassicF |
|---|---|---|---|---|---|---|---|---|
| *2* | GO:0006508 | proteolysis | 594 | 7 | 2.33 | 12 | 0.0064 | 0.00635 |
| *23* | GO:0006807 | nitrogen compound metabolic process | 2967 | 20 | 11.63 | 10 | 1.0000 | 0.00064 |
| *26* | GO:0043170 | macromolecule metabolic process | 2534 | 19 | 9.93 | 8 | 1.0000 | 0.00025 |

| | GO.ID | MFCluster: yellow4 Size: 32 | Annotated | Significant | Expected | Rank in ClassicF | Weight01F | ClassicF |
|---|---|---|---|---|---|---|---|---|
| *7* | GO:0008233 | peptidase activity | 500 | 7 | 1.80 | 4 | 0.085 | 0.00153 |
| *20* | GO:0016787 | hydrolase activity | 1445 | 11 | 5.21 | 6 | 0.786 | 0.00794 |
| *21* | GO:0070011 | peptidase activity, acting on L−amino ac... | 470 | 6 | 1.70 | 5 | 1.000 | 0.00558 |
| *24* | GO:0008144 | drug binding | 721 | 10 | 2.60 | 2 | 1.000 | 0.00011 |

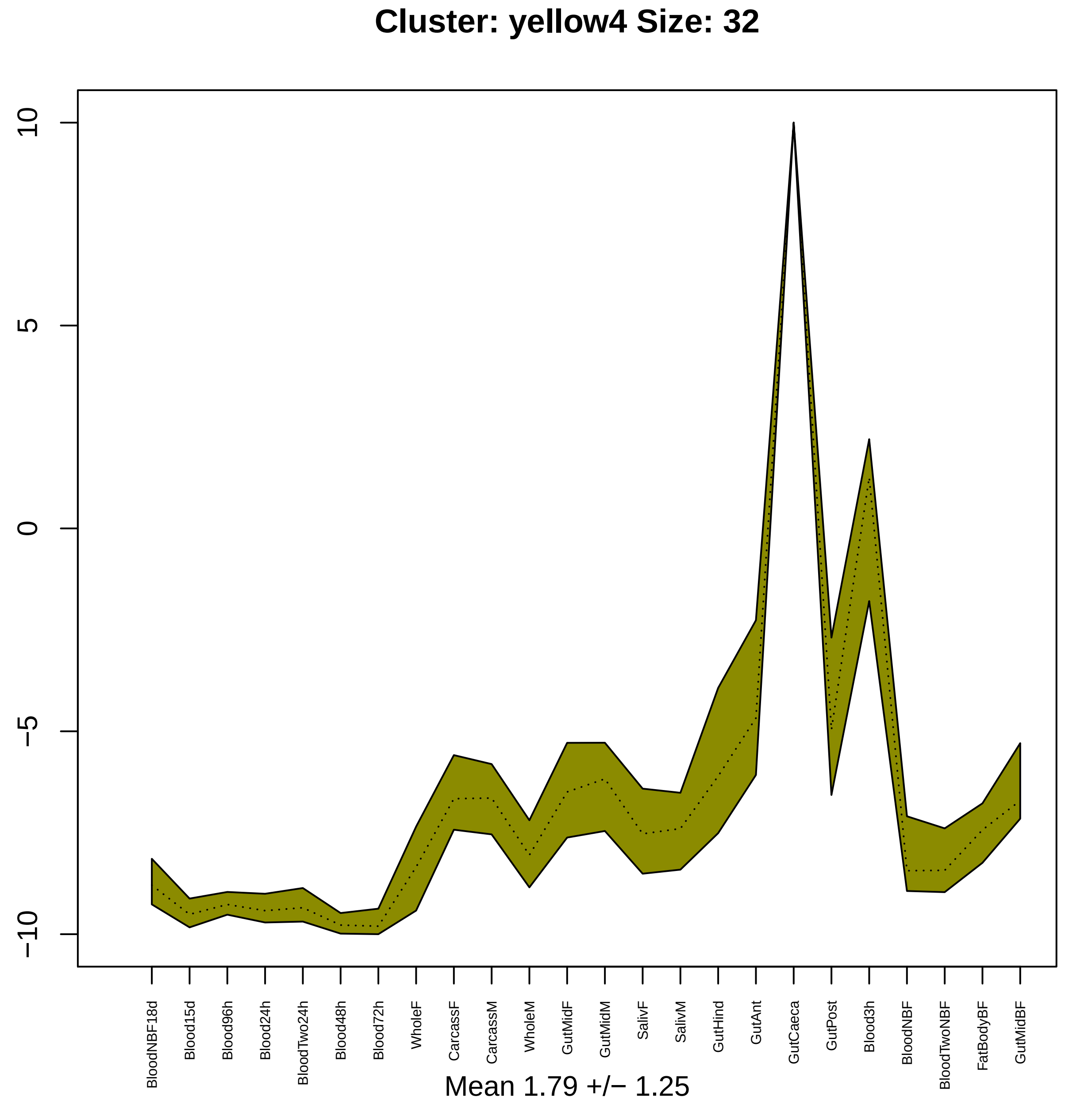
Cluster: yellow4 Size: 32
10
5
0
−5
−10
BloodNBF18d
Blood15d
Blood96h
Blood24h
BloodTwo24h
Blood48h
Blood72h
WholeF
CarcassF
CarcassM
WholeM
GutMidF
GutMidM
SalivF
SalivM
GutHind
GutAnt
GutCaeca
GutPost
Blood3h
BloodNBF
BloodTwoNBF
FatBodyBF
GutMidBF
Mean 1.79 +/− 1.25

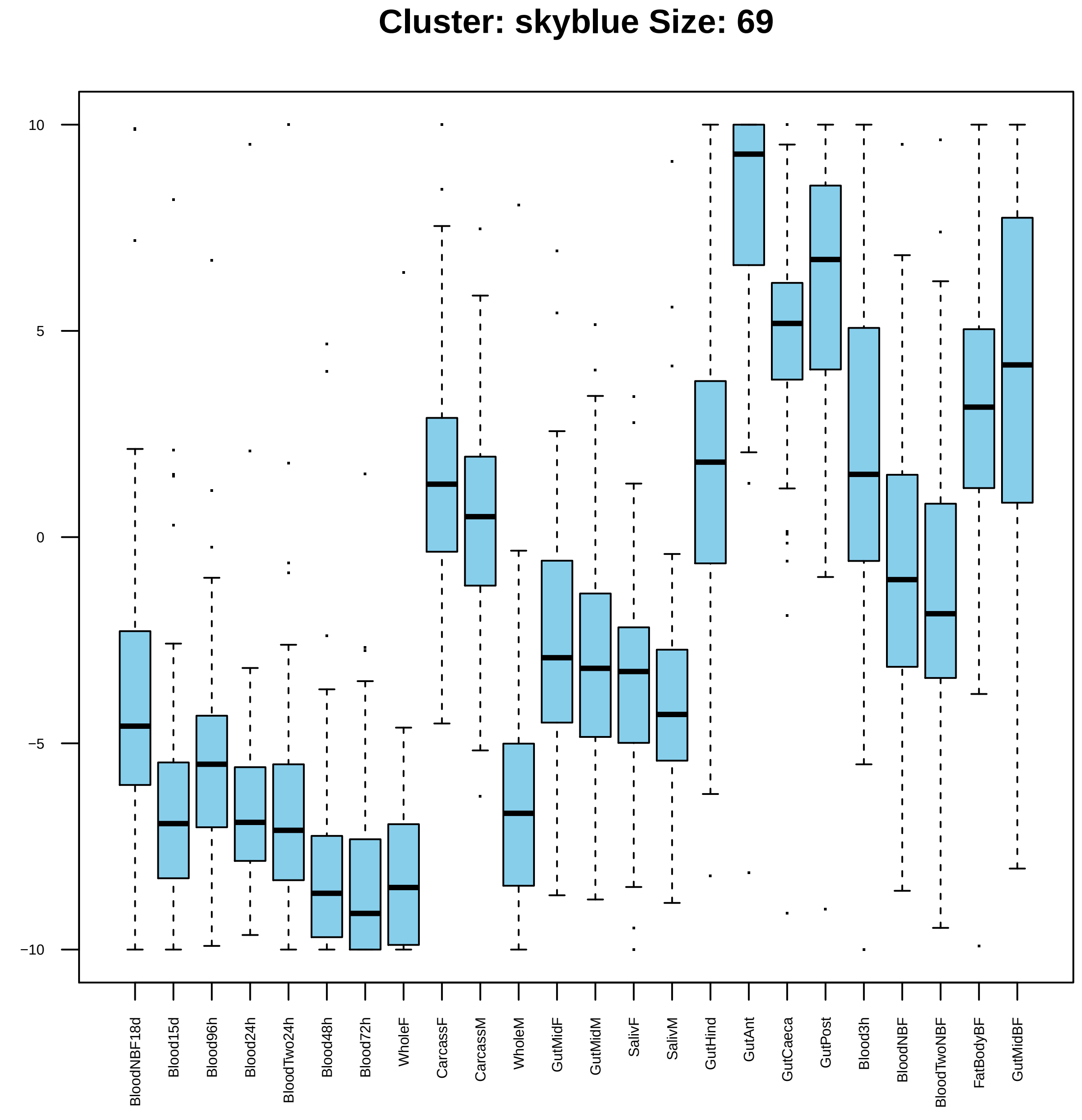

Cluster: skyblue Size: 69
10
5
0
−5
−10
BloodNBF18d
Blood15d
Blood96h
Blood24h
BloodTwo24h
Blood48h
Blood72h
WholeF
CarcassF
CarcassM
WholeM
GutMidF
GutMidM
SalivF
SalivM
GutHind
GutAnt
GutCaeca
GutPost
Blood3h
BloodNBF
BloodTwoNBF
FatBodyBF
GutMidBF

| | GO.ID | BPCluster: skyblue Size: 69 | Annotated | Significant | Expected | Rank in ClassicF | Weight01F | ClassicF |
|---|---|---|---|---|---|---|---|---|
| 1 | GO:0007448 | anterior/posterior pattern specification... | 12 | 2 | 0.08 | 16 | 0.0028 | 0.00281 |
| 2 | GO:0035725 | sodium ion transmembrane transport | 46 | 3 | 0.31 | 17 | 0.0035 | 0.00354 |
| 3 | GO:0007398 | ectoderm development | 14 | 2 | 0.09 | 18 | 0.0038 | 0.00384 |
| 4 | GO:0071310 | cellular response to organic substance | 107 | 5 | 0.72 | 7 | 0.0057 | 0.00070 |
| 11 | GO:0007166 | cell surface receptor signaling pathway | 328 | 8 | 2.21 | 10 | 0.0258 | 0.00134 |
| 23 | GO:0006814 | sodium ion transport | 57 | 4 | 0.38 | 6 | 0.0674 | 0.00055 |

| | GO.ID | MFCluster: skyblue Size: 69 | Annotated | Significant | Expected | Rank in ClassicF | Weight01F | ClassicF |
|---|---|---|---|---|---|---|---|---|
| *1* | GO:0005272 | sodium channel activity | 26 | 3 | 0.17 | 14 | 0.00061 | 0.00061 |
| *2* | GO:0043565 | sequence−specific DNA binding | 212 | 6 | 1.38 | 20 | 0.00381 | 0.00236 |
| *3* | GO:0008188 | neuropeptide receptor activity | 21 | 2 | 0.14 | 27 | 0.00807 | 0.00807 |
| *4* | GO:0003700 | DNA−binding transcription factor activit... | 282 | 7 | 1.84 | 18 | 0.00932 | 0.00213 |
| *5* | GO:0022836 | gated channel activity | 89 | 5 | 0.58 | 10 | 0.01204 | 0.00026 |
| *9* | GO:0008528 | G protein−coupled peptide receptor activ... | 24 | 3 | 0.16 | 13 | 0.01867 | 0.00048 |
| *21* | GO:0005230 | extracellular ligand−gated ion channel a... | 51 | 3 | 0.33 | 22 | 0.12386 | 0.00434 |
| *24* | GO:0022839 | ion gated channel activity | 87 | 4 | 0.57 | 21 | 0.14856 | 0.00238 |

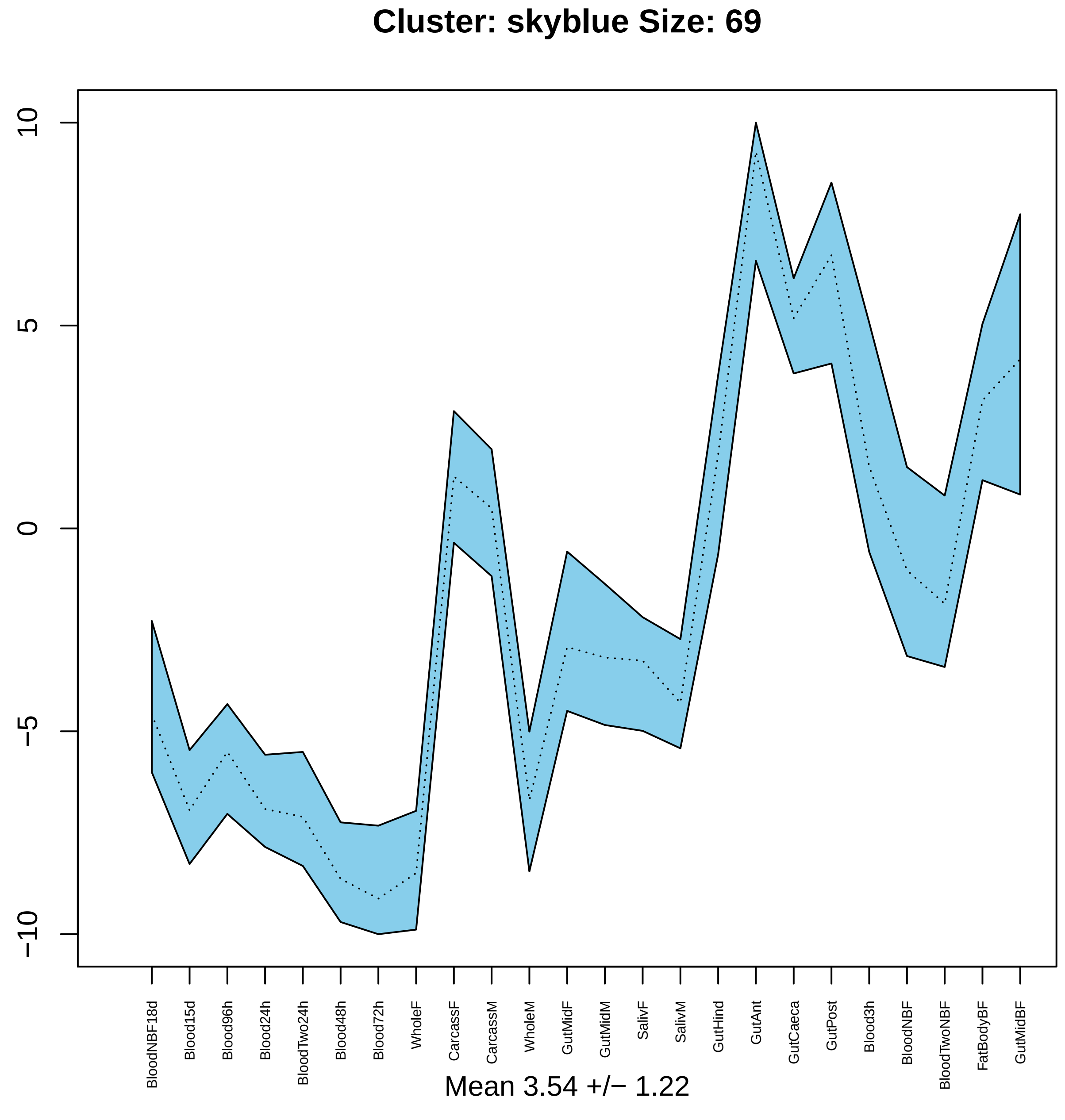
Cluster: skyblue Size: 69
10
5
0
−5
−10
BloodNBF18d
Blood15d
Blood96h
Blood24h
BloodTwo24h
Blood48h
Blood72h
WholeF
CarcassF
CarcassM
WholeM
GutMidF
GutMidM
SalivF
SalivM
GutHind
GutAnt
GutCaeca
GutPost
Blood3h
BloodNBF
BloodTwoNBF
FatBodyBF
GutMidBF
Mean 3.54 +/− 1.22

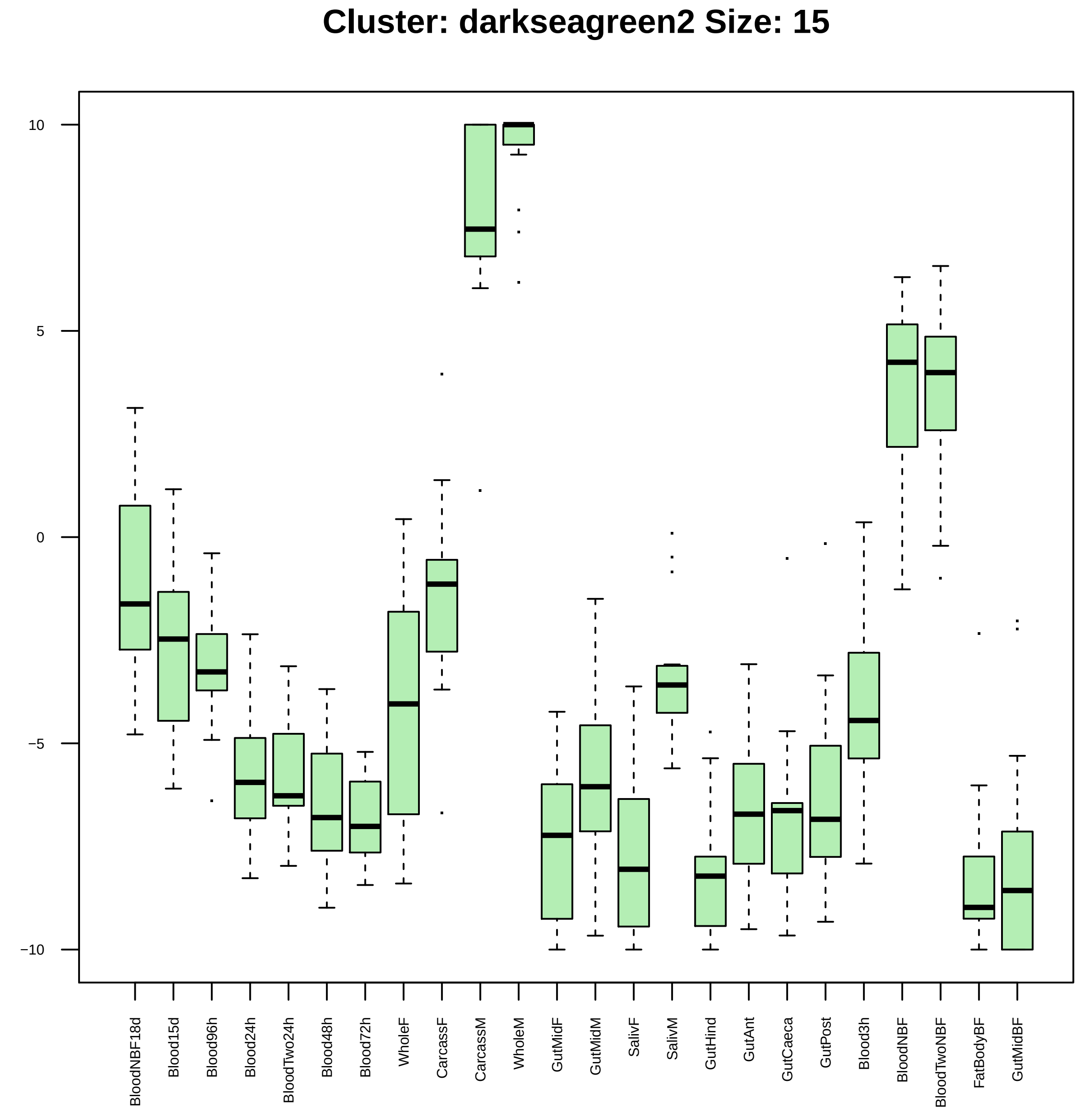
Cluster: darkseagreen2 Size: 15
10
5
0
−5
−10
BloodNBF18d
Blood15d
Blood96h
Blood24h
BloodTwo24h
Blood48h
Blood72h
WholeF
CarcassF
CarcassM
WholeM
GutMidF
GutMidM
SalivF
SalivM
GutHind
GutAnt
GutCaeca
GutPost
Blood3h
BloodNBF
BloodTwoNBF
FatBodyBF
GutMidBF

| | GO.ID | BPCluster: darkseagreen2 Size: 15 | Annotated | Significant | Expected | Rank in ClassicF | Weight01F | ClassicF |
|---|---|---|---|---|---|---|---|---|
| *1* | GO:0006836 | neurotransmitter transport | 81 | 2 | 0.1 | 1 | 0.0042 | 0.0042 |

| | GO.ID | MFCluster: darkseagreen2 Size: 15 | Annotated | Significant | Expected | Rank in ClassicF | Weight01F | ClassicF |
|---|---|---|---|---|---|---|---|---|
| 1 | GO:0017022 | myosin binding | 19 | 2 | 0.03 | 1 | 0.00029 | 0.00029 |
| 2 | GO:0005516 | calmodulin binding | 19 | 2 | 0.03 | 2 | 0.00029 | 0.00029 |

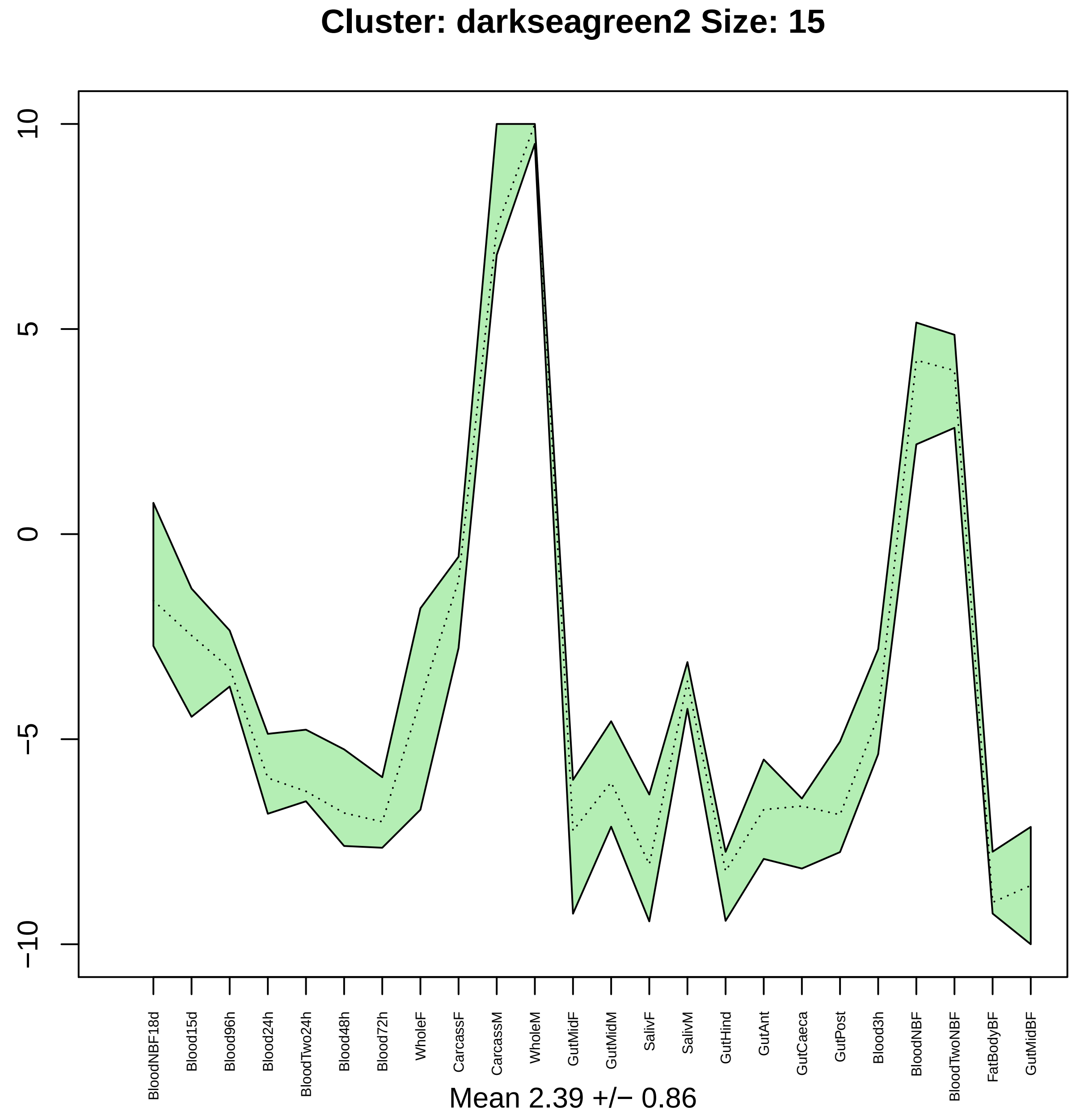

Cluster: darkseagreen2 Size: 15
10
5
0
−5
−10
BloodNBF18d
Blood15d
Blood96h
Blood24h
BloodTwo24h
Blood48h
Blood72h
WholeF
CarcassF
CarcassM
WholeM
GutMidF
GutMidM
SalivF
SalivM
GutHind
GutAnt
GutCaeca
GutPost
Blood3h
BloodNBF
BloodTwoNBF
FatBodyBF
GutMidBF
Mean 2.39 +/− 0.86

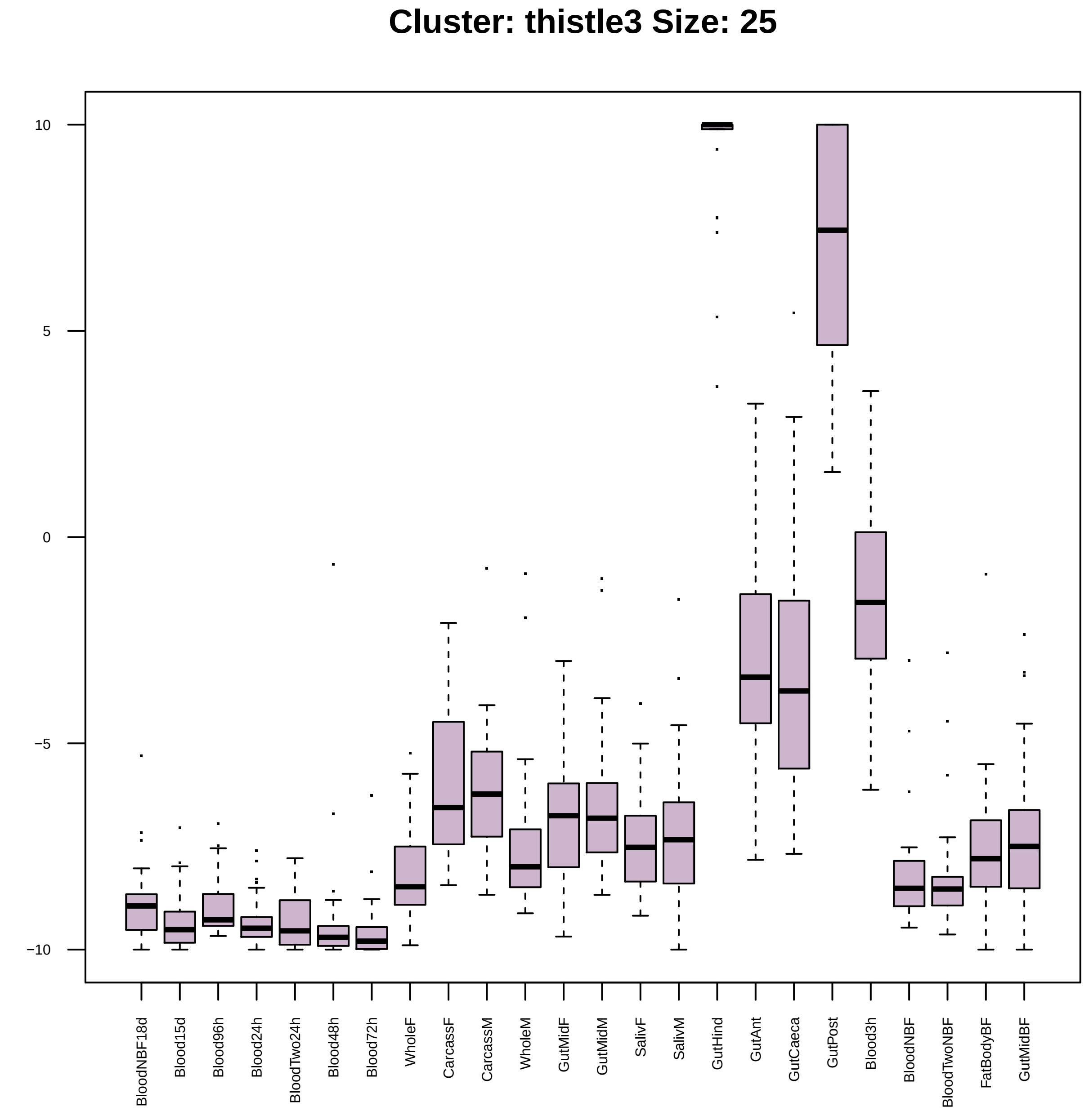

Cluster: thistle3 Size: 25
10
5
0
−5
−10
BloodNBF18d
Blood15d
Blood96h
Blood24h
BloodTwo24h
Blood48h
Blood72h
WholeF
CarcassF
CarcassM
WholeM
GutMidF
GutMidM
SalivF
SalivM
GutHind
GutAnt
GutCaeca
GutPost
Blood3h
BloodNBF
BloodTwoNBF
FatBodyBF
GutMidBF

| | GO.ID | BPCluster: thistle3 Size: 25 | Annotated | Significant | Expected | Rank in ClassicF | Weight01F | ClassicF |
|---|---|---|---|---|---|---|---|---|
| *2* | GO:0006508 | proteolysis | 594 | 7 | 1.86 | 8 | 0.0015 | 0.00155 |
| *18* | GO:1901564 | organonitrogen compound metabolic proces... | 1889 | 14 | 5.92 | 7 | 1.0000 | 0.00022 |

|  | GO.ID | MFCluster: thistle3 Size: 25 | Annotated | Significant | Expected | Rank in ClassicF | Weight01F | ClassicF |
|---|---|---|---|---|---|---|---|---|
| *2* | GO:0004181 | metallocarboxypeptidase activity | 18 | 2 | 0.04 | 4 | 0.00088 | 0.00088 |
| *3* | GO:0008236 | serine−type peptidase activity | 278 | 4 | 0.69 | 8 | 0.00156 | 0.00432 |
| *16* | GO:0017171 | serine hydrolase activity | 279 | 4 | 0.70 | 9 | 1.00000 | 0.00437 |
| *22* | GO:0008233 | peptidase activity | 500 | 7 | 1.25 | 3 | 1.00000 | 0.00012 |
| *24* | GO:0008235 | metalloexopeptidase activity | 29 | 2 | 0.07 | 7 | 1.00000 | 0.00230 |
| *27* | GO:0008237 | metallopeptidase activity | 104 | 3 | 0.26 | 6 | 1.00000 | 0.00203 |
| *29* | GO:0004180 | carboxypeptidase activity | 24 | 2 | 0.06 | 5 | 1.00000 | 0.00157 |

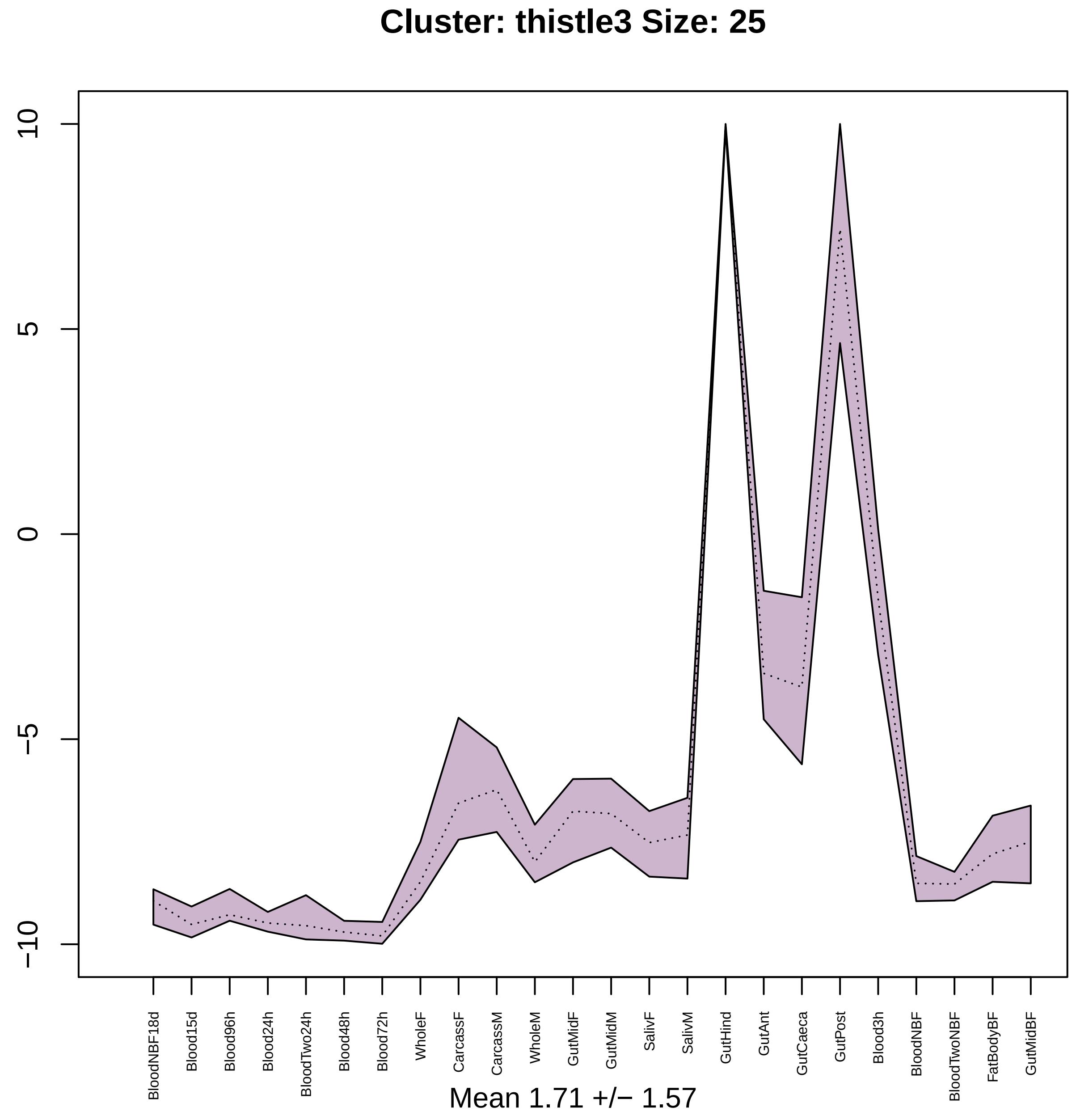
Cluster: thistle3 Size: 25
10
5
0
−5
−10
BloodNBF18d
Blood15d
Blood96h
Blood24h
BloodTwo24h
Blood48h
Blood72h
WholeF
CarcassF
CarcassM
WholeM
GutMidF
GutMidM
SalivF
SalivM
GutHind
GutAnt
GutCaeca
GutPost
Blood3h
BloodNBF
BloodTwoNBF
FatBodyBF
GutMidBF
Mean 1.71 +/− 1.57

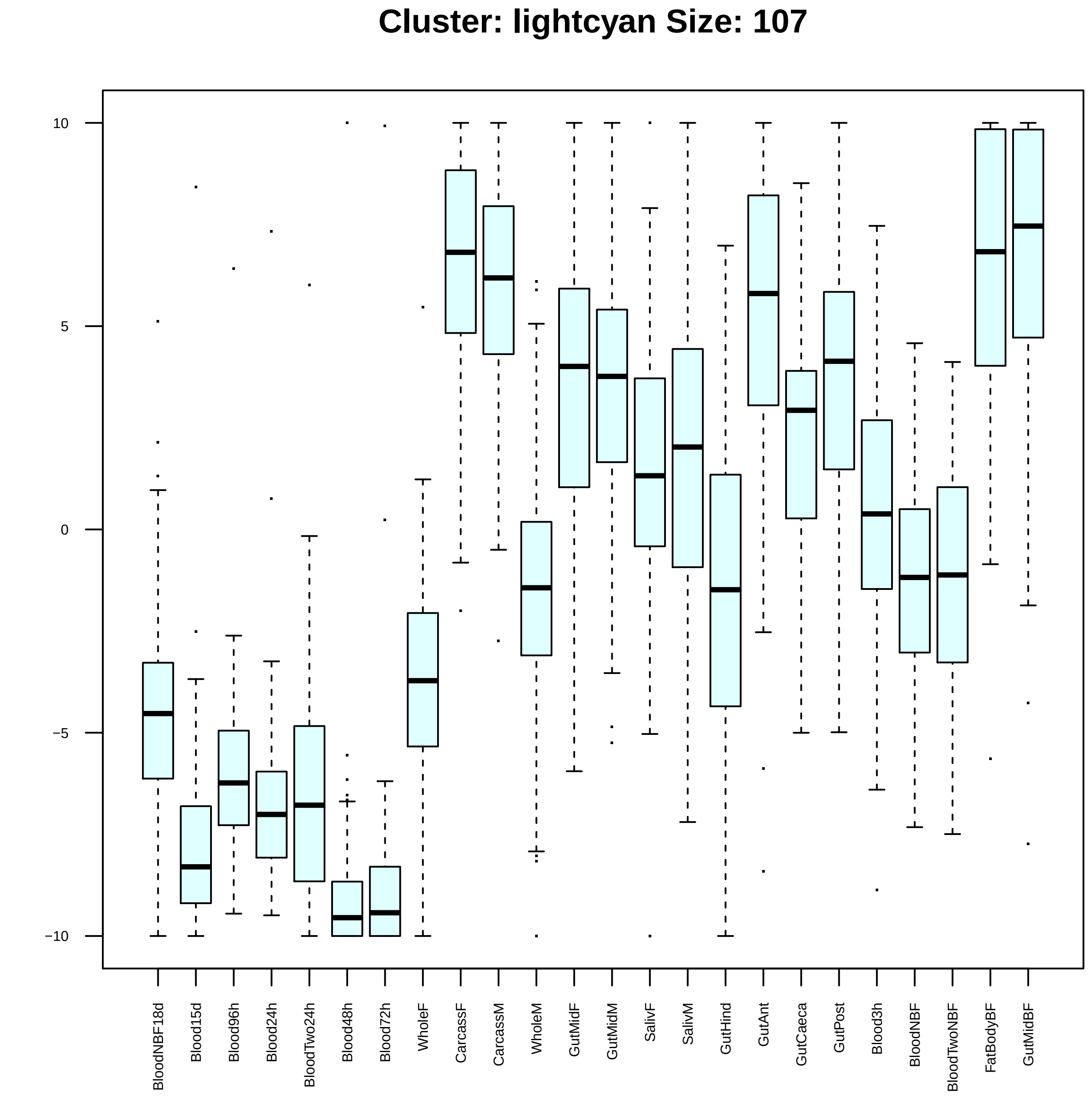

Cluster: lightcyan Size: 107
10
5
0
−5
−10
BloodNBF18d
Blood15d
Blood96h
Blood24h
BloodTwo24h
Blood48h
Blood72h
WholeF
CarcassF
CarcassM
WholeM
GutMidF
GutMidM
SalivF
SalivM
GutHind
GutAnt
GutCaeca
GutPost
Blood3h
BloodNBF
BloodTwoNBF
FatBodyBF
GutMidBF

| | GO.ID | BPCluster: lightcyan Size: 107 | Annotated | Significant | Expected | Rank in ClassicF | Weight01F | ClassicF |
|---|---|---|---|---|---|---|---|---|
| 1 | GO:0050877 | nervous system process | 284 | 10 | 2.54 | 8 | 0.00013 | 0.00018 |
| 2 | GO:0000122 | negative regulation of transcription by ... | 57 | 5 | 0.51 | 6 | 0.00014 | 0.00014 |
| 3 | GO:0051052 | regulation of DNA metabolic process | 13 | 3 | 0.12 | 9 | 0.00018 | 0.00018 |
| 4 | GO:0010948 | negative regulation of cell cycle proces... | 65 | 3 | 0.58 | 90 | 0.00076 | 0.02000 |
| 5 | GO:0007186 | G protein−coupled receptor signaling pat... | 228 | 9 | 2.04 | 7 | 0.00143 | 0.00016 |
| 6 | GO:0007411 | axon guidance | 114 | 5 | 1.02 | 37 | 0.00334 | 0.00334 |
| 7 | GO:0050911 | detection of chemical stimulus involved ... | 71 | 4 | 0.63 | 39 | 0.00357 | 0.00357 |
| 8 | GO:0035147 | branch fusion, open tracheal system | 11 | 2 | 0.10 | 42 | 0.00410 | 0.00410 |
| 9 | GO:0048072 | compound eye pigmentation | 11 | 2 | 0.10 | 43 | 0.00410 | 0.00410 |
| 10 | GO:0042023 | DNA endoreduplication | 12 | 2 | 0.11 | 48 | 0.00489 | 0.00489 |
| 11 | GO:0042048 | olfactory behavior | 43 | 3 | 0.38 | 57 | 0.00649 | 0.00649 |
| 12 | GO:0048070 | regulation of developmental pigmentation | 14 | 2 | 0.13 | 58 | 0.00667 | 0.00667 |
| 13 | GO:0007419 | ventral cord development | 15 | 2 | 0.13 | 61 | 0.00765 | 0.00765 |
| 14 | GO:0007422 | peripheral nervous system development | 48 | 3 | 0.43 | 67 | 0.00881 | 0.00881 |
| 15 | GO:0035017 | cuticle pattern formation | 17 | 2 | 0.15 | 69 | 0.00980 | 0.00980 |
| 17 | GO:0006355 | regulation of transcription, DNA−templat... | 501 | 12 | 4.48 | 23 | 0.01284 | 0.00130 |
| 20 | GO:0043473 | pigmentation | 54 | 4 | 0.48 | 25 | 0.02539 | 0.00130 |
| 22 | GO:2001141 | regulation of RNA biosynthetic process | 505 | 13 | 4.51 | 15 | 0.03029 | 0.00039 |
| 24 | GO:0009593 | detection of chemical stimulus | 87 | 6 | 0.78 | 5 | 0.03265 | 0.00011 |
| 29 | GO:0007417 | central nervous system development | 98 | 5 | 0.88 | 29 | 0.03757 | 0.00172 |

| | GO.ID | MFCluster: lightcyan Size: 107 | Annotated | Significant | Expected | Rank in ClassicF | Weight01F | ClassicF |
|---|---|---|---|---|---|---|---|---|
| *1* | GO:0004930 | G protein−coupled receptor activity | 171 | 9 | 1.61 | 4 | 0.00037 | 2.9e−05 |
| *2* | GO:0043565 | sequence−specific DNA binding | 212 | 8 | 2.00 | 7 | 0.00080 | 0.00079 |
| *3* | GO:0003700 | DNA−binding transcription factor activit... | 282 | 10 | 2.66 | 6 | 0.01027 | 0.00028 |

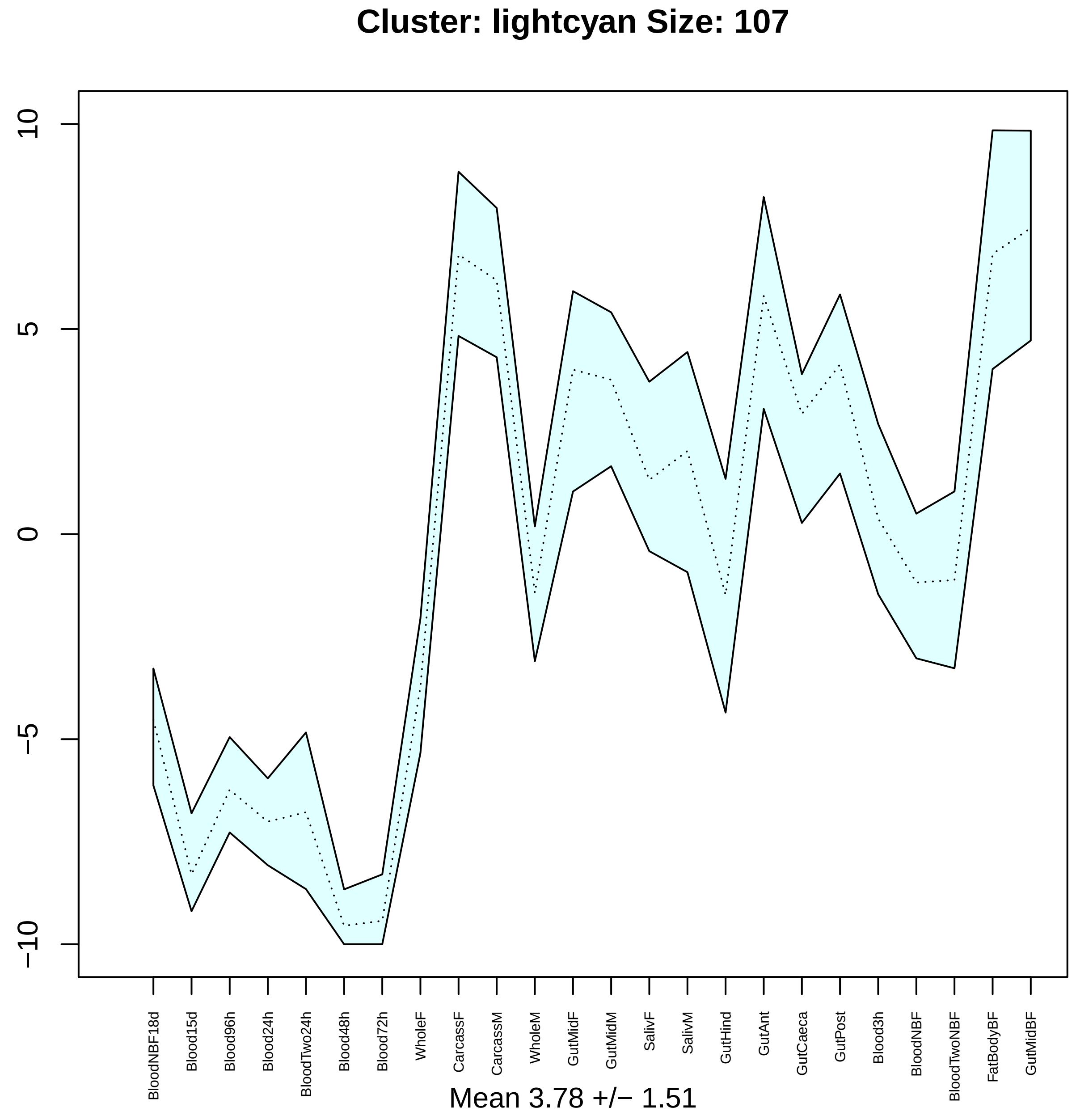

Cluster: lightcyan Size: 107
10
5
0
−5
−10
BloodNBF18d
Blood15d
Blood96h
Blood24h
BloodTwo24h
Blood48h
Blood72h
WholeF
CarcassF
CarcassM
WholeM
GutMidF
GutMidM
SalivF
SalivM
GutHind
GutAnt
GutCaeca
GutPost
Blood3h
BloodNBF
BloodTwoNBF
FatBodyBF
GutMidBF
Mean 3.78 +/− 1.51

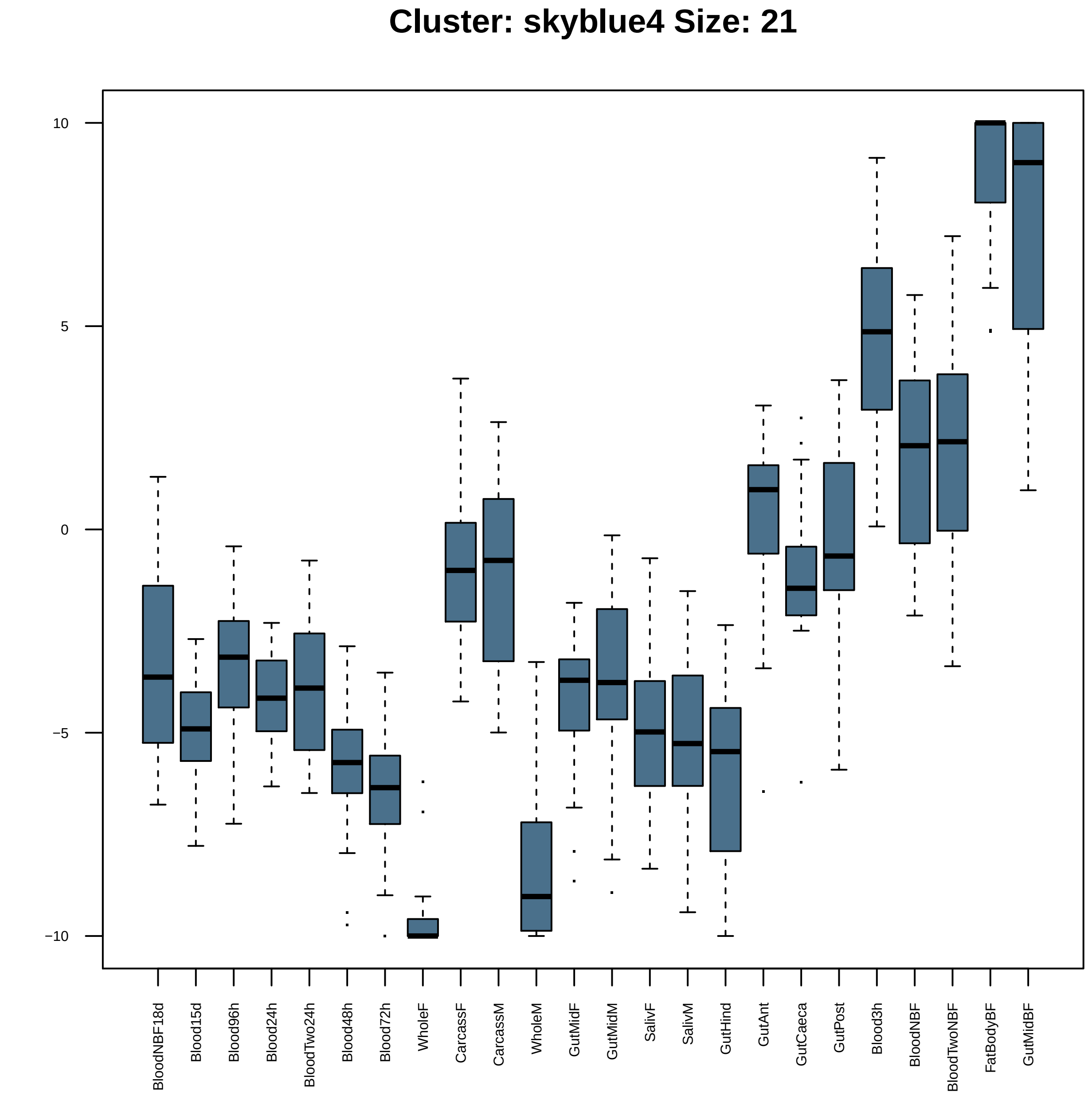

Cluster: skyblue4 Size: 21
10
5
0
−5
−10
BloodNBF18d
Blood15d
Blood96h
Blood24h
BloodTwo24h
Blood48h
Blood72h
WholeF
CarcassF
CarcassM
WholeM
GutMidF
GutMidM
SalivF
SalivM
GutHind
GutAnt
GutCaeca
GutPost
Blood3h
BloodNBF
BloodTwoNBF
FatBodyBF
GutMidBF

| | GO.ID | MFCluster: skyblue4 Size: 21 | Annotated | Significant | Expected | Rank in ClassicF | Weight01F | ClassicF |
|---|---|---|---|---|---|---|---|---|
| *1* | GO:0030545 | receptor regulator activity | 35 | 2 | 0.08 | 1 | 0.0045 | 0.003 |

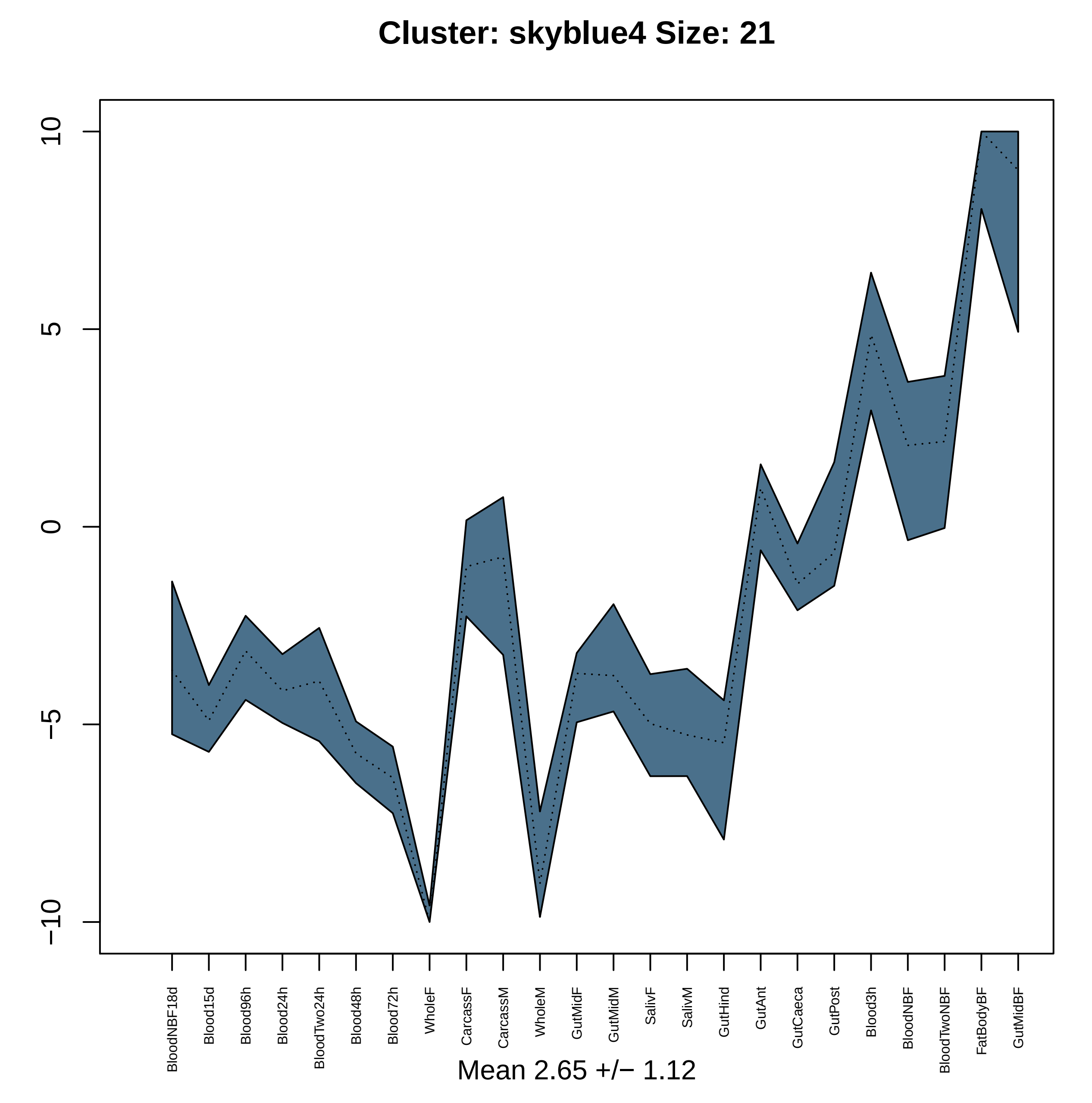

Cluster: skyblue4 Size: 21
10
5
0
−5
−10
BloodNBF18d
Blood15d
Blood96h
Blood24h
BloodTwo24h
Blood48h
Blood72h
WholeF
CarcassF
CarcassM
WholeM
GutMidF
GutMidM
SalivF
SalivM
GutHind
GutAnt
GutCaeca
GutPost
Blood3h
BloodNBF
BloodTwoNBF
FatBodyBF
GutMidBF
Mean 2.65 +/− 1.12

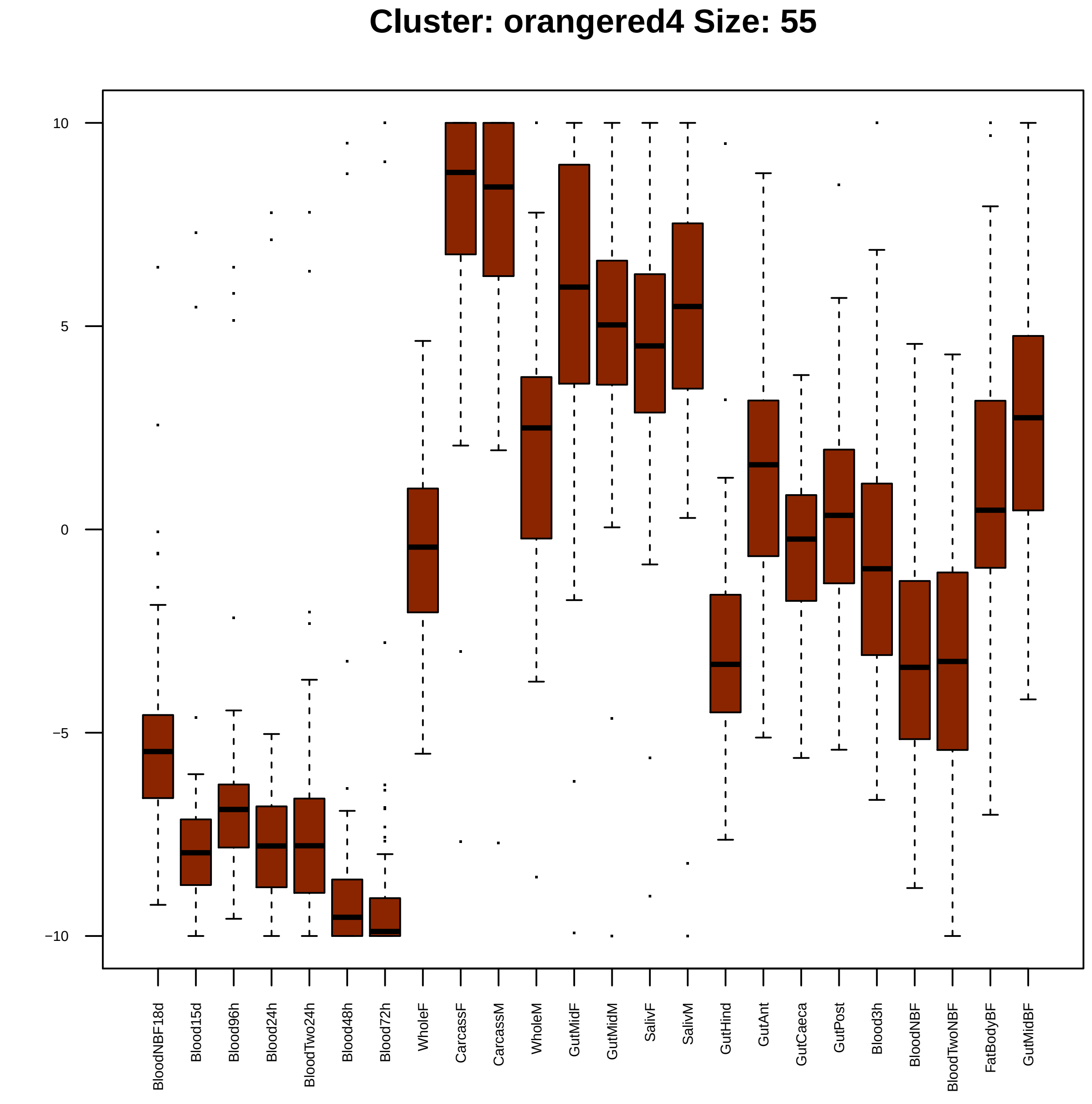

Cluster: orangered4 Size: 55
10
5
0
−5
−10
BloodNBF18d
Blood15d
Blood96h
Blood24h
BloodTwo24h
Blood48h
Blood72h
WholeF
CarcassF
CarcassM
WholeM
GutMidF
GutMidM
SalivF
SalivM
GutHind
GutAnt
GutCaeca
GutPost
Blood3h
BloodNBF
BloodTwoNBF
FatBodyBF
GutMidBF

| | GO.ID | BPCluster: orangered4 Size: 55 | Annotated | Significant | Expected | Rank in ClassicF | Weight01F | ClassicF |
|---|---|---|---|---|---|---|---|---|
| *1* | GO:0007365 | periodic partitioning | 23 | 3 | 0.14 | 1 | 0.00049 | 0.00032 |
| *2* | GO:0035287 | head segmentation | 11 | 2 | 0.07 | 2 | 0.00184 | 0.00184 |
| *4* | GO:0006355 | regulation of transcription, DNA−templat... | 501 | 8 | 2.99 | 5 | 0.01018 | 0.00817 |

| | GO.ID | MFCluster: orangered4 Size: 55 | Annotated | Significant | Expected | Rank in ClassicF | Weight01F | ClassicF |
|---|---|---|---|---|---|---|---|---|
| 1 | GO:0003700 | DNA−binding transcription factor activit... | 282 | 8 | 1.56 | 2 | 0.00071 | 0.00013 |
| 2 | GO:0043565 | sequence−specific DNA binding | 212 | 8 | 1.18 | 1 | 0.00143 | 1.7e−05 |
| 3 | GO:0000976 | transcription regulatory region sequence... | 45 | 3 | 0.25 | 9 | 0.00416 | 0.00191 |
| 12 | GO:0003690 | double−stranded DNA binding | 60 | 4 | 0.33 | 3 | 0.06032 | 0.00032 |
| 24 | GO:0005261 | cation channel activity | 73 | 3 | 0.41 | 16 | 0.21714 | 0.00753 |

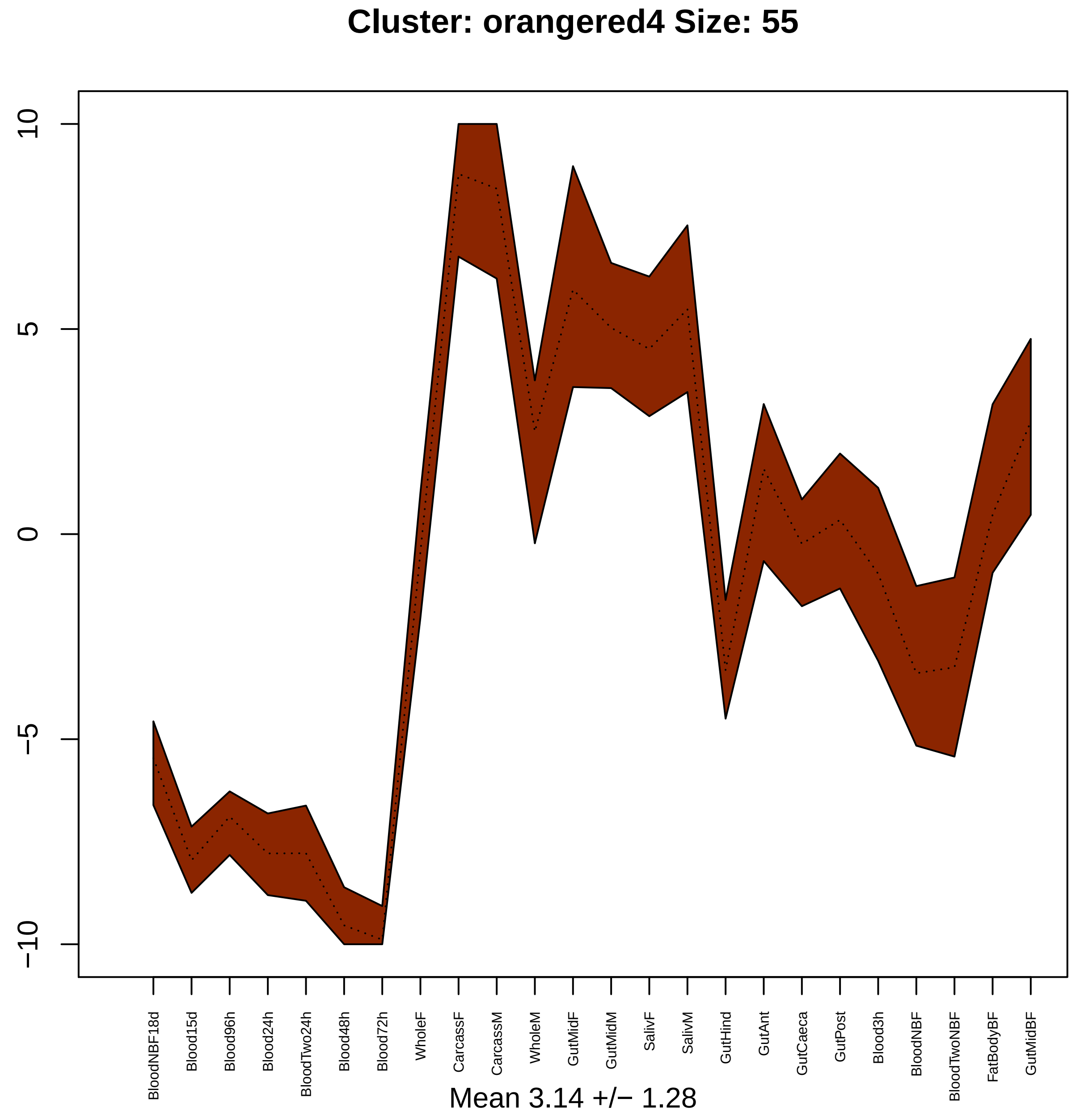

Cluster: orangered4 Size: 55
10
5
0
−5
−10
BloodNBF18d
Blood15d
Blood96h
Blood24h
BloodTwo24h
Blood48h
Blood72h
WholeF
CarcassF
CarcassM
WholeM
GutMidF
GutMidM
SalivF
SalivM
GutHind
GutAnt
GutCaeca
GutPost
Blood3h
BloodNBF
BloodTwoNBF
FatBodyBF
GutMidBF
Mean 3.14 +/− 1.28

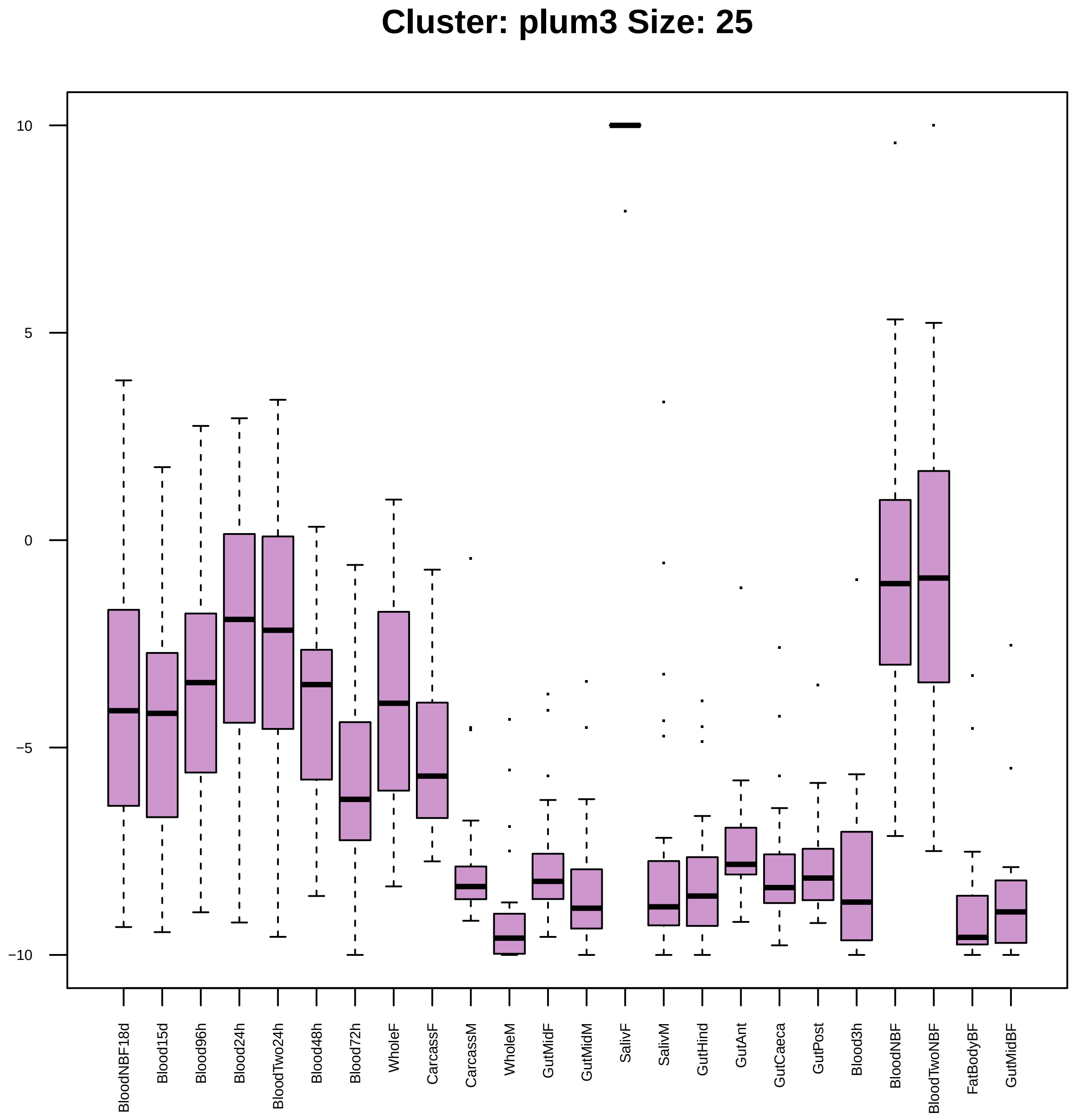
Cluster: plum3 Size: 25
10
5
0
−5
−10
BloodNBF18d
Blood15d
Blood96h
Blood24h
BloodTwo24h
Blood48h
Blood72h
WholeF
CarcassF
CarcassM
WholeM
GutMidF
GutMidM
SalivF
SalivM
GutHind
GutAnt
GutCaeca
GutPost
Blood3h
BloodNBF
BloodTwoNBF
FatBodyBF
GutMidBF

| | GO.ID | BPCluster: plum3 Size: 25 | Annotated | Significant | Expected | Rank in ClassicF | Weight01F | ClassicF |
|---|---|---|---|---|---|---|---|---|
| *1* | GO:0009166 | nucleotide catabolic process | 25 | 2 | 0.02 | 2 | 0.00015 | 0.00015 |
| *12* | GO:0006796 | phosphate−containing compound metabolic ... | 642 | 4 | 0.50 | 9 | 0.09994 | 0.00047 |
| *24* | GO:0044248 | cellular catabolic process | 545 | 3 | 0.43 | 12 | 1.00000 | 0.00545 |
| *28* | GO:0055086 | nucleobase−containing small molecule met... | 193 | 3 | 0.15 | 6 | 1.00000 | 0.00026 |
| *30* | GO:0006753 | nucleoside phosphate metabolic process | 171 | 2 | 0.13 | 14 | 1.00000 | 0.00678 |

| | GO.ID | MFCluster: plum3 Size: 25 | Annotated | Significant | Expected | Rank in ClassicF | Weight01F | ClassicF |
|---|---|---|---|---|---|---|---|---|
| 7 | GO:0016788 | hydrolase activity, acting on ester bond... | 269 | 3 | 0.37 | 2 | 0.114 | 0.0051 |

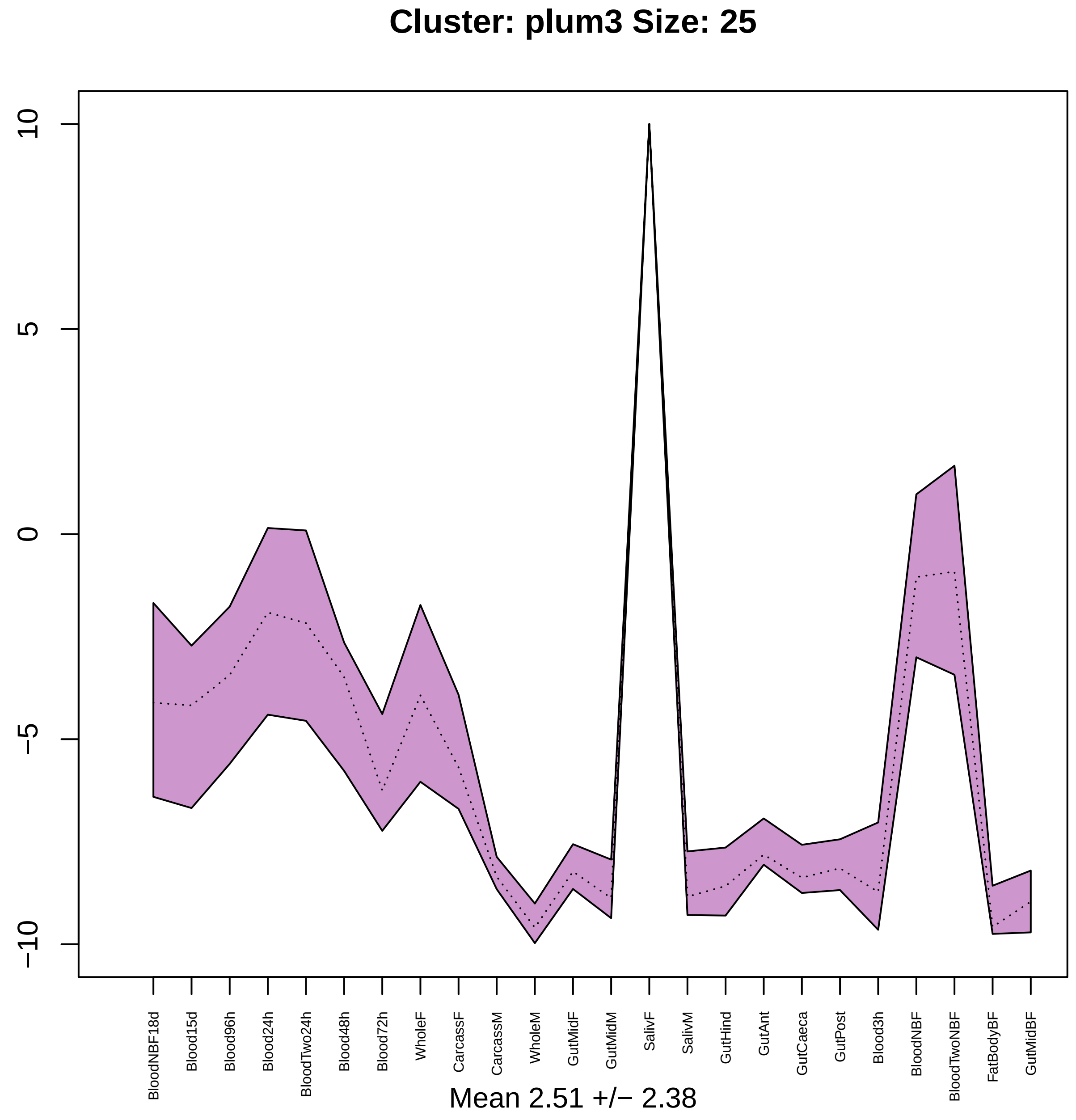
Cluster: plum3 Size: 25
10
5
0
−5
−10
BloodNBF18d
Blood15d
Blood96h
Blood24h
BloodTwo24h
Blood48h
Blood72h
WholeF
CarcassF
CarcassM
WholeM
GutMidF
GutMidM
SalivF
SalivM
GutHind
GutAnt
GutCaeca
GutPost
Blood3h
BloodNBF
BloodTwoNBF
FatBodyBF
GutMidBF
Mean 2.51 +/− 2.38

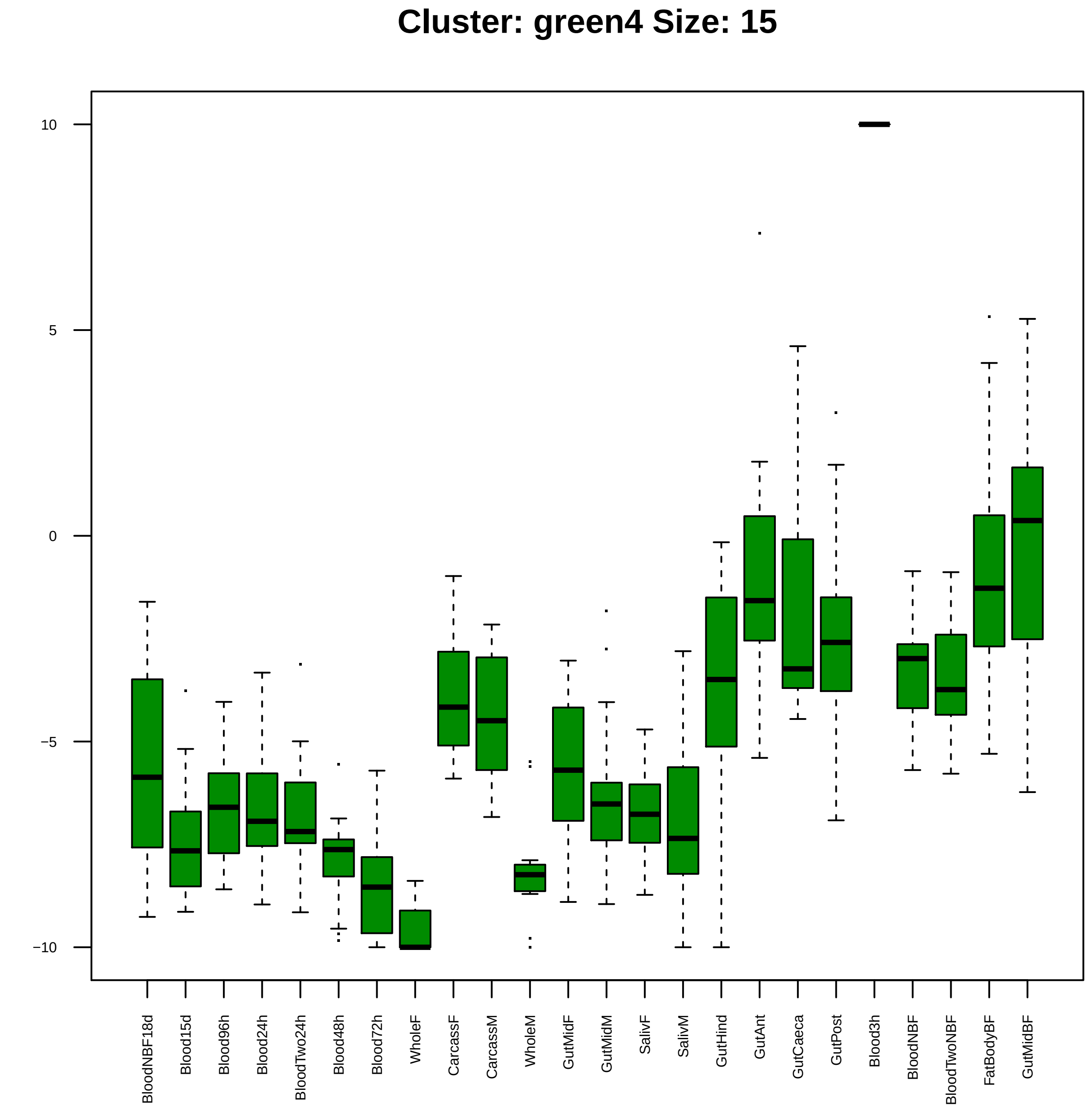

Cluster: green4 Size: 15
10
5
0
−5
−10
BloodNBF18d
Blood15d
Blood96h
Blood24h
BloodTwo24h
Blood48h
Blood72h
WholeF
CarcassF
CarcassM
WholeM
GutMidF
GutMidM
SalivF
SalivM
GutHind
GutAnt
GutCaeca
GutPost
Blood3h
BloodNBF
BloodTwoNBF
FatBodyBF
GutMidBF

| | GO.ID | MFCluster: green4 Size: 15 | Annotated | Significant | Expected | Rank in ClassicF | Weight01F | ClassicF |
|---|---|---|---|---|---|---|---|---|
| *1* | GO:0042302 | structural constituent of cuticle | 99 | 3 | 0.10 | 1 | 0.0017 | 8.4e−05 |
| *10* | GO:0005198 | structural molecule activity | 318 | 3 | 0.31 | 2 | 1.0000 | 0.0026 |

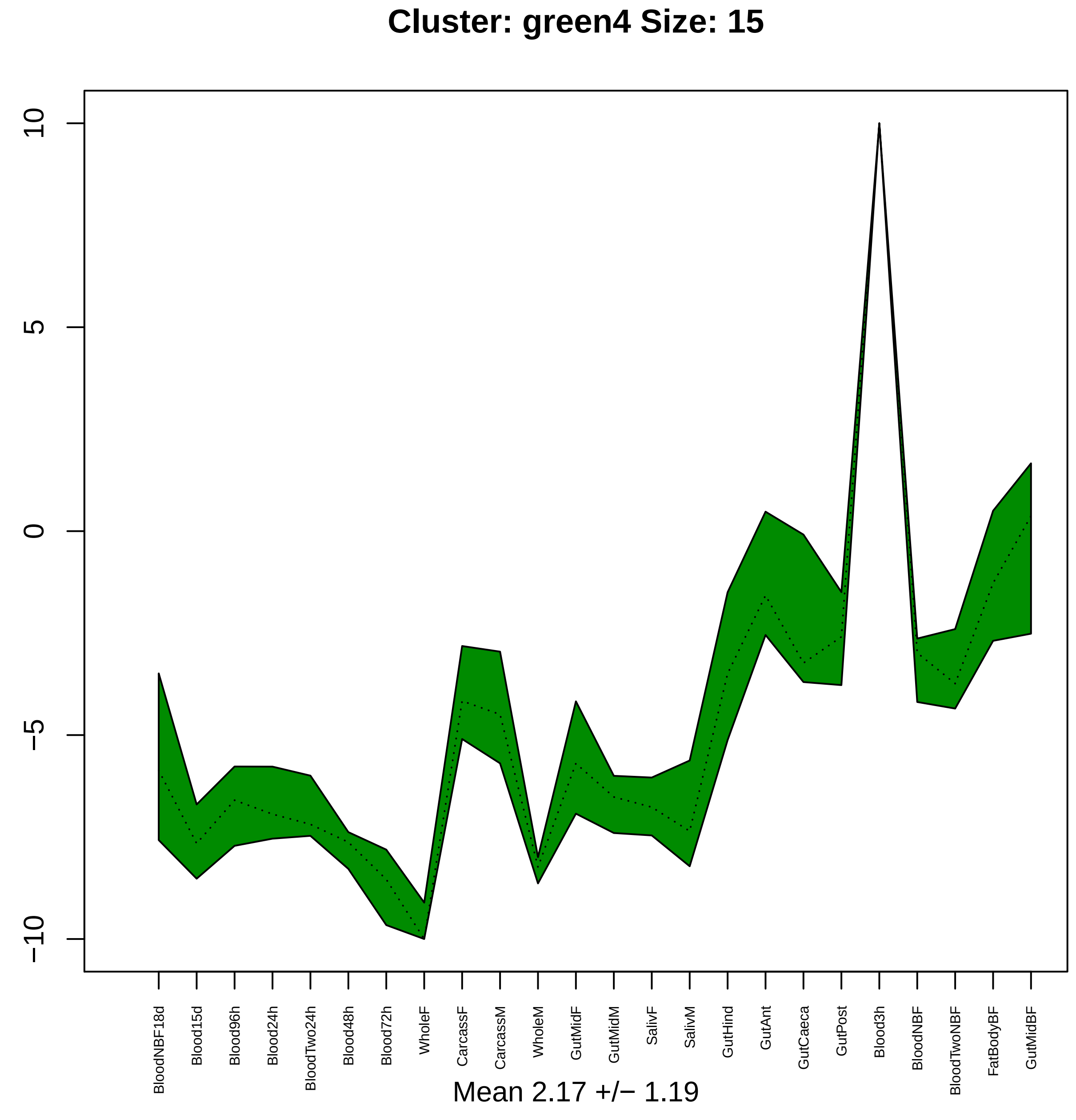

Cluster: green4 Size: 15
10
5
0
−5
−10
BloodNBF18d
Blood15d
Blood96h
Blood24h
BloodTwo24h
Blood48h
Blood72h
WholeF
CarcassF
CarcassM
WholeM
GutMidF
GutMidM
SalivF
SalivM
GutHind
GutAnt
GutCaeca
GutPost
Blood3h
BloodNBF
BloodTwoNBF
FatBodyBF
GutMidBF
Mean 2.17 +/− 1.19

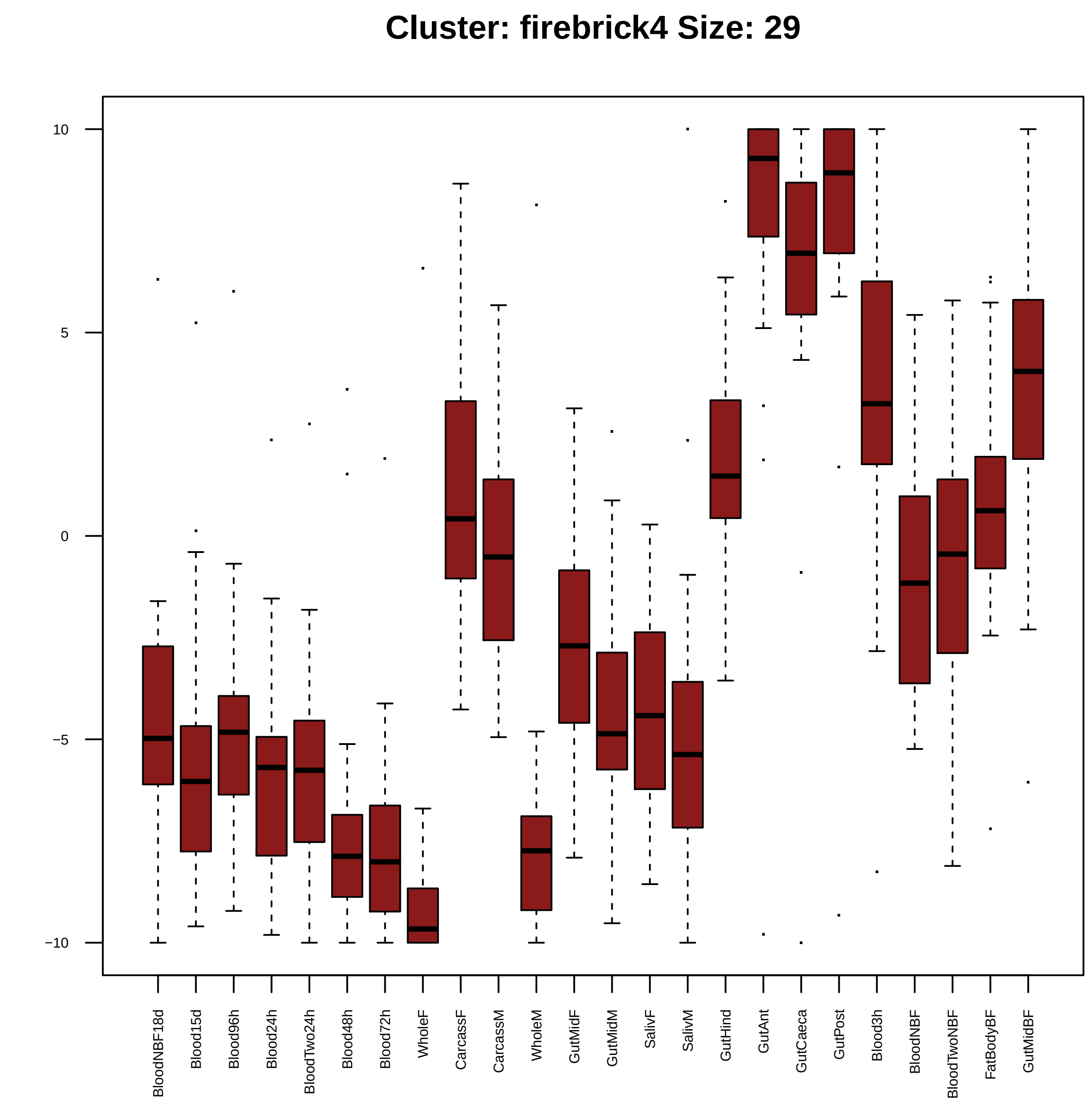
Cluster: firebrick4 Size: 29
10
5
0
−5
−10
BloodNBF18d
Blood15d
Blood96h
Blood24h
BloodTwo24h
Blood48h
Blood72h
WholeF
CarcassF
CarcassM
WholeM
GutMidF
GutMidM
SalivF
SalivM
GutHind
GutAnt
GutCaeca
GutPost
Blood3h
BloodNBF
BloodTwoNBF
FatBodyBF
GutMidBF

| | GO.ID | BPCluster: firebrick4 Size: 29 | Annotated | Significant | Expected | Rank in ClassicF | Weight01F | ClassicF |
|---|---|---|---|---|---|---|---|---|
| 1 | GO:0007186 | G protein−coupled receptor signaling pat... | 228 | 4 | 0.43 | 1 | 0.004 | 0.00063 |

| | GO.ID | MFCluster: firebrick4 Size: 29 | Annotated | Significant | Expected | Rank in ClassicF | Weight01F | ClassicF |
|---|---|---|---|---|---|---|---|---|
| *1* | GO:0004930 | G protein−coupled receptor activity | 171 | 4 | 0.43 | 1 | 0.0046 | 0.00072 |
| *29* | GO:0038023 | signaling receptor activity | 282 | 4 | 0.70 | 3 | 1.0000 | 0.00454 |

#### Cluster: firebrick4 Size: 29

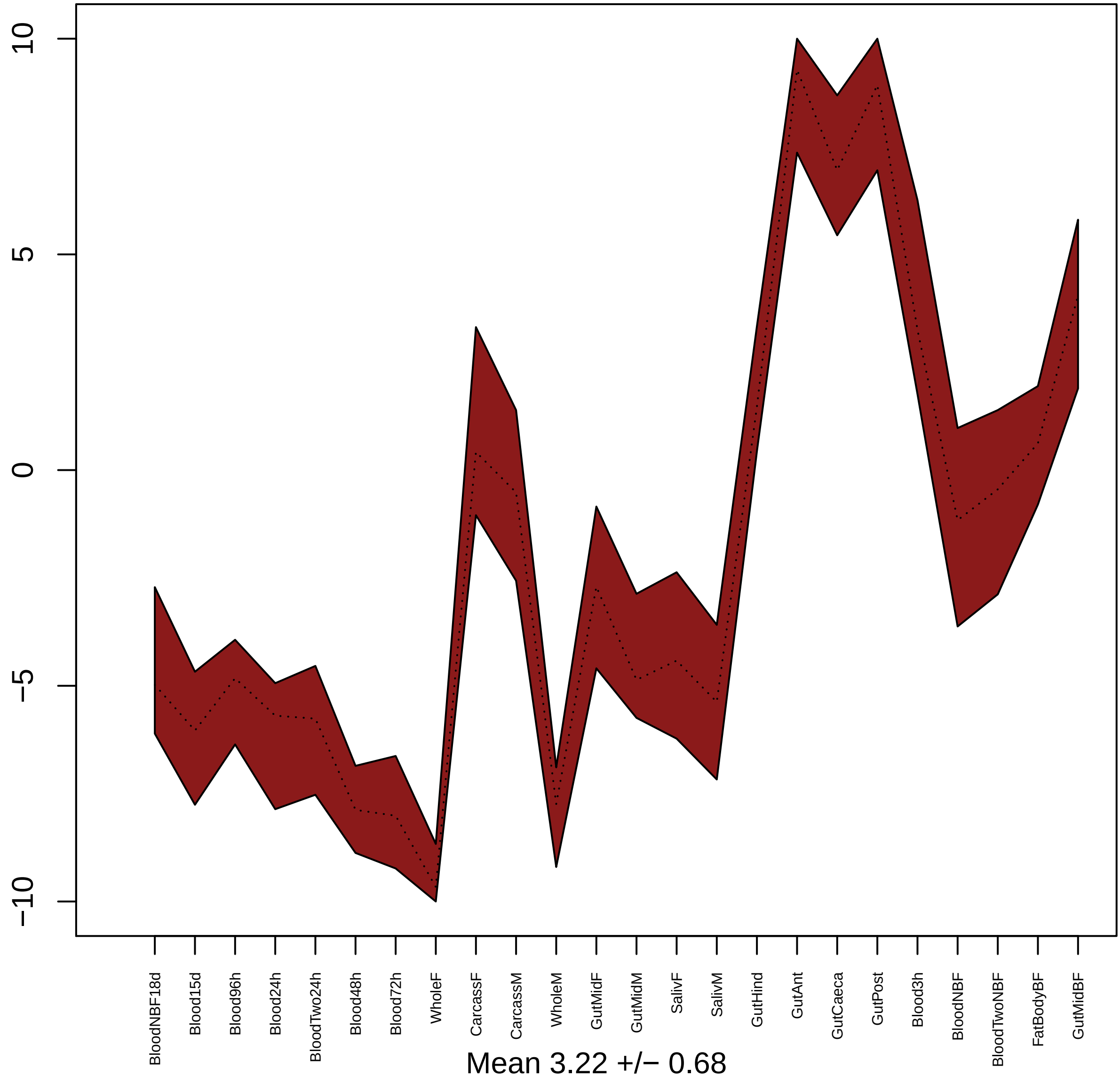


Mean 3.22 +/− 0.68

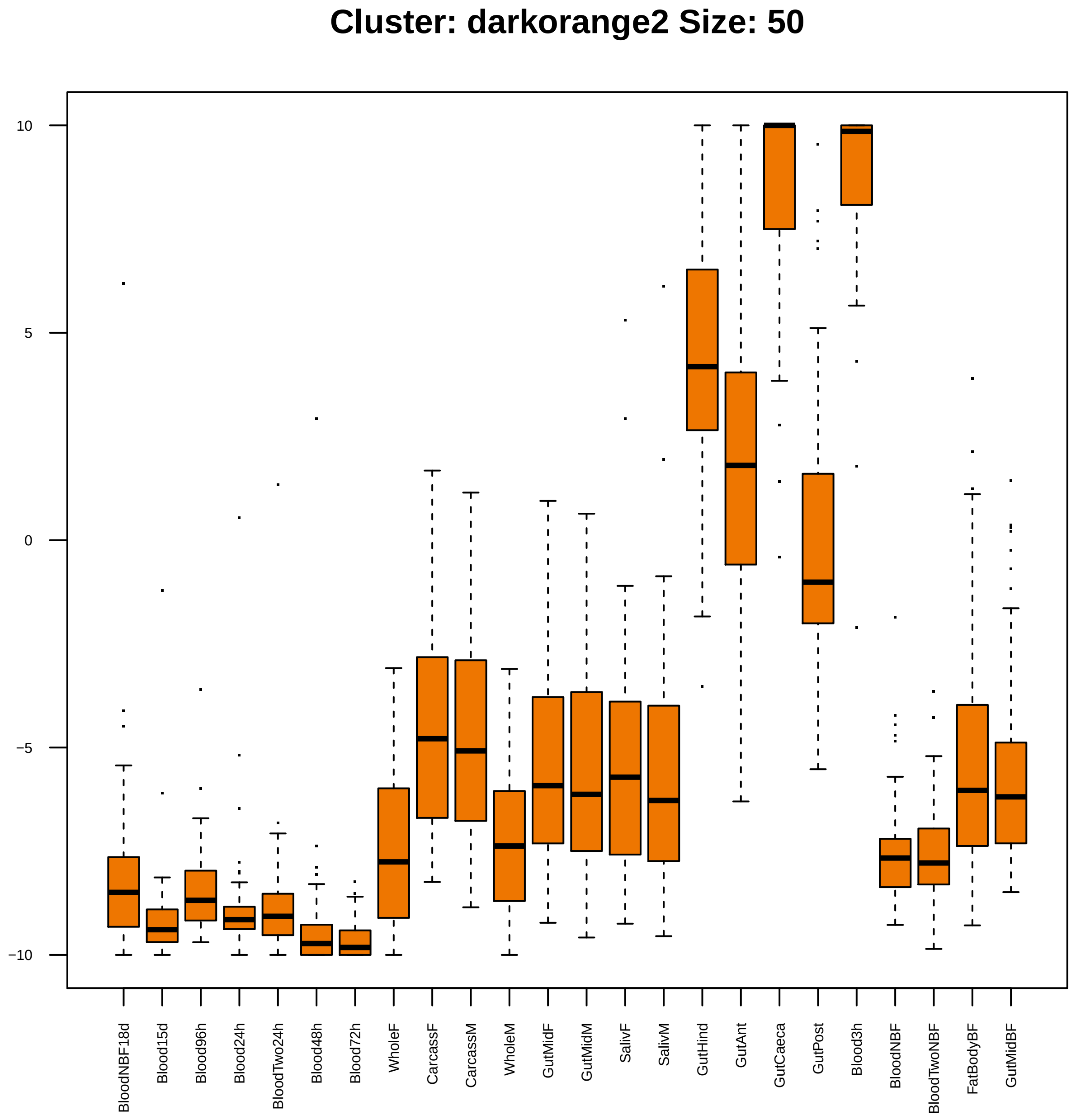

Cluster: darkorange2 Size: 50
10
5
0
−5
−10
BloodNBF18d
Blood15d
Blood96h
Blood24h
BloodTwo24h
Blood48h
Blood72h
WholeF
CarcassF
CarcassM
WholeM
GutMidF
GutMidM
SalivF
SalivM
GutHind
GutAnt
GutCaeca
GutPost
Blood3h
BloodNBF
BloodTwoNBF
FatBodyBF
GutMidBF

| | GO.ID | BPCluster: darkorange2 Size: 50 | Annotated | Significant | Expected | Rank in ClassicF | Weight01F | ClassicF |
|---|---|---|---|---|---|---|---|---|
| 1 | GO:0006979 | response to oxidative stress | 51 | 3 | 0.18 | 1 | 0.00076 | 0.00076 |

| | GO.ID | MFCluster: darkorange2 Size: 50 | Annotated | Significant | Expected | Rank in ClassicF | Weight01F | ClassicF |
|---|---|---|---|---|---|---|---|---|
| *1* | GO:0004601 | peroxidase activity | 26 | 3 | 0.1 | 1 | 0.00012 | 0.00012 |

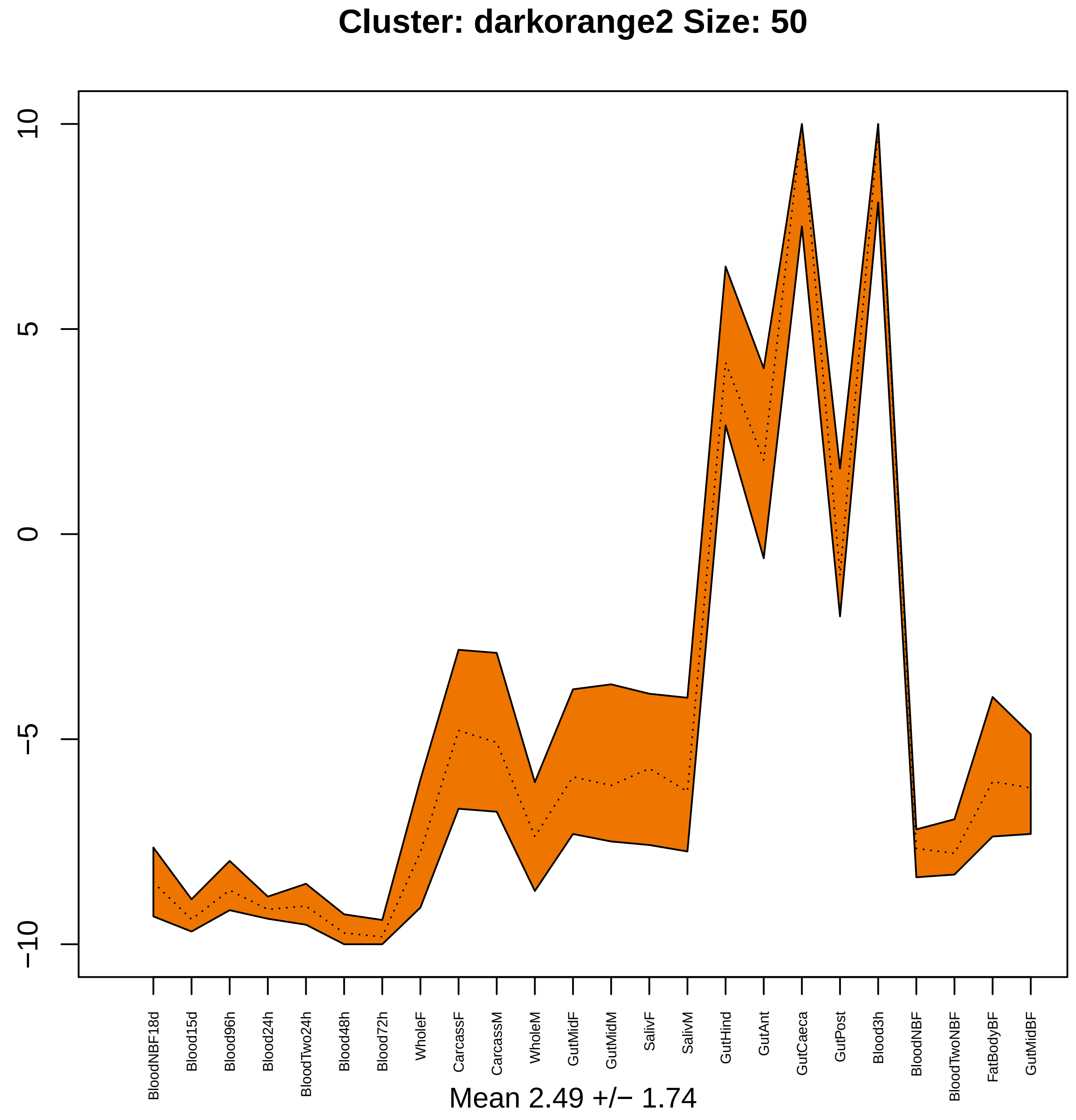

Cluster: darkorange2 Size: 50
10
5
0
−5
−10
BloodNBF18d
Blood15d
Blood96h
Blood24h
BloodTwo24h
Blood48h
Blood72h
WholeF
CarcassF
CarcassM
WholeM
GutMidF
GutMidM
SalivF
SalivM
GutHind
GutAnt
GutCaeca
GutPost
Blood3h
BloodNBF
BloodTwoNBF
FatBodyBF
GutMidBF
Mean 2.49 +/− 1.74

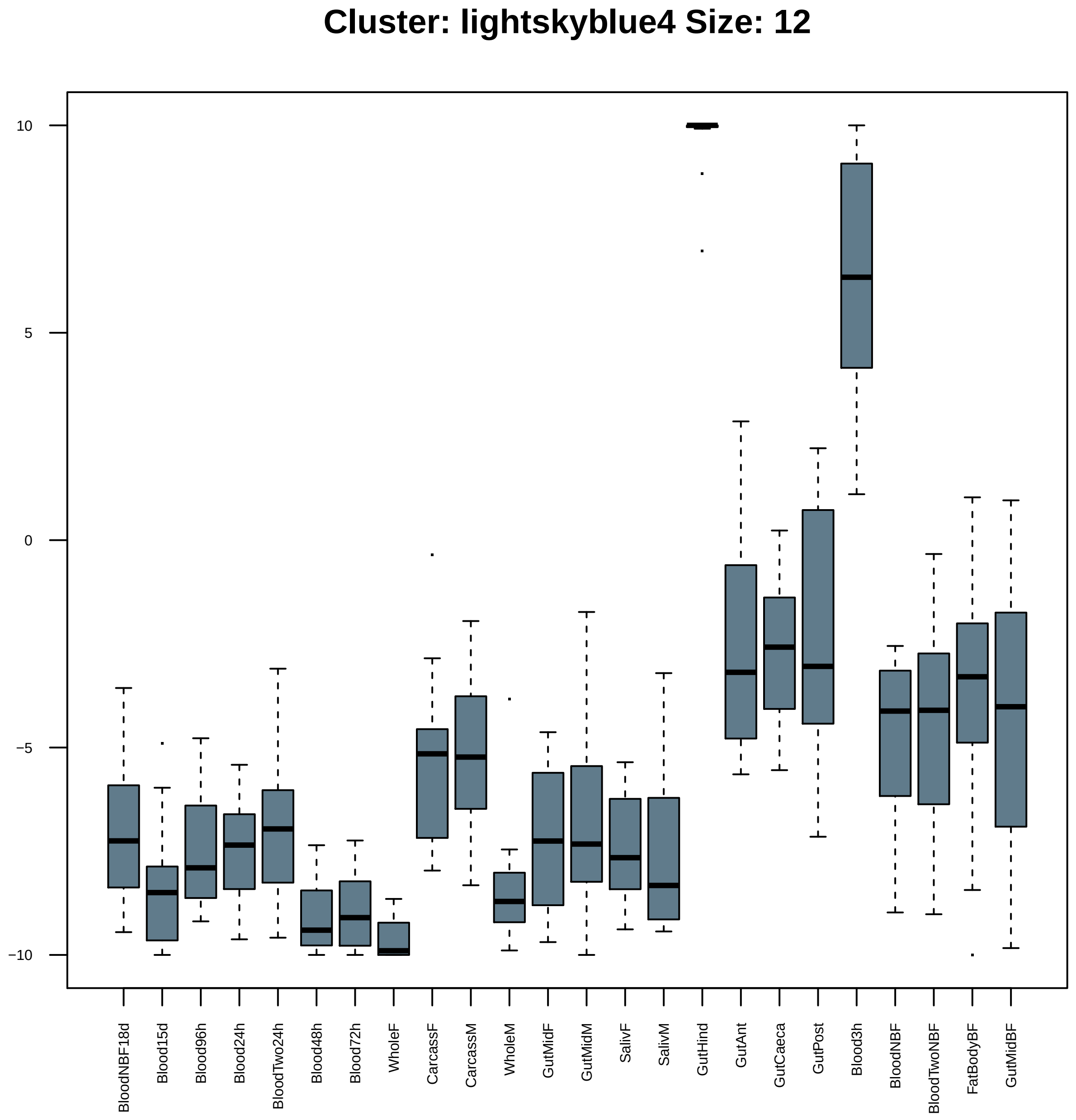
Cluster: lightskyblue4 Size: 12
10
5
0
−5
−10
BloodNBF18d
Blood15d
Blood96h
Blood24h
BloodTwo24h
Blood48h
Blood72h
WholeF
CarcassF
CarcassM
WholeM
GutMidF
GutMidM
SalivF
SalivM
GutHind
GutAnt
GutCaeca
GutPost
Blood3h
BloodNBF
BloodTwoNBF
FatBodyBF
GutMidBF

| | GO.ID | MFCluster: lightskyblue4 Size: 12 | Annotated | Significant | Expected | Rank in ClassicF | Weight01F | ClassicF |
|---|---|---|---|---|---|---|---|---|
| *1* | GO:0042302 | structural constituent of cuticle | 99 | 2 | 0.07 | 1 | 0.0018 | 0.0018 |

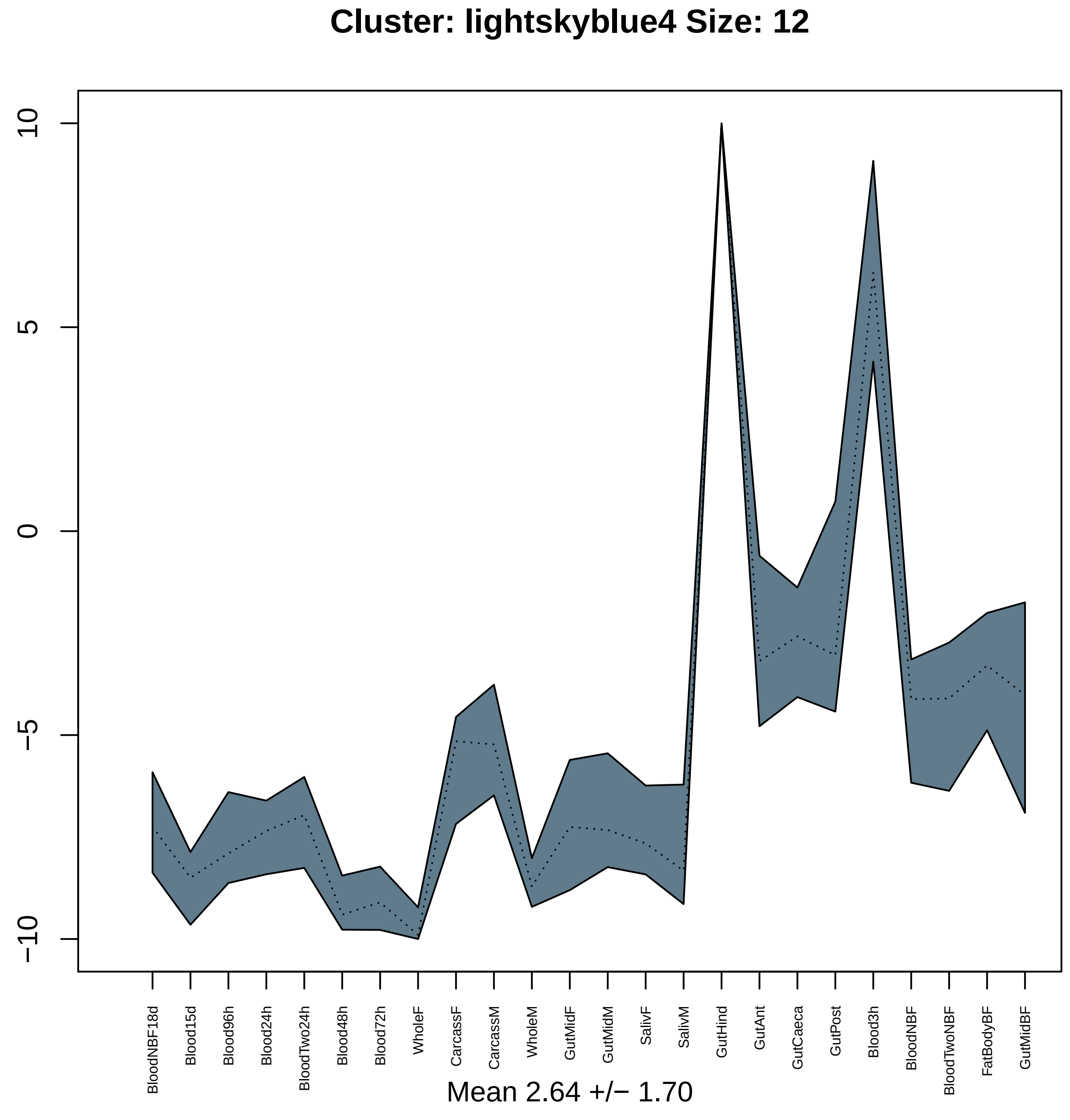

Cluster: lightskyblue4 Size: 12
10
5
0
−5
−10
BloodNBF18d
Blood15d
Blood96h
Blood24h
BloodTwo24h
Blood48h
Blood72h
WholeF
CarcassF
CarcassM
WholeM
GutMidF
GutMidM
SalivF
SalivM
GutHind
GutAnt
GutCaeca
GutPost
Blood3h
BloodNBF
BloodTwoNBF
FatBodyBF
GutMidBF
Mean 2.64 +/− 1.70

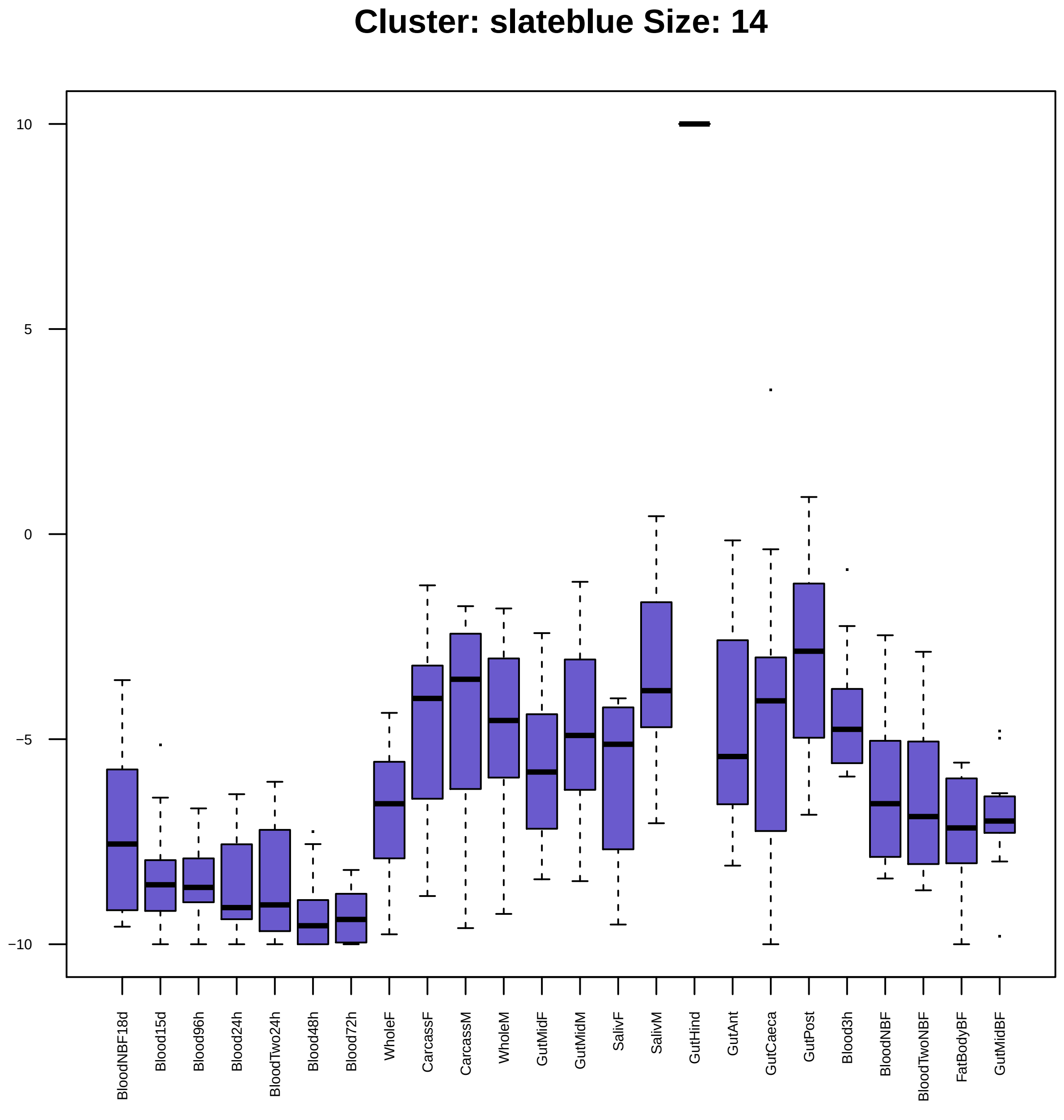

Cluster: slateblue Size: 14
10
5
0
−5
−10
BloodNBF18d
Blood15d
Blood96h
Blood24h
BloodTwo24h
Blood48h
Blood72h
WholeF
CarcassF
CarcassM
WholeM
GutMidF
GutMidM
SalivF
SalivM
GutHind
GutAnt
GutCaeca
GutPost
Blood3h
BloodNBF
BloodTwoNBF
FatBodyBF
GutMidBF

| | GO.ID | MFCluster: slateblue Size: 14 | Annotated | Significant | Expected | Rank in ClassicF | Weight01F | ClassicF |
|---|---|---|---|---|---|---|---|---|
| *1* | GO:0004252 | serine−type endopeptidase activity | 252 | 3 | 0.35 | 1 | 0.0042 | 0.0042 |
| *8* | GO:0017171 | serine hydrolase activity | 279 | 3 | 0.39 | 3 | 1.0000 | 0.0056 |
| *19* | GO:0008236 | serine−type peptidase activity | 278 | 3 | 0.39 | 2 | 1.0000 | 0.0056 |

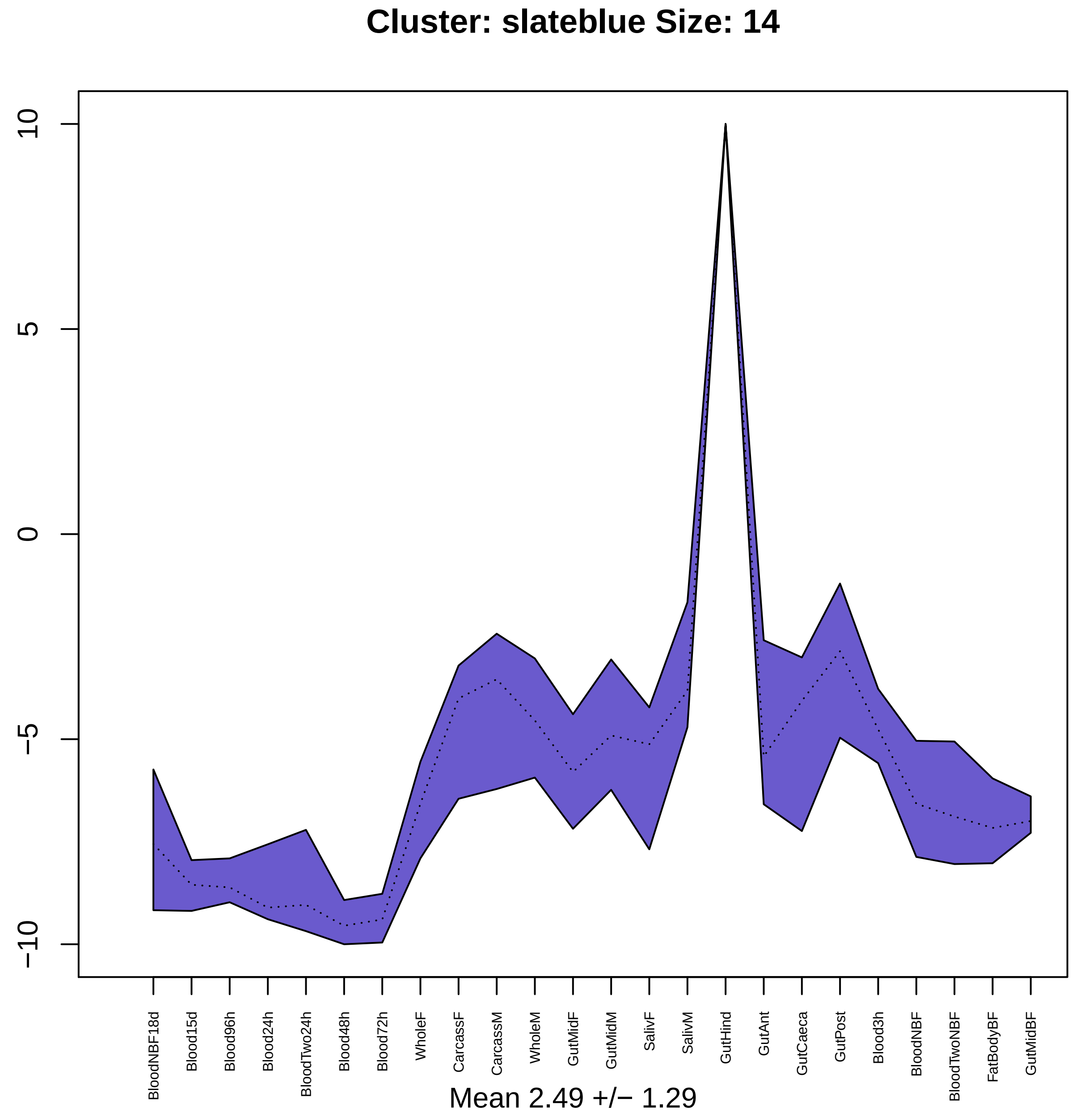
Cluster: slateblue Size: 14
10
5
0
−5
−10
BloodNBF18d
Blood15d
Blood96h
Blood24h
BloodTwo24h
Blood48h
Blood72h
WholeF
CarcassF
CarcassM
WholeM
GutMidF
GutMidM
SalivF
SalivM
GutHind
GutAnt
GutCaeca
GutPost
Blood3h
BloodNBF
BloodTwoNBF
FatBodyBF
GutMidBF
Mean 2.49 +/− 1.29

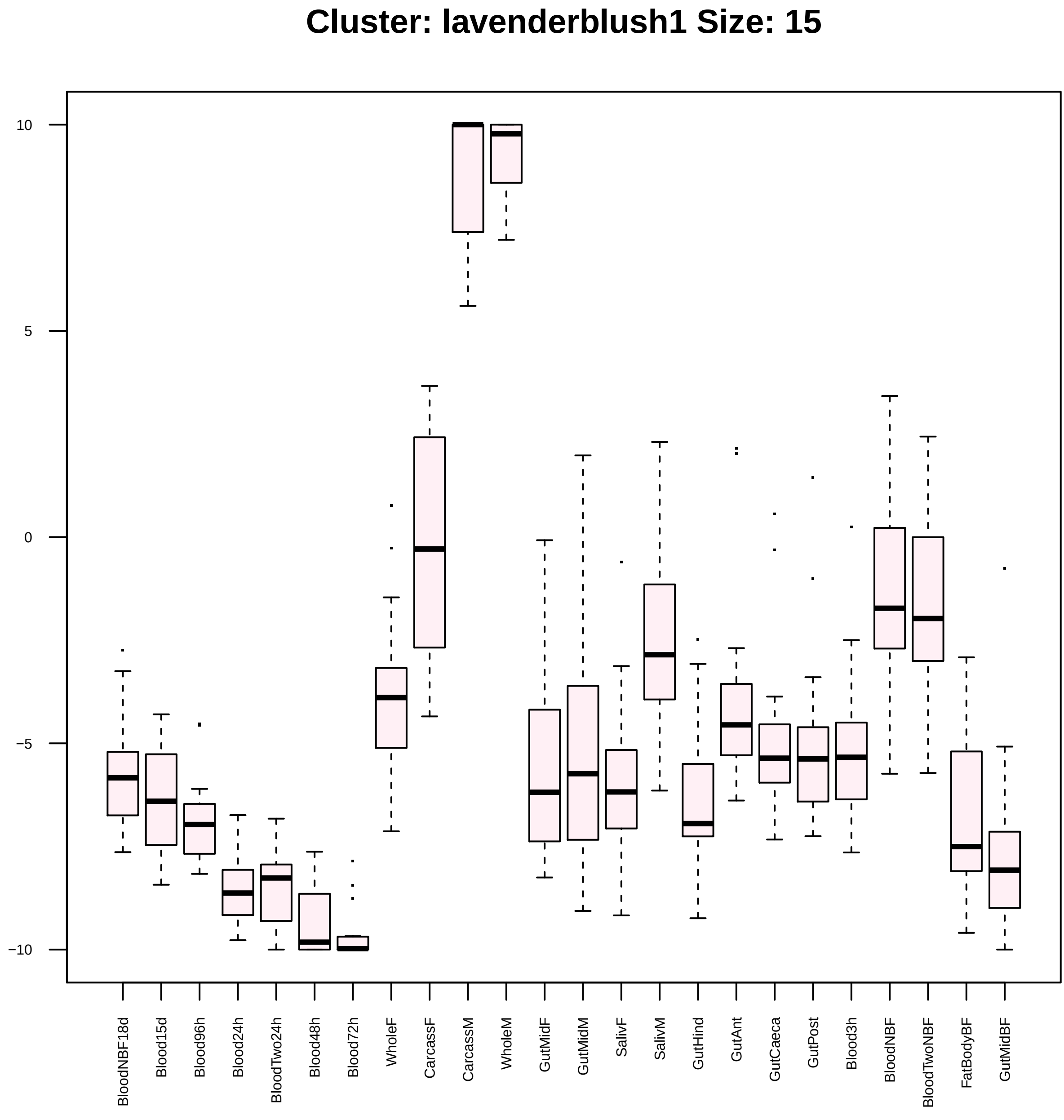
Cluster: lavenderblush1 Size: 15
10
5
0
−5
−10
BloodNBF18d
Blood15d
Blood96h
Blood24h
BloodTwo24h
Blood48h
Blood72h
WholeF
CarcassF
CarcassM
WholeM
GutMidF
GutMidM
SalivF
SalivM
GutHind
GutAnt
GutCaeca
GutPost
Blood3h
BloodNBF
BloodTwoNBF
FatBodyBF
GutMidBF

| | GO.ID | BPCluster: lavenderblush1 Size: 15 | Annotated | Significant | Expected | Rank in ClassicF | Weight01F | ClassicF |
|---|---|---|---|---|---|---|---|---|
| 1 | GO:0007186 | G protein−coupled receptor signaling pat... | 228 | 3 | 0.25 | 1 | 0.0014 | 0.0014 |

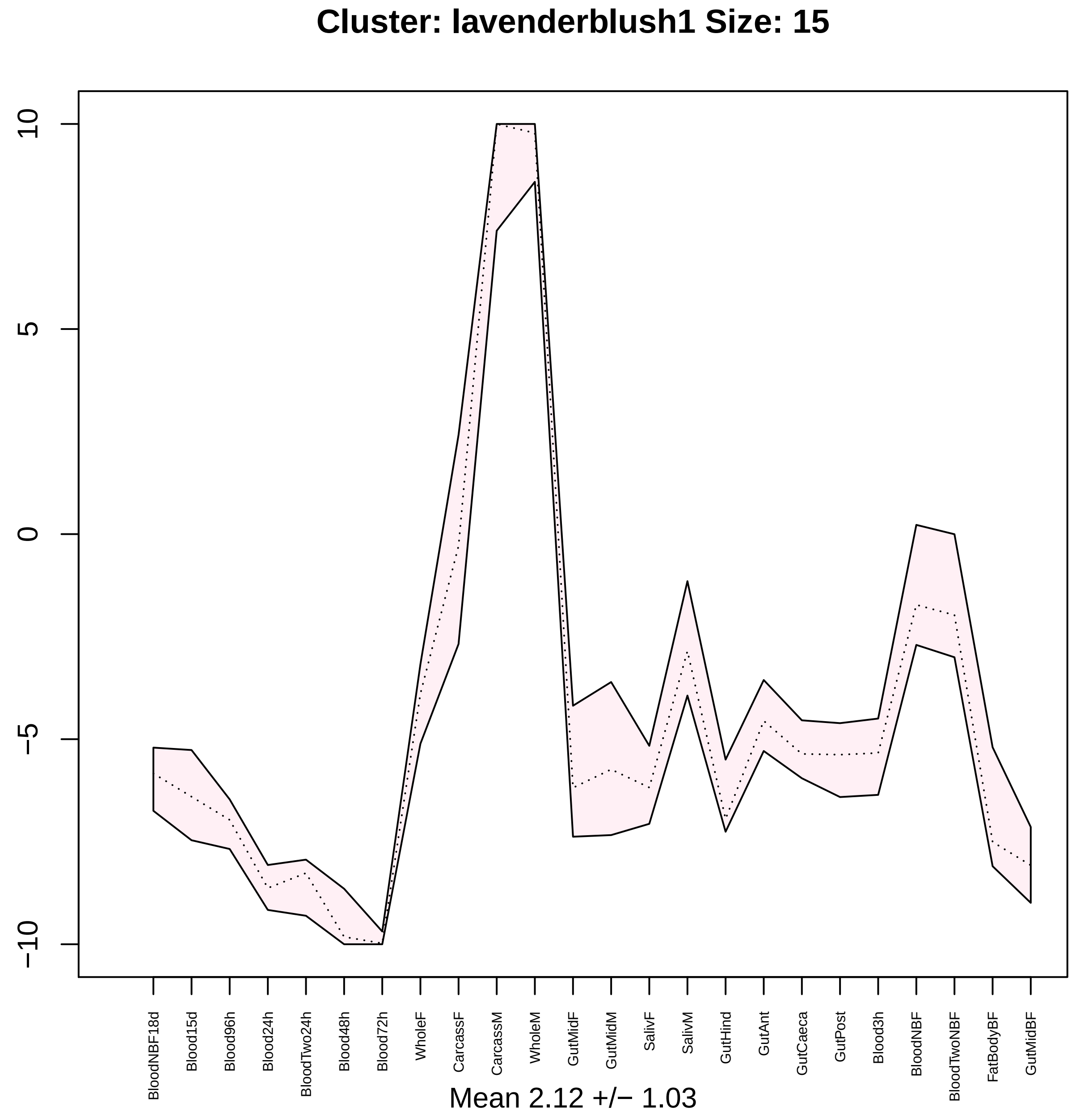
Cluster: lavenderblush1 Size: 15
10
5
0
−5
−10
BloodNBF18d
Blood15d
Blood96h
Blood24h
BloodTwo24h
Blood48h
Blood72h
WholeF
CarcassF
CarcassM
WholeM
GutMidF
GutMidM
SalivF
SalivM
GutHind
GutAnt
GutCaeca
GutPost
Blood3h
BloodNBF
BloodTwoNBF
FatBodyBF
GutMidBF
Mean 2.12 +/− 1.03

### Cluster: yellow3 Size: 21

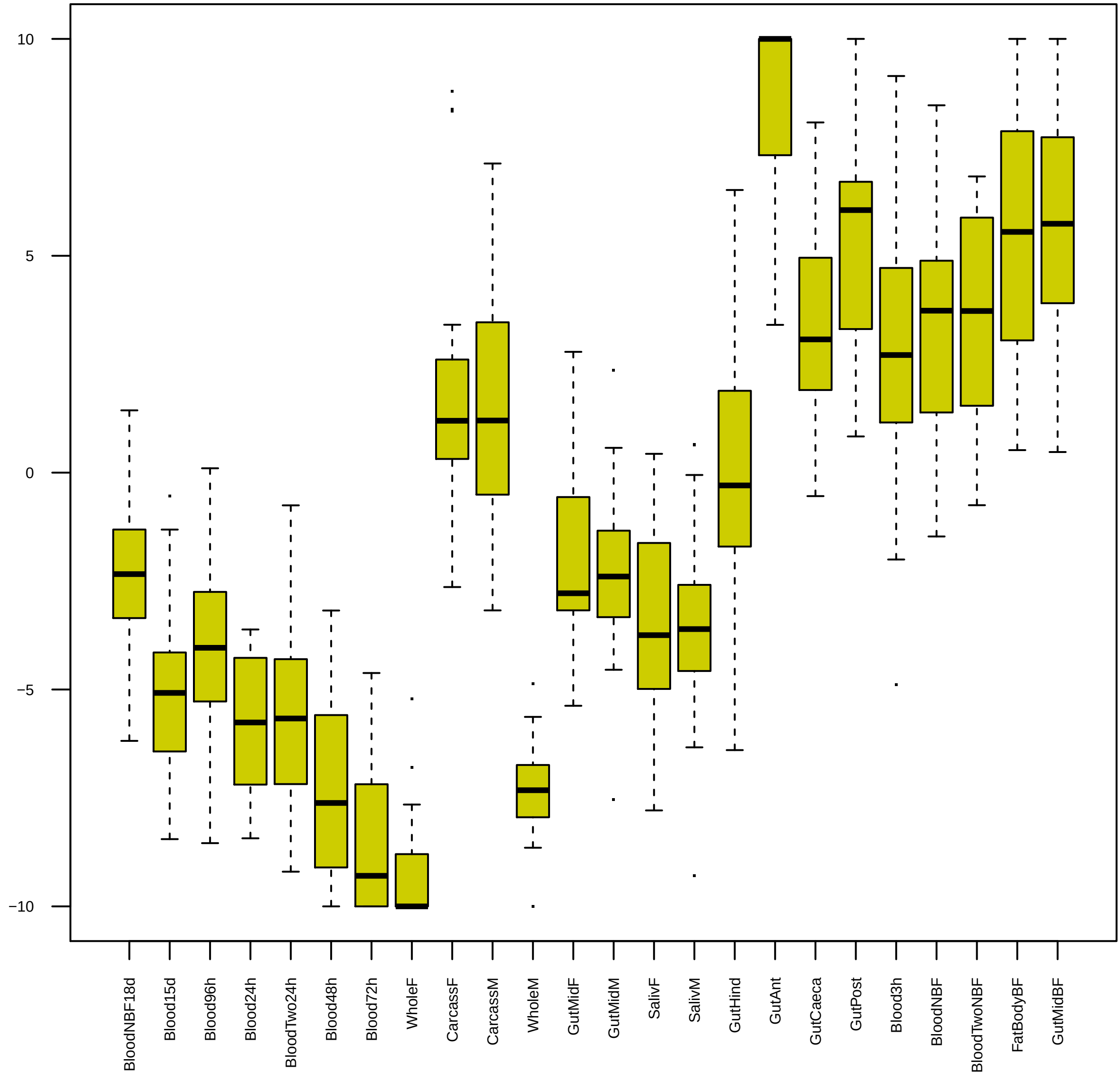

| | GO.ID | BPCluster: yellow3 Size: 21 | Annotated | Significant | Expected | Rank in ClassicF | Weight01F | ClassicF |
|---|---|---|---|---|---|---|---|---|
| *1* | GO:0046068 | cGMP metabolic process | 11 | 2 | 0.02 | 1 | 0.00012 | 0.00012 |
| *2* | GO:0052652 | cyclic purine nucleotide metabolic proce... | 14 | 2 | 0.02 | 2 | 0.00020 | 0.00020 |
| *3* | GO:0009152 | purine ribonucleotide biosynthetic proce... | 67 | 2 | 0.11 | 10 | 0.00463 | 0.00463 |
| *4* | GO:0006468 | protein phosphorylation | 271 | 3 | 0.42 | 15 | 0.00729 | 0.00729 |

| | GO.ID | MFCluster: yellow3 Size: 21 | Annotated | Significant | Expected | Rank in ClassicF | Weight01F | ClassicF |
|---|---|---|---|---|---|---|---|---|
| 1 | GO:0004383 | guanylate cyclase activity | 11 | 2 | 0.03 | 1 | 0.00032 | 0.00032 |
| 2 | GO:0001653 | peptide receptor activity | 26 | 2 | 0.06 | 4 | 0.00473 | 0.00185 |
| 30 | GO:0016849 | phosphorus−oxygen lyase activity | 23 | 2 | 0.06 | 3 | 1.00000 | 0.00144 |

### Cluster: yellow3 Size: 21

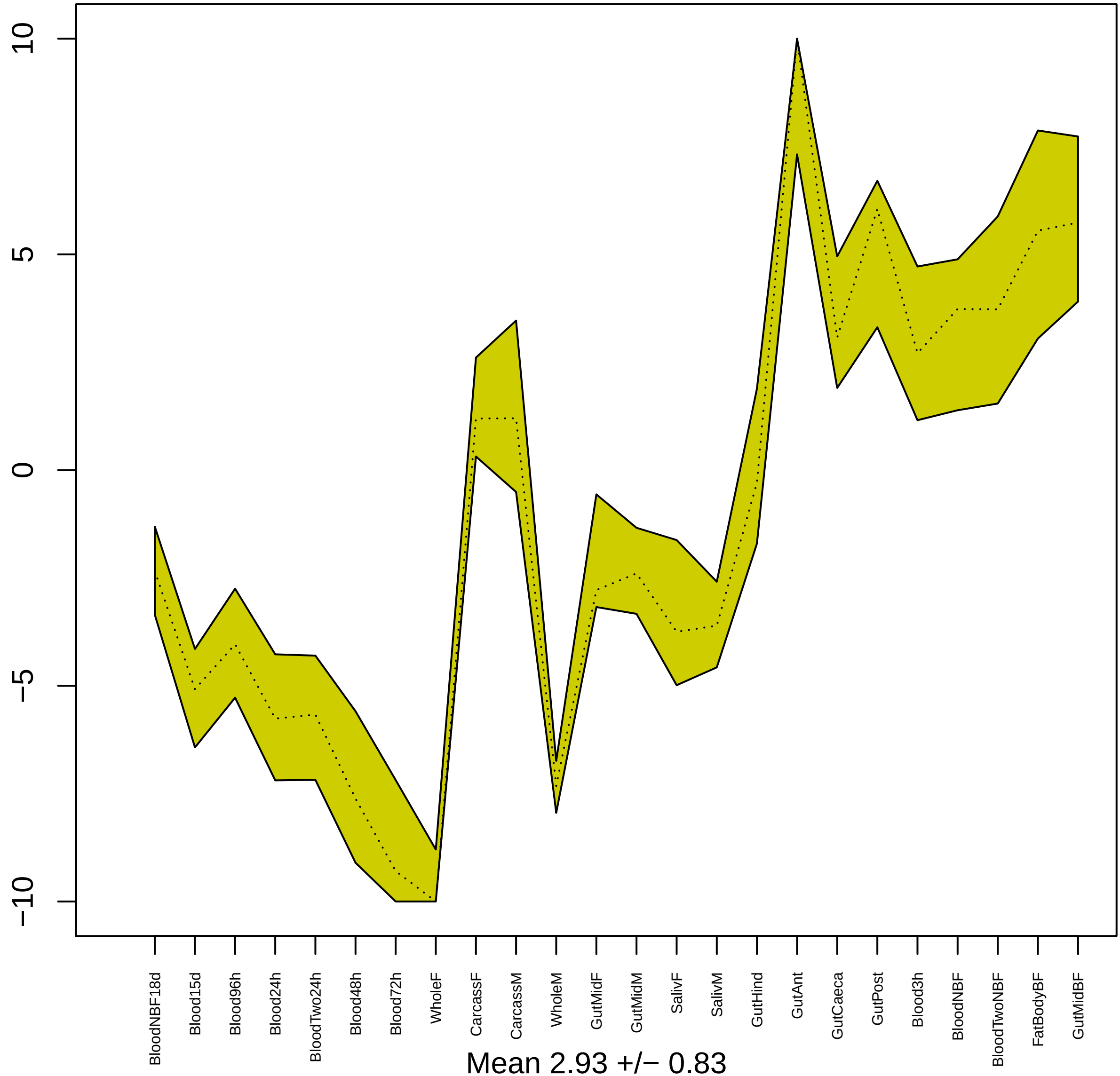


Mean 2.93 +/− 0.83

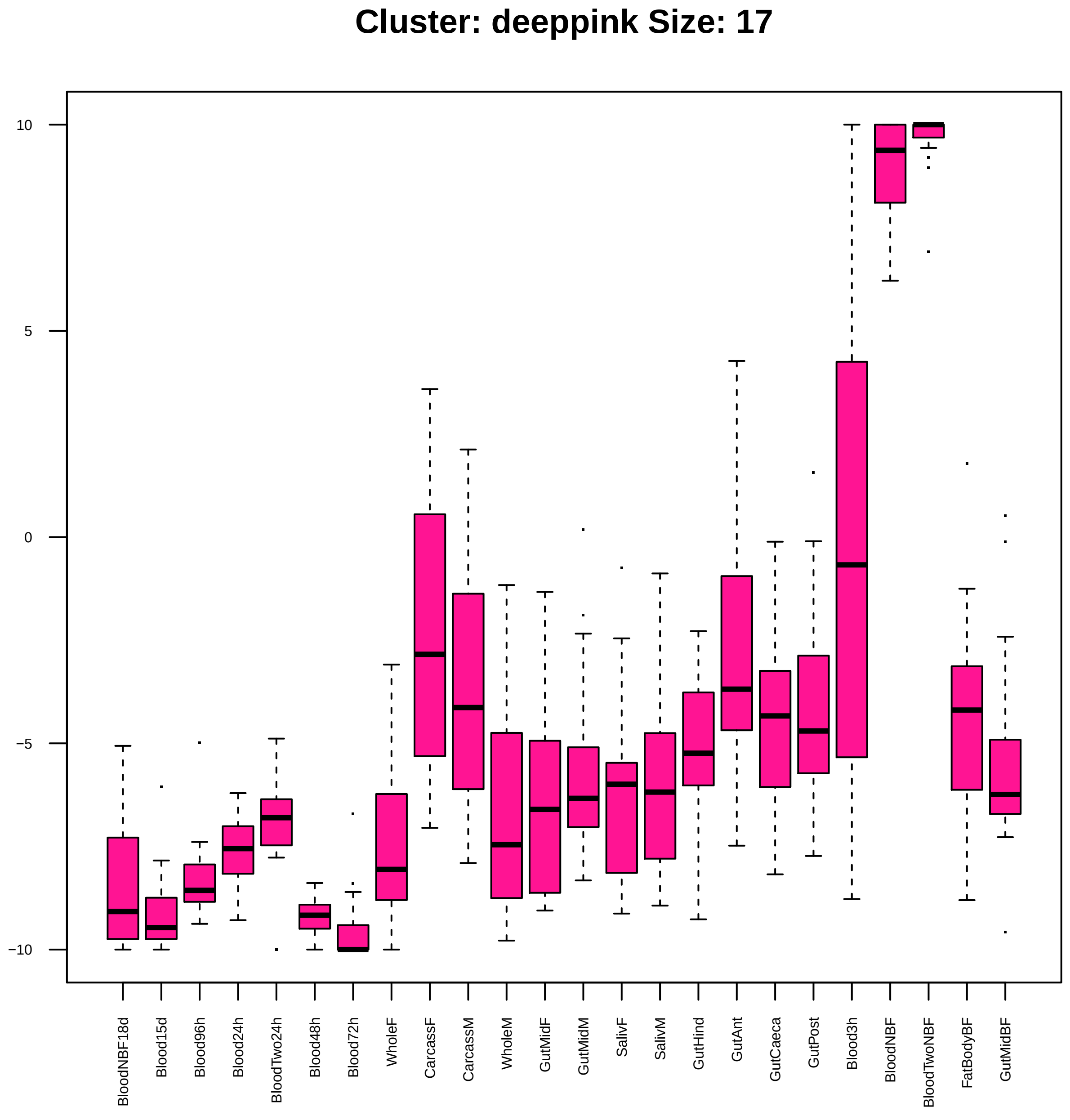
Cluster: deeppink Size: 17
10
5
0
−5
−10
BloodNBF18d
Blood15d
Blood96h
Blood24h
BloodTwo24h
Blood48h
Blood72h
WholeF
CarcassF
CarcassM
WholeM
GutMidF
GutMidM
SalivF
SalivM
GutHind
GutAnt
GutCaeca
GutPost
Blood3h
BloodNBF
BloodTwoNBF
FatBodyBF
GutMidBF

| | GO.ID | BPCluster: deeppink Size: 17 | Annotated | Significant | Expected | Rank in ClassicF | Weight01F | ClassicF |
|---|---|---|---|---|---|---|---|---|
| *1* | GO:0006030 | chitin metabolic process | 95 | 2 | 0.06 | 1 | 0.0013 | 0.0013 |
| *2* | GO:0005975 | carbohydrate metabolic process | 210 | 2 | 0.13 | 5 | 0.0062 | 0.0062 |
| *14* | GO:0017144 | drug metabolic process | 246 | 2 | 0.15 | 6 | 1.0000 | 0.0084 |
| *28* | GO:0006040 | amino sugar metabolic process | 99 | 2 | 0.06 | 2 | 1.0000 | 0.0014 |
| *30* | GO:1901071 | glucosamine−containing compound metaboli... | 99 | 2 | 0.06 | 3 | 1.0000 | 0.0014 |

| | GO.ID | MFCluster: deeppink Size: 17 | Annotated | Significant | Expected | Rank in ClassicF | Weight01F | ClassicF |
|---|---|---|---|---|---|---|---|---|
| *1* | GO:0042302 | structural constituent of cuticle | 99 | 3 | 0.11 | 1 | 0.0023 | 0.00013 |
| *2* | GO:0008061 | chitin binding | 76 | 2 | 0.08 | 3 | 0.0029 | 0.00295 |
| *4* | GO:0016810 | hydrolase activity, acting on carbon−nit... | 56 | 2 | 0.06 | 2 | 0.0232 | 0.00161 |
| *13* | GO:0005198 | structural molecule activity | 318 | 3 | 0.35 | 4 | 1.0000 | 0.00404 |

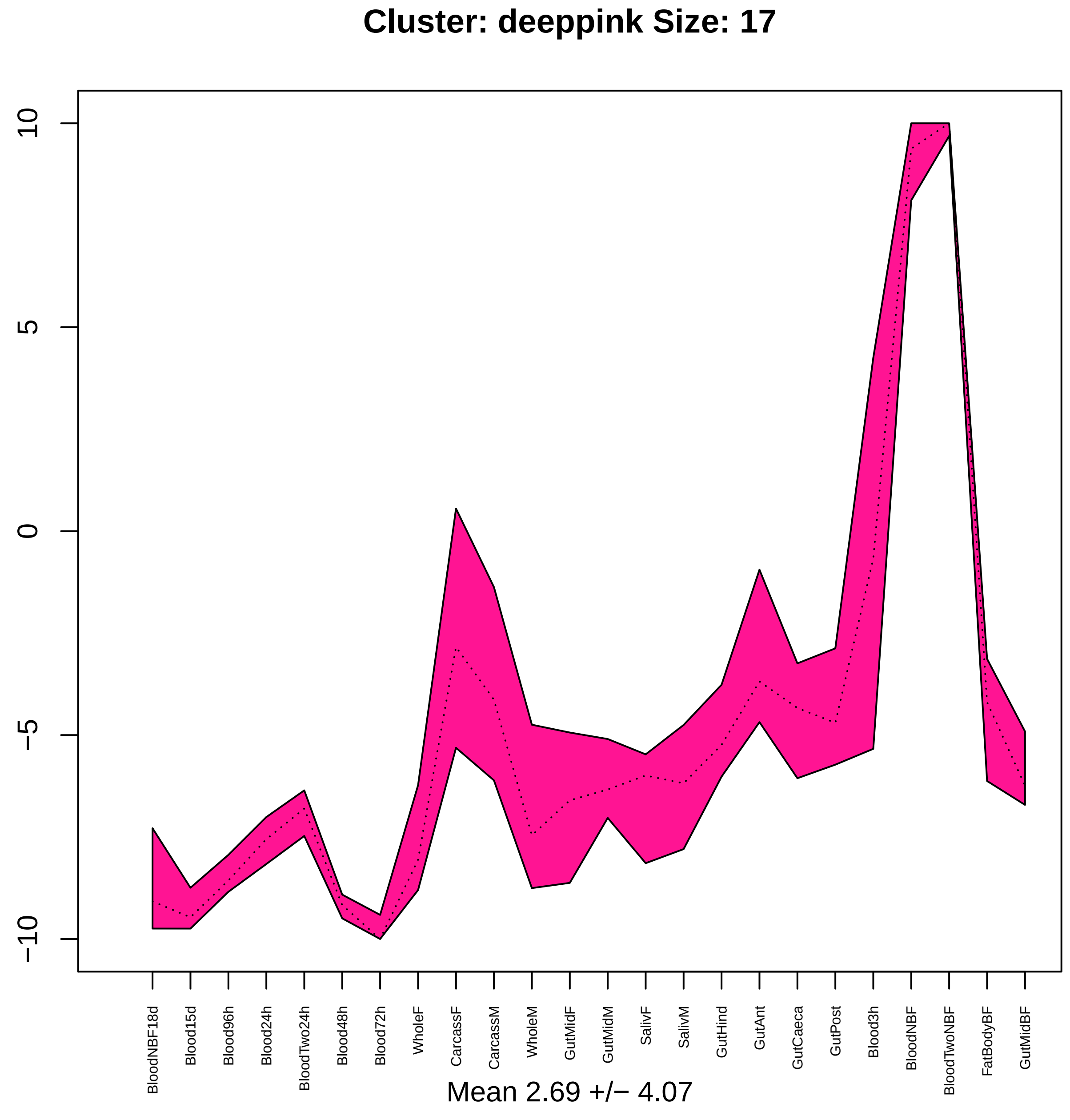
Cluster: deeppink Size: 17
10
5
0
−5
−10
BloodNBF18d
Blood15d
Blood96h
Blood24h
BloodTwo24h
Blood48h
Blood72h
WholeF
CarcassF
CarcassM
WholeM
GutMidF
GutMidM
SalivF
SalivM
GutHind
GutAnt
GutCaeca
GutPost
Blood3h
BloodNBF
BloodTwoNBF
FatBodyBF
GutMidBF
Mean 2.69 +/− 4.07

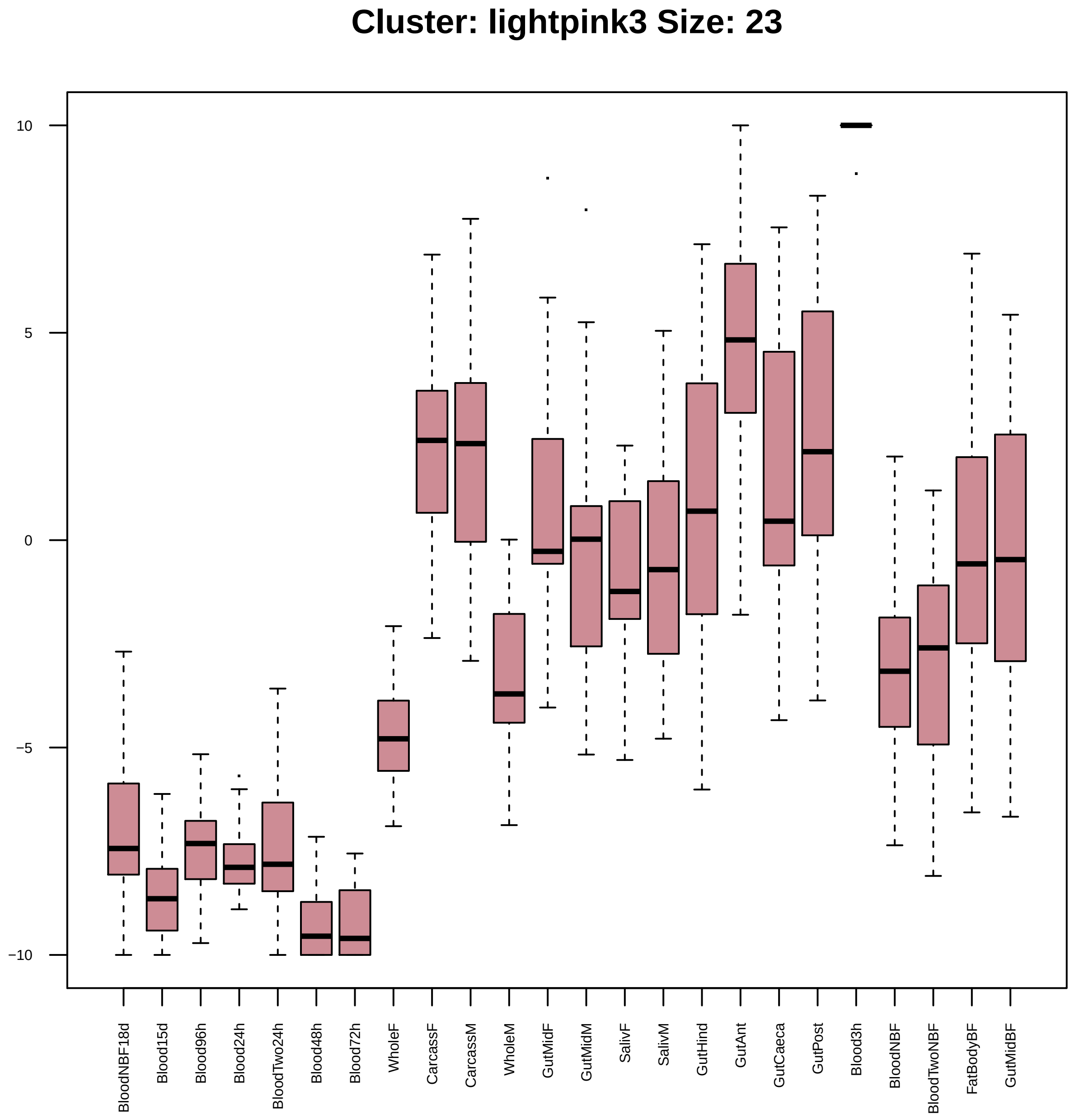

Cluster: lightpink3 Size: 23
10
5
0
−5
−10
BloodNBF18d
Blood15d
Blood96h
Blood24h
BloodTwo24h
Blood48h
Blood72h
WholeF
CarcassF
CarcassM
WholeM
GutMidF
GutMidM
SalivF
SalivM
GutHind
GutAnt
GutCaeca
GutPost
Blood3h
BloodNBF
BloodTwoNBF
FatBodyBF
GutMidBF

| | GO.ID | BPCluster: lightpink3 Size: 23 | Annotated | Significant | Expected | Rank in ClassicF | Weight01F | ClassicF |
|---|---|---|---|---|---|---|---|---|
| *1* | GO:0035159 | regulation of tube length, open tracheal... | 21 | 2 | 0.03 | 1 | 0.00029 | 0.00029 |
| *2* | GO:0006030 | chitin metabolic process | 95 | 2 | 0.12 | 6 | 0.00580 | 0.00580 |

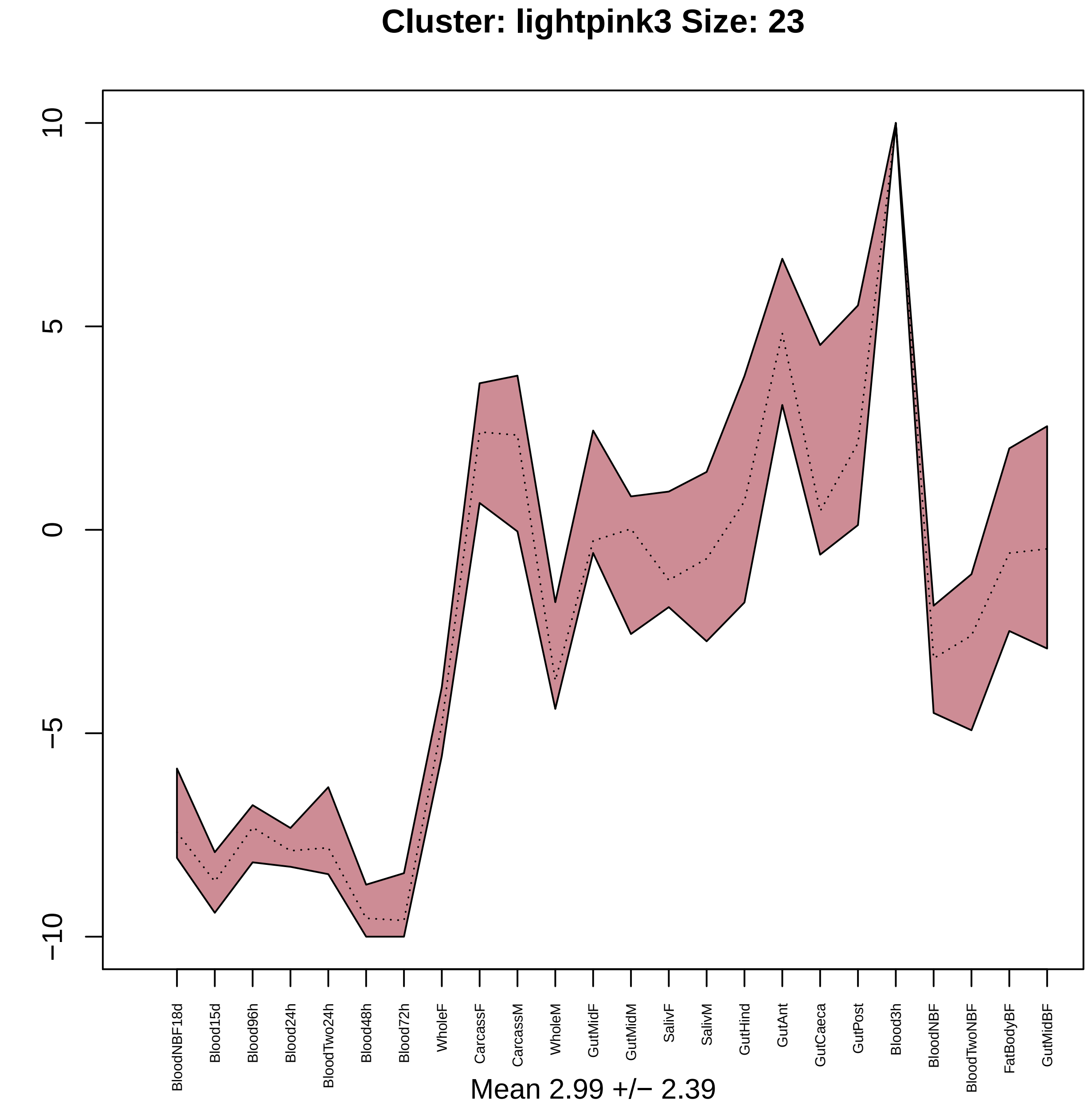
Cluster: lightpink3 Size: 23
10
5
0
−5
−10
BloodNBF18d
Blood15d
Blood96h
Blood24h
BloodTwo24h
Blood48h
Blood72h
WholeF
CarcassF
CarcassM
WholeM
GutMidF
GutMidM
SalivF
SalivM
GutHind
GutAnt
GutCaeca
GutPost
Blood3h
BloodNBF
BloodTwoNBF
FatBodyBF
GutMidBF
Mean 2.99 +/− 2.39

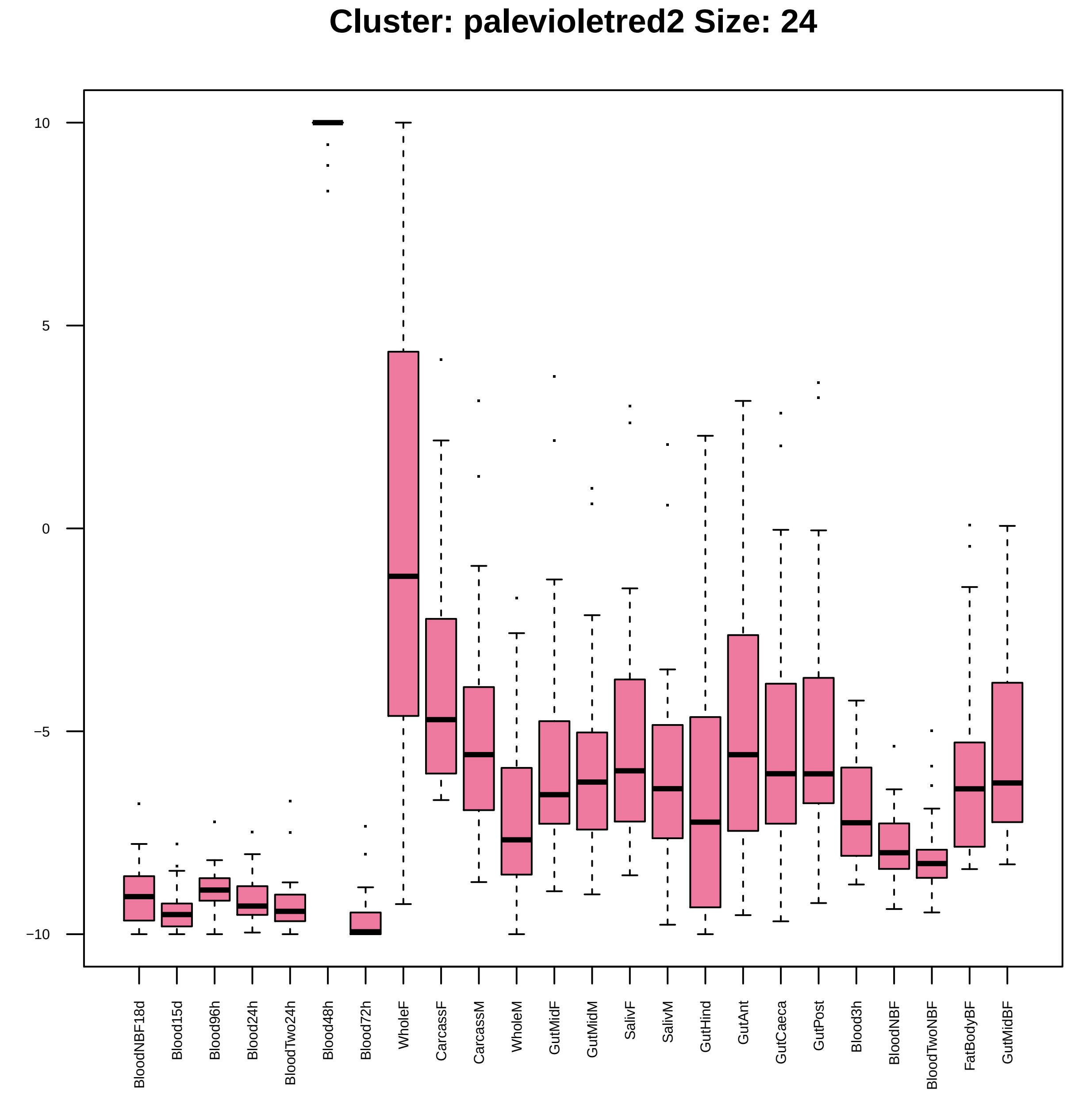
Cluster: palevioletred2 Size: 24
10
5
0
−5
−10
BloodNBF18d
Blood15d
Blood96h
Blood24h
BloodTwo24h
Blood48h
Blood72h
WholeF
CarcassF
CarcassM
WholeM
GutMidF
GutMidM
SalivF
SalivM
GutHind
GutAnt
GutCaeca
GutPost
Blood3h
BloodNBF
BloodTwoNBF
FatBodyBF
GutMidBF

| | GO.ID | BPCluster: palevioletred2 Size: 24 | Annotated | Significant | Expected | Rank in ClassicF | Weight01F | ClassicF |
|---|---|---|---|---|---|---|---|---|
| *1* | GO:0006979 | response to oxidative stress | 51 | 2 | 0.04 | 1 | 0.00062 | 0.00062 |
| *2* | GO:0006030 | chitin metabolic process | 95 | 2 | 0.07 | 2 | 0.00213 | 0.00213 |

| | GO.ID | MFCluster: palevioletred2 Size: 24 | Annotated | Significant | Expected | Rank in ClassicF | Weight01F | ClassicF |
|---|---|---|---|---|---|---|---|---|
| 2 | GO:0004601 | peroxidase activity | 26 | 2 | 0.06 | 2 | 0.0015 | 0.0015 |
| 8 | GO:0005488 | binding | 4627 | 15 | 10.27 | 5 | 1.0000 | 0.0082 |
| 9 | GO:0016684 | oxidoreductase activity, acting on perox... | 26 | 2 | 0.06 | 3 | 1.0000 | 0.0015 |
| 11 | GO:0016209 | antioxidant activity | 33 | 2 | 0.07 | 4 | 1.0000 | 0.0023 |

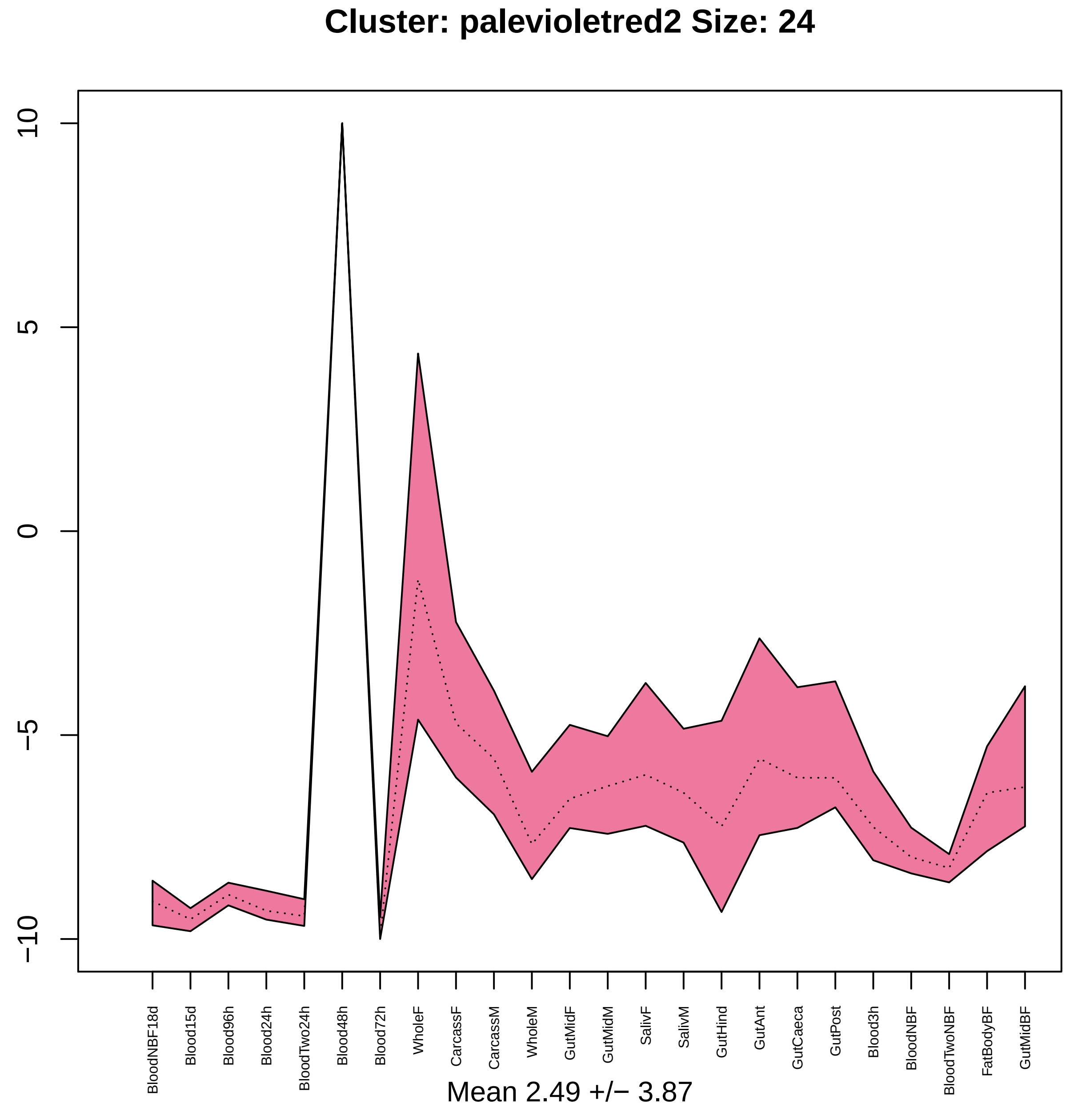
Cluster: palevioletred2 Size: 24
10
5
0
−5
−10
BloodNBF18d
Blood15d
Blood96h
Blood24h
BloodTwo24h
Blood48h
Blood72h
WholeF
CarcassF
CarcassM
WholeM
GutMidF
GutMidM
SalivF
SalivM
GutHind
GutAnt
GutCaeca
GutPost
Blood3h
BloodNBF
BloodTwoNBF
FatBodyBF
GutMidBF
Mean 2.49 +/− 3.87

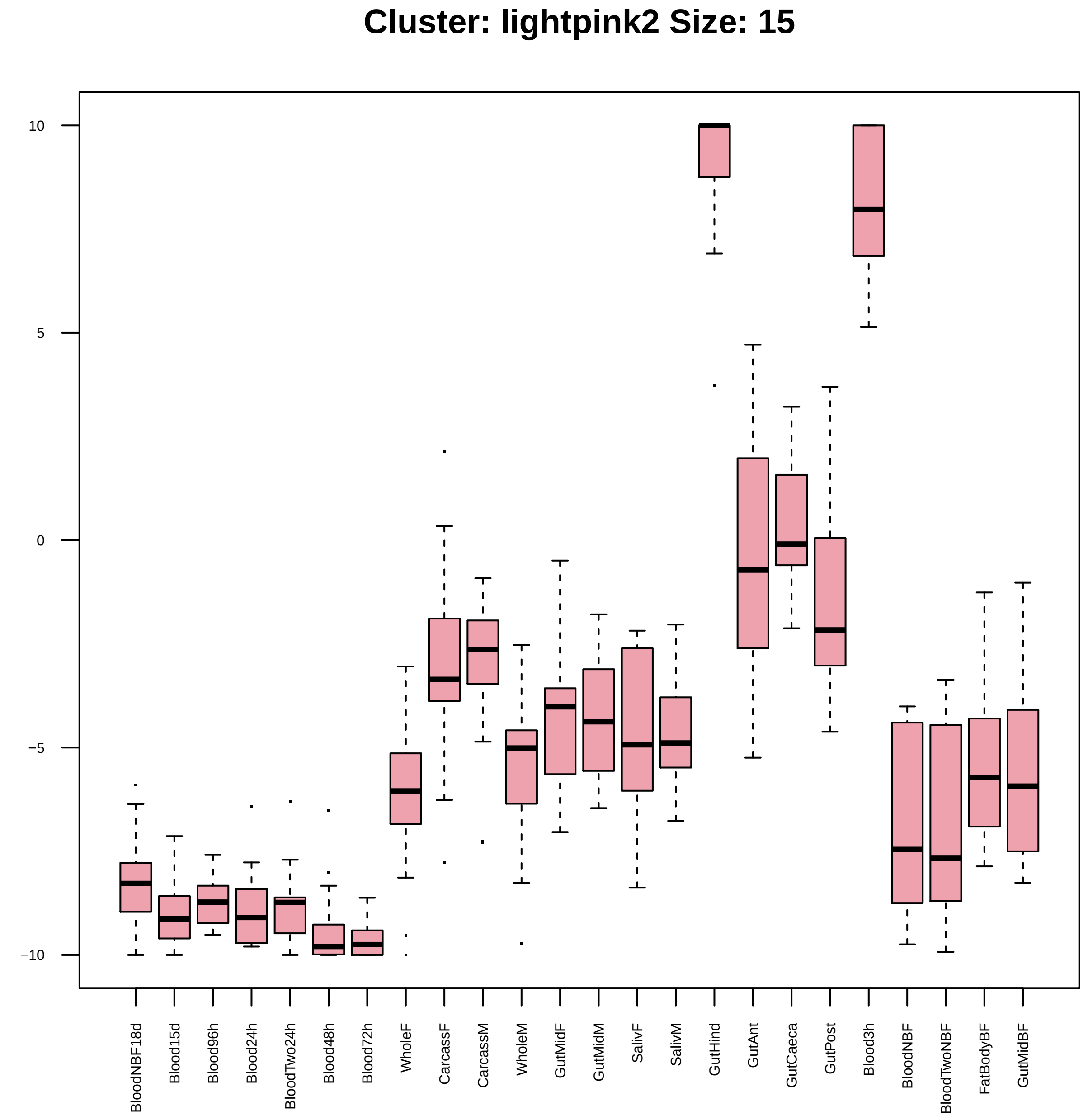

Cluster: lightpink2 Size: 15
10
5
0
−5
−10
BloodNBF18d
Blood15d
Blood96h
Blood24h
BloodTwo24h
Blood48h
Blood72h
WholeF
CarcassF
CarcassM
WholeM
GutMidF
GutMidM
SalivF
SalivM
GutHind
GutAnt
GutCaeca
GutPost
Blood3h
BloodNBF
BloodTwoNBF
FatBodyBF
GutMidBF

| | GO.ID | BPCluster: lightpink2 Size: 15 | Annotated | Significant | Expected | Rank in ClassicF | Weight01F | ClassicF |
|---|---|---|---|---|---|---|---|---|
| 2 | GO:0006030 | chitin metabolic process | 95 | 2 | 0.06 | 1 | 0.033 | 0.0013 |
| 8 | GO:1901071 | glucosamine−containing compound metaboli... | 99 | 2 | 0.06 | 2 | 1.000 | 0.0014 |
| 9 | GO:0006022 | aminoglycan metabolic process | 113 | 2 | 0.07 | 4 | 1.000 | 0.0018 |

| | GO.ID | MFCluster: lightpink2 Size: 15 | Annotated | Significant | Expected | Rank in ClassicF | Weight01F | ClassicF |
|---|---|---|---|---|---|---|---|---|
| *1* | GO:0008061 | chitin binding | 76 | 2 | 0.05 | 1 | 0.0011 | 0.0011 |

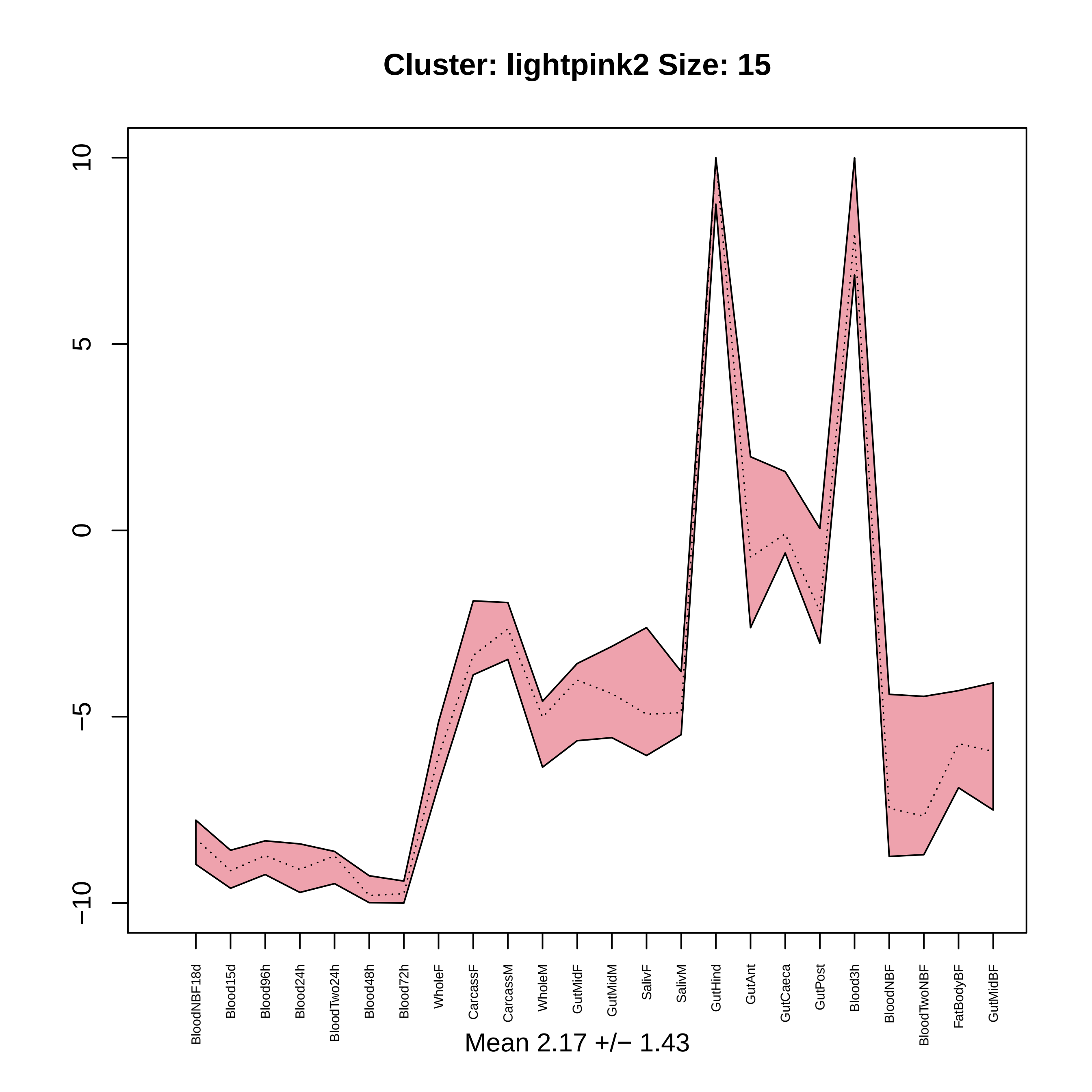
Cluster: lightpink2 Size: 15
10
5
0
−5
−10
BloodNBF18d
Blood15d
Blood96h
Blood24h
BloodTwo24h
Blood48h
Blood72h
WholeF
CarcassF
CarcassM
WholeM
GutMidF
GutMidM
SalivF
SalivM
GutHind
GutAnt
GutCaeca
GutPost
Blood3h
BloodNBF
BloodTwoNBF
FatBodyBF
GutMidBF
Mean 2.17 +/− 1.43

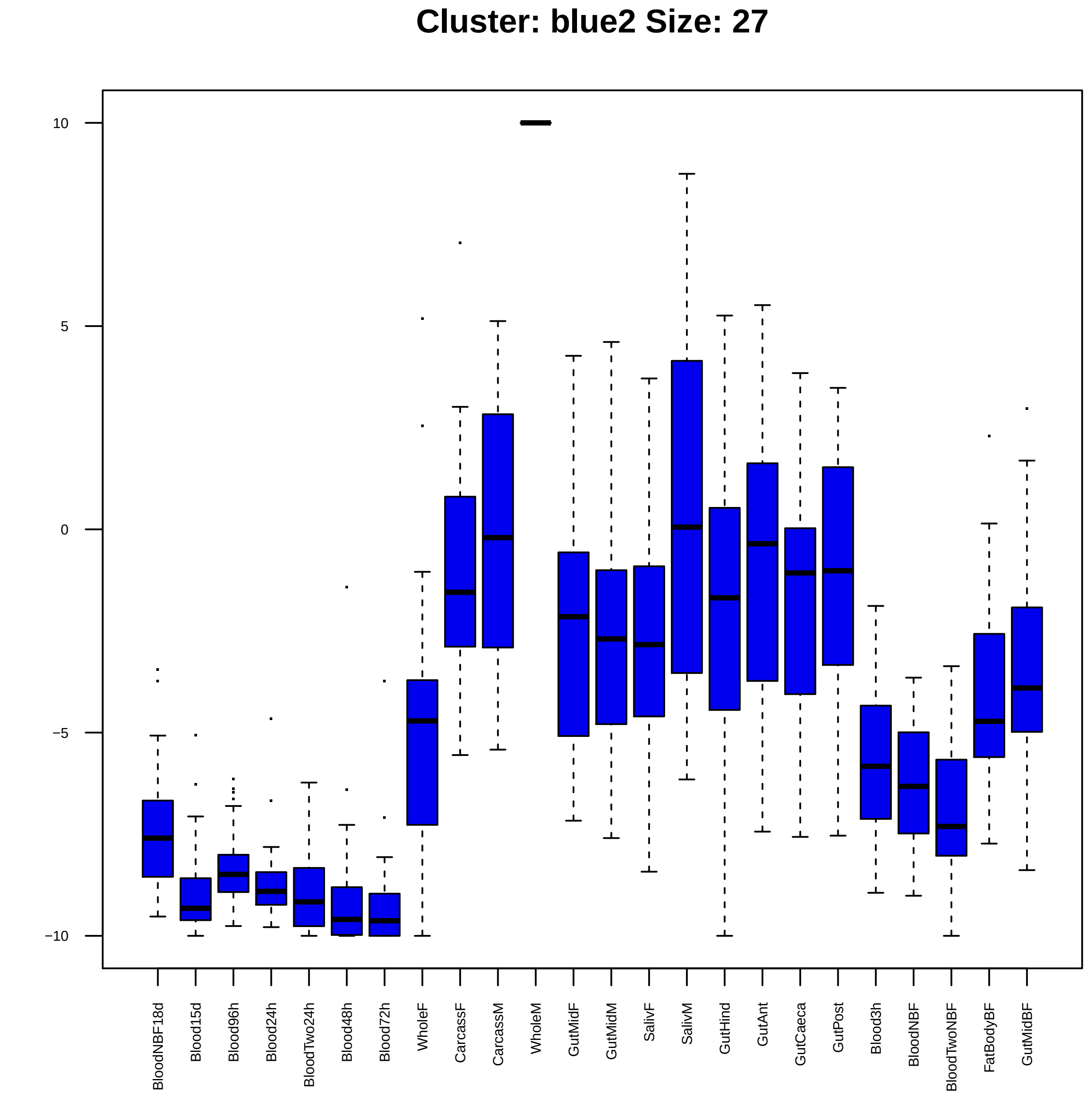

Cluster: blue2 Size: 27
10
5
0
−5
−10
BloodNBF18d
Blood15d
Blood96h
Blood24h
BloodTwo24h
Blood48h
Blood72h
WholeF
CarcassF
CarcassM
WholeM
GutMidF
GutMidM
SalivF
SalivM
GutHind
GutAnt
GutCaeca
GutPost
Blood3h
BloodNBF
BloodTwoNBF
FatBodyBF
GutMidBF

| | GO.ID | BPCluster: blue2 Size: 27 | Annotated | Significant | Expected | Rank in ClassicF | Weight01F | ClassicF |
|---|---|---|---|---|---|---|---|---|
| *1* | GO:0007018 | microtubule−based movement | 65 | 3 | 0.14 | 1 | 0.00034 | 0.00034 |

| | GO.ID | MFCluster: blue2 Size: 27 | Annotated | Significant | Expected | Rank in ClassicF | Weight01F | ClassicF |
|---|---|---|---|---|---|---|---|---|
| 1 | GO:0016887 | ATPase activity | 233 | 4 | 0.52 | 2 | 0.0028 | 0.00142 |
| 2 | GO:0003777 | microtubule motor activity | 40 | 2 | 0.09 | 3 | 0.0034 | 0.00343 |
| 4 | GO:0003774 | motor activity | 60 | 3 | 0.13 | 1 | 0.0384 | 0.00028 |

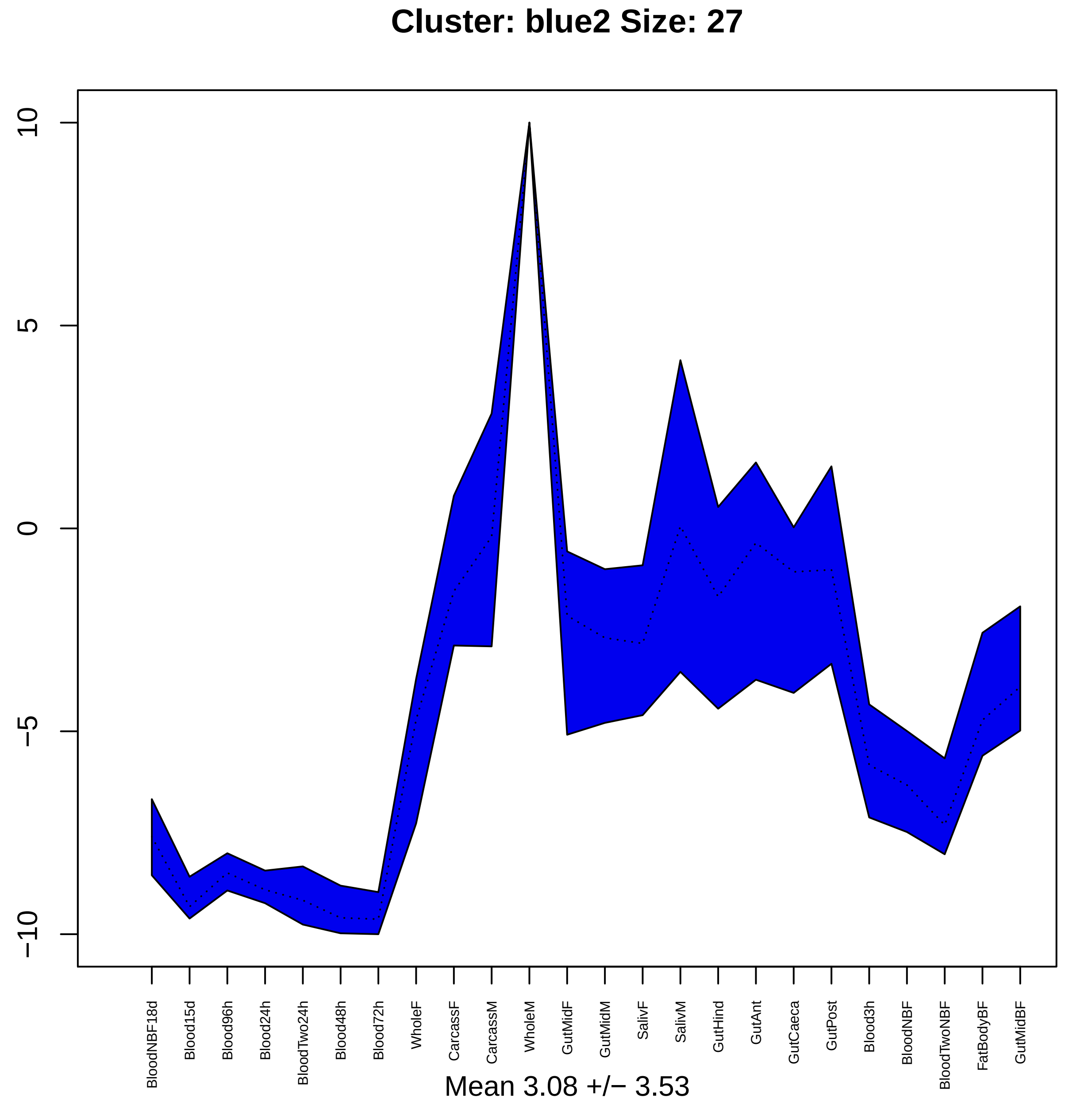
Cluster: blue2 Size: 27
10
5
0
−5
−10
BloodNBF18d
Blood15d
Blood96h
Blood24h
BloodTwo24h
Blood48h
Blood72h
WholeF
CarcassF
CarcassM
WholeM
GutMidF
GutMidM
SalivF
SalivM
GutHind
GutAnt
GutCaeca
GutPost
Blood3h
BloodNBF
BloodTwoNBF
FatBodyBF
GutMidBF
Mean 3.08 +/− 3.53

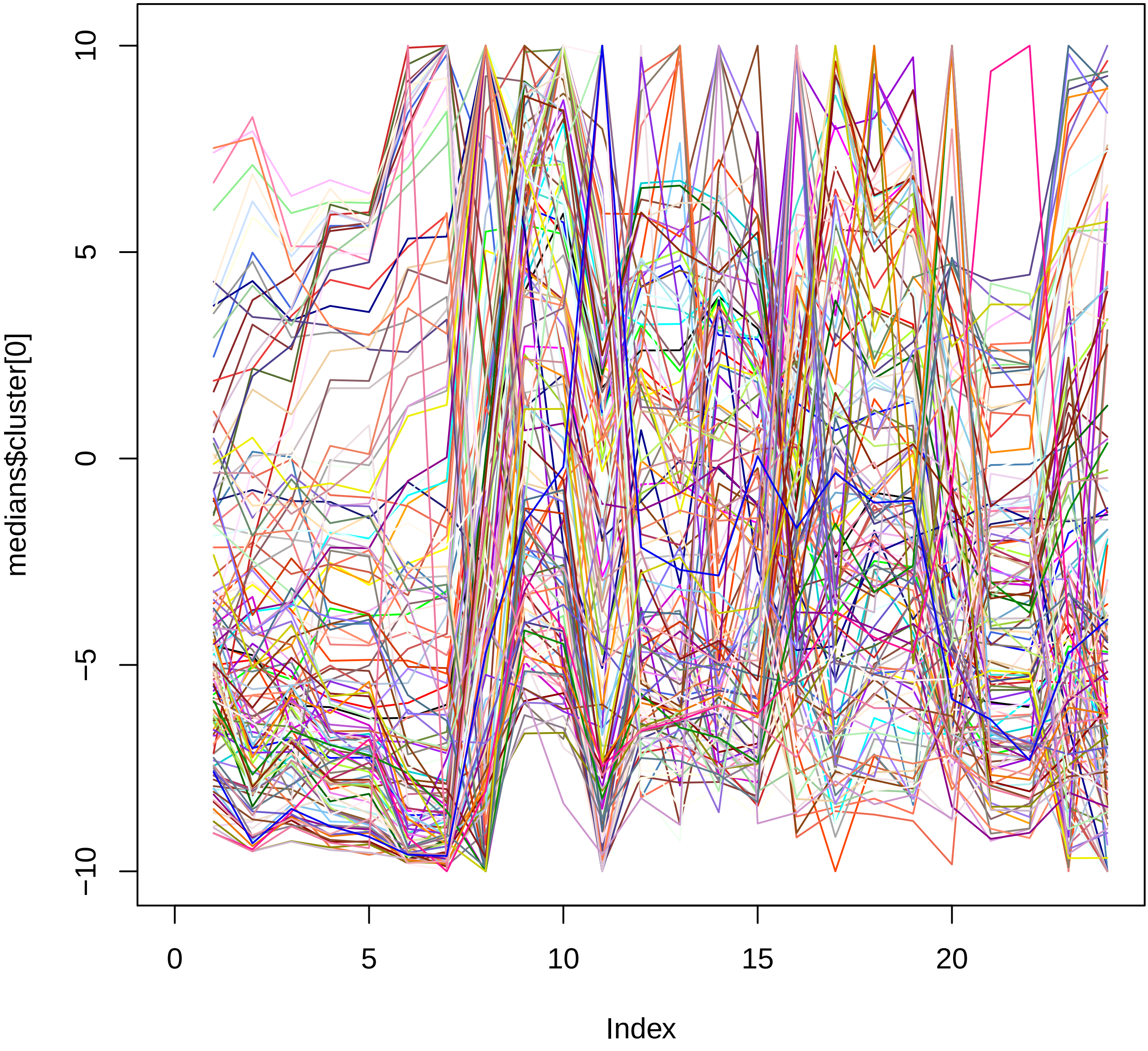

10
5
0
−5
−10
medians$cluster[0]
0
5
10
15
20
Index
