## Additional File 4 for "Functional constraints on insect immune system components govern their evolutionary trajectories"

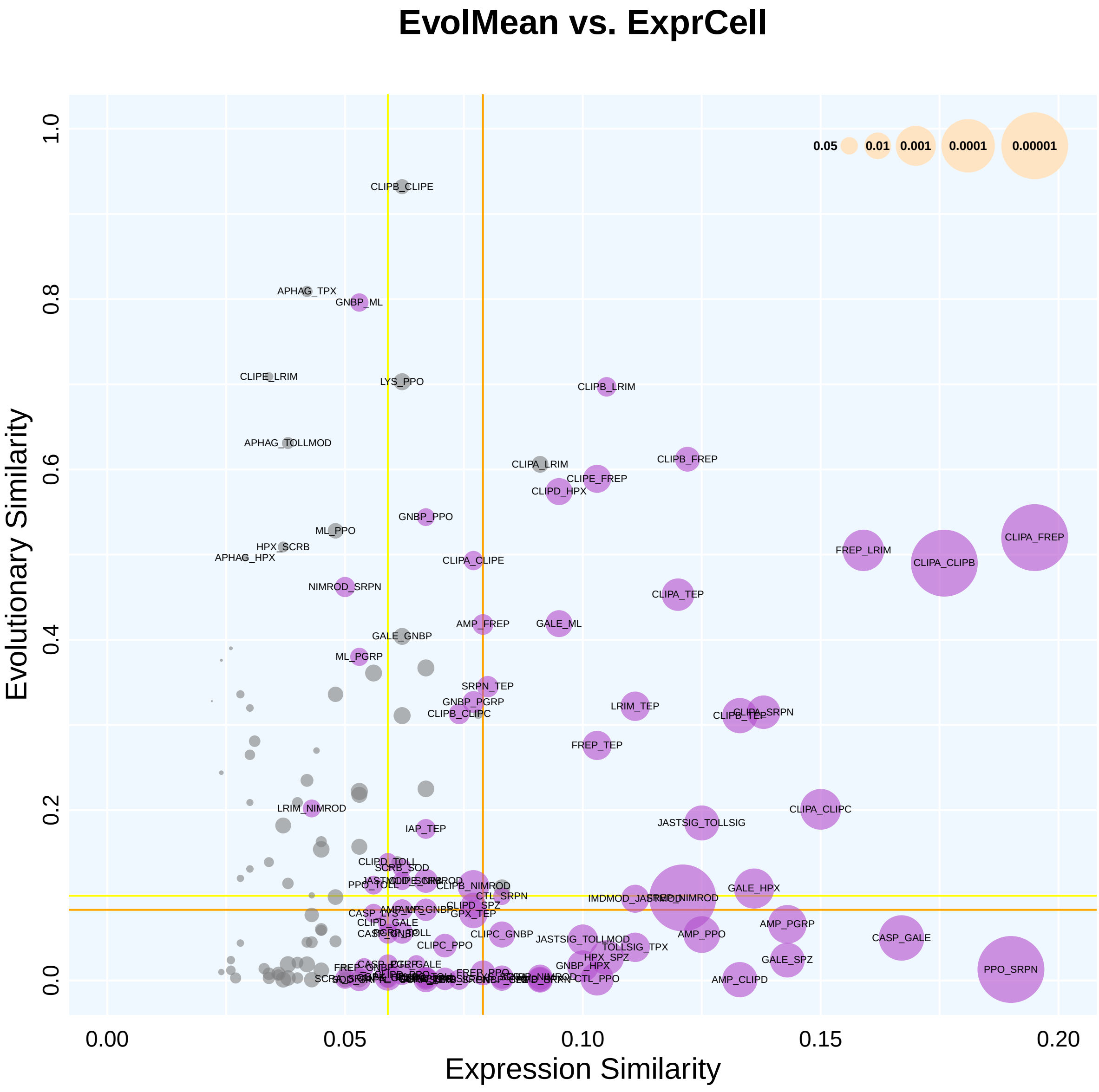
EvolMean vs. ExprCell
Evolutionary Similarity
Expression Similarity
0.0
0.2
0.4
0.6
0.8
1.0
0.00
0.05
0.10
0.15
0.20
0.05
0.01
0.001
0.0001
0.00001
CLIPB_CLIPE
APHAG_TPX
GNBP_ML
CLIPE_LRIM
LYS_PPO
CLIPB_LRIM
APHAG_TOLLMOD
CLIPB_FREP
CLIPA_LRIM
CLIPE_FREP
CLIPD_HPX
GNBP_PPO
ML_PPO
HPX_SCRB
APHAG_HPX
CLIPA_CLIPE
CLIPA_FREP
FREP_LRIM
CLIPA_CLIPB
NIMROD_SRPN
CLIPA_TEP
AMP_FREP
GALE_ML
GALE_GNBP
ML_PGRP
SRPN_TEP
GNBP_PGRP
CLIPB_CLIPC
LRIM_TEP
CLIPB_TEP
CLIPA_SRPN
FREP_TEP
CLIPA_CLIPC
LRIM_NIMROD
JASTSIG_TOLLSIG
IAP_TEP
CLIPD_TOLL
SCRB_SOD
CLIPB_NIMROD
CTL_SRPN
IMDMOD_JASTSIG
GALE_HPX
CLIPD_SPZ
GPX_TEP
CLIPD_GALE
AMP_PGRP
AMP_PPO
CLIPC_GNBP
JASTSIG_TOLLMOD
TOLLSIG_TPX
CASP_GALE
CLIPC_PPO
HPX_SPZ
GALE_SPZ
GNBP_HPX
CTL_PPO
AMP_CLIPD
PPO_SRPN

### EvolMean vs. ExprSupercell

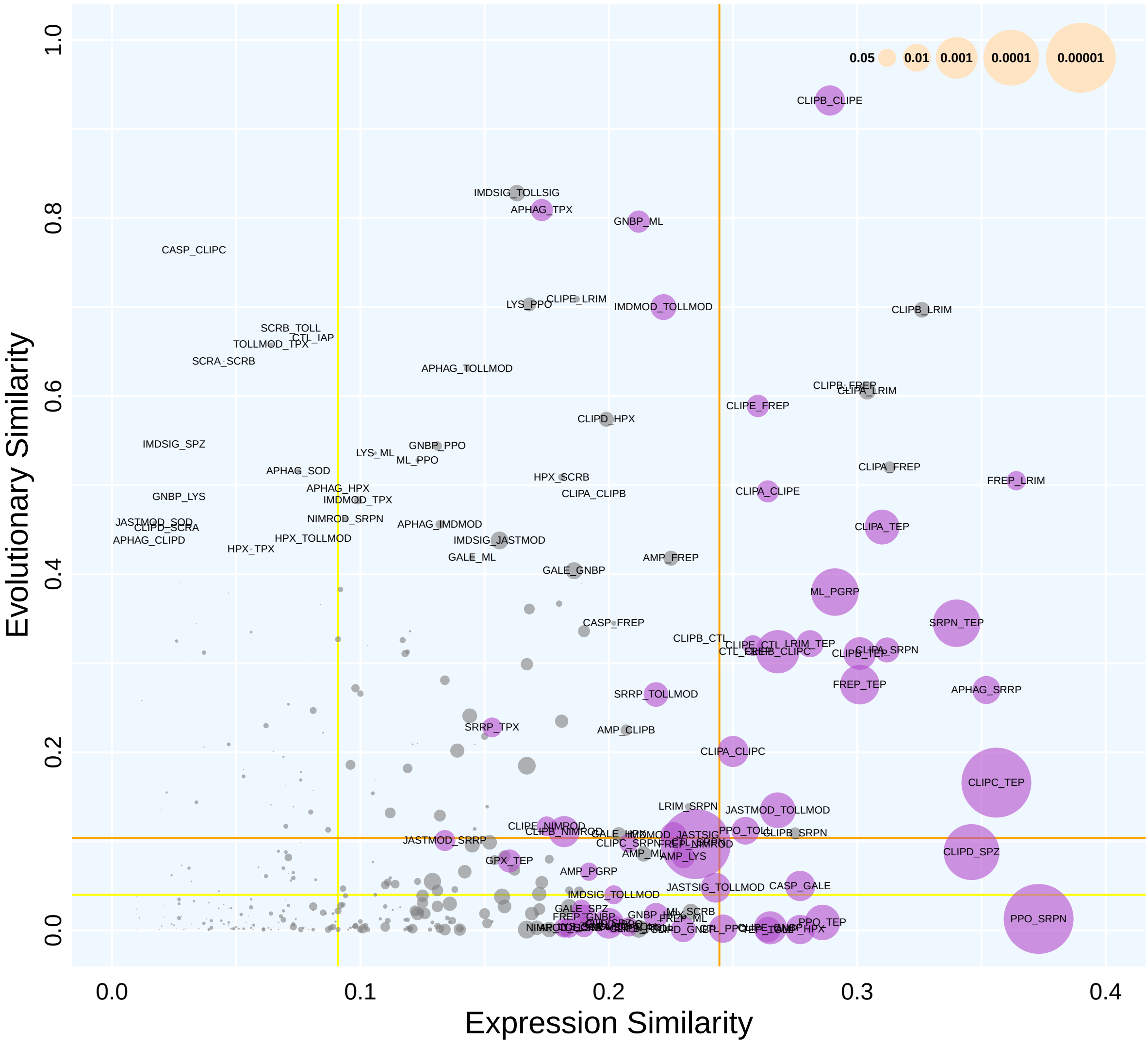

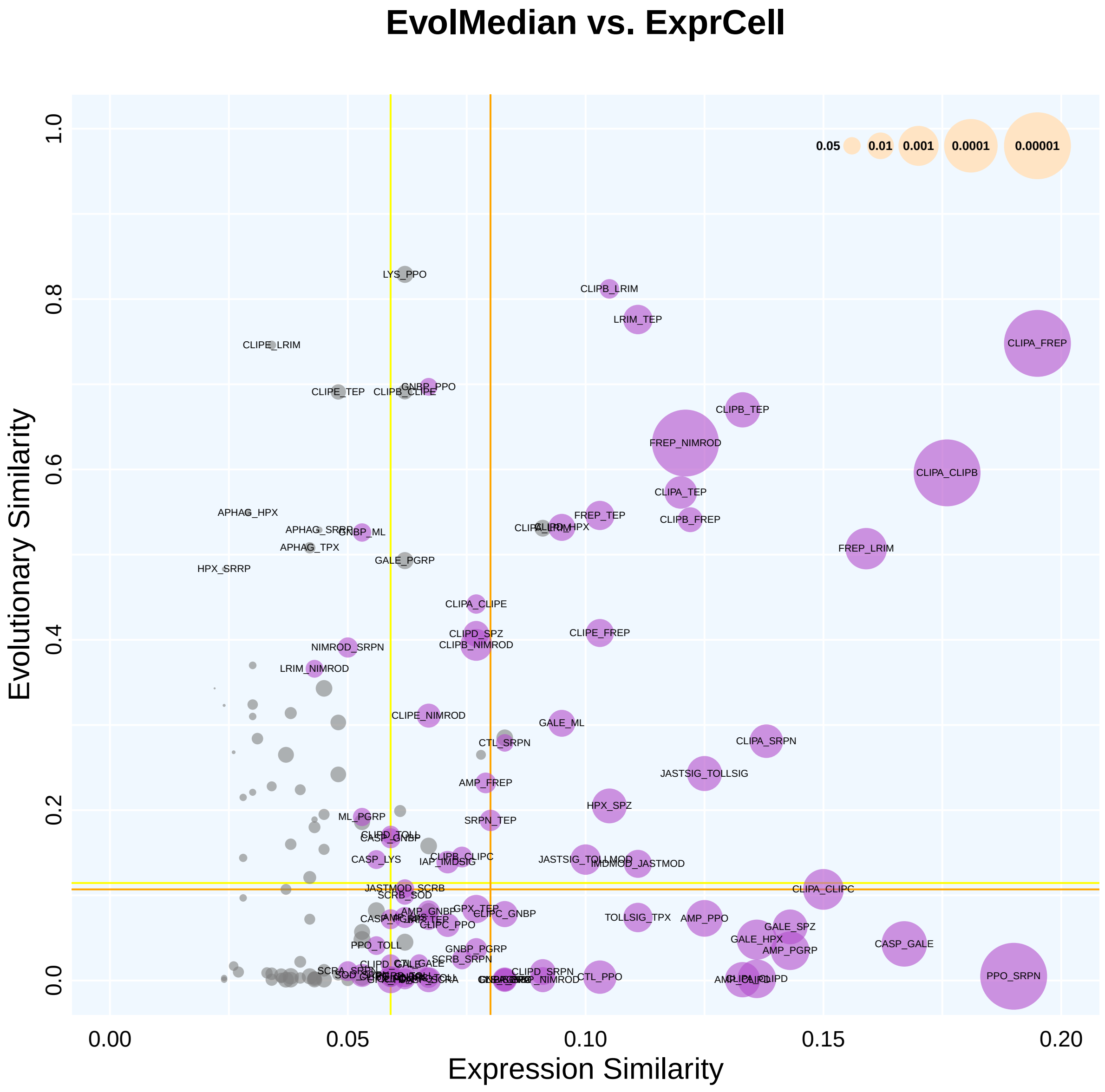
EvolMedian vs. ExprCell
Evolutionary Similarity
Expression Similarity
0.0
0.2
0.4
0.6
0.8
1.0
0.00
0.05
0.10
0.15
0.20
0.05
0.01
0.001
0.0001
0.00001
LYS_PPO
CLIPB_LRIM
LRIM_TEP
CLIPE_LRIM
CLIPA_FREP
CLIPE_TEP
GNBP_PPO
CLIPB_TEP
FREP_NIMROD
CLIPA_CLIPB
CLIPA_TEP
APHAG_HPX
FREP_TEP
CLIPB_FREP
GNBP_ML
APHAG_TPX
FREP_LRIM
HPX_SRRP
GALE_PGRP
CLIPA_CLIPE
CLIPD_SPZ
CLIPE_FREP
NIMROD_SRPN
CLIPB_NIMROD
LRIM_NIMROD
CLIPE_NIMROD
GALE_ML
CTL_SRPN
CLIPA_SRPN
JASTSIG_TOLLSIG
AMP_FREP
HPX_SPZ
ML_PGRP
SRPN_TEP
CASP_GNBP
CASP_LYS
IAP_IMDSIG
CLIPB_CLIPC
IMDMOD_JASTMOD
JASTMOD_SCRB
SCRB_SOD
CLIPA_CLIPC
GPX_TEP
AMP_GNBP
CLIPC_GNBP
TOLLSIG_TPX
AMP_PPO
CLIPC_PPO
GALE_SPZ
GALE_HPX
CASP_GALE
AMP_PGRP
PPO_TOLL
GNBP_PGRP
SCRB_SRPN
CLIPD_SRPN
CTL_PPO
PPO_SRPN

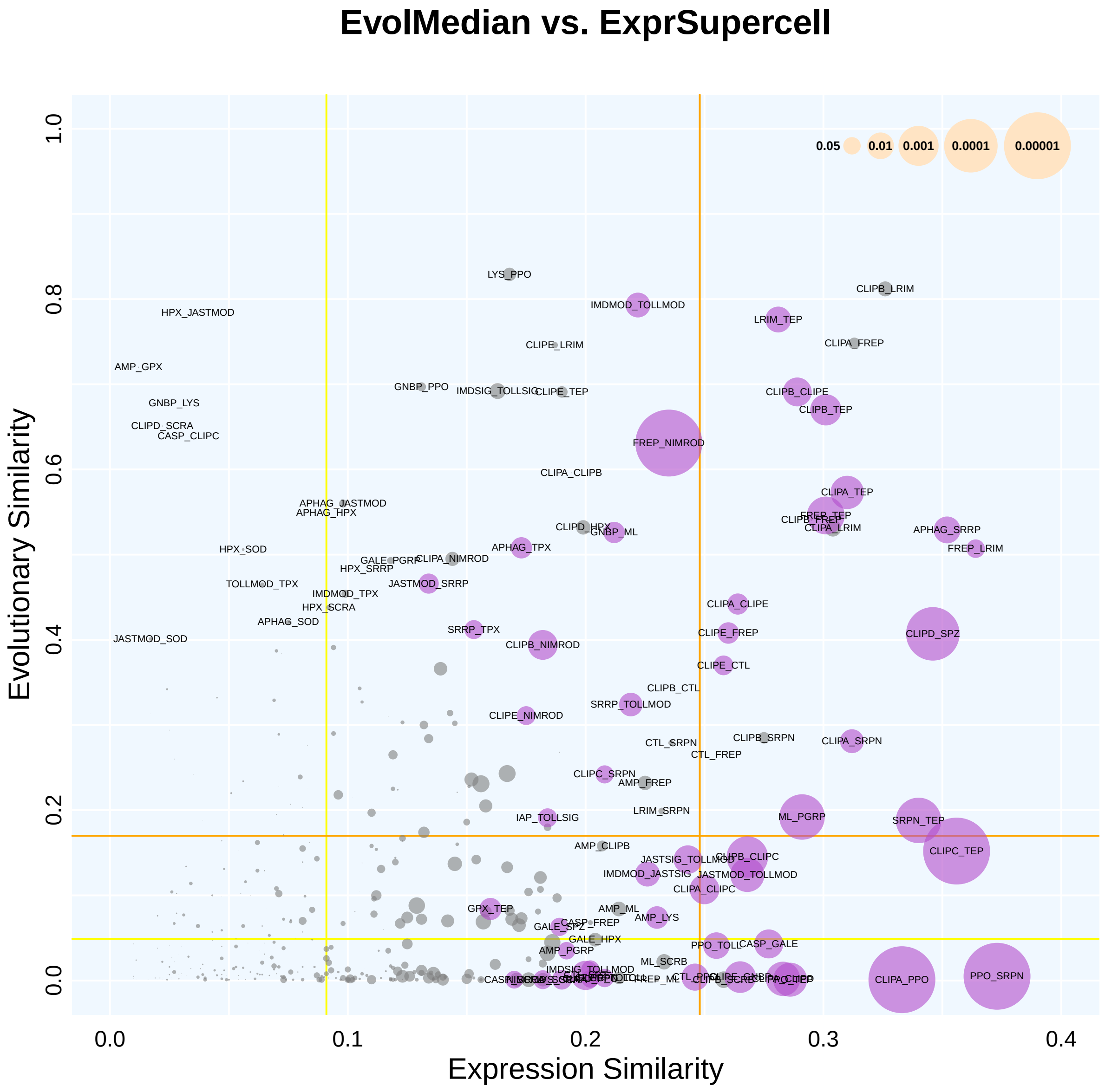
EvolMedian vs. ExprSupercell
Evolutionary Similarity
Expression Similarity
0.0
0.2
0.4
0.6
0.8
1.0
0.1
0.3
0.05
0.01
0.001
0.0001
0.00001
LYS_PPO
CLIPB_LRIM
IMDMOD_TOLLMOD
HPX_JASTMOD
LRIM_TEP
CLIPE_LRIM
CLIPA_FREP
AMP_GPX
GNBP_PPO
IMDSIG_TOLLSIG
CLIPE_TEP
CLIPB_CLIPE
CLIPB_TEP
GNBP_LYS
CLIPD_SCRA
CASP_CLIPC
FREP_NIMROD
CLIPA_CLIPB
CLIPA_TEP
APHAG_JASTMOD
APHAG_HPX
FREP_TEP
CLIPB_FREP
CLIPA_LRIM
CLIPD_HPX
GNBP_ML
APHAG_SRRP
HPX_SOD
APHAG_TPX
FREP_LRIM
GALE_PGRP
CLIPA_NIMROD
HPX_SRRP
JASTMOD_SRRP
TOLLMOD_TPX
IMDMOD_TPX
HPX_SCRA
CLIPA_CLIPE
APHAG_SOD
SRRP_TPX
CLIPE_FREP
CLIPD_SPZ
JASTMOD_SOD
CLIPB_NIMROD
CLIPE_CTL
CLIPB_CTL
SRRP_TOLLMOD
CLIPE_NIMROD
CTL_SRPN
CLIPB_SRPN
CLIPA_SRPN
CTL_FREP
CLIPC_SRPN
AMP_FREP
LRIM_SRPN
ML_PGRP
IAP_TOLLSIG
SRPN_TEP
AMP_CLIPB
CLIPC_TEP
JASTSIG_TOLLMOD
CLIPB_CLIPC
IMDMOD_JASTSIG
JASTMOD_TOLLMOD
CLIPA_CLIPC
GPX_TEP
AMP_ML
AMP_LYS
CASP_FREP
GALE_SPZ
GALE_HPX
PPO_TOLL
CASP_GALE
AMP_PGRP
ML_SCRB
IMDSIG_TOLLMOD
FREP_ML
CLIPA_PPO
PPO_SRPN

### EvolMean vs. ExprModule0

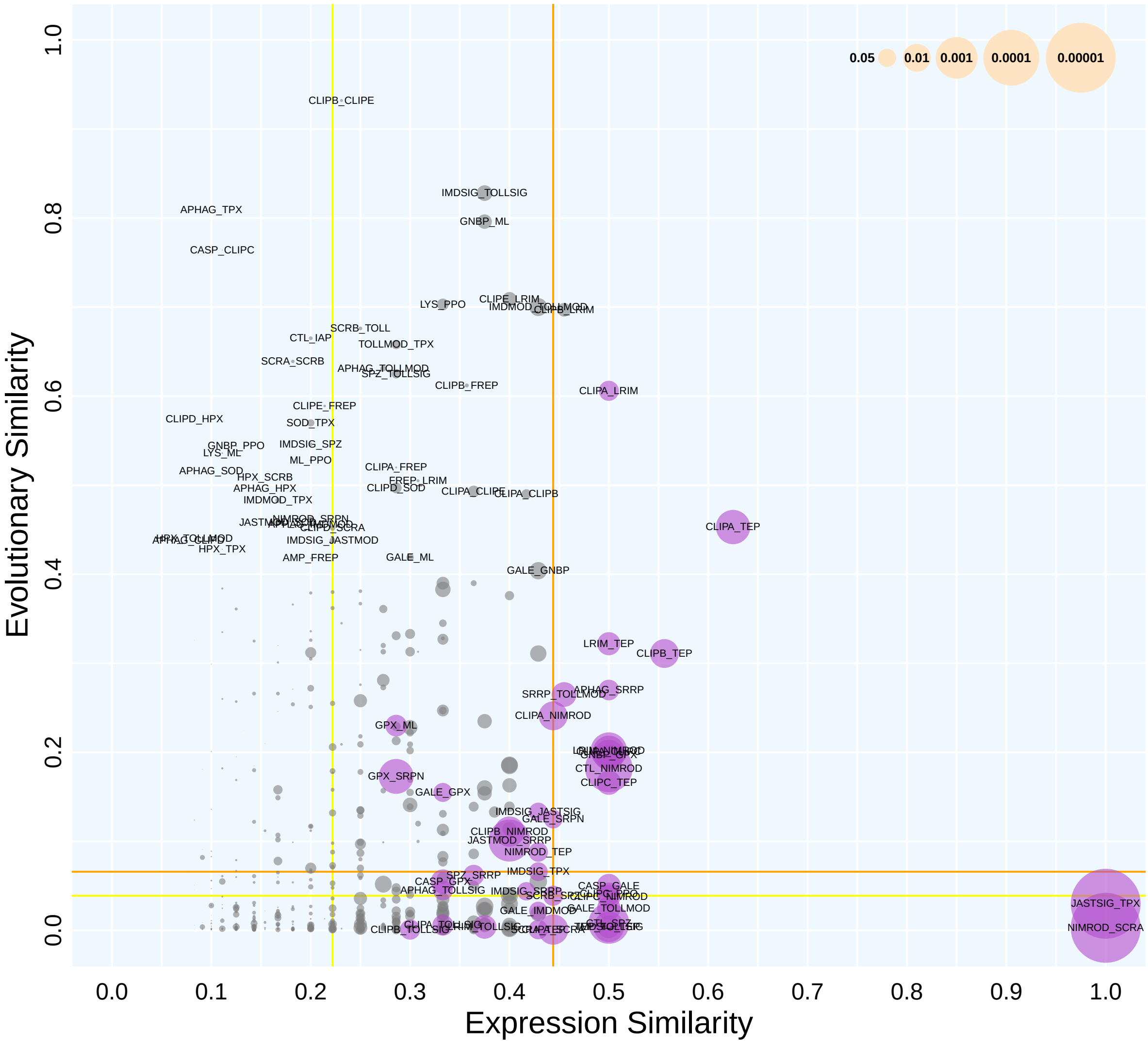

### EvolMean vs. ExprModule1

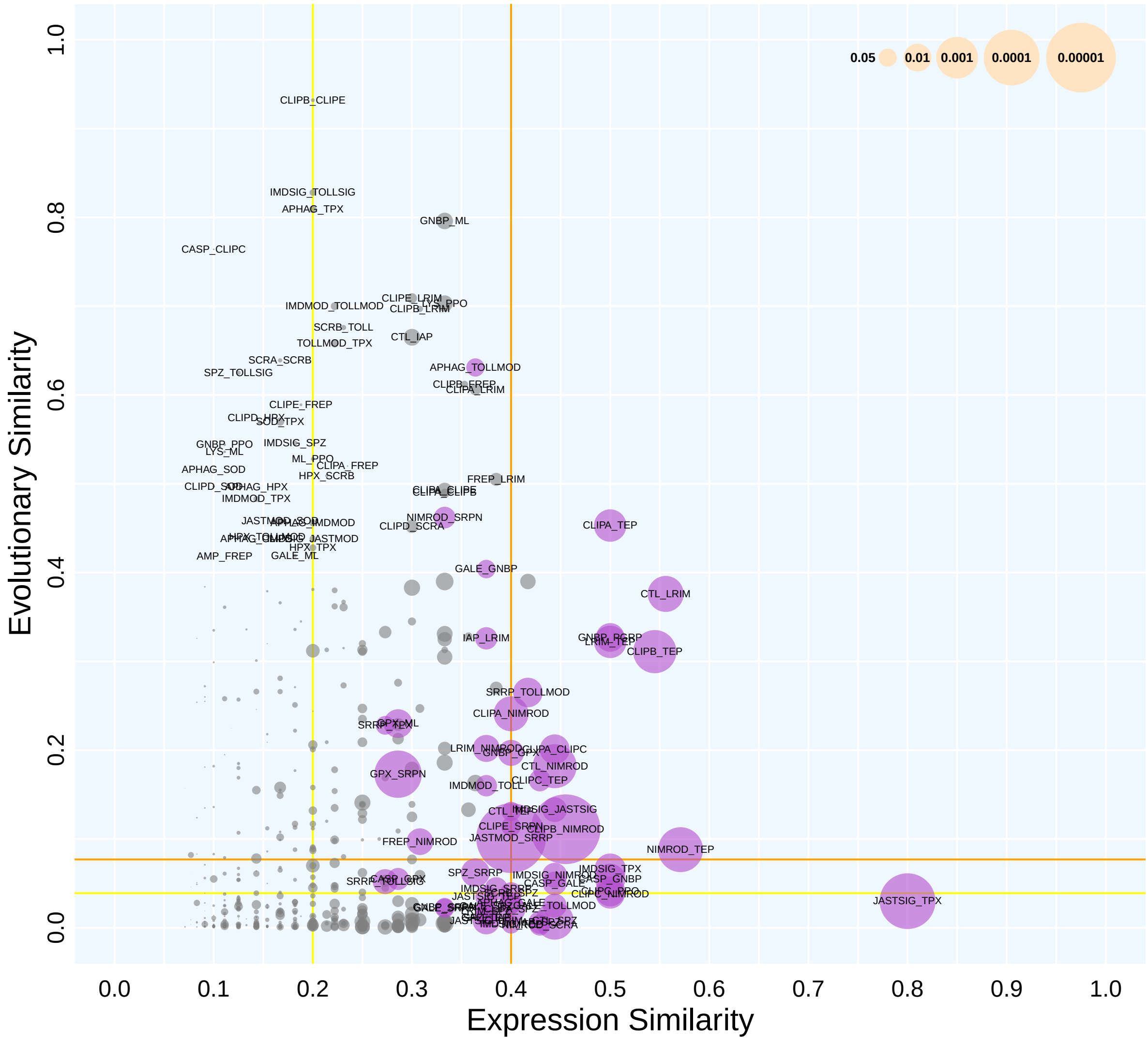

#### EvolMean vs. ExprModule2

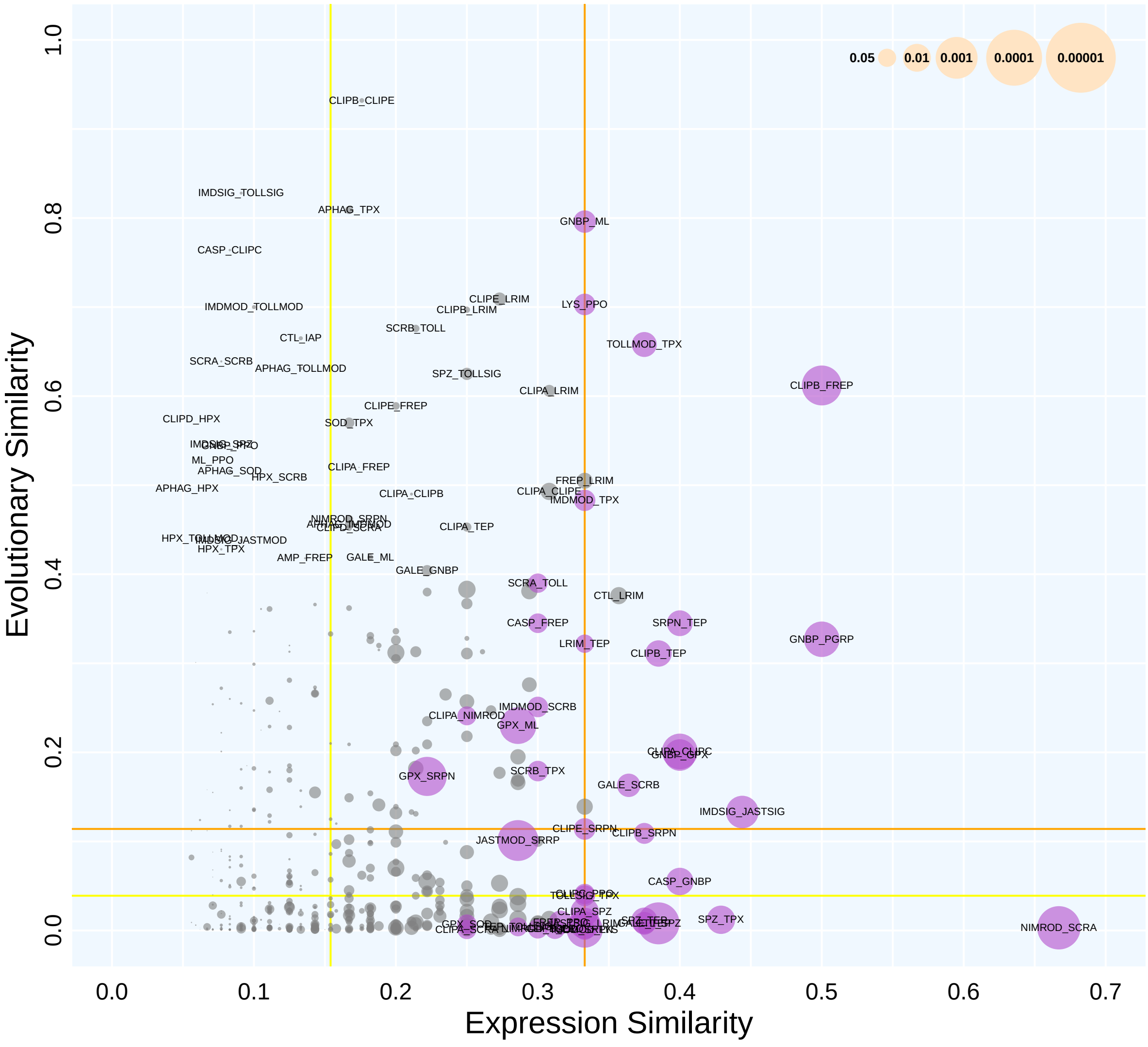

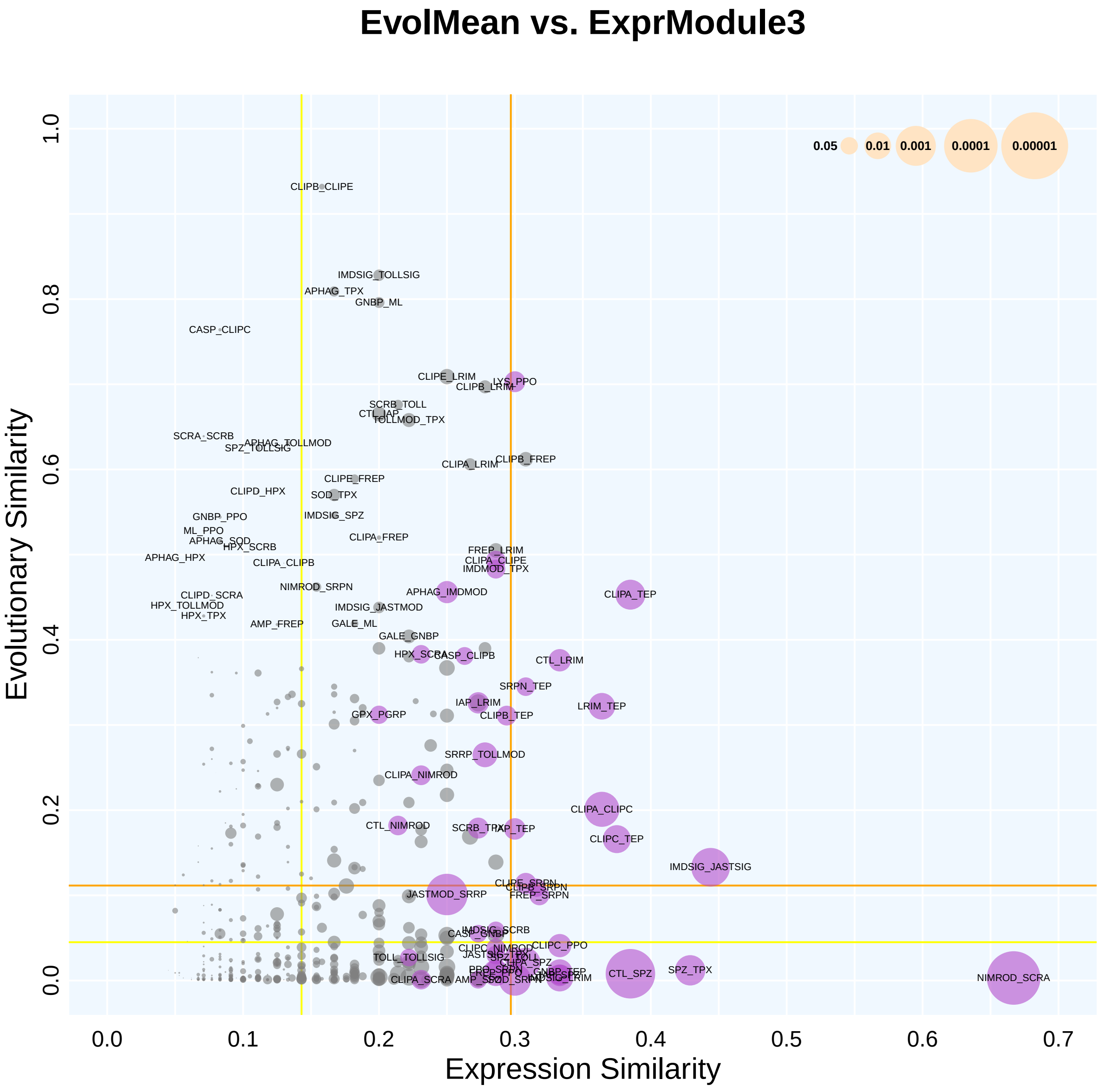
EvolMean vs. ExprModule3
Evolutionary Similarity
Expression Similarity
0.05
0.01
0.001
0.0001
0.00001
0.0
0.2
0.4
0.6
0.8
1.0
0.0
0.1
0.2
0.3
0.4
0.5
0.6
0.7
CLIPB_CLIPE
IMDSIG_TOLLSIG
APHAG_TPX
GNBP_ML
CASP_CLIPC
CLIPE_LRIM
LYS_PPO
CLIPB_LRIM
SCRB_TOLL
CTL_IAP
TOLLMOD_TPX
SCRA_SCRB
APHAG_TOLLMOD
SPZ_TOLLSIG
CLIPA_LRIM
CLIPB_FREP
CLIPE_FREP
CLIPD_HPX
SOD_TPX
GNBP_PPO
IMDSIG_SPZ
ML_PPO
APHAG_SOD
CLIPA_FREP
HPX_SCRB
APHAG_HPX
CLIPA_CLIPB
FREP_LRIM
CLIPA_CLIPE
IMDMOD_TPX
NIMROD_SRPN
APHAG_IMDMOD
CLIPA_TEP
CLIPD_SCRA
HPX_TOLLMOD
IMDSIG_JASTMOD
HPX_TPX
AMP_FREP
GALE_ML
GALE_GNBP
HPX_SCRA
CASP_CLIPB
CTL_LRIM
SRPN_TEP
IAP_LRIM
LRIM_TEP
GPX_PGRP
CLIPB_TEP
SRRP_TOLLMOD
CLIPA_NIMROD
CLIPA_CLIPC
CTL_NIMROD
SCRB_TPX
IAP_TEP
CLIPC_TEP
IMDSIG_JASTSIG
CLIPE_SRPN
CLIPB_SRPN
JASTMOD_SRRP
FREP_SRPN
IMDSIG_SCRB
CASP_GNBP
CLIPC_PPO
CLIPC_NIMROD
TOLL_TOLLSIG
CLIPA_SPZ
CTL_SPZ
SPZ_TPX
GNBP_TEP
IMDSIG_LRIM
CLIPA_SCRA
AMP_SOD
NIMROD_SCRA

#### EvolMean vs. ExprModule4

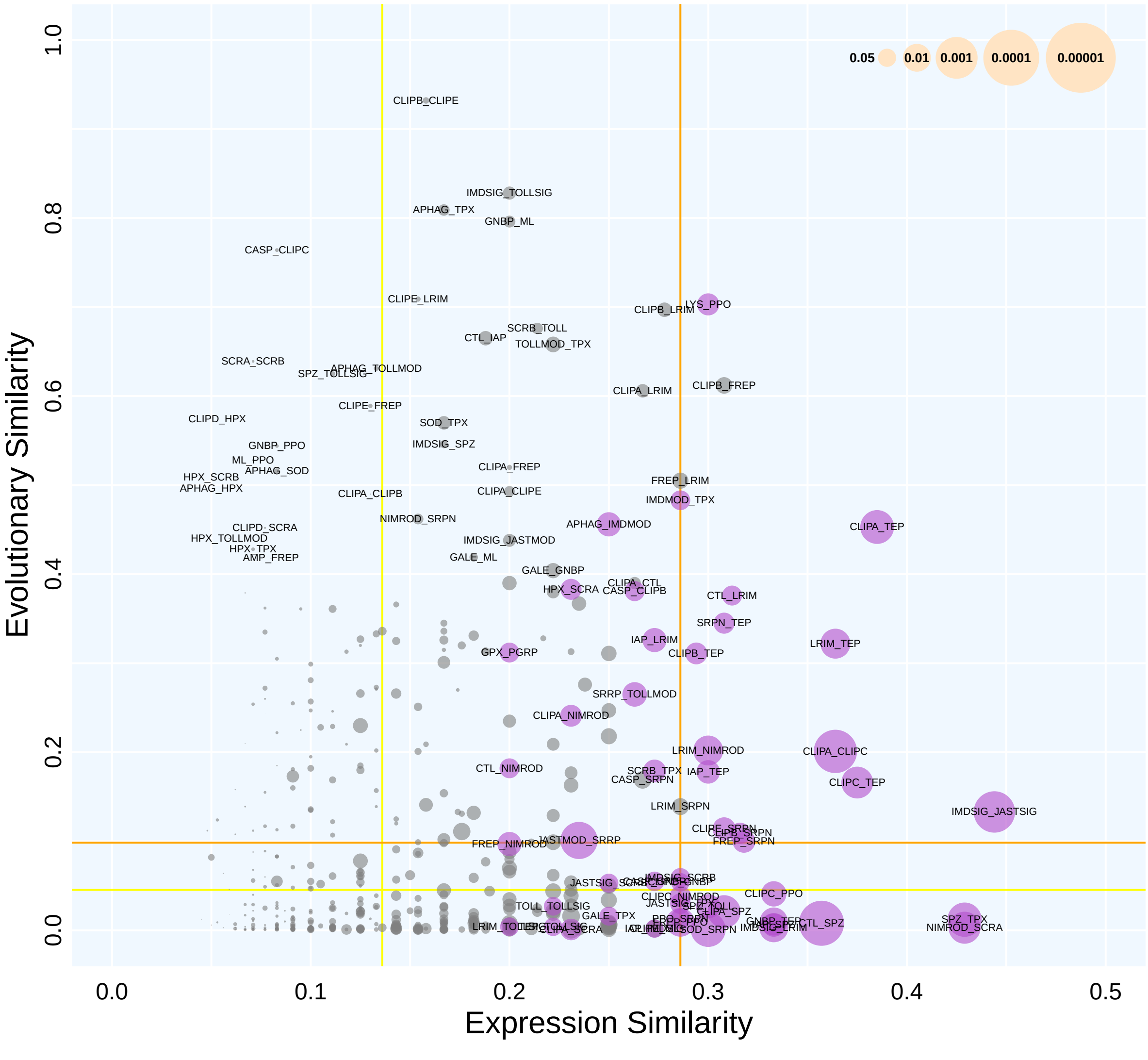

### EvolMedian vs. ExprModule0

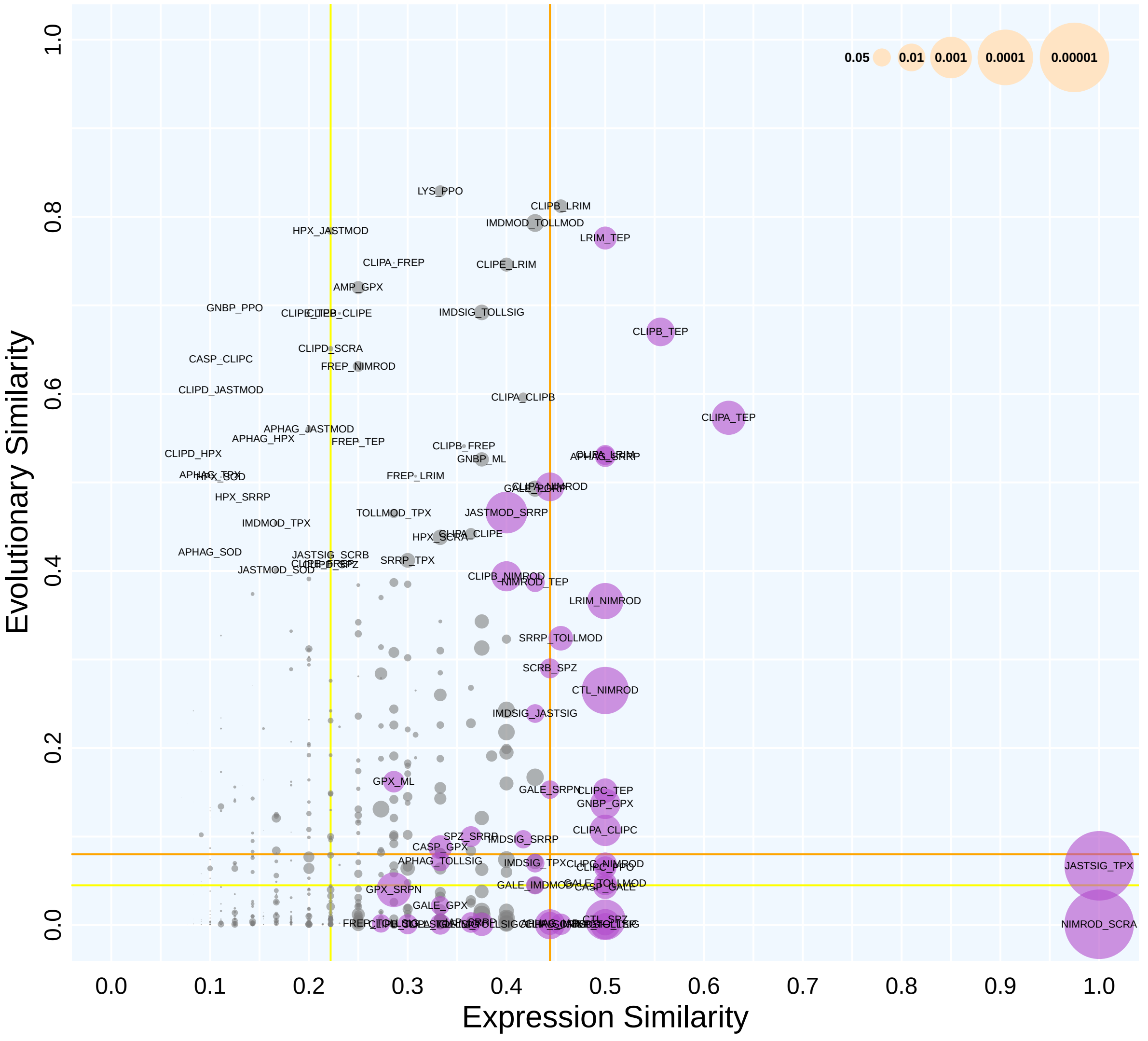

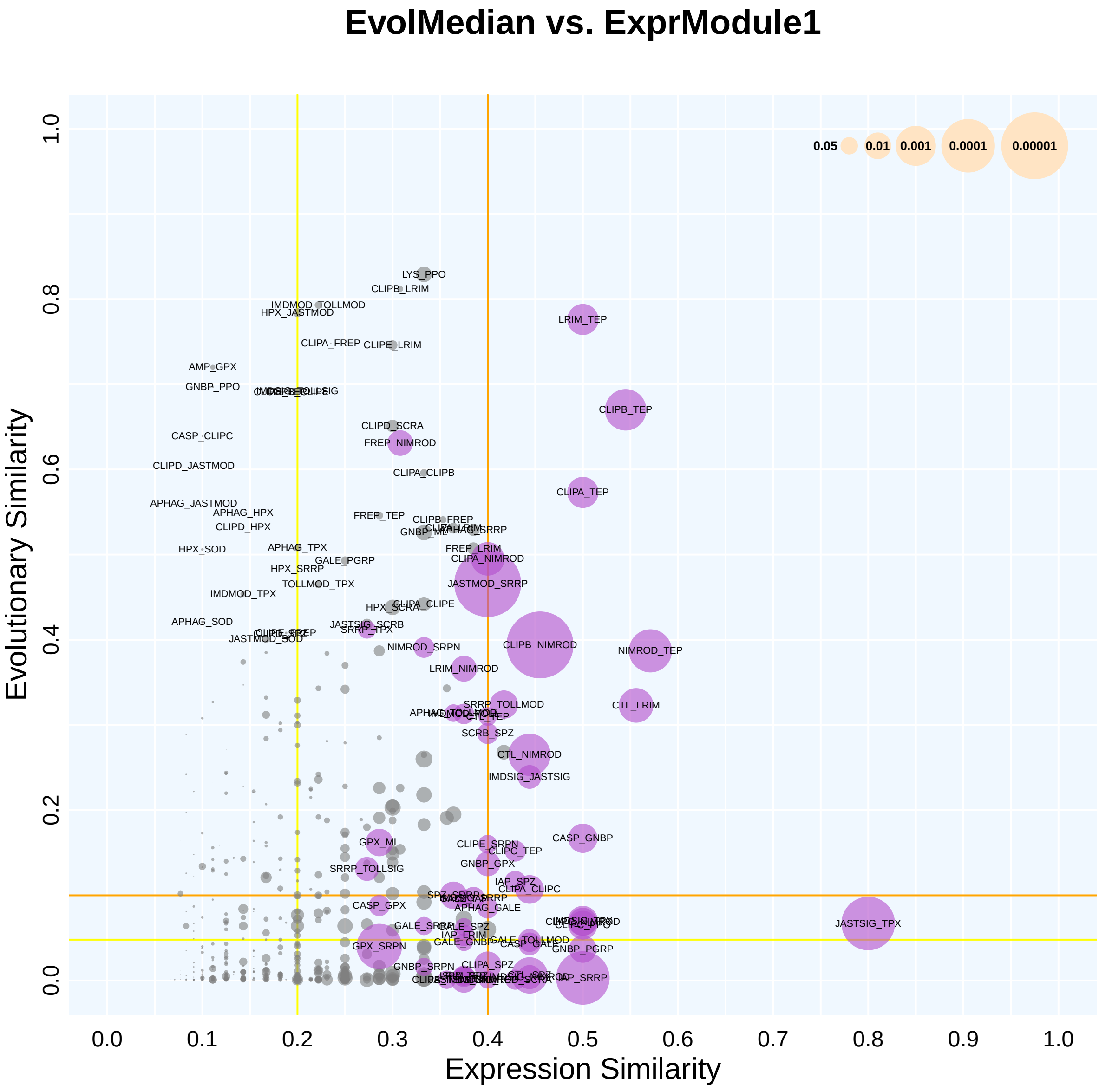
EvolMedian vs. ExprModule1
Evolutionary Similarity
Expression Similarity
0.0
0.2
0.4
0.6
0.8
1.0
0.0
0.1
0.2
0.3
0.4
0.5
0.6
0.7
0.8
0.9
1.0
0.05
0.01
0.001
0.0001
0.00001
LYS_PPO
CLIPB_LRIM
IMDMOD_TOLLMOD
HPX_JASTMOD
LRIM_TEP
CLIPA_FREP
CLIPE_LRIM
AMP_GPX
GNBP_PPO
CLIPB_TEP
CLIPD_SCRA
FREP_NIMROD
CASP_CLIPC
CLIPD_JASTMOD
CLIPA_CLIPB
CLIPA_TEP
APHAG_JASTMOD
APHAG_HPX
CLIPD_HPX
FREP_TEP
CLIPB_FREP
GNBP_ML
HPX_SOD
APHAG_TPX
FREP_LRIM
CLIPA_NIMROD
GALE_PGRP
HPX_SRRP
TOLLMOD_TPX
JASTMOD_SRRP
IMDMOD_TPX
HPX_SCRA
CLIPA_CLIPE
APHAG_SOD
JASTSIG_SCRB
SRRP_TPX
JASTMOD_SOD
CLIPB_NIMROD
NIMROD_TEP
NIMROD_SRPN
LRIM_NIMROD
SRRP_TOLLMOD
CTL_LRIM
CTL_TEP
SCRB_SPZ
CTL_NIMROD
IMDSIG_JASTSIG
CASP_GNBP
GPX_ML
CLIPE_SRPN
CLIPC_TEP
GNBP_GPX
SRRP_TOLLSIG
IAP_SPZ
CLIPA_CLIPC
APHAG_GALE
CASP_GPX
JASTSIG_TPX
GALE_SRPP
GALE_SPZ
IAP_TRIM
CLIPC_PPO
GALE_TOLLMOD
CASP_GALE
GNBP_PGRP
GPX_SRPN
GNBP_SRPN
CLIPA_SPZ
IAP_SRRP

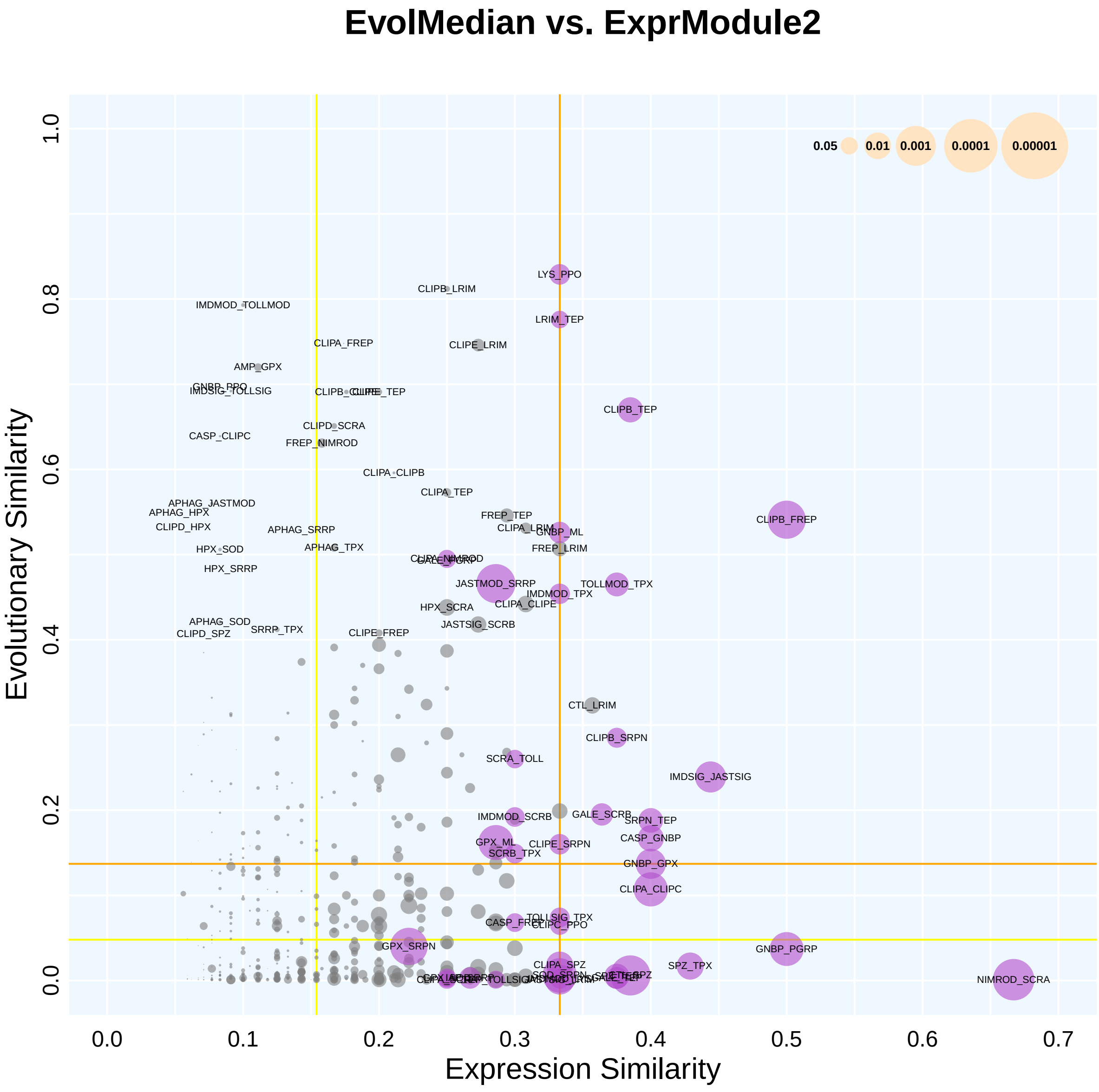
EvolMedian vs. ExprModule2
Evolutionary Similarity
Expression Similarity
0.0
0.2
0.4
0.6
0.8
1.0
0.0
0.1
0.2
0.3
0.4
0.5
0.6
0.7
0.05
0.01
0.001
0.0001
0.00001
LYS_PPO
CLIPB_LRIM
IMDMOD_TOLLMOD
LRIM_TEP
CLIPA_FREP
CLIPE_LRIM
AMP_GPX
IMDSIG_TOLLSIG
CLIPB_TEP
CLIPD_SCRA
CASP_CLIPC
FREP_NIMROD
CLIPA_CLIPB
CLIPA_TEP
APHAG_JASTMOD
APHAG_HPX
CLIPD_HPX
FREP_TEP
CLIPB_FREP
APHAG_SRRP
GNBP_ML
APHAG_TPX
FREP_LRIM
HPX_SOD
HPX_SRRP
JASTMOD_SRRP
TOLLMOD_TPX
IMDMOD_TPX
CLIPA_CLIPE
HPX_SCRA
APHAG_SOD
SRRP_TPX
JASTSIG_SCRB
CLIPD_SPZ
CLIPE_FREP
CTL_LRIM
CLIPB_SRPN
SCRA_TOLL
IMDSIG_JASTSIG
IMDMOD_SCRB
GALE_SCRB
SRPN_TEP
CASP_GNBP
GPX_ML
CLIPE_SRPN
SCRB_TPX
GNBP_GPX
CLIPA_CLIPC
TOLLSIG_TPX
CLIPC_PPO
GPX_SRPN
GNBP_PGRP
CLIPA_SPZ
SPZ_TPX
NIMROD_SCRA

### EvolMedian vs. ExprModule3

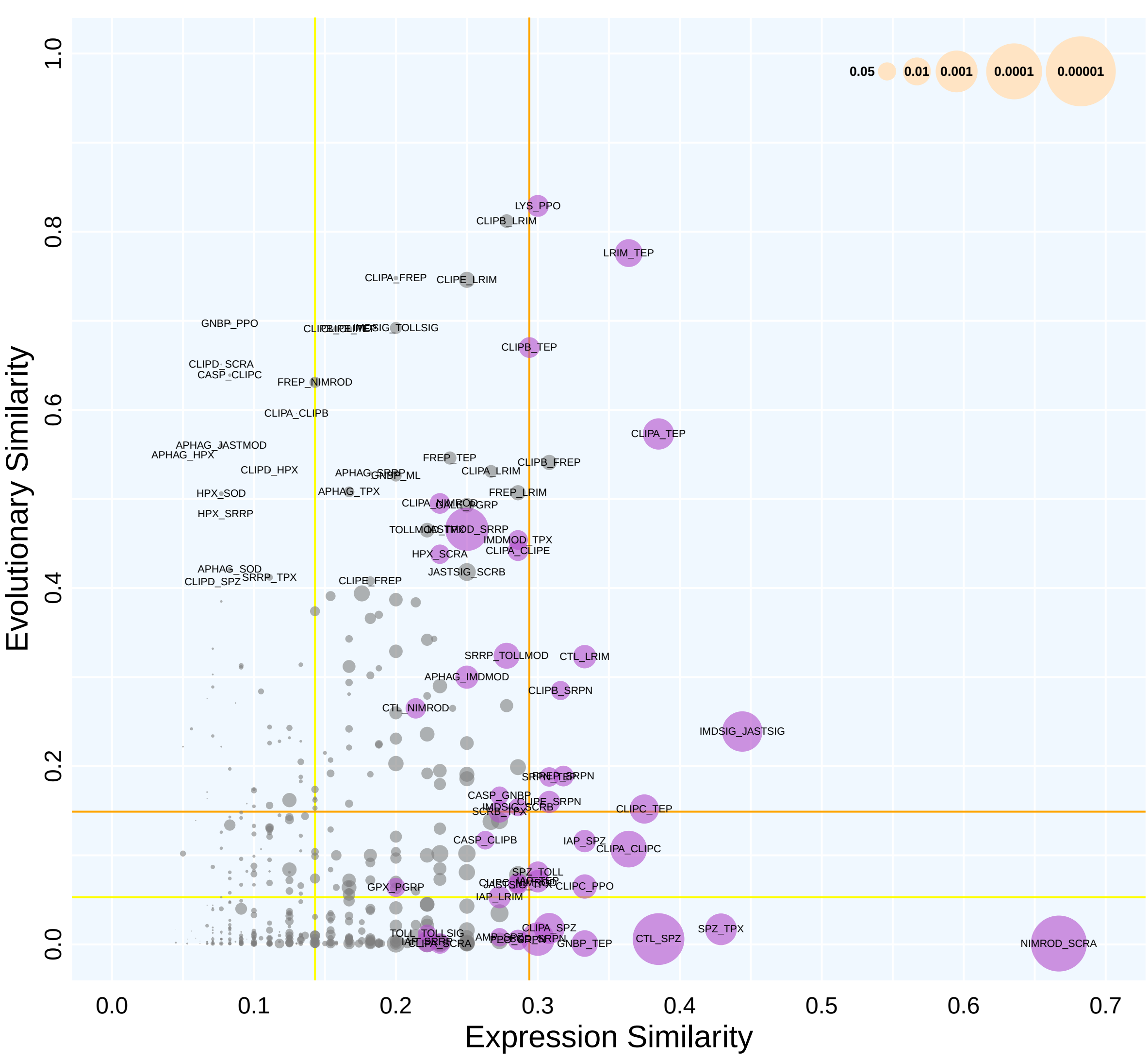

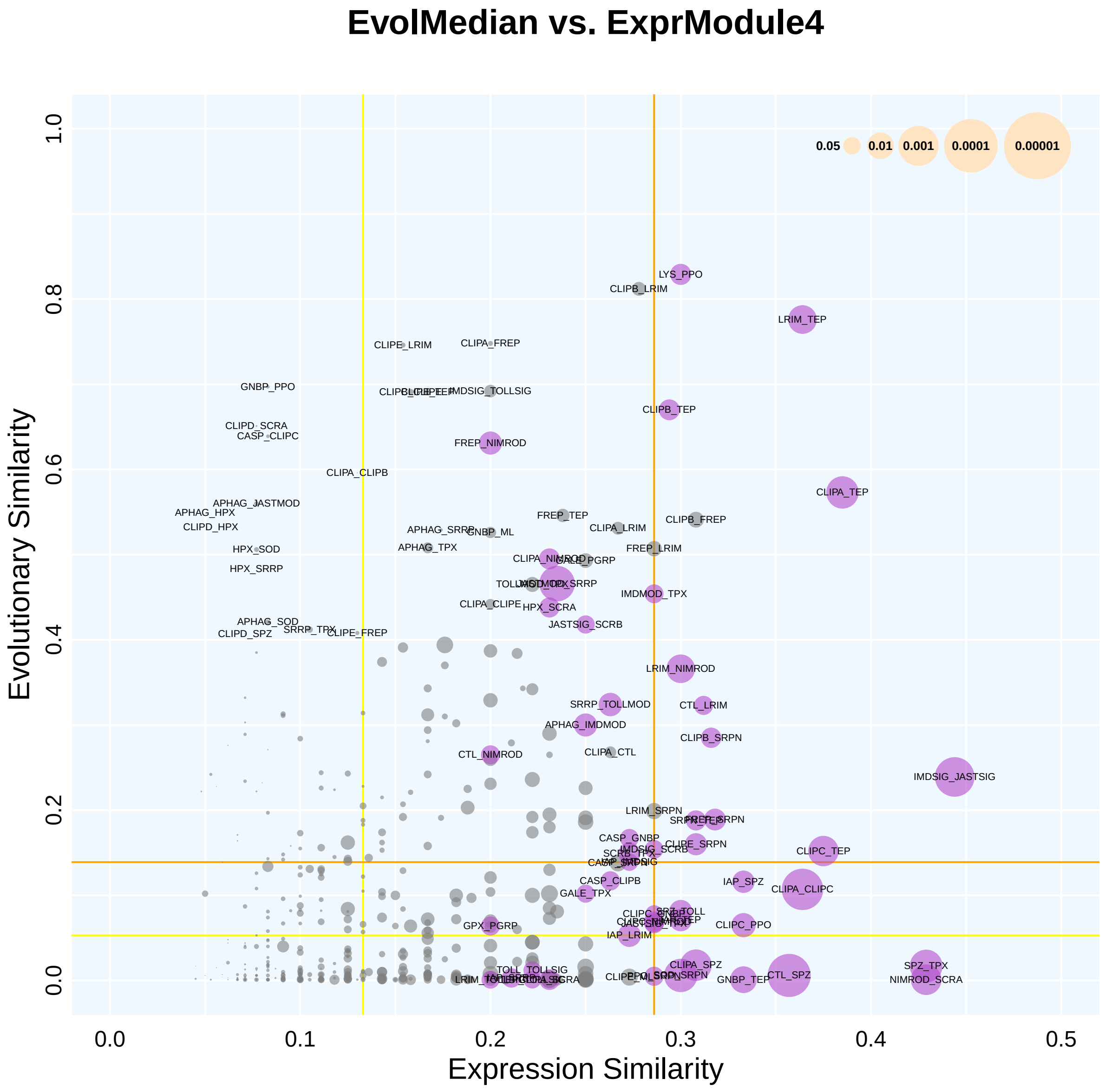
EvolMedian vs. ExprModule4
Evolutionary Similarity
Expression Similarity
0.0
0.1
0.2
0.3
0.4
0.5
0.2
0.4
0.6
0.8
1.0
0.05
0.01
0.001
0.0001
0.00001
LYS_PPO
CLIPB_LRIM
LRIM_TEP
CLIPE_LRIM
CLIPA_FREP
GNBP_PPO
IMDSIG_TOLLSIG
CLIPB_TEP
CLIPD_SCRA
CASP_CLIPC
FREP_NIMROD
CLIPA_CLIPB
CLIPA_TEP
APHAG_JASTMOD
APHAG_HPX
CLIPD_HPX
FREP_TEP
CLIPB_FREP
APHAG_SRRP
GNBP_ML
CLIPA_LRIM
HPX_SOD
APHAG_TPX
FREP_LRIM
HPX_SRRP
GALE_PGRP
IMDMOD_TPX
CLIPA_CLIPE
HPX_SCRA
APHAG_SOD
SRRP_TPX
CLIPD_SPZ
CLIPE_FREP
JASTSIG_SCRB
LRIM_NIMROD
SRRP_TOLLMOD
CTL_LRIM
APHAG_IMDMOD
CLIPB_SRPN
CTL_NIMROD
CLIPA_CTL
IMDSIG_JASTSIG
LRIM_SRPN
CASP_GNBP
CLIPE_SRPN
CLIPC_TEP
CASP_CLIPB
IAP_SPZ
CLIPA_CLIPC
GALE_TPX
CLIPC_PPO
GPX_PGRP
IAP_LRIM
CLIPA_SPZ
SPZ_TPX
NIMROD_SCRA
GNBP_TEP
CTL_SPZ
TOLL_TOLLSIG
